# Single-nucleus chromatin accessibility time course of excisional wound repair in rat skin

**DOI:** 10.64898/2026.10.15.759293

**Authors:** Chaochao Chai, Xinran Ding, Yixu Li, Langchao Liang, Runzhi Huang, Wei Zhang, Ruikang Li, Sujie Xie, Xiaoqi Zeng, Lianrui Zhang, Yue Yuan, Xiaoyu Wei, Jufang Wang, Shuai Wang, Shizhao Ji

**Author notes:** **Corresponding author. E-mail:** (S.J.); (S.W.). These authors contributed equally (Chaochao Chai, Xinran Ding, Yixu Li).

## Abstract

We present a time-series single-nucleus ATAC-seq dataset from full-thickness excisional wounds in Sprague-Dawley rats. Samples were taken from unwounded skin (normal control) and at 1, 2, 5, 10, and 19 days post-wounding, spanning the inflammatory, proliferative, and remodeling phases of skin repair. We extracted nuclei with a two-step tissue dissociation protocol and prepared libraries using the DNBelab C Series Single-Nucleus ATAC Kit, then sequenced on the MGI DNBSEQ-T1 platform. After stringent quality filtering, 37,555 nuclei passed quality control; the resulting chromatin accessibility profiles cover 23 cell types: fibroblast subpopulations, keratinocytes, vascular endothelial cells, immune cells, and skin appendage cell types. This genome-wide chromatin accessibility dataset is a resource for studying regulatory dynamics during excisional wound repair and for comparing results with other wound healing systems.

## Data Description

### Context

Wound healing follows a predictable sequence of inflammation, proliferation, and remodeling[1]. In adult mammals, it usually ends in a scar, with loss of skin appendage function[2]. Single-cell transcriptomics sequencing has sharpened what we know about cellular heterogeneity during skin repair[3–8], but the epigenetic regulatory mechanisms behind wound repair are still poorly mapped, especially at the single-cell level. Chromatin accessibility, or how open specific genomic regions are to transcription factors and other regulatory proteins, is a key layer of epigenetic regulation. For rat skin, single-cell chromatin accessibility data so far come only from burn injury models[9–11]. Burns involve coagulative necrosis and inflammatory responses that differ from excisional wounds, which heal through granulation tissue formation, re-epithelialization, and contraction. To fill this gap, we generated a temporal single- nucleus ATAC-seq (snATAC-seq) dataset from full-thickness excisional wounds in Sprague-Dawley (SD) rats, covering unwounded skin (normal control, NC) and five post-wounding time points: days 1, 2, 5, 10, and 19. The dataset lets researchers explore chromatin accessibility dynamics during excisional wound repair.

## Methods

### Sampling strategy

All animal protocols were approved by the BGI Experimental Animal Ethics Committee. Eight-week-old healthy male SD rats (Vital River Laboratory Animal Technology Co.), 220–280 g were acclimatized for 7 days under controlled conditions (temperature 22 ± 2 °C, 50–70% humidity, 12-hour light/dark cycle) with ad libitum food and water. One day before surgery, the dorsal skin was shaved and depilated. Rats were anesthetized with isoflurane and placed on a thermostatic surgical pad. After skin disinfection, full-thickness excisional wounds (1 cm diameter) were created on both sides of the dorsal midline with sterile instruments, completely removing the epidermis and full-thickness dermis. Hemostasis was achieved and wounds were disinfected with chlorhexidine diacetate and irrigated with sterile saline. The wounds were then covered with sterile petrolatum gauze and wrapped with a breathable compression bandage. Dressings were changed every other day. Post-surgery, rats were housed individually to prevent self-inflicted injury, with animals closely monitored and provided free access to food and water.

Normal uninjured dorsal skin served as the control. Wound tissues were collected at days 1, 2, 5, 10, and 19 post-wounding. At collection, rats were re-anesthetized, and wound center and surrounding repaired skin tissues (approximately 1 cm²) were excised en bloc. Subcutaneous fat and muscle debris were removed, tissues were rinsed with ice-cold sterile PBS (GIBCO, 10010-031) to eliminate blood contamination, and immediately transferred to tissue preservation solution (Miltenyi Biotec, 130-100-008) for temporary storage. Nuclei extraction and subsequent snATAC-seq library construction were completed within 48 hours **(Figure 1)**. For time points in Supplementary Table 1 with two libraries sharing the same AnimalID, the paired libraries were prepared from the same animal-derived sample as technical (library-preparation) replicates. These replicates were used to assess consistency in library preparation, sequencing quality, and downstream data processing, but were not considered independent biological replicates for biological inference.

**Figure 1.**
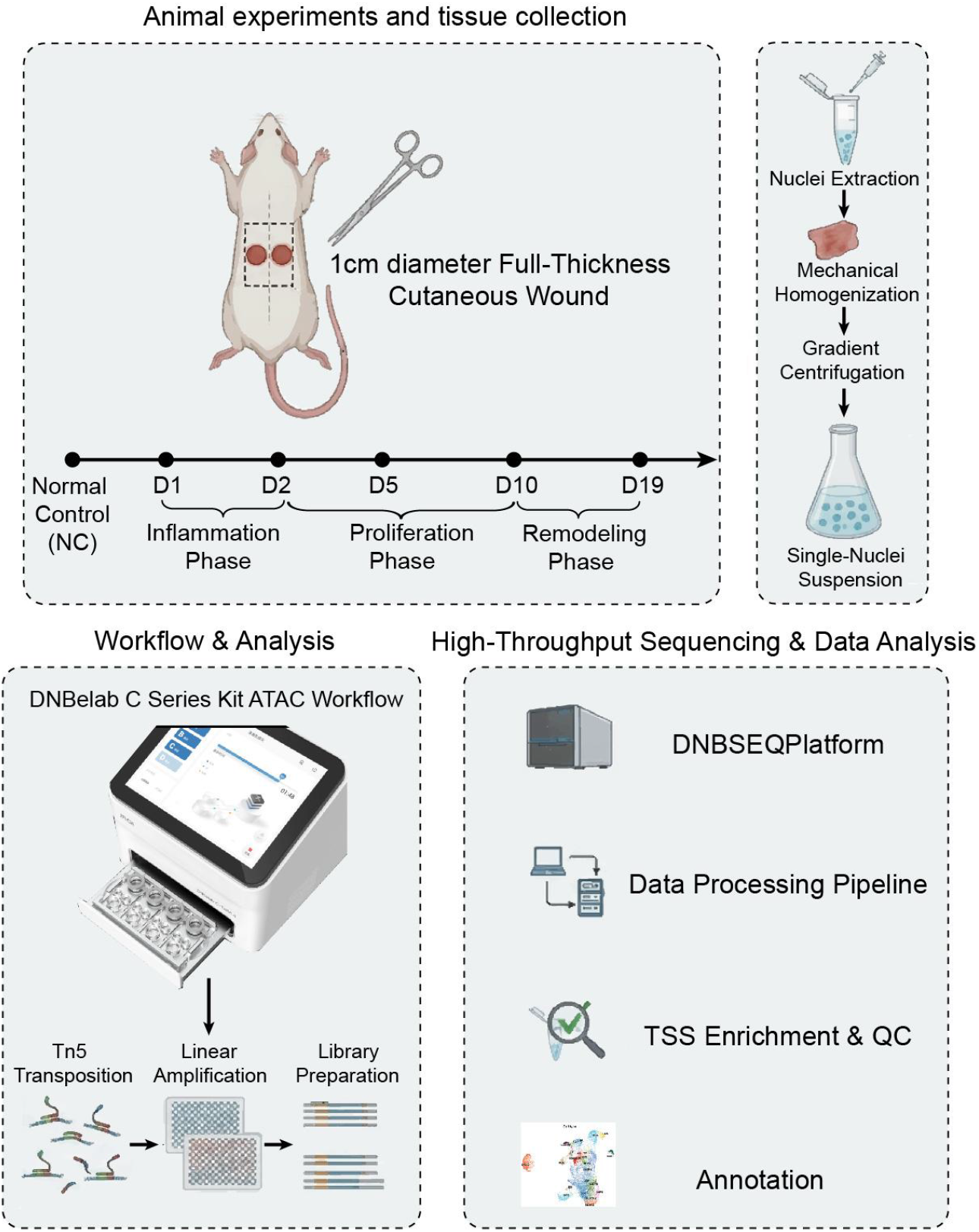
Schematic overview of the experimental design and bioinformatics workflow. Full-thickness cutaneous wounds (1 cm diameter) were created on the dorsal skin of Sprague-Dawley rats. Wound tissues were collected at normal control (NC) and days 1 (1D), 2 (2D), 5 (5D), 10 (10D), and 19 (19D) post-wounding. Single nuclei were extracted using a two-step enzymatic digestion, homogenized, and purified for snATAC-seq library construction using the DNBelab C Series Kit, followed by high-throughput sequencing on the DNBSEQ platform and downstream data processing including demultiplexing, quality control, alignment, and chromatin accessibility analysis.

### Steps

Nuclei extraction. Rat skin specimens were first rinsed in PBS (GIBCO, 10010-031) and then transferred into a 2 mL tube containing 1 mL of digestion buffer 1 (PBS supplemented with 1% penicillin-streptomycin (100× stock, CYTIVA, SV30010) and 0.25% trypsin (GIBCO, 15050065). The tissue was finely minced with scissors in the tube, and the entire contents were transferred to a 50 mL conical tube. The original 2 mL tube was rinsed with additional digestion buffer 1, and the rinse was added to the 50 mL tube. The tube was tightly sealed, placed inside a polyethylene (PE) glove to prevent leakage, and incubated at 37°C with continuous shaking at 130 rpm for 25 min.

The mixture was centrifuged at 500 × g for 5 min, and the supernatant was aspirated. The pellet was resuspended in 10 mL of digestion buffer 2 (DMEM/F-12 (GIBCO, 11320-033) containing collagenase IV (BBI, A004186-0001), collagenase I (BBI, A004194-0001), dispase II (SIGMA, D4693), and 1% penicillin-streptomycin). This suspension was incubated under the same conditions (37°C, 130 rpm) for 60 min. After digestion, the cell suspension was filtered through a 70 μm cell strainer (Falcon, 352350), and the strainer was rinsed with DPBS (BBI, 02-023-1A) supplemented with 2% FBS (CellMax, SA101) to recover additional cells. The filtrate was then passed through a 30 μm filter. Cells were collected by centrifugation at 300 × g for 5 min at 4°C, resuspended in PBS containing 0.04% BSA (Bio-Light Biotech, BL-SJ-0135), and viability was assessed by trypan blue exclusion (INVITROGEN, 15250061) using a Countstar automated cell counter. Qualified nuclei suspensions were immediately used for snATAC-seq library preparation.

#### snATAC-seq library construction and sequencing

Libraries were constructed using the DNBelab C Series Single-Nucleus ATAC Kit (MGI, H-020-000747-00)[12]. For samples with paired libraries from the same AnimalID, paired snATAC-seq libraries were generated from the same animal sample as technical replicates. They were used to assess consistency in library preparation and sequencing and for data integration, but not treated as independent biological replicates. The workflow included nuclei transposition, droplet encapsulation, amplification capture, oil phase removal, magnetic bead enrichment, DNA amplification, and fragment purification. Purified products were quantified using the Qubit fluorometric assay. Qualified samples were indexed with specific sequencing indices. Libraries were sequenced on the MGI DNBSEQ-T1 platform with read parameters: Read 1 at 115 bp, Read 2 at 69 bp, and sample index length of 10 bp.

#### Raw data preprocessing

Demultiplexing of raw sequencing data was performed based on index sequences. Quality filtering was executed using PISA (v1.1)[13] with a base quality threshold of Q >= 20; low-quality reads, adapter-contaminated sequences, and reads with excessive N bases were discarded. Quality-filtered clean reads were mapped to the SD rat reference genome mRatBN7.2 using BWA (v0.7.17-r1188)[14]. The bap2 tool (v0.6.2) was subsequently used to parse BAM files[15], assigning sequencing fragments to individual nuclei based on cell barcodes.

#### Bioinformatics analysis

All single-nucleus chromatin accessibility data analyses were performed using SnapATAC2 (v2.7)[16] in Python.

Quality control and nuclei filtering. Preprocessed BAM files were imported to construct SnapATAC objects. Nuclei with > 3,000 uniquely mapped fragments and TSS enrichment score > 4 were retained. Function snapatac2.pp.scrublet function was used to calculate per-nucleus doublet probabilities, and predicted doublets were removed using the built-in doublet detection algorithm. Dimensionality reduction, integration, and clustering. Qualified fragment matrices were merged, and 500,000 genomic features were selected. Laplacian Eigenmaps were computed for spectral embedding. Harmony(harmonypy, v0.0.10) batch correction was applied to the spectral low-dimensional feature matrix to eliminate batch effects across libraries and time points[17]. The library identifier was used as the batch covariate for integration. This correction was performed after dimensionality reduction and before neighbor-graph construction to align nuclei across experiments while preserving biological variation associated with wound healing progression. Neighborhood graph clustering was performed using the Leiden algorithm (python-igraph v0.11.8)[18], followed by UMAP nonlinear dimensionality reduction for visualization. Gene activity scores were calculated using the function of snapatac2.pp.make_gene_matrix from built-in SnapATAC module with default parameters.

Cell type annotation and sub-clustering. Cell-type-specific accessibility-derived gene activity markers were identified using the FindAllMarkers function in Seurat (v4.3.0). These marker features, together with published rat skin cell marker genes, were used to manually annotate all cell clusters. The clustering resolution was increased using function FindSubCluster for keratinocytes and fibroblasts to further subdivide functionally distinct subpopulations.

Cell-type-specific peak calling and marker peak identification. Pseudo-bulk profiles were constructed for each cell type using snap.tl.macs3 (MACS3, v3.0.2), and peak calling was performed for each cell subpopulation with default parameters. After peak calling, peak-by-nucleus accessibility count matrix was generated using default parameters of snapatac2.pp.make_peak_matrix. Cell-type-specific marker peaks were identified from this matrix using Seurat FindAllMarkers. For each cell type, nuclei assigned to that cell type were compared with all remaining nuclei using the Wilcoxon rank-sum test, followed by Seurat’s default Bonferroni correction for multiple testing. FindAllMarkers was run with only.pos = TRUE, logfc.threshold = 0.25, and return.thresh = 0.05; all other parameters were kept at their default values, including test.use = “wilcox” and min.pct = 0.01. Positively enriched marker peaks with log fold-change > 0.25 and Bonferroni-adjusted P-value < 0.05 were retained for downstream analyses, and coding genes within ±2 kb of marker peak summits were annotated. Accessibility-derived gene activity scores were calculated using the built-in SnapATAC2 module. Cell-type-specific gene activity markers were then identified from the gene activity matrix using Seurat FindAllMarkers with the same statistical test, multiple-testing correction, and filtering thresholds.

Temporal dynamic peak clustering. To identify temporally dynamic accessible peaks, the peak count matrix was first normalized within each cell subpopulation using term frequency–inverse document frequency (TF-IDF) transformation. Nuclei within each time point were then aggregated by computing the mean TF-IDF-normalized accessibility value for each peak using the AverageExpression function in Seurat. Here, AverageExpression was applied to chromatin accessibility values rather than RNA expression values. This generated a pseudobulk accessibility matrix for each cell subpopulation across the six time points (NC, 1D, 2D, 5D, 10D, and 19D). Peaks were retained if they showed detectable accessibility (row mean > 0) and nonzero variance across time points, and the 50,000 most temporally variable peaks were selected for downstream analysis. Peaks with a temporal profile standard deviation below 0.15 were removed using the filter.std function. The remaining peak profiles were then standardized to a mean of zero and a standard deviation of one across time points using the standardise function, so that clustering was driven by temporal pattern rather than absolute accessibility magnitude. The Mfuzz fuzzy c-means algorithm was then applied to the standardized peak profiles. The fuzziness parameter m was estimated from the data using the mestimate function and was set to 1.704. The number of clusters was determined using the minimum centroid distance calculated by the Dmin function across c = 2–12, with three repeated runs for each c. Based on the inflection of the Dmin curve and the interpretability of the resulting temporal modules, c = 8 was selected for downstream analysis. Clustering was performed with a fixed random seed using set.seed(42) to ensure reproducibility[19]. In the main analysis, no membership-based filtering cutoff was applied; because the majority of peaks showed high membership values, each peak was assigned to the cluster for which it had the highest membership value (median membership across all peaks: 0.815). Gene Ontology (GO) functional enrichment was performed on genes associated with each peak cluster.

Motif identify and cell-type-specific motif activity analysis. Cell-type-specific accessible peaks were identified by differential accessibility analysis in the integrated dataset (FindAllMarkers; Benjamini-Hochberg-adjusted P ≤ 0.05, |avg_log2FC| ≥ 0.25). Motif information was added using the JASPAR2020 CORE collection of vertebrate position frequency matrices (all_versions = FALSE). For each peak, GC content and dinucleotide frequencies were computed against the Rattus norvegicus reference genome (mRatBN7.2 assembly) using RegionStats. Motif accessibility deviations were calculated for each nucleus using the RunChromVAR function in Signac (v1.9.0), using previous cell-type-specific differential peaks as the background peak set. This analysis compares observed motif accessibility with accessibility in background peaks matched for GC content and dinucleotide composition, yielding bias-corrected motif deviation scores. To identify cell-type-specific motif activity patterns, differential motif deviation analysis was performed on the chromVAR assay using FindAllMarkers. Motifs with Wilcoxon rank-sum test, Benjamini-Hochberg-adjusted P ≤ 0.05, log2 fold-change ≥ 0.25 were retained, with only.pos = TRUE. For each selected motif, per-cell deviation z-scores were averaged across endothelial cells within each wound-healing time point (normal, days 1, 2, 5, 10, and 19).

Statistical analysis. Cell-type-specific marker peaks and accessibility-derived gene activity markers were identified using the FindAllMarkers function in Seurat. For each cell type, nuclei assigned to that cell type were compared with all remaining nuclei using the Wilcoxon rank-sum test, followed by Seurat’s default Bonferroni correction for multiple testing. FindAllMarkers was run with only.pos = TRUE, logfc.threshold = 0.25, and return.thresh = 0.05. These single-nucleus-level comparisons were used to identify candidate marker features for resource annotation and hypothesis generation. Because paired libraries sharing the same AnimalID represented technical/library replicates rather than independent biological replicates, the results were not interpreted as independent biological replicate-level statistical inference. GO enrichment analysis was performed using clusterProfiler (v4.0)[20], and terms with Benjamini-Hochberg adjusted P-value < 0.05 were considered significant.

### Data Validation and Quality Control

We performed snATAC-seq on nuclei isolated from the wound beds (including wound edges) of full-thickness excisional wounds in SD rats at days 1, 2, 5, 10, and 19 post-wounding, with unwounded skin serving as the normal control (NC). Data generation followed a standardized workflow for stringent quality control and analysis using SnapATAC2 **(Figure 1)**. After excluding low-quality nuclei with transcription start site (TSS) enrichment scores ≤ 4 and unique fragment counts ≤ 3,000, a total of 37,555 high-quality nuclei were retained **(Figure 2A)**. Notably, the yield of nuclei from day 19 (D19) samples was lower than that at other time points **(Supplementary Table 1)**. This drop reflects the dense, fibrotic extracellular matrix (ECM) of late-stage remodeling in full-thickness skin wounds. The tightly cross-linked matrix at this stage makes mechanical homogenization and nuclei extraction difficult, a known challenge in late-stage wound healing studies. Median TSS enrichment scores ranged from 6.18 to 16.77 across libraries, and median unique fragment counts ranged from 4,370 to 25,243, indicating robust technical quality and sufficient sequencing depth **(Figure 2B, Supplementary Table 1)**. Quality control (QC) metrics were generally consistent between technical/library replicates at the corresponding time points **(Figure 2C)**, supporting the reproducibility of library preparation and sequencing. Batch correction after integration effectively reduced library-associated variation, further supporting the technical consistency of the dataset **(Figure 2D)**.

**Figure 2.**
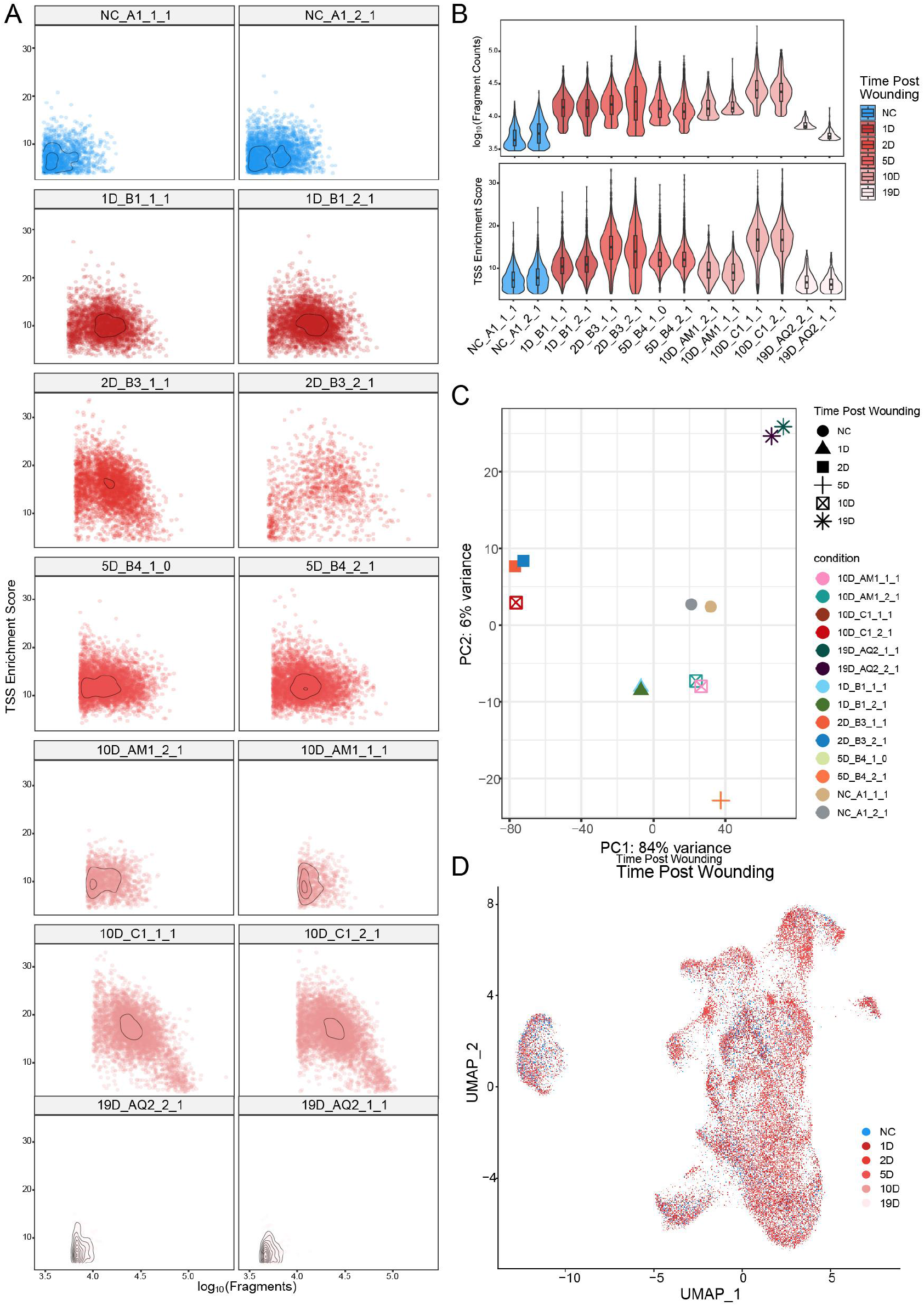
Quality control metrics of the snATAC-seq dataset. (A) Quality control scatter plots showing transcription start site (TSS) enrichment score versus log10(fragments) for each individual sample. Each point represents a single nucleus, with density contours highlighting cell populations. The color of each point represents different post-wounding time periods. (B) Violin plots showing the distribution of log10(fragment counts) (upper panel) and TSS enrichment scores (lower panel) across all samples. (C) Principal component analysis (PCA) of all sequencing libraries. Each point represents an individual library, colored by time point post-wounding. (D) Uniform Manifold Approximation and Projection (UMAP) visualization of all single nuclei colored by time point post-wounding (NC, 1D, 2D, 5D, 10D, and 19D).

The dataset covers all major functional compartments of the skin microenvironment **(Figure 3A)**. Using gene activity scores of canonical markers, we annotated 23 cell types **(Supplementary Table 2)**. Fibroblasts were split into eight subtypes by their chromatin accessibility profiles (*Apcdd1* for papillary fibroblasts (papiFB), *Cxcl12* for reticular fibroblasts (retiFB), *Enpp2* for dermal papilla (DP), etc.)[21], and keratinocytes were resolved into suprabasal and basal subpopulations. The data also captured skin appendage- related cells (sebaceous gland cells (SG), hair follicle stem cells (HFSC), inner root sheath cells (IRS), etc.) and a range of non- epithelial cells: endothelial cells (ENDO), pericytes (PER), muscle cells (MUS), Schwann cells (SCH) and several immune cell types (myeloid cells (MYE), Langerhans cells (LAG), natural killer cells (NK), T lymphocyte (T)) **(Figure 3B)**. These cell types are visible across normal and healing skin **(Figure 3C, Supplementary Table 2)**.

**Figure 3.**
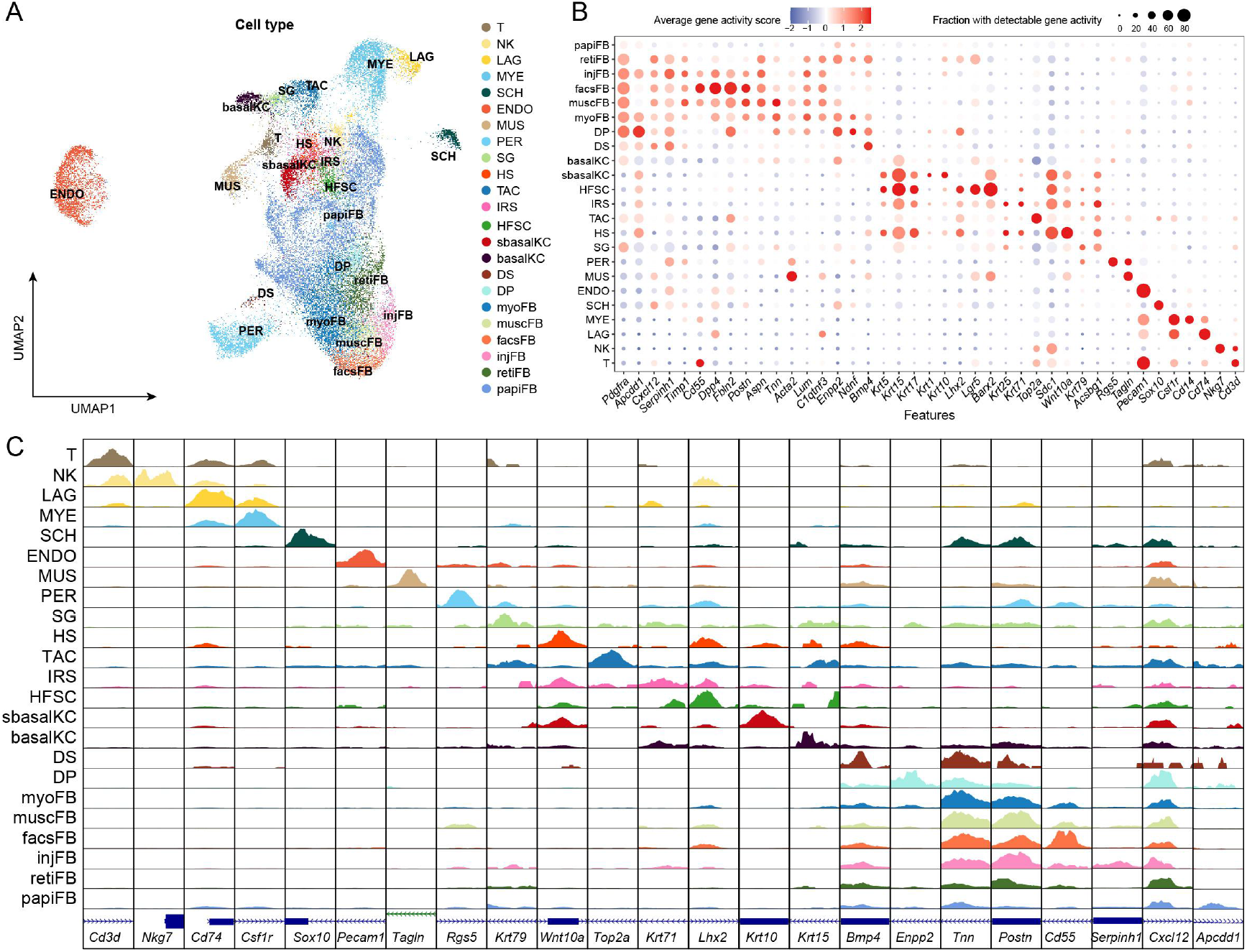
Comprehensive cell type annotation capturing the full skin microenvironment. (A) Uniform Manifold Approximation and Projection (UMAP) visualization of all cell clusters identified from the snATAC-seq data. (B) Dot plot showing the average accessibility-derived gene activity score (color scale) and the fraction of nuclei with detectable gene activity (dot size) for representative marker genes across all identified cell types. Color from blue to red indicates the average gene activity score from low to high. Dot size is proportional to the fraction of nuclei with detectable gene activity. (C) Coverage plots showing the accessibility profiles of selected marker peak regions across the identified cell types.

We further evaluated cell-type identity at the epigenetic level using cell-type-specific marker peaks, accessibility-derived gene activity markers, and chromVAR-derived cell-type-specific motif activity markers. Each cell type showed characteristic chromatin accessibility at marker regions, gene activity patterns, and motif accessibility deviation profiles consistent with its annotated identity. For example, accessibility-derived gene activity scores and chromatin accessibility at *Krt5* and *Krt10* distinguished basal from suprabasal keratinocytes, whereas *Pdgfra* and fibroblast subtype-associated markers showed distinct accessibility-derived gene activity and chromatin accessibility patterns across fibroblast subtypes. In addition, endothelial cells were enriched for vascular-associated motifs such as *FOXC2, HEY1* and *NR2F1*, further supporting their regulatory identity. These results support the cell-type annotations and provide a reference for studying cell-type-specific cis-regulatory elements in cutaneous biology. **(Figure 4, Supplementary Table 3-4)**.

**Figure 4.**
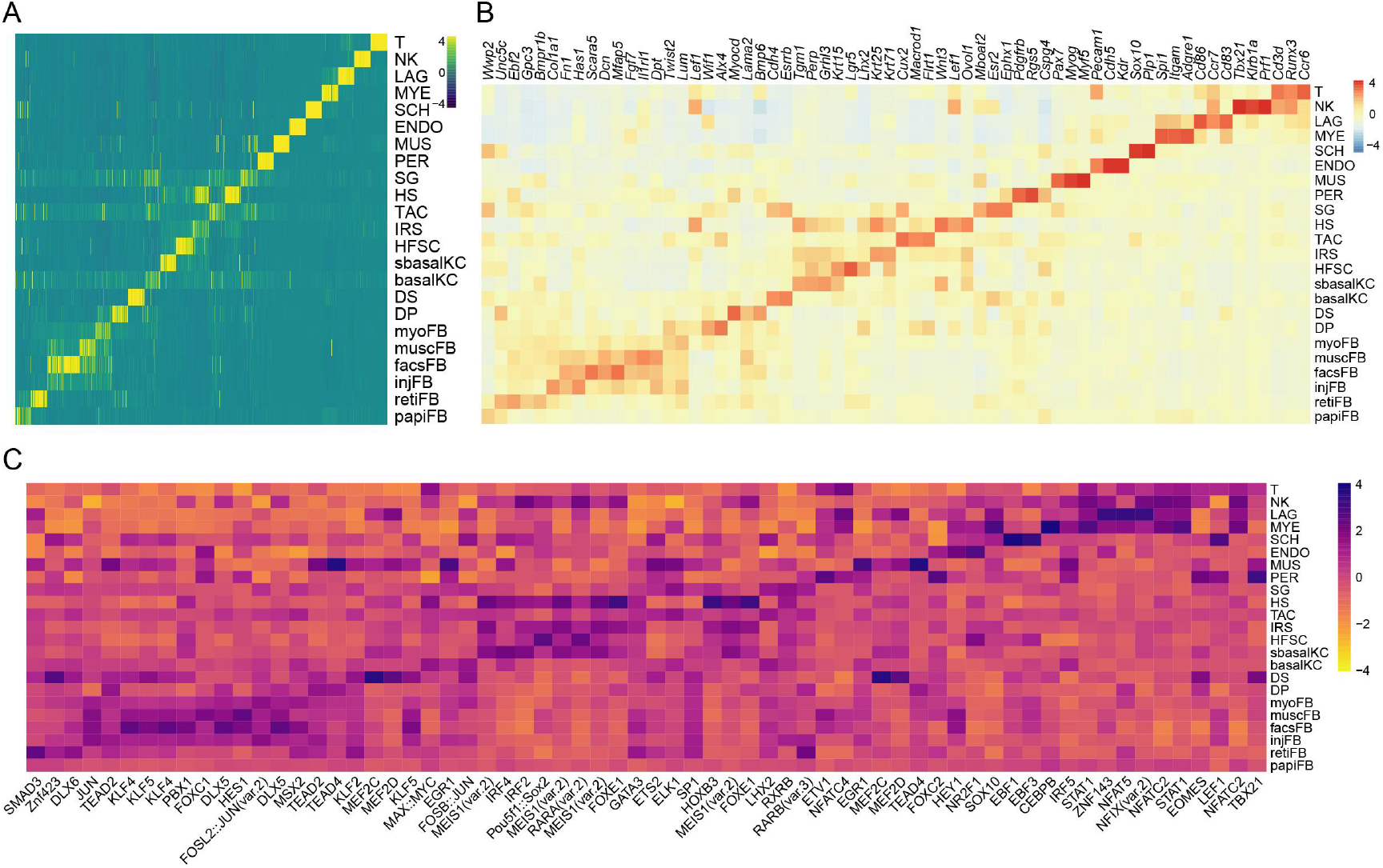
Cell-type-specific chromatin accessibility, gene activity, and transcription factor motif accessibility deviations. (A) Heatmap displaying cell-type-specific marker peaks identified from the peak-by-nucleus chromatin accessibility matrix. (B) Heatmap showing scaled accessibility-derived gene activity scores of representative cell-type-specific marker genes identified from the gene activity matrix. (C) Heatmap showing cell-type-specific chromVAR-derived motif accessibility deviation scores. Marker motifs were identified from the global chromVAR analysis using FindAllMarkers. Heatmap values represent scaled deviation scores across cell types.

Beyond the static cell atlas, the dataset captures how cell states shift over the course of wound healing. Because the D19 samples contained substantially fewer nuclei than other time points, cell-type proportion analyses were restricted to NC through D10 to reduce potential bias from differential nuclei recovery **(Figure 5A)**. Temporal chromatin accessibility patterns were then examined across the wound-healing time course **(Figure 5B, Supplementary Table 5)**. Taking endothelial cells (ENDO) as an example, the data trace a chromatin remodeling timeline through wound angiogenesis: acute stress and barrier disruption on day 1, early angiogenic activation on day 2, endothelial migration and proliferation across days 2 to 5, followed by vascular maturation and remodeling at day 19 **(Figure 5C-E)**. Specifically, the TAL1::TCF3 motif (MA0091.1) had the highest activity at day 1. This is consistent with previous reports implicating it in hypoxia signaling and early vascular remodeling[22]. At day 2, IRF7 motif activity increased, suggesting an interferon response during the early inflammatory phase of wound healing[23]. At day 5, the HEY1/HES5/HES7 motif had the highest accessibility. These Notch effectors regulate vessel sprouting and branching during angiogenesis[24]. After an initial decrease at days 1–2, BHLHE41/DEC2 motif activity slightly above baseline at day 19. This recovery paralleled increased ARNT::HIF1A motif activity and may point to a late hypoxic-adaptive response, with BHLHE41/DEC2 restraining VEGF[25]. Overall, the temporal changes in motif activity outline a regulatory trajectory associated with endothelial maturation during wound healing **(Figure 5F)**. This time-resolved dataset provides an epigenetic reference for generating and prioritizing hypotheses about regulatory mechanisms underlying major cell-state changes during skin repair.

**Figure 5.**
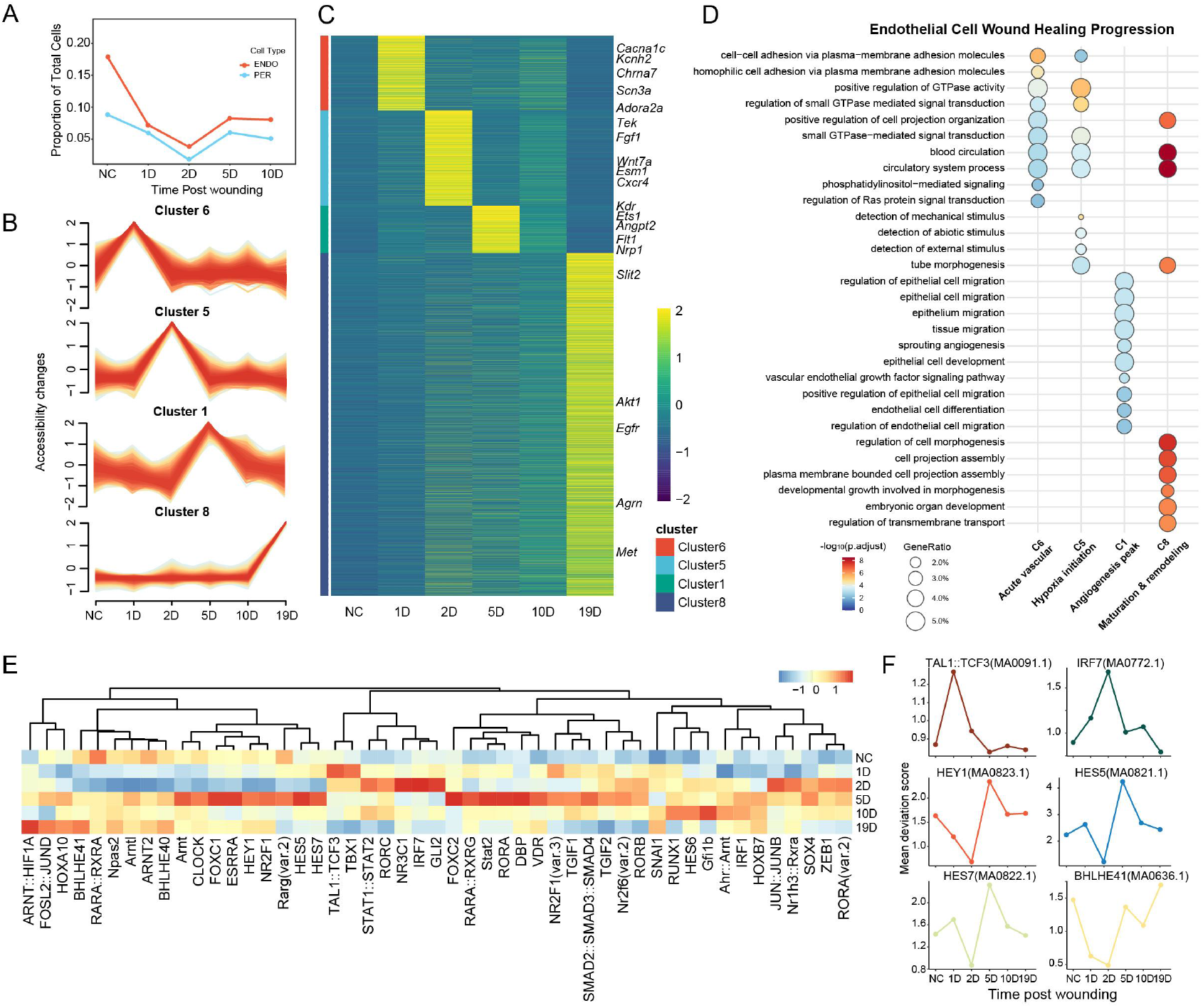
Temporal chromatin dynamics during wound repair. (A) Dynamic changes in the proportion of ENDO and PER cell types during wound healing progression. The line plot shows the relative abundance across time points (NC, 1D, 2D, 5D, 10D). (B) Temporal dynamics of accessible peak clusters in endothelial cells during wound healing. Line plots show accessibility change patterns for four representative clusters (Cluster 6, 5, 1 and 8) from NC to 19D post-wounding. (C) Heatmap showing the temporal accessibility patterns of all accessible peaks identified by Mfuzz clustering in endothelial cells. Peaks are grouped by cluster. For visual clarity, only selected functionally representative genes are annotated on the right. (D) Gene Ontology (GO) enrichment analysis of genes associated with dynamic accessible peak modules in endothelial cells across wound healing stages. Selected biological process terms are shown for four functional modules: C5 (Hypoxia initiation), C6 (Acute vascular response), C1 (Angiogenesis peak), and C8 (Maturation & remodeling). Bubble size represents the gene ratio, and color intensity indicates statistical significance (−log10 adjusted p-value). (E) Heatmap showing mean chromVAR-derived motif accessibility deviation scores in endothelial cells across wound-healing time points. Per-cell deviation z-scores were averaged across endothelial cells within each time point. (F) The line plot shows mean chromVAR deviation z-scores of representative stage-specific endothelial transcription factor motifs across wound-healing time points.

### Limitations and cross-species considerations

Although this dataset provides a time-resolved, single-nucleus chromatin accessibility reference for full-thickness excisional wound repair in rat skin, several species- and model-specific limitations should be considered when applying these data to human wound healing. Rat skin differs from human skin in terms of tissue architecture, hair follicle density, immune composition, and the relative contribution of wound contraction versus re-epithelialization during closure[26]. Specifically, wound closure in rodents is strongly influenced by contraction mediated by the panniculus carnosus, whereas human cutaneous wound repair relies more heavily on granulation tissue formation and re-epithelialization[26,27]. These anatomical and physiological differences can affect the cellular composition, timing of repair phases, and inferred during wound repair. Therefore, the chromatin accessibility patterns identified in rat wounds should not be interpreted as direct equivalents of human regulatory elements without cross-species validation.

However, the rat excisional wound model remains a valuable tool for understanding conserved cellular and regulatory processes involved in mammalian tissue repair, including inflammatory activation, fibroblast remodeling, keratinocyte regeneration, angiogenesis, and extracellular matrix reorganization. This dataset can therefore serve as a reference for identifying conserved and species-specific regulatory programs, particularly when combined with human wound-healing datasets or other animal models. Future studies combining orthologous gene mapping, conserved motif analysis, and cross-species single-cell or single-nucleus multi-omics datasets will be important for evaluating the translational relevance of the regulatory elements and transcription factor programs identified here[28].

### Re-use Potential

This dataset can be used for comparative studies of chromatin accessibility dynamics during excisional skin wound repair under normal conditions. It is a time-resolved epigenetic reference for systematic comparison with other wound healing models, to identify regulatory programs shared across models and those specific to each. It complements existing burn injury snATAC-seq datasets in rats, making it possible to compare mechanical excision directly with thermal injury. The data can be integrated with single-cell transcriptomic atlases of skin wound healing through standard multimodal integration methods to jointly analyze gene expression and chromatin accessibility. More broadly, cell-type-resolved molecular modules have been useful for identifying shared mechanisms across complex biological conditions and diseases[29], and this dataset may support similar module-based comparisons across wound models, injury types, or regenerative contexts. The dataset is also suitable for benchmarking single-cell epigenomic analysis pipelines, especially batch correction and temporal trajectory inference algorithms. Beyond wound repair, the data may be useful for studies of angiogenesis, fibroblast activation, and immune cell recruitment in tissue regeneration and fibrosis. To facilitate reuse by experimental investigators, we provide processed matrices and interactive visualization through CDCP, where users can explore cell-type annotations, marker gene activity, chromatin accessibility profiles, and temporal changes across wound-healing stages.

## Data Availability Statement

The raw snATAC-seq sequencing data generated in this study have been deposited in the CNGBdb-CNSA[30] under accession number CNP0009888.

The processed snATAC-seq data matrices and visualization files have also been deposited in the Cell-omics Data Coordinate Platform (CDCP) under project accession SCDS0000647.

The processed peak-by-nucleus matrix, metadata, and analysis scripts are available in the GigaDB repository at https://gigadb.org/dataset/102833/token/YZxJXRGCUqmK2vBj.

## List of Abbreviations

NC: normal control
SD: Sprague-Dawley
BSA: bovine serum albumin
DPBS: Dulbecco’s phosphate-buffered saline
FBS: fetal bovine serum
GO: Gene Ontology
PBS: phosphate-buffered saline
snATAC-seq: single-nucleus assay for transposase-accessible chromatin using sequencing
TSS: transcription start site
PCA: Principal Component Analysis
UMAP: Uniform Manifold Approximation and Projection
QC: Quality Control
TF-IDF: term frequency–inverse document frequency
papiFB: papillary fibroblasts
retiFB: reticular fibroblasts
DP: dermal papilla fibroblasts
DS: dermal sheath fibroblasts
facsFB: fascia fibroblasts
muscFB: muscular fibroblasts
injFB: injury fibroblasts
myoFB: myofibroblasts
sbasalKC: suprabasal keratinocytes
basalKC: basal keratinocytes
SG: sebaceous gland cells
HFSC: hair follicle stem cells
IRS: inner root sheath cells
TAC: transit-amplifying cells
HS: hair shaft cells
MUS: muscle cells
ENDO: endothelial cells
PER: pericytes
SCH: Schwann cells
MYE: myeloid cells
LAG: Langerhans cells
NK: natural killer cells
T: T lymphocyte
CDCP: Cell-omics Data Coordinate Platform

## Declarations

## Ethics Approval and Consent to Participate

All animal experimental procedures were approved by the BGI Experimental Animal Ethics Committee (Approval Number: BGI-IRB A23035).

## Consent for Publication

Not applicable.

## Authors’ Contributions

Conceptualization: C.C., S.W., X.W.

Methodology: C.C., S.W., X.D., Y.L.

Data Curation: X.D., Y.L., S.X., W.Z., Y.Y., L.Z., S.W., R.H.

Formal Analysis: C.C., X.Z., S.W.

Investigation: X.D., Y.L., S.X., W.Z., Y.Y., C.C., L.L.

Visualization: C.C., R.L., L.L.

Writing – Original Draft: C.C., X.D., Y.L.

Writing – Review & Editing: C.C., S.W., S.J.

Resources: S.W., X.W., S.J., R.H.

Funding Acquisition: S.W., S.J., R.H., X.W.

Supervision: S.W., X.W., J.W., S.J., R.H.

Project Administration: S.W., X.W.

## Acknowledgements

We would like to thank DCS Cloud (https://dcs.cloud) for providing the computational resources and software support necessary for this study. We sincerely thank the China National GeneBank (CNGB) for the support they provided.

