## Supplemental Table 1 for "Single-nucleus chromatin accessibility time course of excisional wound repair in rat skin"

**Supplementary Table1. snATAC-seq metadata and median threshold values per library**

| sample | TimePostInjury | AnimalID | number_of_cells | Median_TSS.enri |
|---|---|---|---|---|
| ATAC_r_Adult_Nor_A1_1_1 | NC | A1 | 1313 | 7.22 |
| ATAC_r_Adult_Nor_A1_2_1 | NC | A1 | 2545 | 7.8 |
| ATAC_r_Adult_1D_B1_1_1 | 1D | B1 | 2896 | 10.45 |
| ATAC_r_Adult_1D_B1_2_1 | 1D | B1 | 3559 | 10.91 |
| ATAC_r_Adult_2D_B3_1_1 | 2D | B3 | 3491 | 14.97 |
| ATAC_r_Adult_2D_B3_2_1 | 2D | B3 | 1096 | 13.99 |
| ATAC_r_Adult_5D_B4_1_0 | 5D | B4 | 4608 | 11.98 |
| ATAC_r_Adult_5D_B4_2_1 | 5D | B4 | 5102 | 11.99 |
| ATAC_r_Adult_10D_AM1_1_1 | 10D | AM1 | 983 | 8.95 |
| ATAC_r_Adult_10D_AM1_2_1 | 10D | AM1 | 1709 | 9.57 |
| ATAC_r_Adult_10D_C1_1_1 | 10D | C1 | 4634 | 16.77 |
| ATAC_r_Adult_10D_C1_2_1 | 10D | C1 | 5118 | 16.72 |
| ATAC_r_Adult_19D_AQ2_1_1 | 19D | AQ2 | 275 | 6.18 |
| ATAC_r_Adult_19D_AQ2_2_1 | 19D | AQ2 | 226 | 6.67 |

Note: Libraries sharing the same AnimalID were generated as technical replicates from the same animal-d

| **Mean_nFragments** | **Median_nFragments** | **Mean_Gene_activity_nCount** |
|---|---|---|
| 5157 | 4370 | 3005.52 |
| 6292 | 5483 | 3717.52 |
| 14880 | 13780 | 9512.41 |
| 14822 | 13608 | 9535.85 |
| 16845 | 15204 | 11823.12 |
| 21644 | 16940 | 15080.36 |
| 15137 | 13084 | 10778.93 |
| 13830 | 11851 | 9859.78 |
| 15130 | 13340 | 9647.77 |
| 15034 | 13164 | 9734.45 |
| 30105 | 25243 | 21207.92 |
| 27243 | 23692 | 19164.25 |
| 5195 | 4932 | 3074.99 |
| 7506 | 7089 | 4485.40 |

erived sample and should not be interpreted as independent biological replicates.

| Median_Gene_activity_nCount | Mean_Gene_activity_nFeature |
|---|---|
| 2595 | 1879.32 |
| 3240 | 2195.34 |
| 8891 | 4279.25 |
| 8812 | 4244.61 |
| 10728 | 5538.93 |
| 12029.5 | 5533.72 |
| 9465.5 | 5067.92 |
| 8575.5 | 4741.13 |
| 8640 | 4588.66 |
| 8570 | 4592.68 |
| 18156.5 | 7649.87 |
| 16843.5 | 7317.13 |
| 2940 | 2193.42 |
| 4322.5 | 2944.63 |

.

| Median_Gene_activity_nFeature | Mean_nCount_peaks | Median_nCount_peaks |
|---|---|---|
| 1717 | 2397.45 | 2103 |
| 2010 | 3037.01 | 2663 |
| 4232 | 8843.37 | 8296 |
| 4167 | 8830.13 | 8238 |
| 5452 | 11534.77 | 10725 |
| 5341 | 14331.40 | 11743 |
| 4811.5 | 11928.34 | 10477 |
| 4500 | 10847.55 | 9473.5 |
| 4440 | 9191.22 | 8334 |
| 4376 | 9332.92 | 8296 |
| 7616 | 21247.47 | 19227.5 |
| 7310.5 | 19374.18 | 17874 |
| 2157 | 2622.24 | 2546 |
| 2930 | 3840.64 | 3744 |

| Mean_nFeature_peaks | Median_nFeature_peaks |
|---|---|
| 2216.64 | 1966 |
| 2753.22 | 2421 |
| 7033.63 | 6652.5 |
| 6926.34 | 6478 |
| 9034.91 | 8513 |
| 9433.80 | 8087 |
| 9230.96 | 8157 |
| 8397.63 | 7420.5 |
| 7666.81 | 7130 |
| 7607.18 | 6937 |
| 15444.41 | 14139.5 |
| 14406.29 | 13445.5 |
| 2515.84 | 2447 |
| 3623.21 | 3552.5 |
