## Supplemental Table 2 for "Single-nucleus chromatin accessibility time course of excisional wound repair in rat skin"

**Supplementary Table2. Marker genes for cell type annotation**

| | |
|---|---|
| FB | *Pdgfra* |
| papiFB | *Apcdd1* |
| RetiFB | *Cxcl12* |
| injFB | *Has1, Serpinh1, Timp1* |
| facsFB | *Cd55, Dpp4, Fbln2* |
| muscFB | *Postn, Aspn* |
| myoFB | *Tnn, Acta2, Lum, C1qtnf3* |
| DP | *Enpp2, Ndnf* |
| DS | *Bmp4* |
| basalKC | *Krt5, Krt15, Krt17* |
| sbasalKC | *Krt1, Krt10* |
| HFSC | *Lhx2, Lgr5, Barx2* |
| IRS | *Krt25, Krt71* |
| TAC | *Top2a, Sdc1* |
| HS | *Wnt10a* |
| SG | *Krt79, Acsbg1* |
| PER | *Rgs5* |
| MUS | *Tagln* |
| ENDO | *Pecam1* |
| SCH | *Sox10* |
| MYE | *Csf1r, Cd14* |
| LAG | *Cd74* |
| NK | *Nkg7* |
| T | *Cd3d* |
