## Supplemental Table 3 for "Single-nucleus chromatin accessibility time course of excisional wound repair in rat skin"

**Supplementary Table 3. Marker peak summary**

| p_val | avg_log2FC | pct.1 | pct.2 | p_val_adj | cell_type | peak |
|---|---|---|---|---|---|---|
| 7.79035558 | 5.15397214523123 | 0.112 | 0.004 | 5.7085077 | DP | chr10-30032234-300 |
| 5.33702057 | 3.82875420162552 | 0.157 | 0.012 | 3.9107872 | DP | chr17-75949135-759 |
| 4.85474043 | 4.11850902510082 | 0.132 | 0.009 | 3.5573887 | DP | chr12-37655669-376 |
| 2.19998378 | 3.95502419150677 | 0.137 | 0.01 | 1.6120733 | DP | chr2-209869752-209 |
| 1.61865350 | 3.79234755427148 | 0.126 | 0.009 | 1.1860942 | DP | chr1-241102109-241 |
| 1.73868020 | 3.12821917872721 | 0.207 | 0.025 | 1.2740457 | DP | chr1-169752388-169 |
| 3.19644881 | 3.63657946167998 | 0.154 | 0.014 | 2.3422490 | DP | chr17-11159716-111 |
| 4.13245326 | 4.5177750046061 | 0.104 | 0.006 | 3.0281212 | DP | chr2-166307253-166 |
| 9.90436030 | 3.6941989969025 | 0.127 | 0.01 | 7.2575784 | DP | chr1-19206048-1920 |
| 7.43017150 | 3.31058091001444 | 0.166 | 0.018 | 5.4445770 | DP | chr4-95222841-9522 |
| 4.68791336 | 4.08277413483551 | 0.1 | 0.007 | 3.4351435 | DP | chr4-126718088-126 |
| 2.53454363 | 3.28237586836539 | 0.172 | 0.02 | 1.8572274 | DP | chr8-97439707-9744 |
| 4.00206019 | 3.698781168 | 0.143 | 0.014 | 2.9325736 | DP | chr3-158317638-158 |
| 3.16002453 | 3.82321583754349 | 0.104 | 0.007 | 2.3155585 | DP | chr19-1032224-1032 |
| 1.59253290 | 3.86772681627365 | 0.103 | 0.007 | 1.1669539 | DP | chr16-67435109-674 |
| 3.56036947 | 3.93997396243746 | 0.112 | 0.009 | 2.6089176 | DP | chr6-10729088-1072 |
| 2.31004517 | 3.45126395590625 | 0.164 | 0.02 | 1.6927225 | DP | chr6-109944601-109 |
| 3.05231013 | 3.98720333422358 | 0.101 | 0.007 | 2.2366290 | DP | chr12-37212560-372 |
| 1.94047631 | 3.62595852708278 | 0.109 | 0.009 | 1.4219150 | DP | chr19-36026668-360 |
| 3.29461558 | 3.05550530721432 | 0.171 | 0.022 | 2.4141822 | DP | chr1-17879274-1787 |
| 3.15753376 | 4.00558254890878 | 0.108 | 0.009 | 2.3137333 | DP | chr2-220777840-220 |
| 4.82634748 | 3.69488893038104 | 0.112 | 0.01 | 3.5365833 | DP | chr3-60486489-6048 |
| 4.63165677 | 3.26424770921152 | 0.135 | 0.014 | 3.3939206 | DP | chr7-71065858-7106 |
| 1.05393653 | 3.41717411237401 | 0.127 | 0.013 | 7.7228886 | DP | chr20-48004032-480 |
| 1.31689887 | 3.34147405654655 | 0.131 | 0.013 | 9.6497871 | DP | chr4-19291454-1929 |
| 5.72667118 | 3.05937318 | 0.171 | 0.023 | 4.1963099 | DP | chr1-87764548-8776 |
| 2.83892025 | 3.24738114634877 | 0.134 | 0.014 | 2.0802642 | DP | chr3-57371452-5737 |
| 8.25997032 | 3.45068286383671 | 0.14 | 0.016 | 6.0526254 | DP | chr6-74209011-7420 |
| 1.96403470 | 3.37318280207457 | 0.126 | 0.013 | 1.4391778 | DP | chr2-45857474-4585 |
| 4.36510859 | 3.69880557742839 | 0.112 | 0.01 | 3.1986031 | DP | chr12-23919568-239 |
| 1.85093160 | 3.32173872081306 | 0.126 | 0.013 | 1.3562997 | DP | chr17-7198139-7198 |
| 2.75445642 | 2.93676901442354 | 0.184 | 0.027 | 2.0183720 | DP | chr1-15336417-1533 |
| 3.24393736 | 3.5787886349267 | 0.114 | 0.011 | 2.3770470 | DP | chr3-28274228-2827 |
| 5.15043642 | 3.68819420453278 | 0.1 | 0.008 | 3.7740646 | DP | chr4-25839819-2584 |
| 8.68294177 | 3.31852843509235 | 0.135 | 0.015 | 6.3625645 | DP | chr3-124288014-124 |
| 1.40484126 | 3.58102128607788 | 0.12 | 0.012 | 1.0294199 | DP | chr1-120717853-120 |
| 2.06674098 | 3.42805033428792 | 0.114 | 0.011 | 1.5144375 | DP | chr6-45493603-4549 |
| 3.16612815 | 3.26642359010507 | 0.143 | 0.017 | 2.3200310 | DP | chr3-161457413-161 |
| 4.32042819 | 3.29883775214389 | 0.131 | 0.014 | 3.1658628 | DP | chr9-63947518-6394 |
| 1.01071704 | 3.07543829333347 | 0.169 | 0.023 | 7.4061908 | DP | chr8-60758591-6075 |
| 1.02813629 | 3.18230090844977 | 0.143 | 0.017 | 7.5338331 | DP | chr5-8548184-85486 |

| | | | | | | | |
|---|---|---|---|---|---|---|---|
| 2.31783695 | 3.53992135511919 | | 0.109 | 0.01 | 1.6984321 | DP | chr8-117376698-117 |
| 2.58268829 | 3.17788748174645 | | 0.149 | 0.019 | 1.8925061 | DP | chr5-97556940-9755 |
| 5.47565339 | 3.24697745082358 | | 0.135 | 0.016 | 4.0123726 | DP | chr18-7562243-7562 |
| 8.02050403 | 2.96671637059628 | | 0.169 | 0.024 | 5.8771526 | DP | chr1-133255922-133 |
| 6.28829111 | 3.32005999611952 | | 0.121 | 0.013 | 4.6078459 | DP | chr1-128341855-128 |
| 4.43986051 | 3.05199384446075 | | 0.147 | 0.019 | 3.2533788 | DP | chr1-168074628-168 |
| 4.39756046 | 3.09958796693174 | | 0.146 | 0.019 | 3.2223827 | DP | chr3-152802841-152 |
| 1.28955940 | 2.98071228199316 | | 0.141 | 0.018 | 9.4494528 | DP | chr4-179173806-179 |
| 1.59341001 | 3.02325262235017 | | 0.152 | 0.02 | 1.1675966 | DP | chr1-95852485-9585 |
| 8.53846882 | 2.9339246673738 | | 0.155 | 0.021 | 6.2566996 | DP | chr13-97519424-975 |
| 9.87798621 | 3.04539591634176 | | 0.163 | 0.023 | 7.2382524 | DP | chr5-150193818-150 |
| 1.74138212 | 3.27150501512914 | | 0.138 | 0.017 | 1.2760256 | DP | chr3-57716527-5771 |
| 2.75764173 | 3.48910307126523 | | 0.126 | 0.014 | 2.0207061 | DP | chr15-82353507-823 |
| 3.69203537 | 3.20027000059959 | | 0.137 | 0.017 | 2.7053979 | DP | chr11-31462376-314 |
| 4.32578667 | 2.99738343731634 | | 0.163 | 0.023 | 3.1697893 | DP | chr5-122429214-122 |
| 5.51552064 | 3.07436543617917 | | 0.14 | 0.017 | 4.0415859 | DP | chr19-36904181-369 |
| 1.95355482 | 3.29973742898423 | | 0.124 | 0.014 | 1.4314985 | DP | chr15-36375256-363 |
| 2.76934622 | 2.96148613238082 | | 0.154 | 0.021 | 2.0292827 | DP | chr6-123972072-123 |
| 2.59222752 | 2.87310954903625 | | 0.141 | 0.018 | 1.8994961 | DP | chr9-17466814-1746 |
| 1.36695727 | 3.03345536997216 | | 0.158 | 0.023 | 1.0016598 | DP | chr18-59628926-596 |
| 1.50930767 | 3.07271088488757 | | 0.151 | 0.021 | 1.1059693 | DP | chr4-116671414-116 |
| 1.91565741 | 3.36385357211435 | | 0.127 | 0.015 | 1.4037286 | DP | chr4-60943696-6094 |
| 1.40383717 | 3.47433251630487 | | 0.106 | 0.011 | 1.0286841 | DP | chr12-37078269-370 |
| 1.48597094 | 3.61693822274318 | | 0.101 | 0.01 | 1.0888689 | DP | chr12-24307078-243 |
| 5.66262817 | 2.94421599132373 | | 0.141 | 0.019 | 4.1493813 | DP | chr10-96535800-965 |
| 1.99549736 | 2.91904438879612 | | 0.154 | 0.022 | 1.4622326 | DP | chr7-91881411-9188 |
| 9.89052473 | 3.30471584359429 | | 0.106 | 0.011 | 7.2474402 | DP | chr12-29411286-294 |
| 1.24641703 | 3.30581869216478 | | 0.104 | 0.011 | 9.1333202 | DP | chr15-38483756-384 |
| 3.10853076 | | 3.141629575 | 0.123 | 0.015 | 2.2778256 | DP | chr3-57077910-5707 |
| 3.00095182 | 3.23368692415047 | | 0.144 | 0.021 | 2.1989954 | DP | chr20-42054349-420 |
| 5.74232985 | 3.40398043341894 | | 0.1 | 0.01 | 4.2077840 | DP | chr17-75414357-754 |
| 5.86480574 | 2.89512978256431 | | 0.134 | 0.018 | 4.2975302 | DP | chr10-35794083-357 |
| 1.27460384 | 3.15434745922791 | | 0.117 | 0.014 | 9.3398636 | DP | chr15-15330819-153 |
| 1.16943535 | 3.30723802686446 | | 0.112 | 0.013 | 8.5692247 | DP | chr1-124088107-124 |
| 1.52984519 | 3.1732064949181 | | 0.114 | 0.013 | 1.1210185 | DP | chr9-102165461-102 |
| 3.20250795 | 3.16582380759963 | | 0.101 | 0.011 | 2.3466889 | DP | chr1-17757904-1775 |
| 1.79521437 | 3.07377500273729 | | 0.14 | 0.02 | 1.3154720 | DP | chr12-33945521-339 |
| 8.83908183 | 3.25709017399841 | | 0.115 | 0.014 | 6.4769786 | DP | chr2-51096630-5109 |
| 2.21108343 | 3.29561821247789 | | 0.104 | 0.012 | 1.6202067 | DP | chr2-236195226-236 |
| 3.34788439 | 3.29706664594786 | | 0.117 | 0.015 | 2.4532158 | DP | chr4-83012107-8301 |
| 7.24914724 | 3.30180480502306 | | 0.115 | 0.014 | 5.3119286 | DP | chr9-49899870-4990 |
| 7.78717291 | 3.35790131443002 | | 0.111 | 0.013 | 5.7061755 | DP | chr18-76480456-764 |
| 1.87822511 | 2.86356639008357 | | 0.138 | 0.02 | 1.3762995 | DP | chr12-22717048-227 |

| | | | | | | |
|---|---|---|---|---|---|---|
| 2.37145060 | 3.08619650481276 | 0.12 | 0.015 | 1.7377183 | DP | chr1-187820441-187 |
| 6.11207477 | 3.20368411002364 | 0.114 | 0.014 | 4.4787205 | DP | chr20-5447601-5448 |
| 2.96761583 | 3.24827001814504 | 0.111 | 0.013 | 2.1745679 | DP | chr17-86100305-861 |
| 8.75842845 | 3.06433904794227 | 0.114 | 0.014 | 6.4178785 | DP | chr18-56231299-562 |
| 2.08496309 | 2.99725215141044 | 0.141 | 0.021 | 1.5277900 | DP | chr1-241885680-241 |
| 2.21984941 | 3.12977149356968 | 0.118 | 0.015 | 1.6266301 | DP | chr3-79642358-7964 |
| 3.32799550 | 3.34297006858937 | 0.1 | 0.011 | 2.4386419 | DP | chr1-88653991-8865 |
| 1.69357789 | 3.25757613567509 | 0.111 | 0.014 | 1.2409963 | DP | chr8-117543463-117 |
| 3.05996908 | 3.04067874587049 | 0.14 | 0.021 | 2.2422413 | DP | chr2-122561191-122 |
| 5.71258260 | 2.95351295892613 | 0.126 | 0.017 | 4.1859863 | DP | chr15-32751391-327 |
| 5.85766385 | 2.94173154663467 | 0.115 | 0.015 | 4.2922969 | DP | chr16-20965169-209 |
| 1.62871642 | 2.90043044999006 | 0.121 | 0.016 | 1.1934680 | DP | chr4-179172759-179 |
| 2.66413021 | 3.30996421251955 | 0.106 | 0.013 | 1.9521840 | DP | chr4-173053243-173 |
| 3.31035835 | 2.95387288154406 | 0.129 | 0.019 | 2.4257180 | DP | chr19-35957971-359 |
| 2.70622141 | 3.15979533940763 | 0.106 | 0.013 | 1.9830270 | DP | chr4-15125464-1512 |
| 3.95437844 | 3.09062928412907 | 0.106 | 0.013 | 2.8976340 | DP | chr1-29414713-2941 |
| 4.73608075 | 3.07894816685691 | 0.118 | 0.016 | 3.4704389 | DP | chr2-150990697-150 |
| 6.73084430 | 3.20531700481915 | 0.103 | 0.012 | 4.9321338 | DP | chr2-201629119-201 |
| 1.41767344 | 3.07781367038734 | 0.109 | 0.014 | 1.0388229 | DP | chr11-54922265-549 |
| 2.39078177 | 2.97071911061385 | 0.129 | 0.019 | 1.7518836 | DP | chr19-14716252-147 |
| 6.52753044 | 3.08576918733362 | 0.101 | 0.012 | 4.7831523 | DP | chr11-29258001-292 |
| 1.18848995 | 2.97953029553011 | 0.115 | 0.016 | 8.7088502 | DP | chr18-42103112-421 |
| 4.09283154 | 3.14380298738478 | 0.123 | 0.018 | 2.9990877 | DP | chr4-81976711-8197 |
| 1.36631857 | 3.08985968618963 | 0.101 | 0.012 | 1.0011917 | DP | chr5-27949043-2794 |
| 1.15910972 | 2.85621441214355 | 0.138 | 0.022 | 8.4935619 | DP | chr2-237217331-237 |
| 4.57048168 | 2.89691578804613 | 0.124 | 0.018 | 3.3490935 | DP | chr16-4633673-4634 |
| 6.85894809 | 2.86024071455345 | 0.138 | 0.023 | 5.0260039 | DP | chr7-124818508-124 |
| 1.18617982 | 3.00785482974606 | 0.103 | 0.013 | 8.6919224 | DP | chr6-125640135-125 |
| 1.82185788 | 2.87303728040066 | 0.127 | 0.02 | 1.3349955 | DP | chr1-2735311-27358 |
| 2.65671464 | 3.01152125253573 | 0.112 | 0.015 | 1.9467501 | DP | chr19-30624686-306 |
| 3.96246592 | 2.84906230728328 | 0.123 | 0.018 | 2.9035603 | DP | chr18-76638945-766 |
| 4.35751512 | 3.05127635715924 | 0.117 | 0.017 | 3.1930389 | DP | chr14-93709688-937 |
| 2.37342224 | 3.17976791198235 | 0.101 | 0.013 | 1.7391631 | DP | chr17-28665811-286 |
| 4.79182031 | 3.20368411002364 | 0.108 | 0.015 | 3.5112830 | DP | chr3-143463858-143 |
| 7.30684622 | 3.09975838413621 | 0.106 | 0.014 | 5.3542084 | DP | chr1-118412107-118 |
| 2.14795263 | 2.9308852387182 | 0.108 | 0.015 | 1.5739466 | DP | chr1-2970613-29711 |
| 4.40085051 | 3.09945321348242 | 0.103 | 0.014 | 3.2247936 | DP | chr17-1845676-1846 |
| 1.29554473 | 2.98226549626063 | 0.132 | 0.022 | 9.4933113 | DP | chr16-62491034-624 |
| 1.52927246 | 2.93470368622539 | 0.118 | 0.018 | 1.1205988 | DP | chr15-98695443-986 |
| 1.79940425 | 2.97033798074874 | 0.121 | 0.019 | 1.3185422 | DP | chr13-47033086-470 |
| 3.64211121 | 3.02856695376992 | 0.1 | 0.013 | 2.6688152 | DP | chr17-25732308-257 |
| 2.65721273 | 3.21596973111548 | 0.101 | 0.013 | 1.9471151 | DP | chr1-21950931-2195 |
| 2.71370013 | 2.88944412852337 | 0.117 | 0.017 | 1.9885071 | DP | chr5-130901883-130 |

| | | | | | | |
|---|---|---|---|---|---|---|
| 2.00927074 | 2.90382120562791 | 0.115 | 0.017 | 1.4723252 | DP | chr3-166100519-166 |
| 4.30777967 | 2.8773373558567 | 0.101 | 0.014 | 3.1565944 | DP | chr6-6313969-63144 |
| 5.49377022 | 2.90053919166032 | 0.121 | 0.019 | 4.0256480 | DP | chr19-36481538-364 |
| 7.16945925 | 2.88746682987761 | 0.103 | 0.014 | 5.2535359 | DP | chr1-84919340-8491 |
| 9.07655744 | 2.94019131332827 | 0.106 | 0.015 | 6.6509926 | DP | chr1-21788431-2178 |
| 2.38662947 | 2.86997690834473 | 0.117 | 0.018 | 1.7488409 | DP | chr11-11583915-115 |
| 1.31570113 | 2.91686967097204 | 0.114 | 0.017 | 9.6410105 | DP | chr8-86231394-8623 |
| 3.00832637 | 3.02767087554175 | 0.103 | 0.014 | 2.2043992 | DP | chr8-106394163-106 |
| 5.27612707 | 2.92604231798698 | 0.104 | 0.015 | 3.8661665 | DP | chr4-134128782-134 |
| 6.34645432 | 2.96752321815611 | 0.109 | 0.016 | 4.6504659 | DP | chr9-75452715-7545 |
| 3.91262046 | 2.99072509847211 | 0.104 | 0.015 | 2.8670352 | DP | chr3-93735475-9373 |
| 5.09210742 | 2.90320661010941 | 0.108 | 0.016 | 3.7313231 | DP | chr18-56365423-563 |
| 6.73225248 | 2.90631275435741 | 0.111 | 0.017 | 4.9331657 | DP | chr18-69040318-690 |
| 1.12651528 | 2.96183711629967 | 0.1 | 0.014 | 8.2547209 | DP | chr10-96639153-966 |
| 5.15220508 | 2.86629581373909 | 0.117 | 0.019 | 3.7753607 | DP | chr13-29640386-296 |
| 1.58587603 | 2.87899224630697 | 0.103 | 0.015 | 1.1620760 | DP | chr4-68273571-6827 |
| 3.81562575 | 2.91061351975465 | 0.103 | 0.015 | 2.7959608 | DP | chr2-76153642-7615 |
| 1.40085069 | 3.07886023228536 | 0.101 | 0.015 | 1.0264957 | DP | chr5-101054169-101 |
| 9.61956741 | 2.95455971584834 | 0.101 | 0.015 | 7.0488919 | DP | chr10-60817227-608 |
| 1.21986832 | 2.86058148360957 | 0.111 | 0.018 | 8.9387803 | DP | chr7-124943087-124 |
| 2.89495503 | 2.96662040995329 | 0.106 | 0.018 | 2.1213246 | DP | chr2-190087977-190 |
| 4.48109861 | 2.85387928980471 | 0.106 | 0.018 | 3.2835967 | DP | chr2-220216722-220 |
| 1.05985677 | 2.88615009408242 | 0.121 | 0.023 | 7.7662700 | DP | chr14-98663214-986 |
| 0 | 7.44633740193694 | 0.131 | 0.001 | 0 | DS | chr1-106438776-106 |
| 0 | 7.04704270510207 | 0.212 | 0.002 | 0 | DS | chr1-173094764-173 |
| 0 | 6.69852919727739 | 0.263 | 0.003 | 0 | DS | chr14-18869449-188 |
| 0 | 6.60601794528792 | 0.273 | 0.003 | 0 | DS | chr2-77607227-7760 |
| 0 | 6.72450587599666 | 0.192 | 0.002 | 0 | DS | chr5-98497407-9849 |
| 3.90902655 | 6.2167536130757 | 0.222 | 0.004 | 2.8644017 | DS | chr1-125550498-125 |
| 2.75612916 | 5.75170981125101 | 0.253 | 0.005 | 2.0195977 | DS | chr14-32463487-324 |
| 3.00671765 | 6.23292186649655 | 0.162 | 0.002 | 2.2032204 | DS | chr15-100001283-10 |
| 9.39235143 | 6.31330812826492 | 0.172 | 0.002 | 6.8823957 | DS | chr1-227951973-227 |
| 8.19514735 | 5.74473102452499 | 0.212 | 0.004 | 6.0051253 | DS | chr3-90411892-9041 |
| 1.62583413 | 5.99554339346253 | 0.212 | 0.004 | 1.1913559 | DS | chr9-85331823-8533 |
| 2.85976537 | 5.88906486088546 | 0.172 | 0.003 | 2.0955388 | DS | chr2-58049513-5805 |
| 2.19722479 | 6.52313063068785 | 0.121 | 0.001 | 1.6100516 | DS | chr2-233183422-233 |
| 3.78071939 | 5.62493473661404 | 0.212 | 0.005 | 2.7703826 | DS | chr14-82569157-825 |
| 3.19236135 | 6.23292186649655 | 0.152 | 0.002 | 2.3392538 | DS | chr5-43570564-4357 |
| 2.70603198 | 5.92883787789757 | 0.182 | 0.003 | 1.9828882 | DS | chr11-56446421-564 |
| 3.40091927 | 5.28890641448667 | 0.232 | 0.006 | 2.4920780 | DS | chr3-139808975-139 |
| 2.66952919 | 6.31058967494483 | 0.162 | 0.003 | 1.9561402 | DS | chr6-114322613-114 |
| 3.46660339 | 6.06081984077545 | 0.162 | 0.003 | 2.5402091 | DS | chr14-6223779-6224 |
| 1.78115705 | 5.44058362215258 | 0.182 | 0.004 | 1.3051713 | DS | chr15-48074217-480 |

| | | | | | | | |
|---|---|---|---|---|---|---|---|
| 2.34946606 | 5.5428049664267 | | 0.192 | 0.004 | 1.7216088 | DS | chr3-63684300-6368 |
| 9.08163348 | 6.2987861796768 | | 0.121 | 0.002 | 6.6547122 | DS | chr6-57932877-5793 |
| 5.80140536 | 5.58711072244066 | | 0.162 | 0.003 | 4.2510726 | DS | chr10-89735898-897 |
| 1.51104357 | 5.91382255976248 | | 0.141 | 0.002 | 1.1072413 | DS | chr4-95656849-9565 |
| 1.31441520 | 6.52313063068785 | | 0.111 | 0.001 | 9.6315877 | DS | chr5-30970320-3097 |
| 1.16052398 | 5.83367441940418 | | 0.152 | 0.003 | 8.5039251 | DS | chr15-99790908-997 |
| 1.69178653 | 5.07000201794983 | | 0.222 | 0.006 | 1.2396836 | DS | chr14-7615693-7616 |
| 1.27303435 | 5.27741552867499 | | 0.212 | 0.006 | 9.3283628 | DS | chr2-152613382-152 |
| 8.52380241 | 5.56527200525595 | | 0.212 | 0.006 | 6.2459526 | DS | chr11-16695597-166 |
| 4.34579060 | 5.40779752649279 | | 0.232 | 0.007 | 3.1844475 | DS | chr1-45968875-4596 |
| 7.07155748 | 5.9177708590289 | | 0.152 | 0.003 | 5.1817968 | DS | chr2-56086721-5608 |
| 2.00918779 | 5.96892255072659 | | 0.131 | 0.002 | 1.4722645 | DS | chr17-22838390-228 |
| 6.93267961 | 5.86074978303128 | | 0.131 | 0.002 | 5.0800319 | DS | chr1-135573711-135 |
| 2.44462637 | 5.76672482951995 | | 0.172 | 0.004 | 1.7913390 | DS | chr19-17864415-178 |
| 9.41467044 | 5.26191628602561 | | 0.182 | 0.004 | 6.8987504 | DS | chr3-146899471-146 |
| 1.41405236 | 5.28905888527649 | | 0.212 | 0.006 | 1.0361694 | DS | chr2-27183069-2718 |
| 8.70287585 | 5.94935331232381 | | 0.172 | 0.004 | 6.3771715 | DS | chr9-91481084-9148 |
| 4.16086357 | 5.56606042649342 | | 0.172 | 0.004 | 3.0489393 | DS | chr19-18025283-180 |
| 8.90021204 | 5.69722062322499 | | 0.131 | 0.002 | 6.5217727 | DS | chr17-26333994-263 |
| 3.71447409 | 4.94948613771963 | | 0.202 | 0.006 | 2.7218403 | DS | chr10-105029224-10 |
| 1.30500836 | 5.54490318676794 | | 0.141 | 0.003 | 9.5626576 | DS | chr9-77949303-7794 |
| 1.87628637 | 5.65129337177356 | | 0.162 | 0.004 | 1.3748788 | DS | chr17-30371205-303 |
| 3.12007563 | 5.47142145041235 | | 0.172 | 0.004 | 2.2862853 | DS | chr2-226430932-226 |
| 1.08167114 | 5.46822907106887 | | 0.172 | 0.004 | 7.9261183 | DS | chr18-28056450-280 |
| 1.71905654 | 5.41521831512975 | | 0.172 | 0.004 | 1.2596661 | DS | chr16-50391969-503 |
| 3.66779208 | 5.24417473771184 | | 0.162 | 0.004 | 2.6876333 | DS | chr4-41185391-4118 |
| 2.77271777 | 5.77062066700286 | | 0.131 | 0.003 | 2.0317533 | DS | chr1-51248233-5124 |
| 8.46375847 | 5.28074772848889 | | 0.162 | 0.004 | 6.2019544 | DS | chr1-252733857-252 |
| 2.82353606 | | 5.460233624 | 0.162 | 0.004 | 2.0689912 | DS | chr18-53930094-539 |
| 3.74112593 | 5.37139900403606 | | 0.152 | 0.004 | 2.7413698 | DS | chr2-20093422-2009 |
| 3.04490447 | 5.0393299253146 | | 0.202 | 0.007 | 2.2312024 | DS | chr1-121456836-121 |
| 2.39937927 | 5.54599292806367 | | 0.131 | 0.003 | 1.7581835 | DS | chr4-150041019-150 |
| 9.31709249 | 5.46333408162531 | | 0.152 | 0.004 | 6.8272486 | DS | chr6-95601549-9560 |
| 2.40508239 | 6.03421234762734 | | 0.121 | 0.002 | 1.7623626 | DS | chr7-39115093-3911 |
| 2.82475731 | 5.54366923661247 | | 0.131 | 0.003 | 2.0698861 | DS | chr7-68673438-6867 |
| 8.78631846 | 5.60417069197831 | | 0.121 | 0.002 | 6.4383154 | DS | chr17-22855020-228 |
| 1.08075133 | 6.29007050755996 | | 0.111 | 0.002 | 7.9193783 | DS | chr6-32730141-3273 |
| 3.20557713 | 5.29783299846873 | | 0.131 | 0.003 | 2.3489379 | DS | chr3-132541822-132 |
| 1.16094421 | 5.50854993948195 | | 0.121 | 0.002 | 8.5070044 | DS | chr5-22334706-2233 |
| 3.12705573 | 5.6105175456932 | | 0.111 | 0.002 | 2.2914001 | DS | chr11-3426412-3426 |
| 2.02950460 | 5.2048746700608 | | 0.141 | 0.004 | 1.4871519 | DS | chr17-30665502-306 |
| 9.14385506 | 5.34536989894305 | | 0.162 | 0.005 | 6.7003060 | DS | chr10-35884931-358 |
| 9.78077342 | 5.2435860461366 | | 0.162 | 0.005 | 7.1670182 | DS | chr4-120191546-120 |

| | | | | | | |
|---|---|---|---|---|---|---|
| 4.53619348 | 5.68902121508737 | 0.131 | 0.003 | 3.3239683 | DS | chr13-78966705-789 |
| 2.52803795 | 5.52404483521047 | 0.141 | 0.004 | 1.8524602 | DS | chr14-82643668-826 |
| 3.42494334 | 5.09225095105626 | 0.152 | 0.004 | 2.5096820 | DS | chr2-100940005-100 |
| 3.76118524 | 5.33856159985367 | 0.141 | 0.004 | 2.7560686 | DS | chr10-12811844-128 |
| 9.35415029 | 6.18025166598743 | 0.101 | 0.002 | 6.8544032 | DS | chr11-27667888-276 |
| 2.63717554 | 5.26021077518506 | 0.192 | 0.007 | 1.9324325 | DS | chr2-46763860-4676 |
| 8.87338273 | 5.13454299841123 | 0.152 | 0.004 | 6.5021131 | DS | chr19-14010246-140 |
| 9.88385062 | 5.41167203878368 | 0.111 | 0.002 | 7.2425496 | DS | chr7-81348079-8134 |
| 1.48559741 | 5.19099621323492 | 0.121 | 0.003 | 1.0885952 | DS | chr15-93576800-935 |
| 7.38930383 | 5.88906486088546 | 0.121 | 0.003 | 5.4146306 | DS | chr16-82293107-822 |
| 1.01635038 | 5.06123630990698 | 0.172 | 0.006 | 7.4474700 | DS | chr13-64295977-642 |
| 2.21776955 | 5.29853877641656 | 0.141 | 0.004 | 1.6251061 | DS | chr2-116388523-116 |
| 2.38293100 | 5.13371956935702 | 0.172 | 0.006 | 1.7461308 | DS | chr8-51605151-5160 |
| 1.50488208 | 5.18294010796649 | 0.131 | 0.003 | 1.1027264 | DS | chr2-157048705-157 |
| 1.18476026 | 5.62910003595945 | 0.111 | 0.002 | 8.6815203 | DS | chr3-35835693-3583 |
| 8.63944701 | 5.01539526281243 | 0.192 | 0.007 | 6.3306930 | DS | chr15-38442030-384 |
| 4.60439252 | 5.09922006648913 | 0.141 | 0.004 | 3.3739422 | DS | chr5-134188412-134 |
| 8.93081137 | 5.25094801106839 | 0.152 | 0.005 | 6.5441949 | DS | chr11-35667330-356 |
| 1.19761408 | 5.52963232871494 | 0.111 | 0.002 | 8.7757088 | DS | chr15-99839873-998 |
| 1.31021563 | 5.30015659025425 | 0.141 | 0.004 | 9.6008147 | DS | chr3-96615944-9661 |
| 1.36216486 | 5.37336032022867 | 0.111 | 0.002 | 9.9814809 | DS | chr17-30603490-306 |
| 5.11753396 | 5.47049005979306 | 0.131 | 0.003 | 3.7499548 | DS | chr10-49878709-498 |
| 6.60177339 | 5.1743138760113 | 0.131 | 0.003 | 4.8375550 | DS | chr18-74558485-745 |
| 1.31669614 | 5.12051882096673 | 0.152 | 0.005 | 9.6483016 | DS | chr16-81352071-813 |
| 8.43465883 | 5.33355936432046 | 0.141 | 0.004 | 6.1806312 | DS | chr3-61950626-6195 |
| 2.95787271 | 5.16258897985645 | 0.162 | 0.005 | 2.1674285 | DS | chr7-43689016-4368 |
| 5.06111315 | 4.96642525388321 | 0.141 | 0.004 | 3.7086116 | DS | chr9-74518107-7451 |
| 5.60971084 | 5.42066416124476 | 0.131 | 0.003 | 4.1106053 | DS | chr19-54317519-543 |
| 9.26056267 | 5.5918657203052 | 0.111 | 0.002 | 6.7858254 | DS | chr9-20535639-2053 |
| 2.62647776 | 4.97616205512628 | 0.202 | 0.008 | 1.9245936 | DS | chr2-195346024-195 |
| 7.37283752 | 5.39863749974118 | 0.101 | 0.002 | 5.4025646 | DS | chr9-79210598-7921 |
| 7.96715190 | 5.09482198964731 | 0.162 | 0.006 | 5.8380580 | DS | chr2-111787109-111 |
| 3.11882136 | 5.3073254327503 | 0.141 | 0.004 | 2.2853662 | DS | chr2-54089410-5408 |
| 7.55927123 | 5.59626797070407 | 0.101 | 0.002 | 5.5391769 | DS | chr1-150662225-150 |
| 7.24520414 | 5.60711413359709 | 0.101 | 0.002 | 5.3090392 | DS | chr5-85855987-8585 |
| 7.40212660 | 5.43281491018667 | 0.121 | 0.003 | 5.4240267 | DS | chr4-17606127-1760 |
| 7.29650974 | 5.30448595860421 | 0.182 | 0.007 | 5.3466342 | DS | chr14-8056642-8057 |
| 4.38359414 | 5.13035845351449 | 0.152 | 0.005 | 3.2121487 | DS | chr5-50588463-5058 |
| 7.32709468 | 5.32737571850373 | 0.101 | 0.002 | 5.3690458 | DS | chr14-32445923-324 |
| 8.93394007 | 4.97322127753219 | 0.131 | 0.004 | 6.5464875 | DS | chr7-47059573-4706 |
| 2.24370242 | 4.99291982038059 | 0.141 | 0.004 | 1.6441088 | DS | chr20-15066389-150 |
| 4.11406191 | 5.19005023295028 | 0.141 | 0.005 | 3.0146446 | DS | chr17-28261202-282 |
| 5.96441031 | 5.44420152936796 | 0.101 | 0.002 | 4.3705170 | DS | chr12-43869998-438 |

| | | | | | | |
|---|---|---|---|---|---|---|
| 1.15588736 | 5.8866872732695 | 0.111 | 0.003 | 8.4699495 | DS | chr4-103619484-103 |
| 5.26144172 | 5.14132551498977 | 0.101 | 0.002 | 3.8554056 | DS | chr14-64795933-647 |
| 1.95254412 | 4.9300508915078 | 0.152 | 0.005 | 1.4307579 | DS | chr12-19196994-191 |
| 2.87167223 | 5.61971389419113 | 0.131 | 0.004 | 2.1042637 | DS | chr18-67499337-674 |
| 6.41464409 | 5.00835361971494 | 0.162 | 0.006 | 4.7004330 | DS | chr8-67767000-6776 |
| 1.77546652 | 4.96014253766178 | 0.131 | 0.004 | 1.3010015 | DS | chr19-1606262-1606 |
| 5.85627203 | 4.89588909125413 | 0.131 | 0.004 | 4.2912770 | DS | chr11-42573666-425 |
| 1.11103712 | 5.83632476492379 | 0.111 | 0.003 | 8.1413023 | DS | chr1-195873100-195 |
| 2.38257001 | 5.17911392375637 | 0.121 | 0.004 | 1.7458662 | DS | chr2-134796837-134 |
| 5.37325181 | 5.36354803456737 | 0.101 | 0.002 | 3.9373362 | DS | chr6-8523896-85243 |
| 6.75132264 | 5.16187109308411 | 0.152 | 0.006 | 4.9471396 | DS | chr6-51300583-5130 |
| 1.27370184 | 5.09891450050342 | 0.121 | 0.004 | 9.3332540 | DS | chr9-13609508-1361 |
| 5.83855616 | 5.43239985236704 | 0.101 | 0.003 | 4.2782954 | DS | chr1-28535798-2853 |
| 4.41692514 | 5.28515854507014 | 0.121 | 0.004 | 3.2365725 | DS | chr9-17792873-1779 |
| 3.62068469 | 5.28201990605026 | 0.131 | 0.004 | 2.6531146 | DS | chr8-51753560-5175 |
| 3.71616961 | 5.46301574181188 | 0.101 | 0.003 | 2.7230827 | DS | chr10-37140750-371 |
| 7.99153293 | 5.11608272892927 | 0.111 | 0.003 | 5.8559236 | DS | chr14-4366850-4367 |
| 3.21123727 | 5.2179702616323 | 0.111 | 0.003 | 2.3530854 | DS | chr8-85189727-8519 |
| 1.25108544 | 4.90299149313144 | 0.131 | 0.005 | 9.1675287 | DS | chr12-24123198-241 |
| 2.85553996 | 5.10676908092452 | 0.121 | 0.004 | 2.0924426 | DS | chr2-154897469-154 |
| 4.69489512 | 5.18294010796649 | 0.111 | 0.003 | 3.4402595 | DS | chr14-99477939-994 |
| 4.92117106 | 5.17911392375637 | 0.111 | 0.003 | 3.6060668 | DS | chr7-63756256-6375 |
| 9.10953865 | 4.92893145665784 | 0.101 | 0.003 | 6.6751602 | DS | chr17-83552606-835 |
| 2.16495842 | 4.96777110836198 | 0.121 | 0.004 | 1.5864079 | DS | chr19-10191919-101 |
| 3.42211920 | 5.10752603928926 | 0.101 | 0.003 | 2.5076126 | DS | chr17-10969364-109 |
| 6.35012629 | 5.11191265474883 | 0.131 | 0.005 | 4.6531566 | DS | chr10-85524740-855 |
| 9.89376894 | 5.58473869684225 | 0.111 | 0.004 | 7.2498174 | DS | chr8-63130434-6313 |
| 2.19058169 | 4.94273347643435 | 0.101 | 0.003 | 1.6051837 | DS | chr4-78611190-7861 |
| 3.68938862 | 5.03872076434681 | 0.101 | 0.003 | 2.7034585 | DS | chr2-34308359-3430 |
| 1.13771172 | 4.93942219804521 | 0.111 | 0.004 | 8.3367646 | DS | chr18-24125553-241 |
| 1.17500636 | 5.19455166866242 | 0.101 | 0.003 | 8.6100471 | DS | chr1-178017748-178 |
| 6.39085890 | 5.18163557766488 | 0.111 | 0.004 | 4.6830041 | DS | chr4-178709412-178 |
| 8.17816761 | 5.05866836335433 | 0.111 | 0.004 | 5.9926831 | DS | chr19-41953689-419 |
| 8.55597966 | 4.91109585650339 | 0.111 | 0.004 | 6.2695309 | DS | chr1-203932862-203 |
| 1.82004047 | 5.10619324108853 | 0.131 | 0.005 | 1.3336637 | DS | chr7-85380977-8538 |
| 4.72186955 | 5.09817277349668 | 0.101 | 0.003 | 3.4600254 | DS | chrX-72807602-7280 |
| 6.71710200 | 4.96774482080142 | 0.152 | 0.007 | 4.9220639 | DS | chr17-73764255-737 |
| 3.49344762 | 4.97280943127804 | 0.101 | 0.003 | 2.5598796 | DS | chr16-63003424-630 |
| 8.92223936 | 5.06677990724737 | 0.131 | 0.006 | 6.5379136 | DS | chr18-62396900-623 |
| 5.74017895 | 5.24462498501911 | 0.101 | 0.004 | 4.2062079 | DS | chr1-155991347-155 |
| 1.45917418 | 5.03912144910117 | 0.111 | 0.004 | 1.0692332 | DS | chr1-189871231-189 |
| 1.07437366 | 5.06364577096969 | 0.101 | 0.004 | 7.8726449 | DS | chr10-5063056-5063 |
| 5.77499350 | 5.10676908092452 | 0.101 | 0.004 | 4.2317188 | DS | chr16-80999354-809 |

| | | | | | | | |
|---|---|---|---|---|---|---|---|
| 3.97443857( | 4.90299149313144 | | 0.101 | 0.005 | 2.9123334 | DS | chr13-103912330-10[illegible] |
| 0 | 6.54691052510458 | | 0.281 | 0.004 | 0 | ENDO | chr1-4343514-43440[illegible] |
| 0 | 6.74831185785323 | | 0.238 | 0.002 | 0 | ENDO | chr1-13196232-1319[illegible] |
| 0 | 6.66252501781112 | | 0.38 | 0.005 | 0 | ENDO | chr1-40237865-4023[illegible] |
| 0 | 5.98695579614925 | | 0.307 | 0.006 | 0 | ENDO | chr1-40307952-4030[illegible] |
| 0 | 6.1655796101813 | | 0.439 | 0.009 | 0 | ENDO | chr1-42016260-4201[illegible] |
| 0 | 6.02226735956464 | | 0.189 | 0.003 | 0 | ENDO | chr1-42017103-4201[illegible] |
| 0 | 6.25540588387924 | | 0.273 | 0.004 | 0 | ENDO | chr1-42017798-4201[illegible] |
| 0 | 5.9344815234609 | | 0.149 | 0.003 | 0 | ENDO | chr1-56090911-5609[illegible] |
| 0 | 5.98963745929049 | | 0.306 | 0.006 | 0 | ENDO | chr1-86399582-8640[illegible] |
| 0 | 6.54230191944047 | | 0.412 | 0.006 | 0 | ENDO | chr1-86882554-8688[illegible] |
| 0 | 6.21752206836636 | | 0.451 | 0.009 | 0 | ENDO | chr1-118979676-118[illegible] |
| 0 | 5.97672733544674 | | 0.207 | 0.004 | 0 | ENDO | chr1-136683648-136[illegible] |
| 0 | 6.10623317907898 | | 0.258 | 0.005 | 0 | ENDO | chr1-147030272-147[illegible] |
| 0 | 6.29793129255004 | | 0.183 | 0.003 | 0 | ENDO | chr1-176753010-176[illegible] |
| 0 | 6.09405418494529 | | 0.126 | 0.002 | 0 | ENDO | chr1-221834445-221[illegible] |
| 0 | 6.3361255016269 | | 0.176 | 0.003 | 0 | ENDO | chr1-224932765-224[illegible] |
| 0 | 6.39978983387505 | | 0.206 | 0.003 | 0 | ENDO | chr1-239437281-239[illegible] |
| 0 | 6.0867510040383 | | 0.194 | 0.003 | 0 | ENDO | chr10-16815130-168[illegible] |
| 0 | 5.97011023019496 | | 0.264 | 0.005 | 0 | ENDO | chr10-35902302-359[illegible] |
| 0 | 6.05994667285427 | | 0.193 | 0.003 | 0 | ENDO | chr10-51620342-516[illegible] |
| 0 | 6.55101404612511 | | 0.33 | 0.004 | 0 | ENDO | chr10-54940464-549[illegible] |
| 0 | 5.94724648005095 | | 0.308 | 0.007 | 0 | ENDO | chr10-64018063-640[illegible] |
| 0 | 6.0750693351194 | | 0.341 | 0.007 | 0 | ENDO | chr10-88321243-883[illegible] |
| 0 | 6.27871641970784 | | 0.223 | 0.004 | 0 | ENDO | chr10-95522828-955[illegible] |
| 0 | 6.0134978422022 | | 0.49 | 0.011 | 0 | ENDO | chr11-30171391-301[illegible] |
| 0 | 6.00415235815272 | | 0.277 | 0.005 | 0 | ENDO | chr11-34761487-347[illegible] |
| 0 | 6.26691439300572 | | 0.339 | 0.006 | 0 | ENDO | chr11-69024565-690[illegible] |
| 0 | 6.13811231383508 | | 0.29 | 0.005 | 0 | ENDO | chr11-77702359-777[illegible] |
| 0 | 5.93580946836003 | | 0.247 | 0.005 | 0 | ENDO | chr11-82539908-825[illegible] |
| 0 | 6.26320058954537 | | 0.308 | 0.006 | 0 | ENDO | chr12-7110938-7111[illegible] |
| 0 | 5.94093152879601 | | 0.191 | 0.004 | 0 | ENDO | chr12-7368485-7368[illegible] |
| 0 | 5.96631621503368 | | 0.204 | 0.004 | 0 | ENDO | chr12-13780803-137[illegible] |
| 0 | 6.21395207650726 | | 0.371 | 0.007 | 0 | ENDO | chr12-19579809-195[illegible] |
| 0 | | 6.169646769 | 0.248 | 0.004 | 0 | ENDO | chr13-21248910-212[illegible] |
| 0 | 6.74447495230536 | | 0.332 | 0.004 | 0 | ENDO | chr13-48302419-483[illegible] |
| 0 | 6.18922546390526 | | 0.178 | 0.003 | 0 | ENDO | chr13-48330629-483[illegible] |
| 0 | 6.12844101619188 | | 0.19 | 0.003 | 0 | ENDO | chr13-85529708-855[illegible] |
| 0 | 6.36531493566767 | | 0.469 | 0.008 | 0 | ENDO | chr13-96415344-964[illegible] |
| 0 | 6.19617726684409 | | 0.295 | 0.005 | 0 | ENDO | chr13-103076766-10[illegible] |
| 0 | 5.94778095485644 | | 0.154 | 0.003 | 0 | ENDO | chr13-106467623-10[illegible] |
| 0 | 6.67275317486424 | | 0.26 | 0.004 | 0 | ENDO | chr14-82411357-824[illegible] |
| 0 | 5.93798095274098 | | 0.212 | 0.004 | 0 | ENDO | chr14-94705611-947[illegible] |

| | | | | | | |
|---|---|---|---|---|---|---|
| 0 | 5.9661250517671 | 0.489 | 0.011 | 0 | ENDO | chr15-4470183-4470 |
| 0 | 6.41567561153664 | 0.413 | 0.007 | 0 | ENDO | chr15-12982176-129 |
| 0 | 6.08846348743496 | 0.251 | 0.004 | 0 | ENDO | chr15-38194565-381 |
| 0 | 6.02465746574706 | 0.268 | 0.005 | 0 | ENDO | chr15-97666018-976 |
| 0 | 6.07720736823738 | 0.163 | 0.002 | 0 | ENDO | chr16-6156287-6156 |
| 0 | 5.97772909028116 | 0.129 | 0.002 | 0 | ENDO | chr16-21080405-210 |
| 0 | 6.11818619706762 | 0.109 | 0.002 | 0 | ENDO | chr16-25461181-254 |
| 0 | 6.00411381489473 | 0.158 | 0.003 | 0 | ENDO | chr16-51380562-513 |
| 0 | 5.97609257635757 | 0.128 | 0.002 | 0 | ENDO | chr16-80616148-806 |
| 0 | 6.04153551067184 | 0.144 | 0.002 | 0 | ENDO | chr16-80624898-806 |
| 0 | 6.02000187451815 | 0.184 | 0.003 | 0 | ENDO | chr16-80645526-806 |
| 0 | 6.20911280440011 | 0.242 | 0.004 | 0 | ENDO | chr17-17813553-178 |
| 0 | 6.3810728677323 | 0.202 | 0.003 | 0 | ENDO | chr17-57444348-574 |
| 0 | 6.3868557408781 | 0.207 | 0.003 | 0 | ENDO | chr17-67850332-678 |
| 0 | 6.15648258636274 | 0.225 | 0.004 | 0 | ENDO | chr17-80154864-801 |
| 0 | 6.00569762388293 | 0.179 | 0.003 | 0 | ENDO | chr17-83521263-835 |
| 0 | 6.0215398751617 | 0.134 | 0.002 | 0 | ENDO | chr18-12228943-122 |
| 0 | 6.05490980953884 | 0.214 | 0.004 | 0 | ENDO | chr18-25252502-252 |
| 0 | 6.75836466491396 | 0.263 | 0.003 | 0 | ENDO | chr19-9478962-9479 |
| 0 | 5.9550239124699 | 0.118 | 0.002 | 0 | ENDO | chr19-31145948-311 |
| 0 | 6.51995542462833 | 0.244 | 0.003 | 0 | ENDO | chr19-46425873-464 |
| 0 | 6.50377225478958 | 0.232 | 0.003 | 0 | ENDO | chr2-23338016-2333 |
| 0 | 6.00770141709709 | 0.124 | 0.002 | 0 | ENDO | chr2-23339424-2333 |
| 0 | 6.14154445384955 | 0.245 | 0.005 | 0 | ENDO | chr2-29475449-2947 |
| 0 | 6.28877733264654 | 0.189 | 0.003 | 0 | ENDO | chr2-40838577-4083 |
| 0 | 6.04729989959073 | 0.141 | 0.003 | 0 | ENDO | chr2-42292692-4229 |
| 0 | 6.23452641416756 | 0.156 | 0.002 | 0 | ENDO | chr2-93119129-9311 |
| 0 | 6.21367466323726 | 0.207 | 0.003 | 0 | ENDO | chr2-93119682-9312 |
| 0 | 5.95996349887204 | 0.121 | 0.002 | 0 | ENDO | chr2-93156177-9315 |
| 0 | 6.03049803230039 | 0.196 | 0.004 | 0 | ENDO | chr2-112953333-112 |
| 0 | 6.29114311379414 | 0.305 | 0.005 | 0 | ENDO | chr2-112961406-112 |
| 0 | 6.08380320253792 | 0.336 | 0.006 | 0 | ENDO | chr2-112965170-112 |
| 0 | 6.24072813669395 | 0.224 | 0.003 | 0 | ENDO | chr2-113223890-113 |
| 0 | 6.65826619548668 | 0.431 | 0.006 | 0 | ENDO | chr2-135166527-135 |
| 0 | 6.87176865114869 | 0.151 | 0.002 | 0 | ENDO | chr2-148523556-148 |
| 0 | 6.39593425916532 | 0.412 | 0.007 | 0 | ENDO | chr2-203623304-203 |
| 0 | 6.49714400028726 | 0.23 | 0.003 | 0 | ENDO | chr2-203681688-203 |
| 0 | 6.52096321669763 | 0.221 | 0.003 | 0 | ENDO | chr2-204903970-204 |
| 0 | 6.05846262813657 | 0.174 | 0.003 | 0 | ENDO | chr2-210221122-210 |
| 0 | 6.83699713992268 | 0.319 | 0.003 | 0 | ENDO | chr2-225875338-225 |
| 0 | 6.13150185836159 | 0.198 | 0.003 | 0 | ENDO | chr2-234764337-234 |
| 0 | 5.93133631535755 | 0.272 | 0.005 | 0 | ENDO | chr2-238080022-238 |
| 0 | 6.25522171690406 | 0.264 | 0.004 | 0 | ENDO | chr2-240339415-240 |

| | | | | | |
|---|---|---|---|---|---|
| 0 | 6.45180385359068 | 0.184 | 0.002 | 0 ENDO | chr2-240384671-240[illegible] |
| 0 | 5.98860717346048 | 0.22 | 0.004 | 0 ENDO | chr20-12358439-123[illegible] |
| 0 | 6.21877677395457 | 0.156 | 0.002 | 0 ENDO | chr20-26403103-264[illegible] |
| 0 | 5.94999903825639 | 0.28 | 0.006 | 0 ENDO | chr20-30162125-301[illegible] |
| 0 | 6.30836867524138 | 0.222 | 0.003 | 0 ENDO | chr20-36052086-360[illegible] |
| 0 | 6.76764473618039 | 0.328 | 0.004 | 0 ENDO | chr20-48026334-480[illegible] |
| 0 | 5.95361190027058 | 0.143 | 0.003 | 0 ENDO | chr3-26778906-2677[illegible] |
| 0 | 6.08248301744346 | 0.278 | 0.005 | 0 ENDO | chr3-28667129-2866[illegible] |
| 0 | 6.46181923182399 | 0.332 | 0.005 | 0 ENDO | chr3-58683791-5868[illegible] |
| 0 | 6.05097408449452 | 0.156 | 0.003 | 0 ENDO | chr3-69060314-6906[illegible] |
| 0 | 6.02378096021006 | 0.183 | 0.003 | 0 ENDO | chr3-69283954-6928[illegible] |
| 0 | 6.57622709564045 | 0.377 | 0.005 | 0 ENDO | chr3-69628306-6962[illegible] |
| 0 | 6.40939946784017 | 0.115 | 0.001 | 0 ENDO | chr3-94590099-9459[illegible] |
| 0 | 6.14878008262513 | 0.226 | 0.004 | 0 ENDO | chr3-101581638-101[illegible] |
| 0 | 5.95574979027067 | 0.103 | 0.002 | 0 ENDO | chr3-147709030-147[illegible] |
| 0 | 5.9377622004686 | 0.314 | 0.007 | 0 ENDO | chr3-153241884-153[illegible] |
| 0 | 6.49998104495263 | 0.169 | 0.002 | 0 ENDO | chr4-17190943-1719[illegible] |
| 0 | 6.32615804438154 | 0.152 | 0.002 | 0 ENDO | chr4-60239556-6024[illegible] |
| 0 | 6.36119411507989 | 0.405 | 0.006 | 0 ENDO | chr4-60240216-6024[illegible] |
| 0 | 6.97316976528126 | 0.324 | 0.004 | 0 ENDO | chr4-60241201-6024[illegible] |
| 0 | 6.98886444267893 | 0.316 | 0.003 | 0 ENDO | chr4-65569978-6557[illegible] |
| 0 | 6.54323686033629 | 0.253 | 0.003 | 0 ENDO | chr4-114632584-114[illegible] |
| 0 | 5.93329168565238 | 0.209 | 0.004 | 0 ENDO | chr4-116536665-116[illegible] |
| 0 | 6.15738089576973 | 0.343 | 0.006 | 0 ENDO | chr4-124490436-124[illegible] |
| 0 | 6.02133821724893 | 0.192 | 0.004 | 0 ENDO | chr4-126716846-126[illegible] |
| 0 | 5.94359052034056 | 0.283 | 0.006 | 0 ENDO | chr4-126727671-126[illegible] |
| 0 | 6.35610145142706 | 0.118 | 0.002 | 0 ENDO | chr4-150949472-150[illegible] |
| 0 | 6.14445803168622 | 0.333 | 0.006 | 0 ENDO | chr4-158331378-158[illegible] |
| 0 | 6.4408993053987 | 0.281 | 0.004 | 0 ENDO | chr4-162876445-162[illegible] |
| 0 | 5.93811397709284 | 0.308 | 0.007 | 0 ENDO | chr4-179864074-179[illegible] |
| 0 | 6.46418530372833 | 0.214 | 0.003 | 0 ENDO | chr5-14565623-1456[illegible] |
| 0 | 5.9362471536961 | 0.108 | 0.002 | 0 ENDO | chr5-14690623-1469[illegible] |
| 0 | 6.7079154102589 | 0.116 | 0.001 | 0 ENDO | chr5-14739018-1473[illegible] |
| 0 | 6.27672324597967 | 0.175 | 0.003 | 0 ENDO | chr5-20471912-2047[illegible] |
| 0 | 6.10306438095765 | 0.136 | 0.002 | 0 ENDO | chr5-29448085-2944[illegible] |
| 0 | 6.02706013904972 | 0.134 | 0.002 | 0 ENDO | chr6-19987848-1998[illegible] |
| 0 | 6.22420746833202 | 0.474 | 0.009 | 0 ENDO | chr6-21680323-2168[illegible] |
| 0 | 6.10374398912065 | 0.223 | 0.004 | 0 ENDO | chr6-45464333-4546[illegible] |
| 0 | 5.98133331008804 | 0.256 | 0.005 | 0 ENDO | chr6-46593026-4659[illegible] |
| 0 | 6.26210944804609 | 0.199 | 0.003 | 0 ENDO | chr6-88522010-8852[illegible] |
| 0 | 6.3239393147366 | 0.231 | 0.003 | 0 ENDO | chr6-88538233-8853[illegible] |
| 0 | 6.25856916648699 | 0.293 | 0.005 | 0 ENDO | chr6-92282371-9228[illegible] |
| 0 | 6.37957497937136 | 0.21 | 0.003 | 0 ENDO | chr6-94080936-9408[illegible] |

| | | | | | | |
|---|---|---|---|---|---|---|
| 0 | 6.16103620637279 | 0.21 | 0.004 | 0 | ENDO | chr6-123924984-123 |
| 0 | 6.05716239711917 | 0.151 | 0.003 | 0 | ENDO | chr6-124983429-124 |
| 0 | 6.24912987379939 | 0.224 | 0.004 | 0 | ENDO | chr7-40986305-4098 |
| 0 | 6.39887947495947 | 0.156 | 0.002 | 0 | ENDO | chr7-51941031-5194 |
| 0 | 6.30640661130102 | 0.4 | 0.007 | 0 | ENDO | chr7-67661225-6766 |
| 0 | 6.10293883113257 | 0.364 | 0.007 | 0 | ENDO | chr7-120568428-120 |
| 0 | 6.05375979199617 | 0.292 | 0.005 | 0 | ENDO | chr8-20371420-2037 |
| 0 | 6.1851246209017 | 0.283 | 0.005 | 0 | ENDO | chr8-31028044-3102 |
| 0 | 6.20340868373767 | 0.484 | 0.01 | 0 | ENDO | chr8-31296210-3129 |
| 0 | 6.12093785290275 | 0.268 | 0.004 | 0 | ENDO | chr8-37119654-3712 |
| 0 | 6.29686838652993 | 0.286 | 0.004 | 0 | ENDO | chr8-42956097-4295 |
| 0 | 6.15765761975861 | 0.139 | 0.002 | 0 | ENDO | chr8-48526015-4852 |
| 0 | 6.00031325320298 | 0.162 | 0.003 | 0 | ENDO | chr8-61316563-6131 |
| 0 | 6.167717319 | 0.339 | 0.006 | 0 | ENDO | chr8-121727346-121 |
| 0 | 6.14216644342116 | 0.18 | 0.003 | 0 | ENDO | chr9-17618384-1761 |
| 0 | 5.94751172073405 | 0.314 | 0.006 | 0 | ENDO | chr9-49824019-4982 |
| 0 | 6.14101988621528 | 0.206 | 0.004 | 0 | ENDO | chr9-49832356-4983 |
| 0 | 6.0801371230911 | 0.148 | 0.003 | 0 | ENDO | chr9-62696719-6269 |
| 0 | 5.918927673 | 0.236 | 0.005 | 0 | ENDO | chr9-98020338-9802 |
| 0 | 7.07065831431474 | 0.225 | 0.002 | 0 | ENDO | chr9-107192047-107 |
| 0 | 6.18408185324543 | 0.249 | 0.004 | 0 | ENDO | chr9-107510569-107 |
| 0 | 6.11208084928058 | 0.176 | 0.003 | 0 | ENDO | chr9-107512314-107 |
| 0 | 4.32910763928749 | 0.315 | 0.017 | 0 | HFSC | chr1-53989805-5399 |
| 0 | 4.54071066348097 | 0.161 | 0.007 | 0 | HFSC | chr1-74207371-7420 |
| 0 | 4.45429405119771 | 0.261 | 0.013 | 0 | HFSC | chr1-94162942-9416 |
| 0 | 4.18177990571334 | 0.186 | 0.011 | 0 | HFSC | chr1-150333919-150 |
| 0 | 5.3408448870152 | 0.12 | 0.003 | 0 | HFSC | chr1-150343973-150 |
| 0 | 4.59038240415361 | 0.165 | 0.007 | 0 | HFSC | chr1-166267608-166 |
| 0 | 4.9058519917465 | 0.128 | 0.005 | 0 | HFSC | chr1-212656294-212 |
| 0 | 4.94968494795594 | 0.162 | 0.006 | 0 | HFSC | chr10-27785254-277 |
| 0 | 5.31451193700698 | 0.104 | 0.003 | 0 | HFSC | chr10-84979708-849 |
| 0 | 5.17532568850307 | 0.118 | 0.004 | 0 | HFSC | chr10-97015463-970 |
| 0 | 4.39952540414015 | 0.285 | 0.017 | 0 | HFSC | chr11-24859663-248 |
| 0 | 4.33721260099684 | 0.197 | 0.01 | 0 | HFSC | chr11-26916112-269 |
| 0 | 4.65483006624103 | 0.205 | 0.009 | 0 | HFSC | chr11-55169658-551 |
| 0 | 4.11059305103013 | 0.272 | 0.016 | 0 | HFSC | chr11-55178023-551 |
| 0 | 4.73425117734678 | 0.193 | 0.008 | 0 | HFSC | chr12-10510230-105 |
| 0 | 4.47755382928425 | 0.238 | 0.013 | 0 | HFSC | chr13-30567833-305 |
| 0 | 4.18398067468317 | 0.175 | 0.01 | 0 | HFSC | chr13-63049663-630 |
| 0 | 5.74390370023646 | 0.193 | 0.004 | 0 | HFSC | chr14-18138570-181 |
| 0 | 4.44983911444209 | 0.145 | 0.007 | 0 | HFSC | chr14-60865822-608 |
| 0 | 4.96315511171889 | 0.169 | 0.006 | 0 | HFSC | chr14-92660602-926 |
| 0 | 4.1233851287922 | 0.193 | 0.012 | 0 | HFSC | chr15-21589943-215 |

| | | | | | | |
|---|---|---|---|---|---|---|
| 0 | 5.20002062856331 | 0.147 | 0.004 | 0 | HFSC | chr15-33849000-338 |
| 0 | 4.14329155661416 | 0.237 | 0.015 | 0 | HFSC | chr16-56675189-566 |
| 0 | 4.44950154184153 | 0.185 | 0.009 | 0 | HFSC | chr16-68473096-684 |
| 0 | 4.20218435357484 | 0.193 | 0.012 | 0 | HFSC | chr16-69781532-697 |
| 0 | 4.31088449643488 | 0.192 | 0.01 | 0 | HFSC | chr17-12442728-124 |
| 0 | 4.34364475946814 | 0.252 | 0.013 | 0 | HFSC | chr17-16642301-166 |
| 0 | 5.08261299767785 | 0.148 | 0.004 | 0 | HFSC | chr17-16644095-166 |
| 0 | 4.59693266330762 | 0.148 | 0.007 | 0 | HFSC | chr18-31451417-314 |
| 0 | 5.3088737182609 | 0.114 | 0.003 | 0 | HFSC | chr18-52401223-524 |
| 0 | 4.94357971266692 | 0.144 | 0.005 | 0 | HFSC | chr2-168297293-168 |
| 0 | 4.39334701883611 | 0.182 | 0.009 | 0 | HFSC | chr2-225052826-225 |
| 0 | 4.56145983873494 | 0.152 | 0.007 | 0 | HFSC | chr20-47585787-475 |
| 0 | 4.14439879689817 | 0.186 | 0.012 | 0 | HFSC | chr3-64428180-6442 |
| 0 | 4.7551317129644 | 0.238 | 0.009 | 0 | HFSC | chr3-124661906-124 |
| 0 | 4.92921549089865 | 0.137 | 0.005 | 0 | HFSC | chr3-129039539-129 |
| 0 | 4.77776607546725 | 0.137 | 0.005 | 0 | HFSC | chr3-150843369-150 |
| 0 | 4.55997654717796 | 0.159 | 0.008 | 0 | HFSC | chr4-44343880-4434 |
| 0 | 5.86091185968703 | 0.12 | 0.002 | 0 | HFSC | chr4-130723672-130 |
| 0 | 5.17789195078967 | 0.12 | 0.003 | 0 | HFSC | chr5-49985326-4998 |
| 0 | 4.74349446456388 | 0.207 | 0.008 | 0 | HFSC | chr5-96751629-9675 |
| 0 | 4.38370528662191 | 0.224 | 0.012 | 0 | HFSC | chr5-98640271-9864 |
| 0 | 4.58760169386658 | 0.141 | 0.005 | 0 | HFSC | chr5-110250567-110 |
| 0 | 4.42900911750029 | 0.193 | 0.01 | 0 | HFSC | chr6-58397679-5839 |
| 0 | 4.83408606769629 | 0.159 | 0.006 | 0 | HFSC | chr7-43168554-4316 |
| 0 | 4.84492528295098 | 0.217 | 0.008 | 0 | HFSC | chr7-51074933-5107 |
| 0 | 5.4277581768194 | 0.104 | 0.003 | 0 | HFSC | chr7-54635910-5463 |
| 0 | 4.14599897321821 | 0.195 | 0.012 | 0 | HFSC | chr7-92836424-9283 |
| 0 | 4.30407547794741 | 0.185 | 0.01 | 0 | HFSC | chr7-122196629-122 |
| 0 | 4.68238878001058 | 0.123 | 0.004 | 0 | HFSC | chr7-132860517-132 |
| 0 | 4.47646851926274 | 0.175 | 0.008 | 0 | HFSC | chr8-11329001-1132 |
| 0 | 4.59120326946979 | 0.15 | 0.007 | 0 | HFSC | chr8-30254335-3025 |
| 0 | 4.20075642418171 | 0.206 | 0.012 | 0 | HFSC | chr8-30282589-3028 |
| 0 | 5.1393448535878 | 0.195 | 0.006 | 0 | HFSC | chr8-46918180-4691 |
| 0 | 4.23858222646552 | 0.172 | 0.01 | 0 | HFSC | chr8-72483562-7248 |
| 0 | 4.59162856021043 | 0.15 | 0.006 | 0 | HFSC | chr8-77380606-7738 |
| 0 | 4.32022176847584 | 0.2 | 0.011 | 0 | HFSC | chr8-77467866-7746 |
| 0 | 4.82128846323876 | 0.114 | 0.004 | 0 | HFSC | chr9-12512479-1251 |
| 0 | 4.78058620084068 | 0.124 | 0.005 | 0 | HFSC | chr9-39783904-3978 |
| 0 | 4.24838928077528 | 0.223 | 0.013 | 0 | HFSC | chr9-40549204-4054 |
| 0 | 4.27893599326793 | 0.157 | 0.008 | 0 | HFSC | chr9-84406107-8440 |
| 0 | 5.36969500705243 | 0.244 | 0.007 | 0 | HFSC | chr9-102212425-102 |
| 3.86586444 | 4.40931147141076 | 0.145 | 0.007 | 2.8327740 | HFSC | chr5-70247768-7024 |
| 5.20144691 | 4.9661893055503 | 0.109 | 0.004 | 3.8114434 | HFSC | chr5-101970568-101 |

| | | | | | | |
|---|---|---|---|---|---|---|
| 4.52773706 | 4.32629985372188 | 0.145 | 0.007 | 3.3177717 | HFSC | chr16-55547557-555 |
| 2.39466293 | 4.16997118388599 | 0.164 | 0.01 | 1.7547275 | HFSC | chr17-31774131-317 |
| 8.46202860 | 4.20026106191925 | 0.15 | 0.008 | 6.2006868 | HFSC | chr8-30043433-3004 |
| 3.45392424 | 4.57897521130136 | 0.121 | 0.005 | 2.5309182 | HFSC | chr10-87571610-875 |
| 7.40352921 | 4.40087249397671 | 0.131 | 0.006 | 5.4250544 | HFSC | chr10-97066003-970 |
| 4.70654578 | 4.21861423109469 | 0.171 | 0.011 | 3.4487967 | HFSC | chr9-78517788-7851 |
| 1.12774401 | 4.35221025355082 | 0.144 | 0.007 | 8.2637247 | HFSC | chr14-81745212-817 |
| 1.71768205 | 4.3973749008642 | 0.144 | 0.007 | 1.2586590 | HFSC | chr15-14387985-143 |
| 2.98494695 | 4.45772783451659 | 0.13 | 0.006 | 2.1872676 | HFSC | chr4-20544788-2054 |
| 1.02811954 | 4.50349707586877 | 0.133 | 0.006 | 7.5337104 | HFSC | chr9-77566123-7756 |
| 1.13568161 | 4.83263828601408 | 0.107 | 0.004 | 8.3218887 | HFSC | chr2-205037733-205 |
| 1.47903494 | 4.17939270914954 | 0.151 | 0.008 | 1.0837865 | HFSC | chr3-22155772-2215 |
| 1.68286736 | 4.65655488504808 | 0.116 | 0.005 | 1.2331479 | HFSC | chr1-126951281-126 |
| 1.48853992 | 4.24019154125859 | 0.141 | 0.008 | 1.0907514 | HFSC | chr10-27703936-277 |
| 3.43324187 | 4.70744018918592 | 0.121 | 0.005 | 2.5157629 | HFSC | chr12-33511511-335 |
| 1.09305188 | 4.5839337322108 | 0.121 | 0.006 | 8.0095125 | HFSC | chr1-149802005-149 |
| 1.10290946 | 4.67590565696208 | 0.111 | 0.005 | 8.0817455 | HFSC | chr17-33157074-331 |
| 2.58121579 | 4.41359012499795 | 0.14 | 0.008 | 1.8914271 | HFSC | chr8-30256016-3025 |
| 4.53942955 | 4.68959352415852 | 0.113 | 0.005 | 3.3263396 | HFSC | chr3-21799240-2179 |
| 6.40146119 | 4.34237217417068 | 0.116 | 0.005 | 4.6907731 | HFSC | chr20-48306126-483 |
| 4.71453665 | 4.33451561747019 | 0.141 | 0.008 | 3.4546521 | HFSC | chr19-25071425-250 |
| 6.47690184 | 4.39088703086956 | 0.134 | 0.007 | 4.7460534 | HFSC | chr14-81737409-817 |
| 1.29734518 | 4.4900014589468 | 0.113 | 0.005 | 9.5065043 | HFSC | chr4-20513299-2051 |
| 4.54251624 | 4.19367643658018 | 0.15 | 0.009 | 3.3286014 | HFSC | chr20-48299322-482 |
| 2.10980620 | 4.3020847406852 | 0.13 | 0.007 | 1.5459942 | HFSC | chr18-49369516-493 |
| 3.41801076 | 4.42287716070182 | 0.117 | 0.005 | 2.5046020 | HFSC | chr17-22261141-222 |
| 4.78327567 | 4.20026106191925 | 0.145 | 0.009 | 3.5050217 | HFSC | chr15-24187359-241 |
| 2.80014573 | 4.43595161047983 | 0.12 | 0.006 | 2.0518515 | HFSC | chr2-46391921-4639 |
| 1.58080379 | 4.13504256112503 | 0.135 | 0.008 | 1.1583592 | HFSC | chr14-67125763-671 |
| 1.15136141 | 4.46280989970342 | 0.126 | 0.006 | 8.4367849 | HFSC | chr2-46392442-4639 |
| 2.79674770 | 4.37998131197479 | 0.121 | 0.006 | 2.0493616 | HFSC | chr17-68871691-688 |
| 4.14193206 | 4.33966859238188 | 0.117 | 0.006 | 3.0350669 | HFSC | chr3-64455801-6445 |
| 4.91375837 | 4.82057368063243 | 0.102 | 0.004 | 3.6006350 | HFSC | chr13-94280878-942 |
| 7.73350948 | 4.70942943041158 | 0.104 | 0.004 | 5.6668528 | HFSC | chr15-24276857-242 |
| 2.40798591 | 4.36706878573184 | 0.117 | 0.006 | 1.7644902 | HFSC | chr5-96052788-9605 |
| 1.10680668 | 4.23909157109559 | 0.128 | 0.007 | 8.1103030 | HFSC | chr10-85007944-850 |
| 9.79290647 | 4.35538662038437 | 0.121 | 0.006 | 7.1759089 | HFSC | chr9-66183203-6618 |
| 1.76852802 | 4.12010764446817 | 0.144 | 0.009 | 1.2959172 | HFSC | chr13-104419072-10 |
| 2.77974381 | 4.61541307285921 | 0.107 | 0.005 | 2.0369017 | HFSC | chr1-12283843-1228 |
| 6.80704330 | 4.33818579142678 | 0.116 | 0.006 | 4.9879698 | HFSC | chr4-58261887-5826 |
| 6.90333820 | 4.6013161987466 | 0.106 | 0.005 | 5.0585315 | HFSC | chr19-55447761-554 |
| 5.43598947 | 4.34393098578048 | 0.118 | 0.006 | 3.9833082 | HFSC | chr2-224160418-224 |
| 8.38154706 | 4.2905267936972 | 0.118 | 0.006 | 6.1417127 | HFSC | chr4-180356383-180 |

| | | | | | | | |
|---|---|---|---|---|---|---|---|
| 8.69108051 | 4.42747553121776 | | 0.11 | 0.005 | 6.3685283 | HFSC | chr1-171706404-171 |
| 2.55544149 | 4.57599265549309 | | 0.104 | 0.005 | 1.8725406 | HFSC | chr10-97250067-972 |
| 8.70933030 | 4.26201473646275 | | 0.123 | 0.007 | 6.3819011 | HFSC | chr7-131863104-131 |
| 2.79518974 | 4.41558599882707 | | 0.109 | 0.005 | 2.0482200 | HFSC | chr17-68236330-682 |
| 8.78341958 | 4.37083325498735 | | 0.103 | 0.005 | 6.4361912 | HFSC | chr6-104189020-104 |
| 9.05021074 | 4.70301554421534 | | 0.1 | 0.004 | 6.6316867 | HFSC | chr1-135922775-135 |
| 1.02374464 | 4.24155482714714 | | 0.114 | 0.006 | 7.5016526 | HFSC | chr10-51638689-516 |
| 1.58037325 | 4.26011949141846 | | 0.118 | 0.006 | 1.1580437 | HFSC | chr1-94163449-9416 |
| 4.27190207 | 4.35378064645856 | | 0.117 | 0.006 | 3.1303045 | HFSC | chr8-12348490-1234 |
| 6.01812590 | 4.28835273157379 | | 0.124 | 0.007 | 4.4098780 | HFSC | chr8-30155784-3015 |
| 6.48101409 | 4.38332930823872 | | 0.11 | 0.005 | 4.7490667 | HFSC | chr17-25022086-250 |
| 6.90376244 | 4.30080666758381 | | 0.117 | 0.006 | 5.0588423 | HFSC | chr1-120549983-120 |
| 2.30989649 | 4.25153817986367 | | 0.12 | 0.006 | 1.6926136 | HFSC | chr9-573554-574055 |
| 4.93276020 | 4.40925845857621 | | 0.124 | 0.007 | 3.6145589 | HFSC | chr3-34309913-3431 |
| 9.32313391 | 4.56797376953628 | | 0.103 | 0.005 | 6.8316755 | HFSC | chr11-55158528-551 |
| 3.81047405 | 4.21461374186264 | | 0.128 | 0.008 | 2.7921858 | HFSC | chr10-81824218-818 |
| 9.84274766 | 4.13302086233065 | | 0.126 | 0.007 | 7.2124308 | HFSC | chr1-17032883-1703 |
| 1.40988409 | 4.39005893851648 | | 0.106 | 0.005 | 1.0331151 | HFSC | chr19-27930995-279 |
| 2.80071985 | 4.21769065224504 | | 0.118 | 0.007 | 2.0522722 | HFSC | chr8-11256257-1125 |
| 4.70658626 | 4.22765114563379 | | 0.113 | 0.006 | 3.4488263 | HFSC | chr1-3900858-39013 |
| 4.06621615 | 4.19115938385959 | | 0.12 | 0.007 | 2.9795849 | HFSC | chr6-114540728-114 |
| 9.62032907 | 4.39835004553156 | | 0.103 | 0.005 | 7.0494500 | HFSC | chr1-186440072-186 |
| 6.30759854 | 4.16470749394315 | | 0.111 | 0.006 | 4.6219937 | HFSC | chr3-124926959-124 |
| 7.98886059 | 4.39102253289638 | | 0.109 | 0.006 | 5.8539654 | HFSC | chr18-66894299-668 |
| 3.05302740 | 4.44549559325615 | | 0.103 | 0.005 | 2.2371546 | HFSC | chr4-69938299-6993 |
| 3.90640501 | 4.11701232985778 | | 0.12 | 0.007 | 2.8624807 | HFSC | chr8-30315470-3031 |
| 2.32221602 | 4.41159870141158 | | 0.104 | 0.005 | 1.7016409 | HFSC | chr10-49691691-496 |
| 7.59630161 | 4.14409407345106 | | 0.121 | 0.007 | 5.5663115 | HFSC | chr18-31451981-314 |
| 5.07940966 | 4.3558330628774 | | 0.1 | 0.005 | 3.7220187 | HFSC | chr1-163139147-163 |
| 3.59493470 | 4.33746456552398 | | 0.1 | 0.005 | 2.6342459 | HFSC | chr10-97935407-979 |
| 3.39542991 | 4.41657646400474 | | 0.103 | 0.005 | 2.4880555 | HFSC | chr15-8779848-8780 |
| 1.61187527 | 4.47182146441791 | | 0.1 | 0.005 | 1.1811274 | HFSC | chr18-72592085-725 |
| 2.48318133 | 4.38628390933919 | | 0.103 | 0.005 | 1.8195908 | HFSC | chr15-36012398-360 |
| 4.71235462 | 4.26201473646275 | | 0.111 | 0.006 | 3.4530532 | HFSC | chr17-69126844-691 |
| 6.38255245 | 4.17467186019457 | | 0.103 | 0.006 | 4.6769174 | HFSC | chr11-16925822-169 |
| 1.62097542 | 4.10825728969129 | | 0.118 | 0.007 | 1.1877956 | HFSC | chr4-59963296-5996 |
| 1.89729408 | 4.18233525996255 | | 0.102 | 0.005 | 1.3902725 | HFSC | chr13-93041890-930 |
| 1.35854151 | 4.16362515083429 | | 0.106 | 0.006 | 9.9549303 | HFSC | chr18-2269770-2270 |
| 1.46636934 | 4.39714855123874 | | 0.102 | 0.006 | 1.0745055 | HFSC | chr18-36021501-360 |
| 2.31987392 | 4.17256361295591 | | 0.107 | 0.006 | 1.6999247 | HFSC | chr5-151810623-151 |
| 1.10193129 | 4.15936782362956 | | 0.102 | 0.006 | 8.0745778 | HFSC | chr6-34111820-3411 |
| 9.68663319 | | 4.132020952 | 0.1 | 0.006 | 7.0980354 | HFSC | chr4-20501670-2050 |
| 1.66454366 | 4.11241600299229 | | 0.104 | 0.007 | 1.2197210 | HFSC | chr2-46442691-4644 |

| | | | | | | |
|---|---|---|---|---|---|---|
| 0 | 4.86052816967822 | 0.15 | 0.005 | 0 | HS | chr1-16050662-1605 |
| 0 | 4.8312694525724 | 0.181 | 0.007 | 0 | HS | chr1-120019199-120 |
| 0 | 5.57494383707381 | 0.119 | 0.002 | 0 | HS | chr1-188497992-188 |
| 0 | 4.81041374698437 | 0.275 | 0.012 | 0 | HS | chr1-202861520-202 |
| 0 | 4.70857874524665 | 0.192 | 0.009 | 0 | HS | chr1-202874660-202 |
| 0 | 5.38020031279101 | 0.133 | 0.004 | 0 | HS | chr1-208896960-208 |
| 0 | 5.51837748488235 | 0.204 | 0.006 | 0 | HS | chr10-61225619-612 |
| 0 | 5.6662909569171 | 0.115 | 0.003 | 0 | HS | chr10-80108017-801 |
| 0 | 4.8494861106841 | 0.131 | 0.004 | 0 | HS | chr10-84449071-844 |
| 0 | 5.67182659115806 | 0.108 | 0.002 | 0 | HS | chr10-85029590-850 |
| 0 | 5.12414192697381 | 0.123 | 0.003 | 0 | HS | chr10-94020678-940 |
| 0 | 5.07852137050918 | 0.219 | 0.007 | 0 | HS | chr11-30105464-301 |
| 0 | 5.49629922908139 | 0.117 | 0.003 | 0 | HS | chr11-48491600-484 |
| 0 | 4.81067605374336 | 0.192 | 0.008 | 0 | HS | chr12-9567381-9567 |
| 0 | 5.08807849885576 | 0.154 | 0.005 | 0 | HS | chr12-20671157-206 |
| 0 | 4.96839226412761 | 0.169 | 0.005 | 0 | HS | chr12-28113238-281 |
| 0 | 5.02650824225593 | 0.188 | 0.006 | 0 | HS | chr13-77642421-776 |
| 0 | 4.90287516619733 | 0.21 | 0.009 | 0 | HS | chr14-32809440-328 |
| 0 | 4.86005412599464 | 0.208 | 0.008 | 0 | HS | chr15-1685479-1685 |
| 0 | 4.72762574237841 | 0.254 | 0.012 | 0 | HS | chr15-31385109-313 |
| 0 | 4.96036417334762 | 0.194 | 0.007 | 0 | HS | chr15-45120246-451 |
| 0 | 5.43755328392953 | 0.131 | 0.003 | 0 | HS | chr15-95069211-950 |
| 0 | 4.95848686853358 | 0.179 | 0.006 | 0 | HS | chr15-95347417-953 |
| 0 | 4.79430023181625 | 0.196 | 0.007 | 0 | HS | chr17-53822139-538 |
| 0 | 5.05651055218386 | 0.156 | 0.006 | 0 | HS | chr17-53836866-538 |
| 0 | 5.2742834413932 | 0.138 | 0.004 | 0 | HS | chr17-53837620-538 |
| 0 | 4.88691581415635 | 0.19 | 0.008 | 0 | HS | chr18-27110584-271 |
| 0 | 5.6672603162941 | 0.121 | 0.003 | 0 | HS | chr18-48662324-486 |
| 0 | 4.95805360720572 | 0.173 | 0.007 | 0 | HS | chr19-10886623-108 |
| 0 | 4.72583517459437 | 0.165 | 0.006 | 0 | HS | chr19-34654316-346 |
| 0 | 4.91143379572634 | 0.198 | 0.007 | 0 | HS | chr19-43978905-439 |
| 0 | 5.76242166857642 | 0.127 | 0.002 | 0 | HS | chr2-42480533-4248 |
| 0 | 4.91518884827236 | 0.194 | 0.007 | 0 | HS | chr2-136715982-136 |
| 0 | 4.85183490798812 | 0.3 | 0.012 | 0 | HS | chr2-221854757-221 |
| 0 | 5.45449784937264 | 0.135 | 0.004 | 0 | HS | chr3-118105250-118 |
| 0 | 5.26818815901577 | 0.19 | 0.006 | 0 | HS | chr3-145262097-145 |
| 0 | 5.34512180681289 | 0.123 | 0.003 | 0 | HS | chr4-34714699-3471 |
| 0 | 4.7098065758268 | 0.212 | 0.01 | 0 | HS | chr4-71264280-7126 |
| 0 | 4.79457999348144 | 0.162 | 0.006 | 0 | HS | chr4-157759447-157 |
| 0 | 5.31098515322408 | 0.179 | 0.005 | 0 | HS | chr5-139681790-139 |
| 0 | 4.95060680933339 | 0.252 | 0.009 | 0 | HS | chr5-145818472-145 |
| 0 | 4.72527475610423 | 0.188 | 0.009 | 0 | HS | chr5-153119985-153 |
| 0 | 5.34251263177694 | 0.14 | 0.004 | 0 | HS | chr5-159492490-159 |

| | | | | | | |
|---|---|---|---|---|---|---|
| 0 | 4.77958157708859 | 0.215 | 0.009 | 0 | HS | chr6-6278425-62789 |
| 0 | 4.73060350176494 | 0.175 | 0.007 | 0 | HS | chr6-11021212-1102 |
| 0 | 5.37921837232322 | 0.123 | 0.003 | 0 | HS | chr6-24517507-2451 |
| 0 | 5.20024214099188 | 0.154 | 0.005 | 0 | HS | chr6-98048013-9804 |
| 0 | 4.76548208845636 | 0.192 | 0.008 | 0 | HS | chr6-124911103-124 |
| 0 | 5.06084262126502 | 0.188 | 0.007 | 0 | HS | chr7-43643082-4364 |
| 0 | 5.04393574662546 | 0.165 | 0.005 | 0 | HS | chr7-117006831-117 |
| 0 | 4.87914857180549 | 0.173 | 0.006 | 0 | HS | chr7-133216488-133 |
| 0 | 4.9755742294393 | 0.156 | 0.006 | 0 | HS | chr8-12058603-1205 |
| 0 | 5.56393459229532 | 0.188 | 0.005 | 0 | HS | chr8-29571109-2957 |
| 0 | 4.70890396711825 | 0.2 | 0.008 | 0 | HS | chr8-59485552-5948 |
| 0 | 4.84345749388785 | 0.258 | 0.01 | 0 | HS | chr9-14374138-1437 |
| 0 | 5.09232241775718 | 0.179 | 0.006 | 0 | HS | chr9-45139604-4514 |
| 5.07263883 | 4.82180445895352 | 0.171 | 0.007 | 3.7170572 | HS | chr1-50895261-5089 |
| 3.52643111 | 4.9198023312164 | 0.152 | 0.005 | 2.5840488 | HS | chr10-103947360-10 |
| 5.31794541 | 4.90800088998368 | 0.169 | 0.007 | 3.8968095 | HS | chr9-91633562-9163 |
| 5.13963284 | 5.34792940533992 | 0.121 | 0.003 | 3.7661481 | HS | chr5-134789636-134 |
| 8.76015740 | 5.02689136954112 | 0.14 | 0.005 | 6.4191454 | HS | chr7-92775348-9277 |
| 2.16012335 | 5.47871643728279 | 0.123 | 0.003 | 1.5828649 | HS | chr14-71504487-715 |
| 6.46763897 | 5.25476812512409 | 0.117 | 0.003 | 4.7392659 | HS | chr15-19606801-196 |
| 9.77468177 | 4.79103588619177 | 0.156 | 0.006 | 7.1625544 | HS | chr11-65377344-653 |
| 8.87964759 | 5.09131393450795 | 0.121 | 0.003 | 6.5067038 | HS | chr14-57814296-578 |
| 8.73248356 | 4.90929710949677 | 0.156 | 0.006 | 6.3988670 | HS | chr4-160050422-160 |
| 5.57140160 | 5.01436280779327 | 0.144 | 0.005 | 4.0825336 | HS | chr3-143910775-143 |
| 6.88185007 | 5.26569903617505 | 0.123 | 0.004 | 5.0427857 | HS | chr11-79722621-797 |
| 1.20967734 | 4.85502129648062 | 0.15 | 0.006 | 8.8641042 | HS | chr17-11036569-110 |
| 1.52667398 | 5.27456736711255 | 0.121 | 0.003 | 1.1186947 | HS | chr7-31500018-3150 |
| 2.22918555 | 5.41014156898841 | 0.1 | 0.002 | 1.6334713 | HS | chr8-111414979-111 |
| 3.39876113 | 4.95177026509057 | 0.142 | 0.005 | 2.4904966 | HS | chr12-28116604-281 |
| 6.92944912 | 5.36613526380509 | 0.1 | 0.002 | 5.0776647 | HS | chr9-74274236-7427 |
| 9.31904177 | 5.4489040995204 | 0.11 | 0.003 | 6.8286769 | HS | chr9-55349372-5534 |
| 1.28983520 | 4.8666232008501 | 0.152 | 0.006 | 9.4514738 | HS | chr4-175848245-175 |
| 8.13070539 | 4.77682874572844 | 0.15 | 0.006 | 5.9579044 | HS | chr8-63979796-6398 |
| 4.64614421 | 4.9169487408664 | 0.15 | 0.006 | 3.4045365 | HS | chr4-114007318-114 |
| 4.55325789 | 4.95513558891323 | 0.135 | 0.005 | 3.3364725 | HS | chr8-21721777-2172 |
| 5.20537405 | 5.44179499666738 | 0.1 | 0.002 | 3.8143211 | HS | chr2-123621648-123 |
| 8.39974642 | 4.98474936641927 | 0.125 | 0.004 | 6.1550485 | HS | chr18-48670403-486 |
| 1.31477695 | 5.22748141308704 | 0.125 | 0.004 | 9.6342384 | HS | chr4-6256469-62569 |
| 3.62442444 | 4.87135357108273 | 0.142 | 0.005 | 2.6558550 | HS | chr14-72254821-722 |
| 2.31226478 | 5.01821822316839 | 0.133 | 0.005 | 1.6943490 | HS | chr15-95096791-950 |
| 3.61533052 | 4.90387715723617 | 0.146 | 0.006 | 2.6491912 | HS | chr17-11038784-110 |
| 1.67484159 | 4.75824521397553 | 0.173 | 0.008 | 1.2272669 | HS | chr9-22039088-2203 |
| 1.69868742 | 5.10270235540625 | 0.112 | 0.003 | 1.2447403 | HS | chr3-117321049-117 |

| | | | | | | | |
|---|---|---|---|---|---|---|---|
| 1.35814302 | 4.85984309447453 | | 0.148 | 0.006 | 9.9520102 | HS | chr16-4072130-4072 |
| 4.74618517 | 4.82824316983098 | | 0.14 | 0.005 | 3.4778431 | HS | chr17-17801761-178 |
| 1.21320534 | 4.80160578317528 | | 0.148 | 0.006 | 8.8899562 | HS | chr5-98031468-9803 |
| 2.18354236 | | 5.019982008 | 0.112 | 0.003 | 1.6000256 | HS | chr14-36605455-366 |
| 9.53525147 | 5.19543729545719 | | 0.119 | 0.004 | 6.9871080 | HS | chr15-2204369-2204 |
| 1.41304687 | 4.98058497267918 | | 0.119 | 0.004 | 1.0354327 | HS | chr9-41643218-4164 |
| 3.98969021 | 5.2305290859359 | | 0.104 | 0.003 | 2.9235093 | HS | chr7-133985233-133 |
| 4.26663143 | 5.17934671865996 | | 0.115 | 0.003 | 3.1264424 | HS | chr2-29164459-2916 |
| 4.39216469 | 4.72657722706035 | | 0.138 | 0.005 | 3.2184289 | HS | chr2-171224342-171 |
| 7.76654703 | 4.72169706562776 | | 0.154 | 0.007 | 5.6910616 | HS | chr17-44524974-445 |
| 1.19884545 | 4.71499189883597 | | 0.144 | 0.006 | 8.7847318 | HS | chr15-39770290-397 |
| 1.71001837 | 5.12632215875661 | | 0.125 | 0.004 | 1.2530433 | HS | chr1-120213375-120 |
| 2.04789360 | 5.00155100357764 | | 0.127 | 0.005 | 1.5006268 | HS | chr3-17075493-1707 |
| 5.60989123 | 5.2412231680807 | | 0.104 | 0.003 | 4.1107375 | HS | chr15-31387073-313 |
| 5.67013487 | 5.28674895725178 | | 0.102 | 0.003 | 4.1548820 | HS | chr12-46564595-465 |
| 1.67379102 | 5.13374789618853 | | 0.11 | 0.003 | 1.2264971 | HS | chr8-46931032-4693 |
| 2.04975422 | 4.94053423722728 | | 0.115 | 0.004 | 1.5019902 | HS | chr9-83881143-8388 |
| 1.82606211 | 5.16139498301242 | | 0.106 | 0.003 | 1.3380762 | HS | chr17-53881908-538 |
| 2.24686728 | 4.99553603860932 | | 0.119 | 0.004 | 1.6464279 | HS | chr17-10950196-109 |
| 7.83652794 | 4.84220983086714 | | 0.14 | 0.006 | 5.7423412 | HS | chr8-67301035-6730 |
| 1.85484985 | 4.93209093298541 | | 0.127 | 0.005 | 1.3591709 | HS | chr1-174598273-174 |
| 2.07665147 | 5.22739066284953 | | 0.125 | 0.005 | 1.5216995 | HS | chr15-78576280-785 |
| 3.72354628 | 4.81985622282077 | | 0.121 | 0.004 | 2.7284881 | HS | chr7-32105083-3210 |
| 6.65682605 | 4.79657229649497 | | 0.138 | 0.006 | 4.8778958 | HS | chr4-180097828-180 |
| 8.95075004 | 5.25565465239506 | | 0.104 | 0.003 | 6.5588053 | HS | chr17-36063560-360 |
| 1.13179556 | 5.10491217191169 | | 0.108 | 0.003 | 8.2934130 | HS | chr1-200412897-200 |
| 1.91380424 | 4.89632097051492 | | 0.108 | 0.003 | 1.4023706 | HS | chr10-64640129-646 |
| 5.51631639 | 4.84056516364007 | | 0.135 | 0.006 | 4.0421690 | HS | chr14-32605632-326 |
| 1.62739869 | 4.75409838367629 | | 0.125 | 0.005 | 1.1925024 | HS | chr18-68380102-683 |
| 6.84228461 | 4.98150669447683 | | 0.117 | 0.004 | 5.0137935 | HS | chr13-104658709-10 |
| 5.12019308 | 5.16376741421604 | | 0.106 | 0.003 | 3.7519034 | HS | chr3-22265400-2226 |
| 5.65577510 | 4.75742189567951 | | 0.129 | 0.005 | 4.1443596 | HS | chr12-31770214-317 |
| 7.12527975 | 4.92752257209913 | | 0.117 | 0.004 | 5.2211627 | HS | chr4-6289325-62898 |
| 3.77522905 | 4.94856432794693 | | 0.104 | 0.003 | 2.7663594 | HS | chr3-121223371-121 |
| 2.26330114 | 4.85822708767603 | | 0.135 | 0.006 | 1.6584701 | HS | chr17-11040593-110 |
| 9.80216639 | 5.08373787787888 | | 0.11 | 0.004 | 7.1826942 | HS | chr1-87254701-8725 |
| 3.69097085 | 4.75508214961377 | | 0.127 | 0.005 | 2.7046179 | HS | chr15-9155599-9156 |
| 5.75911487 | 4.94500300028031 | | 0.115 | 0.004 | 4.2200835 | HS | chr14-72250109-722 |
| 1.78579246 | 4.99654133115035 | | 0.1 | 0.003 | 1.3085680 | HS | chr15-21793371-217 |
| 8.82311441 | 4.91041991399458 | | 0.127 | 0.005 | 6.4652782 | HS | chr13-95031472-950 |
| 1.37504832 | 4.73818906656784 | | 0.127 | 0.005 | 1.0075886 | HS | chr3-7143540-71440 |
| 3.11258389 | 4.81531812908597 | | 0.108 | 0.004 | 2.2807956 | HS | chr5-159675117-159 |
| 4.80483321 | 5.04355341603028 | | 0.104 | 0.004 | 3.5208184 | HS | chr3-148955118-148 |

| | | | | | | | |
|---|---|---|---|---|---|---|---|
| 8.72740498 | 4.80713062904773 | | 0.106 | 0.004 | 6.3951456 | HS | chr4-36269170-3626 |
| 1.89637841 | 4.72798536446741 | | 0.125 | 0.005 | 1.3896016 | HS | chr3-43905408-4390 |
| 1.32891623 | 4.87715208694029 | | 0.104 | 0.004 | 9.7378463 | HS | chr1-96028959-9602 |
| 1.79879172 | 4.70849643179961 | | 0.117 | 0.005 | 1.3180934 | HS | chr13-70131919-701 |
| 2.02411003 | | 4.796485224 | 0.119 | 0.005 | 1.4831990 | HS | chr10-20219954-202 |
| 6.45702187 | 5.08886577998984 | | 0.102 | 0.003 | 4.7314860 | HS | chr14-98879610-988 |
| 1.39943358 | 4.81610399640481 | | 0.121 | 0.005 | 1.0254573 | HS | chr1-239302139-239 |
| 4.45223279 | 4.76057121046024 | | 0.1 | 0.003 | 3.2624448 | HS | chr8-57359755-5736 |
| 4.23988337 | 4.83991263718951 | | 0.108 | 0.004 | 3.1068423 | HS | chr6-55026816-5502 |
| 6.36463059 | 4.78748809335053 | | 0.108 | 0.004 | 4.6637849 | HS | chr1-245380307-245 |
| 5.08107923 | 4.87244179359834 | | 0.108 | 0.004 | 3.7232421 | HS | chr15-45079011-450 |
| 1.01235492 | 4.81840168304598 | | 0.11 | 0.004 | 7.4181926 | HS | chr1-222669746-222 |
| 7.24661395 | 4.74414083425887 | | 0.115 | 0.005 | 5.3100723 | HS | chr4-83732850-8373 |
| 7.43870462 | 4.75208842009223 | | 0.112 | 0.005 | 5.4508298 | HS | chr1-151020759-151 |
| 8.36876980 | 4.77691919922783 | | 0.112 | 0.005 | 6.1323499 | HS | chr13-38271997-382 |
| 4.17916107 | 4.71996114486737 | | 0.121 | 0.006 | 3.0623471 | HS | chr14-34875518-348 |
| 8.41300303 | 4.83664249656767 | | 0.115 | 0.005 | 6.1647625 | HS | chr1-86195457-8619 |
| 1.26118768 | 4.78093515662121 | | 0.11 | 0.005 | 9.2415545 | HS | chr10-84294552-842 |
| 2.79880256 | 4.73984117653433 | | 0.108 | 0.005 | 2.0508673 | HS | chr10-90903846-909 |
| 3.12242292 | 4.73394847823737 | | 0.1 | 0.004 | 2.2880053 | HS | chr9-87973672-8797 |
| 1.40622258 | 4.72652023468876 | | 0.104 | 0.005 | 1.0304320 | HS | chr6-122554373-122 |
| 0 | 3.56764195913206 | | 0.222 | 0.019 | 0 | IRS | chr4-30679350-3067 |
| 2.86154999 | 4.20232055818314 | | 0.151 | 0.009 | 2.0968465 | IRS | chr5-60587560-6058 |
| 2.85786903 | 3.45811917637512 | | 0.254 | 0.026 | 2.0941492 | IRS | chr1-255985881-255 |
| 3.69859939 | 3.51820160787597 | | 0.229 | 0.021 | 2.7102078 | IRS | chr9-18216553-1821 |
| 3.32154144 | 3.4380803029054 | | 0.226 | 0.022 | 2.4339126 | IRS | chr5-109913264-109 |
| 4.34442699 | 4.5393134741522 | | 0.101 | 0.004 | 3.1834483 | IRS | chr19-26830859-268 |
| 3.81813995 | 3.61558050789273 | | 0.18 | 0.014 | 2.7978031 | IRS | chr17-24087432-240 |
| 2.07887056 | 3.47978832251122 | | 0.2 | 0.018 | 1.5233256 | IRS | chr11-74423571-744 |
| 1.68606826 | 3.50513649926691 | | 0.202 | 0.019 | 1.2354934 | IRS | chr1-134695407-134 |
| 1.37626367 | 3.85352746338375 | | 0.159 | 0.012 | 1.0084792 | IRS | chr4-181428915-181 |
| 8.39258559 | 4.32968592897255 | | 0.115 | 0.006 | 6.1498013 | IRS | chr5-136469169-136 |
| 3.00105299 | 3.44296124799256 | | 0.217 | 0.022 | 2.1990695 | IRS | chr5-159218892-159 |
| 1.79385984 | 3.98139123422023 | | 0.134 | 0.008 | 1.3144795 | IRS | chr4-29291445-2929 |
| 2.36715859 | 4.04726734331442 | | 0.141 | 0.009 | 1.7345733 | IRS | chr12-20066319-200 |
| 3.37508534 | 3.44553533654511 | | 0.205 | 0.02 | 2.4731477 | IRS | chr10-101486785-10 |
| 1.01941667 | 4.4315629544899 | | 0.107 | 0.005 | 7.4699388 | IRS | chr1-142241918-142 |
| 3.78831751 | 3.42649612294183 | | 0.202 | 0.02 | 2.7759502 | IRS | chr5-74271472-7427 |
| 2.77578332 | 4.17945915126107 | | 0.117 | 0.007 | 2.0339996 | IRS | chr10-67545097-675 |
| 1.40955961 | 4.03554514320166 | | 0.122 | 0.007 | 1.0328773 | IRS | chr3-148929593-148 |
| 4.32931908 | 3.75722546984222 | | 0.145 | 0.011 | 3.1723778 | IRS | chr7-562102-562603 |
| 3.17661248 | 3.42468653650025 | | 0.191 | 0.019 | 2.3277136 | IRS | chr6-131187627-131 |
| 8.12422829 | 3.96486868218382 | | 0.127 | 0.008 | 5.9531582 | IRS | chr11-43730008-437 |

| | | | | | | | | |
|---|---|---|---|---|---|---|---|---|
| 2.99654044 | 3.50377367072537 | | 0.187 | 0.018 | 2.1957629 | IRS | | chr14-13366328-133 |
| 6.36027946 | 4.1923247286699 | | 0.126 | 0.008 | 4.6605965 | IRS | | chr18-11681051-116 |
| 3.52363848 | 3.54917788658013 | | 0.17 | 0.015 | 2.5820024 | IRS | | chr1-202847505-202 |
| 7.55739117 | 3.42436133118127 | | 0.183 | 0.018 | 5.5377993 | IRS | | chr4-70430180-7043 |
| 1.28424855 | 3.79779258943828 | | 0.132 | 0.009 | 9.4105367 | IRS | | chr17-10999010-109 |
| 9.08176936 | 3.64841799111169 | | 0.159 | 0.014 | 6.6548118 | IRS | | chr4-148063428-148 |
| 2.10069920 | 3.55550616475027 | | 0.151 | 0.012 | 1.5393209 | IRS | | chr5-56871140-5687 |
| 3.59660110 | 3.66644201008425 | | 0.146 | 0.012 | 2.6354670 | IRS | | chr9-64365822-6436 |
| 8.89130767 | 4.0105286395271 | | 0.118 | 0.007 | 6.5152479 | IRS | | chr1-235090970-235 |
| 1.51228957 | 3.50959816055194 | | 0.162 | 0.014 | 1.1081543 | IRS | | chr12-6642284-6642 |
| 1.19490625 | 3.64570433657352 | | 0.155 | 0.013 | 8.7558667 | IRS | | chr20-5176730-5177 |
| 1.36632960 | 3.41389100299563 | | 0.152 | 0.013 | 1.0011998 | IRS | | chr17-10937641-109 |
| 3.27727998 | 3.78396670647053 | | 0.149 | 0.013 | 2.4014793 | IRS | | chr11-24021581-240 |
| 2.57618757 | 3.43466306044564 | | 0.16 | 0.015 | 1.8877426 | IRS | | chr3-43053875-4305 |
| 2.26780181 | 3.59871395243147 | | 0.138 | 0.011 | 1.6617680 | IRS | | chr14-66864926-668 |
| 1.15127665 | 3.94099827652799 | | 0.118 | 0.008 | 8.4361638 | IRS | | chr2-54493189-5449 |
| 4.46608936 | 3.98254426021416 | | 0.13 | 0.01 | 3.2725984 | IRS | | chr7-61738023-6173 |
| 7.93352861 | 3.51469019929352 | | 0.166 | 0.016 | 5.8134200 | IRS | | chr4-158124375-158 |
| 2.96821109 | 3.92110740039566 | | 0.106 | 0.006 | 2.1750041 | IRS | | chr14-33074053-330 |
| 2.31631596 | 3.81284719178821 | | 0.127 | 0.009 | 1.6973175 | IRS | | chr5-56834577-5683 |
| 2.49634050 | 4.07293771649142 | | 0.109 | 0.007 | 1.8292334 | IRS | | chr5-122492397-122 |
| 5.60371531 | 3.41681969107937 | | 0.17 | 0.017 | 4.1062120 | IRS | | chr13-93596397-935 |
| 6.60489131 | 3.47116116011499 | | 0.148 | 0.013 | 4.8398397 | IRS | | chr1-197495181-197 |
| 6.05535338 | 3.51462434498344 | | 0.157 | 0.015 | 4.4371570 | IRS | | chr10-103151100-10 |
| 3.04805405 | 3.68809579303284 | | 0.138 | 0.012 | 2.2335103 | IRS | | chr17-35696949-356 |
| 6.35037488 | 3.70746638694118 | | 0.128 | 0.01 | 4.6533388 | IRS | | chr3-145983448-145 |
| 1.05739264 | 3.74105290141321 | | 0.117 | 0.008 | 7.7482137 | IRS | | chr17-76816960-768 |
| 1.87688972 | | 3.503535856 | 0.141 | 0.012 | 1.3753209 | IRS | | chr8-70039738-7004 |
| 4.18799638 | 3.45629850189031 | | 0.152 | 0.015 | 3.0688213 | IRS | | chr1-117469996-117 |
| 2.82168875 | 3.47725096567893 | | 0.155 | 0.015 | 2.0676375 | IRS | | chr18-3204892-3205 |
| 9.46069530 | 3.88204128798526 | | 0.105 | 0.007 | 6.9324758 | IRS | | chr2-178711120-178 |
| 1.22671222 | 3.57573110218173 | | 0.134 | 0.011 | 8.9889301 | IRS | | chr17-24121050-241 |
| 5.43021665 | 3.78700451922936 | | 0.122 | 0.009 | 3.9790781 | IRS | | chr10-99725473-997 |
| 1.75627395 | 4.01349320759924 | | 0.104 | 0.007 | 1.2869378 | IRS | | chr5-72023346-7202 |
| 1.84130451 | 3.65122588400515 | | 0.139 | 0.012 | 1.3492453 | IRS | | chr18-16228522-162 |
| 2.95398096 | 3.44397577103786 | | 0.144 | 0.013 | 2.1645768 | IRS | | chr2-63099795-6310 |
| 8.02984007 | 3.77916455407582 | | 0.124 | 0.01 | 5.8839937 | IRS | | chr9-102253211-102 |
| 1.61898764 | 3.42113329495177 | | 0.149 | 0.014 | 1.1863391 | IRS | | chr16-74766478-747 |
| 2.87772056 | 3.57226255009381 | | 0.139 | 0.012 | 2.1086957 | IRS | | chr9-64352066-6435 |
| 7.44572620 | 3.55850243984258 | | 0.134 | 0.012 | 5.4559750 | IRS | | chr10-102741645-10 |
| 3.09318774 | 3.46658297546675 | | 0.151 | 0.015 | 2.2665828 | IRS | | chr17-35915836-359 |
| 2.00853850 | 3.38800553026492 | | 0.159 | 0.016 | 1.4717887 | IRS | | chr5-159237201-159 |
| 2.92111882 | 3.38296638858372 | | 0.148 | 0.014 | 2.1404965 | IRS | | chr15-44320606-443 |

| | | | | | | | |
|---|---|---|---|---|---|---|---|
| 1.32952399 | 3.79343122319668 | | 0.117 | 0.009 | 9.7422998 | IRS | chr4-132983672-132 |
| 1.75374685 | 3.62612078157906 | | 0.132 | 0.011 | 1.2850860 | IRS | chr2-78675724-7867 |
| 2.89471426 | 3.52357890406283 | | 0.149 | 0.015 | 2.1211481 | IRS | chr8-108245237-108 |
| 5.43845730 | 3.6868839709441 | | 0.113 | 0.008 | 3.9851166 | IRS | chr1-241060725-241 |
| 1.82716785 | 3.62258754958882 | | 0.124 | 0.01 | 1.3388864 | IRS | chr10-73791346-737 |
| 1.20801834 | 4.02417762168305 | | 0.107 | 0.008 | 8.8519477 | IRS | chr1-153693481-153 |
| 2.63861636 | 3.3966666495496 | | 0.143 | 0.014 | 1.9334883 | IRS | chr11-74522666-745 |
| 9.23404885 | 3.59500130280903 | | 0.121 | 0.01 | 6.7663970 | IRS | chr17-30445644-304 |
| 4.15193689 | 3.65356360275556 | | 0.127 | 0.011 | 3.0423981 | IRS | chr17-30649926-306 |
| 4.80065395 | 3.40637383929527 | | 0.14 | 0.013 | 3.5177559 | IRS | chr5-98822515-9882 |
| 6.63776193 | 3.60258421168526 | | 0.127 | 0.011 | 4.8639262 | IRS | chr10-57088466-570 |
| 3.82742376 | 3.44336264998239 | | 0.146 | 0.015 | 2.8046060 | IRS | chr9-41433727-4143 |
| 1.06797385 | 3.6165566029712 | | 0.124 | 0.011 | 7.8257493 | IRS | chr20-7249390-7249 |
| 2.73826506 | 3.60806273594964 | | 0.124 | 0.011 | 2.0065075 | IRS | chr18-64147756-641 |
| 1.49435813 | | 3.552292188 | 0.134 | 0.013 | 1.0950148 | IRS | chr12-19993972-199 |
| 3.94596543 | 3.51834306646121 | | 0.123 | 0.011 | 2.8914693 | IRS | chr4-71264280-7126 |
| 2.73851930 | 3.43436830383411 | | 0.145 | 0.015 | 2.0066938 | IRS | chr2-178620333-178 |
| 1.90868621 | 3.81399194479411 | | 0.109 | 0.008 | 1.3986203 | IRS | chr5-75825654-7582 |
| 1.08310569 | 3.56630819545444 | | 0.129 | 0.012 | 7.9366303 | IRS | chr5-133734435-133 |
| 2.14415687 | 3.39401110953033 | | 0.14 | 0.014 | 1.5711652 | IRS | chr1-202850167-202 |
| 1.58976738 | 3.66898422067565 | | 0.115 | 0.009 | 1.1649274 | IRS | chr5-130219069-130 |
| 2.36949497 | 3.42820445443456 | | 0.143 | 0.015 | 1.7362853 | IRS | chr5-154355790-154 |
| 5.01921847 | 3.5388980910504 | | 0.118 | 0.01 | 3.6779126 | IRS | chr12-21176107-211 |
| 6.71562433 | 3.58973433674685 | | 0.124 | 0.011 | 4.9209811 | IRS | chr18-12520689-125 |
| 9.96818224 | 3.83992811354222 | | 0.104 | 0.008 | 7.3043450 | IRS | chr3-157053790-157 |
| 1.07467071 | 3.48708583297567 | | 0.128 | 0.012 | 7.8748216 | IRS | chr17-26745556-267 |
| 7.05248483 | 3.59077540258732 | | 0.117 | 0.01 | 5.1678211 | IRS | chr1-241089097-241 |
| 9.21187302 | 3.59494214593676 | | 0.126 | 0.012 | 6.7501473 | IRS | chr5-109922149-109 |
| 9.68787902 | 3.71622139098586 | | 0.107 | 0.008 | 7.0989483 | IRS | chr8-59265110-5926 |
| 3.51685669 | 3.65620783324782 | | 0.116 | 0.01 | 2.5770330 | IRS | chr1-241066560-241 |
| 1.23654443 | 3.53652981011082 | | 0.123 | 0.011 | 9.0609771 | IRS | chr19-53626943-536 |
| 6.38854503 | 3.60050522694244 | | 0.124 | 0.011 | 4.6813085 | IRS | chr5-121331805-121 |
| 2.96193349 | 3.67678931884521 | | 0.11 | 0.009 | 2.1704041 | IRS | chr3-9364836-93653 |
| 3.84100256 | 3.70736485256688 | | 0.106 | 0.008 | 2.8145560 | IRS | chr14-97111656-971 |
| 4.11133899 | 3.3814149412523 | | 0.135 | 0.014 | 3.0126494 | IRS | chr10-84625549-846 |
| 7.14884147 | 3.64946118981355 | | 0.113 | 0.01 | 5.2384279 | IRS | chr15-37624890-376 |
| 8.27232451 | 3.6445100376693 | | 0.107 | 0.009 | 6.0616781 | IRS | chrX-116005483-116 |
| 1.09728650 | 3.39011219420551 | | 0.128 | 0.012 | 8.0405424 | IRS | chr15-31385109-313 |
| 6.55488129 | 3.55006292483253 | | 0.109 | 0.009 | 4.8031941 | IRS | chr1-153714135-153 |
| 1.28794770 | 3.62152240005151 | | 0.113 | 0.01 | 9.4376428 | IRS | chr4-142021348-142 |
| 3.45902308 | 3.37639104198149 | | 0.13 | 0.013 | 2.5346545 | IRS | chr1-199728294-199 |
| 5.79635816 | 3.54959559134107 | | 0.127 | 0.012 | 4.2473741 | IRS | chr6-98866532-9886 |
| 1.91253003 | 3.51747900361224 | | 0.109 | 0.009 | 1.4014369 | IRS | chr14-32809440-328 |

| | | | | | | |
|---|---|---|---|---|---|---|
| 2.78031194 | 3.42051720824171 | 0.134 | 0.014 | 2.0373180 | IRS | chr7-56886301-5688 |
| 1.63977364 | 3.40673201677143 | 0.117 | 0.011 | 1.2015703 | IRS | chr13-90001897-900 |
| 4.22988756 | 3.39772749316825 | 0.124 | 0.012 | 3.0995177 | IRS | chr14-78581123-785 |
| 8.22757249 | 3.39839595270886 | 0.118 | 0.011 | 6.0288853 | IRS | chr8-45449044-4544 |
| 3.83172526 | 3.66362267344062 | 0.113 | 0.01 | 2.8077579 | IRS | chr20-35797908-357 |
| 5.12403964 | 3.47927401953261 | 0.12 | 0.011 | 3.7547220 | IRS | chr1-184401619-184 |
| 5.13018063 | 3.3938517555031 | 0.123 | 0.012 | 3.7592219 | IRS | chr19-52366648-523 |
| 1.28760950 | 3.49173852549184 | 0.123 | 0.012 | 9.4351646 | IRS | chr15-51196065-511 |
| 1.38792081 | 3.69584232849119 | 0.104 | 0.008 | 1.0170211 | IRS | chr12-20067299-200 |
| 1.40404835 | 3.66305337938198 | 0.109 | 0.009 | 1.0288388 | IRS | chr2-45320063-4532 |
| 2.81144209 | 3.48121305057886 | 0.113 | 0.01 | 2.0601291 | IRS | chr15-76377716-763 |
| 4.65607706 | 3.71824725231254 | 0.116 | 0.011 | 3.4118149 | IRS | chr7-91413436-9141 |
| 1.16636824 | 3.36172643263385 | 0.11 | 0.01 | 8.5467499 | IRS | chr10-59311662-593 |
| 2.00425579 | 3.66625926780444 | 0.105 | 0.009 | 1.4686505 | IRS | chr3-21644075-2164 |
| 1.53450295 | 3.45197822206761 | 0.129 | 0.014 | 1.1244315 | IRS | chr19-52973113-529 |
| 3.41224347 | 3.40070412034636 | 0.111 | 0.01 | 2.5003760 | IRS | chr12-33787345-337 |
| 3.84923972 | 3.41287884298561 | 0.115 | 0.011 | 2.8205919 | IRS | chr6-71929783-7193 |
| 5.65262365 | 3.56500280539353 | 0.111 | 0.01 | 4.1420504 | IRS | chr10-84300967-843 |
| 5.94179647 | 3.5772412198247 | 0.101 | 0.008 | 4.3539464 | IRS | chr18-3235934-3236 |
| 1.07246893 | 3.37175583649083 | 0.118 | 0.012 | 7.8586876 | IRS | chr12-45856346-458 |
| 2.87092288 | 3.46455760064717 | 0.101 | 0.009 | 2.1037146 | IRS | chr12-9567381-9567 |
| 2.00552488 | 3.40934964548098 | 0.11 | 0.01 | 1.4695804 | IRS | chr4-125632871-125 |
| 3.03188396 | 3.51419611824139 | 0.106 | 0.01 | 2.2216614 | IRS | chr9-87026241-8702 |
| 3.59702680 | 3.56886417895431 | 0.101 | 0.009 | 2.6357789 | IRS | chr7-61785334-6178 |
| 5.26424869 | 3.44377029763474 | 0.117 | 0.012 | 3.8574624 | IRS | chr2-193330315-193 |
| 5.74220727 | 3.51527994304561 | 0.117 | 0.012 | 4.2076942 | IRS | chr9-54878500-5487 |
| 8.71681647 | 3.42752861298242 | 0.109 | 0.01 | 6.3873867 | IRS | chr1-128453794-128 |
| 7.94979624 | 3.54050279644142 | 0.102 | 0.009 | 5.8253403 | IRS | chr1-14736446-1473 |
| 8.19049148 | 3.45698439133472 | 0.111 | 0.011 | 6.0017136 | IRS | chr9-14374138-1437 |
| 6.67546243 | 3.38074840112436 | 0.118 | 0.012 | 4.8915519 | IRS | chr10-5361397-5361 |
| 5.92477753 | 3.48177276047346 | 0.115 | 0.012 | 4.3414755 | IRS | chr17-10909827-109 |
| 8.59026167 | 3.55797681834892 | 0.11 | 0.011 | 6.2946516 | IRS | chr4-6490529-64910 |
| 5.32286019 | 3.3655212379136 | 0.11 | 0.011 | 3.9004109 | IRS | chr1-2254272-22547 |
| 6.71564720 | 3.44865770189408 | 0.117 | 0.012 | 4.9209979 | IRS | chr2-190494250-190 |
| 1.64305312 | 3.36794129902489 | 0.109 | 0.011 | 1.2039734 | IRS | chr12-43429849-434 |
| 6.19912132 | 3.50639536217327 | 0.1 | 0.009 | 4.5425053 | IRS | chr7-9394373-93948 |
| 8.19001626 | 3.39694073632811 | 0.102 | 0.01 | 6.0013654 | IRS | chr16-69308814-693 |
| 9.05196678 | 3.44835369130281 | 0.1 | 0.009 | 6.6329734 | IRS | chr4-148058229-148 |
| 1.95510267 | 3.42260560071937 | 0.116 | 0.013 | 1.4326327 | IRS | chr5-97796314-9779 |
| 4.60280608 | 3.42888313736076 | 0.102 | 0.01 | 3.3727798 | IRS | chr13-67121342-671 |
| 1.35136827 | 3.40462836472927 | 0.104 | 0.011 | 9.9023672 | IRS | chr9-55446994-5544 |
| 2.48360519 | 3.36809002149318 | 0.102 | 0.011 | 1.8199014 | IRS | chr2-28068342-2806 |
| 0 | 5.53580662781969 | 0.169 | 0.004 | 0 | LAG | chr1-26904153-2690 |

| | | | | | | | |
|---|---|---|---|---|---|---|---|
| 0 | 5.5719232931683 | | 0.127 | 0.003 | 0 | LAG | chr1-79295821-7929 |
| 0 | 4.90279464356255 | | 0.152 | 0.006 | 0 | LAG | chr1-129494326-129 |
| 0 | 5.08527469412707 | | 0.193 | 0.006 | 0 | LAG | chr1-169368861-169 |
| 0 | 5.13367527955533 | | 0.134 | 0.004 | 0 | LAG | chr1-194800621-194 |
| 0 | 5.1215856881428 | | 0.11 | 0.003 | 0 | LAG | chr1-195087181-195 |
| 0 | 5.16282933235453 | | 0.211 | 0.006 | 0 | LAG | chr1-207396978-207 |
| 0 | 4.95402407675903 | | 0.166 | 0.005 | 0 | LAG | chr1-214619304-214 |
| 0 | 5.32334745989689 | | 0.127 | 0.003 | 0 | LAG | chr1-222694219-222 |
| 0 | | 5.067120243 | 0.175 | 0.006 | 0 | LAG | chr1-227124141-227 |
| 0 | 5.12453434164763 | | 0.132 | 0.004 | 0 | LAG | chr1-227154718-227 |
| 0 | 4.91065866985583 | | 0.141 | 0.005 | 0 | LAG | chr1-233579774-233 |
| 0 | 5.66589480160011 | | 0.25 | 0.006 | 0 | LAG | chr10-31187621-311 |
| 0 | 5.55563689941194 | | 0.102 | 0.002 | 0 | LAG | chr10-45657695-456 |
| 0 | 5.55330641186465 | | 0.196 | 0.004 | 0 | LAG | chr10-55228464-552 |
| 0 | 5.19853157510841 | | 0.144 | 0.004 | 0 | LAG | chr10-57623384-576 |
| 0 | 4.97576523700906 | | 0.183 | 0.007 | 0 | LAG | chr10-57735936-577 |
| 0 | 5.88634919107393 | | 0.191 | 0.004 | 0 | LAG | chr10-72404190-724 |
| 0 | 5.02763966767444 | | 0.261 | 0.01 | 0 | LAG | chr10-88229549-882 |
| 0 | 5.09419481133555 | | 0.169 | 0.005 | 0 | LAG | chr10-94236148-942 |
| 0 | 5.13390678345961 | | 0.158 | 0.005 | 0 | LAG | chr11-14529060-145 |
| 0 | 4.85354677329705 | | 0.152 | 0.006 | 0 | LAG | chr11-64155727-641 |
| 0 | 5.03072282394393 | | 0.194 | 0.006 | 0 | LAG | chr11-64168862-641 |
| 0 | 6.30151596305115 | | 0.143 | 0.002 | 0 | LAG | chr11-64267627-642 |
| 0 | 5.98568941524871 | | 0.141 | 0.002 | 0 | LAG | chr11-64289817-642 |
| 0 | 5.2907235918701 | | 0.169 | 0.004 | 0 | LAG | chr11-64291413-642 |
| 0 | 6.39909636530663 | | 0.231 | 0.004 | 0 | LAG | chr11-65506857-655 |
| 0 | 4.9175412789651 | | 0.172 | 0.007 | 0 | LAG | chr11-65517225-655 |
| 0 | 5.87432367349781 | | 0.161 | 0.003 | 0 | LAG | chr11-76834270-768 |
| 0 | 5.55803083080624 | | 0.207 | 0.005 | 0 | LAG | chr12-10188869-101 |
| 0 | 4.95403300464736 | | 0.2 | 0.008 | 0 | LAG | chr12-19430357-194 |
| 0 | 6.14223532272943 | | 0.11 | 0.002 | 0 | LAG | chr13-56074951-560 |
| 0 | 5.62867355044315 | | 0.179 | 0.004 | 0 | LAG | chr13-73703715-737 |
| 0 | 5.23718984371491 | | 0.113 | 0.003 | 0 | LAG | chr13-73706104-737 |
| 0 | 4.9308737067134 | | 0.137 | 0.004 | 0 | LAG | chr13-74137140-741 |
| 0 | 4.8337787284352 | | 0.158 | 0.007 | 0 | LAG | chr13-83696754-836 |
| 0 | 5.80422408664321 | | 0.149 | 0.003 | 0 | LAG | chr14-32150771-321 |
| 0 | 6.19636565519927 | | 0.151 | 0.002 | 0 | LAG | chr14-71626865-716 |
| 0 | 5.31376487521255 | | 0.12 | 0.003 | 0 | LAG | chr14-97771218-977 |
| 0 | 5.50772420943661 | | 0.115 | 0.003 | 0 | LAG | chr15-46690434-466 |
| 0 | 5.84801498190292 | | 0.286 | 0.006 | 0 | LAG | chr15-78277904-782 |
| 0 | 5.23774138702183 | | 0.318 | 0.01 | 0 | LAG | chr15-78279965-782 |
| 0 | 6.15576328501665 | | 0.325 | 0.006 | 0 | LAG | chr15-78294371-782 |
| 0 | 5.98454229707949 | | 0.143 | 0.003 | 0 | LAG | chr16-8595463-8595 |

| | | | | | | |
|---|---|---|---|---|---|---|
| 0 | 5.06853144115423 | 0.154 | 0.005 | 0 | LAG | chr16-8598066-8598 |
| 0 | 5.29177040590368 | 0.311 | 0.01 | 0 | LAG | chr16-8599162-8599 |
| 0 | 5.21596786360015 | 0.137 | 0.004 | 0 | LAG | chr16-24954733-249 |
| 0 | 5.05162906017822 | 0.158 | 0.005 | 0 | LAG | chr16-46146532-461 |
| 0 | 5.00969329057988 | 0.199 | 0.007 | 0 | LAG | chr17-23890829-238 |
| 0 | 5.68074570250292 | 0.102 | 0.002 | 0 | LAG | chr17-40581912-405 |
| 0 | 5.92858812196431 | 0.104 | 0.002 | 0 | LAG | chr17-46994044-469 |
| 0 | 5.06003994679773 | 0.13 | 0.004 | 0 | LAG | chr17-47131746-471 |
| 0 | 5.30330458770172 | 0.144 | 0.004 | 0 | LAG | chr17-51324928-513 |
| 0 | 6.50593397569753 | 0.109 | 0.001 | 0 | LAG | chr17-64654020-646 |
| 0 | 5.45707253699953 | 0.242 | 0.005 | 0 | LAG | chr17-67684096-676 |
| 0 | 5.70453349340133 | 0.211 | 0.005 | 0 | LAG | chr17-74617138-746 |
| 0 | 5.57529546829043 | 0.118 | 0.003 | 0 | LAG | chr18-30825149-308 |
| 0 | 5.02113413054107 | 0.166 | 0.005 | 0 | LAG | chr18-50657065-506 |
| 0 | 4.89427093688167 | 0.21 | 0.007 | 0 | LAG | chr18-60017214-600 |
| 0 | 5.76264619284144 | 0.104 | 0.002 | 0 | LAG | chr18-82406654-824 |
| 0 | 5.42303768771221 | 0.124 | 0.003 | 0 | LAG | chr18-82424902-824 |
| 0 | 4.94699102394531 | 0.166 | 0.006 | 0 | LAG | chr19-10269750-102 |
| 0 | 5.10655080687391 | 0.154 | 0.005 | 0 | LAG | chr19-23933592-239 |
| 0 | 5.14629850218248 | 0.135 | 0.004 | 0 | LAG | chr2-28528632-2852 |
| 0 | 6.11811265750002 | 0.25 | 0.005 | 0 | LAG | chr2-30605629-3060 |
| 0 | 4.98309164789638 | 0.197 | 0.007 | 0 | LAG | chr2-30699137-3069 |
| 0 | 5.13883077586324 | 0.135 | 0.004 | 0 | LAG | chr2-39143364-3914 |
| 0 | 5.14614264622107 | 0.185 | 0.006 | 0 | LAG | chr2-48917816-4891 |
| 0 | 5.01496665228935 | 0.155 | 0.005 | 0 | LAG | chr2-58417320-5841 |
| 0 | 5.37535223205677 | 0.116 | 0.003 | 0 | LAG | chr2-58422001-5842 |
| 0 | 4.92661264648587 | 0.169 | 0.006 | 0 | LAG | chr2-136523004-136 |
| 0 | 5.38385870902089 | 0.129 | 0.003 | 0 | LAG | chr2-188657509-188 |
| 0 | 4.81720437257769 | 0.154 | 0.006 | 0 | LAG | chr2-203642155-203 |
| 0 | 4.9125765601494 | 0.213 | 0.008 | 0 | LAG | chr20-4561230-4561 |
| 0 | 5.11302510458735 | 0.349 | 0.012 | 0 | LAG | chr20-4569119-4569 |
| 0 | 5.16288095138478 | 0.11 | 0.003 | 0 | LAG | chr20-36941484-369 |
| 0 | 5.14226783188507 | 0.104 | 0.003 | 0 | LAG | chr20-45925159-459 |
| 0 | 5.27413277439253 | 0.113 | 0.003 | 0 | LAG | chr3-21480156-2148 |
| 0 | 5.25689965914001 | 0.343 | 0.012 | 0 | LAG | chr3-41903984-4190 |
| 0 | 5.18764854857746 | 0.144 | 0.004 | 0 | LAG | chr3-91071087-9107 |
| 0 | 4.97517994463369 | 0.12 | 0.004 | 0 | LAG | chr3-99048772-9904 |
| 0 | 5.15720743458212 | 0.121 | 0.003 | 0 | LAG | chr3-141142000-141 |
| 0 | 5.72231210960272 | 0.109 | 0.002 | 0 | LAG | chr4-11336856-1133 |
| 0 | 5.18180011759896 | 0.129 | 0.004 | 0 | LAG | chr4-90613653-9061 |
| 0 | 6.03806678832442 | 0.236 | 0.004 | 0 | LAG | chr4-115434205-115 |
| 0 | 5.14460321120977 | 0.171 | 0.006 | 0 | LAG | chr4-141384296-141 |
| 0 | 5.69062989609761 | 0.144 | 0.003 | 0 | LAG | chr4-167167722-167 |

| | | | | | | |
|---|---|---|---|---|---|---|
| 0 | 4.83829857946149 | 0.182 | 0.007 | 0 | LAG | chr5-49916055-4991 |
| 0 | 5.86940925698583 | 0.126 | 0.002 | 0 | LAG | chr5-58858179-5885 |
| 0 | 5.48489030591804 | 0.225 | 0.005 | 0 | LAG | chr5-62051618-6205 |
| 0 | 5.13595954435486 | 0.357 | 0.011 | 0 | LAG | chr5-62053060-6205 |
| 0 | 4.94043518535059 | 0.388 | 0.015 | 0 | LAG | chr5-62055071-6205 |
| 0 | 5.80985345882897 | 0.202 | 0.005 | 0 | LAG | chr5-62062362-6206 |
| 0 | 5.4033245558537 | 0.213 | 0.005 | 0 | LAG | chr5-62114445-6211 |
| 0 | 4.95327388691591 | 0.36 | 0.014 | 0 | LAG | chr5-62134816-6213 |
| 0 | 6.40188941308272 | 0.104 | 0.001 | 0 | LAG | chr5-62146197-6214 |
| 0 | 4.82390009811198 | 0.196 | 0.008 | 0 | LAG | chr5-62176599-6217 |
| 0 | 5.25800425223857 | 0.197 | 0.006 | 0 | LAG | chr5-74103808-7410 |
| 0 | 5.17868222477259 | 0.199 | 0.006 | 0 | LAG | chr5-132687687-132 |
| 0 | 5.40883142718368 | 0.179 | 0.005 | 0 | LAG | chr5-147911812-147 |
| 0 | 5.82277654275354 | 0.132 | 0.003 | 0 | LAG | chr5-160541989-160 |
| 0 | 6.05597920052734 | 0.16 | 0.003 | 0 | LAG | chr5-160542769-160 |
| 0 | 5.31224095111051 | 0.112 | 0.003 | 0 | LAG | chr6-11032983-1103 |
| 0 | 5.1909218936839 | 0.193 | 0.006 | 0 | LAG | chr6-20978938-2097 |
| 0 | 5.32734011404058 | 0.12 | 0.003 | 0 | LAG | chr6-22168128-2216 |
| 0 | 4.91891248227053 | 0.135 | 0.005 | 0 | LAG | chr6-48794967-4879 |
| 0 | 4.86474388557232 | 0.144 | 0.006 | 0 | LAG | chr7-33313878-3331 |
| 0 | 5.71180155964643 | 0.321 | 0.009 | 0 | LAG | chr7-33314610-3331 |
| 0 | 5.6238710394074 | 0.189 | 0.004 | 0 | LAG | chr7-39916726-3991 |
| 0 | 5.38403814251207 | 0.123 | 0.003 | 0 | LAG | chr7-69011494-6901 |
| 0 | 5.11940745463083 | 0.118 | 0.004 | 0 | LAG | chr7-73135914-7313 |
| 0 | 5.0557452167856 | 0.191 | 0.007 | 0 | LAG | chr7-91347244-9134 |
| 0 | 4.93063771994159 | 0.168 | 0.007 | 0 | LAG | chr7-92246463-9224 |
| 0 | 5.45390196441071 | 0.129 | 0.003 | 0 | LAG | chr7-121534232-121 |
| 0 | 5.15341318748226 | 0.127 | 0.004 | 0 | LAG | chr7-133370997-133 |
| 0 | 5.20404904979723 | 0.134 | 0.004 | 0 | LAG | chr8-23973561-2397 |
| 0 | 4.98715820032823 | 0.141 | 0.005 | 0 | LAG | chr9-14569521-1457 |
| 0 | 4.96695267620768 | 0.152 | 0.005 | 0 | LAG | chr9-15290061-1529 |
| 0 | 6.46643427864786 | 0.168 | 0.002 | 0 | LAG | chr9-39177483-3917 |
| 0 | 4.93267231016372 | 0.134 | 0.005 | 0 | LAG | chr9-39486927-3948 |
| 0 | 5.03432840735837 | 0.205 | 0.006 | 0 | LAG | chr9-60970351-6097 |
| 0 | 5.15970874698954 | 0.334 | 0.012 | 0 | LAG | chr9-60970934-6097 |
| 0 | 5.3815891175536 | 0.135 | 0.004 | 0 | LAG | chr9-81909720-8191 |
| 0 | 5.14777661805482 | 0.356 | 0.013 | 0 | LAG | chr9-89297441-8929 |
| 0 | 4.92737867211815 | 0.247 | 0.01 | 0 | LAG | chr9-96003890-9600 |
| 0 | 5.11848714695758 | 0.135 | 0.004 | 0 | LAG | chr9-107965647-107 |
| 0 | 5.92760583060312 | 0.109 | 0.002 | 0 | LAG | chrX-152072221-152 |
| 1.02280736 | 4.8530898512138 | 0.116 | 0.004 | 7.4947845 | LAG | chr5-62128308-6212 |
| 1.09278598 | 4.81576277421339 | 0.137 | 0.006 | 8.0075641 | LAG | chr9-39323878-3932 |
| 7.20308609 | 5.18982675498891 | 0.112 | 0.004 | 5.2781765 | LAG | chr20-41060107-410 |

| | | | | | | |
|---|---|---|---|---|---|---|
| 3.05584265 | 5.17057860098415 | 0.107 | 0.003 | 2.2392175 | LAG | chr13-73079561-730 |
| 2.14850633 | 4.92805717878009 | 0.129 | 0.005 | 1.5743523 | LAG | chr17-64160772-641 |
| 5.47561821 | 4.8963347088202 | 0.129 | 0.005 | 4.0123468 | LAG | chr10-39132155-391 |
| 1.79933710 | 4.90533838079059 | 0.106 | 0.003 | 1.3184930 | LAG | chr3-42977542-4297 |
| 5.65697746 | 4.97423672840443 | 0.115 | 0.004 | 4.1452407 | LAG | chr17-35035714-350 |
| 1.83285777 | 4.82659611033206 | 0.138 | 0.006 | 1.3430558 | LAG | chr3-54124678-5412 |
| 1.13734018 | 5.21799998026847 | 0.106 | 0.003 | 8.3340421 | LAG | chr3-168543784-168 |
| 1.68995752 | 4.84447733063751 | 0.113 | 0.004 | 1.2383434 | LAG | chr4-181908347-181 |
| 1.22884310 | 4.98671486860565 | 0.112 | 0.004 | 9.0045444 | LAG | chr9-43149681-4315 |
| 1.06114947 | 5.16668696545804 | 0.106 | 0.003 | 7.7757425 | LAG | chr4-64949047-6494 |
| 3.64796703 | 5.11534864162299 | 0.104 | 0.003 | 2.6731062 | LAG | chr1-121254697-121 |
| 2.55753639 | 4.82810573548783 | 0.109 | 0.004 | 1.8740757 | LAG | chr1-15197932-1519 |
| 9.91461558 | 4.81346723752365 | 0.127 | 0.005 | 7.2650932 | LAG | chr3-152752083-152 |
| 1.59940676 | 4.88707814534682 | 0.113 | 0.004 | 1.1719909 | LAG | chr4-167164301-167 |
| 1.52205688 | 4.94433504290739 | 0.106 | 0.004 | 1.1153115 | LAG | chr6-57310722-5731 |
| 2.41174850 | 4.97979722741663 | 0.101 | 0.003 | 1.7672473 | LAG | chr17-29710670-297 |
| 2.41647805 | 4.94308171308377 | 0.124 | 0.005 | 1.7707129 | LAG | chr1-43852826-4385 |
| 1.86095974 | 4.94125249285388 | 0.104 | 0.004 | 1.3636480 | LAG | chr10-97847235-978 |
| 1.37716819 | 4.99433859003004 | 0.115 | 0.005 | 1.0091420 | LAG | chr3-7651897-76523 |
| 1.21411328 | 4.82090847227106 | 0.107 | 0.004 | 8.8966093 | LAG | chr14-80650354-806 |
| 0 | 6.27044343759938 | 0.22 | 0.003 | 0 | MUS | chr1-16721936-1672 |
| 0 | 6.6125438801775 | 0.176 | 0.002 | 0 | MUS | chr1-21447344-2144 |
| 0 | 6.32055776670546 | 0.267 | 0.004 | 0 | MUS | chr1-24297978-2429 |
| 0 | 6.20098663119716 | 0.168 | 0.003 | 0 | MUS | chr1-52391485-5239 |
| 0 | 6.36972231447731 | 0.209 | 0.003 | 0 | MUS | chr1-92430387-9243 |
| 0 | 6.17207459764811 | 0.106 | 0.002 | 0 | MUS | chr1-94642930-9464 |
| 0 | 6.20923536632732 | 0.294 | 0.005 | 0 | MUS | chr1-100993894-100 |
| 0 | 6.48196588221375 | 0.301 | 0.004 | 0 | MUS | chr1-129822656-129 |
| 0 | 6.5231525559538 | 0.173 | 0.002 | 0 | MUS | chr1-135784065-135 |
| 0 | 6.76203094861284 | 0.173 | 0.002 | 0 | MUS | chr1-136991684-136 |
| 0 | 6.36751235616428 | 0.31 | 0.005 | 0 | MUS | chr1-144246278-144 |
| 0 | 6.45883102047207 | 0.176 | 0.003 | 0 | MUS | chr1-170006240-170 |
| 0 | 6.65688793077134 | 0.182 | 0.002 | 0 | MUS | chr1-173055944-173 |
| 0 | 6.37370913594462 | 0.163 | 0.003 | 0 | MUS | chr1-175892579-175 |
| 0 | 6.20193199334033 | 0.234 | 0.004 | 0 | MUS | chr1-190747693-190 |
| 0 | 6.37746114760903 | 0.195 | 0.003 | 0 | MUS | chr1-222585402-222 |
| 0 | 6.26837359609988 | 0.153 | 0.002 | 0 | MUS | chr1-231895717-231 |
| 0 | 6.52911101187949 | 0.129 | 0.001 | 0 | MUS | chr10-8472812-8473 |
| 0 | 6.26304947981541 | 0.145 | 0.002 | 0 | MUS | chr10-15971349-159 |
| 0 | 6.46108931844422 | 0.111 | 0.001 | 0 | MUS | chr10-40669756-406 |
| 0 | 6.20928594288162 | 0.108 | 0.002 | 0 | MUS | chr10-51769823-517 |
| 0 | 6.19706866716007 | 0.16 | 0.002 | 0 | MUS | chr11-3804916-3805 |
| 0 | 6.60756463416318 | 0.108 | 0.001 | 0 | MUS | chr11-60972613-609 |

| | | | | | | |
|---|---|---|---|---|---|---|
| 0 | 6.20422582856415 | 0.107 | 0.002 | 0 | MUS | chr11-71482936-714 |
| 0 | 6.24112762474589 | 0.356 | 0.006 | 0 | MUS | chr11-71525409-715 |
| 0 | 6.20302174495294 | 0.141 | 0.002 | 0 | MUS | chr11-75135762-751 |
| 0 | 6.68998105096645 | 0.128 | 0.001 | 0 | MUS | chr11-81194334-811 |
| 0 | 6.64081419799559 | 0.208 | 0.003 | 0 | MUS | chr12-829159-82966 |
| 0 | 6.18754175505104 | 0.146 | 0.002 | 0 | MUS | chr12-39486834-394 |
| 0 | 6.48498578645931 | 0.11 | 0.001 | 0 | MUS | chr13-55041500-550 |
| 0 | 6.21826373844021 | 0.176 | 0.003 | 0 | MUS | chr13-80910145-809 |
| 0 | 6.50573425315536 | 0.145 | 0.002 | 0 | MUS | chr13-81005776-810 |
| 0 | 6.51981472752674 | 0.119 | 0.002 | 0 | MUS | chr13-87538006-875 |
| 0 | 6.34889347082197 | 0.117 | 0.001 | 0 | MUS | chr13-99471559-994 |
| 0 | 6.28811096990302 | 0.21 | 0.003 | 0 | MUS | chr14-13185075-131 |
| 0 | 6.16514386392344 | 0.177 | 0.003 | 0 | MUS | chr14-13520722-135 |
| 0 | 6.32230868466113 | 0.12 | 0.002 | 0 | MUS | chr14-24817153-248 |
| 0 | 6.53738182194249 | 0.184 | 0.002 | 0 | MUS | chr14-28046041-280 |
| 0 | 6.34022060177941 | 0.124 | 0.002 | 0 | MUS | chr14-40816224-408 |
| 0 | 6.6871972490937 | 0.207 | 0.002 | 0 | MUS | chr14-53153714-531 |
| 0 | 6.23963692563613 | 0.117 | 0.002 | 0 | MUS | chr14-67979549-679 |
| 0 | 6.39271210524954 | 0.393 | 0.006 | 0 | MUS | chr14-86814173-868 |
| 0 | 6.26577308188552 | 0.238 | 0.004 | 0 | MUS | chr15-5430697-5431 |
| 0 | 6.52633553857487 | 0.288 | 0.003 | 0 | MUS | chr15-32666151-326 |
| 0 | 6.6564404104887 | 0.116 | 0.001 | 0 | MUS | chr15-34075754-340 |
| 0 | 6.52721083945538 | 0.43 | 0.006 | 0 | MUS | chr15-35276811-352 |
| 0 | 6.21100181359614 | 0.129 | 0.002 | 0 | MUS | chr15-48343298-483 |
| 0 | 6.32549203334796 | 0.147 | 0.002 | 0 | MUS | chr15-65610324-656 |
| 0 | 6.28674939938802 | 0.246 | 0.004 | 0 | MUS | chr16-12795890-127 |
| 0 | 6.84996267731581 | 0.133 | 0.001 | 0 | MUS | chr16-29964318-299 |
| 0 | 6.21724420690236 | 0.182 | 0.003 | 0 | MUS | chr16-71220558-712 |
| 0 | 6.93689200525415 | 0.16 | 0.001 | 0 | MUS | chr16-73672322-736 |
| 0 | 6.19592719197496 | 0.15 | 0.002 | 0 | MUS | chr17-7435931-7436 |
| 0 | 6.52359785391422 | 0.131 | 0.002 | 0 | MUS | chr17-31624470-316 |
| 0 | 6.26292065771917 | 0.126 | 0.002 | 0 | MUS | chr17-33837954-338 |
| 0 | 6.25334659197163 | 0.199 | 0.003 | 0 | MUS | chr17-58169002-581 |
| 0 | 6.44784344437145 | 0.117 | 0.001 | 0 | MUS | chr17-60113814-601 |
| 0 | 6.18337075935968 | 0.151 | 0.003 | 0 | MUS | chr17-66975470-669 |
| 0 | 6.18917210117307 | 0.239 | 0.003 | 0 | MUS | chr17-71417325-714 |
| 0 | 6.33322211074704 | 0.171 | 0.003 | 0 | MUS | chr17-82217842-822 |
| 0 | 6.3563017294045 | 0.154 | 0.002 | 0 | MUS | chr17-84427077-844 |
| 0 | 6.16678868875618 | 0.116 | 0.002 | 0 | MUS | chr18-2000145-2000 |
| 0 | 6.16844916241611 | 0.146 | 0.003 | 0 | MUS | chr18-4289828-4290 |
| 0 | 6.22767448288254 | 0.115 | 0.002 | 0 | MUS | chr18-14008550-140 |
| 0 | 6.37020877980084 | 0.253 | 0.004 | 0 | MUS | chr18-27269679-272 |
| 0 | 6.77741491606518 | 0.157 | 0.002 | 0 | MUS | chr18-28883622-288 |

| | | | | | | |
|---|---|---|---|---|---|---|
| 0 | 6.21082869013252 | 0.199 | 0.003 | 0 | MUS | chr18-55282896-552 |
| 0 | 6.25469108528917 | 0.1 | 0.001 | 0 | MUS | chr18-82185280-821 |
| 0 | 6.5173546117943 | 0.267 | 0.004 | 0 | MUS | chr19-26191358-261 |
| 0 | 6.48017590682614 | 0.167 | 0.002 | 0 | MUS | chr19-30384753-303 |
| 0 | 6.47799840343194 | 0.276 | 0.004 | 0 | MUS | chr19-46670442-466 |
| 0 | 6.31726798267896 | 0.1 | 0.001 | 0 | MUS | chr2-13632919-1363 |
| 0 | 6.46181404920256 | 0.239 | 0.003 | 0 | MUS | chr2-23551061-2355 |
| 0 | 6.19441749073554 | 0.196 | 0.003 | 0 | MUS | chr2-39167084-3916 |
| 0 | 6.22151682502585 | 0.2 | 0.004 | 0 | MUS | chr2-78719948-7872 |
| 0 | 6.48257400398261 | 0.122 | 0.001 | 0 | MUS | chr2-91487250-9148 |
| 0 | 6.25332127710036 | 0.111 | 0.002 | 0 | MUS | chr2-93304285-9330 |
| 0 | 6.47221783035629 | 0.273 | 0.004 | 0 | MUS | chr2-102099847-102 |
| 0 | 6.34202988318369 | 0.166 | 0.002 | 0 | MUS | chr2-125249373-125 |
| 0 | 6.42526048385852 | 0.144 | 0.002 | 0 | MUS | chr2-128496006-128 |
| 0 | 6.48503778770746 | 0.19 | 0.003 | 0 | MUS | chr2-142238245-142 |
| 0 | 6.34966791282175 | 0.147 | 0.002 | 0 | MUS | chr2-217489507-217 |
| 0 | 6.45650363468918 | 0.133 | 0.002 | 0 | MUS | chr2-226566671-226 |
| 0 | 6.28061261766621 | 0.168 | 0.002 | 0 | MUS | chr2-230595492-230 |
| 0 | 6.40333035199516 | 0.242 | 0.003 | 0 | MUS | chr20-21780219-217 |
| 0 | 6.64599791034639 | 0.122 | 0.002 | 0 | MUS | chr20-31138136-311 |
| 0 | 6.68571146878098 | 0.265 | 0.003 | 0 | MUS | chr20-31216430-312 |
| 0 | 6.47592203772266 | 0.205 | 0.003 | 0 | MUS | chr3-27847926-2784 |
| 0 | 6.32733322725359 | 0.134 | 0.002 | 0 | MUS | chr3-38326985-3832 |
| 0 | 6.2796043764194 | 0.217 | 0.003 | 0 | MUS | chr3-61926757-6192 |
| 0 | 6.35389998541148 | 0.247 | 0.004 | 0 | MUS | chr3-62049123-6204 |
| 0 | 6.81070554651217 | 0.204 | 0.002 | 0 | MUS | chr3-63201262-6320 |
| 0 | 6.19152485759192 | 0.136 | 0.002 | 0 | MUS | chr3-63957162-6395 |
| 0 | 6.16703050452699 | 0.169 | 0.003 | 0 | MUS | chr3-90494303-9049 |
| 0 | 6.41916965549183 | 0.214 | 0.003 | 0 | MUS | chr3-99884404-9988 |
| 0 | 6.29311161174449 | 0.312 | 0.004 | 0 | MUS | chr3-100736484-100 |
| 0 | 6.25160333181397 | 0.227 | 0.004 | 0 | MUS | chr3-128642423-128 |
| 0 | 6.63732674107953 | 0.215 | 0.002 | 0 | MUS | chr3-132933479-132 |
| 0 | 6.16517402440316 | 0.337 | 0.005 | 0 | MUS | chr3-132951394-132 |
| 0 | 6.67193227323678 | 0.331 | 0.004 | 0 | MUS | chr3-163056082-163 |
| 0 | 6.4158610595203 | 0.148 | 0.002 | 0 | MUS | chr4-14987418-1498 |
| 0 | 6.23582491541878 | 0.171 | 0.003 | 0 | MUS | chr4-30078791-3007 |
| 0 | 6.19888631633213 | 0.242 | 0.004 | 0 | MUS | chr4-32953954-3295 |
| 0 | 6.21912522916028 | 0.157 | 0.002 | 0 | MUS | chr4-36102563-3610 |
| 0 | 6.2338062776558 | 0.163 | 0.003 | 0 | MUS | chr4-53069199-5306 |
| 0 | 6.1835213396447 | 0.115 | 0.002 | 0 | MUS | chr4-71583657-7158 |
| 0 | 6.17824031037043 | 0.157 | 0.003 | 0 | MUS | chr4-83526016-8352 |
| 0 | 6.25407756195073 | 0.128 | 0.002 | 0 | MUS | chr4-103253056-103 |
| 0 | 6.17428716444354 | 0.165 | 0.003 | 0 | MUS | chr5-21440950-2144 |

| | | | | | | |
|---|---|---|---|---|---|---|
| 0 | 6.23589605963727 | 0.111 | 0.002 | 0 | MUS | chr5-25027450-2502 |
| 0 | 6.2529843685753 | 0.301 | 0.005 | 0 | MUS | chr5-117209515-117 |
| 0 | 6.37985656514858 | 0.18 | 0.003 | 0 | MUS | chr5-161964907-161 |
| 0 | 6.27152239552164 | 0.118 | 0.002 | 0 | MUS | chr6-23435689-2343 |
| 0 | 6.1756076954017 | 0.125 | 0.002 | 0 | MUS | chr6-25636358-2563 |
| 0 | 6.2120431648696 | 0.18 | 0.003 | 0 | MUS | chr6-37684727-3768 |
| 0 | 6.22907810851133 | 0.157 | 0.003 | 0 | MUS | chr6-45327767-4532 |
| 0 | 6.34865159670157 | 0.106 | 0.001 | 0 | MUS | chr6-84630804-8463 |
| 0 | 6.70921083834421 | 0.174 | 0.002 | 0 | MUS | chr6-91370723-9137 |
| 0 | 6.16042007063973 | 0.218 | 0.003 | 0 | MUS | chr6-128464070-128 |
| 0 | 6.17583797393524 | 0.202 | 0.003 | 0 | MUS | chr6-137333811-137 |
| 0 | 6.55339082863358 | 0.293 | 0.003 | 0 | MUS | chr7-7078680-70791 |
| 0 | 6.39614693452029 | 0.163 | 0.002 | 0 | MUS | chr7-54745282-5474 |
| 0 | 6.30599310311026 | 0.343 | 0.005 | 0 | MUS | chr7-56649445-5664 |
| 0 | 6.27743347674469 | 0.141 | 0.002 | 0 | MUS | chr7-72644032-7264 |
| 0 | 6.20385075135727 | 0.179 | 0.002 | 0 | MUS | chr7-84056706-8405 |
| 0 | 6.23961952614717 | 0.184 | 0.003 | 0 | MUS | chr7-94491401-9449 |
| 0 | 6.21739922848468 | 0.148 | 0.002 | 0 | MUS | chr7-99170583-9917 |
| 0 | 6.4553316901801 | 0.192 | 0.002 | 0 | MUS | chr7-115108773-115 |
| 0 | 6.4934052746052 | 0.22 | 0.003 | 0 | MUS | chr7-118296244-118 |
| 0 | 6.53654954929244 | 0.141 | 0.002 | 0 | MUS | chr7-118987698-118 |
| 0 | 6.37011597086455 | 0.16 | 0.002 | 0 | MUS | chr8-40506056-4050 |
| 0 | 6.39967725443119 | 0.114 | 0.001 | 0 | MUS | chr8-54245423-5424 |
| 0 | 6.39039015784201 | 0.292 | 0.004 | 0 | MUS | chr8-112305766-112 |
| 0 | 6.80558241577763 | 0.163 | 0.002 | 0 | MUS | chr8-112310000-112 |
| 0 | 6.39798485255951 | 0.237 | 0.003 | 0 | MUS | chr9-23060094-2306 |
| 0 | 6.19385360670038 | 0.197 | 0.003 | 0 | MUS | chr9-23073684-2307 |
| 0 | 6.54386631556955 | 0.313 | 0.004 | 0 | MUS | chr9-23082924-2308 |
| 0 | 6.76048879901626 | 0.351 | 0.004 | 0 | MUS | chr9-23085440-2308 |
| 0 | 6.55370190751571 | 0.168 | 0.003 | 0 | MUS | chr9-28019377-2801 |
| 0 | 6.39770895908916 | 0.117 | 0.002 | 0 | MUS | chr9-34763040-3476 |
| 0 | 6.42889202524714 | 0.217 | 0.004 | 0 | MUS | chr9-56698598-5669 |
| 0 | 6.37666143509968 | 0.137 | 0.002 | 0 | MUS | chr9-68462388-6846 |
| 0 | 6.2468116305443 | 0.121 | 0.002 | 0 | MUS | chr9-79522873-7952 |
| 0 | 6.26536591644428 | 0.105 | 0.001 | 0 | MUS | chrX-71656113-7165 |
| 0 | 6.44024285137772 | 0.134 | 0.001 | 0 | MUS | chrX-74551424-7455 |
| 0 | 7.45583022642651 | 0.12 | 0.001 | 0 | MUS | chrX-98333117-9833 |
| 0 | 6.39385656997408 | 0.118 | 0.001 | 0 | MUS | chrX-100029964-100 |
| 0 | 6.22099427997985 | 0.148 | 0.002 | 0 | MUS | chrX-107422288-107 |
| 0 | 6.67021220123258 | 0.199 | 0.002 | 0 | MUS | chrX-123974426-123 |
| 0 | 6.31787254632369 | 0.169 | 0.002 | 0 | MUS | chrX-123986152-123 |
| 0 | 3.71768896456221 | 0.209 | 0.017 | 0 | MYE | chr1-8243910-82444 |
| 0 | 3.66182211941734 | 0.144 | 0.013 | 0 | MYE | chr1-18490719-1849 |

| | | | | |
|---|---|---|---|---|
| 0 3.62341363861956 | 0.16 | 0.015 | 0 MYE | chr1-20128141-2012 |
| 0 3.67659979313006 | 0.102 | 0.008 | 0 MYE | chr1-69976120-6997 |
| 0 3.81607889941637 | 0.276 | 0.022 | 0 MYE | chr1-76957847-7695 |
| 0 3.68878780991841 | 0.173 | 0.015 | 0 MYE | chr1-93940162-9394 |
| 0 3.62284509383926 | 0.268 | 0.024 | 0 MYE | chr1-134093168-134 |
| 0 3.74021859356803 | 0.144 | 0.012 | 0 MYE | chr1-159966571-159 |
| 0 3.74397300756856 | 0.107 | 0.008 | 0 MYE | chr1-164961008-164 |
| 0 3.60255327510088 | 0.117 | 0.011 | 0 MYE | chr1-170771927-170 |
| 0 3.60679222109443 | 0.114 | 0.01 | 0 MYE | chr1-181911909-181 |
| 0 3.60733647910947 | 0.127 | 0.011 | 0 MYE | chr1-190471888-190 |
| 0 3.60350115727677 | 0.104 | 0.009 | 0 MYE | chr1-207971751-207 |
| 0 3.61114056996401 | 0.194 | 0.016 | 0 MYE | chr1-209433450-209 |
| 0 3.90762075909901 | 0.125 | 0.01 | 0 MYE | chr1-232199338-232 |
| 0 3.62864887775908 | 0.105 | 0.009 | 0 MYE | chr1-239820303-239 |
| 0 3.60942813103558 | 0.116 | 0.01 | 0 MYE | chr10-15854963-158 |
| 0 3.70219420820195 | 0.174 | 0.015 | 0 MYE | chr10-47157348-471 |
| 0 3.68656354672248 | 0.295 | 0.024 | 0 MYE | chr10-47712655-477 |
| 0 3.6295477357995 | 0.141 | 0.013 | 0 MYE | chr10-47727779-477 |
| 0 3.6535892958321 | 0.15 | 0.013 | 0 MYE | chr10-61304004-613 |
| 0 3.66489454858967 | 0.112 | 0.01 | 0 MYE | chr10-67069758-670 |
| 0 3.67884557916724 | 0.383 | 0.033 | 0 MYE | chr10-86920898-869 |
| 0 3.84295867694462 | 0.114 | 0.009 | 0 MYE | chr10-96199289-961 |
| 0 3.63828124553862 | 0.159 | 0.013 | 0 MYE | chr10-100150487-10 |
| 0 3.84312885231342 | 0.227 | 0.017 | 0 MYE | chr10-100264188-10 |
| 0 3.6025730244912 | 0.209 | 0.017 | 0 MYE | chr11-14586422-145 |
| 0 3.82457253677562 | 0.244 | 0.021 | 0 MYE | chr11-56729677-567 |
| 0 3.78449196693753 | 0.146 | 0.011 | 0 MYE | chr11-77594962-775 |
| 0 3.9324339517661 | 0.109 | 0.008 | 0 MYE | chr12-15617483-156 |
| 0 3.59706345677258 | 0.392 | 0.036 | 0 MYE | chr12-32935473-329 |
| 0 3.64611528399422 | 0.111 | 0.009 | 0 MYE | chr12-33926781-339 |
| 0 3.80317908409023 | 0.273 | 0.022 | 0 MYE | chr13-31192366-311 |
| 0 3.76824406162392 | 0.107 | 0.008 | 0 MYE | chr13-32653707-326 |
| 0 3.89466991136466 | 0.1 | 0.008 | 0 MYE | chr13-42401522-424 |
| 0 3.67159441925241 | 0.145 | 0.013 | 0 MYE | chr13-43118503-431 |
| 0 3.65412876937158 | 0.105 | 0.008 | 0 MYE | chr13-51109244-511 |
| 0 3.86119129694529 | 0.141 | 0.011 | 0 MYE | chr13-62001293-620 |
| 0 3.61774142388643 | 0.164 | 0.014 | 0 MYE | chr13-64647282-646 |
| 0 3.78006858540743 | 0.106 | 0.008 | 0 MYE | chr13-65648837-656 |
| 0 3.6148305706099 | 0.114 | 0.009 | 0 MYE | chr13-66160678-661 |
| 0 3.64410680504957 | 0.186 | 0.018 | 0 MYE | chr13-73274144-732 |
| 0 3.72217517587433 | 0.129 | 0.012 | 0 MYE | chr13-76756144-767 |
| 0 3.58895940665852 | 0.102 | 0.009 | 0 MYE | chr13-83249523-832 |
| 0 3.58932737259612 | 0.137 | 0.012 | 0 MYE | chr13-96173677-961 |

| | | | | | | |
|---|---|---|---|---|---|---|
| 0 | 3.87476430169065 | 0.111 | 0.008 | 0 | MYE | chr14-34575404-345 |
| 0 | 4.05846633250717 | 0.1 | 0.007 | 0 | MYE | chr14-57523109-575 |
| 0 | 3.64587513307745 | 0.15 | 0.012 | 0 | MYE | chr14-67122263-671 |
| 0 | 3.69169532632189 | 0.232 | 0.019 | 0 | MYE | chr14-83340475-833 |
| 0 | 3.74370178809566 | 0.255 | 0.02 | 0 | MYE | chr14-86299787-863 |
| 0 | 3.83905815717031 | 0.159 | 0.012 | 0 | MYE | chr15-24216004-242 |
| 0 | 3.84001973068747 | 0.125 | 0.009 | 0 | MYE | chr16-8497402-8497 |
| 0 | 3.62505873467756 | 0.15 | 0.013 | 0 | MYE | chr16-16798257-167 |
| 0 | 3.64259288195625 | 0.117 | 0.01 | 0 | MYE | chr16-20810457-208 |
| 0 | 3.67917384305131 | 0.131 | 0.01 | 0 | MYE | chr16-51049030-510 |
| 0 | 3.72810986679694 | 0.129 | 0.01 | 0 | MYE | chr16-79497656-794 |
| 0 | 3.63874293287282 | 0.265 | 0.022 | 0 | MYE | chr17-22537938-225 |
| 0 | 3.62382500301413 | 0.265 | 0.023 | 0 | MYE | chr17-27149663-271 |
| 0 | 3.65271503461889 | 0.275 | 0.023 | 0 | MYE | chr17-64632126-646 |
| 0 | 3.69625383884759 | 0.122 | 0.01 | 0 | MYE | chr17-67313984-673 |
| 0 | 3.69425984620648 | 0.122 | 0.01 | 0 | MYE | chr17-77226778-772 |
| 0 | 3.87939047519018 | 0.102 | 0.007 | 0 | MYE | chr17-77228707-772 |
| 0 | 3.60110622928517 | 0.106 | 0.009 | 0 | MYE | chr17-86031940-860 |
| 0 | 3.68696857805621 | 0.154 | 0.013 | 0 | MYE | chr17-86033397-860 |
| 0 | 3.74440559609237 | 0.132 | 0.011 | 0 | MYE | chr18-54563538-545 |
| 0 | 3.64166530470258 | 0.145 | 0.012 | 0 | MYE | chr18-54620716-546 |
| 0 | 3.64822569874142 | 0.113 | 0.009 | 0 | MYE | chr19-13428466-134 |
| 0 | 3.70026963355647 | 0.152 | 0.013 | 0 | MYE | chr19-13453896-134 |
| 0 | 3.9449361991889 | 0.149 | 0.012 | 0 | MYE | chr19-16121526-161 |
| 0 | 3.61044978657542 | 0.189 | 0.016 | 0 | MYE | chr19-23668713-236 |
| 0 | 3.66597393046181 | 0.11 | 0.009 | 0 | MYE | chr19-23670417-236 |
| 0 | 3.65843636716491 | 0.259 | 0.022 | 0 | MYE | chr19-24962526-249 |
| 0 | 3.65277420942723 | 0.103 | 0.008 | 0 | MYE | chr19-34053167-340 |
| 0 | 3.69396265383308 | 0.314 | 0.027 | 0 | MYE | chr19-42846578-428 |
| 0 | 3.77970653115275 | 0.224 | 0.018 | 0 | MYE | chr19-43397376-433 |
| 0 | 3.63748938909504 | 0.199 | 0.017 | 0 | MYE | chr19-50655764-506 |
| 0 | 3.62621131600646 | 0.254 | 0.021 | 0 | MYE | chr2-33121525-3312 |
| 0 | 3.70840547600115 | 0.205 | 0.017 | 0 | MYE | chr2-33856972-3385 |
| 0 | 3.70537030580782 | 0.16 | 0.013 | 0 | MYE | chr2-33857477-3385 |
| 0 | 3.66823320949245 | 0.11 | 0.009 | 0 | MYE | chr2-39552540-3955 |
| 0 | 3.70252858349473 | 0.119 | 0.01 | 0 | MYE | chr2-46960489-4696 |
| 0 | 3.61154638030192 | 0.104 | 0.009 | 0 | MYE | chr2-82342213-8234 |
| 0 | 3.71024239807937 | 0.225 | 0.018 | 0 | MYE | chr2-110190481-110 |
| 0 | 3.66760164869124 | 0.102 | 0.008 | 0 | MYE | chr2-123822427-123 |
| 0 | 3.81888603581225 | 0.115 | 0.008 | 0 | MYE | chr2-145475689-145 |
| 0 | 3.65893752709962 | 0.159 | 0.013 | 0 | MYE | chr2-164391012-164 |
| 0 | 3.63555407852592 | 0.29 | 0.027 | 0 | MYE | chr2-192220756-192 |
| 0 | 3.59275508357471 | 0.238 | 0.022 | 0 | MYE | chr2-231326647-231 |

| | | | | | | |
|---|---|---|---|---|---|---|
| 0 | 3.63535428956808 | 0.304 | 0.026 | 0 | MYE | chr3-29483750-2948 |
| 0 | 3.88989826498411 | 0.102 | 0.007 | 0 | MYE | chr3-62180129-6218 |
| 0 | 3.63654673498981 | 0.12 | 0.01 | 0 | MYE | chr3-113896927-113 |
| 0 | 3.65412166719864 | 0.279 | 0.024 | 0 | MYE | chr3-116726649-116 |
| 0 | 3.7125708983961 | 0.142 | 0.011 | 0 | MYE | chr3-116751457-116 |
| 0 | 3.73768419921441 | 0.131 | 0.011 | 0 | MYE | chr3-118309596-118 |
| 0 | 3.6571877430661 | 0.126 | 0.011 | 0 | MYE | chr3-140011088-140 |
| 0 | 3.59262234619667 | 0.199 | 0.017 | 0 | MYE | chr3-141664395-141 |
| 0 | 3.67989053867856 | 0.136 | 0.012 | 0 | MYE | chr3-148699829-148 |
| 0 | 3.73706640954443 | 0.206 | 0.017 | 0 | MYE | chr3-148709982-148 |
| 0 | 3.93812441031318 | 0.1 | 0.007 | 0 | MYE | chr3-148904708-148 |
| 0 | 3.83083427626246 | 0.242 | 0.019 | 0 | MYE | chr3-155509566-155 |
| 0 | 3.75661098210832 | 0.135 | 0.011 | 0 | MYE | chr4-45174253-4517 |
| 0 | 3.59635527954176 | 0.116 | 0.01 | 0 | MYE | chr4-45300861-4530 |
| 0 | 4.08060288079607 | 0.194 | 0.013 | 0 | MYE | chr4-130029943-130 |
| 0 | 3.68756455682365 | 0.13 | 0.012 | 0 | MYE | chr4-146523855-146 |
| 0 | 3.63983046068224 | 0.203 | 0.018 | 0 | MYE | chr4-155828335-155 |
| 0 | 3.64494944662914 | 0.103 | 0.008 | 0 | MYE | chr4-155866802-155 |
| 0 | 3.60644690132524 | 0.152 | 0.014 | 0 | MYE | chr4-155882842-155 |
| 0 | 3.73293327111621 | 0.108 | 0.009 | 0 | MYE | chr4-156072025-156 |
| 0 | 3.79333232208163 | 0.187 | 0.013 | 0 | MYE | chr4-156265136-156 |
| 0 | 3.76637453164007 | 0.142 | 0.01 | 0 | MYE | chr4-156357885-156 |
| 0 | 3.80343066603889 | 0.17 | 0.013 | 0 | MYE | chr4-169526209-169 |
| 0 | 3.62550774390752 | 0.252 | 0.023 | 0 | MYE | chr5-16723482-1672 |
| 0 | 3.78728857521375 | 0.113 | 0.008 | 0 | MYE | chr5-29577932-2957 |
| 0 | 3.61356072054796 | 0.358 | 0.031 | 0 | MYE | chr5-29597697-2959 |
| 0 | 3.71207033452686 | 0.142 | 0.012 | 0 | MYE | chr5-55264413-5526 |
| 0 | 3.64985231921251 | 0.128 | 0.011 | 0 | MYE | chr5-137271194-137 |
| 0 | 4.19863044859883 | 0.116 | 0.008 | 0 | MYE | chr5-149130736-149 |
| 0 | 3.71647117565414 | 0.232 | 0.018 | 0 | MYE | chr5-157126094-157 |
| 0 | 3.64454875924153 | 0.111 | 0.01 | 0 | MYE | chr6-13444131-1344 |
| 0 | 3.72561700627107 | 0.197 | 0.016 | 0 | MYE | chr6-35025379-3502 |
| 0 | 3.67346033680646 | 0.108 | 0.009 | 0 | MYE | chr6-51410054-5141 |
| 0 | 3.77408968323624 | 0.227 | 0.019 | 0 | MYE | chr6-55774563-5577 |
| 0 | 4.06341804739847 | 0.165 | 0.011 | 0 | MYE | chr6-56083619-5608 |
| 0 | 3.69763267585585 | 0.21 | 0.018 | 0 | MYE | chr6-56084136-5608 |
| 0 | 3.75630142365715 | 0.199 | 0.016 | 0 | MYE | chr6-104396704-104 |
| 0 | 3.72057226423176 | 0.168 | 0.013 | 0 | MYE | chr6-124835248-124 |
| 0 | 4.13555647546095 | 0.141 | 0.009 | 0 | MYE | chr6-138471522-138 |
| 0 | 3.59712851444434 | 0.216 | 0.02 | 0 | MYE | chr6-138476599-138 |
| 0 | 3.6139438865423 | 0.236 | 0.021 | 0 | MYE | chr7-28111411-2811 |
| 0 | 3.58986223450829 | 0.134 | 0.012 | 0 | MYE | chr7-82082570-8208 |
| 0 | 3.69343973902555 | 0.142 | 0.012 | 0 | MYE | chr7-91077657-9107 |

| | | | | | | |
|---|---|---|---|---|---|---|
| 0 | 3.59928140551831 | 0.153 | 0.014 | 0 | MYE | chr7-95952095-9595 |
| 0 | 3.66692720618098 | 0.116 | 0.01 | 0 | MYE | chr7-99992303-9999 |
| 0 | 3.58736108112392 | 0.129 | 0.011 | 0 | MYE | chr7-131786882-131 |
| 0 | 3.58849691223056 | 0.13 | 0.012 | 0 | MYE | chr8-5780496-57809 |
| 0 | 3.63029430925742 | 0.17 | 0.015 | 0 | MYE | chr8-48680846-4868 |
| 0 | 3.62075049242081 | 0.102 | 0.008 | 0 | MYE | chr8-78605719-7860 |
| 0 | 3.63322381819865 | 0.216 | 0.018 | 0 | MYE | chr8-90816702-9081 |
| 0 | 3.73376169449101 | 0.228 | 0.018 | 0 | MYE | chr8-90818625-9081 |
| 0 | 3.81632774381046 | 0.28 | 0.023 | 0 | MYE | chr8-99743737-9974 |
| 0 | 3.76349031497612 | 0.197 | 0.019 | 0 | MYE | chr8-119681874-119 |
| 0 | 3.86025360654253 | 0.231 | 0.017 | 0 | MYE | chr8-123672356-123 |
| 0 | 3.65296703099247 | 0.165 | 0.014 | 0 | MYE | chr9-2229309-22298 |
| 0 | 4.00549798356845 | 0.105 | 0.008 | 0 | MYE | chr9-3671377-36718 |
| 0 | 3.6892659943721 | 0.151 | 0.012 | 0 | MYE | chr9-12692323-1269 |
| 0 | 3.60681966522887 | 0.12 | 0.01 | 0 | MYE | chr9-12692946-1269 |
| 0 | 3.66656005911735 | 0.258 | 0.022 | 0 | MYE | chr9-17398689-1739 |
| 0 | 3.63772859813517 | 0.189 | 0.016 | 0 | MYE | chr9-17402123-1740 |
| 0 | 3.73081561400839 | 0.189 | 0.015 | 0 | MYE | chr9-62068591-6206 |
| 0 | 3.94782686389355 | 0.186 | 0.013 | 0 | MYE | chr9-65765750-6576 |
| 0 | 6.1141447467601 | 0.232 | 0.005 | 0 | NK | chr1-2609947-26104 |
| 0 | 6.39481893278018 | 0.404 | 0.006 | 0 | NK | chr1-13646019-1364 |
| 0 | 6.21938766517511 | 0.577 | 0.011 | 0 | NK | chr1-14421569-1442 |
| 0 | 6.17522358157889 | 0.135 | 0.002 | 0 | NK | chr1-16040060-1604 |
| 0 | 6.53395963696459 | 0.286 | 0.004 | 0 | NK | chr1-43353940-4335 |
| 0 | 6.24373458157857 | 0.171 | 0.003 | 0 | NK | chr1-44124665-4412 |
| 0 | 6.3170565202439 | 0.122 | 0.001 | 0 | NK | chr1-69622362-6962 |
| 0 | 6.40332827458614 | 0.272 | 0.004 | 0 | NK | chr1-93841717-9384 |
| 0 | 6.41729126178043 | 0.368 | 0.004 | 0 | NK | chr1-93843990-9384 |
| 0 | 6.34282153021968 | 0.419 | 0.005 | 0 | NK | chr1-93844495-9384 |
| 0 | 6.13557305021422 | 0.246 | 0.004 | 0 | NK | chr1-94728248-9472 |
| 0 | 6.11907171136844 | 0.278 | 0.005 | 0 | NK | chr1-137784238-137 |
| 0 | 6.51150332355642 | 0.236 | 0.003 | 0 | NK | chr1-176849342-176 |
| 0 | 6.12689842294132 | 0.109 | 0.001 | 0 | NK | chr1-220664365-220 |
| 0 | 6.18409004949098 | 0.425 | 0.007 | 0 | NK | chr1-227206936-227 |
| 0 | 6.52307239421757 | 0.147 | 0.002 | 0 | NK | chr10-12200529-122 |
| 0 | 6.41074936139333 | 0.265 | 0.003 | 0 | NK | chr10-52219464-522 |
| 0 | 6.21360521836036 | 0.309 | 0.005 | 0 | NK | chr10-53142942-531 |
| 0 | 6.19244349858192 | 0.324 | 0.005 | 0 | NK | chr10-80672631-806 |
| 0 | 6.1345585849646 | 0.185 | 0.003 | 0 | NK | chr10-104756193-10 |
| 0 | 6.48067532351138 | 0.122 | 0.002 | 0 | NK | chr11-49053084-490 |
| 0 | 6.45534962123743 | 0.377 | 0.005 | 0 | NK | chr11-49097956-490 |
| 0 | 6.56567151484377 | 0.297 | 0.003 | 0 | NK | chr11-51047890-510 |
| 0 | 6.34428424119163 | 0.12 | 0.002 | 0 | NK | chr11-54829141-548 |

| | | | | | |
|---|---|---|---|---|---|
| 0 | 6.42674018838691 | 0.221 | 0.003 | 0 NK | chr11-67030272-670 |
| 0 | 6.61358457688213 | 0.124 | 0.001 | 0 NK | chr11-68032632-680 |
| 0 | 6.10885043997258 | 0.116 | 0.002 | 0 NK | chr11-68033446-680 |
| 0 | 6.27146153674079 | 0.35 | 0.005 | 0 NK | chr11-76616109-766 |
| 0 | 6.33266090858072 | 0.318 | 0.004 | 0 NK | chr12-15345643-153 |
| 0 | 6.51323698351933 | 0.263 | 0.003 | 0 NK | chr12-17008092-170 |
| 0 | 6.22677783623815 | 0.166 | 0.002 | 0 NK | chr12-17365200-173 |
| 0 | 6.29368161906592 | 0.335 | 0.005 | 0 NK | chr13-39030536-390 |
| 0 | 6.31663845721966 | 0.211 | 0.003 | 0 NK | chr13-40280999-402 |
| 0 | 6.29095698158093 | 0.27 | 0.004 | 0 NK | chr13-65773192-657 |
| 0 | 6.29571719375682 | 0.322 | 0.005 | 0 NK | chr13-74042699-740 |
| 0 | 6.17729924875398 | 0.179 | 0.002 | 0 NK | chr13-74153207-741 |
| 0 | 6.14458464623904 | 0.16 | 0.002 | 0 NK | chr14-32519823-325 |
| 0 | 6.45675454502643 | 0.278 | 0.003 | 0 NK | chr14-43817262-438 |
| 0 | 6.23265505351822 | 0.347 | 0.006 | 0 NK | chr14-74952392-749 |
| 0 | 6.16331158587981 | 0.183 | 0.003 | 0 NK | chr14-95228761-952 |
| 0 | 6.32439064707121 | 0.145 | 0.002 | 0 NK | chr15-29988886-299 |
| 0 | 6.62306245417113 | 0.242 | 0.002 | 0 NK | chr15-30364113-303 |
| 0 | 6.7932374127135 | 0.17 | 0.002 | 0 NK | chr15-30366604-303 |
| 0 | 6.11681315197405 | 0.28 | 0.004 | 0 NK | chr15-47989839-479 |
| 0 | 6.61129697293023 | 0.257 | 0.003 | 0 NK | chr16-6102521-6103 |
| 0 | 6.20284337769911 | 0.225 | 0.003 | 0 NK | chr16-6135130-6135 |
| 0 | 6.38860963993466 | 0.141 | 0.001 | 0 NK | chr16-67341396-673 |
| 0 | 6.12825952915914 | 0.11 | 0.002 | 0 NK | chr17-3837081-3837 |
| 0 | 6.21980699634287 | 0.126 | 0.002 | 0 NK | chr17-3892962-3893 |
| 0 | 6.2793232129683 | 0.314 | 0.004 | 0 NK | chr17-9181286-9181 |
| 0 | 6.30785290390232 | 0.211 | 0.003 | 0 NK | chr17-31370030-313 |
| 0 | 6.19144904443122 | 0.211 | 0.004 | 0 NK | chr17-45616563-456 |
| 0 | 6.28570041731598 | 0.12 | 0.002 | 0 NK | chr17-69447272-694 |
| 0 | 6.12035615454008 | 0.253 | 0.004 | 0 NK | chr17-81200150-812 |
| 0 | 6.50100762367924 | 0.15 | 0.002 | 0 NK | chr17-85889856-858 |
| 0 | 6.51897313833506 | 0.28 | 0.003 | 0 NK | chr17-85890876-858 |
| 0 | 6.46158873639474 | 0.23 | 0.003 | 0 NK | chr17-85892774-858 |
| 0 | 6.23122102464588 | 0.234 | 0.004 | 0 NK | chr18-54369648-543 |
| 0 | 6.23304758281172 | 0.211 | 0.003 | 0 NK | chr18-64136983-641 |
| 0 | 6.49666091080137 | 0.221 | 0.002 | 0 NK | chr18-82476922-824 |
| 0 | 6.48029068029304 | 0.16 | 0.002 | 0 NK | chr19-19302370-193 |
| 0 | 6.51704234824306 | 0.339 | 0.005 | 0 NK | chr19-48841554-488 |
| 0 | 6.13013835044544 | 0.274 | 0.004 | 0 NK | chr19-52324073-523 |
| 0 | 6.20009021072034 | 0.303 | 0.004 | 0 NK | chr19-56560342-565 |
| 0 | 6.60225960704875 | 0.349 | 0.004 | 0 NK | chr2-44675324-4467 |
| 0 | 6.10885043997258 | 0.105 | 0.001 | 0 NK | chr2-86511960-8651 |
| 0 | 6.34545839583196 | 0.107 | 0.001 | 0 NK | chr2-116692657-116 |

| | | | | | | |
|---|---|---|---|---|---|---|
| 0 | 6.20007347988854 | 0.152 | 0.002 | 0 | NK | chr2-169475745-169 |
| 0 | 6.30231072243925 | 0.505 | 0.008 | 0 | NK | chr2-173311735-173 |
| 0 | 6.33267434032834 | 0.303 | 0.004 | 0 | NK | chr2-188712839-188 |
| 0 | 6.40239289848822 | 0.377 | 0.005 | 0 | NK | chr2-193939474-193 |
| 0 | 6.15991201585617 | 0.371 | 0.006 | 0 | NK | chr2-242650006-242 |
| 0 | 6.34428424119163 | 0.122 | 0.002 | 0 | NK | chr2-243705930-243 |
| 0 | 6.46807264654824 | 0.335 | 0.004 | 0 | NK | chr20-29206789-292 |
| 0 | 6.61219810715319 | 0.322 | 0.004 | 0 | NK | chr20-33402297-334 |
| 0 | 6.5867093728414 | 0.177 | 0.002 | 0 | NK | chr20-33788823-337 |
| 0 | 6.64238302980774 | 0.156 | 0.002 | 0 | NK | chr3-28535139-2853 |
| 0 | 6.18966057158903 | 0.446 | 0.008 | 0 | NK | chr3-55484411-5548 |
| 0 | 6.32982978250369 | 0.501 | 0.007 | 0 | NK | chr3-55494382-5549 |
| 0 | 6.17541500462846 | 0.278 | 0.004 | 0 | NK | chr3-104207636-104 |
| 0 | 6.77719712210055 | 0.152 | 0.002 | 0 | NK | chr3-119961773-119 |
| 0 | 6.13314391282643 | 0.423 | 0.007 | 0 | NK | chr3-155444712-155 |
| 0 | 6.8549578518875 | 0.337 | 0.003 | 0 | NK | chr4-36487628-3648 |
| 0 | 6.30335350336608 | 0.255 | 0.004 | 0 | NK | chr4-36664996-3666 |
| 0 | 6.32166304487757 | 0.225 | 0.003 | 0 | NK | chr4-70403739-7040 |
| 0 | 6.64187568721706 | 0.251 | 0.003 | 0 | NK | chr4-70420973-7042 |
| 0 | 6.22662751177547 | 0.24 | 0.003 | 0 | NK | chr4-79476284-7947 |
| 0 | 6.57369355067872 | 0.345 | 0.004 | 0 | NK | chr4-96478955-9647 |
| 0 | 6.42460891405876 | 0.225 | 0.003 | 0 | NK | chr4-96541989-9654 |
| 0 | 6.16030503509124 | 0.204 | 0.003 | 0 | NK | chr4-116272702-116 |
| 0 | 6.69329099428521 | 0.297 | 0.003 | 0 | NK | chr4-141480317-141 |
| 0 | 6.26346674503054 | 0.213 | 0.003 | 0 | NK | chr4-146060766-146 |
| 0 | 6.46175711412027 | 0.318 | 0.004 | 0 | NK | chr4-146702083-146 |
| 0 | 6.35192509736229 | 0.253 | 0.004 | 0 | NK | chr4-151701250-151 |
| 0 | 6.36822419695595 | 0.192 | 0.003 | 0 | NK | chr4-161912296-161 |
| 0 | 6.34510787219272 | 0.168 | 0.002 | 0 | NK | chr4-162562173-162 |
| 0 | 6.17410872690234 | 0.457 | 0.008 | 0 | NK | chr4-162563533-162 |
| 0 | 6.14578167722374 | 0.128 | 0.002 | 0 | NK | chr4-163042427-163 |
| 0 | 6.30471077433597 | 0.137 | 0.002 | 0 | NK | chr4-163055613-163 |
| 0 | 6.55606581302988 | 0.253 | 0.003 | 0 | NK | chr4-163096408-163 |
| 0 | 6.27635492314574 | 0.171 | 0.002 | 0 | NK | chr4-163153863-163 |
| 0 | 6.68660244041497 | 0.219 | 0.002 | 0 | NK | chr4-163306827-163 |
| 0 | 6.30255328787792 | 0.17 | 0.002 | 0 | NK | chr4-164014911-164 |
| 0 | 6.50152857018908 | 0.2 | 0.002 | 0 | NK | chr4-164063614-164 |
| 0 | 6.6354755785748 | 0.25 | 0.002 | 0 | NK | chr4-164249614-164 |
| 0 | 6.63144444047911 | 0.116 | 0.001 | 0 | NK | chr4-164503934-164 |
| 0 | 6.27870889673816 | 0.318 | 0.004 | 0 | NK | chr4-165012914-165 |
| 0 | 6.14513449680675 | 0.156 | 0.002 | 0 | NK | chr5-5110079-51105 |
| 0 | 6.12069566868431 | 0.177 | 0.003 | 0 | NK | chr5-5162207-51627 |
| 0 | 6.30576286382713 | 0.21 | 0.003 | 0 | NK | chr5-6089238-60897 |

| | | | | | | |
|---|---|---|---|---|---|---|
| 0 | 6.21212245534599 | 0.17 | 0.002 | 0 | NK | chr5-47624193-4762 |
| 0 | 6.80855331225319 | 0.301 | 0.003 | 0 | NK | chr5-103002121-103 |
| 0 | 6.12283679283749 | 0.158 | 0.003 | 0 | NK | chr5-135130198-135 |
| 0 | 6.11410934828334 | 0.206 | 0.003 | 0 | NK | chr5-141902901-141 |
| 0 | 6.55824959029743 | 0.358 | 0.005 | 0 | NK | chr5-146420799-146 |
| 0 | 6.61876925451337 | 0.16 | 0.002 | 0 | NK | chr6-19944560-1994 |
| 0 | 6.26987611242492 | 0.32 | 0.005 | 0 | NK | chr6-57168441-5716 |
| 0 | 6.58320773859111 | 0.175 | 0.002 | 0 | NK | chr6-57185098-5718 |
| 0 | 6.33896376093336 | 0.183 | 0.002 | 0 | NK | chr6-121091942-121 |
| 0 | 6.16277485986835 | 0.282 | 0.004 | 0 | NK | chr6-139319759-139 |
| 0 | 6.16706435513985 | 0.278 | 0.005 | 0 | NK | chr7-9117336-91178 |
| 0 | 6.17162875554783 | 0.257 | 0.004 | 0 | NK | chr7-31068874-3106 |
| 0 | 6.23868531027571 | 0.223 | 0.003 | 0 | NK | chr7-53546742-5354 |
| 0 | 6.67540782608767 | 0.371 | 0.004 | 0 | NK | chr7-53903029-5390 |
| 0 | 6.34520674300634 | 0.381 | 0.006 | 0 | NK | chr7-57010093-5701 |
| 0 | 6.57657038470267 | 0.286 | 0.004 | 0 | NK | chr7-86984187-8698 |
| 0 | 6.17890702353141 | 0.269 | 0.004 | 0 | NK | chr7-87053171-8705 |
| 0 | 6.28327046945519 | 0.135 | 0.002 | 0 | NK | chr7-91582111-9158 |
| 0 | 6.49575670643033 | 0.246 | 0.003 | 0 | NK | chr7-91748142-9174 |
| 0 | 6.13301496627589 | 0.389 | 0.006 | 0 | NK | chr7-93802229-9380 |
| 0 | 6.61057393617105 | 0.375 | 0.004 | 0 | NK | chr7-110008146-110 |
| 0 | 6.78097996044474 | 0.267 | 0.003 | 0 | NK | chr7-131761276-131 |
| 0 | 6.55119743242266 | 0.225 | 0.003 | 0 | NK | chr8-31321171-3132 |
| 0 | 6.51323698351933 | 0.248 | 0.003 | 0 | NK | chr8-37350259-3735 |
| 0 | 6.34510787219272 | 0.183 | 0.002 | 0 | NK | chr8-75965399-7596 |
| 0 | 6.2077211315695 | 0.156 | 0.002 | 0 | NK | chr8-95861393-9586 |
| 0 | 6.61876925451337 | 0.139 | 0.002 | 0 | NK | chr8-105540294-105 |
| 0 | 6.72518413801515 | 0.28 | 0.003 | 0 | NK | chr8-110619356-110 |
| 0 | 6.16206206500753 | 0.297 | 0.004 | 0 | NK | chr8-118071979-118 |
| 0 | 6.36127783621361 | 0.171 | 0.002 | 0 | NK | chr8-123435574-123 |
| 0 | 6.54467658962291 | 0.149 | 0.002 | 0 | NK | chr9-4955032-49555 |
| 0 | 6.14679207746327 | 0.112 | 0.002 | 0 | NK | chr9-5104736-51052 |
| 0 | 6.15893606747902 | 0.32 | 0.005 | 0 | NK | chr9-15498472-1549 |
| 0 | 6.69371409098246 | 0.385 | 0.004 | 0 | NK | chr9-39777701-3977 |
| 0 | 6.17224902427633 | 0.156 | 0.002 | 0 | NK | chr9-40337659-4033 |
| 0 | 6.16226323679776 | 0.131 | 0.002 | 0 | NK | chr9-42729912-4273 |
| 0 | 6.48249431605373 | 0.25 | 0.003 | 0 | NK | chr9-45470403-4547 |
| 0 | 6.38428900818047 | 0.143 | 0.002 | 0 | NK | chr9-45490802-4549 |
| 0 | 6.39052891724963 | 0.314 | 0.004 | 0 | NK | chr9-94420800-9442 |
| 0 | 6.27191163529819 | 0.539 | 0.009 | 0 | NK | chr9-94445738-9444 |
| 0 | 5.64903376118374 | 0.149 | 0.003 | 0 | PER | chr1-34652643-3465 |
| 0 | 5.57663741987038 | 0.104 | 0.002 | 0 | PER | chr1-51832219-5183 |
| 0 | 4.98721140606872 | 0.113 | 0.004 | 0 | PER | chr1-133298450-133 |

| | | | | | |
|---|---|---|---|---|---|
| 0 | 5.95694098070172 | 0.331 | 0.006 | 0 PER | chr1-157132177-157 |
| 0 | 5.43217062357808 | 0.149 | 0.004 | 0 PER | chr1-165050601-165 |
| 0 | 5.90900358853233 | 0.229 | 0.004 | 0 PER | chr1-165051180-165 |
| 0 | 5.35801034805976 | 0.124 | 0.003 | 0 PER | chr1-165536882-165 |
| 0 | 5.29978469556034 | 0.203 | 0.005 | 0 PER | chr1-199531146-199 |
| 0 | 4.82592229189461 | 0.208 | 0.008 | 0 PER | chr1-199922094-199 |
| 0 | 5.33501557449052 | 0.121 | 0.003 | 0 PER | chr1-229537140-229 |
| 0 | 5.05503724323171 | 0.164 | 0.006 | 0 PER | chr1-245769806-245 |
| 0 | 5.02237851410981 | 0.256 | 0.009 | 0 PER | chr1-245770444-245 |
| 0 | 5.19337185958345 | 0.11 | 0.003 | 0 PER | chr1-253287832-253 |
| 0 | 4.90112418726611 | 0.157 | 0.006 | 0 PER | chr1-260108012-260 |
| 0 | 4.99248923632896 | 0.161 | 0.006 | 0 PER | chr10-13760364-137 |
| 0 | 5.15859601730564 | 0.153 | 0.005 | 0 PER | chr10-14132639-141 |
| 0 | 5.15888034675901 | 0.289 | 0.01 | 0 PER | chr10-14668349-146 |
| 0 | 4.94992609397925 | 0.123 | 0.005 | 0 PER | chr10-17446297-174 |
| 0 | 5.15726484331752 | 0.106 | 0.003 | 0 PER | chr10-61270185-612 |
| 0 | 5.09823634806158 | 0.218 | 0.009 | 0 PER | chr10-74443201-744 |
| 0 | 5.62900232537913 | 0.356 | 0.008 | 0 PER | chr10-87800369-878 |
| 0 | 5.35716748633889 | 0.258 | 0.007 | 0 PER | chr10-102064731-10 |
| 0 | 4.9531549590618 | 0.179 | 0.006 | 0 PER | chr10-103656155-10 |
| 0 | 4.9439764884654 | 0.146 | 0.005 | 0 PER | chr10-103656773-10 |
| 0 | 5.12685148980746 | 0.184 | 0.006 | 0 PER | chr11-66410541-664 |
| 0 | 4.93144149210117 | 0.1 | 0.004 | 0 PER | chr11-68887334-688 |
| 0 | 5.53013526339181 | 0.172 | 0.004 | 0 PER | chr11-77403051-774 |
| 0 | 5.01507459074744 | 0.169 | 0.005 | 0 PER | chr11-77403780-774 |
| 0 | 5.4878606533898 | 0.102 | 0.002 | 0 PER | chr12-1781922-1782 |
| 0 | 4.95055929772703 | 0.132 | 0.005 | 0 PER | chr12-27507189-275 |
| 0 | 5.39471060005882 | 0.249 | 0.007 | 0 PER | chr12-36462160-364 |
| 0 | 4.85056585682521 | 0.333 | 0.013 | 0 PER | chr12-38245155-382 |
| 0 | 5.34530153031996 | 0.25 | 0.007 | 0 PER | chr12-42082581-420 |
| 0 | 5.3696524778299 | 0.24 | 0.007 | 0 PER | chr13-44715038-447 |
| 0 | 5.60232475498404 | 0.287 | 0.006 | 0 PER | chr13-50301759-503 |
| 0 | 5.27492917571541 | 0.264 | 0.009 | 0 PER | chr13-55518129-555 |
| 0 | 4.98761820249816 | 0.101 | 0.003 | 0 PER | chr13-64420303-644 |
| 0 | 4.82975972570571 | 0.169 | 0.006 | 0 PER | chr13-64965981-649 |
| 0 | 5.02619075801725 | 0.119 | 0.004 | 0 PER | chr13-81877896-818 |
| 0 | 5.10951972939255 | 0.137 | 0.004 | 0 PER | chr13-105073768-10 |
| 0 | 4.93969069088983 | 0.146 | 0.005 | 0 PER | chr14-16384212-163 |
| 0 | 4.97909865165041 | 0.192 | 0.007 | 0 PER | chr14-35672530-356 |
| 0 | 5.25269967150341 | 0.134 | 0.003 | 0 PER | chr14-44125672-441 |
| 0 | 4.95684640354365 | 0.318 | 0.013 | 0 PER | chr14-80808041-808 |
| 0 | 4.83729890218746 | 0.136 | 0.005 | 0 PER | chr14-96750831-967 |
| 0 | 5.46564331896566 | 0.176 | 0.005 | 0 PER | chr15-7225431-7225 |

| | | | | | | |
|---|---|---|---|---|---|---|
| 0 | 5.05010688838274 | 0.142 | 0.005 | 0 | PER | chr15-21179426-211 |
| 0 | 4.98777583274238 | 0.106 | 0.003 | 0 | PER | chr15-40613248-406 |
| 0 | 5.12884112999117 | 0.104 | 0.003 | 0 | PER | chr15-52637767-526 |
| 0 | 4.8746453999349 | 0.154 | 0.005 | 0 | PER | chr15-97657467-976 |
| 0 | 4.98938748672525 | 0.223 | 0.008 | 0 | PER | chr16-57051699-570 |
| 0 | 5.21532980726433 | 0.277 | 0.008 | 0 | PER | chr16-57059058-570 |
| 0 | 4.99263625329122 | 0.12 | 0.004 | 0 | PER | chr16-57424641-574 |
| 0 | 5.40632749517774 | 0.116 | 0.003 | 0 | PER | chr16-64987876-649 |
| 0 | 4.84054180925148 | 0.13 | 0.005 | 0 | PER | chr16-66901182-669 |
| 0 | 5.08399857636682 | 0.164 | 0.006 | 0 | PER | chr16-66904310-669 |
| 0 | 5.06380377213811 | 0.175 | 0.006 | 0 | PER | chr16-71110357-711 |
| 0 | 5.0821429575992 | 0.127 | 0.004 | 0 | PER | chr16-71113822-711 |
| 0 | 4.83752394912932 | 0.129 | 0.005 | 0 | PER | chr16-77800005-778 |
| 0 | 6.56005413882217 | 0.215 | 0.002 | 0 | PER | chr17-10609587-106 |
| 0 | 5.78297495822134 | 0.181 | 0.004 | 0 | PER | chr17-10778257-107 |
| 0 | 5.03409474376902 | 0.281 | 0.009 | 0 | PER | chr17-10783289-107 |
| 0 | 5.65056912174233 | 0.151 | 0.003 | 0 | PER | chr17-31899230-318 |
| 0 | 5.16793213262841 | 0.132 | 0.004 | 0 | PER | chr17-34043098-340 |
| 0 | 5.35461421450818 | 0.134 | 0.003 | 0 | PER | chr17-34621960-346 |
| 0 | 5.66861586744466 | 0.125 | 0.003 | 0 | PER | chr17-63766896-637 |
| 0 | 5.34447897053527 | 0.233 | 0.006 | 0 | PER | chr18-29998395-299 |
| 0 | 4.8671955015517 | 0.188 | 0.008 | 0 | PER | chr18-33501071-335 |
| 0 | 4.84727249209141 | 0.278 | 0.011 | 0 | PER | chr18-54055073-540 |
| 0 | 5.03825840181213 | 0.227 | 0.009 | 0 | PER | chr18-54526310-545 |
| 0 | 5.03336839168323 | 0.2 | 0.007 | 0 | PER | chr18-55107643-551 |
| 0 | 4.8334014473386 | 0.125 | 0.004 | 0 | PER | chr18-59257641-592 |
| 0 | 5.08123265153218 | 0.203 | 0.006 | 0 | PER | chr19-15902014-159 |
| 0 | 5.04622920542117 | 0.116 | 0.004 | 0 | PER | chr19-16024176-160 |
| 0 | 5.06812937122195 | 0.11 | 0.003 | 0 | PER | chr19-37479656-374 |
| 0 | 5.23397287212277 | 0.134 | 0.003 | 0 | PER | chr19-38747686-387 |
| 0 | 5.6050714833482 | 0.116 | 0.003 | 0 | PER | chr19-41252656-412 |
| 0 | 5.60283635900188 | 0.112 | 0.002 | 0 | PER | chr19-44500124-445 |
| 0 | 5.15076488945378 | 0.25 | 0.008 | 0 | PER | chr19-48039186-480 |
| 0 | 4.85723999173859 | 0.113 | 0.004 | 0 | PER | chr19-48862906-488 |
| 0 | 5.25000906248188 | 0.2 | 0.006 | 0 | PER | chr2-12588619-1258 |
| 0 | 5.11211542227535 | 0.118 | 0.004 | 0 | PER | chr2-28311339-2831 |
| 0 | 5.89523979844735 | 0.136 | 0.003 | 0 | PER | chr2-46711378-4671 |
| 0 | 5.47986208586927 | 0.119 | 0.003 | 0 | PER | chr2-115019550-115 |
| 0 | 5.20470048297707 | 0.104 | 0.003 | 0 | PER | chr2-167690603-167 |
| 0 | 4.85157957826943 | 0.225 | 0.008 | 0 | PER | chr2-189758207-189 |
| 0 | 5.28391350290375 | 0.299 | 0.009 | 0 | PER | chr2-210961304-210 |
| 0 | 4.90064390595834 | 0.164 | 0.006 | 0 | PER | chr2-210968006-210 |
| 0 | 4.82903206821506 | 0.209 | 0.009 | 0 | PER | chr2-211264937-211 |

| | | | | | |
|---|---|---|---|---|---|
| 0 | 5.04684759739686 | 0.12 | 0.004 | 0 PER | chr2-221496583-221 |
| 0 | 4.91918535305407 | 0.262 | 0.01 | 0 PER | chr2-236043781-236 |
| 0 | 4.84498173273117 | 0.236 | 0.009 | 0 PER | chr20-13647825-136 |
| 0 | 5.43244749548512 | 0.197 | 0.005 | 0 PER | chr20-22234674-222 |
| 0 | 5.18705372248594 | 0.137 | 0.004 | 0 PER | chr20-48600533-486 |
| 0 | 4.99049318300611 | 0.191 | 0.006 | 0 PER | chr3-11271144-1127 |
| 0 | 4.89953544529613 | 0.222 | 0.008 | 0 PER | chr3-18632721-1863 |
| 0 | 5.52103123090019 | 0.286 | 0.007 | 0 PER | chr3-64813824-6481 |
| 0 | 5.43076395177997 | 0.165 | 0.004 | 0 PER | chr3-64886470-6488 |
| 0 | 5.01796074852502 | 0.181 | 0.006 | 0 PER | chr3-78453189-7845 |
| 0 | 5.10794624626035 | 0.158 | 0.005 | 0 PER | chr3-79845754-7984 |
| 0 | 4.92432739021909 | 0.257 | 0.01 | 0 PER | chr3-88398705-8839 |
| 0 | 5.20021768577003 | 0.224 | 0.007 | 0 PER | chr3-106359966-106 |
| 0 | 4.90278736193869 | 0.307 | 0.013 | 0 PER | chr3-112046477-112 |
| 0 | 4.86762928785904 | 0.114 | 0.004 | 0 PER | chr3-118895532-118 |
| 0 | 5.54232344303687 | 0.255 | 0.006 | 0 PER | chr3-131191295-131 |
| 0 | 5.18797020608675 | 0.123 | 0.003 | 0 PER | chr3-152213587-152 |
| 0 | 4.8532652659084 | 0.141 | 0.006 | 0 PER | chr3-152846565-152 |
| 0 | 4.86613760142597 | 0.223 | 0.008 | 0 PER | chr4-10609230-1060 |
| 0 | 5.23000111471882 | 0.223 | 0.007 | 0 PER | chr4-84281158-8428 |
| 0 | 5.08389550907218 | 0.264 | 0.009 | 0 PER | chr4-148049932-148 |
| 0 | 5.53151841684063 | 0.109 | 0.003 | 0 PER | chr4-175694444-175 |
| 0 | 6.00902687323092 | 0.182 | 0.003 | 0 PER | chr4-175694990-175 |
| 0 | 5.03964470541102 | 0.2 | 0.007 | 0 PER | chr5-72049602-7205 |
| 0 | 4.86453700882449 | 0.146 | 0.005 | 0 PER | chr5-72450807-7245 |
| 0 | 4.86954781159687 | 0.12 | 0.004 | 0 PER | chr5-101192401-101 |
| 0 | 5.0940304419535 | 0.111 | 0.004 | 0 PER | chr5-128113098-128 |
| 0 | 5.61868301540806 | 0.183 | 0.004 | 0 PER | chr5-139637808-139 |
| 0 | 5.56068816325257 | 0.223 | 0.005 | 0 PER | chr5-142469654-142 |
| 0 | 5.75499018457359 | 0.181 | 0.004 | 0 PER | chr5-143487862-143 |
| 0 | 5.09055693156758 | 0.16 | 0.006 | 0 PER | chr5-152903767-152 |
| 0 | 5.1385585226193 | 0.142 | 0.004 | 0 PER | chr5-154227209-154 |
| 0 | 5.03208329710571 | 0.244 | 0.009 | 0 PER | chr5-164849972-164 |
| 0 | 5.02221461589184 | 0.147 | 0.005 | 0 PER | chr6-26377546-2637 |
| 0 | 4.85968020530683 | 0.184 | 0.006 | 0 PER | chr6-89884988-8988 |
| 0 | 4.85019911001397 | 0.12 | 0.004 | 0 PER | chr6-119804002-119 |
| 0 | 4.8737033222085 | 0.165 | 0.005 | 0 PER | chr6-127387052-127 |
| 0 | 4.96025413480604 | 0.122 | 0.004 | 0 PER | chr6-131659079-131 |
| 0 | 5.23899722871079 | 0.108 | 0.003 | 0 PER | chr7-14396619-1439 |
| 0 | 4.83411545899727 | 0.327 | 0.015 | 0 PER | chr7-57854311-5785 |
| 0 | 5.24524510629926 | 0.248 | 0.008 | 0 PER | chr7-65927471-6592 |
| 0 | 5.17511213998319 | 0.191 | 0.006 | 0 PER | chr7-84301221-8430 |
| 0 | 4.85586407861316 | 0.109 | 0.004 | 0 PER | chr7-90192901-9019 |

| | | | | | | |
|---|---|---|---|---|---|---|
| 0 | 5.17630027152998 | 0.279 | 0.009 | 0 | PER | chr7-100987599-100 |
| 0 | 5.67772921734547 | 0.142 | 0.003 | 0 | PER | chr7-103941473-103 |
| 0 | 5.04212438927619 | 0.117 | 0.003 | 0 | PER | chr7-105643583-105 |
| 0 | 5.83510781305218 | 0.206 | 0.004 | 0 | PER | chr8-5466492-54669 |
| 0 | 5.10397403767092 | 0.1 | 0.003 | 0 | PER | chr8-51849445-5184 |
| 0 | 5.23239608798051 | 0.2 | 0.006 | 0 | PER | chr8-62044010-6204 |
| 0 | 5.13960270243309 | 0.189 | 0.006 | 0 | PER | chr8-84028520-8402 |
| 0 | 4.92160238103599 | 0.111 | 0.004 | 0 | PER | chr8-111385684-111 |
| 0 | 5.06961866960032 | 0.174 | 0.006 | 0 | PER | chr8-113775664-113 |
| 0 | 4.95123338293742 | 0.25 | 0.01 | 0 | PER | chr8-121064190-121 |
| 0 | 4.93554247473829 | 0.119 | 0.004 | 0 | PER | chr9-2215855-22163 |
| 0 | 5.3049790520326 | 0.173 | 0.005 | 0 | PER | chr9-23371034-2337 |
| 0 | 4.87555066908913 | 0.111 | 0.004 | 0 | PER | chr9-23396765-2339 |
| 0 | 5.25941407044001 | 0.113 | 0.004 | 0 | PER | chr9-23896101-2389 |
| 0 | 5.20756261545798 | 0.144 | 0.004 | 0 | PER | chr9-49843895-4984 |
| 0 | 4.99447740661356 | 0.145 | 0.005 | 0 | PER | chr9-76844742-7684 |
| 0 | 5.16424851699415 | 0.204 | 0.007 | 0 | PER | chr9-89250961-8925 |
| 0 | 5.1751770868003 | 0.111 | 0.003 | 0 | PER | chrX-127476549-127 |
| 0 | 7.96419735209287 | 0.399 | 0.002 | 0 | SCH | chr1-3096640-30971 |
| 0 | 7.91906954330629 | 0.129 | 0.001 | 0 | SCH | chr1-122941340-122 |
| 0 | 7.55329330794396 | 0.124 | 0.001 | 0 | SCH | chr1-131503120-131 |
| 0 | 7.72425933494091 | 0.175 | 0.001 | 0 | SCH | chr1-148068801-148 |
| 0 | 7.99744100098434 | 0.423 | 0.002 | 0 | SCH | chr1-168997252-168 |
| 0 | 7.55329330794396 | 0.14 | 0.001 | 0 | SCH | chr1-171907474-171 |
| 0 | 7.6008360176851 | 0.111 | 0.001 | 0 | SCH | chr1-177223344-177 |
| 0 | 7.87813587147192 | 0.133 | 0.001 | 0 | SCH | chr1-211256054-211 |
| 0 | 7.49365859755534 | 0.209 | 0.001 | 0 | SCH | chr1-218400675-218 |
| 0 | 7.54871079759505 | 0.138 | 0.001 | 0 | SCH | chr1-231417245-231 |
| 0 | 7.52860963304711 | 0.111 | 0.001 | 0 | SCH | chr1-251245351-251 |
| 0 | 7.59850485951807 | 0.386 | 0.003 | 0 | SCH | chr1-256891987-256 |
| 0 | 7.5045349000721 | 0.168 | 0.001 | 0 | SCH | chr10-21159475-211 |
| 0 | 7.53511779863342 | 0.224 | 0.001 | 0 | SCH | chr10-21900588-219 |
| 0 | 7.61597294083043 | 0.293 | 0.002 | 0 | SCH | chr10-31589921-315 |
| 0 | 7.91221085528681 | 0.22 | 0.001 | 0 | SCH | chr10-76385575-763 |
| 0 | 8.55097103448408 | 0.31 | 0.001 | 0 | SCH | chr11-5391833-5392 |
| 0 | 7.68533794943125 | 0.239 | 0.002 | 0 | SCH | chr11-42155800-421 |
| 0 | 7.48542636251899 | 0.295 | 0.002 | 0 | SCH | chr11-44695098-446 |
| 0 | 7.7198500117236 | 0.302 | 0.002 | 0 | SCH | chr12-6540288-6540 |
| 0 | 7.48741764007417 | 0.12 | 0.001 | 0 | SCH | chr13-3925515-3926 |
| 0 | 8.4655155971818 | 0.344 | 0.001 | 0 | SCH | chr13-12079325-120 |
| 0 | 8.34513045949538 | 0.242 | 0.001 | 0 | SCH | chr13-28047082-280 |
| 0 | 8.35325582722359 | 0.228 | 0.001 | 0 | SCH | chr13-28189473-281 |
| 0 | 8.28288598974062 | 0.417 | 0.002 | 0 | SCH | chr13-62618270-626 |

| | | | | | |
|---|---|---|---|---|---|
| 0 | 8.17578193088429 | 0.235 | 0.001 | 0 SCH | chr13-70549945-705[illegible] |
| 0 | 7.62623106761526 | 0.202 | 0.001 | 0 SCH | chr13-79037520-790[illegible] |
| 0 | 7.81190895786179 | 0.432 | 0.003 | 0 SCH | chr14-2704135-2704[illegible] |
| 0 | 7.4979307549461 | 0.111 | 0.001 | 0 SCH | chr14-24532626-245[illegible] |
| 0 | 7.68898960790834 | 0.279 | 0.002 | 0 SCH | chr14-40927205-409[illegible] |
| 0 | 7.66551113722177 | 0.255 | 0.002 | 0 SCH | chr14-46670664-466[illegible] |
| 0 | 8.00601451375928 | 0.25 | 0.001 | 0 SCH | chr14-47567511-475[illegible] |
| 0 | 7.97100013852177 | 0.179 | 0.001 | 0 SCH | chr14-48647921-486[illegible] |
| 0 | 7.67072139568942 | 0.171 | 0.001 | 0 SCH | chr14-52684183-526[illegible] |
| 0 | 7.44862299883891 | 0.26 | 0.002 | 0 SCH | chr14-104102689-10[illegible] |
| 0 | 8.22981253153695 | 0.315 | 0.001 | 0 SCH | chr15-27240224-272[illegible] |
| 0 | 7.58560410835895 | 0.33 | 0.002 | 0 SCH | chr15-35770555-357[illegible] |
| 0 | 8.23464746476364 | 0.466 | 0.002 | 0 SCH | chr15-46100991-461[illegible] |
| 0 | 8.1687076355153 | 0.24 | 0.001 | 0 SCH | chr15-60460582-604[illegible] |
| 0 | 7.68212121148021 | 0.142 | 0.001 | 0 SCH | chr15-62775513-627[illegible] |
| 0 | 7.85877049717819 | 0.153 | 0.001 | 0 SCH | chr15-63620357-636[illegible] |
| 0 | 7.66633270283377 | 0.233 | 0.001 | 0 SCH | chr15-64637511-646[illegible] |
| 0 | 8.19746928570846 | 0.18 | 0.001 | 0 SCH | chr15-70657368-706[illegible] |
| 0 | 7.46178993338286 | 0.266 | 0.002 | 0 SCH | chr15-79305030-793[illegible] |
| 0 | 7.86110571657921 | 0.279 | 0.002 | 0 SCH | chr15-79716242-797[illegible] |
| 0 | 7.81247482468837 | 0.129 | 0.001 | 0 SCH | chr15-90724490-907[illegible] |
| 0 | 8.15031942167016 | 0.186 | 0.001 | 0 SCH | chr16-14386863-143[illegible] |
| 0 | 7.70681168376595 | 0.133 | 0.001 | 0 SCH | chr16-49681304-496[illegible] |
| 0 | 8.05246852261691 | 0.177 | 0.001 | 0 SCH | chr16-53842654-538[illegible] |
| 0 | 7.51121700859388 | 0.204 | 0.001 | 0 SCH | chr16-67898631-678[illegible] |
| 0 | 7.65562898942717 | 0.321 | 0.003 | 0 SCH | chr17-25305850-253[illegible] |
| 0 | 7.58980394775313 | 0.217 | 0.001 | 0 SCH | chr17-38345188-383[illegible] |
| 0 | 7.69523751259778 | 0.397 | 0.003 | 0 SCH | chr17-38418410-384[illegible] |
| 0 | 7.80823392647267 | 0.419 | 0.003 | 0 SCH | chr17-68280447-682[illegible] |
| 0 | 7.60797201912138 | 0.177 | 0.001 | 0 SCH | chr18-1920262-1920[illegible] |
| 0 | 8.26740763129141 | 0.189 | 0.001 | 0 SCH | chr18-19583207-195[illegible] |
| 0 | 7.44843369206363 | 0.124 | 0.001 | 0 SCH | chr18-33851958-338[illegible] |
| 0 | 7.50703573218397 | 0.268 | 0.002 | 0 SCH | chr18-35211351-352[illegible] |
| 0 | 7.98110369736523 | 0.155 | 0.001 | 0 SCH | chr18-38171314-381[illegible] |
| 0 | 7.82129676946965 | 0.175 | 0.001 | 0 SCH | chr18-75247538-752[illegible] |
| 0 | 8.25716248971767 | 0.279 | 0.001 | 0 SCH | chr18-79461760-794[illegible] |
| 0 | 8.65882304384889 | 0.268 | 0.001 | 0 SCH | chr19-3694619-3695[illegible] |
| 0 | 7.72684702406841 | 0.166 | 0.001 | 0 SCH | chr19-21671827-216[illegible] |
| 0 | 7.67501062863453 | 0.281 | 0.002 | 0 SCH | chr19-42321657-423[illegible] |
| 0 | 7.54708926526218 | 0.129 | 0.001 | 0 SCH | chr2-8919283-89197[illegible] |
| 0 | 7.59357361082689 | 0.311 | 0.003 | 0 SCH | chr2-21518569-2151[illegible] |
| 0 | 7.62017496683249 | 0.37 | 0.002 | 0 SCH | chr2-60496158-6049[illegible] |
| 0 | 8.03585196551484 | 0.248 | 0.001 | 0 SCH | chr2-62636603-6263[illegible] |

| | | | | | | |
|---|---|---|---|---|---|---|
| 0 | 7.95905612931477 | 0.302 | 0.002 | 0 | SCH | chr2-81323497-8132 |
| 0 | 7.51288690566565 | 0.169 | 0.001 | 0 | SCH | chr2-111532259-111 |
| 0 | 7.92529517175204 | 0.308 | 0.002 | 0 | SCH | chr2-135094665-135 |
| 0 | 7.50278026608304 | 0.321 | 0.003 | 0 | SCH | chr2-142588502-142 |
| 0 | 8.33964475963911 | 0.22 | 0.001 | 0 | SCH | chr2-165583152-165 |
| 0 | 7.67909950201172 | 0.273 | 0.002 | 0 | SCH | chr2-198492551-198 |
| 0 | 7.56862525069547 | 0.171 | 0.001 | 0 | SCH | chr2-203283000-203 |
| 0 | 7.74300575213821 | 0.239 | 0.002 | 0 | SCH | chr2-232709774-232 |
| 0 | 7.83784971172414 | 0.191 | 0.001 | 0 | SCH | chr2-245525338-245 |
| 0 | 7.47824772641994 | 0.189 | 0.001 | 0 | SCH | chr20-16267296-162 |
| 0 | 7.90642451155262 | 0.239 | 0.001 | 0 | SCH | chr20-32175740-321 |
| 0 | 7.46575267632688 | 0.188 | 0.001 | 0 | SCH | chr3-28108232-2810 |
| 0 | 7.90813972954852 | 0.373 | 0.002 | 0 | SCH | chr3-28451890-2845 |
| 0 | 7.99985632041378 | 0.302 | 0.002 | 0 | SCH | chr3-29520296-2952 |
| 0 | 7.66915264798566 | 0.182 | 0.001 | 0 | SCH | chr3-30808420-3080 |
| 0 | 8.25091457655516 | 0.264 | 0.001 | 0 | SCH | chr3-31339519-3134 |
| 0 | 7.70263023399055 | 0.166 | 0.001 | 0 | SCH | chr3-32362217-3236 |
| 0 | 7.52020674434073 | 0.23 | 0.002 | 0 | SCH | chr3-32869026-3286 |
| 0 | 7.74703810011869 | 0.381 | 0.002 | 0 | SCH | chr3-36755896-3675 |
| 0 | 8.08026132942908 | 0.317 | 0.002 | 0 | SCH | chr3-40944999-4094 |
| 0 | 8.07020905659718 | 0.242 | 0.001 | 0 | SCH | chr3-49021501-4902 |
| 0 | 7.86463103563395 | 0.244 | 0.001 | 0 | SCH | chr3-50687283-5068 |
| 0 | 7.57353910628599 | 0.268 | 0.002 | 0 | SCH | chr3-50884011-5088 |
| 0 | 7.50536654689876 | 0.162 | 0.001 | 0 | SCH | chr3-51660648-5166 |
| 0 | 7.73574112515115 | 0.14 | 0.001 | 0 | SCH | chr3-66790302-6679 |
| 0 | 7.48807014096727 | 0.14 | 0.001 | 0 | SCH | chr3-70482480-7048 |
| 0 | 7.64475488285199 | 0.126 | 0.001 | 0 | SCH | chr3-70799894-7080 |
| 0 | 7.53496647851011 | 0.279 | 0.002 | 0 | SCH | chr3-118077017-118 |
| 0 | 8.02377924725617 | 0.308 | 0.002 | 0 | SCH | chr3-124826050-124 |
| 0 | 7.45352041725634 | 0.266 | 0.002 | 0 | SCH | chr3-127862508-127 |
| 0 | 7.46248695669058 | 0.337 | 0.003 | 0 | SCH | chr3-127974428-127 |
| 0 | 7.98879476095612 | 0.273 | 0.001 | 0 | SCH | chr3-150700219-150 |
| 0 | 8.18191269052506 | 0.282 | 0.001 | 0 | SCH | chr3-150721705-150 |
| 0 | 7.78819294648155 | 0.251 | 0.001 | 0 | SCH | chr4-12963722-1296 |
| 0 | 7.56434679159329 | 0.115 | 0.001 | 0 | SCH | chr4-31904662-3190 |
| 0 | 8.03160303323135 | 0.237 | 0.001 | 0 | SCH | chr4-38562452-3856 |
| 0 | 8.0538838225779 | 0.191 | 0.001 | 0 | SCH | chr4-74944443-7494 |
| 0 | 7.68438405721844 | 0.404 | 0.003 | 0 | SCH | chr4-89778122-8977 |
| 0 | 7.99194802490133 | 0.155 | 0.001 | 0 | SCH | chr4-91668018-9166 |
| 0 | 7.50221089733079 | 0.149 | 0.001 | 0 | SCH | chr4-97226342-9722 |
| 0 | 8.10661016548045 | 0.202 | 0.001 | 0 | SCH | chr4-129331136-129 |
| 0 | 7.98284418494518 | 0.23 | 0.001 | 0 | SCH | chr4-136938860-136 |
| 0 | 7.59302485771823 | 0.211 | 0.001 | 0 | SCH | chr4-147316922-147 |

| | | | | | | |
|---|---|---|---|---|---|---|
| 0 | 7.92930426344403 | 0.264 | 0.001 | 0 | SCH | chr4-168805845-168 |
| 0 | 7.62734921753629 | 0.26 | 0.002 | 0 | SCH | chr4-177304883-177 |
| 0 | 8.06813836973749 | 0.328 | 0.002 | 0 | SCH | chr5-14231669-1423 |
| 0 | 7.93379180807594 | 0.202 | 0.001 | 0 | SCH | chr5-15560558-1556 |
| 0 | 7.84311779231561 | 0.16 | 0.001 | 0 | SCH | chr5-39619596-3962 |
| 0 | 7.52894465187045 | 0.179 | 0.001 | 0 | SCH | chr5-87864796-8786 |
| 0 | 7.80905331768305 | 0.333 | 0.002 | 0 | SCH | chr5-114210220-114 |
| 0 | 7.62234956456221 | 0.186 | 0.001 | 0 | SCH | chr5-116125925-116 |
| 0 | 7.51991599721833 | 0.353 | 0.003 | 0 | SCH | chr5-163945703-163 |
| 0 | 8.04417871604223 | 0.492 | 0.003 | 0 | SCH | chr6-47881441-4788 |
| 0 | 7.9714274059598 | 0.179 | 0.001 | 0 | SCH | chr6-54896000-5489 |
| 0 | 7.79451122913277 | 0.168 | 0.001 | 0 | SCH | chr6-75621326-7562 |
| 0 | 7.4793385077883 | 0.253 | 0.002 | 0 | SCH | chr6-108192405-108 |
| 0 | 7.72165381708935 | 0.244 | 0.002 | 0 | SCH | chr6-122291417-122 |
| 0 | 7.73057824557987 | 0.193 | 0.001 | 0 | SCH | chr6-140908620-140 |
| 0 | 8.01597369037664 | 0.128 | 0.001 | 0 | SCH | chr7-23978316-2397 |
| 0 | 7.49935599150421 | 0.424 | 0.004 | 0 | SCH | chr7-43355920-4335 |
| 0 | 7.53528038530455 | 0.173 | 0.001 | 0 | SCH | chr7-47789254-4778 |
| 0 | 7.93300134423742 | 0.129 | 0 | 0 | SCH | chr7-83352958-8335 |
| 0 | 7.49169144000996 | 0.18 | 0.001 | 0 | SCH | chr7-93612312-9361 |
| 0 | 7.50028588107318 | 0.257 | 0.002 | 0 | SCH | chr7-99942310-9994 |
| 0 | 8.03993893170842 | 0.197 | 0.001 | 0 | SCH | chr7-117819715-117 |
| 0 | 7.52762215994313 | 0.366 | 0.002 | 0 | SCH | chr7-119285555-119 |
| 0 | 7.88381197935677 | 0.115 | 0.001 | 0 | SCH | chr8-9670833-96713 |
| 0 | 8.05308975907328 | 0.275 | 0.001 | 0 | SCH | chr8-25693349-2569 |
| 0 | 7.59624920475913 | 0.319 | 0.002 | 0 | SCH | chr8-32969268-3296 |
| 0 | 7.73191730063549 | 0.271 | 0.002 | 0 | SCH | chr8-47790407-4779 |
| 0 | 8.05473176881089 | 0.219 | 0.001 | 0 | SCH | chr8-82066106-8206 |
| 0 | 7.82023622860478 | 0.364 | 0.002 | 0 | SCH | chr8-115996271-115 |
| 0 | 8.37010319050876 | 0.199 | 0.001 | 0 | SCH | chr9-5813912-58144 |
| 0 | 7.59135481635718 | 0.244 | 0.002 | 0 | SCH | chr9-22557913-2255 |
| 0 | 7.86101578374613 | 0.22 | 0.001 | 0 | SCH | chr9-72224203-7222 |
| 0 | 7.9284905518381 | 0.326 | 0.002 | 0 | SCH | chr9-79998395-7999 |
| 0 | 8.09625033197756 | 0.33 | 0.002 | 0 | SCH | chr9-94087849-9408 |
| 0 | 7.47109566764362 | 0.257 | 0.001 | 0 | SCH | chrX-11122285-1112 |
| 0 | 7.59778091753072 | 0.253 | 0.001 | 0 | SCH | chrX-68115937-6811 |
| 0 | 7.80213827601529 | 0.222 | 0.001 | 0 | SCH | chrX-100192037-100 |
| 0 | 7.6712708144732 | 0.128 | 0.001 | 0 | SCH | chrX-111405795-111 |
| 0 | 7.71852743965127 | 0.131 | 0.001 | 0 | SCH | chrX-139138364-139 |
| 8.59345026 | 2.95206338934067 | 0.332 | 0.047 | 6.2969881 | SG | chr2-183838714-183 |
| 9.55176825 | 4.29430151499089 | 0.102 | 0.005 | 6.9992110 | SG | chr10-79768401-797 |
| 1.98227576 | 3.62054406896639 | 0.189 | 0.017 | 1.4525442 | SG | chr17-42484916-424 |
| 6.04656682 | 3.23612758731167 | 0.23 | 0.031 | 4.4307185 | SG | chr1-47800563-4780 |

| | | | | | | | |
|---|---|---|---|---|---|---|---|
| 3.33190533 | 3.97960851497684 | | 0.102 | 0.007 | 2.4415069 | SG | chr20-2649385-2649 |
| 4.73513941 | 3.38463360386407 | | 0.161 | 0.017 | 3.4697491 | SG | chr1-47801132-4780 |
| 1.48397947 | 3.59725016415874 | | 0.105 | 0.008 | 1.0874097 | SG | chr20-3567933-3568 |
| 4.85489936 | 2.96800737015657 | | 0.169 | 0.02 | 3.5575051 | SG | chr4-104742920-104 |
| 1.91333771 | 3.84189509290405 | | 0.1 | 0.008 | 1.4020288 | SG | chr17-7777399-7777 |
| 6.03608158 | 3.81682116307277 | | 0.1 | 0.008 | 4.4230353 | SG | chr10-19821273-198 |
| 4.30838520 | 2.74216807332968 | | 0.266 | 0.05 | 3.1570381 | SG | chr17-42477932-424 |
| 1.26280453 | 2.89576423952614 | | 0.148 | 0.017 | 9.2534022 | SG | chr5-103095635-103 |
| 3.28171711 | 3.73086750907774 | | 0.115 | 0.011 | 2.4047307 | SG | chr1-84377013-8437 |
| 1.03916220 | 3.45184261723395 | | 0.105 | 0.009 | 7.6146273 | SG | chr10-19807502-198 |
| 7.58660765 | 3.37413914016662 | | 0.115 | 0.012 | 5.5592081 | SG | chr9-92233086-9223 |
| 5.89515731 | 3.22810614570498 | | 0.113 | 0.011 | 4.3197708 | SG | chr1-153426529-153 |
| 1.33636095 | 2.77229516779001 | | 0.225 | 0.041 | 9.7923987 | SG | chr17-42799404-427 |
| 5.14969940 | | 2.818253454 | 0.187 | 0.03 | 3.7735246 | SG | chr14-74395633-743 |
| 3.90816539 | 2.65648338642986 | | 0.189 | 0.031 | 2.8637707 | SG | chr3-11162065-1116 |
| 3.31486247 | 3.11604290921226 | | 0.133 | 0.017 | 2.4290185 | SG | chr8-121135720-121 |
| 1.07371618 | 3.17120043815751 | | 0.123 | 0.014 | 7.8678271 | SG | chr6-9958250-99587 |
| 2.19547748 | 2.80911007990701 | | 0.166 | 0.025 | 1.6087712 | SG | chr12-44384365-443 |
| 2.90712603 | 2.48892093310099 | | 0.205 | 0.038 | 2.1302431 | SG | chr1-203378884-203 |
| 9.21588360 | 2.89405110641484 | | 0.164 | 0.025 | 6.7530861 | SG | chr6-98480820-9848 |
| 1.74316432 | 3.19901358368784 | | 0.125 | 0.016 | 1.2773315 | SG | chr7-111914973-111 |
| 3.98289248 | 2.80392421488548 | | 0.164 | 0.026 | 2.9185281 | SG | chr12-14065804-140 |
| 2.30059553 | 2.49798194201091 | | 0.246 | 0.054 | 1.6857981 | SG | chr17-41380220-413 |
| 3.10398964 | 3.12647260293575 | | 0.12 | 0.015 | 2.2744980 | SG | chr9-15395696-1539 |
| 4.42159287 | 2.78278237425933 | | 0.151 | 0.023 | 3.2399929 | SG | chr18-56366449-563 |
| 2.64170430 | 3.03842914654795 | | 0.123 | 0.016 | 1.9357510 | SG | chr17-41560145-415 |
| 3.57121309 | 2.77140965863993 | | 0.176 | 0.031 | 2.6168635 | SG | chr17-41535040-415 |
| 1.40657837 | 2.76027378550666 | | 0.166 | 0.028 | 1.0306928 | SG | chr1-96110001-9611 |
| 1.63344970 | 2.99059528173316 | | 0.128 | 0.017 | 1.1969364 | SG | chr10-16543692-165 |
| 2.00119965 | 2.80249466905297 | | 0.151 | 0.024 | 1.4664110 | SG | chr20-7949511-7950 |
| 2.47907385 | 2.75839526250063 | | 0.187 | 0.035 | 1.8165810 | SG | chr8-110010878-110 |
| 1.37695361 | 3.6230934438131 | | 0.1 | 0.011 | 1.0089847 | SG | chr7-105241216-105 |
| 1.15674305 | 2.50763184071096 | | 0.212 | 0.045 | 8.4762198 | SG | chr1-47735193-4773 |
| 1.87982702 | 2.67162089565689 | | 0.174 | 0.032 | 1.3774733 | SG | chr17-1157271-1157 |
| 2.28286533 | 3.23156159929845 | | 0.1 | 0.012 | 1.6728060 | SG | chr20-29523148-295 |
| 1.45619758 | 3.07212633662464 | | 0.1 | 0.012 | 1.0670520 | SG | chr2-173477888-173 |
| 2.13012981 | 2.97565825626563 | | 0.113 | 0.015 | 1.5608867 | SG | chr13-94394411-943 |
| 2.14795168 | 2.67725370723117 | | 0.171 | 0.031 | 1.5739459 | SG | chr19-48326132-483 |
| 3.99375899 | 3.15400199872964 | | 0.102 | 0.012 | 2.9264908 | SG | chr1-166968245-166 |
| 4.97188960 | 3.20514319313885 | | 0.1 | 0.012 | 3.6432316 | SG | chr1-47812932-4781 |
| 8.32188020 | 2.90123555886982 | | 0.133 | 0.02 | 6.0979908 | SG | chr1-99105642-9910 |
| 1.72170214 | 2.77729880906147 | | 0.125 | 0.018 | 1.2616047 | SG | chr17-1112816-1113 |
| 3.20391347 | 2.77311641114984 | | 0.12 | 0.017 | 2.3477188 | SG | chr10-105921002-10 |

| | | | | | | | |
|---|---|---|---|---|---|---|---|
| 4.17573896 | 3.07216633613213 | | 0.102 | 0.013 | 3.0598395 | SG | chr2-219640066-219 |
| 4.92222803 | 2.66880096438383 | | 0.156 | 0.027 | 3.6068413 | SG | chr18-56328732-563 |
| 3.95960713 | 2.83835347826957 | | 0.113 | 0.015 | 2.9014654 | SG | chr20-3412629-3413 |
| 5.83288585 | 3.07343616775531 | | 0.118 | 0.017 | 4.2741404 | SG | chr3-15182008-1518 |
| 1.54943021 | 2.81786233447677 | | 0.118 | 0.017 | 1.1353697 | SG | chr4-134402561-134 |
| 2.55591585 | 3.12382385257911 | | 0.1 | 0.013 | 1.8728882 | SG | chr5-138207226-138 |
| 1.23878679 | 2.99463654291708 | | 0.12 | 0.018 | 9.0774084 | SG | chr3-78942921-7894 |
| 1.72679537 | 2.4762713492806 | | 0.156 | 0.028 | 1.2653369 | SG | chr5-103096427-103 |
| 3.11968187 | 3.07286551983438 | | 0.102 | 0.013 | 2.2859968 | SG | chr19-52257633-522 |
| 5.88102647 | 2.65134980503494 | | 0.148 | 0.026 | 4.3094162 | SG | chr7-56220417-5622 |
| 7.17918409 | 2.58962072682929 | | 0.123 | 0.019 | 5.2606620 | SG | chr5-138850499-138 |
| 1.14465381 | 2.76575511729307 | | 0.13 | 0.021 | 8.3876339 | SG | chr19-15716034-157 |
| 1.54430357 | 2.80592348395493 | | 0.146 | 0.026 | 1.1316131 | SG | chr8-33732768-3373 |
| 2.71709294 | 2.52410042538241 | | 0.169 | 0.033 | 1.9909933 | SG | chr18-56329313-563 |
| 5.45365154 | 2.52702779426327 | | 0.166 | 0.033 | 3.9962504 | SG | chr9-15713984-1571 |
| 4.88568638 | 2.97430239835358 | | 0.113 | 0.017 | 3.5800648 | SG | chr10-35799109-357 |
| 5.04913966 | 2.76461943695507 | | 0.12 | 0.019 | 3.6998378 | SG | chr5-159365688-159 |
| 5.47882302 | 2.64640437074283 | | 0.128 | 0.021 | 4.0146952 | SG | chr18-46221273-462 |
| 5.94903088 | 2.54152138206445 | | 0.143 | 0.025 | 4.3592475 | SG | chr17-21495107-214 |
| 1.60110445 | 2.65447036234964 | | 0.133 | 0.023 | 1.1732349 | SG | chr4-104803178-104 |
| 1.93163376 | 2.55284220384542 | | 0.13 | 0.022 | 1.4154355 | SG | chr12-28083573-280 |
| 4.10844547 | 2.83255021988725 | | 0.107 | 0.015 | 3.0105291 | SG | chr1-47804640-4780 |
| 9.26593926 | 2.64339289775893 | | 0.118 | 0.018 | 6.7897652 | SG | chr1-166987492-166 |
| 9.33477294 | 2.88342901622771 | | 0.1 | 0.014 | 6.8402042 | SG | chr9-91797551-9179 |
| 9.75552830 | 2.69701215775955 | | 0.107 | 0.016 | 7.1485194 | SG | chr3-146497458-146 |
| 1.14208529 | | 2.5143395 | 0.202 | 0.048 | 8.3688127 | SG | chr10-81392483-813 |
| 1.19399711 | 2.72952591690328 | | 0.128 | 0.021 | 8.7492048 | SG | chr11-32176823-321 |
| 1.24722541 | 3.01106790708776 | | 0.1 | 0.014 | 9.1392437 | SG | chr7-1364024-13645 |
| 1.54471188 | 2.57907084206134 | | 0.169 | 0.035 | 1.1319123 | SG | chr4-147657441-147 |
| 1.85621339 | 2.52393835577728 | | 0.143 | 0.026 | 1.3601700 | SG | chr5-56955433-5695 |
| 1.05197727 | 2.74591439581381 | | 0.12 | 0.019 | 7.7085317 | SG | chr10-72089981-720 |
| 1.35567293 | 2.80404293011372 | | 0.118 | 0.019 | 9.9339103 | SG | chr2-183830767-183 |
| 2.81471934 | 2.64598816852433 | | 0.136 | 0.024 | 2.0625306 | SG | chr1-240626120-240 |
| 3.61086385 | 2.51113528005801 | | 0.128 | 0.022 | 2.6459182 | SG | chr4-80012433-8001 |
| 4.45436670 | 2.70337707881957 | | 0.115 | 0.018 | 3.2640084 | SG | chr6-126197181-126 |
| 6.13264760 | 2.60601308446595 | | 0.123 | 0.02 | 4.4937956 | SG | chr17-41386108-413 |
| 1.55606406 | 2.60023577399622 | | 0.113 | 0.018 | 1.1402308 | SG | chr3-60379398-6037 |
| 1.77575210 | 2.8533533629753 | | 0.102 | 0.015 | 1.3012107 | SG | chr5-151956194-151 |
| 1.87392633 | 2.69734590222956 | | 0.12 | 0.02 | 1.3731495 | SG | chr3-149266177-149 |
| 1.96367171 | 2.89316735112478 | | 0.113 | 0.018 | 1.4389118 | SG | chr10-35613406-356 |
| 2.44944095 | 2.72596180802381 | | 0.107 | 0.016 | 1.7948670 | SG | chr18-46264424-462 |
| 2.48105914 | 2.73789077507518 | | 0.102 | 0.015 | 1.8180357 | SG | chr10-71064622-710 |
| 4.66249119 | 2.56338438340802 | | 0.107 | 0.016 | 3.4165150 | SG | chr12-19740512-197 |

| | | | | | | |
|---|---|---|---|---|---|---|
| 1.16723051 | 2.6426602379899 | 0.11 | 0.017 | 8.5530683 | SG | chr7-56087338-5608 |
| 1.28873443 | 2.87861426448955 | 0.102 | 0.015 | 9.4434077 | SG | chr5-103088516-103 |
| 1.37084160 | 2.83048556825764 | 0.1 | 0.015 | 1.0045061 | SG | chr1-197705464-197 |
| 1.55880921 | 2.59616485659262 | 0.113 | 0.018 | 1.1422423 | SG | chr15-40051361-400 |
| 1.73972470 | 2.49404433093659 | 0.128 | 0.023 | 1.2748111 | SG | chr10-81860855-818 |
| 2.25264736 | 2.71590173448783 | 0.105 | 0.016 | 1.6506634 | SG | chr10-99021737-990 |
| 5.11120432 | 2.58722311896768 | 0.115 | 0.019 | 3.7453167 | SG | chr4-10848631-1084 |
| 1.04111875 | 2.69007968662878 | 0.1 | 0.015 | 7.6289642 | SG | chr3-141513029-141 |
| 1.57605337 | 2.82040516748481 | 0.107 | 0.017 | 1.1548783 | SG | chr18-47527869-475 |
| 2.07084823 | 2.51632354364574 | 0.138 | 0.027 | 1.5174471 | SG | chr20-29769276-297 |
| 2.07541163 | 2.92726969690312 | 0.1 | 0.015 | 1.5207910 | SG | chr8-61037922-6103 |
| 2.66413252 | 2.62284506572133 | 0.107 | 0.017 | 1.9521857 | SG | chr9-874466-874967 |
| 8.51343914 | 2.48307127195808 | 0.13 | 0.025 | 6.2383587 | SG | chr8-118588216-118 |
| 9.41189881 | 2.56750809194948 | 0.105 | 0.017 | 6.8967194 | SG | chr5-152042671-152 |
| 3.64170138 | 2.47834600288176 | 0.11 | 0.018 | 2.6685149 | SG | chr7-111898675-111 |
| 9.86575958 | 2.81084753119468 | 0.102 | 0.016 | 7.2292931 | SG | chr5-137232666-137 |
| 1.00231001 | 2.58640111474512 | 0.13 | 0.026 | 7.3445869 | SG | chr6-89426453-8942 |
| 1.41219729 | 2.54291021152614 | 0.123 | 0.023 | 1.0348101 | SG | chr7-120500373-120 |
| 4.91725611 | 2.55215766150768 | 0.105 | 0.018 | 3.6031980 | SG | chr3-161707319-161 |
| 1.18911879 | 2.51944513057796 | 0.105 | 0.018 | 8.7134582 | SG | chr1-85239439-8523 |
| 1.28200629 | 2.5177271577819 | 0.1 | 0.016 | 9.3941062 | SG | chr1-187554669-187 |
| 1.78809842 | 2.62883784783876 | 0.113 | 0.02 | 1.3102577 | SG | chr8-110734146-110 |
| 2.33501193 | 2.46509754915019 | 0.118 | 0.022 | 1.7110173 | SG | chr10-52778810-527 |
| 2.36070679 | 2.62755999937038 | 0.115 | 0.021 | 1.7298456 | SG | chr7-114800866-114 |
| 6.27165009 | 2.49932601656657 | 0.133 | 0.027 | 4.5956519 | SG | chr8-60957435-6095 |
| 7.43185513 | 2.49056892128745 | 0.128 | 0.026 | 5.4458107 | SG | chr7-56071560-5607 |
| 7.88014519 | 2.69120096541522 | 0.102 | 0.017 | 5.7743024 | SG | chr3-8972768-89732 |
| 8.02513817 | 2.50748673812524 | 0.105 | 0.018 | 5.8805484 | SG | chr3-141515286-141 |
| 1.41997746 | 2.54369339389547 | 0.118 | 0.022 | 1.0405112 | SG | chr8-41820453-4182 |
| 3.14910531 | 2.78776270067433 | 0.123 | 0.024 | 2.3075573 | SG | chr12-38530390-385 |
| 3.84034173 | 2.64407625385374 | 0.107 | 0.019 | 2.8140718 | SG | chr10-61155990-611 |
| 4.39798925 | 2.77914554997223 | 0.102 | 0.018 | 3.2226969 | SG | chr14-83496810-834 |
| 4.74382388 | 2.59774380078679 | 0.102 | 0.018 | 3.4761128 | SG | chr8-44200686-4420 |
| 7.79688674 | 2.63015181261136 | 0.136 | 0.029 | 5.7132935 | SG | chr1-152441127-152 |
| 4.02329431 | 2.7674702774361 | 0.107 | 0.02 | 2.9481332 | SG | chr8-44140545-4414 |
| 1.16814351 | 2.85554508407139 | 0.1 | 0.018 | 8.5597585 | SG | chr6-131527560-131 |
| 1.32039501 | 2.75973274317838 | 0.115 | 0.023 | 9.6754057 | SG | chr1-95186669-9518 |
| 2.18595588 | 2.53613152960924 | 0.113 | 0.022 | 1.6017941 | SG | chr18-32044820-320 |
| 2.75065776 | 2.47347994806214 | 0.102 | 0.018 | 2.0155884 | SG | chr5-70591465-7059 |
| 6.56143285 | 2.46681855629397 | 0.128 | 0.027 | 4.8079949 | SG | chr4-146038439-146 |
| 8.51440754 | 2.47537459850644 | 0.115 | 0.023 | 6.2390683 | SG | chr4-96171561-9617 |
| 9.25976787 | 2.48305361114109 | 0.105 | 0.019 | 6.7852430 | SG | chr5-144931008-144 |
| 9.45194542 | 2.5522446759767 | 0.105 | 0.019 | 6.9260642 | SG | chr10-54020314-540 |

| | | | | | | |
|---|---|---|---|---|---|---|
| 5.20644836 | 2.50688879398549 | 0.105 | 0.02 | 3.8151083 | SG | chr10-88023977-880 |
| 8.45242597 | 2.54769169448888 | 0.102 | 0.019 | 6.1936503 | SG | chr4-82955157-8295 |
| 1.00436571 | 2.58954356450234 | 0.102 | 0.019 | 7.3596504 | SG | chr1-166963723-166 |
| 8.26835456 | 2.51616109087626 | 0.105 | 0.02 | 6.0587691 | SG | chr6-10648531-1064 |
| 9.03658992 | 2.53484933783548 | 0.1 | 0.019 | 6.6217058 | SG | chr16-18396178-183 |
| 1.37324532 | 2.59834139257584 | 0.105 | 0.021 | 1.0062674 | SG | chr3-57100878-5710 |
| 1.46937801 | 2.52381952047947 | 0.102 | 0.02 | 1.0767102 | SG | chr5-135279716-135 |
| 2.43116764 | 2.55669054541436 | 0.118 | 0.025 | 1.7814769 | SG | chr4-59319985-5932 |
| 2.76599678 | 2.5806674437733 | 0.115 | 0.024 | 2.0268284 | SG | chr18-48328036-483 |
| 1.05915973 | 2.48175890133453 | 0.11 | 0.023 | 7.7611624 | SG | chr8-41429306-4142 |
| 3.24162458 | 2.55456324520971 | 0.105 | 0.021 | 2.3753522 | SG | chr10-72195744-721 |
| 2.73101876 | 2.48057373416408 | 0.11 | 0.024 | 2.0011976 | SG | chr19-48334548-483 |
| 1.40626405 | 2.52140617153948 | 0.107 | 0.023 | 1.0304624 | SG | chr12-40844546-408 |
| 9.23061746 | 2.56440682915925 | 0.118 | 0.028 | 6.7638826 | SG | chr20-2655372-2655 |
| 4.24522019 | 2.62828270774633 | 0.102 | 0.022 | 3.1107530 | SG | chr16-18393767-183 |
| 4.16008960 | 2.49387055741556 | 0.105 | 0.024 | 3.0483722 | SG | chr20-2641343-2641 |
| 7.05066483 | 2.56270600109138 | 0.1 | 0.022 | 5.1664874 | SG | chr1-133533167-133 |
| 0 | 6.66915529818199 | 0.138 | 0.002 | 0 | T | chr1-10763142-1076 |
| 0 | 6.90056159931331 | 0.209 | 0.002 | 0 | T | chr1-10765127-1076 |
| 0 | 6.72762643446368 | 0.175 | 0.002 | 0 | T | chr1-28599274-2859 |
| 0 | 6.56418827290079 | 0.214 | 0.002 | 0 | T | chr1-41844019-4184 |
| 0 | 6.55679878479777 | 0.315 | 0.004 | 0 | T | chr1-46409245-4640 |
| 0 | 6.71531871511087 | 0.236 | 0.002 | 0 | T | chr1-48078125-4807 |
| 0 | 6.57468335241388 | 0.306 | 0.004 | 0 | T | chr1-48079947-4808 |
| 0 | 6.58289583642354 | 0.253 | 0.003 | 0 | T | chr1-52570619-5257 |
| 0 | 6.64273178462404 | 0.119 | 0.001 | 0 | T | chr1-93985501-9398 |
| 0 | 7.06568694128959 | 0.48 | 0.004 | 0 | T | chr1-93995000-9399 |
| 0 | 6.50449484735566 | 0.323 | 0.004 | 0 | T | chr1-94997360-9499 |
| 0 | 6.8239487114638 | 0.378 | 0.004 | 0 | T | chr1-97535487-9753 |
| 0 | 6.53599401403913 | 0.197 | 0.002 | 0 | T | chr1-106590785-106 |
| 0 | 6.79959745035328 | 0.223 | 0.002 | 0 | T | chr1-106607215-106 |
| 0 | 6.52558205677082 | 0.228 | 0.003 | 0 | T | chr1-151402342-151 |
| 0 | 6.4835283305738 | 0.216 | 0.003 | 0 | T | chr1-163267248-163 |
| 0 | 6.65282449695992 | 0.461 | 0.005 | 0 | T | chr1-207230710-207 |
| 0 | 6.65868491214491 | 0.366 | 0.005 | 0 | T | chr1-239850045-239 |
| 0 | 6.68151794870778 | 0.241 | 0.002 | 0 | T | chr1-239856782-239 |
| 0 | 6.61548324540282 | 0.328 | 0.004 | 0 | T | chr1-241001036-241 |
| 0 | 6.80113887329249 | 0.286 | 0.003 | 0 | T | chr10-1270040-1270 |
| 0 | 6.60630299991536 | 0.15 | 0.001 | 0 | T | chr10-2935217-2935 |
| 0 | 6.84237023865889 | 0.301 | 0.003 | 0 | T | chr10-21569781-215 |
| 0 | 6.67514354540067 | 0.231 | 0.002 | 0 | T | chr10-31076499-310 |
| 0 | 6.53537067939935 | 0.165 | 0.002 | 0 | T | chr10-42584866-425 |
| 0 | 6.51097736692714 | 0.238 | 0.003 | 0 | T | chr10-43829663-438 |

| | | | | | | |
|---|---|---|---|---|---|---|
| 0 | 6.53229985115363 | 0.338 | 0.004 | 0 | T | chr10-45705722-457 |
| 0 | 6.92613955189405 | 0.357 | 0.003 | 0 | T | chr10-49236117-492 |
| 0 | 6.5033361343691 | 0.286 | 0.004 | 0 | T | chr10-56407393-564 |
| 0 | 6.49477651024024 | 0.148 | 0.002 | 0 | T | chr10-67791548-677 |
| 0 | 6.55563008775189 | 0.226 | 0.003 | 0 | T | chr10-101147994-10 |
| 0 | 6.82355940274997 | 0.281 | 0.003 | 0 | T | chr11-81879992-818 |
| 0 | 6.86193743427453 | 0.25 | 0.002 | 0 | T | chr12-1005447-1005 |
| 0 | 6.637963536 | 0.277 | 0.003 | 0 | T | chr12-1787201-1787 |
| 0 | 6.67143470950434 | 0.105 | 0.001 | 0 | T | chr12-1805256-1805 |
| 0 | 6.7734466334625 | 0.162 | 0.001 | 0 | T | chr12-1810267-1810 |
| 0 | 7.14657468062049 | 0.308 | 0.003 | 0 | T | chr12-1871354-1871 |
| 0 | 6.62022470517508 | 0.418 | 0.006 | 0 | T | chr12-6653829-6654 |
| 0 | 6.57154231185569 | 0.306 | 0.004 | 0 | T | chr12-10382850-103 |
| 0 | 6.59090381628421 | 0.32 | 0.004 | 0 | T | chr12-14086969-140 |
| 0 | 6.63578294474469 | 0.141 | 0.002 | 0 | T | chr12-17142442-171 |
| 0 | 6.52765767031946 | 0.294 | 0.004 | 0 | T | chr12-20336763-203 |
| 0 | 6.88626677443149 | 0.379 | 0.003 | 0 | T | chr12-31818157-318 |
| 0 | 6.96610742689072 | 0.408 | 0.004 | 0 | T | chr12-43732235-437 |
| 0 | 6.49223422186372 | 0.468 | 0.007 | 0 | T | chr12-45090040-450 |
| 0 | 6.63496275992295 | 0.207 | 0.002 | 0 | T | chr13-37160047-371 |
| 0 | 6.76472522412215 | 0.202 | 0.002 | 0 | T | chr13-37234507-372 |
| 0 | 6.95884066361116 | 0.148 | 0.001 | 0 | T | chr13-55600065-556 |
| 0 | 6.75924871334415 | 0.315 | 0.003 | 0 | T | chr13-76796580-767 |
| 0 | 6.85302600212036 | 0.129 | 0.001 | 0 | T | chr14-19378948-193 |
| 0 | 6.49152383067812 | 0.378 | 0.005 | 0 | T | chr14-44308364-443 |
| 0 | 6.90485078340268 | 0.218 | 0.002 | 0 | T | chr14-80330078-803 |
| 0 | 6.63841987315137 | 0.185 | 0.002 | 0 | T | chr14-91170349-911 |
| 0 | 6.55064584591972 | 0.23 | 0.002 | 0 | T | chr15-95890-96391 |
| 0 | 6.70647046995312 | 0.265 | 0.003 | 0 | T | chr15-119446-11994 |
| 0 | 6.59311350490522 | 0.252 | 0.003 | 0 | T | chr15-4418739-4419 |
| 0 | 6.6517938583034 | 0.173 | 0.002 | 0 | T | chr15-6748615-6749 |
| 0 | 6.76575694440699 | 0.19 | 0.002 | 0 | T | chr15-27351347-273 |
| 0 | 6.68383813639036 | 0.313 | 0.003 | 0 | T | chr15-36523280-365 |
| 0 | 6.60518273425801 | 0.156 | 0.002 | 0 | T | chr15-50393830-503 |
| 0 | 6.59274634045003 | 0.179 | 0.002 | 0 | T | chr15-90175248-901 |
| 0 | 6.7324237736611 | 0.383 | 0.004 | 0 | T | chr16-7404676-7405 |
| 0 | 6.49877933322187 | 0.301 | 0.003 | 0 | T | chr16-65945782-659 |
| 0 | 7.15695649363466 | 0.31 | 0.002 | 0 | T | chr16-74664600-746 |
| 0 | 6.55563008775189 | 0.235 | 0.003 | 0 | T | chr16-74850990-748 |
| 0 | 6.64409701582637 | 0.197 | 0.002 | 0 | T | chr16-74921542-749 |
| 0 | 6.58429370606352 | 0.207 | 0.002 | 0 | T | chr17-23656949-236 |
| 0 | 6.85076774040615 | 0.122 | 0.001 | 0 | T | chr17-29434034-294 |
| 0 | 6.65618407862203 | 0.427 | 0.005 | 0 | T | chr17-51437886-514 |

| | | | | | | |
|---|---|---|---|---|---|---|
| 0 | 6.48369996480183 | 0.133 | 0.001 | 0 | T | chr17-65741126-657 |
| 0 | 6.60841520138182 | 0.287 | 0.003 | 0 | T | chr17-68721833-687 |
| 0 | 6.7308763234655 | 0.207 | 0.002 | 0 | T | chr17-72371767-723 |
| 0 | 6.52051924383312 | 0.269 | 0.003 | 0 | T | chr18-30952599-309 |
| 0 | 6.77837951429095 | 0.105 | 0.001 | 0 | T | chr18-33011071-330 |
| 0 | 6.60002512822423 | 0.148 | 0.002 | 0 | T | chr18-49894635-498 |
| 0 | 6.60712382221291 | 0.359 | 0.004 | 0 | T | chr18-61351037-613 |
| 0 | 6.55353884558726 | 0.299 | 0.003 | 0 | T | chr18-62306154-623 |
| 0 | 6.72299282490751 | 0.32 | 0.003 | 0 | T | chr18-62307893-623 |
| 0 | 6.66475546394047 | 0.238 | 0.003 | 0 | T | chr18-68165640-681 |
| 0 | 6.75193028457783 | 0.267 | 0.003 | 0 | T | chr18-74650195-746 |
| 0 | 6.68748462269394 | 0.276 | 0.003 | 0 | T | chr19-23633879-236 |
| 0 | 6.5041929382687 | 0.173 | 0.002 | 0 | T | chr19-34534689-345 |
| 0 | 6.71956390682976 | 0.243 | 0.003 | 0 | T | chr2-192620440-192 |
| 0 | 6.58718067523305 | 0.143 | 0.002 | 0 | T | chr2-192765387-192 |
| 0 | 7.01669408240173 | 0.259 | 0.002 | 0 | T | chr2-218018842-218 |
| 0 | 6.89772985393147 | 0.298 | 0.003 | 0 | T | chr2-238310799-238 |
| 0 | 6.96547509053864 | 0.255 | 0.002 | 0 | T | chr20-4526198-4526 |
| 0 | 6.56399590725798 | 0.338 | 0.004 | 0 | T | chr20-9084825-9085 |
| 0 | 6.70339118753689 | 0.294 | 0.004 | 0 | T | chr20-31799879-318 |
| 0 | 6.52013403349026 | 0.26 | 0.003 | 0 | T | chr20-45137904-451 |
| 0 | 6.50615692918517 | 0.209 | 0.002 | 0 | T | chr20-46753857-467 |
| 0 | 6.63111136577605 | 0.24 | 0.003 | 0 | T | chr20-47458857-474 |
| 0 | 6.48795758583277 | 0.173 | 0.002 | 0 | T | chr20-48467305-484 |
| 0 | 6.58579035629274 | 0.417 | 0.006 | 0 | T | chr3-12668345-1266 |
| 0 | 6.51432585241233 | 0.165 | 0.002 | 0 | T | chr3-29387515-2938 |
| 0 | 6.81596500704554 | 0.265 | 0.003 | 0 | T | chr3-33043969-3304 |
| 0 | 6.48629012172942 | 0.16 | 0.002 | 0 | T | chr3-88009062-8800 |
| 0 | 6.76763955911453 | 0.151 | 0.001 | 0 | T | chr3-88215709-8821 |
| 0 | 6.75745848308679 | 0.134 | 0.001 | 0 | T | chr3-95495805-9549 |
| 0 | 6.58869472352264 | 0.352 | 0.005 | 0 | T | chr3-113761224-113 |
| 0 | 6.67286087643026 | 0.281 | 0.003 | 0 | T | chr3-135401831-135 |
| 0 | 6.79815494143199 | 0.386 | 0.004 | 0 | T | chr3-151873960-151 |
| 0 | 6.72147198979348 | 0.173 | 0.002 | 0 | T | chr3-165277641-165 |
| 0 | 6.63562368887618 | 0.286 | 0.004 | 0 | T | chr3-167551327-167 |
| 0 | 6.71238106830363 | 0.344 | 0.004 | 0 | T | chr4-5511115-55116 |
| 0 | 7.26582552816762 | 0.223 | 0.002 | 0 | T | chr4-71343450-7134 |
| 0 | 6.78204843527584 | 0.33 | 0.003 | 0 | T | chr4-77546823-7754 |
| 0 | 6.91179682895601 | 0.361 | 0.004 | 0 | T | chr4-83704882-8370 |
| 0 | 6.64831259354683 | 0.287 | 0.003 | 0 | T | chr4-102697184-102 |
| 0 | 6.51169835641939 | 0.145 | 0.002 | 0 | T | chr4-115843482-115 |
| 0 | 6.65998021440122 | 0.179 | 0.002 | 0 | T | chr4-118197246-118 |
| 0 | 6.75872263150931 | 0.241 | 0.003 | 0 | T | chr4-122820501-122 |

| | | | | | | |
|---|---|---|---|---|---|---|
| 0 | 6.72008426077282 | 0.306 | 0.003 | 0 | T | chr4-129926709-129 |
| 0 | 6.63028191942546 | 0.313 | 0.004 | 0 | T | chr4-146324468-146 |
| 0 | 6.8460789316068 | 0.247 | 0.002 | 0 | T | chr4-156188776-156 |
| 0 | 6.82775402505063 | 0.289 | 0.003 | 0 | T | chr4-166749321-166 |
| 0 | 6.76296500102458 | 0.335 | 0.003 | 0 | T | chr4-173248354-173 |
| 0 | 6.63641371368976 | 0.104 | 0.001 | 0 | T | chr5-25036634-2503 |
| 0 | 6.80460495626099 | 0.294 | 0.003 | 0 | T | chr5-61216235-6121 |
| 0 | 6.51803092328312 | 0.116 | 0.001 | 0 | T | chr5-132780819-132 |
| 0 | 6.49600498548771 | 0.44 | 0.006 | 0 | T | chr5-138405633-138 |
| 0 | 6.7005545554854 | 0.209 | 0.002 | 0 | T | chr5-141269283-141 |
| 0 | 6.48838979662725 | 0.298 | 0.004 | 0 | T | chr5-151187152-151 |
| 0 | 6.61976380385075 | 0.241 | 0.002 | 0 | T | chr5-157048288-157 |
| 0 | 7.02382382904536 | 0.267 | 0.002 | 0 | T | chr6-14054192-1405 |
| 0 | 6.50968071738222 | 0.15 | 0.002 | 0 | T | chr6-52749385-5274 |
| 0 | 6.84056255082606 | 0.34 | 0.003 | 0 | T | chr6-70296713-7029 |
| 0 | 6.5304664369552 | 0.213 | 0.003 | 0 | T | chr6-71899773-7190 |
| 0 | 6.53229578518569 | 0.367 | 0.005 | 0 | T | chr6-92742956-9274 |
| 0 | 6.61697740748886 | 0.141 | 0.002 | 0 | T | chr6-137285495-137 |
| 0 | 6.83988085014912 | 0.451 | 0.005 | 0 | T | chr7-21343328-2134 |
| 0 | 6.63220049337067 | 0.223 | 0.003 | 0 | T | chr7-52558944-5255 |
| 0 | 6.6185024221893 | 0.386 | 0.005 | 0 | T | chr7-65476091-6547 |
| 0 | 6.95204098570766 | 0.315 | 0.003 | 0 | T | chr7-86653271-8665 |
| 0 | 6.70789972392604 | 0.444 | 0.005 | 0 | T | chr7-114518267-114 |
| 0 | 6.51924750197894 | 0.361 | 0.005 | 0 | T | chr7-120371943-120 |
| 0 | 7.17845872734276 | 0.238 | 0.002 | 0 | T | chr7-121879221-121 |
| 0 | 7.14152284093546 | 0.366 | 0.003 | 0 | T | chr8-23993155-2399 |
| 0 | 6.64998597821385 | 0.213 | 0.002 | 0 | T | chr8-25493962-2549 |
| 0 | 6.55938015053272 | 0.189 | 0.002 | 0 | T | chr8-52722967-5272 |
| 0 | 6.50192576502703 | 0.25 | 0.003 | 0 | T | chr8-106280358-106 |
| 0 | 6.60890041878885 | 0.395 | 0.004 | 0 | T | chr8-123395945-123 |
| 0 | 6.91325672494826 | 0.27 | 0.002 | 0 | T | chr9-3222700-32232 |
| 0 | 7.11156475445885 | 0.464 | 0.004 | 0 | T | chr9-15721378-1572 |
| 0 | 6.55945142002483 | 0.121 | 0.001 | 0 | T | chr9-36071953-3607 |
| 0 | 6.57998952421446 | 0.236 | 0.003 | 0 | T | chr9-39313903-3931 |
| 0 | 6.79081174442144 | 0.303 | 0.003 | 0 | T | chr9-81057580-8105 |
| 0 | 6.55935191565301 | 0.223 | 0.002 | 0 | T | chr9-87383638-8738 |
| 0 | 6.59028401616456 | 0.216 | 0.002 | 0 | T | chrX-72165197-7216 |
| 0 | 6.58637979003418 | 0.148 | 0.001 | 0 | T | chrX-100178649-100 |
| 2.12069007 | 3.56722407467296 | 0.18 | 0.029 | 1.5539695 | TAC | chr1-47800563-4780 |
| 2.42816047 | 2.26742675164393 | 0.233 | 0.051 | 1.7792734 | TAC | chr5-103097502-103 |
| 1.03753079 | 2.3408868524293 | 0.219 | 0.048 | 7.6026729 | TAC | chr17-42477932-424 |
| 1.27589459 | 3.00259557724201 | 0.116 | 0.016 | 9.3493217 | TAC | chr17-42484916-424 |
| 1.20993813 | 2.49602722074789 | 0.205 | 0.046 | 8.8660152 | TAC | chr2-183838714-183 |

| | | | | | | |
|---|---|---|---|---|---|---|
| 1.63982175 | 2.97277861543871 | 0.124 | 0.018 | 1.2016056 | TAC | chr17-41386108-413 |
| 3.07989567 | 2.30322342809564 | 0.213 | 0.051 | 2.2568428 | TAC | chr17-41380220-413 |
| 5.02538575 | 2.25182160785547 | 0.215 | 0.052 | 3.6824318 | TAC | chr3-156273140-156 |
| 2.41164604 | 2.53371906880979 | 0.151 | 0.028 | 1.7671722 | TAC | chr10-105446622-10 |
| 4.87733012 | 2.18170912209392 | 0.195 | 0.045 | 3.5739416 | TAC | chr5-103098110-103 |
| 1.42039677 | 2.73782296409688 | 0.128 | 0.022 | 1.0408184 | TAC | chr19-48334548-483 |
| 2.97162730 | 2.20569302880747 | 0.197 | 0.047 | 2.1775074 | TAC | chr10-80521889-805 |
| 1.08316619 | 2.23073798101083 | 0.189 | 0.044 | 7.9370736 | TAC | chr6-11444813-1144 |
| 4.00332818 | 2.90889729184829 | 0.111 | 0.018 | 2.9335027 | TAC | chr10-86801081-868 |
| 6.40388886 | 2.18500429763557 | 0.19 | 0.045 | 4.6925520 | TAC | chr17-41521048-415 |
| 9.10004718 | 2.75427931312283 | 0.104 | 0.015 | 6.6682051 | TAC | chr3-15182008-1518 |
| 6.56093817 | 2.88012878622588 | 0.103 | 0.015 | 4.8076324 | TAC | chr10-103280532-10 |
| 8.06419525 | 2.36403906065513 | 0.157 | 0.033 | 5.9091681 | TAC | chr8-33678604-3367 |
| 9.43508244 | 2.50298364196165 | 0.131 | 0.024 | 6.9137076 | TAC | chr12-14065804-140 |
| 6.60193254 | 2.18020864774928 | 0.192 | 0.047 | 4.8376717 | TAC | chr9-92214271-9221 |
| 3.68303945 | 2.50018944329334 | 0.118 | 0.021 | 2.6988060 | TAC | chr7-8796232-87967 |
| 2.91616611 | 2.41518761879323 | 0.135 | 0.027 | 2.1368673 | TAC | chr5-103096427-103 |
| 9.70966379 | 2.51956837606297 | 0.133 | 0.026 | 7.1149115 | TAC | chr1-96110001-9611 |
| 1.34706136 | 2.8412427513494 | 0.109 | 0.019 | 9.8708076 | TAC | chr10-83905963-839 |
| 2.81099843 | 2.5338953236024 | 0.113 | 0.02 | 2.0598040 | TAC | chr1-95661980-9566 |
| 3.50350988 | 2.63230771615552 | 0.121 | 0.022 | 2.5672529 | TAC | chr8-33679921-3368 |
| 2.06676741 | 2.76456134652773 | 0.109 | 0.019 | 1.5144568 | TAC | chr12-31581021-315 |
| 4.44012737 | 2.37535164700747 | 0.123 | 0.024 | 3.2535743 | TAC | chr10-80528193-805 |
| 7.27396808 | 2.53892461483558 | 0.129 | 0.026 | 5.3301164 | TAC | chr8-19477566-1947 |
| 5.63837588 | 2.59320970525872 | 0.118 | 0.022 | 4.1316101 | TAC | chr16-1314719-1315 |
| 1.74820950 | 2.18616367215724 | 0.163 | 0.039 | 1.2810284 | TAC | chr17-42799404-427 |
| 3.87691755 | 2.68868307382447 | 0.109 | 0.02 | 2.8408733 | TAC | chr10-83883274-838 |
| 1.24876072 | 2.29272441294505 | 0.137 | 0.029 | 9.1504940 | TAC | chr17-41535040-415 |
| 1.99069548 | 2.64071019989036 | 0.107 | 0.019 | 1.4587139 | TAC | chr9-15314934-1531 |
| 1.21507015 | 2.1385839944407 | 0.184 | 0.049 | 8.9036209 | TAC | chr5-130574623-130 |
| 2.22433401 | 2.40026174221832 | 0.132 | 0.028 | 1.6299163 | TAC | chr10-86798644-867 |
| 7.12391424 | 2.64253747653968 | 0.119 | 0.023 | 5.2201621 | TAC | chr20-6357508-6358 |
| 4.77855052 | 2.3992828153734 | 0.128 | 0.027 | 3.5015593 | TAC | chr10-80485169-804 |
| 1.56333380 | 2.61903975060001 | 0.101 | 0.018 | 1.1455578 | TAC | chr12-31520945-315 |
| 3.72804374 | 2.56362196968247 | 0.114 | 0.022 | 2.7317837 | TAC | chr20-6377680-6378 |
| 4.80391512 | 2.34656084232551 | 0.134 | 0.03 | 3.5201456 | TAC | chr10-83925510-839 |
| 6.74990192 | 2.34470799970144 | 0.144 | 0.034 | 4.9460986 | TAC | chr17-41525819-415 |
| 2.88245490 | 2.55421589067008 | 0.104 | 0.019 | 2.1121649 | TAC | chr4-104742920-104 |
| 1.07931298 | 2.25767682581784 | 0.148 | 0.036 | 7.9088386 | TAC | chr19-51573539-515 |
| 1.20196962 | 2.6658536248541 | 0.1 | 0.018 | 8.8076247 | TAC | chr12-31541149-315 |
| 7.42133079 | 2.35918469737762 | 0.111 | 0.022 | 5.4380988 | TAC | chr12-31533143-315 |
| 1.98202260 | 2.28441702452027 | 0.125 | 0.027 | 1.4523587 | TAC | chr8-107089666-107 |
| 5.58442654 | 2.22833829153128 | 0.131 | 0.03 | 4.0920778 | TAC | chr17-42486838-424 |

| | | | | | | | |
|---|---|---|---|---|---|---|---|
| 5.70190751 | 2.3590939016949 | | 0.131 | 0.03 | 4.1781639 | TAC | chr19-48326132-483 |
| 1.14355899 | 2.29829554220225 | | 0.131 | 0.03 | 8.3796114 | TAC | chr3-11162065-1116 |
| 1.21364218 | 2.34692243012661 | | 0.119 | 0.025 | 8.8931572 | TAC | chr5-149197961-149 |
| 9.73977758 | 2.51542400029703 | | 0.114 | 0.024 | 7.1369778 | TAC | chr10-87976856-879 |
| 1.32818477 | 2.18458860003975 | | 0.117 | 0.025 | 9.7324864 | TAC | chr10-102508177-10 |
| 1.07919101 | 2.11847874662006 | | 0.126 | 0.029 | 7.9079448 | TAC | chr1-153021928-153 |
| 3.53982868 | 2.10573634339848 | | 0.149 | 0.038 | 2.5938661 | TAC | chr16-18837708-188 |
| 5.84707006 | 2.43099677502087 | | 0.117 | 0.025 | 4.2845341 | TAC | chr7-9681109-96816 |
| 6.32020574 | 2.44570806865467 | | 0.123 | 0.028 | 4.6312318 | TAC | chr3-16032918-1603 |
| 2.98905174 | 2.28759802557947 | | 0.136 | 0.033 | 2.1902754 | TAC | chr16-18741967-187 |
| 4.64511991 | 2.34729520135068 | | 0.122 | 0.028 | 3.4037859 | TAC | chr12-14040677-140 |
| 1.42248601 | 2.34406500923103 | | 0.108 | 0.023 | 1.0423493 | TAC | chr7-109001108-109 |
| 2.03756985 | 2.31874178495452 | | 0.125 | 0.029 | 1.4930619 | TAC | chr10-13785223-137 |
| 5.24913939 | 2.32697362545579 | | 0.108 | 0.023 | 3.8463908 | TAC | chr2-174702610-174 |
| 1.87380200 | 2.26991689606379 | | 0.124 | 0.029 | 1.3730583 | TAC | chr2-174051842-174 |
| 6.38023062 | 2.1540035792538 | | 0.14 | 0.036 | 4.6752160 | TAC | chr12-41044122-410 |
| 1.05468226 | 2.49085250045506 | | 0.107 | 0.023 | 7.7283530 | TAC | chr1-153166370-153 |
| 1.10996731 | | 2.582288479 | 0.107 | 0.023 | 8.1334631 | TAC | chr7-110482157-110 |
| 1.32306409 | 2.24669207290049 | | 0.115 | 0.026 | 9.6949638 | TAC | chr19-51574462-515 |
| 3.90415146 | 2.44158570485518 | | 0.1 | 0.02 | 2.8608294 | TAC | chr3-156375854-156 |
| 5.31728142 | 2.44181279350858 | | 0.111 | 0.024 | 3.8963230 | TAC | chr10-105559312-10 |
| 1.24299540 | 2.27509582433014 | | 0.118 | 0.027 | 9.1082477 | TAC | chr5-153600507-153 |
| 3.13479757 | 2.33911396447133 | | 0.108 | 0.023 | 2.2970730 | TAC | chr9-92416149-9241 |
| 1.12602033 | 2.3395539208225 | | 0.105 | 0.022 | 8.2510941 | TAC | chr7-134169669-134 |
| 2.45251454 | 2.20625123514545 | | 0.122 | 0.029 | 1.7971192 | TAC | chr7-8279581-82800 |
| 4.12785894 | 2.31081665607552 | | 0.117 | 0.027 | 3.0247546 | TAC | chr5-130577365-130 |
| 4.83407661 | 2.09878546336155 | | 0.125 | 0.031 | 3.5422469 | TAC | chr10-87171788-871 |
| 9.08293136 | 2.12804652987205 | | 0.142 | 0.038 | 6.6556632 | TAC | chr6-104832164-104 |
| 1.24633031 | 2.23270840240563 | | 0.119 | 0.028 | 9.1326847 | TAC | chr1-166617725-166 |
| 2.13527077 | 2.32850639752635 | | 0.111 | 0.025 | 1.5646538 | TAC | chr7-56220417-5622 |
| 2.97612640 | 2.17454336944118 | | 0.104 | 0.022 | 2.1808042 | TAC | chr4-83023343-8302 |
| 4.97202859 | 2.11810860871208 | | 0.12 | 0.029 | 3.6433335 | TAC | chr19-50685929-506 |
| 7.29352142 | 2.38340534145067 | | 0.113 | 0.026 | 5.3444445 | TAC | chr12-31504697-315 |
| 1.16716378 | 2.08554549094647 | | 0.131 | 0.033 | 8.5525793 | TAC | chr10-13749815-137 |
| 1.47051305 | 2.20219200676258 | | 0.109 | 0.025 | 1.0775419 | TAC | chr8-33732768-3373 |
| 3.97627231 | 2.14031742244922 | | 0.133 | 0.034 | 2.9136771 | TAC | chr7-134253502-134 |
| 1.57413192 | 2.20101747543662 | | 0.118 | 0.029 | 1.1534703 | TAC | chr6-10481427-1048 |
| 1.86820266 | 2.33667504593486 | | 0.113 | 0.027 | 1.3689553 | TAC | chr1-166592476-166 |
| 2.04440345 | 2.41636308534535 | | 0.104 | 0.023 | 1.4980693 | TAC | chr1-240626120-240 |
| 5.73518843 | 2.12415810954624 | | 0.108 | 0.025 | 4.2025510 | TAC | chr6-98480820-9848 |
| 6.80047250 | 2.32062457978981 | | 0.102 | 0.022 | 4.9831550 | TAC | chr2-174734078-174 |
| 8.65372098 | 2.36384509506871 | | 0.108 | 0.025 | 6.3411525 | TAC | chr10-102532881-10 |
| 8.68598142 | 2.1310863735422 | | 0.106 | 0.024 | 6.3647918 | TAC | chr4-59319985-5932 |

| | | | | | | |
|---|---|---|---|---|---|---|
| 1.14016340 | 2.24247016359709 | 0.106 | 0.024 | 8.3547298 | TAC | chr1-204330669-204 |
| 4.98337766 | 2.07984971619671 | 0.132 | 0.035 | 3.6516497 | TAC | chr12-34481854-344 |
| 1.06645576 | 2.09481306160544 | 0.109 | 0.026 | 7.8146252 | TAC | chr7-8771770-87722 |
| 1.32451450 | 2.23527428899642 | 0.107 | 0.025 | 9.7055919 | TAC | chr4-59194875-5919 |
| 1.35422792 | 2.11039004046869 | 0.109 | 0.026 | 9.9233218 | TAC | chr20-29769276-297 |
| 1.55941917 | 2.26696405851156 | 0.111 | 0.027 | 1.1426893 | TAC | chr7-134256593-134 |
| 2.62519519 | 2.16648258305629 | 0.13 | 0.035 | 1.9236537 | TAC | chr10-83915678-839 |
| 3.27951160 | 2.09075500129849 | 0.111 | 0.027 | 2.4031146 | TAC | chr8-103374350-103 |
| 4.56385086 | 2.41416054297424 | 0.1 | 0.022 | 3.3442347 | TAC | chr1-77523067-7752 |
| 2.33518321 | 2.23332815752189 | 0.11 | 0.027 | 1.7111428 | TAC | chr17-42698534-426 |
| 2.41610189 | 2.17750701457046 | 0.117 | 0.029 | 1.7704373 | TAC | chr5-153545065-153 |
| 5.61455374 | 2.1279762202191 | 0.125 | 0.033 | 4.1141540 | TAC | chr7-134196267-134 |
| 6.32575704 | 2.19294080645176 | 0.112 | 0.027 | 4.6352996 | TAC | chr19-48336859-483 |
| 9.25285252 | 2.09790276352613 | 0.132 | 0.036 | 6.7801757 | TAC | chr1-95677298-9567 |
| 1.04090328 | 2.18528178014944 | 0.121 | 0.031 | 7.6273853 | TAC | chr12-21986389-219 |
| 1.12150898 | 2.19299602457624 | 0.108 | 0.026 | 8.2180365 | TAC | chr10-86772504-867 |
| 1.30732136 | 2.28147949873929 | 0.1 | 0.023 | 9.5796064 | TAC | chr7-9628209-96287 |
| 2.52783682 | 2.11570650562394 | 0.109 | 0.026 | 1.8523128 | TAC | chr18-56328732-563 |
| 4.32864085 | 2.22465783969188 | 0.126 | 0.034 | 3.1718808 | TAC | chr6-98945924-9894 |
| 4.72826710 | 2.30913741896564 | 0.112 | 0.028 | 3.4647133 | TAC | chr7-8283815-82843 |
| 5.42400739 | 2.14827422695102 | 0.12 | 0.031 | 3.9745282 | TAC | chr2-173991745-173 |
| 5.52490896 | 2.16678341763842 | 0.109 | 0.027 | 4.0484654 | TAC | chr8-118733114-118 |
| 5.80162792 | 2.38600858183494 | 0.108 | 0.026 | 4.2512356 | TAC | chr8-73475347-7347 |
| 3.00032536 | 2.0801026110454 | 0.122 | 0.032 | 2.1985364 | TAC | chr3-156368215-156 |
| 7.64509921 | 2.10581107900951 | 0.105 | 0.025 | 5.6020687 | TAC | chr10-84024669-840 |
| 1.66527612 | 2.13927079318164 | 0.115 | 0.029 | 1.2202577 | TAC | chr3-79711034-7971 |
| 1.24347352 | 2.16047184552306 | 0.119 | 0.032 | 9.1117512 | TAC | chr1-47802557-4780 |
| 2.45073914 | 2.21958634483653 | 0.105 | 0.026 | 1.7958183 | TAC | chr10-104472843-10 |
| 5.15025737 | 2.2299732914084 | 0.101 | 0.024 | 3.7739334 | TAC | chr3-155972177-155 |
| 8.19216710 | 2.09337146904831 | 0.109 | 0.028 | 6.0029415 | TAC | chr11-79768192-797 |
| 2.01631036 | 2.08483151255421 | 0.106 | 0.026 | 1.4774836 | TAC | chr10-83726025-837 |
| 6.77898036 | 2.10660160040083 | 0.122 | 0.033 | 4.9674063 | TAC | chr6-131332685-131 |
| 7.26503749 | 2.15936896052575 | 0.125 | 0.035 | 5.3235724 | TAC | chr12-32582293-325 |
| 1.12799406 | 2.12484350494511 | 0.101 | 0.024 | 8.2655569 | TAC | chr3-15703083-1570 |
| 1.72646421 | 2.18954894249019 | 0.109 | 0.028 | 1.2650942 | TAC | chr1-197729863-197 |
| 1.84882788 | 2.14591588545949 | 0.105 | 0.026 | 1.3547582 | TAC | chr12-31603625-316 |
| 5.92471932 | 2.47621376850055 | 0.109 | 0.029 | 4.3414328 | TAC | chr7-8632449-86329 |
| 6.15059600 | 2.23121675283881 | 0.119 | 0.033 | 4.5069476 | TAC | chr16-19110782-191 |
| 1.59069000 | 2.13700337979012 | 0.113 | 0.03 | 1.1656035 | TAC | chr20-7379874-7380 |
| 1.74711481 | 2.16279934787106 | 0.102 | 0.025 | 1.2802263 | TAC | chr10-80570149-805 |
| 2.73943036 | 2.21265168715805 | 0.1 | 0.025 | 2.0073614 | TAC | chr4-127341785-127 |
| 1.02801789 | 2.10163046718645 | 0.105 | 0.027 | 7.5329656 | TAC | chr18-56365932-563 |
| 1.05762937 | 2.13094527341046 | 0.107 | 0.028 | 7.7499484 | TAC | chr5-160849597-160 |

| | | | | | | | |
|---|---|---|---|---|---|---|---|
| 1.82234501 | 2.08706292348284 | | 0.12 | 0.034 | 1.3353524 | TAC | chr10-83917143-839 |
| 2.03861581 | 2.31074749458232 | | 0.108 | 0.028 | 1.4938283 | TAC | chr19-50562525-505 |
| 4.94835475 | 2.19042764341694 | | 0.108 | 0.029 | 3.6259861 | TAC | chr1-77523621-7752 |
| 5.24326298 | 2.13435032398016 | | 0.101 | 0.025 | 3.8420848 | TAC | chr20-6319027-6319 |
| 8.28457758 | 2.30547973123343 | | 0.105 | 0.027 | 6.0706567 | TAC | chr7-9439064-94395 |
| 5.05332658 | 2.22475091104019 | | 0.104 | 0.027 | 3.7029059 | TAC | chr7-116377886-116 |
| 9.08020986 | 2.0823465588199 | | 0.114 | 0.032 | 6.6536690 | TAC | chr1-95508745-9550 |
| 1.70573824 | 2.09388315310675 | | 0.113 | 0.031 | 1.2499069 | TAC | chr7-9353050-93535 |
| 1.33581595 | | 2.135782077 | 0.105 | 0.028 | 9.7884051 | TAC | chr12-14153420-141 |
| 6.78634288 | 2.10212360918151 | | 0.108 | 0.029 | 4.9728013 | TAC | chr12-11595259-115 |
| 9.05759339 | 2.10799545557577 | | 0.101 | 0.027 | 6.6370964 | TAC | chr17-73930128-739 |
| 5.98095590 | 2.08453933229263 | | 0.103 | 0.029 | 4.3826411 | TAC | chr16-1294751-1295 |
| 1.69271182 | 2.09938908239373 | | 0.1 | 0.027 | 1.2403616 | TAC | chr14-80021196-800 |
| 8.78093925 | 2.13612760300522 | | 0.102 | 0.03 | 6.4343737 | TAC | chr20-2904320-2904 |
| 1.53919847 | 2.19366314498657 | | 0.1 | 0.029 | 1.1278723 | TAC | chr8-118569380-118 |
| 3.98030092 | 2.10574997747426 | | 0.106 | 0.032 | 2.9166291 | TAC | chr10-102308950-10 |
| 1.31235289 | 2.93113962884588 | | 0.285 | 0.046 | 9.6164758 | basalKC | chr2-183838714-183 |
| 1.88731939 | 2.36407728813823 | | 0.237 | 0.053 | 1.3829634 | basalKC | chr17-41380220-413 |
| 1.33989312 | 1.90058541216775 | | 0.462 | 0.178 | 9.8182812 | basalKC | chr2-183831497-183 |
| 5.72202005 | 2.43330011229054 | | 0.213 | 0.05 | 4.1929017 | basalKC | chr17-42477932-424 |
| 1.36235635 | 1.86053711423637 | | 0.288 | 0.087 | 9.9828841 | basalKC | chr9-1134849-11353 |
| 1.03629754 | 2.34627384383686 | | 0.163 | 0.036 | 7.5936360 | basalKC | chr17-41569902-415 |
| 1.78906302 | 2.45996368772713 | | 0.147 | 0.031 | 1.3109645 | basalKC | chr1-47800563-4780 |
| 4.90285688 | 1.71095618238665 | | 0.298 | 0.101 | 3.5926468 | basalKC | chr20-6316683-6317 |
| 5.49626947 | 2.31545996959108 | | 0.187 | 0.047 | 4.0274793 | basalKC | chr17-41521048-415 |
| 4.84586596 | 2.60918488315031 | | 0.108 | 0.018 | 3.5508858 | basalKC | chr2-183830767-183 |
| 2.52758184 | 1.85007740250124 | | 0.263 | 0.085 | 1.8521260 | basalKC | chr17-41425540-414 |
| 8.32249896 | 2.34217370306984 | | 0.139 | 0.031 | 6.0984442 | basalKC | chr17-41535040-415 |
| 4.41481657 | 2.30698666737046 | | 0.125 | 0.026 | 3.2350274 | basalKC | chr20-29769276-297 |
| 5.26725956 | 2.19916534593481 | | 0.161 | 0.041 | 3.8596687 | basalKC | chr17-42799404-427 |
| 6.48289548 | 1.84230881407107 | | 0.191 | 0.054 | 4.7504453 | basalKC | chr5-103097502-103 |
| 5.39472576 | 1.79026486810808 | | 0.224 | 0.071 | 3.9530716 | basalKC | chr17-42707811-427 |
| 1.30009971 | 1.96115103001202 | | 0.163 | 0.042 | 9.5266887 | basalKC | chr19-15700263-157 |
| 1.40410730 | 2.03940392794636 | | 0.169 | 0.045 | 1.0288820 | basalKC | chr1-47735193-4773 |
| 2.99201105 | 1.75799814378348 | | 0.219 | 0.071 | 2.1924439 | basalKC | chr17-41574601-415 |
| 4.22092300 | 2.27552056437476 | | 0.139 | 0.035 | 3.0929488 | basalKC | chr4-147657441-147 |
| 6.19439423 | 2.17683430080728 | | 0.122 | 0.028 | 4.5390414 | basalKC | chr5-103096427-103 |
| 7.21857114 | 2.00353662993306 | | 0.138 | 0.034 | 5.2895235 | basalKC | chr17-41378027-413 |
| 1.40531291 | 1.74980152265616 | | 0.182 | 0.054 | 1.0297655 | basalKC | chr10-102701041-10 |
| 2.76438855 | 2.13043639076251 | | 0.132 | 0.032 | 2.0256499 | basalKC | chr20-17556467-175 |
| 1.28791198 | 1.77349588186461 | | 0.208 | 0.069 | 9.4373811 | basalKC | chr13-47686472-476 |
| 1.20783364 | 2.02200429203448 | | 0.155 | 0.043 | 8.8505942 | basalKC | chr7-29417687-2941 |
| 2.18456675 | 1.98184027400008 | | 0.143 | 0.038 | 1.6007762 | basalKC | chr20-3541669-3542 |

| | | | | | | |
|---|---|---|---|---|---|---|
| 1.19178523 | 1.8197304862194 | 0.163 | 0.048 | 8.7329970 | basalKC | chr18-54076570-540 |
| 1.76104530 | 1.70134798127669 | 0.229 | 0.084 | 1.2904341 | basalKC | chr19-18497393-184 |
| 3.01775537 | 1.91501912790398 | 0.201 | 0.069 | 2.2113085 | basalKC | chr4-133402319-133 |
| 3.04970239 | 1.71524453020569 | 0.191 | 0.063 | 2.2347182 | basalKC | chr6-41338816-4133 |
| 4.14555250 | 1.70061396453744 | 0.188 | 0.061 | 3.0377199 | basalKC | chr18-63213345-632 |
| 1.08510311 | 1.78995578714301 | 0.171 | 0.053 | 7.9512666 | basalKC | chr18-50148547-501 |
| 3.44481122 | 2.40321406922798 | 0.102 | 0.023 | 2.5242405 | basalKC | chr17-19848792-198 |
| 8.47139301 | 2.24801951557424 | 0.119 | 0.03 | 6.2075487 | basalKC | chr19-22487924-224 |
| 1.55525707 | 1.75244639528301 | 0.16 | 0.048 | 1.1396395 | basalKC | chr7-133861299-133 |
| 5.72215553 | 1.79366440614387 | 0.165 | 0.051 | 4.1930010 | basalKC | chr10-94114775-941 |
| 1.94189447 | 1.9790455515608 | 0.124 | 0.033 | 1.4229542 | basalKC | chr6-87643282-8764 |
| 4.86731144 | 2.0357040154532 | 0.124 | 0.033 | 3.5666003 | basalKC | chr7-446683-447184 |
| 1.35383285 | 1.77289600261892 | 0.161 | 0.051 | 9.9204268 | basalKC | chr3-61021762-6102 |
| 1.78713208 | 1.76005628223687 | 0.157 | 0.049 | 1.3095496 | basalKC | chr1-85966531-8596 |
| 4.60596340 | 1.71237843222146 | 0.171 | 0.056 | 3.3750933 | basalKC | chr7-134058109-134 |
| 7.05022287 | 1.90565424843693 | 0.174 | 0.059 | 5.1661636 | basalKC | chr10-90460397-904 |
| 2.55453787 | 2.06234145845731 | 0.127 | 0.035 | 1.8718784 | basalKC | chr17-41525819-415 |
| 6.88828872 | 1.78721148998689 | 0.16 | 0.052 | 5.0475037 | basalKC | chr5-145480624-145 |
| 1.19432640 | 1.86631217688951 | 0.133 | 0.039 | 8.7516178 | basalKC | chr4-127395831-127 |
| 1.25129284 | 1.8185508094936 | 0.172 | 0.059 | 9.1690485 | basalKC | chr8-88885499-8888 |
| 1.85507054 | 1.96290059766063 | 0.119 | 0.032 | 1.3593326 | basalKC | chr4-50820621-5082 |
| 2.01529179 | 1.72868273100285 | 0.174 | 0.06 | 1.4767373 | basalKC | chr5-99044558-9904 |
| 2.05791967 | 1.75617287725011 | 0.139 | 0.042 | 1.5079735 | basalKC | chr20-3566545-3567 |
| 2.21382369 | 1.90881594869171 | 0.124 | 0.034 | 1.6222147 | basalKC | chr6-131332685-131 |
| 3.08048173 | 2.15749538445589 | 0.118 | 0.032 | 2.2572722 | basalKC | chr20-3308264-3308 |
| 3.17065506 | 1.77060849707561 | 0.125 | 0.035 | 2.3233482 | basalKC | chr8-33678604-3367 |
| 5.91021057 | 1.7818305324035 | 0.141 | 0.043 | 4.3308013 | basalKC | chr8-33716843-3371 |
| 6.37844263 | 1.81481248988222 | 0.135 | 0.04 | 4.6739058 | basalKC | chr14-96387952-963 |
| 6.61700165 | 1.9408246817144 | 0.116 | 0.031 | 4.8487138 | basalKC | chr17-42486838-424 |
| 8.73482820 | 1.80339587576774 | 0.129 | 0.037 | 6.4005851 | basalKC | chr15-40022778-400 |
| 1.24893861 | 1.71337448037339 | 0.16 | 0.053 | 9.1517975 | basalKC | chr18-48372338-483 |
| 1.25084990 | 1.95455347223952 | 0.103 | 0.026 | 9.1658027 | basalKC | chr5-56955433-5695 |
| 2.17686318 | 2.23432551826801 | 0.119 | 0.033 | 1.5951313 | basalKC | chr18-56329313-563 |
| 5.25251378 | 1.7741950297229 | 0.147 | 0.047 | 3.8488635 | basalKC | chr1-120703176-120 |
| 6.51958725 | 2.19846183656249 | 0.103 | 0.027 | 4.7773318 | basalKC | chr3-143493860-143 |
| 9.13778291 | 1.90325065392969 | 0.11 | 0.029 | 6.6958566 | basalKC | chr10-61876340-618 |
| 1.31724238 | 1.71444970526157 | 0.161 | 0.055 | 9.6523043 | basalKC | chr1-2689238-26897 |
| 2.42378536 | 1.91033693949006 | 0.119 | 0.034 | 1.7760675 | basalKC | chr4-79744360-7974 |
| 2.66469989 | 1.87166770957753 | 0.183 | 0.068 | 1.9526014 | basalKC | chr8-108394147-108 |
| 3.19773478 | 1.8827147892319 | 0.136 | 0.042 | 2.3431913 | basalKC | chr3-18481879-1848 |
| 5.75763252 | 1.72372890030343 | 0.154 | 0.051 | 4.2189973 | basalKC | chr14-83533120-835 |
| 5.88446269 | 2.05675090524147 | 0.105 | 0.028 | 4.3119341 | basalKC | chr19-22875167-228 |
| 6.60343138 | 2.01568886952719 | 0.105 | 0.028 | 4.8387700 | basalKC | chr4-82902809-8290 |

| | | | | | | | |
|---|---|---|---|---|---|---|---|
| 1.11525805 | 1.80295053956543 | | 0.127 | 0.038 | 8.1722318 | basalKC | chr1-203378884-203 |
| 1.13114827 | 2.04451375079064 | | 0.108 | 0.029 | 8.2886699 | basalKC | chr19-13780469-137 |
| 1.46957080 | 1.71239316137938 | | 0.165 | 0.057 | 1.0768515 | basalKC | chr17-42706910-427 |
| 2.17454024 | | 1.790916558 | 0.147 | 0.048 | 1.5934291 | basalKC | chr9-92182176-9218 |
| 1.64402801 | 1.97423408932814 | | 0.102 | 0.027 | 1.2046878 | basalKC | chr4-146038439-146 |
| 1.90997899 | 1.71106348517808 | | 0.135 | 0.043 | 1.3995676 | basalKC | chr9-75349944-7535 |
| 3.08718621 | 1.96914410560403 | | 0.105 | 0.029 | 2.2621850 | basalKC | chr2-35282576-3528 |
| 3.23476062 | 1.75493894783467 | | 0.136 | 0.044 | 2.3703226 | basalKC | chr1-187835945-187 |
| 3.39268235 | 2.10528161525933 | | 0.133 | 0.043 | 2.4860422 | basalKC | chr19-22876254-228 |
| 3.64553464 | 2.02432305010964 | | 0.122 | 0.037 | 2.6713238 | basalKC | chr7-120272256-120 |
| 3.96431854 | 1.86866764534479 | | 0.121 | 0.036 | 2.9049178 | basalKC | chr17-8696113-8696 |
| 5.93867486 | 1.98128115908701 | | 0.11 | 0.031 | 4.3516590 | basalKC | chr2-243499040-243 |
| 7.17132272 | 1.87362261753511 | | 0.136 | 0.044 | 5.2549014 | basalKC | chr15-2309307-2309 |
| 9.56282622 | 1.94449294637076 | | 0.102 | 0.028 | 7.0073139 | basalKC | chr17-42698534-426 |
| 1.32391062 | 1.80630749025088 | | 0.111 | 0.032 | 9.7011669 | basalKC | chr10-98504612-985 |
| 1.43445463 | 1.75148974045849 | | 0.129 | 0.04 | 1.0511195 | basalKC | chr10-79836070-798 |
| 1.62653848 | 1.82863808215286 | | 0.155 | 0.055 | 1.1918721 | basalKC | chr7-116278504-116 |
| 3.05543377 | 1.76816811806601 | | 0.129 | 0.041 | 2.2389179 | basalKC | chr19-13766304-137 |
| 3.41514468 | 1.71633300411059 | | 0.122 | 0.037 | 2.5025019 | basalKC | chr8-119719505-119 |
| 3.88598992 | 2.03336522002449 | | 0.103 | 0.029 | 2.8475212 | basalKC | chr14-75282234-752 |
| 5.29916202 | 1.94337654443247 | | 0.102 | 0.028 | 3.8830457 | basalKC | chr7-120224932-120 |
| 7.01354202 | 1.77399258041532 | | 0.141 | 0.047 | 5.1392851 | basalKC | chr13-74501165-745 |
| 1.14115892 | 1.85129615600235 | | 0.168 | 0.063 | 8.3620245 | basalKC | chr19-21922169-219 |
| 1.24263068 | 1.92653334050554 | | 0.111 | 0.033 | 9.1055751 | basalKC | chr19-22487224-224 |
| 1.74675402 | 1.74438899742445 | | 0.13 | 0.042 | 1.2799619 | basalKC | chr8-30619886-3062 |
| 2.22935299 | 2.01867878559912 | | 0.108 | 0.032 | 1.6335940 | basalKC | chr19-48326132-483 |
| 4.61059693 | 1.71831731992531 | | 0.141 | 0.048 | 3.3784886 | basalKC | chr2-33543725-3354 |
| 6.93291033 | 1.8621862691557 | | 0.105 | 0.03 | 5.0802009 | basalKC | chr1-135607305-135 |
| 9.02688351 | 1.87588726400533 | | 0.119 | 0.037 | 6.6145933 | basalKC | chr2-120752828-120 |
| 1.02149805 | 1.92903470700851 | | 0.11 | 0.033 | 7.4851904 | basalKC | chr1-47802557-4780 |
| 1.65721965 | 1.93700318775398 | | 0.124 | 0.04 | 1.2143542 | basalKC | chr19-18752687-187 |
| 2.57169804 | 1.72996503290017 | | 0.121 | 0.038 | 1.8844528 | basalKC | chr20-3405758-3406 |
| 5.86138839 | 1.82846051884661 | | 0.114 | 0.035 | 4.2950261 | basalKC | chr4-10423784-1042 |
| 6.95949519 | 1.85066066348288 | | 0.135 | 0.046 | 5.0996814 | basalKC | chr8-110736785-110 |
| 7.72739684 | 1.76992663813417 | | 0.121 | 0.039 | 5.6623736 | basalKC | chr6-70231666-7023 |
| 9.57310981 | | 1.820529312 | 0.15 | 0.055 | 7.0148493 | basalKC | chr5-122289774-122 |
| 1.33272714 | 2.05545393128145 | | 0.102 | 0.03 | 9.7657713 | basalKC | chr9-90095410-9009 |
| 1.37964121 | 1.71053542198868 | | 0.107 | 0.032 | 1.0109541 | basalKC | chr17-1157271-1157 |
| 1.58432455 | 1.88168715239819 | | 0.1 | 0.029 | 1.1609391 | basalKC | chr1-223244062-223 |
| 2.94331865 | 1.81137862814423 | | 0.125 | 0.042 | 2.1567638 | basalKC | chr8-68151851-6815 |
| 2.96318297 | 1.80230588262644 | | 0.118 | 0.038 | 2.1713197 | basalKC | chr10-43732643-437 |
| 8.37153543 | 1.82713247904636 | | 0.121 | 0.04 | 6.1343765 | basalKC | chr3-156606613-156 |
| 8.85988930 | 1.9417799265878 | | 0.1 | 0.03 | 6.4922256 | basalKC | chr19-55252929-552 |

| | | | | | | |
|---|---|---|---|---|---|---|
| 9.59869228 | 1.82972243598091 | 0.119 | 0.039 | 7.0335953 | basalKC | chr3-8194118-81946 |
| 1.29391051 | 1.7795051169751 | 0.108 | 0.033 | 9.4813363 | basalKC | chr2-61552783-6155 |
| 3.00679350 | 1.78971426310695 | 0.108 | 0.034 | 2.2032760 | basalKC | chr10-19608632-196 |
| 4.20414848 | 1.78597320393399 | 0.121 | 0.04 | 3.0806570 | basalKC | chr6-10226590-1022 |
| 7.65169511 | 1.90695626389807 | 0.118 | 0.039 | 5.6069020 | basalKC | chr9-14911303-1491 |
| 7.80444270 | 1.84378381465136 | 0.107 | 0.033 | 5.7188302 | basalKC | chr11-70580946-705 |
| 1.11644159 | 1.76466774112008 | 0.103 | 0.032 | 8.1809044 | basalKC | chr11-11431646-114 |
| 1.45211995 | 1.8314379406097 | 0.105 | 0.033 | 1.0640641 | basalKC | chr19-13775647-137 |
| 1.78499170 | 1.74796117615355 | 0.103 | 0.032 | 1.3079812 | basalKC | chr14-93887307-938 |
| 2.08723347 | 1.91845119652272 | 0.1 | 0.031 | 1.5294537 | basalKC | chr3-79711034-7971 |
| 3.95739666 | 1.70521092800895 | 0.102 | 0.032 | 2.8998457 | basalKC | chr1-242776854-242 |
| 4.09284220 | 1.81168369501271 | 0.114 | 0.038 | 2.9990956 | basalKC | chr9-42270283-4227 |
| 4.37136133 | 1.80592052041017 | 0.114 | 0.038 | 3.2031849 | basalKC | chr7-120571227-120 |
| 6.69989667 | 1.72557989402116 | 0.124 | 0.043 | 4.9094564 | basalKC | chr3-156610283-156 |
| 7.31673075 | 1.8147024100464 | 0.102 | 0.032 | 5.3614515 | basalKC | chr5-131804628-131 |
| 9.99337565 | 2.00696661575576 | 0.1 | 0.031 | 7.3228059 | basalKC | chr6-126599500-126 |
| 1.45620646 | 1.76747303245434 | 0.103 | 0.033 | 1.0670585 | basalKC | chr3-134769600-134 |
| 1.91732841 | 1.80630749025088 | 0.103 | 0.033 | 1.4049530 | basalKC | chr2-30562796-3056 |
| 2.08020792 | 1.75502751405594 | 0.102 | 0.032 | 1.5243056 | basalKC | chr13-69086283-690 |
| 2.29486924 | 1.99836788512761 | 0.118 | 0.041 | 1.6816021 | basalKC | chr19-22875678-228 |
| 2.51911772 | 1.79759238342333 | 0.107 | 0.035 | 1.8459238 | basalKC | chr5-152199010-152 |
| 5.97244939 | 1.83417519275765 | 0.1 | 0.032 | 4.3764078 | basalKC | chr10-90126937-901 |
| 6.58407031 | 1.83478608693142 | 0.102 | 0.033 | 4.8245828 | basalKC | chr10-93605574-936 |
| 1.08753508 | 1.75999213020598 | 0.113 | 0.039 | 7.9690873 | basalKC | chr17-19795073-197 |
| 1.58622410 | 1.87207591851606 | 0.1 | 0.032 | 1.1623310 | basalKC | chr19-49484287-494 |
| 1.84875488 | 1.8501145497429 | 0.102 | 0.033 | 1.3547047 | basalKC | chr19-13755799-137 |
| 6.45806268 | 1.70682187162466 | 0.1 | 0.033 | 4.7322487 | basalKC | chr7-22166452-2216 |
| 6.50610147 | 1.79685516642965 | 0.1 | 0.033 | 4.7674499 | basalKC | chr9-60776215-6077 |
| 8.44946990 | 1.77051336079248 | 0.103 | 0.034 | 6.1914842 | basalKC | chr19-13779906-137 |
| 1.03608779 | 1.74617221448716 | 0.108 | 0.037 | 7.5920990 | basalKC | chr1-31314095-3131 |
| 1.85472385 | 1.75722133162598 | 0.1 | 0.033 | 1.3590785 | basalKC | chr20-3218901-3219 |
| 2.48395835 | 1.70294454501326 | 0.103 | 0.036 | 1.8201602 | basalKC | chr7-134253502-134 |
| 9.58373104 | 1.74091820566702 | 0.105 | 0.038 | 7.0226322 | basalKC | chr1-87718814-8771 |
| 1.93935096 | 1.70775102252613 | 0.105 | 0.039 | 1.4210904 | basalKC | chr9-102478427-102 |
| 5.27744422 | 1.70689478558478 | 0.118 | 0.047 | 3.8671316 | basalKC | chr19-28582567-285 |
| 1.63245221 | 1.70634090659829 | 0.114 | 0.046 | 1.1962054 | basalKC | chr5-154369291-154 |
| 2.86972421 | 1.72673150428604 | 0.103 | 0.039 | 2.1028363 | basalKC | chr4-123694457-123 |
| 0 | 4.59966183092451 | 0.105 | 0.004 | 0 | facsFB | chr1-13077185-1307 |
| 0 | 4.3945107220713 | 0.123 | 0.007 | 0 | facsFB | chr1-20093174-2009 |
| 0 | 5.00069667414454 | 0.124 | 0.004 | 0 | facsFB | chr1-31946882-3194 |
| 0 | 4.71672693581356 | 0.162 | 0.006 | 0 | facsFB | chr1-47553193-4755 |
| 0 | 4.40821321735248 | 0.168 | 0.01 | 0 | facsFB | chr1-198517753-198 |
| 0 | 4.57863343895549 | 0.225 | 0.012 | 0 | facsFB | chr1-207674964-207 |

| | | | | | | |
|---|---|---|---|---|---|---|
| 0 | 4.72272992070722 | 0.11 | 0.004 | 0 | facsFB | chr1-212561910-212 |
| 0 | 4.94187261130502 | 0.146 | 0.005 | 0 | facsFB | chr1-225755224-225 |
| 0 | 4.50350478411028 | 0.144 | 0.007 | 0 | facsFB | chr1-225924541-225 |
| 0 | 4.42167657703132 | 0.127 | 0.006 | 0 | facsFB | chr10-32082414-320 |
| 0 | 4.26455420629279 | 0.157 | 0.009 | 0 | facsFB | chr10-34067534-340 |
| 0 | 4.27847352675718 | 0.116 | 0.006 | 0 | facsFB | chr10-51107760-511 |
| 0 | 4.57454625817421 | 0.111 | 0.005 | 0 | facsFB | chr10-56638802-566 |
| 0 | 4.27539685539986 | 0.141 | 0.008 | 0 | facsFB | chr11-16146275-161 |
| 0 | 4.2723954571236 | 0.16 | 0.008 | 0 | facsFB | chr11-21539437-215 |
| 0 | 4.63808365623534 | 0.12 | 0.005 | 0 | facsFB | chr11-23041821-230 |
| 0 | 4.35912442870216 | 0.145 | 0.008 | 0 | facsFB | chr11-25232526-252 |
| 0 | 4.45356873601657 | 0.157 | 0.007 | 0 | facsFB | chr11-25327779-253 |
| 0 | 4.59310313176937 | 0.104 | 0.005 | 0 | facsFB | chr11-58389015-583 |
| 0 | 4.70864650342139 | 0.131 | 0.006 | 0 | facsFB | chr11-75369229-753 |
| 0 | 5.00483795793186 | 0.103 | 0.004 | 0 | facsFB | chr11-84133919-841 |
| 0 | 4.34646939815274 | 0.242 | 0.014 | 0 | facsFB | chr12-27299470-272 |
| 0 | 4.42504195288774 | 0.153 | 0.007 | 0 | facsFB | chr13-35330648-353 |
| 0 | 4.44652531264759 | 0.105 | 0.005 | 0 | facsFB | chr13-81235615-812 |
| 0 | 4.60218276141644 | 0.17 | 0.008 | 0 | facsFB | chr13-82408460-824 |
| 0 | 4.34091091584655 | 0.124 | 0.007 | 0 | facsFB | chr13-95218355-952 |
| 0 | 4.53962672222111 | 0.116 | 0.005 | 0 | facsFB | chr14-14788856-147 |
| 0 | 4.61568785831369 | 0.104 | 0.004 | 0 | facsFB | chr14-82309947-823 |
| 0 | 4.87076891564209 | 0.113 | 0.005 | 0 | facsFB | chr14-88549291-885 |
| 0 | 4.452373619 | 0.108 | 0.005 | 0 | facsFB | chr14-102514367-10 |
| 0 | 4.611432232 | 0.138 | 0.006 | 0 | facsFB | chr15-11347065-113 |
| 0 | 4.34073559945754 | 0.133 | 0.008 | 0 | facsFB | chr15-19444858-194 |
| 0 | 4.83889966908435 | 0.129 | 0.005 | 0 | facsFB | chr15-33139776-331 |
| 0 | 4.8392530294097 | 0.17 | 0.006 | 0 | facsFB | chr15-77099270-770 |
| 0 | 4.77086585815647 | 0.201 | 0.009 | 0 | facsFB | chr16-1584678-1585 |
| 0 | 4.29174555998492 | 0.124 | 0.007 | 0 | facsFB | chr16-22176242-221 |
| 0 | 4.95346958067581 | 0.116 | 0.004 | 0 | facsFB | chr17-22093918-220 |
| 0 | 4.82935028220752 | 0.108 | 0.004 | 0 | facsFB | chr17-26435507-264 |
| 0 | 4.26842461628832 | 0.133 | 0.007 | 0 | facsFB | chr17-33264539-332 |
| 0 | 4.45186234398395 | 0.123 | 0.006 | 0 | facsFB | chr17-39756329-397 |
| 0 | 4.42240543127645 | 0.124 | 0.007 | 0 | facsFB | chr17-48108877-481 |
| 0 | 4.32222690860891 | 0.137 | 0.007 | 0 | facsFB | chr17-60856591-608 |
| 0 | 4.65633922403788 | 0.167 | 0.008 | 0 | facsFB | chr17-66141389-661 |
| 0 | 4.44440524682327 | 0.121 | 0.006 | 0 | facsFB | chr17-76062768-760 |
| 0 | 4.41012876732847 | 0.127 | 0.006 | 0 | facsFB | chr18-24280199-242 |
| 0 | 4.29116035573836 | 0.161 | 0.009 | 0 | facsFB | chr18-52384461-523 |
| 0 | 4.60780316172174 | 0.103 | 0.004 | 0 | facsFB | chr19-3871288-3871 |
| 0 | 4.29907451557173 | 0.177 | 0.01 | 0 | facsFB | chr19-41414134-414 |
| 0 | 4.27848436813362 | 0.167 | 0.01 | 0 | facsFB | chr19-45733249-457 |

| | | | | | | |
|---|---|---|---|---|---|---|
| 0 | 5.14443266150102 | 0.11 | 0.004 | 0 | facsFB | chr2-1398458-13989 |
| 0 | 4.47605850530838 | 0.121 | 0.007 | 0 | facsFB | chr2-7307568-73080 |
| 0 | 4.77520147137066 | 0.152 | 0.006 | 0 | facsFB | chr2-38500389-3850 |
| 0 | 4.82803775493918 | 0.108 | 0.004 | 0 | facsFB | chr2-41004869-4100 |
| 0 | 4.47633218111219 | 0.147 | 0.007 | 0 | facsFB | chr2-60688716-6068 |
| 0 | 4.35824809627706 | 0.167 | 0.01 | 0 | facsFB | chr2-60777549-6077 |
| 0 | 4.65358708941606 | 0.114 | 0.005 | 0 | facsFB | chr2-63417988-6341 |
| 0 | 4.42182700106603 | 0.116 | 0.006 | 0 | facsFB | chr2-105240828-105 |
| 0 | 4.93217735929442 | 0.139 | 0.006 | 0 | facsFB | chr2-115888735-115 |
| 0 | 4.44054239315428 | 0.15 | 0.008 | 0 | facsFB | chr2-125819900-125 |
| 0 | 4.89818146908519 | 0.121 | 0.004 | 0 | facsFB | chr2-159271853-159 |
| 0 | 4.52342200956282 | 0.127 | 0.007 | 0 | facsFB | chr2-168515260-168 |
| 0 | 4.34048241554126 | 0.113 | 0.006 | 0 | facsFB | chr2-220697693-220 |
| 0 | 5.26027542901981 | 0.113 | 0.003 | 0 | facsFB | chr20-34878387-348 |
| 0 | 4.56956859685268 | 0.148 | 0.007 | 0 | facsFB | chr20-40717176-407 |
| 0 | 4.55944145693095 | 0.102 | 0.005 | 0 | facsFB | chr3-22625120-2262 |
| 0 | 4.28543520277437 | 0.17 | 0.011 | 0 | facsFB | chr3-35477504-3547 |
| 0 | 4.42008485767661 | 0.158 | 0.008 | 0 | facsFB | chr3-35478294-3547 |
| 0 | 4.49276190186398 | 0.104 | 0.005 | 0 | facsFB | chr3-97593169-9759 |
| 0 | 4.79529752191974 | 0.106 | 0.005 | 0 | facsFB | chr3-121738508-121 |
| 0 | 4.45296709089703 | 0.115 | 0.006 | 0 | facsFB | chr3-150816250-150 |
| 0 | 4.28642616419197 | 0.132 | 0.008 | 0 | facsFB | chr4-22426801-2242 |
| 0 | 4.30275847942803 | 0.128 | 0.007 | 0 | facsFB | chr4-23069204-2306 |
| 0 | 4.32346368222846 | 0.157 | 0.009 | 0 | facsFB | chr4-34233228-3423 |
| 0 | 4.37801234813753 | 0.281 | 0.018 | 0 | facsFB | chr4-54606197-5460 |
| 0 | 4.28444511007877 | 0.127 | 0.007 | 0 | facsFB | chr4-131492899-131 |
| 0 | 4.50419766573161 | 0.114 | 0.006 | 0 | facsFB | chr4-142680519-142 |
| 0 | 5.16686307331896 | 0.102 | 0.004 | 0 | facsFB | chr4-150591238-150 |
| 0 | 4.62438129355034 | 0.115 | 0.006 | 0 | facsFB | chr4-150770684-150 |
| 0 | 4.3284045235734 | 0.21 | 0.012 | 0 | facsFB | chr4-155735409-155 |
| 0 | 4.46747567395751 | 0.15 | 0.007 | 0 | facsFB | chr4-180709684-180 |
| 0 | 4.28516715108943 | 0.226 | 0.015 | 0 | facsFB | chr5-9252137-92526 |
| 0 | 4.35452683052121 | 0.153 | 0.008 | 0 | facsFB | chr5-17363071-1736 |
| 0 | 4.33493593363628 | 0.161 | 0.01 | 0 | facsFB | chr5-23967628-2396 |
| 0 | 4.54450592370182 | 0.118 | 0.006 | 0 | facsFB | chr5-29167624-2916 |
| 0 | 4.35577543342954 | 0.145 | 0.008 | 0 | facsFB | chr5-77408656-7740 |
| 0 | 4.55825148087479 | 0.114 | 0.006 | 0 | facsFB | chr5-99264379-9926 |
| 0 | 4.41623773806805 | 0.135 | 0.007 | 0 | facsFB | chr5-120644661-120 |
| 0 | 4.85155412070468 | 0.184 | 0.007 | 0 | facsFB | chr6-14081466-1408 |
| 0 | 5.31912402001788 | 0.194 | 0.005 | 0 | facsFB | chr6-34385379-3438 |
| 0 | 4.26488124727532 | 0.182 | 0.01 | 0 | facsFB | chr6-34386405-3438 |
| 0 | 4.61641243902363 | 0.129 | 0.006 | 0 | facsFB | chr6-37552039-3755 |
| 0 | 4.5083826323361 | 0.162 | 0.008 | 0 | facsFB | chr6-50716291-5071 |

| | | | | | | |
|---|---|---|---|---|---|---|
| 0 | 4.49815240644691 | 0.117 | 0.006 | 0 | facsFB | chr6-52013147-5201 |
| 0 | 4.99642243820177 | 0.142 | 0.005 | 0 | facsFB | chr6-52693991-5269 |
| 0 | 4.30480802776508 | 0.155 | 0.009 | 0 | facsFB | chr6-53552300-5355 |
| 0 | 4.73627755486482 | 0.149 | 0.006 | 0 | facsFB | chr6-71314987-7131 |
| 0 | 4.53106831615046 | 0.137 | 0.007 | 0 | facsFB | chr6-84092007-8409 |
| 0 | 4.30280136531869 | 0.197 | 0.012 | 0 | facsFB | chr6-103436038-103 |
| 0 | 4.3778437180077 | 0.12 | 0.006 | 0 | facsFB | chr7-32809073-3280 |
| 0 | 4.63512545343458 | 0.113 | 0.005 | 0 | facsFB | chr7-40251326-4025 |
| 0 | 4.81619294158173 | 0.291 | 0.012 | 0 | facsFB | chr7-45985187-4598 |
| 0 | 4.74391790097107 | 0.15 | 0.007 | 0 | facsFB | chr7-59180565-5918 |
| 0 | 4.6798079393682 | 0.172 | 0.008 | 0 | facsFB | chr7-71360132-7136 |
| 0 | 4.44893496751148 | 0.212 | 0.011 | 0 | facsFB | chr7-86111914-8611 |
| 0 | 4.7817043407118 | 0.165 | 0.006 | 0 | facsFB | chr7-88376699-8837 |
| 0 | 5.22499665158304 | 0.11 | 0.004 | 0 | facsFB | chr7-95020133-9502 |
| 0 | 5.39784373172385 | 0.1 | 0.003 | 0 | facsFB | chr7-103484246-103 |
| 0 | 4.58438416822383 | 0.137 | 0.006 | 0 | facsFB | chr7-119118037-119 |
| 0 | 5.83654225934328 | 0.127 | 0.003 | 0 | facsFB | chr8-40405666-4040 |
| 0 | 4.27429715340338 | 0.158 | 0.009 | 0 | facsFB | chr8-51911037-5191 |
| 0 | 4.40492826778487 | 0.129 | 0.007 | 0 | facsFB | chr8-52008834-5200 |
| 0 | 5.06851065747435 | 0.164 | 0.007 | 0 | facsFB | chr8-52015278-5201 |
| 0 | 4.66157853394258 | 0.163 | 0.007 | 0 | facsFB | chr8-68676302-6867 |
| 0 | 4.4771033666448 | 0.15 | 0.007 | 0 | facsFB | chr8-69901046-6990 |
| 0 | 4.58770699200992 | 0.144 | 0.006 | 0 | facsFB | chr8-88767912-8876 |
| 0 | 4.61665505397729 | 0.215 | 0.01 | 0 | facsFB | chr8-106361717-106 |
| 0 | 4.30812528569608 | 0.146 | 0.008 | 0 | facsFB | chr8-106440955-106 |
| 0 | 4.50367806500902 | 0.193 | 0.012 | 0 | facsFB | chr8-106486748-106 |
| 0 | 4.48837008743372 | 0.151 | 0.008 | 0 | facsFB | chr9-6585227-65857 |
| 0 | 4.47962814312864 | 0.145 | 0.007 | 0 | facsFB | chr9-50365823-5036 |
| 0 | 4.44710964255497 | 0.123 | 0.006 | 0 | facsFB | chr9-73305062-7330 |
| 0 | 4.46337065301849 | 0.247 | 0.012 | 0 | facsFB | chr9-75398088-7539 |
| 0 | 4.48195173127285 | 0.122 | 0.006 | 0 | facsFB | chr9-81783923-8178 |
| 0 | 4.95222792324534 | 0.144 | 0.005 | 0 | facsFB | chr9-109869360-109 |
| 0 | 4.3440337977656 | 0.11 | 0.005 | 0 | facsFB | chrX-141031448-141 |
| 6.51685796 | 4.56404190597161 | 0.105 | 0.005 | 4.7753319 | facsFB | chr15-91805841-918 |
| 2.88032129 | 4.32448675712353 | 0.121 | 0.007 | 2.1106015 | facsFB | chr8-75660897-7566 |
| 5.36848130 | 4.39321346625791 | 0.112 | 0.006 | 3.9338405 | facsFB | chr2-211875152-211 |
| 1.16309493 | 4.41561565572893 | 0.107 | 0.005 | 8.5227642 | facsFB | chr10-77693542-776 |
| 5.50050595 | 4.33666735021427 | 0.11 | 0.006 | 4.0305837 | facsFB | chr6-33269589-3327 |
| 8.48117436 | 4.32492818518286 | 0.105 | 0.005 | 6.2147162 | facsFB | chr1-42867531-4286 |
| 3.65459047 | 4.34356563639448 | 0.106 | 0.005 | 2.6779596 | facsFB | chr17-19498227-194 |
| 1.63235454 | 4.3119982724136 | 0.125 | 0.008 | 1.1961339 | facsFB | chr13-69747268-697 |
| 2.39024529 | 4.28124658972657 | 0.114 | 0.007 | 1.7514904 | facsFB | chr17-8996544-8997 |
| 3.70979536 | 4.37603115904107 | 0.106 | 0.006 | 2.7184119 | facsFB | chr6-90865004-9086 |

| | | | | | | |
|---|---|---|---|---|---|---|
| 3.94109000 | 4.34927733983451 | 0.11 | 0.006 | 2.8878967 | facsFB | chr18-4852736-4853 |
| 1.40751422 | 4.48169444085504 | 0.115 | 0.007 | 1.0313785 | facsFB | chr9-919749-920250 |
| 6.71359048 | 4.62219213670019 | 0.104 | 0.005 | 4.9194908 | facsFB | chr16-55264977-552 |
| 6.99843943 | 4.30443899781055 | 0.101 | 0.005 | 5.1282184 | facsFB | chr15-3444357-3444 |
| 3.28694947 | 4.35000718352993 | 0.113 | 0.007 | 2.4085648 | facsFB | chr13-32247953-322 |
| 3.12460972 | 4.29598152928447 | 0.103 | 0.005 | 2.2896077 | facsFB | chr17-63246211-632 |
| 1.37825490 | 4.46174945009764 | 0.101 | 0.005 | 1.0099383 | facsFB | chr18-22424995-224 |
| 1.35322994 | 4.32561596109298 | 0.107 | 0.006 | 9.9160089 | facsFB | chr4-139200465-139 |
| 5.64177289 | 4.37004693642395 | 0.103 | 0.006 | 4.1340993 | facsFB | chr1-120634596-120 |
| 6.34196485 | 4.36956927110057 | 0.101 | 0.006 | 4.6471762 | facsFB | chr8-80018151-8001 |
| 8.69769448 | 4.50609868285148 | 0.103 | 0.006 | 6.3733747 | facsFB | chr15-1173075-1173 |
| 6.22360578 | 4.30977676932053 | 0.104 | 0.006 | 4.5604467 | facsFB | chr18-4438315-4438 |
| 5.50966724 | 4.28675899958872 | 0.102 | 0.006 | 4.0372968 | facsFB | chr7-27392466-2739 |
| 9.12215517 | 4.28715273652333 | 0.1 | 0.006 | 6.6844051 | facsFB | chr2-152073374-152 |
| 1.24171303 | 4.26785605140473 | 0.105 | 0.007 | 9.0988509 | facsFB | chr10-51153276-511 |
| 4.17972213 | 2.75193538444286 | 0.213 | 0.033 | 3.0627582 | injFB | chr1-47544980-4754 |
| 4.08713686 | 2.38354667134354 | 0.312 | 0.067 | 2.9949149 | injFB | chr11-67221141-672 |
| 3.22384540 | 2.26577160717833 | 0.291 | 0.062 | 2.3623243 | injFB | chr4-17556868-1755 |
| 2.95277513 | 2.56253791727799 | 0.222 | 0.041 | 2.1636932 | injFB | chr5-160305019-160 |
| 2.08146845 | 2.72592067963228 | 0.186 | 0.031 | 1.5252293 | injFB | chr2-122953365-122 |
| 8.98418431 | 2.2539983869654 | 0.261 | 0.056 | 6.5833048 | injFB | chr12-27635960-276 |
| 3.28372064 | 2.2692045566084 | 0.304 | 0.075 | 2.4061988 | injFB | chr17-12045548-120 |
| 7.35556537 | 2.51274079011333 | 0.198 | 0.035 | 5.3899082 | injFB | chr3-105003005-105 |
| 2.20457346 | 2.4421876358284 | 0.211 | 0.042 | 1.6154364 | injFB | chr12-12915985-129 |
| 3.82657674 | 2.31458092669656 | 0.255 | 0.059 | 2.8039853 | injFB | chr15-97816424-978 |
| 6.51399586 | 2.31254761012845 | 0.24 | 0.054 | 4.7732346 | injFB | chr1-7751410-77519 |
| 9.05629242 | 2.32166521821826 | 0.207 | 0.041 | 6.6361431 | injFB | chr13-95248516-952 |
| 3.89058222 | 2.2931655115292 | 0.226 | 0.049 | 2.8508863 | injFB | chr3-11134066-1113 |
| 5.24820491 | 2.29424073254454 | 0.227 | 0.049 | 3.8457061 | injFB | chr9-108687887-108 |
| 1.26409685 | 2.51602764668505 | 0.171 | 0.03 | 9.2628719 | injFB | chr7-86709293-8670 |
| 4.46928775 | 2.43844875721839 | 0.184 | 0.034 | 3.2749421 | injFB | chr2-204049117-204 |
| 4.56665017 | 2.61376655638384 | 0.178 | 0.033 | 3.3462859 | injFB | chr2-123480945-123 |
| 1.54382710 | 2.51438541226442 | 0.18 | 0.035 | 1.1312640 | injFB | chr1-8257705-82582 |
| 1.01894370 | 2.60902483580408 | 0.165 | 0.03 | 7.4664730 | injFB | chr17-47006755-470 |
| 1.26721560 | 2.28438845964051 | 0.214 | 0.047 | 9.2857250 | injFB | chr2-168106280-168 |
| 1.50675939 | 2.70564432144858 | 0.151 | 0.026 | 1.1041020 | injFB | chr5-120950060-120 |
| 6.69153576 | 2.41726153879095 | 0.166 | 0.031 | 4.9033298 | injFB | chr5-120397462-120 |
| 3.34064122 | 2.44290718959999 | 0.163 | 0.031 | 2.4479083 | injFB | chr13-37101400-371 |
| 4.47846121 | 2.78334459745337 | 0.126 | 0.019 | 3.2816641 | injFB | chr1-47545753-4754 |
| 1.41707580 | 2.94234301187854 | 0.11 | 0.015 | 1.0383849 | injFB | chr6-60666032-6066 |
| 1.43452924 | 2.56569540740873 | 0.151 | 0.028 | 1.0511742 | injFB | chr9-62238004-6223 |
| 2.88002877 | 2.48827292594139 | 0.149 | 0.027 | 2.1103871 | injFB | chr20-12050329-120 |
| 2.41211246 | 2.33373163895259 | 0.162 | 0.032 | 1.7675140 | injFB | chr8-71979729-7198 |

| | | | | | | |
|---|---|---|---|---|---|---|
| 3.65701257 | 2.39752780059397 | 0.156 | 0.03 | 2.6797344 | injFB | chr19-7988357-7988 |
| 4.41977194 | 2.81682305071041 | 0.119 | 0.018 | 3.2386586 | injFB | chr4-175602739-175 |
| 2.36898128 | 2.91693296914385 | 0.105 | 0.014 | 1.7359089 | injFB | chr20-11976389-119 |
| 3.33843007 | 2.56265393209242 | 0.144 | 0.026 | 2.4462880 | injFB | chr8-33647924-3364 |
| 3.39602692 | 2.75162800145324 | 0.133 | 0.023 | 2.4884930 | injFB | chr7-51348493-5134 |
| 5.19321915 | 2.79036979915955 | 0.114 | 0.017 | 3.8054144 | injFB | chr3-143738945-143 |
| 3.89063287 | 2.37328549929117 | 0.171 | 0.035 | 2.8509234 | injFB | chr8-33880936-3388 |
| 7.72120437 | 2.76853265651126 | 0.118 | 0.018 | 5.6578360 | injFB | chr9-85697600-8569 |
| 1.26639364 | 2.49451633406104 | 0.141 | 0.025 | 9.2797020 | injFB | chr2-23960804-2396 |
| 1.64775496 | 2.25273303995728 | 0.198 | 0.046 | 1.2074188 | injFB | chr8-88889218-8888 |
| 3.18849095 | 2.36776833224752 | 0.169 | 0.035 | 2.3364177 | injFB | chr20-40712903-407 |
| 1.24431622 | 2.51422648800858 | 0.14 | 0.025 | 9.1179262 | injFB | chr4-125872192-125 |
| 6.05583539 | 2.34359563688042 | 0.153 | 0.03 | 4.4375102 | injFB | chr9-65284829-6528 |
| 6.38284292 | 2.29362809594248 | 0.173 | 0.037 | 4.6771302 | injFB | chr3-122935397-122 |
| 2.50351890 | 2.46582388107711 | 0.147 | 0.028 | 1.8344935 | injFB | chr3-105003561-105 |
| 4.98667538 | 2.44029424366058 | 0.159 | 0.032 | 3.6540661 | injFB | chr9-61011551-6101 |
| 5.31070328 | 2.27367840187125 | 0.169 | 0.036 | 3.8915028 | injFB | chr5-120654294-120 |
| 6.97767059 | 2.34130700808116 | 0.16 | 0.033 | 5.1129997 | injFB | chr2-21477148-2147 |
| 1.15169458 | 2.26706286766828 | 0.17 | 0.036 | 8.4392263 | injFB | chr1-137730456-137 |
| 7.60170945 | 2.27235333627047 | 0.164 | 0.034 | 5.5702742 | injFB | chr14-96441520-964 |
| 5.23692864 | 2.35334414232374 | 0.172 | 0.038 | 3.8374432 | injFB | chr16-68512741-685 |
| 3.22414988 | 2.2718699068614 | 0.163 | 0.034 | 2.3625474 | injFB | chr18-51801342-518 |
| 9.23642918 | 2.50911895566468 | 0.111 | 0.018 | 6.7681412 | injFB | chr4-150970580-150 |
| 1.28494508 | 2.27759562330366 | 0.159 | 0.034 | 9.4156406 | injFB | chr4-124109363-124 |
| 2.05004338 | 2.61327022767127 | 0.11 | 0.018 | 1.5022020 | injFB | chr1-58712415-5871 |
| 2.88333125 | 2.37104119805063 | 0.154 | 0.032 | 2.1128071 | injFB | chr6-104533683-104 |
| 7.88593905 | 2.26578293017361 | 0.175 | 0.041 | 5.7785480 | injFB | chr11-76072564-760 |
| 1.70973582 | 2.35852618661136 | 0.126 | 0.023 | 1.2528362 | injFB | chr2-187350218-187 |
| 7.15273082 | 2.63428606598927 | 0.121 | 0.021 | 5.2412779 | injFB | chr8-60935721-6093 |
| 5.24359085 | 2.53041474896851 | 0.108 | 0.017 | 3.8423250 | injFB | chr3-158245825-158 |
| 7.52359839 | 2.4308822767372 | 0.147 | 0.03 | 5.5130371 | injFB | chr10-48036841-480 |
| 1.58043784 | 2.54416625443953 | 0.126 | 0.023 | 1.1580911 | injFB | chr18-34998198-349 |
| 1.76039138 | 2.45117030141279 | 0.138 | 0.027 | 1.2899549 | injFB | chr14-63656833-636 |
| 5.36336269 | 2.64923710925963 | 0.106 | 0.017 | 3.9300898 | injFB | chr5-70469385-7046 |
| 6.41786119 | 2.30546423880073 | 0.161 | 0.036 | 4.7027904 | injFB | chr8-80839053-8083 |
| 7.99770922 | 2.3708011162019 | 0.142 | 0.029 | 5.8604493 | injFB | chr13-81923396-819 |
| 8.26066926 | 2.59835449893041 | 0.109 | 0.018 | 6.0531375 | injFB | chrX-112030393-112 |
| 1.36023289 | 2.25181204820208 | 0.163 | 0.037 | 9.9673241 | injFB | chr17-65413165-654 |
| 3.55634771 | 2.83364680580066 | 0.108 | 0.018 | 2.6059706 | injFB | chr12-27602038-276 |
| 4.71761371 | 2.51965785150794 | 0.126 | 0.023 | 3.4569069 | injFB | chr12-38226518-382 |
| 6.57257187 | 2.35338910350767 | 0.141 | 0.029 | 4.8161571 | injFB | chr13-97207053-972 |
| 5.49485230 | 2.6637634401322 | 0.107 | 0.018 | 4.0264409 | injFB | chr2-164814090-164 |
| 2.43187802 | 2.51096405257943 | 0.14 | 0.029 | 1.7819975 | injFB | chr1-257748324-257 |

| | | | | | | |
|---|---|---|---|---|---|---|
| 2.49983113 | 2.47208698349127 | 0.131 | 0.026 | 1.8317912 | injFB | chr9-60002348-6000 |
| 2.87235186 | 2.57758308645376 | 0.121 | 0.023 | 2.1047617 | injFB | chr1-58706754-5870 |
| 3.46722760 | 2.60992843342465 | 0.111 | 0.019 | 2.5406664 | injFB | chr3-97129004-9712 |
| 1.04011102 | 2.62350519332305 | 0.108 | 0.018 | 7.6215799 | injFB | chr7-54471285-5447 |
| 2.45455987 | 2.43844552050969 | 0.124 | 0.024 | 1.7986180 | injFB | chr5-115067323-115 |
| 3.04682336 | 2.61936574973786 | 0.107 | 0.018 | 2.2326085 | injFB | chr2-51026020-5102 |
| 3.77994860 | 2.33731757842324 | 0.138 | 0.029 | 2.7698178 | injFB | chr13-83859822-838 |
| 5.06811692 | 2.66093046560429 | 0.106 | 0.018 | 3.7137437 | injFB | chr6-53549700-5355 |
| 5.12261974 | 2.4494818834937 | 0.122 | 0.023 | 3.7536815 | injFB | chr4-22627876-2262 |
| 1.16131027 | 2.58774029994385 | 0.108 | 0.019 | 8.5096868 | injFB | chr7-45985187-4598 |
| 4.09671130 | 2.34167366905497 | 0.129 | 0.026 | 3.0019307 | injFB | chr11-52981600-529 |
| 4.63463791 | 2.38614978470812 | 0.109 | 0.019 | 3.3961050 | injFB | chr12-301524-30202 |
| 1.23779671 | 2.44731644416447 | 0.133 | 0.027 | 9.0701535 | injFB | chr2-48955523-4895 |
| 1.62347696 | 2.28668669396347 | 0.137 | 0.029 | 1.1896287 | injFB | chr5-126295666-126 |
| 3.83858045 | 2.47592767141195 | 0.12 | 0.023 | 2.8127812 | injFB | chr1-40615873-4061 |
| 7.55171029 | 2.55986973489779 | 0.108 | 0.019 | 5.5336365 | injFB | chr2-32552336-3255 |
| 9.04847644 | 2.33471257406233 | 0.13 | 0.027 | 6.6304158 | injFB | chr3-53169974-5317 |
| 1.16649131 | 2.64117286717711 | 0.105 | 0.018 | 8.5476517 | injFB | chr4-22890319-2289 |
| 6.28257203 | 2.31358176930163 | 0.129 | 0.026 | 4.6036551 | injFB | chr14-104145873-10 |
| 6.32822654 | 2.37714140574383 | 0.121 | 0.024 | 4.6371092 | injFB | chr16-83176716-831 |
| 1.29565142 | 2.58257105892519 | 0.105 | 0.018 | 9.4940931 | injFB | chr1-222822071-222 |
| 2.85140746 | 2.26454669839862 | 0.146 | 0.034 | 2.0894144 | injFB | chr17-67745925-677 |
| 3.30709351 | 2.38650199158886 | 0.127 | 0.026 | 2.4233256 | injFB | chr14-33552561-335 |
| 8.51782559 | 2.29864240559265 | 0.121 | 0.024 | 6.2415729 | injFB | chr5-100924910-100 |
| 1.70695289 | 2.36293213738146 | 0.12 | 0.024 | 1.2507970 | injFB | chr10-19381015-193 |
| 3.77175428 | 2.34018325727621 | 0.135 | 0.03 | 2.7638133 | injFB | chr2-121096441-121 |
| 4.94070972 | 2.48847800901849 | 0.111 | 0.021 | 3.6203841 | injFB | chr9-102720308-102 |
| 5.17282011 | 2.37733808524216 | 0.12 | 0.024 | 3.7904667 | injFB | chr6-33103178-3310 |
| 5.54830606 | 2.26265716762985 | 0.134 | 0.029 | 4.0656100 | injFB | chr13-97704262-977 |
| 4.80551495 | 2.27313043845401 | 0.125 | 0.026 | 3.5213179 | injFB | chr13-74502696-745 |
| 6.81373509 | 2.31323443754575 | 0.117 | 0.023 | 4.9928734 | injFB | chr17-77953642-779 |
| 1.45065730 | 2.48800516541612 | 0.109 | 0.021 | 1.0629923 | injFB | chr3-113475174-113 |
| 1.82746383 | 2.51062629775864 | 0.118 | 0.024 | 1.3391033 | injFB | chr11-3677667-3678 |
| 2.28085751 | 2.33422170759362 | 0.118 | 0.024 | 1.6713348 | injFB | chr6-92793519-9279 |
| 6.35371793 | 2.38338591837429 | 0.107 | 0.02 | 4.6557884 | injFB | chr7-124847827-124 |
| 1.81013036 | 2.41293914320806 | 0.11 | 0.022 | 1.3264019 | injFB | chr3-158933184-158 |
| 4.44925979 | 2.44912696501037 | 0.112 | 0.023 | 3.2602663 | injFB | chr14-75385261-753 |
| 5.07658938 | 2.40168007069127 | 0.105 | 0.02 | 3.7199520 | injFB | chr3-138598529-138 |
| 6.61705067 | 2.38004646926555 | 0.121 | 0.026 | 4.8487497 | injFB | chr4-175557933-175 |
| 9.24133703 | 2.57318036889494 | 0.105 | 0.02 | 6.7717375 | injFB | chr6-6319261-63197 |
| 1.80770807 | 2.38323342150087 | 0.118 | 0.025 | 1.3246270 | injFB | chr5-156870682-156 |
| 2.37376676 | 2.55409773444694 | 0.105 | 0.02 | 1.7394155 | injFB | chr5-120223591-120 |
| 3.19785571 | 2.33340782899395 | 0.113 | 0.023 | 2.3432799 | injFB | chr1-58705733-5870 |

| | | | | | | | |
|---|---|---|---|---|---|---|---|
| 3.82281341 | 2.39778848911448 | | 0.107 | 0.021 | 2.8012276 | injFB | chr11-26790189-267 |
| 9.07881336 | 2.32574228822244 | | 0.111 | 0.022 | 6.6526457 | injFB | chr17-45408176-454 |
| 3.91045341 | 2.29065863351832 | | 0.125 | 0.028 | 2.8654473 | injFB | chr16-42353054-423 |
| 4.20343527 | 2.35646464824561 | | 0.101 | 0.019 | 3.0801344 | injFB | chr7-134994249-134 |
| 1.84741898 | 2.33538379493324 | | 0.121 | 0.027 | 1.3537258 | injFB | chr14-16757520-167 |
| 1.98583872 | 2.3224770622579 | | 0.106 | 0.021 | 1.4551550 | injFB | chr7-56060433-5606 |
| 2.90470478 | 2.25969371342213 | | 0.117 | 0.025 | 2.1284689 | injFB | chr7-127484110-127 |
| 3.36548519 | 2.2665982742912 | | 0.121 | 0.027 | 2.4661131 | injFB | chr3-10529305-1052 |
| 3.83122494 | 2.29634490084914 | | 0.108 | 0.022 | 2.8073913 | injFB | chr5-78952032-7895 |
| 1.43686959 | 2.35436994452232 | | 0.112 | 0.023 | 1.0528891 | injFB | chr17-45340198-453 |
| 1.44233004 | 2.3370767610156 | | 0.108 | 0.022 | 1.0568904 | injFB | chr17-22751390-227 |
| 3.67990026 | | 2.359231984 | 0.112 | 0.023 | 2.6965057 | injFB | chr16-50862122-508 |
| 4.99429548 | 2.3751575159306 | | 0.11 | 0.023 | 3.6596499 | injFB | chr14-102664639-10 |
| 1.13328661 | 2.31934558702292 | | 0.125 | 0.029 | 8.3043390 | injFB | chr13-79878813-798 |
| 1.37184841 | 2.25808287106221 | | 0.114 | 0.024 | 1.0052438 | injFB | chr14-2590030-2590 |
| 3.22496775 | 2.34789561305285 | | 0.1 | 0.019 | 2.3631467 | injFB | chr7-86714224-8671 |
| 3.85211882 | 2.31930831374407 | | 0.112 | 0.024 | 2.8227017 | injFB | chr19-14223378-142 |
| 4.74366333 | 2.37191387213082 | | 0.101 | 0.02 | 3.4759952 | injFB | chr16-3169285-3169 |
| 6.29950151 | 2.35118946301436 | | 0.12 | 0.027 | 4.6160605 | injFB | chr5-7129653-71301 |
| 1.47348641 | 2.25793389302192 | | 0.107 | 0.022 | 1.0797207 | injFB | chr10-19181962-191 |
| 3.73327894 | 2.27381362241409 | | 0.121 | 0.028 | 2.7356198 | injFB | chr14-5382142-5382 |
| 6.54631951 | 2.25563202017611 | | 0.111 | 0.024 | 4.7969203 | injFB | chr14-84300247-843 |
| 1.97387636 | 2.27741252949562 | | 0.109 | 0.023 | 1.4463894 | injFB | chr2-83180333-8318 |
| 7.43247937 | 2.41827568421775 | | 0.106 | 0.022 | 5.4462681 | injFB | chr15-49387156-493 |
| 2.98779095 | 2.31304843685499 | | 0.11 | 0.024 | 2.1893516 | injFB | chr2-55300628-5530 |
| 9.12750109 | 2.26238557034919 | | 0.136 | 0.035 | 6.6883224 | injFB | chr15-19542638-195 |
| 2.43295272 | 2.27957165608269 | | 0.107 | 0.023 | 1.7827850 | injFB | chr9-24577645-2457 |
| 4.77095234 | 2.29287539957385 | | 0.1 | 0.02 | 3.4959916 | injFB | chr3-35712306-3571 |
| 4.51821312 | 2.29436292839842 | | 0.1 | 0.021 | 3.3107929 | injFB | chr4-82548760-8254 |
| 3.67001026 | 2.35185839614379 | | 0.1 | 0.021 | 2.6892587 | injFB | chr3-96941356-9694 |
| 2.69115539 | 2.26974455482688 | | 0.109 | 0.025 | 1.9719871 | injFB | chr6-4347716-43482 |
| 9.81643444 | 2.29339920995133 | | 0.1 | 0.022 | 7.1931494 | injFB | chr1-55131624-5513 |
| 1.41157044 | 2.28572516060229 | | 0.109 | 0.025 | 1.0343508 | injFB | chr9-57061353-5706 |
| 1.96048094 | 2.26305240543218 | | 0.105 | 0.024 | 1.4365737 | injFB | chr20-26581045-265 |
| 3.51126967 | 2.2861004238626 | | 0.101 | 0.022 | 2.5729390 | injFB | chr19-50126440-501 |
| 1.03414704 | 2.38000636941844 | | 0.108 | 0.025 | 7.5778779 | injFB | chr6-61624717-6162 |
| 0 | 4.09679193857777 | | 0.162 | 0.01 | 0 | muscFB | chr1-55278704-5527 |
| 5.00016619 | 3.0225243515796 | | 0.259 | 0.035 | 3.6639517 | muscFB | chr10-19303757-193 |
| 4.28421430 | 3.10941180161315 | | 0.229 | 0.028 | 3.1393265 | muscFB | chr3-113575657-113 |
| 1.07314858 | 3.13223313247701 | | 0.218 | 0.026 | 7.8636679 | muscFB | chr16-65771797-657 |
| 4.90530169 | 3.14055955226219 | | 0.219 | 0.027 | 3.5944382 | muscFB | chr12-27562194-275 |
| 3.75620861 | 3.06147098911747 | | 0.223 | 0.028 | 2.7524219 | muscFB | chr9-83479001-8347 |
| 5.38312433 | 3.06929334012228 | | 0.236 | 0.032 | 3.9445704 | muscFB | chr13-103108657-10 |

| | | | | | | | |
|---|---|---|---|---|---|---|---|
| 1.21389871 | 3.42016763231025 | | 0.168 | 0.016 | 8.8950370 | muscFB | chr9-85697600-8569 |
| 2.99447744 | 3.0317884632506 | | 0.211 | 0.027 | 2.1942512 | muscFB | chr14-67618639-676 |
| 6.81329187 | 2.97645769586421 | | 0.224 | 0.03 | 4.9925486 | muscFB | chr5-98529550-9853 |
| 1.92612098 | 3.32224877852031 | | 0.167 | 0.017 | 1.4113959 | muscFB | chr1-99621724-9962 |
| 2.94868624 | 3.68653869779175 | | 0.133 | 0.011 | 2.1606970 | muscFB | chr14-32188800-321 |
| 1.18723442 | 3.04789592201095 | | 0.202 | 0.026 | 8.6996501 | muscFB | chr20-36629112-366 |
| 1.20114379 | 3.23323012294013 | | 0.155 | 0.015 | 8.8015733 | muscFB | chr17-45347942-453 |
| 1.02757544 | 3.58349712446838 | | 0.138 | 0.012 | 7.5297234 | muscFB | chr11-44092066-440 |
| 1.71491720 | 3.02616196716751 | | 0.173 | 0.02 | 1.2566330 | muscFB | chr8-49226623-4922 |
| 3.61127716 | 2.9879867341121 | | 0.203 | 0.028 | 2.6462211 | muscFB | chr18-25246307-252 |
| 9.26382768 | 3.20310467853392 | | 0.171 | 0.02 | 6.7882179 | muscFB | chr14-33442900-334 |
| 7.27491576 | 3.08924826368264 | | 0.177 | 0.021 | 5.3308109 | muscFB | chr17-30128676-301 |
| 1.45804311 | 3.60176332923864 | | 0.134 | 0.012 | 1.0684044 | muscFB | chr10-17597497-175 |
| 5.24502956 | 3.22533248300886 | | 0.17 | 0.02 | 3.8433793 | muscFB | chr8-106333153-106 |
| 1.20880869 | 3.01422524957908 | | 0.202 | 0.028 | 8.8577391 | muscFB | chr11-56034941-560 |
| 4.58118235 | 3.38485804915066 | | 0.159 | 0.017 | 3.3569346 | muscFB | chr11-76281337-762 |
| 4.37076387 | 2.98466288917537 | | 0.192 | 0.025 | 3.2027471 | muscFB | chr10-29740371-297 |
| 6.26858129 | 3.4796801085103 | | 0.126 | 0.011 | 4.5934032 | muscFB | chr1-226398040-226 |
| 1.61907719 | 3.09153430777732 | | 0.174 | 0.021 | 1.1864047 | muscFB | chr20-17789970-177 |
| 5.08795260 | 3.1671447283682 | | 0.156 | 0.017 | 3.7282786 | muscFB | chr9-91096668-9109 |
| 3.64456703 | | 3.072269299 | 0.175 | 0.021 | 2.6706148 | muscFB | chr2-138706052-138 |
| 1.79305708 | 3.70522550539445 | | 0.117 | 0.01 | 1.3138912 | muscFB | chr1-17912600-1791 |
| 2.67181792 | 3.03190858651029 | | 0.181 | 0.024 | 1.9578173 | muscFB | chr6-33105029-3310 |
| 5.43929256 | 3.24245608106993 | | 0.161 | 0.019 | 3.9857286 | muscFB | chr4-150094882-150 |
| 5.37576210 | 3.00525675533524 | | 0.169 | 0.021 | 3.9391756 | muscFB | chr1-200759127-200 |
| 4.82849036 | 3.28250870582378 | | 0.142 | 0.015 | 3.5381535 | muscFB | chr1-47381762-4738 |
| 2.38797800 | 3.54394641988898 | | 0.122 | 0.011 | 1.7498290 | muscFB | chr9-81844829-8184 |
| 6.55312828 | 3.03946090748236 | | 0.183 | 0.025 | 4.8019096 | muscFB | chr4-178909878-178 |
| 8.22086296 | 3.81243625837531 | | 0.105 | 0.008 | 6.0239688 | muscFB | chr7-117985270-117 |
| 3.94372986 | 3.35348322317844 | | 0.133 | 0.013 | 2.8898311 | muscFB | chr13-77135462-771 |
| 1.34953078 | 2.98750376668298 | | 0.159 | 0.019 | 9.8889027 | muscFB | chr7-99686044-9968 |
| 2.91003617 | 3.26017446082128 | | 0.135 | 0.014 | 2.1323755 | muscFB | chr7-100686612-100 |
| 2.32644748 | 3.48864469499277 | | 0.126 | 0.012 | 1.7047416 | muscFB | chr7-19623885-1962 |
| 1.23425049 | 3.13204735269595 | | 0.147 | 0.017 | 9.0441679 | muscFB | chr9-35911678-3591 |
| 3.22137583 | 3.43125295003176 | | 0.123 | 0.012 | 2.3605146 | muscFB | chr19-55715844-557 |
| 3.60426728 | 3.1165520085128 | | 0.148 | 0.017 | 2.6410845 | muscFB | chr3-132173573-132 |
| 2.02627522 | 2.98331984442814 | | 0.166 | 0.022 | 1.4847855 | muscFB | chr5-122818102-122 |
| 2.10948385 | 3.02788411373099 | | 0.142 | 0.016 | 1.5457580 | muscFB | chr8-21895124-2189 |
| 4.19603980 | 3.16073062335641 | | 0.152 | 0.019 | 3.0747153 | muscFB | chr15-3431240-3431 |
| 1.22990676 | 3.33783910477708 | | 0.127 | 0.013 | 9.0123386 | muscFB | chr7-22308980-2230 |
| 4.29718932 | 3.21373049070432 | | 0.151 | 0.018 | 3.1488342 | muscFB | chr10-66998421-669 |
| 9.22059713 | 3.32135297412247 | | 0.126 | 0.013 | 6.7565400 | muscFB | chr1-188814915-188 |
| 4.63789779 | 3.17479462634783 | | 0.14 | 0.016 | 3.3984938 | muscFB | chr6-67270930-6727 |

| | | | | | | |
|---|---|---|---|---|---|---|
| 1.82003263 | 3.01867595709059 | 0.141 | 0.016 | 1.3336580 | muscFB | chr1-58712415-5871 |
| 6.26523800 | 3.29610765937414 | 0.129 | 0.014 | 4.5909533 | muscFB | chr6-20226753-2022 |
| 6.50423898 | 3.0492369268265 | 0.148 | 0.019 | 4.7660851 | muscFB | chr5-120339489-120 |
| 8.58285446 | 3.42738032773928 | 0.115 | 0.011 | 6.2892239 | muscFB | chr8-29056380-2905 |
| 4.34309976 | 2.97676583494356 | 0.155 | 0.021 | 3.1824758 | muscFB | chr18-50258816-502 |
| 5.16261167 | 3.07683960837258 | 0.148 | 0.019 | 3.7829863 | muscFB | chr7-68184188-6818 |
| 1.17761835 | 3.19330288020635 | 0.136 | 0.016 | 8.6291869 | muscFB | chr6-92520980-9252 |
| 2.32985441 | 3.19836575946962 | 0.129 | 0.015 | 1.7072381 | muscFB | chr1-214924096-214 |
| 1.22618525 | 2.98637944715822 | 0.148 | 0.019 | 8.9850686 | muscFB | chr6-40693697-4069 |
| 2.44306704 | 3.01788572405858 | 0.149 | 0.02 | 1.7901964 | muscFB | chr14-67492699-674 |
| 3.77872820 | 3.39626353208442 | 0.113 | 0.011 | 2.7689235 | muscFB | chr3-36161143-3616 |
| 2.04600365 | 3.38495507185165 | 0.115 | 0.012 | 1.4992419 | muscFB | chr20-17401627-174 |
| 3.30330895 | 3.2831275874334 | 0.121 | 0.013 | 2.4205524 | muscFB | chr9-77603930-7760 |
| 6.92423294 | 2.98018108075535 | 0.148 | 0.02 | 5.0738424 | muscFB | chr8-49224463-4922 |
| 8.72996007 | 3.13083264308229 | 0.131 | 0.015 | 6.3970179 | muscFB | chr1-13137006-1313 |
| 6.75501333 | 3.2296040035085 | 0.132 | 0.016 | 4.9498440 | muscFB | chr13-70953516-709 |
| 1.12773281 | 3.43489138647252 | 0.104 | 0.01 | 8.2636426 | muscFB | chr18-24443029-244 |
| 1.30091944 | 3.01583406386967 | 0.131 | 0.016 | 9.5326953 | muscFB | chr10-19427201-194 |
| 1.45015624 | 3.20322347565983 | 0.12 | 0.013 | 1.0626251 | muscFB | chr18-59654763-596 |
| 1.82970682 | 3.32322008126939 | 0.112 | 0.011 | 1.3407469 | muscFB | chr1-167281096-167 |
| 5.98821574 | 3.08086838610079 | 0.135 | 0.017 | 4.3879608 | muscFB | chr4-172638797-172 |
| 3.71582531 | 3.31195059638996 | 0.107 | 0.011 | 2.7228304 | muscFB | chr15-49908571-499 |
| 1.54652008 | 3.13629938373074 | 0.128 | 0.016 | 1.1332373 | muscFB | chr17-83679919-836 |
| 8.82122313 | 3.28517805748371 | 0.11 | 0.012 | 6.4638923 | muscFB | chr2-44125208-4412 |
| 2.64226473 | 3.09858208844518 | 0.121 | 0.014 | 1.9361617 | muscFB | chr2-227057158-227 |
| 1.25609457 | 3.03944092301423 | 0.128 | 0.016 | 9.2042339 | muscFB | chr14-73033589-730 |
| 2.43035693 | 3.04631127488027 | 0.127 | 0.016 | 1.7808829 | muscFB | chr4-173266831-173 |
| 1.32067419 | 3.20011322119984 | 0.123 | 0.015 | 9.6774514 | muscFB | chr4-132080604-132 |
| 2.69484127 | 3.06238765800858 | 0.132 | 0.017 | 1.9746880 | muscFB | chr1-55234572-5523 |
| 5.98191978 | 2.99878499363317 | 0.125 | 0.015 | 4.3833474 | muscFB | chr1-120637326-120 |
| 8.34420368 | 3.22887984682096 | 0.111 | 0.012 | 6.1143487 | muscFB | chr1-137730962-137 |
| 2.21368391 | 3.36524425893569 | 0.103 | 0.01 | 1.6221123 | muscFB | chr17-40408524-404 |
| 1.29788198 | 2.98000371642917 | 0.14 | 0.019 | 9.5104378 | muscFB | chr4-168242461-168 |
| 2.67242046 | 3.09849031081475 | 0.114 | 0.013 | 1.9582588 | muscFB | chr10-29428563-294 |
| 3.41305396 | 3.27094860646003 | 0.123 | 0.015 | 2.5009698 | muscFB | chr12-10986834-109 |
| 4.48628912 | 3.06446617559188 | 0.119 | 0.014 | 3.2874001 | muscFB | chr13-97255604-972 |
| 1.80383335 | 3.17043748381066 | 0.117 | 0.014 | 1.3217877 | muscFB | chr3-104967113-104 |
| 1.43060444 | 3.2183279539166 | 0.101 | 0.01 | 1.0482982 | muscFB | chr7-22033635-2203 |
| 1.48662727 | 3.29195827794155 | 0.104 | 0.011 | 1.0893499 | muscFB | chr5-98714927-9871 |
| 3.40579364 | 3.32694583569447 | 0.103 | 0.011 | 2.4956497 | muscFB | chr14-71898596-718 |
| 6.03136915 | 3.01903386988019 | 0.134 | 0.018 | 4.4195822 | muscFB | chr14-79129634-791 |
| 8.16795599 | 2.97726596371661 | 0.128 | 0.017 | 5.9852004 | muscFB | chr15-39911335-399 |
| 1.65299372 | 3.29837412275503 | 0.114 | 0.013 | 1.2112576 | muscFB | chr4-150745742-150 |

| | | | | | | | |
|---|---|---|---|---|---|---|---|
| 8.07027745( | 3.09539012142409 | | 0.11 | 0.013 | 5.9136249 | muscFB | chr2-150605590-150 |
| 9.41838727 | 2.98020686210738 | | 0.15 | 0.023 | 6.9014739 | muscFB | chr16-60776274-607 |
| 7.95253671 | 3.14194493093519 | | 0.104 | 0.011 | 5.8273485 | muscFB | chr1-220521288-220 |
| 1.42617014 | 3.18454371699063 | | 0.111 | 0.013 | 1.0450489 | muscFB | chr2-5482326-54828 |
| 1.96110696 | 3.09696182732562 | | 0.107 | 0.012 | 1.4370325 | muscFB | chr19-13141048-131 |
| 1.99041387 | 3.17685670541416 | | 0.105 | 0.012 | 1.4585076 | muscFB | chr9-41045724-4104 |
| 3.15616357 | 3.18949621429681 | | 0.103 | 0.011 | 2.3127293 | muscFB | chr5-115692462-115 |
| 5.86429348 | 3.16406749091088 | | 0.107 | 0.012 | 4.2971548 | muscFB | chr7-72019806-7202 |
| 7.48733156( | 3.11624968484294 | | 0.109 | 0.013 | 5.4864619 | muscFB | chr16-65772452-657 |
| 1.27654393 | 3.00558335613415 | | 0.115 | 0.014 | 9.3540799 | muscFB | chr2-150320224-150 |
| 4.18171220 | 2.96759547425138 | | 0.124 | 0.016 | 3.0642165 | muscFB | chr14-18919988-189 |
| 3.58003765 | 3.10153132025275 | | 0.111 | 0.014 | 2.6233298 | muscFB | chr1-154894025-154 |
| 3.98340273 | 3.04524214100935 | | 0.115 | 0.015 | 2.9189020 | muscFB | chr2-45548094-4554 |
| 4.69213383 | 3.17982629749358 | | 0.11 | 0.013 | 3.4382361 | muscFB | chr6-121384027-121 |
| 1.51081876 | 3.00564978261582 | | 0.113 | 0.014 | 1.1070766 | muscFB | chr9-36172595-3617 |
| 4.14735537 | 3.06916905746878 | | 0.109 | 0.013 | 3.0390410 | muscFB | chr17-5849459-5849 |
| 9.60150279 | 3.06036842757188 | | 0.115 | 0.015 | 7.0356547 | muscFB | chr13-97186762-971 |
| 3.82890232( | 3.07454173340759 | | 0.104 | 0.012 | 2.8056894 | muscFB | chr10-107022129-10 |
| 9.68916490 | 3.05168057438509 | | 0.112 | 0.014 | 7.0998906 | muscFB | chr9-74707605-7470 |
| 3.15131520 | 3.10018594956973 | | 0.104 | 0.013 | 2.3091766 | muscFB | chr5-25321300-2532 |
| 4.02713181 | 3.08735182054227 | | 0.115 | 0.015 | 2.9509452 | muscFB | chr8-21804404-2180 |
| 5.53606823 | 3.09381294822015 | | 0.106 | 0.013 | 4.0566425 | muscFB | chr6-20175987-2017 |
| 8.95636789 | 3.17801079795319 | | 0.105 | 0.013 | 6.5629218 | muscFB | chr7-99147972-9914 |
| 1.40904187 | 3.05155477570745 | | 0.104 | 0.012 | 1.0324979 | muscFB | chr10-34058895-340 |
| 1.41557813( | 3.1168922571958 | | 0.101 | 0.012 | 1.0372875 | muscFB | chr17-76503480-765 |
| 4.78216953 | 2.97141257319949 | | 0.114 | 0.015 | 3.5042112 | muscFB | chr13-66203652-662 |
| 3.12852685 | 2.99074127147423 | | 0.112 | 0.015 | 2.2924781 | muscFB | chr11-24962769-249 |
| 4.00532917 | 3.13304781391388 | | 0.108 | 0.014 | 2.9349690 | muscFB | chr16-923443-92394 |
| 8.86541159 | 3.02858508212352 | | 0.115 | 0.015 | 6.4962721 | muscFB | chr16-28512165-285 |
| 1.07826637 | 3.01720180210516 | | 0.114 | 0.015 | 7.9011694 | muscFB | chr11-16625665-166 |
| 5.53570592 | 2.97501023500294 | | 0.13 | 0.02 | 4.0563770 | muscFB | chr1-227668482-227 |
| 1.07640480 | 3.09203790351449 | | 0.106 | 0.014 | 7.8875284 | muscFB | chr7-112601177-112 |
| 3.17985649 | 3.00131279065984 | | 0.108 | 0.014 | 2.3300907 | muscFB | chr2-138852968-138 |
| 2.54788002( | | 3.138573092 | 0.106 | 0.014 | 1.8669998 | muscFB | chr12-8799685-8800 |
| 1.11460415 | 2.99826890386667 | | 0.119 | 0.017 | 8.1674403 | muscFB | chr17-82775328-827 |
| 1.12550105 | 3.0640449087852 | | 0.114 | 0.016 | 8.2472890 | muscFB | chr3-112585313-112 |
| 3.63532717 | 3.05539941533507 | | 0.102 | 0.013 | 2.6638441 | muscFB | chr18-706714-70721 |
| 8.72896173 | 2.96701634348538 | | 0.123 | 0.018 | 6.3962863 | muscFB | chr4-93425844-9342 |
| 2.56770592 | 2.9748566261524 | | 0.104 | 0.013 | 1.8815275 | muscFB | chr1-13870516-1387 |
| 1.57534162( | 3.04452397452823 | | 0.108 | 0.015 | 1.1543567 | muscFB | chr11-44316439-443 |
| 3.48509218 | 3.0997001395783 | | 0.104 | 0.014 | 2.5537570 | muscFB | chr1-185530127-185 |
| 7.88520636 | 2.97189252571328 | | 0.101 | 0.013 | 5.7780111 | muscFB | chr19-55362328-553 |
| 8.04417231( | 2.99143552479102 | | 0.103 | 0.013 | 5.8944959 | muscFB | chr15-37661779-376 |

| | | | | | | |
|---|---|---|---|---|---|---|
| 2.13087793 | 2.97604505930118 | 0.101 | 0.013 | 1.5614349 | muscFB | chr1-126385016-126 |
| 2.37004113 | 3.13667896900086 | 0.102 | 0.013 | 1.7366855 | muscFB | chr9-81763745-8176 |
| 1.70597839 | 3.05207830983527 | 0.113 | 0.016 | 1.2500829 | muscFB | chr11-76492033-764 |
| 1.19458466 | 2.99320221601681 | 0.104 | 0.014 | 8.7535102 | muscFB | chr9-54999711-5500 |
| 5.52297557 | 2.99860092375498 | 0.101 | 0.013 | 4.0470487 | muscFB | chr14-75111699-751 |
| 1.96170897 | 3.1604455252727 | 0.105 | 0.015 | 1.4374736 | muscFB | chr6-32451675-3245 |
| 2.02985678 | 2.97706523337807 | 0.107 | 0.015 | 1.4874100 | muscFB | chr1-41112155-4111 |
| 2.72596707 | 2.98848777486829 | 0.111 | 0.016 | 1.9974959 | muscFB | chr6-15361954-1536 |
| 1.43426261 | 2.98016084935144 | 0.107 | 0.015 | 1.0509788 | muscFB | chr9-90965807-9096 |
| 2.57772222 | 2.98715766213141 | 0.105 | 0.015 | 1.8888672 | muscFB | chr1-171139356-171 |
| 6.44866851 | 2.98000097591811 | 0.1 | 0.014 | 4.7253650 | muscFB | chr8-42296729-4229 |
| 6.59633989 | 2.99149552728495 | 0.1 | 0.014 | 4.8335735 | muscFB | chr1-129097850-129 |
| 8.90301534 | 2.95975215880473 | 0.102 | 0.014 | 6.5238269 | muscFB | chr5-141054430-141 |
| 6.69448281 | 2.96101015512996 | 0.102 | 0.015 | 4.9054893 | muscFB | chr1-179662387-179 |
| 0 | 2.32973693086231 | 0.201 | 0.043 | 0 | myoFB | chr1-225285654-225 |
| 0 | 2.34375954778916 | 0.183 | 0.038 | 0 | myoFB | chr15-36406870-364 |
| 0 | 2.25265283612737 | 0.197 | 0.045 | 0 | myoFB | chr19-36480026-364 |
| 0 | 2.2561776994076 | 0.229 | 0.053 | 0 | myoFB | chr2-171306689-171 |
| 2.77576615 | 2.37261021705203 | 0.163 | 0.032 | 2.0339870 | myoFB | chr18-43698964-436 |
| 4.88351484 | 2.35131053350788 | 0.166 | 0.034 | 3.5784736 | myoFB | chr5-48732248-4873 |
| 6.35896086 | 2.23914934599706 | 0.18 | 0.04 | 4.6596303 | myoFB | chr6-15855588-1585 |
| 2.06981989 | 2.28939298749103 | 0.167 | 0.036 | 1.5166936 | myoFB | chr6-38311523-3831 |
| 3.21358611 | 2.24047558321056 | 0.172 | 0.039 | 2.3548066 | myoFB | chr5-95303257-9530 |
| 2.92869163 | 2.25188894940118 | 0.173 | 0.039 | 2.1460456 | myoFB | chr15-2721213-2721 |
| 8.56413591 | 2.2602671680047 | 0.164 | 0.035 | 6.2755076 | myoFB | chr1-186964501-186 |
| 2.18348240 | 2.40600933661908 | 0.15 | 0.03 | 1.5999816 | myoFB | chr15-78051972-780 |
| 1.22782760 | 2.30477484448659 | 0.155 | 0.032 | 8.9971032 | myoFB | chr17-49018264-490 |
| 3.08948139 | 2.68443761211602 | 0.119 | 0.019 | 2.2638669 | myoFB | chr7-75897160-7589 |
| 1.77004926 | 2.39530782997427 | 0.147 | 0.029 | 1.2970319 | myoFB | chr2-57775945-5777 |
| 1.72253426 | 2.94103892129215 | 0.104 | 0.014 | 1.2622145 | myoFB | chr9-80588556-8058 |
| 2.25046416 | 2.5828739953197 | 0.123 | 0.02 | 1.6490636 | myoFB | chr3-144514041-144 |
| 8.27711398 | 2.36122089393095 | 0.15 | 0.031 | 6.0651877 | myoFB | chr2-7021409-70219 |
| 1.74018280 | 2.53174764926892 | 0.148 | 0.03 | 1.2751467 | myoFB | chr9-102304461-102 |
| 1.08402019 | 2.41211893843275 | 0.146 | 0.029 | 7.9433314 | myoFB | chr9-82651035-8265 |
| 1.33067627 | 2.87032699949472 | 0.111 | 0.017 | 9.7507433 | myoFB | chr9-56918401-5691 |
| 1.00170313 | 2.50719311136112 | 0.136 | 0.025 | 7.3401399 | myoFB | chr15-42400230-424 |
| 2.17863394 | 2.34620966984243 | 0.148 | 0.03 | 1.5964288 | myoFB | chr6-92239304-9223 |
| 8.20017664 | 2.25028663355477 | 0.167 | 0.038 | 6.0088106 | myoFB | chr7-10431854-1043 |
| 3.70815740 | 2.32838653246777 | 0.15 | 0.031 | 2.7172116 | myoFB | chr6-55355473-5535 |
| 3.88167147 | 2.46648451965449 | 0.143 | 0.029 | 2.8443568 | myoFB | chr2-186537569-186 |
| 5.54866283 | 2.48401582416483 | 0.132 | 0.025 | 4.0658714 | myoFB | chr3-159776703-159 |
| 3.91108902 | 2.40983417630844 | 0.14 | 0.028 | 2.8659130 | myoFB | chr3-62683437-6268 |
| 6.85543538 | 2.5626338759377 | 0.133 | 0.025 | 5.0234299 | myoFB | chr6-33278785-3327 |

| | | | | | | | |
|---|---|---|---|---|---|---|---|
| 5.20994051 | 2.27643528909081 | | 0.158 | 0.035 | 3.8176672 | myoFB | chr7-35532105-3553 |
| 1.51692575 | 2.52098456923746 | | 0.124 | 0.022 | 1.1115516 | myoFB | chr3-55093059-5509 |
| 7.68581293 | 2.35275528451158 | | 0.143 | 0.029 | 5.6319024 | myoFB | chr7-103357767-103 |
| 3.31450269 | 2.53032798949339 | | 0.133 | 0.025 | 2.4287548 | myoFB | chr4-30905238-3090 |
| 5.13951403 | 2.28266823822786 | | 0.149 | 0.032 | 3.7660611 | myoFB | chr8-27134989-2713 |
| 1.04760994 | | 2.284909385 | 0.143 | 0.03 | 7.6765294 | myoFB | chr8-41010516-4101 |
| 1.17181853 | 2.44011852172398 | | 0.13 | 0.025 | 8.5866877 | myoFB | chr2-116077908-116 |
| 1.44035577 | 2.35305032500899 | | 0.142 | 0.03 | 1.0554437 | myoFB | chr8-41725107-4172 |
| 3.74477616 | 2.3676709993463 | | 0.14 | 0.029 | 2.7440446 | myoFB | chr15-52646612-526 |
| 1.76478840 | 2.25379198206812 | | 0.148 | 0.032 | 1.2931769 | myoFB | chr9-70870904-7087 |
| 3.42860803 | 2.42854318998181 | | 0.134 | 0.027 | 2.5123673 | myoFB | chr4-116939055-116 |
| 5.69258466 | 2.76681178416353 | | 0.104 | 0.016 | 4.1713324 | myoFB | chr16-50506409-505 |
| 3.48254880 | 2.72952584137677 | | 0.105 | 0.016 | 2.5518933 | myoFB | chr5-77511530-7751 |
| 3.26571569 | 2.40163985973278 | | 0.137 | 0.028 | 2.3930054 | myoFB | chr7-120843419-120 |
| 1.30428869 | 2.27301465129012 | | 0.145 | 0.032 | 9.5573840 | myoFB | chr11-75839176-758 |
| 1.36122696 | 2.24710285883853 | | 0.152 | 0.035 | 9.9746083 | myoFB | chr2-220423622-220 |
| 1.37564180 | 2.26449458441613 | | 0.148 | 0.033 | 1.0080235 | myoFB | chr18-39882311-398 |
| 9.87252787 | 2.67608428257048 | | 0.108 | 0.018 | 7.2342527 | myoFB | chr7-123898834-123 |
| 2.90371167 | 2.36786774809408 | | 0.132 | 0.027 | 2.1277411 | myoFB | chr8-29296973-2929 |
| 3.49669747 | 2.44135657037659 | | 0.127 | 0.025 | 2.5622610 | myoFB | chr19-2903407-2903 |
| 1.39765052 | 2.48833205188671 | | 0.12 | 0.023 | 1.0241507 | myoFB | chr14-61506153-615 |
| 4.78362814 | 2.46446242271136 | | 0.13 | 0.026 | 3.5052800 | myoFB | chr2-13568456-1356 |
| 6.58396760 | 2.49872876093467 | | 0.117 | 0.022 | 4.8245076 | myoFB | chr19-15742842-157 |
| 4.14716855 | 2.3764975187755 | | 0.126 | 0.025 | 3.0389041 | myoFB | chr5-98899573-9890 |
| 1.83110963 | 2.53460261919526 | | 0.114 | 0.021 | 1.3417748 | myoFB | chr8-92183266-9218 |
| 2.81312607 | 2.27102909610504 | | 0.132 | 0.028 | 2.0613631 | myoFB | chr3-101918482-101 |
| 1.06680769 | 2.67715686526056 | | 0.11 | 0.02 | 7.8172040 | myoFB | chr9-111695888-111 |
| 1.02238154 | 2.36679529152293 | | 0.119 | 0.023 | 7.4916643 | myoFB | chr19-8881642-8882 |
| 1.05941625 | 2.36523839049424 | | 0.138 | 0.031 | 7.7630420 | myoFB | chr17-65340473-653 |
| 2.20374384 | 2.33762061759566 | | 0.126 | 0.026 | 1.6148285 | myoFB | chr13-95494887-954 |
| 3.62927460 | 2.32763028193822 | | 0.126 | 0.026 | 2.6594090 | myoFB | chr14-93387638-933 |
| 4.01354826 | 2.53283840814832 | | 0.113 | 0.021 | 2.9409917 | myoFB | chr14-11015075-110 |
| 3.41397757 | 2.60602915005665 | | 0.109 | 0.02 | 2.5016466 | myoFB | chr2-150787746-150 |
| 4.47938742 | 2.42273307553405 | | 0.119 | 0.023 | 3.2823428 | myoFB | chr4-15351250-1535 |
| 1.77515518 | 2.45777691056114 | | 0.116 | 0.022 | 1.3007733 | myoFB | chr14-38837302-388 |
| 3.43431950 | 2.44817812919539 | | 0.119 | 0.023 | 2.5165525 | myoFB | chr7-27234961-2723 |
| 1.12953213 | 2.44277144225303 | | 0.126 | 0.026 | 8.2768274 | myoFB | chr17-16093729-160 |
| 4.01806865 | 2.32553346472478 | | 0.133 | 0.029 | 2.9443040 | myoFB | chr1-8173975-81744 |
| 1.41323473 | 2.42482174710082 | | 0.115 | 0.022 | 1.0355703 | myoFB | chr6-36581590-3658 |
| 6.68174212 | 2.37691509077474 | | 0.118 | 0.023 | 4.8961534 | myoFB | chr2-143670987-143 |
| 4.23690303 | 2.57103302094257 | | 0.105 | 0.018 | 3.1046584 | myoFB | chr2-119688869-119 |
| 4.61012237 | 2.36198438245438 | | 0.121 | 0.024 | 3.3781409 | myoFB | chr4-100829992-100 |
| 1.06325075 | 2.44156945804337 | | 0.126 | 0.027 | 7.7911400 | myoFB | chr18-6647616-6648 |

| | | | | | | |
|---|---|---|---|---|---|---|
| 1.55118316 | 2.32420254237539 | 0.13 | 0.028 | 1.1366542 | myoFB | chr2-33562823-3356 |
| 3.32648909 | 2.32978212386143 | 0.123 | 0.026 | 2.4375381 | myoFB | chr8-14186206-1418 |
| 4.97543888 | 2.29981519260191 | 0.115 | 0.022 | 3.6458324 | myoFB | chr19-26525746-265 |
| 7.40870924 | 2.28185313711635 | 0.13 | 0.029 | 5.4288502 | myoFB | chr8-88069612-8807 |
| 1.84648976 | 2.47171282435704 | 0.106 | 0.019 | 1.3530449 | myoFB | chr6-67935608-6793 |
| 2.01193813 | 2.5279354602351 | 0.106 | 0.019 | 1.4742798 | myoFB | chr3-62695600-6269 |
| 1.03488431 | 2.2807998505428 | 0.13 | 0.029 | 7.5832803 | myoFB | chr18-56626188-566 |
| 1.68086885 | 2.24504024713275 | 0.127 | 0.028 | 1.2316835 | myoFB | chr4-40488984-4048 |
| 3.64713674 | 2.56024374278384 | 0.103 | 0.018 | 2.6724978 | myoFB | chr2-124565700-124 |
| 9.87310966 | 2.39525096226404 | 0.11 | 0.021 | 7.2346790 | myoFB | chr4-141862154-141 |
| 1.27249621 | 2.51901874020541 | 0.101 | 0.018 | 9.3244195 | myoFB | chr6-18973921-1897 |
| 2.55813842 | 2.36921266962224 | 0.113 | 0.022 | 1.8745168 | myoFB | chr10-48016521-480 |
| 2.82046847 | 2.28239701003316 | 0.123 | 0.027 | 2.0667434 | myoFB | chr8-7465451-74659 |
| 3.06645163 | 2.33073022262472 | 0.109 | 0.021 | 2.2469914 | myoFB | chr15-14368291-143 |
| 4.99754059 | 2.47446309870503 | 0.109 | 0.021 | 3.6620278 | myoFB | chr5-8263656-82641 |
| 7.18938978 | 2.39713052497417 | 0.119 | 0.025 | 5.2681403 | myoFB | chr19-25393113-253 |
| 1.29411504 | 2.39782809349772 | 0.11 | 0.022 | 9.4828350 | myoFB | chr11-32122492-321 |
| 2.40857477 | 2.33189646754118 | 0.112 | 0.022 | 1.7649217 | myoFB | chr8-84932800-8493 |
| 2.81043554 | 2.25939769471141 | 0.122 | 0.026 | 2.0593916 | myoFB | chr8-79907437-7990 |
| 9.04578033 | 2.38342988038919 | 0.106 | 0.02 | 6.6284402 | myoFB | chr3-109124324-109 |
| 2.52336835 | 2.34220869166673 | 0.117 | 0.024 | 1.8490385 | myoFB | chr7-85397133-8539 |
| 2.57469621 | 2.38710023831837 | 0.11 | 0.022 | 1.8866498 | myoFB | chr1-101674187-101 |
| 3.00522345 | 2.48914475509001 | 0.105 | 0.02 | 2.2021255 | myoFB | chr6-27538761-2753 |
| 2.33932183 | 2.39544733995775 | 0.117 | 0.025 | 1.7141755 | myoFB | chr6-98874021-9887 |
| 1.04153269 | 2.35228819602423 | 0.114 | 0.023 | 7.6319975 | myoFB | chr5-27278251-2727 |
| 1.32202067 | 2.29595774658113 | 0.112 | 0.023 | 9.6873180 | myoFB | chr14-71354324-713 |
| 4.85032455 | 2.48293574902269 | 0.105 | 0.02 | 3.5541529 | myoFB | chr3-70066813-7006 |
| 4.61397308 | 2.44018349245099 | 0.106 | 0.021 | 3.3809626 | myoFB | chr17-67541523-675 |
| 5.38766851 | 2.24535693657058 | 0.121 | 0.027 | 3.9479003 | myoFB | chr3-37565316-3756 |
| 5.31073687 | 2.27295583065831 | 0.115 | 0.025 | 3.8915274 | myoFB | chr10-70847071-708 |
| 1.69750686 | 2.29789162318389 | 0.111 | 0.023 | 1.2438753 | myoFB | chr20-42370390-423 |
| 5.11539933 | 2.30638351984958 | 0.109 | 0.022 | 3.7483907 | myoFB | chr17-17749744-177 |
| 1.27059234 | 2.42006006746695 | 0.111 | 0.023 | 9.3104687 | myoFB | chr6-68091886-6809 |
| 2.84891499 | 2.2906236365754 | 0.126 | 0.029 | 2.0875880 | myoFB | chr4-96266160-9626 |
| 6.65390032 | 2.41143948436956 | 0.101 | 0.019 | 4.8757519 | myoFB | chr9-70885822-7088 |
| 6.93367255 | 2.32580880473339 | 0.109 | 0.022 | 5.0807595 | myoFB | chr19-1871797-1872 |
| 1.61062158 | 2.35985944851344 | 0.108 | 0.022 | 1.1802087 | myoFB | chr20-36091575-360 |
| 2.13401487 | 2.24895714761562 | 0.121 | 0.027 | 1.5637335 | myoFB | chr4-125887738-125 |
| 2.59207437 | 2.36638835493598 | 0.105 | 0.021 | 1.8993839 | myoFB | chr9-63215013-6321 |
| 3.44100754 | 2.24488202947487 | 0.118 | 0.026 | 2.5214533 | myoFB | chr10-21579912-215 |
| 1.28315592 | 2.28974495727916 | 0.11 | 0.023 | 9.4025303 | myoFB | chr2-83600720-8360 |
| 1.59329977 | 2.41861231776738 | 0.1 | 0.019 | 1.1675159 | myoFB | chr16-79789626-797 |
| 2.83379289 | 2.4006586694628 | 0.109 | 0.023 | 2.0765070 | myoFB | chr1-56209041-5620 |

| | | | | | | |
|---|---|---|---|---|---|---|
| 3.48195617 | 2.26804340989356 | 0.116 | 0.026 | 2.5514591 | myoFB | chr1-232785233-232 |
| 1.12927604 | 2.39751808818148 | 0.108 | 0.022 | 8.2749509 | myoFB | chr4-12252042-1225 |
| 2.74547121 | 2.42632638055273 | 0.104 | 0.021 | 2.0117879 | myoFB | chr16-81230722-812 |
| 6.61887796 | 2.42535794291199 | 0.104 | 0.021 | 4.8500887 | myoFB | chr2-25100399-2510 |
| 8.50984072 | 2.38475463408889 | 0.101 | 0.019 | 6.2357219 | myoFB | chr10-54874387-548 |
| 8.52372276 | 2.45303415732928 | 0.101 | 0.02 | 6.2458942 | myoFB | chr16-60726793-607 |
| 5.58249024 | 2.42320763773169 | 0.104 | 0.021 | 4.0906590 | myoFB | chr8-91493990-9149 |
| 7.23858679 | 2.3466152980235 | 0.102 | 0.02 | 5.3041902 | myoFB | chr11-79529853-795 |
| 2.46919778 | 2.33851890547892 | 0.105 | 0.021 | 1.8093441 | myoFB | chr18-47237766-472 |
| 4.66135511 | 2.28814229642269 | 0.113 | 0.025 | 3.4156825 | myoFB | chr2-52985864-5298 |
| 1.97426465 | 2.24936350206131 | 0.113 | 0.025 | 1.4466740 | myoFB | chr8-25789362-2578 |
| 2.63133023 | 2.30244326919463 | 0.113 | 0.025 | 1.9281493 | myoFB | chr19-16600038-166 |
| 3.78586112 | 2.30177033651295 | 0.11 | 0.024 | 2.7741503 | myoFB | chr19-55576310-555 |
| 1.44169532 | 2.30414469818344 | 0.106 | 0.022 | 1.0564253 | myoFB | chr9-101907212-101 |
| 2.48721794 | 2.33546992783031 | 0.102 | 0.021 | 1.8225487 | myoFB | chr5-32603797-3260 |
| 4.20526421 | 2.35517298625007 | 0.104 | 0.022 | 3.0814746 | myoFB | chr4-150849220-150 |
| 9.66420297 | 2.29102062050339 | 0.109 | 0.024 | 7.0815993 | myoFB | chr7-57252330-5725 |
| 1.30391260 | 2.28329617467919 | 0.103 | 0.022 | 9.5546282 | myoFB | chr13-2228215-2228 |
| 1.78138715 | 2.44085675999415 | 0.1 | 0.02 | 1.3053399 | myoFB | chr18-44458096-444 |
| 7.94109136 | 2.2636198208716 | 0.105 | 0.022 | 5.8189617 | myoFB | chr2-20966744-2096 |
| 1.02927578 | 2.28759552495547 | 0.105 | 0.022 | 7.5421830 | myoFB | chr2-25238025-2523 |
| 6.46322338 | 2.3425362719585 | 0.105 | 0.023 | 4.7360303 | myoFB | chr9-99151599-9915 |
| 1.51274165 | 2.2725844334767 | 0.109 | 0.024 | 1.1084856 | myoFB | chr11-78141834-781 |
| 8.73541526 | 2.26416332797555 | 0.1 | 0.021 | 6.4010152 | myoFB | chr3-53169974-5317 |
| 1.00935189 | 2.30593673892097 | 0.1 | 0.021 | 7.3961875 | myoFB | chr8-9908685-99091 |
| 8.56419963 | 2.35471973890518 | 0.101 | 0.021 | 6.2755543 | myoFB | chr3-127370241-127 |
| 2.20322629 | 2.26001377653044 | 0.101 | 0.021 | 1.6144493 | myoFB | chr3-53949154-5394 |
| 3.80816441 | 2.35055413265388 | 0.102 | 0.022 | 2.7904934 | myoFB | chr1-91430188-9143 |
| 2.14082530 | 2.24559627012173 | 0.103 | 0.022 | 1.5687239 | myoFB | chr15-35235085-352 |
| 8.69461043 | 2.24142107567786 | 0.104 | 0.023 | 6.3711149 | myoFB | chr13-63808570-638 |
| 9.38489588 | 2.29474971625806 | 0.1 | 0.021 | 6.8769326 | myoFB | chr8-79713832-7971 |
| 2.45534163 | 2.27868343307072 | 0.108 | 0.025 | 1.7991908 | myoFB | chr1-11672640-1167 |
| 4.74078742 | 2.24351125999198 | 0.107 | 0.024 | 3.4738878 | myoFB | chr12-26572979-265 |
| 7.41035924 | 2.26246387467286 | 0.103 | 0.023 | 5.4300593 | myoFB | chr2-230211279-230 |
| 4.38081892 | 2.27757873897134 | 0.1 | 0.022 | 3.2101151 | myoFB | chr17-79564983-795 |
| 0 | 3.60428817192785 | 0.1 | 0.011 | 0 | papiFB | chr1-82475445-8247 |
| 0 | 2.90333620627359 | 0.103 | 0.016 | 0 | papiFB | chr1-169618685-169 |
| 0 | 3.09444792973296 | 0.101 | 0.014 | 0 | papiFB | chr1-184767035-184 |
| 0 | 2.78044752332241 | 0.113 | 0.021 | 0 | papiFB | chr1-197725487-197 |
| 0 | 2.81915715160045 | 0.138 | 0.03 | 0 | papiFB | chr10-98937229-989 |
| 0 | 3.14364865365655 | 0.108 | 0.018 | 0 | papiFB | chr11-32646931-326 |
| 0 | 2.89531092311031 | 0.111 | 0.019 | 0 | papiFB | chr15-55109889-551 |
| 0 | 2.78168312247642 | 0.114 | 0.021 | 0 | papiFB | chr2-196382949-196 |

| | | | | | | | |
|---|---|---|---|---|---|---|---|
| 0 | 2.71675010740135 | | 0.157 | 0.034 | 0 | papiFB | chr3-152822382-152 |
| 0 | 3.01316821985378 | | 0.108 | 0.018 | 0 | papiFB | chr4-64034604-6403 |
| 0 | 3.16073455738911 | | 0.112 | 0.014 | 0 | papiFB | chr4-176523616-176 |
| 0 | 2.97138887790899 | | 0.122 | 0.02 | 0 | papiFB | chr5-76612428-7661 |
| 0 | 3.12811103109363 | | 0.101 | 0.016 | 0 | papiFB | chr7-133236587-133 |
| 0 | 3.47289845891586 | | 0.118 | 0.014 | 0 | papiFB | chr9-11369288-1136 |
| 0 | 2.87384765747824 | | 0.104 | 0.016 | 0 | papiFB | chr9-13118259-1311 |
| 0 | 3.57107768004723 | | 0.105 | 0.011 | 0 | papiFB | chr9-68104484-6810 |
| 2.85977661 | 2.6860328859473 | | 0.108 | 0.02 | 2.0955470 | papiFB | chr8-103262927-103 |
| 6.88183722 | 2.85316460203556 | | 0.1 | 0.017 | 5.0427763 | papiFB | chr15-98280979-982 |
| 3.39906147 | 2.97352632773012 | | 0.1 | 0.017 | 2.4907166 | papiFB | chr9-26605796-2660 |
| 6.14238109 | 2.48293628538659 | | 0.114 | 0.024 | 4.5009280 | papiFB | chr4-33108076-3310 |
| 1.52998525 | 2.70036119902446 | | 0.103 | 0.019 | 1.1211211 | papiFB | chr3-89976936-8997 |
| 5.67672263 | 2.67719154490888 | | 0.115 | 0.025 | 4.1597093 | papiFB | chr4-177004825-177 |
| 7.45815802 | 2.52980020357499 | | 0.112 | 0.023 | 5.4650846 | papiFB | chr5-122366504-122 |
| 5.62738592 | | 2.738423824 | 0.113 | 0.025 | 4.1235570 | papiFB | chr4-63940927-6394 |
| 4.58548441 | 2.5600005764724 | | 0.116 | 0.026 | 3.3600870 | papiFB | chr19-35385798-353 |
| 1.75016483 | 2.49847081813912 | | 0.112 | 0.024 | 1.2824612 | papiFB | chr10-98250914-982 |
| 1.20299889 | 2.50257754087052 | | 0.121 | 0.029 | 8.8151668 | papiFB | chr15-20588428-205 |
| 8.08105827 | 2.70459143786402 | | 0.102 | 0.02 | 5.9215247 | papiFB | chr8-97515735-9751 |
| 3.27945502 | 2.67084362781174 | | 0.108 | 0.024 | 2.4030731 | papiFB | chr7-20593533-2059 |
| 1.13477658 | 2.5699625650748 | | 0.101 | 0.021 | 8.3152569 | papiFB | chr7-88634529-8863 |
| 2.56321187 | 2.55522937459969 | | 0.107 | 0.024 | 1.8782345 | papiFB | chr13-66329854-663 |
| 6.54718253 | 2.49077111487111 | | 0.114 | 0.028 | 4.7975527 | papiFB | chr8-57281265-5728 |
| 5.03404453 | 2.51548611403787 | | 0.105 | 0.023 | 3.6887766 | papiFB | chr8-114041268-114 |
| 4.38222732 | 2.1427952994731 | | 0.136 | 0.04 | 3.2111471 | papiFB | chr2-78571321-7857 |
| 2.11984798 | | 2.412708418 | 0.11 | 0.026 | 1.5533525 | papiFB | chr12-31652902-316 |
| 9.30102591 | 2.31253769967024 | | 0.12 | 0.031 | 6.8154755 | papiFB | chr1-29842913-2984 |
| 1.03007379 | 2.0447042024613 | | 0.137 | 0.042 | 7.5480305 | papiFB | chr3-141182001-141 |
| 6.31191448 | 2.64389299634125 | | 0.1 | 0.022 | 4.6251563 | papiFB | chr1-65874800-6587 |
| 2.20934941 | 2.14921663047674 | | 0.124 | 0.035 | 1.6189361 | papiFB | chr18-73985213-739 |
| 2.06547725 | 2.17206944399627 | | 0.119 | 0.033 | 1.5135115 | papiFB | chr16-78331547-783 |
| 9.86758296 | 2.07753451270631 | | 0.145 | 0.048 | 7.2306292 | papiFB | chr19-35458936-354 |
| 6.26623416 | 2.53359878978316 | | 0.1 | 0.023 | 4.5916833 | papiFB | chr12-8724048-8724 |
| 3.49335681 | 2.24021886292633 | | 0.112 | 0.031 | 2.5598131 | papiFB | chr18-74081686-740 |
| 2.03376159 | 2.29338853093968 | | 0.1 | 0.024 | 1.4902713 | papiFB | chr4-177924309-177 |
| 9.17123474 | 2.25351956494303 | | 0.11 | 0.03 | 6.7203689 | papiFB | chr7-129127472-129 |
| 6.34622616 | 1.88991714374901 | | 0.151 | 0.054 | 4.6502987 | papiFB | chr1-95158868-9515 |
| 2.63785222 | 2.05534820197827 | | 0.125 | 0.039 | 1.9329284 | papiFB | chr7-133388225-133 |
| 5.04574187 | 2.06115304332001 | | 0.124 | 0.038 | 3.6973480 | papiFB | chr1-58336941-5833 |
| 1.48766165 | 2.30364421088448 | | 0.104 | 0.027 | 1.0901078 | papiFB | chr1-254986322-254 |
| 5.75341942 | 1.76427496194844 | | 0.141 | 0.049 | 4.2159101 | papiFB | chr12-24852800-248 |
| 1.11884681 | 1.92496436046541 | | 0.112 | 0.034 | 8.1985290 | papiFB | chr3-155435431-155 |

| | | | | | | | |
|---|---|---|---|---|---|---|---|
| 1.07561507 | 2.01238695474404 | | 0.104 | 0.03 | 7.8817415 | papiFB | chr1-184381983-184 |
| 1.04815026 | 2.15593855067707 | | 0.111 | 0.034 | 7.6804887 | papiFB | chr14-77030313-770 |
| 1.15919410 | 1.8248021087445 | | 0.113 | 0.035 | 8.4941802 | papiFB | chr19-53993055-539 |
| 3.75010206 | 1.77536890670536 | | 0.116 | 0.038 | 2.7479472 | papiFB | chr14-83637796-836 |
| 5.93685743 | 1.83897943967488 | | 0.131 | 0.048 | 4.3503272 | papiFB | chr8-103309881-103 |
| 1.72676793 | 1.91456954336569 | | 0.101 | 0.03 | 1.2653168 | papiFB | chr14-83278795-832 |
| 4.38809265 | 1.82445017000898 | | 0.123 | 0.043 | 3.2154450 | papiFB | chr11-35159810-351 |
| 1.49250961 | 1.75967635845364 | | 0.114 | 0.04 | 1.0936603 | papiFB | chr11-34143290-341 |
| 2.88348322 | 1.81216322584138 | | 0.1 | 0.033 | 2.1129184 | papiFB | chr2-196406182-196 |
| 3.75609084 | 1.64085072375898 | | 0.116 | 0.043 | 2.7523356 | papiFB | chr5-21737834-2173 |
| 4.12741348 | 1.2624282973847 | | 0.153 | 0.068 | 3.0244282 | papiFB | chr18-51456411-514 |
| 1.92383517 | 1.77917462832631 | | 0.125 | 0.05 | 1.4097210 | papiFB | chr1-187161773-187 |
| 5.81438170 | 1.65627779997713 | | 0.102 | 0.037 | 4.2605812 | papiFB | chr17-12000919-120 |
| 1.37207299 | 1.65724497785868 | | 0.125 | 0.051 | 1.0054084 | papiFB | chr4-179814695-179 |
| 4.14487291 | 1.73785900421063 | | 0.113 | 0.044 | 3.0372219 | papiFB | chr1-223035874-223 |
| 6.39377354 | 1.44444231903418 | | 0.143 | 0.064 | 4.6851398 | papiFB | chr3-8915267-89157 |
| 2.44224136 | 1.62291917011682 | | 0.107 | 0.042 | 1.7895914 | papiFB | chr16-2529022-2529 |
| 5.56838169 | | 1.435146427 | 0.111 | 0.044 | 4.0803207 | papiFB | chr6-8707830-87083 |
| 5.81717159 | 1.67230772700879 | | 0.134 | 0.062 | 4.2626255 | papiFB | chr2-110513158-110 |
| 4.81592624 | 1.48108268815973 | | 0.106 | 0.042 | 3.5289470 | papiFB | chr15-17022377-170 |
| 1.30791069 | 1.39930409032272 | | 0.129 | 0.058 | 9.5839249 | papiFB | chr20-5999524-6000 |
| 1.25706170 | 1.5529579893467 | | 0.108 | 0.044 | 9.2113207 | papiFB | chr17-13129846-131 |
| 2.96487173 | 1.48034667737564 | | 0.136 | 0.064 | 2.1725572 | papiFB | chr17-80480332-804 |
| 6.49181915 | 1.43949986184607 | | 0.105 | 0.043 | 4.7569843 | papiFB | chr4-133871789-133 |
| 2.57363606 | 1.59153819007256 | | 0.115 | 0.051 | 1.8858730 | papiFB | chr1-230418212-230 |
| 6.87630721 | 1.46995947132012 | | 0.13 | 0.061 | 5.0387241 | papiFB | chr2-234601071-234 |
| 7.48544709 | 1.30847448974209 | | 0.122 | 0.055 | 5.4850811 | papiFB | chr18-69105147-691 |
| 7.57982126 | 1.38553468858459 | | 0.11 | 0.048 | 5.5542353 | papiFB | chr7-53271875-5327 |
| 9.96453251 | 1.62805075409552 | | 0.101 | 0.042 | 7.3016706 | papiFB | chr15-17086559-170 |
| 7.37187155 | 1.55682321070119 | | 0.112 | 0.05 | 5.4018568 | papiFB | chr11-34719132-347 |
| 3.42785342 | 1.43522692520774 | | 0.11 | 0.048 | 2.5118144 | papiFB | chr17-8563005-8563 |
| 1.86635162 | 1.28332960021873 | | 0.13 | 0.062 | 1.3675990 | papiFB | chr7-53117918-5311 |
| 1.10326697 | 1.48499907093132 | | 0.102 | 0.043 | 8.0843653 | papiFB | chr8-48080882-4808 |
| 3.61873350 | 1.48861199157661 | | 0.114 | 0.052 | 2.6516848 | papiFB | chr11-31892825-318 |
| 6.39091570 | 1.23095781958341 | | 0.133 | 0.066 | 4.6830457 | papiFB | chr9-102452199-102 |
| 1.19073958 | 1.36856675973922 | | 0.101 | 0.044 | 8.7253348 | papiFB | chr8-118459496-118 |
| 4.53809015 | 1.24608706744971 | | 0.117 | 0.055 | 3.3253581 | papiFB | chr5-147538244-147 |
| 1.27354163 | 1.26171625122541 | | 0.118 | 0.056 | 9.3320800 | papiFB | chr18-74077110-740 |
| 4.22393421 | 1.17411219296007 | | 0.158 | 0.086 | 3.0951553 | papiFB | chr10-10916368-109 |
| 1.05693423 | 1.37346813837028 | | 0.102 | 0.045 | 7.7448547 | papiFB | chr15-36360367-363 |
| 1.02236986 | 1.12781258481953 | | 0.147 | 0.078 | 7.4915787 | papiFB | chr11-78618415-786 |
| 9.67964901 | 1.29243954887541 | | 0.114 | 0.054 | 7.0929176 | papiFB | chr2-219427322-219 |
| 7.43336034 | 1.08140640585372 | | 0.196 | 0.118 | 5.4469137 | papiFB | chr10-47637129-476 |

| | | | | | | |
|---|---|---|---|---|---|---|
| 1.44011013 | 1.10820433949661 | 0.127 | 0.064 | 1.0552637 | papiFB | chr2-55406883-5540 |
| 4.56210793 | 1.33276905798234 | 0.105 | 0.049 | 3.3429575 | papiFB | chr5-138576143-138 |
| 5.96115215 | 1.42620713695556 | 0.114 | 0.055 | 4.3681296 | papiFB | chr11-76547183-765 |
| 4.68147489 | 1.32310820665305 | 0.107 | 0.051 | 3.4304256 | papiFB | chr13-39007055-390 |
| 1.95652632 | 1.14776711547365 | 0.138 | 0.074 | 1.4336759 | papiFB | chr15-29331619-293 |
| 1.06517238 | 1.14929330745089 | 0.115 | 0.058 | 7.8052210 | papiFB | chr1-137405451-137 |
| 8.28994809 | 1.19409622775368 | 0.127 | 0.069 | 6.0745921 | papiFB | chr7-56382797-5638 |
| 3.88154633 | 1.14586064765922 | 0.147 | 0.085 | 2.8442651 | papiFB | chr3-157268803-157 |
| 3.34098486 | 1.06413633852309 | 0.116 | 0.061 | 2.4481601 | papiFB | chr5-133237989-133 |
| 3.81598674 | 0.885506883118284 | 0.17 | 0.104 | 2.7962253 | papiFB | chr12-38048656-380 |
| 6.47210381 | 1.25641664081829 | 0.103 | 0.052 | 4.7425376 | papiFB | chr20-45607495-456 |
| 2.39325893 | 1.04962884819439 | 0.128 | 0.071 | 1.7536987 | papiFB | chr7-69628578-6962 |
| 5.87377143 | 1.01504775738945 | 0.13 | 0.073 | 4.3040999 | papiFB | chr1-136950640-136 |
| 1.28650558 | 1.09903988003591 | 0.123 | 0.068 | 9.4270755 | papiFB | chr13-74389057-743 |
| 3.06171234 | 1.10566361487194 | 0.136 | 0.079 | 2.2435187 | papiFB | chr6-11529513-1153 |
| 3.29804238 | 1.0212406483972 | 0.133 | 0.076 | 2.4166933 | papiFB | chr17-15472171-154 |
| 1.54931749 | 1.10828585975414 | 0.112 | 0.06 | 1.1352871 | papiFB | chr13-42571497-425 |
| 8.89924683 | 0.988198265874724 | 0.121 | 0.068 | 6.5210655 | papiFB | chr1-80835862-8083 |
| 3.54812103 | 1.13621486407871 | 0.113 | 0.062 | 2.5999424 | papiFB | chr6-95036103-9503 |
| 3.84384932 | 0.805751594963119 | 0.159 | 0.098 | 2.8166420 | papiFB | chr2-201820348-201 |
| 1.69986242 | 1.10642250445351 | 0.127 | 0.073 | 1.2456013 | papiFB | chr17-1953736-1954 |
| 3.05298703 | 1.04925235343883 | 0.102 | 0.054 | 2.2371251 | papiFB | chr12-31698395-316 |
| 7.01415922 | 1.02644861330367 | 0.109 | 0.059 | 5.1397373 | papiFB | chr8-103976786-103 |
| 8.35106841 | 1.00648106402071 | 0.136 | 0.081 | 6.1193789 | papiFB | chr3-156306864-156 |
| 1.61070150 | 1.14871582558221 | 0.111 | 0.062 | 1.1802672 | papiFB | chr3-78397078-7839 |
| 1.84709258 | 0.806396026108635 | 0.155 | 0.096 | 1.3534866 | papiFB | chr1-95080823-9508 |
| 4.55244948 | 1.01058552243948 | 0.15 | 0.093 | 3.3358801 | papiFB | chr2-182806658-182 |
| 1.55374246 | 1.02445511672496 | 0.119 | 0.068 | 1.1385296 | papiFB | chr9-13005985-1300 |
| 5.60709669 | 1.14929000034713 | 0.108 | 0.06 | 4.1086898 | papiFB | chr15-97510850-975 |
| 6.40046002 | 0.975406556497782 | 0.107 | 0.059 | 4.6900394 | papiFB | chr14-91236035-912 |
| 1.09917522 | 0.848920428856934 | 0.161 | 0.102 | 8.0543823 | papiFB | chr8-61890578-6189 |
| 3.65200493 | 0.969872469403864 | 0.117 | 0.067 | 2.6760650 | papiFB | chr1-184252762-184 |
| 4.27660278 | 0.94241747230547 | 0.121 | 0.07 | 3.1337491 | papiFB | chr3-155141233-155 |
| 9.00792802 | 0.974281756823358 | 0.107 | 0.059 | 6.6007033 | papiFB | chr2-195803264-195 |
| 2.95253445 | 0.914407350495132 | 0.118 | 0.069 | 2.1635168 | papiFB | chr15-52222077-522 |
| 4.71476128 | 1.08325501694698 | 0.137 | 0.085 | 3.4548167 | papiFB | chr18-61812310-618 |
| 1.07931337 | 0.905761818766965 | 0.139 | 0.086 | 7.9088414 | papiFB | chr1-200369740-200 |
| 5.63403062 | 0.954471415594429 | 0.108 | 0.061 | 4.1284260 | papiFB | chr1-197669786-197 |
| 9.00050163 | 0.971280972056925 | 0.162 | 0.107 | 6.5952615 | papiFB | chr7-62758095-6275 |
| 4.66631802 | 0.982965851666447 | 0.103 | 0.058 | 3.4193191 | papiFB | chr16-66529316-665 |
| 2.25778772 | 1.13945430221685 | 0.106 | 0.062 | 1.6544300 | papiFB | chr1-44046908-4404 |
| 2.73234911 | 0.928590910302789 | 0.1 | 0.057 | 2.0021725 | papiFB | chr7-125560584-125 |
| 1.15867996 | 0.957075646786264 | 0.103 | 0.06 | 8.4904128 | papiFB | chr17-20882838-208 |

| | | | | | | |
|---|---|---|---|---|---|---|
| 1.02991242 | 0.826537532742186 | 0.122 | 0.076 | 7.5468480 | papiFB | chr6-126853787-126 |
| 3.27871692 | 0.923627835611041 | 0.121 | 0.075 | 2.4025322 | papiFB | chr2-234616947-234 |
| 3.99034748 | 0.910690735874423 | 0.133 | 0.085 | 2.9239909 | papiFB | chr7-124656849-124 |
| 7.29539770 | 0.832767146939881 | 0.134 | 0.086 | 5.3458193 | papiFB | chr1-213219288-213 |
| 4.75029318 | 0.865622215829152 | 0.107 | 0.064 | 3.4808533 | papiFB | chr9-18839342-1883 |
| 5.81475460 | 0.845338423761918 | 0.113 | 0.069 | 4.2608544 | papiFB | chr19-16134847-161 |
| 3.10545803 | 0.846786669013324 | 0.119 | 0.074 | 2.2755740 | papiFB | chr1-197640990-197 |
| 1.12219591 | 0.860649498585175 | 0.14 | 0.092 | 8.2230701 | papiFB | chr18-75892903-758 |
| 1.03495936 | 0.85655907674695 | 0.1 | 0.06 | 7.5838303 | papiFB | chr3-156268778-156 |
| 1.86677437 | 0.801985232796907 | 0.14 | 0.095 | 1.3679087 | papiFB | chr20-12777536-127 |
| 9.23592368 | 0.838089566997748 | 0.108 | 0.071 | 6.7677708 | papiFB | chr10-64657975-646 |
| 1.03454870 | 0.911035293977972 | 0.1 | 0.065 | 7.5808211 | papiFB | chr11-70127294-701 |
| 3.65062442 | 0.812512676922639 | 0.118 | 0.084 | 2.6750534 | papiFB | chr18-68999341-689 |
| 0 | 3.65684292674712 | 0.174 | 0.016 | 0 | retiFB | chr1-2463198-24636 |
| 0 | 3.58014139034865 | 0.14 | 0.013 | 0 | retiFB | chr1-101310449-101 |
| 0 | 3.72401743868796 | 0.139 | 0.012 | 0 | retiFB | chr1-146305853-146 |
| 0 | 4.38861801244023 | 0.127 | 0.007 | 0 | retiFB | chr1-163884032-163 |
| 0 | 3.6246959302059 | 0.143 | 0.013 | 0 | retiFB | chr1-215118857-215 |
| 0 | 4.19084659919674 | 0.136 | 0.008 | 0 | retiFB | chr10-7876605-7877 |
| 0 | 3.65582164713889 | 0.152 | 0.013 | 0 | retiFB | chr10-34453491-344 |
| 0 | 3.68652035148743 | 0.159 | 0.013 | 0 | retiFB | chr10-44153004-441 |
| 0 | 4.64397275292701 | 0.109 | 0.004 | 0 | retiFB | chr11-16482679-164 |
| 0 | 3.75950807697469 | 0.114 | 0.009 | 0 | retiFB | chr11-26570186-265 |
| 0 | 3.58655803123395 | 0.197 | 0.017 | 0 | retiFB | chr11-73812715-738 |
| 0 | 3.65614185665207 | 0.163 | 0.013 | 0 | retiFB | chr12-37169819-371 |
| 0 | 3.79675352853856 | 0.112 | 0.008 | 0 | retiFB | chr13-40828150-408 |
| 0 | 3.7898813316335 | 0.168 | 0.015 | 0 | retiFB | chr13-96167935-961 |
| 0 | 4.30167765435388 | 0.115 | 0.006 | 0 | retiFB | chr13-97720236-977 |
| 0 | 3.91852387508311 | 0.112 | 0.008 | 0 | retiFB | chr14-11963584-119 |
| 0 | 3.68625765776842 | 0.135 | 0.012 | 0 | retiFB | chr14-15520412-155 |
| 0 | 4.14102693725461 | 0.152 | 0.01 | 0 | retiFB | chr14-33517754-335 |
| 0 | 3.66684739033515 | 0.17 | 0.015 | 0 | retiFB | chr14-34450314-344 |
| 0 | 3.87965835379733 | 0.134 | 0.01 | 0 | retiFB | chr14-71944699-719 |
| 0 | 3.68327054035351 | 0.127 | 0.011 | 0 | retiFB | chr14-75104517-751 |
| 0 | 3.56931831405207 | 0.132 | 0.012 | 0 | retiFB | chr15-930992-93149 |
| 0 | 3.67055014371736 | 0.132 | 0.012 | 0 | retiFB | chr15-1763396-1763 |
| 0 | 3.62189009284786 | 0.144 | 0.012 | 0 | retiFB | chr15-19869639-198 |
| 0 | 3.64345323475487 | 0.168 | 0.015 | 0 | retiFB | chr15-41585101-415 |
| 0 | 3.74690890710281 | 0.128 | 0.01 | 0 | retiFB | chr15-41775884-417 |
| 0 | 3.66082962206937 | 0.2 | 0.019 | 0 | retiFB | chr15-83218672-832 |
| 0 | 3.71408406167692 | 0.15 | 0.012 | 0 | retiFB | chr16-32961281-329 |
| 0 | 4.09895487669956 | 0.219 | 0.015 | 0 | retiFB | chr16-44912290-449 |
| 0 | 3.84504092584003 | 0.151 | 0.011 | 0 | retiFB | chr16-82694996-826 |

| | | | | | | | |
|---|---|---|---|---|---|---|---|
| 0 | 3.89655108497216 | | 0.139 | 0.01 | 0 | retiFB | chr17-31727621-317 |
| 0 | 3.72070928853349 | | 0.126 | 0.011 | 0 | retiFB | chr17-48322659-483 |
| 0 | 3.62259835149426 | | 0.134 | 0.012 | 0 | retiFB | chr17-74675054-746 |
| 0 | 3.62475112541497 | | 0.142 | 0.013 | 0 | retiFB | chr18-26982621-269 |
| 0 | 3.60818566923053 | | 0.134 | 0.012 | 0 | retiFB | chr18-40341294-403 |
| 0 | 3.59850534747002 | | 0.231 | 0.023 | 0 | retiFB | chr18-40380451-403 |
| 0 | 3.72731437636904 | | 0.122 | 0.01 | 0 | retiFB | chr18-56646392-566 |
| 0 | 3.7611368301353 | | 0.142 | 0.011 | 0 | retiFB | chr18-81966005-819 |
| 0 | 3.62103141321336 | | 0.13 | 0.011 | 0 | retiFB | chr19-17945885-179 |
| 0 | 3.96211035341446 | | 0.147 | 0.01 | 0 | retiFB | chr19-28733184-287 |
| 0 | 3.94937223683534 | | 0.15 | 0.011 | 0 | retiFB | chr19-34388200-343 |
| 0 | 4.06882540824844 | | 0.117 | 0.008 | 0 | retiFB | chr2-24100969-2410 |
| 0 | 3.85498639760067 | | 0.127 | 0.011 | 0 | retiFB | chr2-55185569-5518 |
| 0 | 4.00229091398125 | | 0.12 | 0.009 | 0 | retiFB | chr2-75327340-7532 |
| 0 | 3.69642232704583 | | 0.117 | 0.01 | 0 | retiFB | chr2-77738190-7773 |
| 0 | 4.26362044885148 | | 0.195 | 0.013 | 0 | retiFB | chr2-93141764-9314 |
| 0 | 3.7949320832011 | | 0.119 | 0.009 | 0 | retiFB | chr2-122245444-122 |
| 0 | 4.10707769308018 | | 0.103 | 0.007 | 0 | retiFB | chr2-123356935-123 |
| 0 | 3.81109827560831 | | 0.139 | 0.01 | 0 | retiFB | chr2-133958473-133 |
| 0 | 4.30732597085692 | | 0.105 | 0.006 | 0 | retiFB | chr2-168783457-168 |
| 0 | 3.7857264024106 | | 0.115 | 0.008 | 0 | retiFB | chr2-172645007-172 |
| 0 | 4.3024629530711 | | 0.104 | 0.006 | 0 | retiFB | chr20-14567281-145 |
| 0 | 3.61631036274456 | | 0.142 | 0.013 | 0 | retiFB | chr20-40080536-400 |
| 0 | 3.6086723705266 | | 0.133 | 0.011 | 0 | retiFB | chr3-88438429-8843 |
| 0 | 3.68603075924038 | | 0.15 | 0.013 | 0 | retiFB | chr3-131059812-131 |
| 0 | 3.61956596086625 | | 0.129 | 0.011 | 0 | retiFB | chr3-134961278-134 |
| 0 | 3.79185692117248 | | 0.154 | 0.012 | 0 | retiFB | chr3-135060945-135 |
| 0 | 3.81974993892896 | | 0.217 | 0.018 | 0 | retiFB | chr3-163668405-163 |
| 0 | 4.39286118487608 | | 0.187 | 0.011 | 0 | retiFB | chr4-22265497-2226 |
| 0 | 3.6778623972097 | | 0.139 | 0.012 | 0 | retiFB | chr4-32498811-3249 |
| 0 | 3.98177718030949 | | 0.199 | 0.015 | 0 | retiFB | chr4-133327445-133 |
| 0 | 3.88238149567658 | | 0.113 | 0.009 | 0 | retiFB | chr4-176484109-176 |
| 0 | 4.13091125976893 | | 0.22 | 0.013 | 0 | retiFB | chr5-5010036-50105 |
| 0 | 3.69914521900834 | | 0.149 | 0.012 | 0 | retiFB | chr5-5141205-51417 |
| 0 | 3.73715004310085 | | 0.121 | 0.01 | 0 | retiFB | chr5-5445361-54458 |
| 0 | 3.95300436105954 | | 0.126 | 0.009 | 0 | retiFB | chr5-6952079-69525 |
| 0 | 4.86192915661614 | | 0.123 | 0.005 | 0 | retiFB | chr5-26257086-2625 |
| 0 | 3.62782561461521 | | 0.171 | 0.015 | 0 | retiFB | chr5-26587005-2658 |
| 0 | 3.5817712946684 | | 0.143 | 0.014 | 0 | retiFB | chr5-69542947-6954 |
| 0 | 3.67007393018393 | | 0.162 | 0.014 | 0 | retiFB | chr5-73525931-7352 |
| 0 | | 3.620591575 | 0.185 | 0.016 | 0 | retiFB | chr5-112136134-112 |
| 0 | 3.58933721245415 | | 0.135 | 0.013 | 0 | retiFB | chr5-113224092-113 |
| 0 | 4.40300667061413 | | 0.131 | 0.007 | 0 | retiFB | chr5-122808730-122 |

| | | | | | | | |
|---|---|---|---|---|---|---|---|
| 0 | 3.60376634288272 | | 0.198 | 0.017 | 0 | retiFB | chr5-143650221-143 |
| 0 | 4.03249314312645 | | 0.145 | 0.01 | 0 | retiFB | chr6-30798251-3079 |
| 0 | 3.85651923304986 | | 0.136 | 0.01 | 0 | retiFB | chr6-33084088-3308 |
| 0 | 3.75371414608508 | | 0.136 | 0.011 | 0 | retiFB | chr6-50209346-5020 |
| 0 | 3.7141575312093 | | 0.128 | 0.011 | 0 | retiFB | chr6-53262637-5326 |
| 0 | 4.58120158643129 | | 0.102 | 0.005 | 0 | retiFB | chr6-53654862-5365 |
| 0 | 3.91810348907562 | | 0.161 | 0.012 | 0 | retiFB | chr6-90184905-9018 |
| 0 | 3.74764176468135 | | 0.141 | 0.013 | 0 | retiFB | chr6-92215401-9221 |
| 0 | | 3.78820797 | 0.113 | 0.009 | 0 | retiFB | chr6-92243854-9224 |
| 0 | 3.65461790977707 | | 0.153 | 0.013 | 0 | retiFB | chr6-112705021-112 |
| 0 | 3.61207030703697 | | 0.164 | 0.014 | 0 | retiFB | chr6-112759694-112 |
| 0 | 3.88653698788177 | | 0.114 | 0.009 | 0 | retiFB | chr6-114012692-114 |
| 0 | 3.89595101107401 | | 0.115 | 0.008 | 0 | retiFB | chr6-115064838-115 |
| 0 | 3.85978958266293 | | 0.121 | 0.009 | 0 | retiFB | chr6-115691801-115 |
| 0 | 4.5363540236932 | | 0.108 | 0.005 | 0 | retiFB | chr7-45428641-4542 |
| 0 | 3.97298611753807 | | 0.102 | 0.007 | 0 | retiFB | chr7-55509581-5551 |
| 0 | 4.09430769734123 | | 0.101 | 0.006 | 0 | retiFB | chr7-82595685-8259 |
| 0 | 3.74038320690685 | | 0.122 | 0.01 | 0 | retiFB | chr8-28955399-2895 |
| 0 | 3.91802978114664 | | 0.131 | 0.01 | 0 | retiFB | chr8-33849417-3384 |
| 0 | 3.62957851916887 | | 0.143 | 0.012 | 0 | retiFB | chr8-76801148-7680 |
| 0 | 3.98473352782784 | | 0.104 | 0.007 | 0 | retiFB | chr8-80636743-8063 |
| 0 | 3.6362909837237 | | 0.128 | 0.012 | 0 | retiFB | chr8-92889331-9288 |
| 0 | 3.87165663705085 | | 0.109 | 0.008 | 0 | retiFB | chr8-116545349-116 |
| 0 | 4.07635038148077 | | 0.106 | 0.007 | 0 | retiFB | chr8-116761407-116 |
| 0 | 3.81257661848046 | | 0.139 | 0.011 | 0 | retiFB | chr9-11721945-1172 |
| 0 | 4.17899277892335 | | 0.103 | 0.006 | 0 | retiFB | chr9-16705290-1670 |
| 0 | 3.74555222298101 | | 0.123 | 0.01 | 0 | retiFB | chr9-46654568-4665 |
| 0 | 3.89621703550609 | | 0.113 | 0.008 | 0 | retiFB | chr9-62188494-6218 |
| 0 | 3.56997815715892 | | 0.17 | 0.017 | 0 | retiFB | chr9-74957530-7495 |
| 0 | 3.59911053325079 | | 0.12 | 0.01 | 0 | retiFB | chr9-77544604-7754 |
| 0 | 3.85393153938808 | | 0.11 | 0.008 | 0 | retiFB | chr9-99103020-9910 |
| 0 | 3.74227403984665 | | 0.113 | 0.009 | 0 | retiFB | chr9-107049502-107 |
| 1.15056440 | 3.6558280080992 | | 0.115 | 0.01 | 8.4309447 | retiFB | chr2-138594028-138 |
| 1.23344096 | 3.83578256365613 | | 0.11 | 0.009 | 9.0382360 | retiFB | chr3-59259811-5926 |
| 3.33209113 | 3.76531881550506 | | 0.105 | 0.008 | 2.4416430 | retiFB | chr2-53479171-5347 |
| 2.94550905 | 3.72015476951818 | | 0.127 | 0.012 | 2.1583688 | retiFB | chr17-27420228-274 |
| 5.58734835 | 3.89278568427658 | | 0.109 | 0.009 | 4.0942189 | retiFB | chr3-29459527-2946 |
| 2.17272531 | 3.61312752993378 | | 0.12 | 0.011 | 1.5920992 | retiFB | chr20-41915667-419 |
| 4.32123496 | 3.71711816742926 | | 0.111 | 0.009 | 3.1664540 | retiFB | chr7-50598155-5059 |
| 3.17537309 | 3.61597426995712 | | 0.116 | 0.01 | 2.3268054 | retiFB | chr1-189265711-189 |
| 4.02655711 | 3.65715667505927 | | 0.115 | 0.01 | 2.9505241 | retiFB | chr5-71975274-7197 |
| 6.33731336 | 3.73297010138735 | | 0.109 | 0.009 | 4.6437677 | retiFB | chr5-46315141-4631 |
| 2.49408215 | 3.68055587772136 | | 0.118 | 0.011 | 1.8275786 | retiFB | chr7-55958481-5595 |

| | | | | | | | |
|---|---|---|---|---|---|---|---|
| 2.63200573( | 3.64799849796662 | | 0.118 | 0.011 | 1.9286443 | retiFB | chr6-92228589-9222 |
| 7.15381227( | 3.69269297162988 | | 0.112 | 0.009 | 5.2420704 | retiFB | chr6-98130871-9813 |
| 1.25569221: | 3.78478742775394 | | 0.115 | 0.01 | 9.2012855 | retiFB | chr3-127024881-127 |
| 1.93347554( | 3.89983907251693 | | 0.103 | 0.008 | 1.4167851 | retiFB | chr10-75644546-756 |
| 9.61585763( | 3.86799826988969 | | 0.107 | 0.008 | 7.0461735 | retiFB | chr8-12732534-1273 |
| 4.16285723: | 3.95570424556447 | | 0.103 | 0.008 | 3.0504002 | retiFB | chr15-83218163-832 |
| 9.48879233( | 3.65345749607597 | | 0.113 | 0.01 | 6.9530644 | retiFB | chr2-49697754-4969 |
| 1.28789333: | 3.81534427436581 | | 0.11 | 0.009 | 9.4372444 | retiFB | chr2-34542706-3454 |
| 2.75545332 | 3.70354368642308 | | 0.111 | 0.009 | 2.0191025 | retiFB | chr17-31994718-319 |
| 3.79117511( | | 3.751019848 | 0.107 | 0.009 | 2.7780442 | retiFB | chr16-61350953-613 |
| 1.03418954 | 3.58307201662055 | | 0.113 | 0.01 | 7.5781893 | retiFB | chr16-51837206-518 |
| 2.35503765 | 3.6778036775091 | | 0.115 | 0.01 | 1.7256915 | retiFB | chr11-42720493-427 |
| 3.26780080( | 3.91632776971626 | | 0.109 | 0.009 | 2.3945333 | retiFB | chr17-37985814-379 |
| 4.36426158: | 3.69370424816159 | | 0.106 | 0.008 | 3.1979825 | retiFB | chr3-120763275-120 |
| 1.85028139: | 4.05732190664208 | | 0.101 | 0.008 | 1.3558232 | retiFB | chr8-59085408-5908 |
| 2.40901562( | 3.71322328415619 | | 0.112 | 0.01 | 1.7652447 | retiFB | chr2-226033784-226 |
| 2.59155160( | 3.62468588979236 | | 0.109 | 0.009 | 1.8990009 | retiFB | chr18-16816432-168 |
| 6.13306158 | 3.71911469142583 | | 0.109 | 0.009 | 4.4940990 | retiFB | chr13-41893015-418 |
| 2.65870214 | 3.68671427617146 | | 0.104 | 0.009 | 1.9482065 | retiFB | chr9-40433071-4043 |
| 4.31682068( | 3.82682719171454 | | 0.101 | 0.008 | 3.1632194 | retiFB | chr5-119797631-119 |
| 3.03106775( | 3.79147847182319 | | 0.102 | 0.008 | 2.2210633 | retiFB | chr14-40619369-406 |
| 4.42993102( | 3.64541182173788 | | 0.106 | 0.009 | 3.2461028 | retiFB | chr8-47902608-4790 |
| 1.37826997( | 3.58452389841684 | | 0.107 | 0.009 | 1.0099493 | retiFB | chr19-35979764-359 |
| 5.81937556( | 3.65583308563813 | | 0.102 | 0.008 | 4.2642405 | retiFB | chr2-34586971-3458 |
| 9.60792360 | 3.62461466344695 | | 0.106 | 0.009 | 7.0403597 | retiFB | chr10-93077093-930 |
| 1.99245422( | 3.68355654054866 | | 0.109 | 0.01 | 1.4600027 | retiFB | chr16-47796039-477 |
| 1.28185474( | 3.63057951623002 | | 0.113 | 0.011 | 9.3929957 | retiFB | chr4-59293674-5929 |
| 7.78111802( | 3.58164779365481 | | 0.107 | 0.01 | 5.7017387 | retiFB | chr10-97005759-970 |
| 7.65570954( | 3.7903924175375 | | 0.101 | 0.009 | 5.6098436 | retiFB | chr14-34229111-342 |
| 8.28485770( | 3.58089175989552 | | 0.101 | 0.009 | 6.0708620 | retiFB | chr9-57264790-5726 |
| 1.76122738( | 3.71778347082691 | | 0.1 | 0.009 | 1.2905675 | retiFB | chr16-42469363-424 |
| 7.19907420 | 3.61387243136446 | | 0.1 | 0.009 | 5.2752368 | retiFB | chr7-67036599-6703 |
| 1.18908491 | 3.64078309054399 | | 0.101 | 0.01 | 8.7132099 | retiFB | chr9-62481288-6248 |
| 2.33035018 | 3.57845830589672 | | 0.101 | 0.01 | 1.7076013 | retiFB | chr16-28707229-287 |
| 0 | 5.26078339872396 | | 0.106 | 0.003 | 0 | sbasalKC | chr1-9635389-96358 |
| 0 | 4.42923797798847 | | 0.112 | 0.005 | 0 | sbasalKC | chr1-17320830-1732 |
| 0 | 4.47585681626778 | | 0.161 | 0.008 | 0 | sbasalKC | chr1-32603033-3260 |
| 0 | 4.44231082355267 | | 0.123 | 0.006 | 0 | sbasalKC | chr1-66053764-6605 |
| 0 | 4.36065697380238 | | 0.129 | 0.007 | 0 | sbasalKC | chr1-94274079-9427 |
| 0 | 4.61856537477135 | | 0.205 | 0.01 | 0 | sbasalKC | chr1-169570230-169 |
| 0 | 4.6697977826758 | | 0.112 | 0.005 | 0 | sbasalKC | chr1-204405999-204 |
| 0 | 4.48490859757056 | | 0.208 | 0.011 | 0 | sbasalKC | chr10-84510453-845 |
| 0 | 4.65442698101552 | | 0.147 | 0.007 | 0 | sbasalKC | chr10-87678111-876 |

| | | | | | |
|---|---|---|---|---|---|
| 0 4.71836355559061 | | 0.157 | 0.007 | 0 sbasalKC | chr10-88743457-887 |
| 0 4.70378519603816 | | 0.104 | 0.004 | 0 sbasalKC | chr10-95606086-956 |
| 0 4.89091112171847 | | 0.111 | 0.005 | 0 sbasalKC | chr11-25423866-254 |
| 0 4.98433214585734 | | 0.12 | 0.004 | 0 sbasalKC | chr11-42792705-427 |
| 0 4.67474954607063 | | 0.159 | 0.007 | 0 sbasalKC | chr11-45127349-451 |
| 0 5.09391373192584 | | 0.1 | 0.003 | 0 sbasalKC | chr11-45144381-451 |
| 0 4.98501215232293 | | 0.129 | 0.004 | 0 sbasalKC | chr11-64344933-643 |
| 0 4.84384543122057 | | 0.1 | 0.004 | 0 sbasalKC | chr11-64353283-643 |
| 0 4.58743299235712 | | 0.104 | 0.005 | 0 sbasalKC | chr11-64400448-644 |
| 0 5.00831339343089 | | 0.186 | 0.007 | 0 sbasalKC | chr11-64514334-645 |
| 0 4.38068673649883 | | 0.111 | 0.006 | 0 sbasalKC | chr11-64617848-646 |
| 0 4.55986603271798 | | 0.135 | 0.007 | 0 sbasalKC | chr11-75351374-753 |
| 0 4.65881024836278 | | 0.12 | 0.006 | 0 sbasalKC | chr12-6876410-6876 |
| 0 4.58743299235712 | | 0.104 | 0.005 | 0 sbasalKC | chr12-16869588-168 |
| 0 4.31499211143507 | | 0.137 | 0.007 | 0 sbasalKC | chr12-16870330-168 |
| 0 4.41024318342965 | | 0.161 | 0.01 | 0 sbasalKC | chr12-45837053-458 |
| 0 4.99995630115833 | | 0.144 | 0.005 | 0 sbasalKC | chr13-23475717-234 |
| 0 5.01445394226381 | | 0.145 | 0.005 | 0 sbasalKC | chr14-1100805-1101 |
| 0 4.39344376545825 | | 0.213 | 0.011 | 0 sbasalKC | chr14-21285666-212 |
| 0 4.41121435372783 | | 0.144 | 0.007 | 0 sbasalKC | chr14-21474636-214 |
| 0 4.67993845150775 | | 0.154 | 0.007 | 0 sbasalKC | chr14-44492254-444 |
| 0 4.36453337354089 | | 0.152 | 0.007 | 0 sbasalKC | chr14-82254843-822 |
| 0 4.48243982462964 | | 0.188 | 0.01 | 0 sbasalKC | chr14-82420107-824 |
| 0 5.26222423571329 | | 0.133 | 0.004 | 0 sbasalKC | chr14-92395266-923 |
| 0 4.68143409683724 | | 0.155 | 0.007 | 0 sbasalKC | chr15-1577166-1577 |
| 0 4.49119034429036 | | 0.115 | 0.005 | 0 sbasalKC | chr15-3754772-3755 |
| 0 4.34149895594734 | | 0.181 | 0.01 | 0 sbasalKC | chr15-35191274-351 |
| 0 4.45090162494315 | | 0.154 | 0.008 | 0 sbasalKC | chr15-76237934-762 |
| 0 4.4483417390941 | | 0.108 | 0.005 | 0 sbasalKC | chr15-76287836-762 |
| 0 4.42628585800003 | | 0.157 | 0.008 | 0 sbasalKC | chr15-95560685-955 |
| 0 4.55242882008258 | | 0.126 | 0.006 | 0 sbasalKC | chr16-20850225-208 |
| 0 | 4.68428892 | 0.108 | 0.005 | 0 sbasalKC | chr16-28439551-284 |
| 0 4.31031239299311 | | 0.128 | 0.007 | 0 sbasalKC | chr16-43672772-436 |
| 0 4.55209381281517 | | 0.106 | 0.005 | 0 sbasalKC | chr16-56780709-567 |
| 0 4.32416142786849 | | 0.174 | 0.01 | 0 sbasalKC | chr16-77655157-776 |
| 0 4.66341187774182 | | 0.115 | 0.005 | 0 sbasalKC | chr17-22772780-227 |
| 0 4.50541941343365 | | 0.138 | 0.007 | 0 sbasalKC | chr17-23305437-233 |
| 0 4.60244146937625 | | 0.101 | 0.004 | 0 sbasalKC | chr17-24446702-244 |
| 0 4.88691655699242 | | 0.118 | 0.005 | 0 sbasalKC | chr17-25694335-256 |
| 0 4.92962497059841 | | 0.175 | 0.007 | 0 sbasalKC | chr17-29681202-296 |
| 0 4.67867371628342 | | 0.112 | 0.005 | 0 sbasalKC | chr17-68991510-689 |
| 0 4.3915732333392 | | 0.133 | 0.007 | 0 sbasalKC | chr17-69523317-695 |
| 0 4.65741619537307 | | 0.125 | 0.006 | 0 sbasalKC | chr17-69737813-697 |

| | | | | | | |
|---|---|---|---|---|---|---|
| 0 | 4.61624158591619 | 0.111 | 0.005 | 0 | sbasalKC | chr17-75597369-755[illegible] |
| 0 | 5.01860269443553 | 0.15 | 0.006 | 0 | sbasalKC | chr17-80495413-804[illegible] |
| 0 | 5.31608822877253 | 0.186 | 0.006 | 0 | sbasalKC | chr17-80636845-806[illegible] |
| 0 | 4.37570033280787 | 0.117 | 0.006 | 0 | sbasalKC | chr18-25703010-257[illegible] |
| 0 | 4.52080069971066 | 0.123 | 0.007 | 0 | sbasalKC | chr18-32118236-321[illegible] |
| 0 | 4.37493982559802 | 0.171 | 0.009 | 0 | sbasalKC | chr18-36264199-362[illegible] |
| 0 | 4.35280843347905 | 0.213 | 0.012 | 0 | sbasalKC | chr18-75870373-758[illegible] |
| 0 | 4.68004214441542 | 0.146 | 0.007 | 0 | sbasalKC | chr19-14611820-146[illegible] |
| 0 | 4.30075398775821 | 0.198 | 0.013 | 0 | sbasalKC | chr19-14626073-146[illegible] |
| 0 | 4.41889644359089 | 0.134 | 0.006 | 0 | sbasalKC | chr19-15234214-152[illegible] |
| 0 | 4.34402586418836 | 0.13 | 0.007 | 0 | sbasalKC | chr19-41988186-419[illegible] |
| 0 | 5.04996288494669 | 0.149 | 0.005 | 0 | sbasalKC | chr2-10874206-1087[illegible] |
| 0 | 4.31554148848174 | 0.197 | 0.012 | 0 | sbasalKC | chr2-30750044-3075[illegible] |
| 0 | 4.52978021297152 | 0.134 | 0.007 | 0 | sbasalKC | chr2-33917722-3391[illegible] |
| 0 | 4.56980115142973 | 0.12 | 0.005 | 0 | sbasalKC | chr2-45106772-4510[illegible] |
| 0 | 4.85987248248435 | 0.107 | 0.004 | 0 | sbasalKC | chr2-57458621-5745[illegible] |
| 0 | 4.57900719458132 | 0.122 | 0.006 | 0 | sbasalKC | chr2-118947780-118[illegible] |
| 0 | 4.71700431935154 | 0.157 | 0.007 | 0 | sbasalKC | chr2-121712193-121[illegible] |
| 0 | 4.6895799750222 | 0.104 | 0.004 | 0 | sbasalKC | chr2-136613588-136[illegible] |
| 0 | 4.74810294284593 | 0.163 | 0.007 | 0 | sbasalKC | chr2-195950399-195[illegible] |
| 0 | 4.51864797818659 | 0.189 | 0.009 | 0 | sbasalKC | chr2-205121224-205[illegible] |
| 0 | 4.38069974157816 | 0.136 | 0.007 | 0 | sbasalKC | chr2-205460341-205[illegible] |
| 0 | 4.41294975866175 | 0.14 | 0.007 | 0 | sbasalKC | chr2-218239239-218[illegible] |
| 0 | 4.51703108664677 | 0.131 | 0.006 | 0 | sbasalKC | chr2-218242047-218[illegible] |
| 0 | 4.63133932464104 | 0.104 | 0.005 | 0 | sbasalKC | chr2-219609411-219[illegible] |
| 0 | 4.42830974245627 | 0.114 | 0.006 | 0 | sbasalKC | chr20-4448970-4449[illegible] |
| 0 | 4.37583689358131 | 0.206 | 0.012 | 0 | sbasalKC | chr20-10943999-109[illegible] |
| 0 | 4.64012388648636 | 0.127 | 0.006 | 0 | sbasalKC | chr3-32414045-3241[illegible] |
| 0 | 4.77649794493354 | 0.124 | 0.005 | 0 | sbasalKC | chr3-34158289-3415[illegible] |
| 0 | 4.77766265268036 | 0.108 | 0.004 | 0 | sbasalKC | chr3-45847878-4584[illegible] |
| 0 | 5.06185685975663 | 0.167 | 0.006 | 0 | sbasalKC | chr3-47351051-4735[illegible] |
| 0 | 4.99935026366959 | 0.13 | 0.005 | 0 | sbasalKC | chr3-102228098-102[illegible] |
| 0 | 4.86261726598178 | 0.161 | 0.006 | 0 | sbasalKC | chr3-102422747-102[illegible] |
| 0 | 4.96471204201017 | 0.138 | 0.005 | 0 | sbasalKC | chr3-109256105-109[illegible] |
| 0 | 4.71702673184229 | 0.109 | 0.005 | 0 | sbasalKC | chr3-126847774-126[illegible] |
| 0 | 4.36763000538874 | 0.121 | 0.006 | 0 | sbasalKC | chr3-135910347-135[illegible] |
| 0 | 4.48222391651894 | 0.151 | 0.007 | 0 | sbasalKC | chr3-162740449-162[illegible] |
| 0 | 4.34203898561355 | 0.116 | 0.006 | 0 | sbasalKC | chr4-46040821-4604[illegible] |
| 0 | 4.39042136449465 | 0.218 | 0.013 | 0 | sbasalKC | chr4-46786063-4678[illegible] |
| 0 | 4.44275649345705 | 0.113 | 0.005 | 0 | sbasalKC | chr4-51253552-5125[illegible] |
| 0 | 4.57292053003271 | 0.104 | 0.005 | 0 | sbasalKC | chr4-70366982-7036[illegible] |
| 0 | 4.77906259013945 | 0.168 | 0.007 | 0 | sbasalKC | chr4-104552227-104[illegible] |
| 0 | 4.73600941454256 | 0.123 | 0.005 | 0 | sbasalKC | chr4-153844420-153[illegible] |

| | | | | | | |
|---|---|---|---|---|---|---|
| 0 | 4.44346334898871 | 0.13 | 0.007 | 0 | sbasalKC | chr4-158224987-158 |
| 0 | 4.72802052071257 | 0.108 | 0.004 | 0 | sbasalKC | chr5-13728363-1372 |
| 0 | 4.37365478675187 | 0.118 | 0.006 | 0 | sbasalKC | chr5-28243283-2824 |
| 0 | 4.4279595899272 | 0.114 | 0.005 | 0 | sbasalKC | chr5-28630903-2863 |
| 0 | 4.42155401745017 | 0.112 | 0.006 | 0 | sbasalKC | chr5-39306729-3930 |
| 0 | 5.15728327115593 | 0.113 | 0.003 | 0 | sbasalKC | chr5-48537696-4853 |
| 0 | 4.53843175030364 | 0.139 | 0.008 | 0 | sbasalKC | chr5-98182752-9818 |
| 0 | 4.51626933930833 | 0.114 | 0.005 | 0 | sbasalKC | chr5-136418267-136 |
| 0 | 4.38242973298486 | 0.207 | 0.011 | 0 | sbasalKC | chr5-147030661-147 |
| 0 | 4.39862635239643 | 0.125 | 0.007 | 0 | sbasalKC | chr5-147643060-147 |
| 0 | 4.70109495478177 | 0.154 | 0.007 | 0 | sbasalKC | chr5-156188974-156 |
| 0 | 4.52936660745685 | 0.251 | 0.014 | 0 | sbasalKC | chr5-162239745-162 |
| 0 | 4.72455709594657 | 0.106 | 0.005 | 0 | sbasalKC | chr6-53028960-5302 |
| 0 | 4.39727413259005 | 0.116 | 0.006 | 0 | sbasalKC | chr6-53220941-5322 |
| 0 | 4.91868991674228 | 0.117 | 0.004 | 0 | sbasalKC | chr6-60965982-6096 |
| 0 | 4.95392800459043 | 0.142 | 0.005 | 0 | sbasalKC | chr6-109767882-109 |
| 0 | 4.60493981164226 | 0.12 | 0.005 | 0 | sbasalKC | chr7-22021942-2202 |
| 0 | 4.69536122526295 | 0.148 | 0.007 | 0 | sbasalKC | chr7-30253102-3025 |
| 0 | 4.51151024232934 | 0.182 | 0.009 | 0 | sbasalKC | chr7-68438091-6843 |
| 0 | 4.32545607709604 | 0.123 | 0.006 | 0 | sbasalKC | chr7-70056786-7005 |
| 0 | 4.85971456538212 | 0.105 | 0.004 | 0 | sbasalKC | chr7-82176862-8217 |
| 0 | 4.43104811488676 | 0.118 | 0.006 | 0 | sbasalKC | chr7-125453730-125 |
| 0 | 4.83092631402771 | 0.118 | 0.004 | 0 | sbasalKC | chr7-130856031-130 |
| 0 | 4.4371594850406 | 0.153 | 0.008 | 0 | sbasalKC | chr8-4867165-48676 |
| 0 | 4.72079970168095 | 0.149 | 0.007 | 0 | sbasalKC | chr8-27916993-2791 |
| 0 | 4.49718550385511 | 0.119 | 0.006 | 0 | sbasalKC | chr8-65840766-6584 |
| 0 | 4.29742644139762 | 0.176 | 0.011 | 0 | sbasalKC | chr8-83604492-8360 |
| 0 | 5.01406331697795 | 0.14 | 0.005 | 0 | sbasalKC | chr8-122385134-122 |
| 0 | 4.88920850576092 | 0.108 | 0.004 | 0 | sbasalKC | chr9-4890557-48910 |
| 0 | 4.3014387103106 | 0.186 | 0.01 | 0 | sbasalKC | chr9-66542941-6654 |
| 0 | 4.59690767433902 | 0.114 | 0.005 | 0 | sbasalKC | chr9-71116556-7111 |
| 0 | 5.17598411885064 | 0.244 | 0.008 | 0 | sbasalKC | chr9-79937551-7993 |
| 0 | 4.37411012186888 | 0.169 | 0.009 | 0 | sbasalKC | chr9-89183563-8918 |
| 0 | 4.94308859326943 | 0.105 | 0.004 | 0 | sbasalKC | chr9-98001565-9800 |
| 0 | 4.4302841853117 | 0.141 | 0.007 | 0 | sbasalKC | chrX-94317165-9431 |
| 7.57913850 | 4.46329276512714 | 0.112 | 0.006 | 5.5537350 | sbasalKC | chr5-71873774-7187 |
| 1.10511267 | 4.52584775170085 | 0.102 | 0.005 | 8.0978899 | sbasalKC | chr1-56098602-5609 |
| 1.71529036 | 4.31872021917618 | 0.111 | 0.006 | 1.2569064 | sbasalKC | chr13-23495816-234 |
| 6.31369113 | 4.67973377058551 | 0.104 | 0.005 | 4.6264581 | sbasalKC | chr3-101558243-101 |
| 4.10553531 | 4.71171248681715 | 0.1 | 0.005 | 3.0083966 | sbasalKC | chr5-124721529-124 |
| 4.82019921 | 4.51083837929859 | 0.104 | 0.005 | 3.5320780 | sbasalKC | chr10-42530944-425 |
| 1.39721488 | 4.50451463401977 | 0.103 | 0.005 | 1.0238315 | sbasalKC | chr2-184746614-184 |
| 4.85925755 | 4.65871156255536 | 0.104 | 0.005 | 3.5606987 | sbasalKC | chr1-33704214-3370 |

| | | | | | | |
|---|---|---|---|---|---|---|
| 2.30459372 | 4.29769787848789 | 0.121 | 0.007 | 1.6887279 | sbasalKC | chr2-197256614-197 |
| 6.84111410 | 4.29474582482424 | 0.113 | 0.006 | 5.0129358 | sbasalKC | chr16-594364-59486 |
| 4.00302915 | 4.39900839822815 | 0.113 | 0.006 | 2.9332836 | sbasalKC | chr1-56605674-5660 |
| 7.24461617 | 4.41806716836199 | 0.104 | 0.005 | 5.3086084 | sbasalKC | chr15-96948937-969 |
| 1.05427856 | 4.2977431607184 | 0.112 | 0.006 | 7.7253948 | sbasalKC | chr7-31668367-3166 |
| 6.51001980 | 4.50749949432416 | 0.104 | 0.005 | 4.7703211 | sbasalKC | chr17-69367471-693 |
| 1.22976002 | 4.47792362511971 | 0.104 | 0.005 | 9.0112633 | sbasalKC | chr5-55488965-5548 |
| 1.11625386 | 4.49997818548627 | 0.105 | 0.006 | 8.1795288 | sbasalKC | chr1-218726874-218 |
| 1.70865422 | 4.33722834167922 | 0.105 | 0.006 | 1.2520437 | sbasalKC | chr8-63449303-6344 |
| 7.05371182 | 4.37067197901069 | 0.117 | 0.007 | 5.1687201 | sbasalKC | chr1-126461520-126 |
| 1.47701325 | 4.45409359087684 | 0.1 | 0.005 | 1.0823050 | sbasalKC | chr14-21638334-216 |
| 2.71892917 | 4.32262851855852 | 0.106 | 0.006 | 1.9923388 | sbasalKC | chr1-198927185-198 |

32735
49636
56170
870253
102610
752889
60217
307754
6549
3342
718589
0208
318139
725
35610
9589
945102
13061
27169
9775
778341
6990
6359
04533
1955
5049
1953
9512
7975
20069
640
6918
4729
0320
288515
718354
4104
457914
8019
9092
85

377199
7441
744
256423
342356
075129
803342
174307
2986
19925
194319
7028
54008
62877
429715
04682
75757
972573
7315
29427
671915
4197
78770
07579
36301
1912
11787
84257
8411
54850
14858
94584
31320
088608
165962
8405
46022
7131
195727
2608
0371
80957
17549

820942
102
00806
31800
886181
2859
4492
543964
561692
51892
65670
173260
053744
58472
5965
5214
991198
629620
22766
16753
58502
03613
7212
9544
217832
174
819009
640636
12
25187
39446
10189
66312
464359
412608
14
177
91535
95944
33587
32809
1432
902384

101020
70
82039
9841
8932
84416
1895
394664
129283
3216
5976
65924
40819
39654
40887
4072
4143
054670
17728
943588
088478
217223
63715
439277
095265
69950
7728
7908
550999
63988
0001784
952474
2393
2324
0014
183923
69658
1065
46922
809476
323114
280
74718

4801
3378
36399
7350
0821
91409
194
613883
96098
9376
7222
38891
574212
64916
899972
3570
1585
25784
34495
5029725
9804
71706
431433
56951
92470
5892
8734
734358
30595
3923
457337
041520
2050
5594
3939
55521
0642
542323
5207
913
66003
85432
192047

67206
44169
940506
12345
68389
4361
10747
8580
77301
93608
96478
389024
5652
049206
6194
42531
188913
67831
40374
6445
03991
79210
58986
52572
1127
9517
8608
18020
6140
346525
1099
787610
9911
662726
6488
6628
143
8964
46424
0074
66890
61703
70499

619985
96434
97495
99838
7501
763
74167
873601
797338
97
1084
0009
6299
3374
4061
41251
351
0228
23699
897970
78440
6757
53107
92420
69865
25241
0935
1691
8860
26054
018249
709913
54190
933363
1478
[illegible]8103
64756
03925
97401
991848
871732
557
99855

3912831
15
6733
8366
8453
6761
7604
8299
1412
0083
3055
980177
684149
030773
753511
834946
933266
437782
15631
02803
20843
40965
18564
21744
23329
71892
61988
25066
02860
40409
439
986
81304
80310
49411
02920
31130
30209
15845
3077267
6468124
11858
06112

684
82677
95066
66519
788
80906
61682
81063
16649
25399
46027
14054
44849
50833
55365
21764
29444
53003
463
46449
26374
8517
9925
5950
9078
3193
9630
0183
6678
953834
961907
965671
224391
167028
524057
623805
682189
904471
221623
875839
764838
080523
339916

385172
58940
03604
62626
52587
26835
9407
7630
4292
0815
4455
8807
0600
582139
709531
242385
1444
0057
0717
1702
0479
633085
537166
490937
717347
728172
949973
331879
876946
864575
6124
1124
9519
2413
8586
8349
0824
4834
3527
2511
8734
2872
1437

925485
983930
6806
1532
1726
568929
1921
8545
6711
0155
6598
6516
7064
727847
8885
4520
2857
7220
0839
192548
511070
512815
0306
7872
3443
334420
344474
268109
656795
85755
80209
15964
60164
16613
70159
78524
10731
68334
50164
39071
66323
61103
90444

49501
75690
73597
82033
43229
42802
44596
51918
01724
297794
053327
86288
8681
662407
040040
843870
4381
724173
5827
2130
0772
251068
8180
9055
5434
6411
6925
197130
861018
9502
4836
3090
8681
4063
1107
8367
2980
4405
9705
6608
212926
8269
971069

48058
74632
3934
72111
66504
8289
45713
88486
5289
6624
038234
6273
951782
04437
12012
802506
57575
6517
9741
06627
71926
37910
3800
99823
70017
61642
87860
2422
26264
2943
72192
6302
81379
77358
3289
08445
3704
4419573
4344
2388
48262
160919
356884

706905
50568
863605
36831
189521
923276
39190
3950
8991
6285
22587
550484

0414
59029
24719
3384
31496
6758
59
541229
440573
927460
94800
8800
5971
92192
52482
139648
35908
349
92586
12899
27345
26323
3797
42391
271
22002
811124
2321
2171
3192

1163
019700
498493
862021
875161
897461
26120
08518
49572
30091
21179
05965
92101
882
71658
13739
42922
09941
980
85610
20747
69712
47918
22640
37367
38121
11085
62825
87124
54817
79406
1034
716483
855258
105751
262598
5200
4781
759948
682291
818973
120486
492991

26
1713
8008
8514
911604
3583
007332
216989
9104
1610
6053
4639
0105
5762
3947861
4063
790137
5849
04988
07302
77845
14797
050923
911276
23122
37070
0519
415480
17105
4737
9873
848746
0297
007819
2278
622149
70904
70
55322
97292
39285
9589
321550

631
02262
1969
05956
870
3719
985734
4960
224843
25475
70791
213876
5994
87574
65096
1533
1644
82409
50697
1536
598774
76781
5584
098329
64061
413398
40630
06133
80603
4659210
5901
70715
26
223872
41094
5202
100
50610
93872
31973
41
675618
955619

9671
5909
9460
32420
20455
80111
302640
0256
7317
380808
79512
670247
3351
021260
72498
76019
5958
95053
04347
4173
554874
9851
8061
986382
7054
913765
31360
87933
24072
695908
429416
469670
219393
1946
66820
1487286
242419
1973
45598
930094

188128
30509

66829
81552
848006
0681
99511
063929
1641
6323
091471
785
231
38142
22082
4376
65427
3690
8524
124876
74554
5078
492898
96898
495682
3151601
97450
983949
17461
0239
470497
393
711621
21551
25974
3847
29023
0296
253712
66979
2567
2742146
16337
237702
21107

984173
6225
245738
061226
91847
693982
23167
46145
50427
3016
88967
4228
891
48257
94473
4781
620834
6155
734936
850668
219570
356291
76608
21190
054291
46057
089598
922650
5611
067061
27444
332306
37
12157
26050
25391
[illegible]005984
85610
714636
021849
728795
7033
09941

6802
02398
81624
9545
98409
402120
67149
96566
67800
0564
78217
3937
12163
4576
73614
87846
0284
01468
435
56847
882
633372
6742
5835
330816
9001
454295
6947
4639
898
10328
30
73
494751
30350
74
09315
058730
6815
21843
7495
8843
4654

6322
494827
369362
801122
087682
397479
619805
694720
124642
155219
580275
88122
58196
28965
23885
36437
04691
30050
36649
29561
56228
69363
68128
90318
91914
07358
17726
34771
89370
30858
75452
04216
06605
37641
97255
51272
27366
71719
90935
78405
80466
94872
964

567
663
55234
47033
91330
82413
94545
32247
25429
54521
84597
17639
25650
57566
17715
07155
25403
70251
34093
9133
6130
9638
3865
8317
7821
2502
523505
658010
642656
731
620
41985
25660
0657
4485
1588
9273
142501
7357
4154
434706
384797
168223

6556
8680
2119
3561
5572
2863
4946
5317
6698
7100
4309
688188
912313
542490
543270
3484
9439
8629
5468
4379
5111
7227
1995
6415
7745
6964
534733
371498
4062
0022
0562
7984
7428
0852
1435
0221
7942
4391
966148
2072722
8809
4379
60608

80062
61273
32656
8043
36215
5179
544285
908848
0182
9548
255198
8433
752584
164802
1223
11171
3327
47736
98
50855
2437
7845
8479
1986
0888
3431
994395
823157
784566
992185
246779
006741
056445
893080
748194
585903
896218
313
71850
70257
70324
417
73114

83437
25910
36263
94835
0
87335
42001
10646
06277
38507
72060
85576
21223
17654
46542
16725
54215
80050
14674
198
66652
76255
77312
43799
10825
96391
64819
21059
72823
432
24971
38455
69503
14315
75971
17826
18343
27578
646
329
09051
70180
84123

83397
85781
91859
85254
70943
3420
1562
7585
0449
7751
4786
100348
249874
496507
238746
490008
567172
595993
80720
38637
16931
8427
7486
7258
9624
1763
7663
4804
4905
736985
642924
933980
951895
056583
7919
9292
4455
3064
9700
4158
6517
253557
1451

7951
210016
965408
6190
6859
5228
8268
1305
1224
464571
334312
81
5783
9946
4533
7207
1902
1084
109274
296745
988199
6557
5924
306267
310501
0595
4185
3425
5941
9878
3541
9099
2889
3374
6614
1925
3618
030465
422789
974927
986653
11
1220

8642
6621
8348
0663
093669
967072
961509
772428
912410
472389
972252
433951
199839
820804
55464
57849
13156
28280
04505
70259
21399
99790
0150988
0264689
86923
30178
95463
17984
35974
27282
92867
54208
02023
19004
09745
01794
47783
49338
61179
74645
56645
50024
74178

75905
23610
22764
40976
00288
16505
903
98758
10958
49531
98157
38439
50164
32627
14485
27279
29208
32441
33898
64039
21217
28967
54397
22027
69214
70918
63027
53668
47079
97877
56265
2026
7473
7978
3041
0990
2714
190982
822928
476190
391513
221257
327148

4251
0630
897428
727150
751958
310097
011589
664896
700330
710483
905209
510067
4754
1362
030444
524356
828836
867303
883343
072526
265637
358386
526710
3983
8433
8198
4914
271695
131237
126595
4632
5880
0555
5064
4120
4637
397205
835749
472023
477100
1912
3071
8158

2596
2804
787383
97
1347
6220
7203
9126
4238
682375
672857
10
78
2824
3447
9190
2624
9092
6251
48
6520
2070
0561
4441
5166
2863
2218
4491
4996
8749
784739
849843
664866
207437
01030
19965
43443
73132
4756694
53585
98457
48391
29642

30773
33133
33947
16610
46144
08593
65701
31037
81500
73693
43200
53708
20324
17763
52893
29262
89387
64614
67105
90340
022
631
41897
582
463
787
70531
17064
47773
00651
90357
91377
93275
70149
37484
77423
02871
42055
24574
60843
5825
2461
693158

476246
312236
713340
939975
650507
706431
07290
02798
89324
5640
4912
4883
208137
962274
445213
8129
5497
4240
1474
6785
9456
2490
273203
480818
061267
702584
701751
912797
562674
564034
042928
056114
096909
154364
307328
015412
064115
250115
504435
013415
80
08
39

4694
002622
130699
903402
421300
5061
8942
5599
092443
320260
37
9375
7243
3530
0594
4688
3672
2612
8643
2730
008647
761777
1672
0760
5900
1894
540795
619857
072480
436075
33
37
8973
8202
8160
0413
0904
1303
1301
6239
3144
2720
298951

132678
051102
051681
537383
531647
922595
537641
770307
770945
288333
108513
60865
33140
68850
46798
70686
43702
00870
2065232
3656656
3657274
11042
87835
03552
04281
423
07690
62661
45656
83082
15539
02260
18630
20804
66482
78397
5074269
84713
73031
26173
08542
51332
932

79927
13749
38268
57968
52200
59559
25142
88377
01683
04811
10858
14323
00506
10088
78758
83790
99731
43599
22461
67397
98896
01572
55574
26811
08144
58142
02515
24677
80157
48187
53157
00625
39687
63407
9120
1840
1879
020051
691104
758708
961805
968507
265438

497084
044282
48326
35175
01034
1645
3222
4325
6971
3690
6255
9206
360467
046978
896033
191796
214088
847066
9731
1659
050433
694945
695491
0103
1308
192902
113599
638309
470155
488363
904268
227710
850473
8047
5489
804503
387553
659580
7120
4812
7972
1722
3402

988100
941974
644084
93
9946
4511
9021
386185
776165
064691
56
1535
7266
6602
4396
5243
1462
7477050
41
941841
503621
069302
997753
907975
223845
256555
401176
417746
245852
892488
59976
01089
90422
86076
334
56301
95599
789
016
79826
47583
89974
18771

50446
38021
636
33127
27706
71165
68012
48422
84684
4103190
40725
71056
01492
61083
76014
20858
38012
57869
05531
16743
24991
87364
81805
43155
99132
06351
45689
18911
80948
763
83708
52459
11852
71815
48039
62261
120
72328
22158
84
9070
6659
7104

3998
532760
095166
589003
583653
493052
283501
710275
525839
67797
76241
8733
2391
0797
8921
0020
2718
9527
6397
5500
2002
7784
4512
1149
0803
2981
0395
077518
826551
863009
974929
700720
722206
4223
5163
2953
4944
8623
8519
6843
331637
939361
317423

806346
305384
2170
1059
0097
5297
210721
126426
946204
1942
6501
1827
192906
291918
909121
8817
6421
9755
3459
2813
2811
820216
286056
34
3850
9769
0908
6607
996772
13
8414
4704
8896
8350
22786
6438
0192538
406296
9138865
839215
68902
85417
1064

886
1633
434
743421
900
21774
78433
096136
7514
08003
3587
427030
99905
96134
2566
136221
51
84866
379385
1321
915474
66305
80721
6197
66950
60646
35541
0502
44193
012
011379
241717
5694
772
23649
478389
94912
26633
968746
3433
6143
317
5921503

640567
29233
130
2509
403062
207727
3422
096928
58134
0918
851000
16535
3269
29814
4485
99610
366189
21774
95608
803679
84074
5141
987993
8052
497959
92984
77324
25
657942
5934
90482
831268
626621
2934
197682
86609
9899
956695
266678
13907
64925
65123
41013

7839
089017
705965
51862
61356
22238
9132
513530
28370
69777
8423

588717
043172
899176
233167
6954
500874
707820
9940
555170
734647
79311
801367
7936
2061
69
515787
0954
30891
56491
97311
1187
441628
1046
528061
7170
45321
1966
038940
2062
931509
20815

24478
5658
964224
9032
96679
1379
280217
0486
28537
9807
96245
35049
45047
873
94268
844
533668
3643
5628
9775
4520
9746
8626
0448
1120
6002
5501
7861
5988
591286
607716
402843
267749
231211
850546
857283
001537
541
718
70282
77000
85367
30164

06223
36618
07894
92049
1148495
80493
948
702
757
768
855
330
83351
87470
42943
37264
18658
32736
90541
60548
35008
00566
97081
79449
08865
30579
70850

7
240
116
51848
23781
94331
75749
177
46283
65101
51491
22043
57450
34535
38387

41627
22334
72268
53100
11572
95136
51538
06655
08394
66141
50696
34380
35190
620941
765888
019343
311300
699
326
00380
38405
54358
59358
67806
8846
8016
4470
9563
6210
6306
761725
402332
874461
278142
551828
16
3951
7324
5383
697685
843983
197747
821002

927210
324969
189277
749822
248855
7135
6736
781320
406134
269784
187653
048789
4693
9886
7214
0274
3457
285996
3829
9445
6592
3772
518768
372444
879722
3656
4463
3468
280859
396446
01
1879
2454
4404
8081
4139
[illegible]5698
[illegible]179150
1064
098003
78433
85417
839215

86609
80721
273641
5447123
098611
35049
22390
5314
01582
21549
2509
3281033
9105
66305
4772
33
096928
0502
06464
2481
0422
81522
28694
8067
220
99905
83775
35541
5435
575124
99145
009
85670
21446
181
26011
26320
743421
74040
41650
33644
090167
87339

26633
2566
198462
77357
2508678
022429
38209
10
3419
42468
41178
001609
85724
703111
052343
44623
166871
482658
74963
376355
5559813
601008
6650
170170
82
577866
72289
832665
618226
0918
3844
86430
05198
50316
3269
254003
1928
592977
626621
1321
734579
2533382
0486

331170
82355
71
5376
69777
257094
16179
374851
3568
99035
545566
196768
37360
7799
86890
73005
10
29233
6425
16
992246
733615
5848
368716
25170
1535
3058
4473344
972678
68693
26526
333186
82794
3584
730364
04126
50
11283
375
70650
342286
66433
850098

17644
63026
4122
528
65
378387
9246
51
53921
95760
30629
252
21697
821
569881
2309451
839215
80721
831998
78433
50
70403
1064
184
21549
831268
26041
35541
69777
99905
098003
08312
00764
5694
75102
657942
096928
78528
2701542
56968
86973
8188
170

77071
97894
402820
9317
13846
49048
49293
88425
861800
15276
3783

2263
7032
058610
60898
26320
481125
396332
6000
1122
5059
046
333186
765
9105
7344
88453
87339
23279
72839
5934
29814
703677
494361
76841
39
4861
394648
2380
33621
75668
3310

379385
80970
07411
2677
038940
0445
3077
836446
76755
272757
614
499541
808
99035
05113
36571
279005
66805
720006
82735
225433
01666
22670
87725
0387
26633
4226
607806
753329
3058
53188
259
4285
737286
2167
290275
5911
772
244563
2352
33144
607114
53430

19
3284
09133
7091
1804
81447
32147
76148
87808
1535
777355
0784
571728
610784
805129
600001
770101
3297
86784
76179
199511
27438
06075
95574
84788
56300
6953
6716
80407
4596
402
254003
9315
478928
83068
369792
694958
7686
3675
7383
3694
518254
675465

562411
755725
925042
82915
68035
08261
39303
46776
39938
42322
33027
28280
89516
69730
34420
99971
31149
36116
08961
18856
89357
10448
49792
2514868
47566
45359
40277
99771
179
76743
94419
36008
65040
56830
09378
57092
41890
63269
80700
84962
789
14635
33750

59
69
0890
5370
9217
8050
8489
241329
889236
820401
272354
515761
698194
78888
17677
5621
8005
8795
3670
739009
816751
7302
9705
3729
6698
493400
681020
591739
771185
735910
710185
38
3572
8129
8125
9157
4880
645162
1967
5880
6906
2540
6792

3648
4492
2801
5488
2508
436539
9574
1827
5688
1066
0633
2415
7200
0634
484747
118538
6167
1538
9335
5779
6803
1547
8413
362218
441456
487249
28
6324
5563
8589
4424
869861
[illegible]031949
06342
1398
875653
94043
0090
8032
98728
47769
045
5505

237

65478
858
48454
46712
25496
200966
635097
8652
576
816
2967
073875
53777
5481
21642
7369
305520
953866
36461
46049
003506
16486
16925
11
49017
4567
688388
9794
049618
481446
06
07256
106781
950561
397963
01901
6254
6533
8505
50830
0230

858
603240
76890
8425
8994
739446
1437
8101
1305
9719
13404
872693
5330
935898
004062
2052
654795
7649
730957
42021
13242
01843
971081
109864
2916
534184
73065
350719
6222
246326
37342
98699
57334
9886
9554
23897
2030894
13666
02539
27019
07554
814591
748825

2849
7255
9505
1786
067824
6521
60323
0201
8377
5688
82101
5
6024
296167
6374
2837
0475
0820
4146374
77217
822572
46426
53062
925411
81516
096942
720809
3679
04763
03197
54143
475675
168
4020
848328
933685
85762
599030
558434
62
871183
224092
6234

90690
08677
53555
994750
58021
0934
484611
9806
2533
40699
51891
62623
2665140
79314
531
4725
23879
786
54
82463
643
00748
0834
87657
1129
43139
8146
2807
9261
1857
17
2125
1854
81546
26941
5218
9205
04258
576158
72298
62695
9502
3109158

8101
19140
0051
2225
89301
29613
48443
92567
7124
46808
43401
29177
97998
333654
35442
81838
40872
398541
90471
7169
706553
3101
5530
095383
759628
2263
5330
910379
985771
35963
6545
687113
4386
2179
16345
174074
818603
5625
741
9481
98922
815416
1431

2916
7254
339990
6881
59317
4689
1481
924597
4198
93200
1644
02128
4431
4964
7507
54017
43530
27702
55264
281597
639298
09072
80420
5709
057659
34090
267332
081105
5073
637827
731463
09025
242962
29064
87335
56105
967614
4136
5428
99097
30135
11836
746243

606091
76775
521789
27
41549
6225
692963
0307
72953
320725
20489
894526
8595
384528
3096
960
87263
7022630
8106
1801
4905
6488
8473
59396
03981
04153
63270
4
12666
26166
668983
601678
853469
186
75829
585814
5
6345
1017
16940
530628
62829
62280

385517
4246
92534
0212
12200
2176
2656
2455
6308
139857
7230
098351
054931
662888
286155
07371
80527
307190
99465
2749
6089
2024
3758
714
965002
52473
18765
7661
6446
9057
514542
10
304962
1536
8902
00731
9805
2355
5974
538070
777204
3938
9286

2606
3560
358268
5739
5490
1017
078409
5608
47113
1405
939556
06910
2031
843920
39677
424123
82812
899335
7474
908
06654
8957
43343
0074
3767
918983
696389
143
40974
95388
88139
15576
788247
1751
37803
5462
94230
76
2091
671488
689370
830493
117

3324
6707
26247
0113
6109
6101
26689
9485
566201
862655
4422
17022
52
68792
57
93614
22993
3301
7938
124825
7634
674688
9262
4522
8752
54825
7314
42024
5817
47572
70891
50245
2387
6661
6323
298
92076
888239
5514
80413
1221
90127
9542

785734
2543
31223
0900
74888
27294
4491
30354
38267
6365
9863
00539
76811
907713
4298
849721
2831
716
58597
7245
8526
2100
42335
0475
86
370742
9655
0689
35586
09071
4333
3141
73480
211780
65484
5946
619186
767536
725988
37730
47432
10390
383450

822883
5105
524117
2929
237088
9789
8760
4985
263428
81480
6297
8577
7437
005326
367005
1428
86299
51415
88929
6236
4034
5030
30355
1766
041769
1822
53403
3414
182502
5301
85714
32048
59437
549
82187
924810
127973
9369
388726
7442
986823
53301
435932

382484
30814
93556
38297
310382
79296
60311
43791
406683
8335
56912
162274
01420
815196
036375
68
523
31
513659
22878
025
30347
80833
872290
418713
601572
05648
2376
87060
19633
506
8419
1383
93326
452700
459997
538745
77611
16869
60868
18916
427823
37630

7384
576644
47684
07556
32120
405952
3298
269304
238490
49157
07996
9079
951141
89558
0014
72672
71998
6363
6604
820849
237
98896
977287
307365
7579
1324
807159
6486
11351
36536
1079
253263
141734
803765
22578
12811
370241
670287
8596
29817
7409
561085
83339

854288
617448
657350
219789
9843
35348
641491
93404
269279
78037
58476
27795
99842
99
310950
306354
884533
119358
106
53992
53505
83180
70687
13216
70320
28651
68436
20737
64085
20913
18255
50815
45200
05018
3
897
70140
85602
76385
19173
61782
12791
95497

28122
23160
75555
83122
41795
80952
46893
66506
46386
33685
88701
1470
6070
7841
8691
2265
245945
357436
958974
783958
645508
67782
81037
8930
060313
961779
061446
668906
5998
9312
327946
484610
37
06
62
80
7587
7506
3448
6432
136635
224593
809231

650722
8752
4589
9847
3138
5363
5406
5902
4355
705522
760195
013193
065339
692302
9142
0082
6186
5900
9918
1649
7244
9832
545850
761908
2446
5791
5069
8995
8031
5105
3521
050003
594529
0312
9672
20729
0028
16168
8656
266212
5775
5642
8982

9090
1372
025382
45047
3035
18664
8255
3207
95219
51454
37707
20994
86315
763776
5909
034285
16933
93516
3572
798132
19870
3109
80265
7472
77594
96540
4175
06260
29612
5291
69864
7100
1789
07730
90
1331
3534
4265
4580
570731
406500
10954
78612

43958
06587
24367
93206
27850
44882
45434
53784
00949
14835
18349
51875
911
70089
70831
37554
76218
306
86167
75137
92755
55344
20608
95767
667
273
91775
38435
88337
61186
50726
40052
73273
81210
55658
73281
05938
47203
94836
81703
92011
23818
38314

97870
95914
37346
03511
18737
64700
70874
12321
26574
34715
88687
4707
0545
8223
7273
9122
948281
712694
614089
950900
121725
460842
239740
242548
609912
471
44500
4546
8790
8379
1552
228599
423248
256606
848275
910848
740950
1322
6564
4053
7483
552728
844921

225488
8864
3784
1404
7230
8197
3253
418768
031162
643561
189475
240246
9461
1442
6483
768383
2443
3603
8592
7287
7363
454231
856532
66
7494
1267
4993
385635
58
3442
7057
8052
4064
2066
7666
4275
9103
96317
558744
722030
31445
747115
4715

257115
5
6175
49438
8868
67972
9466
727375
9804
462021
38835
927686
