## Supplemental Table 4 for "Single-nucleus chromatin accessibility time course of excisional wound repair in rat skin"

**Supplementary Table 4. Marker motif summary**

| p_val | avg_log2FC | pct.1 | pct.2 | p_val_adj | cell_type | motifID |
|---|---|---|---|---|---|---|
| 3.26771454 | 0.376389033195772 | 0.608 | 0.485 | 2.44E-16 | TAC | MA1594.1 |
| 5.81807724 | 0.337356113299777 | 0.589 | 0.482 | 4.34E-13 | TAC | MA0125.1 |
| 4.50149054 | 0.332561303816619 | 0.577 | 0.474 | 3.36E-13 | TAC | MA1515.1 |
| 6.42987358 | 0.33065189778241 | 0.58 | 0.487 | 4.80E-11 | TAC | MA0742.1 |
| 6.67738824 | 0.329060987057898 | 0.588 | 0.494 | 4.98E-12 | TAC | MA1640.1 |
| 1.45071367 | 0.327481680508504 | 0.591 | 0.488 | 1.08E-14 | TAC | MA0685.1 |
| 2.75884360 | 0.323487455217585 | 0.588 | 0.487 | 2.06E-12 | TAC | MA0076.2 |
| 3.04403949 | 0.3219318035567 | 0.6 | 0.497 | 2.27E-12 | TAC | MA1419.1 |
| 3.35635223 | 0.320508045042565 | 0.603 | 0.483 | 2.50E-14 | TAC | MA0079.4 |
| 1.19881653 | 0.31224483043823 | 0.564 | 0.48 | 8.94E-10 | TAC | MA1484.1 |
| 2.25827706 | 0.309644115216894 | 0.561 | 0.477 | 1.68E-08 | TAC | MA0882.1 |
| 1.88142708 | 0.308021714346628 | 0.572 | 0.484 | 1.40E-09 | TAC | MA1552.1 |
| 3.95301114 | 0.305719427185998 | 0.591 | 0.49 | 2.95E-10 | TAC | MA0028.2 |
| 2.06684139 | 0.301964712124456 | 0.553 | 0.484 | 1.54E-07 | TAC | MA0700.2 |
| 5.55629508 | 0.301534273643064 | 0.566 | 0.483 | 4.14E-09 | TAC | MA0799.1 |
| 5.55406410 | 0.30073083650548 | 0.581 | 0.492 | 4.14E-10 | TAC | MA0752.1 |
| 4.08181782 | 0.299070720707743 | 0.595 | 0.496 | 3.05E-09 | TAC | MA1639.1 |
| 5.25849250 | 0.298253506302772 | 0.565 | 0.476 | 3.92E-09 | TAC | MA1533.1 |
| 5.33702426 | 0.294373667516275 | 0.571 | 0.492 | 3.98E-09 | TAC | MA0887.1 |
| 5.07190718 | 0.292991394103729 | 0.584 | 0.486 | 3.78E-11 | TAC | MA0746.2 |
| 2.20641337 | 0.292272921231902 | 0.591 | 0.48 | 1.65E-10 | TAC | MA1127.1 |
| 1.25529337 | 0.292191528685697 | 0.578 | 0.488 | 9.36E-10 | TAC | MA1136.1 |
| 7.99521160 | 0.287897676034417 | 0.575 | 0.487 | 5.96E-09 | TAC | MA0888.1 |
| 1.94294163 | 0.284968728826429 | 0.555 | 0.474 | 1.45E-08 | TAC | MA1517.1 |
| 1.20495498 | 0.284314581834223 | 0.57 | 0.488 | 8.99E-08 | TAC | MA1470.1 |
| 2.58447391 | 0.282873556251147 | 0.554 | 0.482 | 1.93E-07 | TAC | MA0007.3 |
| 4.81644056 | 0.280689403709847 | 0.563 | 0.484 | 3.59E-08 | TAC | MA0510.2 |
| 2.14089978 | 0.279070824675032 | 0.58 | 0.487 | 1.60E-08 | TAC | MA0722.1 |
| 2.14089978 | 0.279070824675032 | 0.58 | 0.487 | 1.60E-08 | TAC | MA0723.1 |
| 9.84820286 | 0.278683794275554 | 0.582 | 0.488 | 7.35E-08 | TAC | MA1501.1 |
| 2.68010669 | 0.272442778198927 | 0.578 | 0.488 | 2.00E-08 | TAC | MA1126.1 |
| 5.38756239 | 0.27193402104635 | 0.572 | 0.487 | 4.02E-07 | TAC | MA1129.1 |
| 3.24929045 | 0.270053693045348 | 0.566 | 0.489 | 2.42E-08 | TAC | MA1632.1 |
| 7.23164192 | 0.268635433743855 | 0.57 | 0.493 | 5.39E-07 | TAC | MA1113.2 |
| 1.05713153 | 0.266710841570415 | 0.561 | 0.485 | 7.89E-07 | TAC | MA0600.2 |
| 7.00263900 | 0.266115055433672 | 0.585 | 0.488 | 5.22E-08 | TAC | MA0132.2 |
| 2.88564545 | 0.266013013434681 | 0.587 | 0.489 | 2.15E-08 | TAC | MA0901.2 |
| 8.63527758 | 0.261706142165517 | 0.568 | 0.485 | 6.44E-08 | TAC | MA0747.1 |
| 3.81169155 | 0.26087309896908 | 0.565 | 0.482 | 2.84E-07 | TAC | MA1511.1 |
| 5.49604849 | 0.259680218828707 | 0.572 | 0.485 | 4.10E-08 | TAC | MA1133.1 |
| 5.81901528 | 0.259058334543406 | 0.573 | 0.487 | 4.34E-08 | TAC | MA0018.4 |

| | | | | | | |
|---|---|---|---|---|---|---|
| 2.67513017 | 0.257844463910246 | 0.575 | 0.49 | 2.00E-08 | TAC | MA1652.1 |
| 2.51901808 | 0.25538942736523 | 0.566 | 0.492 | 1.88E-06 | TAC | MA1117.1 |
| 4.24582760 | 0.255263431011306 | 0.587 | 0.488 | 3.17E-07 | TAC | MA0109.1 |
| 2.48250622 | 0.25231704537906 | 0.57 | 0.491 | 1.85E-06 | TAC | MA0475.2 |
| 9.84945014 | 0.252049253166817 | 0.56 | 0.486 | 7.35E-06 | TAC | MA0037.3 |
| 5.16830880 | 0.364482069527691 | 0.599 | 0.487 | 3.86E-09 | basalKC | MA0138.2 |
| 5.93600715 | 0.33957605727771 | 0.575 | 0.483 | 4.43E-06 | basalKC | MA0007.3 |
| 4.19504579 | 0.335398305310944 | 0.61 | 0.49 | 3.13E-07 | basalKC | MA0059.1 |
| 2.75015484 | 0.333052694882175 | 0.6 | 0.486 | 2.05E-07 | basalKC | MA1594.1 |
| 8.25612913 | 0.330126961457693 | 0.575 | 0.478 | 6.16E-06 | basalKC | MA0882.1 |
| 6.33494491 | 0.328566622725457 | 0.582 | 0.488 | 4.73E-06 | basalKC | MA0722.1 |
| 6.33494491 | 0.328566622725457 | 0.582 | 0.488 | 4.73E-06 | basalKC | MA0723.1 |
| 8.52688461 | 0.322782701550217 | 0.588 | 0.494 | 6.36E-07 | basalKC | MA1125.1 |
| 2.23580609 | 0.322451320682499 | 0.577 | 0.485 | 0.000166791 | basalKC | MA0116.1 |
| 1.88594931 | 0.319532966778161 | 0.564 | 0.482 | 1.41E-05 | basalKC | MA1127.1 |
| 2.77342513 | 0.31634830942467 | 0.575 | 0.485 | 0.000206898 | basalKC | MA0700.2 |
| 1.73355390 | 0.307515053913181 | 0.582 | 0.489 | 1.29E-05 | basalKC | MA0132.2 |
| 4.57843401 | 0.296556716895062 | 0.58 | 0.488 | 3.42E-05 | basalKC | MA0742.1 |
| 1.88585903 | 0.291078898154672 | 0.575 | 0.475 | 1.41E-06 | basalKC | MA1515.1 |
| 4.07842017 | 0.289554756963434 | 0.592 | 0.483 | 3.04E-05 | basalKC | MA0799.1 |
| 8.89544264 | 0.283516129641114 | 0.578 | 0.491 | 6.64E-05 | basalKC | MA1481.1 |
| 3.25176227 | 0.278594891320147 | 0.558 | 0.483 | 0.000242581 | basalKC | MA0668.1 |
| 6.89878766 | 0.277688430267737 | 0.564 | 0.485 | 0.00051465 | basalKC | MA1549.1 |
| 4.68518030 | 0.27713142817566 | 0.589 | 0.498 | 3.50E-05 | basalKC | MA0851.1 |
| 8.85373175 | 0.277061063654804 | 0.561 | 0.474 | 6.60E-05 | basalKC | MA1517.1 |
| 1.72959550 | 0.275183420228915 | 0.592 | 0.492 | 0.000129028 | basalKC | MA0887.1 |
| 1.59489838 | 0.27462382826619 | 0.585 | 0.489 | 0.001189794 | basalKC | MA1470.1 |
| 6.08752273 | 0.274615907592432 | 0.574 | 0.489 | 0.000454129 | basalKC | MA1136.1 |
| 4.19217556 | 0.274114204645097 | 0.561 | 0.482 | 0.000312736 | basalKC | MA1618.1 |
| 2.20040354 | 0.263508815283886 | 0.569 | 0.491 | 0.001641501 | basalKC | MA0706.1 |
| 7.32504646 | 0.261701320244894 | 0.575 | 0.484 | 0.000546448 | basalKC | MA0493.1 |
| 2.35182446 | 0.261224468045331 | 0.577 | 0.488 | 0.000175446 | basalKC | MA0162.4 |
| 7.75989076 | 0.260504936212194 | 0.567 | 0.489 | 0.000578888 | basalKC | MA0842.2 |
| 2.53520563 | 0.259936292248027 | 0.578 | 0.495 | 0.000189126 | basalKC | MA0855.1 |
| 1.50270695 | 0.255328419328871 | 0.552 | 0.482 | 0.011210194 | basalKC | MA1554.1 |
| 3.61285369 | 0.255012707109733 | 0.594 | 0.491 | 0.000269519 | basalKC | MA1478.1 |
| 7.57700837 | 0.251012337263365 | 0.611 | 0.499 | 5.65E-06 | basalKC | MA1627.1 |
| 1.19842767 | 0.456864976649551 | 0.621 | 0.487 | 8.94E-09 | SG | MA1594.1 |
| 2.70682006 | 0.45067808234255 | 0.627 | 0.485 | 2.02E-08 | SG | MA0700.2 |
| 2.98096206 | 0.427600775171049 | 0.614 | 0.474 | 2.22E-08 | SG | MA1517.1 |
| 4.51720698 | 0.411888304653746 | 0.629 | 0.489 | 3.37E-08 | SG | MA0132.2 |
| 2.71762998 | 0.40588230488209 | 0.591 | 0.486 | 0.000202735 | SG | MA0116.1 |
| 1.30384116 | 0.401849124068664 | 0.593 | 0.483 | 9.73E-05 | SG | MA0668.1 |

| | | | | | | |
|---|---|---|---|---|---|---|
| 1.18275982 | 0.395689915392845 | 0.578 | 0.483 | 8.82E-06 | SG | MA0007.3 |
| 2.03321025 | 0.394894560790541 | 0.624 | 0.488 | 1.52E-07 | SG | MA0722.1 |
| 2.03321025 | 0.394894560790541 | 0.624 | 0.488 | 1.52E-07 | SG | MA0723.1 |
| 2.83046875 | 0.394384043749952 | 0.601 | 0.493 | 2.11E-05 | SG | MA0887.1 |
| 1.02230436 | 0.392336751360291 | 0.601 | 0.496 | 7.63E-07 | SG | MA0855.1 |
| 3.00744766 | 0.391619097033095 | 0.604 | 0.478 | 2.24E-06 | SG | MA0882.1 |
| 6.39658959 | 0.391594153718926 | 0.621 | 0.483 | 4.77E-07 | SG | MA0799.1 |
| 3.74331029 | 0.389795148242341 | 0.619 | 0.499 | 2.79E-06 | SG | MA1542.1 |
| 1.04871145 | 0.386906372353231 | 0.614 | 0.489 | 7.82E-06 | SG | MA0888.1 |
| 2.62279340 | 0.376090006198779 | 0.598 | 0.487 | 1.96E-05 | SG | MA0903.1 |
| 3.44187019 | 0.373506453996657 | 0.634 | 0.489 | 2.57E-06 | SG | MA0109.1 |
| 9.96232060 | 0.371172373758653 | 0.601 | 0.486 | 7.43E-06 | SG | MA0600.2 |
| 5.60150272 | 0.370783921632667 | 0.588 | 0.489 | 0.000417872 | SG | MA0742.1 |
| 2.52957880 | 0.368738044049883 | 0.601 | 0.485 | 1.89E-05 | SG | MA0510.2 |
| 6.59920280 | 0.355603703250845 | 0.588 | 0.486 | 0.000492301 | SG | MA1552.1 |
| 2.15420794 | 0.355497161973533 | 0.616 | 0.485 | 1.61E-05 | SG | MA1105.2 |
| 6.15637207 | 0.354046708295574 | 0.581 | 0.482 | 0.000459265 | SG | MA1618.1 |
| 4.31345164 | 0.351220982942132 | 0.601 | 0.495 | 3.22E-05 | SG | MA1125.1 |
| 3.77224962 | 0.350181410677442 | 0.593 | 0.49 | 0.00028141 | SG | MA1136.1 |
| 3.30940813 | 0.343884486097951 | 0.575 | 0.484 | 0.000246882 | SG | MA1516.1 |
| 7.30079824 | 0.339875485898601 | 0.588 | 0.492 | 5.45E-05 | SG | MA1478.1 |
| 2.12094059 | 0.338287964453032 | 0.601 | 0.486 | 0.000158222 | SG | MA0647.1 |
| 3.59471474 | 0.337243675271754 | 0.586 | 0.489 | 0.002681657 | SG | MA1470.1 |
| 3.11444985 | 0.335955996085839 | 0.598 | 0.492 | 0.000232338 | SG | MA0706.1 |
| 2.75588194 | 0.333939957788883 | 0.611 | 0.484 | 0.000205589 | SG | MA0125.1 |
| 8.52806292 | 0.332357900341022 | 0.575 | 0.476 | 6.36E-05 | SG | MA1515.1 |
| 8.12040435 | 0.330874766548376 | 0.583 | 0.491 | 0.000605782 | SG | MA1481.1 |
| 5.73653024 | 0.329168623634652 | 0.606 | 0.486 | 0.000427945 | SG | MA1549.1 |
| 7.48822242 | 0.32844361272962 | 0.573 | 0.493 | 0.005586214 | SG | MA0461.2 |
| 2.08903418 | 0.321422260945294 | 0.588 | 0.49 | 0.00155842 | SG | MA1501.1 |
| 7.23593884 | 0.316804496743974 | 0.575 | 0.49 | 0.00539801 | SG | MA1535.1 |
| 1.17434213 | 0.313732743875106 | 0.591 | 0.488 | 0.000876059 | SG | MA1479.1 |
| 6.29052728 | 0.313576501418831 | 0.581 | 0.485 | 0.000469273 | SG | MA0079.4 |
| 4.16410365 | 0.310014681057417 | 0.598 | 0.499 | 0.003106421 | SG | MA0869.1 |
| 1.04174499 | 0.307925803072091 | 0.596 | 0.486 | 0.007771418 | SG | MA0678.1 |
| 5.76375922 | 0.307911782776165 | 0.583 | 0.481 | 0.004299764 | SG | MA1484.1 |
| 1.10326034 | 0.302814875258614 | 0.593 | 0.492 | 0.000823032 | SG | MA1471.1 |
| 4.59619628 | 0.299596937610732 | 0.563 | 0.489 | 0.034287624 | SG | MA1519.1 |
| 2.39646683 | 0.29388721695582 | 0.583 | 0.482 | 0.017877643 | SG | MA1127.1 |
| 1.16169396 | 0.288791761468119 | 0.583 | 0.482 | 0.008666237 | SG | MA1554.1 |
| 6.50886092 | 0.28441650775817 | 0.573 | 0.487 | 0.048556103 | SG | MA1585.1 |
| 1.90805342 | 0.2842453844894 | 0.583 | 0.49 | 0.014234079 | SG | MA0091.1 |
| 2.86141892 | 0.280649483003649 | 0.57 | 0.495 | 0.021346185 | SG | MA0681.2 |

| | | | | | | |
|---|---|---|---|---|---|---|
| 3.21782201[illegible] | 0.278111631943372 | 0.57 | 0.49 | 0.024004952 | SG | MA0685.1 |
| 1.10796625[illegible] | 0.278017498191602 | 0.609 | 0.498 | 0.008265428 | SG | MA1639.1 |
| 4.99339166[illegible] | 0.277935281208817 | 0.563 | 0.49 | 0.037250702 | SG | MA0842.[illegible] |
| 3.03790100[illegible] | 0.274374712416657 | 0.578 | 0.489 | 0.022662742 | SG | MA1129.1 |
| 2.63433760[illegible] | 0.273060051625932 | 0.583 | 0.492 | 0.019652159 | SG | MA0675.1 |
| 2.76084838[illegible] | 0.267406542121213 | 0.573 | 0.491 | 0.020595929 | SG | MA0083.[illegible] |
| 4.55285964[illegible] | 0.267219569794615 | 0.598 | 0.485 | 0.033964333 | SG | MA0739.1 |
| 5.93432777[illegible] | 0.264839063672033 | 0.575 | 0.502 | 0.044270085 | SG | MA0597.1 |
| 6.49843728[illegible] | 0.256875624574798 | 0.573 | 0.494 | 0.048478342 | SG | MA0674.1 |
| 3.64873972[illegible] | 0.256757587943776 | 0.586 | 0.491 | 0.027219598 | SG | MA0059.1 |
| 3.50679531[illegible] | 0.254330131221307 | 0.581 | 0.499 | 0.026160693 | SG | MA0632.[illegible] |
| 0 | 1.07772751349466 | 0.781 | 0.454 | 0.00 | ENDO | MA1560.1 |
| 0 | 0.883099900678904 | 0.772 | 0.467 | 0.00 | ENDO | MA0006.1 |
| 0 | 0.876777593432043 | 0.759 | 0.466 | 0.00 | ENDO | MA1497.1 |
| 7.26216952[illegible] | 0.844667976359255 | 0.737 | 0.464 | 5.42E-299 | ENDO | MA0071.1 |
| 1.07023940[illegible] | 0.842679983147396 | 0.75 | 0.475 | 7.98E-289 | ENDO | MA0821.1 |
| 0 | 0.842348831726205 | 0.747 | 0.477 | 0.00 | ENDO | MA0004.1 |
| 6.23259749[illegible] | 0.80472330997221 | 0.741 | 0.468 | 4.65E-294 | ENDO | MA0513.1 |
| 7.56470857[illegible] | 0.801652068597709 | 0.733 | 0.469 | 5.64E-273 | ENDO | MA0737.1 |
| 6.56046331[illegible] | 0.790537951101599 | 0.731 | 0.471 | 4.89E-271 | ENDO | MA1474.1 |
| 2.46524563[illegible] | 0.788577134056997 | 0.739 | 0.47 | 1.84E-281 | ENDO | MA0040.1 |
| 1.53939196[illegible] | 0.778383837359885 | 0.747 | 0.47 | 1.15E-274 | ENDO | MA0771.1 |
| 6.98450330[illegible] | 0.708656790836235 | 0.734 | 0.474 | 5.21E-271 | ENDO | MA1623.1 |
| 3.37757917[illegible] | 0.696255842963051 | 0.712 | 0.476 | 2.52E-212 | ENDO | MA0135.1 |
| 2.06054444[illegible] | 0.695677193628927 | 0.711 | 0.468 | 1.54E-220 | ENDO | MA0522.[illegible] |
| 7.83781630[illegible] | 0.688266456151856 | 0.706 | 0.473 | 5.85E-209 | ENDO | MA0844.1 |
| 4.09461966[illegible] | 0.680635175560587 | 0.691 | 0.478 | 3.05E-178 | ENDO | MA0680.1 |
| 1.55451521[illegible] | 0.654723929713725 | 0.686 | 0.466 | 1.16E-180 | ENDO | MA0693.[illegible] |
| 7.61704875[illegible] | 0.653849452621439 | 0.679 | 0.477 | 5.68E-176 | ENDO | MA0072.1 |
| 2.49235498[illegible] | 0.64814957389491 | 0.687 | 0.464 | 1.86E-180 | ENDO | MA0103.[illegible] |
| 1.09236165[illegible] | 0.635089594871508 | 0.683 | 0.468 | 8.15E-178 | ENDO | MA1648.1 |
| 1.55402178[illegible] | 0.633437781411603 | 0.699 | 0.484 | 1.16E-171 | ENDO | MA1464.1 |
| 2.24359838[illegible] | 0.630297770661139 | 0.681 | 0.477 | 1.67E-159 | ENDO | MA0628.1 |
| 1.96203020[illegible] | 0.623216151001675 | 0.694 | 0.474 | 1.46E-173 | ENDO | MA1587.1 |
| 4.15748532[illegible] | 0.6192508912995 | 0.697 | 0.476 | 3.10E-166 | ENDO | MA0486.[illegible] |
| 2.29279626[illegible] | 0.618583651288121 | 0.69 | 0.471 | 1.71E-190 | ENDO | MA1107.[illegible] |
| 6.35390918[illegible] | 0.618552428665551 | 0.682 | 0.485 | 4.74E-159 | ENDO | MA0058.[illegible] |
| 5.35057511[illegible] | 0.601886728747506 | 0.686 | 0.483 | 3.99E-150 | ENDO | MA0067.1 |
| 7.64520679[illegible] | 0.591305082813868 | 0.689 | 0.478 | 5.70E-152 | ENDO | MA0845.1 |
| 1.82520048[illegible] | 0.587749998210957 | 0.663 | 0.482 | 1.36E-131 | ENDO | MA0819.1 |
| 3.13280452[illegible] | 0.587583870197213 | 0.672 | 0.481 | 2.34E-139 | ENDO | MA1571.1 |
| 4.59670333[illegible] | 0.587103058135369 | 0.674 | 0.482 | 3.43E-140 | ENDO | MA0608.1 |
| 1.05962539[illegible] | 0.587102148873337 | 0.687 | 0.481 | 7.90E-149 | ENDO | MA1467.1 |

| | | | | | | |
|---|---|---|---|---|---|---|
| 1.49847049 | 0.586712797542059 | 0.674 | 0.477 | 1.12E-134 | ENDO | MA0822.1 |
| 3.84300368 | 0.586619953655244 | 0.662 | 0.472 | 2.87E-134 | ENDO | MA0755.1 |
| 8.16018703 | 0.582795679267284 | 0.678 | 0.488 | 6.09E-145 | ENDO | MA1489.1 |
| 1.12640885 | 0.581314777969877 | 0.687 | 0.48 | 8.40E-150 | ENDO | MA1152.1 |
| 2.50877568 | 0.578932308478138 | 0.653 | 0.475 | 1.87E-128 | ENDO | MA1151.1 |
| 4.97134157 | 0.577436877876346 | 0.681 | 0.5 | 3.71E-128 | ENDO | MA0093.3 |
| 2.45268161 | 0.575221880343902 | 0.666 | 0.471 | 1.83E-141 | ENDO | MA0687.1 |
| 7.44215602 | 0.568235820740402 | 0.67 | 0.476 | 5.55E-129 | ENDO | MA0525.2 |
| 1.40781227 | 0.564037823879808 | 0.66 | 0.473 | 1.05E-125 | ENDO | MA0046.2 |
| 7.81255497 | 0.563250472725259 | 0.663 | 0.484 | 5.83E-121 | ENDO | MA0780.1 |
| 3.89929905 | 0.561022030128717 | 0.654 | 0.477 | 2.91E-126 | ENDO | MA1150.1 |
| 2.23326335 | 0.547947359512727 | 0.646 | 0.47 | 1.67E-119 | ENDO | MA0673.1 |
| 6.73909228 | 0.545940837798823 | 0.655 | 0.484 | 5.03E-118 | ENDO | MA0464.2 |
| 4.06960778 | 0.542736306001511 | 0.659 | 0.471 | 3.04E-144 | ENDO | MA0073.1 |
| 4.02021896 | 0.541812491009878 | 0.68 | 0.474 | 3.00E-160 | ENDO | MA0050.2 |
| 2.61017446 | 0.540147428823449 | 0.64 | 0.474 | 1.95E-107 | ENDO | MA0111.1 |
| 2.12919945 | 0.53854707893008 | 0.665 | 0.49 | 1.59E-118 | ENDO | MA0831.2 |
| 5.15417372 | 0.537825306533101 | 0.671 | 0.502 | 3.85E-110 | ENDO | MA0526.3 |
| 4.72392331 | 0.533641642632193 | 0.664 | 0.489 | 3.52E-120 | ENDO | MA0017.2 |
| 3.68614773 | 0.531617206869815 | 0.657 | 0.477 | 2.75E-109 | ENDO | MA0604.1 |
| 2.48920980 | 0.531566171854044 | 0.646 | 0.489 | 1.86E-100 | ENDO | MA0756.1 |
| 5.84192682 | 0.530258187619521 | 0.66 | 0.48 | 4.36E-114 | ENDO | MA0679.2 |
| 4.91611735 | 0.527540944927567 | 0.641 | 0.475 | 3.67E-106 | ENDO | MA0629.1 |
| 9.67408510 | 0.527425338553764 | 0.642 | 0.474 | 7.22E-108 | ENDO | MA1559.1 |
| 1.18443773 | 0.526896058180872 | 0.658 | 0.487 | 8.84E-110 | ENDO | MA0603.1 |
| 1.25865696 | 0.526000442898236 | 0.657 | 0.485 | 9.39E-110 | ENDO | MA1572.1 |
| 2.02897381 | 0.525303042149857 | 0.646 | 0.482 | 1.51E-113 | ENDO | MA1114.1 |
| 1.71772685 | 0.523357269513215 | 0.649 | 0.482 | 1.28E-110 | ENDO | MA1108.2 |
| 5.55786595 | 0.518778014642997 | 0.65 | 0.481 | 4.15E-111 | ENDO | MA0483.1 |
| 7.16540218 | 0.518444406834879 | 0.664 | 0.483 | 5.35E-117 | ENDO | MA0032.2 |
| 7.69489925 | 0.509933753105059 | 0.636 | 0.473 | 5.74E-104 | ENDO | MA0622.1 |
| 2.44453114 | 0.506576617050141 | 0.634 | 0.472 | 1.82E-104 | ENDO | MA0002.2 |
| 1.33119808 | 0.505807212778715 | 0.667 | 0.489 | 9.93E-106 | ENDO | MA1524.1 |
| 4.21679569 | 0.504336283126507 | 0.645 | 0.484 | 3.15E-96 | ENDO | MA0823.1 |
| 9.51822124 | 0.503694365224216 | 0.653 | 0.489 | 7.10E-99 | ENDO | MA0914.1 |
| 9.09352427 | 0.502008136569741 | 0.646 | 0.482 | 6.78E-101 | ENDO | MA1110.1 |
| 8.05726494 | 0.500308787906945 | 0.653 | 0.493 | 6.01E-106 | ENDO | MA1485.1 |
| 4.90132511 | 0.499292445704071 | 0.641 | 0.471 | 3.66E-100 | ENDO | MA0024.3 |
| 2.77859808 | 0.496055871490791 | 0.631 | 0.469 | 2.07E-91 | ENDO | MA1554.1 |
| 1.57531682 | 0.491556043739556 | 0.621 | 0.475 | 1.18E-89 | ENDO | MA0896.1 |
| 3.72448167 | 0.489928638392864 | 0.618 | 0.473 | 2.78E-88 | ENDO | MA0897.1 |
| 9.73055074 | 0.487068661563736 | 0.651 | 0.489 | 7.26E-98 | ENDO | MA0797.1 |
| 3.64711133 | 0.486645429055355 | 0.644 | 0.484 | 2.72E-94 | ENDO | MA0106.3 |

| | | | | | | |
|---|---|---|---|---|---|---|
| 3.64819943 | 0.480336538452673 | 0.632 | 0.476 | 2.72E-86 | ENDO | MA1645.1 |
| 5.38058261 | 0.476645122327488 | 0.627 | 0.478 | 4.01E-84 | ENDO | MA0727.1 |
| 1.47215015 | 0.476426055279875 | 0.629 | 0.477 | 1.10E-87 | ENDO | MA0864. |
| 1.71590549 | 0.470482001988088 | 0.65 | 0.486 | 1.28E-91 | ENDO | MA1491.1 |
| 2.46881465 | 0.468953057400259 | 0.638 | 0.481 | 1.84E-88 | ENDO | MA0697.1 |
| 4.02559352 | 0.468702309266521 | 0.652 | 0.491 | 3.00E-93 | ENDO | MA0514.1 |
| 2.91673536 | 0.465484241023502 | 0.622 | 0.48 | 2.18E-85 | ENDO | MA0505.1 |
| 3.16742629 | 0.463235816978225 | 0.643 | 0.49 | 2.36E-90 | ENDO | MA1628.1 |
| 6.65293082 | 0.461397369672396 | 0.633 | 0.491 | 4.96E-84 | ENDO | MA0796.1 |
| 3.18196399 | 0.461362687064074 | 0.639 | 0.484 | 2.37E-86 | ENDO | MA0825.1 |
| 2.86697826 | 0.460872576312334 | 0.643 | 0.488 | 2.14E-90 | ENDO | MA1155.1 |
| 4.46705414 | 0.457389383186154 | 0.634 | 0.484 | 3.33E-81 | ENDO | MA1583.1 |
| 3.12800358 | 0.455096294015083 | 0.642 | 0.494 | 2.33E-86 | ENDO | MA0783.1 |
| 4.49721608 | 0.455070702953054 | 0.618 | 0.475 | 3.35E-78 | ENDO | MA1503.1 |
| 9.41605053 | 0.450730074625301 | 0.636 | 0.477 | 7.02E-88 | ENDO | MA0830. |
| 1.46330614 | 0.45069456683308 | 0.608 | 0.474 | 1.09E-73 | ENDO | MA0736.1 |
| 1.16136361 | 0.45060339638637 | 0.619 | 0.476 | 8.66E-78 | ENDO | MA1479.1 |
| 1.20655941 | 0.449791482930533 | 0.63 | 0.481 | 9.00E-78 | ENDO | MA0683.1 |
| 3.77507969 | 0.447834354261867 | 0.617 | 0.479 | 2.82E-76 | ENDO | MA0029.1 |
| 9.85572980 | 0.447640971880588 | 0.611 | 0.473 | 7.35E-78 | ENDO | MA0696.1 |
| 9.70666406 | 0.447584100780601 | 0.621 | 0.483 | 7.24E-75 | ENDO | MA0643.1 |
| 5.39792007 | 0.446244166577423 | 0.626 | 0.478 | 4.03E-77 | ENDO | MA0905.1 |
| 9.85017631 | 0.444846137339934 | 0.627 | 0.492 | 7.35E-75 | ENDO | MA0871. |
| 5.69245051 | 0.444323616104433 | 0.639 | 0.485 | 4.25E-90 | ENDO | MA0041.1 |
| 1.12828514 | 0.444175912447718 | 0.609 | 0.475 | 8.42E-71 | ENDO | MA0898.1 |
| 2.28128565 | 0.443168880842306 | 0.634 | 0.49 | 1.70E-80 | ENDO | MA1132.1 |
| 3.16241901 | 0.441074563516805 | 0.626 | 0.481 | 2.36E-76 | ENDO | MA1468.1 |
| 2.78476362 | 0.440575644683214 | 0.606 | 0.474 | 2.08E-67 | ENDO | MA0493.1 |
| 1.85452799 | 0.439764021063093 | 0.627 | 0.478 | 1.38E-78 | ENDO | MA0770.1 |
| 4.62556362 | 0.43834888690652 | 0.638 | 0.487 | 3.45E-81 | ENDO | MA0785.1 |
| 1.15175021 | 0.430778111675304 | 0.625 | 0.483 | 8.59E-72 | ENDO | MA0715.1 |
| 2.38782819 | 0.430074119474456 | 0.612 | 0.473 | 1.78E-68 | ENDO | MA1538.1 |
| 5.67450637 | 0.429389878235047 | 0.628 | 0.484 | 4.23E-76 | ENDO | MA0772.1 |
| 4.18313244 | 0.423349558349434 | 0.618 | 0.477 | 3.12E-72 | ENDO | MA1631.1 |
| 3.45213871 | 0.419908445938966 | 0.618 | 0.481 | 2.58E-68 | ENDO | MA0153. |
| 1.60148178 | 0.419150736326526 | 0.635 | 0.491 | 1.19E-70 | ENDO | MA1466.1 |
| 4.95358362 | 0.419101840324187 | 0.61 | 0.485 | 3.70E-66 | ENDO | MA1570.1 |
| 4.01975174 | 0.417759519454139 | 0.626 | 0.488 | 3.00E-72 | ENDO | MA1579.1 |
| 1.09721392 | 0.415855179722305 | 0.63 | 0.49 | 8.19E-80 | ENDO | MA0517.1 |
| 3.73840392 | 0.411106536508182 | 0.609 | 0.478 | 2.79E-66 | ENDO | MA0751.1 |
| 2.02066901 | 0.410458094447558 | 0.604 | 0.477 | 1.51E-59 | ENDO | MA0862.1 |
| 6.85026959 | 0.399648766489724 | 0.625 | 0.489 | 5.11E-65 | ENDO | MA0463. |
| 1.11248208 | 0.398575174596392 | 0.603 | 0.48 | 8.30E-57 | ENDO | MA0713.1 |

| | | | | | | |
|---|---|---|---|---|---|---|
| 5.52689104 | 0.394344041875473 | 0.626 | 0.495 | 4.12E-63 | ENDO | MA0782.2 |
| 1.31975324 | 0.394021475330564 | 0.602 | 0.48 | 9.85E-55 | ENDO | MA0626.1 |
| 2.15919846 | 0.389856972566486 | 0.621 | 0.495 | 1.61E-59 | ENDO | MA0088.2 |
| 1.33471522 | 0.388913631679749 | 0.607 | 0.493 | 9.96E-54 | ENDO | MA0664.1 |
| 8.12857248 | 0.383427203959168 | 0.623 | 0.484 | 6.06E-66 | ENDO | MA1125.1 |
| 1.09412381 | 0.380581947940612 | 0.606 | 0.487 | 8.16E-54 | ENDO | MA1500.1 |
| 6.93580533 | 0.379964859905384 | 0.599 | 0.48 | 5.17E-52 | ENDO | MA1470.1 |
| 1.68740455 | 0.370446156200025 | 0.598 | 0.48 | 1.26E-47 | ENDO | MA0113.3 |
| 1.45782348 | 0.369797234879691 | 0.602 | 0.48 | 1.09E-50 | ENDO | MA1535.1 |
| 2.21398864 | 0.364448242304997 | 0.59 | 0.479 | 1.65E-49 | ENDO | MA1561.1 |
| 2.26552911 | 0.359898221773232 | 0.599 | 0.487 | 1.69E-48 | ENDO | MA1640.1 |
| 1.68109445 | 0.359466653652852 | 0.585 | 0.475 | 1.25E-45 | ENDO | MA0056.2 |
| 3.43915302 | 0.358583423819535 | 0.591 | 0.482 | 2.57E-47 | ENDO | MA0091.1 |
| 8.32385798 | 0.358054026182913 | 0.591 | 0.477 | 6.21E-46 | ENDO | MA1633.1 |
| 1.27486419 | 0.356520111758229 | 0.603 | 0.487 | 9.51E-50 | ENDO | MA1601.1 |
| 5.44803681 | 0.353195360979515 | 0.603 | 0.478 | 4.06E-49 | ENDO | MA0899.1 |
| 7.76002848 | 0.352135649659357 | 0.597 | 0.481 | 5.79E-44 | ENDO | MA1501.1 |
| 8.01732256 | 0.349283416709281 | 0.601 | 0.482 | 5.98E-48 | ENDO | MA1119.1 |
| 1.79659230 | 0.34924860993398 | 0.614 | 0.493 | 1.34E-48 | ENDO | MA0884.1 |
| 1.50312696 | 0.348203663008743 | 0.597 | 0.482 | 1.12E-44 | ENDO | MA1506.1 |
| 2.18320790 | 0.345395761913778 | 0.594 | 0.484 | 1.63E-41 | ENDO | MA0860.1 |
| 2.98325568 | 0.345319089716017 | 0.601 | 0.486 | 2.23E-45 | ENDO | MA0719.1 |
| 3.56085064 | 0.3445074253754 | 0.588 | 0.486 | 2.66E-40 | ENDO | MA0104.4 |
| 2.59371087 | 0.339691763139305 | 0.618 | 0.493 | 1.93E-54 | ENDO | MA1596.1 |
| 2.90038859 | 0.337382079672722 | 0.581 | 0.474 | 2.16E-40 | ENDO | MA0470.2 |
| 1.97266791 | 0.332144609473779 | 0.603 | 0.493 | 1.47E-42 | ENDO | MA0119.1 |
| 1.68528981 | 0.32948249197532 | 0.591 | 0.484 | 1.26E-42 | ENDO | MA1478.1 |
| 4.27967565 | 0.327026364085277 | 0.599 | 0.498 | 3.19E-41 | ENDO | MA0619.1 |
| 1.72542396 | 0.324558383080294 | 0.584 | 0.475 | 1.29E-39 | ENDO | MA0728.1 |
| 4.49792238 | 0.324162740003551 | 0.591 | 0.496 | 3.36E-36 | ENDO | MA0620.3 |
| 2.17679615 | 0.320257934702226 | 0.6 | 0.49 | 1.62E-40 | ENDO | MA1135.1 |
| 6.02460181 | 0.318690394884863 | 0.589 | 0.489 | 4.49E-36 | ENDO | MA0779.1 |
| 7.93986143 | 0.317731983555813 | 0.606 | 0.503 | 5.92E-38 | ENDO | MA0692.1 |
| 1.77032354 | 0.314263806865419 | 0.594 | 0.486 | 1.32E-36 | ENDO | MA0596.1 |
| 4.36544408 | 0.314000776128822 | 0.58 | 0.486 | 3.26E-35 | ENDO | MA1540.1 |
| 2.79011058 | 0.313863333962525 | 0.59 | 0.487 | 2.08E-37 | ENDO | MA0839.1 |
| 7.78379681 | 0.313452926549706 | 0.601 | 0.488 | 5.81E-41 | ENDO | MA0508.3 |
| 6.46054818 | 0.310943402407111 | 0.582 | 0.479 | 4.82E-34 | ENDO | MA0815.1 |
| 2.65088876 | 0.310199430883769 | 0.582 | 0.491 | 1.98E-38 | ENDO | MA0732.1 |
| 6.18332005 | 0.309355328116943 | 0.588 | 0.483 | 4.61E-37 | ENDO | MA0867.2 |
| 5.47453306 | 0.309035033230766 | 0.573 | 0.481 | 4.08E-35 | ENDO | MA0499.2 |
| 1.16946103 | 0.30889491108647 | 0.576 | 0.482 | 8.72E-34 | ENDO | MA1558.1 |
| 4.10151557 | 0.307683775757756 | 0.576 | 0.482 | 3.06E-34 | ENDO | MA1656.1 |

| | | | | | | |
|---|---|---|---|---|---|---|
| 5.484962479 | 0.303927075496323 | 0.589 | 0.493 | 4.09E-34 | ENDO | MA1149.1 |
| 6.523836817 | 0.30213733126591 | 0.582 | 0.486 | 4.87E-36 | ENDO | MA0774.1 |
| 6.697114594 | 0.298827888835557 | 0.573 | 0.481 | 5.00E-32 | ENDO | MA1642.1 |
| 4.317709693 | 0.297132440795052 | 0.581 | 0.489 | 3.22E-30 | ENDO | MA0879.1 |
| 2.597346625 | 0.296897880491209 | 0.584 | 0.482 | 1.94E-33 | ENDO | MA1472.1 |
| 4.490979407 | 0.296453260000846 | 0.598 | 0.49 | 3.35E-37 | ENDO | MA0846.1 |
| 2.287690180 | 0.295254271583123 | 0.576 | 0.477 | 1.71E-32 | ENDO | MA0139.1 |
| 1.082826408 | 0.294844266032411 | 0.574 | 0.49 | 8.08E-29 | ENDO | MA0259.1 |
| 2.384572869 | 0.290306434106972 | 0.582 | 0.489 | 1.78E-29 | ENDO | MA0592.3 |
| 5.704987687 | 0.28893877915031 | 0.58 | 0.488 | 4.26E-30 | ENDO | MA0659.2 |
| 8.094369480 | 0.287858787075286 | 0.574 | 0.485 | 6.04E-30 | ENDO | MA0754.1 |
| 7.070246532 | 0.287833725320677 | 0.572 | 0.487 | 5.27E-29 | ENDO | MA0147.3 |
| 2.267007115 | 0.287695729844716 | 0.583 | 0.494 | 1.69E-30 | ENDO | MA0524.2 |
| 8.072165227 | 0.283666610797901 | 0.58 | 0.487 | 6.02E-30 | ENDO | MA0805.1 |
| 8.158358560 | 0.28225795617123 | 0.561 | 0.483 | 6.09E-25 | ENDO | MA1493.1 |
| 3.614341732 | 0.28063740263132 | 0.57 | 0.484 | 2.70E-26 | ENDO | MA0639.1 |
| 2.956594792 | 0.280135939093266 | 0.579 | 0.485 | 2.21E-28 | ENDO | MA0913.2 |
| 6.595004060 | 0.278804601939165 | 0.574 | 0.487 | 4.92E-27 | ENDO | MA0161.2 |
| 3.675847150 | 0.276517299890665 | 0.574 | 0.479 | 2.74E-32 | ENDO | MA0747.1 |
| 2.392332810 | 0.275642802953995 | 0.581 | 0.494 | 1.78E-26 | ENDO | MA0494.1 |
| 1.350418833 | 0.271024535704133 | 0.573 | 0.485 | 1.01E-26 | ENDO | MA0806.1 |
| 4.832955023 | 0.270656814205195 | 0.568 | 0.494 | 3.61E-24 | ENDO | MA0159.1 |
| 1.182962022 | 0.270466285887836 | 0.576 | 0.491 | 8.82E-27 | ENDO | MA0734.2 |
| 4.006766268 | 0.265392109411999 | 0.57 | 0.486 | 2.99E-25 | ENDO | MA0461.2 |
| 1.159329047 | 0.265284227924337 | 0.58 | 0.496 | 8.65E-26 | ENDO | MA0788.1 |
| 1.264540879 | 0.263976656973197 | 0.56 | 0.483 | 9.43E-22 | ENDO | MA0651.1 |
| 1.137920097 | 0.261594406434291 | 0.582 | 0.481 | 8.49E-27 | ENDO | MA1607.1 |
| 9.957293989 | 0.2601326223446 | 0.584 | 0.493 | 7.43E-26 | ENDO | MA1144.1 |
| 3.742733259 | 0.259830061858944 | 0.565 | 0.485 | 2.79E-25 | ENDO | MA0688.1 |
| 8.918928607 | 0.259600581114984 | 0.586 | 0.485 | 6.65E-31 | ENDO | MA0081.2 |
| 4.465081464 | 0.257140562033685 | 0.577 | 0.497 | 3.33E-23 | ENDO | MA0636.1 |
| 1.141501268 | 0.255232814880462 | 0.569 | 0.492 | 8.52E-22 | ENDO | MA1639.1 |
| 5.050850429 | 0.252094816452126 | 0.578 | 0.502 | 3.77E-21 | ENDO | MA0142.1 |
| 2.092631088 | 0.250585390256147 | 0.572 | 0.495 | 1.56E-22 | ENDO | MA0758.1 |
| 4.832538594 | 0.579823996389889 | 0.674 | 0.482 | 3.61E-44 | injFB | MA1573.1 |
| 9.337664489 | 0.53514694028642 | 0.683 | 0.482 | 6.97E-45 | injFB | MA0747.1 |
| 4.555677794 | 0.527777643113276 | 0.671 | 0.485 | 3.40E-42 | injFB | MA0740.1 |
| 8.684650643 | 0.514856798709241 | 0.649 | 0.485 | 6.48E-38 | injFB | MA0746.2 |
| 2.730308699 | 0.466415752134569 | 0.628 | 0.485 | 2.04E-25 | injFB | MA0813.1 |
| 5.139639962 | 0.46094846306405 | 0.636 | 0.487 | 3.83E-29 | injFB | MA0685.1 |
| 6.290140677 | 0.451999153945697 | 0.611 | 0.485 | 4.69E-23 | injFB | MA0815.1 |
| 1.406736827 | 0.447185172351301 | 0.598 | 0.488 | 1.05E-22 | injFB | MA0109.1 |
| 1.015813231 | 0.445035135272664 | 0.631 | 0.485 | 7.58E-24 | injFB | MA0163.1 |

| | | | | | | |
|---|---|---|---|---|---|---|
| 6.17272002 | 0.444654519152094 | 0.632 | 0.482 | 4.60E-22 | injFB | MA1608.1 |
| 2.34488924 | 0.44322207552615 | 0.625 | 0.489 | 1.75E-23 | injFB | MA1121.1 |
| 4.62443765 | 0.441725817219756 | 0.612 | 0.486 | 3.45E-20 | injFB | MA0742.1 |
| 3.68172410 | 0.434935770456285 | 0.637 | 0.486 | 2.75E-28 | injFB | MA1564.1 |
| 4.95622827 | 0.430865227384038 | 0.611 | 0.484 | 3.70E-17 | injFB | MA0116.1 |
| 8.04957856 | 0.426183298106508 | 0.65 | 0.488 | 6.00E-33 | injFB | MA1513.1 |
| 5.17632531 | 0.424399716881595 | 0.634 | 0.483 | 3.86E-26 | injFB | MA1512.1 |
| 9.55485851 | 0.416303954209402 | 0.6 | 0.482 | 7.13E-17 | injFB | MA0493.1 |
| 7.88140270 | 0.410927952456583 | 0.605 | 0.482 | 5.88E-18 | injFB | MA0880.1 |
| 7.88140270 | 0.410927952456583 | 0.605 | 0.482 | 5.88E-18 | injFB | MA0881.1 |
| 6.41404895 | 0.409445502580562 | 0.602 | 0.493 | 4.78E-18 | injFB | MA0681.2 |
| 3.01890757 | 0.404012885952986 | 0.619 | 0.483 | 2.25E-21 | injFB | MA0079.4 |
| 1.11359848 | 0.401454037890978 | 0.653 | 0.491 | 8.31E-29 | injFB | MA1653.1 |
| 8.91899419 | 0.401020014582131 | 0.626 | 0.484 | 6.65E-20 | injFB | MA1584.1 |
| 1.94092580 | 0.400714768127699 | 0.614 | 0.492 | 1.45E-18 | injFB | MA0070.1 |
| 5.96605072 | 0.395536226691905 | 0.605 | 0.487 | 4.45E-19 | injFB | MA1607.1 |
| 1.93557817 | 0.392729829949179 | 0.612 | 0.484 | 1.44E-18 | injFB | MA1483.1 |
| 6.47393124 | 0.391544410844831 | 0.614 | 0.489 | 4.83E-18 | injFB | MA0778.1 |
| 5.16952047 | 0.388877694826338 | 0.6 | 0.493 | 3.86E-17 | injFB | MA0596.1 |
| 2.54991984 | 0.38814503768461 | 0.597 | 0.487 | 1.90E-16 | injFB | MA0872.1 |
| 2.14711052 | 0.385895251523369 | 0.627 | 0.483 | 1.60E-24 | injFB | MA0599.1 |
| 4.05465954 | 0.381957861971645 | 0.588 | 0.481 | 3.02E-15 | injFB | MA1554.1 |
| 4.21371608 | 0.379061277277565 | 0.598 | 0.485 | 3.14E-18 | injFB | MA0057.1 |
| 1.31297418 | 0.374766682607877 | 0.619 | 0.489 | 9.79E-18 | injFB | MA0645.1 |
| 6.74240772 | 0.369939962328146 | 0.595 | 0.488 | 5.03E-14 | injFB | MA0063.2 |
| 1.88188604 | 0.368824300608776 | 0.607 | 0.483 | 1.40E-15 | injFB | MA1476.1 |
| 4.70727086 | 0.364786024023422 | 0.6 | 0.487 | 3.51E-16 | injFB | MA0076.2 |
| 1.41744636 | 0.353960183994577 | 0.584 | 0.476 | 1.06E-13 | injFB | MA1545.1 |
| 8.16939604 | 0.343759686831321 | 0.577 | 0.474 | 6.09E-13 | injFB | MA1515.1 |
| 1.57975303 | 0.340848988552888 | 0.606 | 0.493 | 1.18E-14 | injFB | MA0845.1 |
| 2.94244494 | 0.339488411423608 | 0.6 | 0.484 | 2.20E-14 | injFB | MA1522.1 |
| 5.18841041 | 0.339304820931489 | 0.597 | 0.481 | 3.87E-17 | injFB | MA0039.4 |
| 3.52619133 | 0.334488657920814 | 0.582 | 0.48 | 2.63E-13 | injFB | MA0750.2 |
| 1.04361348 | 0.329581186588614 | 0.589 | 0.494 | 7.79E-12 | injFB | MA1103.2 |
| 2.30139490 | 0.327986563362601 | 0.608 | 0.491 | 1.72E-12 | injFB | MA0104.4 |
| 3.73410943 | 0.321869190506412 | 0.617 | 0.5 | 2.79E-13 | injFB | MA0655.1 |
| 1.05403191 | 0.321224993753721 | 0.611 | 0.493 | 7.86E-14 | injFB | MA1125.1 |
| 2.44901439 | 0.31757808638289 | 0.588 | 0.487 | 1.83E-11 | injFB | MA0488.1 |
| 6.43298862 | 0.313677596572929 | 0.593 | 0.488 | 4.80E-11 | injFB | MA0607.1 |
| 1.59559504 | 0.3128040438588 | 0.575 | 0.484 | 1.19E-08 | injFB | MA1552.1 |
| 1.06943435 | 0.31263470288263 | 0.575 | 0.489 | 7.98E-10 | injFB | MA0657.1 |
| 5.57837862 | 0.312389046883374 | 0.585 | 0.49 | 4.16E-09 | injFB | MA0083.3 |
| 1.73988938 | 0.311046200430108 | 0.59 | 0.491 | 1.30E-10 | injFB | MA0641.1 |

| | | | | | | |
|---|---|---|---|---|---|---|
| 4.82621620[illegible] | 0.308684106415362 | 0.584 | 0.492 | 3.60E-10 | injFB | MA0066.1 |
| 1.73315475[illegible] | 0.308613944294804 | 0.597 | 0.486 | 1.29E-13 | injFB | MA0741.1 |
| 1.59390786[illegible] | 0.306189156366931 | 0.576 | 0.488 | 1.19E-10 | injFB | MA0136.2 |
| 8.89030160[illegible] | 0.304630587529269 | 0.596 | 0.496 | 6.63E-12 | injFB | MA0462.2 |
| 2.41957642[illegible] | 0.298771471389411 | 0.577 | 0.491 | 1.81E-09 | injFB | MA1550.1 |
| 5.01938410[illegible] | 0.298433479449756 | 0.606 | 0.497 | 3.74E-11 | injFB | MA1634.1 |
| 3.82885296[illegible] | 0.29542852535809 | 0.586 | 0.488 | 2.86E-09 | injFB | MA1632.1 |
| 7.97812255[illegible] | 0.295295153547139 | 0.587 | 0.491 | 5.95E-09 | injFB | MA1657.1 |
| 1.15327369[illegible] | 0.293845906407025 | 0.585 | 0.49 | 8.60E-10 | injFB | MA0258.2 |
| 3.71606308[illegible] | 0.290318710926285 | 0.559 | 0.477 | 2.77E-07 | injFB | MA0882.1 |
| 1.85053834[illegible] | 0.288750219873838 | 0.586 | 0.482 | 1.38E-09 | injFB | MA1516.1 |
| 3.31847356[illegible] | 0.286049079156407 | 0.58 | 0.486 | 2.48E-08 | injFB | MA1131.1 |
| 1.08662262[illegible] | 0.284985626489513 | 0.57 | 0.486 | 8.11E-08 | injFB | MA0609.2 |
| 6.96759664[illegible] | 0.283226871972861 | 0.589 | 0.489 | 5.20E-09 | injFB | MA0829.2 |
| 5.53229045[illegible] | 0.282897322557858 | 0.578 | 0.49 | 4.13E-08 | injFB | MA1124.1 |
| 1.39467440[illegible] | 0.282362968448718 | 0.557 | 0.486 | 1.04E-06 | injFB | MA1594.1 |
| 1.59199222[illegible] | 0.281694159099235 | 0.581 | 0.496 | 1.19E-08 | injFB | MA0032.2 |
| 8.27320412[illegible] | 0.281508632646864 | 0.583 | 0.487 | 6.17E-09 | injFB | MA0751.1 |
| 6.24691556[illegible] | 0.278275544889895 | 0.561 | 0.483 | 4.66E-07 | injFB | MA1556.1 |
| 5.07839254[illegible] | 0.276226804892743 | 0.58 | 0.487 | 3.79E-07 | injFB | MA0840.1 |
| 4.07669235[illegible] | 0.275194597773888 | 0.57 | 0.5 | 3.04E-07 | injFB | MA1132.1 |
| 1.57365583[illegible] | 0.272947833244125 | 0.558 | 0.485 | 1.17E-05 | injFB | MA0666.1 |
| 1.57365583[illegible] | 0.272947833244125 | 0.558 | 0.485 | 1.17E-05 | injFB | MA0708.1 |
| 1.57365583[illegible] | 0.272947833244125 | 0.558 | 0.485 | 1.17E-05 | injFB | MA0876.1 |
| 9.48837187[illegible] | 0.271160724715166 | 0.574 | 0.482 | 7.08E-07 | injFB | MA0056.2 |
| 1.70368365[illegible] | 0.268066179169477 | 0.571 | 0.493 | 1.27E-07 | injFB | MA1683.1 |
| 5.39576429[illegible] | 0.264753025033812 | 0.564 | 0.482 | 4.03E-06 | injFB | MA1534.1 |
| 1.11244974[illegible] | 0.262726563159783 | 0.578 | 0.494 | 8.30E-06 | injFB | MA0855.1 |
| 5.46889756[illegible] | 0.261348895343618 | 0.586 | 0.498 | 4.08E-08 | injFB | MA0835.2 |
| 5.13537773[illegible] | 0.260544642041717 | 0.582 | 0.492 | 3.83E-07 | injFB | MA0769.2 |
| 2.10535600[illegible] | 0.260471383395308 | 0.581 | 0.49 | 1.57E-06 | injFB | MA0525.2 |
| 6.84066637[illegible] | 0.260122170779429 | 0.574 | 0.488 | 5.10E-07 | injFB | MA1518.1 |
| 7.99210109[illegible] | 0.258614088626526 | 0.574 | 0.488 | 5.96E-07 | injFB | MA0492.1 |
| 7.24546630[illegible] | 0.257671754828062 | 0.599 | 0.495 | 5.41E-08 | injFB | MA1651.1 |
| 8.14019731[illegible] | 0.256495220098969 | 0.592 | 0.503 | 6.07E-07 | injFB | MA0472.2 |
| 4.94691421[illegible] | 0.256385333635223 | 0.583 | 0.499 | 3.69E-07 | injFB | MA0491.2 |
| 2.22508193[illegible] | 0.254608602624682 | 0.563 | 0.484 | 1.66E-06 | injFB | MA1105.2 |
| 2.20205534[illegible] | 0.252292273495176 | 0.571 | 0.49 | 1.64E-07 | injFB | MA0496.3 |
| 5.47357028[illegible] | 0.251703754414836 | 0.543 | 0.485 | 4.08E-05 | injFB | MA0700.2 |
| 1.20053169[illegible] | 0.251604110502198 | 0.56 | 0.489 | 8.96E-06 | injFB | MA1126.1 |
| 6.91505874[illegible] | 0.251418597910866 | 0.58 | 0.493 | 5.16E-07 | injFB | MA0503.1 |
| 2.05419072[illegible] | 0.25054157301681 | 0.551 | 0.487 | 1.53E-05 | injFB | MA1539.1 |
| 7.09708393[illegible] | 0.735616651272794 | 0.746 | 0.489 | 5.29E-170 | PER | MA0761.2 |

| | | | | | | |
|---|---|---|---|---|---|---|
| 2.24805493 | 0.677633550316843 | 0.706 | 0.479 | 1.68E-132 | PER | MA0770.1 |
| 1.02765172 | 0.673274127041109 | 0.707 | 0.478 | 7.67E-129 | PER | MA0801.1 |
| 1.07717883 | 0.666274905657308 | 0.715 | 0.488 | 8.04E-133 | PER | MA0868.2 |
| 2.01188121 | 0.665707962948205 | 0.724 | 0.498 | 1.50E-135 | PER | MA0065.2 |
| 4.89669343 | 0.660279227926744 | 0.694 | 0.478 | 3.65E-119 | PER | MA1567.1 |
| 8.76325538 | 0.657904298768779 | 0.707 | 0.488 | 6.54E-123 | PER | MA1491.1 |
| 1.77649629 | 0.656040919265072 | 0.679 | 0.465 | 1.33E-110 | PER | MA0711.1 |
| 2.66639057 | 0.654521859600932 | 0.701 | 0.487 | 1.99E-123 | PER | MA0734.2 |
| 1.03657227 | 0.654106995411837 | 0.713 | 0.492 | 7.73E-131 | PER | MA0514.1 |
| 2.84569307 | 0.642627894663018 | 0.714 | 0.483 | 2.12E-132 | PER | MA0764.2 |
| 2.23485616 | 0.610881962111093 | 0.691 | 0.487 | 1.67E-108 | PER | MA1152.1 |
| 5.45982282 | 0.60945256484036 | 0.686 | 0.483 | 4.07E-109 | PER | MA0484.2 |
| 5.12402680 | 0.603330249404271 | 0.668 | 0.481 | 3.82E-94 | PER | MA0807.1 |
| 6.27659771 | 0.599555432203962 | 0.672 | 0.485 | 4.68E-96 | PER | MA0486.2 |
| 1.46671185 | 0.595146169300996 | 0.678 | 0.492 | 1.09E-90 | PER | MA0756.1 |
| 1.35886924 | 0.586038142109528 | 0.671 | 0.484 | 1.01E-90 | PER | MA0771.1 |
| 2.01433457 | 0.58376453703694 | 0.696 | 0.484 | 1.50E-106 | PER | MA0473.3 |
| 1.29161605 | 0.577352837593682 | 0.698 | 0.492 | 9.64E-104 | PER | MA0062.3 |
| 5.56128410 | 0.57716350790244 | 0.69 | 0.481 | 4.15E-104 | PER | MA0598.3 |
| 3.67077544 | 0.573788497382892 | 0.669 | 0.483 | 2.74E-90 | PER | MA0645.1 |
| 7.35030469 | 0.560428672560783 | 0.66 | 0.491 | 5.48E-84 | PER | MA0048.2 |
| 1.11504615 | 0.552553660067675 | 0.672 | 0.485 | 8.32E-81 | PER | MA0679.2 |
| 5.30610081 | 0.546101869555431 | 0.662 | 0.479 | 3.96E-78 | PER | MA0751.1 |
| 3.05441020 | 0.541049992075794 | 0.648 | 0.475 | 2.28E-75 | PER | MA0696.1 |
| 1.78874726 | 0.537951056130876 | 0.667 | 0.484 | 1.33E-76 | PER | MA0474.2 |
| 1.39799808 | 0.536942029032462 | 0.666 | 0.485 | 1.04E-80 | PER | MA0689.1 |
| 4.60743293 | 0.527284515715995 | 0.657 | 0.477 | 3.44E-85 | PER | MA0640.2 |
| 9.42019841 | 0.523511113671285 | 0.648 | 0.48 | 7.03E-69 | PER | MA0690.1 |
| 5.01720824 | 0.522677024479485 | 0.666 | 0.487 | 3.74E-79 | PER | MA0114.4 |
| 9.43565284 | 0.505675604548616 | 0.663 | 0.485 | 7.04E-77 | PER | MA1615.1 |
| 1.15809925 | 0.503993717252285 | 0.64 | 0.486 | 8.64E-70 | PER | MA0803.1 |
| 2.72767308 | 0.501089415387155 | 0.659 | 0.507 | 2.03E-62 | PER | MA0093.3 |
| 4.07588267 | 0.493729878042986 | 0.643 | 0.495 | 3.04E-63 | PER | MA0797.1 |
| 6.70496159 | 0.485958629973236 | 0.647 | 0.488 | 5.00E-65 | PER | MA0515.1 |
| 2.61462749 | 0.485106596101424 | 0.648 | 0.487 | 1.95E-61 | PER | MA0845.1 |
| 1.81006221 | 0.48261568554957 | 0.649 | 0.497 | 1.35E-60 | PER | MA0831.2 |
| 2.85380670 | 0.480401577838279 | 0.661 | 0.489 | 2.13E-71 | PER | MA0041.1 |
| 1.93382599 | 0.479054115700613 | 0.639 | 0.491 | 1.44E-61 | PER | MA1421.1 |
| 4.83790699 | 0.474583414775586 | 0.642 | 0.489 | 3.61E-58 | PER | MA1571.1 |
| 1.52787382 | 0.474431960826843 | 0.645 | 0.495 | 1.14E-57 | PER | MA0796.1 |
| 1.20317302 | 0.466372996387945 | 0.621 | 0.478 | 8.98E-55 | PER | MA1566.1 |
| 3.21467687 | 0.460843031310806 | 0.657 | 0.489 | 2.40E-61 | PER | MA0032.2 |
| 3.80267533 | 0.460530930084118 | 0.633 | 0.473 | 2.84E-55 | PER | MA1484.1 |

| | | | | | | |
|---|---|---|---|---|---|---|
| 4.35098566 | 0.459420102658411 | 0.641 | 0.49 | 3.25E-55 | PER | MA0608.1 |
| 6.28034177 | 0.457282459821142 | 0.622 | 0.485 | 4.69E-53 | PER | MA0806.1 |
| 1.14394659 | 0.454548059966986 | 0.631 | 0.479 | 8.53E-55 | PER | MA0744.2 |
| 6.82810332 | 0.449398098325962 | 0.65 | 0.49 | 5.09E-58 | PER | MA0846.1 |
| 2.59712776 | 0.447950117788111 | 0.616 | 0.48 | 1.94E-51 | PER | MA1584.1 |
| 9.88001964 | 0.446229221378622 | 0.626 | 0.481 | 7.37E-50 | PER | MA1559.1 |
| 2.02244987 | 0.445387738362525 | 0.624 | 0.478 | 1.51E-55 | PER | MA0002.2 |
| 4.97469962 | 0.443352763104695 | 0.637 | 0.495 | 3.71E-53 | PER | MA0597.1 |
| 4.00570393 | 0.440830670821322 | 0.615 | 0.48 | 2.99E-46 | PER | MA1483.1 |
| 4.19567828 | 0.435388239061035 | 0.606 | 0.476 | 3.13E-46 | PER | MA0103.3 |
| 7.73853763 | 0.433136255938319 | 0.625 | 0.489 | 5.77E-47 | PER | MA0768.1 |
| 6.05775356 | 0.432083409326472 | 0.615 | 0.484 | 4.52E-43 | PER | MA0800.1 |
| 6.61463861 | 0.430484105771871 | 0.62 | 0.483 | 4.93E-48 | PER | MA0607.1 |
| 3.26128475 | 0.424057253102519 | 0.615 | 0.483 | 2.43E-45 | PER | MA1558.1 |
| 2.20605847 | 0.418085252158164 | 0.61 | 0.483 | 1.65E-42 | PER | MA1546.1 |
| 5.14337882 | 0.417877423119555 | 0.634 | 0.485 | 3.84E-51 | PER | MA1102.2 |
| 8.90924631 | 0.417300973927375 | 0.621 | 0.487 | 6.65E-44 | PER | MA0697.1 |
| 7.59066223 | 0.414737234576748 | 0.629 | 0.485 | 5.66E-55 | PER | MA0081.2 |
| 2.57608942 | 0.414691227293048 | 0.629 | 0.496 | 1.92E-43 | PER | MA1525.1 |
| 3.95116256 | 0.413678409988385 | 0.621 | 0.486 | 2.95E-43 | PER | MA0475.2 |
| 1.29136174 | 0.413113143129721 | 0.621 | 0.484 | 9.63E-46 | PER | MA1621.1 |
| 2.40652413 | 0.409669445803097 | 0.617 | 0.496 | 1.80E-42 | PER | MA1529.1 |
| 1.38719296 | 0.409263659366909 | 0.594 | 0.477 | 1.03E-35 | PER | MA1554.1 |
| 7.30507613 | 0.408980477890682 | 0.599 | 0.482 | 5.45E-40 | PER | MA0802.1 |
| 3.63134177 | 0.408871530036346 | 0.617 | 0.493 | 2.71E-39 | PER | MA1572.1 |
| 1.21831758 | 0.406592960313124 | 0.636 | 0.481 | 9.09E-52 | PER | MA0162.4 |
| 4.72815611 | 0.40286660142556 | 0.619 | 0.501 | 3.53E-38 | PER | MA1154.1 |
| 9.83491639 | 0.397568240044743 | 0.635 | 0.496 | 7.34E-50 | PER | MA1596.1 |
| 8.70898893 | 0.395017413506029 | 0.603 | 0.479 | 6.50E-38 | PER | MA0139.1 |
| 6.93348273 | 0.393227128560971 | 0.617 | 0.486 | 5.17E-38 | PER | MA0156.2 |
| 4.66878143 | 0.391603360625905 | 0.634 | 0.495 | 3.48E-51 | PER | MA0080.5 |
| 4.34147745 | 0.38835005818491 | 0.619 | 0.493 | 3.24E-39 | PER | MA0851.1 |
| 2.58348773 | 0.388013318152798 | 0.607 | 0.491 | 1.93E-36 | PER | MA0624.1 |
| 1.59555555 | 0.387572060163005 | 0.617 | 0.501 | 1.19E-38 | PER | MA0783.1 |
| 6.81826208 | 0.386573377831063 | 0.619 | 0.493 | 5.09E-34 | PER | MA0780.1 |
| 6.54001010 | 0.382167227488482 | 0.618 | 0.495 | 4.88E-37 | PER | MA1635.1 |
| 7.23303507 | 0.372142334396902 | 0.6 | 0.482 | 5.40E-31 | PER | MA1553.1 |
| 6.38876171 | 0.368606532998395 | 0.612 | 0.49 | 4.77E-36 | PER | MA0523.1 |
| 7.81240809 | 0.368115732199038 | 0.572 | 0.482 | 5.83E-23 | PER | MA0116.1 |
| 1.65913739 | 0.359690670688214 | 0.633 | 0.495 | 1.24E-42 | PER | MA0149.1 |
| 5.10542315 | 0.359596559971839 | 0.6 | 0.484 | 3.81E-31 | PER | MA0076.2 |
| 2.78492708 | 0.357476465692688 | 0.602 | 0.487 | 2.08E-30 | PER | MA0641.1 |
| 2.45783659 | 0.35209286323588 | 0.605 | 0.49 | 1.83E-34 | PER | MA1125.1 |

| | | | | | | |
|---|---|---|---|---|---|---|
| 3.77839421 | 0.35181371244485 | 0.597 | 0.493 | 2.82E-28 | PER | MA0625.1 |
| 2.12726824 | 0.340983092907294 | 0.614 | 0.511 | 1.59E-26 | PER | MA0526.3 |
| 5.97825227 | 0.340750130040117 | 0.606 | 0.49 | 4.46E-29 | PER | MA0772.1 |
| 5.51030467 | 0.339495167755045 | 0.6 | 0.489 | 4.11E-28 | PER | MA0124.2 |
| 1.55394969 | 0.339451309245888 | 0.585 | 0.485 | 1.16E-26 | PER | MA0113.3 |
| 6.85275366 | 0.339333876515319 | 0.591 | 0.481 | 5.11E-27 | PER | MA1648.1 |
| 6.92203337 | 0.338721477309371 | 0.618 | 0.493 | 5.16E-31 | PER | MA0504.1 |
| 3.36789191 | 0.335598482912778 | 0.58 | 0.483 | 2.51E-24 | PER | MA0883.1 |
| 1.12984477 | 0.330675300219312 | 0.605 | 0.496 | 8.43E-28 | PER | MA1122.1 |
| 2.49292279 | 0.327950139786952 | 0.615 | 0.498 | 1.86E-27 | PER | MA0627.2 |
| 3.66609422 | 0.326992026299086 | 0.59 | 0.485 | 2.73E-25 | PER | MA0724.1 |
| 1.33678095 | 0.32386032761016 | 0.596 | 0.49 | 9.97E-25 | PER | MA0743.2 |
| 4.35453352 | 0.322947092669641 | 0.567 | 0.478 | 3.25E-23 | PER | MA0594.2 |
| 7.83247775 | 0.32279636638127 | 0.609 | 0.496 | 5.84E-29 | PER | MA1142.1 |
| 3.65722870 | 0.321970128741388 | 0.595 | 0.484 | 2.73E-28 | PER | MA1508.1 |
| 3.65856644 | 0.320409663941488 | 0.608 | 0.49 | 2.73E-28 | PER | MA0847.2 |
| 8.03781861 | 0.320216902449682 | 0.589 | 0.487 | 6.00E-23 | PER | MA0688.1 |
| 3.54829396 | 0.320162817536388 | 0.571 | 0.478 | 2.65E-22 | PER | MA0750.2 |
| 1.91429385 | 0.318915457590203 | 0.596 | 0.497 | 1.43E-25 | PER | MA1132.1 |
| 2.75857238 | 0.318825556332319 | 0.581 | 0.48 | 2.06E-23 | PER | MA0777.1 |
| 2.22155590 | 0.317713066898538 | 0.596 | 0.5 | 1.66E-23 | PER | MA0100.3 |
| 1.23148407 | 0.308980403847203 | 0.6 | 0.496 | 9.19E-23 | PER | MA0865.1 |
| 7.87273545 | 0.308297271923582 | 0.59 | 0.492 | 5.87E-21 | PER | MA1575.1 |
| 1.05714255 | 0.307104041157929 | 0.575 | 0.481 | 7.89E-21 | PER | MA1146.1 |
| 6.22998759 | 0.305898359189626 | 0.582 | 0.494 | 4.65E-21 | PER | MA0467.1 |
| 5.53962517 | 0.305460397920288 | 0.595 | 0.495 | 4.13E-19 | PER | MA0067.1 |
| 8.94941596 | 0.303285641070317 | 0.588 | 0.49 | 6.68E-21 | PER | MA0098.3 |
| 1.15345668 | 0.302257545391572 | 0.583 | 0.484 | 8.60E-21 | PER | MA0522.3 |
| 8.01605584 | 0.30157518484385 | 0.584 | 0.488 | 5.98E-22 | PER | MA1587.1 |
| 2.67733024 | 0.298672136083446 | 0.594 | 0.495 | 2.00E-21 | PER | MA1588.1 |
| 1.14003121 | 0.298076027964194 | 0.571 | 0.482 | 8.50E-20 | PER | MA0745.2 |
| 1.18107278 | 0.294301386067052 | 0.592 | 0.495 | 8.81E-21 | PER | MA1120.1 |
| 5.58646437 | 0.294298520070607 | 0.597 | 0.502 | 4.17E-24 | PER | MA0787.1 |
| 3.95142624 | 0.293636995155721 | 0.573 | 0.488 | 2.95E-22 | PER | MA1652.1 |
| 9.73403983 | 0.29293492931003 | 0.596 | 0.498 | 7.26E-22 | PER | MA0143.4 |
| 2.38850133 | 0.290885111326352 | 0.582 | 0.49 | 1.78E-20 | PER | MA0760.1 |
| 8.73796562 | 0.289794668668254 | 0.588 | 0.497 | 6.52E-20 | PER | MA1099.2 |
| 2.11584402 | 0.28389195818504 | 0.565 | 0.491 | 1.58E-17 | PER | MA0805.1 |
| 6.57276659 | 0.283748551906947 | 0.583 | 0.497 | 4.90E-19 | PER | MA0087.1 |
| 6.18599821 | 0.281541812056577 | 0.577 | 0.49 | 4.61E-17 | PER | MA0131.2 |
| 5.74616614 | 0.281050938115326 | 0.578 | 0.493 | 4.29E-17 | PER | MA1565.1 |
| 6.14818764 | 0.280652054219193 | 0.577 | 0.489 | 4.59E-17 | PER | MA1657.1 |
| 5.93416003 | 0.278721283451197 | 0.58 | 0.493 | 4.43E-18 | PER | MA0759.1 |

| | | | | | | |
|---|---|---|---|---|---|---|
| 1.13134262 | 0.276147309564526 | 0.578 | 0.491 | 8.44E-19 | PER | MA1620.1 |
| 6.90247192 | 0.27595365145823 | 0.573 | 0.486 | 5.15E-17 | PER | MA0091.1 |
| 1.62366174 | 0.275847846579194 | 0.566 | 0.489 | 1.21E-16 | PER | MA1471.1 |
| 1.18652919 | 0.26826347857159 | 0.575 | 0.491 | 8.85E-17 | PER | MA0155.1 |
| 3.68057855 | 0.268176800912834 | 0.567 | 0.482 | 2.75E-14 | PER | MA1552.1 |
| 6.98396287 | 0.268003761585978 | 0.595 | 0.485 | 5.21E-24 | PER | MA0050.2 |
| 1.92751071 | 0.266137935701391 | 0.585 | 0.491 | 1.44E-16 | PER | MA0069.1 |
| 9.11433603 | 0.265732869581754 | 0.574 | 0.484 | 6.80E-16 | PER | MA1562.1 |
| 2.82373555 | 0.265580492164438 | 0.588 | 0.497 | 2.11E-18 | PER | MA0517.1 |
| 1.10516128 | 0.262757538460934 | 0.557 | 0.487 | 8.24E-16 | PER | MA0520.1 |
| 4.59368227 | 0.259789014063024 | 0.566 | 0.497 | 3.43E-14 | PER | MA0695.1 |
| 2.43854611 | 0.258279343373915 | 0.577 | 0.495 | 1.82E-15 | PER | MA0077.1 |
| 1.60091411 | 0.255814002857887 | 0.561 | 0.493 | 1.19E-15 | PER | MA0698.1 |
| 7.82730395 | 0.252655465987887 | 0.581 | 0.497 | 5.84E-16 | PER | MA1418.1 |
| 1.53175768 | 0.251681420222198 | 0.566 | 0.492 | 1.14E-12 | PER | MA1420.1 |
| 2.45499462 | 0.250167019994487 | 0.569 | 0.487 | 1.83E-14 | PER | MA0767.1 |
| 6.91180175 | 0.616646971743035 | 0.658 | 0.477 | 5.16E-76 | retiFB | MA0116.1 |
| 9.01001886 | 0.607842264830627 | 0.673 | 0.479 | 6.72E-87 | retiFB | MA0742.1 |
| 1.56717592 | 0.581762895253803 | 0.654 | 0.469 | 1.17E-80 | retiFB | MA1533.1 |
| 2.97084998 | 0.579581963052594 | 0.668 | 0.475 | 2.22E-81 | retiFB | MA0493.1 |
| 3.55142921 | 0.573196770491797 | 0.654 | 0.467 | 2.65E-88 | retiFB | MA1515.1 |
| 1.79551935 | 0.555977210874797 | 0.638 | 0.47 | 1.34E-68 | retiFB | MA0882.1 |
| 6.48632095 | 0.545903137001929 | 0.639 | 0.476 | 4.84E-65 | retiFB | MA0880.1 |
| 6.48632095 | 0.545903137001929 | 0.639 | 0.476 | 4.84E-65 | retiFB | MA0881.1 |
| 1.52136108 | 0.537890213462436 | 0.653 | 0.479 | 1.13E-68 | retiFB | MA1594.1 |
| 1.01653955 | 0.528012700371967 | 0.654 | 0.478 | 7.58E-69 | retiFB | MA1573.1 |
| 1.13750964 | 0.512333384043124 | 0.641 | 0.484 | 8.49E-60 | retiFB | MA0795.1 |
| 7.26151633 | 0.498948941782861 | 0.643 | 0.478 | 5.42E-59 | retiFB | MA1552.1 |
| 4.97338590 | 0.481454370012909 | 0.65 | 0.482 | 3.71E-64 | retiFB | MA0685.1 |
| 2.18233922 | 0.478217581733417 | 0.653 | 0.48 | 1.63E-59 | retiFB | MA0138.2 |
| 8.66948739 | 0.466421117351168 | 0.64 | 0.474 | 6.47E-56 | retiFB | MA1618.1 |
| 4.07862217 | 0.461434500046332 | 0.62 | 0.468 | 3.04E-52 | retiFB | MA1517.1 |
| 2.06219438 | 0.458125073722212 | 0.636 | 0.478 | 1.54E-60 | retiFB | MA0079.4 |
| 2.06252747 | 0.438537090247064 | 0.626 | 0.48 | 1.54E-46 | retiFB | MA0691.1 |
| 6.32742158 | 0.435505297411864 | 0.614 | 0.481 | 4.72E-45 | retiFB | MA1578.1 |
| 3.00125639 | 0.429593958183861 | 0.621 | 0.484 | 2.24E-45 | retiFB | MA0790.1 |
| 2.69808669 | 0.428441228369728 | 0.621 | 0.481 | 2.01E-47 | retiFB | MA0057.1 |
| 5.74038003 | 0.421788113304011 | 0.615 | 0.479 | 4.28E-42 | retiFB | MA0139.1 |
| 1.79319466 | 0.420790397162672 | 0.61 | 0.484 | 1.34E-37 | retiFB | MA1470.1 |
| 4.50441409 | 0.40531098946394 | 0.614 | 0.484 | 3.36E-38 | retiFB | MA0063.2 |
| 7.18703199 | 0.396126553108333 | 0.602 | 0.482 | 5.36E-37 | retiFB | MA0037.3 |
| 3.16907527 | 0.395542454433991 | 0.609 | 0.478 | 2.36E-37 | retiFB | MA1556.1 |
| 2.59582530 | 0.392926427270691 | 0.606 | 0.48 | 1.94E-34 | retiFB | MA0666.1 |

| | | | | | | |
|---|---|---|---|---|---|---|
| 2.59582530 | 0.392926427270691 | 0.606 | 0.48 | 1.94E-34 | retiFB | MA0708.1 |
| 2.59582530 | 0.392926427270691 | 0.606 | 0.48 | 1.94E-34 | retiFB | MA0876.1 |
| 1.19884085 | 0.386858537185877 | 0.607 | 0.478 | 8.94E-38 | retiFB | MA1516.1 |
| 3.52675332 | 0.384211649791934 | 0.606 | 0.487 | 2.63E-34 | retiFB | MA0602.1 |
| 3.46215217 | 0.380649834031618 | 0.601 | 0.48 | 2.58E-31 | retiFB | MA1549.1 |
| 6.31404398 | 0.371346296032244 | 0.6 | 0.489 | 4.71E-31 | retiFB | MA0105.4 |
| 4.18117230 | 0.368473735238009 | 0.609 | 0.482 | 3.12E-36 | retiFB | MA0746.2 |
| 6.94374530 | 0.365450365483773 | 0.614 | 0.491 | 5.18E-33 | retiFB | MA0465.2 |
| 1.82501475 | 0.355571469221496 | 0.602 | 0.485 | 1.36E-29 | retiFB | MA1641.1 |
| 1.26152256 | 0.350286499451585 | 0.604 | 0.483 | 9.41E-30 | retiFB | MA0498.2 |
| 4.49484726 | 0.349593220894801 | 0.603 | 0.486 | 3.35E-28 | retiFB | MA0258.2 |
| 4.19893794 | 0.349175629292355 | 0.598 | 0.488 | 3.13E-29 | retiFB | MA0066.1 |
| 3.42205132 | 0.347211603351327 | 0.592 | 0.487 | 2.55E-26 | retiFB | MA1121.1 |
| 9.89152092 | 0.3446447893036 | 0.59 | 0.486 | 7.38E-27 | retiFB | MA1619.1 |
| 7.50060275 | 0.341642732772567 | 0.582 | 0.478 | 5.60E-24 | retiFB | MA1568.1 |
| 8.71237118 | 0.340186644080634 | 0.602 | 0.488 | 6.50E-27 | retiFB | MA1657.1 |
| 1.57696094 | 0.338125913894464 | 0.586 | 0.484 | 1.18E-24 | retiFB | MA1645.1 |
| 6.17918000 | 0.334998043959824 | 0.579 | 0.48 | 4.61E-25 | retiFB | MA0019.1 |
| 5.89563639 | 0.328771237294503 | 0.596 | 0.483 | 4.40E-26 | retiFB | MA0163.1 |
| 2.06525274 | 0.325701254217389 | 0.603 | 0.489 | 1.54E-25 | retiFB | MA1113.2 |
| 4.29806985 | 0.322582481540878 | 0.58 | 0.482 | 3.21E-26 | retiFB | MA1512.1 |
| 5.46443620 | 0.316258226631954 | 0.592 | 0.48 | 4.08E-23 | retiFB | MA0700.2 |
| 9.20276740 | 0.316003416005711 | 0.584 | 0.487 | 6.87E-23 | retiFB | MA1600.1 |
| 1.05294469 | 0.315969877623743 | 0.593 | 0.486 | 7.85E-23 | retiFB | MA0770.1 |
| 1.01137253 | 0.315489939413905 | 0.585 | 0.485 | 7.54E-24 | retiFB | MA0684.2 |
| 9.59229802 | 0.312909869083352 | 0.586 | 0.481 | 7.16E-23 | retiFB | MA0509.2 |
| 6.01807606 | 0.31156441635454 | 0.585 | 0.49 | 4.49E-20 | retiFB | MA0117.2 |
| 3.20229527 | 0.305670175538064 | 0.57 | 0.477 | 2.39E-18 | retiFB | MA1484.1 |
| 7.86476112 | 0.300014229226359 | 0.577 | 0.484 | 5.87E-19 | retiFB | MA0899.1 |
| 4.43896883 | 0.29871772091065 | 0.574 | 0.483 | 3.31E-21 | retiFB | MA0747.1 |
| 6.63587534 | 0.292904289510364 | 0.578 | 0.486 | 4.95E-18 | retiFB | MA0091.1 |
| 5.78578815 | 0.283041134970635 | 0.578 | 0.489 | 4.32E-20 | retiFB | MA1102.2 |
| 3.32767388 | 0.282070858115571 | 0.585 | 0.491 | 2.48E-17 | retiFB | MA0789.1 |
| 4.41817559 | 0.281714122984709 | 0.56 | 0.474 | 3.30E-16 | retiFB | MA1545.1 |
| 3.59392586 | 0.277951207502163 | 0.578 | 0.485 | 2.68E-18 | retiFB | MA0740.1 |
| 1.57979194 | 0.277707593260123 | 0.574 | 0.487 | 1.18E-16 | retiFB | MA0778.1 |
| 6.70329393 | 0.275333282407185 | 0.585 | 0.495 | 5.00E-18 | retiFB | MA0729.1 |
| 4.73345442 | 0.271118643942264 | 0.55 | 0.487 | 3.53E-13 | retiFB | MA0873.1 |
| 5.21801176 | 0.270656499741543 | 0.56 | 0.478 | 3.89E-14 | retiFB | MA0615.1 |
| 6.94425225 | 0.269622066721153 | 0.562 | 0.48 | 5.18E-16 | retiFB | MA1511.1 |
| 1.68301049 | 0.267732733612307 | 0.566 | 0.486 | 1.26E-14 | retiFB | MA0842.2 |
| 1.27403019 | 0.265611501336832 | 0.562 | 0.483 | 9.50E-14 | retiFB | MA0678.1 |
| 8.40435083 | 0.265242172965577 | 0.565 | 0.485 | 6.27E-14 | retiFB | MA1523.1 |

| | | | | | | |
|---|---|---|---|---|---|---|
| 6.25418087 | 0.264293010634128 | 0.565 | 0.488 | 4.67E-15 | retiFB | MA0901.2 |
| 8.57885219 | 0.263847427881234 | 0.569 | 0.483 | 6.40E-14 | retiFB | MA0511.2 |
| 6.26242000 | 0.263325523535242 | 0.564 | 0.481 | 4.67E-13 | retiFB | MA0739.1 |
| 1.83631222 | 0.263313007202125 | 0.566 | 0.482 | 1.37E-14 | retiFB | MA1532.1 |
| 1.17945589 | 0.262111644919282 | 0.561 | 0.487 | 8.80E-15 | retiFB | MA0775.1 |
| 7.17866516 | 0.257227019017319 | 0.579 | 0.49 | 5.36E-15 | retiFB | MA0146.2 |
| 1.59062566 | 0.252932376192438 | 0.579 | 0.479 | 1.19E-19 | retiFB | MA0039.4 |
| 7.85221872 | 0.500436078476949 | 0.618 | 0.474 | 5.86E-100 | myoFB | MA0742.1 |
| 8.49140168 | 0.471713896948894 | 0.622 | 0.471 | 6.33E-98 | myoFB | MA1573.1 |
| 1.40820527 | 0.471438875079906 | 0.617 | 0.473 | 1.05E-101 | myoFB | MA0163.1 |
| 6.81517160 | 0.460649029072298 | 0.609 | 0.469 | 5.08E-91 | myoFB | MA1534.1 |
| 4.85656637 | 0.453848845364424 | 0.601 | 0.471 | 3.62E-79 | myoFB | MA0880.1 |
| 4.85656637 | 0.453848845364424 | 0.601 | 0.471 | 3.62E-79 | myoFB | MA0881.1 |
| 7.39114500 | 0.419779102309405 | 0.586 | 0.474 | 5.51E-69 | myoFB | MA0666.1 |
| 7.39114500 | 0.419779102309405 | 0.586 | 0.474 | 5.51E-69 | myoFB | MA0708.1 |
| 7.39114500 | 0.419779102309405 | 0.586 | 0.474 | 5.51E-69 | myoFB | MA0876.1 |
| 7.47929690 | 0.419002001860181 | 0.609 | 0.473 | 5.58E-80 | myoFB | MA0057.1 |
| 1.05499328 | 0.417762501290134 | 0.604 | 0.47 | 7.87E-80 | myoFB | MA1516.1 |
| 1.67635219 | 0.416870701799596 | 0.615 | 0.475 | 1.25E-86 | myoFB | MA0740.1 |
| 4.03092977 | 0.414201962274381 | 0.595 | 0.472 | 3.01E-69 | myoFB | MA0493.1 |
| 2.69638024 | 0.413979831144436 | 0.607 | 0.475 | 2.01E-84 | myoFB | MA0746.2 |
| 1.69461235 | 0.412625968539949 | 0.601 | 0.483 | 1.26E-74 | myoFB | MA0681.2 |
| 3.29424840 | 0.411025927759095 | 0.605 | 0.477 | 2.46E-72 | myoFB | MA0657.1 |
| 4.22991993 | 0.406403635239903 | 0.606 | 0.472 | 3.16E-83 | myoFB | MA1512.1 |
| 9.18975432 | 0.402449114409346 | 0.596 | 0.475 | 6.86E-68 | myoFB | MA1131.1 |
| 1.35103346 | 0.392952889749515 | 0.604 | 0.468 | 1.01E-70 | myoFB | MA0750.2 |
| 3.03687914 | 0.387098799237434 | 0.593 | 0.473 | 2.27E-63 | myoFB | MA1476.1 |
| 6.03315613 | 0.374931469985981 | 0.583 | 0.475 | 4.50E-48 | myoFB | MA0116.1 |
| 2.13636244 | 0.368415292270223 | 0.591 | 0.476 | 1.59E-56 | myoFB | MA0609.2 |
| 3.06754966 | 0.355509270516241 | 0.614 | 0.478 | 2.29E-78 | myoFB | MA1513.1 |
| 2.67761405 | 0.354587445705301 | 0.572 | 0.475 | 2.00E-48 | myoFB | MA0637.1 |
| 5.17021044 | 0.350431765406156 | 0.597 | 0.478 | 3.86E-61 | myoFB | MA0685.1 |
| 3.27030049 | 0.347231398215638 | 0.571 | 0.464 | 2.44E-52 | myoFB | MA1517.1 |
| 2.67364566 | 0.34649895132397 | 0.58 | 0.476 | 1.99E-57 | myoFB | MA1522.1 |
| 5.70545471 | 0.346086871398091 | 0.588 | 0.474 | 4.26E-58 | myoFB | MA0079.4 |
| 6.69437199 | 0.344159798656562 | 0.593 | 0.478 | 4.99E-52 | myoFB | MA0076.2 |
| 9.21829564 | 0.343484111842185 | 0.579 | 0.474 | 6.88E-49 | myoFB | MA0673.1 |
| 3.28604090 | 0.342416805079121 | 0.572 | 0.475 | 2.45E-46 | myoFB | MA0739.1 |
| 1.45497823 | 0.340666936585259 | 0.58 | 0.478 | 1.09E-49 | myoFB | MA0498.2 |
| 4.42381549 | 0.337546274125037 | 0.573 | 0.478 | 3.30E-44 | myoFB | MA1594.1 |
| 2.42430920 | 0.334650478758316 | 0.571 | 0.465 | 1.81E-51 | myoFB | MA1515.1 |
| 4.85785753 | 0.333795474086288 | 0.588 | 0.481 | 3.62E-48 | myoFB | MA1121.1 |
| 4.73414828 | 0.329338456633415 | 0.59 | 0.477 | 3.53E-55 | myoFB | MA1650.1 |

| | | | | | | |
|---|---|---|---|---|---|---|
| 3.30454652 | 0.327994509676449 | 0.583 | 0.478 | 2.47E-47 | myoFB | MA0018. |
| 1.66797931 | 0.327336882418743 | 0.586 | 0.484 | 1.24E-46 | myoFB | MA0503. |
| 1.71358948 | 0.323708337869528 | 0.576 | 0.48 | 1.28E-40 | myoFB | MA0820. |
| 7.97781805 | 0.318737708878949 | 0.576 | 0.477 | 5.95E-47 | myoFB | MA0747. |
| 4.63144969 | 0.317715703688175 | 0.561 | 0.475 | 3.46E-37 | myoFB | MA0056. |
| 1.96253597 | 0.315796291713535 | 0.558 | 0.47 | 1.46E-34 | myoFB | MA0882. |
| 1.18095623 | 0.315359934097919 | 0.581 | 0.482 | 8.81E-45 | myoFB | MA0645. |
| 9.47901140 | 0.31347107747413 | 0.576 | 0.479 | 7.07E-50 | myoFB | MA1564. |
| 1.10851572 | 0.312999227206363 | 0.579 | 0.485 | 8.27E-40 | myoFB | MA1145. |
| 3.99210865 | 0.312916926544134 | 0.565 | 0.477 | 2.98E-37 | myoFB | MA0700. |
| 5.22836099 | 0.309794352744982 | 0.593 | 0.483 | 3.90E-57 | myoFB | MA1653. |
| 1.01158693 | 0.30489265143428 | 0.562 | 0.476 | 7.55E-35 | myoFB | MA0019. |
| 3.39534707 | 0.304463410541821 | 0.579 | 0.479 | 2.53E-43 | myoFB | MA1607. |
| 2.09133588 | 0.302534047440661 | 0.574 | 0.481 | 1.56E-37 | myoFB | MA1126. |
| 1.05449940 | 0.300221120892119 | 0.575 | 0.483 | 7.87E-39 | myoFB | MA1124. |
| 5.29787107 | 0.300162351591649 | 0.557 | 0.474 | 3.95E-34 | myoFB | MA1127. |
| 3.61365636 | 0.297425735389524 | 0.566 | 0.476 | 2.70E-35 | myoFB | MA0656. |
| 1.06360909 | 0.297036771869471 | 0.564 | 0.478 | 7.93E-37 | myoFB | MA1483. |
| 2.44124694 | 0.296166333709703 | 0.559 | 0.482 | 1.82E-33 | myoFB | MA1518. |
| 6.84179195 | 0.28865068214572 | 0.581 | 0.488 | 5.10E-36 | myoFB | MA0032. |
| 1.20023628 | 0.28602676532549 | 0.573 | 0.473 | 8.95E-45 | myoFB | MA0039. |
| 2.02753755 | 0.283076164427505 | 0.568 | 0.475 | 1.51E-38 | myoFB | MA1511. |
| 2.46619375 | 0.279160431298664 | 0.569 | 0.479 | 1.84E-33 | myoFB | MA0815. |
| 2.55437328 | 0.275335334100463 | 0.558 | 0.479 | 1.91E-28 | myoFB | MA1133. |
| 2.52235330 | 0.273939201026695 | 0.563 | 0.481 | 1.88E-31 | myoFB | MA0488. |
| 3.03349239 | 0.27100974228542 | 0.557 | 0.485 | 2.26E-31 | myoFB | MA0258. |
| 2.87813078 | 0.270974430199156 | 0.568 | 0.486 | 2.15E-30 | myoFB | MA0070. |
| 4.61943533 | 0.26900701111631 | 0.567 | 0.479 | 3.45E-32 | myoFB | MA0762. |
| 5.99042955 | 0.267623684450388 | 0.572 | 0.483 | 4.47E-31 | myoFB | MA0809. |
| 4.25999355 | 0.264780980411579 | 0.57 | 0.48 | 3.18E-36 | myoFB | MA0162. |
| 1.76776117 | 0.263717464172631 | 0.558 | 0.482 | 1.32E-26 | myoFB | MA0164. |
| 1.02057522 | 0.254857599073139 | 0.551 | 0.477 | 7.61E-24 | myoFB | MA1556. |
| 1.15784209 | 0.253837724826771 | 0.564 | 0.482 | 8.64E-27 | myoFB | MA0605. |
| 9.07406943 | 0.253069659538572 | 0.556 | 0.484 | 6.77E-26 | myoFB | MA0790. |
| 4.48273831 | 0.252518101386183 | 0.564 | 0.48 | 3.34E-26 | myoFB | MA0840. |
| 7.38113123 | 0.250589199375603 | 0.555 | 0.482 | 5.51E-25 | myoFB | MA0626. |
| 1.69973015 | 0.860981502464559 | 0.774 | 0.493 | 1.27E-70 | T | MA0508. |
| 5.46868956 | 0.853345843485257 | 0.793 | 0.486 | 4.08E-67 | T | MA0727. |
| 1.00910917 | 0.84508032346838 | 0.787 | 0.492 | 7.53E-72 | T | MA0501. |
| 4.58115674 | 0.770150188475965 | 0.767 | 0.483 | 3.42E-56 | T | MA0814. |
| 8.01095072 | 0.758865066687577 | 0.772 | 0.498 | 5.98E-56 | T | MA1494. |
| 1.63181355 | 0.739003133273961 | 0.747 | 0.481 | 1.22E-48 | T | MA0799. |
| 1.76702968 | 0.72131001559889 | 0.748 | 0.488 | 1.32E-45 | T | MA0907. |

| | | | | | | |
|---|---|---|---|---|---|---|
| 8.74635503 | 0.703529182333729 | 0.728 | 0.498 | 6.52E-44 | T | MA0524.2 |
| 1.82867509 | 0.700877498613904 | 0.714 | 0.483 | 1.36E-39 | T | MA0812.1 |
| 4.06458864 | 0.687353421041204 | 0.738 | 0.494 | 3.03E-41 | T | MA1565.1 |
| 3.81189557 | 0.682900254835537 | 0.714 | 0.49 | 2.84E-40 | T | MA0101.1 |
| 6.47416861 | 0.67708743017865 | 0.743 | 0.5 | 4.83E-46 | T | MA0062.3 |
| 1.00017033 | 0.670013252077077 | 0.728 | 0.486 | 7.46E-40 | T | MA0113.3 |
| 3.00632470 | 0.664808814250844 | 0.728 | 0.499 | 2.24E-45 | T | MA0149.1 |
| 9.83558879 | 0.66415062498601 | 0.714 | 0.494 | 7.34E-39 | T | MA1563.1 |
| 2.06734644 | 0.648014267193685 | 0.692 | 0.496 | 1.54E-35 | T | MA0003.4 |
| 5.37225500 | 0.624232756538594 | 0.714 | 0.494 | 4.01E-36 | T | MA0150.2 |
| 1.55256542 | 0.617623176796468 | 0.694 | 0.483 | 1.16E-31 | T | MA1105.2 |
| 8.80185725 | 0.616484946138052 | 0.723 | 0.491 | 6.57E-35 | T | MA0689.1 |
| 2.95127401 | 0.616157739300369 | 0.721 | 0.494 | 2.20E-36 | T | MA1647.1 |
| 4.50034096 | 0.606378768345712 | 0.653 | 0.477 | 3.36E-25 | T | MA0776.1 |
| 6.19252543 | 0.603210632718666 | 0.673 | 0.481 | 4.62E-27 | T | MA0007.3 |
| 2.38490456 | 0.597973106720737 | 0.721 | 0.5 | 1.78E-32 | T | MA1525.1 |
| 8.86514352 | 0.579830104769956 | 0.689 | 0.494 | 6.61E-28 | T | MA0811.1 |
| 9.46319323 | 0.572554935447009 | 0.701 | 0.493 | 7.06E-32 | T | MA0473.3 |
| 7.65926710 | 0.562507523632295 | 0.66 | 0.488 | 5.71E-26 | T | MA0892.1 |
| 3.00875837 | 0.546158920035464 | 0.677 | 0.492 | 2.24E-24 | T | MA1123.2 |
| 2.67860828 | 0.545298630921892 | 0.689 | 0.486 | 2.00E-25 | T | MA0802.1 |
| 1.99099640 | 0.541304017628837 | 0.69 | 0.497 | 1.49E-24 | T | MA1654.1 |
| 1.06552702 | 0.538324701430962 | 0.673 | 0.489 | 7.95E-26 | T | MA0810.1 |
| 5.29935371 | 0.529172850378338 | 0.66 | 0.49 | 3.95E-21 | T | MA0604.1 |
| 8.20951207 | 0.525042989169966 | 0.689 | 0.489 | 6.12E-25 | T | MA0089.2 |
| 4.25784157 | 0.519622832718445 | 0.66 | 0.489 | 3.18E-21 | T | MA0794.1 |
| 1.69011389 | 0.519531260838621 | 0.694 | 0.514 | 1.26E-20 | T | MA0526.3 |
| 6.67644519 | 0.51661022905798 | 0.689 | 0.495 | 4.98E-23 | T | MA0625.1 |
| 1.23079404 | 0.510006927669851 | 0.682 | 0.494 | 9.18E-23 | T | MA0698.1 |
| 2.88294542 | 0.508004734528402 | 0.653 | 0.496 | 2.15E-20 | T | MA0663.1 |
| 2.25888197 | 0.507091121001485 | 0.646 | 0.492 | 1.69E-18 | T | MA0066.1 |
| 4.48176022 | 0.505446135543064 | 0.668 | 0.484 | 3.34E-20 | T | MA0600.2 |
| 2.88516474 | 0.501769073654444 | 0.651 | 0.501 | 2.15E-20 | T | MA1616.1 |
| 7.63564997 | 0.501494566078618 | 0.651 | 0.498 | 5.70E-22 | T | MA1138.1 |
| 1.48099717 | 0.496107375933313 | 0.65 | 0.489 | 1.10E-18 | T | MA0639.1 |
| 1.07390346 | 0.492609662153594 | 0.687 | 0.5 | 8.01E-22 | T | MA0761.2 |
| 1.73340528 | 0.484839792903082 | 0.675 | 0.493 | 1.29E-20 | T | MA1143.1 |
| 9.27973915 | 0.48468729142988 | 0.662 | 0.49 | 6.92E-18 | T | MA0806.1 |
| 3.26393803 | 0.473842520074362 | 0.707 | 0.502 | 2.43E-21 | T | MA0152.1 |
| 1.74101084 | 0.47339281232258 | 0.655 | 0.502 | 1.30E-17 | T | MA0137.3 |
| 1.03303172 | 0.470473656048425 | 0.648 | 0.498 | 7.71E-17 | T | MA0159.1 |
| 7.94033126 | 0.469345780397713 | 0.636 | 0.491 | 5.92E-17 | T | MA1521.1 |
| 1.64431069 | 0.466676767981627 | 0.65 | 0.489 | 1.23E-16 | T | MA0843.1 |

| | | | | | | |
|---|---|---|---|---|---|---|
| 3.47484608 | 0.459306700658246 | 0.65 | 0.501 | 2.59E-15 | T | MA0884.1 |
| 1.79961108 | 0.45688504032701 | 0.687 | 0.499 | 1.34E-22 | T | MA0080.5 |
| 2.22304107 | 0.455405496550632 | 0.634 | 0.492 | 1.66E-16 | T | MA0803.1 |
| 1.40869327 | 0.452128620911496 | 0.653 | 0.495 | 1.05E-16 | T | MA0624.1 |
| 4.96873987 | 0.452124213839018 | 0.66 | 0.498 | 3.71E-16 | T | MA0863.1 |
| 8.21468882 | 0.446572936055124 | 0.656 | 0.498 | 6.13E-17 | T | MA1579.1 |
| 2.99681327 | 0.443581684751928 | 0.65 | 0.494 | 2.24E-15 | T | MA1520.1 |
| 2.22458638 | 0.430214529115933 | 0.65 | 0.499 | 1.66E-17 | T | MA1622.1 |
| 1.06889056 | 0.428100580514901 | 0.648 | 0.492 | 7.97E-14 | T | MA0715.1 |
| 2.24682890 | 0.427327628516388 | 0.633 | 0.493 | 1.68E-14 | T | MA0518.1 |
| 5.61728648 | 0.42496428756251 | 0.651 | 0.499 | 4.19E-17 | T | MA0528.2 |
| 2.72341745 | 0.422997316946472 | 0.631 | 0.49 | 2.03E-14 | T | MA0598.3 |
| 3.40323134 | 0.422608568503241 | 0.673 | 0.501 | 2.54E-17 | T | MA0476.1 |
| 7.15246877 | 0.420660720391377 | 0.641 | 0.484 | 5.34E-13 | T | MA0612.2 |
| 5.25935997 | 0.413665969574544 | 0.651 | 0.492 | 3.92E-15 | T | MA0146.2 |
| 1.86836242 | 0.405813144812666 | 0.631 | 0.491 | 1.39E-12 | T | MA1124.1 |
| 1.13930595 | 0.402273438309575 | 0.597 | 0.49 | 8.50E-11 | T | MA0059.1 |
| 3.49993560 | 0.399948710562102 | 0.616 | 0.484 | 2.61E-11 | T | MA1566.1 |
| 5.75085593 | 0.396578808713757 | 0.626 | 0.493 | 4.29E-11 | T | MA1683.1 |
| 9.26349694 | 0.394328391493504 | 0.599 | 0.489 | 6.91E-11 | T | MA0136.2 |
| 8.83925133 | 0.393141038877079 | 0.634 | 0.496 | 6.59E-14 | T | MA0489.1 |
| 1.05334087 | 0.392972096796051 | 0.604 | 0.484 | 7.86E-11 | T | MA0510.2 |
| 2.02817242 | 0.391678777403685 | 0.648 | 0.502 | 1.51E-10 | T | MA0620.3 |
| 1.41701535 | 0.391235215449219 | 0.634 | 0.492 | 1.06E-10 | T | MA0805.1 |
| 9.55103794 | 0.388654240906821 | 0.617 | 0.489 | 7.13E-11 | T | MA0145.3 |
| 2.57370097 | 0.386892584449799 | 0.624 | 0.486 | 1.92E-11 | T | MA1133.1 |
| 2.29747594 | 0.38626101930104 | 0.668 | 0.499 | 1.71E-15 | T | MA1627.1 |
| 1.02974666 | 0.384724933667786 | 0.616 | 0.49 | 7.68E-11 | T | MA0781.1 |
| 2.09827439 | 0.383851039479442 | 0.634 | 0.499 | 1.57E-11 | T | MA0463.2 |
| 6.61919215 | 0.383263047381151 | 0.629 | 0.499 | 4.94E-12 | T | MA0835.2 |
| 1.79617317 | 0.382458741944815 | 0.628 | 0.498 | 1.34E-11 | T | MA1130.1 |
| 6.82042510 | 0.379510170951983 | 0.634 | 0.493 | 5.09E-12 | T | MA0155.1 |
| 4.77396119 | 0.378803110222095 | 0.634 | 0.491 | 3.56E-10 | T | MA0683.1 |
| 1.34400927 | 0.376182280128892 | 0.631 | 0.496 | 1.00E-11 | T | MA0606.1 |
| 1.05303544 | 0.374074186877732 | 0.631 | 0.499 | 7.86E-11 | T | MA1137.1 |
| 2.93914190 | 0.372834575617466 | 0.617 | 0.498 | 2.19E-09 | T | MA1527.1 |
| 4.74887651 | 0.372673152237484 | 0.609 | 0.49 | 3.54E-09 | T | MA0807.1 |
| 1.34204861 | 0.372236758188673 | 0.634 | 0.502 | 1.00E-09 | T | MA0519.1 |
| 5.62220506 | 0.370622833881707 | 0.622 | 0.498 | 4.19E-11 | T | MA1634.1 |
| 1.02450181 | 0.369540674499638 | 0.616 | 0.497 | 7.64E-11 | T | MA0462.2 |
| 2.49207830 | 0.368021090448936 | 0.611 | 0.495 | 1.86E-09 | T | MA1601.1 |
| 3.65225212 | 0.367843635544993 | 0.611 | 0.5 | 2.72E-09 | T | MA0494.1 |
| 1.52971563 | 0.366219676662492 | 0.597 | 0.492 | 1.14E-08 | T | MA0035.4 |

| | | | | | | |
|---|---|---|---|---|---|---|
| 4.25538790 | 0.365658403318197 | 0.616 | 0.501 | 3.17E-10 | T | MA0099.3 |
| 1.95215875 | 0.364496992449172 | 0.614 | 0.503 | 1.46E-08 | T | MA0636.1 |
| 9.45957720 | 0.364243032582421 | 0.611 | 0.495 | 7.06E-09 | T | MA1569.1 |
| 1.44192229 | 0.363626840516095 | 0.626 | 0.494 | 1.08E-10 | T | MA0768.1 |
| 6.48207037 | 0.360113520374609 | 0.634 | 0.497 | 4.84E-11 | T | MA0478.1 |
| 4.22582458 | 0.359412212345173 | 0.624 | 0.493 | 3.15E-09 | T | MA0124.2 |
| 4.32133454 | 0.35924503952058 | 0.6 | 0.485 | 3.22E-08 | T | MA0900.2 |
| 5.81397605 | 0.350181001669553 | 0.599 | 0.495 | 4.34E-08 | T | MA0592.3 |
| 3.38969572 | 0.34971636059548 | 0.614 | 0.499 | 2.53E-08 | T | MA1542.1 |
| 8.87057515 | 0.348247922021388 | 0.592 | 0.49 | 6.62E-08 | T | MA0800.1 |
| 5.04315201 | 0.347297176761528 | 0.604 | 0.499 | 3.76E-09 | T | MA1144.1 |
| 2.55835317 | 0.346635618643937 | 0.605 | 0.494 | 1.91E-07 | T | MA0107.1 |
| 5.04039035 | 0.346236589395689 | 0.599 | 0.501 | 3.76E-08 | T | MA1142.1 |
| 6.03512234 | 0.345369717939326 | 0.612 | 0.492 | 4.50E-08 | T | MA1475.1 |
| 1.36717824 | 0.344729040848767 | 0.607 | 0.489 | 1.02E-07 | T | MA1151.1 |
| 7.20115270 | 0.343107214927675 | 0.621 | 0.504 | 5.37E-09 | T | MA1655.1 |
| 1.38847926 | 0.342155768858484 | 0.609 | 0.495 | 1.04E-07 | T | MA0825.1 |
| 1.32382925 | 0.342005286777314 | 0.609 | 0.485 | 9.88E-09 | T | MA1633.1 |
| 4.52131495 | 0.339982476413893 | 0.624 | 0.497 | 3.37E-08 | T | MA1638.1 |
| 2.82505637 | 0.338548578952254 | 0.626 | 0.493 | 2.11E-09 | T | MA0131.2 |
| 4.98913793 | 0.337438173566795 | 0.605 | 0.495 | 3.72E-07 | T | MA0481.3 |
| 9.29149657 | 0.332560272595691 | 0.619 | 0.489 | 6.93E-08 | T | MA0605.2 |
| 4.28514919 | 0.331044883742182 | 0.602 | 0.488 | 3.20E-07 | T | MA1567.1 |
| 1.39190898 | 0.328574331005847 | 0.617 | 0.496 | 1.04E-07 | T | MA1651.1 |
| 2.11810908 | 0.328233954536729 | 0.605 | 0.494 | 1.58E-06 | T | MA1541.1 |
| 2.51414050 | 0.327147529041918 | 0.616 | 0.499 | 1.88E-08 | T | MA1580.1 |
| 1.13278548 | 0.326966053400079 | 0.602 | 0.499 | 8.45E-08 | T | MA0593.1 |
| 1.63855814 | 0.325874403798874 | 0.599 | 0.491 | 1.22E-06 | T | MA0688.1 |
| 4.09153248 | 0.325492627422852 | 0.607 | 0.503 | 3.05E-08 | T | MA0490.2 |
| 4.62204883 | 0.32486106305079 | 0.628 | 0.487 | 3.45E-08 | T | MA0522.3 |
| 5.35925699 | 0.321710228190604 | 0.619 | 0.504 | 4.00E-08 | T | MA1624.1 |
| 3.76716148 | 0.319468373541143 | 0.605 | 0.489 | 2.81E-06 | T | MA0895.1 |
| 6.98917928 | 0.317135711060764 | 0.607 | 0.487 | 5.21E-07 | T | MA1562.1 |
| 9.52796320 | 0.316734949322188 | 0.597 | 0.486 | 7.11E-07 | T | MA0691.1 |
| 9.65829441 | 0.313063554793681 | 0.609 | 0.489 | 7.21E-07 | T | MA0132.2 |
| 1.11988592 | 0.310481459104936 | 0.607 | 0.491 | 8.35E-07 | T | MA0611.1 |
| 3.96294525 | 0.306001515860028 | 0.583 | 0.491 | 2.96E-05 | T | MA0889.1 |
| 2.71765816 | 0.30483915859657 | 0.583 | 0.492 | 2.03E-05 | T | MA1657.1 |
| 4.60510773 | 0.304276531070334 | 0.616 | 0.5 | 3.44E-07 | T | MA0491.2 |
| 2.65818908 | 0.302473781418189 | 0.599 | 0.5 | 1.98E-06 | T | MA0480.1 |
| 1.10670922 | 0.301523272315403 | 0.605 | 0.496 | 8.26E-06 | T | MA0759.1 |
| 4.66175377 | 0.297973632502739 | 0.609 | 0.502 | 3.48E-06 | T | MA1134.1 |
| 7.48193750 | 0.291346917499621 | 0.578 | 0.497 | 5.58E-06 | T | MA0468.1 |

| | | | | | | |
|---|---|---|---|---|---|---|
| 4.46319652 | 0.289802360985028 | 0.588 | 0.501 | 3.33E-05 | T | MA0058.3 |
| 2.75571590 | 0.288987780886184 | 0.58 | 0.497 | 0.000205576 | T | MA0031.1 |
| 2.75571590 | 0.288987780886184 | 0.58 | 0.497 | 0.000205576 | T | MA0157.2 |
| 2.26705862 | 0.288269710906381 | 0.607 | 0.495 | 1.69E-06 | T | MA0515.1 |
| 6.46468363 | 0.286791608418433 | 0.582 | 0.486 | 4.82E-05 | T | MA0705.1 |
| 2.70390864 | 0.286211233840277 | 0.607 | 0.503 | 2.02E-06 | T | MA0477.2 |
| 8.13287391 | 0.283632064756469 | 0.6 | 0.5 | 6.07E-06 | T | MA1630.1 |
| 3.56519445 | 0.280421016113162 | 0.566 | 0.488 | 0.000265964 | T | MA0798.2 |
| 3.33214878 | 0.27741197158374 | 0.595 | 0.489 | 0.000248578 | T | MA1555.1 |
| 5.38941938 | 0.276510387653698 | 0.566 | 0.49 | 0.000402051 | T | MA0801.1 |
| 1.36334611 | 0.272241473707342 | 0.561 | 0.495 | 0.001017056 | T | MA0671.1 |
| 2.26758649 | 0.270303711765382 | 0.585 | 0.502 | 0.000169162 | T | MA0614.1 |
| 7.26608460 | 0.26947594844769 | 0.565 | 0.496 | 0.00054205 | T | MA0523.1 |
| 4.49936330 | 0.26505307056363 | 0.568 | 0.488 | 0.000335653 | T | MA0690.1 |
| 2.50226777 | 0.263530055803364 | 0.59 | 0.495 | 0.000186669 | T | MA0659.2 |
| 2.01480558 | 0.263469470562897 | 0.57 | 0.491 | 0.001503045 | T | MA1119.1 |
| 1.02317029 | 0.261834780767806 | 0.57 | 0.509 | 0.000763285 | T | MA0730.1 |
| 5.14819091 | 0.260342901436592 | 0.57 | 0.493 | 0.00384055 | T | MA0870.1 |
| 9.28614425 | 0.257159071760882 | 0.561 | 0.49 | 0.006927464 | T | MA0109.1 |
| 3.72248777 | 0.257017617397516 | 0.571 | 0.487 | 0.002776976 | T | MA1479.1 |
| 3.96561646 | 0.253034913368689 | 0.575 | 0.496 | 0.00295835 | T | MA0122.3 |
| 4.52240145 | 0.252873661087478 | 0.558 | 0.491 | 0.003373711 | T | MA0686.1 |
| 1.83718596 | 0.252707062008114 | 0.607 | 0.514 | 0.001370541 | T | MA0093.3 |
| 5.68567116 | 0.25234933472346 | 0.56 | 0.495 | 0.004241511 | T | MA1649.1 |
| 1.96347353 | 0.250702774922047 | 0.594 | 0.5 | 0.001464751 | T | MA0738.1 |
| 2.89299438 | 0.250444805468905 | 0.578 | 0.487 | 0.002158174 | T | MA1603.1 |
| 4.72369370 | 0.805055505041124 | 0.726 | 0.481 | 3.52E-76 | MUS | MA0612.2 |
| 8.57540297 | 0.782744322830123 | 0.758 | 0.494 | 6.40E-84 | MUS | MA0653.1 |
| 2.86012716 | 0.735784343501611 | 0.731 | 0.485 | 2.13E-69 | MUS | MA0892.1 |
| 3.13672191 | 0.719368853431056 | 0.694 | 0.488 | 2.34E-60 | MUS | MA0809.2 |
| 1.74566097 | 0.705206189197777 | 0.717 | 0.482 | 1.30E-61 | MUS | MA0705.1 |
| 2.69868476 | 0.660883697112816 | 0.695 | 0.484 | 2.01E-53 | MUS | MA0027.2 |
| 6.83310492 | 0.630742071449642 | 0.677 | 0.482 | 5.10E-46 | MUS | MA0900.2 |
| 1.02724811 | 0.62686197575837 | 0.676 | 0.492 | 7.66E-45 | MUS | MA0808.1 |
| 2.78516464 | 0.625895639795651 | 0.684 | 0.477 | 2.08E-46 | MUS | MA1484.1 |
| 6.52235579 | 0.606793954849507 | 0.677 | 0.491 | 4.87E-43 | MUS | MA0068.2 |
| 2.74264271 | 0.604109215756361 | 0.676 | 0.485 | 2.05E-41 | MUS | MA0893.2 |
| 2.74264271 | 0.604109215756361 | 0.676 | 0.485 | 2.05E-41 | MUS | MA0912.2 |
| 8.99201906 | 0.601272875493723 | 0.708 | 0.496 | 6.71E-51 | MUS | MA1418.1 |
| 2.20624458 | 0.59254104802209 | 0.695 | 0.496 | 1.65E-45 | MUS | MA0866.1 |
| 2.08821264 | 0.58557704202625 | 0.69 | 0.489 | 1.56E-40 | MUS | MA0654.1 |
| 2.08821264 | 0.58557704202625 | 0.69 | 0.489 | 1.56E-40 | MUS | MA0720.1 |
| 2.08821264 | 0.58557704202625 | 0.69 | 0.489 | 1.56E-40 | MUS | MA0701.2 |

| | | | | | | |
|---|---|---|---|---|---|---|
| 3.543816739 | 0.58109726167919 | 0.675 | 0.488 | 2.64E-40 | MUS | MA1592.1 |
| 1.323737427 | 0.574108109708434 | 0.663 | 0.489 | 9.88E-37 | MUS | MA0610.1 |
| 1.148922659 | 0.56625259983093 | 0.666 | 0.483 | 8.57E-38 | MUS | MA0600.2 |
| 2.248154651 | 0.565529254172685 | 0.675 | 0.495 | 1.68E-38 | MUS | MA0084.1 |
| 2.889090630 | 0.556042671324539 | 0.658 | 0.481 | 2.16E-35 | MUS | MA0125.1 |
| 5.689038942 | 0.552747582464707 | 0.658 | 0.49 | 4.24E-36 | MUS | MA0887.1 |
| 1.232931698 | 0.551297531458539 | 0.645 | 0.482 | 9.20E-34 | MUS | MA0736.1 |
| 3.683788990 | 0.539452031110058 | 0.65 | 0.48 | 2.75E-31 | MUS | MA0007.3 |
| 5.439966950 | 0.532543942819652 | 0.663 | 0.483 | 4.06E-33 | MUS | MA0647.1 |
| 1.039113420 | 0.520932601140144 | 0.646 | 0.486 | 7.75E-31 | MUS | MA0662.1 |
| 1.427159381 | 0.519821638511782 | 0.668 | 0.489 | 1.06E-31 | MUS | MA0630.1 |
| 5.382138854 | 0.518212952586573 | 0.657 | 0.486 | 4.02E-36 | MUS | MA0162.4 |
| 1.010884869 | 0.517779492531517 | 0.664 | 0.494 | 7.54E-35 | MUS | MA1153.1 |
| 4.756197551 | 0.501767468569566 | 0.65 | 0.481 | 3.55E-31 | MUS | MA0799.1 |
| 1.713561041 | 0.499484017706965 | 0.643 | 0.484 | 1.28E-27 | MUS | MA1578.1 |
| 1.001599479 | 0.49717686721439 | 0.651 | 0.497 | 7.47E-27 | MUS | MA1419.1 |
| 9.442941904 | 0.493391924408516 | 0.654 | 0.489 | 7.04E-30 | MUS | MA0090.3 |
| 7.592015730 | 0.482713555139095 | 0.65 | 0.487 | 5.66E-26 | MUS | MA0886.1 |
| 2.990016274 | 0.480012384017345 | 0.64 | 0.489 | 2.23E-27 | MUS | MA0710.1 |
| 1.319494221 | 0.479914084369636 | 0.622 | 0.489 | 9.84E-24 | MUS | MA1121.1 |
| 1.292256114 | 0.478392908057077 | 0.65 | 0.482 | 9.64E-27 | MUS | MA0510.2 |
| 9.952682513 | 0.474391278912607 | 0.667 | 0.491 | 7.42E-28 | MUS | MA0902.2 |
| 1.537153717 | 0.471715307285033 | 0.679 | 0.488 | 1.15E-32 | MUS | MA0516.2 |
| 9.462486885 | 0.466082898029283 | 0.638 | 0.482 | 7.06E-24 | MUS | MA0637.1 |
| 5.023622059 | 0.465562595497528 | 0.644 | 0.494 | 3.75E-25 | MUS | MA0773.1 |
| 1.067095472 | 0.463882051608931 | 0.661 | 0.489 | 7.96E-28 | MUS | MA0611.1 |
| 2.373591648 | 0.463648381279476 | 0.639 | 0.486 | 1.77E-23 | MUS | MA1577.1 |
| 2.911278783 | 0.463252900641927 | 0.65 | 0.489 | 2.17E-32 | MUS | MA1513.1 |
| 2.670947905 | 0.461401760893388 | 0.655 | 0.489 | 1.99E-28 | MUS | MA1652.1 |
| 1.328467527 | 0.4593882939123 | 0.629 | 0.485 | 9.91E-25 | MUS | MA0753.2 |
| 4.739748981 | 0.45061286704579 | 0.643 | 0.487 | 3.54E-22 | MUS | MA0725.1 |
| 4.739748981 | 0.45061286704579 | 0.643 | 0.487 | 3.54E-22 | MUS | MA0726.1 |
| 9.185368075 | 0.450330830052471 | 0.619 | 0.483 | 6.85E-23 | MUS | MA0509.2 |
| 5.638127805 | 0.439693250403653 | 0.616 | 0.473 | 4.21E-22 | MUS | MA1515.1 |
| 1.079080649 | 0.439314775671731 | 0.633 | 0.493 | 8.05E-22 | MUS | MA0793.1 |
| 2.978989708 | 0.438910644344849 | 0.644 | 0.488 | 2.22E-28 | MUS | MA0050.2 |
| 2.586699025 | 0.436222597495247 | 0.646 | 0.49 | 1.93E-25 | MUS | MA0471.2 |
| 7.821656277 | 0.431461463538849 | 0.588 | 0.474 | 5.83E-19 | MUS | MA0711.1 |
| 2.248882494 | 0.423824364984261 | 0.64 | 0.489 | 1.68E-20 | MUS | MA0028.2 |
| 2.212840474 | 0.419562097176426 | 0.624 | 0.493 | 1.65E-19 | MUS | MA0772.1 |
| 1.240574519 | 0.417193944285556 | 0.618 | 0.489 | 9.25E-20 | MUS | MA0078.1 |
| 1.163837354 | 0.413061069421995 | 0.605 | 0.476 | 8.68E-17 | MUS | MA0776.1 |
| 3.821826501 | 0.406205586144563 | 0.612 | 0.486 | 2.85E-16 | MUS | MA1509.1 |

| | | | | | | |
|---|---|---|---|---|---|---|
| 1.73015799 | 0.405069252736877 | 0.634 | 0.491 | 1.29E-18 | MUS | MA1117.1 |
| 7.18083898 | 0.399859406930147 | 0.61 | 0.487 | 5.36E-16 | MUS | MA0888.1 |
| 8.67882777 | 0.393960281046007 | 0.618 | 0.489 | 6.47E-18 | MUS | MA0781.1 |
| 2.90902431 | 0.392959887963455 | 0.61 | 0.491 | 2.17E-16 | MUS | MA0634.1 |
| 5.83810405 | 0.390541870472521 | 0.62 | 0.487 | 4.36E-20 | MUS | MA1564.1 |
| 1.21831912 | 0.387577658126283 | 0.608 | 0.489 | 9.09E-17 | MUS | MA1481.1 |
| 3.47253929 | 0.384738105295828 | 0.622 | 0.488 | 2.59E-15 | MUS | MA0063. |
| 1.93073160 | 0.383094459106768 | 0.587 | 0.486 | 1.44E-12 | MUS | MA1479.1 |
| 2.39787495 | 0.380376217340027 | 0.63 | 0.495 | 1.79E-15 | MUS | MA0894.1 |
| 1.13781514 | 0.379932666392465 | 0.606 | 0.489 | 8.49E-15 | MUS | MA0889.1 |
| 9.05075800 | 0.379101867961494 | 0.617 | 0.495 | 6.75E-17 | MUS | MA0759.1 |
| 2.61088396 | 0.37855682138938 | 0.609 | 0.487 | 1.95E-18 | MUS | MA1650.1 |
| 3.60720890 | 0.377510194561693 | 0.607 | 0.487 | 2.69E-15 | MUS | MA0908.1 |
| 1.55881218 | 0.374822906153776 | 0.617 | 0.488 | 1.16E-16 | MUS | MA0685.1 |
| 8.46721636 | 0.374035184945747 | 0.596 | 0.476 | 6.32E-13 | MUS | MA0882.1 |
| 2.66222177 | 0.369496552093592 | 0.597 | 0.484 | 1.99E-15 | MUS | MA0079. |
| 3.51911028 | 0.36510005561765 | 0.598 | 0.493 | 2.63E-13 | MUS | MA1420.1 |
| 4.43371078 | 0.364038214999139 | 0.61 | 0.494 | 3.31E-13 | MUS | MA0114. |
| 2.41958745 | 0.362512278637098 | 0.606 | 0.486 | 1.81E-15 | MUS | MA0690.1 |
| 4.12353458 | 0.356576994111747 | 0.62 | 0.495 | 3.08E-15 | MUS | MA0853.1 |
| 1.34748608 | 0.355369785187404 | 0.616 | 0.492 | 1.01E-13 | MUS | MA0699.1 |
| 4.57660049 | 0.354699826467715 | 0.597 | 0.488 | 3.41E-13 | MUS | MA0132. |
| 1.73383996 | 0.353075580673887 | 0.598 | 0.486 | 1.29E-13 | MUS | MA1559.1 |
| 1.16294040 | 0.352601396535083 | 0.59 | 0.487 | 8.68E-14 | MUS | MA0746. |
| 8.12244027 | 0.352377798860648 | 0.604 | 0.497 | 6.06E-14 | MUS | MA0869.1 |
| 2.96215230 | 0.35007724346066 | 0.604 | 0.49 | 2.21E-13 | MUS | MA1521.1 |
| 1.27307145 | 0.345808222560058 | 0.612 | 0.484 | 9.50E-18 | MUS | MA0599.1 |
| 8.49929311 | 0.34433327129523 | 0.621 | 0.49 | 6.34E-14 | MUS | MA0101.1 |
| 5.87615013 | 0.341414651285455 | 0.6 | 0.492 | 4.38E-12 | MUS | MA1475.1 |
| 6.22267765 | 0.339912644137467 | 0.592 | 0.493 | 4.64E-12 | MUS | MA0131. |
| 3.47538701 | 0.339511867890227 | 0.626 | 0.498 | 2.59E-13 | MUS | MA1630.1 |
| 3.19928503 | 0.33804200593141 | 0.611 | 0.492 | 2.39E-13 | MUS | MA1463.1 |
| 4.16125783 | 0.337790561955048 | 0.59 | 0.488 | 3.10E-10 | MUS | MA0113. |
| 8.00500139 | 0.337075483946697 | 0.614 | 0.493 | 5.97E-13 | MUS | MA0107.1 |
| 7.16548340 | 0.335904682195191 | 0.585 | 0.491 | 5.35E-12 | MUS | MA1471.1 |
| 9.54487933 | 0.333968461996502 | 0.604 | 0.489 | 7.12E-13 | MUS | MA0867. |
| 1.33384988 | 0.331877420994076 | 0.633 | 0.499 | 9.95E-17 | MUS | MA0149.1 |
| 4.12029669 | 0.331803053817652 | 0.58 | 0.49 | 3.07E-10 | MUS | MA0795.1 |
| 1.92274518 | 0.328065209898479 | 0.602 | 0.494 | 1.43E-10 | MUS | MA0845.1 |
| 3.51670730 | 0.324881360814347 | 0.587 | 0.488 | 2.62E-10 | MUS | MA0076. |
| 8.89983813 | 0.324766599550661 | 0.578 | 0.489 | 6.64E-10 | MUS | MA0644.1 |
| 3.58104548 | 0.324557488671457 | 0.587 | 0.486 | 2.67E-11 | MUS | MA0798. |
| 5.86426063 | 0.32248430465261 | 0.58 | 0.476 | 4.37E-10 | MUS | MA1545.1 |

| | | | | | | |
|---|---|---|---|---|---|---|
| 4.61559796 | 0.319755820124291 | 0.59 | 0.494 | 3.44E-10 | MUS | MA0658.1 |
| 3.74426303 | 0.31780206707079 | 0.589 | 0.49 | 2.79E-10 | MUS | MA0806.1 |
| 7.77932571 | 0.313577474379042 | 0.598 | 0.491 | 5.80E-11 | MUS | MA0591.1 |
| 6.19905224 | 0.313087636243328 | 0.581 | 0.48 | 4.62E-09 | MUS | MA1495.1 |
| 6.19905224 | 0.313087636243328 | 0.581 | 0.48 | 4.62E-09 | MUS | MA0904.2 |
| 3.83616274 | 0.312351620384751 | 0.586 | 0.487 | 2.86E-12 | MUS | MA0741.1 |
| 9.02404131 | 0.310241484693919 | 0.589 | 0.485 | 6.73E-10 | MUS | MA0744.2 |
| 3.17939926 | 0.308106555932181 | 0.609 | 0.492 | 2.37E-11 | MUS | MA0497.1 |
| 6.55570851 | 0.306038630362421 | 0.605 | 0.489 | 4.89E-09 | MUS | MA0874.1 |
| 4.71198756 | 0.301176048221585 | 0.582 | 0.491 | 3.52E-08 | MUS | MA0604.1 |
| 7.73905433 | 0.298595038681645 | 0.595 | 0.488 | 5.77E-09 | MUS | MA0145.3 |
| 1.26205430 | 0.297930857173917 | 0.587 | 0.494 | 9.41E-09 | MUS | MA0069.1 |
| 4.97690172 | 0.297515504712149 | 0.581 | 0.492 | 3.71E-08 | MUS | MA0484.2 |
| 1.48271752 | 0.292179878948556 | 0.595 | 0.487 | 1.11E-07 | MUS | MA1567.1 |
| 5.60625576 | 0.288091567294525 | 0.581 | 0.489 | 4.18E-08 | MUS | MA1535.1 |
| 2.12029480 | 0.287753441739334 | 0.595 | 0.493 | 1.58E-08 | MUS | MA0075.3 |
| 9.70998695 | 0.284536913743209 | 0.576 | 0.489 | 7.24E-09 | MUS | MA0830.2 |
| 3.21727812 | 0.283895100358277 | 0.577 | 0.489 | 2.40E-08 | MUS | MA0724.1 |
| 3.36400509 | 0.283527710073637 | 0.601 | 0.498 | 2.51E-08 | MUS | MA1542.1 |
| 1.57200345 | 0.282870617735517 | 0.577 | 0.5 | 1.17E-08 | MUS | MA0517.1 |
| 9.62397990 | 0.282259821374698 | 0.547 | 0.484 | 7.18E-07 | MUS | MA1566.1 |
| 1.37092377 | 0.281700617402345 | 0.605 | 0.5 | 1.02E-10 | MUS | MA0080.5 |
| 7.20034310 | 0.281479482267918 | 0.567 | 0.484 | 5.37E-07 | MUS | MA1105.2 |
| 7.24260435 | 0.281001491219398 | 0.575 | 0.488 | 5.40E-08 | MUS | MA0740.1 |
| 2.83453765 | 0.279524487714111 | 0.561 | 0.486 | 2.11E-06 | MUS | MA0903.1 |
| 7.06364554 | 0.276325494830572 | 0.579 | 0.49 | 5.27E-08 | MUS | MA0800.1 |
| 1.81647335 | 0.274397128478467 | 0.576 | 0.493 | 1.36E-07 | MUS | MA0689.1 |
| 9.27490103 | 0.274328090408355 | 0.599 | 0.493 | 6.92E-08 | MUS | MA0743.2 |
| 3.09858593 | 0.272698150262193 | 0.573 | 0.482 | 2.31E-09 | MUS | MA0039.4 |
| 7.43827598 | 0.263475991507656 | 0.563 | 0.489 | 5.55E-05 | MUS | MA0014.3 |
| 6.73012018 | 0.259025706563834 | 0.558 | 0.496 | 5.02E-06 | MUS | MA1565.1 |
| 1.53893527 | 0.258200105130761 | 0.59 | 0.488 | 1.15E-06 | MUS | MA0684.2 |
| 5.04613706 | 0.258091455000412 | 0.561 | 0.484 | 3.76E-06 | MUS | MA1538.1 |
| 8.08673779 | 0.256823424940897 | 0.57 | 0.488 | 6.03E-07 | MUS | MA1631.1 |
| 2.02120159 | 0.252808881005839 | 0.569 | 0.486 | 1.51E-05 | MUS | MA1594.1 |
| 1.33974103 | 0.250168571840902 | 0.584 | 0.497 | 9.99E-07 | MUS | MA0660.1 |
| 9.90828159 | 0.838622693445856 | 0.787 | 0.495 | 7.39E-55 | HS | MA1639.1 |
| 7.15951742 | 0.762965021343768 | 0.754 | 0.493 | 5.34E-44 | HS | MA1487.1 |
| 1.97702611 | 0.742606866644918 | 0.698 | 0.485 | 1.47E-35 | HS | MA0903.1 |
| 1.42710776 | 0.63282265710908 | 0.731 | 0.494 | 1.06E-28 | HS | MA1640.1 |
| 1.07293338 | 0.604987611693217 | 0.69 | 0.49 | 8.00E-24 | HS | MA0706.1 |
| 2.15753562 | 0.603044501228608 | 0.7 | 0.498 | 1.61E-24 | HS | MA1542.1 |
| 1.21558156 | 0.586623293150469 | 0.694 | 0.509 | 9.07E-27 | HS | MA0051.1 |

| | | | | | | |
|---|---|---|---|---|---|---|
| 3.39701772 | 0.566887854722929 | 0.669 | 0.491 | 2.53E-20 | HS | MA0618.1 |
| 1.52592312 | 0.563568267859747 | 0.688 | 0.494 | 1.14E-20 | HS | MA0658.1 |
| 2.85133941 | 0.558329609298417 | 0.704 | 0.504 | 2.13E-21 | HS | MA1624.1 |
| 2.50983900 | 0.555704072853518 | 0.673 | 0.496 | 1.87E-18 | HS | MA0894.1 |
| 8.69992186 | 0.533324214792599 | 0.644 | 0.498 | 6.49E-16 | HS | MA1419.1 |
| 1.08734765 | 0.519020071380812 | 0.677 | 0.498 | 8.11E-17 | HS | MA0875.1 |
| 5.77402639 | 0.507518071747241 | 0.702 | 0.496 | 4.31E-17 | HS | MA0852.2 |
| 9.23062200 | 0.496470725644838 | 0.644 | 0.483 | 6.89E-13 | HS | MA0125.1 |
| 7.63042645 | 0.486594918409357 | 0.646 | 0.501 | 5.69E-14 | HS | MA0108.2 |
| 1.42612347 | 0.486159889033632 | 0.619 | 0.491 | 1.06E-12 | HS | MA0661.1 |
| 3.28532355 | 0.485540993160425 | 0.646 | 0.493 | 2.45E-15 | HS | MA1113.2 |
| 1.24000097 | 0.485362623079696 | 0.613 | 0.488 | 9.25E-14 | HS | MA0888.1 |
| 1.57047311 | 0.483484071118369 | 0.644 | 0.493 | 1.17E-14 | HS | MA1530.1 |
| 2.96463483 | 0.481026411722476 | 0.646 | 0.496 | 2.21E-12 | HS | MA0259.1 |
| 2.34185599 | 0.473670704601788 | 0.644 | 0.502 | 1.75E-13 | HS | MA0601.1 |
| 9.34711087 | 0.466066318092169 | 0.667 | 0.506 | 6.97E-14 | HS | MA0142.1 |
| 2.57703814 | 0.461264178231614 | 0.681 | 0.498 | 1.92E-12 | HS | MA0766.2 |
| 4.61192594 | 0.457190425206967 | 0.656 | 0.5 | 3.44E-13 | HS | MA0479.1 |
| 1.17013196 | 0.455385063443387 | 0.617 | 0.492 | 8.73E-11 | HS | MA0887.1 |
| 1.10178124 | 0.447999610747112 | 0.656 | 0.503 | 8.22E-13 | HS | MA0052.4 |
| 2.11711799 | 0.439097364309398 | 0.627 | 0.501 | 1.58E-10 | HS | MA1106.1 |
| 9.40333588 | 0.437907441232465 | 0.65 | 0.494 | 7.01E-11 | HS | MA0075.3 |
| 7.11473476 | 0.435757676840927 | 0.633 | 0.486 | 5.31E-11 | HS | MA0037.3 |
| 5.71497675 | 0.434117120861342 | 0.644 | 0.49 | 4.26E-11 | HS | MA0781.1 |
| 2.50326774 | 0.431619653827715 | 0.629 | 0.498 | 1.87E-11 | HS | MA0851.1 |
| 1.98499000 | 0.431187727701443 | 0.627 | 0.496 | 1.48E-10 | HS | MA0909.2 |
| 1.65048051 | 0.424183599888301 | 0.658 | 0.502 | 1.23E-11 | HS | MA0527.1 |
| 2.95464597 | 0.422567514985169 | 0.642 | 0.492 | 2.20E-11 | HS | MA1475.1 |
| 3.05823070 | 0.420143055238456 | 0.6 | 0.497 | 2.28E-09 | HS | MA0031.1 |
| 3.05823070 | 0.420143055238456 | 0.6 | 0.497 | 2.28E-09 | HS | MA0157.2 |
| 1.32521471 | 0.417996234925123 | 0.644 | 0.508 | 9.89E-12 | HS | MA0730.1 |
| 1.97253091 | 0.408186118856566 | 0.629 | 0.495 | 1.47E-09 | HS | MA0793.1 |
| 9.02835913 | 0.407918455642237 | 0.623 | 0.494 | 6.74E-09 | HS | MA1110.1 |
| 3.34380282 | 0.395757970502041 | 0.604 | 0.492 | 2.49E-08 | HS | MA1478.1 |
| 1.17129883 | 0.392784114314045 | 0.613 | 0.495 | 8.74E-10 | HS | MA0847.2 |
| 4.89963326 | 0.39051360164217 | 0.6 | 0.494 | 3.66E-07 | HS | MA0902.2 |
| 7.49713899 | 0.386524438186357 | 0.617 | 0.489 | 5.59E-08 | HS | MA1576.1 |
| 3.01497896 | 0.383081418901318 | 0.625 | 0.51 | 2.25E-07 | HS | MA0692.1 |
| 6.67078999 | 0.382745595852627 | 0.602 | 0.491 | 4.98E-07 | HS | MA0059.1 |
| 2.10386659 | 0.382623326923023 | 0.652 | 0.496 | 1.57E-09 | HS | MA0038.2 |
| 1.29963254 | 0.381457595464888 | 0.621 | 0.493 | 9.70E-08 | HS | MA0890.1 |
| 1.62279659 | 0.380058841222833 | 0.642 | 0.498 | 1.21E-09 | HS | MA0869.1 |
| 1.19283900 | 0.370697078564671 | 0.642 | 0.501 | 8.90E-09 | HS | MA0613.1 |

| | | | | | | |
|---|---|---|---|---|---|---|
| 2.48522313 | 0.367976644778596 | 0.61 | 0.493 | 1.85E-06 | HS | MA0070.1 |
| 2.26174547 | 0.365244998995975 | 0.608 | 0.493 | 1.69E-06 | HS | MA0717.1 |
| 1.43182275 | 0.364922665802106 | 0.656 | 0.502 | 1.07E-08 | HS | MA0858.1 |
| 1.85299168 | 0.361002399962798 | 0.615 | 0.489 | 1.38E-06 | HS | MA0886.1 |
| 1.23166941 | 0.357944742479599 | 0.594 | 0.5 | 9.19E-06 | HS | MA0495.3 |
| 8.93911728 | 0.352912497615016 | 0.621 | 0.504 | 6.67E-06 | HS | MA0100.3 |
| 2.06212226 | 0.35285342920147 | 0.59 | 0.492 | 1.54E-05 | HS | MA0654.1 |
| 2.06212226 | 0.35285342920147 | 0.59 | 0.492 | 1.54E-05 | HS | MA0720.1 |
| 2.06212226 | 0.35285342920147 | 0.59 | 0.492 | 1.54E-05 | HS | MA0701.2 |
| 1.24580648 | 0.35225650451037 | 0.596 | 0.502 | 9.29E-07 | HS | MA0614.1 |
| 1.47972401 | 0.350450780869831 | 0.588 | 0.495 | 1.10E-05 | HS | MA0855.1 |
| 2.07250579 | 0.339841059595575 | 0.585 | 0.49 | 0.000154609 | HS | MA1501.1 |
| 2.31980235 | 0.339367960238231 | 0.61 | 0.498 | 1.73E-06 | HS | MA1537.1 |
| 9.97455523 | 0.33784412040236 | 0.6 | 0.496 | 7.44E-06 | HS | MA1583.1 |
| 9.80671237 | 0.337276961521373 | 0.588 | 0.501 | 7.32E-05 | HS | MA0494.1 |
| 2.23484831 | 0.33696851592189 | 0.621 | 0.494 | 1.67E-06 | HS | MA1114.1 |
| 3.35536307 | 0.336903538223072 | 0.577 | 0.503 | 2.50E-05 | HS | MA1466.1 |
| 1.21776637 | 0.336096220565686 | 0.6 | 0.501 | 9.08E-06 | HS | MA0058.3 |
| 4.72255289 | 0.335317066798611 | 0.594 | 0.495 | 3.52E-05 | HS | MA0638.1 |
| 1.04455823 | 0.333028464696198 | 0.606 | 0.492 | 7.79E-06 | HS | MA1496.1 |
| 6.42618118 | 0.331020325346274 | 0.623 | 0.495 | 4.79E-06 | HS | MA0030.1 |
| 3.56304713 | 0.328145318880977 | 0.608 | 0.494 | 2.66E-05 | HS | MA0042.2 |
| 3.56304713 | 0.328145318880977 | 0.608 | 0.494 | 2.66E-05 | HS | MA0033.2 |
| 3.56304713 | 0.328145318880977 | 0.608 | 0.494 | 2.66E-05 | HS | MA0848.1 |
| 3.56304713 | 0.328145318880977 | 0.608 | 0.494 | 2.66E-05 | HS | MA0849.1 |
| 3.56304713 | 0.328145318880977 | 0.608 | 0.494 | 2.66E-05 | HS | MA0850.1 |
| 2.41842329 | 0.328075300389784 | 0.619 | 0.494 | 1.80E-06 | HS | MA0036.3 |
| 5.91560999 | 0.325833872320428 | 0.581 | 0.486 | 4.41E-05 | HS | MA1549.1 |
| 3.98167940 | 0.325261644322575 | 0.619 | 0.502 | 2.97E-08 | HS | MA0151.1 |
| 1.70869270 | 0.324008559609978 | 0.592 | 0.494 | 0.000127468 | HS | MA0068.2 |
| 1.44156560 | 0.323949328516084 | 0.588 | 0.489 | 0.000107541 | HS | MA0725.1 |
| 1.44156560 | 0.323949328516084 | 0.588 | 0.489 | 0.000107541 | HS | MA0726.1 |
| 1.22068464 | 0.322032337410518 | 0.604 | 0.505 | 9.11E-05 | HS | MA0782.2 |
| 5.60105475 | 0.32005772896387 | 0.577 | 0.487 | 0.000417839 | HS | MA0705.1 |
| 6.24425730 | 0.31999542352189 | 0.598 | 0.487 | 0.000465822 | HS | MA1594.1 |
| 7.89154497 | 0.316670592828321 | 0.581 | 0.488 | 0.000588709 | HS | MA0722.1 |
| 7.89154497 | 0.316670592828321 | 0.581 | 0.488 | 0.000588709 | HS | MA0723.1 |
| 1.16186073 | 0.316523859355981 | 0.575 | 0.493 | 8.67E-05 | HS | MA0628.1 |
| 8.76856289 | 0.315751403135674 | 0.577 | 0.497 | 6.54E-05 | HS | MA0608.1 |
| 9.03586540 | 0.31289150622514 | 0.606 | 0.504 | 6.74E-05 | HS | MA1655.1 |
| 1.99554571 | 0.311810985950017 | 0.588 | 0.491 | 0.000148868 | HS | MA0710.1 |
| 1.81289964 | 0.30553949394619 | 0.594 | 0.499 | 0.000135242 | HS | MA0004.1 |
| 7.07947967 | 0.303490303798001 | 0.594 | 0.494 | 0.000528129 | HS | MA0699.1 |

| | | | | | | |
|---|---|---|---|---|---|---|
| 4.97059840 | 0.302048347658248 | 0.598 | 0.499 | 3.71E-05 | HS | MA0653.1 |
| 5.21602109 | 0.299879589631277 | 0.573 | 0.496 | 0.003891152 | HS | MA0804.1 |
| 3.99756772 | 0.293906766800485 | 0.629 | 0.505 | 0.000298219 | HS | MA1485.1 |
| 8.95591713 | 0.293338144874286 | 0.6 | 0.49 | 0.000668111 | HS | MA0775.1 |
| 1.43158965 | 0.292064597417052 | 0.588 | 0.503 | 0.001067966 | HS | MA0519.1 |
| 3.17620689 | 0.289355359222415 | 0.615 | 0.488 | 0.000236945 | HS | MA0662.1 |
| 6.80690167 | 0.287722069407379 | 0.592 | 0.495 | 0.000507795 | HS | MA0845.1 |
| 2.04939270 | 0.286233574422991 | 0.596 | 0.492 | 0.001528847 | HS | MA0646.1 |
| 4.49880659 | 0.285578501077195 | 0.592 | 0.492 | 0.00335611 | HS | MA0630.1 |
| 1.39606767 | 0.285427497588754 | 0.59 | 0.501 | 0.000104147 | HS | MA1122.1 |
| 4.92948921 | 0.285184263945532 | 0.585 | 0.496 | 0.003677399 | HS | MA0707.1 |
| 2.43173648 | 0.285017975390914 | 0.577 | 0.494 | 0.001814075 | HS | MA1683.1 |
| 7.67112918 | 0.283067160931821 | 0.615 | 0.499 | 5.72E-05 | HS | MA0650.2 |
| 9.25793374 | 0.282434980800332 | 0.571 | 0.507 | 0.006906419 | HS | MA1597.1 |
| 1.47527860 | 0.27956313469824 | 0.581 | 0.491 | 0.001100558 | HS | MA0874.1 |
| 7.70582773 | 0.279040332465748 | 0.581 | 0.494 | 0.005748547 | HS | MA0674.1 |
| 3.46088110 | 0.277773067569547 | 0.567 | 0.493 | 0.002581817 | HS | MA0147.3 |
| 4.47504524 | 0.277258157889093 | 0.579 | 0.497 | 0.003338384 | HS | MA0773.1 |
| 9.23299641 | 0.277117161323093 | 0.583 | 0.488 | 0.000688782 | HS | MA1553.1 |
| 8.08044273 | 0.26622005954397 | 0.573 | 0.491 | 0.00602801 | HS | MA0083.3 |
| 1.12029058 | 0.265052169717046 | 0.575 | 0.499 | 0.008357368 | HS | MA1120.1 |
| 2.24070800 | 0.263347112474474 | 0.577 | 0.495 | 0.016715682 | HS | MA0621.1 |
| 3.33612832 | 0.262033405970595 | 0.562 | 0.499 | 0.024887517 | HS | MA0660.1 |
| 1.76180047 | 0.261378064718369 | 0.594 | 0.5 | 0.001314303 | HS | MA1627.1 |
| 5.61651570 | 0.261194946008812 | 0.581 | 0.491 | 0.004189921 | HS | MA1652.1 |
| 3.59750034 | 0.256525919999366 | 0.573 | 0.503 | 0.026837353 | HS | MA0636.1 |
| 6.04703780 | 0.25410253664733 | 0.577 | 0.495 | 0.000451109 | HS | MA1125.1 |
| 1.97842545 | 0.252843460532437 | 0.596 | 0.515 | 0.014759054 | HS | MA0526.3 |
| 1.72581448 | 0.251668384803685 | 0.588 | 0.495 | 0.012874576 | HS | MA0069.1 |
| 9.35818052 | 0.87253591008054 | 0.763 | 0.485 | 6.98E-64 | SCH | MA0046.2 |
| 3.65245029 | 0.864389540829831 | 0.752 | 0.489 | 2.72E-58 | SCH | MA0706.1 |
| 1.22310585 | 0.846618041394488 | 0.75 | 0.481 | 9.12E-58 | SCH | MA0728.1 |
| 1.68527375 | 0.823023231056862 | 0.736 | 0.48 | 1.26E-51 | SCH | MA0056.2 |
| 2.25746456 | 0.798257655589531 | 0.77 | 0.495 | 1.68E-53 | SCH | MA0857.1 |
| 6.19395728 | 0.767272296171647 | 0.749 | 0.488 | 4.62E-50 | SCH | MA0810.1 |
| 8.31937002 | 0.765836639257995 | 0.741 | 0.493 | 6.21E-45 | SCH | MA0859.1 |
| 2.32122806 | 0.761742346590695 | 0.727 | 0.482 | 1.73E-45 | SCH | MA1556.1 |
| 6.32799419 | 0.72594275584982 | 0.714 | 0.483 | 4.72E-40 | SCH | MA1146.1 |
| 1.69799497 | 0.725169752690719 | 0.74 | 0.487 | 1.27E-43 | SCH | MA1656.1 |
| 4.74250625 | 0.723894660319959 | 0.723 | 0.487 | 3.54E-41 | SCH | MA1555.1 |
| 2.14719256 | 0.711015274887342 | 0.703 | 0.482 | 1.60E-36 | SCH | MA0693.2 |
| 9.83553788 | 0.707429278714567 | 0.719 | 0.487 | 7.34E-40 | SCH | MA0886.1 |
| 8.22321253 | 0.699199271811972 | 0.719 | 0.485 | 6.13E-40 | SCH | MA0798.2 |

| | | | | | | |
|---|---|---|---|---|---|---|
| 1.04824296 | 0.685390932634059 | 0.719 | 0.497 | 7.82E-38 | SCH | MA0653.1 |
| 6.03263605 | 0.677913760177694 | 0.719 | 0.488 | 4.50E-37 | SCH | MA1641.1 |
| 2.75219776 | 0.67522138491371 | 0.719 | 0.494 | 2.05E-38 | SCH | MA0160.1 |
| 2.78622729 | 0.673697119950367 | 0.736 | 0.49 | 2.08E-38 | SCH | MA0702. |
| 4.64419090 | 0.663138623280261 | 0.714 | 0.488 | 3.46E-35 | SCH | MA1619.1 |
| 2.56477720 | 0.661665251274591 | 0.736 | 0.502 | 1.91E-38 | SCH | MA1604.1 |
| 2.92728217 | 0.658071242624963 | 0.716 | 0.494 | 2.18E-36 | SCH | MA1637.1 |
| 2.66025770 | 0.648628552764439 | 0.694 | 0.49 | 1.98E-33 | SCH | MA0154. |
| 1.27364269 | 0.64655811920242 | 0.714 | 0.497 | 9.50E-35 | SCH | MA1536.1 |
| 3.31093559 | 0.643305218765722 | 0.707 | 0.492 | 2.47E-33 | SCH | MA1463.1 |
| 2.53193479 | 0.64281147326055 | 0.712 | 0.489 | 1.89E-34 | SCH | MA0675.1 |
| 1.30653017 | 0.636397733432964 | 0.667 | 0.49 | 9.75E-28 | SCH | MA1531.1 |
| 7.17422908 | 0.63509878171172 | 0.707 | 0.487 | 5.35E-31 | SCH | MA1519.1 |
| 8.36940202 | 0.632799564453546 | 0.687 | 0.493 | 6.24E-31 | SCH | MA1589.1 |
| 1.12270580 | 0.627514716609811 | 0.707 | 0.494 | 8.38E-31 | SCH | MA0811.1 |
| 9.19314266 | 0.624681686569475 | 0.687 | 0.493 | 6.86E-30 | SCH | MA0719.1 |
| 3.36575489 | 0.622161417611328 | 0.718 | 0.494 | 2.51E-31 | SCH | MA0141. |
| 2.10165460 | 0.620855125977024 | 0.699 | 0.491 | 1.57E-29 | SCH | MA0642.1 |
| 4.32058241 | 0.614549452687422 | 0.678 | 0.487 | 3.22E-27 | SCH | MA0722.1 |
| 4.32058241 | 0.614549452687422 | 0.678 | 0.487 | 3.22E-27 | SCH | MA0723.1 |
| 1.60463860 | 0.611948279081233 | 0.73 | 0.499 | 1.20E-34 | SCH | MA0479.1 |
| 7.09105730 | 0.589279149198266 | 0.707 | 0.49 | 5.29E-28 | SCH | MA0634.1 |
| 5.81635110 | 0.587500488802488 | 0.69 | 0.488 | 4.34E-25 | SCH | MA0724.1 |
| 2.05106854 | 0.573300666295783 | 0.65 | 0.484 | 1.53E-17 | SCH | MA0116.1 |
| 1.23807438 | 0.570997934925852 | 0.692 | 0.492 | 9.24E-28 | SCH | MA1615.1 |
| 1.93681999 | 0.570038477136454 | 0.663 | 0.494 | 1.44E-24 | SCH | MA0768.1 |
| 8.96097445 | 0.568058389197664 | 0.679 | 0.487 | 6.68E-28 | SCH | MA1508.1 |
| 1.51141816 | 0.56540756822908 | 0.687 | 0.5 | 1.13E-27 | SCH | MA0816.1 |
| 8.53722126 | 0.561955610972325 | 0.679 | 0.49 | 6.37E-24 | SCH | MA0153. |
| 1.09199690 | 0.560345668600186 | 0.674 | 0.489 | 8.15E-24 | SCH | MA1481.1 |
| 5.95329088 | 0.554365876323738 | 0.696 | 0.498 | 4.44E-25 | SCH | MA0524. |
| 1.48611289 | 0.552407139982051 | 0.667 | 0.488 | 1.11E-23 | SCH | MA0713.1 |
| 8.41680895 | 0.53993566292694 | 0.665 | 0.487 | 6.28E-20 | SCH | MA0662.1 |
| 8.05201464 | 0.534753575299769 | 0.648 | 0.477 | 6.01E-20 | SCH | MA0776.1 |
| 3.11851307 | 0.530419075069882 | 0.674 | 0.491 | 2.33E-22 | SCH | MA0704.1 |
| 8.80190447 | 0.52298343187256 | 0.658 | 0.485 | 6.57E-21 | SCH | MA0071.1 |
| 4.92889647 | 0.522789459275922 | 0.667 | 0.492 | 3.68E-19 | SCH | MA0628.1 |
| 1.11317815 | 0.51896682607314 | 0.659 | 0.497 | 8.30E-20 | SCH | MA0115.1 |
| 5.84043959 | 0.517692770136191 | 0.652 | 0.485 | 4.36E-18 | SCH | MA1505.1 |
| 9.95555229 | 0.516996059665724 | 0.645 | 0.499 | 7.43E-21 | SCH | MA0521.1 |
| 9.26595406 | 0.516677176381227 | 0.65 | 0.49 | 6.91E-19 | SCH | MA0889.1 |
| 2.24106163 | 0.515515885073187 | 0.654 | 0.49 | 1.67E-18 | SCH | MA0525. |
| 5.51687576 | 0.505949821953771 | 0.627 | 0.485 | 4.12E-17 | SCH | MA1547.1 |

| | | | | | | |
|---|---|---|---|---|---|---|
| 1.239380769 | 0.50350696308461 | 0.639 | 0.48 | 9.25E-18 | SCH | MA1484.1 |
| 4.082081693 | 0.501288727911435 | 0.663 | 0.494 | 3.05E-18 | SCH | MA0680.1 |
| 3.062517737 | 0.493194143474592 | 0.648 | 0.489 | 2.28E-17 | SCH | MA0716.1 |
| 3.062517737 | 0.493194143474592 | 0.648 | 0.489 | 2.28E-17 | SCH | MA0721.1 |
| 2.668163730 | 0.491909467834244 | 0.65 | 0.489 | 1.99E-18 | SCH | MA1472.1 |
| 5.515611932 | 0.48714349328831 | 0.681 | 0.502 | 4.11E-19 | SCH | MA1629.1 |
| 5.606286911 | 0.484616861949285 | 0.65 | 0.494 | 4.18E-16 | SCH | MA0712.2 |
| 4.485049740 | 0.482265870469485 | 0.659 | 0.49 | 3.35E-19 | SCH | MA0598.3 |
| 5.979602299 | 0.480762201906659 | 0.667 | 0.484 | 4.46E-19 | SCH | MA0640.2 |
| 3.023922880 | 0.480261242224976 | 0.661 | 0.489 | 2.26E-17 | SCH | MA1636.1 |
| 6.166091178 | 0.477856758872301 | 0.641 | 0.483 | 4.60E-14 | SCH | MA0656.1 |
| 4.602539139 | 0.475491909489955 | 0.65 | 0.495 | 3.43E-16 | SCH | MA0707.1 |
| 4.612605088 | 0.475481979450128 | 0.654 | 0.498 | 3.44E-16 | SCH | MA0780.1 |
| 8.202490617 | 0.469171328615471 | 0.619 | 0.485 | 6.12E-14 | SCH | MA0600.2 |
| 2.276877400 | 0.468836683932624 | 0.63 | 0.498 | 1.70E-14 | SCH | MA1419.1 |
| 1.936506977 | 0.468639193251867 | 0.668 | 0.483 | 1.44E-17 | SCH | MA0509.2 |
| 9.684873237 | 0.466291988531098 | 0.63 | 0.49 | 7.22E-14 | SCH | MA0661.1 |
| 1.233331154 | 0.461876384824227 | 0.676 | 0.496 | 9.20E-18 | SCH | MA1421.1 |
| 6.717491060 | 0.461587150781636 | 0.605 | 0.482 | 5.01E-12 | SCH | MA0007.3 |
| 1.336966484 | 0.459629704283743 | 0.658 | 0.5 | 9.97E-16 | SCH | MA1111.1 |
| 1.381151365 | 0.45907274565054 | 0.656 | 0.5 | 1.03E-15 | SCH | MA1635.1 |
| 1.428652340 | 0.458392682157205 | 0.659 | 0.491 | 1.07E-17 | SCH | MA0471.2 |
| 3.498992622 | 0.455846053411217 | 0.625 | 0.497 | 2.61E-14 | SCH | MA0670.1 |
| 2.066864713 | 0.455422373932708 | 0.674 | 0.494 | 1.54E-16 | SCH | MA1581.1 |
| 2.064443829 | 0.453345690719104 | 0.654 | 0.492 | 1.54E-15 | SCH | MA0735.1 |
| 1.181308765 | 0.449236021748059 | 0.661 | 0.499 | 8.81E-16 | SCH | MA1100.2 |
| 1.005243492 | 0.449104927849335 | 0.639 | 0.491 | 7.50E-15 | SCH | MA1471.1 |
| 5.918616390 | 0.44558226845127 | 0.659 | 0.486 | 4.42E-15 | SCH | MA0499.2 |
| 5.563696472 | 0.443636969806602 | 0.647 | 0.502 | 4.15E-15 | SCH | MA0477.2 |
| 7.939020867 | 0.441503798892873 | 0.656 | 0.49 | 5.92E-16 | SCH | MA1652.1 |
| 1.597616251 | 0.437296110916832 | 0.636 | 0.491 | 1.19E-13 | SCH | MA0618.1 |
| 2.127433625 | 0.431404361215661 | 0.625 | 0.489 | 1.59E-12 | SCH | MA0132.2 |
| 1.071184467 | 0.429691191994852 | 0.658 | 0.5 | 7.99E-15 | SCH | MA1529.1 |
| 2.963263270 | 0.429196018089051 | 0.639 | 0.486 | 2.21E-14 | SCH | MA0753.2 |
| 7.318710529 | 0.427094689603191 | 0.627 | 0.494 | 5.46E-12 | SCH | MA0772.1 |
| 2.503228717 | 0.42557164117996 | 0.608 | 0.484 | 1.87E-11 | SCH | MA0510.2 |
| 5.207044387 | 0.424294842829344 | 0.634 | 0.494 | 3.88E-14 | SCH | MA0764.2 |
| 1.673363578 | 0.421432893769066 | 0.641 | 0.489 | 1.25E-12 | SCH | MA1535.1 |
| 2.246314498 | 0.418785779336315 | 0.645 | 0.501 | 1.68E-13 | SCH | MA1628.1 |
| 2.124483088 | 0.414428303714706 | 0.652 | 0.494 | 1.58E-13 | SCH | MA0698.1 |
| 2.867321639 | 0.405593863989978 | 0.636 | 0.489 | 2.14E-10 | SCH | MA0727.1 |
| 2.772469831 | 0.401061062866158 | 0.656 | 0.5 | 2.07E-15 | SCH | MA0080.5 |
| 1.366715591 | 0.398770173770281 | 0.597 | 0.482 | 1.02E-08 | SCH | MA1127.1 |

| | | | | | | |
|---|---|---|---|---|---|---|
| 5.29469749 | 0.396904064012349 | 0.596 | 0.489 | 3.95E-09 | SCH | MA0714.1 |
| 2.30967379 | 0.395940111309829 | 0.612 | 0.482 | 1.72E-10 | SCH | MA0103.3 |
| 2.66672113 | 0.395760434882475 | 0.638 | 0.491 | 1.99E-12 | SCH | MA0832.1 |
| 7.44258050 | 0.392292522614633 | 0.639 | 0.496 | 5.55E-11 | SCH | MA0106.3 |
| 9.61041889 | 0.391883222153593 | 0.643 | 0.501 | 7.17E-12 | SCH | MA0761.2 |
| 6.65347961 | 0.391797998099699 | 0.608 | 0.495 | 4.96E-10 | SCH | MA0523.1 |
| 2.80376482 | 0.390346277273474 | 0.596 | 0.486 | 2.09E-08 | SCH | MA1147.1 |
| 1.93083338 | 0.38825406395301 | 0.617 | 0.483 | 1.44E-09 | SCH | MA0799.1 |
| 9.81069634 | 0.387123810717853 | 0.636 | 0.5 | 7.32E-12 | SCH | MA1630.1 |
| 2.60361245 | 0.386683146584122 | 0.599 | 0.488 | 1.94E-08 | SCH | MA1645.1 |
| 5.95073375 | 0.386254433172365 | 0.617 | 0.49 | 4.44E-10 | SCH | MA0790.1 |
| 4.57116680 | 0.385654619841295 | 0.596 | 0.487 | 3.41E-08 | SCH | MA0883.1 |
| 4.21827409 | 0.384085204293397 | 0.614 | 0.486 | 3.15E-09 | SCH | MA0903.1 |
| 6.17521570 | 0.382108014376609 | 0.612 | 0.489 | 4.61E-09 | SCH | MA0833.2 |
| 6.34479431 | 0.38193345904744 | 0.63 | 0.493 | 4.73E-09 | SCH | MA0674.1 |
| 2.63488024 | 0.381759634558031 | 0.605 | 0.491 | 1.97E-08 | SCH | MA0907.1 |
| 1.07473523 | 0.381705215835494 | 0.621 | 0.51 | 8.02E-09 | SCH | MA0051.1 |
| 4.87495340 | 0.381645889271833 | 0.61 | 0.49 | 3.64E-10 | SCH | MA0737.1 |
| 2.42480379 | 0.38035114736538 | 0.612 | 0.488 | 1.81E-08 | SCH | MA1546.1 |
| 1.01345153 | 0.379491970169466 | 0.632 | 0.502 | 7.56E-10 | SCH | MA0676.1 |
| 3.31517479 | 0.376222581050103 | 0.643 | 0.498 | 2.47E-10 | SCH | MA0048.2 |
| 1.35546353 | 0.375671148086986 | 0.603 | 0.477 | 1.01E-07 | SCH | MA0882.1 |
| 3.12771692 | 0.373848801119822 | 0.576 | 0.489 | 2.33E-07 | SCH | MA1470.1 |
| 7.29443833 | 0.373630961049909 | 0.607 | 0.491 | 5.44E-09 | SCH | MA0078.1 |
| 6.67921150 | 0.370514195412397 | 0.599 | 0.485 | 4.98E-08 | SCH | MA0700.2 |
| 1.43481505 | 0.369893510435695 | 0.621 | 0.498 | 1.07E-08 | SCH | MA1499.1 |
| 1.43481505 | 0.369893510435695 | 0.621 | 0.498 | 1.07E-08 | SCH | MA1507.1 |
| 4.41900737 | 0.36903331343478 | 0.645 | 0.493 | 3.30E-13 | SCH | MA1653.1 |
| 7.63577212 | 0.36852984615101 | 0.619 | 0.502 | 5.70E-09 | SCH | MA0601.1 |
| 8.58802215 | 0.36761475571275 | 0.605 | 0.487 | 6.41E-08 | SCH | MA0138.2 |
| 6.43802574 | 0.367191282834274 | 0.597 | 0.494 | 4.80E-07 | SCH | MA1649.1 |
| 5.74952875 | 0.366133746231692 | 0.63 | 0.49 | 4.29E-10 | SCH | MA0505.1 |
| 7.69032748 | 0.365372260427629 | 0.605 | 0.487 | 5.74E-09 | SCH | MA0163.1 |
| 5.94449552 | 0.364077414019424 | 0.607 | 0.493 | 4.43E-08 | SCH | MA0643.1 |
| 1.01181922 | 0.362759121458393 | 0.574 | 0.477 | 7.55E-07 | SCH | MA1533.1 |
| 4.47502334 | 0.36107323502798 | 0.616 | 0.489 | 3.34E-09 | SCH | MA0091.1 |
| 7.75720594 | 0.357537536826823 | 0.59 | 0.482 | 5.79E-07 | SCH | MA1554.1 |
| 2.89149782 | 0.357051213655927 | 0.607 | 0.487 | 2.16E-07 | SCH | MA0609.2 |
| 3.63837969 | 0.355817483383475 | 0.621 | 0.493 | 2.71E-09 | SCH | MA0665.1 |
| 2.17330114 | 0.355384757062957 | 0.599 | 0.488 | 1.62E-07 | SCH | MA0840.1 |
| 5.30427017 | 0.355323227686822 | 0.587 | 0.489 | 3.96E-06 | SCH | MA0605.2 |
| 7.94971764 | 0.35402858711893 | 0.621 | 0.488 | 5.93E-09 | SCH | MA0164.1 |
| 1.56199356 | 0.353354576738815 | 0.61 | 0.489 | 1.17E-07 | SCH | MA1497.1 |

| | | | | | | |
|---|---|---|---|---|---|---|
| 3.09413429[illegible] | 0.34835510692071 | 0.561 | 0.484 | 2.31E-06 | SCH | MA1538.1 |
| 3.07774514[illegible] | 0.347517789558673 | 0.579 | 0.489 | 2.30E-06 | SCH | MA1567.1 |
| 5.33892381[illegible] | 0.347084571477747 | 0.603 | 0.5 | 3.98E-08 | SCH | MA1137.1 |
| 4.90413996[illegible] | 0.344778638645608 | 0.61 | 0.487 | 3.66E-07 | SCH | MA1553.1 |
| 8.64526246[illegible] | 0.341732823048183 | 0.579 | 0.489 | 6.45E-06 | SCH | MA0682.[illegible] |
| 6.55103992[illegible] | 0.340530110120969 | 0.585 | 0.491 | 4.89E-06 | SCH | MA1521.1 |
| 1.02252022[illegible] | 0.340283488432518 | 0.599 | 0.486 | 7.63E-06 | SCH | MA1585.1 |
| 1.08369082[illegible] | 0.340033581070885 | 0.596 | 0.494 | 8.08E-07 | SCH | MA0679.[illegible] |
| 8.16992281[illegible] | 0.338993366631055 | 0.614 | 0.497 | 6.09E-08 | SCH | MA1128.1 |
| 1.04075615[illegible] | 0.336952312854095 | 0.608 | 0.5 | 7.76E-08 | SCH | MA1141.1 |
| 1.44937194[illegible] | 0.336889731785814 | 0.625 | 0.492 | 1.08E-07 | SCH | MA0591.1 |
| 5.33164217[illegible] | 0.335437103912926 | 0.601 | 0.486 | 3.98E-07 | SCH | MA0705.1 |
| 1.14299374[illegible] | 0.334458992631938 | 0.59 | 0.489 | 8.53E-07 | SCH | MA1518.1 |
| 2.51608340[illegible] | 0.331127259779653 | 0.588 | 0.485 | 1.88E-07 | SCH | MA1633.1 |
| 2.63590664[illegible] | 0.330346524627032 | 0.587 | 0.484 | 1.97E-06 | SCH | MA1608.1 |
| 5.46867155[illegible] | 0.327630423262336 | 0.605 | 0.493 | 4.08E-07 | SCH | MA0703.[illegible] |
| 1.55451971[illegible] | 0.326571178645587 | 0.594 | 0.494 | 1.16E-06 | SCH | MA0075.[illegible] |
| 4.08512064[illegible] | 0.325337528315735 | 0.612 | 0.482 | 3.05E-08 | SCH | MA0039.[illegible] |
| 6.40753353[illegible] | 0.324265350997821 | 0.616 | 0.498 | 4.78E-07 | SCH | MA0478.1 |
| 2.21768699[illegible] | 0.322653661237162 | 0.612 | 0.489 | 1.65E-06 | SCH | MA0607.1 |
| 1.48657838[illegible] | 0.322464448015687 | 0.581 | 0.488 | 1.11E-05 | SCH | MA0018.[illegible] |
| 6.32221455[illegible] | 0.320097569027257 | 0.614 | 0.493 | 4.72E-08 | SCH | MA0146.[illegible] |
| 1.01687564[illegible] | 0.317717792523148 | 0.581 | 0.481 | 7.59E-05 | SCH | MA0594.[illegible] |
| 4.11423430[illegible] | 0.314882818937441 | 0.623 | 0.496 | 3.07E-06 | SCH | MA0853.1 |
| 4.83375242[illegible] | 0.313736539530782 | 0.587 | 0.493 | 3.61E-06 | SCH | MA0484.[illegible] |
| 5.63764903[illegible] | 0.313339958163951 | 0.61 | 0.5 | 4.21E-08 | SCH | MA1627.1 |
| 6.90144386[illegible] | 0.312983528328584 | 0.588 | 0.49 | 5.15E-08 | SCH | MA0050.[illegible] |
| 8.73522253[illegible] | 0.310133296060606 | 0.599 | 0.489 | 6.52E-06 | SCH | MA0113.[illegible] |
| 1.16123984[illegible] | 0.309401956021527 | 0.596 | 0.492 | 8.66E-06 | SCH | MA1657.1 |
| 9.34024443[illegible] | 0.308944741033224 | 0.596 | 0.499 | 6.97E-06 | SCH | MA1491.1 |
| 1.82499496[illegible] | 0.308202580718322 | 0.597 | 0.494 | 1.36E-05 | SCH | MA1530.1 |
| 7.12448301[illegible] | 0.307488142671083 | 0.581 | 0.493 | 5.31E-05 | SCH | MA0765.[illegible] |
| 2.28260044[illegible] | 0.306281403458051 | 0.592 | 0.499 | 1.70E-05 | SCH | MA0159.1 |
| 2.71862088[illegible] | 0.298901137989521 | 0.61 | 0.502 | 2.03E-05 | SCH | MA1134.1 |
| 1.17488151[illegible] | 0.298089827459489 | 0.612 | 0.503 | 8.76E-05 | SCH | MA0514.1 |
| 1.41781459[illegible] | 0.296778720521344 | 0.587 | 0.491 | 1.06E-05 | SCH | MA0645.1 |
| 3.51350820[illegible] | 0.293825901333974 | 0.579 | 0.492 | 0.000262108 | SCH | MA0654.1 |
| 3.51350820[illegible] | 0.293825901333974 | 0.579 | 0.492 | 0.000262108 | SCH | MA0720.1 |
| 3.51350820[illegible] | 0.293825901333974 | 0.579 | 0.492 | 0.000262108 | SCH | MA0701.[illegible] |
| 1.67730721[illegible] | 0.29345870598463 | 0.601 | 0.488 | 1.25E-05 | SCH | MA1607.1 |
| 3.67653769[illegible] | 0.290493568450671 | 0.59 | 0.488 | 0.00027427 | SCH | MA0888.1 |
| 8.00428444[illegible] | 0.288408050836274 | 0.601 | 0.489 | 5.97E-05 | SCH | MA0667.1 |
| 1.95319790[illegible] | 0.288301279783945 | 0.605 | 0.494 | 1.46E-06 | SCH | MA1125.1 |

| | | | | | | |
|---|---|---|---|---|---|---|
| 5.240656418 | 0.286617126216204 | 0.581 | 0.49 | 0.000390953 | SCH | MA0767.1 |
| 9.749707920 | 0.286547395139799 | 0.605 | 0.496 | 7.27E-05 | SCH | MA0508.3 |
| 1.821650235 | 0.283595554277278 | 0.572 | 0.489 | 0.000135895 | SCH | MA0892.1 |
| 3.268201330 | 0.280572225605069 | 0.607 | 0.499 | 0.000243808 | SCH | MA0863.1 |
| 3.344963184 | 0.279739798382117 | 0.579 | 0.481 | 0.000249534 | SCH | MA0750.2 |
| 5.159940490 | 0.277577033582216 | 0.594 | 0.494 | 0.000384932 | SCH | MA0621.1 |
| 1.458120838 | 0.273959890886791 | 0.592 | 0.485 | 0.000108776 | SCH | MA1552.1 |
| 9.903138149 | 0.273775583991494 | 0.588 | 0.501 | 0.000738774 | SCH | MA1132.1 |
| 5.608521130 | 0.273410692633354 | 0.605 | 0.494 | 4.18E-05 | SCH | MA0473.3 |
| 2.632410459 | 0.271364607955346 | 0.577 | 0.491 | 0.001963778 | SCH | MA1150.1 |
| 2.465849991 | 0.271205297129734 | 0.574 | 0.493 | 0.001839524 | SCH | MA0822.1 |
| 5.721398020 | 0.270833178294793 | 0.587 | 0.497 | 0.000426816 | SCH | MA0467.1 |
| 1.172476764 | 0.269994698803054 | 0.583 | 0.505 | 0.000874668 | SCH | MA0442.2 |
| 8.795034751 | 0.268660079248894 | 0.59 | 0.485 | 0.00065611 | SCH | MA0485.2 |
| 1.194684907 | 0.266968813871349 | 0.59 | 0.495 | 0.000891235 | SCH | MA0515.1 |
| 1.075641580 | 0.263628268706966 | 0.572 | 0.491 | 0.000802429 | SCH | MA1548.1 |
| 2.655706645 | 0.263171468692618 | 0.599 | 0.502 | 0.001981157 | SCH | MA1148.1 |
| 5.940967328 | 0.263009637166274 | 0.592 | 0.5 | 0.000443196 | SCH | MA1622.1 |
| 3.831722265 | 0.262943495558346 | 0.57 | 0.5 | 0.002858465 | SCH | MA1144.1 |
| 6.902083340 | 0.262175299831683 | 0.559 | 0.499 | 0.005148954 | SCH | MA0067.1 |
| 7.420046350 | 0.261812728017814 | 0.581 | 0.493 | 0.000553535 | SCH | MA0161.2 |
| 7.134142067 | 0.259383903791893 | 0.554 | 0.493 | 0.00532207 | SCH | MA0860.1 |
| 1.273084477 | 0.258279812672991 | 0.576 | 0.483 | 0.00949721 | SCH | MA1568.1 |
| 1.273569560 | 0.255139861994463 | 0.574 | 0.507 | 0.009500829 | SCH | MA0142.1 |
| 2.342599775 | 0.253645094825424 | 0.561 | 0.482 | 0.001747579 | SCH | MA1618.1 |
| 6.647274653 | 0.2520981052106 | 0.57 | 0.498 | 0.004958867 | SCH | MA0861.1 |
| 1.223742730 | 0.251838087733464 | 0.576 | 0.495 | 0.009129121 | SCH | MA0793.1 |
| 2.583994724 | 0.251476632568957 | 0.585 | 0.495 | 0.00192766 | SCH | MA0671.1 |
| 2.301138182 | 0.251095935985688 | 0.566 | 0.492 | 0.017166491 | SCH | MA0710.1 |
| 2.740053513 | 0.671283013261262 | 0.702 | 0.48 | 2.04E-71 | facsFB | MA1573.1 |
| 3.669888288 | 0.656496548401696 | 0.686 | 0.482 | 2.74E-72 | facsFB | MA0163.1 |
| 1.241236271 | 0.649062842104603 | 0.709 | 0.48 | 9.26E-81 | facsFB | MA0747.1 |
| 2.119140419 | 0.639201575798104 | 0.707 | 0.481 | 1.58E-68 | facsFB | MA0815.1 |
| 2.032584533 | 0.624658374669264 | 0.685 | 0.48 | 1.52E-59 | facsFB | MA1608.1 |
| 4.783523220 | 0.623855706033071 | 0.697 | 0.482 | 3.57E-62 | facsFB | MA0813.1 |
| 4.499932412 | 0.622853399311483 | 0.698 | 0.488 | 3.36E-67 | facsFB | MA0070.1 |
| 5.320980283 | 0.619751860234745 | 0.719 | 0.477 | 3.97E-83 | facsFB | MA0039.4 |
| 2.861873910 | 0.600484258667634 | 0.7 | 0.483 | 2.13E-67 | facsFB | MA0740.1 |
| 1.242878229 | 0.600296349842266 | 0.671 | 0.482 | 9.27E-58 | facsFB | MA1584.1 |
| 1.110002538 | 0.587385591345959 | 0.688 | 0.482 | 8.28E-58 | facsFB | MA0057.1 |
| 7.706285404 | 0.55762066221581 | 0.652 | 0.481 | 5.75E-42 | facsFB | MA1476.1 |
| 3.621777815 | 0.552200109206222 | 0.681 | 0.487 | 2.70E-48 | facsFB | MA0645.1 |
| 3.483544610 | 0.549487923210249 | 0.68 | 0.484 | 2.60E-57 | facsFB | MA1564.1 |

| | | | | | | |
|---|---|---|---|---|---|---|
| 1.873200529 | 0.549367089693875 | 0.674 | 0.483 | 1.40E-54 | facsFB | MA0746.2 |
| 2.394958250 | 0.544776385142042 | 0.686 | 0.48 | 1.79E-63 | facsFB | MA0599.1 |
| 2.978241360 | 0.543702812044224 | 0.686 | 0.497 | 2.22E-51 | facsFB | MA0655.1 |
| 2.760179683 | 0.529124470135728 | 0.652 | 0.48 | 2.06E-38 | facsFB | MA0493.1 |
| 6.060518541 | 0.526973487158649 | 0.666 | 0.49 | 4.52E-44 | facsFB | MA0596.1 |
| 6.220421617 | 0.511505771561454 | 0.664 | 0.498 | 4.64E-42 | facsFB | MA1646.1 |
| 2.546264160 | 0.50811508327287 | 0.622 | 0.482 | 1.90E-34 | facsFB | MA0666.1 |
| 2.546264160 | 0.50811508327287 | 0.622 | 0.482 | 1.90E-34 | facsFB | MA0708.1 |
| 2.546264160 | 0.50811508327287 | 0.622 | 0.482 | 1.90E-34 | facsFB | MA0876.1 |
| 9.702982280 | 0.500720354544478 | 0.658 | 0.481 | 7.24E-45 | facsFB | MA1512.1 |
| 7.563200537 | 0.492248196195518 | 0.653 | 0.483 | 5.64E-38 | facsFB | MA0762.1 |
| 1.641322485 | 0.488878967130242 | 0.655 | 0.491 | 1.22E-39 | facsFB | MA0764.2 |
| 3.227800719 | 0.487835383881825 | 0.652 | 0.49 | 2.41E-37 | facsFB | MA1104.2 |
| 3.649879248 | 0.484573001555652 | 0.642 | 0.491 | 2.72E-34 | facsFB | MA0679.2 |
| 9.477148751 | 0.479522647654056 | 0.636 | 0.483 | 7.07E-33 | facsFB | MA0691.1 |
| 3.584036401 | 0.478424423299011 | 0.646 | 0.491 | 2.67E-36 | facsFB | MA0681.2 |
| 9.385763770 | 0.478422907712624 | 0.659 | 0.496 | 7.00E-37 | facsFB | MA1099.2 |
| 8.408759244 | 0.477088450437472 | 0.639 | 0.486 | 6.27E-34 | facsFB | MA0014.3 |
| 9.744858921 | 0.474862147161262 | 0.626 | 0.48 | 7.27E-28 | facsFB | MA0880.1 |
| 9.744858921 | 0.474862147161262 | 0.626 | 0.48 | 7.27E-28 | facsFB | MA0881.1 |
| 1.098493874 | 0.472994930978399 | 0.635 | 0.486 | 8.19E-33 | facsFB | MA0136.2 |
| 1.471575011 | 0.469225179630902 | 0.644 | 0.484 | 1.10E-33 | facsFB | MA0872.1 |
| 9.941495888 | 0.464247264443658 | 0.656 | 0.49 | 7.42E-47 | facsFB | MA1653.1 |
| 8.714042171 | 0.459089464713592 | 0.636 | 0.489 | 6.50E-33 | facsFB | MA0709.1 |
| 3.458112131 | 0.457029246143089 | 0.628 | 0.483 | 2.58E-33 | facsFB | MA1522.1 |
| 5.146064125 | 0.455009210450726 | 0.63 | 0.49 | 3.84E-29 | facsFB | MA0104.4 |
| 2.198291749 | 0.454842662111238 | 0.627 | 0.483 | 1.64E-28 | facsFB | MA1483.1 |
| 7.791920727 | 0.451678976000123 | 0.633 | 0.49 | 5.81E-29 | facsFB | MA0717.1 |
| 1.648189210 | 0.450257250956213 | 0.636 | 0.482 | 1.23E-34 | facsFB | MA0640.2 |
| 9.635545844 | 0.448320840717079 | 0.671 | 0.496 | 7.19E-36 | facsFB | MA0791.1 |
| 2.185595142 | 0.447910943784542 | 0.619 | 0.49 | 1.63E-29 | facsFB | MA0774.1 |
| 1.188639312 | 0.444246516020244 | 0.61 | 0.487 | 8.87E-27 | facsFB | MA0063.2 |
| 1.713300657 | 0.443289474985682 | 0.618 | 0.491 | 1.28E-27 | facsFB | MA0699.1 |
| 3.861379239 | 0.441257331558384 | 0.634 | 0.483 | 2.88E-28 | facsFB | MA0744.2 |
| 3.297235970 | 0.440534068189575 | 0.641 | 0.486 | 2.46E-31 | facsFB | MA0685.1 |
| 1.506294161 | 0.439843948829426 | 0.648 | 0.497 | 1.12E-30 | facsFB | MA1132.1 |
| 5.867978671 | 0.434963552405597 | 0.619 | 0.493 | 4.38E-28 | facsFB | MA1575.1 |
| 2.644544681 | 0.43281481410072 | 0.617 | 0.481 | 1.97E-27 | facsFB | MA0696.1 |
| 5.826221860 | 0.43114471346942 | 0.642 | 0.484 | 4.35E-26 | facsFB | MA0609.2 |
| 1.167897598 | 0.427080208289745 | 0.631 | 0.486 | 8.71E-31 | facsFB | MA1508.1 |
| 1.886269437 | 0.417637857408891 | 0.627 | 0.5 | 1.41E-24 | facsFB | MA0627.2 |
| 7.837891340 | 0.414609961657611 | 0.642 | 0.488 | 5.85E-34 | facsFB | MA1513.1 |
| 1.311233931 | 0.412496364984653 | 0.601 | 0.475 | 9.78E-21 | facsFB | MA0882.1 |

| | | | | | | |
|---|---|---|---|---|---|---|
| 1.08312827 | 0.41110382784861 | 0.646 | 0.484 | 8.08E-34 | facsFB | MA0741.1 |
| 1.40546905 | 0.40923075480986 | 0.609 | 0.488 | 1.05E-26 | facsFB | MA0598.3 |
| 9.63157414 | 0.40867407056031 | 0.603 | 0.479 | 7.19E-24 | facsFB | MA0750.2 |
| 1.76458967 | 0.398368757336485 | 0.616 | 0.479 | 1.32E-21 | facsFB | MA1554.1 |
| 8.44291877 | 0.3959223970674 | 0.6 | 0.486 | 6.30E-22 | facsFB | MA0840.1 |
| 4.38502120 | 0.388896405835484 | 0.636 | 0.493 | 3.27E-24 | facsFB | MA1101.2 |
| 6.06430164 | 0.388584757447532 | 0.606 | 0.486 | 4.52E-21 | facsFB | MA0488.1 |
| 9.40098891 | 0.384740471780535 | 0.602 | 0.487 | 7.01E-19 | facsFB | MA0657.1 |
| 2.72994287 | 0.380436439821553 | 0.598 | 0.482 | 2.04E-19 | facsFB | MA0656.1 |
| 1.54856484 | 0.379570908374651 | 0.613 | 0.489 | 1.16E-21 | facsFB | MA0475.2 |
| 5.08432643 | 0.375640119785157 | 0.591 | 0.488 | 3.79E-18 | facsFB | MA0029.1 |
| 5.59664782 | 0.374313353567876 | 0.63 | 0.494 | 4.18E-23 | facsFB | MA0462.2 |
| 5.08852822 | 0.372024528401135 | 0.619 | 0.494 | 3.80E-19 | facsFB | MA0823.1 |
| 1.84794161 | 0.368859757637965 | 0.636 | 0.491 | 1.38E-23 | facsFB | MA1125.1 |
| 6.71313612 | 0.366704065141504 | 0.617 | 0.491 | 5.01E-21 | facsFB | MA0769.2 |
| 3.01111455 | 0.363890606215904 | 0.641 | 0.501 | 2.25E-20 | facsFB | MA0472.2 |
| 3.81002996 | 0.361867355170262 | 0.588 | 0.473 | 2.84E-16 | facsFB | MA0711.1 |
| 3.07689684 | 0.360512802949036 | 0.629 | 0.5 | 2.30E-23 | facsFB | MA1596.1 |
| 1.87265496 | 0.357345779128363 | 0.611 | 0.494 | 1.40E-17 | facsFB | MA0894.1 |
| 2.74602846 | 0.356351439354037 | 0.585 | 0.481 | 2.05E-16 | facsFB | MA0056.2 |
| 4.72293573 | 0.355031709880249 | 0.577 | 0.491 | 3.52E-16 | facsFB | MA0630.1 |
| 3.18487361 | 0.352612915266536 | 0.589 | 0.489 | 2.38E-15 | facsFB | MA0829.2 |
| 1.42503937 | 0.352321989780574 | 0.624 | 0.498 | 1.06E-20 | facsFB | MA1142.1 |
| 2.32414584 | 0.351982853577985 | 0.578 | 0.48 | 1.73E-15 | facsFB | MA0594.2 |
| 2.16547311 | 0.351334589457186 | 0.608 | 0.505 | 1.62E-15 | facsFB | MA1154.1 |
| 9.15127716 | 0.351323725008537 | 0.586 | 0.485 | 6.83E-16 | facsFB | MA0037.3 |
| 1.67553530 | 0.35030664986847 | 0.609 | 0.496 | 1.25E-19 | facsFB | MA1634.1 |
| 2.34643191 | 0.35025359770929 | 0.598 | 0.491 | 1.75E-16 | facsFB | MA0066.1 |
| 6.31917937 | 0.349739478709859 | 0.607 | 0.49 | 4.71E-15 | facsFB | MA0641.1 |
| 4.16528445 | 0.347396585025464 | 0.612 | 0.49 | 3.11E-20 | facsFB | MA0081.2 |
| 1.22313773 | 0.344717849407895 | 0.59 | 0.488 | 9.12E-16 | facsFB | MA0716.1 |
| 1.22313773 | 0.344717849407895 | 0.59 | 0.488 | 9.12E-16 | facsFB | MA0721.1 |
| 1.93698057 | 0.343764559097986 | 0.609 | 0.502 | 1.44E-15 | facsFB | MA0831.2 |
| 7.84879240 | 0.33971034732947 | 0.6 | 0.495 | 5.86E-16 | facsFB | MA0608.1 |
| 4.41613658 | 0.339446225554842 | 0.593 | 0.488 | 3.29E-15 | facsFB | MA0605.2 |
| 1.89512034 | 0.339213630286305 | 0.6 | 0.491 | 1.41E-16 | facsFB | MA0697.1 |
| 2.10515446 | 0.336700401206959 | 0.585 | 0.483 | 1.57E-16 | facsFB | MA0079.4 |
| 9.13569967 | 0.335717200949818 | 0.595 | 0.475 | 6.82E-15 | facsFB | MA1545.1 |
| 1.38647206 | 0.334028315237326 | 0.599 | 0.494 | 1.03E-15 | facsFB | MA0114.4 |
| 5.44297074 | 0.333599811437594 | 0.594 | 0.486 | 4.06E-15 | facsFB | MA0751.1 |
| 4.07638469 | 0.330161305340179 | 0.602 | 0.496 | 3.04E-15 | facsFB | MA0595.1 |
| 9.40475164 | 0.329954570956013 | 0.6 | 0.493 | 7.02E-16 | facsFB | MA1581.1 |
| 9.56090182 | 0.328360761743967 | 0.584 | 0.483 | 7.13E-13 | facsFB | MA0700.2 |

| | | | | | | |
|---|---|---|---|---|---|---|
| 7.99449030 | 0.328249254763151 | 0.588 | 0.496 | 5.96E-14 | facsFB | MA0482.2 |
| 1.15329745 | 0.327730985123576 | 0.596 | 0.492 | 8.60E-15 | facsFB | MA0098.3 |
| 4.86041299 | 0.326229998883869 | 0.596 | 0.487 | 3.63E-14 | facsFB | MA0492.1 |
| 1.67561277 | 0.326128939610742 | 0.602 | 0.488 | 1.25E-13 | facsFB | MA1619.1 |
| 1.76158970 | 0.324537074200227 | 0.596 | 0.493 | 1.31E-14 | facsFB | MA0671.1 |
| 1.23748090 | 0.324504771294615 | 0.588 | 0.493 | 9.23E-13 | facsFB | MA0845.1 |
| 2.42710210 | 0.323961477528662 | 0.615 | 0.499 | 1.81E-15 | facsFB | MA0507.1 |
| 4.50366398 | 0.322659539999678 | 0.59 | 0.491 | 3.36E-14 | facsFB | MA0484.2 |
| 4.74829961 | 0.318453426451202 | 0.583 | 0.485 | 3.54E-12 | facsFB | MA0111.1 |
| 1.19115886 | 0.317778595329375 | 0.6 | 0.486 | 8.89E-13 | facsFB | MA0893.2 |
| 1.19115886 | 0.317778595329375 | 0.6 | 0.486 | 8.89E-13 | facsFB | MA0912.2 |
| 2.57296855 | 0.317209107065119 | 0.582 | 0.488 | 1.92E-13 | facsFB | MA0607.1 |
| 1.95346305 | 0.315853122329824 | 0.602 | 0.489 | 1.46E-14 | facsFB | MA0258.2 |
| 2.87960872 | 0.315776094174956 | 0.597 | 0.491 | 2.15E-14 | facsFB | MA0642.1 |
| 4.20314498 | 0.313763563185553 | 0.587 | 0.491 | 3.14E-13 | facsFB | MA1657.1 |
| 6.79739321 | 0.313677150616398 | 0.587 | 0.5 | 5.07E-13 | facsFB | MA0761.2 |
| 2.58154008 | 0.311523450058835 | 0.568 | 0.488 | 1.93E-11 | facsFB | MA0109.1 |
| 4.21359705 | 0.311133662802814 | 0.598 | 0.5 | 3.14E-12 | facsFB | MA0871.2 |
| 7.83677541 | 0.308557328855552 | 0.589 | 0.49 | 5.85E-13 | facsFB | MA0860.1 |
| 6.71946578 | 0.306082779732294 | 0.593 | 0.49 | 5.01E-12 | facsFB | MA0634.1 |
| 3.11012720 | 0.304819684786596 | 0.574 | 0.49 | 2.32E-11 | facsFB | MA1124.1 |
| 8.29297996 | 0.304380726517583 | 0.601 | 0.493 | 6.19E-13 | facsFB | MA1103.2 |
| 1.34290201 | 0.304229467528383 | 0.596 | 0.495 | 1.00E-12 | facsFB | MA0032.2 |
| 1.86421198 | 0.303211620880358 | 0.589 | 0.486 | 1.39E-12 | facsFB | MA0651.1 |
| 5.82200059 | 0.302944661837205 | 0.589 | 0.495 | 4.34E-11 | facsFB | MA1651.1 |
| 5.12473232 | 0.301003999654756 | 0.585 | 0.485 | 3.82E-11 | facsFB | MA0074.1 |
| 4.99824904 | 0.299989702928061 | 0.591 | 0.49 | 3.73E-12 | facsFB | MA0156.2 |
| 5.27667472 | 0.299189585285613 | 0.59 | 0.489 | 3.94E-13 | facsFB | MA0496.3 |
| 1.65304165 | 0.298681121508397 | 0.586 | 0.487 | 1.23E-11 | facsFB | MA0076.2 |
| 7.83889681 | 0.297979210739523 | 0.585 | 0.49 | 5.85E-12 | facsFB | MA1587.1 |
| 7.86782166 | 0.29778272092398 | 0.569 | 0.489 | 5.87E-13 | facsFB | MA0790.1 |
| 7.54244270 | 0.297605822276257 | 0.59 | 0.492 | 5.63E-12 | facsFB | MA0757.1 |
| 2.17315651 | 0.297369977313375 | 0.605 | 0.496 | 1.62E-12 | facsFB | MA0077.1 |
| 2.75829140 | 0.297263514682076 | 0.575 | 0.49 | 2.06E-09 | facsFB | MA0646.1 |
| 1.00811923 | 0.29644100573545 | 0.577 | 0.486 | 7.52E-12 | facsFB | MA0027.2 |
| 1.02666679 | 0.294480062090043 | 0.573 | 0.492 | 7.66E-11 | facsFB | MA0474.2 |
| 3.73807317 | 0.294311558077694 | 0.609 | 0.498 | 2.79E-14 | facsFB | MA0491.2 |
| 6.79971396 | 0.292578277752376 | 0.574 | 0.483 | 5.07E-11 | facsFB | MA0509.2 |
| 1.73753903 | 0.290430814624849 | 0.573 | 0.488 | 1.30E-10 | facsFB | MA1656.1 |
| 4.84035233 | 0.29020080169032 | 0.581 | 0.492 | 3.61E-11 | facsFB | MA1683.1 |
| 4.51380436 | 0.288920771851733 | 0.589 | 0.495 | 3.37E-11 | facsFB | MA0106.3 |
| 2.07211799 | 0.287320667200873 | 0.561 | 0.48 | 1.55E-10 | facsFB | MA1495.1 |
| 2.07211799 | 0.287320667200873 | 0.561 | 0.48 | 1.55E-10 | facsFB | MA0904.2 |

| | | | | | | |
|---|---|---|---|---|---|---|
| 2.93905214 | 0.286509816861931 | 0.589 | 0.499 | 2.19E-10 | facsFB | MA1115.1 |
| 4.81453234 | 0.286060274228082 | 0.566 | 0.488 | 3.59E-10 | facsFB | MA1607.1 |
| 1.83930714 | 0.284810903757281 | 0.577 | 0.488 | 1.37E-10 | facsFB | MA1632.1 |
| 3.14066362 | 0.283845458677962 | 0.567 | 0.486 | 2.34E-07 | facsFB | MA0877.2 |
| 2.25426132 | 0.283051378412918 | 0.573 | 0.497 | 1.68E-08 | facsFB | MA1572.1 |
| 2.79968649 | 0.282037746442303 | 0.601 | 0.489 | 2.09E-13 | facsFB | MA0516.2 |
| 8.03748911 | 0.281965426671685 | 0.571 | 0.483 | 6.00E-10 | facsFB | MA1105.2 |
| 3.66857452 | 0.281240324169459 | 0.585 | 0.493 | 2.74E-10 | facsFB | MA0155.1 |
| 2.03377688 | 0.280726744019397 | 0.573 | 0.486 | 1.52E-08 | facsFB | MA1594.1 |
| 1.45652036 | 0.276999807728899 | 0.58 | 0.498 | 1.09E-10 | facsFB | MA0835.2 |
| 1.95863524 | 0.276246254427698 | 0.55 | 0.475 | 1.46E-08 | facsFB | MA1515.1 |
| 5.07296434 | 0.27469714059387 | 0.57 | 0.489 | 3.78E-08 | facsFB | MA0009.2 |
| 2.15546936 | 0.274499227902113 | 0.603 | 0.5 | 1.61E-13 | facsFB | MA0151.1 |
| 2.67194633 | 0.27341594984546 | 0.576 | 0.495 | 1.99E-09 | facsFB | MA0649.1 |
| 1.42362088 | 0.272657651653857 | 0.568 | 0.489 | 1.06E-08 | facsFB | MA1641.1 |
| 8.41991381 | 0.271949897473705 | 0.573 | 0.487 | 6.28E-09 | facsFB | MA0662.1 |
| 3.44628221 | 0.271395043021597 | 0.593 | 0.498 | 2.57E-11 | facsFB | MA0521.1 |
| 1.05537116 | 0.267827847387582 | 0.556 | 0.486 | 7.87E-07 | facsFB | MA1131.1 |
| 3.20870121 | 0.265984486263235 | 0.569 | 0.492 | 2.39E-09 | facsFB | MA0503.1 |
| 5.64509442 | 0.263919550585037 | 0.552 | 0.486 | 4.21E-07 | facsFB | MA1603.1 |
| 5.13301197 | 0.263641512013923 | 0.596 | 0.493 | 3.83E-10 | facsFB | MA0047.3 |
| 9.09887856 | 0.261626892968493 | 0.589 | 0.498 | 6.79E-08 | facsFB | MA0616.2 |
| 3.05650468 | 0.258279881227458 | 0.587 | 0.501 | 2.28E-09 | facsFB | MA0062.3 |
| 1.94863768 | 0.257623019086899 | 0.589 | 0.499 | 1.45E-10 | facsFB | MA0149.1 |
| 3.36473067 | 0.257545890298867 | 0.589 | 0.504 | 2.51E-09 | facsFB | MA0782.2 |
| 6.58195635 | 0.257346997595146 | 0.553 | 0.489 | 4.91E-08 | facsFB | MA1472.1 |
| 2.24674270 | 0.257310375837593 | 0.568 | 0.489 | 1.68E-08 | facsFB | MA1621.1 |
| 8.74366538 | 0.255789395524366 | 0.557 | 0.497 | 6.52E-07 | facsFB | MA0861.1 |
| 1.07924336 | 0.252924072510545 | 0.549 | 0.485 | 8.05E-07 | facsFB | MA1547.1 |
| 6.73922648 | 0.252660876223958 | 0.579 | 0.49 | 5.03E-08 | facsFB | MA0525.2 |
| 2.00258387 | 0.250619889607036 | 0.567 | 0.493 | 1.49E-07 | facsFB | MA0760.1 |
| 0 | 0.906534418304133 | 0.764 | 0.46 | 0.00 | MYE | MA0748.2 |
| 0 | 0.874454546772652 | 0.741 | 0.465 | 0.00 | MYE | MA0639.1 |
| 0 | 0.84719632640224 | 0.771 | 0.475 | 0.00 | MYE | MA0137.3 |
| 0 | 0.816875697527332 | 0.732 | 0.467 | 0.00 | MYE | MA0907.1 |
| 2.49581454 | 0.80609374807374 | 0.732 | 0.462 | 1.86E-301 | MYE | MA0466.2 |
| 8.43048019 | 0.805780925002299 | 0.723 | 0.466 | 6.29E-299 | MYE | MA0833.2 |
| 1.10967553 | 0.798731680132939 | 0.734 | 0.474 | 8.28E-288 | MYE | MA0821.1 |
| 2.58078431 | 0.79699232460387 | 0.726 | 0.468 | 1.93E-298 | MYE | MA0710.1 |
| 0 | 0.782818305213845 | 0.765 | 0.481 | 0.00 | MYE | MA1116.1 |
| 2.12911130 | 0.781244810213173 | 0.733 | 0.471 | 1.59E-290 | MYE | MA0879.1 |
| 1.19747811 | 0.753755484419924 | 0.713 | 0.463 | 8.93E-258 | MYE | MA0837.1 |
| 2.60599113 | 0.737803155223675 | 0.714 | 0.47 | 1.94E-236 | MYE | MA0822.1 |

| | | | | | | |
|---|---|---|---|---|---|---|
| 1.13504187 | 0.697037027080643 | 0.692 | 0.468 | 8.47E-216 | MYE | MA1577.1 |
| 1.26177605 | 0.683567417586789 | 0.713 | 0.483 | 9.41E-239 | MYE | MA0152.1 |
| 1.41170219 | 0.680304206455815 | 0.71 | 0.478 | 1.05E-230 | MYE | MA1654.1 |
| 5.97796113 | 0.667681393053997 | 0.713 | 0.481 | 4.46E-211 | MYE | MA0884.1 |
| 2.39639575 | 0.663455644120349 | 0.681 | 0.474 | 1.79E-194 | MYE | MA0817.1 |
| 5.82034120 | 0.654020770210638 | 0.684 | 0.476 | 4.34E-182 | MYE | MA0680.1 |
| 6.05691079 | 0.651072116231585 | 0.696 | 0.477 | 4.52E-198 | MYE | MA0625.1 |
| 1.25203946 | 0.647593879845438 | 0.686 | 0.475 | 9.34E-184 | MYE | MA0068.2 |
| 4.28776275 | 0.625845243181024 | 0.673 | 0.471 | 3.20E-169 | MYE | MA0644.1 |
| 1.35915904 | 0.623451288752768 | 0.675 | 0.482 | 1.01E-174 | MYE | MA1580.1 |
| 2.45087914 | 0.611616635090362 | 0.681 | 0.48 | 1.83E-170 | MYE | MA0854.1 |
| 8.13383949 | 0.605959067858238 | 0.677 | 0.476 | 6.07E-169 | MYE | MA1480.1 |
| 2.03968951 | 0.603917121408225 | 0.693 | 0.486 | 1.52E-185 | MYE | MA0442.2 |
| 2.20556216 | 0.598884691294375 | 0.676 | 0.482 | 1.65E-165 | MYE | MA1112.2 |
| 9.47625035 | 0.596129737238049 | 0.662 | 0.479 | 7.07E-155 | MYE | MA1500.1 |
| 5.28251499 | 0.59273669523952 | 0.687 | 0.488 | 3.94E-167 | MYE | MA1154.1 |
| 6.32121492 | 0.590923324004093 | 0.674 | 0.477 | 4.72E-158 | MYE | MA0075.3 |
| 1.15325328 | 0.586721910709761 | 0.655 | 0.472 | 8.60E-148 | MYE | MA0838.1 |
| 3.04812534 | 0.585950019703789 | 0.688 | 0.486 | 2.27E-162 | MYE | MA0088.2 |
| 4.49108934 | 0.5746148441138 | 0.67 | 0.481 | 3.35E-153 | MYE | MA0084.1 |
| 1.55612778 | 0.567594954971029 | 0.673 | 0.485 | 1.16E-149 | MYE | MA1525.1 |
| 4.02785759 | 0.563696706650403 | 0.677 | 0.49 | 3.00E-150 | MYE | MA1597.1 |
| 1.22567290 | 0.561920987112276 | 0.666 | 0.473 | 9.14E-143 | MYE | MA0718.1 |
| 3.45558042 | 0.551255576148733 | 0.667 | 0.477 | 2.58E-147 | MYE | MA1123.2 |
| 2.47597229 | 0.548969711385353 | 0.666 | 0.479 | 1.85E-145 | MYE | MA0508.3 |
| 1.80632418 | 0.54358985096448 | 0.651 | 0.478 | 1.35E-124 | MYE | MA0699.1 |
| 4.19476687 | 0.543070761914625 | 0.66 | 0.48 | 3.13E-132 | MYE | MA0140.2 |
| 2.44597444 | 0.540405750276556 | 0.661 | 0.484 | 1.82E-126 | MYE | MA0863.1 |
| 3.30244596 | 0.531414797659186 | 0.665 | 0.486 | 2.46E-131 | MYE | MA0527.1 |
| 2.62402510 | 0.528788494062739 | 0.649 | 0.479 | 1.96E-117 | MYE | MA0674.1 |
| 1.29986839 | 0.52715263535626 | 0.665 | 0.485 | 9.70E-128 | MYE | MA1106.1 |
| 1.55672207 | 0.519979285798198 | 0.653 | 0.487 | 1.16E-112 | MYE | MA0914.1 |
| 1.23923964 | 0.519026780616929 | 0.64 | 0.476 | 9.24E-121 | MYE | MA0686.1 |
| 4.40149347 | 0.504556290845092 | 0.666 | 0.5 | 3.28E-111 | MYE | MA0526.3 |
| 5.03515602 | 0.503251170401176 | 0.661 | 0.5 | 3.76E-110 | MYE | MA0093.3 |
| 8.72589599 | 0.502544107107735 | 0.647 | 0.484 | 6.51E-104 | MYE | MA0875.1 |
| 3.73138340 | 0.501889345092693 | 0.639 | 0.478 | 2.78E-107 | MYE | MA0630.1 |
| 5.38323067 | 0.500775872039869 | 0.653 | 0.488 | 4.02E-109 | MYE | MA1524.1 |
| 2.20118907 | 0.490674739394863 | 0.647 | 0.479 | 1.64E-112 | MYE | MA0518.1 |
| 2.82692250 | 0.489536372206883 | 0.653 | 0.486 | 2.11E-110 | MYE | MA0119.1 |
| 5.54749380 | 0.487783206196466 | 0.63 | 0.483 | 4.14E-98 | MYE | MA1504.1 |
| 3.68621463 | 0.485919339775635 | 0.632 | 0.48 | 2.75E-98 | MYE | MA1463.1 |
| 3.78093778 | 0.483897409970943 | 0.644 | 0.483 | 2.82E-107 | MYE | MA1467.1 |

| | | | | | | |
|---|---|---|---|---|---|---|
| 1.52302998 | 0.483599254570167 | 0.643 | 0.491 | 1.14E-103 | MYE | MA1528.1 |
| 1.21559395 | 0.481503945979353 | 0.641 | 0.487 | 9.07E-103 | MYE | MA1111.1 |
| 7.74513872 | 0.47916322370802 | 0.631 | 0.483 | 5.78E-97 | MYE | MA0853.1 |
| 6.45480303 | 0.477738845600575 | 0.638 | 0.486 | 4.82E-100 | MYE | MA0512.2 |
| 4.50567797 | 0.470570723271169 | 0.641 | 0.487 | 3.36E-97 | MYE | MA0865.1 |
| 1.14137939 | 0.457384113071183 | 0.653 | 0.489 | 8.51E-101 | MYE | MA1604.1 |
| 5.44230433 | 0.45658606107734 | 0.633 | 0.483 | 4.06E-95 | MYE | MA0707.1 |
| 2.93654011 | 0.455053684459817 | 0.617 | 0.479 | 2.19E-87 | MYE | MA1119.1 |
| 9.98692077 | 0.453662951791994 | 0.63 | 0.484 | 7.45E-91 | MYE | MA0606.1 |
| 6.74764508 | 0.452002517662054 | 0.639 | 0.486 | 5.03E-92 | MYE | MA0524.2 |
| 6.92082326 | 0.450913753090828 | 0.642 | 0.486 | 5.16E-97 | MYE | MA1418.1 |
| 2.76916176 | 0.450664955082077 | 0.618 | 0.478 | 2.07E-82 | MYE | MA0843.1 |
| 3.23535975 | 0.448610989996562 | 0.627 | 0.483 | 2.41E-90 | MYE | MA0624.1 |
| 3.29584781 | 0.448544790567148 | 0.623 | 0.479 | 2.46E-82 | MYE | MA0604.1 |
| 1.47910107 | 0.44662660991532 | 0.602 | 0.466 | 1.10E-79 | MYE | MA0776.1 |
| 8.51904113 | 0.440529215850901 | 0.617 | 0.486 | 6.36E-80 | MYE | MA0828.1 |
| 6.10206586 | 0.436174483573227 | 0.618 | 0.483 | 4.55E-78 | MYE | MA0719.1 |
| 1.33923415 | 0.434572581674778 | 0.638 | 0.49 | 9.99E-85 | MYE | MA0620.3 |
| 2.55066854 | 0.432311172213789 | 0.643 | 0.484 | 1.90E-88 | MYE | MA1638.1 |
| 9.95582146 | 0.430731510986288 | 0.631 | 0.482 | 7.43E-83 | MYE | MA0122.3 |
| 3.46151540 | 0.430640607774649 | 0.616 | 0.479 | 2.58E-78 | MYE | MA0874.1 |
| 1.66927967 | 0.428131867797095 | 0.643 | 0.492 | 1.25E-88 | MYE | MA0619.1 |
| 4.37934854 | 0.420953402713753 | 0.612 | 0.477 | 3.27E-74 | MYE | MA0892.1 |
| 1.47078178 | 0.415680395884429 | 0.606 | 0.476 | 1.10E-71 | MYE | MA0883.1 |
| 1.97679258 | 0.414617095670101 | 0.613 | 0.488 | 1.47E-65 | MYE | MA0780.1 |
| 2.02028259 | 0.412060113196859 | 0.618 | 0.484 | 1.51E-74 | MYE | MA0141.3 |
| 7.66742436 | 0.41026697339835 | 0.602 | 0.473 | 5.72E-69 | MYE | MA0897.1 |
| 2.63802908 | 0.410245803944664 | 0.602 | 0.48 | 1.97E-65 | MYE | MA0889.1 |
| 4.36586716 | 0.407212417449575 | 0.621 | 0.487 | 3.26E-73 | MYE | MA0048.2 |
| 1.46786551 | 0.405213459247716 | 0.611 | 0.483 | 1.10E-66 | MYE | MA0648.1 |
| 4.14732221 | 0.404848043368933 | 0.614 | 0.478 | 3.09E-72 | MYE | MA0667.1 |
| 2.29315244 | 0.404122395548651 | 0.6 | 0.48 | 1.71E-62 | MYE | MA1497.1 |
| 1.27929560 | 0.403822038030764 | 0.607 | 0.485 | 9.54E-65 | MYE | MA1140.2 |
| 3.18557879 | 0.402411556838996 | 0.623 | 0.489 | 2.38E-66 | MYE | MA0756.1 |
| 5.71876554 | 0.40050327027849 | 0.617 | 0.481 | 4.27E-68 | MYE | MA0703.2 |
| 6.99646502 | 0.396485861982221 | 0.612 | 0.483 | 5.22E-69 | MYE | MA0621.1 |
| 1.65379189 | 0.39457451736317 | 0.619 | 0.484 | 1.23E-65 | MYE | MA0259.1 |
| 1.56375895 | 0.391367041960749 | 0.614 | 0.478 | 1.17E-64 | MYE | MA0827.1 |
| 6.30585140 | 0.38944851894907 | 0.633 | 0.491 | 4.70E-71 | MYE | MA0514.1 |
| 6.20850152 | 0.385658471340794 | 0.625 | 0.493 | 4.63E-69 | MYE | MA1624.1 |
| 1.51548398 | 0.384561081708613 | 0.6 | 0.479 | 1.13E-61 | MYE | MA0492.1 |
| 1.15183845 | 0.384090788238266 | 0.604 | 0.481 | 8.59E-60 | MYE | MA0654.1 |
| 1.15183845 | 0.384090788238266 | 0.604 | 0.481 | 8.59E-60 | MYE | MA0720.1 |

| | | | | | | |
|---|---|---|---|---|---|---|
| 1.15183845 | 0.384090788238266 | 0.604 | 0.481 | 8.59E-60 | MYE | MA0701. |
| 5.26885367 | 0.377256288953636 | 0.624 | 0.49 | 3.93E-67 | MYE | MA0761. |
| 3.73979685 | 0.37605511004973 | 0.614 | 0.485 | 2.79E-59 | MYE | MA0823.1 |
| 8.33073824 | 0.370348009946625 | 0.622 | 0.5 | 6.21E-59 | MYE | MA0692.1 |
| 1.29849128 | 0.368648170295123 | 0.625 | 0.489 | 9.69E-73 | MYE | MA0149.1 |
| 5.73979281 | 0.368134912185463 | 0.603 | 0.481 | 4.28E-58 | MYE | MA1109.1 |
| 5.61136683 | 0.367855084386261 | 0.609 | 0.485 | 4.19E-58 | MYE | MA0712. |
| 3.35173690 | 0.367317127286877 | 0.611 | 0.489 | 2.50E-58 | MYE | MA0836. |
| 3.61593044 | 0.363503745373963 | 0.584 | 0.479 | 2.70E-51 | MYE | MA0662.1 |
| 8.54862389 | 0.363262178890428 | 0.599 | 0.488 | 6.38E-57 | MYE | MA0115.1 |
| 2.36825115 | 0.36286500399422 | 0.596 | 0.483 | 1.77E-54 | MYE | MA0735.1 |
| 5.84242931 | 0.360060953285512 | 0.603 | 0.488 | 4.36E-53 | MYE | MA0595.1 |
| 9.08737165 | 0.359359478923127 | 0.611 | 0.484 | 6.78E-61 | MYE | MA0515.1 |
| 4.03206480 | 0.358720498247117 | 0.603 | 0.486 | 3.01E-56 | MYE | MA0150. |
| 3.35027011 | 0.358294185389811 | 0.596 | 0.479 | 2.50E-53 | MYE | MA0763.1 |
| 9.11069931 | 0.357433422917105 | 0.593 | 0.479 | 6.80E-51 | MYE | MA0714.1 |
| 1.71984363 | 0.352248247048946 | 0.587 | 0.48 | 1.28E-45 | MYE | MA0895.1 |
| 1.30714825 | 0.350962427290608 | 0.608 | 0.489 | 9.75E-53 | MYE | MA0866.1 |
| 1.31044292 | 0.349508360286993 | 0.59 | 0.478 | 9.78E-47 | MYE | MA0669.1 |
| 6.01535665 | 0.34845273897228 | 0.61 | 0.487 | 4.49E-56 | MYE | MA0003. |
| 9.09618303 | 0.348422085931226 | 0.602 | 0.485 | 6.79E-53 | MYE | MA0592. |
| 4.16424406 | 0.345782202088029 | 0.592 | 0.487 | 3.11E-50 | MYE | MA1520.1 |
| 6.24293722 | 0.345631771185195 | 0.585 | 0.485 | 4.66E-48 | MYE | MA0890.1 |
| 2.34853635 | 0.345056540890477 | 0.602 | 0.49 | 1.75E-52 | MYE | MA0677.1 |
| 6.84670474 | 0.343367508133039 | 0.599 | 0.484 | 5.11E-48 | MYE | MA0717.1 |
| 1.60756662 | 0.343196926694451 | 0.597 | 0.483 | 1.20E-49 | MYE | MA0628.1 |
| 3.30767498 | 0.341547543556652 | 0.603 | 0.49 | 2.47E-50 | MYE | MA0463. |
| 2.48920104 | 0.341460843855533 | 0.594 | 0.49 | 1.86E-48 | MYE | MA1579.1 |
| 2.98142656 | 0.340696485486431 | 0.594 | 0.487 | 2.22E-47 | MYE | MA0759.1 |
| 6.01512896 | 0.339969636684672 | 0.573 | 0.475 | 4.49E-41 | MYE | MA0668.1 |
| 1.30591989 | 0.338131695504781 | 0.587 | 0.478 | 9.74E-46 | MYE | MA1603.1 |
| 1.93544871 | 0.335614956998001 | 0.6 | 0.493 | 1.44E-49 | MYE | MA0519.1 |
| 2.91817558 | 0.335008112503215 | 0.587 | 0.483 | 2.18E-44 | MYE | MA0101.1 |
| 4.19299816 | 0.334640878888739 | 0.583 | 0.481 | 3.13E-42 | MYE | MA0886.1 |
| 3.49726506 | 0.331855910880245 | 0.593 | 0.486 | 2.61E-45 | MYE | MA0658.1 |
| 3.45247291 | 0.330276372341502 | 0.599 | 0.486 | 2.58E-50 | MYE | MA1101. |
| 2.72913998 | 0.323020704721369 | 0.588 | 0.484 | 2.04E-40 | MYE | MA0709.1 |
| 2.24169749 | 0.322593308845161 | 0.581 | 0.488 | 1.67E-41 | MYE | MA0160.1 |
| 9.84744246 | 0.321644325752717 | 0.581 | 0.486 | 7.35E-40 | MYE | MA1420.1 |
| 8.68300635 | 0.319524409577762 | 0.602 | 0.493 | 6.48E-44 | MYE | MA1466.1 |
| 1.50320552 | 0.317432093026068 | 0.597 | 0.494 | 1.12E-45 | MYE | MA1134.1 |
| 1.29182218 | 0.316246768004602 | 0.579 | 0.479 | 9.64E-40 | MYE | MA0877. |
| 2.82481995 | 0.313993595817768 | 0.576 | 0.481 | 2.11E-36 | MYE | MA0682. |

| | | | | | | |
|---|---|---|---|---|---|---|
| 7.374791585 | 0.313293631458952 | 0.585 | 0.488 | 5.50E-38 | MYE | MA0804.1 |
| 6.033021179 | 0.313174878750716 | 0.588 | 0.487 | 4.50E-43 | MYE | MA1601.1 |
| 6.862773440 | 0.312483900436673 | 0.589 | 0.491 | 5.12E-39 | MYE | MA1491.1 |
| 1.812776865 | 0.309265695615682 | 0.585 | 0.487 | 1.35E-40 | MYE | MA0144.2 |
| 4.337752799 | 0.306051031130627 | 0.581 | 0.482 | 3.24E-37 | MYE | MA0009.2 |
| 5.107987448 | 0.301686084217605 | 0.578 | 0.485 | 3.81E-36 | MYE | MA0643.1 |
| 2.241553757 | 0.301268644685579 | 0.603 | 0.494 | 1.67E-44 | MYE | MA0052.4 |
| 1.563461931 | 0.300665317325522 | 0.595 | 0.49 | 1.17E-37 | MYE | MA1588.1 |
| 3.506038854 | 0.297641330848829 | 0.596 | 0.499 | 2.62E-36 | MYE | MA0142.1 |
| 9.056273280 | 0.297370743459906 | 0.583 | 0.495 | 6.76E-36 | MYE | MA0143.4 |
| 1.574132651 | 0.297295290788304 | 0.596 | 0.49 | 1.17E-41 | MYE | MA0521.1 |
| 1.057972775 | 0.2968370126221 | 0.569 | 0.482 | 7.89E-35 | MYE | MA0908.1 |
| 1.481168458 | 0.296063327551267 | 0.571 | 0.485 | 1.10E-33 | MYE | MA0704.1 |
| 3.951877779 | 0.289380040053191 | 0.589 | 0.493 | 2.95E-35 | MYE | MA1635.1 |
| 7.435462719 | 0.287805952726066 | 0.578 | 0.486 | 5.55E-35 | MYE | MA0131.2 |
| 1.595088773 | 0.286111404152766 | 0.572 | 0.482 | 1.19E-29 | MYE | MA0724.1 |
| 1.014361909 | 0.285080460943157 | 0.569 | 0.488 | 7.57E-32 | MYE | MA0092.1 |
| 3.753841330 | 0.282594365853849 | 0.564 | 0.475 | 2.80E-30 | MYE | MA1568.1 |
| 2.749067217 | 0.27944056876939 | 0.588 | 0.495 | 2.05E-32 | MYE | MA0017.2 |
| 5.542470190 | 0.277590512831911 | 0.587 | 0.495 | 4.13E-32 | MYE | MA1616.1 |
| 4.877435460 | 0.27605818101759 | 0.581 | 0.485 | 3.64E-36 | MYE | MA0130.1 |
| 8.718911590 | 0.273954941593333 | 0.588 | 0.493 | 6.50E-34 | MYE | MA0099.3 |
| 3.022931605 | 0.27357091953078 | 0.58 | 0.492 | 2.26E-31 | MYE | MA0650.2 |
| 2.637332213 | 0.273338150705896 | 0.564 | 0.483 | 1.97E-27 | MYE | MA0911.1 |
| 6.860872403 | 0.272488637014285 | 0.581 | 0.49 | 5.12E-31 | MYE | MA1421.1 |
| 3.254353574 | 0.267513427795821 | 0.584 | 0.491 | 2.43E-32 | MYE | MA1130.1 |
| 5.285651125 | 0.267123603605088 | 0.55 | 0.483 | 3.94E-27 | MYE | MA0891.1 |
| 1.483216027 | 0.266635092357356 | 0.587 | 0.497 | 1.11E-29 | MYE | MA0100.3 |
| 2.055506565 | 0.265994600263655 | 0.586 | 0.492 | 1.53E-31 | MYE | MA0480.1 |
| 9.685447509 | 0.26585072414441 | 0.572 | 0.49 | 7.23E-29 | MYE | MA0468.1 |
| 9.972737073 | 0.265773326061588 | 0.579 | 0.492 | 7.44E-31 | MYE | MA1144.1 |
| 6.246553470 | 0.26175880457584 | 0.579 | 0.492 | 4.66E-27 | MYE | MA0694.1 |
| 1.551112810 | 0.257861411159345 | 0.575 | 0.495 | 1.16E-28 | MYE | MA1628.1 |
| 9.985004798 | 0.25782354796284 | 0.578 | 0.496 | 7.45E-29 | MYE | MA0062.3 |
| 1.095622869 | 0.255322967450269 | 0.578 | 0.489 | 8.17E-28 | MYE | MA0779.1 |
| 1.888066134 | 0.255040512603908 | 0.559 | 0.486 | 1.41E-23 | MYE | MA0826.1 |
| 1.083455887 | 0.255028177806122 | 0.566 | 0.488 | 8.08E-26 | MYE | MA0473.3 |
| 4.229889062 | 0.254782058808677 | 0.578 | 0.492 | 3.16E-28 | MYE | MA1138.1 |
| 5.667050654 | 0.252963715890832 | 0.562 | 0.486 | 4.23E-22 | MYE | MA1139.1 |
| 2.009811509 | 0.251918200466258 | 0.593 | 0.497 | 1.50E-26 | MYE | MA1485.1 |
| 7.842816790 | 0.250426998214873 | 0.565 | 0.489 | 5.85E-24 | MYE | MA0671.1 |
| 6.447617260 | 0.752875407264307 | 0.755 | 0.491 | 4.81E-98 | sbasalKC | MA1639.1 |
| 4.768180837 | 0.648379631622451 | 0.704 | 0.494 | 3.56E-67 | sbasalKC | MA1419.1 |

| | | | | | | |
|---|---|---|---|---|---|---|
| 2.19943631 | 0.623219491600251 | 0.722 | 0.505 | 1.64E-67 | sbasalKC | MA0051.1 |
| 6.00462165 | 0.621755176946956 | 0.698 | 0.487 | 4.48E-63 | sbasalKC | MA1478.1 |
| 4.73744761 | 0.598215193815724 | 0.691 | 0.496 | 3.53E-58 | sbasalKC | MA0479.1 |
| 2.40592167 | 0.559589380526154 | 0.668 | 0.489 | 1.79E-48 | sbasalKC | MA1475.1 |
| 2.72463204 | 0.546117899523468 | 0.675 | 0.498 | 2.03E-45 | sbasalKC | MA0527.1 |
| 2.97108825 | 0.525820262132447 | 0.655 | 0.484 | 2.22E-40 | sbasalKC | MA1509.1 |
| 7.25518274 | 0.510959212082664 | 0.654 | 0.496 | 5.41E-41 | sbasalKC | MA1542.1 |
| 2.29975764 | 0.510956225551921 | 0.671 | 0.493 | 1.72E-42 | sbasalKC | MA0038.2 |
| 2.75375706 | 0.495163284081409 | 0.654 | 0.496 | 2.05E-38 | sbasalKC | MA0495.3 |
| 1.47955527 | 0.493222981102925 | 0.683 | 0.49 | 1.10E-43 | sbasalKC | MA1487.1 |
| 1.30255495 | 0.488684839992382 | 0.643 | 0.491 | 9.72E-33 | sbasalKC | MA1530.1 |
| 1.63315435 | 0.481758670868383 | 0.663 | 0.497 | 1.22E-34 | sbasalKC | MA0108.2 |
| 9.72854783 | 0.46728507450567 | 0.65 | 0.505 | 7.26E-32 | sbasalKC | MA0730.1 |
| 3.37195490 | 0.445575397183469 | 0.644 | 0.5 | 2.52E-29 | sbasalKC | MA0601.1 |
| 1.43495612 | 0.442123034160229 | 0.638 | 0.493 | 1.07E-28 | sbasalKC | MA0095.2 |
| 4.11068324 | 0.43349728547029 | 0.647 | 0.495 | 3.07E-32 | sbasalKC | MA0851.1 |
| 3.83408185 | 0.432034843549084 | 0.637 | 0.5 | 2.86E-28 | sbasalKC | MA0858.1 |
| 1.17176012 | 0.425984764430664 | 0.629 | 0.484 | 8.74E-27 | sbasalKC | MA1479.1 |
| 5.21867359 | 0.422189712935724 | 0.649 | 0.495 | 3.89E-32 | sbasalKC | MA0506.1 |
| 2.14631309 | 0.413455561202037 | 0.625 | 0.488 | 1.60E-23 | sbasalKC | MA0778.1 |
| 2.79050676 | 0.397037643324573 | 0.626 | 0.499 | 2.08E-23 | sbasalKC | MA0613.1 |
| 1.06036051 | 0.391450327868927 | 0.635 | 0.496 | 7.91E-24 | sbasalKC | MA0653.1 |
| 6.49967554 | 0.389016219408245 | 0.622 | 0.5 | 4.85E-23 | sbasalKC | MA0614.1 |
| 1.15637823 | 0.38633025825701 | 0.622 | 0.502 | 8.63E-22 | sbasalKC | MA1624.1 |
| 1.91842799 | 0.373337204699122 | 0.608 | 0.493 | 1.43E-18 | sbasalKC | MA0712.2 |
| 1.95662835 | 0.367488305672319 | 0.607 | 0.487 | 1.46E-17 | sbasalKC | MA0112.3 |
| 2.34108721 | 0.362792540078179 | 0.617 | 0.49 | 1.75E-19 | sbasalKC | MA0618.1 |
| 8.89502674 | 0.356702531310892 | 0.623 | 0.495 | 6.64E-22 | sbasalKC | MA0732.1 |
| 3.52250590 | 0.356313587206975 | 0.605 | 0.489 | 2.63E-17 | sbasalKC | MA0083.3 |
| 3.96844284 | 0.344405902892566 | 0.611 | 0.496 | 2.96E-16 | sbasalKC | MA1527.1 |
| 3.57569174 | 0.343048954706882 | 0.6 | 0.496 | 2.67E-15 | sbasalKC | MA0660.1 |
| 6.44536747 | 0.340079479097708 | 0.606 | 0.492 | 4.81E-16 | sbasalKC | MA0036.3 |
| 5.30471376 | 0.336343531734951 | 0.585 | 0.491 | 3.96E-14 | sbasalKC | MA0887.1 |
| 1.37772562 | 0.334999665866802 | 0.611 | 0.489 | 1.03E-15 | sbasalKC | MA0913.2 |
| 3.57968096 | 0.32660545712057 | 0.606 | 0.505 | 2.67E-15 | sbasalKC | MA1597.1 |
| 3.01678067 | 0.325032931933935 | 0.592 | 0.492 | 2.25E-13 | sbasalKC | MA0070.1 |
| 4.26562797 | 0.322633054924529 | 0.599 | 0.494 | 3.18E-13 | sbasalKC | MA1583.1 |
| 3.05804948 | 0.316829719153572 | 0.594 | 0.495 | 2.28E-14 | sbasalKC | MA0852.2 |
| 6.22523913 | 0.313082806307291 | 0.588 | 0.495 | 4.64E-13 | sbasalKC | MA0773.1 |
| 1.44782012 | 0.305048138691245 | 0.585 | 0.495 | 1.08E-11 | sbasalKC | MA1563.1 |
| 9.54076779 | 0.304110778835902 | 0.606 | 0.498 | 7.12E-15 | sbasalKC | MA1627.1 |
| 2.33122089 | 0.303691012710367 | 0.592 | 0.494 | 1.74E-11 | sbasalKC | MA1640.1 |
| 3.02095479 | 0.303151808824835 | 0.589 | 0.493 | 2.25E-12 | sbasalKC | MA0772.1 |

| | | | | | | |
|---|---|---|---|---|---|---|
| 1.80954692 | 0.299326940629861 | 0.591 | 0.496 | 1.35E-10 | sbasalKC | MA0482.2 |
| 3.00055911 | 0.298474699304084 | 0.579 | 0.497 | 2.24E-10 | sbasalKC | MA0504.1 |
| 1.15704838 | 0.291073801680696 | 0.566 | 0.488 | 8.63E-09 | sbasalKC | MA0888.1 |
| 2.12013557 | 0.286922450273706 | 0.574 | 0.484 | 1.58E-11 | sbasalKC | MA0079.4 |
| 9.91068744 | 0.284581113316285 | 0.578 | 0.492 | 7.39E-11 | sbasalKC | MA0497.1 |
| 1.41585260 | 0.283989008364304 | 0.597 | 0.503 | 1.06E-10 | sbasalKC | MA1655.1 |
| 2.91403136 | 0.283357729277073 | 0.603 | 0.503 | 2.17E-11 | sbasalKC | MA0782.2 |
| 1.52284581 | 0.277784689846351 | 0.571 | 0.487 | 1.14E-07 | sbasalKC | MA0742.1 |
| 1.61437846 | 0.276085694303579 | 0.581 | 0.493 | 1.20E-10 | sbasalKC | MA0652.1 |
| 7.66302294 | 0.275780805252743 | 0.578 | 0.488 | 5.72E-09 | sbasalKC | MA0657.1 |
| 1.27543705 | 0.275171814496642 | 0.59 | 0.494 | 9.51E-11 | sbasalKC | MA0030.1 |
| 1.97337980 | 0.27031570314352 | 0.598 | 0.496 | 1.47E-10 | sbasalKC | MA1537.1 |
| 2.41364395 | 0.269692417541925 | 0.584 | 0.498 | 1.80E-09 | sbasalKC | MA1155.1 |
| 3.96478075 | 0.26935351943422 | 0.59 | 0.502 | 2.96E-10 | sbasalKC | MA0052.4 |
| 4.83835153 | 0.267950780043081 | 0.559 | 0.486 | 3.61E-07 | sbasalKC | MA1594.1 |
| 1.43890337 | 0.266109846663471 | 0.583 | 0.495 | 1.07E-08 | sbasalKC | MA0909.2 |
| 3.84716515 | 0.259744143916467 | 0.567 | 0.494 | 2.87E-07 | sbasalKC | MA0060.3 |
| 9.00419695 | 0.257700961040305 | 0.565 | 0.492 | 6.72E-08 | sbasalKC | MA1496.1 |
| 5.56123133 | 0.257612084894968 | 0.585 | 0.504 | 4.15E-08 | sbasalKC | MA0619.1 |
| 3.97168412 | 0.255496182002631 | 0.58 | 0.493 | 2.96E-09 | sbasalKC | MA0483.1 |
| 4.40494065 | 0.25279712602136 | 0.583 | 0.503 | 3.29E-09 | sbasalKC | MA0472.2 |
| 5.12625538 | 0.252145289689213 | 0.58 | 0.506 | 3.82E-07 | sbasalKC | MA0142.1 |
| 3.35471636 | 0.251221308326831 | 0.574 | 0.493 | 2.50E-07 | sbasalKC | MA0596.1 |
| 2.26055969 | 0.25048906400962 | 0.57 | 0.496 | 1.69E-06 | sbasalKC | MA0467.1 |
| 4.45220675 | 0.25005361278145 | 0.594 | 0.489 | 3.32E-12 | sbasalKC | MA1513.1 |
| 1.34979643 | 0.654138508483461 | 0.711 | 0.484 | 1.01E-72 | muscFB | MA0740.1 |
| 1.23415826 | 0.558043863790697 | 0.675 | 0.482 | 9.21E-42 | muscFB | MA1573.1 |
| 9.03799875 | 0.552721633631506 | 0.676 | 0.493 | 6.74E-50 | muscFB | MA0032.2 |
| 1.30805381 | 0.521903047875199 | 0.673 | 0.496 | 9.76E-40 | muscFB | MA1099.2 |
| 2.87419451 | 0.518343996687758 | 0.658 | 0.483 | 2.14E-41 | muscFB | MA0057.1 |
| 3.71398743 | 0.512620828817648 | 0.672 | 0.481 | 2.77E-42 | muscFB | MA1512.1 |
| 1.76502069 | 0.51089747196921 | 0.651 | 0.483 | 1.32E-38 | muscFB | MA0747.1 |
| 7.36186821 | 0.509315544313346 | 0.659 | 0.486 | 5.49E-37 | muscFB | MA0136.2 |
| 3.78409175 | 0.506511877506313 | 0.658 | 0.491 | 2.82E-37 | muscFB | MA0596.1 |
| 1.82989033 | 0.505697539144908 | 0.654 | 0.485 | 1.37E-38 | muscFB | MA1607.1 |
| 5.00827501 | 0.498175877800492 | 0.682 | 0.478 | 3.74E-50 | muscFB | MA0039.4 |
| 3.28469848 | 0.480894740155295 | 0.64 | 0.484 | 2.45E-34 | muscFB | MA0163.1 |
| 5.34714823 | 0.480518481116917 | 0.649 | 0.488 | 3.99E-35 | muscFB | MA0645.1 |
| 6.95132347 | 0.472189105746116 | 0.64 | 0.482 | 5.19E-35 | muscFB | MA0640.2 |
| 6.19189495 | 0.466110943098348 | 0.643 | 0.491 | 4.62E-31 | muscFB | MA0679.2 |
| 1.48792609 | 0.465099223750127 | 0.632 | 0.485 | 1.11E-32 | muscFB | MA0746.2 |
| 2.15301544 | 0.463831947189778 | 0.634 | 0.482 | 1.61E-29 | muscFB | MA1476.1 |
| 2.26338227 | 0.463718850123093 | 0.683 | 0.481 | 1.69E-44 | muscFB | MA0599.1 |

| | | | | | | |
|---|---|---|---|---|---|---|
| 1.20549480 | 0.451665793070224 | 0.628 | 0.483 | 8.99E-27 | muscFB | MA0744.2 |
| 5.43624366 | 0.439173871639018 | 0.644 | 0.494 | 4.06E-31 | muscFB | MA0462.2 |
| 2.18145181 | 0.43871943056406 | 0.632 | 0.483 | 1.63E-29 | muscFB | MA1584.1 |
| 6.48197546 | 0.435948894835726 | 0.643 | 0.492 | 4.84E-27 | muscFB | MA0681.2 |
| 2.03834775 | 0.435389660447841 | 0.632 | 0.486 | 1.52E-29 | muscFB | MA1564.1 |
| 1.76410926 | 0.434723098372578 | 0.634 | 0.489 | 1.32E-26 | muscFB | MA0641.1 |
| 5.80675279 | 0.432102680097651 | 0.623 | 0.491 | 4.33E-25 | muscFB | MA0070.1 |
| 5.89193840 | 0.431935902702834 | 0.643 | 0.488 | 4.40E-30 | muscFB | MA0598.3 |
| 3.13845836 | 0.426309247080341 | 0.638 | 0.484 | 2.34E-24 | muscFB | MA0815.1 |
| 2.70604264 | 0.420784697933932 | 0.638 | 0.5 | 2.02E-25 | muscFB | MA0627.2 |
| 1.63717115 | 0.420420487352208 | 0.618 | 0.479 | 1.22E-20 | muscFB | MA1554.1 |
| 4.26588434 | 0.418888079249818 | 0.635 | 0.499 | 3.18E-25 | muscFB | MA1646.1 |
| 2.83713900 | 0.416380802323705 | 0.644 | 0.496 | 2.12E-27 | muscFB | MA1634.1 |
| 7.18804546 | 0.412040066440143 | 0.632 | 0.498 | 5.36E-25 | muscFB | MA1132.1 |
| 7.69864702 | 0.412004626536878 | 0.642 | 0.499 | 5.74E-26 | muscFB | MA0655.1 |
| 1.01206559 | 0.410438369929333 | 0.621 | 0.493 | 7.55E-22 | muscFB | MA0649.1 |
| 4.78842489 | 0.40749670055052 | 0.588 | 0.482 | 3.57E-18 | muscFB | MA0493.1 |
| 1.11446895 | 0.406237505250262 | 0.618 | 0.493 | 8.31E-22 | muscFB | MA0845.1 |
| 7.27164414 | 0.405385856449214 | 0.608 | 0.479 | 5.42E-21 | muscFB | MA0750.2 |
| 5.58025634 | 0.402383414234803 | 0.619 | 0.498 | 4.16E-22 | muscFB | MA0791.1 |
| 2.50160015 | 0.402122264278569 | 0.622 | 0.491 | 1.87E-22 | muscFB | MA0503.1 |
| 1.26326423 | 0.393572645669057 | 0.611 | 0.494 | 9.42E-19 | muscFB | MA0823.1 |
| 6.44209647 | 0.391201257722282 | 0.613 | 0.492 | 4.81E-20 | muscFB | MA0743.2 |
| 1.95161037 | 0.387522852778814 | 0.624 | 0.485 | 1.46E-25 | muscFB | MA0741.1 |
| 6.94256603 | 0.38717566630348 | 0.633 | 0.488 | 5.18E-29 | muscFB | MA1513.1 |
| 1.11901793 | 0.386295732565905 | 0.61 | 0.491 | 8.35E-21 | muscFB | MA0697.1 |
| 1.12287416 | 0.385170201191649 | 0.631 | 0.491 | 8.38E-27 | muscFB | MA1653.1 |
| 8.48019878 | 0.38494425685504 | 0.583 | 0.474 | 6.33E-17 | muscFB | MA0711.1 |
| 1.30182424 | 0.379992502772974 | 0.607 | 0.485 | 9.71E-19 | muscFB | MA0813.1 |
| 3.65312136 | 0.377965941821089 | 0.624 | 0.499 | 2.73E-18 | muscFB | MA0761.2 |
| 4.51218783 | 0.377745949360437 | 0.608 | 0.488 | 3.37E-19 | muscFB | MA0790.1 |
| 2.11886571 | 0.373825230651211 | 0.615 | 0.489 | 1.58E-19 | muscFB | MA0258.2 |
| 2.27871913 | 0.371021839481392 | 0.61 | 0.483 | 1.70E-19 | muscFB | MA1522.1 |
| 1.23173054 | 0.369915794245632 | 0.594 | 0.485 | 9.19E-15 | muscFB | MA1594.1 |
| 3.52542227 | 0.369630351254107 | 0.614 | 0.496 | 2.63E-18 | muscFB | MA1572.1 |
| 1.52660945 | 0.36932487409348 | 0.603 | 0.487 | 1.14E-15 | muscFB | MA0014.3 |
| 2.20468832 | 0.367027542331487 | 0.611 | 0.486 | 1.64E-17 | muscFB | MA0488.1 |
| 6.91537971 | 0.365551740058085 | 0.588 | 0.488 | 5.16E-14 | muscFB | MA0063.2 |
| 1.59593703 | 0.357007265049108 | 0.62 | 0.497 | 1.19E-19 | muscFB | MA0835.2 |
| 5.36525587 | 0.356575088990929 | 0.621 | 0.5 | 4.00E-21 | muscFB | MA1596.1 |
| 4.86660268 | 0.355958360575556 | 0.588 | 0.486 | 3.63E-16 | muscFB | MA0762.1 |
| 1.76748758 | 0.355396240784092 | 0.603 | 0.481 | 1.32E-17 | muscFB | MA0696.1 |
| 2.17531905 | 0.354077209907962 | 0.592 | 0.486 | 1.62E-15 | muscFB | MA0893.2 |

| | | | | | | |
|---|---|---|---|---|---|---|
| 2.17531905 | 0.354077209907962 | 0.592 | 0.486 | 1.62E-15 | muscFB | MA0912.2 |
| 1.95070551 | 0.353927211821889 | 0.585 | 0.487 | 1.46E-14 | muscFB | MA0076.2 |
| 4.13476935 | 0.353230656344579 | 0.602 | 0.487 | 3.08E-17 | muscFB | MA1508.1 |
| 2.20989807 | 0.353061256667703 | 0.631 | 0.501 | 1.65E-17 | muscFB | MA0786.1 |
| 8.15850381 | 0.352761068653621 | 0.602 | 0.488 | 6.09E-15 | muscFB | MA0657.1 |
| 5.27577274 | 0.351783210109407 | 0.601 | 0.487 | 3.94E-17 | muscFB | MA0685.1 |
| 6.53534582 | 0.351775767458875 | 0.599 | 0.484 | 4.88E-16 | muscFB | MA1483.1 |
| 6.58189572 | 0.350834347595193 | 0.609 | 0.494 | 4.91E-17 | muscFB | MA1101.2 |
| 6.72419658 | 0.34750575449482 | 0.606 | 0.493 | 5.02E-16 | muscFB | MA0764.2 |
| 5.65020763 | 0.344553229135921 | 0.571 | 0.487 | 4.22E-10 | muscFB | MA0742.1 |
| 1.93386534 | 0.341516087828656 | 0.609 | 0.5 | 1.44E-16 | muscFB | MA0476.1 |
| 1.75289826 | 0.336264275940813 | 0.603 | 0.498 | 1.31E-14 | muscFB | MA0792.1 |
| 1.59346534 | 0.335219973201886 | 0.59 | 0.497 | 1.19E-12 | muscFB | MA0861.1 |
| 1.71299783 | 0.335034987509422 | 0.599 | 0.493 | 1.28E-14 | muscFB | MA0047.3 |
| 4.92749084 | 0.331810760169014 | 0.593 | 0.486 | 3.68E-13 | muscFB | MA0018.4 |
| 4.42074229 | 0.331577919632607 | 0.568 | 0.476 | 3.30E-11 | muscFB | MA1545.1 |
| 1.02491985 | 0.330034361881414 | 0.593 | 0.486 | 7.65E-13 | muscFB | MA0872.1 |
| 2.61839509 | 0.329062293323618 | 0.588 | 0.481 | 1.95E-12 | muscFB | MA0056.2 |
| 7.93074090 | 0.327950295512145 | 0.584 | 0.495 | 5.92E-13 | muscFB | MA1575.1 |
| 1.82235250 | 0.326307544063145 | 0.584 | 0.483 | 1.36E-12 | muscFB | MA1608.1 |
| 1.63307660 | 0.325945273820978 | 0.59 | 0.49 | 1.22E-12 | muscFB | MA1124.1 |
| 7.56484556 | 0.324356505712963 | 0.621 | 0.5 | 5.64E-14 | muscFB | MA0871.2 |
| 9.15913742 | 0.321311448892984 | 0.61 | 0.499 | 6.83E-14 | muscFB | MA1115.1 |
| 5.22230790 | 0.318958046077133 | 0.591 | 0.503 | 3.90E-12 | muscFB | MA0831.2 |
| 2.79809425 | 0.318392956434998 | 0.58 | 0.483 | 2.09E-12 | muscFB | MA0656.1 |
| 5.81576517 | 0.316683718721703 | 0.6 | 0.493 | 4.34E-13 | muscFB | MA0789.1 |
| 6.14547065 | 0.313397222224164 | 0.598 | 0.493 | 4.58E-13 | muscFB | MA0155.1 |
| 8.98063140 | 0.309839476423393 | 0.575 | 0.488 | 6.70E-11 | muscFB | MA0626.1 |
| 1.50208783 | 0.309039750377358 | 0.603 | 0.501 | 1.12E-11 | muscFB | MA1134.1 |
| 1.59199078 | 0.306682746933617 | 0.58 | 0.486 | 1.19E-10 | muscFB | MA0609.2 |
| 2.82622045 | 0.306118724334407 | 0.582 | 0.497 | 2.11E-11 | muscFB | MA0595.1 |
| 7.00556299 | 0.302496875600665 | 0.566 | 0.485 | 5.23E-10 | muscFB | MA0691.1 |
| 1.61528341 | 0.298943251462364 | 0.582 | 0.49 | 1.21E-09 | muscFB | MA0525.2 |
| 1.23396542 | 0.298671932084475 | 0.584 | 0.488 | 9.21E-10 | muscFB | MA1632.1 |
| 3.97483235 | 0.297895438636123 | 0.588 | 0.492 | 2.97E-10 | muscFB | MA1683.1 |
| 1.83616761 | 0.297446084229273 | 0.583 | 0.486 | 1.37E-11 | muscFB | MA0751.1 |
| 2.07542543 | 0.297303239134075 | 0.571 | 0.494 | 1.55E-10 | muscFB | MA1103.2 |
| 1.03670723 | 0.295980483095885 | 0.603 | 0.503 | 7.73E-11 | muscFB | MA0100.3 |
| 2.00915956 | 0.295794425142273 | 0.568 | 0.492 | 1.50E-09 | muscFB | MA1657.1 |
| 8.39631807 | 0.295336762971226 | 0.591 | 0.487 | 6.26E-11 | muscFB | MA0492.1 |
| 1.49837388 | 0.295251962046676 | 0.577 | 0.483 | 1.12E-09 | muscFB | MA0509.2 |
| 8.30037179 | 0.295033813980408 | 0.605 | 0.496 | 6.19E-11 | muscFB | MA1499.1 |
| 8.30037179 | 0.295033813980408 | 0.605 | 0.496 | 6.19E-11 | muscFB | MA1507.1 |

| | | | | | | |
|---|---|---|---|---|---|---|
| 5.02717294 | 0.292595752421288 | 0.571 | 0.481 | 3.75E-09 | muscFB | MA1534.1 |
| 5.80036408 | 0.287572287030603 | 0.593 | 0.498 | 4.33E-12 | muscFB | MA0521.1 |
| 1.88227663 | 0.286102983444632 | 0.566 | 0.495 | 1.40E-09 | muscFB | MA0909.2 |
| 7.16991218 | 0.285458650815635 | 0.585 | 0.498 | 5.35E-10 | muscFB | MA0616.2 |
| 2.93058296 | 0.285372408656389 | 0.609 | 0.501 | 2.19E-11 | muscFB | MA0062.3 |
| 1.18303221 | 0.283837705616259 | 0.58 | 0.482 | 8.83E-09 | muscFB | MA1556.1 |
| 2.97541685 | 0.281955277423794 | 0.581 | 0.488 | 2.22E-09 | muscFB | MA0109.1 |
| 8.73260959 | 0.280853084321265 | 0.581 | 0.496 | 6.51E-10 | muscFB | MA0106.3 |
| 3.91706449 | 0.279014256639756 | 0.579 | 0.495 | 2.92E-09 | muscFB | MA1571.1 |
| 8.84613161 | 0.2732151420104 | 0.577 | 0.497 | 6.60E-09 | muscFB | MA0077.1 |
| 4.01917786 | 0.272668981111604 | 0.587 | 0.501 | 3.00E-11 | muscFB | MA0151.1 |
| 3.96827904 | 0.2720116530441 | 0.553 | 0.484 | 2.96E-08 | muscFB | MA0079.4 |
| 4.30812275 | 0.268848198503097 | 0.566 | 0.489 | 3.21E-07 | muscFB | MA0605.2 |
| 7.01853337 | 0.26829428643462 | 0.585 | 0.5 | 5.24E-09 | muscFB | MA1142.1 |
| 9.32311895 | 0.268025426550164 | 0.568 | 0.492 | 6.96E-08 | muscFB | MA0104.4 |
| 9.19059045 | 0.267829494509467 | 0.58 | 0.491 | 6.86E-09 | muscFB | MA0860.1 |
| 1.21569945 | 0.266870586880634 | 0.594 | 0.493 | 9.07E-10 | muscFB | MA1125.1 |
| 1.63519418 | 0.263161474686971 | 0.572 | 0.493 | 1.22E-07 | muscFB | MA0098.3 |
| 3.27350566 | 0.26209617157101 | 0.556 | 0.483 | 2.44E-06 | muscFB | MA0880.1 |
| 3.27350566 | 0.26209617157101 | 0.556 | 0.483 | 2.44E-06 | muscFB | MA0881.1 |
| 5.60885640 | 0.260752035189572 | 0.552 | 0.486 | 4.18E-06 | muscFB | MA0111.1 |
| 2.89590051 | 0.260581646670358 | 0.58 | 0.494 | 2.16E-08 | muscFB | MA0135.1 |
| 3.15631754 | 0.260230636790903 | 0.569 | 0.485 | 2.35E-07 | muscFB | MA0037.3 |
| 2.27763046 | 0.25987030048512 | 0.576 | 0.5 | 1.70E-07 | muscFB | MA0099.3 |
| 5.36087068 | 0.259489015476583 | 0.571 | 0.486 | 4.00E-07 | muscFB | MA0027.2 |
| 2.07024942 | 0.259303569215583 | 0.551 | 0.491 | 1.54E-06 | muscFB | MA1121.1 |
| 1.58700165 | 0.259199809499227 | 0.562 | 0.482 | 1.18E-08 | muscFB | MA1511.1 |
| 7.31722601 | 0.258338256175307 | 0.578 | 0.497 | 5.46E-08 | muscFB | MA1128.1 |
| 3.02125351 | 0.256821196740703 | 0.572 | 0.484 | 2.25E-07 | muscFB | MA0666.1 |
| 3.02125351 | 0.256821196740703 | 0.572 | 0.484 | 2.25E-07 | muscFB | MA0708.1 |
| 3.02125351 | 0.256821196740703 | 0.572 | 0.484 | 2.25E-07 | muscFB | MA0876.1 |
| 4.28180952 | 0.256783631104073 | 0.559 | 0.491 | 3.19E-07 | muscFB | MA0475.2 |
| 7.98037241 | 0.255462369127774 | 0.573 | 0.492 | 5.95E-08 | muscFB | MA0774.1 |
| 2.20967379 | 0.254703445127403 | 0.594 | 0.499 | 1.65E-08 | muscFB | MA0491.2 |
| 1.12981824 | 0.251989187354108 | 0.557 | 0.49 | 8.43E-06 | muscFB | MA0829.2 |
| 1.48820257 | 0.251721084916477 | 0.563 | 0.489 | 1.11E-05 | muscFB | MA0029.1 |
| 5.00482115 | 0.251022240232092 | 0.595 | 0.505 | 3.73E-08 | muscFB | MA0787.1 |
| 1.07131909 | 0.65597806434966 | 0.699 | 0.494 | 7.99E-45 | IRS | MA1639.1 |
| 3.48601360 | 0.511181300729205 | 0.662 | 0.492 | 2.60E-29 | IRS | MA1487.1 |
| 2.47507942 | 0.460123444956574 | 0.665 | 0.508 | 1.85E-23 | IRS | MA0051.1 |
| 3.34732750 | 0.449466201071616 | 0.63 | 0.495 | 2.50E-19 | IRS | MA0038.2 |
| 6.40471657 | 0.440087245113531 | 0.645 | 0.503 | 4.78E-20 | IRS | MA1624.1 |
| 2.96275548 | 0.426164608641939 | 0.654 | 0.506 | 2.21E-19 | IRS | MA0730.1 |

| | | | | | | |
|---|---|---|---|---|---|---|
| 2.57787933 | 0.419347829183208 | 0.646 | 0.505 | 1.92E-18 | IRS | MA0142.1 |
| 2.01175806 | 0.415648567310875 | 0.643 | 0.501 | 1.50E-20 | IRS | MA0601.1 |
| 4.74050690 | 0.411019608165611 | 0.613 | 0.498 | 3.54E-15 | IRS | MA1542.1 |
| 6.75943047 | 0.409342705025579 | 0.615 | 0.485 | 5.04E-15 | IRS | MA0903.1 |
| 2.93434081 | 0.408605456649265 | 0.641 | 0.498 | 2.19E-16 | IRS | MA0495.3 |
| 6.64883812 | 0.397443011998029 | 0.613 | 0.494 | 4.96E-15 | IRS | MA1640.1 |
| 1.47741494 | 0.394474796556651 | 0.607 | 0.495 | 1.10E-13 | IRS | MA0894.1 |
| 8.49654133 | 0.392978048960719 | 0.606 | 0.49 | 6.34E-14 | IRS | MA0706.1 |
| 2.85347634 | 0.386532712187263 | 0.611 | 0.5 | 2.13E-14 | IRS | MA0479.1 |
| 1.50437613 | 0.384299159587523 | 0.62 | 0.495 | 1.12E-13 | IRS | MA0259.1 |
| 4.26176885 | 0.379742279040109 | 0.628 | 0.506 | 3.18E-14 | IRS | MA1597.1 |
| 1.01973060 | 0.372470815138986 | 0.615 | 0.498 | 7.61E-11 | IRS | MA0875.1 |
| 5.40940120 | 0.368365934596961 | 0.612 | 0.498 | 4.04E-14 | IRS | MA0869.1 |
| 1.34249664 | 0.365495537671193 | 0.609 | 0.495 | 1.00E-13 | IRS | MA0909.2 |
| 3.56861817 | 0.359914013343288 | 0.623 | 0.502 | 2.66E-13 | IRS | MA0858.1 |
| 4.85368425 | 0.359438203184724 | 0.627 | 0.5 | 3.62E-13 | IRS | MA0108.2 |
| 7.10949830 | 0.35725851137224 | 0.605 | 0.501 | 5.30E-11 | IRS | MA0884.1 |
| 8.19095102 | 0.352419477426217 | 0.628 | 0.5 | 6.11E-12 | IRS | MA1106.1 |
| 2.84032210 | 0.339037080103791 | 0.568 | 0.486 | 2.12E-08 | IRS | MA0037.3 |
| 1.46823409 | 0.338680194174717 | 0.611 | 0.49 | 1.10E-10 | IRS | MA0913.2 |
| 5.69273551 | 0.338587971188419 | 0.616 | 0.494 | 4.25E-11 | IRS | MA1583.1 |
| 1.08763927 | 0.334029293519623 | 0.607 | 0.501 | 8.11E-11 | IRS | MA0527.1 |
| 1.15550796 | 0.326850220496511 | 0.572 | 0.489 | 8.62E-08 | IRS | MA0112.3 |
| 3.54122915 | 0.325597122263987 | 0.591 | 0.515 | 2.64E-08 | IRS | MA0526.3 |
| 1.90109853 | 0.3234940224718 | 0.598 | 0.494 | 1.42E-10 | IRS | MA0847.2 |
| 9.01627023 | 0.321292772572388 | 0.612 | 0.499 | 6.73E-12 | IRS | MA0087.1 |
| 1.28606380 | 0.315785547403841 | 0.601 | 0.492 | 9.59E-09 | IRS | MA1475.1 |
| 5.84391247 | 0.315152227104679 | 0.61 | 0.503 | 4.36E-10 | IRS | MA1655.1 |
| 4.18697426 | 0.314598562983302 | 0.583 | 0.499 | 3.12E-08 | IRS | MA0766.2 |
| 7.00760661 | 0.312852510357694 | 0.579 | 0.493 | 5.23E-07 | IRS | MA0902.2 |
| 4.36585988 | 0.308031450409097 | 0.58 | 0.493 | 3.26E-08 | IRS | MA1530.1 |
| 2.87613349 | 0.305385205061594 | 0.595 | 0.499 | 2.15E-10 | IRS | MA1627.1 |
| 6.77107190 | 0.301900566803932 | 0.617 | 0.498 | 5.05E-10 | IRS | MA0791.1 |
| 1.07945576 | 0.301515851113144 | 0.58 | 0.499 | 8.05E-08 | IRS | MA0650.2 |
| 1.29250684 | 0.300246894140774 | 0.609 | 0.504 | 9.64E-09 | IRS | MA0782.2 |
| 5.94064145 | 0.300202464055816 | 0.571 | 0.494 | 4.43E-07 | IRS | MA0658.1 |
| 1.42460828 | 0.29389378370032 | 0.576 | 0.491 | 1.06E-06 | IRS | MA1478.1 |
| 1.42024576 | 0.292720068760042 | 0.588 | 0.498 | 1.06E-08 | IRS | MA0851.1 |
| 6.19122468 | 0.286479071216225 | 0.576 | 0.489 | 4.62E-07 | IRS | MA1501.1 |
| 1.32612152 | 0.28499133181976 | 0.573 | 0.493 | 9.89E-06 | IRS | MA0890.1 |
| 4.41362438 | 0.278645130998115 | 0.613 | 0.501 | 3.29E-08 | IRS | MA0614.1 |
| 1.34595625 | 0.277962322065344 | 0.58 | 0.493 | 1.00E-06 | IRS | MA0070.1 |
| 5.02826670 | 0.272466425562535 | 0.595 | 0.501 | 3.75E-09 | IRS | MA0151.1 |

| | | | | | | |
|---|---|---|---|---|---|---|
| 9.57425833 | 0.269512249463474 | 0.578 | 0.497 | 7.14E-06 | IRS | MA0031.1 |
| 9.57425833 | 0.269512249463474 | 0.578 | 0.497 | 7.14E-06 | IRS | MA0157.2 |
| 7.24198397 | 0.26754608493989 | 0.579 | 0.492 | 5.40E-06 | IRS | MA0618.1 |
| 2.78216755 | 0.267194788371608 | 0.574 | 0.488 | 2.08E-05 | IRS | MA0888.1 |
| 2.95883645 | 0.26676614467562 | 0.578 | 0.489 | 2.21E-06 | IRS | MA1576.1 |
| 2.83985019 | 0.266646321169711 | 0.562 | 0.503 | 2.12E-05 | IRS | MA0786.1 |
| 2.79578704 | 0.265147019288354 | 0.562 | 0.502 | 0.000208566 | IRS | MA1466.1 |
| 6.69798135 | 0.260344105165137 | 0.585 | 0.51 | 5.00E-05 | IRS | MA0692.1 |
| 2.97489299 | 0.258846328434114 | 0.576 | 0.493 | 2.22E-05 | IRS | MA1113.2 |
| 9.50498330 | 0.256439553515763 | 0.568 | 0.498 | 7.09E-05 | IRS | MA0732.1 |
| 6.82868460 | 0.254278380070828 | 0.576 | 0.494 | 5.09E-05 | IRS | MA1589.1 |
| 2.30040567 | 0.252787862447496 | 0.58 | 0.494 | 1.72E-05 | IRS | MA0036.3 |
| 3.08481721 | 0.252649236900978 | 0.596 | 0.503 | 2.30E-06 | IRS | MA0052.4 |
| 2.48169307 | 0.251327113886276 | 0.587 | 0.501 | 1.85E-06 | IRS | MA0613.1 |
| 3.97419723 | 0.883891088326707 | 0.789 | 0.482 | 2.96E-62 | NK | MA0897.1 |
| 1.66112461 | 0.816743827274948 | 0.785 | 0.493 | 1.24E-59 | NK | MA0501.1 |
| 4.82695679 | 0.815819926436163 | 0.789 | 0.5 | 3.60E-59 | NK | MA0858.1 |
| 1.13237109 | 0.794353532003096 | 0.796 | 0.497 | 8.45E-55 | NK | MA1149.1 |
| 8.42361273 | 0.793507958236528 | 0.771 | 0.5 | 6.28E-54 | NK | MA1616.1 |
| 1.24050964 | 0.790810550479214 | 0.726 | 0.489 | 9.25E-45 | NK | MA1521.1 |
| 1.90470158 | 0.788335140911993 | 0.781 | 0.488 | 1.42E-52 | NK | MA0907.1 |
| 1.16717474 | 0.776192615297807 | 0.749 | 0.485 | 8.71E-46 | NK | MA0074.1 |
| 2.09143039 | 0.744873338693624 | 0.75 | 0.492 | 1.56E-45 | NK | MA1541.1 |
| 3.15794602 | 0.744788856737305 | 0.745 | 0.494 | 2.36E-48 | NK | MA0150.2 |
| 1.46867211 | 0.742646030510218 | 0.737 | 0.494 | 1.10E-43 | NK | MA0804.1 |
| 7.13328716 | 0.728665097395368 | 0.726 | 0.483 | 5.32E-40 | NK | MA0898.1 |
| 1.85595265 | 0.719621961603452 | 0.724 | 0.494 | 1.38E-34 | NK | MA1520.1 |
| 9.83079904 | 0.704243336540909 | 0.752 | 0.495 | 7.33E-43 | NK | MA0734.2 |
| 2.59817527 | 0.698832074996964 | 0.735 | 0.492 | 1.94E-39 | NK | MA0689.1 |
| 2.04748189 | 0.693962200496974 | 0.726 | 0.501 | 1.53E-36 | NK | MA1466.1 |
| 2.73105560 | 0.688572996468857 | 0.749 | 0.496 | 2.04E-45 | NK | MA0732.1 |
| 4.03352303 | 0.687223112504256 | 0.768 | 0.501 | 3.01E-44 | NK | MA0152.1 |
| 8.62482253 | 0.684550952609693 | 0.716 | 0.487 | 6.43E-36 | NK | MA0895.1 |
| 1.16285018 | 0.680382352747977 | 0.72 | 0.498 | 8.67E-38 | NK | MA0480.1 |
| 3.79258859 | 0.666121227207513 | 0.693 | 0.489 | 2.83E-32 | NK | MA0767.1 |
| 7.76847669 | 0.663275904656492 | 0.716 | 0.489 | 5.80E-36 | NK | MA0089.2 |
| 4.01402222 | 0.661832776178417 | 0.741 | 0.506 | 2.99E-36 | NK | MA0730.1 |
| 5.03230455 | 0.655878083691758 | 0.752 | 0.501 | 3.75E-41 | NK | MA0490.2 |
| 1.14849660 | 0.646735264284885 | 0.731 | 0.503 | 8.57E-35 | NK | MA1528.1 |
| 7.32541673 | 0.633049279660313 | 0.716 | 0.499 | 5.46E-31 | NK | MA1494.1 |
| 5.66782498 | 0.632923091509536 | 0.703 | 0.489 | 4.23E-34 | NK | MA0496.3 |
| 5.19733808 | 0.629291538250097 | 0.714 | 0.491 | 3.88E-34 | NK | MA0146.2 |
| 4.11663969 | 0.627620420019913 | 0.73 | 0.501 | 3.07E-31 | NK | MA0137.3 |

| | | | | | | |
|---|---|---|---|---|---|---|
| 3.12180565 | 0.624556758499587 | 0.71 | 0.509 | 2.33E-30 | NK | MA0051.1 |
| 6.30361409 | 0.622676347795273 | 0.718 | 0.497 | 4.70E-31 | NK | MA1491.1 |
| 5.83440331 | 0.614853096401736 | 0.724 | 0.493 | 4.35E-30 | NK | MA0719.1 |
| 2.93428314 | 0.613238006005671 | 0.693 | 0.491 | 2.19E-25 | NK | MA0628.1 |
| 2.05785982 | 0.605722538789134 | 0.705 | 0.491 | 1.54E-29 | NK | MA0591.1 |
| 3.90798681 | 0.604678245644248 | 0.691 | 0.485 | 2.92E-30 | NK | MA0814.2 |
| 9.67285414 | 0.594239636858019 | 0.71 | 0.495 | 7.22E-29 | NK | MA1565.1 |
| 1.69706609 | 0.592995327072595 | 0.672 | 0.494 | 1.27E-20 | NK | MA0680.1 |
| 1.41724724 | 0.588398029669099 | 0.693 | 0.502 | 1.06E-25 | NK | MA0636.1 |
| 8.34430350 | 0.588162049462004 | 0.703 | 0.487 | 6.22E-26 | NK | MA0714.1 |
| 1.71839207 | 0.582094121767577 | 0.682 | 0.491 | 1.28E-25 | NK | MA0702.2 |
| 8.05689586 | 0.580141807708373 | 0.69 | 0.493 | 6.01E-27 | NK | MA0092.1 |
| 8.99066750 | 0.577375472036267 | 0.693 | 0.502 | 6.71E-25 | NK | MA0831.2 |
| 1.29859786 | 0.575049986517886 | 0.684 | 0.494 | 9.69E-24 | NK | MA0638.1 |
| 1.80508333 | 0.562024272653508 | 0.695 | 0.499 | 1.35E-22 | NK | MA0756.1 |
| 7.14574437 | 0.558742651272376 | 0.684 | 0.493 | 5.33E-24 | NK | MA1487.1 |
| 4.74016083 | 0.556238474858174 | 0.697 | 0.504 | 3.54E-24 | NK | MA1485.1 |
| 8.06219618 | 0.554055354823385 | 0.684 | 0.484 | 6.01E-23 | NK | MA0812.1 |
| 5.49998154 | 0.552069956115949 | 0.684 | 0.496 | 4.10E-21 | NK | MA0828.1 |
| 2.11035523 | 0.551408291453809 | 0.712 | 0.497 | 1.57E-27 | NK | MA1527.1 |
| 1.50241376 | 0.54620689370647 | 0.67 | 0.488 | 1.12E-21 | NK | MA0672.1 |
| 1.04541550 | 0.545544598533752 | 0.638 | 0.485 | 7.80E-19 | NK | MA0896.1 |
| 8.44419534 | 0.542847073883732 | 0.676 | 0.489 | 6.30E-22 | NK | MA0790.1 |
| 5.34241216 | 0.541638029331884 | 0.686 | 0.509 | 3.99E-21 | NK | MA0692.1 |
| 1.11972036 | 0.540600724373203 | 0.686 | 0.499 | 8.35E-26 | NK | MA0491.2 |
| 5.32612435 | 0.540250597740793 | 0.65 | 0.491 | 3.97E-19 | NK | MA1474.1 |
| 3.76933665 | 0.538946662763828 | 0.697 | 0.506 | 2.81E-23 | NK | MA0142.1 |
| 8.04541657 | 0.535433107228871 | 0.682 | 0.494 | 6.00E-21 | NK | MA1108.2 |
| 6.04250262 | 0.531943794676558 | 0.667 | 0.495 | 4.51E-18 | NK | MA0853.1 |
| 1.91051941 | 0.530962415747109 | 0.674 | 0.495 | 1.43E-19 | NK | MA1644.1 |
| 9.85473649 | 0.526341095097709 | 0.651 | 0.488 | 7.35E-19 | NK | MA1519.1 |
| 1.17456070 | 0.525478699258082 | 0.663 | 0.493 | 8.76E-18 | NK | MA0674.1 |
| 1.46118080 | 0.522206468160089 | 0.67 | 0.486 | 1.09E-18 | NK | MA0755.1 |
| 4.93693749 | 0.521047529298775 | 0.69 | 0.503 | 3.68E-21 | NK | MA0088.2 |
| 1.36775651 | 0.520982544419168 | 0.672 | 0.491 | 1.02E-18 | NK | MA0704.1 |
| 1.17809078 | 0.515650592875473 | 0.638 | 0.49 | 8.79E-17 | NK | MA0874.1 |
| 4.99982085 | 0.515438920114733 | 0.674 | 0.498 | 3.73E-19 | NK | MA0863.1 |
| 9.72061404 | 0.512943753328437 | 0.672 | 0.484 | 7.25E-21 | NK | MA1633.1 |
| 1.19289486 | 0.500128984845199 | 0.646 | 0.494 | 8.90E-15 | NK | MA0839.1 |
| 1.54823433 | 0.499752541264068 | 0.678 | 0.499 | 1.15E-17 | NK | MA0836.2 |
| 2.64034885 | 0.497450824398586 | 0.67 | 0.496 | 1.97E-17 | NK | MA1639.1 |
| 2.22969732 | 0.497441873126775 | 0.684 | 0.499 | 1.66E-19 | NK | MA0119.1 |
| 1.91535371 | 0.497057921654004 | 0.678 | 0.5 | 1.43E-16 | NK | MA1106.1 |

| | | | | | | |
|---|---|---|---|---|---|---|
| 1.42698645 | 0.487379453103529 | 0.663 | 0.495 | 1.06E-15 | NK | MA0259.1 |
| 3.54831330 | 0.486657010264894 | 0.68 | 0.489 | 2.65E-15 | NK | MA0639.1 |
| 1.43261347 | 0.484754155705732 | 0.657 | 0.487 | 1.07E-15 | NK | MA0682.2 |
| 1.93572778 | 0.483924062973123 | 0.665 | 0.49 | 1.44E-16 | NK | MA0806.1 |
| 1.15834859 | 0.483456490069934 | 0.648 | 0.485 | 8.64E-16 | NK | MA1547.1 |
| 2.69960798 | 0.483149434635384 | 0.672 | 0.498 | 2.01E-16 | NK | MA0653.1 |
| 1.87530856 | 0.482345528249035 | 0.659 | 0.498 | 1.40E-16 | NK | MA0159.1 |
| 1.84667055 | 0.480346385465097 | 0.644 | 0.483 | 1.38E-15 | NK | MA0739.1 |
| 1.90393146 | 0.479732180312201 | 0.676 | 0.497 | 1.42E-17 | NK | MA0003.4 |
| 6.49996339 | 0.478995565445373 | 0.659 | 0.49 | 4.85E-16 | NK | MA1481.1 |
| 3.23301508 | 0.478259218747735 | 0.678 | 0.497 | 2.41E-17 | NK | MA0729.1 |
| 2.44922379 | 0.472034303403408 | 0.667 | 0.503 | 1.83E-16 | NK | MA1655.1 |
| 1.01536328 | 0.471184587082907 | 0.636 | 0.498 | 7.57E-13 | NK | MA0780.1 |
| 7.91762242 | 0.469004609411503 | 0.674 | 0.494 | 5.91E-16 | NK | MA0652.1 |
| 2.42326977 | 0.468666887231844 | 0.669 | 0.498 | 1.81E-17 | NK | MA0733.1 |
| 3.39197630 | 0.467081651807927 | 0.661 | 0.499 | 2.53E-15 | NK | MA1580.1 |
| 3.14691662 | 0.465219182005213 | 0.644 | 0.495 | 2.35E-14 | NK | MA0160.1 |
| 5.26297530 | 0.464051979530525 | 0.63 | 0.489 | 3.93E-12 | NK | MA1555.1 |
| 2.44391703 | 0.461484126243845 | 0.648 | 0.498 | 1.82E-14 | NK | MA0854.1 |
| 2.01726065 | 0.454970445978674 | 0.636 | 0.489 | 1.50E-13 | NK | MA1576.1 |
| 5.48188760 | 0.451406125209762 | 0.661 | 0.491 | 4.09E-15 | NK | MA0006.1 |
| 1.83494254 | 0.449220804411008 | 0.642 | 0.489 | 1.37E-13 | NK | MA0513.1 |
| 1.32890545 | 0.44607682484334 | 0.623 | 0.495 | 9.91E-11 | NK | MA0135.1 |
| 3.23444206 | 0.446068422626229 | 0.667 | 0.492 | 2.41E-14 | NK | MA0774.1 |
| 9.09578091 | 0.445749680069475 | 0.665 | 0.504 | 6.79E-15 | NK | MA0619.1 |
| 5.58378450 | 0.442303607886742 | 0.632 | 0.494 | 4.17E-12 | NK | MA0743.2 |
| 5.05507209 | 0.440325889949678 | 0.625 | 0.49 | 3.77E-12 | NK | MA0716.1 |
| 5.05507209 | 0.440325889949678 | 0.625 | 0.49 | 3.77E-12 | NK | MA0721.1 |
| 1.62344585 | 0.433667805104632 | 0.61 | 0.488 | 1.21E-10 | NK | MA0891.1 |
| 1.49830620 | 0.433151837451676 | 0.634 | 0.498 | 1.12E-12 | NK | MA1572.1 |
| 1.35035797 | 0.432304327780002 | 0.629 | 0.489 | 1.01E-12 | NK | MA0892.1 |
| 1.48477674 | 0.429691707866083 | 0.638 | 0.486 | 1.11E-11 | NK | MA0705.1 |
| 1.48026140 | 0.428906823043323 | 0.63 | 0.496 | 1.10E-12 | NK | MA1643.1 |
| 5.81029515 | 0.424854096864906 | 0.638 | 0.496 | 4.33E-13 | NK | MA1571.1 |
| 4.63818260 | 0.418205280995249 | 0.619 | 0.488 | 3.46E-11 | NK | MA1470.1 |
| 1.08668033 | 0.413598193350596 | 0.608 | 0.492 | 8.11E-09 | NK | MA0752.1 |
| 3.23696569 | 0.410173777086347 | 0.615 | 0.499 | 2.41E-10 | NK | MA1155.1 |
| 7.24053938 | 0.408286397261047 | 0.634 | 0.505 | 5.40E-10 | NK | MA0782.2 |
| 2.63869158 | 0.405544264496645 | 0.613 | 0.499 | 1.97E-10 | NK | MA1654.1 |
| 2.84438034 | 0.403673821626125 | 0.629 | 0.493 | 2.12E-10 | NK | MA0805.1 |
| 1.62785673 | 0.403622876802813 | 0.627 | 0.492 | 1.21E-11 | NK | MA0860.1 |
| 1.04647751 | 0.402895043703376 | 0.623 | 0.484 | 7.81E-10 | NK | MA0777.1 |
| 7.36363660 | 0.402859425088534 | 0.629 | 0.485 | 5.49E-10 | NK | MA1532.1 |

| | | | | | | |
|---|---|---|---|---|---|---|
| 2.11098297 | 0.39841831809433 | 0.606 | 0.478 | 1.57E-09 | NK | MA0776.1 |
| 1.41363110 | 0.395389531152379 | 0.623 | 0.49 | 1.05E-09 | NK | MA0810.1 |
| 5.14427916 | 0.393160069353423 | 0.636 | 0.499 | 3.84E-10 | NK | MA0633.1 |
| 1.53139598 | 0.391759742665703 | 0.613 | 0.495 | 1.14E-08 | NK | MA1500.1 |
| 3.04550816 | 0.390582377960718 | 0.625 | 0.494 | 2.27E-09 | NK | MA0671.1 |
| 5.99453178 | 0.388505320490637 | 0.634 | 0.495 | 4.47E-10 | NK | MA0712.2 |
| 1.77954850 | 0.386476176137738 | 0.611 | 0.492 | 1.33E-08 | NK | MA0822.1 |
| 2.52406784 | 0.38625336875261 | 0.602 | 0.482 | 1.88E-09 | NK | MA1534.1 |
| 3.28194387 | 0.385785533430101 | 0.636 | 0.49 | 2.45E-10 | NK | MA0688.1 |
| 4.68401497 | 0.384825226557533 | 0.623 | 0.49 | 3.49E-09 | NK | MA0807.1 |
| 8.16882982 | 0.383874464134219 | 0.636 | 0.497 | 6.09E-12 | NK | MA0462.2 |
| 9.71594183 | 0.383855312610494 | 0.621 | 0.489 | 7.25E-10 | NK | MA1535.1 |
| 2.85875011 | 0.383619033254807 | 0.632 | 0.495 | 2.13E-09 | NK | MA0859.1 |
| 4.62881733 | 0.37983166883722 | 0.627 | 0.5 | 3.45E-11 | NK | MA1142.1 |
| 5.63648913 | 0.379222591669696 | 0.629 | 0.502 | 4.20E-10 | NK | MA0476.1 |
| 2.22467118 | 0.377912270868317 | 0.65 | 0.494 | 1.66E-09 | NK | MA1143.1 |
| 2.91862416 | 0.376225674047438 | 0.615 | 0.499 | 2.18E-09 | NK | MA0593.1 |
| 9.46262993 | 0.372558618273806 | 0.632 | 0.497 | 7.06E-11 | NK | MA0489.1 |
| 6.11269038 | 0.3722934729636 | 0.621 | 0.487 | 4.56E-08 | NK | MA0802.1 |
| 1.83412666 | 0.371796602 | 0.619 | 0.496 | 1.37E-09 | NK | MA0038.2 |
| 3.64370889 | 0.371335399443528 | 0.589 | 0.49 | 2.72E-08 | NK | MA0644.1 |
| 3.84043465 | 0.370947543393939 | 0.627 | 0.492 | 2.86E-08 | NK | MA0735.1 |
| 2.47293304 | 0.364820792696767 | 0.568 | 0.481 | 1.84E-06 | NK | MA1495.1 |
| 2.47293304 | 0.364820792696767 | 0.568 | 0.481 | 1.84E-06 | NK | MA0904.2 |
| 2.44137998 | 0.363071387829735 | 0.606 | 0.496 | 1.82E-07 | NK | MA1504.1 |
| 1.55180284 | 0.362324010334695 | 0.604 | 0.496 | 1.16E-07 | NK | MA0508.3 |
| 5.06093330 | 0.361978395461327 | 0.61 | 0.496 | 3.78E-07 | NK | MA0468.1 |
| 3.92494510 | 0.360846067065488 | 0.623 | 0.489 | 2.93E-08 | NK | MA1501.1 |
| 1.17739856 | 0.359963280346819 | 0.619 | 0.5 | 8.78E-08 | NK | MA0603.1 |
| 8.58341776 | 0.358952490649406 | 0.638 | 0.499 | 6.40E-08 | NK | MA0524.2 |
| 6.88024570 | 0.358882139013242 | 0.592 | 0.491 | 5.13E-07 | NK | MA0844.1 |
| 2.02290910 | 0.356732914514109 | 0.61 | 0.494 | 1.51E-07 | NK | MA0648.1 |
| 4.50630117 | 0.355761096622505 | 0.623 | 0.5 | 3.36E-07 | NK | MA1111.1 |
| 1.19135239 | 0.355200620071268 | 0.604 | 0.495 | 8.89E-08 | NK | MA0060.3 |
| 1.81880051 | 0.353912702630159 | 0.627 | 0.501 | 1.36E-08 | NK | MA0655.1 |
| 4.99108619 | 0.350969403597582 | 0.608 | 0.502 | 3.72E-07 | NK | MA0884.1 |
| 1.54514170 | 0.350821384832874 | 0.617 | 0.495 | 1.15E-07 | NK | MA0659.2 |
| 3.21470870 | 0.349792445980908 | 0.589 | 0.486 | 2.40E-06 | NK | MA0885.1 |
| 2.56582624 | 0.347773519980678 | 0.632 | 0.498 | 1.91E-09 | NK | MA1634.1 |
| 6.21922858 | 0.347732735232832 | 0.619 | 0.499 | 4.64E-09 | NK | MA1622.1 |
| 1.22519455 | 0.345508500343521 | 0.632 | 0.515 | 9.14E-07 | NK | MA0526.3 |
| 9.53889157 | 0.343916515005349 | 0.634 | 0.5 | 7.12E-09 | NK | MA1630.1 |
| 5.41326174 | 0.342867138804761 | 0.636 | 0.499 | 4.04E-09 | NK | MA0835.2 |

| | | | | | | |
|---|---|---|---|---|---|---|
| 6.72465837 | 0.340025839215912 | 0.611 | 0.491 | 5.02E-07 | NK | MA0675.1 |
| 1.00376651 | 0.339844577331559 | 0.61 | 0.503 | 7.49E-06 | NK | MA0620.3 |
| 5.55104735 | 0.338837551111581 | 0.617 | 0.502 | 4.14E-07 | NK | MA0797.1 |
| 8.50752682 | 0.335360364351019 | 0.634 | 0.489 | 6.35E-08 | NK | MA0132.2 |
| 1.83008815 | 0.334954815161584 | 0.6 | 0.49 | 1.37E-06 | NK | MA0667.1 |
| 1.68349760 | 0.332283803476958 | 0.61 | 0.495 | 1.26E-06 | NK | MA1103.2 |
| 4.91427032 | 0.331617137772064 | 0.632 | 0.494 | 3.67E-08 | NK | MA1623.1 |
| 2.04490092 | 0.330861058357354 | 0.61 | 0.491 | 1.53E-06 | NK | MA1600.1 |
| 2.49370335 | 0.330347697158466 | 0.61 | 0.494 | 1.86E-06 | NK | MA1114.1 |
| 1.51190844 | 0.330030485027214 | 0.583 | 0.481 | 1.13E-05 | NK | MA1560.1 |
| 2.94343997 | 0.328964694998643 | 0.611 | 0.488 | 2.20E-06 | NK | MA1567.1 |
| 4.06126381 | 0.328287371298345 | 0.585 | 0.49 | 3.03E-05 | NK | MA0775.1 |
| 3.51708628 | 0.327810797744092 | 0.589 | 0.491 | 2.62E-06 | NK | MA1531.1 |
| 1.97361834 | 0.327261534075241 | 0.613 | 0.49 | 1.47E-05 | NK | MA0737.1 |
| 8.01440098 | 0.325214797316825 | 0.594 | 0.498 | 5.98E-06 | NK | MA0504.1 |
| 1.51680814 | 0.320406494418279 | 0.562 | 0.493 | 0.000113154 | NK | MA1496.1 |
| 8.65941938 | 0.317626379652384 | 0.592 | 0.49 | 6.46E-06 | NK | MA0800.1 |
| 1.34364097 | 0.317520693572629 | 0.602 | 0.486 | 1.00E-05 | NK | MA1585.1 |
| 1.41629572 | 0.317285012684698 | 0.598 | 0.493 | 1.06E-06 | NK | MA0130.1 |
| 4.50690775 | 0.315244805881934 | 0.556 | 0.49 | 0.000336215 | NK | MA0713.1 |
| 2.46759839 | 0.313570593513881 | 0.606 | 0.496 | 1.84E-06 | NK | MA1637.1 |
| 9.22131634 | 0.312985669374068 | 0.594 | 0.491 | 6.88E-06 | NK | MA0901.2 |
| 2.97714348 | 0.309169938201742 | 0.611 | 0.501 | 2.22E-05 | NK | MA0058.3 |
| 6.54265135 | 0.309005159566676 | 0.598 | 0.49 | 4.88E-05 | NK | MA1497.1 |
| 1.55430708 | 0.303115281700988 | 0.598 | 0.49 | 0.000115951 | NK | MA0843.1 |
| 4.09541630 | 0.30257299300503 | 0.596 | 0.492 | 3.06E-05 | NK | MA0913.2 |
| 2.27120771 | 0.30181906562098 | 0.573 | 0.502 | 0.000169432 | NK | MA0597.1 |
| 2.28471054 | 0.299671353722014 | 0.59 | 0.5 | 1.70E-05 | NK | MA0650.2 |
| 1.22425937 | 0.299082751926877 | 0.573 | 0.491 | 9.13E-05 | NK | MA0829.2 |
| 1.00774494 | 0.295310284538587 | 0.581 | 0.492 | 7.52E-05 | NK | MA1109.1 |
| 2.30588756 | 0.294906359098421 | 0.587 | 0.495 | 0.000172019 | NK | MA0658.1 |
| 1.20859137 | 0.293849514519678 | 0.6 | 0.496 | 9.02E-05 | NK | MA1575.1 |
| 3.77499191 | 0.292637086334037 | 0.583 | 0.493 | 0.000281614 | NK | MA0642.1 |
| 1.12108095 | 0.292026875116344 | 0.636 | 0.502 | 8.36E-06 | NK | MA0017.2 |
| 5.91703435 | 0.291296627694665 | 0.592 | 0.498 | 4.41E-05 | NK | MA1499.1 |
| 5.91703435 | 0.291296627694665 | 0.592 | 0.498 | 4.41E-05 | NK | MA1507.1 |
| 2.54712919 | 0.29129568815136 | 0.598 | 0.494 | 0.000190016 | NK | MA1480.1 |
| 1.11224652 | 0.291287180812496 | 0.549 | 0.485 | 0.000829736 | NK | MA1476.1 |
| 5.27671956 | 0.290119923087813 | 0.611 | 0.501 | 3.94E-06 | NK | MA1141.1 |
| 3.08681222 | 0.2888828763262 | 0.604 | 0.506 | 0.000230276 | NK | MA0783.1 |
| 4.14784280 | 0.287813278543117 | 0.6 | 0.493 | 0.000309429 | NK | MA0817.1 |
| 3.44150754 | 0.287413079982848 | 0.577 | 0.484 | 0.000256736 | NK | MA1566.1 |
| 4.80412448 | 0.285332210735713 | 0.57 | 0.499 | 0.003583877 | NK | MA1419.1 |

| | | | | | | |
|---|---|---|---|---|---|---|
| 8.55863116 | 0.285191205916891 | 0.568 | 0.493 | 0.000638474 | NK | MA0803.1 |
| 1.63736070 | 0.284534642760053 | 0.59 | 0.496 | 0.000122147 | NK | MA1101.2 |
| 2.32801111 | 0.280134335184891 | 0.604 | 0.497 | 0.00017367 | NK | MA0106.3 |
| 5.64186634 | 0.27780472959644 | 0.602 | 0.495 | 0.000420883 | NK | MA0811.1 |
| 7.30135333 | 0.276940270510995 | 0.577 | 0.484 | 0.000544681 | NK | MA0799.1 |
| 1.12653045 | 0.276612063741646 | 0.56 | 0.491 | 0.008403917 | NK | MA0686.1 |
| 1.04074612 | 0.276424999925905 | 0.592 | 0.486 | 0.000776397 | NK | MA0622.1 |
| 2.65667542 | 0.275933688685712 | 0.592 | 0.5 | 0.000198188 | NK | MA0791.1 |
| 6.99279711 | 0.275200986502748 | 0.575 | 0.494 | 0.000521663 | NK | MA0036.3 |
| 2.37784106 | 0.274817841371165 | 0.566 | 0.488 | 0.001773869 | NK | MA1642.1 |
| 4.64084896 | 0.273951312456638 | 0.573 | 0.491 | 0.003462073 | NK | MA0059.1 |
| 7.58346676 | 0.273255300643002 | 0.6 | 0.501 | 5.66E-05 | NK | MA0149.1 |
| 4.64927281 | 0.271662443028757 | 0.613 | 0.503 | 3.47E-05 | NK | MA0477.2 |
| 5.00468250 | 0.271185099639194 | 0.579 | 0.495 | 0.000373349 | NK | MA1569.1 |
| 1.32731593 | 0.270057208671356 | 0.55 | 0.485 | 0.009901777 | NK | MA1608.1 |
| 1.39391415 | 0.269713728752468 | 0.589 | 0.493 | 0.00103986 | NK | MA0035.4 |
| 1.58394032 | 0.268881023819536 | 0.632 | 0.503 | 0.000118162 | NK | MA1489.1 |
| 7.00201622 | 0.268419024602229 | 0.566 | 0.5 | 0.005223504 | NK | MA0495.3 |
| 1.02973188 | 0.267992207515344 | 0.615 | 0.499 | 7.68E-05 | NK | MA1138.1 |
| 1.01781824 | 0.267370515597343 | 0.602 | 0.498 | 7.59E-05 | NK | MA0478.1 |
| 8.56910086 | 0.265842087564995 | 0.57 | 0.487 | 0.000639255 | NK | MA1479.1 |
| 9.20113733 | 0.265234209129718 | 0.575 | 0.488 | 0.000686405 | NK | MA0522.3 |
| 1.03822277 | 0.265149230555863 | 0.596 | 0.493 | 0.000774514 | NK | MA0070.1 |
| 2.02419506 | 0.265130225721464 | 0.585 | 0.502 | 0.00151005 | NK | MA1464.1 |
| 1.62465660 | 0.2639497630837 | 0.566 | 0.486 | 0.012119938 | NK | MA0900.2 |
| 1.84430634 | 0.26091299381425 | 0.583 | 0.486 | 0.001375853 | NK | MA1505.1 |
| 3.88469959 | 0.259846885943358 | 0.541 | 0.496 | 0.028979859 | NK | MA0069.1 |
| 5.23443289 | 0.256285918316009 | 0.611 | 0.502 | 0.000390489 | NK | MA1596.1 |
| 5.93702447 | 0.255725144276637 | 0.611 | 0.495 | 0.000442902 | NK | MA0144.2 |
| 3.36337804 | 0.255556085567463 | 0.566 | 0.493 | 0.00250908 | NK | MA0161.2 |
| 1.22283779 | 0.253816917239183 | 0.554 | 0.493 | 0.00912237 | NK | MA0101.1 |
| 1.36396946 | 0.251170068628367 | 0.606 | 0.505 | 0.001017521 | NK | MA1624.1 |
| 1.85498682 | 0.250502465263231 | 0.577 | 0.495 | 0.013838202 | NK | MA1420.1 |
| 6.90177322 | 0.685822643392188 | 0.697 | 0.486 | 5.15E-38 | DP | MA0742.1 |
| 5.02580619 | 0.544879427774269 | 0.659 | 0.481 | 3.75E-23 | DP | MA1534.1 |
| 3.31277814 | 0.4683985656615 | 0.648 | 0.49 | 2.47E-18 | DP | MA1121.1 |
| 6.60826752 | 0.458225820848507 | 0.61 | 0.485 | 4.93E-14 | DP | MA1573.1 |
| 1.28185141 | 0.448002185822533 | 0.644 | 0.482 | 9.56E-18 | DP | MA1516.1 |
| 2.12534864 | 0.441387531968698 | 0.63 | 0.491 | 1.59E-15 | DP | MA1468.1 |
| 7.04362199 | 0.432399818338386 | 0.641 | 0.487 | 5.25E-15 | DP | MA1607.1 |
| 1.82989320 | 0.430313570891774 | 0.621 | 0.473 | 1.37E-15 | DP | MA1517.1 |
| 3.46875016 | 0.407440862850602 | 0.594 | 0.477 | 2.59E-11 | DP | MA1533.1 |
| 3.57406507 | 0.405849151135043 | 0.622 | 0.49 | 2.67E-13 | DP | MA0809.2 |

| | | | | | | |
|---|---|---|---|---|---|---|
| 4.04309105 | 0.404458969666364 | 0.613 | 0.485 | 3.02E-11 | DP | MA1549.1 |
| 1.19582680 | 0.39398080790516 | 0.608 | 0.484 | 8.92E-11 | DP | MA0510.2 |
| 3.78952096 | 0.387319699586004 | 0.61 | 0.491 | 2.83E-11 | DP | MA0090.3 |
| 4.94815730 | 0.383884295491126 | 0.633 | 0.474 | 3.69E-14 | DP | MA1515.1 |
| 6.94252986 | 0.383506752749732 | 0.604 | 0.489 | 5.18E-11 | DP | MA0685.1 |
| 1.36906496 | 0.381365435395886 | 0.582 | 0.485 | 1.02E-07 | DP | MA0116.1 |
| 9.22637020 | 0.377669768818256 | 0.624 | 0.485 | 6.88E-13 | DP | MA1512.1 |
| 4.69167419 | 0.372968144263429 | 0.605 | 0.485 | 3.50E-10 | DP | MA0647.1 |
| 4.28259978 | 0.363792277113264 | 0.593 | 0.484 | 3.19E-09 | DP | MA0739.1 |
| 3.59893482 | 0.362356782352889 | 0.599 | 0.489 | 2.68E-08 | DP | MA1535.1 |
| 1.53227041 | 0.358439117820643 | 0.605 | 0.485 | 1.14E-08 | DP | MA0600.2 |
| 1.15338849 | 0.355545610101804 | 0.59 | 0.484 | 8.60E-09 | DP | MA0019.1 |
| 1.09204161 | 0.347570931178379 | 0.59 | 0.484 | 8.15E-08 | DP | MA0493.1 |
| 1.02750109 | 0.342086301881437 | 0.616 | 0.493 | 7.67E-10 | DP | MA0148.4 |
| 4.90303798 | 0.340762247954435 | 0.578 | 0.489 | 3.66E-07 | DP | MA0657.1 |
| 5.24162812 | 0.339453663625458 | 0.59 | 0.485 | 3.91E-08 | DP | MA0678.1 |
| 1.44185978 | 0.337926138330338 | 0.591 | 0.483 | 1.08E-07 | DP | MA0668.1 |
| 8.00219483 | 0.33391278104253 | 0.591 | 0.485 | 5.97E-08 | DP | MA0079.4 |
| 9.75344475 | 0.333527781226037 | 0.602 | 0.487 | 7.28E-08 | DP | MA0498.2 |
| 1.16190298 | 0.331518763504182 | 0.585 | 0.493 | 8.67E-07 | DP | MA1117.1 |
| 8.00309108 | 0.331284390651416 | 0.575 | 0.485 | 5.97E-07 | DP | MA0509.2 |
| 3.94255091 | 0.328459789246895 | 0.611 | 0.491 | 2.94E-11 | DP | MA1513.1 |
| 1.03657249 | 0.317231358939457 | 0.585 | 0.484 | 7.73E-07 | DP | MA0637.1 |
| 1.31942964 | 0.314791936634473 | 0.593 | 0.487 | 9.84E-08 | DP | MA0746.2 |
| 2.67596858 | 0.314788293804619 | 0.582 | 0.492 | 2.00E-06 | DP | MA0602.1 |
| 8.30797795 | 0.308614121118397 | 0.579 | 0.483 | 6.20E-05 | DP | MA0799.1 |
| 8.10273013 | 0.303905110943219 | 0.584 | 0.49 | 6.04E-06 | DP | MA1641.1 |
| 1.44491232 | 0.300760636304576 | 0.587 | 0.499 | 1.08E-05 | DP | MA0633.1 |
| 2.84310241 | 0.296901400596127 | 0.584 | 0.495 | 2.12E-05 | DP | MA1103.2 |
| 6.46668626 | 0.296163717607412 | 0.575 | 0.494 | 4.82E-05 | DP | MA1683.1 |
| 7.37123469 | 0.295657421012447 | 0.601 | 0.494 | 5.50E-07 | DP | MA1487.1 |
| 5.97534546 | 0.291186097946472 | 0.593 | 0.492 | 4.46E-07 | DP | MA1102.2 |
| 2.36732983 | 0.288096983320896 | 0.568 | 0.485 | 1.77E-05 | DP | MA1105.2 |
| 4.33106545 | 0.287131629827849 | 0.596 | 0.483 | 3.23E-07 | DP | MA1511.1 |
| 1.94972158 | 0.286589944457397 | 0.584 | 0.483 | 0.000145449 | DP | MA0880.1 |
| 1.94972158 | 0.286589944457397 | 0.584 | 0.483 | 0.000145449 | DP | MA0881.1 |
| 1.06303417 | 0.285489209914419 | 0.556 | 0.478 | 7.93E-05 | DP | MA0882.1 |
| 4.77117006 | 0.28208167502613 | 0.555 | 0.486 | 0.000355929 | DP | MA1483.1 |
| 9.18722912 | 0.281857347605148 | 0.593 | 0.502 | 6.85E-05 | DP | MA0143.4 |
| 1.80862310 | 0.280829644777193 | 0.573 | 0.495 | 0.000134923 | DP | MA0069.1 |
| 3.03299824 | 0.280058905327065 | 0.579 | 0.495 | 2.26E-05 | DP | MA0107.1 |
| 8.90548687 | 0.279872011936341 | 0.581 | 0.492 | 6.64E-05 | DP | MA0461.2 |
| 7.63310261 | 0.279499017267247 | 0.573 | 0.499 | 5.69E-05 | DP | MA1120.1 |

| | | | | | | |
|---|---|---|---|---|---|---|
| 1.53186061 | 0.278304134047044 | 0.558 | 0.488 | 0.000114277 | DP | MA0163.1 |
| 1.88665856 | 0.277558714467868 | 0.558 | 0.497 | 0.001407447 | DP | MA0031.1 |
| 1.88665856 | 0.277558714467868 | 0.558 | 0.497 | 0.001407447 | DP | MA0157.2 |
| 5.46170870 | 0.276704087708745 | 0.596 | 0.494 | 4.07E-05 | DP | MA0808.1 |
| 8.69901460 | 0.269939570652967 | 0.57 | 0.482 | 0.000648946 | DP | MA1618.1 |
| 6.89164233 | 0.267091968993861 | 0.585 | 0.487 | 5.14E-05 | DP | MA0818.1 |
| 4.14858576 | 0.264803318925854 | 0.545 | 0.489 | 0.003094845 | DP | MA1470.1 |
| 1.20558556 | 0.263747338981596 | 0.573 | 0.488 | 0.000899367 | DP | MA0623.2 |
| 1.52767368 | 0.262134280151044 | 0.564 | 0.494 | 0.001139645 | DP | MA0042.2 |
| 1.52767368 | 0.262134280151044 | 0.564 | 0.494 | 0.001139645 | DP | MA0033.2 |
| 1.52767368 | 0.262134280151044 | 0.564 | 0.494 | 0.001139645 | DP | MA0848.1 |
| 1.52767368 | 0.262134280151044 | 0.564 | 0.494 | 0.001139645 | DP | MA0849.1 |
| 1.52767368 | 0.262134280151044 | 0.564 | 0.494 | 0.001139645 | DP | MA0850.1 |
| 4.38553949 | 0.25966221596038 | 0.575 | 0.495 | 0.000327161 | DP | MA0060.3 |
| 1.65156326 | 0.257742787299458 | 0.587 | 0.497 | 0.000123207 | DP | MA0852.2 |
| 1.02482998 | 0.253628167160776 | 0.538 | 0.486 | 0.007645232 | DP | MA1552.1 |
| 4.64588216 | 0.252973222030848 | 0.585 | 0.489 | 0.000346583 | DP | MA0607.1 |
| 4.53034843 | 0.645772812252065 | 0.711 | 0.505 | 3.38E-42 | HFSC | MA0142.1 |
| 7.36062397 | 0.631401724302947 | 0.726 | 0.494 | 5.49E-42 | HFSC | MA1639.1 |
| 9.85426385 | 0.624517332738541 | 0.712 | 0.506 | 7.35E-42 | HFSC | MA0730.1 |
| 8.51454099 | 0.582323654643954 | 0.677 | 0.5 | 6.35E-33 | HFSC | MA0527.1 |
| 4.26363073 | 0.56807435077993 | 0.685 | 0.501 | 3.18E-33 | HFSC | MA0601.1 |
| 6.55693151 | 0.561994597896207 | 0.656 | 0.489 | 4.89E-26 | HFSC | MA0083.3 |
| 6.80928927 | 0.533412563945198 | 0.663 | 0.494 | 5.08E-28 | HFSC | MA0259.1 |
| 7.60627521 | 0.518669265836201 | 0.649 | 0.49 | 5.67E-24 | HFSC | MA1478.1 |
| 2.08992051 | 0.493570203579303 | 0.629 | 0.488 | 1.56E-21 | HFSC | MA0112.3 |
| 2.99254039 | 0.488534438695755 | 0.645 | 0.502 | 2.23E-20 | HFSC | MA0858.1 |
| 6.57272931 | 0.480786944183445 | 0.654 | 0.496 | 4.90E-20 | HFSC | MA0670.1 |
| 3.76726329 | 0.47945674609327 | 0.647 | 0.495 | 2.81E-19 | HFSC | MA0038.2 |
| 2.22697156 | 0.462155415929142 | 0.642 | 0.501 | 1.66E-18 | HFSC | MA1466.1 |
| 3.98234504 | 0.454996814523588 | 0.652 | 0.502 | 2.97E-19 | HFSC | MA0786.1 |
| 5.48344300 | 0.43977753665591 | 0.622 | 0.492 | 4.09E-15 | HFSC | MA1475.1 |
| 4.38317534 | 0.436588364875251 | 0.64 | 0.499 | 3.27E-17 | HFSC | MA1149.1 |
| 4.06378071 | 0.43550427334213 | 0.635 | 0.496 | 3.03E-16 | HFSC | MA0106.3 |
| 3.04631038 | 0.43352265899227 | 0.656 | 0.501 | 2.27E-18 | HFSC | MA1593.1 |
| 1.17028212 | 0.430557161691258 | 0.652 | 0.5 | 8.73E-17 | HFSC | MA0108.2 |
| 2.08136290 | 0.425770920322731 | 0.646 | 0.509 | 1.55E-16 | HFSC | MA0051.1 |
| 6.14972338 | 0.421693868487645 | 0.645 | 0.489 | 4.59E-17 | HFSC | MA0901.2 |
| 2.33381097 | 0.415430947987306 | 0.642 | 0.505 | 1.74E-15 | HFSC | MA0787.1 |
| 4.16664276 | 0.411160175833831 | 0.628 | 0.498 | 3.11E-14 | HFSC | MA1527.1 |
| 5.31655178 | 0.409283404265849 | 0.619 | 0.501 | 3.97E-14 | HFSC | MA1106.1 |
| 1.06631798 | 0.407276168501272 | 0.623 | 0.504 | 7.95E-15 | HFSC | MA1624.1 |
| 3.27467021 | 0.406539342908185 | 0.606 | 0.5 | 2.44E-12 | HFSC | MA0479.1 |

| | | | | | | |
|---|---|---|---|---|---|---|
| 5.29594999 | 0.400736015173674 | 0.633 | 0.495 | 3.95E-13 | HFSC | MA0909.2 |
| 1.91654740 | 0.395217140254704 | 0.673 | 0.505 | 1.43E-16 | HFSC | MA1597.1 |
| 1.50997536 | 0.393508814411618 | 0.632 | 0.492 | 1.13E-13 | HFSC | MA0769.2 |
| 6.09874532 | 0.390035674965656 | 0.629 | 0.501 | 4.55E-14 | HFSC | MA0614.1 |
| 3.41608023 | 0.389399006146798 | 0.597 | 0.495 | 2.55E-11 | HFSC | MA0638.1 |
| 4.27815371 | 0.386371354239041 | 0.614 | 0.493 | 3.19E-12 | HFSC | MA0070.1 |
| 1.13530109 | 0.384482674676112 | 0.626 | 0.499 | 8.47E-13 | HFSC | MA0495.3 |
| 1.19277548 | 0.380712632301157 | 0.614 | 0.498 | 8.90E-13 | HFSC | MA0506.1 |
| 8.54018301 | 0.378468676010833 | 0.599 | 0.492 | 6.37E-12 | HFSC | MA0618.1 |
| 1.81164296 | 0.378233732803932 | 0.623 | 0.501 | 1.35E-12 | HFSC | MA0784.1 |
| 4.81288984 | 0.375692190723351 | 0.608 | 0.502 | 3.59E-11 | HFSC | MA0884.1 |
| 1.99972364 | 0.36657495773624 | 0.626 | 0.499 | 1.49E-11 | HFSC | MA0087.1 |
| 7.14006740 | 0.365663505667384 | 0.616 | 0.488 | 5.33E-11 | HFSC | MA0672.1 |
| 3.68437020 | 0.358691791436944 | 0.598 | 0.483 | 2.75E-10 | HFSC | MA0880.1 |
| 3.68437020 | 0.358691791436944 | 0.598 | 0.483 | 2.75E-10 | HFSC | MA0881.1 |
| 2.54971291 | 0.348668454511838 | 0.575 | 0.487 | 1.90E-09 | HFSC | MA1479.1 |
| 6.59535755 | 0.343836757732425 | 0.612 | 0.494 | 4.92E-10 | HFSC | MA0811.1 |
| 5.43952842 | 0.34293474044525 | 0.595 | 0.494 | 4.06E-09 | HFSC | MA0483.1 |
| 1.72219099 | 0.342256700478259 | 0.605 | 0.499 | 1.28E-09 | HFSC | MA0524.2 |
| 1.13905408 | 0.341220513819064 | 0.609 | 0.494 | 8.50E-10 | HFSC | MA0855.1 |
| 2.65039960 | 0.339775209674207 | 0.605 | 0.502 | 1.98E-09 | HFSC | MA0627.2 |
| 1.59317484 | 0.336163741379769 | 0.605 | 0.501 | 1.19E-08 | HFSC | MA0796.1 |
| 3.90921989 | 0.335702810738853 | 0.616 | 0.493 | 2.92E-09 | HFSC | MA1589.1 |
| 8.35410462 | 0.335653120903631 | 0.585 | 0.499 | 6.23E-08 | HFSC | MA0766.2 |
| 4.08072317 | 0.329985761731763 | 0.598 | 0.496 | 3.04E-09 | HFSC | MA0852.2 |
| 5.02816101 | 0.329520856607109 | 0.601 | 0.499 | 3.75E-08 | HFSC | MA1542.1 |
| 1.36527687 | 0.328442604800684 | 0.605 | 0.497 | 1.02E-08 | HFSC | MA0467.1 |
| 2.73070585 | 0.326291206750184 | 0.602 | 0.499 | 2.04E-08 | HFSC | MA0791.1 |
| 3.06454468 | 0.324935921207135 | 0.598 | 0.492 | 2.29E-07 | HFSC | MA0887.1 |
| 6.40032242 | 0.323211004090197 | 0.568 | 0.491 | 4.77E-06 | HFSC | MA0706.1 |
| 2.34886933 | 0.32190033172438 | 0.592 | 0.499 | 1.75E-07 | HFSC | MA0650.2 |
| 1.48843978 | 0.321732729816456 | 0.584 | 0.493 | 1.11E-07 | HFSC | MA0715.1 |
| 1.42805945 | 0.321137594734538 | 0.615 | 0.501 | 1.07E-08 | HFSC | MA0613.1 |
| 5.84698507 | 0.320738771253839 | 0.606 | 0.497 | 4.36E-08 | HFSC | MA0861.1 |
| 1.85494135 | 0.318389323243377 | 0.599 | 0.502 | 1.38E-07 | HFSC | MA0017.2 |
| 8.09875695 | 0.315683006363649 | 0.601 | 0.493 | 6.04E-08 | HFSC | MA1113.2 |
| 1.36191591 | 0.315363147330899 | 0.591 | 0.489 | 1.02E-06 | HFSC | MA1557.1 |
| 3.10571859 | 0.312977340049246 | 0.609 | 0.505 | 2.32E-07 | HFSC | MA0783.1 |
| 6.33614752 | 0.312545084027154 | 0.606 | 0.494 | 4.73E-08 | HFSC | MA1640.1 |
| 8.60431224 | 0.310758769883846 | 0.619 | 0.504 | 6.42E-09 | HFSC | MA0782.2 |
| 2.97543394 | 0.309980799952241 | 0.568 | 0.488 | 2.22E-06 | HFSC | MA0888.1 |
| 7.01307059 | 0.305098571846723 | 0.567 | 0.486 | 5.23E-06 | HFSC | MA0903.1 |
| 1.08509131 | 0.304392692210005 | 0.573 | 0.494 | 8.09E-07 | HFSC | MA0596.1 |

| | | | | | | |
|---|---|---|---|---|---|---|
| 1.55466004 | 0.304272524721487 | 0.594 | 0.495 | 1.16E-06 | HFSC | MA1583.1 |
| 5.93358219 | 0.304199650487466 | 0.592 | 0.497 | 4.43E-07 | HFSC | MA0031.1 |
| 5.93358219 | 0.304199650487466 | 0.592 | 0.497 | 4.43E-07 | HFSC | MA0157.2 |
| 2.09590287 | 0.302828423058442 | 0.611 | 0.504 | 1.56E-08 | HFSC | MA1655.1 |
| 5.55513518 | 0.300247285377795 | 0.601 | 0.505 | 4.14E-06 | HFSC | MA1485.1 |
| 1.05721052 | 0.29557619797045 | 0.616 | 0.503 | 7.89E-07 | HFSC | MA0100.3 |
| 1.75148172 | 0.292943974566555 | 0.588 | 0.488 | 1.31E-05 | HFSC | MA0742.1 |
| 1.30826128 | 0.292093467652649 | 0.592 | 0.502 | 9.76E-06 | HFSC | MA0502.2 |
| 4.43204091 | 0.286265881163512 | 0.568 | 0.495 | 3.31E-05 | HFSC | MA0859.1 |
| 3.58527750 | 0.28619861128814 | 0.616 | 0.501 | 2.67E-09 | HFSC | MA0151.1 |
| 3.62836353 | 0.28221069445927 | 0.557 | 0.491 | 2.71E-05 | HFSC | MA0844.1 |
| 6.24694641 | 0.281868318295334 | 0.563 | 0.491 | 4.66E-05 | HFSC | MA1592.1 |
| 1.62772902 | 0.280259504126328 | 0.58 | 0.504 | 1.21E-05 | HFSC | MA1604.1 |
| 1.33323544 | 0.280193710385703 | 0.574 | 0.494 | 9.95E-05 | HFSC | MA1110.1 |
| 7.01949918 | 0.27988911608029 | 0.588 | 0.501 | 5.24E-06 | HFSC | MA0507.1 |
| 5.25078313 | 0.278560093909126 | 0.578 | 0.503 | 3.92E-05 | HFSC | MA0636.1 |
| 2.79541656 | 0.27740643252843 | 0.553 | 0.492 | 0.000208538 | HFSC | MA0913.2 |
| 1.81731191 | 0.275711246550474 | 0.606 | 0.495 | 1.36E-06 | HFSC | MA0523.1 |
| 2.23963195 | 0.274958675910697 | 0.585 | 0.498 | 1.67E-05 | HFSC | MA0504.1 |
| 2.36097750 | 0.272226662997542 | 0.566 | 0.488 | 0.000176129 | HFSC | MA1493.1 |
| 1.25220622 | 0.271359870341932 | 0.566 | 0.487 | 9.34E-05 | HFSC | MA0813.1 |
| 4.70220815 | 0.26849163068164 | 0.583 | 0.496 | 3.51E-05 | HFSC | MA0468.1 |
| 3.08299996 | 0.266339244060331 | 0.549 | 0.49 | 0.000229992 | HFSC | MA1501.1 |
| 2.49489561 | 0.265002464896383 | 0.587 | 0.502 | 1.86E-06 | HFSC | MA1596.1 |
| 4.66911550 | 0.263455797609781 | 0.563 | 0.49 | 0.000348316 | HFSC | MA0810.1 |
| 4.79677950 | 0.261352866163455 | 0.57 | 0.498 | 0.00035784 | HFSC | MA0084.1 |
| 5.76165048 | 0.25824715975623 | 0.571 | 0.502 | 0.000429819 | HFSC | MA0797.1 |
| 2.08991413 | 0.257577764470436 | 0.574 | 0.496 | 0.000155908 | HFSC | MA0906.1 |
| 9.11712971 | 0.256701843519551 | 0.567 | 0.502 | 0.000680138 | HFSC | MA0676.1 |
| 2.12155235 | 0.256053961963604 | 0.578 | 0.499 | 0.000158268 | HFSC | MA0695.1 |
| 6.10502629 | 0.253877911725372 | 0.577 | 0.504 | 4.55E-05 | HFSC | MA0472.2 |
| 3.25275346 | 0.253219275079517 | 0.559 | 0.481 | 0.002426554 | HFSC | MA0470.2 |
| 8.07296533 | 0.252405364296497 | 0.573 | 0.502 | 6.02E-05 | HFSC | MA1628.1 |
| 3.10216334 | 0.252287972761617 | 0.584 | 0.494 | 2.31E-05 | HFSC | MA1114.1 |
| 1.65045094 | 0.250809089731123 | 0.592 | 0.499 | 0.000123124 | HFSC | MA1419.1 |
| 1.01876075 | 0.250204370353759 | 0.55 | 0.484 | 0.000759996 | HFSC | MA0777.1 |
| 3.98931909 | 0.824812735082504 | 0.77 | 0.491 | 2.98E-65 | LAG | MA0068.2 |
| 4.47734384 | 0.795718576567081 | 0.752 | 0.486 | 3.34E-59 | LAG | MA1577.1 |
| 1.52533797 | 0.789936292873966 | 0.78 | 0.501 | 1.14E-66 | LAG | MA0088.2 |
| 8.85267123 | 0.789505171796616 | 0.747 | 0.487 | 6.60E-54 | LAG | MA0639.1 |
| 8.10189550 | 0.775415130502333 | 0.77 | 0.501 | 6.04E-62 | LAG | MA1528.1 |
| 6.40046979 | 0.774314135545689 | 0.762 | 0.486 | 4.77E-60 | LAG | MA0892.1 |
| 4.36726432 | 0.767978459022396 | 0.761 | 0.495 | 3.26E-57 | LAG | MA0854.1 |

| | | | | | | |
|---|---|---|---|---|---|---|
| 4.77056553 | 0.740197954175557 | 0.748 | 0.498 | 3.56E-53 | LAG | MA0119.1 |
| 4.30702299 | 0.703910943850051 | 0.73 | 0.488 | 3.21E-45 | LAG | MA0907.1 |
| 1.57971642 | 0.674831398980284 | 0.72 | 0.486 | 1.18E-41 | LAG | MA0748. |
| 6.22581875 | 0.665074371699717 | 0.714 | 0.494 | 4.64E-44 | LAG | MA1643.1 |
| 2.22547276 | 0.659929009487644 | 0.734 | 0.488 | 1.66E-39 | LAG | MA0710.1 |
| 2.14773640 | 0.652251352253396 | 0.733 | 0.49 | 1.60E-39 | LAG | MA0630.1 |
| 5.58284945 | 0.649955154404399 | 0.711 | 0.492 | 4.16E-39 | LAG | MA0075. |
| 2.86734683 | 0.649421566366091 | 0.714 | 0.491 | 2.14E-39 | LAG | MA0699.1 |
| 3.15756131 | 0.636762589122223 | 0.696 | 0.491 | 2.36E-37 | LAG | MA0817.1 |
| 5.18969947 | 0.636598159368723 | 0.705 | 0.491 | 3.87E-36 | LAG | MA1463.1 |
| 1.52195281 | 0.63533575382082 | 0.708 | 0.484 | 1.14E-37 | LAG | MA0705.1 |
| 1.36565990 | 0.632174677092822 | 0.7 | 0.495 | 1.02E-37 | LAG | MA0606.1 |
| 2.59261633 | 0.628233141065433 | 0.727 | 0.489 | 1.93E-36 | LAG | MA0654.1 |
| 2.59261633 | 0.628233141065433 | 0.727 | 0.489 | 1.93E-36 | LAG | MA0720.1 |
| 2.59261633 | 0.628233141065433 | 0.727 | 0.489 | 1.93E-36 | LAG | MA0701. |
| 9.56765856 | 0.614615971609989 | 0.71 | 0.488 | 7.14E-36 | LAG | MA0874.1 |
| 5.38141062 | 0.6106216300254 | 0.688 | 0.486 | 4.01E-32 | LAG | MA0662.1 |
| 1.92735529 | 0.603450218341128 | 0.699 | 0.493 | 1.44E-33 | LAG | MA0879.1 |
| 8.68498190 | 0.591149492480029 | 0.696 | 0.495 | 6.48E-32 | LAG | MA0625.1 |
| 1.28443217 | 0.590918790087114 | 0.717 | 0.513 | 9.58E-33 | LAG | MA0526. |
| 1.57326875 | 0.583977286895304 | 0.672 | 0.496 | 1.17E-29 | LAG | MA0660.1 |
| 9.17749055 | 0.580519691163032 | 0.711 | 0.493 | 6.85E-34 | LAG | MA0501.1 |
| 2.95625494 | 0.57548466096885 | 0.705 | 0.501 | 2.21E-31 | LAG | MA1524.1 |
| 1.80768519 | 0.571810616799342 | 0.675 | 0.501 | 1.35E-26 | LAG | MA0884.1 |
| 6.04186305 | 0.571538630174731 | 0.66 | 0.484 | 4.51E-28 | LAG | MA0900. |
| 2.17654945 | 0.570373771290204 | 0.693 | 0.496 | 1.62E-26 | LAG | MA0821.1 |
| 6.60513524 | 0.569397043895068 | 0.658 | 0.476 | 4.93E-25 | LAG | MA0776.1 |
| 8.33200945 | 0.569320586476552 | 0.696 | 0.494 | 6.22E-30 | LAG | MA0853.1 |
| 7.60190101 | 0.568670446815749 | 0.688 | 0.498 | 5.67E-29 | LAG | MA1580.1 |
| 2.44054242 | 0.568608015756039 | 0.697 | 0.488 | 1.82E-29 | LAG | MA0718.1 |
| 1.07716301 | 0.56108233238351 | 0.691 | 0.488 | 8.04E-28 | LAG | MA0644.1 |
| 1.65508387 | 0.559634570047203 | 0.677 | 0.5 | 1.23E-26 | LAG | MA1525.1 |
| 1.59567660 | 0.550778913450703 | 0.677 | 0.488 | 1.19E-26 | LAG | MA0886.1 |
| 9.50542076 | 0.54985584363207 | 0.671 | 0.493 | 7.09E-27 | LAG | MA1541.1 |
| 6.09123018 | 0.545722085206729 | 0.696 | 0.494 | 4.54E-29 | LAG | MA0508. |
| 1.23646511 | 0.545449396032351 | 0.663 | 0.498 | 9.22E-26 | LAG | MA1654.1 |
| 2.90692208 | 0.544446550277907 | 0.657 | 0.498 | 2.17E-27 | LAG | MA1155.1 |
| 1.31231244 | 0.539964928156216 | 0.7 | 0.502 | 9.79E-29 | LAG | MA0152.1 |
| 2.95269239 | 0.531525419658506 | 0.674 | 0.499 | 2.20E-25 | LAG | MA1494.1 |
| 8.76393087 | 0.531021714297067 | 0.691 | 0.498 | 6.54E-27 | LAG | MA1418.1 |
| 8.12350346 | 0.522080847670897 | 0.668 | 0.501 | 6.06E-23 | LAG | MA0137. |
| 1.13471838 | 0.516108692614043 | 0.677 | 0.498 | 8.46E-25 | LAG | MA0524. |
| 6.95069857 | 0.513990633614294 | 0.677 | 0.485 | 5.19E-24 | LAG | MA0027. |

| | | | | | | |
|---|---|---|---|---|---|---|
| 2.23860912 | 0.512958489598498 | 0.668 | 0.494 | 1.67E-23 | LAG | MA0624.1 |
| 3.82771909 | 0.506688983230347 | 0.658 | 0.491 | 2.86E-21 | LAG | MA0634.1 |
| 2.87078264 | 0.50359322277581 | 0.658 | 0.494 | 2.14E-20 | LAG | MA0804.1 |
| 3.64231872 | 0.49967895152627 | 0.672 | 0.494 | 2.72E-23 | LAG | MA0150.2 |
| 5.25792731 | 0.498628249798483 | 0.672 | 0.498 | 3.92E-24 | LAG | MA1138.1 |
| 1.75555467 | 0.489675127404784 | 0.644 | 0.484 | 1.31E-20 | LAG | MA0612.2 |
| 8.12760837 | 0.48580348216285 | 0.671 | 0.497 | 6.06E-23 | LAG | MA1135.1 |
| 2.31439110 | 0.485183439303151 | 0.654 | 0.489 | 1.73E-20 | LAG | MA0889.1 |
| 4.33674593 | 0.479597695284995 | 0.644 | 0.493 | 3.24E-19 | LAG | MA0671.1 |
| 3.51255800 | 0.479314038502258 | 0.64 | 0.489 | 2.62E-18 | LAG | MA0843.1 |
| 3.40749399 | 0.47685924782255 | 0.641 | 0.486 | 2.54E-17 | LAG | MA0669.1 |
| 1.05010160 | 0.469565389655934 | 0.652 | 0.495 | 7.83E-18 | LAG | MA0773.1 |
| 1.14604813 | 0.465323539002176 | 0.635 | 0.485 | 8.55E-19 | LAG | MA0814.2 |
| 8.37730205 | 0.46443367576277 | 0.672 | 0.497 | 6.25E-20 | LAG | MA1491.1 |
| 1.46643316 | 0.464205171956475 | 0.663 | 0.496 | 1.09E-20 | LAG | MA0003.4 |
| 1.09910418 | 0.461667273164798 | 0.671 | 0.499 | 8.20E-23 | LAG | MA0149.1 |
| 1.34328089 | 0.457481555420174 | 0.668 | 0.495 | 1.00E-18 | LAG | MA1565.1 |
| 1.94323622 | 0.45634326807365 | 0.641 | 0.499 | 1.45E-16 | LAG | MA0865.1 |
| 3.25373551 | 0.444844489800139 | 0.663 | 0.513 | 2.43E-16 | LAG | MA0093.3 |
| 2.15870646 | 0.442460268411893 | 0.644 | 0.499 | 1.61E-15 | LAG | MA0758.1 |
| 6.10944161 | 0.442094507717435 | 0.64 | 0.495 | 4.56E-15 | LAG | MA0894.1 |
| 7.50426377 | 0.436454052805918 | 0.654 | 0.488 | 5.60E-17 | LAG | MA0667.1 |
| 1.55512581 | 0.429250427390849 | 0.655 | 0.493 | 1.16E-16 | LAG | MA1123.2 |
| 1.02394403 | 0.427775563448706 | 0.626 | 0.488 | 7.64E-14 | LAG | MA0895.1 |
| 5.74233836 | 0.427106157279481 | 0.641 | 0.502 | 4.28E-17 | LAG | MA0490.2 |
| 5.14743305 | 0.426840425677345 | 0.649 | 0.495 | 3.84E-16 | LAG | MA1101.2 |
| 5.92923678 | 0.425877903656603 | 0.615 | 0.492 | 4.42E-13 | LAG | MA0822.1 |
| 3.99184288 | 0.42346935563441 | 0.641 | 0.489 | 2.98E-14 | LAG | MA1481.1 |
| 3.65389886 | 0.422985187257623 | 0.651 | 0.498 | 2.73E-16 | LAG | MA1144.1 |
| 3.22388623 | 0.416832485209287 | 0.629 | 0.497 | 2.41E-13 | LAG | MA0595.1 |
| 2.18062411 | 0.413398070104651 | 0.64 | 0.499 | 1.63E-15 | LAG | MA1137.1 |
| 5.08186557 | 0.412333612310416 | 0.643 | 0.496 | 3.79E-15 | LAG | MA1647.1 |
| 2.80614243 | 0.405435166907594 | 0.635 | 0.495 | 2.09E-12 | LAG | MA1563.1 |
| 2.73266200 | 0.403645133597856 | 0.641 | 0.494 | 2.04E-12 | LAG | MA1140.2 |
| 4.97266154 | 0.395325996911336 | 0.635 | 0.498 | 3.71E-13 | LAG | MA1579.1 |
| 1.80542360 | 0.394273766230024 | 0.609 | 0.489 | 1.35E-10 | LAG | MA1557.1 |
| 4.92351220 | 0.392899101792765 | 0.643 | 0.507 | 3.67E-14 | LAG | MA1116.1 |
| 5.94987053 | 0.391108275496886 | 0.615 | 0.5 | 4.44E-12 | LAG | MA1630.1 |
| 8.00720904 | 0.387411499864144 | 0.626 | 0.503 | 5.97E-11 | LAG | MA0100.3 |
| 4.65678759 | 0.385366979312123 | 0.624 | 0.496 | 3.47E-12 | LAG | MA0828.1 |
| 1.37420644 | 0.383564046477649 | 0.596 | 0.495 | 1.03E-10 | LAG | MA0140.2 |
| 1.54661926 | 0.382507068011487 | 0.632 | 0.494 | 1.15E-11 | LAG | MA0658.1 |
| 3.17178093 | 0.380864133300308 | 0.609 | 0.489 | 2.37E-11 | LAG | MA0827.1 |

| | | | | | | |
|---|---|---|---|---|---|---|
| 2.214627409 | 0.379937842625308 | 0.61 | 0.49 | 1.65E-10 | LAG | MA0810.1 |
| 4.142375221 | 0.379118665163051 | 0.643 | 0.504 | 3.09E-13 | LAG | MA0442.2 |
| 1.870357831 | 0.375412307531198 | 0.623 | 0.495 | 1.40E-11 | LAG | MA0811.1 |
| 1.667936283 | 0.375077830500979 | 0.596 | 0.498 | 1.24E-10 | LAG | MA1638.1 |
| 4.947200390 | 0.374148978121956 | 0.623 | 0.495 | 3.69E-12 | LAG | MA0122.3 |
| 9.734668809 | 0.373392040298818 | 0.638 | 0.5 | 7.26E-12 | LAG | MA0761.2 |
| 1.371364192 | 0.372832895749751 | 0.616 | 0.491 | 1.02E-10 | LAG | MA0101.1 |
| 1.821815551 | 0.369373941636752 | 0.629 | 0.493 | 1.36E-10 | LAG | MA0092.1 |
| 4.400202447 | 0.36662077713229 | 0.606 | 0.502 | 3.28E-10 | LAG | MA1616.1 |
| 1.711805530 | 0.365723725386542 | 0.637 | 0.498 | 1.28E-11 | LAG | MA1527.1 |
| 3.497384215 | 0.36365744762334 | 0.626 | 0.499 | 2.61E-11 | LAG | MA0491.2 |
| 3.831011648 | 0.363643652916811 | 0.607 | 0.493 | 2.86E-10 | LAG | MA0902.2 |
| 3.086230071 | 0.362339222319941 | 0.634 | 0.507 | 2.30E-11 | LAG | MA0730.1 |
| 3.758660050 | 0.360586055512247 | 0.64 | 0.502 | 2.80E-12 | LAG | MA0477.2 |
| 3.368528155 | 0.357150926762558 | 0.607 | 0.493 | 2.51E-09 | LAG | MA0131.2 |
| 6.048994600 | 0.356477278532432 | 0.612 | 0.492 | 4.51E-09 | LAG | MA0072.1 |
| 2.228535635 | 0.352900833814974 | 0.635 | 0.502 | 1.66E-11 | LAG | MA1134.1 |
| 4.413430337 | 0.352648098863498 | 0.616 | 0.49 | 3.29E-09 | LAG | MA0089.2 |
| 1.522217975 | 0.351212389883429 | 0.613 | 0.493 | 1.14E-09 | LAG | MA0717.1 |
| 2.961559187 | 0.347569030777551 | 0.598 | 0.496 | 2.21E-08 | LAG | MA0468.1 |
| 3.293291810 | 0.345722841116444 | 0.595 | 0.484 | 2.46E-08 | LAG | MA0897.1 |
| 3.063113448 | 0.344360351498708 | 0.606 | 0.503 | 2.29E-09 | LAG | MA0052.4 |
| 8.895986491 | 0.340603491578419 | 0.621 | 0.504 | 6.64E-10 | LAG | MA1624.1 |
| 4.558791105 | 0.339104742137688 | 0.602 | 0.49 | 3.40E-08 | LAG | MA0806.1 |
| 5.849126271 | 0.338296426445901 | 0.602 | 0.49 | 4.36E-08 | LAG | MA0661.1 |
| 9.564097424 | 0.337459571500874 | 0.598 | 0.492 | 7.13E-08 | LAG | MA0752.1 |
| 3.971387445 | 0.333379492345717 | 0.589 | 0.485 | 2.96E-08 | LAG | MA0812.1 |
| 2.714126092 | 0.332474942132822 | 0.616 | 0.502 | 2.02E-08 | LAG | MA0620.3 |
| 2.768360251 | 0.33159356927163 | 0.635 | 0.5 | 2.07E-09 | LAG | MA0099.3 |
| 2.444307895 | 0.329808782454538 | 0.612 | 0.499 | 1.82E-08 | LAG | MA0694.1 |
| 1.959659979 | 0.326012324838063 | 0.599 | 0.493 | 1.46E-07 | LAG | MA0674.1 |
| 4.055295057 | 0.317078729061238 | 0.606 | 0.498 | 3.03E-08 | LAG | MA1130.1 |
| 4.175200008 | 0.314629794165768 | 0.59 | 0.491 | 3.11E-06 | LAG | MA0795.1 |
| 2.472706318 | 0.313241759959802 | 0.599 | 0.499 | 1.84E-08 | LAG | MA1622.1 |
| 2.024643511 | 0.313175783325646 | 0.584 | 0.495 | 1.51E-06 | LAG | MA0652.1 |
| 3.609937420 | 0.311565101168781 | 0.592 | 0.488 | 2.69E-06 | LAG | MA0908.1 |
| 2.873811610 | 0.311267725461489 | 0.576 | 0.49 | 2.14E-06 | LAG | MA0713.1 |
| 6.893081210 | 0.31099130450594 | 0.582 | 0.493 | 5.14E-07 | LAG | MA0715.1 |
| 7.121871825 | 0.310718094930441 | 0.584 | 0.483 | 5.31E-06 | LAG | MA0668.1 |
| 1.104942262 | 0.308491950045552 | 0.601 | 0.495 | 8.24E-07 | LAG | MA0698.1 |
| 3.170456581 | 0.30822131371594 | 0.578 | 0.489 | 2.37E-05 | LAG | MA0725.1 |
| 3.170456581 | 0.30822131371594 | 0.578 | 0.489 | 2.37E-05 | LAG | MA0726.1 |
| 6.277411117 | 0.308080987433283 | 0.584 | 0.495 | 4.68E-06 | LAG | MA1108.2 |

| | | | | | | |
|---|---|---|---|---|---|---|
| 1.70663101 | 0.307652464233416 | 0.606 | 0.493 | 1.27E-07 | LAG | MA0497.1 |
| 1.74415853 | 0.30700930484013 | 0.61 | 0.495 | 1.30E-07 | LAG | MA1569.1 |
| 7.40927332 | 0.306326351350273 | 0.585 | 0.492 | 5.53E-06 | LAG | MA1474.1 |
| 7.25348386 | 0.306308333909079 | 0.616 | 0.501 | 5.41E-08 | LAG | MA0476.1 |
| 3.60637283 | 0.305145477420941 | 0.561 | 0.495 | 2.69E-05 | LAG | MA0621.1 |
| 2.06522244 | 0.304021688299816 | 0.618 | 0.497 | 1.54E-07 | LAG | MA0478.1 |
| 2.24738946 | 0.303607774497426 | 0.596 | 0.506 | 1.68E-06 | LAG | MA1154.1 |
| 2.75080300 | 0.303301392935678 | 0.564 | 0.487 | 2.05E-05 | LAG | MA1603.1 |
| 1.01819205 | 0.301830979405569 | 0.587 | 0.491 | 7.60E-06 | LAG | MA0826.1 |
| 1.76982366 | 0.30089828505323 | 0.595 | 0.495 | 1.32E-05 | LAG | MA0680.1 |
| 5.42341352 | 0.299901810568257 | 0.601 | 0.501 | 4.05E-06 | LAG | MA0914.1 |
| 1.41696199 | 0.299658728765603 | 0.584 | 0.492 | 1.06E-05 | LAG | MA0704.1 |
| 5.35036085 | 0.298504458129508 | 0.589 | 0.501 | 3.99E-05 | LAG | MA0756.1 |
| 6.82369480 | 0.297075854010635 | 0.623 | 0.499 | 5.09E-09 | LAG | MA1627.1 |
| 5.24014120 | 0.295651675467215 | 0.607 | 0.495 | 3.91E-06 | LAG | MA0160.1 |
| 1.16002005 | 0.291712739432739 | 0.573 | 0.495 | 8.65E-06 | LAG | MA0515.1 |
| 4.33439130 | 0.290852868658387 | 0.601 | 0.497 | 3.23E-06 | LAG | MA0734.2 |
| 1.68698522 | 0.290802240115161 | 0.612 | 0.489 | 1.26E-06 | LAG | MA0833.2 |
| 6.57707216 | 0.288486796472726 | 0.596 | 0.489 | 4.91E-06 | LAG | MA1497.1 |
| 2.88485848 | 0.286819421182633 | 0.598 | 0.502 | 2.15E-05 | LAG | MA0143.4 |
| 9.31028682 | 0.286245798573278 | 0.612 | 0.49 | 6.95E-06 | LAG | MA0794.1 |
| 8.40718559 | 0.286189595465563 | 0.573 | 0.502 | 6.27E-05 | LAG | MA1466.1 |
| 1.13903146 | 0.282632572322059 | 0.609 | 0.501 | 8.50E-06 | LAG | MA1646.1 |
| 2.86550702 | 0.279738191514779 | 0.584 | 0.485 | 2.14E-05 | LAG | MA0678.1 |
| 4.96393483 | 0.274344065196298 | 0.59 | 0.492 | 3.70E-05 | LAG | MA1468.1 |
| 3.36532188 | 0.269000229806015 | 0.557 | 0.491 | 0.00251053 | LAG | MA0781.1 |
| 1.28741927 | 0.266932557292967 | 0.568 | 0.485 | 0.000960415 | LAG | MA0898.1 |
| 5.59177186 | 0.266215506142974 | 0.578 | 0.489 | 0.000417146 | LAG | MA0492.1 |
| 9.11731000 | 0.26480048871189 | 0.582 | 0.489 | 6.80E-05 | LAG | MA0132.2 |
| 4.93082412 | 0.26465282314807 | 0.559 | 0.495 | 0.003678395 | LAG | MA0839.1 |
| 1.68420585 | 0.261528040141537 | 0.568 | 0.497 | 0.001256418 | LAG | MA0467.1 |
| 6.20710743 | 0.260648185787496 | 0.59 | 0.501 | 4.63E-05 | LAG | MA1141.1 |
| 1.06748887 | 0.259421743275316 | 0.589 | 0.494 | 0.000796347 | LAG | MA0648.1 |
| 1.17823019 | 0.258979824270556 | 0.584 | 0.495 | 8.79E-05 | LAG | MA0030.1 |
| 3.37861335 | 0.258022014394858 | 0.578 | 0.495 | 0.000252045 | LAG | MA1500.1 |
| 9.99564685 | 0.257628427706153 | 0.564 | 0.495 | 0.000745675 | LAG | MA0592.3 |
| 6.93866264 | 0.256575577461865 | 0.592 | 0.502 | 5.18E-05 | LAG | MA0062.3 |
| 5.57880254 | 0.255339354353239 | 0.581 | 0.489 | 0.000416179 | LAG | MA1151.1 |
| 4.51252606 | 0.255240931103089 | 0.567 | 0.488 | 0.000336634 | LAG | MA1546.1 |
| 1.48853673 | 0.253993305760852 | 0.562 | 0.49 | 0.011104484 | LAG | MA0911.1 |
| 6.28351263 | 0.251337551764284 | 0.582 | 0.497 | 0.00046875 | LAG | MA0489.1 |
| 3.12064875 | 0.2510842590769 | 0.557 | 0.482 | 0.002328004 | LAG | MA1127.1 |
| 1.26454789 | 0.743287379117298 | 0.677 | 0.486 | 9.43E-05 | DS | MA0116.1 |

| | | | | | | |
|---|---|---|---|---|---|---|
| 2.22985994 | 0.654498527158438 | 0.747 | 0.494 | 1.66E-06 | DS | MA1615.1 |
| 1.91179391 | 0.577838207871873 | 0.697 | 0.499 | 0.001426198 | DS | MA1120.1 |
| 1.84391602 | 0.562045154163516 | 0.657 | 0.495 | 0.013755614 | DS | MA0596.1 |
| 1.11369232 | 0.548790941466062 | 0.667 | 0.476 | 0.008308145 | DS | MA1515.1 |
| 4.63621735 | 0.530843204153894 | 0.667 | 0.494 | 0.034586181 | DS | MA0497.1 |
| 3.85485018 | 0.508008702740592 | 0.636 | 0.486 | 0.002875718 | DS | MA0599.1 |
| 3.15632688 | 0.495175069377948 | 0.747 | 0.495 | 0.00235462 | DS | MA0155.1 |
| 2.67076850 | 0.491581407755616 | 0.636 | 0.488 | 0.019923933 | DS | MA0057.1 |
| 4.18869736 | 0.476057506209721 | 0.636 | 0.484 | 0.003124768 | DS | MA0039.4 |
| 4.06213060 | 0.449057560052153 | 0.667 | 0.496 | 0.030303494 | DS | MA1625.1 |
| 6.55707990 | 0.446546644793286 | 0.657 | 0.496 | 0.048915816 | DS | MA1125.1 |

**MotifName**
ZNF382
Nobox
KLF2
Klf12
MEIS2(var.2)
SP4
ELK4
IRF4
SP1
ETS2
DLX6
RARB(var.3)
ELK1
LHX2
RFX4
ZNF410
MEIS1(var.2)
NR1I2
EVX1
SP3
FOSB::JUN
FOSB::JUNB(var.2)
EVX2
KLF6
BACH2(var.2)
Ar
RFX5
VAX1
VAX2
HOXB7
FOS::JUN(var.2)
FOSL1::JUN(var.2)
ATF2
PBX2
RFX2
PDX1
HOXB13
SP8
KLF10
JUN::JUNB(var.2)
CREB1

ZKSCAN5
RELB
HLTF
FLI1
GATA3
REST
Ar
MAX::MYC
ZNF382
DLX6
VAX1
VAX2
ZNF384
Znf423
FOSB::JUN
LHX2
PDX1
Klf12
KLF2
RFX4
DRGX
NEUROD2
POU6F1(var.2)
Foxj3
KLF6
EVX1
BACH2(var.2)
FOSB::JUNB(var.2)
Ptf1a
MEOX2
Klf1
EGR1
NRL
RXRB
RFX7
DMRTA2
Wt1
ZNF382
LHX2
KLF6
PDX1
Znf423
NEUROD2

Ar
VAX1
VAX2
EVX1
RXRB
DLX6
RFX4
OSR1
EVX2
HOXB3
HLTF
RFX2
Klf12
RFX5
RARB(var.3)
GRHL2
Ptf1a
ZNF384
FOSB::JUNB(var.2)
KLF3
DMRTA2
GRHL1
BACH2(var.2)
MEOX2
Nobox
KLF2
DRGX
POU6F1(var.2)
Atoh1
HOXB7
NR2C1
DMRTC2
SP1
Sox11
OLIG2
ETS2
BARX2
LHX5
FOSB::JUN
RFX7
ZKSCAN1
TAL1::TCF3
PHOX2B

SP4
MEIS1(var.2)
NRL
FOSL1::JUN(var.2)
NKX6-2
SRF
Hic1
THAP1
NKX6-1
MAX::MYC
TCFL5
SOHLH2
Ahr::Arnt
HOXA6
RORA
HES5
Arnt
SMAD2::SMAD3::SMAD4
GLIS3
CREB3L4
Foxq1
HSF4
Stat2
Lhx3
TCF3
XBP1
PAX7
VDR
RORA(var.2)
ZEB1
TCF12(var.2)
ARNT2
POU6F1
ZNF135
HSF1
KLF9
MAX
Pax2
FOXB1
CLOCK
TGIF2LX
Creb3l2
ATOH1(var.2)

HES7
CUX2
FOXN3
SOX15
RORC
USF1
SPIC
TP63
HNF1A
PAX3
RORB
NKX2-8
BHLHE40
RREB1
IRF1
Spz1
TFE3
USF2
NR2F1
Atf1
ONECUT2
ONECUT1
Rhox11
SNAI3
Arntl
TGIF2LY
PBX3
MXI1
Gfi1b
FOXC1
Mlxip
RUNX1
MSGN1
HEY1
ISL2
NR1H4
FERD3L
E2F1
RFX7
Hmx1
Hmx2
TGIF2
TP53

NKX2-2
NR3C2
E2F2
GLI3
ZIC3
Sox3
Nr5a2
Zic1::Zic2
TGIF1
MNT
ZSCAN4
ZFP57
PKNOX2
HOXB9
TCF4
GLIS2
DMRTC2
POU4F2
Mecom
ZIC1
Esrrg
HOXC10
TFEC
Foxd3
Hmx3
JUN::JUNB
ATOH7
Klf1
HSF2
POU2F1
PROP1
NR2F1(var.3)
IRF7
ASCL1(var.2)
HNF1B
ATF6
TFAP4(var.2)
ZBTB26
STAT1::STAT2
ZIC4
GMEB2
BCL6
PHOX2A

PKNOX1
Npas2
ZNF143
MLXIPL
ZNF384
HOXB6
BACH2(var.2)
NR3C1
NR2C1
SOX12
MEIS2(var.2)
MZF1
TAL1::TCF3
BACH1
ZNF75D
HOXA10
HOXB7
SIX2
DUXA
HOXD10
Rarg(var.2)
RHOXF1
MYCN
ZNF460
E2F4
NFIC::TLX1
DMRTA2
LIN54
Nr2f6(var.2)
MITF
FOSB::JUNB
PAX1
TFEB
SREBF2
NR5A1
CREB3L1
PRDM1
TFAP2C(var.3)
EGR3
SOX4
MYOD1
SNAI1
ZNF449

RARA::RXRG
MEIS2
NEUROG2(var.2)
Dlx1
BHLHA15(var.2)
FOXC2
CTCF
ARNT::HIF1A
ESRRA
MAFG
CUX1
MYC
TFAP2C
TBX1
HES6
DBP
HOXD9
NFIC
SP8
Nr1h3::Rxra
TBX4
RARA::RXRA
GLI2
Atoh1
POU3F3
HOXC11
Foxl2
FOSL2::JUND
TBX2
SPIB
BHLHE41
MEIS1(var.2)
Pou5f1::Sox2
E2F7
THAP11
SP8
KLF14
SP3
TFAP2B(var.3)
SP4
TFAP2C(var.3)
HLTF
PLAG1

Isl1
TEAD2
Klf12
SP9
Znf423
KLF15
KLF11
Klf1
Dlx3
Dlx4
PHOX2B
SP1
ZNF148
ZIC5
PBX1
Foxl2
ELF2
NFKB2
SREBF2
TFAP2A(var.3)
KLF5
RFX7
MZF1(var.2)
ETV6
NKX2-5
DLX5
ELK4
OVOL2
KLF2
FOXB1
MAZ
KLF4
ZBTB7A
FOXK2
MYCN
JDP2
ZNF384
JUN
Bhlha15
RARB(var.3)
KLF13
SRF
ELF4

PPARG
KLF16
ELF5
BATF::JUN
PPARD
BATF
ATF2
ZNF652
ESR2
DLX6
KLF3
FOSL2::JUN(var.2)
CREM
SREBF1(var.2)
ZNF24
ZNF382
FOXC1
ZIC4
RXRG(var.2)
Creb5
JUN::JUNB
MSX1
MSX2
BSX
MZF1
FOXA3
NR1I3
RXRB
BATF3
TCF7
TP63
LHX1
JUND(var.2)
ZFP42
EGR2
JUND
GRHL2
MAFK
LHX2
FOS::JUN(var.2)
Nkx2-5(var.2)
NR2F6(var.3)
ETV1

HSF2
MGA
SOX8
Pparg::Rxra
TBX6
GLI3
OTX1
GLI2
Sox3
ETV4
SOX15
HNF4G
TBX5
HSF1
ONECUT2
HSF4
ELF1
GABPA
EHF
ETV6
NHLH1
ONECUT1
ZIC4
ZIC1
ERG
TBX20
ELF3
TBX21
HNF4A
Plagl1
TBX15
USF1
TGIF2
Sox6
FOXB1
TFE3
Foxd3
TCF7L1
TGIF2LX
TGIF1
TBX3
FOXC1
ETS2

Creb3l2
TBX4
SCRT2
FOXC2
ZIC5
SNAI3
RUNX1
THAP1
ELF2
ZEB1
LEF1
EOMES
Bhlha15
SNAI1
PAX3(var.2)
CTCFL
ZIC3
SPIB
NFATC4
FLI1
Rbpjl
NHLH2
RFX7
TBR1
TGIF2LY
EGR1
ZNF282
ZNF460
CTCF
FEV
SPI1
Foxj3
NFATC1
PKNOX2
PAX3
BHLHE22(var.2)
RARG(var.3)
TCF7L2
Znf423
EWSR1-FLI1
ELK4
ELF4
ZNF384

NFATC3
USF2
IRF7
Nkx3-1
NR3C1
TCF12(var.2)
NR2C2
Dmbx1
TFDP1
POU2F3
VENTX
SCRT1
HOXA9
FOSL1::JUND
IKZF1
FOXD2
TBX2
ZBTB7A
JUN::JUNB
MYBL2
MYB
E2F8
THRB(var.2)
NR1H4::RXRA
Crx
Pax2
ETS1
TCF3
ZNF135
ZNF136
SNAI2
SOX13
POU3F2
ZKSCAN5
SOX2
ERF
HES1
TBX1
Sox5
HINFP
TBX18
ZNF652
ELK3

Ptf1a(var.3)
TAL1::TCF3
BARX2
INSM1
RARB(var.3)
IRF1
PAX6
SOX14
STAT1::STAT2
Stat6
ZBTB7C
SOX9
ZBTB18
IRF3
IRF5
GCM2
Znf423
Klf12
NR1I2
Klf1
KLF2
DLX6
Dlx3
Dlx4
ZNF382
THAP11
SMAD3
RARB(var.3)
SP4
REST
Ptf1a
KLF6
SP1
TFAP4
VEZF1
POU4F1
MZF1(var.2)
CTCF
BACH2(var.2)
NKX2-5
GATA3
RXRG(var.2)
MSX1

MSX2
BSX
KLF3
Arid5a
POU6F1(var.2)
NFKB1
SP3
CDX2
MYF5
MEIS1
ESR2
PPARG
TEAD2
Ptf1a(var.2)
TCF21(var.2)
ZNF652
NKX2-2
Ddit3::Cebpa
PLAG1
PBX2
KLF11
LHX2
ZNF684
HSF2
RUNX3
RFX1
Mafb
ETS2
HOXA10
SP8
TAL1::TCF3
CTCFL
POU3F4
OVOL2
KLF14
NFKB2
RARA
HOXD12
Gmeb1
KLF10
NRL
OLIG2
MSANTD3

HOXB13
RUNX2
Hic1
NR1D2
MEIS3
Zfx
KLF4
Klf12
THAP11
PLAG1
NR1I3
Dlx3
Dlx4
MSX1
MSX2
BSX
MZF1(var.2)
KLF3
KLF14
Klf1
SP3
PHOX2B
KLF13
KLF11
FOSL2::JUN(var.2)
ZBTB7A
DLX5
Znf423
CREM
KLF15
CENPB
SP4
KLF6
MAZ
SP1
ELK4
NKX2-8
Hic1
MEIS1
ZNF382
KLF2
TEAD2
ZBTB14

CREB1
Nkx2-5(var.2)
FIGLA
SP8
MZF1
DLX6
ETV6
SP9
FOSL2::JUND(var.2)
LHX2
ZNF148
Ddit3::Cebpa
Foxl2
FOS::JUN(var.2)
ZNF24
FOSB::JUN
JDP2(var.2)
ELF2
LHX1
FOXC1
KLF4
KLF10
TFAP2C(var.3)
JUN::JUNB(var.2)
JUN
ESR2
PBX1
ETV2
TEAD4
EGR1
Nr2e3
RXRG(var.2)
ATF3
POU4F1
Creb5
Npas2
PRDM1
NR3C2
MAF::NFE2
TFAP2C(var.2)
HNF4A(var.2)
RFX4
HOXC13

TFAP2C
TFAP2B(var.2)
TBX18
REL
GABPA
NR3C1
EWSR1-FLI1
SOX18
TFAP2A
Nfe2l2
GRHL2
TBX20
PRDM4
MYBL1
Ar
NFATC4
TFAP2B
ELF1
GSX1
TWIST1
TBR1
ZNF16
TFAP2A(var.2)
Atf1
NFE2L1
PROX1
USF2
NFATC3
ZBTB18
MLX
PPARG
RFX2
Prdm15
FOSL2::JUNB
DBP
ETV1
FOSL1::JUND(var.2)
TBX4
NFATC2
STAT1
RARA::RXRA
MAFA
TEF

DUXA
SPI1
TBX15
NFATC1
MTF1
ZBTB26
MAF
Smad2::Smad3
PROP1
Stat4
ZNF263
EHF
FOS
EMX1
Zfx
ZNF24
MAX::MYC
TBX3
FOXA3
ELF5
JUN(var.2)
RFX5
MITF
TBX1
TFCP2
JUN::JUNB(var.2)
Wt1
PAX9
BCL6
BATF3
FOSL2::JUN
INSM1
POU4F2
NFAT5
FOSL1::JUNB
NFIC(var.2)
TBX5
Stat5a::Stat5b
BATF
BATF::JUN
ZNF75D
Nr1h3::Rxra
GATA1

FOS::JUN
BHLHE41
TFAP2E
LEF1
FOSL2
Nkx3-1
HOXA2
ESRRA
OSR1
EOMES
FOSL2::JUND
RELA
FOSL1::JUND
CREB3L4(var.2)
RORC
ZNF341
MNT
BACH1
HAND2
HINFP
FOXP1
ATF3
TBX6
ZFP42
NR6A1
ZBTB32
FOXP2
TBX2
JUNB
TCF3
Stat5a
HMBOX1
SOX14
TFAP4
PDX1
Dux
GBX1
ZNF652
JUND
Foxo1
ELK3
FOS::JUNB
DUX4

MAX
FOXD1
FOXO3
Sox6
Lhx8
FOSL1
Znf281
RFX3
RXRB(var.2)
MGA
NFIX
Foxj2
TCF7L2
TBX21
MAFG
SIX2
RARA(var.2)
Sox1
HLTF
DMRTC2
Nkx3-2
SPDEF
USF1
ZBTB12
HIC2
Dmrt1
EMX1
IRF9
GSX1
TEAD4
Lhx8
EN1
HOXA2
TEAD3
ETS2
PAX4
GSX2
HOXD3
IRF3
SOX21
ISX
Shox2
LHX9

ZNF274
DMRT3
RFX2
SRY
Nobox
EVX1
GLIS2
Ar
GRHL1
MIXL1
SHOX
EGR1
Smad4
RFX4
VEZF1
IRF4
TEAD1
EMX2
NOTO
TEAD2
RFX5
HOXB2
SP2
CENPB
MEF2D
Dux
TLX2
KLF15
ZKSCAN5
ZNF740
VSX1
VSX2
RFX1
KLF2
POU6F2
IRF1
E2F6
OTX1
ELK1
IRF7
Sox17
MYBL1
IRF6

RELB
EVX2
PAX9
ALX3
SP9
DRGX
NKX2-5
DMRTC2
HESX1
GBX1
ELK3
ZBTB14
HOXD11
SP4
DLX6
SP1
IRF5
HNF4A
TBX21
Alx4
LBX2
PDX1
SNAI3
SP3
Sox11
MAFA
KLF5
REL
CREB3L4(var.2)
HINFP
Znf281
ARGFX
NR3C1
RELA
BARX2
SOX4
EWSR1-FLI1
SMAD3
FOXB1
ELK4
ESX1
RFX3
OVOL2

LHX6
TBX4
Bach1::Mafk
HOXA1
HOXB5
KLF16
SCRT2
MEF2C
Arx
Atf1
TFCP2
PAX6
HNF4G
TBX6
NR2C1
PRRX2
TCF4
VENTX
OSR1
STAT1::STAT2
TBX3
SPI1
GRHL2
KLF14
HOXB3
EOMES
TBX20
SCRT1
KLF4
PAX5
TBX18
RUNX3
NR2F1(var.3)
ASCL1(var.2)
ZNF382
MEF2B
MEIS1(var.2)
FOXE1
HOXB3
MEIS2(var.2)
MEOX2
OSR1
IRF2

LBX1
LHX6
Stat5a
HESX1
IRF4
BARX1
FOXK1
Nobox
TBP
MEOX1
PBX2
EVX2
NKX6-3
ARNT::HIF1A
Arid3b
Pou5f1::Sox2
GATA5
FOXH1
EVX1
MEF2A
HIF1A
PRRX2
GATA3
PAX9
Foxj3
HOXD13
ZBTB33
CREB3L4(var.2)
FOXD1
FOXO3
RARA(var.2)
POU6F2
NR1H4
DMRTA2
FOXD2
HOXB2
THRB(var.3)
TFEB
MAX::MYC
GFI1
GBX2
Sox11
FOXG1

PBX1
RAX2
Rarb(var.2)
EMX2
MAFF
MYB
ISX
Shox2
LHX9
Foxj2
RXRB
HOXB7
NR2F1(var.2)
ZFP57
Nr1h3::Rxra
PBX3
ATF6
MAX
CREB3
HOXA4
FOXF2
FOXI1
FOXL1
FOXO4
FOXO6
FOXP3
GATA2
POU6F1(var.2)
Arid3a
PAX4
VSX1
VSX2
PKNOX1
Lhx8
ZNF382
VAX1
VAX2
POU6F1
Creb3l2
ZNF341
NOTO
Arnt
LBX2

IRF9
TBX19
FERD3L
MEIS3
Stat5a::Stat5b
MIXL1
FOXB1
GCM1
SHOX
TFDP1
MNX1
FOXA3
HOXA13
ZNF528
Arx
NKX6-1
MYC
MEF2D
RARG(var.3)
SRF
SOX13
mix-a
MEF2B
Wt1
ZKSCAN5
BHLHE41
ZNF384
USF2
PAX6
HNF1A
MEOX2
Nr2f6(var.2)
MZF1
Rarb
TFAP2A(var.2)
Rarg
RXRG(var.2)
NR1H4::RXRA
ZNF449
RXRB(var.2)
VDR
EMX2
RFX3

IRF9
MYF5
NR4A2
LMX1A
Ptf1a(var.2)
Ebf2
EBF3
EBF1
NR2C2(var.2)
ARGFX
NKX6-2
NR1D1
LHX5
ZNF140
TFAP2B
RHOXF1
ESRRB
EN2
VAX1
VAX2
FOXH1
ALX3
VENTX
Znf423
Plagl1
LEF1
IKZF1
Ascl2
HNF1B
DRGX
TFAP2C
PHOX2A
MIXL1
MYBL1
Lhx4
RORA
POU6F1
NR1H2::RXRA
HOXC8
Tcf12
GBX1
TP63
PITX2

ETS2
PAX7
PRRX1
UNCX
BHLHA15(var.2)
Zic2
OTX2
EHF
ELF3
CEBPG(var.2)
JDP2(var.2)
MNX1
PAX3
RFX2
IRF4
RFX1
MEOX1
TCF7L1
Ar
NR2F2
BHLHE22(var.2)
E2F6
NFIA
ZBTB6
GLIS1
ASCL1
BARX2
MYOD1
FOSL1
ZKSCAN5
LBX1
PDX1
NHLH2
ZNF740
IRF7
RFX5
ETV4
NR2C1
Zic1::Zic2
ZBTB18
NR3C2
SPI1
FOSB::JUN

PITX3
ZEB1
Tcf21
TP53
ETV1
TCF7L2
NR4A2::RXRA
RFX4
Znf281
NKX2-2
POU4F1
Dmbx1
HOXB3
ATF4
NKX6-1
HOXC13
IRF2
GLIS3
PAX3(var.2)
Nr2e1
NHLH1
DLX6
BACH2(var.2)
Sox17
LHX2
HOXB4
HOXD4
ZNF148
Arid3b
REST
ZBTB12
Nr5a2
PLAG1
Esrrg
NR1I2
TAL1::TCF3
RFX7
CREM
MSC
Creb5
ATF3
Nr2e3
HOXA6

NR2F1(var.3)
TBX6
FOSL1::JUNB
RARG(var.3)
PITX1
MAFA
ZKSCAN1
ONECUT1
FOSL1::JUN
FOS::JUND
Bach1::Mafk
Lhx8
LHX1
BACH1
Isl1
LMX1B
PRRX2
KLF4
FOSL2
Bhlha15
CREB1
Zfx
HOXA9
Alx4
HNF4G
Wt1
IRF1
NR3C1
ZNF652
GLI3
NKX6-3
ETV5
RARA::RXRA
FOS::JUNB
Sox3
ETV6
ISX
Shox2
LHX9
Foxl2
EVX2
MYF6
ZNF384

GCM2
PRDM1
GSX1
MTF1
ZBTB7A
mix-a
RARB(var.3)
JUN::JUNB
ELF1
RORB
HES7
Crx
SOX10
HOXC9
Sox6
PLAGL2
PPARA::RXRA
Smad2::Smad3
FOSL2::JUND
Pax2
NFIC
Rarg(var.2)
TCF21(var.2)
Pou5f1::Sox2
Ptf1a
TP73
POU6F2
NFIX
NOTO
THAP11
PLAG1
SP8
TFAP2C(var.3)
Isl1
TFAP2B(var.3)
PBX1
KLF4
KLF14
ZIC5
MZF1(var.2)
DLX5
ETV6
SP9

SP3
KLF5
JDP2
Klf1
SREBF2
OSR2
MSX1
MSX2
BSX
KLF11
ETV2
ETV4
GATA6
ONECUT1
TFAP4
PHOX2B
HES1
PAX5
Dlx3
Dlx4
ELF5
TFAP2A(var.3)
ZNF148
Msx3
MAZ
MYCN
ELF2
RAX2
ELF3
POU4F3
MEIS2
NKX2-5
LBX2
SCRT2
SP4
JUN::JUNB
THRB(var.2)
ZIC1
CREM
IKZF1
POU2F3
KLF15
DLX6

KLF16
EHF
ZBTB7A
RFX7
Creb5
BACH2
JUN
KLF13
JDP2(var.2)
FLI1
Mecom
BATF::JUN
HEY1
ZNF384
TCF7
EGR2
OTX1
ZNF460
HESX1
MZF1
SHOX
SREBF1(var.2)
FOSL1::JUND
HOXA9
ZNF282
GATA3
BATF
PPARG
ELF4
SPIB
PRRX1
UNCX
TFE3
Creb3l2
ATF3
ZIC3
SP1
OVOL2
HNF4A
ZIC4
SREBF1
ZBTB6
LHX2

GATA4
ETS1
JUND(var.2)
Ptf1a(var.2)
NFIX
FOXB1
POU2F2
HNF4G
Spz1
GSX2
HOXD3
Bhlha15
ESR2
EN2
ZNF652
ETV1
HLTF
TFEC
Rarg(var.2)
ALX3
ZNF24
FOXK2
FOXC1
HOXC11
ZFP42
RXRA::VDR
FEV
MAFK
ELK4
ZNF135
POU4F1
ONECUT3
SOX9
GCM1
EN1
ERG
JUND
RFX1
ZNF449
FOXA3
TP53
HOXA1
HOXB5

POU5F1
Foxl2
ATF2
BARHL1
TGIF2LY
SP2
GRHL2
INSM1
ZNF382
BATF3
KLF2
TBXT
Arid3a
HEY2
MYF5
MIXL1
Tcf12
FOSL2::JUN(var.2)
Nkx2-5(var.2)
Dmrt1
FOXA2
HES2
GABPA
EWSR1-FLI1
PKNOX1
BHLHA15(var.2)
Rbpjl
TP73
PITX2
TP63
ERF
YY2
DBP
STAT1
HOXC13
CEBPB
ATF4
HES5
NOTO
RBPJ
Dlx1
CEBPE
HES7

TLX2
NFATC2
ZNF16
DUXA
BHLHE23
PAX7
NFATC3
PAX4
ESX1
ZBTB32
Alx1
DPRX
SOX10
NR4A1
HOXB6
ZNF282
PRRX2
CEBPG
ZNF143
SRY
NFATC4
ZNF528
RAX
TWIST1
PRDM1
LBX2
GATA1::TAL1
MTF1
ZBTB33
NKX6-1
HIF1A
ISL2
SPDEF
USF2
USF1
BARX1
SHOX
MSGN1
Stat4
NFIC::TLX1
HOXC4
ARGFX
ATOH1(var.2)

NFIX(var.2)
NR2F2
Alx4
Rxra
E2F8
Ebf2
MNX1
SIX2
NFAT5
TFAP2C
IRF3
TEF
NFATC1
Atf1
MYBL1
SREBF2(var.2)
RHOXF1
MITF
HAND2
Nkx3-2
Arx
LIN54
GSX1
Dmbx1
PAX3
ESRRB
Hmx2
GBX1
NHLH1
GSC
MYF6
HOXA6
JUNB(var.2)
ONECUT2
LMX1B
mix-a
ARNT::HIF1A
OLIG3
Sox3
Stat5a
JUND(var.2)
ISX
Shox2

LHX9
ETV1
HEY1
TFEB
EWSR1-FLI1
NEUROD1
OTX2
CEBPD
MIXL1
NR1H2::RXRA
GLIS1
SREBF1
Sox6
Nfe2l2
ETV3
PITX3
HMBOX1
SOX21
NEUROG2
TFAP2A
ESRRA
MAF
GBX2
Nr2f6
RAX2
POU6F1
BCL6
ZBTB26
ELK3
NEUROD2
Dmrt1
Stat5a::Stat5b
REL
EMX2
LHX6
BACH2
Msx3
NR4A2
IRF5
ATF6
FOS::JUNB
BARHL1
PITX1

TBX19
ZNF75D
GLI3
STAT3
TBXT
Esrrg
MEF2A
ZNF136
Pou5f1::Sox2
SOX2
Tcf12
HOXD11
Lhx4
BHLHE22(var.2)
HINFP
VENTX
Hand1::Tcf3
TCF21(var.2)
NR2F1
Prdm15
ZNF354C
FOS::JUN
HOXA13
Hoxa11
TCF7L1
FOSL2::JUN
GSC2
MYB
Foxo1
DUX4
FOSL2::JUND
ZBTB7B
Zic1::Zic2
GABPA
PAX1
OLIG1
ELF1
FOSL2::JUNB
FOSL2::JUNB(var.2)
FERD3L
NFIX
MEIS1(var.2)
IRF4

IRF2
DMRTA2
FOXH1
CREB3L4(var.2)
ZBTB33
IRF6
OSR1
GFI1
MAFF
FOXE1
NKX6-3
TBP
RARA(var.2)
Arid3b
YY1
Foxj3
Rarb(var.2)
DMRTC2
NRF1
NFKB2
FOXG1
IRF9
Foxj2
Stat5a
OTX2
ESR1
LBX1
EGR3
SRF
NFIC(var.2)
MEF2B
GATA2
EVX1
HOXD9
ZNF528
PBX1
ZFP57
FOXK1
MEF2D
SOX18
Wt1
MEIS2(var.2)
IRF7

GATA4
NR2C2
EVX2
SP1
MEF2C
ZNF341
PKNOX1
Klf12
IRF8
KLF13
FOXF2
NR2F1(var.2)
ZSCAN4
MEF2A
ZNF382
HOXD13
NFYA
HOXA4
LIN54
Gfi1b
EGR2
Pou5f1::Sox2
SREBF2
Crx
KLF15
KLF14
THAP11
FOXC1
HES1
MZF1(var.2)
KLF11
SP8
ELF5
SREBF2
Foxl2
KLF4
PLAG1
ETV6
ELF3
ONECUT1
SP3
DLX5
KLF5

SCRT2
BATF::JUN
ZIC5
PHOX2B
SP9
ELF4
PBX1
EHF
TFAP2C(var.3)
POU2F3
RFX7
OSR2
BATF
JUN::JUNB
JDP2
HEY2
Klf1
FOXB1
ZBTB7A
POU4F3
Nkx2-5(var.2)
HEY1
SCRT1
KLF16
KLF15
ZIC3
ZNF148
OTX1
TFAP2B(var.3)
ETV1
POU4F1
ESR2
MAZ
ZNF382
TGIF2LY
PAX5
JUN
NKX2-5
BATF3
ZNF460
ETV2
ZIC1
GSX2

HOXD3
ELK4
IKZF1
POU3F1
KLF13
SP4
ELF2
BACH2
ETV4
Klf12
FOS
POU5F1B
TP73
FOXA2
CREB1
OVOL2
TFAP2A(var.3)
MZF1
THRB(var.2)
Isl1
ZNF24
TFEC
POU5F1
TFE3
JDP2(var.2)
POU3F4
INSM1
Npas2
FOS::JUNB
CREM
SREBF1
TFAP4
TP63
ATF2
FOXA3
ZIC4
FOXK2
MYB
ZNF652
JUND(var.2)
RFX1
HOXB4
HOXD4

NR1I3
Tcf12
HOXD13
HES2
GABPA
RXRG(var.2)
HLTF
TP53
TGIF2LX
SOX9
Arid3a
SP1
ATF3
FOSL1::JUND
MYCN
Rarg(var.2)
ZNF384
ETS1
Dlx3
Dlx4
Spz1
Lhx3
GATA3
FOS::JUN
EN1
TEAD2
KLF10
FOSL1::JUN
MSX1
MSX2
BSX
FLI1
MEIS2
JUND
SREBF1(var.2)
Mecom
POU3F2
MEIS1(var.2)
FOXE1
IRF2
GFI1
Stat5a
RARA(var.2)

Pou5f1::Sox2
Arid3b
OSR1
HOXB3
MAFF
MEIS2(var.2)
HESX1
MEOX2
FOXH1
ARNT::HIF1A
ZNF528
BARX1
Sox11
HOXD13
Rarb(var.2)
TBP
DUXA
HIF1A
GATA3
HOXD9
ZFP57
ZBTB33
ESR1
USF2
FOXD2
Sox5
CREB3L4(var.2)
ZNF341
GATA5
HOXB2
NKX6-3
Wt1
POU4F3
HOXA13
PKNOX1
LHX6
DMRTA2
Foxj3
HOXB7
GBX2
Foxj2
PBX1
Arid3a

FOXD1
FOXO3
LBX1
EVX2
THRB(var.3)
POU3F1
ATF6
TFEB
PBX2
EGR3
ZNF140
GATA2
MEF2A
FOXG1
Hmx2
MAF::NFE2
Rarb(var.2)
RARA::RXRG
Prdm15
MAFA
HOXC13
RXRA::VDR
NR6A1
Nfe2l2
TBX19
Hmx3
MAF
GLI2
TBX20
ATF6
EGR3
NFATC2
HMBOX1
Foxo1
GCM2
NFE2L1
RARA(var.2)
JUNB
NFIX(var.2)
HNF4A(var.2)
MAFK
Zfx
STAT1

IRF2
GLI3
RHOXF1
POU6F1
Bach1::Mafk
TFAP2C(var.2)
TBX18
PAX7
BHLHE41
PITX3
LMX1A
Hand1::Tcf3
TFE3
CREB3
ONECUT2
FOXE1
FERD3L
TFAP2B(var.2)
SREBF2(var.2)
NFIC(var.2)
NKX2-3
Hmx1
POU4F1
TFEB
JUND
CREB3L4
Pou5f1::Sox2
MXI1
Alx4
NFYC
LHX5
NKX6-1
CUX2
ZNF143
Lhx4
Arx
MTF1
BACH1
CREB3L1
CEBPD
MEIS1(var.2)
NFIC::TLX1
HIF1A

ARNT::HIF1A
DBP
PITX1
TBX4
PITX2
IRF9
RARA::RXRA
Hic1
TFAP2A
DRGX
RARA
ZNF341
PAX3
IRF8
EGR4
ZBTB32
NR4A2
RXRB(var.2)
Alx1
THRB(var.3)
Ahr::Arnt
SMAD2::SMAD3::SMAD4
Lhx3
MEIS2
LIN54
SCRT1
PRRX1
UNCX
GSC2
TGIF2LY
GSX1
Lhx8
NFIB
TGIF2LX
BACH2(var.2)
ZNF410
ZSCAN4
PKNOX1
ZNF16
TBX1
Rarg(var.2)
MYBL2
NR1D2

MYBL1
TFAP2A(var.2)
Twist2
HOXB6
NFIX
OTX2
HES7
NR1I3
TBX2
TBX5
BATF::JUN
NR2C1
Rarg
FOSL1::JUND
FOS
FOSL1::JUND(var.2)
FOXP2
JUN(var.2)
TBR1
GFI1
ESX1
GLIS1
HOXA1
HOXB5
HOXC4
PRDM1
DUX4
HOXB7
Arntl
TFAP2C
XBP1
GSC
NR2F2
NFYA
JDP2
DUXA
MAFG
Dlx2
BATF
Smad2::Smad3
USF2
Znf281
BATF3

NKX6-2
MITF
TGIF2
PDX1
MYF6
FOXK2
Stat2
ZNF684
PBX3
SOHLH2
TBX6
MEIS3
NR1D1
GLIS3
NR2C2
HOXA4
EOMES
ZKSCAN1
ZNF354C
PHOX2A
EBF3
HOXB13
MAX
HOXA6
TEF
HOXD9
THAP1
HOXA13
SREBF1(var.2)
NEUROD1
LHX6
THRB(var.2)
EN2
NR2F1
HOXB4
HOXD4
DPRX
DLX5
FOS::JUND
PKNOX2
BHLHE23
TBX3
IRF4

TBX15
BACH2
TP53
TFAP2B
RFX4
SPDEF
Mlxip
POU4F3
GATA2
NEUROG2(var.2)
MAX::MYC
EWSR1-FLI1
FOSL1
TFAP2E
Isl1
GATA1
FOXN3
MAFF
FOSL2::JUNB
FOSL2
DMRTC2
TCF3
PBX1
ARNT2
HOXA2
HOXC8
PAX6
ZNF460
STAT3
NFIC
REL
Stat5a
IRF5
Klf12
NR1I3
TEAD2
THAP11
KLF3
ATOH7
Foxl2
KLF6
NR1I2
TEAD4

POU6F1(var.2)
RFX5
TEAD1
KLF2
SP4
Znf423
KLF11
GRHL1
Hic1
NR2C1
RFX2
Ddit3::Cebpa
Klf1
FOXA1
KLF13
OLIG2
NEUROD2
SP1
MEIS1
RELB
RFX1
KLF15
CENPB
SP3
Arid5a
RFX4
MYF5
Twist2
FOXK2
FOXA3
FOXE1
CTCFL
GRHL2
KLF10
Dlx3
Dlx4
DLX6
ELF2
SOX2
PAX6
RELA
Atoh1
SOX13

PLAG1
FOXD1
FOXO3
TEAD3
Ptf1a
BHLHE22
BACH2(var.2)
NEUROG1
FOXI1
FOXL1
FOXO4
FOXO6
FOXP3
NFYA
FOXK1
RARB(var.3)
Bhlha15
Pou5f1::Sox2
MEIS1(var.2)
RARA(var.2)
ZBTB33
Arid3b
SRF
ARNT::HIF1A
DMRTA2
ESR1
Rarb(var.2)
NFIA
GFI1
ATF6
POU3F1
CREB3L4(var.2)
RARA::RXRG
TP53
ZNF317
TBP
IRF2
HOXB13
POU3F2
NFIC(var.2)
HIF1A
Stat5a
FOXH1

HOXD13
ZNF528
TCF7
Foxj2
CREB3
PBX1
MAFF
NRF1
LBX1
POU1F1
DUXA
Sox5
NKX2-3
Dlx3
Dlx4
DMRTC2
TFAP2B
Gfi1b
TFAP2C
RXRB
POU2F3
TGIF1
ZNF140
GATA5
FOXK1
OSR1
Crx
POU4F3
EVX1
MEOX2
HOXA13
PROP1
FOXG1
TP73
NR2F1
PBX2
SMAD5
PKNOX2
MEIS2(var.2)
PKNOX1
EVX2
HOXB3
SREBF2

ZFP57
FOXD1
FOXO3
ZNF341
FERD3L
MYB
Klf12
NFYB
Rarg
Arid3a
XBP1
ZNF274
Ebf2
NR1H4
POU2F2
BHLHE41
HOXD9
TCF7L2
NR2C2
HES6
TFAP2B(var.3)
DUX4
HOXB7
ZNF460
TFAP2A(var.2)
SRY
TGIF2
HOXC12
Nr2e1
ZBTB7C
EGR2
E2F4
Zic1::Zic2
PBX3
IRF4
MYBL2
PAX4
TLX2
ZNF143
DBP
NFIX(var.2)
GSX1
Alx1

NFIC::TLX1
HOXC13
YY2
NFIB
NOTO
SHOX
PRRX2
LBX2
BHLHE23
ARGFX
Lhx8
NFAT5
ISX
Shox2
LHX9
Arx
MIXL1
Dlx1
NFATC3
USF2
MEF2B
MAF::NFE2
MSGN1
DUXA
HOXA2
HES5
MYBL1
Alx4
ZBTB32
RAX
ESX1
NFATC4
EMX2
NR6A1
PRDM1
ZNF16
ZSCAN4
NFATC2
HNF4A(var.2)
IRF3
STAT1
TFAP2C
EN1

NFATC1
ALX3
TBX19
Nfe2l2
FOSL2::JUNB
EMX1
FOSB::JUNB
GBX1
NFIX
TEF
NEUROG2
MEF2D
TFAP2C(var.2)
GLI3
TFAP2A
EWSR1-FLI1
TBX18
E2F8
USF1
E2F7
HESX1
MYF6
TWIST1
HMBOX1
JUNB
BACH2
HES7
DRGX
FOSL2::JUND
SREBF1
FOSL1::JUNB
PRDM4
SOX18
JUNB(var.2)
ZBTB26
SMAD5
RBPJ
Znf281
MYB
SREBF2(var.2)
GATA1::TAL1
LHX6
OLIG3

TFAP2A(var.2)
SOX10
TFAP2B
HAND2
Nkx3-2
ETV1
REL
Hand1::Tcf3
Prdm15
NFIC(var.2)
JUND
HOXB2
RARA(var.2)
FOSL1
HINFP
RORA(var.2)
FOS::JUNB
NFE2L1
RAX2
DUX4
Hmx2
MEF2A
Stat5a
TBX4
MEOX1
ZNF410
TFAP2B(var.2)
MITF
FOS::JUN
ZBTB7B
NKX6-1
FOSL2::JUN
SMAD3
Smad2::Smad3
IRF8
HOXD11
PHOX2A
PROP1
NEUROD2
ZBTB18
VSX1
VSX2
MXI1

MEF2C
TFAP2E
CREB3L4
FOS
mix-a
FOSL2
ZNF282
Dmrt1
OLIG1
PAX7
ISL2
Lhx4
ONECUT2
Wt1
NR4A2
Sox6
GLI2
ATF4
HOXA6
SOX2
PROX1
ATF6
OSR2
OLIG2
ATOH7
PAX9
Hmx3
JUND(var.2)
PDX1
CREB3L1
Crx
FOS::JUND
GSC
FOXF2
HOXB6
ESRRA
GABPA
RORC
PAX3(var.2)
Hoxa11
JUN(var.2)
FOSB::JUN
Znf423

Plagl1
SOX13
SREBF2
KLF2
MEF2C
KLF5
INSM1
MZF1(var.2)
KLF4
Stat5b
ZNF384
