## Supplemental Table 5 for "Single-nucleus chromatin accessibility time course of excisional wound repair in rat skin"

**Supplementary Table 5. Mfuzz peak clustering summary**

**ENDO**

| peak | cluster | membership |
|---|---|---|
| chrY-15708174-15708675 | 8 | 0.993493681128311 |
| chr5-109300264-109300765 | 8 | 0.958795561668776 |
| chr14-93059757-93060258 | 5 | 0.984125437458406 |
| chr17-74392201-74392702 | 5 | 0.952628690465112 |
| chr2-93160756-93161257 | 5 | 0.982955507049856 |
| chr5-86587369-86587870 | 8 | 0.992091901304046 |
| chrX-47075428-47075929 | 8 | 0.993493681128311 |
| chr5-155798265-155798766 | 8 | 0.993493681128311 |
| chr15-65971693-65972194 | 8 | 0.867848118035522 |
| chr1-70958154-70958655 | 8 | 0.967546 |
| chrX-76223124-76223625 | 8 | 0.993493681128311 |
| chr9-64926917-64927418 | 5 | 0.983919978840557 |
| chr17-22268457-22268958 | 5 | 0.983998773265116 |
| chr2-127137359-127137860 | 8 | 0.906063465787594 |
| chr17-64877927-64878428 | 8 | 0.99138883239321 |
| chrY-17444779-17445280 | 8 | 0.993493681128311 |
| chr15-64540981-64541482 | 8 | 0.993493681128311 |
| chrX-18647967-18648468 | 8 | 0.987275677472124 |
| chrX-48199468-48199969 | 8 | 0.995225398424137 |
| chr3-38521288-38521789 | 8 | 0.993493681128311 |
| chr14-28815779-28816280 | 8 | 0.845027696230136 |
| chr14-70966794-70967295 | 5 | 0.916398680298389 |
| chr18-62634044-62634545 | 5 | 0.986807519013635 |
| chr4-16769428-16769929 | 8 | 0.991144838533906 |
| chr3-70680181-70680682 | 8 | 0.993528980243982 |
| chr7-10597046-10597547 | 8 | 0.974440705691469 |
| chr6-55998788-55999289 | 8 | 0.995880449035049 |
| chrY-1113509-1114010 | 8 | 0.829526399016336 |
| chr14-71115264-71115765 | 8 | 0.984852812071021 |
| chr9-52769884-52770385 | 8 | 0.99477257662208 |
| chr2-77091998-77092499 | 8 | 0.947236118966203 |
| chr17-17311755-17312256 | 5 | 0.983919978840557 |
| chr16-35107564-35108065 | 8 | 0.933510676523364 |
| chr16-52862837-52863338 | 8 | 0.978045030081617 |
| chr6-5142490-5142991 | 8 | 0.970344546792251 |
| chr11-797907-798408 | 8 | 0.993670321117823 |
| chr4-36890459-36890960 | 8 | 0.935825120689795 |
| chr9-6010395-6010896 | 8 | 0.992390819996944 |
| chr9-46370901-46371402 | 8 | 0.880463788724506 |
| chr4-8665173-8665674 | 4 | 0.957739732233066 |

| | | |
|---|---|---|
| chr4-8213386-8213887 | 8 | 0.994799057899685 |
| chr2-97890202-97890703 | 8 | 0.91174241488962 |
| chr3-27079361-27079862 | 8 | 0.993493681128311 |
| chr14-89358642-89359143 | 8 | 0.853039788996951 |
| chr7-36000264-36000765 | 8 | 0.942353914208759 |
| chr4-64915077-64915578 | 8 | 0.991757281709867 |
| chr11-32730648-32731149 | 5 | 0.98385936764366 |
| chr6-13270171-13270672 | 8 | 0.987101230570104 |
| chr15-58535337-58535838 | 8 | 0.848618750540606 |
| chr9-80903638-80904139 | 8 | 0.972899690150627 |
| chr5-107660068-107660569 | 8 | 0.96816738852861 |
| chr18-6806627-6807128 | 8 | 0.991617235038676 |
| chr8-4330421-4330922 | 8 | 0.920408389678508 |
| chr5-103542636-103543137 | 8 | 0.993493681128311 |
| chr4-56213323-56213824 | 5 | 0.983484956868441 |
| chr4-164115260-164115761 | 8 | 0.993461541847334 |
| chr1-131605436-131605937 | 8 | 0.967911902701974 |
| chr8-46900431-46900932 | 8 | 0.9764092961245 |
| chr3-121686982-121687483 | 8 | 0.993493681128311 |
| chr10-40690655-40691156 | 8 | 0.971565564404526 |
| chr6-78156813-78157314 | 8 | 0.98417898813506 |
| chr20-17993410-17993911 | 5 | 0.98249146735986 |
| chrY-9309548-9310049 | 4 | 0.826800721109367 |
| chr5-77607784-77608285 | 8 | 0.989133345792246 |
| chr4-9116742-9117243 | 8 | 0.982590082425067 |
| chr15-75504504-75505005 | 8 | 0.993868566808207 |
| chr16-16745976-16746477 | 8 | 0.910521155827123 |
| chr9-112022386-112022887 | 8 | 0.975590158627678 |
| chr20-26276904-26277405 | 8 | 0.850850641026862 |
| chr6-128861738-128862239 | 8 | 0.969506930406598 |
| chr1-123639832-123640333 | 5 | 0.9825422569829 |
| chr4-5190372-5190873 | 8 | 0.982318174898235 |
| chr16-65812048-65812549 | 8 | 0.901276686067279 |
| chr9-22265860-22266361 | 8 | 0.902632203825318 |
| chr16-47689923-47690424 | 8 | 0.995181567786079 |
| chr1-54852526-54853027 | 8 | 0.993493681128311 |
| chr1-149276749-149277250 | 8 | 0.994809665558215 |
| chr2-44233165-44233666 | 5 | 0.986208354640174 |
| chr16-15720760-15721261 | 8 | 0.995110642229674 |
| chr8-22852769-22853270 | 8 | 0.973294076279874 |
| chr1-115999962-116000463 | 8 | 0.994312408306141 |
| chr5-26979459-26979960 | 8 | 0.995484316995599 |
| chr3-75232185-75232686 | 8 | 0.99265150419425 |

| | | |
|---|---|---|
| chr15-86146101-86146602 | 8 | 0.959869162754413 |
| chr19-3802433-3802934 | 8 | 0.994591081305626 |
| chr9-72071819-72072320 | 8 | 0.993493681128311 |
| chr8-8233256-8233757 | 8 | 0.967952759220017 |
| chr19-3558838-3559339 | 8 | 0.994755151441339 |
| chr6-32965905-32966406 | 8 | 0.978347237170794 |
| chr4-30400426-30400927 | 8 | 0.849824171593444 |
| chr6-12859741-12860242 | 8 | 0.99458532813998 |
| chr19-13306356-13306857 | 8 | 0.98115757137448 |
| chr15-40240276-40240777 | 5 | 0.989839324383249 |
| chr4-42312935-42313436 | 8 | 0.980882170817118 |
| chr7-102686367-102686868 | 8 | 0.86734096279813 |
| chr3-2941236-2941737 | 8 | 0.993910176423607 |
| chr7-104450921-104451422 | 8 | 0.993493681128311 |
| chr14-64099569-64100070 | 8 | 0.88577278802974 |
| chr4-27044255-27044756 | 8 | 0.944450446424014 |
| chr19-43125110-43125611 | 8 | 0.993493681128311 |
| chr6-35486470-35486971 | 8 | 0.945001040299025 |
| chr20-52406134-52406635 | 8 | 0.87497053559258 |
| chr13-21430262-21430763 | 8 | 0.993493681128311 |
| chr17-15365255-15365756 | 5 | 0.981271632672218 |
| chr1-112673706-112674207 | 8 | 0.883777491152626 |
| chr2-238970154-238970655 | 8 | 0.981018596069797 |
| chr18-77726148-77726649 | 5 | 0.980568395128997 |
| chr9-3905494-3905995 | 2 | 0.92554808419864 |
| chr14-36316231-36316732 | 5 | 0.981874880937004 |
| chr3-139557809-139558310 | 8 | 0.983422749688893 |
| chr10-24071134-24071635 | 8 | 0.821173125857289 |
| chr19-16820040-16820541 | 8 | 0.991839828358793 |
| chr5-53931527-53932028 | 8 | 0.931106138964787 |
| chr8-74608147-74608648 | 8 | 0.904478650913658 |
| chr14-66328052-66328553 | 5 | 0.991079822539052 |
| chr19-20825416-20825917 | 8 | 0.993493681128311 |
| chr2-238583326-238583827 | 8 | 0.993203390477383 |
| chr3-65145925-65146426 | 8 | 0.979011246605684 |
| chr9-70022355-70022856 | 8 | 0.976215649478478 |
| chr11-49072555-49073056 | 8 | 0.915487812139952 |
| chr3-43209743-43210244 | 5 | 0.850231787467224 |
| chrX-124224758-124225259 | 8 | 0.990230744725132 |
| chr9-4501793-4502294 | 8 | 0.993493681128311 |
| chr5-122670171-122670672 | 5 | 0.983919978840557 |
| chr17-67230592-67231093 | 8 | 0.986185525328834 |
| chr5-66698531-66699032 | 8 | 0.996667400829965 |

| | | |
|---|---|---|
| chr2-66667841-66668342 | 8 | 0.973869970327606 |
| chr5-47988604-47989105 | 8 | 0.946441284463128 |
| chr13-53510761-53511262 | 8 | 0.98219216533091 |
| chr14-23187580-23188081 | 2 | 0.947074083661991 |
| chr17-41100473-41100974 | 8 | 0.991850261858557 |
| chr9-60315627-60316128 | 8 | 0.974054330862821 |
| chr1-162199083-162199584 | 8 | 0.993493681128311 |
| chr17-57384322-57384823 | 8 | 0.984072247074317 |
| chr4-35088942-35089443 | 8 | 0.93197550631642 |
| chr5-132190478-132190979 | 8 | 0.995248699227914 |
| chr9-111254621-111255122 | 5 | 0.985894069839047 |
| chr14-24637198-24637699 | 8 | 0.891344725390005 |
| chr5-26820569-26821070 | 8 | 0.917306940494404 |
| chr5-42249354-42249855 | 4 | 0.875804945412341 |
| chr8-9246081-9246582 | 8 | 0.995228231784032 |
| chr9-27311241-27311742 | 8 | 0.949912339358975 |
| chr17-75704437-75704938 | 8 | 0.991429721225465 |
| chr10-78663782-78664283 | 8 | 0.993493681128311 |
| chr10-77870145-77870646 | 8 | 0.993493681128311 |
| chr19-38018601-38019102 | 5 | 0.98569915425709 |
| chr7-40230877-40231378 | 8 | 0.92620292095317 |
| chr17-14619470-14619971 | 8 | 0.998104301778241 |
| chr17-29848257-29848758 | 5 | 0.987068176633204 |
| chr8-54665942-54666443 | 5 | 0.987216081252406 |
| chr6-52646918-52647419 | 8 | 0.882499354297941 |
| chr2-4860066-4860567 | 2 | 0.892669950137788 |
| chr6-94850835-94851336 | 8 | 0.973633028462932 |
| chr13-31954400-31954901 | 8 | 0.982880012121234 |
| chr13-16024268-16024769 | 8 | 0.993493681128311 |
| chr5-32451321-32451822 | 8 | 0.993493681128311 |
| chr3-123001940-123002441 | 8 | 0.984816575361036 |
| chr15-74388327-74388828 | 8 | 0.876870011794064 |
| chr2-231787856-231788357 | 8 | 0.987274710883997 |
| chr10-28740632-28741133 | 8 | 0.96398457225449 |
| chr15-72266998-72267499 | 8 | 0.825999994614731 |
| chr5-17501091-17501592 | 8 | 0.994886656951694 |
| chr5-118485856-118486357 | 8 | 0.89128353535207 |
| chr7-23680052-23680553 | 8 | 0.983616030238817 |
| chr1-36961684-36962185 | 8 | 0.986247376793702 |
| chr2-37406373-37406874 | 8 | 0.994824562733065 |
| chr9-34642809-34643310 | 8 | 0.977399554552729 |
| chr1-163194157-163194658 | 8 | 0.956394400682913 |
| chr14-102339054-102339555 | 8 | 0.895637521506452 |

| | | |
|---|---|---|
| chr18-51017116-51017617 | 8 | 0.981522166370674 |
| chr14-84972469-84972970 | 5 | 0.985225306888484 |
| chr1-5510320-5510821 | 8 | 0.990304537074715 |
| chr19-907601-908102 | 5 | 0.983919978840557 |
| chr4-31168307-31168808 | 8 | 0.979728768971533 |
| chr7-115473306-115473807 | 8 | 0.994087746335188 |
| chr9-34716127-34716628 | 8 | 0.993493681128311 |
| chr18-1266882-1267383 | 8 | 0.954629244647072 |
| chr13-23304020-23304521 | 8 | 0.991231024460415 |
| chr7-105942547-105943048 | 8 | 0.974978881940353 |
| chr14-82874828-82875329 | 5 | 0.984744164692953 |
| chr18-40321498-40321999 | 4 | 0.971295282624218 |
| chr6-51613936-51614437 | 8 | 0.991028506399272 |
| chr2-130201967-130202468 | 8 | 0.993493681128311 |
| chr2-20446095-20446596 | 8 | 0.943941739141448 |
| chr8-116268674-116269175 | 8 | 0.995177853940886 |
| chr6-111062642-111063143 | 8 | 0.993493681128311 |
| chr2-159554158-159554659 | 8 | 0.993419775558755 |
| chr13-37168454-37168955 | 5 | 0.978922939968277 |
| chrX-120224911-120225412 | 4 | 0.903865676061069 |
| chr6-54559908-54560409 | 5 | 0.972335126007772 |
| chr9-104979460-104979961 | 4 | 0.842128240513108 |
| chrX-30041359-30041860 | 8 | 0.93574571607651 |
| chr14-46553310-46553811 | 8 | 0.964946225097359 |
| chr18-42191509-42192010 | 8 | 0.983309908473751 |
| chr5-89143986-89144487 | 8 | 0.993493681128311 |
| chr20-26616688-26617189 | 5 | 0.986034921731301 |
| chr5-14995211-14995712 | 8 | 0.983198160270592 |
| chr15-62158126-62158627 | 8 | 0.993493681128311 |
| chr13-49987570-49988071 | 8 | 0.994601960949552 |
| chr2-99846395-99846896 | 5 | 0.957804452989393 |
| chr4-108547182-108547683 | 8 | 0.932804868436361 |
| chr2-99925261-99925762 | 8 | 0.994738259641846 |
| chr14-46905988-46906489 | 8 | 0.994583408033587 |
| chr12-23395877-23396378 | 8 | 0.980126124987432 |
| chr2-147703040-147703541 | 8 | 0.953002401800277 |
| chr6-12519146-12519647 | 8 | 0.993671730396514 |
| chr4-85870293-85870794 | 8 | 0.879230731511026 |
| chr7-64913682-64914183 | 8 | 0.993493681128311 |
| chr20-18091580-18092081 | 5 | 0.98677682712155 |
| chr14-63936785-63937286 | 8 | 0.993493681128311 |
| chr14-74456831-74457332 | 5 | 0.962303729898066 |
| chr7-23971439-23971940 | 8 | 0.993493681128311 |

| | | |
|---|---|---|
| chr8-1082536-1083037 | 8 | 0.990274186870349 |
| chr16-62953427-62953928 | 8 | 0.893216430369647 |
| chr8-27053574-27054075 | 4 | 0.842061902296986 |
| chr14-41160153-41160654 | 5 | 0.987479089299243 |
| chr2-26200884-26201385 | 8 | 0.935684763494005 |
| chrX-107956121-107956622 | 8 | 0.984153155370198 |
| chr6-1816038-1816539 | 8 | 0.976392727294192 |
| chr10-56881833-56882334 | 4 | 0.928753222752955 |
| chr13-34829981-34830482 | 8 | 0.967318679486366 |
| chr4-161460413-161460914 | 8 | 0.985601121741654 |
| chr13-42973886-42974387 | 5 | 0.968898889481903 |
| chrX-108139844-108140345 | 8 | 0.98006123025597 |
| chrY-2288755-2289256 | 8 | 0.993493681128311 |
| chr4-70581650-70582151 | 8 | 0.992368155958371 |
| chrX-92344760-92345261 | 5 | 0.983919978840557 |
| chr13-65992634-65993135 | 8 | 0.991136079758481 |
| chr5-53793601-53794102 | 8 | 0.993493681128311 |
| chr3-21742203-21742704 | 5 | 0.98329053983582 |
| chr3-8525944-8526445 | 5 | 0.979063939412483 |
| chr5-13566440-13566941 | 8 | 0.948169218247352 |
| chr4-89434381-89434882 | 8 | 0.994717003641562 |
| chr14-17609016-17609517 | 8 | 0.929913503202556 |
| chr2-207881142-207881643 | 8 | 0.991591141490171 |
| chr5-73774938-73775439 | 8 | 0.855281398906843 |
| chr15-7313490-7313991 | 8 | 0.987901044080497 |
| chr19-38122691-38123192 | 5 | 0.983919978840557 |
| chr6-135635299-135635800 | 6 | 0.991206925481043 |
| chr7-5584203-5584704 | 8 | 0.985065975368391 |
| chr12-45749021-45749522 | 8 | 0.949276794216377 |
| chr15-60311164-60311665 | 8 | 0.847020418686207 |
| chr6-28746835-28747336 | 8 | 0.924682507602561 |
| chr2-157964679-157965180 | 8 | 0.862466951128242 |
| chr3-80828646-80829147 | 8 | 0.843801979942107 |
| chr8-40499257-40499758 | 2 | 0.911361906890065 |
| chr2-47051206-47051707 | 8 | 0.914875124215162 |
| chr4-161471111-161471612 | 8 | 0.8719494223464 |
| chr15-51730147-51730648 | 8 | 0.96334323715835 |
| chrX-39299351-39299852 | 3 | 0.994448246298691 |
| chr14-1866659-1867160 | 8 | 0.854970985537885 |
| chr13-40737122-40737623 | 8 | 0.994980764625701 |
| chr8-8104788-8105289 | 8 | 0.991131247975656 |
| chr5-26793391-26793892 | 8 | 0.993493681128311 |
| chr14-32764594-32765095 | 5 | 0.985065996616026 |

| | | |
|---|---|---|
| chr2-14443969-14444470 | 8 | 0.983438843264657 |
| chr18-7754207-7754708 | 8 | 0.99251286743303 |
| chr14-55083264-55083765 | 8 | 0.974556728248236 |
| chr16-65872365-65872866 | 8 | 0.968056311815145 |
| chr9-50327986-50328487 | 2 | 0.860752838234212 |
| chr1-124144874-124145375 | 8 | 0.944331821960144 |
| chr1-149925570-149926071 | 8 | 0.990359558447425 |
| chr17-64754284-64754785 | 8 | 0.843629599593488 |
| chr20-21840287-21840788 | 5 | 0.984518426166223 |
| chr12-34931648-34932149 | 8 | 0.987526912702572 |
| chr12-1651181-1651682 | 5 | 0.990557417570851 |
| chr9-52546232-52546733 | 4 | 0.875773422711318 |
| chr13-37918433-37918934 | 5 | 0.986093060104065 |
| chr3-80609177-80609678 | 8 | 0.853000042444797 |
| chr11-51921894-51922395 | 4 | 0.911945271368724 |
| chr9-56002787-56003288 | 4 | 0.8612552064817 |
| chr1-102842716-102843217 | 8 | 0.984004411727376 |
| chr2-136874898-136875399 | 8 | 0.976794708727254 |
| chrX-115747293-115747794 | 8 | 0.956347744055521 |
| chr17-64153826-64154327 | 5 | 0.983789417427831 |
| chr6-82141934-82142435 | 8 | 0.897811173981676 |
| chr2-92853936-92854437 | 8 | 0.981960490015543 |
| chr10-496341-496842 | 8 | 0.98161261650018 |
| chr3-151566558-151567059 | 8 | 0.979522416799372 |
| chr19-3926698-3927199 | 8 | 0.995070868281434 |
| chr18-25668825-25669326 | 8 | 0.988590287603198 |
| chr1-36183304-36183805 | 8 | 0.993493681128311 |
| chr4-108273672-108274173 | 8 | 0.993493681128311 |
| chr13-38959494-38959995 | 5 | 0.932399063232293 |
| chr20-2727453-2727954 | 5 | 0.984883371491804 |
| chr10-3551029-3551530 | 8 | 0.833899552662919 |
| chr17-52384395-52384896 | 5 | 0.85595074125913 |
| chr11-58457813-58458314 | 8 | 0.982356465419085 |
| chr4-179654206-179654707 | 8 | 0.924719572181884 |
| chr1-123626343-123626844 | 8 | 0.990465237914372 |
| chr15-89635976-89636477 | 8 | 0.830995246826293 |
| chr4-95975177-95975678 | 8 | 0.996002922378557 |
| chr7-97126241-97126742 | 8 | 0.972699590178749 |
| chr7-72964215-72964716 | 8 | 0.937104476730396 |
| chrX-21900768-21901269 | 8 | 0.994864406059897 |
| chr6-12265190-12265691 | 8 | 0.8997885196193 |
| chr1-58310200-58310701 | 8 | 0.991689139497546 |
| chr3-129941292-129941793 | 8 | 0.993493681128311 |

| | | |
|---|---|---|
| chr6-110478956-110479457 | 8 | 0.823720096893868 |
| chr7-92915427-92915928 | 8 | 0.946828335059818 |
| chr1-106284671-106285172 | 3 | 0.90106439003347 |
| chr11-9489794-9490295 | 8 | 0.979760617220649 |
| chr4-49086062-49086563 | 8 | 0.994150229790403 |
| chr20-32546427-32546928 | 8 | 0.876265156127024 |
| chr16-72238036-72238537 | 8 | 0.986334331553816 |
| chr5-165914997-165915498 | 8 | 0.931787124582649 |
| chr4-17259269-17259770 | 8 | 0.993962797121121 |
| chrX-27354673-27355174 | 2 | 0.96655529441928 |
| chr4-9713596-9714097 | 5 | 0.984657618632882 |
| chr6-102910185-102910686 | 5 | 0.984650288884347 |
| chr8-15029923-15030424 | 5 | 0.988205666187535 |
| chr5-64815822-64816323 | 8 | 0.949550738571963 |
| chr7-119398643-119399144 | 8 | 0.995050011598543 |
| chr16-70809548-70810049 | 8 | 0.993385599123466 |
| chr20-938914-939415 | 8 | 0.99295920654811 |
| chr17-74134217-74134718 | 5 | 0.908310972470566 |
| chr1-108563338-108563839 | 8 | 0.930280108164802 |
| chr7-58794091-58794592 | 8 | 0.887451036946515 |
| chr2-33312636-33313137 | 4 | 0.888963957601604 |
| chr8-119613031-119613532 | 8 | 0.940673769473372 |
| chr17-20612319-20612820 | 5 | 0.990008320073211 |
| chr13-84859050-84859551 | 8 | 0.957403670176395 |
| chr19-29695502-29696003 | 8 | 0.995137098546366 |
| chr2-108760806-108761307 | 2 | 0.875769100744413 |
| chr6-73576940-73577441 | 8 | 0.987249242028566 |
| chr18-67628559-67629060 | 5 | 0.983919978840557 |
| chr19-16692290-16692791 | 8 | 0.86018610428088 |
| chr11-5696514-5697015 | 8 | 0.951766273679108 |
| chr10-64501483-64501984 | 8 | 0.993066848235726 |
| chr3-95055453-95055954 | 8 | 0.976558384729152 |
| chr17-82980587-82981088 | 8 | 0.956353717930165 |
| chr8-52966936-52967437 | 8 | 0.888073542415142 |
| chr18-9079089-9079590 | 8 | 0.90206281269326 |
| chr14-27118636-27119137 | 8 | 0.90420407582461 |
| chr5-86717432-86717933 | 8 | 0.993493681128311 |
| chr2-50865701-50866202 | 8 | 0.95266651641169 |
| chr16-22503096-22503597 | 3 | 0.996845172378378 |
| chr6-46027697-46028198 | 8 | 0.994666460947865 |
| chr4-68383025-68383526 | 8 | 0.976476650857761 |
| chr10-32812972-32813473 | 8 | 0.941743941909019 |
| chr4-3832104-3832605 | 8 | 0.944451608663439 |

| | | |
|---|---|---|
| chr7-60805496-60805997 | 8 | 0.942927913908722 |
| chr7-72102408-72102909 | 8 | 0.991719268972281 |
| chr4-63939150-63939651 | 8 | 0.986100692542952 |
| chr9-57514701-57515202 | 8 | 0.99384277273131 |
| chr17-28786758-28787259 | 8 | 0.994901050099397 |
| chr3-20104096-20104597 | 5 | 0.986648592778108 |
| chr13-36521228-36521729 | 5 | 0.984653991748425 |
| chr18-4711533-4712034 | 8 | 0.962560758881249 |
| chr8-42254880-42255381 | 8 | 0.995229596585266 |
| chr2-194758132-194758633 | 8 | 0.89489788256503 |
| chr13-58986119-58986620 | 8 | 0.967613161857545 |
| chr5-20251921-20252422 | 4 | 0.863633654629993 |
| chr16-40860290-40860791 | 8 | 0.919812783473456 |
| chr4-148085535-148086036 | 5 | 0.987499641800644 |
| chr2-146021021-146021522 | 3 | 0.996907617023073 |
| chr15-55914418-55914919 | 3 | 0.99593553545041 |
| chr11-7298572-7299073 | 8 | 0.995090167347922 |
| chr4-93269470-93269971 | 8 | 0.989607101822331 |
| chr16-10534224-10534725 | 2 | 0.948909750267931 |
| chr9-190637-191138 | 5 | 0.984678751375717 |
| chr6-93698320-93698821 | 8 | 0.860669336863346 |
| chr11-28156141-28156642 | 8 | 0.921283901293079 |
| chr14-68331377-68331878 | 8 | 0.915620674341279 |
| chr19-6541890-6542391 | 8 | 0.91120778222089 |
| chr3-133881659-133882160 | 5 | 0.839456230327144 |
| chr10-9981627-9982128 | 8 | 0.928697748362073 |
| chr20-17395279-17395780 | 8 | 0.888627835983104 |
| chr2-48823371-48823872 | 8 | 0.869255409526004 |
| chr11-20659186-20659687 | 8 | 0.944439366715963 |
| chr2-4929027-4929528 | 8 | 0.990730862510927 |
| chr12-7022618-7023119 | 8 | 0.993890847506996 |
| chr17-52470192-52470693 | 5 | 0.983864 |
| chr18-34141841-34142342 | 8 | 0.985071890096964 |
| chr2-164169093-164169594 | 8 | 0.990990426931407 |
| chr18-63946344-63946845 | 8 | 0.994041033618807 |
| chr15-90400810-90401311 | 8 | 0.94940676698491 |
| chr14-52431878-52432379 | 8 | 0.978768234648044 |
| chr3-123363606-123364107 | 8 | 0.993493681128311 |
| chr16-26159608-26160109 | 8 | 0.993493681128311 |
| chr5-43211722-43212223 | 8 | 0.989231373134666 |
| chr16-70046350-70046851 | 8 | 0.978577291915795 |
| chr17-33471862-33472363 | 8 | 0.968799814227143 |
| chr2-38472362-38472863 | 5 | 0.983433895649931 |

| | | |
|---|---|---|
| chr3-75485914-75486415 | 8 | 0.993493681128311 |
| chr4-3904151-3904652 | 4 | 0.937911885708954 |
| chr7-39123641-39124142 | 8 | 0.861916092266972 |
| chr1-109906390-109906891 | 4 | 0.844742546010523 |
| chr3-50417540-50418041 | 8 | 0.990074226372889 |
| chr19-42639397-42639898 | 5 | 0.876579228019518 |
| chr19-41488356-41488857 | 8 | 0.99238914730466 |
| chr1-161161778-161162279 | 8 | 0.984192354842239 |
| chr14-93624661-93625162 | 5 | 0.979406333792032 |
| chr14-41470789-41471290 | 8 | 0.992846279864699 |
| chr12-30438960-30439461 | 8 | 0.919331253491037 |
| chr3-103499319-103499820 | 8 | 0.891760249029318 |
| chrX-32243011-32243512 | 8 | 0.890201501760445 |
| chr7-45922992-45923493 | 8 | 0.877682576261248 |
| chr5-13844824-13845325 | 4 | 0.971875957608875 |
| chr14-81649693-81650194 | 8 | 0.98299610455106 |
| chr17-14818471-14818972 | 5 | 0.987011924700892 |
| chr7-27985716-27986217 | 8 | 0.974326432439152 |
| chr7-27330960-27331461 | 8 | 0.896083946067046 |
| chr3-101070139-101070640 | 8 | 0.942243116958345 |
| chr17-66116574-66117075 | 4 | 0.953420724010681 |
| chr1-25930811-25931312 | 5 | 0.99663839728598 |
| chr8-12764265-12764766 | 4 | 0.858387944820328 |
| chr15-16308824-16309325 | 8 | 0.993493681128311 |
| chrX-88115935-88116436 | 8 | 0.993493681128311 |
| chr1-222438081-222438582 | 2 | 0.884298233412806 |
| chr4-60729107-60729608 | 8 | 0.916290279319513 |
| chr10-74607319-74607820 | 8 | 0.993493681128311 |
| chr5-152446566-152447067 | 8 | 0.993493681128311 |
| chr2-185460977-185461478 | 8 | 0.980456469070269 |
| chr9-101548981-101549482 | 8 | 0.892692185924578 |
| chr9-71269361-71269862 | 2 | 0.90215506793593 |
| chr15-13603194-13603695 | 5 | 0.939421501669618 |
| chr13-40686142-40686643 | 8 | 0.920420887333955 |
| chr3-111442151-111442652 | 5 | 0.944092043166001 |
| chr9-36857129-36857630 | 8 | 0.992443269749564 |
| chrX-2357493-2357994 | 3 | 0.945165438438565 |
| chr10-75642936-75643437 | 8 | 0.986090827809511 |
| chr1-130200081-130200582 | 8 | 0.987652080603029 |
| chr20-47635022-47635523 | 8 | 0.964954554524962 |
| chr15-49939718-49940219 | 5 | 0.984311673678588 |
| chr19-19575643-19576144 | 5 | 0.989074415916574 |
| chr3-51128210-51128711 | 8 | 0.993493681128311 |

| | | |
|---|---|---|
| chr14-11910716-11911217 | 5 | 0.98446566417477 |
| chr4-1699980-1700481 | 8 | 0.895018136892462 |
| chr19-42869319-42869820 | 8 | 0.988258670933178 |
| chr18-75732358-75732859 | 8 | 0.995047702950434 |
| chr1-60036993-60037494 | 8 | 0.993493681128311 |
| chrX-4412142-4412643 | 8 | 0.90846368010814 |
| chr17-73529669-73530170 | 5 | 0.983919978840557 |
| chr8-33183363-33183864 | 8 | 0.995247910271004 |
| chr2-180157914-180158415 | 8 | 0.982371875146801 |
| chr4-88010842-88011343 | 8 | 0.995133877401295 |
| chr14-7302779-7303280 | 8 | 0.993493681128311 |
| chr6-50633812-50634313 | 8 | 0.993493681128311 |
| chr7-132913037-132913538 | 8 | 0.993493681128311 |
| chr2-217593843-217594344 | 8 | 0.940685488909854 |
| chr13-9713410-9713911 | 8 | 0.986348139447226 |
| chr11-81387960-81388461 | 8 | 0.985642585221884 |
| chr9-109340808-109341309 | 4 | 0.910012211317569 |
| chrX-22015191-22015692 | 5 | 0.839394985620991 |
| chr6-19784775-19785276 | 8 | 0.993493681128311 |
| chr18-92060-92561 | 8 | 0.993493681128311 |
| chr10-18877633-18878134 | 8 | 0.997343558410418 |
| chr18-12143856-12144357 | 8 | 0.992456123484421 |
| chr13-8176215-8176716 | 8 | 0.99394888872103 |
| chr6-30171242-30171743 | 6 | 0.991206925481043 |
| chr4-100220870-100221371 | 4 | 0.905962641479051 |
| chr2-147098678-147099179 | 8 | 0.992948213618544 |
| chr6-19090779-19091280 | 8 | 0.991417528375208 |
| chr13-45664710-45665211 | 5 | 0.96768675613389 |
| chr7-12443555-12444056 | 8 | 0.93834044055028 |
| chr1-176423304-176423805 | 3 | 0.987818522888539 |
| chr9-42045810-42046311 | 8 | 0.932628609195726 |
| chr18-17216504-17217005 | 8 | 0.958356384565137 |
| chr4-65245546-65246047 | 8 | 0.992715675217498 |
| chr3-103555862-103556363 | 8 | 0.973774349597735 |
| chr4-105640726-105641227 | 8 | 0.993493681128311 |
| chr3-38375884-38376385 | 8 | 0.993560943267215 |
| chr2-148511672-148512173 | 8 | 0.896449172463115 |
| chr6-120271591-120272092 | 8 | 0.970050119886828 |
| chr6-6809417-6809918 | 8 | 0.954254237150567 |
| chr4-136380652-136381153 | 2 | 0.952034439695635 |
| chr3-157842327-157842828 | 4 | 0.908932993437046 |
| chr2-199301713-199302214 | 8 | 0.974368102374728 |
| chr4-177816552-177817053 | 5 | 0.992172685644717 |

| | | |
|---|---|---|
| chr4-64764219-64764720 | 8 | 0.907581496258744 |
| chr4-22646198-22646699 | 8 | 0.975063773909594 |
| chr18-80647564-80648065 | 8 | 0.992953955776914 |
| chr5-92138780-92139281 | 8 | 0.950065682988647 |
| chrX-123553870-123554371 | 8 | 0.99523063766179 |
| chr8-65512849-65513350 | 5 | 0.98639252364494 |
| chr13-52407294-52407795 | 8 | 0.971594255374082 |
| chr3-157988588-157989089 | 8 | 0.976963716108602 |
| chr2-213631469-213631970 | 8 | 0.965076074801576 |
| chr14-92041039-92041540 | 5 | 0.982634423262301 |
| chr15-47316039-47316540 | 8 | 0.971361507495561 |
| chr20-24583985-24584486 | 8 | 0.994421289787373 |
| chr7-19136583-19137084 | 8 | 0.870023572289631 |
| chr2-15430613-15431114 | 8 | 0.968486826766069 |
| chr4-23895795-23896296 | 8 | 0.81863357695741 |
| chr17-69945272-69945773 | 8 | 0.882512734645728 |
| chr11-60011847-60012348 | 8 | 0.987015118279105 |
| chr10-2032544-2033045 | 8 | 0.99335536799112 |
| chr10-59738252-59738753 | 5 | 0.988629864857282 |
| chr14-72772428-72772929 | 8 | 0.994623468176911 |
| chr6-14775775-14776276 | 8 | 0.962002317960396 |
| chr11-81401264-81401765 | 8 | 0.991127334275943 |
| chr14-3056370-3056871 | 8 | 0.975062445696013 |
| chr14-92702865-92703366 | 5 | 0.990668856236964 |
| chr20-44699376-44699877 | 8 | 0.975294190652175 |
| chr14-7191950-7192451 | 8 | 0.993493681128311 |
| chr4-55532512-55533013 | 8 | 0.973047379354752 |
| chr14-12432220-12432721 | 8 | 0.992320348324544 |
| chr5-88905557-88906058 | 8 | 0.994387825985785 |
| chr19-27695435-27695936 | 5 | 0.97422058603545 |
| chr18-19253372-19253873 | 2 | 0.843206162160751 |
| chr14-41298123-41298624 | 5 | 0.985537109402296 |
| chr5-12242223-12242724 | 8 | 0.824912230973074 |
| chr19-38115771-38116272 | 5 | 0.981129528882881 |
| chr18-11933698-11934199 | 8 | 0.994684881880515 |
| chr20-45143561-45144062 | 8 | 0.984571796538737 |
| chr19-28489289-28489790 | 8 | 0.97063008337741 |
| chr16-34153085-34153586 | 8 | 0.900109500647998 |
| chr17-40569887-40570388 | 8 | 0.973478046389917 |
| chr4-180349731-180350232 | 5 | 0.980009202285355 |
| chr14-43138692-43139193 | 5 | 0.989147528009268 |
| chr2-10338400-10338901 | 8 | 0.913962878669206 |
| chr19-53333599-53334100 | 5 | 0.985118777504461 |

| | | |
|---|---|---|
| chr8-95528787-95529288 | 8 | 0.951512946324413 |
| chr9-94507941-94508442 | 8 | 0.89854610280074 |
| chr2-13086847-13087348 | 8 | 0.993493681128311 |
| chr5-58450707-58451208 | 8 | 0.959530703138499 |
| chr6-6833103-6833604 | 8 | 0.937703732281071 |
| chr20-17903308-17903809 | 8 | 0.979334961107553 |
| chr2-123977504-123978005 | 8 | 0.995046446959392 |
| chr14-52838413-52838914 | 5 | 0.974261664871737 |
| chr1-74041264-74041765 | 3 | 0.99593553545041 |
| chr7-101449866-101450367 | 8 | 0.930696198907069 |
| chr16-62969922-62970423 | 8 | 0.991220996182931 |
| chr9-54231088-54231589 | 8 | 0.993493681128311 |
| chrX-151953995-151954496 | 8 | 0.980716529343873 |
| chr9-104940515-104941016 | 8 | 0.990989682063535 |
| chr14-10075499-10076000 | 8 | 0.993440845668619 |
| chrX-13453257-13453758 | 8 | 0.908676681986607 |
| chr4-42277577-42278078 | 8 | 0.98858897347442 |
| chr9-53894413-53894914 | 8 | 0.994686472038261 |
| chr15-22055161-22055662 | 8 | 0.926179404240985 |
| chr6-5166966-5167467 | 8 | 0.991418793462765 |
| chrX-70983041-70983542 | 8 | 0.900053746913625 |
| chr20-1439546-1440047 | 8 | 0.853690428232109 |
| chr15-75626844-75627345 | 8 | 0.954654716331999 |
| chr2-167499594-167500095 | 8 | 0.959442413634607 |
| chr12-9584431-9584932 | 8 | 0.888807982991845 |
| chr8-39441694-39442195 | 2 | 0.876272758212718 |
| chr16-72072467-72072968 | 8 | 0.995063295136995 |
| chr2-2316596-2317097 | 3 | 0.99593553545041 |
| chr14-27315778-27316279 | 8 | 0.871388842071184 |
| chr18-78748221-78748722 | 4 | 0.888980058364144 |
| chr16-37858248-37858749 | 8 | 0.888274447774135 |
| chrX-49350474-49350975 | 8 | 0.983914308338795 |
| chr13-63829495-63829996 | 8 | 0.878890255473578 |
| chr2-165268231-165268732 | 8 | 0.964294500889175 |
| chr15-71306343-71306844 | 8 | 0.993678840961586 |
| chr3-34144216-34144717 | 8 | 0.99487492345303 |
| chrX-97259874-97260375 | 5 | 0.985027016496224 |
| chrX-102112547-102113048 | 8 | 0.992414517059163 |
| chr5-161879903-161880404 | 8 | 0.944696812490336 |
| chr11-6728650-6729151 | 8 | 0.913028649390816 |
| chr7-91177784-91178285 | 5 | 0.984161557303414 |
| chr12-13130461-13130962 | 8 | 0.96705710813439 |
| chrX-36402845-36403346 | 8 | 0.993493681128311 |

| | | |
|---|---|---|
| chr4-66742923-66743424 | 8 | 0.938110645437481 |
| chr11-56318086-56318587 | 8 | 0.828677230393154 |
| chr19-43666992-43667493 | 5 | 0.978287040002891 |
| chr15-16317210-16317711 | 8 | 0.966124696839106 |
| chr5-50815530-50816031 | 8 | 0.971141868770213 |
| chr1-252386220-252386721 | 8 | 0.977211859698852 |
| chr3-37552591-37553092 | 8 | 0.93688700420938 |
| chr13-42338518-42339019 | 5 | 0.991068897516395 |
| chr20-47920013-47920514 | 8 | 0.821850693826789 |
| chr17-63543014-63543515 | 8 | 0.942093073344099 |
| chr19-25288138-25288639 | 8 | 0.959047119331059 |
| chr17-20321097-20321598 | 5 | 0.983320984437584 |
| chr3-124283967-124284468 | 8 | 0.984910047435939 |
| chr2-106292654-106293155 | 8 | 0.99522038671293 |
| chr1-40811650-40812151 | 8 | 0.978439078571176 |
| chr7-102661105-102661606 | 8 | 0.955950350225572 |
| chr14-64521243-64521744 | 8 | 0.997194974287713 |
| chr6-57425557-57426058 | 8 | 0.964624281116662 |
| chr14-57975779-57976280 | 8 | 0.992153334474173 |
| chr8-77140517-77141018 | 8 | 0.95770107106046 |
| chr9-18884898-18885399 | 8 | 0.931235783232786 |
| chr13-56126188-56126689 | 8 | 0.986849078795314 |
| chr13-37348740-37349241 | 5 | 0.987090639339659 |
| chr6-75990154-75990655 | 8 | 0.961348245899488 |
| chr6-119719854-119720355 | 8 | 0.98846715601519 |
| chr2-221470426-221470927 | 8 | 0.965446823716433 |
| chr7-88162521-88163022 | 8 | 0.987068627814344 |
| chr9-94481939-94482440 | 8 | 0.990811913087671 |
| chr2-119177633-119178134 | 8 | 0.987173088558843 |
| chr2-236383084-236383585 | 8 | 0.993493681128311 |
| chr3-119979530-119980031 | 4 | 0.868719037719811 |
| chr1-81161030-81161531 | 8 | 0.986127247316426 |
| chr15-24020663-24021164 | 4 | 0.827048164844289 |
| chr20-22140504-22141005 | 8 | 0.98688033112396 |
| chr6-85624822-85625323 | 8 | 0.993493681128311 |
| chr16-37695481-37695982 | 8 | 0.993493681128311 |
| chr7-121931873-121932374 | 8 | 0.982595725699459 |
| chr15-17262896-17263397 | 8 | 0.995127293467158 |
| chr17-74393942-74394443 | 5 | 0.990454200367775 |
| chr19-41441877-41442378 | 5 | 0.973057314982056 |
| chr17-31326144-31326645 | 5 | 0.954398065679967 |
| chr7-42675675-42676176 | 8 | 0.979625986385977 |
| chr15-54277190-54277691 | 5 | 0.98465516611801 |

| | | |
|---|---|---|
| chr13-38860384-38860885 | 5 | 0.984935891776628 |
| chr15-77812518-77813019 | 3 | 0.996029300266591 |
| chr2-233247183-233247684 | 8 | 0.81977643708529 |
| chr2-235348702-235349203 | 8 | 0.993493681128311 |
| chr5-10375790-10376291 | 5 | 0.986946731321932 |
| chr5-137627521-137628022 | 8 | 0.913834587762004 |
| chr2-54370234-54370735 | 8 | 0.994889994553721 |
| chr1-58242539-58243040 | 8 | 0.952373780579279 |
| chr18-23728663-23729164 | 8 | 0.865886510167857 |
| chr5-156441963-156442464 | 2 | 0.927277881267851 |
| chr1-13515299-13515800 | 5 | 0.992067013269252 |
| chr14-61654808-61655309 | 5 | 0.962409493321408 |
| chr10-55952504-55953005 | 8 | 0.987000022344518 |
| chr2-245160596-245161097 | 8 | 0.993493681128311 |
| chr14-61645241-61645742 | 5 | 0.988530207496788 |
| chr14-45187034-45187535 | 2 | 0.852374454642221 |
| chr1-89317782-89318283 | 8 | 0.981611226653395 |
| chr1-31502813-31503314 | 8 | 0.896049915543724 |
| chr17-37398164-37398665 | 8 | 0.992739393846238 |
| chr8-29083902-29084403 | 8 | 0.9950032593941 |
| chr6-136442699-136443200 | 8 | 0.951533092775826 |
| chr4-70639644-70640145 | 8 | 0.952540966461495 |
| chrX-25902025-25902526 | 8 | 0.993493681128311 |
| chr2-87981230-87981731 | 8 | 0.814968164236936 |
| chr15-54125862-54126363 | 8 | 0.871099918648584 |
| chr2-38864953-38865454 | 3 | 0.987524316202017 |
| chr1-102827635-102828136 | 8 | 0.940039509741059 |
| chr17-51361239-51361740 | 5 | 0.984870631674003 |
| chr1-220363470-220363971 | 5 | 0.979900030304435 |
| chr4-19153200-19153701 | 8 | 0.993493681128311 |
| chr3-83123446-83123947 | 8 | 0.993493681128311 |
| chr19-45035175-45035676 | 5 | 0.987338949069726 |
| chr10-24703501-24704002 | 8 | 0.974990130823969 |
| chr5-139478709-139479210 | 8 | 0.846939946548442 |
| chr19-56136951-56137452 | 8 | 0.993402784925696 |
| chr3-50047118-50047619 | 8 | 0.910099790151461 |
| chr18-68135227-68135728 | 5 | 0.982447255183763 |
| chr3-99788431-99788932 | 8 | 0.983206667092837 |
| chr14-64550035-64550536 | 4 | 0.898242809749694 |
| chr9-12717693-12718194 | 8 | 0.898801227934997 |
| chr4-47835233-47835734 | 8 | 0.990017507055953 |
| chrX-99212132-99212633 | 3 | 0.99593553545041 |
| chr10-77858017-77858518 | 8 | 0.993493681128311 |

| | | |
|---|---|---|
| chr6-67476144-67476645 | 8 | 0.989509323831931 |
| chr2-93744627-93745128 | 5 | 0.985064656307852 |
| chr16-57686375-57686876 | 8 | 0.994555480194337 |
| chr18-40799808-40800309 | 8 | 0.993493681128311 |
| chr12-27707646-27708147 | 8 | 0.982051618915653 |
| chrX-138854375-138854876 | 4 | 0.867275187012815 |
| chr1-27595318-27595819 | 4 | 0.904273380606711 |
| chr4-181225608-181226109 | 8 | 0.985264049933053 |
| chr5-5746173-5746674 | 8 | 0.995212618969045 |
| chr4-110699226-110699727 | 8 | 0.993493681128311 |
| chr2-129526159-129526660 | 8 | 0.946881150740183 |
| chr4-55235513-55236014 | 4 | 0.886874941589403 |
| chr5-104517756-104518257 | 8 | 0.831558772905392 |
| chrX-59245676-59246177 | 8 | 0.993493681128311 |
| chr11-49031669-49032170 | 8 | 0.995002554249718 |
| chr13-76976664-76977165 | 4 | 0.971283767287289 |
| chrX-18670696-18671197 | 3 | 0.99593553545041 |
| chr2-45133810-45134311 | 5 | 0.985536115023645 |
| chr14-60514634-60515135 | 5 | 0.971361380176588 |
| chr15-100084152-100084653 | 8 | 0.847572339978268 |
| chr7-35325627-35326128 | 8 | 0.982127056583129 |
| chr11-52037362-52037863 | 8 | 0.993428598357844 |
| chr16-50609276-50609777 | 8 | 0.978537792771696 |
| chr18-13066992-13067493 | 3 | 0.995298272094907 |
| chr17-44381052-44381553 | 8 | 0.984498008550107 |
| chr3-48489984-48490485 | 8 | 0.97830363148949 |
| chr2-8133518-8134019 | 4 | 0.957907409277358 |
| chr7-124603503-124604004 | 5 | 0.985667161043358 |
| chr6-125699875-125700376 | 8 | 0.99427810035862 |
| chr14-76934546-76935047 | 5 | 0.983919978840557 |
| chr6-21512042-21512543 | 8 | 0.992839879717005 |
| chr3-66336651-66337152 | 8 | 0.857568745370504 |
| chr4-109996704-109997205 | 2 | 0.89972213069722 |
| chr5-71640283-71640784 | 4 | 0.905522857376162 |
| chr2-98461240-98461741 | 4 | 0.829742453138415 |
| chr7-48750797-48751298 | 8 | 0.949495404251681 |
| chr14-45182345-45182846 | 8 | 0.899600865226899 |
| chr15-31716911-31717412 | 8 | 0.994019196638823 |
| chr18-57359101-57359602 | 8 | 0.994850352785399 |
| chr2-81711178-81711679 | 8 | 0.927194558991201 |
| chr3-27078455-27078956 | 6 | 0.979015124136161 |
| chr2-19036522-19037023 | 4 | 0.945676862135884 |
| chr4-87983909-87984410 | 8 | 0.868705309409554 |

| | | |
|---|---|---|
| chr4-87812210-87812711 | 6 | 0.906569438020934 |
| chr2-67201252-67201753 | 8 | 0.983059849762968 |
| chr3-92649207-92649708 | 4 | 0.967643966289726 |
| chr15-64375936-64376437 | 8 | 0.901832904357458 |
| chr8-32855323-32855824 | 8 | 0.956996933673818 |
| chr3-75325586-75326087 | 8 | 0.993493681128311 |
| chr2-111736584-111737085 | 8 | 0.901853555162252 |
| chr1-58321584-58322085 | 8 | 0.850173229278221 |
| chr14-71287543-71288044 | 2 | 0.865122991855198 |
| chr8-122238341-122238842 | 8 | 0.976439388871263 |
| chr6-98289907-98290408 | 5 | 0.985425634403433 |
| chr8-7155304-7155805 | 8 | 0.985442055789036 |
| chr17-75488615-75489116 | 5 | 0.983919978840557 |
| chr20-47024074-47024575 | 8 | 0.990061627424703 |
| chr1-211266432-211266933 | 8 | 0.900005190520082 |
| chr5-110387174-110387675 | 8 | 0.938817606330209 |
| chr7-49419362-49419863 | 8 | 0.993493681128311 |
| chr4-91216553-91217054 | 8 | 0.989971004584666 |
| chr14-2380355-2380856 | 2 | 0.855369149084919 |
| chr4-32442442-32442943 | 8 | 0.988253744191672 |
| chr1-193958045-193958546 | 8 | 0.876265830366337 |
| chr4-8822215-8822716 | 8 | 0.966280866844418 |
| chr17-48502859-48503360 | 8 | 0.987374390259428 |
| chr8-46065171-46065672 | 8 | 0.934094288519954 |
| chrX-122251431-122251932 | 3 | 0.985868880858728 |
| chr14-22477830-22478331 | 3 | 0.99593553545041 |
| chr4-38972542-38973043 | 8 | 0.993493681128311 |
| chr2-95050554-95051055 | 5 | 0.976383608199297 |
| chr11-17685435-17685936 | 4 | 0.917297664791758 |
| chr15-57120346-57120847 | 3 | 0.99593553545041 |
| chr5-85187429-85187930 | 8 | 0.988123550668737 |
| chr7-117808497-117808998 | 8 | 0.99523487186945 |
| chr4-61110291-61110792 | 8 | 0.986277065143885 |
| chr2-54737853-54738354 | 8 | 0.991845601408615 |
| chr16-14493215-14493716 | 8 | 0.90306266757198 |
| chr2-94888070-94888571 | 4 | 0.870410855525735 |
| chr7-12228045-12228546 | 8 | 0.971531479615758 |
| chr15-80747744-80748245 | 5 | 0.976738611609732 |
| chr10-24877008-24877509 | 5 | 0.994089838771986 |
| chr8-4371888-4372389 | 8 | 0.992154511783741 |
| chr14-69611902-69612403 | 5 | 0.982020350484814 |
| chr7-80007133-80007634 | 5 | 0.994696557097708 |
| chr7-7482503-7483004 | 8 | 0.977923655445152 |

| | | |
|---|---|---|
| chr2-208101223-208101724 | 8 | 0.882754387949985 |
| chr2-28439588-28440089 | 8 | 0.994469081205348 |
| chr7-85556146-85556647 | 8 | 0.964619817962969 |
| chr5-158960206-158960707 | 8 | 0.974687017626075 |
| chr6-30842663-30843164 | 8 | 0.82836459195252 |
| chr1-163354323-163354824 | 5 | 0.935546629549649 |
| chr1-148764745-148765246 | 8 | 0.911671484820117 |
| chr11-12984823-12985324 | 2 | 0.951922306582257 |
| chr2-184915133-184915634 | 8 | 0.992567413184449 |
| chr9-57543640-57544141 | 8 | 0.937176579224337 |
| chr5-15131289-15131790 | 8 | 0.991001243036754 |
| chr10-43509556-43510057 | 8 | 0.989640684394753 |
| chr14-21872916-21873417 | 8 | 0.836852049210625 |
| chr8-33223534-33224035 | 8 | 0.993493681128311 |
| chr3-111743065-111743566 | 8 | 0.945512156661032 |
| chr10-75544097-75544598 | 4 | 0.944881353152785 |
| chr2-98136438-98136939 | 8 | 0.926079607610678 |
| chr9-7323676-7324177 | 4 | 0.832929611700823 |
| chr6-118452986-118453487 | 5 | 0.986531335813044 |
| chr16-59858044-59858545 | 8 | 0.94010860389612 |
| chr4-84374548-84375049 | 3 | 0.860525576321434 |
| chr13-42330415-42330916 | 8 | 0.986097048954795 |
| chr17-48493221-48493722 | 4 | 0.973397568270299 |
| chr14-31687309-31687810 | 8 | 0.993493681128311 |
| chr20-14736337-14736838 | 8 | 0.932978276186207 |
| chr4-118112321-118112822 | 8 | 0.985657104578974 |
| chr11-24258685-24259186 | 8 | 0.833110137504546 |
| chr6-34097478-34097979 | 8 | 0.856836545127713 |
| chr4-165155578-165156079 | 8 | 0.994455204865993 |
| chr2-111354507-111355008 | 8 | 0.964181608179758 |
| chr1-69753884-69754385 | 2 | 0.958462083712826 |
| chr15-1018041-1018542 | 8 | 0.873647420709457 |
| chr16-48744046-48744547 | 4 | 0.966718239024761 |
| chr1-192873586-192874087 | 8 | 0.889458990221628 |
| chr4-19938861-19939362 | 5 | 0.983919978840557 |
| chr14-61978899-61979400 | 8 | 0.880878935488927 |
| chr3-150765657-150766158 | 8 | 0.97019157045169 |
| chr8-107357046-107357547 | 5 | 0.986688459835739 |
| chr2-70133755-70134256 | 3 | 0.995379604745381 |
| chr4-53668267-53668768 | 8 | 0.993493681128311 |
| chr5-116589986-116590487 | 8 | 0.995240488399014 |
| chr7-104407064-104407565 | 2 | 0.915233374235136 |
| chr3-95277672-95278173 | 2 | 0.943461663709379 |

| | | |
|---|---|---|
| chr15-14748734-14749235 | 8 | 0.986190030466495 |
| chr18-48491192-48491693 | 4 | 0.95157869599776 |
| chr1-146966036-146966537 | 8 | 0.946293274187002 |
| chr5-68596771-68597272 | 5 | 0.986094352092618 |
| chr15-2838315-2838816 | 8 | 0.926798914514935 |
| chr9-60054380-60054881 | 8 | 0.983288896847642 |
| chr18-13557903-13558404 | 8 | 0.922618912927321 |
| chr9-111259195-111259696 | 5 | 0.986255077761451 |
| chr15-86644503-86645004 | 8 | 0.914534450957915 |
| chr7-54128507-54129008 | 8 | 0.872693321976086 |
| chr14-89438321-89438822 | 8 | 0.97175159034337 |
| chr13-43471885-43472386 | 8 | 0.904174788143346 |
| chr10-41945909-41946410 | 8 | 0.919569649537039 |
| chr14-44036403-44036904 | 2 | 0.960114335463223 |
| chrX-134316415-134316916 | 5 | 0.983919978840557 |
| chr8-74616971-74617472 | 3 | 0.95939945751509 |
| chr6-17704336-17704837 | 8 | 0.995011222593086 |
| chr5-25028340-25028841 | 8 | 0.991003663692337 |
| chr10-57333418-57333919 | 8 | 0.930110003035003 |
| chr10-60210358-60210859 | 8 | 0.926566068899991 |
| chr3-50184320-50184821 | 5 | 0.991124547806326 |
| chr2-44834835-44835336 | 8 | 0.87716053875135 |
| chr2-94745465-94745966 | 5 | 0.985181655289304 |
| chr5-97746003-97746504 | 8 | 0.984442060186403 |
| chr2-13058020-13058521 | 8 | 0.994660266484063 |
| chr1-136649039-136649540 | 8 | 0.970786800279242 |
| chr6-129172289-129172790 | 8 | 0.994705647729275 |
| chr20-21815381-21815882 | 5 | 0.985111753112849 |
| chrX-78443272-78443773 | 8 | 0.956466925942202 |
| chr4-11244768-11245269 | 8 | 0.952018073146877 |
| chr2-204554677-204555178 | 8 | 0.932147679474301 |
| chr8-42087566-42088067 | 8 | 0.913083406073455 |
| chr15-82842142-82842643 | 8 | 0.98472036771142 |
| chr17-49752671-49753172 | 4 | 0.913806884811101 |
| chr6-63274338-63274839 | 8 | 0.977620798351087 |
| chr4-112927250-112927751 | 2 | 0.91908739508431 |
| chr7-1399086-1399587 | 8 | 0.961318176183942 |
| chr20-39860680-39861181 | 8 | 0.975564131773831 |
| chr3-59463287-59463788 | 8 | 0.988264729165997 |
| chr11-4462089-4462590 | 8 | 0.887068803250546 |
| chr4-151605480-151605981 | 8 | 0.968600995902348 |
| chr6-68492565-68493066 | 5 | 0.972685723814447 |
| chr7-2075703-2076204 | 8 | 0.948215872588302 |

| | | |
|---|---|---|
| chrX-8182519-8183020 | 4 | 0.829741128451803 |
| chr6-89926791-89927292 | 8 | 0.994954955984735 |
| chr2-142320799-142321300 | 8 | 0.856575504880533 |
| chr13-80719406-80719907 | 8 | 0.99031679132149 |
| chr2-32209782-32210283 | 8 | 0.979351794482648 |
| chr6-3473117-3473618 | 8 | 0.993493681128311 |
| chrX-74693675-74694176 | 3 | 0.996061400506539 |
| chr12-42734161-42734662 | 8 | 0.963765836077746 |
| chr18-82578290-82578791 | 8 | 0.936087005468634 |
| chr5-69318163-69318664 | 8 | 0.980759758109749 |
| chr8-85586425-85586926 | 8 | 0.992069226996886 |
| chr14-84734711-84735212 | 8 | 0.944204 |
| chr18-6920710-6921211 | 8 | 0.993493681128311 |
| chr3-67540956-67541457 | 8 | 0.993934417537384 |
| chr8-47580396-47580897 | 8 | 0.994256798529914 |
| chr5-101084568-101085069 | 8 | 0.995000872861776 |
| chr3-150061664-150062165 | 8 | 0.989323537454916 |
| chr2-228931870-228932371 | 8 | 0.930806498472479 |
| chr8-56301803-56302304 | 5 | 0.991118761683287 |
| chr2-109012835-109013336 | 2 | 0.902805471214698 |
| chr3-21350978-21351479 | 5 | 0.980281812288097 |
| chr6-19828801-19829302 | 8 | 0.832414354589221 |
| chr13-70796430-70796931 | 8 | 0.977032844394893 |
| chr16-25058242-25058743 | 8 | 0.96650323623828 |
| chr1-106387580-106388081 | 8 | 0.952502711317134 |
| chr14-9692005-9692506 | 8 | 0.993493681128311 |
| chr11-49422444-49422945 | 8 | 0.993493681128311 |
| chr3-131078401-131078902 | 8 | 0.988705825952497 |
| chr15-76112357-76112858 | 8 | 0.997252359503695 |
| chr4-66745395-66745896 | 8 | 0.917444469758073 |
| chr14-65858791-65859292 | 4 | 0.965165156003239 |
| chr6-84789775-84790276 | 2 | 0.880086051815961 |
| chr14-18587933-18588434 | 8 | 0.985565638813207 |
| chr13-36523699-36524200 | 8 | 0.967926449515497 |
| chr8-76602208-76602709 | 4 | 0.931789167537322 |
| chrX-25850962-25851463 | 8 | 0.994461885187643 |
| chr14-71204016-71204517 | 5 | 0.98114427944303 |
| chr10-74728380-74728881 | 8 | 0.993493681128311 |
| chr14-71116919-71117420 | 8 | 0.815826128304042 |
| chrX-15009815-15010316 | 8 | 0.994653459306499 |
| chr7-11083029-11083530 | 8 | 0.990195116613192 |
| chr4-41256521-41257022 | 3 | 0.997112736324796 |
| chr7-125038635-125039136 | 8 | 0.926931579826391 |

| | | |
|---|---|---|
| chr4-102452975-102453476 | 3 | 0.998181304234569 |
| chr16-80100346-80100847 | 8 | 0.993493681128311 |
| chr7-97569497-97569998 | 8 | 0.993493681128311 |
| chr18-35412617-35413118 | 8 | 0.948796909957276 |
| chr5-66933827-66934328 | 4 | 0.885857988765272 |
| chr12-40405221-40405722 | 4 | 0.828167897014567 |
| chr9-50833370-50833871 | 5 | 0.980206392324627 |
| chr11-187004-187505 | 8 | 0.928359958508571 |
| chr5-112780703-112781204 | 8 | 0.964104100319628 |
| chrX-129512973-129513474 | 8 | 0.933670560913156 |
| chr8-99498355-99498856 | 5 | 0.993924781044932 |
| chrX-15892691-15893192 | 3 | 0.978998184176029 |
| chr7-44832004-44832505 | 8 | 0.941142725655823 |
| chr12-317136-317637 | 4 | 0.931446626609316 |
| chr4-67183138-67183639 | 8 | 0.990915658239245 |
| chr5-92570342-92570843 | 8 | 0.992025673438624 |
| chr15-79525035-79525536 | 8 | 0.977844301905454 |
| chr17-6649324-6649825 | 8 | 0.890811274505891 |
| chr4-174532017-174532518 | 4 | 0.829656375901026 |
| chr6-52172972-52173473 | 8 | 0.825347388268019 |
| chr20-11589391-11589892 | 8 | 0.937779879640291 |
| chr4-161166576-161167077 | 8 | 0.995105511407896 |
| chrX-49879395-49879896 | 8 | 0.958220308456442 |
| chr1-254191965-254192466 | 8 | 0.916810025862594 |
| chr2-53330137-53330638 | 8 | 0.992465242597502 |
| chr2-144931372-144931873 | 8 | 0.970328583588345 |
| chr4-12591378-12591879 | 5 | 0.862526326007828 |
| chr3-6937473-6937974 | 8 | 0.832304734064022 |
| chr5-57585109-57585610 | 8 | 0.966748130862215 |
| chrX-12986465-12986966 | 8 | 0.936246538915876 |
| chr4-27200418-27200919 | 3 | 0.996895178774349 |
| chr14-23460255-23460756 | 4 | 0.935790167802269 |
| chr9-45856595-45857096 | 8 | 0.995176168443575 |
| chr9-40427424-40427925 | 8 | 0.986662462586028 |
| chr1-41395998-41396499 | 8 | 0.990369862241751 |
| chr9-20363480-20363981 | 3 | 0.99593553545041 |
| chr7-72620655-72621156 | 8 | 0.992388369582457 |
| chr6-11085351-11085852 | 8 | 0.965018226016277 |
| chr20-49197756-49198257 | 5 | 0.994036799479741 |
| chr14-82095984-82096485 | 5 | 0.996334377101778 |
| chr10-65685813-65686314 | 8 | 0.859329818718958 |
| chr2-4745764-4746265 | 8 | 0.990668944505951 |
| chr13-854475-854976 | 8 | 0.917333757472013 |

| | | |
|---|---|---|
| chr2-32838098-32838599 | 8 | 0.990488985684703 |
| chr15-95767319-95767820 | 8 | 0.966416207803054 |
| chr2-64918021-64918522 | 8 | 0.964337864158648 |
| chr11-78129673-78130174 | 8 | 0.927373876405553 |
| chr15-61304826-61305327 | 8 | 0.987720225934866 |
| chr14-91289061-91289562 | 5 | 0.971086875741378 |
| chr15-33196529-33197030 | 8 | 0.993493681128311 |
| chr4-50471992-50472493 | 5 | 0.983919978840557 |
| chr6-83077530-83078031 | 8 | 0.992238199832554 |
| chr2-161840524-161841025 | 8 | 0.993493681128311 |
| chr13-74734197-74734698 | 2 | 0.901906580264639 |
| chr6-3303159-3303660 | 4 | 0.844396677444958 |
| chr1-253450328-253450829 | 8 | 0.903710667570106 |
| chr9-19643181-19643682 | 8 | 0.838048410867459 |
| chr13-33759090-33759591 | 8 | 0.984189840523678 |
| chr4-110886976-110887477 | 8 | 0.993493681128311 |
| chr5-127803494-127803995 | 5 | 0.842862379551058 |
| chr10-28703776-28704277 | 8 | 0.966195035387742 |
| chr2-33150351-33150852 | 8 | 0.994112077364357 |
| chr5-99708562-99709063 | 8 | 0.827333596340005 |
| chrX-33299504-33300005 | 3 | 0.99593553545041 |
| chr4-90040973-90041474 | 4 | 0.972757748504182 |
| chr13-7214507-7215008 | 8 | 0.926714414443257 |
| chr20-47127461-47127962 | 8 | 0.99509502316003 |
| chr2-105705786-105706287 | 8 | 0.847281771172701 |
| chr2-183120178-183120679 | 8 | 0.847077878261102 |
| chr2-202043122-202043623 | 6 | 0.979691304787456 |
| chr2-194070919-194071420 | 8 | 0.947957656686691 |
| chr8-42981748-42982249 | 8 | 0.99583364439854 |
| chr11-62474426-62474927 | 8 | 0.976831911213207 |
| chr11-60826096-60826597 | 8 | 0.83750596423484 |
| chr4-24125839-24126340 | 8 | 0.957311502158411 |
| chr2-79514611-79515112 | 8 | 0.990508358231202 |
| chr4-15120602-15121103 | 8 | 0.92891840675744 |
| chr9-6761384-6761885 | 8 | 0.970168433361204 |
| chr3-44418130-44418631 | 5 | 0.988072334551933 |
| chr1-67258006-67258507 | 8 | 0.993786259962096 |
| chr10-74106366-74106867 | 8 | 0.944006724641884 |
| chr15-37357358-37357859 | 8 | 0.862450606414589 |
| chr1-185461206-185461707 | 8 | 0.939638899987449 |
| chr19-26508021-26508522 | 5 | 0.982574700531149 |
| chr4-162964439-162964940 | 3 | 0.954149950148901 |
| chr11-1654495-1654996 | 8 | 0.993493681128311 |

| | | |
|---|---|---|
| chr7-81052378-81052879 | 8 | 0.990494478275356 |
| chr4-7046347-7046848 | 8 | 0.818698874165614 |
| chr8-82388572-82389073 | 5 | 0.988469790427456 |
| chr1-219147589-219148090 | 8 | 0.90062413186884 |
| chr6-123323291-123323792 | 3 | 0.952742333280438 |
| chr3-102007071-102007572 | 8 | 0.993493681128311 |
| chr18-66932732-66933233 | 8 | 0.89504774343386 |
| chr4-161524447-161524948 | 4 | 0.936115534319124 |
| chr3-159902943-159903444 | 2 | 0.914248705536848 |
| chr4-133744321-133744822 | 8 | 0.824136725801869 |
| chr4-47566436-47566937 | 8 | 0.959231533192705 |
| chr5-135548128-135548629 | 8 | 0.991063986121632 |
| chr13-26390976-26391477 | 3 | 0.99593553545041 |
| chr18-3395298-3395799 | 8 | 0.864716372853176 |
| chr3-30349880-30350381 | 5 | 0.894307594161031 |
| chr10-46420846-46421347 | 8 | 0.996890047346617 |
| chr3-94931089-94931590 | 8 | 0.993217426520818 |
| chr2-6295531-6296032 | 8 | 0.890461328859653 |
| chr9-6734543-6735044 | 8 | 0.989332628006244 |
| chr1-229690657-229691158 | 8 | 0.911373147520247 |
| chr6-76162057-76162558 | 8 | 0.993493681128311 |
| chr5-33039529-33040030 | 8 | 0.984749638799678 |
| chr18-7794557-7795058 | 8 | 0.973667605147215 |
| chr16-16475182-16475683 | 8 | 0.993493681128311 |
| chr4-128361706-128362207 | 3 | 0.996843349345123 |
| chr8-113087492-113087993 | 5 | 0.985565489243383 |
| chr9-27408603-27409104 | 8 | 0.995042027523625 |
| chr2-190701431-190701932 | 8 | 0.86772718370182 |
| chr3-77778263-77778764 | 8 | 0.983933172432389 |
| chr1-46664022-46664523 | 8 | 0.993493681128311 |
| chr18-1973501-1974002 | 8 | 0.910171597345958 |
| chr3-52989842-52990343 | 8 | 0.980882838042977 |
| chr12-23213493-23213994 | 8 | 0.994385440416784 |
| chr7-104182563-104183064 | 6 | 0.991206925481043 |
| chr14-99290600-99291101 | 5 | 0.966194625520766 |
| chr6-48658199-48658700 | 8 | 0.993493681128311 |
| chr14-34597314-34597815 | 8 | 0.951674969344249 |
| chr15-68767623-68768124 | 8 | 0.971862162721119 |
| chr8-23335714-23336215 | 8 | 0.991946644810532 |
| chr1-241943256-241943757 | 8 | 0.993493681128311 |
| chr1-74139466-74139967 | 4 | 0.924882866569429 |
| chr4-57548710-57549211 | 8 | 0.993493681128311 |
| chr15-93357103-93357604 | 8 | 0.993008335471676 |

| | | |
|---|---|---|
| chr3-95068598-95069099 | 8 | 0.926839496155669 |
| chr15-71460285-71460786 | 8 | 0.946286371380171 |
| chr6-110394058-110394559 | 8 | 0.993493681128311 |
| chr18-66959755-66960256 | 5 | 0.985208053805335 |
| chr17-23600738-23601239 | 8 | 0.980033244070676 |
| chr8-82042842-82043343 | 3 | 0.996665865406983 |
| chr17-12961491-12961992 | 8 | 0.923513505213219 |
| chr15-41793652-41794153 | 8 | 0.993493681128311 |
| chr11-77431937-77432438 | 3 | 0.990082461912823 |
| chr13-39062412-39062913 | 8 | 0.991991558420429 |
| chrX-112232112-112232613 | 8 | 0.993493681128311 |
| chr17-18450676-18451177 | 8 | 0.941534263404603 |
| chr11-4679650-4680151 | 8 | 0.983743174517184 |
| chr13-102670859-102671360 | 8 | 0.832292657818371 |
| chr2-105883675-105884176 | 8 | 0.886541808202571 |
| chr7-120310406-120310907 | 8 | 0.981871000295872 |
| chr2-245572871-245573372 | 2 | 0.954840872352381 |
| chr8-123411203-123411704 | 8 | 0.993340587462737 |
| chr15-84552148-84552649 | 8 | 0.828831249140431 |
| chr3-27696983-27697484 | 8 | 0.988022034699196 |
| chr10-72751126-72751627 | 4 | 0.944459207280391 |
| chr9-31381621-31382122 | 8 | 0.979184323222099 |
| chr13-39924885-39925386 | 8 | 0.952338746064164 |
| chr19-42947825-42948326 | 5 | 0.983919978840557 |
| chr16-7827401-7827902 | 8 | 0.825602209392534 |
| chr5-112206119-112206620 | 4 | 0.972560122325923 |
| chr1-260435378-260435879 | 8 | 0.993493681128311 |
| chr1-5743423-5743924 | 8 | 0.942695908624858 |
| chr5-4000531-4001032 | 8 | 0.943751073557159 |
| chr4-47762000-47762501 | 8 | 0.995247515291731 |
| chr5-103543366-103543867 | 8 | 0.967550640560614 |
| chr6-19434164-19434665 | 8 | 0.993493681128311 |
| chr3-35105705-35106206 | 8 | 0.940092276738566 |
| chr17-39290503-39291004 | 8 | 0.81693680706505 |
| chr8-122998961-122999462 | 8 | 0.968351944485611 |
| chr1-253217924-253218425 | 8 | 0.994640666776155 |
| chr18-16995103-16995604 | 4 | 0.957110274576756 |
| chr19-42037699-42038200 | 8 | 0.991108949838297 |
| chr3-31850166-31850667 | 8 | 0.993493681128311 |
| chr14-21419194-21419695 | 8 | 0.97814843312854 |
| chr17-25289867-25290368 | 3 | 0.996226927758663 |
| chr16-75263913-75264414 | 8 | 0.946249068362909 |
| chr7-86590279-86590780 | 2 | 0.871667943779517 |

| | | |
|---|---|---|
| chr6-58865051-58865552 | 8 | 0.955653207196452 |
| chr19-21671827-21672328 | 8 | 0.987133424975876 |
| chr17-47536077-47536578 | 8 | 0.868485532671574 |
| chr17-40484280-40484781 | 8 | 0.937692998597465 |
| chr13-3152585-3153086 | 3 | 0.874767276257279 |
| chr8-96655362-96655863 | 8 | 0.934964908242977 |
| chr3-46725882-46726383 | 5 | 0.988532015134439 |
| chr4-51635573-51636074 | 8 | 0.857178814300884 |
| chr17-26676598-26677099 | 5 | 0.964062605437354 |
| chr16-5679533-5680034 | 8 | 0.884539299061455 |
| chr7-123940159-123940660 | 8 | 0.868097610552928 |
| chr4-129940314-129940815 | 8 | 0.913468607271288 |
| chr16-47941426-47941927 | 8 | 0.941884033509124 |
| chr6-113978911-113979412 | 8 | 0.993493681128311 |
| chr15-69300499-69301000 | 2 | 0.878589639841494 |
| chr13-94315664-94316165 | 8 | 0.936239018063082 |
| chr2-6532574-6533075 | 8 | 0.926215211303593 |
| chr9-63607171-63607672 | 5 | 0.95048107158427 |
| chr11-56250025-56250526 | 8 | 0.851210443631903 |
| chr17-74349693-74350194 | 8 | 0.865410143975951 |
| chr14-78584388-78584889 | 8 | 0.987856481678271 |
| chr3-127957496-127957997 | 4 | 0.88268905034566 |
| chr13-203180-203681 | 3 | 0.99656058915697 |
| chr5-4069031-4069532 | 3 | 0.988063292278031 |
| chr3-97188618-97189119 | 8 | 0.993493681128311 |
| chr13-74764454-74764955 | 8 | 0.994714248093522 |
| chr20-12696510-12697011 | 5 | 0.985732172121559 |
| chr1-39809374-39809875 | 6 | 0.990347253549817 |
| chr15-32124294-32124795 | 5 | 0.987065864442063 |
| chr5-27069047-27069548 | 8 | 0.816741754336205 |
| chr10-100524983-100525484 | 8 | 0.965106054997324 |
| chr10-27596797-27597298 | 5 | 0.903200739901558 |
| chr10-27523237-27523738 | 5 | 0.972410452638997 |
| chr4-71053774-71054275 | 5 | 0.964538196288622 |
| chr1-33916882-33917383 | 5 | 0.986002714229935 |
| chr4-94892227-94892728 | 8 | 0.921986811994114 |
| chr6-3603386-3603887 | 8 | 0.991372144745136 |
| chr4-150558097-150558598 | 3 | 0.93449167270807 |
| chr8-14017997-14018498 | 8 | 0.971566617007126 |
| chr7-6505788-6506289 | 8 | 0.990724998188507 |
| chr14-57307712-57308213 | 3 | 0.965369442697368 |
| chr16-66159856-66160357 | 8 | 0.995241525387029 |
| chr5-93382233-93382734 | 8 | 0.882507307705961 |

| | | |
|---|---|---|
| chr11-36696579-36697080 | 8 | 0.926426718409396 |
| chr6-19296343-19296844 | 8 | 0.976062339885255 |
| chr3-31211163-31211664 | 8 | 0.963130886094044 |
| chr9-55659503-55660004 | 4 | 0.964026234170608 |
| chrX-115041517-115042018 | 3 | 0.99593553545041 |
| chr3-30712312-30712813 | 3 | 0.996064766489511 |
| chr15-41424802-41425303 | 8 | 0.976592832887736 |
| chr15-13176737-13177238 | 8 | 0.990582868599389 |
| chr17-14711513-14712014 | 3 | 0.99593553545041 |
| chr18-22075484-22075985 | 4 | 0.935574501296419 |
| chr2-133035350-133035851 | 2 | 0.959661332352553 |
| chr19-42011653-42012154 | 5 | 0.983919978840557 |
| chr5-27110415-27110916 | 2 | 0.822931404187049 |
| chr4-14612344-14612845 | 8 | 0.993493681128311 |
| chr5-34179038-34179539 | 8 | 0.861305017702982 |
| chrY-9170888-9171389 | 3 | 0.99593553545041 |
| chr1-161252037-161252538 | 8 | 0.942435927401623 |
| chr17-22417666-22418167 | 8 | 0.993010499339294 |
| chr16-33346184-33346685 | 8 | 0.993493681128311 |
| chr6-120640550-120641051 | 8 | 0.993493681128311 |
| chr6-77316135-77316636 | 8 | 0.936560067114332 |
| chr3-95335985-95336486 | 8 | 0.994889513597133 |
| chr18-27056969-27057470 | 8 | 0.994713848557932 |
| chr2-3979519-3980020 | 8 | 0.977192149165623 |
| chr1-199804792-199805293 | 8 | 0.993791225228663 |
| chr1-16688707-16689208 | 8 | 0.869538972492352 |
| chr5-26876168-26876669 | 8 | 0.984732501471004 |
| chr2-111346857-111347358 | 8 | 0.85227649861588 |
| chr5-148758312-148758813 | 8 | 0.943119527065313 |
| chr14-5038442-5038943 | 4 | 0.943311673867224 |
| chr10-5459625-5460126 | 8 | 0.992793077516239 |
| chr10-23647824-23648325 | 8 | 0.994533524616984 |
| chr7-68071108-68071609 | 8 | 0.995875824253337 |
| chr8-14485626-14486127 | 8 | 0.943336379023998 |
| chrX-15952696-15953197 | 8 | 0.917854452083101 |
| chr6-21607922-21608423 | 5 | 0.989508418409783 |
| chr1-191105700-191106201 | 8 | 0.945397690836677 |
| chr14-86625431-86625932 | 5 | 0.981983095931775 |
| chr10-2081746-2082247 | 8 | 0.927923509460542 |
| chr6-76428642-76429143 | 2 | 0.841624835098517 |
| chr11-4382179-4382680 | 2 | 0.957201542666599 |
| chrX-64293288-64293789 | 8 | 0.949399851433396 |
| chr15-30321909-30322410 | 8 | 0.985632568654935 |

| | | |
|---|---|---|
| chr10-7084913-7085414 | 8 | 0.982851961062287 |
| chr2-20131381-20131882 | 5 | 0.967099490033941 |
| chr15-101364933-101365434 | 8 | 0.921750000014368 |
| chr6-44972035-44972536 | 8 | 0.994292696360619 |
| chr16-49376924-49377425 | 8 | 0.867252464649647 |
| chr5-53808837-53809338 | 8 | 0.985233390194791 |
| chr8-31598703-31599204 | 8 | 0.990930507617173 |
| chr4-92928000-92928501 | 8 | 0.990176419898784 |
| chr6-24681314-24681815 | 8 | 0.937138458830754 |
| chr15-63869296-63869797 | 8 | 0.993493681128311 |
| chr2-79464614-79465115 | 8 | 0.932200731292322 |
| chr15-72574455-72574956 | 8 | 0.993493681128311 |
| chr7-39738122-39738623 | 3 | 0.99593553545041 |
| chr1-62821469-62821970 | 6 | 0.991206925481043 |
| chr20-2725013-2725514 | 5 | 0.980450000497173 |
| chr7-110652623-110653124 | 4 | 0.973916679972112 |
| chr17-63379318-63379819 | 8 | 0.973089654084368 |
| chr18-82275311-82275812 | 8 | 0.98664863988755 |
| chr5-13395947-13396448 | 3 | 0.99593553545041 |
| chr1-75704507-75705008 | 8 | 0.940007612699913 |
| chr18-40908667-40909168 | 8 | 0.993493681128311 |
| chr19-46033745-46034246 | 8 | 0.990964107899172 |
| chr8-17233020-17233521 | 3 | 0.99593553545041 |
| chr11-50322432-50322933 | 8 | 0.913840323541042 |
| chr3-64585159-64585660 | 8 | 0.994614055266958 |
| chr3-70763851-70764352 | 5 | 0.974312514724936 |
| chr8-56128910-56129411 | 5 | 0.979124263372752 |
| chr6-4697542-4698043 | 6 | 0.991664601547014 |
| chr16-32400774-32401275 | 8 | 0.854900417033813 |
| chr7-59449992-59450493 | 8 | 0.937238302713537 |
| chr6-68941129-68941630 | 2 | 0.946747325365878 |
| chr6-93690436-93690937 | 8 | 0.993493681128311 |
| chr10-13139093-13139594 | 8 | 0.874806942079962 |
| chr3-47180498-47180999 | 8 | 0.987593802007577 |
| chr17-19491357-19491858 | 5 | 0.982517593585508 |
| chr1-102208979-102209480 | 8 | 0.957468682540416 |
| chr8-34221435-34221936 | 3 | 0.996730337445286 |
| chr19-46208027-46208528 | 8 | 0.920945100879956 |
| chr3-45047520-45048021 | 8 | 0.988559624220086 |
| chr13-22799151-22799652 | 8 | 0.875634735166886 |
| chr5-111753029-111753530 | 8 | 0.844993342836555 |
| chr18-74581109-74581610 | 8 | 0.861840573716959 |
| chr14-74806449-74806950 | 8 | 0.920169185201239 |

| | | |
|---|---|---|
| chr17-46470155-46470656 | 8 | 0.988874033909871 |
| chr10-60232466-60232967 | 8 | 0.994285598578067 |
| chr2-158867062-158867563 | 8 | 0.966616646389574 |
| chr1-177216854-177217355 | 8 | 0.927163135002042 |
| chr12-16901288-16901789 | 8 | 0.893283925983994 |
| chr2-243511658-243512159 | 8 | 0.856561200033793 |
| chr12-28061506-28062007 | 5 | 0.977048982331363 |
| chr6-65967375-65967876 | 8 | 0.815924933779466 |
| chr10-48955677-48956178 | 8 | 0.968011054762307 |
| chr2-19067285-19067786 | 5 | 0.983919978840557 |
| chr2-49027376-49027877 | 8 | 0.983610270489322 |
| chr20-40252268-40252769 | 8 | 0.928778668011941 |
| chr2-169805879-169806380 | 8 | 0.994921117611929 |
| chr17-68776556-68777057 | 8 | 0.990424421011169 |
| chrX-111399027-111399528 | 8 | 0.990710659724297 |
| chr12-21123613-21124114 | 8 | 0.993493681128311 |
| chr8-13762999-13763500 | 8 | 0.903367464113289 |
| chr6-45090418-45090919 | 8 | 0.99499712583495 |
| chr4-116732534-116733035 | 4 | 0.966049038807111 |
| chr13-92852831-92853332 | 8 | 0.974282592792876 |
| chr8-111911734-111912235 | 2 | 0.952487626731345 |
| chr6-66203772-66204273 | 6 | 0.991206925481043 |
| chr13-76443555-76444056 | 3 | 0.942063220284421 |
| chr17-3548451-3548952 | 3 | 0.992931624382266 |
| chr1-126586961-126587462 | 6 | 0.990037978957712 |
| chr10-43049843-43050344 | 2 | 0.924851188664817 |
| chr13-48378681-48379182 | 2 | 0.868346390423502 |
| chr16-60866515-60867016 | 8 | 0.856421267156695 |
| chr13-70522421-70522922 | 8 | 0.995200677649412 |
| chr2-36630077-36630578 | 8 | 0.99164669177565 |
| chr11-66472209-66472710 | 8 | 0.905895094647524 |
| chr12-21812107-21812608 | 3 | 0.971998334737636 |
| chr2-20368363-20368864 | 5 | 0.921257914433029 |
| chr2-41331782-41332283 | 8 | 0.904703990376865 |
| chr1-57921773-57922274 | 2 | 0.849100954591544 |
| chr18-67515893-67516394 | 5 | 0.992925694285931 |
| chr8-7370201-7370702 | 8 | 0.992183863849833 |
| chr3-14988148-14988649 | 8 | 0.991643478390544 |
| chr1-60600290-60600791 | 4 | 0.935301312549095 |
| chr14-85822579-85823080 | 8 | 0.993493681128311 |
| chr13-29621834-29622335 | 8 | 0.976629679087542 |
| chr20-34000534-34001035 | 8 | 0.912228156713647 |
| chr7-42789287-42789788 | 8 | 0.845988916815449 |

| | | |
|---|---|---|
| chr1-188357568-188358069 | 8 | 0.953427841767224 |
| chr13-64583958-64584459 | 5 | 0.985216081450784 |
| chr9-56641096-56641597 | 5 | 0.988644998622669 |
| chr3-165747921-165748422 | 8 | 0.989741089675806 |
| chr5-70940072-70940573 | 2 | 0.956456599130832 |
| chr1-125098783-125099284 | 5 | 0.9195625864095 |
| chrX-5034163-5034664 | 4 | 0.962368008893024 |
| chr13-56632628-56633129 | 3 | 0.99593553545041 |
| chr8-30226919-30227420 | 8 | 0.923496175389641 |
| chr2-86583605-86584106 | 2 | 0.951156449516551 |
| chrX-117072812-117073313 | 8 | 0.844955044618237 |
| chr1-26866138-26866639 | 8 | 0.983236551302289 |
| chr7-72668012-72668513 | 8 | 0.941279161496484 |
| chr5-91193344-91193845 | 8 | 0.995232223646634 |
| chr4-141942105-141942606 | 8 | 0.971280163066778 |
| chr4-101285134-101285635 | 8 | 0.978277650337999 |
| chr17-60243983-60244484 | 3 | 0.996562808992728 |
| chr14-30593496-30593997 | 8 | 0.993493681128311 |
| chr1-130480256-130480757 | 2 | 0.938648137587212 |
| chr15-47713336-47713837 | 8 | 0.993120854069974 |
| chr16-63192304-63192805 | 8 | 0.916470574658108 |
| chr5-38717316-38717817 | 5 | 0.900461408856839 |
| chr18-7611423-7611924 | 3 | 0.996745076548304 |
| chr11-44452535-44453036 | 8 | 0.985581012231639 |
| chr13-16192199-16192700 | 8 | 0.980761674789145 |
| chr2-86800863-86801364 | 8 | 0.857824208978751 |
| chr17-68531272-68531773 | 8 | 0.878527333887553 |
| chr8-70201419-70201920 | 8 | 0.995230067838608 |
| chr2-148888344-148888845 | 2 | 0.961441428823662 |
| chr14-41138118-41138619 | 8 | 0.87674386549794 |
| chr5-111777984-111778485 | 8 | 0.843087838955329 |
| chr8-12040470-12040971 | 5 | 0.981093426794824 |
| chr9-34403242-34403743 | 3 | 0.99572440815138 |
| chr1-176346888-176347389 | 8 | 0.87626321396014 |
| chr6-13511086-13511587 | 8 | 0.985308074159424 |
| chr14-12930339-12930840 | 8 | 0.993493681128311 |
| chr14-72380620-72381121 | 5 | 0.981474289435069 |
| chr18-55433778-55434279 | 8 | 0.944797522974101 |
| chr17-22633335-22633836 | 5 | 0.985080946212701 |
| chr17-29777378-29777879 | 8 | 0.989663707659493 |
| chr3-11740943-11741444 | 8 | 0.917472657931053 |
| chr3-83647257-83647758 | 2 | 0.873798367766238 |
| chr18-56968407-56968908 | 4 | 0.941425438431482 |

| | | |
|---|---|---|
| chr4-52818367-52818868 | 4 | 0.876527709235167 |
| chr7-42675168-42675669 | 8 | 0.97525004605797 |
| chr5-26434679-26435180 | 8 | 0.990446173623535 |
| chr13-47305701-47306202 | 8 | 0.994160210437686 |
| chr7-21601038-21601539 | 8 | 0.966253510767562 |
| chr8-60305469-60305970 | 5 | 0.988916861912264 |
| chr7-57303498-57303999 | 2 | 0.854436180981356 |
| chr2-82469689-82470190 | 8 | 0.956743214522201 |
| chr14-31973446-31973947 | 4 | 0.817565319134616 |
| chr1-61382172-61382673 | 8 | 0.95166804768171 |
| chr14-62523288-62523789 | 8 | 0.97068142421688 |
| chr14-3296298-3296799 | 8 | 0.916373846199901 |
| chrX-32744959-32745460 | 8 | 0.993493681128311 |
| chr3-39818229-39818730 | 8 | 0.994988438461864 |
| chr13-848912-849413 | 8 | 0.944798911443964 |
| chr4-4728187-4728688 | 2 | 0.934320103118172 |
| chr18-11082437-11082938 | 8 | 0.993493681128311 |
| chr11-59339973-59340474 | 8 | 0.994756617571875 |
| chr6-47444861-47445362 | 8 | 0.945075416658885 |
| chr6-35502541-35503042 | 8 | 0.995054845859395 |
| chrX-141207498-141207999 | 8 | 0.950056048121141 |
| chrX-12440732-12441233 | 2 | 0.86197929073262 |
| chr3-138038372-138038873 | 8 | 0.994885566122561 |
| chr20-16701244-16701745 | 8 | 0.868693031709106 |
| chr13-49983322-49983823 | 8 | 0.982719497430057 |
| chr13-29475839-29476340 | 5 | 0.980584433121916 |
| chr5-22842512-22843013 | 8 | 0.969030230052208 |
| chr5-47008369-47008870 | 5 | 0.938842485763551 |
| chr5-110285099-110285600 | 8 | 0.997359579080816 |
| chr9-56739376-56739877 | 8 | 0.946729280909662 |
| chr8-85587119-85587620 | 8 | 0.971892878328289 |
| chr2-68793056-68793557 | 8 | 0.995249043812321 |
| chr2-235949391-235949892 | 8 | 0.954148960081241 |
| chr1-258566659-258567160 | 8 | 0.987848220898863 |
| chr15-38115788-38116289 | 8 | 0.994309456151706 |
| chr2-114082500-114083001 | 8 | 0.956687187458537 |
| chr3-6588723-6589224 | 8 | 0.961622765683111 |
| chr13-32790980-32791481 | 8 | 0.993493681128311 |
| chr10-8280021-8280522 | 8 | 0.974227617868351 |
| chr1-109405477-109405978 | 8 | 0.995033197447012 |
| chr10-95620619-95621120 | 4 | 0.969623766795383 |
| chr20-12624571-12625072 | 8 | 0.994153772241156 |
| chr8-20803087-20803588 | 8 | 0.929807521675946 |

| | | |
|---|---|---|
| chr9-107524295-107524796 | 8 | 0.992455499829394 |
| chr14-12042463-12042964 | 8 | 0.97295948570109 |
| chr5-119464172-119464673 | 8 | 0.983852150128998 |
| chr2-29415918-29416419 | 8 | 0.990531113014523 |
| chr1-54859290-54859791 | 8 | 0.890720571762608 |
| chr10-40347174-40347675 | 6 | 0.972457248308294 |
| chr5-47031430-47031931 | 8 | 0.994901744095275 |
| chr16-41163106-41163607 | 8 | 0.993493681128311 |
| chr11-52902132-52902633 | 8 | 0.993493681128311 |
| chr9-55519195-55519696 | 5 | 0.931132576419713 |
| chr5-96596393-96596894 | 8 | 0.994868944140708 |
| chr3-114610685-114611186 | 8 | 0.893859104311373 |
| chr5-86643375-86643876 | 4 | 0.827987384176697 |
| chr3-106499462-106499963 | 8 | 0.971323177427264 |
| chr2-187762566-187763067 | 4 | 0.815322520495401 |
| chr13-66053933-66054434 | 8 | 0.842047039382297 |
| chr4-36235885-36236386 | 8 | 0.995135667989376 |
| chr19-4146923-4147424 | 8 | 0.997337976639521 |
| chr7-19844645-19845146 | 3 | 0.967037680816764 |
| chrX-77823434-77823935 | 8 | 0.926485706822216 |
| chr5-17430289-17430790 | 8 | 0.933759647561799 |
| chr9-48692351-48692852 | 8 | 0.900982760957104 |
| chr3-114477855-114478356 | 8 | 0.958538687947639 |
| chr13-66607597-66608098 | 8 | 0.955698504352989 |
| chr14-43017261-43017762 | 5 | 0.992503346077064 |
| chr18-11610304-11610805 | 8 | 0.97182315096361 |
| chr12-13325809-13326310 | 2 | 0.933514442530624 |
| chr1-169077121-169077622 | 8 | 0.984454435012217 |
| chr1-161028468-161028969 | 2 | 0.940698643399343 |
| chr6-114175801-114176302 | 5 | 0.935822389511084 |
| chr19-40142682-40143183 | 5 | 0.983586054096943 |
| chr17-84388729-84389230 | 8 | 0.979461530516077 |
| chr13-55826551-55827052 | 5 | 0.987218259650357 |
| chr12-21377882-21378383 | 8 | 0.948098479800059 |
| chr20-49523298-49523799 | 2 | 0.918756581575161 |
| chr2-48215458-48215959 | 3 | 0.997245792935381 |
| chr5-42700613-42701114 | 8 | 0.886537736339476 |
| chr6-59852037-59852538 | 6 | 0.991206925481043 |
| chr10-77094732-77095233 | 8 | 0.902156828729271 |
| chr17-43682412-43682913 | 8 | 0.98449065165798 |
| chr7-115118922-115119423 | 8 | 0.990346666018226 |
| chr4-94591308-94591809 | 8 | 0.905404091852561 |
| chr9-4947401-4947902 | 3 | 0.995935535450410 |

| | | |
|---|---|---|
| chr15-96344967-96345468 | 8 | 0.930479709010391 |
| chrX-24394302-24394803 | 8 | 0.868378560313145 |
| chr1-168762559-168763060 | 8 | 0.880828945421053 |
| chr2-67810370-67810871 | 8 | 0.994217723155897 |
| chr2-95012886-95013387 | 5 | 0.984009213642532 |
| chr19-33553542-33554043 | 8 | 0.957926326054848 |
| chr10-72486734-72487235 | 6 | 0.94357998338413 |
| chr2-31719540-31720041 | 5 | 0.983410236157418 |
| chr9-56118264-56118765 | 8 | 0.993493681128311 |
| chr3-109122112-109122613 | 8 | 0.842231624590348 |
| chr1-95231868-95232369 | 8 | 0.994339915295084 |
| chr11-54633998-54634499 | 8 | 0.968608597080302 |
| chrX-71669023-71669524 | 8 | 0.994713553299228 |
| chr2-45166019-45166520 | 5 | 0.98420708462735 |
| chr8-84744396-84744897 | 8 | 0.972984273421807 |
| chr3-91081522-91082023 | 8 | 0.995167907344419 |
| chr7-72636521-72637022 | 8 | 0.958000809170856 |
| chr11-53063553-53064054 | 8 | 0.983792163299938 |
| chr8-93050571-93051072 | 8 | 0.965386418901799 |
| chr14-4740856-4741357 | 5 | 0.974785642894519 |
| chr6-95979555-95980056 | 8 | 0.99510356668579 |
| chr9-81627216-81627717 | 8 | 0.957885074201898 |
| chr12-39944877-39945378 | 8 | 0.975892218276147 |
| chr5-30233956-30234457 | 8 | 0.99417797930858 |
| chr19-42861227-42861728 | 2 | 0.869051988824723 |
| chr9-97142722-97143223 | 2 | 0.894480172174511 |
| chr1-127824356-127824857 | 8 | 0.849354934302867 |
| chr5-67368921-67369422 | 8 | 0.993493681128311 |
| chr4-48948382-48948883 | 6 | 0.990731725323329 |
| chr1-187065010-187065511 | 8 | 0.94068811048398 |
| chr16-49547552-49548053 | 3 | 0.964244298512495 |
| chr15-15710386-15710887 | 5 | 0.970202731070428 |
| chr11-21294448-21294949 | 8 | 0.978932127907697 |
| chr11-14690596-14691097 | 8 | 0.969062999105133 |
| chr15-75050377-75050878 | 8 | 0.896174770496897 |
| chr6-40400780-40401281 | 8 | 0.995112947364898 |
| chr2-81842408-81842909 | 8 | 0.903003867047076 |
| chr4-141136965-141137466 | 8 | 0.993493681128311 |
| chr3-43445040-43445541 | 4 | 0.887323076471129 |
| chr15-26740639-26741140 | 8 | 0.993574194173503 |
| chr2-234835851-234836352 | 8 | 0.971602968667986 |
| chr2-247336352-247336853 | 8 | 0.993493681128311 |
| chr9-57443263-57443764 | 8 | 0.8833587655396 |

| | | |
|---|---|---|
| chr1-131199405-131199906 | 8 | 0.993493681128311 |
| chr15-33560959-33561460 | 8 | 0.989207120246659 |
| chr4-2819314-2819815 | 8 | 0.98221218263118 |
| chr4-109159647-109160148 | 8 | 0.849183449133172 |
| chr5-155067327-155067828 | 8 | 0.992433216545936 |
| chr16-82750663-82751164 | 8 | 0.966281940648309 |
| chr15-7610384-7610885 | 5 | 0.984365890884591 |
| chr4-26331991-26332492 | 5 | 0.987863102688507 |
| chr6-137290334-137290835 | 8 | 0.983452933385626 |
| chr15-6557877-6558378 | 8 | 0.993493681128311 |
| chr18-14871873-14872374 | 8 | 0.993493681128311 |
| chr9-111249450-111249951 | 5 | 0.987863882678954 |
| chrX-135213923-135214424 | 6 | 0.979609788993225 |
| chr3-78078916-78079417 | 8 | 0.994530224534823 |
| chr8-117892961-117893462 | 4 | 0.966640678755617 |
| chr6-13471825-13472326 | 8 | 0.9899554754131 |
| chr2-104888859-104889360 | 8 | 0.834803084014605 |
| chr10-20431028-20431529 | 8 | 0.934882957209801 |
| chr7-7922572-7923073 | 2 | 0.937921818540004 |
| chr17-36779392-36779893 | 2 | 0.932622175253159 |
| chr7-105616302-105616803 | 8 | 0.952826227099226 |
| chr16-3174968-3175469 | 6 | 0.913750822841457 |
| chr16-58149521-58150022 | 8 | 0.904834757119542 |
| chr5-144486599-144487100 | 8 | 0.940422254836342 |
| chr6-107364754-107365255 | 8 | 0.990312440359765 |
| chr5-78528460-78528961 | 6 | 0.987160401856127 |
| chr15-74753111-74753612 | 8 | 0.860236793466906 |
| chr4-113858925-113859426 | 8 | 0.973984154270808 |
| chr2-35240128-35240629 | 8 | 0.846718218868367 |
| chr19-12161075-12161576 | 8 | 0.981583841544016 |
| chr5-51169511-51170012 | 5 | 0.989122571201697 |
| chr2-58626061-58626562 | 8 | 0.96254022291739 |
| chr1-63634106-63634607 | 4 | 0.948072201876568 |
| chr17-36753496-36753997 | 8 | 0.864486372805012 |
| chr5-5338922-5339423 | 8 | 0.986832404463189 |
| chr11-69820489-69820990 | 5 | 0.833337537536445 |
| chr1-171566035-171566536 | 8 | 0.834027610486801 |
| chr1-34750322-34750823 | 8 | 0.993054802276672 |
| chr3-95143187-95143688 | 3 | 0.996853952543348 |
| chr2-178155834-178156335 | 8 | 0.874493446238586 |
| chr4-136975867-136976368 | 8 | 0.993493681128311 |
| chr4-116473037-116473538 | 8 | 0.993832326559451 |
| chr2-205326215-205326716 | 4 | 0.899574708979834 |

| | | |
|---|---|---|
| chr3-38910226-38910727 | 8 | 0.835735547627913 |
| chr18-25843771-25844272 | 8 | 0.986470792403894 |
| chr1-201262005-201262506 | 8 | 0.900158269969735 |
| chr2-247227048-247227549 | 5 | 0.972735180813071 |
| chr15-68999911-69000412 | 8 | 0.99524474010621 |
| chr11-67341148-67341649 | 8 | 0.993493681128311 |
| chr14-8583470-8583971 | 4 | 0.960855349647128 |
| chr11-28387411-28387912 | 3 | 0.996630359699831 |
| chr19-56305648-56306149 | 5 | 0.983675689419132 |
| chr1-56033185-56033686 | 3 | 0.954092203897818 |
| chr7-106174376-106174877 | 8 | 0.914810980973154 |
| chr1-89412465-89412966 | 8 | 0.881956386285539 |
| chr11-36923879-36924380 | 8 | 0.955610374923174 |
| chr4-24892414-24892915 | 4 | 0.909609779800201 |
| chr15-84329743-84330244 | 8 | 0.884797549147876 |
| chr5-47805990-47806491 | 8 | 0.832832581065725 |
| chr17-79501184-79501685 | 8 | 0.984995467402033 |
| chr6-59215810-59216311 | 2 | 0.90490850818178 |
| chr11-65474553-65475054 | 8 | 0.945958882873334 |
| chr1-36446731-36447232 | 8 | 0.985202824843595 |
| chr3-154367400-154367901 | 8 | 0.993846102491298 |
| chr6-23512937-23513438 | 8 | 0.993493681128311 |
| chr8-15311250-15311751 | 8 | 0.983823275649524 |
| chr20-22212137-22212638 | 8 | 0.900545397698535 |
| chr2-22280924-22281425 | 8 | 0.993658400598247 |
| chr10-32312824-32313325 | 8 | 0.984186001789033 |
| chr14-92411809-92412310 | 8 | 0.865544388082655 |
| chr20-12652288-12652789 | 8 | 0.858468215772908 |
| chr18-46568985-46569486 | 8 | 0.950074717207754 |
| chr4-58722237-58722738 | 8 | 0.967330999161925 |
| chr4-51089928-51090429 | 8 | 0.994841108899519 |
| chr20-17015089-17015590 | 8 | 0.9334534436915 |
| chr6-17562141-17562642 | 3 | 0.99593553545041 |
| chr19-43686720-43687221 | 5 | 0.986752504701627 |
| chr15-33051630-33052131 | 8 | 0.947006602413671 |
| chr1-185720791-185721292 | 3 | 0.879395099269473 |
| chr14-86206710-86207211 | 8 | 0.908902958709894 |
| chrX-30933867-30934368 | 8 | 0.963285633914194 |
| chr8-26842105-26842606 | 4 | 0.952202717609137 |
| chr7-11898652-11899153 | 8 | 0.887746961556044 |
| chr16-8224668-8225169 | 5 | 0.975820690293828 |
| chr8-78733854-78734355 | 8 | 0.97625841936901 |
| chr6-87571306-87571807 | 8 | 0.837625418536755 |

| | | |
|---|---|---|
| chr8-102574659-102575160 | 8 | 0.928623933672917 |
| chrX-120341564-120342065 | 5 | 0.852147024917739 |
| chr11-22073293-22073794 | 8 | 0.815739840056888 |
| chr14-91070923-91071424 | 5 | 0.952075984990487 |
| chr5-119205839-119206340 | 8 | 0.941133551142384 |
| chr5-3283739-3284240 | 4 | 0.975092241928153 |
| chr15-6382117-6382618 | 8 | 0.984462382241813 |
| chr4-108131339-108131840 | 8 | 0.982090579620694 |
| chrX-40593049-40593550 | 6 | 0.964588045469806 |
| chr2-65608346-65608847 | 5 | 0.983919978840557 |
| chr17-27811282-27811783 | 8 | 0.992039453130244 |
| chr3-99729367-99729868 | 8 | 0.993493681128311 |
| chr2-126955566-126956067 | 8 | 0.905147235332277 |
| chrX-72731757-72732258 | 3 | 0.996529461801484 |
| chr6-96654072-96654573 | 8 | 0.993983455601648 |
| chr6-42436609-42437110 | 8 | 0.936139073613964 |
| chr8-7769829-7770330 | 3 | 0.99593553545041 |
| chr15-38198073-38198574 | 3 | 0.990772636142748 |
| chr14-9498838-9499339 | 5 | 0.981805879241072 |
| chr2-72074605-72075106 | 6 | 0.895306648761824 |
| chr11-72437313-72437814 | 4 | 0.823394652323381 |
| chr4-154508297-154508798 | 8 | 0.944839046685539 |
| chr19-6116066-6116567 | 3 | 0.99593553545041 |
| chr2-169954866-169955367 | 8 | 0.945874280224615 |
| chr3-95518991-95519492 | 8 | 0.993493681128311 |
| chr9-89805570-89806071 | 8 | 0.900082901227466 |
| chr20-9580165-9580666 | 8 | 0.968922752119929 |
| chr17-19842836-19843337 | 5 | 0.984405298803915 |
| chr2-200216423-200216924 | 3 | 0.996517498717827 |
| chr4-112382907-112383408 | 4 | 0.878786699692627 |
| chr14-41285949-41286450 | 5 | 0.989095320903917 |
| chr7-70871446-70871947 | 4 | 0.84779797741966 |
| chr5-118571920-118572421 | 8 | 0.880720804789444 |
| chr7-83752977-83753478 | 8 | 0.988462128531996 |
| chr17-55784820-55785321 | 8 | 0.961911444182179 |
| chr14-66033920-66034421 | 3 | 0.99593553545041 |
| chr2-237608106-237608607 | 8 | 0.991651579026843 |
| chr18-48999456-48999957 | 4 | 0.829045969784559 |
| chr13-53350257-53350758 | 8 | 0.892160976385642 |
| chr11-79128485-79128986 | 8 | 0.989285483910275 |
| chr9-112056728-112057229 | 6 | 0.993983561854994 |
| chr1-9053536-9054037 | 8 | 0.909888390153511 |
| chr14-7673915-7674416 | 8 | 0.967042382236728 |

| | | |
|---|---|---|
| chr20-34088228-34088729 | 8 | 0.977565933149753 |
| chr1-213783785-213784286 | 8 | 0.95751047588578 |
| chr17-26276598-26277099 | 8 | 0.937881885131993 |
| chr8-55051402-55051903 | 5 | 0.972011867790618 |
| chr8-30063306-30063807 | 4 | 0.957962161775347 |
| chr12-40672569-40673070 | 8 | 0.982893727324205 |
| chr15-45963916-45964417 | 8 | 0.90908417229282 |
| chr20-23596091-23596592 | 8 | 0.992017449666919 |
| chr7-47696903-47697404 | 5 | 0.987769361352635 |
| chr5-91911979-91912480 | 8 | 0.911040660907357 |
| chr13-60373636-60374137 | 3 | 0.99593553545041 |
| chr16-32179144-32179645 | 8 | 0.982844010295132 |
| chr2-232607656-232608157 | 8 | 0.98812721509521 |
| chr14-46398000-46398501 | 8 | 0.956242806473819 |
| chr5-32617097-32617598 | 5 | 0.943835435931758 |
| chr4-167175173-167175674 | 5 | 0.983919978840557 |
| chr13-88888841-88889342 | 8 | 0.953087878651423 |
| chr6-74861327-74861828 | 8 | 0.993493681128311 |
| chr11-75037796-75038297 | 8 | 0.981659739563868 |
| chr19-25753314-25753815 | 5 | 0.990761960145384 |
| chr4-77688591-77689092 | 4 | 0.862308692210385 |
| chr2-237045001-237045502 | 8 | 0.993493681128311 |
| chr13-39610052-39610553 | 8 | 0.920500759025931 |
| chr18-1952347-1952848 | 8 | 0.983854749138161 |
| chr5-130314488-130314989 | 8 | 0.885030377569433 |
| chr18-74489607-74490108 | 5 | 0.992734785795061 |
| chr10-88391536-88392037 | 8 | 0.948202393050142 |
| chr2-80849768-80850269 | 4 | 0.968840259299856 |
| chrX-110879596-110880097 | 8 | 0.990795574801164 |
| chr13-89953932-89954433 | 8 | 0.995248948481242 |
| chr18-73984217-73984718 | 8 | 0.993493681128311 |
| chr6-127634353-127634854 | 8 | 0.849808808686763 |
| chr1-177310109-177310610 | 8 | 0.991286823451053 |
| chr7-91186789-91187290 | 4 | 0.918248665800859 |
| chr10-75413139-75413640 | 2 | 0.89450189764474 |
| chr14-62411265-62411766 | 8 | 0.953869692347185 |
| chr3-134294435-134294936 | 8 | 0.990318018490365 |
| chr3-43909096-43909597 | 3 | 0.844667219477427 |
| chr15-62705273-62705774 | 8 | 0.987798505925019 |
| chr6-121261025-121261526 | 8 | 0.920529677112066 |
| chr4-50336605-50337106 | 3 | 0.962609831195296 |
| chr2-105054484-105054985 | 2 | 0.851688154605298 |
| chr7-57351519-57352020 | 8 | 0.994949188390155 |

| | | |
|---|---|---|
| chr7-29030464-29030965 | 2 | 0.937805646563704 |
| chr7-44302069-44302570 | 8 | 0.99473753353112 |
| chr1-196815199-196815700 | 4 | 0.911806498529276 |
| chr8-14176848-14177349 | 8 | 0.88712649023845 |
| chrX-36209989-36210490 | 8 | 0.994956616830076 |
| chr17-63740247-63740748 | 8 | 0.990240563301755 |
| chr17-51281043-51281544 | 8 | 0.864332418374618 |
| chr20-32422572-32423073 | 2 | 0.854646249651848 |
| chr1-45332579-45333080 | 8 | 0.975357055608943 |
| chr13-66542730-66543231 | 5 | 0.882406579381502 |
| chr2-133452850-133453351 | 5 | 0.862753250691555 |
| chr14-60318303-60318804 | 3 | 0.99593553545041 |
| chr3-94250796-94251297 | 4 | 0.881435470662308 |
| chr13-89772141-89772642 | 8 | 0.971366996085458 |
| chr7-97826958-97827459 | 3 | 0.99593553545041 |
| chr5-6796764-6797265 | 2 | 0.90320500974233 |
| chr1-91589020-91589521 | 8 | 0.897335542102537 |
| chr7-35932816-35933317 | 6 | 0.992818086189911 |
| chr6-31072521-31073022 | 8 | 0.994964730052285 |
| chr3-36323335-36323836 | 8 | 0.874086774595232 |
| chr16-76858166-76858667 | 8 | 0.978138495707787 |
| chr14-20573116-20573617 | 8 | 0.846258887170255 |
| chr2-53647601-53648102 | 5 | 0.996709927733102 |
| chr17-71970690-71971191 | 8 | 0.981812835491153 |
| chr16-40599597-40600098 | 8 | 0.964532027469195 |
| chr6-108502071-108502572 | 4 | 0.957338855394469 |
| chr11-9577183-9577684 | 4 | 0.879063714505219 |
| chr10-103807609-103808110 | 8 | 0.980956155757085 |
| chr3-111366020-111366521 | 5 | 0.979440494248143 |
| chr4-165074504-165075005 | 8 | 0.980918166793766 |
| chr18-23656337-23656838 | 8 | 0.99264037758058 |
| chr19-16344950-16345451 | 8 | 0.987045090298393 |
| chr11-67715555-67716056 | 8 | 0.865877056856674 |
| chr6-28068282-28068783 | 8 | 0.992160226371337 |
| chr2-187989941-187990442 | 8 | 0.964129087732508 |
| chr2-95090789-95091290 | 8 | 0.865784572438557 |
| chr3-90897850-90898351 | 5 | 0.993761857952802 |
| chr7-117454714-117455215 | 4 | 0.964415631687865 |
| chr20-51061948-51062449 | 2 | 0.820228668903141 |
| chr12-37516628-37517129 | 5 | 0.872644798541784 |
| chr2-143115830-143116331 | 8 | 0.993493681128311 |
| chr15-554360-554861 | 8 | 0.993493681128311 |
| chr2-245955208-245955709 | 2 | 0.934201763694055 |

| | | |
|---|---|---|
| chr9-93752351-93752852 | 8 | 0.860975411271875 |
| chr20-21836261-21836762 | 5 | 0.984412841607147 |
| chr8-46867503-46868004 | 8 | 0.942330326165701 |
| chr3-33145871-33146372 | 4 | 0.881444808927451 |
| chr2-205463206-205463707 | 3 | 0.996740924997324 |
| chr15-59658980-59659481 | 5 | 0.98146658357413 |
| chr4-51205064-51205565 | 8 | 0.912526070488271 |
| chr16-12290096-12290597 | 5 | 0.983919978840557 |
| chr3-139321476-139321977 | 8 | 0.993493681128311 |
| chr11-55632418-55632919 | 8 | 0.962230277922087 |
| chr8-95315529-95316030 | 8 | 0.827531699973049 |
| chr9-55734891-55735392 | 5 | 0.908627935190068 |
| chr12-1361575-1362076 | 3 | 0.871664303580802 |
| chr5-130315991-130316492 | 8 | 0.924482849392744 |
| chr8-13210133-13210634 | 2 | 0.87087526618636 |
| chr17-54622054-54622555 | 8 | 0.852213247449471 |
| chr13-6204576-6205077 | 5 | 0.959716948486908 |
| chr11-33810684-33811185 | 8 | 0.968524755973731 |
| chr17-46199409-46199910 | 8 | 0.98240325631936 |
| chr6-36942343-36942844 | 8 | 0.970862814001562 |
| chr5-107501597-107502098 | 6 | 0.898550872893516 |
| chr9-18702847-18703348 | 3 | 0.991979930819446 |
| chr2-86850540-86851041 | 8 | 0.964403210239104 |
| chr5-79067167-79067668 | 2 | 0.919438853100519 |
| chr7-72511278-72511779 | 8 | 0.941796207182078 |
| chr1-220314744-220315245 | 8 | 0.99514181418919 |
| chrX-132614170-132614671 | 8 | 0.993493681128311 |
| chr3-32478436-32478937 | 8 | 0.943318565165519 |
| chr18-43108922-43109423 | 8 | 0.864281209102523 |
| chr2-92664944-92665445 | 8 | 0.899262683047215 |
| chr2-5114089-5114590 | 4 | 0.958851712898465 |
| chrX-16683060-16683561 | 4 | 0.971161935658001 |
| chr1-144812905-144813406 | 4 | 0.970583 |
| chr2-83917839-83918340 | 8 | 0.824929484296973 |
| chr2-130288392-130288893 | 5 | 0.841201326995145 |
| chr1-241793643-241794144 | 8 | 0.904192188017643 |
| chr15-55375377-55375878 | 8 | 0.941943482000389 |
| chr9-31729232-31729733 | 3 | 0.99593553545041 |
| chr14-92572559-92573060 | 8 | 0.92653565235632 |
| chr12-41849453-41849954 | 8 | 0.975230855338178 |
| chr6-4852030-4852531 | 8 | 0.976123100455842 |
| chr5-66635701-66636202 | 2 | 0.901269824144629 |
| chr6-68929491-68929992 | 8 | 0.90148835128637 |

| | | |
|---|---|---|
| chr1-234322995-234323496 | 8 | 0.883834790281617 |
| chr1-220924657-220925158 | 5 | 0.952620297522591 |
| chr14-37508192-37508693 | 8 | 0.817794306135174 |
| chr11-22185380-22185881 | 8 | 0.985769055523761 |
| chr18-42623591-42624092 | 8 | 0.948416102450783 |
| chr20-47266921-47267422 | 8 | 0.927597272490192 |
| chr8-33285623-33286124 | 8 | 0.983191931600584 |
| chr1-257926840-257927341 | 8 | 0.970042903356928 |
| chr2-142617425-142617926 | 8 | 0.992743686677312 |
| chr14-94909234-94909735 | 5 | 0.985549403462906 |
| chr1-56352598-56353099 | 8 | 0.867342863728189 |
| chr1-208079090-208079591 | 8 | 0.894223764052503 |
| chr2-134555736-134556237 | 3 | 0.987725375823558 |
| chr4-30306451-30306952 | 8 | 0.83710275608969 |
| chr1-157644136-157644637 | 5 | 0.962434569029585 |
| chr2-40058561-40059062 | 8 | 0.944736034574215 |
| chr1-70208663-70209164 | 8 | 0.993493681128311 |
| chr18-17257395-17257896 | 8 | 0.975537720790067 |
| chr1-178097451-178097952 | 8 | 0.867274958117814 |
| chr17-14720675-14721176 | 5 | 0.983831901959103 |
| chr13-53992700-53993201 | 8 | 0.994852406473016 |
| chr9-84437782-84438283 | 8 | 0.995239112841556 |
| chr8-80699900-80700401 | 5 | 0.945472490474888 |
| chr15-47842874-47843375 | 8 | 0.960285381120839 |
| chr13-25364050-25364551 | 8 | 0.993493681128311 |
| chr16-159553-160054 | 8 | 0.913757526787242 |
| chr5-53991626-53992127 | 8 | 0.987522073220648 |
| chr11-10873351-10873852 | 8 | 0.948798285488718 |
| chr9-62724668-62725169 | 8 | 0.886610339511214 |
| chr1-87330189-87330690 | 8 | 0.9611187897118 |
| chrX-13464953-13465454 | 8 | 0.994748729656265 |
| chr3-132296597-132297098 | 8 | 0.954723774613094 |
| chr13-78399286-78399787 | 2 | 0.934748698926973 |
| chr1-49983613-49984114 | 8 | 0.931393546483255 |
| chr1-259248291-259248792 | 5 | 0.979044569412639 |
| chr5-28280065-28280566 | 8 | 0.993493681128311 |
| chr2-15821630-15822131 | 8 | 0.9432037460665 |
| chr8-113523618-113524119 | 4 | 0.906586897624001 |
| chr6-34344801-34345302 | 8 | 0.879460662301045 |
| chr13-85551258-85551759 | 4 | 0.878995945100862 |
| chr1-90368874-90369375 | 8 | 0.911807688216994 |
| chr14-88552806-88553307 | 6 | 0.989019769451024 |
| chr19-42072316-42072817 | 8 | 0.978417532531634 |

| | | |
|---|---|---|
| chr9-4160991-4161492 | 8 | 0.993493681128311 |
| chr3-136350930-136351431 | 3 | 0.996506270843571 |
| chr9-37732177-37732678 | 8 | 0.928378810447498 |
| chr7-78964699-78965200 | 2 | 0.834532976148482 |
| chr13-87586149-87586650 | 8 | 0.935599107631163 |
| chr13-71860710-71861211 | 8 | 0.954590447732174 |
| chr15-97115581-97116082 | 8 | 0.986758595397502 |
| chr3-8543645-8544146 | 5 | 0.98277491640377 |
| chr15-5836219-5836720 | 5 | 0.983919978840557 |
| chr2-57291236-57291737 | 8 | 0.855703495064951 |
| chr5-138869406-138869907 | 8 | 0.888102824362327 |
| chr4-33200968-33201469 | 3 | 0.996422011553898 |
| chr20-35452750-35453251 | 8 | 0.993493681128311 |
| chr6-9242838-9243339 | 4 | 0.92516545344993 |
| chr10-93352826-93353327 | 8 | 0.969645838085954 |
| chr3-17320837-17321338 | 8 | 0.929888962368966 |
| chr20-51353686-51354187 | 3 | 0.989358188506365 |
| chr4-34620494-34620995 | 2 | 0.940700629803965 |
| chr1-132075514-132076015 | 8 | 0.975436854896684 |
| chr3-55662637-55663138 | 8 | 0.992139509088781 |
| chr9-81933623-81934124 | 8 | 0.993493681128311 |
| chr20-21807710-21808211 | 8 | 0.876250854186368 |
| chr1-57947539-57948040 | 8 | 0.995215722716292 |
| chr11-47549820-47550321 | 8 | 0.840762720196169 |
| chr14-20368586-20369087 | 2 | 0.857820774326493 |
| chr10-48964123-48964624 | 8 | 0.994355808698783 |
| chr9-23274666-23275167 | 8 | 0.992621212111107 |
| chr2-16676752-16677253 | 5 | 0.948486056798046 |
| chr2-27395612-27396113 | 8 | 0.985712838304567 |
| chr2-19340658-19341159 | 8 | 0.989473112668604 |
| chr13-41179167-41179668 | 8 | 0.995228456485316 |
| chr2-28101104-28101605 | 4 | 0.925559623428393 |
| chr4-150841024-150841525 | 8 | 0.983177329158133 |
| chr9-45652275-45652776 | 8 | 0.971250339811625 |
| chr5-66536071-66536572 | 3 | 0.996556678855797 |
| chr4-67099353-67099854 | 2 | 0.939453985413214 |
| chr16-5222725-5223226 | 5 | 0.986358340572996 |
| chr3-160707899-160708400 | 8 | 0.914526923330897 |
| chr10-90296144-90296645 | 8 | 0.952880945023927 |
| chr19-44143159-44143660 | 5 | 0.987177929150826 |
| chr8-28624092-28624593 | 8 | 0.857258484151183 |
| chr14-61267129-61267630 | 5 | 0.989862611645446 |
| chr3-127344514-127345015 | 4 | 0.880849086479206 |

| | | |
|---|---|---|
| chr2-192833488-192833989 | 6 | 0.989826224610225 |
| chr10-93001238-93001739 | 8 | 0.914350384884487 |
| chr15-53281695-53282196 | 8 | 0.98551334288576 |
| chr6-58509414-58509915 | 8 | 0.990424757457286 |
| chr1-61706577-61707078 | 2 | 0.832723375277497 |
| chr17-32749932-32750433 | 3 | 0.996010175689606 |
| chr1-257112087-257112588 | 8 | 0.912775582833359 |
| chr1-108407064-108407565 | 8 | 0.927366413780615 |
| chr8-53691933-53692434 | 5 | 0.973232222000574 |
| chr19-9626151-9626652 | 8 | 0.969558209272614 |
| chr10-63523193-63523694 | 8 | 0.986502685556437 |
| chr2-34288188-34288689 | 8 | 0.994857775152304 |
| chr10-18281026-18281527 | 8 | 0.929125540462807 |
| chr3-6138329-6138830 | 8 | 0.936367330397977 |
| chr17-23365325-23365826 | 8 | 0.951677982773785 |
| chr13-85122935-85123436 | 3 | 0.936653820620358 |
| chr2-123721883-123722384 | 5 | 0.889433250532685 |
| chr5-40497626-40498127 | 3 | 0.99593553545041 |
| chr8-84184992-84185493 | 8 | 0.993493681128311 |
| chr6-30747646-30748147 | 4 | 0.965672335274956 |
| chr2-31615335-31615836 | 8 | 0.995019792821361 |
| chr10-1328055-1328556 | 2 | 0.866696843397399 |
| chr3-124053654-124054155 | 8 | 0.887498513453891 |
| chr7-92887287-92887788 | 8 | 0.994579746484556 |
| chr5-34303532-34304033 | 8 | 0.917912606415657 |
| chr8-29924502-29925003 | 2 | 0.891096423655405 |
| chrX-46926887-46927388 | 5 | 0.938154426210383 |
| chr17-33672448-33672949 | 8 | 0.841878763676235 |
| chr6-107683873-107684374 | 2 | 0.879241531178447 |
| chr3-77649236-77649737 | 8 | 0.955463279820051 |
| chr4-46171751-46172252 | 8 | 0.993493681128311 |
| chr14-67767905-67768406 | 4 | 0.948237799861832 |
| chr18-63083785-63084286 | 8 | 0.919416534781003 |
| chr9-65056410-65056911 | 4 | 0.941339959090672 |
| chr11-294262-294763 | 3 | 0.996593327301283 |
| chr4-173838576-173839077 | 8 | 0.979183608361724 |
| chr2-41733750-41734251 | 8 | 0.993493681128311 |
| chr7-134729422-134729923 | 8 | 0.993493681128311 |
| chr20-28452951-28453452 | 5 | 0.984714769964966 |
| chr15-49879491-49879992 | 8 | 0.900717262557439 |
| chr12-7736113-7736614 | 8 | 0.993493681128311 |
| chr2-144491242-144491743 | 8 | 0.884873837557025 |
| chr2-125010123-125010624 | 8 | 0.911397018878338 |

| | | |
|---|---|---|
| chr16-35628986-35629487 | 2 | 0.889123648826062 |
| chr15-73727678-73728179 | 6 | 0.991206925481043 |
| chr18-16575146-16575647 | 8 | 0.994895674226769 |
| chr1-82014433-82014934 | 8 | 0.989625746847602 |
| chr4-51099958-51100459 | 5 | 0.942441338379027 |
| chr9-85286808-85287309 | 8 | 0.981085544909134 |
| chr2-22245089-22245590 | 8 | 0.993493681128311 |
| chr4-106589652-106590153 | 8 | 0.925787572027984 |
| chr10-769526-770027 | 8 | 0.865050028427559 |
| chr20-13658177-13658678 | 5 | 0.840282405417152 |
| chr14-1525040-1525541 | 2 | 0.941447025394202 |
| chr1-117771961-117772462 | 8 | 0.971510069107104 |
| chr2-60447091-60447592 | 8 | 0.991401014570431 |
| chr7-16718439-16718940 | 8 | 0.823790587966443 |
| chr7-20124143-20124644 | 8 | 0.994234087391888 |
| chr7-49095591-49096092 | 3 | 0.99593553545041 |
| chr1-236991440-236991941 | 3 | 0.995177517242999 |
| chr19-11404528-11405029 | 8 | 0.979275275894188 |
| chr5-25956140-25956641 | 8 | 0.938775099949301 |
| chr19-20044189-20044690 | 5 | 0.991174972383896 |
| chr13-81552871-81553372 | 8 | 0.993493681128311 |
| chr13-54212334-54212835 | 8 | 0.908382696180244 |
| chr2-100440641-100441142 | 8 | 0.899292458650313 |
| chr14-93497625-93498126 | 5 | 0.982750200199782 |
| chr3-106169527-106170028 | 8 | 0.978366851831808 |
| chr6-139037915-139038416 | 8 | 0.966103366555783 |
| chr3-28616771-28617272 | 8 | 0.994938704146882 |
| chr8-104641352-104641853 | 8 | 0.995158732890697 |
| chr9-34665345-34665846 | 8 | 0.906128117587942 |
| chr17-374521-375022 | 3 | 0.996480676287119 |
| chr5-1289226-1289727 | 8 | 0.914727078191225 |
| chr15-91686462-91686963 | 6 | 0.988267749586541 |
| chr14-30813729-30814230 | 8 | 0.85637623347739 |
| chr19-16265117-16265618 | 8 | 0.992876561950919 |
| chr14-70253540-70254041 | 8 | 0.993493681128311 |
| chr17-51573415-51573916 | 5 | 0.858222274060042 |
| chr8-7426481-7426982 | 8 | 0.994931965486057 |
| chr1-160772816-160773317 | 8 | 0.986526042249749 |
| chr14-94282831-94283332 | 8 | 0.964099382089892 |
| chr13-38943848-38944349 | 4 | 0.963508107240602 |
| chr1-185724445-185724946 | 3 | 0.846805386131182 |
| chr2-37472025-37472526 | 4 | 0.975653035145407 |
| chr17-38522951-38523452 | 8 | 0.968747253575784 |

| | | |
|---|---|---|
| chr14-97984883-97985384 | 8 | 0.995208821972034 |
| chr2-208155019-208155520 | 8 | 0.876063201992109 |
| chr6-39884399-39884900 | 8 | 0.96931957153628 |
| chr11-74739089-74739590 | 8 | 0.972068045238098 |
| chr5-73068778-73069279 | 8 | 0.917041021614765 |
| chr3-142845905-142846406 | 8 | 0.993853600357835 |
| chr14-94116590-94117091 | 8 | 0.981963771983075 |
| chr11-59572862-59573363 | 8 | 0.993493681128311 |
| chr13-55660227-55660728 | 8 | 0.995167462057486 |
| chr17-42855246-42855747 | 8 | 0.995083870385015 |
| chr2-203581471-203581972 | 4 | 0.84661990710265 |
| chr3-168989206-168989707 | 8 | 0.894585324561307 |
| chr19-34151050-34151551 | 8 | 0.977631168336614 |
| chr2-95228252-95228753 | 5 | 0.946164145701858 |
| chr3-142844849-142845350 | 8 | 0.942160883354721 |
| chr15-97185558-97186059 | 8 | 0.816220173834143 |
| chr11-72076095-72076596 | 8 | 0.982247705719901 |
| chr13-8875138-8875639 | 3 | 0.977787144549883 |
| chr7-109493691-109494192 | 3 | 0.845709255164296 |
| chr11-40781473-40781974 | 8 | 0.906496686937198 |
| chr13-64508881-64509382 | 2 | 0.950803492681149 |
| chr19-3954479-3954980 | 8 | 0.991861977181758 |
| chr1-109519166-109519667 | 8 | 0.947290745190977 |
| chr3-107822699-107823200 | 4 | 0.850519619986508 |
| chr2-122854282-122854783 | 8 | 0.954062824789165 |
| chr20-39322408-39322909 | 8 | 0.993493681128311 |
| chr2-245800700-245801201 | 8 | 0.993493681128311 |
| chr18-18333320-18333821 | 6 | 0.993142785485641 |
| chr2-172883690-172884191 | 8 | 0.877829296907783 |
| chr14-66339538-66340039 | 5 | 0.979024 |
| chr7-115910543-115911044 | 5 | 0.983091086994974 |
| chr18-70252941-70253442 | 8 | 0.993493681128311 |
| chr6-112783615-112784116 | 2 | 0.918079853717854 |
| chr6-5118092-5118593 | 8 | 0.993493681128311 |
| chr1-30120362-30120863 | 8 | 0.995157797119419 |
| chr4-37490902-37491403 | 8 | 0.993493681128311 |
| chr20-32018426-32018927 | 8 | 0.987241044774065 |
| chr5-23671317-23671818 | 5 | 0.90024295006384 |
| chr10-84543337-84543838 | 8 | 0.970840008997962 |
| chr16-70736839-70737340 | 8 | 0.993493681128311 |
| chr9-586066-586567 | 5 | 0.981369196022631 |
| chr20-14032037-14032538 | 8 | 0.964089625912185 |
| chr17-6887924-6888425 | 8 | 0.995152125479146 |

| | | |
|---|---|---|
| chr8-25014775-25015276 | 6 | 0.993883685387372 |
| chr1-22512546-22513047 | 8 | 0.993493681128311 |
| chr4-168407360-168407861 | 8 | 0.983713029389087 |
| chr6-134634946-134635447 | 6 | 0.994317762156821 |
| chr17-51957765-51958266 | 8 | 0.867078183637635 |
| chr15-51319253-51319754 | 8 | 0.951770294498242 |
| chr8-61401461-61401962 | 8 | 0.839704972511411 |
| chr7-120615899-120616400 | 8 | 0.948080033128891 |
| chrX-101433566-101434067 | 3 | 0.99593553545041 |
| chr6-86982524-86983025 | 8 | 0.993493681128311 |
| chrX-107724025-107724526 | 8 | 0.995237229816187 |
| chr4-176350604-176351105 | 8 | 0.882104986116167 |
| chr16-75616744-75617245 | 5 | 0.991682976789121 |
| chr5-94037213-94037714 | 2 | 0.931245833497375 |
| chr4-133083268-133083769 | 8 | 0.893781453548582 |
| chr17-6359364-6359865 | 8 | 0.993493681128311 |
| chr12-38061982-38062483 | 4 | 0.96470906682287 |
| chr18-77504329-77504830 | 8 | 0.911602415313684 |
| chr18-36664041-36664542 | 8 | 0.979069835721612 |
| chr6-125080599-125081100 | 4 | 0.971399103078162 |
| chr19-48675810-48676311 | 8 | 0.949628397644323 |
| chr4-130160415-130160916 | 8 | 0.963069130973609 |
| chr11-16200965-16201466 | 4 | 0.966632992579084 |
| chr15-81642264-81642765 | 8 | 0.847667861898514 |
| chr17-23102585-23103086 | 5 | 0.983829319974392 |
| chr1-152926306-152926807 | 8 | 0.97186087650759 |
| chr5-62843366-62843867 | 8 | 0.993493681128311 |
| chr14-73054620-73055121 | 8 | 0.995162427452917 |
| chr9-69347555-69348056 | 8 | 0.994054750149055 |
| chr2-3473359-3473860 | 8 | 0.993444552077129 |
| chr13-100611835-100612336 | 8 | 0.989473835796193 |
| chr15-14162785-14163286 | 2 | 0.921536688174424 |
| chr5-112988650-112989151 | 8 | 0.933374686610217 |
| chr5-36386527-36387028 | 3 | 0.996806562664598 |
| chr2-13908578-13909079 | 8 | 0.929620585293235 |
| chr1-132727392-132727893 | 4 | 0.837098501982645 |
| chr18-33112838-33113339 | 8 | 0.981514943330782 |
| chr4-33172242-33172743 | 5 | 0.948535297484119 |
| chr19-51190665-51191166 | 8 | 0.92967247642568 |
| chr13-38827167-38827668 | 5 | 0.968819074306956 |
| chr9-69672229-69672730 | 5 | 0.990287713346974 |
| chr17-51671338-51671839 | 5 | 0.970858933297317 |
| chr11-63843820-63844321 | 8 | 0.993935136559023 |

| | | |
|---|---|---|
| chr3-120478050-120478551 | 8 | 0.983374696615179 |
| chr4-123305734-123306235 | 8 | 0.852045642757836 |
| chr3-107708931-107709432 | 8 | 0.995133756690036 |
| chr11-60834906-60835407 | 8 | 0.974468590534309 |
| chr2-45158846-45159347 | 5 | 0.962056766442214 |
| chrX-127038870-127039371 | 8 | 0.995171316462112 |
| chr14-39682122-39682623 | 8 | 0.940624117723646 |
| chr2-129681144-129681645 | 3 | 0.852294468979861 |
| chr6-39460691-39461192 | 5 | 0.983797309876391 |
| chr7-82685072-82685573 | 8 | 0.979220945210529 |
| chr1-227633460-227633961 | 8 | 0.920008463785785 |
| chr6-34233465-34233966 | 6 | 0.996067010145428 |
| chr1-26381045-26381546 | 8 | 0.990403320542889 |
| chr10-57305551-57306052 | 8 | 0.983549304455529 |
| chr18-53722423-53722924 | 8 | 0.828554942464781 |
| chr14-36471192-36471693 | 3 | 0.99593553545041 |
| chr5-126803401-126803902 | 3 | 0.907392267770651 |
| chr2-125564877-125565378 | 5 | 0.830974775110235 |
| chr1-53870757-53871258 | 8 | 0.900639723193353 |
| chr1-181760277-181760778 | 5 | 0.988495138862221 |
| chr3-34208179-34208680 | 8 | 0.963068820768495 |
| chr4-71791155-71791656 | 6 | 0.983839197678157 |
| chr6-31500157-31500658 | 8 | 0.919281349083244 |
| chr17-45964787-45965288 | 5 | 0.969324001279411 |
| chr3-109199148-109199649 | 8 | 0.973169907060996 |
| chr3-158889193-158889694 | 8 | 0.990706997261383 |
| chr7-123498330-123498831 | 8 | 0.976514054499396 |
| chr13-76380688-76381189 | 3 | 0.850322684096424 |
| chr10-6084729-6085230 | 8 | 0.994925517278108 |
| chr6-20665462-20665963 | 8 | 0.962666141630367 |
| chr1-208754744-208755245 | 6 | 0.981813941417684 |
| chr8-18886134-18886635 | 8 | 0.989518746116865 |
| chr13-57600768-57601269 | 5 | 0.88490858234429 |
| chr10-67256220-67256721 | 5 | 0.915555727589381 |
| chr15-1941843-1942344 | 8 | 0.967788640896847 |
| chr14-1588860-1589361 | 8 | 0.907001784921001 |
| chr8-94460804-94461305 | 8 | 0.970347042312481 |
| chr9-31256817-31257318 | 8 | 0.993493681128311 |
| chr11-56494868-56495369 | 8 | 0.994830610006959 |
| chr5-29755198-29755699 | 6 | 0.991206925481043 |
| chr1-190004294-190004795 | 8 | 0.994018505645738 |
| chr15-41422661-41423162 | 8 | 0.980403770735135 |
| chrX-101249089-101249590 | 3 | 0.99593553545041 |

| | | |
|---|---|---|
| chr5-79675602-79676103 | 8 | 0.862848549915538 |
| chr2-100609450-100609951 | 4 | 0.835575673287758 |
| chr15-72353753-72354254 | 8 | 0.994084306725228 |
| chr15-12031839-12032340 | 3 | 0.99593553545041 |
| chr2-201473641-201474142 | 8 | 0.957644254024847 |
| chr8-103017239-103017740 | 8 | 0.994720501882136 |
| chr5-69777727-69778228 | 8 | 0.970994857619808 |
| chr18-63540594-63541095 | 5 | 0.987924697335486 |
| chr13-42729281-42729782 | 5 | 0.991251080961947 |
| chr12-30285846-30286347 | 8 | 0.993493681128311 |
| chr2-27606812-27607313 | 4 | 0.844471213653937 |
| chr4-17257782-17258283 | 8 | 0.923492954474552 |
| chr1-23270893-23271394 | 3 | 0.95961507694222 |
| chr16-23694457-23694958 | 8 | 0.994796801855559 |
| chr2-86849946-86850447 | 8 | 0.895641061696849 |
| chr6-90549973-90550474 | 8 | 0.961348398623274 |
| chr6-14575437-14575938 | 5 | 0.96717329238051 |
| chr8-78296766-78297267 | 8 | 0.882692504887271 |
| chr16-39444264-39444765 | 3 | 0.996049710896441 |
| chr3-21680559-21681060 | 8 | 0.9948397615597 |
| chr6-81068887-81069388 | 6 | 0.994139513444534 |
| chr9-94237396-94237897 | 8 | 0.968353218608345 |
| chr18-48035756-48036257 | 5 | 0.949616341209516 |
| chr5-118909609-118910110 | 8 | 0.960062129469484 |
| chr6-89578987-89579488 | 8 | 0.986705802604756 |
| chr4-75479853-75480354 | 8 | 0.977240556516645 |
| chr2-153720055-153720556 | 8 | 0.966702115550522 |
| chr4-180260834-180261335 | 8 | 0.882437165345654 |
| chr17-38333417-38333918 | 8 | 0.945341307668411 |
| chr1-131818806-131819307 | 8 | 0.815639202772745 |
| chr18-21585477-21585978 | 5 | 0.893073037888539 |
| chr3-73941580-73942081 | 8 | 0.843864749264266 |
| chr2-241469904-241470405 | 8 | 0.987450422820446 |
| chr19-42262088-42262589 | 4 | 0.894380182514138 |
| chr10-89887167-89887668 | 5 | 0.861573673755081 |
| chr11-12414194-12414695 | 2 | 0.820042020163277 |
| chr3-111787472-111787973 | 5 | 0.982780477650746 |
| chr5-12938941-12939442 | 8 | 0.990775691641441 |
| chr8-7837111-7837612 | 5 | 0.961788043166246 |
| chr2-129838872-129839373 | 3 | 0.987451211744771 |
| chr2-46088551-46089052 | 5 | 0.987954497578507 |
| chr6-54349648-54350149 | 3 | 0.976110659811787 |
| chr13-56956754-56957255 | 5 | 0.950663996626096 |

| | | |
|---|---|---|
| chr9-37349744-37350245 | 5 | 0.923445522959172 |
| chr13-37645258-37645759 | 8 | 0.969558991518373 |
| chr12-37591145-37591646 | 8 | 0.944106481554159 |
| chr17-23772607-23773108 | 4 | 0.944118697163797 |
| chr9-54226504-54227005 | 8 | 0.903470069011424 |
| chr2-182962983-182963484 | 2 | 0.964788236404462 |
| chrX-7143113-7143614 | 3 | 0.99593553545041 |
| chr3-28807410-28807911 | 8 | 0.925854830527572 |
| chr6-67520985-67521486 | 6 | 0.980812216167615 |
| chr18-201794-202295 | 8 | 0.880341136251767 |
| chr17-72773-73274 | 8 | 0.938092841430186 |
| chr18-28465125-28465626 | 8 | 0.92860884059815 |
| chr11-55784697-55785198 | 8 | 0.993493681128311 |
| chr3-32721766-32722267 | 8 | 0.923427918492657 |
| chr6-112252009-112252510 | 3 | 0.993680393698198 |
| chr3-90664160-90664661 | 8 | 0.974004532906701 |
| chr14-100240619-100241120 | 3 | 0.988330992110747 |
| chr8-103143162-103143663 | 8 | 0.82386560424351 |
| chr7-93374605-93375106 | 2 | 0.93961746277618 |
| chr4-72115411-72115912 | 8 | 0.975251146588962 |
| chr8-22605224-22605725 | 8 | 0.829472176756599 |
| chr14-11224191-11224692 | 8 | 0.874575349617978 |
| chr19-48096398-48096899 | 5 | 0.960802005380817 |
| chr8-21379897-21380398 | 8 | 0.841785259218916 |
| chr1-238733202-238733703 | 6 | 0.992702714410038 |
| chr4-87908968-87909469 | 3 | 0.99689781428656 |
| chr6-125663661-125664162 | 8 | 0.981613226324338 |
| chr14-72780702-72781203 | 8 | 0.991601199938105 |
| chr20-50489606-50490107 | 3 | 0.995972220873445 |
| chr2-216512343-216512844 | 3 | 0.993799382231378 |
| chr13-94946151-94946652 | 8 | 0.993794509153169 |
| chr9-93723518-93724019 | 8 | 0.882325204915229 |
| chr10-6162871-6163372 | 8 | 0.938975707477434 |
| chr2-102498365-102498866 | 3 | 0.963129678547411 |
| chr14-52418209-52418710 | 8 | 0.95541615764494 |
| chr2-188737955-188738456 | 8 | 0.993493681128311 |
| chr7-99359684-99360185 | 8 | 0.987146693691707 |
| chr18-9328906-9329407 | 6 | 0.975583034409997 |
| chr7-84761434-84761935 | 8 | 0.94402372517457 |
| chr18-56954712-56955213 | 8 | 0.909141519802567 |
| chr2-186026604-186027105 | 8 | 0.964599379189664 |
| chr13-72593150-72593651 | 8 | 0.855038255454204 |
| chr20-16904672-16905173 | 4 | 0.919419381871989 |

| | | |
|---|---|---|
| chr11-41998858-41999359 | 8 | 0.993493681128311 |
| chr2-229694115-229694616 | 8 | 0.966619063097161 |
| chr2-34356369-34356870 | 8 | 0.989552314811044 |
| chr6-106322518-106323019 | 5 | 0.848353109487089 |
| chr13-78639438-78639939 | 8 | 0.836149080710009 |
| chr13-76412103-76412604 | 3 | 0.975829932713134 |
| chr1-79742738-79743239 | 8 | 0.972699680427589 |
| chr3-109448715-109449216 | 8 | 0.855310438355824 |
| chr3-84250156-84250657 | 8 | 0.988858741285514 |
| chr10-39673384-39673885 | 8 | 0.994133294647586 |
| chr20-18036316-18036817 | 8 | 0.962136645760672 |
| chr8-12374137-12374638 | 8 | 0.839177183730422 |
| chr6-38224015-38224516 | 4 | 0.962822760460194 |
| chr2-229785813-229786314 | 3 | 0.879180175774476 |
| chr2-223777864-223778365 | 8 | 0.988221473548537 |
| chr19-22556332-22556833 | 8 | 0.979917847040546 |
| chr6-113306368-113306869 | 3 | 0.996060356654772 |
| chr8-3174519-3175020 | 8 | 0.993493681128311 |
| chr1-7996567-7997068 | 4 | 0.893027877636661 |
| chr4-25895903-25896404 | 8 | 0.993493681128311 |
| chr17-59478859-59479360 | 8 | 0.870144541693004 |
| chr16-17411362-17411863 | 8 | 0.952155757355218 |
| chr1-191509359-191509860 | 8 | 0.915673553233473 |
| chr13-18176660-18177161 | 3 | 0.99593553545041 |
| chr18-15552617-15553118 | 8 | 0.940724416071068 |
| chr9-48657300-48657801 | 8 | 0.972108627395133 |
| chr15-64185487-64185988 | 4 | 0.921728715065239 |
| chr15-69682762-69683263 | 8 | 0.893304090743883 |
| chr11-56619499-56620000 | 8 | 0.936713439521274 |
| chr11-63954281-63954782 | 8 | 0.925848010201269 |
| chr5-58333301-58333802 | 4 | 0.851527611226018 |
| chr5-131083130-131083631 | 8 | 0.938450641406009 |
| chr6-13147945-13148446 | 8 | 0.987837560228051 |
| chr2-79779402-79779903 | 8 | 0.995015706432784 |
| chr15-70432427-70432928 | 8 | 0.923898146137302 |
| chr7-72261354-72261855 | 2 | 0.905684947036112 |
| chr13-104106985-104107486 | 8 | 0.976272097289204 |
| chr6-93116544-93117045 | 8 | 0.919340254662774 |
| chr20-54319475-54319976 | 8 | 0.993493681128311 |
| chr7-126656050-126656551 | 8 | 0.992872786739094 |
| chr3-28912609-28913110 | 8 | 0.990524754790697 |
| chr9-6702855-6703356 | 8 | 0.870274361223829 |
| chr3-48750671-48751172 | 8 | 0.990120832112877 |

| | | |
|---|---|---|
| chr14-37710582-37711083 | 8 | 0.922390396588812 |
| chr4-21309903-21310404 | 8 | 0.957806307008467 |
| chr4-73025406-73025907 | 8 | 0.923842083678173 |
| chr3-41365098-41365599 | 8 | 0.838062511203761 |
| chr14-75405898-75406399 | 5 | 0.984351135457025 |
| chr1-224747084-224747585 | 4 | 0.911584289410683 |
| chr17-70322443-70322944 | 8 | 0.993995770073954 |
| chr1-41622388-41622889 | 8 | 0.97694979807945 |
| chr7-59293532-59294033 | 8 | 0.988770211062489 |
| chr13-85589626-85590127 | 3 | 0.836081360254323 |
| chr8-2078788-2079289 | 8 | 0.854643828082799 |
| chr13-67435808-67436309 | 8 | 0.887068882585919 |
| chr6-72607459-72607960 | 8 | 0.9902752663987 |
| chr2-12323073-12323574 | 8 | 0.950570780645061 |
| chr1-39707903-39708404 | 4 | 0.973022666890532 |
| chr1-221719684-221720185 | 8 | 0.980424033961082 |
| chr13-106489862-106490363 | 5 | 0.935937093713039 |
| chr4-41249825-41250326 | 3 | 0.996771608620903 |
| chr15-27609538-27610039 | 8 | 0.995006724630431 |
| chr14-42148100-42148601 | 8 | 0.912386508001668 |
| chr1-127712753-127713254 | 8 | 0.841175704629083 |
| chr9-73483724-73484225 | 8 | 0.82313135352471 |
| chr11-33830444-33830945 | 8 | 0.969873627419825 |
| chr4-24753215-24753716 | 8 | 0.88200064008739 |
| chr3-163715328-163715829 | 8 | 0.985680032938776 |
| chr3-131754681-131755182 | 8 | 0.889220147439051 |
| chr4-158430129-158430630 | 8 | 0.932757653300519 |
| chr16-16817913-16818414 | 6 | 0.9926749499583 |
| chr8-9359711-9360212 | 8 | 0.870759167956548 |
| chr8-45867352-45867853 | 8 | 0.977675580536389 |
| chr2-55218871-55219372 | 8 | 0.986725515585226 |
| chr2-130949579-130950080 | 8 | 0.988883884002719 |
| chr2-44745877-44746378 | 3 | 0.995071793379106 |
| chr9-45781110-45781611 | 8 | 0.986031291157864 |
| chrX-79574843-79575344 | 7 | 0.867336525173023 |
| chr1-158760853-158761354 | 4 | 0.86012654376318 |
| chr9-58896017-58896518 | 8 | 0.89708752649639 |
| chr9-18008619-18009120 | 8 | 0.885413102816186 |
| chr11-3708343-3708844 | 3 | 0.996645486746834 |
| chr8-115998714-115999215 | 8 | 0.851642706840159 |
| chr3-54526647-54527148 | 8 | 0.993872396017076 |
| chr14-98702560-98703061 | 8 | 0.99439655018738 |
| chr18-63079438-63079939 | 8 | 0.980190537248579 |

| | | |
|---|---|---|
| chr3-148728967-148729468 | 4 | 0.890341522203082 |
| chr5-2462507-2463008 | 4 | 0.941588191942702 |
| chr2-86399553-86400054 | 8 | 0.921664030177377 |
| chrX-51020959-51021460 | 8 | 0.991032803564843 |
| chr2-187473369-187473870 | 4 | 0.958134877942557 |
| chr5-4533265-4533766 | 8 | 0.883519191760941 |
| chr17-43339364-43339865 | 8 | 0.993493681128311 |
| chr2-232977698-232978199 | 8 | 0.994152721255473 |
| chr4-77641162-77641663 | 3 | 0.956102781107349 |
| chr6-38851038-38851539 | 8 | 0.993275976195114 |
| chr2-145609195-145609696 | 8 | 0.828409550419406 |
| chr8-4562137-4562638 | 8 | 0.870652165159746 |
| chr6-38383196-38383697 | 8 | 0.989842014074088 |
| chr17-22361240-22361741 | 4 | 0.948827354996298 |
| chr8-51627213-51627714 | 8 | 0.914879085489029 |
| chr7-32853764-32854265 | 5 | 0.893569440141584 |
| chr2-51120099-51120600 | 3 | 0.996309491415905 |
| chr13-86425285-86425786 | 8 | 0.957351995371159 |
| chr9-100335983-100336484 | 8 | 0.928534930706344 |
| chr11-85867448-85867949 | 8 | 0.912138982166989 |
| chr19-40159156-40159657 | 8 | 0.992981422347697 |
| chr5-50366811-50367312 | 5 | 0.963610093862764 |
| chr3-41807724-41808225 | 5 | 0.986327198145913 |
| chr2-156919868-156920369 | 8 | 0.81870901053018 |
| chr15-86213043-86213544 | 8 | 0.952435804459851 |
| chr10-26372159-26372660 | 5 | 0.983505931419811 |
| chr14-55380020-55380521 | 8 | 0.986158504341387 |
| chr13-88618786-88619287 | 8 | 0.927032537180684 |
| chr9-63585966-63586467 | 5 | 0.984288144408718 |
| chr11-41788120-41788621 | 2 | 0.930940790191457 |
| chrX-9962938-9963439 | 8 | 0.892020333659268 |
| chr16-20594326-20594827 | 8 | 0.845465591171773 |
| chr1-219154983-219155484 | 8 | 0.865011761685338 |
| chr20-47717914-47718415 | 3 | 0.99593553545041 |
| chr11-72324050-72324551 | 3 | 0.99593553545041 |
| chr13-31968386-31968887 | 8 | 0.894800072815243 |
| chr4-106451004-106451505 | 8 | 0.991285720766338 |
| chr14-87580178-87580679 | 8 | 0.896830238560491 |
| chr2-205103085-205103586 | 3 | 0.978607886272588 |
| chr8-12297604-12298105 | 8 | 0.950394598655922 |
| chr13-30741167-30741668 | 8 | 0.993493681128311 |
| chr17-26708272-26708773 | 8 | 0.912034081335636 |
| chr4-74694775-74695276 | 2 | 0.838138763840913 |

| | | |
|---|---|---|
| chr9-57244304-57244805 | 8 | 0.91497430777667 |
| chr1-100295485-100295986 | 3 | 0.996485585210058 |
| chr2-118537497-118537998 | 8 | 0.951740140278265 |
| chr2-157557351-157557852 | 8 | 0.947498176319213 |
| chr7-79029762-79030263 | 8 | 0.995030963010212 |
| chr2-23600639-23601140 | 8 | 0.934046744554531 |
| chr3-137959382-137959883 | 3 | 0.99593553545041 |
| chr3-107078769-107079270 | 3 | 0.910768501421184 |
| chr14-28682370-28682871 | 8 | 0.892807427130353 |
| chr1-108779309-108779810 | 4 | 0.940329830401745 |
| chr10-15664817-15665318 | 6 | 0.87055465908865 |
| chr10-25473994-25474495 | 8 | 0.985491634984061 |
| chr9-7143543-7144044 | 8 | 0.994742861956766 |
| chr20-21824282-21824783 | 5 | 0.971323329100829 |
| chr15-61979744-61980245 | 2 | 0.827006829285413 |
| chr1-122008857-122009358 | 4 | 0.936038932709753 |
| chr13-102052464-102052965 | 5 | 0.864691685374004 |
| chr4-143699524-143700025 | 8 | 0.945783406704929 |
| chr10-60194023-60194524 | 8 | 0.987972876173623 |
| chr1-146651077-146651578 | 8 | 0.995229933350768 |
| chr6-12172033-12172534 | 8 | 0.985359760099216 |
| chr8-86878438-86878939 | 8 | 0.897277836125346 |
| chr2-29332858-29333359 | 8 | 0.990113025195682 |
| chr2-24814113-24814614 | 8 | 0.877464458985931 |
| chr2-160007140-160007641 | 8 | 0.993493681128311 |
| chr18-49839636-49840137 | 8 | 0.966677787642736 |
| chr4-162777292-162777793 | 8 | 0.935898218376797 |
| chr6-97845069-97845570 | 5 | 0.975728330439376 |
| chr9-27804731-27805232 | 8 | 0.963251301492153 |
| chr20-16421126-16421627 | 8 | 0.846739808669687 |
| chr1-44594694-44595195 | 8 | 0.827326746952574 |
| chr14-68367376-68367877 | 5 | 0.970196580198354 |
| chr9-101327022-101327523 | 8 | 0.981761125690453 |
| chr4-74560714-74561215 | 8 | 0.993493681128311 |
| chr1-236611136-236611637 | 8 | 0.995240163995656 |
| chr16-33543220-33543721 | 8 | 0.990254874431356 |
| chr15-48317183-48317684 | 8 | 0.912690517525398 |
| chr13-36641335-36641836 | 8 | 0.908460301545773 |
| chr6-101373362-101373863 | 2 | 0.938051996420258 |
| chr1-45695783-45696284 | 8 | 0.869193053498221 |
| chr9-20452995-20453496 | 8 | 0.890040910358325 |
| chr6-137094161-137094662 | 8 | 0.995125031297208 |
| chr16-70638078-70638579 | 8 | 0.91163885699652 |

| | | |
|---|---|---|
| chr6-110507913-110508414 | 8 | 0.914113821279403 |
| chr7-81074024-81074525 | 3 | 0.996469105275485 |
| chr18-10335290-10335791 | 8 | 0.993493681128311 |
| chr16-35291670-35292171 | 2 | 0.922299193463973 |
| chr15-46546834-46547335 | 8 | 0.993493681128311 |
| chr19-42175299-42175800 | 5 | 0.985088217021137 |
| chr13-77169331-77169832 | 2 | 0.952109149253198 |
| chr19-12087943-12088444 | 8 | 0.959881747732898 |
| chr4-92977826-92978327 | 5 | 0.983919978840557 |
| chr12-45507480-45507981 | 8 | 0.947588663451582 |
| chr13-34862400-34862901 | 2 | 0.887157014029639 |
| chr1-43004059-43004560 | 8 | 0.965364623138114 |
| chr16-80981908-80982409 | 4 | 0.870132326111117 |
| chr13-38507954-38508455 | 8 | 0.993493681128311 |
| chr2-145818868-145819369 | 3 | 0.99593553545041 |
| chr15-27599506-27600007 | 8 | 0.993463721656645 |
| chr2-141508416-141508917 | 8 | 0.993493681128311 |
| chr13-76488082-76488583 | 3 | 0.994782371243408 |
| chr4-84496586-84497087 | 2 | 0.898189428766592 |
| chr6-55092054-55092555 | 8 | 0.986290950453957 |
| chr10-19055820-19056321 | 8 | 0.961563305480631 |
| chr7-54964662-54965163 | 8 | 0.969246782933024 |
| chr12-22837181-22837682 | 8 | 0.980561892242614 |
| chr7-128825593-128826094 | 5 | 0.990098465141472 |
| chr2-103407417-103407918 | 6 | 0.993203200233644 |
| chr6-117144865-117145366 | 8 | 0.992914204697333 |
| chr15-39125909-39126410 | 8 | 0.865696847917932 |
| chr17-24485773-24486274 | 8 | 0.982649002264084 |
| chr20-10224794-10225295 | 8 | 0.958427440764738 |
| chr8-99036778-99037279 | 8 | 0.957805505824525 |
| chr10-70322183-70322684 | 8 | 0.959355700108181 |
| chr13-93400800-93401301 | 8 | 0.994156118271119 |
| chr14-75341906-75342407 | 8 | 0.996400496480735 |
| chr5-141133663-141134164 | 8 | 0.954894831447466 |
| chr8-24841340-24841841 | 3 | 0.996550858119724 |
| chr4-54228406-54228907 | 8 | 0.989653527772219 |
| chr1-259557631-259558132 | 5 | 0.950792571945133 |
| chr4-39139788-39140289 | 6 | 0.936351697992068 |
| chr1-231037784-231038285 | 8 | 0.932444059587634 |
| chr14-77473207-77473708 | 5 | 0.852291938327507 |
| chr2-1357007-1357508 | 6 | 0.9912029564407 |
| chr1-53324797-53325298 | 2 | 0.83068375029827 |
| chr3-130034342-130034843 | 8 | 0.890083611714933 |

| | | |
|---|---|---|
| chr2-38524913-38525414 | 8 | 0.905013208982405 |
| chr1-247055322-247055823 | 4 | 0.949079812536643 |
| chr19-27381505-27382006 | 2 | 0.861703352633873 |
| chr5-113315074-113315575 | 8 | 0.983002884598746 |
| chr18-43376928-43377429 | 8 | 0.937836121454364 |
| chr3-20856761-20857262 | 4 | 0.838657357678357 |
| chr20-52381905-52382406 | 8 | 0.964581991166926 |
| chr18-32311448-32311949 | 8 | 0.967438607537423 |
| chr15-16431944-16432445 | 8 | 0.946599918601086 |
| chr18-62096663-62097164 | 8 | 0.876570137241184 |
| chr11-17184733-17185234 | 5 | 0.939719864650504 |
| chr3-126814161-126814662 | 8 | 0.977613169055244 |
| chr6-87218029-87218530 | 8 | 0.827780779081532 |
| chr6-65584068-65584569 | 2 | 0.842728687768133 |
| chr8-50833082-50833583 | 8 | 0.922501156017815 |
| chr17-48539351-48539852 | 8 | 0.933292604566843 |
| chr6-37975066-37975567 | 8 | 0.993493681128311 |
| chr17-40661852-40662353 | 8 | 0.979313559290869 |
| chr10-47860210-47860711 | 8 | 0.995246892385832 |
| chr3-139829622-139830123 | 5 | 0.983154044604575 |
| chr9-98781427-98781928 | 4 | 0.963151526417526 |
| chr14-42663292-42663793 | 5 | 0.861947834080635 |
| chr11-52471456-52471957 | 5 | 0.880908398249048 |
| chrX-74182199-74182700 | 8 | 0.994312822276918 |
| chrX-71682728-71683229 | 2 | 0.929836643334293 |
| chr4-161513751-161514252 | 4 | 0.969463199577865 |
| chrX-76364738-76365239 | 8 | 0.903319850805698 |
| chr13-78737244-78737745 | 8 | 0.993493681128311 |
| chr1-191890186-191890687 | 6 | 0.819498667305392 |
| chr16-43441460-43441961 | 8 | 0.993493681128311 |
| chr7-70797194-70797695 | 8 | 0.838805024526645 |
| chr9-73493488-73493989 | 5 | 0.957005638397425 |
| chr18-49268473-49268974 | 8 | 0.935440259639195 |
| chr18-48880208-48880709 | 8 | 0.989036123275624 |
| chr2-145957088-145957589 | 8 | 0.995156888937635 |
| chr11-56670538-56671039 | 8 | 0.903799567308459 |
| chr15-61000247-61000748 | 6 | 0.988256065925967 |
| chr1-178302054-178302555 | 8 | 0.993493681128311 |
| chr5-85209375-85209876 | 8 | 0.97249681599313 |
| chr3-161277980-161278481 | 5 | 0.836716922563708 |
| chr5-5557563-5558064 | 8 | 0.97419527686608 |
| chr4-26665738-26666239 | 5 | 0.976252392067405 |
| chr5-68423514-68424015 | 4 | 0.95972715586615 |

| | | |
|---|---|---|
| chr2-134427278-134427779 | 8 | 0.991189490178842 |
| chr5-29761441-29761942 | 6 | 0.991206925481043 |
| chr3-63166097-63166598 | 8 | 0.866539537279683 |
| chr7-123967265-123967766 | 6 | 0.997196405585353 |
| chr18-30863471-30863972 | 2 | 0.948857350305129 |
| chr10-35474032-35474533 | 3 | 0.99704927413992 |
| chr15-7964766-7965267 | 8 | 0.884246852596489 |
| chr18-57839501-57840002 | 8 | 0.982504869948168 |
| chr6-104016067-104016568 | 8 | 0.959296016156022 |
| chr6-101740155-101740656 | 5 | 0.983054733265381 |
| chr2-145180568-145181069 | 3 | 0.99593553545041 |
| chr4-103095960-103096461 | 8 | 0.960424572103912 |
| chr3-155982568-155983069 | 8 | 0.995170723636494 |
| chrX-103029057-103029558 | 6 | 0.996838829013077 |
| chr1-4535642-4536143 | 8 | 0.957626310834722 |
| chr6-38497020-38497521 | 8 | 0.9481755062532 |
| chr13-106156038-106156539 | 2 | 0.952065931350897 |
| chr3-166817049-166817550 | 8 | 0.885012863595992 |
| chr11-38855289-38855790 | 3 | 0.995689553571162 |
| chr2-81618292-81618793 | 3 | 0.960246678423689 |
| chr2-238312657-238313158 | 4 | 0.929176293510561 |
| chr3-159405613-159406114 | 8 | 0.855374884481532 |
| chr2-31643397-31643898 | 8 | 0.865693416967682 |
| chr16-72873803-72874304 | 4 | 0.821096710407996 |
| chr5-154680433-154680934 | 8 | 0.957524663355419 |
| chr18-8282164-8282665 | 8 | 0.991957349474945 |
| chr4-123683653-123684154 | 5 | 0.985459261244837 |
| chr2-239238909-239239410 | 8 | 0.964096895756778 |
| chr19-44554346-44554847 | 8 | 0.994735000386962 |
| chr11-18344563-18345064 | 4 | 0.972167864812415 |
| chr1-107955622-107956123 | 3 | 0.996775047660982 |
| chr4-123290616-123291117 | 8 | 0.83625738211359 |
| chr6-106275121-106275622 | 5 | 0.938216549397843 |
| chr5-58912761-58913262 | 5 | 0.982840170117634 |
| chr9-44891603-44892104 | 5 | 0.941810857980401 |
| chr4-64351825-64352326 | 5 | 0.959883596188341 |
| chr1-226738092-226738593 | 8 | 0.927791530083863 |
| chr4-40810481-40810982 | 8 | 0.979666286141895 |
| chr7-16462244-16462745 | 8 | 0.976666401570935 |
| chr2-236504459-236504960 | 8 | 0.993299781845005 |
| chrX-11905988-11906489 | 6 | 0.835221676005733 |
| chr5-81449416-81449917 | 8 | 0.932554318852143 |
| chr10-8962201-8962702 | 3 | 0.913320074311212 |

| | | |
|---|---|---|
| chr15-28635647-28636148 | 5 | 0.981453483845704 |
| chr11-23900100-23900601 | 8 | 0.837684953274511 |
| chr4-125367499-125368000 | 8 | 0.982552797858093 |
| chr17-51555758-51556259 | 8 | 0.930939936902242 |
| chr4-163875842-163876343 | 8 | 0.993650677690947 |
| chr3-76842164-76842665 | 8 | 0.835653252271163 |
| chr17-69798427-69798928 | 8 | 0.993493681128311 |
| chrX-102657038-102657539 | 3 | 0.99593553545041 |
| chr7-118389085-118389586 | 6 | 0.995602780938691 |
| chr13-7143824-7144325 | 4 | 0.952150749870741 |
| chr6-37355965-37356466 | 4 | 0.873444862733521 |
| chr8-111685341-111685842 | 8 | 0.882179842080656 |
| chr14-65877533-65878034 | 2 | 0.943045878537543 |
| chr14-42149107-42149608 | 8 | 0.822697440519898 |
| chr2-17980330-17980831 | 6 | 0.989124390782645 |
| chr2-118575202-118575703 | 8 | 0.984977287133672 |
| chr15-27615684-27616185 | 8 | 0.830761315800367 |
| chr10-24597743-24598244 | 4 | 0.961413220394097 |
| chr17-53856936-53857437 | 8 | 0.934279527935419 |
| chr6-35402492-35402993 | 8 | 0.971805712113175 |
| chr11-35985559-35986060 | 8 | 0.819232819151064 |
| chr13-94078018-94078519 | 4 | 0.969349359794206 |
| chr14-28521198-28521699 | 3 | 0.982139766051255 |
| chr10-43545419-43545920 | 8 | 0.994305261896508 |
| chr18-51527603-51528104 | 8 | 0.997901445013752 |
| chr4-67158495-67158996 | 8 | 0.848582015493614 |
| chr14-1413624-1414125 | 8 | 0.991377126029442 |
| chr5-30464916-30465417 | 8 | 0.992501866344583 |
| chr11-82214840-82215341 | 5 | 0.87329683052741 |
| chr11-6267690-6268191 | 8 | 0.973544873376274 |
| chr5-94597354-94597855 | 2 | 0.892063380262092 |
| chr13-49922215-49922716 | 8 | 0.834254239342624 |
| chr17-34274794-34275295 | 2 | 0.946286039098552 |
| chr6-21953834-21954335 | 8 | 0.925649374270014 |
| chr8-115893396-115893897 | 8 | 0.985786299301869 |
| chr15-37615404-37615905 | 8 | 0.994817145138023 |
| chr4-72865733-72866234 | 3 | 0.990155416547952 |
| chr18-77798065-77798566 | 2 | 0.91917637668174 |
| chr13-84158460-84158961 | 3 | 0.996060835675658 |
| chr15-88347186-88347687 | 8 | 0.993493681128311 |
| chr13-27329490-27329991 | 5 | 0.983103370939676 |
| chr2-67958719-67959220 | 4 | 0.965576449187428 |
| chr16-73849264-73849765 | 8 | 0.994674004458806 |

| | | |
|---|---|---|
| chr5-73026567-73027068 | 8 | 0.982702469528719 |
| chr5-109673795-109674296 | 8 | 0.957549738224773 |
| chr9-109338649-109339150 | 8 | 0.976284837632142 |
| chrX-152278125-152278626 | 5 | 0.969073219991104 |
| chr13-101610647-101611148 | 4 | 0.948582828548828 |
| chr8-1893936-1894437 | 8 | 0.956396476008257 |
| chr9-1891008-1891509 | 4 | 0.844446285304642 |
| chr5-89911556-89912057 | 6 | 0.993205399456801 |
| chr19-44691380-44691881 | 8 | 0.858727058010454 |
| chr4-69961813-69962314 | 5 | 0.991707796606258 |
| chr14-43763618-43764119 | 8 | 0.993493681128311 |
| chr3-161219402-161219903 | 4 | 0.83048980130554 |
| chr10-92196723-92197224 | 8 | 0.992436410248775 |
| chr2-99594028-99594529 | 3 | 0.997061148523527 |
| chr3-150920039-150920540 | 8 | 0.870067755865649 |
| chr4-96668825-96669326 | 8 | 0.983978218136538 |
| chr18-16499030-16499531 | 8 | 0.957937963013649 |
| chr1-192155629-192156130 | 8 | 0.909260874583653 |
| chr2-22749165-22749666 | 8 | 0.995246929644488 |
| chr10-29435813-29436314 | 3 | 0.845204133644928 |
| chr4-172450575-172451076 | 2 | 0.953620924203409 |
| chr19-52934314-52934815 | 8 | 0.907122041780664 |
| chr15-29844841-29845342 | 8 | 0.947455199020388 |
| chr2-190344361-190344862 | 5 | 0.981630019047871 |
| chr5-115201888-115202389 | 4 | 0.968632336103674 |
| chr8-56020066-56020567 | 5 | 0.990171605703717 |
| chr15-61541248-61541749 | 8 | 0.964945609743557 |
| chr14-46800879-46801380 | 5 | 0.992227484613492 |
| chr10-3512237-3512738 | 8 | 0.935808007071891 |
| chr1-45333230-45333731 | 8 | 0.988437404228312 |
| chr1-216201937-216202438 | 6 | 0.988600693604801 |
| chr2-247420873-247421374 | 2 | 0.909211985615622 |
| chrX-67409200-67409701 | 8 | 0.855472550512714 |
| chr18-51233658-51234159 | 8 | 0.993493681128311 |
| chr3-7511734-7512235 | 4 | 0.903477337004575 |
| chr3-13469006-13469507 | 8 | 0.989187276756855 |
| chr6-111632126-111632627 | 4 | 0.939534682633471 |
| chr3-13915263-13915764 | 8 | 0.991419068857337 |
| chr7-123941893-123942394 | 8 | 0.972916867603784 |
| chr5-15934501-15935002 | 2 | 0.857973388244785 |
| chr7-124243049-124243550 | 8 | 0.896889858291748 |
| chr15-79915270-79915771 | 8 | 0.893609232200673 |
| chr9-61373990-61374491 | 8 | 0.876577390303109 |

| | | |
|---|---|---|
| chr4-93070553-93071054 | 6 | 0.991206925481043 |
| chr18-55633049-55633550 | 8 | 0.915206584078825 |
| chr17-67148452-67148953 | 8 | 0.995066084825362 |
| chr1-80417215-80417716 | 8 | 0.984932365809099 |
| chr5-134284799-134285300 | 8 | 0.989255161876277 |
| chr19-56139024-56139525 | 8 | 0.9156025252398 |
| chr2-212090843-212091344 | 5 | 0.919583290581567 |
| chr8-72431139-72431640 | 5 | 0.87459074295622 |
| chr6-282690-283191 | 8 | 0.884717932974527 |
| chr2-77339969-77340470 | 8 | 0.935219984786696 |
| chr6-13767449-13767950 | 8 | 0.991577034879481 |
| chr7-47352279-47352780 | 5 | 0.991648821556596 |
| chr9-53077906-53078407 | 4 | 0.974506194796166 |
| chr14-79647285-79647786 | 8 | 0.865401051533596 |
| chr7-54663668-54664169 | 8 | 0.992667463552896 |
| chr11-23825714-23826215 | 3 | 0.840156208092743 |
| chr9-90194802-90195303 | 5 | 0.837520725762553 |
| chr3-75280511-75281012 | 2 | 0.927986544927769 |
| chr4-154838969-154839470 | 8 | 0.90126147901223 |
| chr9-91248534-91249035 | 8 | 0.941952934476476 |
| chr20-30066900-30067401 | 8 | 0.903656674742902 |
| chr20-31435382-31435883 | 5 | 0.966045530322553 |
| chr13-67009466-67009967 | 8 | 0.982999166293239 |
| chr11-2831433-2831934 | 2 | 0.879814733070421 |
| chr5-2934679-2935180 | 8 | 0.985833882903458 |
| chr3-90858756-90859257 | 8 | 0.913886369477643 |
| chr5-116617369-116617870 | 8 | 0.971646604575515 |
| chr8-55962352-55962853 | 8 | 0.993493681128311 |
| chr9-110919084-110919585 | 8 | 0.975381247150711 |
| chr11-47364071-47364572 | 8 | 0.988096710565118 |
| chrX-98706723-98707224 | 2 | 0.839948435238286 |
| chr1-132372170-132372671 | 8 | 0.983915972004766 |
| chr16-46409034-46409535 | 8 | 0.91263883043623 |
| chr8-68406322-68406823 | 2 | 0.96981886351952 |
| chr7-45262992-45263493 | 4 | 0.936676266286099 |
| chr8-15021444-15021945 | 5 | 0.986184973477738 |
| chr13-93110725-93111226 | 4 | 0.963727975253502 |
| chr1-183762793-183763294 | 8 | 0.989746763989658 |
| chr20-46413416-46413917 | 8 | 0.929318543669017 |
| chr9-44776692-44777193 | 8 | 0.856800043168943 |
| chr1-254693333-254693834 | 8 | 0.986930460452749 |
| chr8-15168948-15169449 | 5 | 0.950599967906255 |
| chr3-57954830-57955331 | 8 | 0.906949018085868 |

| | | |
|---|---|---|
| chr3-132729367-132729868 | 8 | 0.975847904792036 |
| chr3-96021137-96021638 | 6 | 0.993138516776475 |
| chr18-22728968-22729469 | 5 | 0.983919978840557 |
| chr10-16422129-16422630 | 8 | 0.868382392301498 |
| chr14-59718592-59719093 | 8 | 0.992843531949819 |
| chr18-1605262-1605763 | 5 | 0.984379488551694 |
| chr7-127921681-127922182 | 8 | 0.816500099282619 |
| chr1-41068343-41068844 | 3 | 0.996611071121329 |
| chr3-48159158-48159659 | 8 | 0.968299711647246 |
| chr6-140068516-140069017 | 8 | 0.981487937878676 |
| chr17-76955435-76955936 | 8 | 0.871423753701977 |
| chr14-60770895-60771396 | 5 | 0.982594145818312 |
| chr8-72282873-72283374 | 8 | 0.995242307374012 |
| chr8-27924654-27925155 | 8 | 0.994758817154017 |
| chr11-8911620-8912121 | 2 | 0.913724228423954 |
| chr9-25923612-25924113 | 4 | 0.960824967135992 |
| chr17-77341694-77342195 | 8 | 0.990718435291765 |
| chr11-70285922-70286423 | 8 | 0.920558589453654 |
| chrX-100788566-100789067 | 3 | 0.99593553545041 |
| chr1-156061727-156062228 | 2 | 0.887854984666121 |
| chr4-29346817-29347318 | 8 | 0.947514311078532 |
| chr3-64369662-64370163 | 8 | 0.980872087149056 |
| chrX-14236820-14237321 | 5 | 0.984637969830615 |
| chrX-28030677-28031178 | 8 | 0.993950346198923 |
| chr6-81254354-81254855 | 4 | 0.869626629122143 |
| chr2-239798678-239799179 | 3 | 0.832186285992914 |
| chrX-11443393-11443894 | 8 | 0.993493681128311 |
| chr1-132933424-132933925 | 8 | 0.990388163848343 |
| chr8-102157539-102158040 | 8 | 0.993493681128311 |
| chr20-38406311-38406812 | 8 | 0.925566528167146 |
| chr8-11685898-11686399 | 8 | 0.903751794070932 |
| chr17-81453208-81453709 | 6 | 0.992368834330493 |
| chr8-63547672-63548173 | 8 | 0.846947753698232 |
| chr16-21931949-21932450 | 5 | 0.991592227924182 |
| chr15-28508225-28508726 | 8 | 0.990253549824106 |
| chr16-32245724-32246225 | 2 | 0.951865305739957 |
| chr1-117360640-117361141 | 4 | 0.866721349530746 |
| chr14-57920614-57921115 | 8 | 0.843630438518823 |
| chrX-37754251-37754752 | 8 | 0.956813321866659 |
| chr2-58880405-58880906 | 6 | 0.968241176639702 |
| chr1-178725779-178726280 | 8 | 0.991798967780809 |
| chr1-115780536-115781037 | 8 | 0.993493681128311 |
| chr15-89267929-89268430 | 2 | 0.882930749978073 |

| | | |
|---|---|---|
| chr1-247116064-247116565 | 6 | 0.993090745253039 |
| chr20-52433360-52433861 | 8 | 0.993493681128311 |
| chr3-28163294-28163795 | 4 | 0.946451526468923 |
| chr11-68502913-68503414 | 8 | 0.883471171459667 |
| chr3-32140850-32141351 | 2 | 0.919841287662645 |
| chrX-60969512-60970013 | 3 | 0.921858582746028 |
| chr5-33027558-33028059 | 8 | 0.849575070255551 |
| chr7-132434405-132434906 | 8 | 0.829737039815891 |
| chr19-14741809-14742310 | 8 | 0.995186822476193 |
| chr1-235690428-235690929 | 2 | 0.919220446844096 |
| chr5-34320342-34320843 | 8 | 0.977288084630346 |
| chr7-21399543-21400044 | 8 | 0.96886654314751 |
| chr1-127790693-127791194 | 6 | 0.994019582609882 |
| chr1-21485853-21486354 | 2 | 0.946312777198037 |
| chrX-126904482-126904983 | 8 | 0.823236492370677 |
| chr2-113521528-113522029 | 5 | 0.852434936721769 |
| chr9-63462939-63463440 | 8 | 0.898733110084311 |
| chr2-173570256-173570757 | 8 | 0.954223289692765 |
| chr10-85970026-85970527 | 5 | 0.987619564596778 |
| chr1-170251085-170251586 | 8 | 0.989278829435063 |
| chr8-59368579-59369080 | 8 | 0.954707075682428 |
| chr6-17885770-17886271 | 2 | 0.831085324535785 |
| chr15-85997875-85998376 | 3 | 0.981278348940375 |
| chr18-77056096-77056597 | 5 | 0.944092434160148 |
| chr13-72122526-72123027 | 8 | 0.889790641058404 |
| chr7-377960-378461 | 8 | 0.963410300597643 |
| chr13-16573351-16573852 | 2 | 0.945303591814731 |
| chr14-41367655-41368156 | 5 | 0.973040379404325 |
| chr10-101783196-101783697 | 8 | 0.979433418287612 |
| chr10-21954151-21954652 | 2 | 0.902935431857142 |
| chr1-252940118-252940619 | 8 | 0.959258803323611 |
| chr6-47616569-47617070 | 5 | 0.9713609234759 |
| chr6-10175056-10175557 | 8 | 0.993766401990558 |
| chr3-103307822-103308323 | 8 | 0.85888923868486 |
| chr2-362432-362933 | 8 | 0.830499007946514 |
| chr5-118577144-118577645 | 8 | 0.983139928636128 |
| chr6-120944412-120944913 | 3 | 0.996678719028962 |
| chr2-164124955-164125456 | 2 | 0.957992879705029 |
| chr5-26736839-26737340 | 4 | 0.911613246233726 |
| chr17-15358094-15358595 | 8 | 0.988782207205614 |
| chr4-115096295-115096796 | 3 | 0.945508737520729 |
| chr2-104899433-104899934 | 8 | 0.984481072650659 |
| chr1-86066152-86066653 | 6 | 0.98376688483154 |

| | | |
|---|---|---|
| chr7-84070955-84071456 | 8 | 0.887151982211937 |
| chr8-71379539-71380040 | 8 | 0.930325887424794 |
| chr14-10482044-10482545 | 8 | 0.990458671070658 |
| chrX-78339202-78339703 | 4 | 0.903784972477483 |
| chr8-77621832-77622333 | 8 | 0.942242267278882 |
| chr17-63985720-63986221 | 5 | 0.845222984981792 |
| chr12-17415675-17416176 | 8 | 0.988278832165668 |
| chr12-32190126-32190627 | 8 | 0.9799263830371 |
| chr10-49133548-49134049 | 8 | 0.968355459848413 |
| chr3-69887650-69888151 | 8 | 0.994566169186611 |
| chr2-77096747-77097248 | 5 | 0.99002625841514 |
| chr12-5967018-5967519 | 8 | 0.920858663155681 |
| chr4-126342461-126342962 | 8 | 0.994959312834952 |
| chr5-22783422-22783923 | 8 | 0.977808595564189 |
| chr11-61131039-61131540 | 3 | 0.989833164675522 |
| chr1-242695031-242695532 | 8 | 0.993493681128311 |
| chr14-70865532-70866033 | 8 | 0.980264623688554 |
| chr17-50090841-50091342 | 8 | 0.873265517161971 |
| chr18-58965687-58966188 | 8 | 0.882027864300226 |
| chr9-50945145-50945646 | 8 | 0.971989363117766 |
| chr4-54012763-54013264 | 8 | 0.977805013311446 |
| chr13-18979511-18980012 | 8 | 0.978206606562519 |
| chr2-41146980-41147481 | 8 | 0.899755090376053 |
| chr7-120072561-120073062 | 8 | 0.99188991854134 |
| chr13-1680725-1681226 | 8 | 0.993493681128311 |
| chr2-240416283-240416784 | 8 | 0.973467443923701 |
| chr10-59878777-59879278 | 8 | 0.996941259510879 |
| chrX-107831545-107832046 | 8 | 0.992364388125674 |
| chr1-218663880-218664381 | 4 | 0.852512752667088 |
| chr1-83074672-83075173 | 2 | 0.835379758383052 |
| chr1-136298317-136298818 | 6 | 0.98414539809173 |
| chr4-17467456-17467957 | 8 | 0.9151383540851 |
| chr1-13444309-13444810 | 5 | 0.974140920390446 |
| chr18-21982259-21982760 | 2 | 0.934515321616918 |
| chr1-5084033-5084534 | 2 | 0.893042029609269 |
| chr5-155367563-155368064 | 2 | 0.911508713546957 |
| chr15-26699844-26700345 | 8 | 0.968838859164462 |
| chr19-37873277-37873778 | 8 | 0.84297395662237 |
| chrX-97442908-97443409 | 5 | 0.982248206482803 |
| chr4-61179327-61179828 | 8 | 0.935904489049317 |
| chr20-21178179-21178680 | 5 | 0.836119918256984 |
| chr1-214150996-214151497 | 8 | 0.985157281116639 |
| chr1-241282487-241282988 | 8 | 0.980822119769576 |

| | | |
|---|---|---|
| chr1-180422721-180423222 | 2 | 0.897586226554632 |
| chr1-105969142-105969643 | 3 | 0.903464994090447 |
| chr18-80296052-80296553 | 8 | 0.971035347640006 |
| chr2-233861298-233861799 | 8 | 0.924290035970327 |
| chr2-22390703-22391204 | 8 | 0.981445448671116 |
| chr10-23695221-23695722 | 5 | 0.988599320596797 |
| chr2-130157913-130158414 | 8 | 0.962980908242879 |
| chr2-214473203-214473704 | 8 | 0.908970451207639 |
| chr14-70359851-70360352 | 5 | 0.916692658094469 |
| chr13-36644536-36645037 | 8 | 0.965603960278244 |
| chr13-105307011-105307512 | 8 | 0.883659410868499 |
| chr7-40198989-40199490 | 8 | 0.993369498590065 |
| chr2-244605161-244605662 | 5 | 0.845417111766451 |
| chr10-22367886-22368387 | 8 | 0.963230531438159 |
| chr8-83726847-83727348 | 8 | 0.871584546067416 |
| chr5-86833272-86833773 | 8 | 0.864215632606162 |
| chr14-4281346-4281847 | 3 | 0.977583364417773 |
| chr5-29492120-29492621 | 8 | 0.984420015359435 |
| chr1-153717441-153717942 | 8 | 0.969059204264374 |
| chr7-21173796-21174297 | 8 | 0.947197891872483 |
| chr19-21870782-21871283 | 5 | 0.990114758959076 |
| chr10-74413232-74413733 | 8 | 0.853991504218094 |
| chr18-21504436-21504937 | 8 | 0.877424928862597 |
| chr10-62050805-62051306 | 8 | 0.989565543756811 |
| chr14-35482682-35483183 | 8 | 0.943071489207876 |
| chr9-92113032-92113533 | 8 | 0.913353858319463 |
| chr7-114991149-114991650 | 8 | 0.985439216853705 |
| chr2-144242581-144243082 | 8 | 0.977505647849692 |
| chr7-128892965-128893466 | 3 | 0.984144553925945 |
| chr8-109164798-109165299 | 7 | 0.844220048873617 |
| chr14-53999050-53999551 | 4 | 0.857127996015633 |
| chr1-80920613-80921114 | 8 | 0.984082198308599 |
| chr6-136517639-136518140 | 3 | 0.99593553545041 |
| chr9-42816376-42816877 | 8 | 0.889782450532572 |
| chr16-61150915-61151416 | 3 | 0.97415598861245 |
| chr1-106940480-106940981 | 8 | 0.911584346826012 |
| chr17-52154160-52154661 | 8 | 0.831955225703413 |
| chr13-87514264-87514765 | 4 | 0.936867840341869 |
| chr2-191904988-191905489 | 5 | 0.966098206023066 |
| chr1-222820519-222821020 | 5 | 0.984926741859751 |
| chr10-33325574-33326075 | 8 | 0.887795214714735 |
| chr3-92698094-92698595 | 4 | 0.966051235436852 |
| chr10-57722135-57722636 | 4 | 0.824171806448423 |

| | | |
|---|---|---|
| chr3-20106226-20106727 | 3 | 0.99593553545041 |
| chr9-109442088-109442589 | 4 | 0.8629170764058 |
| chr2-10732110-10732611 | 2 | 0.838988588274641 |
| chr14-77583686-77584187 | 2 | 0.846146336422362 |
| chr15-100488504-100489005 | 5 | 0.988904654088043 |
| chr14-37182901-37183402 | 3 | 0.910506768722639 |
| chr4-149501821-149502322 | 5 | 0.937852246967196 |
| chr10-82412071-82412572 | 8 | 0.902309720364212 |
| chr18-30058497-30058998 | 8 | 0.841890127382253 |
| chr5-69499303-69499804 | 8 | 0.994525566468023 |
| chr11-24580049-24580550 | 8 | 0.862493112829485 |
| chr8-76702707-76703208 | 1 | 0.924625937969885 |
| chr20-30472795-30473296 | 5 | 0.976900491245174 |
| chr2-144078379-144078880 | 8 | 0.938694231438668 |
| chr11-61105961-61106462 | 4 | 0.924286048891529 |
| chr15-89465184-89465685 | 8 | 0.990825724049001 |
| chr11-49332732-49333233 | 8 | 0.916685091408599 |
| chr7-71868440-71868941 | 8 | 0.993493681128311 |
| chr7-113424158-113424659 | 8 | 0.941612250325106 |
| chr11-3868285-3868786 | 5 | 0.997315245166056 |
| chr7-117716820-117717321 | 6 | 0.982608127817147 |
| chr17-17309932-17310433 | 5 | 0.985165259944053 |
| chr13-26678792-26679293 | 3 | 0.995813947972373 |
| chr14-1693560-1694061 | 8 | 0.979989256530584 |
| chr13-21629559-21630060 | 6 | 0.991206925481043 |
| chr8-43426900-43427401 | 5 | 0.953943159318512 |
| chr3-146745982-146746483 | 8 | 0.839741792417189 |
| chr15-10216036-10216537 | 8 | 0.991375287507165 |
| chr15-62604469-62604970 | 8 | 0.976027332493749 |
| chr4-48132265-48132766 | 3 | 0.996813868930661 |
| chr20-28648198-28648699 | 4 | 0.959009005586619 |
| chr11-81136721-81137222 | 8 | 0.995223473432878 |
| chr13-100141039-100141540 | 3 | 0.996411171665916 |
| chr18-70861688-70862189 | 8 | 0.882693193232056 |
| chr5-109160373-109160874 | 8 | 0.824361364022089 |
| chr9-101358459-101358960 | 8 | 0.934751657679517 |
| chr2-145814920-145815421 | 8 | 0.838688573122002 |
| chr5-50200995-50201496 | 5 | 0.963417998112561 |
| chr1-204747036-204747537 | 8 | 0.9952225572195 |
| chr1-211863424-211863925 | 8 | 0.961837777867204 |
| chr1-51958572-51959073 | 8 | 0.903121318920115 |
| chr18-62161555-62162056 | 8 | 0.991693707077956 |
| chr16-21633757-21634258 | 4 | 0.908078543306173 |

| | | |
|---|---|---|
| chr4-60430102-60430603 | 8 | 0.973633695863248 |
| chr19-42666002-42666503 | 5 | 0.992443289459217 |
| chr11-38014227-38014728 | 8 | 0.993493681128311 |
| chr14-15641428-15641929 | 5 | 0.865748528601282 |
| chrX-131705928-131706429 | 8 | 0.995232771087614 |
| chr8-22728694-22729195 | 3 | 0.996063734280174 |
| chr9-24719374-24719875 | 3 | 0.969499328206638 |
| chr12-23814788-23815289 | 8 | 0.991874669170842 |
| chrX-14940083-14940584 | 8 | 0.995244421099686 |
| chr7-63643010-63643511 | 8 | 0.961574923606779 |
| chr5-26336478-26336979 | 5 | 0.981275884075226 |
| chr10-20370469-20370970 | 8 | 0.963485451110699 |
| chr6-69240162-69240663 | 8 | 0.974746549868513 |
| chr9-68180724-68181225 | 8 | 0.982856274208102 |
| chr18-19784819-19785320 | 8 | 0.993493681128311 |
| chr7-15542056-15542557 | 5 | 0.986828270197735 |
| chr6-11131340-11131841 | 8 | 0.961565847684106 |
| chr11-73375906-73376407 | 8 | 0.96232688879118 |
| chr3-95146035-95146536 | 8 | 0.826914115719034 |
| chr16-46978895-46979396 | 8 | 0.975581818816158 |
| chr10-62655893-62656394 | 3 | 0.842693462780315 |
| chr10-68353970-68354471 | 8 | 0.987483159006094 |
| chr1-8747171-8747672 | 8 | 0.981531681230432 |
| chr15-15642913-15643414 | 8 | 0.947998491620628 |
| chr3-126195854-126196355 | 6 | 0.990398939533156 |
| chr16-32860325-32860826 | 8 | 0.991167850126737 |
| chr2-169506417-169506918 | 8 | 0.94445162636224 |
| chr17-31215400-31215901 | 7 | 0.907792738372986 |
| chr1-96926201-96926702 | 8 | 0.924508793816151 |
| chr1-193390578-193391079 | 8 | 0.858924897410028 |
| chr5-85629441-85629942 | 8 | 0.988049812744182 |
| chr1-171572463-171572964 | 8 | 0.935481461738583 |
| chr14-90304463-90304964 | 8 | 0.950298954174729 |
| chr6-84610991-84611492 | 5 | 0.987999730645623 |
| chr4-152886843-152887344 | 8 | 0.986877832766871 |
| chr8-30233788-30234289 | 8 | 0.979275129478128 |
| chr8-86001973-86002474 | 2 | 0.945118576157448 |
| chr1-58168313-58168814 | 8 | 0.977607825010042 |
| chr5-15373931-15374432 | 8 | 0.95626449785948 |
| chr2-210142952-210143453 | 8 | 0.886290383101679 |
| chr11-54765315-54765816 | 8 | 0.992169368652202 |
| chr4-10123283-10123784 | 4 | 0.899769920471852 |
| chr15-20959807-20960308 | 8 | 0.987528075871519 |

| | | |
|---|---|---|
| chr12-1548535-1549036 | 8 | 0.897598103968133 |
| chr5-106827003-106827504 | 3 | 0.996424275847704 |
| chr6-44247141-44247642 | 8 | 0.980338692625114 |
| chr1-181266210-181266711 | 8 | 0.939438535708213 |
| chr9-48423451-48423952 | 4 | 0.895039 |
| chr3-105753274-105753775 | 8 | 0.996047745722891 |
| chr11-67834819-67835320 | 2 | 0.904275530938368 |
| chr2-31352006-31352507 | 8 | 0.823763011772386 |
| chr4-50982095-50982596 | 8 | 0.972717326140042 |
| chr4-86196862-86197363 | 8 | 0.856527636165406 |
| chr10-77560943-77561444 | 6 | 0.990810019734339 |
| chr17-28935997-28936498 | 8 | 0.988500992938497 |
| chr5-9309044-9309545 | 8 | 0.990436696615206 |
| chr10-63596670-63597171 | 8 | 0.827046031409898 |
| chr2-178092269-178092770 | 8 | 0.920611109586828 |
| chr10-5344715-5345216 | 8 | 0.987096061394998 |
| chr6-131385634-131386135 | 8 | 0.820673508156328 |
| chr17-1277979-1278480 | 8 | 0.965380563368546 |
| chr6-11229161-11229662 | 8 | 0.993131667947583 |
| chr2-218399564-218400065 | 4 | 0.826506077664007 |
| chr1-28395273-28395774 | 8 | 0.974282227319828 |
| chr2-146073918-146074419 | 3 | 0.996897078639686 |
| chr4-137480781-137481282 | 5 | 0.918897898321722 |
| chr16-83389220-83389721 | 4 | 0.850336369872125 |
| chr19-37220511-37221012 | 8 | 0.901507368457302 |
| chr2-156250910-156251411 | 3 | 0.99593553545041 |
| chr16-33139688-33140189 | 8 | 0.993493681128311 |
| chr1-28056798-28057299 | 8 | 0.970310729277909 |
| chr1-20583672-20584173 | 8 | 0.841405513151173 |
| chr13-76794206-76794707 | 4 | 0.845088207317927 |
| chr15-74950984-74951485 | 8 | 0.81740815249395 |
| chr14-10045794-10046295 | 8 | 0.965058219955098 |
| chr4-72767980-72768481 | 8 | 0.91816082808645 |
| chr19-47918588-47919089 | 8 | 0.896681301554861 |
| chr19-47200636-47201137 | 5 | 0.983919978840557 |
| chr18-74324339-74324840 | 8 | 0.918514531355154 |
| chr1-168200885-168201386 | 6 | 0.992013381455057 |
| chr4-137447821-137448322 | 2 | 0.888113723822417 |
| chr11-74925164-74925665 | 8 | 0.897849959262234 |
| chr5-12676085-12676586 | 8 | 0.970555854006168 |
| chr7-7933580-7934081 | 2 | 0.89274629743538 |
| chr14-38358849-38359350 | 8 | 0.993493681128311 |
| chr11-78065716-78066217 | 8 | 0.852846473778877 |

| | | |
|---|---|---|
| chr11-8629197-8629698 | 8 | 0.976649873681972 |
| chr10-62020653-62021154 | 4 | 0.956174725417591 |
| chr16-74081253-74081754 | 8 | 0.969203800645491 |
| chr9-67597713-67598214 | 8 | 0.899824639580169 |
| chr15-2387792-2388293 | 8 | 0.985994927256259 |
| chr3-125883430-125883931 | 8 | 0.993493681128311 |
| chr15-38066232-38066733 | 5 | 0.989037921149977 |
| chr1-215482265-215482766 | 2 | 0.82698736191867 |
| chr1-99861341-99861842 | 8 | 0.993493681128311 |
| chr13-28738330-28738831 | 8 | 0.993493681128311 |
| chr11-28542068-28542569 | 8 | 0.994319586381897 |
| chr16-1441035-1441536 | 8 | 0.906714335113281 |
| chr17-71147828-71148329 | 8 | 0.922757390981949 |
| chr2-36865375-36865876 | 8 | 0.956657218260518 |
| chr2-38707504-38708005 | 8 | 0.836980019118053 |
| chr1-258715141-258715642 | 8 | 0.994988551252575 |
| chr4-89960841-89961342 | 8 | 0.951589523365706 |
| chr15-70339658-70340159 | 3 | 0.99593553545041 |
| chr19-4514607-4515108 | 5 | 0.984102673972219 |
| chr1-26243627-26244128 | 8 | 0.918281308038734 |
| chr18-56247427-56247928 | 4 | 0.950251737198429 |
| chr14-42911529-42912030 | 5 | 0.98628186412573 |
| chr18-48538062-48538563 | 8 | 0.908526456786742 |
| chr10-35288198-35288699 | 8 | 0.962302638238048 |
| chr4-166705507-166706008 | 8 | 0.98685982903136 |
| chr4-131060967-131061468 | 8 | 0.858234413770871 |
| chr4-91263837-91264338 | 8 | 0.96497259710845 |
| chr10-67674024-67674525 | 8 | 0.971516173242954 |
| chr5-94856259-94856760 | 6 | 0.926959642487253 |
| chr13-16490443-16490944 | 5 | 0.990872022641459 |
| chr18-55298531-55299032 | 4 | 0.942020565997943 |
| chr15-20868480-20868981 | 8 | 0.993493681128311 |
| chr15-91705365-91705866 | 6 | 0.989915069243535 |
| chr6-51313512-51314013 | 8 | 0.902900383324629 |
| chr14-61905549-61906050 | 5 | 0.984407465827089 |
| chr8-31605687-31606188 | 2 | 0.825050680970524 |
| chr20-46901832-46902333 | 8 | 0.970450483638584 |
| chr6-48037253-48037754 | 8 | 0.928901312708897 |
| chr20-28478803-28479304 | 8 | 0.942713785190509 |
| chr17-23568549-23569050 | 3 | 0.987292020402564 |
| chr11-81554750-81555251 | 2 | 0.857134977561126 |
| chr11-19790294-19790795 | 8 | 0.993493681128311 |
| chr1-191009325-191009826 | 8 | 0.858761688991017 |

| | | |
|---|---|---|
| chr13-37457472-37457973 | 4 | 0.945037047775778 |
| chr18-31668561-31669062 | 8 | 0.993493681128311 |
| chr5-71167814-71168315 | 6 | 0.966888797425089 |
| chr1-40175922-40176423 | 5 | 0.997379548738742 |
| chr5-62745537-62746038 | 8 | 0.935753685116018 |
| chr13-42286113-42286614 | 5 | 0.985184529216181 |
| chr17-48312183-48312684 | 8 | 0.995600749811455 |
| chr20-163501-164002 | 4 | 0.890605956288854 |
| chr5-135358936-135359437 | 8 | 0.959043297246387 |
| chr11-27795717-27796218 | 5 | 0.90174821596689 |
| chr1-185465017-185465518 | 8 | 0.93371889992486 |
| chr2-240246668-240247169 | 3 | 0.930652434088442 |
| chr8-109003742-109004243 | 5 | 0.987358871574276 |
| chr9-25525113-25525614 | 5 | 0.968720179217277 |
| chr17-27968484-27968985 | 2 | 0.870609992000707 |
| chr6-36621504-36622005 | 8 | 0.993493681128311 |
| chr19-48521611-48522112 | 5 | 0.895612141116342 |
| chr7-120648300-120648801 | 8 | 0.878157919615206 |
| chr14-65363493-65363994 | 4 | 0.913625813661527 |
| chr6-100244407-100244908 | 5 | 0.987955082953025 |
| chr13-86928741-86929242 | 8 | 0.982000450416856 |
| chr12-30175252-30175753 | 8 | 0.990985201139131 |
| chr13-81863822-81864323 | 2 | 0.954927962389896 |
| chr14-32041779-32042280 | 5 | 0.966585606195729 |
| chr15-53568171-53568672 | 8 | 0.868314649708927 |
| chr9-69880249-69880750 | 6 | 0.991206925481043 |
| chr14-85292615-85293116 | 8 | 0.815187249189707 |
| chr6-10245221-10245722 | 8 | 0.993493681128311 |
| chr19-32845525-32846026 | 4 | 0.936366737569799 |
| chr14-94099484-94099985 | 2 | 0.953331665511972 |
| chr1-33534562-33535063 | 8 | 0.965173215620408 |
| chr3-3080663-3081164 | 8 | 0.989304599523209 |
| chr6-61061668-61062169 | 4 | 0.864234695696331 |
| chr16-245135-245636 | 8 | 0.81854291367877 |
| chr7-120909259-120909760 | 8 | 0.968080764227002 |
| chr17-83157345-83157846 | 8 | 0.863666127881632 |
| chr9-47183103-47183604 | 4 | 0.971894629494366 |
| chr2-109167740-109168241 | 2 | 0.954670921722255 |
| chr19-3382358-3382859 | 8 | 0.895193136928077 |
| chr16-57869270-57869771 | 5 | 0.974538829200325 |
| chr13-4352903-4353404 | 3 | 0.995947762143795 |
| chr9-47969021-47969522 | 8 | 0.990037800637993 |
| chr4-25163267-25163768 | 8 | 0.995244906164227 |

| | | |
|---|---|---|
| chr7-27493846-27494347 | 8 | 0.993493681128311 |
| chr20-34306078-34306579 | 8 | 0.906372196989749 |
| chr5-80163547-80164048 | 3 | 0.99695673311666 |
| chr18-79540216-79540717 | 6 | 0.993225434010423 |
| chr13-52662191-52662692 | 8 | 0.904441154255449 |
| chr4-85581038-85581539 | 6 | 0.979833442857279 |
| chr13-101039957-101040458 | 8 | 0.880475816814103 |
| chr5-24309866-24310367 | 8 | 0.92649122927912 |
| chr19-16995551-16996052 | 8 | 0.993493681128311 |
| chr6-76970908-76971409 | 8 | 0.847486815186739 |
| chr14-97138330-97138831 | 8 | 0.981765837585107 |
| chr1-219393030-219393531 | 4 | 0.89130984180555 |
| chr11-27246891-27247392 | 5 | 0.920931867769427 |
| chr17-71132955-71133456 | 8 | 0.957405980176203 |
| chr6-136446419-136446920 | 8 | 0.909963722063231 |
| chr11-20138234-20138735 | 6 | 0.98746357445481 |
| chr8-77103054-77103555 | 8 | 0.992462761365299 |
| chr5-134687687-134688188 | 8 | 0.951311093156733 |
| chr16-72786245-72786746 | 8 | 0.954163999353915 |
| chr2-217281822-217282323 | 3 | 0.989576253862945 |
| chr3-132956098-132956599 | 8 | 0.994999812523373 |
| chr2-39193065-39193566 | 8 | 0.994239217842914 |
| chr9-43721846-43722347 | 8 | 0.984388923305876 |
| chr3-166301415-166301916 | 6 | 0.985293806752441 |
| chr2-60666717-60667218 | 8 | 0.993493681128311 |
| chr14-32558815-32559316 | 4 | 0.957987175371272 |
| chr1-149271043-149271544 | 3 | 0.855492737763497 |
| chr4-112170947-112171448 | 3 | 0.996862704513852 |
| chr16-29999067-29999568 | 4 | 0.836648917814187 |
| chr13-85035923-85036424 | 2 | 0.874186836048912 |
| chr20-47220075-47220576 | 2 | 0.913486516256522 |
| chr16-69818296-69818797 | 5 | 0.815951337388559 |
| chr13-82702002-82702503 | 5 | 0.890340444219311 |
| chr6-60221046-60221547 | 3 | 0.995142354471272 |
| chr6-47597103-47597604 | 8 | 0.995188531072963 |
| chr5-100677712-100678213 | 8 | 0.990813246351662 |
| chr3-119102750-119103251 | 4 | 0.95273230053469 |
| chr5-132139381-132139882 | 8 | 0.907245772880897 |
| chr16-63279651-63280152 | 3 | 0.984345891073135 |
| chr20-26002967-26003468 | 5 | 0.842221352101162 |
| chr1-211952443-211952944 | 8 | 0.995103607303339 |
| chr3-153897236-153897737 | 8 | 0.824574345202823 |
| chr5-140480205-140480706 | 2 | 0.964826019612594 |

| | | |
|---|---|---|
| chr2-124104981-124105482 | 2 | 0.818230038640111 |
| chr7-124309669-124310170 | 8 | 0.951548977897989 |
| chr1-230884073-230884574 | 8 | 0.984120447596585 |
| chr18-69497914-69498415 | 5 | 0.977619064890986 |
| chrX-41725830-41726331 | 6 | 0.982227196204876 |
| chr16-57658076-57658577 | 3 | 0.998760760385039 |
| chr6-109070870-109071371 | 4 | 0.944074018330671 |
| chr13-21170515-21171016 | 8 | 0.949261864606493 |
| chr3-135505326-135505827 | 8 | 0.933744512411312 |
| chr11-69158040-69158541 | 8 | 0.861813347435259 |
| chr6-44740539-44741040 | 8 | 0.984608150428301 |
| chr1-107181641-107182142 | 8 | 0.931938507271437 |
| chr9-24033664-24034165 | 2 | 0.831188402318947 |
| chr6-48713515-48714016 | 8 | 0.911281707748947 |
| chr19-19773892-19774393 | 5 | 0.962988791674124 |
| chr9-10714911-10715412 | 8 | 0.896628858896529 |
| chr14-18963553-18964054 | 8 | 0.984889388977053 |
| chr2-97839419-97839920 | 3 | 0.993527288499859 |
| chr10-92995719-92996220 | 2 | 0.925726928462602 |
| chr5-3823458-3823959 | 6 | 0.991006593894086 |
| chr9-20028121-20028622 | 4 | 0.908597178928057 |
| chr13-96424805-96425306 | 2 | 0.948269250839525 |
| chr14-52448833-52449334 | 8 | 0.873767305043176 |
| chr11-17583439-17583940 | 8 | 0.852706645562289 |
| chr4-171489882-171490383 | 8 | 0.88365577510369 |
| chr9-74316566-74317067 | 8 | 0.976746938078182 |
| chr5-163904255-163904756 | 8 | 0.884149885561044 |
| chr1-124787784-124788285 | 8 | 0.962428532442376 |
| chr1-125894543-125895044 | 8 | 0.898799369897916 |
| chr1-186472338-186472839 | 8 | 0.977768689069198 |
| chr18-14663955-14664456 | 8 | 0.992923562964106 |
| chr1-54469681-54470182 | 3 | 0.99689770654266 |
| chr10-51821477-51821978 | 8 | 0.993493681128311 |
| chr9-81051721-81052222 | 3 | 0.863471856574317 |
| chr16-13538764-13539265 | 5 | 0.914464537982053 |
| chr4-123145960-123146461 | 2 | 0.921258407185644 |
| chr3-43907200-43907701 | 3 | 0.971227084010861 |
| chr16-57928516-57929017 | 8 | 0.956579869720724 |
| chr1-17660476-17660977 | 3 | 0.997113211105914 |
| chr1-227009705-227010206 | 2 | 0.912754868188012 |
| chr8-62595965-62596466 | 8 | 0.938132390199625 |
| chr4-53622823-53623324 | 8 | 0.993493681128311 |
| chr14-22221562-22222063 | 3 | 0.994027022474555 |

| | | |
|---|---|---|
| chr1-222223086-222223587 | 6 | 0.935713155231411 |
| chr3-122921516-122922017 | 5 | 0.835263522512289 |
| chr15-12213812-12214313 | 2 | 0.950444262981793 |
| chr15-96089213-96089714 | 8 | 0.860093865190881 |
| chr4-141244429-141244930 | 8 | 0.856708072230463 |
| chr9-77929946-77930447 | 8 | 0.993493681128311 |
| chr2-38041742-38042243 | 3 | 0.996679156206257 |
| chr10-85792344-85792845 | 4 | 0.936293921270848 |
| chr8-65812024-65812525 | 8 | 0.817295490332597 |
| chr5-121151372-121151873 | 8 | 0.993440739651749 |
| chr10-49418949-49419450 | 8 | 0.990629497698724 |
| chr4-172093090-172093591 | 8 | 0.993493681128311 |
| chr2-176997922-176998423 | 6 | 0.992801181100416 |
| chrX-1264753-1265254 | 8 | 0.972705667877846 |
| chr2-20435707-20436208 | 8 | 0.993493681128311 |
| chr8-117841336-117841837 | 8 | 0.950229173675354 |
| chr17-77616805-77617306 | 8 | 0.995046874110491 |
| chr1-136530246-136530747 | 8 | 0.889630479575534 |
| chr19-10640762-10641263 | 8 | 0.993493681128311 |
| chr4-16799852-16800353 | 4 | 0.854641352753205 |
| chr1-242326715-242327216 | 8 | 0.987621158318613 |
| chr4-125536504-125537005 | 8 | 0.932870385913966 |
| chr4-88009915-88010416 | 8 | 0.990919541411095 |
| chr4-5597441-5597942 | 8 | 0.993209713023881 |
| chr10-61944534-61945035 | 8 | 0.858623785116914 |
| chr2-90709761-90710262 | 8 | 0.980488690764143 |
| chr3-43656333-43656834 | 4 | 0.824683902182466 |
| chr4-35917798-35918299 | 5 | 0.957194855208252 |
| chr14-37555468-37555969 | 3 | 0.993753619691921 |
| chr9-3185770-3186271 | 8 | 0.823425271166468 |
| chr10-48624890-48625391 | 4 | 0.968442238650765 |
| chr10-103988020-103988521 | 5 | 0.98047687035089 |
| chr4-34770263-34770764 | 8 | 0.870235918908996 |
| chr2-109694007-109694508 | 8 | 0.875726644150854 |
| chr5-146472411-146472912 | 3 | 0.897183925700248 |
| chr1-212674053-212674554 | 2 | 0.832151843303225 |
| chr1-43418554-43419055 | 8 | 0.993493681128311 |
| chr14-63031815-63032316 | 5 | 0.983919978840557 |
| chr5-80532677-80533178 | 8 | 0.993493681128311 |
| chr1-28719600-28720101 | 2 | 0.927321771880439 |
| chr14-66241624-66242125 | 8 | 0.913017107849978 |
| chr18-62661112-62661613 | 8 | 0.987489679192223 |
| chr14-8076663-8077164 | 5 | 0.987816952269942 |

| | | |
|---|---|---|
| chr2-192345124-192345625 | 8 | 0.864243937973836 |
| chr2-102200454-102200955 | 8 | 0.993493681128311 |
| chr3-126746505-126747006 | 5 | 0.861429652963465 |
| chr13-66939725-66940226 | 5 | 0.974189967319131 |
| chr14-50537932-50538433 | 3 | 0.996876295485722 |
| chr14-69146181-69146682 | 8 | 0.959507971649049 |
| chrX-28023782-28024283 | 8 | 0.993493681128311 |
| chr13-16588762-16589263 | 8 | 0.973291773744345 |
| chr9-62727673-62728174 | 5 | 0.991075951537062 |
| chr8-119289756-119290257 | 8 | 0.94299508682537 |
| chr12-13287100-13287601 | 8 | 0.960756415154465 |
| chr12-10642953-10643454 | 8 | 0.991705920922553 |
| chr6-96014858-96015359 | 5 | 0.964162539258159 |
| chr7-17269105-17269606 | 4 | 0.886112215531337 |
| chr5-158766113-158766614 | 5 | 0.884346399352881 |
| chr19-39257040-39257541 | 8 | 0.943093742588293 |
| chr14-9915134-9915635 | 4 | 0.848963269503871 |
| chr16-10112804-10113305 | 4 | 0.862643403630646 |
| chr19-9908169-9908670 | 4 | 0.945576886989155 |
| chr20-38057274-38057775 | 8 | 0.846134897415863 |
| chr1-27933692-27934193 | 8 | 0.91653732463462 |
| chr19-19992981-19993482 | 6 | 0.982532324907013 |
| chr4-46155525-46156026 | 8 | 0.943781484750964 |
| chr9-20764486-20764987 | 8 | 0.826147712755857 |
| chr9-32604602-32605103 | 3 | 0.99593553545041 |
| chr4-131221491-131221992 | 8 | 0.986791345768077 |
| chr2-576201-576702 | 8 | 0.855692938817137 |
| chrX-106103439-106103940 | 8 | 0.993493681128311 |
| chr5-135754140-135754641 | 8 | 0.946892104431799 |
| chr14-74021358-74021859 | 8 | 0.986966526431052 |
| chr14-100530201-100530702 | 3 | 0.907869172111357 |
| chr18-39578867-39579368 | 5 | 0.984171172873593 |
| chr5-71580436-71580937 | 8 | 0.978787023722922 |
| chr4-155297145-155297646 | 8 | 0.96554976892212 |
| chr16-81419878-81420379 | 8 | 0.986779976117133 |
| chr16-37849887-37850388 | 8 | 0.839926358416285 |
| chr19-30676881-30677382 | 8 | 0.988107684417426 |
| chr9-50330680-50331181 | 5 | 0.964905459198054 |
| chr9-98469576-98470077 | 5 | 0.954140913015271 |
| chr2-152651985-152652486 | 8 | 0.939955153641885 |
| chr5-9002946-9003447 | 4 | 0.868490683079588 |
| chr5-90854101-90854602 | 3 | 0.99593553545041 |
| chr15-13394858-13395359 | 8 | 0.930101280613078 |

| | | |
|---|---|---|
| chr6-140900603-140901104 | 8 | 0.972256378987402 |
| chr7-7966257-7966758 | 8 | 0.967287890084351 |
| chr17-19057184-19057685 | 8 | 0.93074730306878 |
| chr17-28307205-28307706 | 8 | 0.94587342119757 |
| chr1-21186827-21187328 | 8 | 0.936706837631061 |
| chr7-110880126-110880627 | 8 | 0.884162330605652 |
| chr8-12831124-12831625 | 6 | 0.991694581827495 |
| chr1-6717436-6717937 | 5 | 0.950964733201672 |
| chr4-72553197-72553698 | 8 | 0.961344129657067 |
| chrX-143856559-143857060 | 3 | 0.99593553545041 |
| chr6-107378736-107379237 | 8 | 0.989545480298285 |
| chr3-121337811-121338312 | 8 | 0.979718698011316 |
| chr9-49184742-49185243 | 8 | 0.977667441246592 |
| chr13-77904867-77905368 | 8 | 0.884329183795625 |
| chr19-23991024-23991525 | 8 | 0.927596729815243 |
| chr5-129392492-129392993 | 8 | 0.928993286143756 |
| chr16-40882415-40882916 | 6 | 0.991956996908927 |
| chr9-65723523-65724024 | 8 | 0.938050266238568 |
| chr8-4137976-4138477 | 8 | 0.993493681128311 |
| chr14-91886165-91886666 | 8 | 0.993493681128311 |
| chr14-92189155-92189656 | 5 | 0.983267694403368 |
| chrX-4313616-4314117 | 4 | 0.953602782633932 |
| chr5-8798539-8799040 | 8 | 0.936908428553166 |
| chr16-67666767-67667268 | 2 | 0.942096791561934 |
| chr14-22394173-22394674 | 3 | 0.932312905351396 |
| chr10-19550507-19551008 | 8 | 0.940770927082448 |
| chr1-104607646-104608147 | 8 | 0.968495514341372 |
| chr17-28464904-28465405 | 8 | 0.835765819735784 |
| chr5-34899108-34899609 | 3 | 0.987983787292858 |
| chr14-76615240-76615741 | 8 | 0.888147295406647 |
| chr4-115213368-115213869 | 8 | 0.980411424525491 |
| chr11-84752773-84753274 | 8 | 0.993493681128311 |
| chr8-101893158-101893659 | 8 | 0.993493681128311 |
| chr7-27341623-27342124 | 2 | 0.946318296933773 |
| chr1-33584949-33585450 | 7 | 0.839119093913876 |
| chr4-103510368-103510869 | 8 | 0.898184889481181 |
| chr8-77996060-77996561 | 8 | 0.995237546085082 |
| chr18-64229711-64230212 | 5 | 0.988311181097935 |
| chr14-20359024-20359525 | 6 | 0.991505595161678 |
| chr1-205961552-205962053 | 8 | 0.988621187517149 |
| chr11-53849328-53849829 | 8 | 0.872519532861159 |
| chr7-83796087-83796588 | 8 | 0.988294176489961 |
| chr11-71567841-71568342 | 5 | 0.842882383561281 |

| | | |
|---|---|---|
| chr5-114130744-114131245 | 8 | 0.935846685572196 |
| chr2-73063774-73064275 | 3 | 0.964739056708248 |
| chr4-35048864-35049365 | 2 | 0.91993458669193 |
| chrX-64942650-64943151 | 4 | 0.945110696161547 |
| chr19-33664678-33665179 | 8 | 0.993493681128311 |
| chr13-85494101-85494602 | 8 | 0.825427713827813 |
| chr7-121157500-121158001 | 2 | 0.968165620152929 |
| chr4-103739527-103740028 | 8 | 0.946435764348313 |
| chr1-203550546-203551047 | 8 | 0.993147520800234 |
| chr3-142489159-142489660 | 6 | 0.866835276430645 |
| chr10-8741341-8741842 | 8 | 0.968885124372923 |
| chr3-77097087-77097588 | 8 | 0.973153929821932 |
| chr9-61135073-61135574 | 8 | 0.97492746643844 |
| chr5-123124672-123125173 | 3 | 0.821892069041021 |
| chr6-55129849-55130350 | 6 | 0.989180329738271 |
| chr1-41427411-41427912 | 8 | 0.994507949371037 |
| chr3-19642715-19643216 | 8 | 0.943467453768975 |
| chr6-42167451-42167952 | 8 | 0.995147168837396 |
| chr12-11075011-11075512 | 8 | 0.834609648789347 |
| chr20-41406747-41407248 | 5 | 0.975314662594929 |
| chr1-5421335-5421836 | 8 | 0.978144339878986 |
| chr11-70782722-70783223 | 8 | 0.973236515197985 |
| chr17-48126188-48126689 | 4 | 0.903544075341896 |
| chr8-91297857-91298358 | 8 | 0.985899154964731 |
| chr3-109910670-109911171 | 8 | 0.989247618618533 |
| chr2-115556949-115557450 | 8 | 0.993493681128311 |
| chr9-87968862-87969363 | 5 | 0.986445615994135 |
| chr1-5341799-5342300 | 3 | 0.99593553545041 |
| chr15-75256968-75257469 | 8 | 0.880635416917446 |
| chr3-130310724-130311225 | 8 | 0.821964883090125 |
| chr13-62118300-62118801 | 8 | 0.886661949113959 |
| chr18-32118945-32119446 | 8 | 0.979470401489864 |
| chr11-57711069-57711570 | 3 | 0.996600890583421 |
| chr14-80229649-80230150 | 8 | 0.882809569127696 |
| chr8-12285085-12285586 | 3 | 0.894393593412135 |
| chr6-47549932-47550433 | 5 | 0.939160899079722 |
| chr1-28069809-28070310 | 8 | 0.993493681128311 |
| chr5-101872024-101872525 | 8 | 0.831555937408033 |
| chr18-44587905-44588406 | 2 | 0.952851627182224 |
| chr2-204552320-204552821 | 8 | 0.975455268306984 |
| chr11-21213814-21214315 | 8 | 0.898348862901156 |
| chr1-194110871-194111372 | 4 | 0.964787916977443 |
| chr8-34693450-34693951 | 6 | 0.9883998327532 |

| | | |
|---|---|---|
| chr7-40789623-40790124 | 8 | 0.936216544842436 |
| chr1-174579378-174579879 | 8 | 0.823106094251828 |
| chr6-58105706-58106207 | 3 | 0.99593553545041 |
| chr13-64391683-64392184 | 6 | 0.996039536364788 |
| chr3-11740360-11740861 | 8 | 0.886880673117906 |
| chr6-39671530-39672031 | 3 | 0.896308864472345 |
| chr17-78999084-78999585 | 8 | 0.993493681128311 |
| chr14-40341649-40342150 | 3 | 0.99655793164552 |
| chr7-5354696-5355197 | 3 | 0.99593553545041 |
| chrX-17153631-17154132 | 5 | 0.876577837227983 |
| chr4-28366209-28366710 | 8 | 0.828010908252052 |
| chr2-173197910-173198411 | 8 | 0.946789566128173 |
| chr2-42292123-42292624 | 5 | 0.934596384736782 |
| chr4-77842415-77842916 | 3 | 0.947099321194878 |
| chr9-37712220-37712721 | 8 | 0.921609320509686 |
| chr15-45506873-45507374 | 8 | 0.984518982030802 |
| chr2-3070835-3071336 | 8 | 0.964129788231073 |
| chr9-47921695-47922196 | 8 | 0.966457420600969 |
| chr2-165217923-165218424 | 2 | 0.859763510865749 |
| chr18-71503234-71503735 | 8 | 0.990707251652272 |
| chr4-14857249-14857750 | 6 | 0.968875394012006 |
| chr9-54403667-54404168 | 8 | 0.907332279985824 |
| chr18-71582344-71582845 | 8 | 0.913632321713851 |
| chr13-3301361-3301862 | 5 | 0.891901659985945 |
| chr14-72781422-72781923 | 8 | 0.866752470003713 |
| chr8-123613919-123614420 | 8 | 0.98928201860798 |
| chrX-29457477-29457978 | 2 | 0.846952796955674 |
| chr18-29537206-29537707 | 8 | 0.863661235896148 |
| chr11-56312590-56313091 | 8 | 0.990886369047724 |
| chr5-135087923-135088424 | 8 | 0.834204102296327 |
| chr1-175519182-175519683 | 8 | 0.973582441906804 |
| chr17-9368031-9368532 | 8 | 0.926037127606633 |
| chr4-117002224-117002725 | 4 | 0.820561079576132 |
| chr12-25738172-25738673 | 4 | 0.929542136962074 |
| chr18-77010270-77010771 | 5 | 0.921953163973173 |
| chrX-148211724-148212225 | 8 | 0.993493681128311 |
| chr18-68136252-68136753 | 5 | 0.983691919728677 |
| chr7-47671866-47672367 | 8 | 0.982116930957381 |
| chr1-83017289-83017790 | 8 | 0.995207519079835 |
| chr10-66045391-66045892 | 2 | 0.819151309394524 |
| chr16-62339395-62339896 | 5 | 0.965160342397569 |
| chr2-153021441-153021942 | 7 | 0.883730274301254 |
| chr5-154853728-154854229 | 8 | 0.993161577860684 |

| | | |
|---|---|---|
| chr13-64255788-64256289 | 8 | 0.995146034563486 |
| chrX-42401385-42401886 | 8 | 0.93832819122155 |
| chr2-210109866-210110367 | 8 | 0.888022936692348 |
| chr4-162241134-162241635 | 8 | 0.9131315935088 |
| chr19-28172495-28172996 | 8 | 0.917475600582628 |
| chr2-70250894-70251395 | 6 | 0.991206925481043 |
| chr2-128193309-128193810 | 8 | 0.985522267443283 |
| chr1-69245727-69246228 | 8 | 0.995035593329136 |
| chr1-49321826-49322327 | 8 | 0.988480868949242 |
| chr8-7776082-7776583 | 6 | 0.996142781715748 |
| chr8-98321884-98322385 | 8 | 0.993493681128311 |
| chr17-6563683-6564184 | 5 | 0.971343946644105 |
| chr16-76247775-76248276 | 8 | 0.957631778708132 |
| chr1-29559315-29559816 | 3 | 0.860249129458553 |
| chr2-53328306-53328807 | 8 | 0.994609703392564 |
| chrX-93781849-93782350 | 6 | 0.991206925481043 |
| chr2-89831079-89831580 | 6 | 0.990159016463835 |
| chr6-140403185-140403686 | 8 | 0.919539741564742 |
| chr10-96292932-96293433 | 6 | 0.991206925481043 |
| chr4-135165112-135165613 | 8 | 0.994640183229032 |
| chr14-56369092-56369593 | 8 | 0.989735847560747 |
| chr2-169018639-169019140 | 5 | 0.980632503928967 |
| chr15-10170673-10171174 | 8 | 0.825834156175479 |
| chr7-65442553-65443054 | 5 | 0.990462376237491 |
| chr12-28060991-28061492 | 5 | 0.994805393726649 |
| chr9-22270168-22270669 | 8 | 0.992458582484008 |
| chr12-29579440-29579941 | 8 | 0.85131046239366 |
| chr2-66593015-66593516 | 8 | 0.893540993894364 |
| chr2-224396433-224396934 | 8 | 0.868541204786813 |
| chr16-38470692-38471193 | 8 | 0.994914447873055 |
| chrX-14472236-14472737 | 8 | 0.993169200416582 |
| chr5-56427103-56427604 | 2 | 0.919628673604809 |
| chr6-41386135-41386636 | 8 | 0.968258552771968 |
| chr18-52851781-52852282 | 8 | 0.960832510306686 |
| chr1-171652162-171652663 | 8 | 0.91544995752746 |
| chr17-29301870-29302371 | 8 | 0.993493681128311 |
| chr18-5343210-5343711 | 8 | 0.952952723841568 |
| chr6-57876088-57876589 | 8 | 0.968482723068585 |
| chr3-36595147-36595648 | 8 | 0.88736786626255 |
| chr15-74372743-74373244 | 8 | 0.902008767367807 |
| chr16-2710874-2711375 | 5 | 0.985944199363384 |
| chr14-92122799-92123300 | 3 | 0.996049364343712 |
| chr18-67125354-67125855 | 8 | 0.993493681128311 |

| | | |
|---|---|---|
| chr14-22088480-22088981 | 8 | 0.951014208386934 |
| chr20-8610112-8610613 | 8 | 0.975333125922317 |
| chr3-47602698-47603199 | 2 | 0.923691212329323 |
| chr9-76659445-76659946 | 8 | 0.934838019345195 |
| chrX-49599895-49600396 | 8 | 0.991945302935704 |
| chr11-69805597-69806098 | 5 | 0.991311211149829 |
| chr16-76263798-76264299 | 8 | 0.948387327744991 |
| chr6-59821181-59821682 | 3 | 0.996843404958555 |
| chr4-141598341-141598842 | 8 | 0.939083722381048 |
| chr7-99291949-99292450 | 8 | 0.991673732921907 |
| chr4-144825772-144826273 | 8 | 0.814823154145977 |
| chr1-133423287-133423788 | 8 | 0.989771804289965 |
| chr3-152228304-152228805 | 8 | 0.981580425251802 |
| chr6-114307744-114308245 | 4 | 0.879054921622921 |
| chr4-55620202-55620703 | 8 | 0.987996480966568 |
| chr10-70925011-70925512 | 2 | 0.839500691491622 |
| chr5-114396011-114396512 | 3 | 0.995740078372208 |
| chrX-36108357-36108858 | 3 | 0.995682555285556 |
| chr5-74766543-74767044 | 3 | 0.955280351785869 |
| chr14-29387165-29387666 | 6 | 0.869022365286058 |
| chr5-112972667-112973168 | 8 | 0.994706091712634 |
| chr1-91468369-91468870 | 4 | 0.853800526190194 |
| chr7-55582864-55583365 | 2 | 0.826880275371394 |
| chr2-87038884-87039385 | 8 | 0.86643516290357 |
| chr2-28145020-28145521 | 8 | 0.891789691370568 |
| chr18-51913743-51914244 | 8 | 0.915007109219825 |
| chrX-35496832-35497333 | 8 | 0.966673305392185 |
| chr15-91475953-91476454 | 6 | 0.869529995803348 |
| chr5-109059611-109060112 | 6 | 0.991206925481043 |
| chr2-23362658-23363159 | 8 | 0.99522049144512 |
| chr15-12384920-12385421 | 8 | 0.990590233531422 |
| chr5-35974734-35975235 | 2 | 0.832211274984041 |
| chr4-148839441-148839942 | 8 | 0.920347977690561 |
| chr5-144759747-144760248 | 3 | 0.965086926342767 |
| chr16-21237122-21237623 | 8 | 0.928951354086938 |
| chr20-46060628-46061129 | 5 | 0.986126315701142 |
| chr6-45563203-45563704 | 2 | 0.910466599273502 |
| chr4-117587254-117587755 | 8 | 0.987580401839554 |
| chr5-13485675-13486176 | 5 | 0.962181922519799 |
| chr2-205821816-205822317 | 4 | 0.954199608376007 |
| chr4-53238061-53238562 | 5 | 0.983919978840557 |
| chrX-16678954-16679455 | 6 | 0.991958894365489 |
| chr17-76849321-76849822 | 8 | 0.965875098239707 |

| | | |
|---|---|---|
| chr19-44666932-44667433 | 5 | 0.941730751660085 |
| chr19-15370852-15371353 | 5 | 0.987002438579752 |
| chr2-34291629-34292130 | 8 | 0.989760729440672 |
| chr17-19414768-19415269 | 5 | 0.981186438474786 |
| chr8-81143795-81144296 | 5 | 0.895025942170369 |
| chr3-45362491-45362992 | 5 | 0.970941617201187 |
| chr4-176572129-176572630 | 8 | 0.994070159459317 |
| chr8-38160666-38161167 | 8 | 0.844066604027171 |
| chr3-25176653-25177154 | 8 | 0.913031490221779 |
| chr14-73050788-73051289 | 5 | 0.854526178049817 |
| chr3-100329976-100330477 | 8 | 0.959223928942107 |
| chr4-178541776-178542277 | 4 | 0.968818989647277 |
| chr15-82667000-82667501 | 8 | 0.884584818595793 |
| chr4-9429881-9430382 | 8 | 0.941965353190069 |
| chr16-51534647-51535148 | 8 | 0.980393708637057 |
| chr1-143042771-143043272 | 8 | 0.993277992900797 |
| chr13-70109605-70110106 | 8 | 0.984559999192428 |
| chr13-67928907-67929408 | 8 | 0.945578340780983 |
| chr2-116210698-116211199 | 3 | 0.99593553545041 |
| chr1-247462606-247463107 | 8 | 0.91513550492176 |
| chr16-14693282-14693783 | 8 | 0.936470770580308 |
| chr17-63657593-63658094 | 8 | 0.994500052200718 |
| chr1-239174787-239175288 | 3 | 0.951658989459245 |
| chr3-110837351-110837852 | 8 | 0.990936056689123 |
| chr7-100596819-100597320 | 8 | 0.977190868657191 |
| chr4-32301424-32301925 | 8 | 0.904543006134088 |
| chr14-32076607-32077108 | 8 | 0.858132881063716 |
| chr1-146963001-146963502 | 4 | 0.940399035925682 |
| chr2-238829027-238829528 | 8 | 0.91418075225274 |
| chr5-148763846-148764347 | 8 | 0.930417836130851 |
| chr16-76546498-76546999 | 8 | 0.880736000291748 |
| chr14-42393954-42394455 | 8 | 0.979797063804961 |
| chr6-13943300-13943801 | 8 | 0.914102851386615 |
| chrX-54608905-54609406 | 5 | 0.986777460434848 |
| chr1-225055655-225056156 | 8 | 0.933525306718001 |
| chr11-43375720-43376221 | 4 | 0.899237118857099 |
| chr16-80061078-80061579 | 8 | 0.95999701733107 |
| chr15-88140008-88140509 | 3 | 0.892832663394907 |
| chr16-56917583-56918084 | 8 | 0.905884787573695 |
| chrX-65393487-65393988 | 3 | 0.991828022189999 |
| chr1-189642603-189643104 | 8 | 0.870654114894606 |
| chr17-60798175-60798676 | 4 | 0.847735919723699 |
| chr6-2511889-2512390 | 8 | 0.986023618889664 |

| | | |
|---|---|---|
| chr10-93630158-93630659 | 8 | 0.949716532376256 |
| chr12-17561666-17562167 | 6 | 0.945120680033821 |
| chr8-17784394-17784895 | 6 | 0.991206925481043 |
| chr10-90228615-90229116 | 5 | 0.988555210033766 |
| chr1-199461668-199462169 | 8 | 0.974880774407411 |
| chr8-115033060-115033561 | 4 | 0.970243868359488 |
| chr6-46227177-46227678 | 6 | 0.992913412867226 |
| chr8-29908636-29909137 | 2 | 0.825639177180101 |
| chr8-94451343-94451844 | 3 | 0.997397722971115 |
| chr5-8917001-8917502 | 8 | 0.958660184883628 |
| chr18-73952998-73953499 | 4 | 0.973123257064614 |
| chr10-10014426-10014927 | 4 | 0.922158487156604 |
| chr6-38665691-38666192 | 8 | 0.893673054668957 |
| chr4-136296455-136296956 | 8 | 0.835196648739784 |
| chr4-84634233-84634734 | 8 | 0.9717546881823 |
| chr3-80716413-80716914 | 8 | 0.95374090760049 |
| chr13-101656624-101657125 | 8 | 0.982050072628535 |
| chr3-55915707-55916208 | 8 | 0.869584765277452 |
| chr4-90055321-90055822 | 8 | 0.852064122182399 |
| chr11-16051401-16051902 | 8 | 0.903956749465518 |
| chr18-54454921-54455422 | 8 | 0.860834191068907 |
| chr15-49124483-49124984 | 8 | 0.945838303538339 |
| chr8-26804311-26804812 | 8 | 0.992981303858412 |
| chr12-28759273-28759774 | 5 | 0.989755730703127 |
| chr10-97443870-97444371 | 8 | 0.875355138355801 |
| chr15-15460909-15461410 | 8 | 0.936428153448315 |
| chr3-48748654-48749155 | 8 | 0.94491708839723 |
| chr9-74630699-74631200 | 5 | 0.98402539006307 |
| chr1-206228250-206228751 | 8 | 0.963450150701575 |
| chr18-69504580-69505081 | 8 | 0.831082934269058 |
| chr13-37481638-37482139 | 5 | 0.907045191953409 |
| chr3-21443413-21443914 | 8 | 0.983721546283381 |
| chr18-62637188-62637689 | 5 | 0.980848466091965 |
| chr4-150241300-150241801 | 8 | 0.990515477704645 |
| chr1-236144739-236145240 | 8 | 0.988609384598972 |
| chr7-105044932-105045433 | 8 | 0.981328168039732 |
| chr1-228457451-228457952 | 8 | 0.960878969485037 |
| chr1-132635114-132635615 | 8 | 0.885920230634088 |
| chr13-32053139-32053640 | 8 | 0.962021885668255 |
| chr8-4350139-4350640 | 8 | 0.993493681128311 |
| chr2-206206266-206206767 | 6 | 0.992230940860871 |
| chr1-41259029-41259530 | 8 | 0.966530333247552 |
| chr8-76946428-76946929 | 4 | 0.972439558908341 |

| | | |
|---|---|---|
| chr5-123545690-123546191 | 3 | 0.997080183847033 |
| chr20-38190062-38190563 | 8 | 0.993493681128311 |
| chr2-59309601-59310102 | 8 | 0.990553583905545 |
| chr10-24292167-24292668 | 8 | 0.982318800444689 |
| chr1-25830415-25830916 | 3 | 0.993021865221865 |
| chr19-34420196-34420697 | 8 | 0.953737939161358 |
| chr8-90257260-90257761 | 4 | 0.934185387302101 |
| chr7-8894906-8895407 | 8 | 0.98589038565185 |
| chr11-55912211-55912712 | 8 | 0.994106697304172 |
| chr4-78756207-78756708 | 8 | 0.995206965575063 |
| chr6-118562622-118563123 | 5 | 0.978171480878669 |
| chr14-63750131-63750632 | 8 | 0.952005224670611 |
| chr5-36109136-36109637 | 3 | 0.99593553545041 |
| chr2-189412192-189412693 | 8 | 0.976798813499269 |
| chr4-13825794-13826295 | 8 | 0.990393351410162 |
| chr7-87837813-87838314 | 2 | 0.865409072400684 |
| chr3-53829292-53829793 | 8 | 0.992553855365972 |
| chr2-188370441-188370942 | 8 | 0.990311600390029 |
| chr2-32862155-32862656 | 8 | 0.987739864215209 |
| chr3-109155498-109155999 | 4 | 0.85582281496901 |
| chr10-29502760-29503261 | 8 | 0.970924381762691 |
| chr18-50582912-50583413 | 8 | 0.991442355645143 |
| chr3-114099007-114099508 | 8 | 0.942289802955673 |
| chr4-135665654-135666155 | 8 | 0.952160521032617 |
| chr10-39916732-39917233 | 6 | 0.993620311696463 |
| chr4-6445858-6446359 | 8 | 0.992607698891893 |
| chr5-114270392-114270893 | 2 | 0.843266996964401 |
| chr3-160105531-160106032 | 5 | 0.974891568774817 |
| chr18-79736236-79736737 | 4 | 0.900435111819016 |
| chr3-38693776-38694277 | 8 | 0.993493681128311 |
| chr2-167569248-167569749 | 3 | 0.995557515375718 |
| chr14-77163869-77164370 | 5 | 0.963786562842065 |
| chr2-247692662-247693163 | 6 | 0.992775468950514 |
| chr1-3941911-3942412 | 5 | 0.978500279027929 |
| chr17-9692533-9693034 | 8 | 0.991089441689485 |
| chr13-85588062-85588563 | 8 | 0.911649345219233 |
| chr18-52311545-52312046 | 8 | 0.856547876747214 |
| chrX-12387685-12388186 | 2 | 0.867268153206718 |
| chr3-167782263-167782764 | 8 | 0.968709670965735 |
| chr2-214631019-214631520 | 8 | 0.952522355468164 |
| chr2-246189785-246190286 | 8 | 0.961473521199801 |
| chr19-19404735-19405236 | 3 | 0.933537026399096 |
| chr11-71881100-71881601 | 8 | 0.822994741721951 |

| | | |
|---|---|---|
| chr18-11429992-11430493 | 8 | 0.995240693336561 |
| chr6-137460119-137460620 | 8 | 0.92338541781983 |
| chr2-102274194-102274695 | 7 | 0.817040690007907 |
| chr4-176320061-176320562 | 3 | 0.996871375875055 |
| chr3-21878045-21878546 | 8 | 0.912042057103036 |
| chr5-66637938-66638439 | 3 | 0.979362540612683 |
| chrX-148370900-148371401 | 3 | 0.993497615584239 |
| chr9-107102749-107103250 | 5 | 0.991324779207301 |
| chr19-5020019-5020520 | 8 | 0.840667823021491 |
| chr1-242877505-242878006 | 4 | 0.957678662376031 |
| chr3-127867012-127867513 | 4 | 0.906259898786908 |
| chr16-155155-155656 | 8 | 0.988924814944943 |
| chr2-236419043-236419544 | 8 | 0.992462396734171 |
| chr4-15397174-15397675 | 3 | 0.996007483511832 |
| chr13-20994590-20995091 | 2 | 0.840901802210051 |
| chr5-88190539-88191040 | 4 | 0.958023594317841 |
| chr3-30064619-30065120 | 6 | 0.981956957979166 |
| chr13-83950789-83951290 | 8 | 0.839894570389116 |
| chr14-52013410-52013911 | 8 | 0.925697467438628 |
| chr2-10112305-10112806 | 8 | 0.899593001707478 |
| chr8-28549683-28550184 | 8 | 0.9951414094303 |
| chr1-149242089-149242590 | 8 | 0.990738497683269 |
| chr20-11028714-11029215 | 3 | 0.973754179540028 |
| chr11-39912603-39913104 | 8 | 0.887805587757329 |
| chr14-26261968-26262469 | 8 | 0.98292787193607 |
| chr16-64138999-64139500 | 8 | 0.92889155776098 |
| chr1-109542451-109542952 | 8 | 0.993493681128311 |
| chr4-34418013-34418514 | 4 | 0.952427092775276 |
| chr9-23192103-23192604 | 4 | 0.973066116616585 |
| chr19-26295812-26296313 | 4 | 0.892487314069853 |
| chr6-96113157-96113658 | 8 | 0.980876901649335 |
| chr13-98818515-98819016 | 8 | 0.919410410218886 |
| chr7-37950805-37951306 | 8 | 0.895446500436136 |
| chr8-50835550-50836051 | 8 | 0.99519737988234 |
| chr7-112318021-112318522 | 8 | 0.951958150308544 |
| chr2-66563947-66564448 | 3 | 0.99593553545041 |
| chrX-22648489-22648990 | 5 | 0.983919978840557 |
| chr13-32037667-32038168 | 8 | 0.82137951421754 |
| chr1-213498599-213499100 | 8 | 0.890953228761755 |
| chr6-137937896-137938397 | 8 | 0.996286409775243 |
| chr14-12610724-12611225 | 8 | 0.943583788457294 |
| chr6-26057972-26058473 | 8 | 0.991580025999627 |
| chr4-5755583-5756084 | 8 | 0.980226284873215 |

| | | |
|---|---|---|
| chr18-19552838-19553339 | 3 | 0.973568544672772 |
| chr1-121137111-121137612 | 8 | 0.936818987281929 |
| chr17-26549259-26549760 | 8 | 0.949990574514554 |
| chr2-230934172-230934673 | 8 | 0.838214287376669 |
| chr19-7502622-7503123 | 5 | 0.938133581126541 |
| chr20-259646-260147 | 1 | 0.906447245155031 |
| chr10-18679110-18679611 | 8 | 0.960502596797966 |
| chr5-11950473-11950974 | 4 | 0.952970859545559 |
| chr8-62744084-62744585 | 2 | 0.831441530403203 |
| chr12-6192507-6193008 | 8 | 0.831689882000548 |
| chr3-152219100-152219601 | 8 | 0.959477887960899 |
| chr1-167807555-167808056 | 3 | 0.996341542140004 |
| chr12-27510918-27511419 | 8 | 0.987282392662589 |
| chr2-213761334-213761835 | 3 | 0.9965186869524 |
| chr4-160195899-160196400 | 8 | 0.994251726558087 |
| chr17-70162563-70163064 | 8 | 0.980620650646941 |
| chr15-48012300-48012801 | 8 | 0.904196287511019 |
| chr20-10309793-10310294 | 5 | 0.849128838919177 |
| chr9-81803738-81804239 | 8 | 0.871466944303946 |
| chr9-18344561-18345062 | 3 | 0.996849201090957 |
| chr1-100274079-100274580 | 4 | 0.941803539919164 |
| chr11-31522576-31523077 | 4 | 0.926461702008649 |
| chr9-21832613-21833114 | 8 | 0.915257769887422 |
| chr20-36574297-36574798 | 5 | 0.986360593326061 |
| chr7-49566290-49566791 | 3 | 0.987734349763719 |
| chr7-93312089-93312590 | 8 | 0.98942047601309 |
| chr1-106660951-106661452 | 8 | 0.992991391361525 |
| chr6-114743441-114743942 | 3 | 0.99593553545041 |
| chrX-130818880-130819381 | 5 | 0.832329116101055 |
| chrX-110623314-110623815 | 8 | 0.814764720396916 |
| chr20-45193668-45194169 | 5 | 0.984962289121057 |
| chr1-23292704-23293205 | 5 | 0.842694967101614 |
| chr5-26727238-26727739 | 8 | 0.881200361813605 |
| chr1-174154086-174154587 | 8 | 0.915204117150281 |
| chrX-115740348-115740849 | 3 | 0.996057927811504 |
| chr1-34858006-34858507 | 4 | 0.939434780744063 |
| chr4-129216107-129216608 | 8 | 0.942013135341834 |
| chr15-62202397-62202898 | 5 | 0.960076322238473 |
| chr2-156109166-156109667 | 3 | 0.99593553545041 |
| chr2-147626640-147627141 | 8 | 0.886004958071468 |
| chr16-60274354-60274855 | 8 | 0.988939170869361 |
| chr15-81226146-81226647 | 8 | 0.994907766741338 |
| chr7-121423392-121423893 | 4 | 0.962929343063843 |

| | | |
|---|---|---|
| chr5-58930740-58931241 | 8 | 0.945371029731997 |
| chr3-41856558-41857059 | 8 | 0.988605052099277 |
| chr9-78580914-78581415 | 5 | 0.990682284803498 |
| chr2-232067921-232068422 | 8 | 0.994530552308799 |
| chr10-10059144-10059645 | 8 | 0.943957795252576 |
| chr14-11667336-11667837 | 5 | 0.955569421642611 |
| chr14-78839921-78840422 | 4 | 0.856799277881176 |
| chr6-116749195-116749696 | 3 | 0.986842511766666 |
| chr9-104352890-104353391 | 8 | 0.996845473808236 |
| chr2-98370538-98371039 | 4 | 0.8416923420378 |
| chr17-655767-656268 | 8 | 0.928246973498906 |
| chr2-167489885-167490386 | 8 | 0.821706419930459 |
| chr10-55730193-55730694 | 8 | 0.823997488070409 |
| chr17-14260636-14261137 | 8 | 0.977821696598609 |
| chr18-77647422-77647923 | 8 | 0.978228069873462 |
| chr1-67146553-67147054 | 8 | 0.919827840931405 |
| chr19-31627918-31628419 | 6 | 0.988363105300438 |
| chr8-5812733-5813234 | 8 | 0.979288255379467 |
| chr15-1033141-1033642 | 8 | 0.979583656945881 |
| chr2-44553691-44554192 | 8 | 0.984183016727331 |
| chr11-9276568-9277069 | 8 | 0.971708182925526 |
| chr5-111070663-111071164 | 8 | 0.993493681128311 |
| chrX-79688433-79688934 | 7 | 0.868204171866734 |
| chrY-5758181-5758682 | 6 | 0.852056704743975 |
| chr2-64872569-64873070 | 5 | 0.915450426149226 |
| chr17-20052556-20053057 | 8 | 0.9271037596446 |
| chr5-21519947-21520448 | 5 | 0.96473600397852 |
| chr1-200139900-200140401 | 3 | 0.898682281958431 |
| chr9-76133061-76133562 | 8 | 0.970203998469065 |
| chr15-35048156-35048657 | 8 | 0.989819634690337 |
| chr6-3077873-3078374 | 8 | 0.916627918903963 |
| chr11-67802535-67803036 | 8 | 0.981111391880657 |
| chr5-38278261-38278762 | 4 | 0.851939701653624 |
| chr6-123173003-123173504 | 8 | 0.897042636476782 |
| chr2-143407065-143407566 | 8 | 0.995113792040608 |
| chr3-125524409-125524910 | 5 | 0.874228204354204 |
| chr7-63542959-63543460 | 8 | 0.89075425732114 |
| chr4-180137362-180137863 | 5 | 0.988223468989799 |
| chr5-58772766-58773267 | 5 | 0.987947353879595 |
| chr18-6413100-6413601 | 3 | 0.953883418760933 |
| chr5-78385057-78385558 | 4 | 0.853097623027925 |
| chr10-48562578-48563079 | 8 | 0.991495655412136 |
| chr11-48975391-48975892 | 5 | 0.873449493099418 |

| | | |
|---|---|---|
| chr15-62189732-62190233 | 5 | 0.83284384287457 |
| chr3-37942992-37943493 | 5 | 0.983919978840557 |
| chr1-224359852-224360353 | 8 | 0.979910769488646 |
| chr4-157367510-157368011 | 8 | 0.994429514371815 |
| chr10-64955873-64956374 | 4 | 0.835418854917691 |
| chr2-208219462-208219963 | 2 | 0.964577919398351 |
| chr19-48253193-48253694 | 2 | 0.839916999885623 |
| chr14-31679228-31679729 | 8 | 0.864047621642754 |
| chr1-135934734-135935235 | 8 | 0.955898931934306 |
| chr14-186562-187063 | 2 | 0.863809924679762 |
| chr13-55000758-55001259 | 8 | 0.821326081391467 |
| chr7-97610024-97610525 | 8 | 0.997565587346911 |
| chr13-61841508-61842009 | 2 | 0.940416753782491 |
| chr20-18824406-18824907 | 8 | 0.962411743325566 |
| chr14-28147235-28147736 | 8 | 0.993976091653589 |
| chr19-34439066-34439567 | 8 | 0.964710196395766 |
| chr4-182167920-182168421 | 3 | 0.976808077598835 |
| chr10-74763584-74764085 | 5 | 0.989435962526293 |
| chr15-83912454-83912955 | 8 | 0.916126942086789 |
| chr6-27773564-27774065 | 8 | 0.955081536054246 |
| chrX-31107522-31108023 | 8 | 0.995153179329754 |
| chr4-74647372-74647873 | 8 | 0.982900511465296 |
| chr1-127911458-127911959 | 8 | 0.850664604661421 |
| chr10-22269815-22270316 | 8 | 0.831305646452959 |
| chr2-125641135-125641636 | 6 | 0.988148134959496 |
| chr14-75624354-75624855 | 5 | 0.986078300660748 |
| chr6-83382872-83383373 | 3 | 0.99593553545041 |
| chr16-62656516-62657017 | 2 | 0.831801740199866 |
| chr4-79373609-79374110 | 5 | 0.978834427121022 |
| chr13-93472839-93473340 | 8 | 0.994742731178189 |
| chrX-4237733-4238234 | 8 | 0.993493681128311 |
| chr16-74288858-74289359 | 8 | 0.968703510080896 |
| chr8-102261385-102261886 | 8 | 0.961195964031978 |
| chr9-46157672-46158173 | 8 | 0.957937205607858 |
| chr12-23287016-23287517 | 8 | 0.993735773584839 |
| chr1-48428460-48428961 | 5 | 0.916380230209981 |
| chr6-20741350-20741851 | 8 | 0.979000701921469 |
| chr5-155122980-155123481 | 8 | 0.956922605536545 |
| chr5-152565849-152566350 | 8 | 0.991379426629717 |
| chr3-36136741-36137242 | 8 | 0.898443748246846 |
| chr12-30991291-30991792 | 8 | 0.967969223079188 |
| chr2-10737064-10737565 | 8 | 0.993493681128311 |
| chr1-568473-568974 | 3 | 0.993992022791908 |

| | | |
|---|---|---|
| chr1-106056517-106057018 | 8 | 0.971651067297681 |
| chr17-23823030-23823531 | 8 | 0.985085839601815 |
| chr13-33017056-33017557 | 8 | 0.915910137131804 |
| chr14-55504873-55505374 | 8 | 0.993493681128311 |
| chr2-65131108-65131609 | 8 | 0.995163235390143 |
| chr5-43295246-43295747 | 8 | 0.954786412390278 |
| chr15-26470059-26470560 | 8 | 0.829344099506427 |
| chr13-103612486-103612987 | 5 | 0.98179373438936 |
| chr20-1027799-1028300 | 8 | 0.987826590079807 |
| chr8-19107060-19107561 | 8 | 0.83064962566909 |
| chr13-40977915-40978416 | 8 | 0.847239491978563 |
| chrX-150847421-150847922 | 8 | 0.989648259088772 |
| chr7-112310922-112311423 | 4 | 0.95530211805384 |
| chr14-58929167-58929668 | 5 | 0.988741972646273 |
| chr2-21235745-21236246 | 8 | 0.957835714742638 |
| chr14-57681547-57682048 | 6 | 0.992222664191107 |
| chr11-64632905-64633406 | 8 | 0.993493681128311 |
| chr8-59318645-59319146 | 8 | 0.992617032912616 |
| chr2-132370427-132370928 | 8 | 0.886268911035437 |
| chrX-133221825-133222326 | 5 | 0.991206160475241 |
| chr17-69246342-69246843 | 8 | 0.907504472758137 |
| chr5-79492597-79493098 | 8 | 0.979755533088474 |
| chr18-12403299-12403800 | 8 | 0.955642784336213 |
| chr2-226650045-226650546 | 3 | 0.988057186092265 |
| chr6-19613122-19613623 | 4 | 0.959386220672609 |
| chr4-139842761-139843262 | 6 | 0.994031352842656 |
| chr10-26899406-26899907 | 3 | 0.993482064979321 |
| chr12-6304143-6304644 | 8 | 0.99428949688746 |
| chr16-73143670-73144171 | 5 | 0.990332032641779 |
| chr1-133926386-133926887 | 8 | 0.983200419959398 |
| chr15-24160669-24161170 | 8 | 0.86736358103669 |
| chr11-21730740-21731241 | 6 | 0.991206925481043 |
| chr12-38748212-38748713 | 2 | 0.950535590425129 |
| chr13-73675783-73676284 | 8 | 0.869406345467923 |
| chr8-99472085-99472586 | 8 | 0.862198961760803 |
| chr17-81046993-81047494 | 8 | 0.870439597890846 |
| chr1-94124927-94125428 | 3 | 0.954565675802226 |
| chr19-37583184-37583685 | 8 | 0.983950233233345 |
| chr15-35578160-35578661 | 8 | 0.873383384641436 |
| chr14-15785939-15786440 | 8 | 0.988449554038087 |
| chr1-26844860-26845361 | 8 | 0.960476534933033 |
| chr1-249904875-249905376 | 8 | 0.993633755209933 |
| chr6-28896610-28897111 | 8 | 0.992167163888172 |

| | | |
|---|---|---|
| chrX-68354710-68355211 | 3 | 0.990906037971537 |
| chr3-41327419-41327920 | 8 | 0.868469789664169 |
| chr16-42766063-42766564 | 4 | 0.972549333839349 |
| chr6-3512477-3512978 | 8 | 0.934814430516613 |
| chr11-5993131-5993632 | 6 | 0.988309758285319 |
| chr6-116031371-116031872 | 8 | 0.993493681128311 |
| chr17-81881416-81881917 | 5 | 0.965207945661319 |
| chr13-76459783-76460284 | 3 | 0.869442106266787 |
| chr5-119397355-119397856 | 5 | 0.97874046367686 |
| chr13-45481973-45482474 | 4 | 0.928514286165737 |
| chr10-41766766-41767267 | 8 | 0.891898398234194 |
| chr12-1476862-1477363 | 8 | 0.880471321619327 |
| chr17-30690984-30691485 | 8 | 0.995089028143569 |
| chr8-43667449-43667950 | 4 | 0.902130623388477 |
| chr14-58130185-58130686 | 3 | 0.996851427205213 |
| chr2-122109403-122109904 | 8 | 0.920504305722323 |
| chr16-58495454-58495955 | 8 | 0.959954114215097 |
| chr6-67775744-67776245 | 4 | 0.963581677527661 |
| chr2-214105946-214106447 | 8 | 0.986231780569827 |
| chr5-106904631-106905132 | 6 | 0.98319961352094 |
| chr6-75113471-75113972 | 3 | 0.996312260958674 |
| chr4-11156515-11157016 | 4 | 0.817942419237714 |
| chr6-90654305-90654806 | 2 | 0.918168897065971 |
| chr7-27332134-27332635 | 8 | 0.928565257855622 |
| chr7-16390708-16391209 | 3 | 0.98558886808213 |
| chr9-61746378-61746879 | 8 | 0.991513023298187 |
| chr8-76399073-76399574 | 8 | 0.97136545135833 |
| chr3-157792031-157792532 | 8 | 0.996289742977285 |
| chr8-87344696-87345197 | 8 | 0.921596622948749 |
| chr7-57289603-57290104 | 8 | 0.990026291246426 |
| chr19-9644034-9644535 | 7 | 0.879873886115864 |
| chr10-55026240-55026741 | 5 | 0.861009390657141 |
| chr15-7309552-7310053 | 4 | 0.884591959091785 |
| chr3-103894443-103894944 | 8 | 0.907071017035518 |
| chr4-67946416-67946917 | 8 | 0.830190527156669 |
| chr15-67223459-67223960 | 8 | 0.853274694185567 |
| chr8-24873131-24873632 | 8 | 0.873526521177816 |
| chr10-59738960-59739461 | 5 | 0.987685192198018 |
| chr14-47503919-47504420 | 8 | 0.979005297199441 |
| chr1-125309262-125309763 | 3 | 0.996685720724764 |
| chr1-40226999-40227500 | 8 | 0.982374660525766 |
| chr2-55693070-55693571 | 8 | 0.990279813276089 |
| chr16-67765533-67766034 | 8 | 0.973219672048486 |

| | | |
|---|---|---|
| chr3-87781763-87782264 | 8 | 0.929154522133201 |
| chr19-51409048-51409549 | 8 | 0.971333503513025 |
| chr2-44551312-44551813 | 8 | 0.991673005733594 |
| chr10-26173083-26173584 | 8 | 0.993493681128311 |
| chrX-116488328-116488829 | 5 | 0.872785486766107 |
| chr16-80546484-80546985 | 4 | 0.845264692369925 |
| chr11-66421722-66422223 | 3 | 0.893290529110801 |
| chr1-75519633-75520134 | 3 | 0.866345261238269 |
| chr17-27709404-27709905 | 8 | 0.879460015384316 |
| chr14-93884673-93885174 | 8 | 0.990806059089869 |
| chr2-118950761-118951262 | 8 | 0.995600184406168 |
| chr14-68602012-68602513 | 8 | 0.876028073987474 |
| chr4-77887860-77888361 | 8 | 0.993493681128311 |
| chr8-8079601-8080102 | 3 | 0.995951941465573 |
| chr5-36297382-36297883 | 8 | 0.982271519659038 |
| chr7-71094215-71094716 | 8 | 0.935871572317095 |
| chr4-155626674-155627175 | 6 | 0.96038411414131 |
| chr4-75886663-75887164 | 3 | 0.99593553545041 |
| chr16-55697673-55698174 | 5 | 0.956640979869554 |
| chr3-122814273-122814774 | 2 | 0.951026360746587 |
| chr5-29491351-29491852 | 8 | 0.993493681128311 |
| chr1-6592227-6592728 | 8 | 0.983515210222612 |
| chr5-115845965-115846466 | 2 | 0.852528816591986 |
| chr1-135923962-135924463 | 4 | 0.945616381637409 |
| chr2-110765462-110765963 | 8 | 0.940666614988774 |
| chr14-95595722-95596223 | 5 | 0.990285232795708 |
| chr14-72639880-72640381 | 5 | 0.974072979571148 |
| chr1-248163225-248163726 | 8 | 0.871081274948753 |
| chr20-14466586-14467087 | 6 | 0.993201167145366 |
| chr1-143669480-143669981 | 3 | 0.993819850005127 |
| chr10-41688350-41688851 | 4 | 0.880647463478756 |
| chr20-16564582-16565083 | 3 | 0.996217935981319 |
| chr3-107630509-107631010 | 8 | 0.878979193977552 |
| chr5-109634975-109635476 | 5 | 0.932326059185982 |
| chr5-16110684-16111185 | 8 | 0.936729071773112 |
| chr1-86471950-86472451 | 5 | 0.953715798506803 |
| chr9-68148154-68148655 | 4 | 0.8734977484048 |
| chr6-78167673-78168174 | 8 | 0.983740036153411 |
| chr3-148505612-148506113 | 2 | 0.943522139782594 |
| chr1-96565948-96566449 | 1 | 0.852108347256898 |
| chr8-96728511-96729012 | 5 | 0.992164953580715 |
| chr8-118116880-118117381 | 4 | 0.87339378605388 |
| chr5-69808754-69809255 | 8 | 0.985884159526385 |

| | | |
|---|---|---|
| chr6-48945398-48945899 | 8 | 0.935435378894262 |
| chr1-220957084-220957585 | 7 | 0.891782986271661 |
| chr18-26892255-26892756 | 8 | 0.994029100980513 |
| chr1-253724550-253725051 | 2 | 0.839288999150634 |
| chr10-35407594-35408095 | 3 | 0.997862480007738 |
| chr6-140095120-140095621 | 8 | 0.987686725295283 |
| chr5-151005269-151005770 | 8 | 0.967779956619568 |
| chr1-241926194-241926695 | 8 | 0.953699025521457 |
| chr2-167988549-167989050 | 3 | 0.983433363067625 |
| chr8-120899360-120899861 | 8 | 0.980281816950161 |
| chr16-75258352-75258853 | 8 | 0.986540457457607 |
| chr6-47793815-47794316 | 4 | 0.96263215455938 |
| chrX-25014798-25015299 | 8 | 0.883407934374703 |
| chr10-20310786-20311287 | 8 | 0.985394672641985 |
| chr12-965595-966096 | 8 | 0.992536176924543 |
| chr5-68661620-68662121 | 5 | 0.993286217774106 |
| chr5-3261534-3262035 | 6 | 0.983312441442335 |
| chr3-139561549-139562050 | 8 | 0.896597447088339 |
| chr4-153336825-153337326 | 4 | 0.897730737191797 |
| chr1-103461652-103462153 | 6 | 0.989894881229319 |
| chr1-54026179-54026680 | 2 | 0.966113131794544 |
| chr1-198737498-198737999 | 8 | 0.83688219197322 |
| chr7-11702833-11703334 | 8 | 0.985526230756456 |
| chr16-83157119-83157620 | 6 | 0.991471404888775 |
| chr3-22187270-22187771 | 8 | 0.911716569070207 |
| chr3-78551165-78551666 | 8 | 0.826944122501481 |
| chr19-36547473-36547974 | 8 | 0.9949366340577774 |
| chr18-55278098-55278599 | 8 | 0.892032671832712 |
| chr4-168153772-168154273 | 8 | 0.99379318123855 |
| chr12-33015990-33016491 | 8 | 0.961689670691115 |
| chr3-141547574-141548075 | 8 | 0.921160338740411 |
| chr4-123095548-123096049 | 8 | 0.993493681128311 |
| chr4-29094384-29094885 | 8 | 0.824142265878751 |
| chr3-38530634-38531135 | 2 | 0.964724702782464 |
| chr5-41396181-41396682 | 2 | 0.959316407271058 |
| chr5-139258677-139259178 | 8 | 0.912343277632486 |
| chr3-10324663-10325164 | 8 | 0.990293138079875 |
| chr4-26132777-26133278 | 8 | 0.979330619420946 |
| chr16-43005474-43005975 | 8 | 0.885883800612054 |
| chr16-58709350-58709851 | 8 | 0.889677137448723 |
| chr14-26944430-26944931 | 8 | 0.993493681128311 |
| chr5-55166060-55166561 | 6 | 0.991206925481043 |
| chr1-177804867-177805368 | 8 | 0.834576991401949 |

| | | |
|---|---|---|
| chr5-113845065-113845566 | 4 | 0.934384043746915 |
| chr9-67446462-67446963 | 8 | 0.994414945403355 |
| chr10-50686852-50687353 | 8 | 0.974917285913012 |
| chr2-31002014-31002515 | 8 | 0.989927202354624 |
| chr20-26188632-26189133 | 5 | 0.977614839222443 |
| chr3-32293255-32293756 | 8 | 0.877056345528274 |
| chr7-105736348-105736849 | 8 | 0.992997170885802 |
| chr1-23667910-23668411 | 5 | 0.958198801395714 |
| chr14-10108099-10108600 | 4 | 0.90895816348187 |
| chr14-62995949-62996450 | 8 | 0.815712370104256 |
| chrX-110061532-110062033 | 4 | 0.860557490570523 |
| chr4-51655776-51656277 | 8 | 0.972988494187783 |
| chr9-74948445-74948946 | 8 | 0.995209302253885 |
| chr7-128221600-128222101 | 8 | 0.915159881378893 |
| chr4-178561267-178561768 | 8 | 0.924658420749534 |
| chr6-102789901-102790402 | 8 | 0.838204408462171 |
| chr18-15403585-15404086 | 8 | 0.936225086891032 |
| chr17-170092-170593 | 2 | 0.881607679304144 |
| chr14-104671144-104671645 | 4 | 0.893304487669299 |
| chr16-776337-776838 | 3 | 0.915176241089321 |
| chr5-78247638-78248139 | 6 | 0.992483522539783 |
| chr17-27782821-27783322 | 8 | 0.993493681128311 |
| chr15-9506467-9506968 | 3 | 0.927315764073622 |
| chr8-108581673-108582174 | 8 | 0.963721910291173 |
| chr9-22908688-22909189 | 2 | 0.951380054211941 |
| chr2-224187930-224188431 | 8 | 0.834594868251031 |
| chr13-81871970-81872471 | 3 | 0.83538600875349 |
| chr7-42387501-42388002 | 8 | 0.995133174528612 |
| chr4-173658736-173659237 | 8 | 0.95832960742693 |
| chr8-70855256-70855757 | 8 | 0.988315073376127 |
| chr7-106568549-106569050 | 8 | 0.855329676246757 |
| chr13-49055319-49055820 | 8 | 0.853317197274788 |
| chr14-31481494-31481995 | 4 | 0.83207643871342 |
| chr5-88655746-88656247 | 8 | 0.975424501375619 |
| chr13-100217602-100218103 | 5 | 0.943247109634814 |
| chr2-222280347-222280848 | 8 | 0.98432146260992 |
| chr13-43321309-43321810 | 5 | 0.987315722422795 |
| chr8-79196698-79197199 | 5 | 0.980569097441728 |
| chr16-29171928-29172429 | 8 | 0.93195902780643 |
| chr2-56167141-56167642 | 3 | 0.987621386882311 |
| chr8-73944747-73945248 | 4 | 0.934183890918455 |
| chr2-184061816-184062317 | 3 | 0.99593553545041 |
| chr7-75705674-75706175 | 2 | 0.956928417472273 |

| | | |
|---|---|---|
| chr5-66052648-66053149 | 3 | 0.962532829754025 |
| chr7-27869756-27870257 | 4 | 0.885498472790252 |
| chr3-110548690-110549191 | 8 | 0.976205670988115 |
| chr5-120043287-120043788 | 4 | 0.923598656968117 |
| chr5-150872849-150873350 | 5 | 0.93860413892271 |
| chr15-42655495-42655996 | 8 | 0.826077062265398 |
| chr15-39987056-39987557 | 8 | 0.969812831732135 |
| chr4-76329433-76329934 | 8 | 0.874468656525705 |
| chr8-120118704-120119205 | 8 | 0.989210825838748 |
| chrX-10817791-10818292 | 5 | 0.958429496145956 |
| chr18-34724931-34725432 | 8 | 0.889102763696362 |
| chr2-102639114-102639615 | 8 | 0.980524944908328 |
| chrX-126418482-126418983 | 2 | 0.922370413234189 |
| chr2-61920377-61920878 | 8 | 0.931570865981956 |
| chr19-29829571-29830072 | 8 | 0.953992597984898 |
| chr4-132173194-132173695 | 8 | 0.986755127022684 |
| chr2-143025774-143026275 | 8 | 0.990845532556797 |
| chr4-80299246-80299747 | 8 | 0.953664967310204 |
| chr2-68539562-68540063 | 5 | 0.891097314812115 |
| chr15-61964074-61964575 | 8 | 0.837186229719463 |
| chr6-137702110-137702611 | 3 | 0.99593553545041 |
| chr3-101616776-101617277 | 3 | 0.998543787870459 |
| chr9-46241274-46241775 | 8 | 0.920926342784931 |
| chr16-7033974-7034475 | 8 | 0.914121875246853 |
| chr9-87501198-87501699 | 8 | 0.98132688866414 |
| chr4-159428941-159429442 | 5 | 0.986515046142745 |
| chr9-80360325-80360826 | 8 | 0.927747734632044 |
| chr5-48077127-48077628 | 6 | 0.98787477509209 |
| chr15-72691665-72692166 | 8 | 0.838819220261224 |
| chr8-60289550-60290051 | 8 | 0.986388113632232 |
| chr10-2206289-2206790 | 8 | 0.993493681128311 |
| chr3-38031124-38031625 | 5 | 0.972620589962599 |
| chr16-71792557-71793058 | 6 | 0.934285907342789 |
| chr3-63193617-63194118 | 8 | 0.903795023898252 |
| chr5-67141024-67141525 | 8 | 0.92094780883743 |
| chrX-70738290-70738791 | 6 | 0.991206925481043 |
| chr6-71249288-71249789 | 8 | 0.9870056278277 |
| chr1-119158940-119159441 | 3 | 0.997704084502303 |
| chr18-14445147-14445648 | 5 | 0.980530442003346 |
| chr15-50722986-50723487 | 8 | 0.934862622921996 |
| chr2-140801400-140801901 | 5 | 0.982617676687013 |
| chr3-117511075-117511576 | 8 | 0.989440514601466 |
| chr4-71365654-71366155 | 8 | 0.866562791968294 |

| | | |
|---|---|---|
| chr15-98735919-98736420 | 2 | 0.815351972762079 |
| chr15-15229961-15230462 | 8 | 0.971519938496611 |
| chr4-158710002-158710503 | 8 | 0.993493681128311 |
| chr1-164358440-164358941 | 2 | 0.948465631959252 |
| chr15-29858769-29859270 | 6 | 0.992490965495286 |
| chr10-8683776-8684277 | 4 | 0.958084231615706 |
| chr1-98847721-98848222 | 5 | 0.976181575285954 |
| chr9-98813861-98814362 | 8 | 0.918054720272046 |
| chr17-59923849-59924350 | 8 | 0.993493681128311 |
| chr6-137291332-137291833 | 8 | 0.993493681128311 |
| chr9-16918058-16918559 | 8 | 0.992916724214439 |
| chr18-78703167-78703668 | 3 | 0.976456268273376 |
| chr1-157389035-157389536 | 4 | 0.854426948705124 |
| chr1-81638126-81638627 | 3 | 0.99061685301606 |
| chr1-41661447-41661948 | 2 | 0.935977442194775 |
| chr10-25153279-25153780 | 6 | 0.9931807605013 |
| chr2-52448447-52448948 | 5 | 0.935274973665351 |
| chr6-51599026-51599527 | 8 | 0.930627675173395 |
| chr18-63464022-63464523 | 8 | 0.911841572478293 |
| chr20-26713787-26714288 | 8 | 0.891392618929006 |
| chr4-103169497-103169998 | 8 | 0.993006739627703 |
| chr2-58458598-58459099 | 8 | 0.987905804188757 |
| chr18-19058560-19059061 | 8 | 0.84017437866554 |
| chrX-24416361-24416862 | 8 | 0.853031422978276 |
| chr6-25874065-25874566 | 5 | 0.965481088235254 |
| chr5-1582070-1582571 | 8 | 0.993493681128311 |
| chr7-47045864-47046365 | 5 | 0.940659461219864 |
| chr14-40172475-40172976 | 4 | 0.868506548564125 |
| chr15-86458788-86459289 | 8 | 0.99519355507753 |
| chr7-54773849-54774350 | 8 | 0.905713513373235 |
| chr6-82931483-82931984 | 8 | 0.993493681128311 |
| chr5-148499887-148500388 | 3 | 0.963663828622579 |
| chr9-57697833-57698334 | 2 | 0.934538683914776 |
| chr5-93943568-93944069 | 3 | 0.996557904694748 |
| chr2-122835164-122835665 | 8 | 0.953740368043111 |
| chr2-190886081-190886582 | 8 | 0.823019220389405 |
| chr5-12171255-12171756 | 2 | 0.86898673405384 |
| chr5-44694750-44695251 | 8 | 0.904931081070325 |
| chr3-141493116-141493617 | 8 | 0.946657479620538 |
| chr6-59144192-59144693 | 8 | 0.834651647213692 |
| chr20-38602631-38603132 | 4 | 0.871420900899657 |
| chr17-70394475-70394976 | 5 | 0.910389148333808 |
| chr17-55432065-55432566 | 5 | 0.917344851099771 |

| | | |
|---|---|---|
| chr16-12477645-12478146 | 4 | 0.939490169175155 |
| chr11-35849716-35850217 | 8 | 0.933752275220788 |
| chr1-91623656-91624157 | 3 | 0.99263802352731 |
| chr9-67517938-67518439 | 8 | 0.993493681128311 |
| chr18-25525556-25526057 | 8 | 0.879819026533795 |
| chr9-33480569-33481070 | 2 | 0.824734651248943 |
| chr17-39689034-39689535 | 2 | 0.909315609740003 |
| chr4-116150992-116151493 | 8 | 0.917123907970221 |
| chr2-87972955-87973456 | 8 | 0.880251545820898 |
| chr8-101352223-101352724 | 6 | 0.992274034649511 |
| chr19-39901809-39902310 | 5 | 0.995085264342889 |
| chrX-29395545-29396046 | 5 | 0.848986364976839 |
| chr19-18957021-18957522 | 8 | 0.889977256150003 |
| chr18-59515925-59516426 | 8 | 0.904860668847666 |
| chr7-21784491-21784992 | 2 | 0.871415235995452 |
| chr16-72123444-72123945 | 5 | 0.993792221153439 |
| chr8-95394670-95395171 | 4 | 0.943514248594481 |
| chr3-52430722-52431223 | 4 | 0.858299358676793 |
| chr17-21539845-21540346 | 5 | 0.990439523329323 |
| chr14-20492463-20492964 | 8 | 0.985167968348652 |
| chr5-149118240-149118741 | 4 | 0.969777617530282 |
| chr14-41707595-41708096 | 8 | 0.944235504371804 |
| chr6-76080290-76080791 | 8 | 0.819043846858039 |
| chr11-85119865-85120366 | 8 | 0.883757326710225 |
| chr5-123077206-123077707 | 8 | 0.986436001474854 |
| chr3-148373849-148374350 | 8 | 0.995165327665047 |
| chr9-5194299-5194800 | 8 | 0.958454831974274 |
| chr4-104834151-104834652 | 8 | 0.985814227607062 |
| chr1-243058447-243058948 | 8 | 0.930765146680948 |
| chr10-8438986-8439487 | 5 | 0.954312649489857 |
| chr3-138147600-138148101 | 8 | 0.880432342860105 |
| chr4-135045771-135046272 | 8 | 0.995239210046358 |
| chr10-52082608-52083109 | 8 | 0.974519644681608 |
| chr3-102301461-102301962 | 8 | 0.927771243764169 |
| chr11-4594120-4594621 | 8 | 0.943070861415812 |
| chr8-107286091-107286592 | 8 | 0.994900297127154 |
| chr2-39752927-39753428 | 8 | 0.995028910426534 |
| chr15-90435380-90435881 | 8 | 0.993493681128311 |
| chr14-61561897-61562398 | 8 | 0.925215914165712 |
| chr6-6478129-6478630 | 8 | 0.899344825391178 |
| chrX-73621968-73622469 | 4 | 0.969832619320354 |
| chr2-200217512-200218013 | 8 | 0.993565855122315 |
| chr18-9729619-9730120 | 3 | 0.996284971196036 |

| | | |
|---|---|---|
| chr6-102289694-102290195 | 8 | 0.867691631137541 |
| chr1-190930520-190931021 | 8 | 0.848345945390761 |
| chr3-103847328-103847829 | 8 | 0.990077775697339 |
| chr19-26898567-26899068 | 2 | 0.846471074665282 |
| chr1-118864036-118864537 | 8 | 0.972251693917693 |
| chr13-31078829-31079330 | 8 | 0.93060612257034 |
| chr2-218231917-218232418 | 8 | 0.94828462622455 |
| chr9-72666439-72666940 | 3 | 0.99662554764663 |
| chr3-101827754-101828255 | 8 | 0.940200857502643 |
| chr20-21819099-21819600 | 5 | 0.952429859858153 |
| chr4-88230940-88231441 | 8 | 0.965397882932012 |
| chr5-125965346-125965847 | 8 | 0.993329583219388 |
| chr16-47124094-47124595 | 8 | 0.829907238469611 |
| chr4-22430307-22430808 | 8 | 0.987304728390917 |
| chr5-166085213-166085714 | 8 | 0.968753583670107 |
| chr10-43079911-43080412 | 3 | 0.99593553545041 |
| chr13-33345376-33345877 | 3 | 0.996889194532436 |
| chrX-56989739-56990240 | 3 | 0.99593553545041 |
| chr20-16692868-16693369 | 8 | 0.994298698387272 |
| chr1-121006494-121006995 | 8 | 0.957709921933779 |
| chr16-73842475-73842976 | 8 | 0.974796610410736 |
| chr3-143753549-143754050 | 8 | 0.945312614356119 |
| chr14-33197984-33198485 | 4 | 0.950400553255401 |
| chr6-125369945-125370446 | 8 | 0.988828266895251 |
| chr9-85819478-85819979 | 8 | 0.940884711274443 |
| chr17-50388927-50389428 | 8 | 0.983933018209293 |
| chr7-64653696-64654197 | 3 | 0.988915084330434 |
| chr4-175467371-175467872 | 5 | 0.991003188055346 |
| chr2-216507424-216507925 | 8 | 0.968448634042534 |
| chr7-50610936-50611437 | 8 | 0.90260343866271 |
| chr19-52504480-52504981 | 6 | 0.990226728692557 |
| chr11-78174943-78175444 | 8 | 0.955014485652641 |
| chr5-166489393-166489894 | 8 | 0.993175641632548 |
| chr2-170665731-170666232 | 4 | 0.97287406017879 |
| chr9-95757707-95758208 | 4 | 0.964052962355378 |
| chr10-57719762-57720263 | 8 | 0.955103695260154 |
| chr7-81784375-81784876 | 4 | 0.969078753973921 |
| chr3-32037293-32037794 | 8 | 0.850493044421447 |
| chr16-66058098-66058599 | 5 | 0.947304713156784 |
| chr14-43254149-43254650 | 8 | 0.836380471786306 |
| chr15-30467103-30467604 | 8 | 0.8927924094834 |
| chr11-26736936-26737437 | 5 | 0.987804249137695 |
| chr6-124820931-124821432 | 8 | 0.81639685393698 |

| | | |
|---|---|---|
| chr17-27066712-27067213 | 8 | 0.963808719792509 |
| chr8-114719485-114719986 | 8 | 0.950381058919036 |
| chr10-48458637-48459138 | 8 | 0.994908142359356 |
| chr8-105326868-105327369 | 8 | 0.928973213972712 |
| chr3-122532717-122533218 | 3 | 0.947925717551948 |
| chr12-35252717-35253218 | 8 | 0.860375702884479 |
| chr9-86100027-86100528 | 4 | 0.864061070660303 |
| chr3-102721447-102721948 | 8 | 0.879866817496217 |
| chr3-50873335-50873836 | 8 | 0.890026196719816 |
| chr6-91981236-91981737 | 8 | 0.987294678644909 |
| chr4-70684270-70684771 | 8 | 0.993217656380162 |
| chr1-168491114-168491615 | 8 | 0.99313235747249 |
| chr14-28272251-28272752 | 8 | 0.99519404165065 |
| chr10-96756401-96756902 | 4 | 0.895417349569256 |
| chr17-7548862-7549363 | 8 | 0.955292571245059 |
| chr11-297337-297838 | 3 | 0.996406111673441 |
| chr2-17954205-17954706 | 6 | 0.88077256423409 |
| chr17-9625210-9625711 | 4 | 0.939967617162789 |
| chr9-65399933-65400434 | 3 | 0.932808788260829 |
| chr10-1180607-1181108 | 2 | 0.848083868608435 |
| chr8-81680158-81680659 | 8 | 0.993493681128311 |
| chr14-62723970-62724471 | 5 | 0.940527201948543 |
| chr2-191779809-191780310 | 8 | 0.99533578105651 |
| chr2-241645147-241645648 | 8 | 0.973753370379871 |
| chr11-54660447-54660948 | 3 | 0.99593553545041 |
| chr17-26552278-26552779 | 8 | 0.983614396989559 |
| chr17-71641722-71642223 | 8 | 0.990033508345616 |
| chr20-48521911-48522412 | 2 | 0.898117297614947 |
| chrX-96913318-96913819 | 3 | 0.991494487781362 |
| chrX-116841301-116841802 | 8 | 0.877529958182219 |
| chr9-13812673-13813174 | 8 | 0.965111079603077 |
| chr14-55392848-55393349 | 2 | 0.833663952563337 |
| chr1-192592061-192592562 | 4 | 0.856754444997303 |
| chr18-73220435-73220936 | 8 | 0.994058862127475 |
| chr16-40667268-40667769 | 3 | 0.99593553545041 |
| chr1-9192444-9192945 | 8 | 0.966959975126768 |
| chr13-82540155-82540656 | 5 | 0.989400624339308 |
| chr9-69496260-69496761 | 8 | 0.865334636106373 |
| chr9-89369426-89369927 | 8 | 0.943819564635749 |
| chr1-192159547-192160048 | 8 | 0.989153637871512 |
| chr1-183338588-183339089 | 4 | 0.961479801033472 |
| chr14-31239802-31240303 | 5 | 0.949099247707591 |
| chr8-31892962-31893463 | 5 | 0.939889040413698 |

| | | |
|---|---|---|
| chr11-3405251-3405752 | 8 | 0.882194570480049 |
| chr20-14797935-14798436 | 8 | 0.975268604602198 |
| chr9-50756207-50756708 | 8 | 0.983106510268216 |
| chr19-13989126-13989627 | 3 | 0.992416725365231 |
| chr10-43010621-43011122 | 6 | 0.937868746025932 |
| chr15-54103675-54104176 | 5 | 0.982651100669321 |
| chr1-125889519-125890020 | 8 | 0.865069067523506 |
| chr12-31843628-31844129 | 4 | 0.89766067552295 |
| chrX-73273899-73274400 | 8 | 0.865867006350294 |
| chr18-33818209-33818710 | 5 | 0.991360855404516 |
| chr5-125147842-125148343 | 8 | 0.993493681128311 |
| chr5-110591230-110591731 | 8 | 0.97872370525008 |
| chr8-98622040-98622541 | 4 | 0.946355453860108 |
| chr17-82679976-82680477 | 8 | 0.993493681128311 |
| chr1-222987202-222987703 | 3 | 0.855946053803333 |
| chr1-50363732-50364233 | 5 | 0.865761796628513 |
| chrX-14466672-14467173 | 8 | 0.987974451270006 |
| chr14-6674341-6674842 | 4 | 0.887100444418251 |
| chr16-10630633-10631134 | 8 | 0.829003378573666 |
| chr2-223559875-223560376 | 8 | 0.962667318185833 |
| chr5-23566729-23567230 | 4 | 0.965187205293408 |
| chr4-51142617-51143118 | 6 | 0.992670670813826 |
| chr5-117898698-117899199 | 2 | 0.857360440465611 |
| chr2-62686832-62687333 | 6 | 0.991206925481043 |
| chr1-259558215-259558716 | 5 | 0.92061836070115 |
| chr2-81202695-81203196 | 8 | 0.979819740849478 |
| chr3-110068390-110068891 | 8 | 0.990636827364566 |
| chr17-18230840-18231341 | 8 | 0.894341907674817 |
| chr15-13714484-13714985 | 8 | 0.92265891939543 |
| chr10-96273033-96273534 | 4 | 0.954833489372769 |
| chr4-66442616-66443117 | 8 | 0.995151491848204 |
| chr18-15906083-15906584 | 8 | 0.845931398685075 |
| chr13-70380088-70380589 | 8 | 0.959575049304947 |
| chr4-131025393-131025894 | 8 | 0.95253825926256 |
| chrX-67023883-67024384 | 8 | 0.984269587573275 |
| chr15-92219908-92220409 | 8 | 0.952756621722308 |
| chr7-59413787-59414288 | 4 | 0.935468772375058 |
| chr4-24889933-24890434 | 6 | 0.991103085436289 |
| chr2-83772449-83772950 | 8 | 0.995226891360446 |
| chr5-68734636-68735137 | 8 | 0.829669198947815 |
| chr4-161501518-161502019 | 8 | 0.955355588690831 |
| chr1-172947868-172948369 | 4 | 0.904982166289167 |
| chr2-102429610-102430111 | 8 | 0.823155288657577 |

| | | |
|---|---|---|
| chr3-168536898-168537399 | 8 | 0.975800165735953 |
| chrX-115513703-115514204 | 5 | 0.983919978840557 |
| chr7-100785244-100785745 | 3 | 0.996873185655658 |
| chr3-102178530-102179031 | 5 | 0.930496952958563 |
| chr15-101338858-101339359 | 3 | 0.957134895458988 |
| chr20-42373110-42373611 | 8 | 0.948277355032351 |
| chr11-24220225-24220726 | 8 | 0.925219933127724 |
| chr10-104648472-104648973 | 8 | 0.863995608966473 |
| chr1-15040522-15041023 | 5 | 0.885697547361881 |
| chr7-33058206-33058707 | 8 | 0.973198962145375 |
| chr7-84077125-84077626 | 8 | 0.991400357178849 |
| chr8-5891186-5891687 | 6 | 0.98817122174991 |
| chr14-56703167-56703668 | 2 | 0.900629295224964 |
| chr15-10645511-10646012 | 4 | 0.84443463234913 |
| chr6-3544509-3545010 | 4 | 0.830213264774964 |
| chr5-77604575-77605076 | 8 | 0.985857688294869 |
| chr9-1927968-1928469 | 3 | 0.996771930095526 |
| chrX-127011494-127011995 | 4 | 0.973145005967885 |
| chr18-79490115-79490616 | 3 | 0.997277541491651 |
| chr6-43218481-43218982 | 8 | 0.994017859632465 |
| chr14-21752885-21753386 | 3 | 0.883957582120988 |
| chr4-144843247-144843748 | 8 | 0.939009729669961 |
| chr7-117551165-117551666 | 8 | 0.824168719973069 |
| chr3-76492066-76492567 | 5 | 0.988821783681582 |
| chr4-35991728-35992229 | 8 | 0.985319779087921 |
| chr8-81254751-81255252 | 8 | 0.986730996821119 |
| chr17-70746911-70747412 | 8 | 0.965645158748088 |
| chr12-7908279-7908780 | 8 | 0.985321419677807 |
| chr14-72575359-72575860 | 8 | 0.845338495922613 |
| chr2-52707504-52708005 | 6 | 0.94418771329116 |
| chr3-61643409-61643910 | 3 | 0.889211954315389 |
| chr7-83882602-83883103 | 6 | 0.99010194141 89 |
| chr15-81623693-81624194 | 8 | 0.969509786618143 |
| chr17-45828511-45829012 | 8 | 0.895816996849651 |
| chr5-45867153-45867654 | 6 | 0.903258736291049 |
| chr4-36690940-36691441 | 8 | 0.992063126166675 |
| chr7-9487532-9488033 | 8 | 0.976059645791644 |
| chr17-45803129-45803630 | 8 | 0.986755926440394 |
| chr4-46956209-46956710 | 2 | 0.875056326627141 |
| chr2-20500341-20500842 | 5 | 0.990425420160053 |
| chr9-3005619-3006120 | 8 | 0.991949157016063 |
| chr19-25204669-25205170 | 8 | 0.995916165266917 |
| chrX-124774118-124774619 | 3 | 0.996661199661046 |

| | | |
|---|---|---|
| chr3-121143724-121144225 | 8 | 0.944610404332621 |
| chr17-16217373-16217874 | 3 | 0.942074379645717 |
| chr14-43023825-43024326 | 8 | 0.99262266699874 |
| chr8-37781452-37781953 | 8 | 0.98360618119945 |
| chr5-87201337-87201838 | 3 | 0.994846511346013 |
| chr15-32400412-32400913 | 8 | 0.932361003593333 |
| chr20-49121016-49121517 | 5 | 0.968350260842793 |
| chr6-42760458-42760959 | 8 | 0.993376111800624 |
| chr13-34150147-34150648 | 6 | 0.928551759384217 |
| chr17-5414057-5414558 | 8 | 0.993493681128311 |
| chr11-82253228-82253729 | 6 | 0.834752608456563 |
| chr5-53656037-53656538 | 3 | 0.99593553545041 |
| chr2-154983604-154984105 | 3 | 0.99593553545041 |
| chr3-134465301-134465802 | 8 | 0.969380887567886 |
| chr2-110764467-110764968 | 8 | 0.995205957267471 |
| chr2-207443339-207443840 | 8 | 0.920358076458343 |
| chr15-55461915-55462416 | 4 | 0.917935044738934 |
| chr14-63857451-63857952 | 3 | 0.99593553545041 |
| chr2-46270166-46270667 | 4 | 0.921513703442939 |
| chr10-73540090-73540591 | 6 | 0.991206925481043 |
| chrX-120791637-120792138 | 8 | 0.953148264530752 |
| chr2-42289074-42289575 | 5 | 0.922790678920351 |
| chrX-10925607-10926108 | 5 | 0.910171291421988 |
| chr7-87505208-87505709 | 8 | 0.984462349758629 |
| chr19-54982614-54983115 | 8 | 0.972630510256996 |
| chrX-7679473-7679974 | 8 | 0.822315080722979 |
| chr18-35969249-35969750 | 4 | 0.954240088260527 |
| chr1-78026524-78027025 | 3 | 0.993165462269914 |
| chr14-65934186-65934687 | 8 | 0.940211602351308 |
| chr18-24427850-24428351 | 1 | 0.938189258701975 |
| chr19-2229216-2229717 | 5 | 0.912346300273086 |
| chr8-48669415-48669916 | 2 | 0.843067545798833 |
| chr12-46061695-46062196 | 8 | 0.948725310034176 |
| chr13-48508184-48508685 | 2 | 0.872276805755981 |
| chr13-98508486-98508987 | 8 | 0.982206550757032 |
| chr1-39527345-39527846 | 8 | 0.993493681128311 |
| chrX-51919332-51919833 | 3 | 0.970400054723722 |
| chr10-41371842-41372343 | 8 | 0.856505833097567 |
| chr2-66256695-66257196 | 8 | 0.994914606559955 |
| chr6-16728772-16729273 | 8 | 0.898859914572342 |
| chr17-44457726-44458227 | 8 | 0.855319623039782 |
| chr5-40597331-40597832 | 8 | 0.947595163337757 |
| chr10-22449829-22450330 | 6 | 0.991206925481043 |

| | | |
|---|---|---|
| chr16-48885153-48885654 | 6 | 0.980780894852084 |
| chr2-36790199-36790700 | 8 | 0.912987283292974 |
| chr8-99795255-99795756 | 8 | 0.958294312901557 |
| chr10-2458292-2458793 | 8 | 0.994426506656723 |
| chr5-79173859-79174360 | 8 | 0.986041652987846 |
| chr3-135565571-135566072 | 3 | 0.903299754506643 |
| chr2-177520735-177521236 | 7 | 0.8552978800742 |
| chr4-117470390-117470891 | 8 | 0.82606874387558 |
| chr7-119726102-119726603 | 8 | 0.971325007127966 |
| chr17-64321943-64322444 | 4 | 0.973149391957152 |
| chr7-130461000-130461501 | 8 | 0.926118505883385 |
| chr14-23304629-23305130 | 6 | 0.98275668422135 |
| chr1-235425849-235426350 | 4 | 0.972782509735644 |
| chr3-154458079-154458580 | 8 | 0.98644954682325 |
| chr3-32466800-32467301 | 3 | 0.89973382655992 |
| chr17-3923026-3923527 | 8 | 0.995226110969451 |
| chr4-91397474-91397975 | 8 | 0.900441404644014 |
| chr3-109859939-109860440 | 8 | 0.994448894157423 |
| chr2-204291591-204292092 | 8 | 0.977784636118179 |
| chr19-39725861-39726362 | 5 | 0.977047201116214 |
| chr10-3382776-3383277 | 8 | 0.947265407641069 |
| chr16-42070982-42071483 | 8 | 0.955293143284732 |
| chr5-94127041-94127542 | 8 | 0.993493681128311 |
| chr3-99367347-99367848 | 3 | 0.995115267086474 |
| chr11-82028651-82029152 | 3 | 0.99593553545041 |
| chr19-5990591-5991092 | 6 | 0.991206925481043 |
| chr4-73785333-73785834 | 8 | 0.833452151875774 |
| chr16-80049729-80050230 | 8 | 0.853016682004633 |
| chr1-187188847-187189348 | 4 | 0.93380416315498 |
| chr5-112233058-112233559 | 8 | 0.984344004488571 |
| chrX-62068197-62068698 | 3 | 0.982432984636016 |
| chr1-31554236-31554737 | 8 | 0.981031546971843 |
| chr1-241349181-241349682 | 8 | 0.986671165955835 |
| chr14-76730701-76731202 | 8 | 0.994009473154952 |
| chr5-113816378-113816879 | 8 | 0.976796226832313 |
| chr8-58755518-58756019 | 8 | 0.995136137150926 |
| chr7-134728569-134729070 | 5 | 0.983919978840557 |
| chr1-255882782-255883283 | 8 | 0.969720553769812 |
| chr20-43340343-43340844 | 8 | 0.993493681128311 |
| chr5-38310697-38311198 | 5 | 0.972715181061944 |
| chr13-39194877-39195378 | 8 | 0.854512391287587 |
| chr9-75838830-75839331 | 8 | 0.948059132326914 |
| chr10-52365031-52365532 | 8 | 0.938935001113004 |

| | | |
|---|---|---|
| chr9-35871262-35871763 | 3 | 0.980041133669309 |
| chr2-31349443-31349944 | 8 | 0.991023974067153 |
| chr9-25207136-25207637 | 6 | 0.992960515348647 |
| chr10-19406447-19406948 | 8 | 0.887219075987935 |
| chr7-25989507-25990008 | 3 | 0.968506019889791 |
| chr19-40227906-40228407 | 8 | 0.981351294786044 |
| chr3-148659985-148660486 | 2 | 0.852169998663572 |
| chr18-50157708-50158209 | 5 | 0.955510438965104 |
| chr2-47932623-47933124 | 8 | 0.879837446782094 |
| chr8-50640374-50640875 | 8 | 0.978020666372727 |
| chr13-72119218-72119719 | 8 | 0.99150442681455 |
| chr15-47272265-47272766 | 8 | 0.984594349932007 |
| chr1-185285716-185286217 | 8 | 0.997593491668901 |
| chr11-77255006-77255507 | 4 | 0.887529313996155 |
| chr16-36514698-36515199 | 3 | 0.995589215489615 |
| chr2-42148941-42149442 | 5 | 0.937099494030916 |
| chr4-26335380-26335881 | 8 | 0.993493681128311 |
| chr10-20080984-20081485 | 8 | 0.958669097824102 |
| chr19-37821801-37822302 | 8 | 0.877615332936822 |
| chr7-19048516-19049017 | 2 | 0.877387274306605 |
| chr5-146427581-146428082 | 8 | 0.91478639653139 |
| chr2-103364367-103364868 | 5 | 0.955425046197898 |
| chr6-132345627-132346128 | 8 | 0.91346320620339 |
| chr3-29633570-29634071 | 8 | 0.93838376695166 |
| chr1-118204556-118205057 | 6 | 0.988668779295247 |
| chr2-86852865-86853366 | 8 | 0.976328826726365 |
| chr4-146245697-146246198 | 5 | 0.978388164560919 |
| chr17-32998434-32998935 | 5 | 0.862050308684708 |
| chr7-22019574-22020075 | 8 | 0.870169456943639 |
| chr3-131156072-131156573 | 4 | 0.881778591603411 |
| chr8-87793396-87793897 | 8 | 0.982378550172312 |
| chr2-194003781-194004282 | 6 | 0.976011313778062 |
| chr2-104811724-104812225 | 4 | 0.849845256005865 |
| chr13-85783894-85784395 | 3 | 0.886713336683204 |
| chr5-59114165-59114666 | 8 | 0.989949518853892 |
| chr4-25443853-25444354 | 5 | 0.988798815601399 |
| chr4-94137568-94138069 | 5 | 0.932122334570572 |
| chr2-32976305-32976806 | 8 | 0.969567079760546 |
| chr3-157088255-157088756 | 8 | 0.898330951508556 |
| chr1-213155088-213155589 | 4 | 0.972018307182889 |
| chr10-28534315-28534816 | 2 | 0.929391378092201 |
| chr7-107815099-107815600 | 8 | 0.923203819968444 |
| chr15-8991234-8991735 | 8 | 0.848480381697882 |

| | | |
|---|---|---|
| chr14-8861931-8862432 | 5 | 0.989742261725034 |
| chr9-17769117-17769618 | 8 | 0.851907925776301 |
| chr1-220292926-220293427 | 4 | 0.902891327549414 |
| chr5-40732828-40733329 | 6 | 0.988749658348134 |
| chr4-134246122-134246623 | 8 | 0.952007890349538 |
| chr14-39760530-39761031 | 5 | 0.939582256475751 |
| chr6-140907443-140907944 | 6 | 0.987943431927432 |
| chr14-71959083-71959584 | 8 | 0.993493681128311 |
| chr17-23747141-23747642 | 8 | 0.987386471994442 |
| chr9-94093032-94093533 | 8 | 0.98938942803412 |
| chr2-242361466-242361967 | 1 | 0.936571849083948 |
| chr3-8541202-8541703 | 5 | 0.986624364273103 |
| chr4-110937810-110938311 | 6 | 0.984243028497153 |
| chr16-60254206-60254707 | 8 | 0.961273653604726 |
| chr5-83136001-83136502 | 2 | 0.831470045323679 |
| chr1-175114636-175115137 | 5 | 0.9042352212095 |
| chr4-175850724-175851225 | 5 | 0.978389486027638 |
| chr12-9614747-9615248 | 4 | 0.919064472167296 |
| chr16-79147085-79147586 | 8 | 0.993062315004375 |
| chr2-231077399-231077900 | 8 | 0.926102367750125 |
| chr2-40095941-40096442 | 5 | 0.991107236136677 |
| chr7-124354020-124354521 | 8 | 0.957081518666892 |
| chr2-196348189-196348690 | 8 | 0.892567802918393 |
| chr1-194360381-194360882 | 8 | 0.980282878183449 |
| chr4-53464351-53464852 | 8 | 0.965029801739878 |
| chr1-25101530-25102031 | 8 | 0.995234619729435 |
| chr7-47235810-47236311 | 8 | 0.910953602858625 |
| chr2-23569719-23570220 | 8 | 0.885694329095467 |
| chr5-71771290-71771791 | 8 | 0.986133971448093 |
| chr5-46055692-46056193 | 3 | 0.996889473266117 |
| chr1-195381027-195381528 | 3 | 0.994024728731511 |
| chr14-23386752-23387253 | 2 | 0.90197781323599 |
| chr2-175472013-175472514 | 8 | 0.868821433732564 |
| chr1-153791992-153792493 | 8 | 0.951399067076782 |
| chr4-140187988-140188489 | 8 | 0.902980187814802 |
| chr4-100872502-100873003 | 3 | 0.997208801479563 |
| chr5-72819700-72820201 | 8 | 0.965185661000527 |
| chr3-120554827-120555328 | 8 | 0.928084502963217 |
| chr3-89892738-89893239 | 8 | 0.977490468702875 |
| chrX-68580814-68581315 | 8 | 0.977449139477779 |
| chr1-26620482-26620983 | 5 | 0.940844824501542 |
| chr5-118944497-118944998 | 5 | 0.990990210321709 |
| chr1-195957793-195958294 | 8 | 0.832430472593866 |

| | | |
|---|---|---|
| chr6-58198855-58199356 | 3 | 0.862478216502733 |
| chr6-48714563-48715064 | 8 | 0.972417375178179 |
| chr17-66250290-66250791 | 8 | 0.95236037096816 |
| chr14-9244089-9244590 | 8 | 0.988715575166801 |
| chr17-23279411-23279912 | 8 | 0.972434576109368 |
| chr3-44481254-44481755 | 5 | 0.987480416501996 |
| chr3-138021717-138022218 | 8 | 0.989370741344494 |
| chr13-71537052-71537553 | 8 | 0.990855547396907 |
| chr2-4724381-4724882 | 3 | 0.997727862294445 |
| chr1-219590588-219591089 | 4 | 0.882591485944827 |
| chr11-5619295-5619796 | 3 | 0.996449610030369 |
| chr3-52302748-52303249 | 8 | 0.915105313741886 |
| chr2-211998377-211998878 | 5 | 0.872957764623972 |
| chr14-8654746-8655247 | 3 | 0.983283015404493 |
| chr13-47740949-47741450 | 8 | 0.953552588981615 |
| chr5-20835754-20836255 | 2 | 0.856085506726653 |
| chr2-6685346-6685847 | 3 | 0.994351800634578 |
| chrX-59046805-59047306 | 4 | 0.956639551667551 |
| chr4-80737580-80738081 | 8 | 0.976960965714375 |
| chr9-61486591-61487092 | 8 | 0.97158506802466 |
| chr2-5764546-5765047 | 4 | 0.903954461607674 |
| chr5-5092096-5092597 | 8 | 0.939872053477993 |
| chr7-75015567-75016068 | 3 | 0.997117461780804 |
| chr10-97754827-97755328 | 8 | 0.993493681128311 |
| chr1-136237282-136237783 | 2 | 0.951018945679233 |
| chr8-69418215-69418716 | 3 | 0.996601686512012 |
| chr1-150977487-150977988 | 8 | 0.973547933758175 |
| chr5-27067712-27068213 | 8 | 0.993493681128311 |
| chr17-22180481-22180982 | 5 | 0.862298715190674 |
| chr16-42911036-42911537 | 5 | 0.987486397981072 |
| chr8-95171710-95172211 | 8 | 0.833018473781452 |
| chr4-21546383-21546884 | 8 | 0.87596222481436 |
| chr16-73381613-73382114 | 8 | 0.99255130732886 |
| chr1-199322718-199323219 | 8 | 0.886727769391796 |
| chr15-40147035-40147536 | 8 | 0.96802081975027 |
| chr1-119292408-119292909 | 8 | 0.976611620034739 |
| chr1-49566234-49566735 | 8 | 0.993493681128311 |
| chr3-148740149-148740650 | 8 | 0.878366742218583 |
| chr1-127508227-127508728 | 8 | 0.978716446007393 |
| chr13-72577060-72577561 | 4 | 0.824695525194131 |
| chr15-93345104-93345605 | 8 | 0.889409929345478 |
| chr18-32069706-32070207 | 2 | 0.95619394413052 |
| chr3-79924339-79924840 | 3 | 0.975644816920201 |

| | | |
|---|---|---|
| chr13-73351142-73351643 | 8 | 0.949449967105639 |
| chr19-17905208-17905709 | 8 | 0.981947273445979 |
| chr9-4077887-4078388 | 8 | 0.840295543048791 |
| chr8-123746202-123746703 | 6 | 0.973580216150989 |
| chrX-115031102-115031603 | 3 | 0.940603413540857 |
| chr14-92247865-92248366 | 8 | 0.916919326525097 |
| chr17-40546429-40546930 | 5 | 0.987007714760693 |
| chr16-13831437-13831938 | 3 | 0.996796548641911 |
| chr18-12925962-12926463 | 8 | 0.956965979279883 |
| chr15-37483874-37484375 | 5 | 0.989756427929816 |
| chr4-134247549-134248050 | 8 | 0.944412116880129 |
| chr10-79757939-79758440 | 8 | 0.985476561882346 |
| chr18-28047063-28047564 | 4 | 0.917700443887957 |
| chr6-51024161-51024662 | 8 | 0.993493681128311 |
| chr13-88592279-88592780 | 2 | 0.857957955545819 |
| chr5-144080036-144080537 | 8 | 0.906339213343649 |
| chr11-3014249-3014750 | 3 | 0.996587718391151 |
| chr5-47274444-47274945 | 8 | 0.98438661779634 |
| chr7-4094849-4095350 | 3 | 0.99593553545041 |
| chr7-123459291-123459792 | 8 | 0.995170737596269 |
| chr8-123632707-123633208 | 3 | 0.994554104199563 |
| chr9-72589128-72589629 | 8 | 0.916653752573961 |
| chr7-118140739-118141240 | 8 | 0.90124589934167 |
| chr8-14599670-14600171 | 4 | 0.967718893084666 |
| chr6-41871738-41872239 | 8 | 0.926598130600096 |
| chr18-10852387-10852888 | 3 | 0.997653774424685 |
| chr1-2358646-2359147 | 4 | 0.973197030329316 |
| chr9-30092883-30093384 | 6 | 0.96330426076648 |
| chr9-26698690-26699191 | 8 | 0.825560571587631 |
| chr3-88927633-88928134 | 8 | 0.993493681128311 |
| chr6-113992676-113993177 | 5 | 0.986763118717602 |
| chr7-114993731-114994232 | 8 | 0.98692451748583 |
| chr6-76857853-76858354 | 8 | 0.888252758755245 |
| chr4-110600613-110601114 | 4 | 0.973092766944795 |
| chr16-75277793-75278294 | 8 | 0.994708154386378 |
| chr8-112663316-112663817 | 5 | 0.943481565831245 |
| chr3-103600060-103600561 | 8 | 0.923715906875068 |
| chr20-18068773-18069274 | 8 | 0.985629072950892 |
| chr7-99637446-99637947 | 4 | 0.899940168094775 |
| chr11-56657747-56658248 | 4 | 0.838227501871757 |
| chr15-95732483-95732984 | 6 | 0.991891400993514 |
| chr13-103714473-103714974 | 8 | 0.989681989547947 |
| chr7-94792872-94793373 | 8 | 0.990268304282225 |

| | | |
|---|---|---|
| chr13-88147925-88148426 | 5 | 0.984744001217947 |
| chr6-124360216-124360717 | 8 | 0.98968317204492 |
| chr3-141362511-141363012 | 8 | 0.98254858904949 |
| chr17-33175221-33175722 | 8 | 0.910117760393661 |
| chr1-172171523-172172024 | 8 | 0.910087481750054 |
| chr1-172887183-172887684 | 7 | 0.885388513997223 |
| chr2-203795086-203795587 | 8 | 0.995244857904211 |
| chr15-35995079-35995580 | 8 | 0.993493681128311 |
| chr1-228145854-228146355 | 8 | 0.994707616137687 |
| chr9-51286831-51287332 | 3 | 0.998356947255655 |
| chr14-2638095-2638596 | 3 | 0.990815619435517 |
| chr9-59084499-59085000 | 5 | 0.98753988353701 |
| chr2-111490077-111490578 | 8 | 0.976371324718715 |
| chr11-154378-154879 | 3 | 0.985426333951999 |
| chr9-84694707-84695208 | 1 | 0.899847559095236 |
| chr4-33286114-33286615 | 7 | 0.840518383371527 |
| chr6-128986068-128986569 | 2 | 0.841581778160342 |
| chr4-46004068-46004569 | 5 | 0.956141854914371 |
| chr14-14748539-14749040 | 3 | 0.969524114334655 |
| chr1-254307907-254308408 | 8 | 0.952504895006161 |
| chr18-69340003-69340504 | 5 | 0.917389305078691 |
| chr9-18736580-18737081 | 8 | 0.994306336293576 |
| chr16-73185262-73185763 | 8 | 0.815659997387167 |
| chr6-62627813-62628314 | 6 | 0.991206925481043 |
| chr14-68619652-68620153 | 8 | 0.85917359426711 |
| chr6-103513616-103514117 | 5 | 0.847773498974898 |
| chr6-112114020-112114521 | 8 | 0.993784770701893 |
| chr13-37203405-37203906 | 8 | 0.974615465179256 |
| chr12-34607772-34608273 | 5 | 0.969336332188318 |
| chr7-86763360-86763861 | 8 | 0.927525409783515 |
| chr12-37757848-37758349 | 2 | 0.89381381159146 |
| chr3-41648104-41648605 | 8 | 0.936741863270745 |
| chr13-35794538-35795039 | 5 | 0.968371431787072 |
| chr1-154768907-154769408 | 8 | 0.98749450095236 |
| chr13-92213181-92213682 | 8 | 0.889466502166666 |
| chr5-23466467-23466968 | 3 | 0.992555423711431 |
| chr3-1187336-1187837 | 4 | 0.904632277934376 |
| chr4-108796760-108797261 | 5 | 0.969679304991991 |
| chr2-23298883-23299384 | 1 | 0.910019057002859 |
| chr14-8780622-8781123 | 8 | 0.869878398247442 |
| chr13-100656929-100657430 | 8 | 0.980092649880924 |
| chr2-239390285-239390786 | 5 | 0.957048675889161 |
| chr3-22379958-22380459 | 8 | 0.951799275848654 |

| | | |
|---|---|---|
| chr7-125208913-125209414 | 8 | 0.991069898949423 |
| chr16-80079994-80080495 | 3 | 0.994916135020141 |
| chr14-80919525-80920026 | 8 | 0.970343295635986 |
| chr5-15084261-15084762 | 5 | 0.843238409102244 |
| chr18-10510015-10510516 | 3 | 0.99593553545041 |
| chr1-66731627-66732128 | 6 | 0.991206925481043 |
| chr14-20860006-20860507 | 2 | 0.938516135080557 |
| chr14-1510353-1510854 | 8 | 0.993493681128311 |
| chr2-170142488-170142989 | 5 | 0.915888785121843 |
| chr9-11451115-11451616 | 4 | 0.858009830986946 |
| chr12-44114194-44114695 | 4 | 0.929479965389506 |
| chr1-231212606-231213107 | 8 | 0.929896509340891 |
| chr6-132495804-132496305 | 4 | 0.942404321074179 |
| chr10-100198611-100199112 | 3 | 0.996409351223389 |
| chrX-134235279-134235780 | 5 | 0.836074690779095 |
| chr14-62340163-62340664 | 5 | 0.983058240430088 |
| chr3-148785658-148786159 | 8 | 0.992353365074201 |
| chr4-127850839-127851340 | 4 | 0.946430511809989 |
| chr8-62335889-62336390 | 5 | 0.982913148282404 |
| chr3-117052599-117053100 | 6 | 0.991127779725222 |
| chr19-34216442-34216943 | 2 | 0.909382519954645 |
| chr2-215969534-215970035 | 8 | 0.916088731263242 |
| chr18-52206144-52206645 | 8 | 0.852463613661319 |
| chr2-22287043-22287544 | 8 | 0.968440822362375 |
| chr2-153228774-153229275 | 8 | 0.948073158845294 |
| chr13-99235014-99235515 | 8 | 0.90593990182315 |
| chr4-61847963-61848464 | 4 | 0.892662921419228 |
| chr11-61344743-61345244 | 6 | 0.981078908458333 |
| chr14-15492211-15492712 | 8 | 0.918610357517895 |
| chr16-21514127-21514628 | 8 | 0.896250037758315 |
| chr6-18927190-18927691 | 5 | 0.983919978840557 |
| chr2-202929952-202930453 | 8 | 0.982146399603691 |
| chr17-48555639-48556140 | 8 | 0.952509663215233 |
| chr14-72678155-72678656 | 2 | 0.84422051380926 |
| chr1-132229674-132230175 | 8 | 0.827184577501792 |
| chr12-40149244-40149745 | 5 | 0.878568420431571 |
| chr1-54075401-54075902 | 4 | 0.845243503582362 |
| chr2-34410517-34411018 | 5 | 0.985669747112596 |
| chr11-31378488-31378989 | 2 | 0.969105328079536 |
| chr6-124546032-124546533 | 8 | 0.970372584961884 |
| chr11-72736093-72736594 | 8 | 0.979137050558308 |
| chr15-82485098-82485599 | 2 | 0.955795030047071 |
| chr11-9145077-9145578 | 8 | 0.993493681128311 |

| | | |
|---|---|---|
| chr15-38529789-38530290 | 8 | 0.974176538376917 |
| chr4-93161132-93161633 | 6 | 0.990795609206762 |
| chr20-10030719-10031220 | 3 | 0.975340377886964 |
| chr17-69062341-69062842 | 4 | 0.96290167937564 |
| chr13-59628643-59629144 | 8 | 0.882114104846671 |
| chr10-69088269-69088770 | 8 | 0.995045275869992 |
| chr2-202505854-202506355 | 8 | 0.993493681128311 |
| chr10-61471244-61471745 | 4 | 0.929556858634928 |
| chr9-57131418-57131919 | 8 | 0.993493681128311 |
| chr18-5365355-5365856 | 8 | 0.970612395890182 |
| chr8-118272961-118273462 | 8 | 0.860633152111011 |
| chr8-119549685-119550186 | 5 | 0.974683798334283 |
| chr5-26665029-26665530 | 4 | 0.920851444054682 |
| chr17-23082837-23083338 | 3 | 0.982928698107196 |
| chr1-20770960-20771461 | 8 | 0.866325343191192 |
| chr1-129872346-129872847 | 8 | 0.993493681128311 |
| chr6-34786267-34786768 | 3 | 0.847312084641007 |
| chr8-119668017-119668518 | 8 | 0.937182489033448 |
| chr6-89186510-89187011 | 7 | 0.945063235489398 |
| chr15-12814707-12815208 | 5 | 0.94692672723917 |
| chr2-21012795-21013296 | 8 | 0.95050947806329 |
| chr3-79580638-79581139 | 6 | 0.995092248910754 |
| chr6-102144090-102144591 | 5 | 0.962231213738644 |
| chr12-12847677-12848178 | 8 | 0.834072872257019 |
| chr1-180864927-180865428 | 8 | 0.892768424724505 |
| chr1-260181944-260182445 | 8 | 0.905889159912882 |
| chr5-135655301-135655802 | 4 | 0.906173054624618 |
| chr19-40135306-40135807 | 5 | 0.979394383132112 |
| chr8-111744455-111744956 | 3 | 0.996892952554903 |
| chr15-34750170-34750671 | 8 | 0.978916303859272 |
| chr5-71087857-71088358 | 8 | 0.993493681128311 |
| chr4-84209462-84209963 | 8 | 0.966144884470605 |
| chr17-1637753-1638254 | 8 | 0.847400676207783 |
| chr9-35861987-35862488 | 4 | 0.837565141570708 |
| chr9-49662747-49663248 | 5 | 0.991509713922577 |
| chr4-179646469-179646970 | 8 | 0.844378294820399 |
| chr4-173220714-173221215 | 8 | 0.938207003342306 |
| chr1-191157343-191157844 | 8 | 0.915761248564122 |
| chr1-195332323-195332824 | 8 | 0.98329844599469 |
| chr9-55918651-55919152 | 8 | 0.941683422283655 |
| chr16-32498442-32498943 | 4 | 0.874551452978789 |
| chr18-67827557-67828058 | 8 | 0.994903199620014 |
| chr17-39443297-39443798 | 8 | 0.98994066383429 |

| | | |
|---|---|---|
| chr4-150339696-150340197 | 8 | 0.989000117425161 |
| chr13-87865545-87866046 | 2 | 0.900144735352913 |
| chr18-69784984-69785485 | 5 | 0.99509930211202 |
| chr3-69621715-69622216 | 8 | 0.857451481476837 |
| chr14-32462509-32463010 | 8 | 0.819291341262136 |
| chr17-23027119-23027620 | 8 | 0.990193187481381 |
| chr14-6093686-6094187 | 8 | 0.982264923225342 |
| chr9-49068195-49068696 | 4 | 0.930852781646341 |
| chr3-129896027-129896528 | 8 | 0.827798810445192 |
| chr8-22167181-22167682 | 8 | 0.992709380162339 |
| chr1-161498064-161498565 | 8 | 0.995197615524698 |
| chr9-83796321-83796822 | 8 | 0.993493681128311 |
| chr12-27537499-27538000 | 8 | 0.993493681128311 |
| chr9-112770511-112771012 | 8 | 0.860333145332024 |
| chr2-171697352-171697853 | 8 | 0.94005866634093 |
| chr4-131216149-131216650 | 8 | 0.989681809562172 |
| chr3-104304440-104304941 | 8 | 0.97913799573341 |
| chr5-73719685-73720186 | 8 | 0.993493681128311 |
| chr6-119669544-119670045 | 5 | 0.990097300329069 |
| chr8-50554498-50554999 | 8 | 0.91648689515937 |
| chr17-70974533-70975034 | 4 | 0.949355507873186 |
| chr15-38255260-38255761 | 6 | 0.973492575843857 |
| chr19-41470912-41471413 | 5 | 0.914227435968239 |
| chr4-126862317-126862818 | 8 | 0.955536017578567 |
| chr3-69552464-69552965 | 8 | 0.972505178969434 |
| chr10-62000034-62000535 | 8 | 0.985814535137423 |
| chr12-11190685-11191186 | 3 | 0.99867141494322 |
| chr2-108554239-108554740 | 3 | 0.99593553545041 |
| chr11-15153914-15154415 | 8 | 0.993493681128311 |
| chr13-70337763-70338264 | 5 | 0.868553386677645 |
| chr6-4052837-4053338 | 3 | 0.996619482834718 |
| chr11-65389053-65389554 | 8 | 0.892224927364163 |
| chr5-33557081-33557582 | 3 | 0.992724327069012 |
| chr2-142029867-142030368 | 4 | 0.966844425863033 |
| chr3-32440421-32440922 | 8 | 0.975413001064816 |
| chr5-62055965-62056466 | 3 | 0.937951411280092 |
| chr18-62343952-62344453 | 8 | 0.822207468585637 |
| chr2-107695077-107695578 | 8 | 0.988400388893997 |
| chr8-32070332-32070833 | 3 | 0.998172036046529 |
| chr1-102908983-102909484 | 3 | 0.99593553545041 |
| chr2-58737693-58738194 | 6 | 0.993242287861103 |
| chr2-53519889-53520390 | 8 | 0.92184327377745 |
| chr2-125244737-125245238 | 8 | 0.958242023403238 |

| | | |
|---|---|---|
| chr1-195366489-195366990 | 8 | 0.993493681128311 |
| chr18-22262021-22262522 | 8 | 0.87853327279702 |
| chr11-77292685-77293186 | 2 | 0.828057245044854 |
| chr12-45833832-45834333 | 3 | 0.928061982603035 |
| chr1-49038671-49039172 | 8 | 0.833354284399151 |
| chr3-64659812-64660313 | 8 | 0.969298231937199 |
| chr4-37018854-37019355 | 8 | 0.896772415196684 |
| chr3-92023687-92024188 | 8 | 0.962210579718716 |
| chr11-27958044-27958545 | 8 | 0.917616825583045 |
| chrX-129056015-129056516 | 8 | 0.948783283262217 |
| chr7-53887929-53888430 | 7 | 0.934125205006976 |
| chr9-35269339-35269840 | 5 | 0.983272801910197 |
| chr7-92134382-92134883 | 8 | 0.931358494868734 |
| chr5-11143708-11144209 | 5 | 0.965319794858862 |
| chr14-95160824-95161325 | 8 | 0.840805138743124 |
| chr7-24717748-24718249 | 3 | 0.989828543975978 |
| chr11-56869211-56869712 | 8 | 0.993925559987382 |
| chr3-29948495-29948996 | 3 | 0.994496280723504 |
| chr10-6862170-6862671 | 8 | 0.961458597320476 |
| chr2-57319275-57319776 | 4 | 0.839094555314917 |
| chr5-14881388-14881889 | 5 | 0.983665748898355 |
| chr18-57731664-57732165 | 6 | 0.939054843547942 |
| chr6-14670837-14671338 | 8 | 0.993493681128311 |
| chr3-1530060-1530561 | 2 | 0.83500572578278 |
| chrX-3975044-3975545 | 7 | 0.897365615566022 |
| chr16-14289842-14290343 | 8 | 0.920325301183423 |
| chr6-5244315-5244816 | 3 | 0.996194099168881 |
| chr2-103516760-103517261 | 6 | 0.951596203530361 |
| chr11-54780723-54781224 | 2 | 0.838698235277628 |
| chr12-8117437-8117938 | 8 | 0.940573063716861 |
| chr10-64973084-64973585 | 8 | 0.938386415662008 |
| chr1-73578704-73579205 | 8 | 0.888538923536057 |
| chr9-66737740-66738241 | 8 | 0.995245322370131 |
| chr1-225519179-225519680 | 8 | 0.877298766907466 |
| chr1-157877385-157877886 | 8 | 0.898032570230078 |
| chr15-74659882-74660383 | 4 | 0.831402877543975 |
| chr13-39426908-39427409 | 8 | 0.854550508090931 |
| chr5-124021616-124022117 | 5 | 0.992686274712139 |
| chr4-144294634-144295135 | 8 | 0.992864364161055 |
| chr4-67489935-67490436 | 8 | 0.824926367422406 |
| chr1-244958505-244959006 | 8 | 0.979454522207512 |
| chr12-40453149-40453650 | 7 | 0.823698560124588 |
| chr6-137542674-137543175 | 3 | 0.999052256289986 |

| | | |
|---|---|---|
| chr14-41059397-41059898 | 5 | 0.983919978840557 |
| chr15-80080645-80081146 | 8 | 0.988885920287034 |
| chr6-5216333-5216834 | 8 | 0.963659656798271 |
| chr10-50235232-50235733 | 8 | 0.993493681128311 |
| chr10-100419303-100419804 | 3 | 0.923571566439035 |
| chr13-76502753-76503254 | 2 | 0.949934186155363 |
| chr4-156598659-156599160 | 8 | 0.964244498817228 |
| chr6-21235948-21236449 | 8 | 0.928798719096868 |
| chr9-111896685-111897186 | 8 | 0.834636208375751 |
| chr1-257362145-257362646 | 8 | 0.987748236647807 |
| chr6-122016889-122017390 | 8 | 0.962159156419904 |
| chr8-37339414-37339915 | 6 | 0.996636619220719 |
| chr4-41515084-41515585 | 4 | 0.899862573668634 |
| chr17-16215242-16215743 | 8 | 0.856099925972898 |
| chr4-177639728-177640229 | 8 | 0.840352409881026 |
| chr4-123061713-123062214 | 8 | 0.883839671321301 |
| chrX-83850997-83851498 | 3 | 0.99593553545041 |
| chr5-77103699-77104200 | 8 | 0.981244940190898 |
| chr8-53073149-53073650 | 3 | 0.984806403350213 |
| chrX-107734214-107734715 | 8 | 0.967289424033164 |
| chr1-6283922-6284423 | 3 | 0.942921305527315 |
| chr14-18978374-18978875 | 8 | 0.953844699112577 |
| chr7-75685059-75685560 | 8 | 0.839861772590642 |
| chr1-91159889-91160390 | 8 | 0.990468313256826 |
| chr11-13060273-13060774 | 4 | 0.818320285988664 |
| chr8-16801714-16802215 | 3 | 0.99593553545041 |
| chr11-49482009-49482510 | 3 | 0.931727448083058 |
| chr9-75768478-75768979 | 8 | 0.973391281740109 |
| chr4-48006885-48007386 | 8 | 0.907586411042137 |
| chr1-43090598-43091099 | 2 | 0.945841185305884 |
| chr14-3781094-3781595 | 8 | 0.995145671125415 |
| chrX-52122274-52122775 | 6 | 0.977356706395403 |
| chr3-162736519-162737020 | 8 | 0.986933720465936 |
| chr6-128044017-128044518 | 8 | 0.962162278539638 |
| chrX-129523908-129524409 | 3 | 0.99593553545041 |
| chr8-58821211-58821712 | 8 | 0.826929114767576 |
| chr7-123756801-123757302 | 3 | 0.996608256386519 |
| chr2-57585800-57586301 | 8 | 0.991209460854585 |
| chr4-159544810-159545311 | 4 | 0.945059956531649 |
| chr8-9256579-9257080 | 3 | 0.977435949493353 |
| chr8-112093752-112094253 | 5 | 0.902759077650403 |
| chr4-120197992-120198493 | 5 | 0.984711939008854 |
| chr13-22664595-22665096 | 8 | 0.972431807833106 |

| | | |
|---|---|---|
| chr10-16903863-16904364 | 8 | 0.973187069314769 |
| chr2-22142053-22142554 | 8 | 0.957664755787784 |
| chr1-100758176-100758677 | 8 | 0.993493681128311 |
| chr4-47654635-47655136 | 2 | 0.891582635944856 |
| chr8-47150516-47151017 | 2 | 0.973152500549613 |
| chr8-11694751-11695252 | 8 | 0.907405121108934 |
| chr2-197105894-197106395 | 8 | 0.980518827363234 |
| chr13-30287968-30288469 | 6 | 0.996701694359551 |
| chr8-5806373-5806874 | 8 | 0.898277143561298 |
| chr8-122040843-122041344 | 8 | 0.989706322974721 |
| chr10-53889220-53889721 | 8 | 0.908121471616483 |
| chr20-22725723-22726224 | 3 | 0.99593553545041 |
| chr1-121504771-121505272 | 2 | 0.951774112407649 |
| chr14-35350815-35351316 | 3 | 0.827142502712153 |
| chr13-61699202-61699703 | 4 | 0.938904919941668 |
| chr14-102636731-102637232 | 8 | 0.993493681128311 |
| chr15-8360304-8360805 | 8 | 0.993493681128311 |
| chr19-25400285-25400786 | 3 | 0.99032631466901 |
| chrX-4242152-4242653 | 3 | 0.99318846791886 |
| chr5-46252410-46252911 | 2 | 0.95374006666477 |
| chr7-104558706-104559207 | 8 | 0.851119295777883 |
| chr8-60261030-60261531 | 8 | 0.990766940764856 |
| chr1-186850902-186851403 | 2 | 0.921147424802032 |
| chr17-44679467-44679968 | 8 | 0.925987692990005 |
| chr19-55347108-55347609 | 8 | 0.99406923527042 |
| chr18-44931622-44932123 | 3 | 0.99593553545041 |
| chr4-169530373-169530874 | 5 | 0.973643277956719 |
| chr9-14540795-14541296 | 8 | 0.971473281532358 |
| chr16-47729626-47730127 | 8 | 0.986071196854441 |
| chrX-58038928-58039429 | 6 | 0.987310746232134 |
| chr4-22604336-22604837 | 2 | 0.853444480834495 |
| chr3-36661478-36661979 | 8 | 0.946637374707044 |
| chr16-23587530-23588031 | 3 | 0.99593553545041 |
| chr7-99874954-99875455 | 8 | 0.949649169390065 |
| chr3-53124220-53124721 | 8 | 0.948734035385746 |
| chr8-85664104-85664605 | 3 | 0.996802242460906 |
| chr7-40471593-40472094 | 8 | 0.898637374835651 |
| chr6-94675622-94676123 | 8 | 0.833528339818309 |
| chr3-161017475-161017976 | 8 | 0.944719621087493 |
| chrX-127489629-127490130 | 5 | 0.905437890535992 |
| chr7-12362527-12363028 | 3 | 0.99593553545041 |
| chr6-135512793-135513294 | 6 | 0.89462280603418 |
| chrX-111148079-111148580 | 3 | 0.858252961361629 |

| | | |
|---|---|---|
| chr11-29056283-29056784 | 7 | 0.866975522904234 |
| chr18-40994718-40995219 | 8 | 0.989532571026236 |
| chr19-10128783-10129284 | 5 | 0.820658437498207 |
| chr3-148348073-148348574 | 8 | 0.884733259954218 |
| chr15-37474441-37474942 | 8 | 0.994665066948594 |
| chr17-33103238-33103739 | 8 | 0.964045897248492 |
| chr7-83748934-83749435 | 6 | 0.991206925481043 |
| chr9-66623450-66623951 | 5 | 0.990282391407581 |
| chr4-60308040-60308541 | 5 | 0.990989533140349 |
| chr1-218611889-218612390 | 5 | 0.98434476758332 |
| chr20-20027021-20027522 | 5 | 0.983433621527859 |
| chr9-23053464-23053965 | 8 | 0.991976761527474 |
| chr2-172378082-172378583 | 7 | 0.94995153207147 |
| chr14-16343925-16344426 | 8 | 0.967434290439184 |
| chr7-26984565-26985066 | 8 | 0.993493681128311 |
| chr18-40001573-40002074 | 8 | 0.897826096937321 |
| chr8-59989960-59990461 | 8 | 0.914540177428346 |
| chr14-46697328-46697829 | 3 | 0.99593553545041 |
| chr4-26112733-26113234 | 8 | 0.870012352549767 |
| chr1-138407994-138408495 | 8 | 0.985368744453869 |
| chr13-50991022-50991523 | 5 | 0.987351597337955 |
| chr5-99385226-99385727 | 2 | 0.891234654800528 |
| chr6-5006377-5006878 | 3 | 0.99755963528053 |
| chr8-54901689-54902190 | 8 | 0.920905298829047 |
| chr4-176860926-176861427 | 6 | 0.991206925481043 |
| chr4-19988604-19989105 | 3 | 0.99593553545041 |
| chr7-76608107-76608608 | 5 | 0.930192192775451 |
| chr8-88117583-88118084 | 3 | 0.996849883710898 |
| chrX-124723991-124724492 | 3 | 0.99593553545041 |
| chr18-42061330-42061831 | 8 | 0.934438489752981 |
| chr9-24677203-24677704 | 8 | 0.974809923649819 |
| chr4-119320629-119321130 | 4 | 0.855418045941988 |
| chr9-63346385-63346886 | 8 | 0.967739317657001 |
| chr3-125407816-125408317 | 3 | 0.949660864328081 |
| chr15-41748976-41749477 | 3 | 0.90708570962275 |
| chr12-22282895-22283396 | 8 | 0.883615073113611 |
| chr2-125563707-125564208 | 8 | 0.973739307535086 |
| chr18-49702696-49703197 | 8 | 0.934205417481226 |
| chr20-28543694-28544195 | 4 | 0.877172318855155 |
| chr7-55980605-55981106 | 5 | 0.977177461218926 |
| chr2-138887701-138888202 | 4 | 0.90530583296605 |
| chr5-128272393-128272894 | 8 | 0.905371203299457 |
| chr18-60222572-60223073 | 8 | 0.991845607646511 |

| | | |
|---|---|---|
| chr16-52120808-52121309 | 6 | 0.991941923653329 |
| chr15-75613659-75614160 | 5 | 0.819581360755676 |
| chr15-1545594-1546095 | 6 | 0.994289520462387 |
| chr3-27969598-27970099 | 4 | 0.930349892878729 |
| chr7-89145904-89146405 | 8 | 0.956961481944757 |
| chr3-165947551-165948052 | 8 | 0.856852670205685 |
| chr1-120792522-120793023 | 8 | 0.98438637325481 |
| chr13-101848426-101848927 | 6 | 0.855454655341511 |
| chr5-28359715-28360216 | 2 | 0.922859801583443 |
| chr14-74586558-74587059 | 5 | 0.968354826481914 |
| chr1-23311944-23312445 | 4 | 0.823970806377373 |
| chr1-220758514-220759015 | 8 | 0.970585226375135 |
| chr7-115718469-115718970 | 8 | 0.868765942768361 |
| chr20-13307580-13308081 | 8 | 0.970914659347722 |
| chr15-56363585-56364086 | 5 | 0.987522498715189 |
| chr9-104776188-104776689 | 5 | 0.979868961910535 |
| chr3-146332985-146333486 | 8 | 0.821327208237109 |
| chr6-44325651-44326152 | 5 | 0.982821768246152 |
| chr4-108203254-108203755 | 2 | 0.938820302178407 |
| chr4-61688292-61688793 | 5 | 0.990250557893385 |
| chr2-20594525-20595026 | 5 | 0.837225917851817 |
| chr3-61329697-61330198 | 8 | 0.94783661861659 |
| chr8-113338048-113338549 | 8 | 0.892394588443247 |
| chr1-230860657-230861158 | 3 | 0.995986919380324 |
| chr4-130113012-130113513 | 8 | 0.995017563066434 |
| chr16-72334172-72334673 | 5 | 0.920589797815804 |
| chr13-26456629-26457130 | 3 | 0.99593553545041 |
| chr4-134454776-134455277 | 8 | 0.913415367771402 |
| chr11-77173070-77173571 | 6 | 0.990763001076158 |
| chr8-102215535-102216036 | 8 | 0.851076857273742 |
| chr15-71109941-71110442 | 2 | 0.924843132995854 |
| chr7-92206313-92206814 | 6 | 0.98683697692648 |
| chr8-107375724-107376225 | 8 | 0.993068989559867 |
| chr10-14125508-14126009 | 8 | 0.83695871804945 |
| chr7-85327482-85327983 | 3 | 0.996300131498452 |
| chr3-94126538-94127039 | 3 | 0.994860452106951 |
| chr10-22707638-22708139 | 3 | 0.996895202935007 |
| chr12-29469571-29470072 | 3 | 0.996885982657031 |
| chr1-17125494-17125995 | 8 | 0.896366401219033 |
| chr5-55188622-55189123 | 5 | 0.83249921742284 |
| chr6-23147154-23147655 | 8 | 0.912951252636793 |
| chr14-67419602-67420103 | 8 | 0.906655185161975 |
| chr7-31277421-31277922 | 3 | 0.99593553545041 |

| | | |
|---|---|---|
| chr7-43654623-43655124 | 5 | 0.899404770315813 |
| chr2-131734805-131735306 | 3 | 0.995328506845203 |
| chr15-25917451-25917952 | 5 | 0.916614631396624 |
| chr17-40591571-40592072 | 8 | 0.915017425969477 |
| chr5-55276072-55276573 | 8 | 0.954997498210187 |
| chr18-75876266-75876767 | 8 | 0.904854611166107 |
| chr19-9247344-9247845 | 8 | 0.882785599871785 |
| chr3-165896281-165896782 | 8 | 0.986999843336353 |
| chr2-165475747-165476248 | 8 | 0.97052233099161 |
| chr14-3061979-3062480 | 8 | 0.994630001201913 |
| chr19-47005423-47005924 | 5 | 0.972605361752505 |
| chr4-177695773-177696274 | 8 | 0.966630549243237 |
| chr2-19467939-19468440 | 3 | 0.975839522693296 |
| chr11-73792402-73792903 | 6 | 0.992972276095607 |
| chr2-107638324-107638825 | 2 | 0.916417535640213 |
| chr3-108775608-108776109 | 5 | 0.962836017238821 |
| chr17-66878841-66879342 | 8 | 0.993493681128311 |
| chr5-138641348-138641849 | 8 | 0.918321452538902 |
| chr8-80362354-80362855 | 8 | 0.87508369151014 |
| chr2-105802874-105803375 | 2 | 0.951026563573827 |
| chr9-20963221-20963722 | 3 | 0.992091318817476 |
| chr4-109812657-109813158 | 8 | 0.993493681128311 |
| chr10-35990814-35991315 | 6 | 0.991507233268228 |
| chr9-56270674-56271175 | 8 | 0.993493681128311 |
| chr6-44710483-44710984 | 4 | 0.8738591286582 |
| chr6-13291727-13292228 | 8 | 0.855536080892514 |
| chr10-72278451-72278952 | 8 | 0.951755304935738 |
| chr17-55457548-55458049 | 5 | 0.910738873112518 |
| chr2-40245752-40246253 | 8 | 0.978520516380676 |
| chr7-8008565-8009066 | 5 | 0.988984222522986 |
| chr16-27780041-27780542 | 8 | 0.897680506441217 |
| chr3-18010804-18011305 | 8 | 0.862309983531445 |
| chr7-34543405-34543906 | 2 | 0.931454383806756 |
| chr5-59198159-59198660 | 8 | 0.863965387682961 |
| chr18-44730828-44731329 | 3 | 0.996689420327779 |
| chr19-14358055-14358556 | 5 | 0.983919978840557 |
| chr3-32289146-32289647 | 8 | 0.993612001487948 |
| chr20-36041610-36042111 | 8 | 0.940820321269508 |
| chr2-112804743-112805244 | 8 | 0.993493681128311 |
| chr10-15878481-15878982 | 8 | 0.942167037263489 |
| chr2-43278372-43278873 | 8 | 0.991334112693885 |
| chr5-134812269-134812770 | 8 | 0.91344533493137 |
| chr1-253578684-253579185 | 8 | 0.980593684549631 |

| | | |
|---|---|---|
| chr18-50782190-50782691 | 8 | 0.989665330992189 |
| chr8-86247326-86247827 | 8 | 0.984993909004663 |
| chr11-64872671-64873172 | 4 | 0.97358866435118 |
| chr1-104873344-104873845 | 8 | 0.956118275874588 |
| chr7-112399966-112400467 | 5 | 0.990420356477973 |
| chr11-48013006-48013507 | 8 | 0.993493681128311 |
| chr5-54127100-54127601 | 2 | 0.94601660859439 |
| chr14-53611354-53611855 | 3 | 0.992409555405229 |
| chr8-96230463-96230964 | 8 | 0.958118629435981 |
| chr6-103568936-103569437 | 3 | 0.851437996738516 |
| chr7-10041941-10042442 | 5 | 0.953150936491584 |
| chr7-65269341-65269842 | 8 | 0.88505978862511 |
| chr1-197570290-197570791 | 2 | 0.856416667879679 |
| chr5-3141096-3141597 | 5 | 0.95346868331908 |
| chr2-76766958-76767459 | 3 | 0.999018529337669 |
| chr10-13322830-13323331 | 8 | 0.993493681128311 |
| chr3-93213022-93213523 | 8 | 0.994650352671266 |
| chr14-92322378-92322879 | 6 | 0.869593153755344 |
| chr8-67544861-67545362 | 8 | 0.951083630136502 |
| chr6-128997028-128997529 | 8 | 0.865479377981008 |
| chr18-790991-791492 | 4 | 0.909484246295585 |
| chr3-55968303-55968804 | 8 | 0.941569725172694 |
| chr13-106468375-106468876 | 5 | 0.977258548371429 |
| chr2-118652681-118653182 | 8 | 0.994774148512581 |
| chr11-34264918-34265419 | 8 | 0.944577525957757 |
| chr15-37584151-37584652 | 4 | 0.858141703700354 |
| chr20-42584051-42584552 | 5 | 0.923191107182878 |
| chr16-83289256-83289757 | 8 | 0.993493681128311 |
| chr11-59614015-59614516 | 2 | 0.903590873257284 |
| chr7-124191589-124192090 | 4 | 0.963149194437725 |
| chr11-75890263-75890764 | 5 | 0.87419844583006 |
| chr7-80805160-80805661 | 4 | 0.923536100063878 |
| chr10-42361253-42361754 | 8 | 0.984291152556328 |
| chr10-65676791-65677292 | 8 | 0.983274211431073 |
| chr11-15410736-15411237 | 8 | 0.938375626713971 |
| chr14-73135185-73135686 | 3 | 0.994681513058542 |
| chr1-113907569-113908070 | 8 | 0.88758093757485 |
| chr7-132740163-132740664 | 8 | 0.991601425411411 |
| chr1-218147822-218148323 | 8 | 0.973112631009339 |
| chr5-137599756-137600257 | 8 | 0.995205569750179 |
| chr9-10996426-10996927 | 8 | 0.82889369858712 |
| chr1-243476132-243476633 | 8 | 0.868761961052702 |
| chr12-43196137-43196638 | 3 | 0.99593553545041 |

| | | |
|---|---|---|
| chr9-55505452-55505953 | 8 | 0.830839544571388 |
| chr15-73160197-73160698 | 8 | 0.853927514623426 |
| chr7-123908062-123908563 | 5 | 0.989013242391662 |
| chr5-51735690-51736191 | 3 | 0.995825700061236 |
| chr2-57208346-57208847 | 4 | 0.934584177965112 |
| chrX-106976622-106977123 | 5 | 0.914677050504023 |
| chr15-33084148-33084649 | 8 | 0.884949164370557 |
| chr5-86952380-86952881 | 8 | 0.976162454991195 |
| chr5-46199156-46199657 | 6 | 0.983635943140885 |
| chr8-83169986-83170487 | 6 | 0.991206925481043 |
| chr19-20012320-20012821 | 4 | 0.960094370822618 |
| chr4-110571213-110571714 | 8 | 0.993084281105463 |
| chr5-26921337-26921838 | 8 | 0.995168126771719 |
| chr1-225790445-225790946 | 8 | 0.874058466943169 |
| chr1-32947054-32947555 | 8 | 0.995245381850691 |
| chr11-58259657-58260158 | 6 | 0.927966935473018 |
| chr16-79743172-79743673 | 8 | 0.858192765322786 |
| chr7-61295307-61295808 | 8 | 0.948325679550791 |
| chr18-11605000-11605501 | 8 | 0.957300285305672 |
| chr8-82854697-82855198 | 8 | 0.995249028520937 |
| chr2-231755329-231755830 | 5 | 0.868608291007303 |
| chr5-21095591-21096092 | 5 | 0.984745180736076 |
| chr4-175357545-175358046 | 8 | 0.896121813213407 |
| chr2-12130334-12130835 | 3 | 0.99593553545041 |
| chr7-10141388-10141889 | 8 | 0.9592710932252 |
| chr4-33211478-33211979 | 8 | 0.989807776939764 |
| chr7-37895082-37895583 | 4 | 0.851884302857702 |
| chr9-66429519-66430020 | 3 | 0.996879256694783 |
| chr20-6730854-6731355 | 8 | 0.989163323115131 |
| chr19-47849053-47849554 | 5 | 0.980766079427035 |
| chr1-223690303-223690804 | 4 | 0.975648750998006 |
| chr13-74890790-74891291 | 4 | 0.950290418363093 |
| chr2-105734855-105735356 | 8 | 0.993493681128311 |
| chr11-72149348-72149849 | 8 | 0.863579679370846 |
| chr5-32079511-32080012 | 8 | 0.895827790905729 |
| chr1-134800833-134801334 | 8 | 0.988336321433798 |
| chrX-91334164-91334665 | 3 | 0.99593553545041 |
| chr8-95683298-95683799 | 8 | 0.984799465283518 |
| chr10-98887718-98888219 | 6 | 0.9918067352214 |
| chr16-22524826-22525327 | 3 | 0.971791032077035 |
| chr1-88924304-88924805 | 8 | 0.957477232321468 |
| chr3-18412535-18413036 | 8 | 0.977034146783868 |
| chr11-63214767-63215268 | 8 | 0.857240575838203 |

| | | |
|---|---|---|
| chr16-43229656-43230157 | 8 | 0.90212761979629 |
| chr7-93417144-93417645 | 8 | 0.988743921038464 |
| chr2-248255885-248256386 | 8 | 0.990384674795114 |
| chrX-115409572-115410073 | 8 | 0.996059184965677 |
| chr15-76307768-76308269 | 8 | 0.978639961849407 |
| chr4-13277075-13277576 | 8 | 0.973040562576037 |
| chr6-51954917-51955418 | 2 | 0.962345564636915 |
| chr1-248898702-248899203 | 3 | 0.996853697748199 |
| chr13-2992102-2992603 | 5 | 0.886253353585551 |
| chr16-84562105-84562606 | 8 | 0.979263870274301 |
| chr4-147623345-147623846 | 3 | 0.919249887775882 |
| chr16-42809840-42810341 | 5 | 0.98875259719235 |
| chr6-84469484-84469985 | 8 | 0.994669704690393 |
| chr4-58729352-58729853 | 8 | 0.925105880480322 |
| chr3-35376827-35377328 | 6 | 0.986774482257358 |
| chr1-155591428-155591929 | 5 | 0.991468184458678 |
| chr9-28400228-28400729 | 3 | 0.982853044185104 |
| chr15-26565893-26566394 | 5 | 0.818945333495736 |
| chr16-44770925-44771426 | 5 | 0.985458744050545 |
| chr11-63028534-63029035 | 8 | 0.978169731029542 |
| chr17-70029849-70030350 | 3 | 0.990917093475424 |
| chr4-62694004-62694505 | 8 | 0.993493681128311 |
| chr14-39941085-39941586 | 5 | 0.938543302722327 |
| chr7-17863925-17864426 | 8 | 0.936619180651488 |
| chr4-85167692-85168193 | 8 | 0.941878039226562 |
| chr1-6541924-6542425 | 8 | 0.821734295975143 |
| chr1-127715748-127716249 | 8 | 0.993493681128311 |
| chr20-50411990-50412491 | 3 | 0.984088189543057 |
| chr8-108621706-108622207 | 2 | 0.887816402493904 |
| chr1-5974303-5974804 | 3 | 0.993678231975639 |
| chr15-54240903-54241404 | 5 | 0.968173222585535 |
| chr6-86097924-86098425 | 3 | 0.99593553545041 |
| chr2-136169260-136169761 | 8 | 0.913234098014071 |
| chr12-630801-631302 | 5 | 0.907539835772812 |
| chr14-61392265-61392766 | 5 | 0.943971924922113 |
| chr18-14374536-14375037 | 5 | 0.968261628557768 |
| chr1-251027722-251028223 | 3 | 0.996756906018556 |
| chr20-44225898-44226399 | 2 | 0.828806924435906 |
| chr6-25857868-25858369 | 8 | 0.984086609034068 |
| chr3-116990144-116990645 | 4 | 0.894345102371072 |
| chr2-38679757-38680258 | 6 | 0.991206925481043 |
| chr2-44573971-44574472 | 8 | 0.993493681128311 |
| chr18-75604318-75604819 | 8 | 0.883851645358078 |

| | | |
|---|---|---|
| chr14-64166896-64167397 | 8 | 0.837060398103395 |
| chr3-159380521-159381022 | 8 | 0.842071710312802 |
| chr2-195665518-195666019 | 8 | 0.958514903094979 |
| chr15-3040099-3040600 | 8 | 0.984825663440291 |
| chr5-109994378-109994879 | 8 | 0.995204218071908 |
| chr4-67204139-67204640 | 8 | 0.933484276090158 |
| chr19-55438748-55439249 | 8 | 0.940924879172365 |
| chr5-25545338-25545839 | 5 | 0.893063051263274 |
| chr6-89000061-89000562 | 8 | 0.993493681128311 |
| chr9-95964837-95965338 | 8 | 0.909422264345885 |
| chr20-17962073-17962574 | 5 | 0.975645121740319 |
| chr9-31154506-31155007 | 3 | 0.930116569294761 |
| chr8-109155595-109156096 | 5 | 0.962315505986283 |
| chr18-63982704-63983205 | 8 | 0.890368835873729 |
| chr2-152097186-152097687 | 8 | 0.902596369132572 |
| chr8-81052563-81053064 | 8 | 0.970290807868831 |
| chr4-39949252-39949753 | 6 | 0.991206925481043 |
| chr2-202322330-202322831 | 8 | 0.945629956045405 |
| chr9-71579839-71580340 | 2 | 0.953594481307634 |
| chr7-103805824-103806325 | 3 | 0.991549977714271 |
| chr15-39728113-39728614 | 8 | 0.951049029333779 |
| chr13-35512750-35513251 | 8 | 0.931628996283551 |
| chr15-4207749-4208250 | 7 | 0.815758486924117 |
| chr13-81593507-81594008 | 5 | 0.844362170810583 |
| chr2-25131553-25132054 | 3 | 0.875097936693183 |
| chr10-78796843-78797344 | 8 | 0.95396014366951 |
| chr2-74754514-74755015 | 8 | 0.947304763733429 |
| chr4-147591450-147591951 | 8 | 0.98665730941764 |
| chr9-42841050-42841551 | 8 | 0.993493681128311 |
| chr16-21202850-21203351 | 4 | 0.863605985891798 |
| chr2-31345195-31345696 | 8 | 0.981778955539409 |
| chr1-122734698-122735199 | 6 | 0.990223860089079 |
| chr18-52477941-52478442 | 8 | 0.944502993292951 |
| chr9-96716027-96716528 | 8 | 0.989267009157293 |
| chr1-114902883-114903384 | 8 | 0.994611018128268 |
| chrX-41714369-41714870 | 6 | 0.991206925481043 |
| chr9-65796567-65797068 | 8 | 0.822332239875129 |
| chr16-65498326-65498827 | 8 | 0.958985889320428 |
| chr3-97728448-97728949 | 3 | 0.99593553545041 |
| chr10-20606213-20606714 | 3 | 0.996423961532884 |
| chr15-8113040-8113541 | 8 | 0.974222959667429 |
| chr20-43445866-43446367 | 8 | 0.981855839275967 |
| chr17-44656989-44657490 | 8 | 0.964024119792981 |

| | | |
|---|---|---|
| chr16-70118872-70119373 | 3 | 0.996870254180838 |
| chr1-9551577-9552078 | 8 | 0.994488299140469 |
| chr5-120304498-120304999 | 3 | 0.996896238932032 |
| chr4-137170795-137171296 | 5 | 0.965237505027635 |
| chr7-111741858-111742359 | 4 | 0.855905630907399 |
| chr2-118043252-118043753 | 8 | 0.962803034305423 |
| chr9-41417688-41418189 | 8 | 0.930506804199047 |
| chr20-42029272-42029773 | 8 | 0.985977404284351 |
| chr4-83867744-83868245 | 8 | 0.93778765829277 |
| chr2-174024942-174025443 | 5 | 0.873465805956537 |
| chr14-38723874-38724375 | 8 | 0.962248103655691 |
| chr1-125885090-125885591 | 8 | 0.98049669822948 |
| chr17-85095136-85095637 | 8 | 0.929515854418081 |
| chrX-93858063-93858564 | 8 | 0.843064454497763 |
| chr3-58127644-58128145 | 8 | 0.902710399211979 |
| chr11-46544912-46545413 | 8 | 0.993983826662569 |
| chr1-189028518-189029019 | 8 | 0.94605734835967 |
| chr18-39772613-39773114 | 5 | 0.991969565685702 |
| chr14-41882495-41882996 | 8 | 0.819080729248125 |
| chr3-23493968-23494469 | 6 | 0.992737044412295 |
| chr16-41260926-41261427 | 3 | 0.996961179814231 |
| chr8-73606020-73606521 | 8 | 0.976478613082797 |
| chr6-117141430-117141931 | 4 | 0.900890554643784 |
| chr2-194942598-194943099 | 8 | 0.955996195364758 |
| chr4-63151567-63152068 | 8 | 0.994166165426562 |
| chr2-141867022-141867523 | 6 | 0.992828798760242 |
| chr20-30620398-30620899 | 8 | 0.958257372368166 |
| chr18-61104624-61105125 | 8 | 0.977171691028099 |
| chr8-120616152-120616653 | 8 | 0.887134613521843 |
| chr1-192755973-192756474 | 8 | 0.885630107016399 |
| chr10-60078216-60078717 | 8 | 0.994208877918839 |
| chr2-111507360-111507861 | 8 | 0.933635393796479 |
| chr1-56407898-56408399 | 8 | 0.983207217977992 |
| chr2-135477866-135478367 | 8 | 0.933890693621299 |
| chr4-158890609-158891110 | 8 | 0.992186598012668 |
| chr20-12631166-12631667 | 4 | 0.869731531497823 |
| chr12-1422679-1423180 | 4 | 0.887034302135247 |
| chr5-54961088-54961589 | 3 | 0.99593553545041 |
| chr18-30845273-30845774 | 8 | 0.831164996673716 |
| chr2-239054328-239054829 | 5 | 0.842808638418885 |
| chr9-79428923-79429424 | 8 | 0.990631609569205 |
| chr4-166816739-166817240 | 8 | 0.918306345631857 |
| chr13-48570458-48570959 | 6 | 0.991692735274593 |

| | | |
|---|---|---|
| chr17-84840074-84840575 | 4 | 0.89901599614076 |
| chr2-243033041-243033542 | 8 | 0.872088244673509 |
| chr12-19586500-19587001 | 1 | 0.932591875065237 |
| chr10-16408790-16409291 | 8 | 0.932685895166046 |
| chr5-98219328-98219829 | 4 | 0.912773526411941 |
| chr20-51710656-51711157 | 3 | 0.995147741889478 |
| chr15-74895579-74896080 | 8 | 0.962254461173722 |
| chr4-172171506-172172007 | 8 | 0.842123703429337 |
| chr5-12157118-12157619 | 2 | 0.917489350717517 |
| chr15-75704920-75705421 | 6 | 0.983935094489804 |
| chr16-74085726-74086227 | 3 | 0.99265170654043 |
| chr13-95785812-95786313 | 8 | 0.973914062414582 |
| chr1-146747236-146747737 | 8 | 0.891834506609125 |
| chr13-30734190-30734691 | 8 | 0.907554772506748 |
| chr1-79134273-79134774 | 2 | 0.877145096376516 |
| chr18-12501137-12501638 | 8 | 0.956069613658022 |
| chr2-196253607-196254108 | 2 | 0.906729594101372 |
| chr15-32829248-32829749 | 4 | 0.8244408375789 |
| chr13-93966373-93966874 | 6 | 0.996479153362823 |
| chr1-118790730-118791231 | 5 | 0.830879220931961 |
| chr17-59301955-59302456 | 3 | 0.99054720969867 |
| chr11-81076088-81076589 | 8 | 0.98324624185096 |
| chr9-112949765-112950266 | 2 | 0.916105603508271 |
| chr17-29756703-29757204 | 2 | 0.884469667771125 |
| chr13-100853616-100854117 | 8 | 0.997346119817848 |
| chr1-93675921-93676422 | 6 | 0.991206925481043 |
| chr2-104315529-104316030 | 8 | 0.976524540072503 |
| chr7-56779284-56779785 | 7 | 0.918186679369154 |
| chr17-69994206-69994707 | 2 | 0.830649123844304 |
| chr1-49685384-49685885 | 8 | 0.952313091283783 |
| chr14-51526852-51527353 | 2 | 0.911428689349559 |
| chr13-81701034-81701535 | 8 | 0.933292428270411 |
| chr2-53153591-53154092 | 4 | 0.928900755149925 |
| chr6-58233883-58234384 | 3 | 0.994605457401308 |
| chr5-1435542-1436043 | 6 | 0.993100122673275 |
| chr1-251445942-251446443 | 3 | 0.99661800737006 |
| chr4-37888498-37888999 | 6 | 0.992972779258414 |
| chr8-29521406-29521907 | 5 | 0.989718155066798 |
| chr4-162116667-162117168 | 3 | 0.98716337971059 |
| chr1-123129379-123129880 | 8 | 0.912593061648879 |
| chr18-68655296-68655797 | 8 | 0.984816710795558 |
| chr9-34655903-34656404 | 8 | 0.9145594599258 |
| chr4-135198257-135198758 | 8 | 0.957156217125137 |

| | | |
|---|---|---|
| chr20-32730614-32731115 | 6 | 0.960206627876325 |
| chr19-9739372-9739873 | 8 | 0.993856408858614 |
| chr10-39553327-39553828 | 8 | 0.960211173638235 |
| chr12-35813941-35814442 | 5 | 0.986793121856088 |
| chr8-888796-889297 | 4 | 0.952298775530372 |
| chr7-59816328-59816829 | 3 | 0.987132042413046 |
| chr14-25800156-25800657 | 8 | 0.994551208148499 |
| chr4-9433370-9433871 | 3 | 0.998693138056803 |
| chr6-42805434-42805935 | 3 | 0.994838006911226 |
| chr5-113627398-113627899 | 8 | 0.927193574924966 |
| chr11-55786095-55786596 | 8 | 0.916010858033731 |
| chr9-81623709-81624210 | 8 | 0.933675298998008 |
| chr1-88648247-88648748 | 4 | 0.898456716305013 |
| chr4-149024846-149025347 | 8 | 0.884419923666772 |
| chr14-32041199-32041700 | 5 | 0.988931803502401 |
| chr13-76485808-76486309 | 3 | 0.973638668861244 |
| chr1-22435306-22435807 | 2 | 0.859496791678308 |
| chr14-100362452-100362953 | 6 | 0.92260649742443 |
| chr15-46940134-46940635 | 8 | 0.916122500833829 |
| chr17-77879512-77880013 | 8 | 0.873280073642929 |
| chr10-24412181-24412682 | 3 | 0.995058657003496 |
| chr8-77864832-77865333 | 8 | 0.99230491751269 |
| chr13-103192137-103192638 | 8 | 0.993493681128311 |
| chr19-4888622-4889123 | 3 | 0.99380077314351 |
| chr9-113403872-113404373 | 2 | 0.906585929308423 |
| chr7-61536878-61537379 | 8 | 0.993493681128311 |
| chr11-61034671-61035172 | 4 | 0.847894863488535 |
| chr14-1546183-1546684 | 3 | 0.982851282451178 |
| chr4-139853022-139853523 | 8 | 0.993493681128311 |
| chr15-37598844-37599345 | 8 | 0.950386031247857 |
| chr5-9592499-9593000 | 3 | 0.99593553545041 |
| chr2-242268068-242268569 | 4 | 0.896523252144512 |
| chr14-1943373-1943874 | 8 | 0.988097271193359 |
| chr3-100478131-100478632 | 4 | 0.924754805847244 |
| chr2-91247684-91248185 | 6 | 0.988395737716461 |
| chr19-39090078-39090579 | 2 | 0.870207289622867 |
| chr1-223337050-223337551 | 4 | 0.975440721726164 |
| chr6-63718632-63719133 | 8 | 0.99048831569753 |
| chr4-14909205-14909706 | 4 | 0.954382449773578 |
| chr8-11793460-11793961 | 5 | 0.94082742758704 |
| chr15-80233971-80234472 | 8 | 0.947245670125477 |
| chr19-637867-638368 | 5 | 0.974339423953821 |
| chr1-163429904-163430405 | 2 | 0.949261519306242 |

| | | |
|---|---|---|
| chr7-58965622-58966123 | 2 | 0.950326724369851 |
| chrX-117090219-117090720 | 8 | 0.920291334789282 |
| chr1-129485286-129485787 | 8 | 0.937950074596203 |
| chr18-47186265-47186766 | 8 | 0.854733045035062 |
| chr3-100709518-100710019 | 8 | 0.942595823047018 |
| chr2-90914174-90914675 | 3 | 0.99593553545041 |
| chr19-3262483-3262984 | 2 | 0.899787415757923 |
| chr7-26882706-26883207 | 8 | 0.926047104781595 |
| chr2-150548745-150549246 | 5 | 0.968146340841871 |
| chr5-144759123-144759624 | 8 | 0.982664202136874 |
| chr2-123280055-123280556 | 8 | 0.932146921331176 |
| chr8-91893466-91893967 | 8 | 0.943749073682507 |
| chr7-95409908-95410409 | 3 | 0.9984138494557 |
| chr7-80130070-80130571 | 3 | 0.991300891099598 |
| chr14-32964456-32964957 | 4 | 0.972906256184799 |
| chr18-72067650-72068151 | 8 | 0.964101912347797 |
| chr4-141134894-141135395 | 8 | 0.847443815847373 |
| chr20-20363357-20363858 | 5 | 0.988160990740297 |
| chr3-33932166-33932667 | 8 | 0.990946838622942 |
| chr2-104715619-104716120 | 8 | 0.936461070106543 |
| chr13-29777092-29777593 | 8 | 0.993493681128311 |
| chr13-58222551-58223052 | 5 | 0.910198922123202 |
| chr19-50479262-50479763 | 8 | 0.945921024422014 |
| chr3-105821138-105821639 | 4 | 0.875944826877806 |
| chr17-32473215-32473716 | 5 | 0.979548090371877 |
| chr6-125737552-125738053 | 8 | 0.981720310866054 |
| chr19-54265443-54265944 | 5 | 0.907391154340923 |
| chr4-8978503-8979004 | 3 | 0.889157369950489 |
| chr7-112179896-112180397 | 8 | 0.871770026934382 |
| chr4-140250305-140250806 | 4 | 0.967754591272685 |
| chr16-68671841-68672342 | 3 | 0.99689648701724 |
| chr7-68851532-68852033 | 5 | 0.850526878770848 |
| chr3-127584425-127584926 | 4 | 0.930661649784842 |
| chr1-33523114-33523615 | 3 | 0.994377976561524 |
| chr4-161418341-161418842 | 8 | 0.966922966966481 |
| chr15-14580582-14581083 | 8 | 0.909451594347382 |
| chr12-12659505-12660006 | 2 | 0.965552822694561 |
| chr2-192876488-192876989 | 5 | 0.982163883854664 |
| chr19-31631910-31632411 | 6 | 0.992968484122683 |
| chr9-24017517-24018018 | 3 | 0.995329857129311 |
| chr10-90326973-90327474 | 2 | 0.912699046004978 |
| chr17-78992541-78993042 | 4 | 0.868311014827937 |
| chr6-139091912-139092413 | 3 | 0.816768171036839 |

| | | |
|---|---|---|
| chr5-94195870-94196371 | 8 | 0.847811649096725 |
| chr7-129652385-129652886 | 8 | 0.983307588982048 |
| chrX-117749590-117750091 | 3 | 0.99593553545041 |
| chr17-51558992-51559493 | 8 | 0.954380217137415 |
| chr5-1284933-1285434 | 8 | 0.993493681128311 |
| chr2-120543066-120543567 | 8 | 0.882042302681822 |
| chr8-96634651-96635152 | 4 | 0.911806526848652 |
| chr2-220457584-220458085 | 5 | 0.834481013488279 |
| chr1-226925572-226926073 | 8 | 0.994767613107324 |
| chr13-14233395-14233896 | 3 | 0.981755436354677 |
| chr4-171635822-171636323 | 8 | 0.993493681128311 |
| chr6-110859616-110860117 | 8 | 0.99503636160739 |
| chr13-62395987-62396488 | 4 | 0.95487523706124 |
| chr2-155177557-155178058 | 3 | 0.99593553545041 |
| chr6-106738686-106739187 | 8 | 0.95768941778056 |
| chr6-126945519-126946020 | 8 | 0.954894652149539 |
| chr1-192174850-192175351 | 8 | 0.993493681128311 |
| chr8-64655214-64655715 | 8 | 0.99331127503647 |
| chr5-150022470-150022971 | 8 | 0.838731492151134 |
| chr3-64436369-64436870 | 8 | 0.965835698434809 |
| chr3-61533719-61534220 | 7 | 0.912433844473394 |
| chr5-150530747-150531248 | 5 | 0.81611179623134 |
| chr1-4254884-4255385 | 5 | 0.873378189908146 |
| chr1-4981690-4982191 | 2 | 0.899755465365317 |
| chr6-121270646-121271147 | 8 | 0.967821544713787 |
| chr14-87686892-87687393 | 8 | 0.993493681128311 |
| chr8-53833068-53833569 | 7 | 0.815993369154051 |
| chr6-130779323-130779824 | 2 | 0.859097583076075 |
| chr1-216526101-216526602 | 3 | 0.99593553545041 |
| chr18-7917640-7918141 | 5 | 0.881551747729683 |
| chr3-118074227-118074728 | 4 | 0.903073657324356 |
| chr9-67712291-67712792 | 6 | 0.995904879438955 |
| chr2-140596378-140596879 | 7 | 0.906307978267741 |
| chr2-146202947-146203448 | 4 | 0.815883424854515 |
| chr17-77580528-77581029 | 8 | 0.926671475006359 |
| chr15-24083411-24083912 | 2 | 0.924311276142825 |
| chr7-74549969-74550470 | 8 | 0.885162228042694 |
| chr17-23226164-23226665 | 2 | 0.934639722799503 |
| chr15-22918975-22919476 | 8 | 0.944813631595372 |
| chr6-119065122-119065623 | 8 | 0.913656334739461 |
| chr6-133796565-133797066 | 3 | 0.970335829500331 |
| chr3-18713912-18714413 | 8 | 0.966440162531879 |
| chr3-97389215-97389716 | 3 | 0.99593553545041 |

| | | |
|---|---|---|
| chr18-23296174-23296675 | 5 | 0.859467127776517 |
| chr2-164163501-164164002 | 2 | 0.882183851272895 |
| chr4-126228654-126229155 | 8 | 0.968772867040247 |
| chr2-229900724-229901225 | 8 | 0.95069586709059 |
| chr5-150834953-150835454 | 4 | 0.952229830637291 |
| chrX-110497274-110497775 | 3 | 0.99593553545041 |
| chr17-23597702-23598203 | 8 | 0.930635138582619 |
| chr8-84816092-84816593 | 6 | 0.942377088743347 |
| chr3-100582163-100582664 | 2 | 0.885036585687223 |
| chr2-8598204-8598705 | 3 | 0.998367750893833 |
| chr13-55458449-55458950 | 4 | 0.965164564778641 |
| chr5-83417720-83418221 | 5 | 0.932689507122742 |
| chr4-144070020-144070521 | 3 | 0.996898207451623 |
| chr8-102859595-102860096 | 4 | 0.825424558725132 |
| chr6-54405473-54405974 | 8 | 0.884994237167679 |
| chr17-43587973-43588474 | 3 | 0.99593553545041 |
| chr18-11497084-11497585 | 8 | 0.978966490537692 |
| chr8-121654702-121655203 | 2 | 0.971803627528326 |
| chr17-4256801-4257302 | 5 | 0.964007277440784 |
| chr2-118931589-118932090 | 5 | 0.938607010309749 |
| chr2-115419061-115419562 | 4 | 0.966168657564298 |
| chr8-101464009-101464510 | 8 | 0.880803504915238 |
| chr2-91910219-91910720 | 4 | 0.967907353502118 |
| chr9-82021555-82022056 | 8 | 0.879077958247667 |
| chr15-80023131-80023632 | 5 | 0.966052789384209 |
| chr6-6267189-6267690 | 8 | 0.886677728549963 |
| chr13-100797760-100798261 | 4 | 0.968296214702836 |
| chr13-78282025-78282526 | 8 | 0.945193757272077 |
| chr18-70919755-70920256 | 5 | 0.990600128541711 |
| chr6-26797983-26798484 | 8 | 0.98080919653164 |
| chr14-8556168-8556669 | 3 | 0.971686556100043 |
| chr16-73634194-73634695 | 3 | 0.98815494674442 |
| chr1-102255634-102256135 | 3 | 0.99185714421491 |
| chr20-21527313-21527814 | 8 | 0.838509551937851 |
| chr12-31062256-31062757 | 8 | 0.990248703731876 |
| chr7-39532688-39533189 | 4 | 0.951692109311033 |
| chrX-136536337-136536838 | 8 | 0.9703451401921 |
| chr5-1603759-1604260 | 8 | 0.99108224047086 |
| chr1-181580672-181581173 | 7 | 0.907643796018719 |
| chr18-21120704-21121205 | 8 | 0.837847019741793 |
| chr2-60867395-60867896 | 5 | 0.986978911703681 |
| chrX-40447315-40447816 | 3 | 0.966998573210703 |
| chr1-28942925-28943426 | 8 | 0.879360744670605 |

| | | |
|---|---|---|
| chr7-32486495-32486996 | 8 | 0.855711283079707 |
| chr4-141745859-141746360 | 8 | 0.994390786964602 |
| chr4-10935061-10935562 | 3 | 0.989502680231845 |
| chrX-91009292-91009793 | 6 | 0.990182700965599 |
| chr8-62978322-62978823 | 7 | 0.832502731629511 |
| chr12-1619546-1620047 | 8 | 0.915149403526501 |
| chr11-57209346-57209847 | 8 | 0.979511648085709 |
| chr2-225742170-225742671 | 5 | 0.983919978840557 |
| chr4-25546894-25547395 | 4 | 0.924318693003462 |
| chr1-185290840-185291341 | 4 | 0.875347101711803 |
| chr2-25983251-25983752 | 4 | 0.974110428843221 |
| chr15-81891485-81891986 | 8 | 0.824378192958655 |
| chr13-88128051-88128552 | 2 | 0.899674589747744 |
| chr19-40957236-40957737 | 8 | 0.823285516873077 |
| chr2-78019050-78019551 | 6 | 0.944272828020432 |
| chr5-163273538-163274039 | 8 | 0.962422848280879 |
| chr17-77841496-77841997 | 8 | 0.98456955306584 |
| chr13-50121708-50122209 | 8 | 0.96530267880617 |
| chr16-46725480-46725981 | 5 | 0.966797413419514 |
| chr13-106336951-106337452 | 8 | 0.922222611990985 |
| chr7-79819980-79820481 | 8 | 0.967651548184208 |
| chr12-30869524-30870025 | 3 | 0.996568 |
| chr18-58359165-58359666 | 3 | 0.897586258125166 |
| chr11-35584999-35585500 | 4 | 0.917451869859657 |
| chr2-194506289-194506790 | 4 | 0.956612904278731 |
| chr7-74507486-74507987 | 8 | 0.850610618859805 |
| chr10-100574100-100574601 | 8 | 0.920058297638851 |
| chr14-79545526-79546027 | 5 | 0.987140529264921 |
| chr13-58379181-58379682 | 6 | 0.991206925481043 |
| chr6-137461605-137462106 | 2 | 0.830097291770656 |
| chr11-77735427-77735928 | 2 | 0.920573702390106 |
| chr17-62689332-62689833 | 3 | 0.995919291284384 |
| chrX-11345738-11346239 | 8 | 0.908224742781747 |
| chr8-102269936-102270437 | 2 | 0.94698987298046 |
| chr1-165213346-165213847 | 8 | 0.859627841901254 |
| chr16-16293717-16294218 | 8 | 0.995249027740187 |
| chr14-43858024-43858525 | 5 | 0.985146218578146 |
| chr10-17272383-17272884 | 8 | 0.994004115725641 |
| chr18-27817322-27817823 | 8 | 0.948119470339018 |
| chr20-44118035-44118536 | 8 | 0.834076051532072 |
| chr3-147852263-147852764 | 4 | 0.962766848947063 |
| chr7-24836593-24837094 | 4 | 0.866858008820775 |
| chr13-13660604-13661105 | 8 | 0.904045493162867 |

| | | |
|---|---|---|
| chr2-204115775-204116276 | 2 | 0.842575826456093 |
| chr2-123438625-123439126 | 8 | 0.94958599478903 |
| chr8-107581275-107581776 | 8 | 0.919387955559958 |
| chr4-102447433-102447934 | 3 | 0.842462636433274 |
| chr8-121209702-121210203 | 8 | 0.943700718917748 |
| chr18-60582922-60583423 | 8 | 0.993493681128311 |
| chr19-31842619-31843120 | 8 | 0.943059068272605 |
| chr7-60194841-60195342 | 5 | 0.985419343545262 |
| chr4-2912138-2912639 | 8 | 0.962877798134187 |
| chr2-107421714-107422215 | 3 | 0.973802783076161 |
| chrX-68618402-68618903 | 4 | 0.974659243008902 |
| chr2-89795269-89795770 | 3 | 0.995937228456398 |
| chr2-185869883-185870384 | 4 | 0.950719988643558 |
| chr2-220634606-220635107 | 5 | 0.859067634584631 |
| chr1-132838395-132838896 | 5 | 0.984389476454259 |
| chr7-52087427-52087928 | 7 | 0.852372726052838 |
| chr6-89592026-89592527 | 8 | 0.920466867406457 |
| chr3-122481134-122481635 | 1 | 0.957137942838351 |
| chr14-13499590-13500091 | 2 | 0.912378636890541 |
| chr4-134244032-134244533 | 8 | 0.990079303569139 |
| chr19-16005735-16006236 | 8 | 0.941627508523138 |
| chr18-80308812-80309313 | 4 | 0.937806591063394 |
| chr7-101098184-101098685 | 4 | 0.973145041060463 |
| chr15-100894766-100895267 | 8 | 0.95392664962489 |
| chr1-135544211-135544712 | 8 | 0.990296696753506 |
| chr13-104084618-104085119 | 8 | 0.974952315399465 |
| chr3-26674814-26675315 | 8 | 0.992371604199031 |
| chr18-82290935-82291436 | 3 | 0.9968317450116 |
| chr15-79989138-79989639 | 2 | 0.875845963851861 |
| chr15-12882005-12882506 | 4 | 0.872066910895768 |
| chr5-58817053-58817554 | 6 | 0.974402155570065 |
| chr5-21760543-21761044 | 4 | 0.917919014634606 |
| chr6-137809830-137810331 | 8 | 0.849063620838937 |
| chr17-82221128-82221629 | 8 | 0.982825249213111 |
| chr16-68960914-68961415 | 8 | 0.991936173913269 |
| chr6-136957089-136957590 | 8 | 0.993509962204576 |
| chr8-25619525-25620026 | 8 | 0.893483035715893 |
| chr18-69322794-69323295 | 5 | 0.984531415933021 |
| chr17-64383741-64384242 | 5 | 0.986395456954295 |
| chr1-216505219-216505720 | 3 | 0.990466666819488 |
| chr2-50554803-50555304 | 4 | 0.862644042395694 |
| chr12-10088409-10088910 | 4 | 0.853866449526309 |
| chr9-678150-678651 | 8 | 0.959138733498658 |

| | | |
|---|---|---|
| chr7-117350450-117350951 | 8 | 0.940507798168866 |
| chr10-21201705-21202206 | 8 | 0.993493681128311 |
| chr5-32107985-32108486 | 2 | 0.839672132000334 |
| chr1-184439178-184439679 | 4 | 0.856960266505572 |
| chrX-17848382-17848883 | 5 | 0.826403110935264 |
| chr1-145697056-145697557 | 6 | 0.955937721477928 |
| chr18-48541554-48542055 | 8 | 0.947354647632658 |
| chr6-45749597-45750098 | 8 | 0.861949777720739 |
| chr12-17102201-17102702 | 6 | 0.991276261035644 |
| chr4-105452823-105453324 | 8 | 0.883939328041958 |
| chr5-38086094-38086595 | 8 | 0.964630176697527 |
| chr5-5295825-5296326 | 8 | 0.956549069501358 |
| chr19-46418103-46418604 | 1 | 0.901715508182914 |
| chr16-16864076-16864577 | 8 | 0.897896836929879 |
| chr13-83054513-83055014 | 8 | 0.882139338225576 |
| chr15-10510761-10511262 | 8 | 0.977262974654418 |
| chr1-197884346-197884847 | 5 | 0.925639492459787 |
| chr2-56366928-56367429 | 8 | 0.993890489829705 |
| chr2-227247603-227248104 | 3 | 0.967698457945079 |
| chr17-59515109-59515610 | 8 | 0.953247573220858 |
| chr19-47253114-47253615 | 4 | 0.935003114126405 |
| chr8-27739849-27740350 | 6 | 0.994316753776642 |
| chr14-22734897-22735398 | 8 | 0.993493681128311 |
| chr7-20856891-20857392 | 8 | 0.834684235399679 |
| chr18-70820982-70821483 | 8 | 0.84244013453637 |
| chr4-53256423-53256924 | 8 | 0.993493681128311 |
| chr15-12579878-12580379 | 5 | 0.896154895440442 |
| chr7-91632066-91632567 | 8 | 0.839390341711112 |
| chr5-117323017-117323518 | 8 | 0.837629164141128 |
| chr11-50992135-50992636 | 8 | 0.880459955569337 |
| chr17-14977140-14977641 | 8 | 0.98798578586298 |
| chr5-27488350-27488851 | 8 | 0.994133554771285 |
| chr5-112359880-112360381 | 8 | 0.990283356634072 |
| chr2-114735486-114735987 | 3 | 0.996869613685166 |
| chr11-79915916-79916417 | 8 | 0.931675565787438 |
| chr3-76987451-76987952 | 8 | 0.885129540316658 |
| chr2-240003054-240003555 | 2 | 0.869401608886621 |
| chr9-110907688-110908189 | 8 | 0.845224480373722 |
| chr1-29597905-29598406 | 8 | 0.988393359054139 |
| chr13-103946329-103946830 | 6 | 0.992893529872837 |
| chrX-65794821-65795322 | 6 | 0.991206925481043 |
| chr3-100598302-100598803 | 8 | 0.984702370281754 |
| chr2-117939026-117939527 | 8 | 0.889533824553527 |

| | | |
|---|---|---|
| chr10-96943511-96944012 | 8 | 0.979318221729125 |
| chr10-22971305-22971806 | 2 | 0.911101959648278 |
| chr19-23811742-23812243 | 8 | 0.971778957667893 |
| chr13-49033257-49033758 | 8 | 0.968189335932525 |
| chr3-156891771-156892272 | 8 | 0.994437871820216 |
| chr1-88267709-88268210 | 8 | 0.993493681128311 |
| chr8-46808224-46808725 | 5 | 0.836911899689662 |
| chr18-1156680-1157181 | 8 | 0.91875339636097 |
| chr17-54619683-54620184 | 8 | 0.981320664108251 |
| chr9-99623425-99623926 | 3 | 0.99593553545041 |
| chrX-147684247-147684748 | 3 | 0.99593553545041 |
| chr15-23648448-23648949 | 3 | 0.996722434130234 |
| chr3-43271200-43271701 | 5 | 0.969138412061152 |
| chr7-132259452-132259953 | 3 | 0.892845722292776 |
| chr10-82590823-82591324 | 4 | 0.95762159578147 |
| chr15-76349867-76350368 | 8 | 0.993493681128311 |
| chr18-11792041-11792542 | 4 | 0.969939177978641 |
| chr9-86308992-86309493 | 3 | 0.875256268316299 |
| chr18-68901705-68902206 | 8 | 0.957325475095427 |
| chr2-171096326-171096827 | 4 | 0.939609687840599 |
| chr11-16314640-16315141 | 8 | 0.84876091547247 |
| chr4-25555453-25555954 | 3 | 0.997499159093579 |
| chr16-57929323-57929824 | 8 | 0.973633787375275 |
| chr2-155952635-155953136 | 3 | 0.991839544782184 |
| chr2-113493637-113494138 | 1 | 0.925963562589578 |
| chr10-51806433-51806934 | 6 | 0.990104727672885 |
| chr14-32978572-32979073 | 8 | 0.994022600951386 |
| chr15-15455461-15455962 | 8 | 0.993493681128311 |
| chr6-120012286-120012787 | 8 | 0.99223550145008 |
| chr2-167414492-167414993 | 6 | 0.990691138647139 |
| chr9-43171455-43171956 | 8 | 0.915364147838407 |
| chr1-185050874-185051375 | 8 | 0.929305128123277 |
| chr2-222355237-222355738 | 8 | 0.951087478096926 |
| chr1-82567658-82568159 | 8 | 0.970102563132277 |
| chr1-88009835-88010336 | 8 | 0.931371486516808 |
| chr8-49817277-49817778 | 8 | 0.905489081869708 |
| chr6-134486155-134486656 | 3 | 0.99593553545041 |
| chr2-102757849-102758350 | 8 | 0.893845839379425 |
| chr13-14386946-14387447 | 8 | 0.899664155638504 |
| chr4-60967741-60968242 | 8 | 0.993493681128311 |
| chr5-33142379-33142880 | 6 | 0.985720305359233 |
| chr15-64768159-64768660 | 3 | 0.978771532623204 |
| chr10-69986279-69986780 | 4 | 0.903960933807331 |

| | | |
|---|---|---|
| chr2-229742987-229743488 | 8 | 0.911259654717331 |
| chr3-32289679-32290180 | 8 | 0.99520202680543 |
| chr17-71641203-71641704 | 8 | 0.935100922071746 |
| chr15-14911659-14912160 | 8 | 0.976132975585264 |
| chrX-3623368-3623869 | 1 | 0.890479082205539 |
| chr9-40132194-40132695 | 8 | 0.867316460660818 |
| chr13-106383257-106383758 | 6 | 0.991995192436103 |
| chr5-14815457-14815958 | 5 | 0.959588077986122 |
| chr10-32196204-32196705 | 8 | 0.993493681128311 |
| chr19-47564451-47564952 | 7 | 0.936578898973567 |
| chr10-35278815-35279316 | 2 | 0.933487561363997 |
| chr1-255694120-255694621 | 6 | 0.980688227273692 |
| chr14-50216055-50216556 | 6 | 0.992629621977625 |
| chr2-31497146-31497647 | 8 | 0.974463007803407 |
| chr2-42292692-42293193 | 7 | 0.927707137888663 |
| chrX-72399965-72400466 | 8 | 0.969856338132034 |
| chr17-51133142-51133643 | 5 | 0.954242393113038 |
| chrX-44375528-44376029 | 6 | 0.991206925481043 |
| chrX-16332472-16332973 | 6 | 0.992437756891354 |
| chr2-58723949-58724450 | 4 | 0.962994668558714 |
| chr6-115001882-115002383 | 2 | 0.93590252227291 |
| chr15-98924852-98925353 | 5 | 0.952771490785092 |
| chr9-25246144-25246645 | 3 | 0.987656782823182 |
| chr2-87599308-87599809 | 5 | 0.913710007874851 |
| chr13-65276012-65276513 | 8 | 0.831263245772217 |
| chr15-15936346-15936847 | 4 | 0.970370831882719 |
| chr12-35832363-35832864 | 8 | 0.954846041197543 |
| chr1-146749255-146749756 | 4 | 0.895597597617723 |
| chr12-36622815-36623316 | 8 | 0.966434814255296 |
| chr2-57773853-57774354 | 5 | 0.960169550283368 |
| chr12-31050566-31051067 | 8 | 0.966717735155517 |
| chr5-31496689-31497190 | 5 | 0.97191513758328 |
| chr1-137757307-137757808 | 8 | 0.869726620042696 |
| chr5-74411876-74412377 | 3 | 0.892031824609207 |
| chr17-5324391-5324892 | 5 | 0.984937275808109 |
| chr1-14058310-14058811 | 8 | 0.952393994176148 |
| chr13-76395330-76395831 | 2 | 0.934112231022577 |
| chr1-185613057-185613558 | 3 | 0.995977576383622 |
| chr6-138835494-138835995 | 5 | 0.984440555408468 |
| chr2-9291290-9291791 | 8 | 0.898182129853766 |
| chr14-22043673-22044174 | 4 | 0.920539918859549 |
| chr11-12484570-12485071 | 2 | 0.949376892905926 |
| chr5-11585995-11586496 | 3 | 0.909055323348722 |

| | | |
|---|---|---|
| chr13-91374526-91375027 | 8 | 0.915384999857769 |
| chr11-55819665-55820166 | 8 | 0.889066107308986 |
| chr9-112044244-112044745 | 2 | 0.937734260982197 |
| chr17-63969035-63969536 | 1 | 0.857165558059847 |
| chr1-141354426-141354927 | 3 | 0.994803010395517 |
| chr15-18705490-18705991 | 8 | 0.830522923152255 |
| chr2-52966265-52966766 | 2 | 0.866270178145996 |
| chr12-35655248-35655749 | 4 | 0.896622197208798 |
| chr5-67993374-67993875 | 8 | 0.993493681128311 |
| chr6-26062765-26063266 | 8 | 0.956592251196482 |
| chr18-12902932-12903433 | 5 | 0.984007791480802 |
| chr4-84635842-84636343 | 8 | 0.993493681128311 |
| chr5-93051571-93052072 | 3 | 0.99593553545041 |
| chr1-188218300-188218801 | 8 | 0.989655977181958 |
| chr17-10017168-10017669 | 8 | 0.906915784392284 |
| chr2-23930579-23931080 | 8 | 0.857615297318948 |
| chr4-172574055-172574556 | 8 | 0.936039876047516 |
| chr1-51396160-51396661 | 2 | 0.922170661906992 |
| chr1-28896218-28896719 | 6 | 0.819237128282671 |
| chr6-121298746-121299247 | 6 | 0.984748208560898 |
| chr8-62742082-62742583 | 8 | 0.966814269800856 |
| chr15-82146606-82147107 | 8 | 0.886667888876182 |
| chr6-30832614-30833115 | 8 | 0.945764080944429 |
| chr6-53447462-53447963 | 4 | 0.949853789728228 |
| chr4-76238479-76238980 | 8 | 0.944028489875334 |
| chr10-94194992-94195493 | 8 | 0.962279656525581 |
| chr2-58005400-58005901 | 8 | 0.918702334293255 |
| chr10-41640332-41640833 | 8 | 0.867435510552546 |
| chr8-116768313-116768814 | 8 | 0.923660337510571 |
| chr7-89532400-89532901 | 8 | 0.860960298299562 |
| chr12-27309502-27310003 | 8 | 0.985894642719241 |
| chr16-13912713-13913214 | 8 | 0.957326371610385 |
| chr10-16759647-16760148 | 8 | 0.962535943502171 |
| chr1-49226872-49227373 | 5 | 0.924670741040411 |
| chr13-67764556-67765057 | 8 | 0.928339738502452 |
| chr14-73566628-73567129 | 6 | 0.985082421183837 |
| chr6-85374099-85374600 | 8 | 0.993493681128311 |
| chr17-52184368-52184869 | 5 | 0.862245117358936 |
| chr15-32844534-32845035 | 2 | 0.836415445145485 |
| chr13-92778920-92779421 | 8 | 0.986595248946575 |
| chr1-148121782-148122283 | 4 | 0.970754907743508 |
| chr9-104326666-104327167 | 8 | 0.843795218036479 |
| chr7-122515954-122516455 | 8 | 0.844613169696954 |

| | | |
|---|---|---|
| chr15-37938367-37938868 | 5 | 0.987149305689683 |
| chr3-25795745-25796246 | 5 | 0.988250937418204 |
| chr17-17068731-17069232 | 4 | 0.964370855262903 |
| chr2-68551575-68552076 | 8 | 0.968230045305965 |
| chr1-50713969-50714470 | 8 | 0.925263671594078 |
| chr4-174437307-174437808 | 5 | 0.84396298836854 |
| chr20-28929561-28930062 | 8 | 0.942308507983391 |
| chr18-31045374-31045875 | 2 | 0.90320482592195 |
| chr10-75346549-75347050 | 8 | 0.889146048902372 |
| chr20-19621987-19622488 | 8 | 0.992597677794638 |
| chrX-137806639-137807140 | 5 | 0.945290036075884 |
| chr4-134462085-134462586 | 2 | 0.888539403405556 |
| chr17-77002264-77002765 | 8 | 0.99524647393726 |
| chr4-173467258-173467759 | 8 | 0.823241306568593 |
| chr6-15969286-15969787 | 8 | 0.973882017231675 |
| chr1-92298838-92299339 | 8 | 0.993493681128311 |
| chr2-187833002-187833503 | 8 | 0.884206200177852 |
| chr15-67600990-67601491 | 3 | 0.99593553545041 |
| chrX-117562568-117563069 | 3 | 0.99688933200803 |
| chr11-59780741-59781242 | 2 | 0.949207513481008 |
| chr5-50185757-50186258 | 3 | 0.976397582285555 |
| chrX-36360836-36361337 | 3 | 0.992584457022762 |
| chr3-19450561-19451062 | 3 | 0.866576376325582 |
| chr6-99047655-99048156 | 3 | 0.875725879312594 |
| chr2-242376779-242377280 | 1 | 0.912518474165776 |
| chr3-162549409-162549910 | 8 | 0.96999329558835 |
| chr11-32812257-32812758 | 8 | 0.88545739275759 |
| chr14-58959701-58960202 | 4 | 0.960523716290896 |
| chr10-63663324-63663825 | 8 | 0.837653101441288 |
| chr1-163878049-163878550 | 8 | 0.986181761823166 |
| chr5-14985521-14986022 | 3 | 0.819388492984671 |
| chr5-32004205-32004706 | 8 | 0.984797168223701 |
| chr18-45328785-45329286 | 5 | 0.975265618667206 |
| chr11-70649432-70649933 | 5 | 0.948887992264035 |
| chr8-118357133-118357634 | 5 | 0.899565930101448 |
| chr15-785435-785936 | 8 | 0.953460170775353 |
| chr16-5674006-5674507 | 8 | 0.966979759870878 |
| chr17-36099830-36100331 | 8 | 0.947756991829269 |
| chr12-37741233-37741734 | 4 | 0.903495789888059 |
| chr16-20270429-20270930 | 2 | 0.964756581646693 |
| chr17-24570676-24571177 | 6 | 0.930181503165862 |
| chr8-53888154-53888655 | 8 | 0.993493681128311 |
| chr15-44625298-44625799 | 5 | 0.852172595639452 |

| | | |
|---|---|---|
| chr6-101693178-101693679 | 8 | 0.964290464910776 |
| chr8-14089850-14090351 | 3 | 0.945722698403007 |
| chr7-111152287-111152788 | 8 | 0.988300601700551 |
| chr4-143325802-143326303 | 8 | 0.923999792877836 |
| chr5-164239347-164239848 | 8 | 0.962395177399404 |
| chr5-46711257-46711758 | 8 | 0.896831161486541 |
| chr13-61556970-61557471 | 8 | 0.847695412302444 |
| chr2-189703046-189703547 | 8 | 0.948452838742064 |
| chr11-40447190-40447691 | 8 | 0.994430052890247 |
| chr17-50794587-50795088 | 8 | 0.968259856439685 |
| chr6-29288466-29288967 | 4 | 0.972266254695081 |
| chrX-104389182-104389683 | 7 | 0.963335339622303 |
| chr3-165633420-165633921 | 8 | 0.983927959264766 |
| chr20-33433994-33434495 | 5 | 0.918917356256598 |
| chr7-66208584-66209085 | 2 | 0.885825600099112 |
| chr7-111257894-111258395 | 5 | 0.893360937351884 |
| chr1-138138069-138138570 | 3 | 0.861640707755108 |
| chr6-14549456-14549957 | 4 | 0.923644206712695 |
| chr6-48710639-48711140 | 6 | 0.99270598750199 |
| chr5-31247841-31248342 | 3 | 0.995902535318766 |
| chr14-66314675-66315176 | 5 | 0.848208939964579 |
| chr15-22715971-22716472 | 5 | 0.958018400313964 |
| chr8-57790811-57791312 | 7 | 0.847196134787106 |
| chr18-45044332-45044833 | 3 | 0.998359600844999 |
| chr3-166653674-166654175 | 6 | 0.964779658640071 |
| chr2-100500512-100501013 | 8 | 0.85496529873792 |
| chr6-100513871-100514372 | 3 | 0.98198479367026 |
| chr16-38311766-38312267 | 8 | 0.838153537393783 |
| chr18-1402550-1403051 | 8 | 0.993493681128311 |
| chr18-70849803-70850304 | 8 | 0.830962913430834 |
| chr4-106410235-106410736 | 3 | 0.995473699726095 |
| chr2-133688536-133689037 | 3 | 0.995285855128359 |
| chr6-21112430-21112931 | 8 | 0.831816639279373 |
| chr1-185801161-185801662 | 3 | 0.873044136085511 |
| chr15-97697580-97698081 | 5 | 0.900933718736446 |
| chr15-85083890-85084391 | 3 | 0.99593553545041 |
| chr1-106747469-106747970 | 2 | 0.956212376471038 |
| chr5-63506472-63506973 | 8 | 0.989967225712666 |
| chr6-91648888-91649389 | 2 | 0.90092594635216 |
| chr13-10712328-10712829 | 3 | 0.996888936397445 |
| chr3-159999437-159999938 | 8 | 0.86840843663192 |
| chr2-137657798-137658299 | 8 | 0.995114068487181 |
| chr19-43630150-43630651 | 3 | 0.966612623584057 |

| | | |
|---|---|---|
| chr9-110413301-110413802 | 4 | 0.875521216860633 |
| chr1-152833217-152833718 | 7 | 0.90176094594206 |
| chr13-60716392-60716893 | 6 | 0.988053231246808 |
| chr2-81222734-81223235 | 3 | 0.939229958545742 |
| chr12-11406432-11406933 | 8 | 0.99518638098032 |
| chr2-19644952-19645453 | 5 | 0.980602275986873 |
| chr18-1658380-1658881 | 3 | 0.998417937919578 |
| chr1-155077102-155077603 | 6 | 0.991206925481043 |
| chr18-59495437-59495938 | 4 | 0.918754110645048 |
| chr13-68476952-68477453 | 8 | 0.990793856588887 |
| chr2-108407478-108407979 | 8 | 0.897411 |
| chr13-22111449-22111950 | 3 | 0.98609579434502 |
| chr5-72447883-72448384 | 4 | 0.904001379517192 |
| chr10-48335549-48336050 | 8 | 0.991762172259581 |
| chr17-27716742-27717243 | 8 | 0.898670176371914 |
| chr2-19993839-19994340 | 5 | 0.840280486629991 |
| chr11-65310316-65310817 | 5 | 0.942882861538627 |
| chr6-70406553-70407054 | 8 | 0.925154289451813 |
| chr5-47808813-47809314 | 8 | 0.893137088296111 |
| chr7-109658276-109658777 | 4 | 0.889591672715437 |
| chr14-74673765-74674266 | 8 | 0.994195238911928 |
| chr13-98594776-98595277 | 8 | 0.850410969826985 |
| chr3-50798480-50798981 | 8 | 0.886046056660858 |
| chr2-41510334-41510835 | 8 | 0.95859711052423 |
| chr3-19703915-19704416 | 3 | 0.993022258575835 |
| chr3-37080047-37080548 | 8 | 0.980239710537562 |
| chr5-21763971-21764472 | 8 | 0.969765062803166 |
| chr4-113305006-113305507 | 3 | 0.990383591120275 |
| chr14-41260069-41260570 | 5 | 0.94549271495939 |
| chr11-9290198-9290699 | 3 | 0.996924517637504 |
| chr8-33152443-33152944 | 8 | 0.994637020417243 |
| chr14-68298450-68298951 | 3 | 0.99593553545041 |
| chr3-76437768-76438269 | 4 | 0.973151935627728 |
| chr17-15542991-15543492 | 5 | 0.81553517998169 |
| chr8-78999438-78999939 | 8 | 0.993493681128311 |
| chr16-67190786-67191287 | 4 | 0.963544348218018 |
| chr17-30030051-30030552 | 4 | 0.854207893431236 |
| chr13-14522779-14523280 | 3 | 0.996877251275193 |
| chr12-2665487-2665988 | 8 | 0.977307007701046 |
| chr16-4261549-4262050 | 4 | 0.905929754875679 |
| chr3-22779125-22779626 | 8 | 0.910705792200844 |
| chr7-41537633-41538134 | 8 | 0.992214473113101 |
| chr12-13280847-13281348 | 8 | 0.94201064904902 |

| | | |
|---|---|---|
| chr9-63615627-63616128 | 6 | 0.910497753665805 |
| chr2-231785630-231786131 | 8 | 0.990379671439465 |
| chr16-3517213-3517714 | 8 | 0.978437539931457 |
| chr15-49196897-49197398 | 6 | 0.993220230884313 |
| chr10-38496443-38496944 | 8 | 0.881795838142357 |
| chr3-43408409-43408910 | 3 | 0.998156592995334 |
| chr7-41148241-41148742 | 5 | 0.984192396687372 |
| chr3-132090524-132091025 | 8 | 0.840831600190259 |
| chr11-86039391-86039892 | 8 | 0.908808292102147 |
| chr2-211568015-211568516 | 4 | 0.89556973589948 |
| chr20-9169376-9169877 | 7 | 0.845065156009206 |
| chr13-106478144-106478645 | 5 | 0.931349803675673 |
| chr10-25683834-25684335 | 6 | 0.993204693131168 |
| chr17-18666277-18666778 | 8 | 0.959604806312091 |
| chr13-23897541-23898042 | 8 | 0.991147357867191 |
| chr2-45787394-45787895 | 8 | 0.993493681128311 |
| chr10-50695623-50696124 | 8 | 0.968495855311329 |
| chr9-94125958-94126459 | 8 | 0.943533196579039 |
| chr4-18435242-18435743 | 5 | 0.935510616805426 |
| chr12-6207731-6208232 | 5 | 0.980072367121374 |
| chr2-11900694-11901195 | 4 | 0.927893258498918 |
| chr12-16576828-16577329 | 6 | 0.85634810275922 |
| chr8-23516581-23517082 | 8 | 0.814993102461489 |
| chrX-129926085-129926586 | 8 | 0.993493681128311 |
| chr3-128997089-128997590 | 4 | 0.969228498380026 |
| chr2-37622306-37622807 | 1 | 0.937458987992403 |
| chr13-8306721-8307222 | 3 | 0.993954853239168 |
| chrX-115170704-115171205 | 8 | 0.927230816314656 |
| chr3-120134833-120135334 | 6 | 0.983543759446478 |
| chr1-180476856-180477357 | 8 | 0.987847201520378 |
| chrX-84521558-84522059 | 3 | 0.99593553545041 |
| chr15-97850542-97851043 | 8 | 0.981938969300203 |
| chr1-165805983-165806484 | 8 | 0.986329617509387 |
| chr1-135571735-135572236 | 8 | 0.994296462019365 |
| chr5-67467134-67467635 | 3 | 0.993157460751185 |
| chr14-22470653-22471154 | 3 | 0.99593553545041 |
| chr11-80946270-80946771 | 8 | 0.983832479105409 |
| chr15-73439864-73440365 | 6 | 0.991206925481043 |
| chr9-108368580-108369081 | 3 | 0.996830518792233 |
| chr10-37533027-37533528 | 4 | 0.929110457704936 |
| chr17-20542485-20542986 | 5 | 0.988709977874896 |
| chr2-30281293-30281794 | 8 | 0.992856 |
| chr1-245547626-245548127 | 8 | 0.916020096160684 |

| | | |
|---|---|---|
| chr10-60436334-60436835 | 8 | 0.903437867122145 |
| chr15-25678769-25679270 | 5 | 0.921034053685751 |
| chrX-41708858-41709359 | 6 | 0.991206925481043 |
| chr13-17831800-17832301 | 3 | 0.983288950342167 |
| chr4-42911022-42911523 | 6 | 0.992976833309741 |
| chrX-32063154-32063655 | 7 | 0.84260585329386 |
| chr18-75769526-75770027 | 8 | 0.992746566452193 |
| chr20-45633438-45633939 | 8 | 0.936417168476928 |
| chr15-82448705-82449206 | 8 | 0.93830422554699 |
| chr3-73360309-73360810 | 3 | 0.99593553545041 |
| chr15-72041300-72041801 | 6 | 0.99017709123176 |
| chr5-4221398-4221899 | 8 | 0.832161437246452 |
| chr7-15076350-15076851 | 5 | 0.851756941891377 |
| chr3-234494-234995 | 3 | 0.998125329976753 |
| chr10-97448563-97449064 | 3 | 0.944697301680969 |
| chr2-84844320-84844821 | 8 | 0.946167342921881 |
| chr8-31594013-31594514 | 8 | 0.990283252237456 |
| chr10-28951697-28952198 | 8 | 0.917405825441058 |
| chr15-92754162-92754663 | 2 | 0.937280788268108 |
| chr3-81171520-81172021 | 3 | 0.945631649854227 |
| chr4-159234604-159235105 | 6 | 0.831851554120515 |
| chr2-218372556-218373057 | 8 | 0.962692989538917 |
| chr1-257438826-257439327 | 8 | 0.95572716326022 |
| chr10-21776903-21777404 | 3 | 0.99593553545041 |
| chr14-37349557-37350058 | 8 | 0.956935045102719 |
| chr8-107712636-107713137 | 6 | 0.921945618986065 |
| chrX-127359684-127360185 | 4 | 0.923111988391995 |
| chr16-17735195-17735696 | 8 | 0.992267852587414 |
| chr1-207212720-207213221 | 5 | 0.989036814268094 |
| chr14-72537198-72537699 | 2 | 0.890130897304188 |
| chr18-61319586-61320087 | 3 | 0.883265448136511 |
| chr1-224373941-224374442 | 2 | 0.886261143209271 |
| chr8-106812304-106812805 | 8 | 0.862713392133986 |
| chr17-55038976-55039477 | 8 | 0.982528737134855 |
| chr19-2424397-2424898 | 8 | 0.817437765890404 |
| chr17-77201620-77202121 | 8 | 0.968532062468172 |
| chr6-88547576-88548077 | 8 | 0.896201664373203 |
| chr15-64948194-64948695 | 3 | 0.971993095531189 |
| chr4-48461408-48461909 | 3 | 0.995985108671256 |
| chr1-109791172-109791673 | 3 | 0.99593553545041 |
| chr13-98562301-98562802 | 8 | 0.994755574294915 |
| chr10-19238492-19238993 | 8 | 0.909364029675612 |
| chr20-52684091-52684592 | 3 | 0.990786305659848 |

| | | |
|---|---|---|
| chrX-71719981-71720482 | 8 | 0.923512153670708 |
| chr5-157472182-157472683 | 5 | 0.817200646861635 |
| chr3-54847101-54847602 | 4 | 0.914542416803789 |
| chr8-20983957-20984458 | 2 | 0.888853967431646 |
| chr5-79466046-79466547 | 8 | 0.965247222358363 |
| chr8-76090746-76091247 | 8 | 0.994031529053921 |
| chr5-119110110-119110611 | 5 | 0.965024444090399 |
| chr6-88143448-88143949 | 5 | 0.984807161803357 |
| chr8-1620527-1621028 | 8 | 0.970707191269404 |
| chr11-51171367-51171868 | 5 | 0.933522972141827 |
| chr8-11771391-11771892 | 4 | 0.822568603370442 |
| chrX-10356332-10356833 | 8 | 0.971360796220374 |
| chr20-32971637-32972138 | 8 | 0.821538449414193 |
| chr14-36988871-36989372 | 3 | 0.977099773758058 |
| chr15-54273609-54274110 | 5 | 0.972863847426132 |
| chr5-47443719-47444220 | 8 | 0.894669774911058 |
| chr15-22882928-22883429 | 8 | 0.8512312390056 |
| chr4-31506978-31507479 | 8 | 0.997018614254541 |
| chr2-144552475-144552976 | 2 | 0.958192550790846 |
| chr4-166117967-166118468 | 3 | 0.993351059771578 |
| chr19-48927208-48927709 | 5 | 0.9435613300113 |
| chr7-83245552-83246053 | 8 | 0.994644603378238 |
| chr6-75569096-75569597 | 8 | 0.85792205790154 |
| chr5-111524243-111524744 | 8 | 0.992551489252134 |
| chr10-41705525-41706026 | 8 | 0.902498567939239 |
| chr16-39575969-39576470 | 5 | 0.963207485745519 |
| chr1-57134611-57135112 | 3 | 0.994265655661363 |
| chr4-13686278-13686779 | 8 | 0.943761443321413 |
| chr19-31915623-31916124 | 3 | 0.991987866520917 |
| chr16-12716585-12717086 | 2 | 0.92669932537568 |
| chr1-172949901-172950402 | 5 | 0.983919978840557 |
| chr2-182852588-182853089 | 8 | 0.997980312745461 |
| chr8-85247238-85247739 | 8 | 0.941967458016381 |
| chr5-24038875-24039376 | 6 | 0.971098395950658 |
| chr16-2344826-2345327 | 8 | 0.966824176331563 |
| chr4-101495325-101495826 | 8 | 0.948716610114594 |
| chr7-67545654-67546155 | 4 | 0.846761500356959 |
| chr19-31634478-31634979 | 3 | 0.99593553545041 |
| chr14-64234329-64234830 | 3 | 0.9577966340204 |
| chrX-10071289-10071790 | 4 | 0.902137006925115 |
| chr10-104931992-104932493 | 5 | 0.959859313964551 |
| chr5-5321909-5322410 | 8 | 0.880169726117139 |
| chr8-68055207-68055708 | 2 | 0.904022944648819 |

| | | |
|---|---|---|
| chr1-16317924-16318425 | 8 | 0.993493681128311 |
| chr5-6145535-6146036 | 8 | 0.898142934997495 |
| chr20-43320833-43321334 | 6 | 0.994404744978373 |
| chr6-137286299-137286800 | 8 | 0.872886328310731 |
| chr2-189371522-189372023 | 8 | 0.864985114805055 |
| chr6-27093200-27093701 | 6 | 0.975909491270428 |
| chr2-93721592-93722093 | 5 | 0.912724109207079 |
| chr9-20205213-20205714 | 2 | 0.833693018459751 |
| chr17-16172181-16172682 | 6 | 0.991206925481043 |
| chr5-101355442-101355943 | 8 | 0.876779295286508 |
| chr5-513559-514060 | 3 | 0.99593553545041 |
| chr16-20105438-20105939 | 4 | 0.904568996219079 |
| chr10-85354780-85355281 | 8 | 0.958866311657789 |
| chr1-183664594-183665095 | 8 | 0.815017893108768 |
| chr1-5127430-5127931 | 8 | 0.920667851385397 |
| chr6-135676506-135677007 | 2 | 0.950699374380374 |
| chr6-39714616-39715117 | 4 | 0.853450691126838 |
| chr6-110581705-110582206 | 8 | 0.993845456698259 |
| chr3-64333037-64333538 | 8 | 0.97761794786805 |
| chr15-875860-876361 | 8 | 0.993493681128311 |
| chr17-16345838-16346339 | 8 | 0.993493681128311 |
| chr3-167007487-167007988 | 8 | 0.909328290174468 |
| chr4-160526854-160527355 | 8 | 0.929536383130626 |
| chr8-112246740-112247241 | 3 | 0.969821780776216 |
| chr3-22783691-22784192 | 8 | 0.985090374653054 |
| chr13-31239739-31240240 | 8 | 0.976930754908549 |
| chr16-57116778-57117279 | 7 | 0.819769243181803 |
| chr8-27062749-27063250 | 8 | 0.937370215547999 |
| chr5-114424776-114425277 | 5 | 0.990044560274322 |
| chr9-12972988-12973489 | 8 | 0.992247294569569 |
| chr1-140974567-140975068 | 8 | 0.967908629075266 |
| chr13-14661984-14662485 | 8 | 0.990019164427187 |
| chr8-30888064-30888565 | 2 | 0.936663363376309 |
| chr6-22732788-22733289 | 4 | 0.871902194824631 |
| chr14-81181239-81181740 | 8 | 0.945321993860081 |
| chr4-159516330-159516831 | 8 | 0.991578400411074 |
| chr16-58883212-58883713 | 6 | 0.981503763664956 |
| chr12-98867-99368 | 8 | 0.98924219186184 |
| chr4-48463282-48463783 | 4 | 0.854777192730038 |
| chr5-134728508-134729009 | 8 | 0.943901715416771 |
| chr9-94513499-94514000 | 8 | 0.983047468724344 |
| chr20-27257291-27257792 | 4 | 0.932388837156581 |
| chr1-160518574-160519075 | 3 | 0.99593553545041 |

| | | |
|---|---|---|
| chr4-77735825-77736326 | 1 | 0.822405084424757 |
| chr10-20438799-20439300 | 8 | 0.989028052731712 |
| chr5-112974444-112974945 | 8 | 0.992433646632817 |
| chr3-30152534-30153035 | 8 | 0.993363127112632 |
| chr14-21835195-21835696 | 6 | 0.992975065906704 |
| chr3-122901783-122902284 | 4 | 0.864519950366103 |
| chr4-85827090-85827591 | 4 | 0.928258014556008 |
| chrX-16253232-16253733 | 6 | 0.993225807301246 |
| chr8-23674357-23674858 | 4 | 0.91794452055642 |
| chr12-17119495-17119996 | 1 | 0.904634971143357 |
| chr15-64554176-64554677 | 8 | 0.929237504016176 |
| chr2-102000793-102001294 | 8 | 0.962348344628562 |
| chr12-12466639-12467140 | 8 | 0.864071748013586 |
| chr10-23826376-23826877 | 4 | 0.924752238315697 |
| chr12-44043155-44043656 | 8 | 0.995099016069688 |
| chr15-34712436-34712937 | 8 | 0.880520787950957 |
| chr4-99446912-99447413 | 5 | 0.98204458076016 |
| chr1-44138333-44138834 | 8 | 0.936783180611329 |
| chr4-181952848-181953349 | 8 | 0.966662554035977 |
| chr11-28000721-28001222 | 8 | 0.994981678210092 |
| chr4-179572103-179572604 | 4 | 0.878422185531748 |
| chr14-46292534-46293035 | 5 | 0.865393065473839 |
| chr7-102805493-102805994 | 3 | 0.99593553545041 |
| chr3-129689954-129690455 | 4 | 0.928550818576408 |
| chr7-121505800-121506301 | 3 | 0.996329162129106 |
| chr14-11731859-11732360 | 3 | 0.995703208734469 |
| chr4-124743970-124744471 | 8 | 0.900674234189678 |
| chr7-38831380-38831881 | 8 | 0.989506797838036 |
| chr1-47351065-47351566 | 6 | 0.953984338257072 |
| chr19-12197000-12197501 | 8 | 0.993493681128311 |
| chr2-248654707-248655208 | 8 | 0.993493681128311 |
| chr4-129382248-129382749 | 3 | 0.962823169457562 |
| chr4-138493987-138494488 | 8 | 0.98695619905038 |
| chr5-115996540-115997041 | 3 | 0.99705191070287 |
| chr12-39452717-39453218 | 8 | 0.9948539807826 |
| chr1-7411419-7411920 | 5 | 0.989558414373156 |
| chrX-24593964-24594465 | 3 | 0.99593553545041 |
| chr2-12354663-12355164 | 6 | 0.990701741106924 |
| chr5-15306094-15306595 | 1 | 0.924855642264818 |
| chr16-35426872-35427373 | 2 | 0.81952928520327 |
| chr6-26037087-26037588 | 4 | 0.936197237240584 |
| chr3-55409211-55409712 | 8 | 0.995122741051457 |
| chr6-90661338-90661839 | 8 | 0.992239662139458 |

| | | |
|---|---|---|
| chr2-78651776-78652277 | 5 | 0.989230368319611 |
| chr15-45884908-45885409 | 8 | 0.930755717897302 |
| chr10-29921988-29922489 | 8 | 0.986833331509959 |
| chr15-88883693-88884194 | 4 | 0.964246179486336 |
| chr5-134412720-134413221 | 8 | 0.856074521520418 |
| chr8-121656243-121656744 | 8 | 0.861678913075289 |
| chr2-222277652-222278153 | 8 | 0.994208050253726 |
| chr11-62204044-62204545 | 8 | 0.994265234665313 |
| chr7-64589538-64590039 | 3 | 0.98372438467232 |
| chr11-58913719-58914220 | 8 | 0.9399361844873 |
| chr13-69067831-69068332 | 8 | 0.908163715241944 |
| chr4-130114886-130115387 | 4 | 0.87541853098802 |
| chr3-59970665-59971166 | 2 | 0.850877632950563 |
| chr2-134066413-134066914 | 3 | 0.99593553545041 |
| chr17-57025131-57025632 | 5 | 0.892854658097884 |
| chr8-106375745-106376246 | 4 | 0.883004751072131 |
| chr1-117691698-117692199 | 8 | 0.981113853171549 |
| chr4-89917991-89918492 | 8 | 0.968716270273838 |
| chr7-68810210-68810711 | 8 | 0.907877263001369 |
| chr9-45499113-45499614 | 4 | 0.885582763636469 |
| chr13-72565859-72566360 | 6 | 0.954438515373582 |
| chr14-78834101-78834602 | 8 | 0.854641857316069 |
| chr4-82819984-82820485 | 8 | 0.956759473674618 |
| chr2-105250684-105251185 | 8 | 0.947926212479318 |
| chr14-35039876-35040377 | 2 | 0.819642204932446 |
| chr2-56862611-56863112 | 8 | 0.991161091989804 |
| chr1-81786961-81787462 | 3 | 0.994225714696583 |
| chr9-111377740-111378241 | 8 | 0.993067310434477 |
| chr15-12042071-12042572 | 3 | 0.99593553545041 |
| chr15-58018947-58019448 | 3 | 0.99687427298051 |
| chr3-114287544-114288045 | 8 | 0.995226424380244 |
| chr10-32175039-32175540 | 8 | 0.856288055913715 |
| chr4-65253797-65254298 | 8 | 0.995137688777624 |
| chr13-102029854-102030355 | 3 | 0.993094747707409 |
| chr1-185588283-185588784 | 1 | 0.91564674087281 |
| chr7-45011030-45011531 | 2 | 0.835728250470849 |
| chr4-62846764-62847265 | 3 | 0.996038474020631 |
| chr8-83333983-83334484 | 3 | 0.99593553545041 |
| chr4-95306451-95306952 | 4 | 0.951132394374438 |
| chr17-7752729-7753230 | 1 | 0.954954243233974 |
| chr14-85121915-85122416 | 5 | 0.840367885230193 |
| chr7-71127302-71127803 | 8 | 0.928015454949062 |
| chr13-12988606-12989107 | 6 | 0.995641115397235 |

| | | |
|---|---|---|
| chrX-96672274-96672775 | 7 | 0.928356986218617 |
| chr7-94160567-94161068 | 8 | 0.94962263269011 |
| chr3-24828632-24829133 | 3 | 0.99593553545041 |
| chr13-40633970-40634471 | 8 | 0.955578013720714 |
| chr17-35591830-35592331 | 8 | 0.993149610600295 |
| chr5-93378721-93379222 | 3 | 0.995551838788367 |
| chr8-22592203-22592704 | 2 | 0.95476753206763 |
| chr2-208951475-208951976 | 8 | 0.989041256204258 |
| chr10-91855773-91856274 | 8 | 0.943481076529856 |
| chr11-56401668-56402169 | 8 | 0.985165227604934 |
| chr16-12061499-12062000 | 3 | 0.997055471715014 |
| chr14-13702677-13703178 | 8 | 0.977282456616142 |
| chr10-571654-572155 | 8 | 0.870734192019332 |
| chr18-11463502-11464003 | 8 | 0.895370410852345 |
| chr2-142699450-142699951 | 8 | 0.929658708549057 |
| chr7-28515227-28515728 | 8 | 0.975904049758248 |
| chr9-66422309-66422810 | 5 | 0.971672532096382 |
| chr2-113426932-113427433 | 5 | 0.932245504981016 |
| chr10-44658180-44658681 | 5 | 0.961130642578016 |
| chr3-62456528-62457029 | 8 | 0.993493681128311 |
| chr10-41586563-41587064 | 2 | 0.974784235658751 |
| chr3-37016094-37016595 | 5 | 0.874676653507987 |
| chr12-27232193-27232694 | 8 | 0.992853073011724 |
| chr2-145197696-145198197 | 5 | 0.986044611772821 |
| chr18-83710883-83711384 | 8 | 0.819186479699482 |
| chr15-39690725-39691226 | 2 | 0.892476586428043 |
| chr4-13440918-13441419 | 8 | 0.985543003638176 |
| chr10-39900628-39901129 | 8 | 0.912595222676391 |
| chr19-53779846-53780347 | 8 | 0.856372384724929 |
| chr11-26904032-26904533 | 6 | 0.96529074817265 |
| chr1-49872113-49872614 | 2 | 0.946265703717442 |
| chr19-7825897-7826398 | 8 | 0.993493681128311 |
| chr13-98919580-98920081 | 7 | 0.866396962998201 |
| chr1-98416994-98417495 | 8 | 0.891585357878055 |
| chr7-68121608-68122109 | 5 | 0.904258666119633 |
| chr3-107817783-107818284 | 5 | 0.977801983078223 |
| chr8-110469953-110470454 | 2 | 0.929923289865744 |
| chr6-87084165-87084666 | 8 | 0.979280342814427 |
| chr7-30768146-30768647 | 8 | 0.905902492321819 |
| chr15-97765675-97766176 | 5 | 0.888763254448494 |
| chr2-217965068-217965569 | 2 | 0.875205633576735 |
| chr1-256503313-256503814 | 8 | 0.846951385965621 |
| chr16-76267168-76267669 | 8 | 0.929337391120951 |

| | | |
|---|---|---|
| chr3-35809515-35810016 | 8 | 0.876471578148967 |
| chr1-219814679-219815180 | 4 | 0.861263875987729 |
| chr8-19527556-19528057 | 8 | 0.85599935640999 |
| chr11-42744666-42745167 | 5 | 0.987383909683482 |
| chr10-28188505-28189006 | 8 | 0.990533140946837 |
| chr1-253739446-253739947 | 3 | 0.970812327966874 |
| chr2-140520966-140521467 | 8 | 0.864650326532951 |
| chr5-128776282-128776783 | 8 | 0.985346371703614 |
| chr9-61841009-61841510 | 8 | 0.984474857353793 |
| chr7-101259360-101259861 | 8 | 0.909109119309696 |
| chr4-122402398-122402899 | 1 | 0.929753908500511 |
| chr13-73221107-73221608 | 8 | 0.859900271326264 |
| chr4-50114761-50115262 | 4 | 0.921467303870167 |
| chr10-99536207-99536708 | 8 | 0.899317639099316 |
| chr2-61218113-61218614 | 8 | 0.993493681128311 |
| chr12-24084096-24084597 | 8 | 0.909501062914464 |
| chr6-51957127-51957628 | 6 | 0.994165204815094 |
| chr4-39885503-39886004 | 3 | 0.99593553545041 |
| chr1-245104022-245104523 | 8 | 0.877839529175295 |
| chr3-122162318-122162819 | 3 | 0.84002174341866 |
| chr19-53096903-53097404 | 5 | 0.960445332273805 |
| chr13-64292691-64293192 | 8 | 0.953714007337347 |
| chr13-40676749-40677250 | 3 | 0.928910536755154 |
| chr3-77589878-77590379 | 8 | 0.968182681571632 |
| chr11-42919356-42919857 | 8 | 0.970484371190564 |
| chr1-62724336-62724837 | 3 | 0.900738975693553 |
| chr6-35267502-35268003 | 5 | 0.990450902102111 |
| chr17-23675655-23676156 | 4 | 0.920563017912237 |
| chr14-98004384-98004885 | 8 | 0.945239083426788 |
| chr4-46350987-46351488 | 8 | 0.981960576771875 |
| chr20-30964050-30964551 | 6 | 0.977481519573619 |
| chr7-34539676-34540177 | 8 | 0.844582713932757 |
| chr5-72839033-72839534 | 8 | 0.957982099428438 |
| chr1-89581597-89582098 | 8 | 0.993493681128311 |
| chr15-16095882-16096383 | 8 | 0.956186616389051 |
| chr5-83839355-83839856 | 3 | 0.994220700457178 |
| chr6-25826823-25827324 | 8 | 0.817494014151758 |
| chr15-32496992-32497493 | 2 | 0.884630510353382 |
| chr7-14633335-14633836 | 5 | 0.845715452770598 |
| chr13-79467193-79467694 | 5 | 0.863190013197038 |
| chr2-132591211-132591712 | 6 | 0.986870289372645 |
| chr17-33166248-33166749 | 5 | 0.942173581340259 |
| chr4-150949472-150949973 | 7 | 0.900512567587723 |

| | | |
|---|---|---|
| chr13-51955896-51956397 | 8 | 0.868603216228949 |
| chr1-211880782-211881283 | 4 | 0.932415064551393 |
| chr3-90768387-90768888 | 2 | 0.929479240651568 |
| chr13-2431088-2431589 | 6 | 0.961790346186765 |
| chr1-212703535-212704036 | 5 | 0.910779644871656 |
| chr2-47587554-47588055 | 8 | 0.993493681128311 |
| chr4-159718445-159718946 | 4 | 0.873998113980525 |
| chr3-149866424-149866925 | 8 | 0.992273987306036 |
| chr9-31784505-31785006 | 8 | 0.993493681128311 |
| chr6-132567177-132567678 | 3 | 0.981522563117611 |
| chr7-40594935-40595436 | 4 | 0.862975190619698 |
| chr7-121128945-121129446 | 8 | 0.825340149939183 |
| chr9-50928357-50928858 | 8 | 0.995247730940124 |
| chr16-71335824-71336325 | 5 | 0.924936379975551 |
| chr17-22899622-22900123 | 5 | 0.985999771738139 |
| chr9-67835284-67835785 | 8 | 0.987824656599234 |
| chr5-47917735-47918236 | 5 | 0.983919978840557 |
| chr16-45324059-45324560 | 8 | 0.927837916507529 |
| chrX-67378129-67378630 | 8 | 0.972956081478939 |
| chr3-91145775-91146276 | 8 | 0.878711990561661 |
| chr18-7517125-7517626 | 6 | 0.989220598672262 |
| chr5-110975939-110976440 | 8 | 0.992252066891198 |
| chr3-97644636-97645137 | 3 | 0.901750790625179 |
| chr4-79513098-79513599 | 8 | 0.97708512800804 |
| chr3-34528796-34529297 | 3 | 0.998352597780663 |
| chr4-177503336-177503837 | 6 | 0.992003034484701 |
| chr1-243699008-243699509 | 8 | 0.967769492061905 |
| chr7-75298063-75298564 | 3 | 0.986123600970662 |
| chr15-16728602-16729103 | 8 | 0.98228858632949 |
| chr7-115893716-115894217 | 5 | 0.987126909261268 |
| chr4-123506375-123506876 | 8 | 0.995242554009859 |
| chr6-20570882-20571383 | 6 | 0.866618002680258 |
| chr17-51521488-51521989 | 8 | 0.890618404704635 |
| chr15-12430864-12431365 | 3 | 0.998234325218797 |
| chrY-9344336-9344837 | 2 | 0.941124164640853 |
| chr14-21704637-21705138 | 1 | 0.849333113182426 |
| chr6-33168868-33169369 | 7 | 0.882594907083549 |
| chrX-49090492-49090993 | 3 | 0.996311898327289 |
| chr1-15699597-15700098 | 5 | 0.945939921197617 |
| chr1-167176698-167177199 | 4 | 0.937947190958135 |
| chr5-55186721-55187222 | 8 | 0.97218403330018 |
| chr6-46298559-46299060 | 8 | 0.963754161701317 |
| chr14-16050213-16050714 | 8 | 0.837477432136268 |

| | | |
|---|---|---|
| chr6-95423408-95423909 | 8 | 0.964635836602079 |
| chr14-83713632-83714133 | 8 | 0.957033930155655 |
| chr8-48279216-48279717 | 8 | 0.986451920397916 |
| chr19-54171629-54172130 | 5 | 0.970697722219137 |
| chr15-51779845-51780346 | 2 | 0.935143051298949 |
| chr20-13118712-13119213 | 8 | 0.962872837424585 |
| chr17-35241856-35242357 | 1 | 0.896585074171377 |
| chr4-76483989-76484490 | 8 | 0.970396980908357 |
| chr1-23303479-23303980 | 3 | 0.915695216673671 |
| chr2-220701450-220701951 | 8 | 0.983626156872773 |
| chr3-37376251-37376752 | 5 | 0.98811551528828 |
| chr8-9774850-9775351 | 3 | 0.996742479584812 |
| chr8-51574485-51574986 | 8 | 0.932628595403573 |
| chr5-161233236-161233737 | 8 | 0.933404107866453 |
| chrX-127141924-127142425 | 8 | 0.990115596051677 |
| chr16-2675206-2675707 | 8 | 0.976433402336582 |
| chr2-31100636-31101137 | 8 | 0.988479538699444 |
| chr7-53766954-53767455 | 5 | 0.962793117243694 |
| chr19-52654823-52655324 | 3 | 0.985013967512222 |
| chr8-30578765-30579266 | 8 | 0.972611680324224 |
| chr18-62632572-62633073 | 5 | 0.988480932610774 |
| chr4-17991773-17992274 | 6 | 0.99212074216976 |
| chr3-127162854-127163355 | 8 | 0.994078075626983 |
| chr2-94873787-94874288 | 2 | 0.951857472644196 |
| chr15-13185225-13185726 | 8 | 0.933890598798111 |
| chr8-84918981-84919482 | 8 | 0.915698516087809 |
| chr3-144615015-144615516 | 8 | 0.991943820265802 |
| chr2-175901300-175901801 | 8 | 0.890716317292154 |
| chr11-3803521-3804022 | 3 | 0.984169519129509 |
| chr4-170643561-170644062 | 3 | 0.980683331849808 |
| chr15-49550596-49551097 | 3 | 0.998537723541153 |
| chr6-84873369-84873870 | 5 | 0.828701387000542 |
| chr2-94293139-94293640 | 3 | 0.99593553545041 |
| chr4-38289855-38290356 | 5 | 0.983919978840557 |
| chr16-16162641-16163142 | 8 | 0.994644511731713 |
| chr2-225920809-225921310 | 3 | 0.873298246947951 |
| chr2-55223356-55223857 | 8 | 0.994650697051862 |
| chr11-73368190-73368691 | 8 | 0.945614630546792 |
| chr16-46576194-46576695 | 3 | 0.946688694817223 |
| chr14-32289403-32289904 | 1 | 0.858049345655409 |
| chr10-20303512-20304013 | 8 | 0.995152124827188 |
| chr1-41830196-41830697 | 8 | 0.886237172977904 |
| chr3-135026915-135027416 | 4 | 0.922874463155954 |

| | | |
|---|---|---|
| chr15-50618756-50619257 | 8 | 0.993493681128311 |
| chr1-222162170-222162671 | 4 | 0.943119915863453 |
| chr1-193389549-193390050 | 3 | 0.997201411227417 |
| chr1-12076922-12077423 | 8 | 0.98655989709126 |
| chr1-127693788-127694289 | 2 | 0.879994092246801 |
| chr6-98741969-98742470 | 4 | 0.880344673815701 |
| chr6-132461141-132461642 | 8 | 0.953623883199739 |
| chr10-92859471-92859972 | 4 | 0.835057513549254 |
| chr3-87066953-87067454 | 8 | 0.993487451646849 |
| chr9-40205069-40205570 | 8 | 0.916166288778683 |
| chr1-163280340-163280841 | 8 | 0.989088503976573 |
| chr11-47347322-47347823 | 3 | 0.863031435032861 |
| chr8-49439845-49440346 | 8 | 0.974561311100152 |
| chrX-5667547-5668048 | 3 | 0.960136913294406 |
| chr6-31672085-31672586 | 3 | 0.99499482224617 |
| chr15-379201-379702 | 8 | 0.833891919518783 |
| chr11-74457415-74457916 | 3 | 0.996890368565238 |
| chr18-64193309-64193810 | 4 | 0.853743740631545 |
| chr7-104236511-104237012 | 3 | 0.96570432106807 |
| chr8-50113291-50113792 | 8 | 0.869504665418461 |
| chr13-99189196-99189697 | 2 | 0.869515400926404 |
| chr3-78197346-78197847 | 8 | 0.991098863952343 |
| chr16-75410048-75410549 | 5 | 0.970132870282124 |
| chrX-32063718-32064219 | 3 | 0.905674347390903 |
| chr6-78557805-78558306 | 8 | 0.993493681128311 |
| chr20-4811998-4812499 | 8 | 0.880306225798541 |
| chr1-41371216-41371717 | 3 | 0.918487959808392 |
| chr15-44849588-44850089 | 8 | 0.889953021631617 |
| chr18-30295419-30295920 | 8 | 0.916078099607265 |
| chr18-63078892-63079393 | 8 | 0.93806275545924 |
| chr4-9983678-9984179 | 8 | 0.929806080538062 |
| chrX-13061892-13062393 | 5 | 0.950550207503897 |
| chr3-60790184-60790685 | 8 | 0.93278000681334 |
| chr19-46577640-46578141 | 3 | 0.821406712985735 |
| chr3-165938219-165938720 | 2 | 0.82624910912441 |
| chr1-18453323-18453824 | 8 | 0.938561879488347 |
| chr7-22496120-22496621 | 8 | 0.993493681128311 |
| chr5-132678611-132679112 | 5 | 0.919204649596223 |
| chr17-51786095-51786596 | 3 | 0.928056283596262 |
| chr3-99628707-99629208 | 3 | 0.997021409833849 |
| chr1-136425553-136426054 | 8 | 0.994930919725897 |
| chr5-126815035-126815536 | 8 | 0.993493681128311 |
| chr2-161608865-161609366 | 6 | 0.992399013344154 |

| | | |
|---|---|---|
| chr18-54386404-54386905 | 8 | 0.853004012334919 |
| chr2-217762513-217763014 | 6 | 0.992566313494204 |
| chr16-9334319-9334820 | 8 | 0.831762617675355 |
| chr8-30761710-30762211 | 4 | 0.963567905603678 |
| chr17-76306912-76307413 | 4 | 0.830687726981193 |
| chr1-188299655-188300156 | 7 | 0.906552872570719 |
| chr18-77625504-77626005 | 4 | 0.875325744567345 |
| chr18-74488406-74488907 | 5 | 0.817621136539963 |
| chr13-105280230-105280731 | 3 | 0.968970270182404 |
| chr2-34777205-34777706 | 8 | 0.952104936648658 |
| chr19-43488606-43489107 | 5 | 0.979523846731063 |
| chr1-233993208-233993709 | 5 | 0.992893768269729 |
| chr9-1271854-1272355 | 8 | 0.95372890232123 |
| chr11-37797246-37797747 | 6 | 0.991206925481043 |
| chr13-95079383-95079884 | 2 | 0.926962289530644 |
| chr17-82224347-82224848 | 3 | 0.907355951886536 |
| chr8-113286865-113287366 | 6 | 0.886721990165721 |
| chr9-91602699-91603200 | 8 | 0.984297111913814 |
| chr6-8640918-8641419 | 2 | 0.824092240244333 |
| chr18-62360171-62360672 | 8 | 0.992973417049328 |
| chr4-116289798-116290299 | 4 | 0.855089165432041 |
| chr2-204952031-204952532 | 1 | 0.831067506549894 |
| chr14-18462356-18462857 | 6 | 0.972441003939558 |
| chr1-69703000-69703501 | 8 | 0.99504169270871 |
| chr15-48545614-48546115 | 2 | 0.960056347803731 |
| chr2-222367015-222367516 | 4 | 0.816985799312641 |
| chr2-77161168-77161669 | 8 | 0.989144138277558 |
| chr4-57571708-57572209 | 8 | 0.988971625828024 |
| chr13-69380330-69380831 | 8 | 0.833317443452106 |
| chr3-82899114-82899615 | 6 | 0.994294595840649 |
| chr5-82433462-82433963 | 4 | 0.843139486515306 |
| chr2-53494853-53495354 | 2 | 0.921014199852849 |
| chr16-69999049-69999550 | 5 | 0.978482529185307 |
| chr4-155377786-155378287 | 8 | 0.911360818661073 |
| chr5-31919293-31919794 | 5 | 0.983919978840557 |
| chrX-42433832-42434333 | 3 | 0.99593553545041 |
| chr2-7280855-7281356 | 6 | 0.950167368503173 |
| chr13-7416014-7416515 | 2 | 0.944835428971977 |
| chr11-43692631-43693132 | 8 | 0.995227652465629 |
| chr10-46108823-46109324 | 8 | 0.992422315700759 |
| chr3-21386573-21387074 | 8 | 0.895377993110154 |
| chr2-235105454-235105955 | 8 | 0.980998381351022 |
| chr1-119825402-119825903 | 8 | 0.841905867726415 |

| | | |
|---|---|---|
| chr10-7758889-7759390 | 2 | 0.907093547820992 |
| chr15-74749301-74749802 | 1 | 0.935818540431776 |
| chr8-61011093-61011594 | 6 | 0.995768590368317 |
| chr14-7511774-7512275 | 8 | 0.977022375691894 |
| chr3-148566446-148566947 | 2 | 0.842044858213042 |
| chr1-236465087-236465588 | 5 | 0.835025891581239 |
| chr4-162402666-162403167 | 8 | 0.955056403939272 |
| chr1-130504580-130505081 | 6 | 0.991206925481043 |
| chr19-12152731-12153232 | 3 | 0.996833840677754 |
| chr7-16446224-16446725 | 2 | 0.900157634273744 |
| chr20-23945716-23946217 | 5 | 0.988576643201478 |
| chr17-5648930-5649431 | 8 | 0.969568261654931 |
| chr19-49978082-49978583 | 5 | 0.994081403390187 |
| chr15-7063020-7063521 | 8 | 0.991879569458236 |
| chr1-32661248-32661749 | 8 | 0.816873810510097 |
| chr8-26922188-26922689 | 8 | 0.99474460361588 |
| chr1-218420241-218420742 | 6 | 0.9906416649797 |
| chr14-42157273-42157774 | 8 | 0.945597338989425 |
| chr8-94757888-94758389 | 5 | 0.86222248014828 |
| chr13-28153114-28153615 | 3 | 0.998190592989031 |
| chr4-160688136-160688637 | 7 | 0.942826915773363 |
| chr12-38725289-38725790 | 3 | 0.999078924087786 |
| chr7-3805533-3806034 | 6 | 0.991206925481043 |
| chr2-230069832-230070333 | 8 | 0.935807177575222 |
| chr13-91828873-91829374 | 8 | 0.921753180554068 |
| chr6-70772531-70773032 | 8 | 0.949102908195967 |
| chr20-9353757-9354258 | 6 | 0.993584335650041 |
| chr14-72669835-72670336 | 5 | 0.915522702143552 |
| chr3-131952227-131952728 | 8 | 0.991459023575478 |
| chr13-30188089-30188590 | 5 | 0.978193160666514 |
| chr2-148860953-148861454 | 3 | 0.99593553545041 |
| chr10-28976250-28976751 | 8 | 0.968644607562247 |
| chr15-37094800-37095301 | 5 | 0.988495474698944 |
| chr19-46152619-46153120 | 5 | 0.993176227299953 |
| chr5-30563354-30563855 | 8 | 0.993493681128311 |
| chr2-167583764-167584265 | 8 | 0.983802054489955 |
| chr14-40350594-40351095 | 8 | 0.993820973997171 |
| chr1-125949531-125950032 | 3 | 0.996667776960005 |
| chr15-24077056-24077557 | 3 | 0.99301289163896 |
| chr1-50745274-50745775 | 5 | 0.959822336864696 |
| chr15-89203964-89204465 | 6 | 0.991206925481043 |
| chr9-99342639-99343140 | 8 | 0.993325318256308 |
| chr7-30004351-30004852 | 8 | 0.91775649505481 |

| | | |
|---|---|---|
| chr2-34555221-34555722 | 8 | 0.936769989224188 |
| chr15-101686929-101687430 | 8 | 0.991671057101124 |
| chr2-166412902-166413403 | 5 | 0.988846004121352 |
| chr1-124833055-124833556 | 8 | 0.970160381069792 |
| chr1-201270326-201270827 | 8 | 0.946555855192978 |
| chr1-21507348-21507849 | 3 | 0.981732162837516 |
| chr12-29787110-29787611 | 4 | 0.94381634994859 |
| chr11-23596654-23597155 | 4 | 0.914236400951373 |
| chr2-95093589-95094090 | 8 | 0.942918392197877 |
| chr7-64543380-64543881 | 4 | 0.965790139845904 |
| chr4-146938157-146938658 | 8 | 0.915831599361688 |
| chr1-123905619-123906120 | 4 | 0.917941726435764 |
| chr14-62905415-62905916 | 5 | 0.873083712420792 |
| chr5-10004227-10004728 | 8 | 0.993752292283022 |
| chr18-73385513-73386014 | 3 | 0.995285268176266 |
| chr2-195021024-195021525 | 8 | 0.894151159794341 |
| chr10-31519153-31519654 | 8 | 0.881039108137687 |
| chr1-149770803-149771304 | 3 | 0.991307984357349 |
| chr7-39228377-39228878 | 4 | 0.929757940590217 |
| chr13-63226188-63226689 | 2 | 0.932642349830048 |
| chr5-98962725-98963226 | 8 | 0.957577 |
| chr12-5667282-5667783 | 6 | 0.976565985074093 |
| chr7-121823353-121823854 | 8 | 0.938941886776059 |
| chr15-76419091-76419592 | 8 | 0.995009320061857 |
| chr2-100657389-100657890 | 8 | 0.902650644491188 |
| chr16-63392509-63393010 | 8 | 0.940134599782831 |
| chr2-177586453-177586954 | 6 | 0.945565894371161 |
| chr18-69401855-69402356 | 4 | 0.953295561335618 |
| chr1-132677245-132677746 | 8 | 0.870793781757415 |
| chr5-19879207-19879708 | 8 | 0.988932093684959 |
| chr1-233951876-233952377 | 5 | 0.936994061290635 |
| chr19-46676999-46677500 | 1 | 0.942209109317341 |
| chr10-4101062-4101563 | 8 | 0.920715421686376 |
| chr2-239739392-239739893 | 1 | 0.947529198009474 |
| chr6-138743946-138744447 | 5 | 0.946813180519377 |
| chr17-47562595-47563096 | 3 | 0.99593553545041 |
| chr15-19223420-19223921 | 4 | 0.850032857075361 |
| chr19-52841835-52842336 | 7 | 0.879154843729586 |
| chr5-62481074-62481575 | 8 | 0.955833950012795 |
| chr8-81266457-81266958 | 8 | 0.957457541947146 |
| chr8-95678550-95679051 | 8 | 0.943811893182417 |
| chr8-72292842-72293343 | 5 | 0.960904711839665 |
| chr4-120625541-120626042 | 6 | 0.9918495753406 |

| | | |
|---|---|---|
| chr3-155554017-155554518 | 8 | 0.930560045749235 |
| chr8-51248973-51249474 | 8 | 0.987333675248883 |
| chr13-36161046-36161547 | 3 | 0.956737689599296 |
| chr17-26566077-26566578 | 7 | 0.839361189676603 |
| chr4-36032176-36032677 | 8 | 0.845412311436035 |
| chr6-77159573-77160074 | 8 | 0.986315142858189 |
| chr2-209460835-209461336 | 2 | 0.857121609300733 |
| chr2-121855792-121856293 | 3 | 0.93674910229734 |
| chr18-11625308-11625809 | 4 | 0.928040691143606 |
| chr13-87288514-87289015 | 8 | 0.930189476095419 |
| chr4-50723113-50723614 | 8 | 0.985504336305875 |
| chr20-16048197-16048698 | 8 | 0.969807233216212 |
| chr6-55893452-55893953 | 3 | 0.99593553545041 |
| chr5-2079026-2079527 | 7 | 0.905254894407622 |
| chr7-123155913-123156414 | 8 | 0.917435882890159 |
| chr8-21623918-21624419 | 8 | 0.992974887244263 |
| chr17-40078792-40079293 | 8 | 0.973517365265993 |
| chr8-22991238-22991739 | 8 | 0.993724087580987 |
| chr7-125437239-125437740 | 4 | 0.908605210746801 |
| chr20-5116214-5116715 | 3 | 0.821792418095133 |
| chr3-156323670-156324171 | 8 | 0.932231809664963 |
| chr19-38414167-38414668 | 4 | 0.942700109872414 |
| chr7-44655679-44656180 | 8 | 0.986431148657699 |
| chr11-49646083-49646584 | 8 | 0.975936544366439 |
| chr5-32556547-32557048 | 8 | 0.819774191362306 |
| chr3-82320557-82321058 | 5 | 0.88099021671523 |
| chrX-121968767-121969268 | 8 | 0.990248419907732 |
| chr19-25750999-25751500 | 5 | 0.990616475112707 |
| chr13-24037146-24037647 | 7 | 0.971469705308953 |
| chr8-30963590-30964091 | 5 | 0.917896531994911 |
| chr4-24569637-24570138 | 8 | 0.993493681128311 |
| chr6-15216602-15217103 | 5 | 0.987841412253606 |
| chr11-22797857-22798358 | 3 | 0.996916404532109 |
| chr18-13055777-13056278 | 5 | 0.991478006634951 |
| chr8-95426786-95427287 | 6 | 0.975506030031305 |
| chr14-31706371-31706872 | 8 | 0.993493681128311 |
| chr2-71652581-71653082 | 8 | 0.993493681128311 |
| chr11-37341608-37342109 | 5 | 0.986565 |
| chr7-100229216-100229717 | 3 | 0.996895430625708 |
| chr9-111293352-111293853 | 8 | 0.992653048694697 |
| chr11-47725287-47725788 | 8 | 0.994790686121259 |
| chr15-79748977-79749478 | 8 | 0.988552522464575 |
| chr15-85974577-85975078 | 3 | 0.99659073279801 |

| | | |
|---|---|---|
| chr3-88712659-88713160 | 4 | 0.867961683698588 |
| chr1-150712500-150713001 | 8 | 0.974568625117435 |
| chr10-46783336-46783837 | 8 | 0.955917115656037 |
| chr20-19131103-19131604 | 4 | 0.891470660326413 |
| chr15-21213120-21213621 | 7 | 0.891684219383693 |
| chr7-64526468-64526969 | 4 | 0.839852034828684 |
| chr3-97197445-97197946 | 8 | 0.958914193118472 |
| chr9-22907218-22907719 | 3 | 0.877132505330721 |
| chr1-90575652-90576153 | 8 | 0.945231547163649 |
| chr20-43437033-43437534 | 8 | 0.961645793972259 |
| chr14-47403547-47404048 | 8 | 0.995246498129885 |
| chrX-57108556-57109057 | 5 | 0.911152473262651 |
| chr3-131730711-131731212 | 5 | 0.84788141876093 |
| chr5-29417755-29418256 | 3 | 0.99604893145714 |
| chr1-218995569-218996070 | 8 | 0.897068068895618 |
| chr9-55760536-55761037 | 8 | 0.874924556260578 |
| chr5-140710669-140711170 | 4 | 0.885183289406245 |
| chr1-48788639-48789140 | 5 | 0.995983134520921 |
| chr8-85140611-85141112 | 6 | 0.993195996339609 |
| chr13-6694077-6694578 | 3 | 0.860456550616267 |
| chr5-163435468-163435969 | 8 | 0.887656635209972 |
| chr3-129856872-129857373 | 8 | 0.938472977060998 |
| chr6-137062774-137063275 | 4 | 0.825595916877982 |
| chr15-8481352-8481853 | 5 | 0.993766940347589 |
| chr19-37997719-37998220 | 1 | 0.86032897753555 |
| chr3-19416080-19416581 | 8 | 0.952858569773212 |
| chr16-19463106-19463607 | 8 | 0.992417119264827 |
| chr12-44715026-44715527 | 8 | 0.957101961736094 |
| chr13-90577739-90578240 | 8 | 0.924111981035625 |
| chr1-244269420-244269921 | 8 | 0.99339294162405 |
| chr4-50704886-50705387 | 3 | 0.99593553545041 |
| chr11-33733635-33734136 | 4 | 0.965230178279489 |
| chr18-39206120-39206621 | 6 | 0.99051317042739 |
| chr3-157618636-157619137 | 8 | 0.897140837058973 |
| chr7-74438780-74439281 | 5 | 0.918076062885422 |
| chr9-105533278-105533779 | 8 | 0.838054872668244 |
| chr16-17457061-17457562 | 3 | 0.995190258553411 |
| chr2-170223443-170223944 | 8 | 0.980317072377818 |
| chr19-4206047-4206548 | 8 | 0.984813299086722 |
| chr5-68284648-68285149 | 2 | 0.860281158533769 |
| chr3-47135270-47135771 | 4 | 0.873153134193503 |
| chr8-49566910-49567411 | 8 | 0.991984220257273 |
| chr4-165362456-165362957 | 6 | 0.991206925481043 |

| | | |
|---|---|---|
| chr1-242676838-242677339 | 5 | 0.986632000906444 |
| chr2-168395094-168395595 | 8 | 0.945477941946569 |
| chr7-70889504-70890005 | 8 | 0.995224979787774 |
| chr16-10043370-10043871 | 8 | 0.941709874610392 |
| chr7-67840993-67841494 | 8 | 0.993493681128311 |
| chr1-41063879-41064380 | 3 | 0.996652438311952 |
| chr3-41725093-41725594 | 4 | 0.947056266337099 |
| chr11-54870395-54870896 | 8 | 0.941335899154718 |
| chr4-124053357-124053858 | 8 | 0.993011094124345 |
| chr16-49284652-49285153 | 8 | 0.926439434844906 |
| chr8-45895284-45895785 | 4 | 0.880679177185242 |
| chr9-85919052-85919553 | 7 | 0.842864529188246 |
| chr2-76753286-76753787 | 7 | 0.912664178860039 |
| chr4-40205184-40205685 | 8 | 0.847789211512319 |
| chr3-130242027-130242528 | 3 | 0.900236883245843 |
| chr9-34533921-34534422 | 8 | 0.991789312571695 |
| chr2-31364282-31364783 | 8 | 0.975524186937137 |
| chr10-10103878-10104379 | 8 | 0.993493681128311 |
| chr6-33267885-33268386 | 5 | 0.967156336590946 |
| chr1-133390121-133390622 | 8 | 0.931167367032028 |
| chr4-103146573-103147074 | 8 | 0.983888115320631 |
| chr13-106204373-106204874 | 2 | 0.868928220336447 |
| chr14-72652298-72652799 | 5 | 0.918957734030452 |
| chr14-41987792-41988293 | 5 | 0.930989094023887 |
| chr16-72046656-72047157 | 6 | 0.993038192396238 |
| chr16-12302950-12303451 | 3 | 0.983525488965752 |
| chr1-27131251-27131752 | 8 | 0.942587396807222 |
| chr5-163098469-163098970 | 8 | 0.993950864628619 |
| chr3-23901234-23901735 | 3 | 0.996766160564982 |
| chr4-118984406-118984907 | 8 | 0.91084604270616 |
| chr9-98535843-98536344 | 3 | 0.99593553545041 |
| chr7-112502699-112503200 | 8 | 0.88507851678519 |
| chr7-124381679-124382180 | 5 | 0.975301873384382 |
| chr4-52163923-52164424 | 1 | 0.927006194359888 |
| chr3-145114917-145115418 | 8 | 0.993763148056435 |
| chr8-8083930-8084431 | 3 | 0.997238421665524 |
| chr15-78225372-78225873 | 1 | 0.827572443096883 |
| chr7-46821989-46822490 | 8 | 0.919830358771444 |
| chrX-107209915-107210416 | 8 | 0.995052005997306 |
| chr17-55909042-55909543 | 8 | 0.989806490383903 |
| chr14-70195456-70195957 | 3 | 0.99593553545041 |
| chr9-17615830-17616331 | 3 | 0.825502172720554 |
| chrX-66857948-66858449 | 4 | 0.872970961573324 |

| | | |
|---|---|---|
| chr19-22606708-22607209 | 2 | 0.841102403496086 |
| chr17-28438323-28438824 | 8 | 0.859320864082955 |
| chr7-118599264-118599765 | 3 | 0.994110621445356 |
| chr3-34450423-34450924 | 4 | 0.911347099628928 |
| chr13-74238119-74238620 | 8 | 0.900122390516966 |
| chr1-96407412-96407913 | 4 | 0.912988661267657 |
| chr10-22565729-22566230 | 3 | 0.952215176748276 |
| chr5-139471720-139472221 | 8 | 0.987526267699681 |
| chr11-9374972-9375473 | 3 | 0.997129351482363 |
| chr5-118544284-118544785 | 3 | 0.99593553545041 |
| chr4-69608898-69609399 | 6 | 0.992767084317855 |
| chr14-26414733-26415234 | 8 | 0.994628996146081 |
| chr5-47181789-47182290 | 5 | 0.972826702477456 |
| chr4-28933755-28934256 | 8 | 0.964915518050291 |
| chr6-25759475-25759976 | 8 | 0.897582012810224 |
| chr8-71523140-71523641 | 8 | 0.958088505633423 |
| chr5-60378782-60379283 | 8 | 0.918501671850009 |
| chr14-11663752-11664253 | 4 | 0.880035526512036 |
| chr1-4516797-4517298 | 2 | 0.961132738368366 |
| chr9-74291854-74292355 | 4 | 0.945826521624899 |
| chr18-51330398-51330899 | 5 | 0.965302599454848 |
| chr7-98056847-98057348 | 8 | 0.995163248720867 |
| chr15-69379309-69379810 | 8 | 0.994984570797748 |
| chr6-124729164-124729665 | 6 | 0.992632268373842 |
| chr9-54937079-54937580 | 8 | 0.894412161326734 |
| chr10-16811019-16811520 | 3 | 0.946059233515496 |
| chr14-50309682-50310183 | 3 | 0.925208025569064 |
| chr2-107837402-107837903 | 6 | 0.921012405492034 |
| chr3-42927844-42928345 | 5 | 0.986833956728175 |
| chr15-81359155-81359656 | 8 | 0.97836804998301 |
| chr16-51058115-51058616 | 6 | 0.992975776407721 |
| chr19-19068629-19069130 | 5 | 0.981239708170695 |
| chr9-83159007-83159508 | 8 | 0.971587872196102 |
| chr14-72582342-72582843 | 8 | 0.870977646666463 |
| chr2-96524191-96524692 | 8 | 0.937302773337502 |
| chr3-79958596-79959097 | 2 | 0.945107474391192 |
| chr8-76265409-76265910 | 8 | 0.883482941545581 |
| chr5-91268083-91268584 | 3 | 0.893233372813222 |
| chr3-104715258-104715759 | 8 | 0.993493681128311 |
| chr9-20634056-20634557 | 6 | 0.934478743584344 |
| chr8-79201086-79201587 | 3 | 0.996431649522412 |
| chr3-50136317-50136818 | 7 | 0.857586992831754 |
| chr10-21185558-21186059 | 3 | 0.990705200503806 |

| | | |
|---|---|---|
| chr17-25239008-25239509 | 5 | 0.828852394804459 |
| chr10-11842572-11843073 | 8 | 0.986343424516356 |
| chr8-24987829-24988330 | 6 | 0.991206925481043 |
| chr2-196902488-196902989 | 8 | 0.971223982624876 |
| chr8-54464821-54465322 | 8 | 0.993493681128311 |
| chr13-28591349-28591850 | 3 | 0.996886547761662 |
| chr13-55937272-55937773 | 8 | 0.965007827717601 |
| chr7-116584515-116585016 | 4 | 0.958764678393689 |
| chr10-21428896-21429397 | 2 | 0.819437740162254 |
| chr4-89889615-89890116 | 7 | 0.927879312931833 |
| chr8-109019798-109020299 | 5 | 0.820294069188645 |
| chr5-24064908-24065409 | 8 | 0.888589697611554 |
| chr4-6046525-6047026 | 5 | 0.991018852324318 |
| chr1-120924939-120925440 | 8 | 0.967980496686923 |
| chr2-6002397-6002898 | 8 | 0.993493681128311 |
| chr2-75950414-75950915 | 6 | 0.991206925481043 |
| chr8-59292644-59293145 | 5 | 0.972705741226798 |
| chr2-196773624-196774125 | 8 | 0.984442048546935 |
| chr8-40608928-40609429 | 7 | 0.96593639626933 |
| chr16-76105630-76106131 | 3 | 0.892999179304051 |
| chr19-777133-777634 | 5 | 0.974092719834373 |
| chr6-45428572-45429073 | 8 | 0.835072611372555 |
| chr9-109023669-109024170 | 4 | 0.967695859871374 |
| chr15-32524228-32524729 | 8 | 0.972140123146653 |
| chr3-146051753-146052254 | 5 | 0.893364687615358 |
| chr9-18524489-18524990 | 6 | 0.891100364296639 |
| chr4-116629269-116629770 | 8 | 0.969622836267471 |
| chr18-64519812-64520313 | 3 | 0.979268695013076 |
| chr18-63688419-63688920 | 3 | 0.99593553545041 |
| chr19-2932128-2932629 | 6 | 0.987604155305306 |
| chr1-26868648-26869149 | 8 | 0.994733198873361 |
| chr13-80665257-80665758 | 8 | 0.982559562584619 |
| chr1-231763200-231763701 | 4 | 0.863452598690049 |
| chr17-32117681-32118182 | 8 | 0.98937985468512 |
| chr19-36993317-36993818 | 3 | 0.996178664084164 |
| chr13-70284402-70284903 | 6 | 0.82067412866545 |
| chr7-41990809-41991310 | 2 | 0.94207443804137 |
| chr13-55075326-55075827 | 8 | 0.977497307670861 |
| chr18-38833435-38833936 | 8 | 0.994903371866194 |
| chr5-1115460-1115961 | 3 | 0.99593553545041 |
| chr16-67956349-67956850 | 4 | 0.950919104712486 |
| chr17-65442867-65443368 | 5 | 0.978452389567946 |
| chrX-102521134-102521635 | 8 | 0.985903027075156 |

| | | |
|---|---|---|
| chr7-94914804-94915305 | 6 | 0.991206925481043 |
| chr14-79419475-79419976 | 8 | 0.967685702445267 |
| chr18-18024899-18025400 | 3 | 0.99593553545041 |
| chr6-68920147-68920648 | 8 | 0.861192582089687 |
| chrX-106050925-106051426 | 8 | 0.995002793209412 |
| chr2-211802014-211802515 | 8 | 0.933866628694948 |
| chr14-98404193-98404694 | 8 | 0.900090031337882 |
| chr1-94001714-94002215 | 8 | 0.983047046360687 |
| chr12-35934190-35934691 | 2 | 0.825212919192565 |
| chrX-149688313-149688814 | 7 | 0.87545570410039 |
| chr17-74969717-74970218 | 8 | 0.946409167358261 |
| chr9-54039004-54039505 | 2 | 0.868241713397028 |
| chrX-25517707-25518208 | 3 | 0.99593553545041 |
| chrY-12236949-12237450 | 3 | 0.99593553545041 |
| chr14-55277833-55278334 | 5 | 0.828681447513633 |
| chr20-18111990-18112491 | 5 | 0.865218121507898 |
| chr4-20682976-20683477 | 8 | 0.931528205269964 |
| chr1-200775629-200776130 | 8 | 0.818234542614071 |
| chrX-22363072-22363573 | 3 | 0.996887232321431 |
| chr13-23896553-23897054 | 6 | 0.992460022488544 |
| chr7-30345946-30346447 | 5 | 0.912402794018495 |
| chr1-123433866-123434367 | 8 | 0.947909263317323 |
| chr20-29134029-29134530 | 8 | 0.900591785326662 |
| chr11-32832627-32833128 | 8 | 0.963479540247309 |
| chr14-74223225-74223726 | 8 | 0.871627796572032 |
| chr3-21426854-21427355 | 4 | 0.972606998052729 |
| chr3-98020712-98021213 | 7 | 0.858477055117039 |
| chr17-51360635-51361136 | 5 | 0.980783542083334 |
| chr4-166777379-166777880 | 8 | 0.943667015580145 |
| chr3-120569177-120569678 | 2 | 0.947876111482309 |
| chr18-59637199-59637700 | 8 | 0.9646496032933 |
| chr4-112775923-112776424 | 6 | 0.992177371131923 |
| chr1-200256976-200257477 | 1 | 0.901317991119978 |
| chr5-13943466-13943967 | 2 | 0.90233642450812 |
| chr3-133668013-133668514 | 2 | 0.849819971297052 |
| chr6-35571909-35572410 | 8 | 0.993493681128311 |
| chr13-90297591-90298092 | 5 | 0.981231670357095 |
| chr14-1542252-1542753 | 2 | 0.90449467632867 |
| chr2-34624915-34625416 | 8 | 0.993493681128311 |
| chr14-83246128-83246629 | 8 | 0.826333742202405 |
| chr17-23074842-23075343 | 2 | 0.956924206382789 |
| chr7-51574324-51574825 | 8 | 0.982525953530884 |
| chr5-104546466-104546967 | 6 | 0.991206925481043 |

| | | |
|---|---|---|
| chr15-4151102-4151603 | 7 | 0.916489889401311 |
| chr4-142926884-142927385 | 8 | 0.879000257369991 |
| chr5-7233629-7234130 | 4 | 0.867943125517013 |
| chr3-168896270-168896771 | 8 | 0.987640982261403 |
| chr13-79980966-79981467 | 3 | 0.996346183696429 |
| chr10-76129922-76130423 | 6 | 0.89693839654183 |
| chr20-46209881-46210382 | 8 | 0.995094111543492 |
| chr1-243107373-243107874 | 5 | 0.953389151169571 |
| chr18-39050151-39050652 | 8 | 0.966672005653966 |
| chr16-84201525-84202026 | 3 | 0.996845570573741 |
| chr8-89895226-89895727 | 2 | 0.859401356142694 |
| chr8-115343813-115344314 | 8 | 0.990002590506226 |
| chr8-114629586-114630087 | 8 | 0.958206710296252 |
| chr4-148219985-148220486 | 4 | 0.947974041088069 |
| chr11-50793638-50794139 | 8 | 0.902405183854313 |
| chr4-37140439-37140940 | 8 | 0.931128596716989 |
| chr16-79666541-79667042 | 2 | 0.907471890428137 |
| chr9-61914949-61915450 | 4 | 0.870529301330633 |
| chr2-95427133-95427634 | 8 | 0.996742047047748 |
| chr6-137932587-137933088 | 8 | 0.975354499728617 |
| chr8-41937269-41937770 | 5 | 0.988088840414551 |
| chr5-58423251-58423752 | 8 | 0.852544579231162 |
| chr7-51959681-51960182 | 1 | 0.940878437142135 |
| chr1-194640612-194641113 | 1 | 0.972701068759264 |
| chr15-5467159-5467660 | 4 | 0.913362221755064 |
| chr8-2695593-2696094 | 3 | 0.839600441487201 |
| chr2-72093450-72093951 | 6 | 0.991206925481043 |
| chr15-91473263-91473764 | 6 | 0.968680138257616 |
| chr9-19701269-19701770 | 3 | 0.952214946216998 |
| chr2-23354854-23355355 | 1 | 0.870280302238532 |
| chr4-132342064-132342565 | 8 | 0.934172051971441 |
| chrX-136464266-136464767 | 8 | 0.994913456169717 |
| chr2-87348095-87348596 | 3 | 0.99593553545041 |
| chr6-94464394-94464895 | 8 | 0.83290550793177 |
| chr11-44891494-44891995 | 5 | 0.975808278198907 |
| chr19-53754123-53754624 | 5 | 0.955563601873066 |
| chr1-94317391-94317892 | 8 | 0.99521684145085 |
| chr19-8253546-8254047 | 5 | 0.833143574504918 |
| chr7-58518124-58518625 | 5 | 0.858561018882884 |
| chr6-47036224-47036725 | 8 | 0.861251820464816 |
| chr1-88386876-88387377 | 3 | 0.996877909999139 |
| chr7-121257020-121257521 | 8 | 0.979801020418994 |
| chr3-90233216-90233717 | 8 | 0.992936427154476 |

| | | |
|---|---|---|
| chr1-81149303-81149804 | 8 | 0.995111815745234 |
| chr9-79875397-79875898 | 3 | 0.859096198992981 |
| chr15-47107851-47108352 | 5 | 0.91793698298068 |
| chr4-128841968-128842469 | 8 | 0.965772062948387 |
| chr3-62718347-62718848 | 8 | 0.944617686679635 |
| chr10-38314913-38315414 | 1 | 0.918668708254196 |
| chr9-26313388-26313889 | 8 | 0.951407714366759 |
| chr10-9674891-9675392 | 8 | 0.993493681128311 |
| chr3-167977716-167978217 | 4 | 0.92604235238454 |
| chr4-52697101-52697602 | 1 | 0.819795085220409 |
| chr10-81224605-81225106 | 3 | 0.847189758557809 |
| chr7-121316171-121316672 | 2 | 0.90193505200734 |
| chr10-72037045-72037546 | 4 | 0.961789734567964 |
| chr4-30299735-30300236 | 6 | 0.988182675486441 |
| chr7-22602021-22602522 | 2 | 0.940062945052445 |
| chr14-59016082-59016583 | 3 | 0.978339161100204 |
| chr3-12120692-12121193 | 4 | 0.82057477453474 |
| chr12-6092345-6092846 | 8 | 0.904964798581101 |
| chr11-66471005-66471506 | 4 | 0.928791402600714 |
| chr1-73422772-73423273 | 8 | 0.8785344833579 |
| chr6-39834472-39834973 | 2 | 0.960119516354666 |
| chr18-66574560-66575061 | 6 | 0.991206925481043 |
| chr5-19873398-19873899 | 5 | 0.924982452635765 |
| chr7-93910638-93911139 | 4 | 0.963915894100042 |
| chr17-32244236-32244737 | 8 | 0.890133920390801 |
| chr9-37131319-37131820 | 1 | 0.950236517196528 |
| chr13-73984493-73984994 | 1 | 0.839645369075434 |
| chr6-42218373-42218874 | 3 | 0.996730749916106 |
| chr18-2980155-2980656 | 8 | 0.940391063209479 |
| chr9-77804673-77805174 | 5 | 0.978401504166141 |
| chr11-72992051-72992552 | 3 | 0.996063492422171 |
| chr17-72670059-72670560 | 4 | 0.936927005992552 |
| chr5-79168571-79169072 | 8 | 0.986632870948215 |
| chr4-168702923-168703424 | 7 | 0.844837711675514 |
| chr19-55992160-55992661 | 5 | 0.968343603783954 |
| chr14-54899530-54900031 | 6 | 0.991206925481043 |
| chr5-15112192-15112693 | 5 | 0.918527321147466 |
| chr13-99003989-99004490 | 7 | 0.917219297934109 |
| chr7-31495965-31496466 | 4 | 0.934711985520245 |
| chr14-61845379-61845880 | 8 | 0.96655374914056 |
| chr3-7833386-7833887 | 8 | 0.862236110191537 |
| chr18-56918149-56918650 | 8 | 0.826837528218908 |
| chr11-24876310-24876811 | 8 | 0.946855192782086 |

| | | |
|---|---|---|
| chr10-73710476-73710977 | 8 | 0.944499487736801 |
| chr20-29050033-29050534 | 8 | 0.845672529613259 |
| chr3-72320744-72321245 | 3 | 0.990232081472943 |
| chr2-114393644-114394145 | 3 | 0.987198598015373 |
| chr6-61225601-61226102 | 8 | 0.833212412500923 |
| chr1-145258336-145258837 | 4 | 0.842200258687162 |
| chr3-140105329-140105830 | 2 | 0.974319637669607 |
| chr19-44765405-44765906 | 4 | 0.972676112190696 |
| chr5-140625280-140625781 | 8 | 0.916847571385935 |
| chr10-50210886-50211387 | 8 | 0.949301511762816 |
| chrX-69575412-69575913 | 8 | 0.993493681128311 |
| chr12-27838203-27838704 | 8 | 0.969756664781529 |
| chr15-47212685-47213186 | 6 | 0.983843837951854 |
| chr3-126252486-126252987 | 3 | 0.91653616173721 |
| chr6-48185679-48186180 | 8 | 0.993493681128311 |
| chr3-30743830-30744331 | 5 | 0.988772208613533 |
| chr10-37168171-37168672 | 1 | 0.889607666864063 |
| chr3-84894629-84895130 | 3 | 0.995937706095473 |
| chr5-90342102-90342603 | 8 | 0.99524742175704 |
| chr14-43822028-43822529 | 5 | 0.976780138180194 |
| chr6-25855734-25856235 | 5 | 0.978208767523242 |
| chr7-103765631-103766132 | 3 | 0.99593553545041 |
| chr3-89652709-89653210 | 8 | 0.848274912358663 |
| chr1-227180014-227180515 | 6 | 0.974860877204188 |
| chr5-154053632-154054133 | 3 | 0.879570587403465 |
| chr14-7926496-7926997 | 8 | 0.995107512676359 |
| chr1-45180503-45181004 | 8 | 0.874013198696659 |
| chr3-117839903-117840404 | 4 | 0.825606886192917 |
| chr2-174922761-174923262 | 7 | 0.934906333001482 |
| chr8-62444344-62444845 | 8 | 0.960215952628669 |
| chr3-53624289-53624790 | 8 | 0.915332127779345 |
| chr2-111504296-111504797 | 8 | 0.99289072927114 |
| chr4-50977237-50977738 | 8 | 0.947387146731972 |
| chr11-3451456-3451957 | 8 | 0.97659266619405 |
| chr9-110119670-110120171 | 8 | 0.993493681128311 |
| chr9-13964018-13964519 | 2 | 0.841591918386799 |
| chr20-1860567-1861068 | 3 | 0.99593553545041 |
| chr8-11006412-11006913 | 8 | 0.99518723978315 |
| chr13-69499585-69500086 | 8 | 0.872410560593003 |
| chr3-149655750-149656251 | 8 | 0.958703772046057 |
| chr10-35359750-35360251 | 3 | 0.997724576019213 |
| chr7-92300309-92300810 | 8 | 0.989936145363549 |
| chr15-81636177-81636678 | 5 | 0.889713157804559 |

| | | |
|---|---|---|
| chr20-39157855-39158356 | 2 | 0.956449007786013 |
| chr14-76495380-76495881 | 8 | 0.993921764774891 |
| chr4-112976929-112977430 | 8 | 0.938155916801122 |
| chr4-143819149-143819650 | 8 | 0.990369355243816 |
| chr2-38934363-38934864 | 7 | 0.914424613505648 |
| chr5-134263944-134264445 | 8 | 0.90304829317776 |
| chr11-29630534-29631035 | 8 | 0.994875531526589 |
| chr11-56784210-56784711 | 8 | 0.874415127528318 |
| chr1-176466701-176467202 | 8 | 0.921986621638167 |
| chr3-120745171-120745672 | 8 | 0.950102524141776 |
| chr13-35948514-35949015 | 8 | 0.909316477415961 |
| chr3-102081246-102081747 | 3 | 0.987713782575738 |
| chr1-152929802-152930303 | 4 | 0.970640713589019 |
| chr2-59491383-59491884 | 8 | 0.993493681128311 |
| chr19-10273978-10274479 | 2 | 0.878909527273962 |
| chr5-80286206-80286707 | 8 | 0.97627338531855 |
| chr11-60893487-60893988 | 3 | 0.99593553545041 |
| chr1-27013564-27014065 | 3 | 0.838252445875821 |
| chr8-66162585-66163086 | 8 | 0.875414553432745 |
| chr7-36220250-36220751 | 3 | 0.977330797520905 |
| chr10-77402545-77403046 | 7 | 0.879727955970771 |
| chr3-157747580-157748081 | 4 | 0.937249510202195 |
| chr9-106361128-106361629 | 5 | 0.976767898491311 |
| chr1-186799302-186799803 | 8 | 0.944037097646307 |
| chr10-70421172-70421673 | 7 | 0.938296088905008 |
| chr19-46823783-46824284 | 1 | 0.905182844436066 |
| chr8-63914573-63915074 | 6 | 0.991206925481043 |
| chr9-111973604-111974105 | 8 | 0.850503658857209 |
| chr4-33932689-33933190 | 8 | 0.9700944388532 |
| chr16-34825635-34826136 | 3 | 0.99593553545041 |
| chr13-18888730-18889231 | 3 | 0.996585154230841 |
| chr13-31435196-31435697 | 4 | 0.973089858123216 |
| chr4-16379485-16379986 | 3 | 0.991610140907243 |
| chr14-95282718-95283219 | 5 | 0.978921268264901 |
| chr3-147125769-147126270 | 4 | 0.89207649533636 |
| chr3-94672368-94672869 | 6 | 0.982446337528536 |
| chr9-110967558-110968059 | 8 | 0.865899005010746 |
| chr4-51835641-51836142 | 5 | 0.824128855643414 |
| chr6-33891993-33892494 | 8 | 0.962290772493823 |
| chr12-7850240-7850741 | 7 | 0.860897880257941 |
| chr4-72571036-72571537 | 4 | 0.857267664702097 |
| chr8-80480970-80481471 | 8 | 0.993493681128311 |
| chr19-30612479-30612980 | 8 | 0.985164712679928 |

| | | |
|---|---|---|
| chr3-48785437-48785938 | 7 | 0.830776118950477 |
| chr3-18639468-18639969 | 6 | 0.950037304232177 |
| chr10-57340148-57340649 | 8 | 0.929747798406893 |
| chr14-44569040-44569541 | 1 | 0.929424167066804 |
| chr4-36348922-36349423 | 8 | 0.96869148165688 |
| chr18-18079477-18079978 | 8 | 0.994230550633762 |
| chr19-52317230-52317731 | 6 | 0.99203331887105 |
| chr14-77281926-77282427 | 8 | 0.851941016396359 |
| chr4-135574926-135575427 | 8 | 0.895939769642019 |
| chr17-6057701-6058202 | 4 | 0.870640033820552 |
| chr14-71647004-71647505 | 3 | 0.996567021923991 |
| chr5-19017678-19018179 | 2 | 0.911833915454793 |
| chr3-166113568-166114069 | 4 | 0.853542850451787 |
| chr7-42228917-42229418 | 6 | 0.989596012842031 |
| chr8-54668639-54669140 | 2 | 0.955064886781931 |
| chr15-8990438-8990939 | 4 | 0.870113138018362 |
| chr2-16216900-16217401 | 4 | 0.839607383555552 |
| chr9-105765370-105765871 | 7 | 0.824212056542669 |
| chr14-9314240-9314741 | 8 | 0.971999681030056 |
| chr5-80095673-80096174 | 8 | 0.994006380158232 |
| chr1-178402824-178403325 | 8 | 0.994486910796559 |
| chr8-115837003-115837504 | 4 | 0.933477269206838 |
| chr6-139320433-139320934 | 8 | 0.911798317759194 |
| chr7-21606092-21606593 | 3 | 0.993142402154794 |
| chr4-24940719-24941220 | 8 | 0.907233281698052 |
| chr14-2110011-2110512 | 8 | 0.864750365852919 |
| chr14-79942599-79943100 | 8 | 0.854454896899751 |
| chr6-31665738-31666239 | 8 | 0.998107093033321 |
| chr11-63140480-63140981 | 6 | 0.959533159842673 |
| chr6-83883856-83884357 | 8 | 0.987703775947184 |
| chr18-71701258-71701759 | 8 | 0.991214599743551 |
| chr13-38962958-38963459 | 5 | 0.987813856560703 |
| chr15-49115784-49116285 | 1 | 0.818011986775912 |
| chr2-23337419-23337920 | 1 | 0.960541213262735 |
| chr1-183517053-183517554 | 8 | 0.993493681128311 |
| chr3-16606391-16606892 | 8 | 0.913796831049161 |
| chr6-45866820-45867321 | 8 | 0.95235203111621 |
| chr14-43453956-43454457 | 3 | 0.847284412021467 |
| chr12-26775418-26775919 | 8 | 0.970798408297815 |
| chr7-129389600-129390101 | 2 | 0.948033260364545 |
| chrX-103444042-103444543 | 7 | 0.935341590191523 |
| chr1-155734717-155735218 | 3 | 0.996157485144357 |
| chr2-128571903-128572404 | 6 | 0.974387489065291 |

| | | |
|---|---|---|
| chr10-27563797-27564298 | 8 | 0.993896513430885 |
| chr8-24052860-24053361 | 8 | 0.947755951924717 |
| chr18-57109940-57110441 | 8 | 0.989920783426058 |
| chr4-176503127-176503628 | 8 | 0.968422473916842 |
| chr17-54616352-54616853 | 7 | 0.904741479320616 |
| chr7-32065467-32065968 | 8 | 0.897053938901297 |
| chr16-7028932-7029433 | 8 | 0.886852192803825 |
| chr5-151006955-151007456 | 8 | 0.993493681128311 |
| chr10-60455058-60455559 | 8 | 0.832939643357165 |
| chr18-38964740-38965241 | 5 | 0.955893268447577 |
| chr4-128505337-128505838 | 8 | 0.993493681128311 |
| chr1-125151088-125151589 | 4 | 0.967321125230246 |
| chr18-59497643-59498144 | 4 | 0.825797953425729 |
| chr6-117027841-117028342 | 3 | 0.982429104635777 |
| chr5-103739923-103740424 | 3 | 0.994700856712992 |
| chr15-78792074-78792575 | 8 | 0.968617241998961 |
| chr7-72594875-72595376 | 8 | 0.99280484068256 |
| chr3-128199751-128200252 | 8 | 0.87074085545445 |
| chr19-46572467-46572968 | 5 | 0.973403558942707 |
| chr16-56395805-56396306 | 8 | 0.992047229664038 |
| chr9-36025503-36026004 | 4 | 0.835091470186415 |
| chr1-246050590-246051091 | 8 | 0.898631711972861 |
| chr13-51679807-51680308 | 8 | 0.873087532065401 |
| chr18-34406313-34406814 | 8 | 0.974209946571518 |
| chr19-12991551-12992052 | 8 | 0.881561952650116 |
| chr11-78839349-78839850 | 2 | 0.888831597266077 |
| chr19-32019349-32019850 | 8 | 0.954983608428712 |
| chr2-244803476-244803977 | 8 | 0.993493681128311 |
| chrX-26735131-26735632 | 8 | 0.982113822082181 |
| chr4-12012027-12012528 | 8 | 0.994914661026048 |
| chr7-52757521-52758022 | 8 | 0.974247084510951 |
| chr2-149380260-149380761 | 6 | 0.839886086574814 |
| chr18-49203469-49203970 | 8 | 0.869420284873777 |
| chr2-155902670-155903171 | 6 | 0.99179439619202 |
| chr1-188461786-188462287 | 8 | 0.988004046449179 |
| chr9-67171145-67171646 | 1 | 0.903429628823564 |
| chr4-33543416-33543917 | 5 | 0.987145484628869 |
| chr1-172262584-172263085 | 5 | 0.984054222551975 |
| chr19-44141651-44142152 | 8 | 0.92091962483278 |
| chr16-71580698-71581199 | 4 | 0.877249518813051 |
| chr1-17842999-17843500 | 8 | 0.96972838608704 |
| chr15-22648848-22649349 | 3 | 0.959502479000444 |
| chr17-21934698-21935199 | 8 | 0.837943326502625 |

| | | |
|---|---|---|
| chr20-16775131-16775632 | 2 | 0.918574220713809 |
| chr13-34846884-34847385 | 8 | 0.885973357416985 |
| chr5-11717786-11718287 | 3 | 0.99567456827829 |
| chr4-14068654-14069155 | 8 | 0.946406040161232 |
| chr2-220729003-220729504 | 8 | 0.989399359235427 |
| chr2-246727520-246728021 | 2 | 0.95821889934936 |
| chrX-20008904-20009405 | 5 | 0.969332086951009 |
| chr6-100664874-100665375 | 2 | 0.963828468188677 |
| chr7-393211-393712 | 8 | 0.906270911151635 |
| chr2-224664622-224665123 | 8 | 0.908735653460186 |
| chr1-188544953-188545454 | 5 | 0.937493605415985 |
| chr4-72226740-72227241 | 1 | 0.828095478379759 |
| chr10-95610647-95611148 | 3 | 0.99593553545041 |
| chr18-5125966-5126467 | 6 | 0.941343356631818 |
| chr10-33488782-33489283 | 6 | 0.867435776341337 |
| chr5-68127574-68128075 | 2 | 0.928657385602376 |
| chr18-27781531-27782032 | 8 | 0.993493681128311 |
| chr19-47086166-47086667 | 4 | 0.857805892588374 |
| chr16-37291724-37292225 | 8 | 0.993493681128311 |
| chr5-138688954-138689455 | 8 | 0.994251746827974 |
| chr4-27906655-27907156 | 8 | 0.89904268739642 |
| chr2-11227606-11228107 | 4 | 0.950890432105568 |
| chr1-235880903-235881404 | 8 | 0.99164362429081 |
| chr8-3379622-3380123 | 6 | 0.869881279536551 |
| chr7-34476517-34477018 | 3 | 0.986890921190558 |
| chr17-15155459-15155960 | 8 | 0.847661891003124 |
| chr5-143309126-143309627 | 8 | 0.835098036069465 |
| chr1-67683751-67684252 | 8 | 0.995158731736159 |
| chr10-85650665-85651166 | 8 | 0.892935441165834 |
| chr7-15514697-15515198 | 3 | 0.984660441832894 |
| chr3-122587960-122588461 | 6 | 0.937085982723313 |
| chr2-35991289-35991790 | 8 | 0.979450462921702 |
| chr6-16051456-16051957 | 4 | 0.92197538161902 |
| chr18-4699299-4699800 | 1 | 0.948348043530613 |
| chr14-37580455-37580956 | 3 | 0.98602390986318 |
| chr13-6669233-6669734 | 4 | 0.828702286381286 |
| chr19-26316450-26316951 | 5 | 0.929323881904949 |
| chr4-153767842-153768343 | 8 | 0.953728426260234 |
| chr16-46163785-46164286 | 8 | 0.835966410901623 |
| chr5-60187259-60187760 | 8 | 0.88825719074631 |
| chr1-51830249-51830750 | 5 | 0.976302652615347 |
| chr4-7078789-7079290 | 8 | 0.989775219768734 |
| chr1-205042581-205043082 | 2 | 0.8765250887728 |

| | | |
|---|---|---|
| chr16-48348596-48349097 | 6 | 0.914181936587034 |
| chr5-23574683-23575184 | 8 | 0.961084739757412 |
| chr10-40670769-40671270 | 3 | 0.85232283786196 |
| chr18-61183578-61184079 | 8 | 0.90589236561959 |
| chr9-24369022-24369523 | 8 | 0.891110619372995 |
| chr2-237104235-237104736 | 8 | 0.880401622512193 |
| chr2-37684070-37684571 | 3 | 0.996035706029003 |
| chr10-35019258-35019759 | 8 | 0.993493681128311 |
| chr4-102326511-102327012 | 8 | 0.89900125169073 |
| chr11-75103584-75104085 | 1 | 0.874434283148653 |
| chr8-102933472-102933973 | 4 | 0.880936110626557 |
| chr6-98296005-98296506 | 5 | 0.993015856640976 |
| chr15-36074249-36074750 | 8 | 0.995570543968209 |
| chr9-57855885-57856386 | 8 | 0.993203362225316 |
| chr7-104809072-104809573 | 8 | 0.906513868628543 |
| chr6-128035367-128035868 | 4 | 0.850870954931833 |
| chr5-79359810-79360311 | 3 | 0.996108644244694 |
| chrX-72041382-72041883 | 3 | 0.99677360755463 |
| chr14-77301826-77302327 | 8 | 0.872681874716974 |
| chr2-55142522-55143023 | 3 | 0.976500945870765 |
| chr20-18766036-18766537 | 5 | 0.932953535312518 |
| chr1-40320903-40321404 | 4 | 0.844625390381248 |
| chr11-16560329-16560830 | 8 | 0.993697487815005 |
| chr14-88359299-88359800 | 3 | 0.991717071735003 |
| chr14-87170263-87170764 | 6 | 0.992561246409448 |
| chr17-82021938-82022439 | 8 | 0.931690241411613 |
| chr5-121202739-121203240 | 8 | 0.9366269892866 |
| chr18-32558468-32558969 | 8 | 0.993493681128311 |
| chr6-87070011-87070512 | 6 | 0.98018434925698 |
| chr4-149902548-149903049 | 8 | 0.886444756886497 |
| chr4-31955382-31955883 | 8 | 0.987059438411618 |
| chr6-128798619-128799120 | 8 | 0.922136152783683 |
| chr8-66932539-66933040 | 8 | 0.913491491731395 |
| chr10-20967214-20967715 | 8 | 0.995079192360108 |
| chr4-70010404-70010905 | 8 | 0.917203724567812 |
| chr8-21846830-21847331 | 4 | 0.963780532376067 |
| chr16-11494446-11494947 | 6 | 0.991206925481043 |
| chr8-82014657-82015158 | 8 | 0.903908996197609 |
| chr11-16067689-16068190 | 8 | 0.994877979334923 |
| chr7-83628544-83629045 | 3 | 0.908589892649537 |
| chr12-16661263-16661764 | 8 | 0.988790031085178 |
| chr10-76451071-76451572 | 8 | 0.935087145411644 |
| chr8-114938975-114939476 | 8 | 0.970433797800041 |

| | | |
|---|---|---|
| chrX-104975261-104975762 | 8 | 0.994329179517048 |
| chr10-26130462-26130963 | 2 | 0.874550374773849 |
| chr1-122167497-122167998 | 2 | 0.858831369066013 |
| chr4-18666438-18666939 | 6 | 0.991206925481043 |
| chr14-54734199-54734700 | 8 | 0.819215272008919 |
| chr3-120005009-120005510 | 5 | 0.989908134703167 |
| chr14-7564699-7565200 | 8 | 0.819679818678789 |
| chr5-156364482-156364983 | 8 | 0.993493681128311 |
| chr19-7930377-7930878 | 3 | 0.988720054089416 |
| chr1-256271271-256271772 | 3 | 0.92407127762409 |
| chr9-949639-950140 | 3 | 0.817239186598749 |
| chr4-96311315-96311816 | 3 | 0.891019471910362 |
| chr17-53605566-53606067 | 8 | 0.861282725572867 |
| chr7-75773874-75774375 | 3 | 0.989375927807545 |
| chr3-109373677-109374178 | 8 | 0.932137539228284 |
| chr1-171858917-171859418 | 8 | 0.951033708671526 |
| chr19-909913-910414 | 5 | 0.983919978840557 |
| chr7-133168849-133169350 | 8 | 0.937697124514833 |
| chr17-19571473-19571974 | 5 | 0.99292592043532 |
| chr10-21076532-21077033 | 4 | 0.926700810547934 |
| chr10-59780170-59780671 | 8 | 0.994217469813083 |
| chr17-60125110-60125611 | 5 | 0.846553915314378 |
| chr10-26670160-26670661 | 8 | 0.897656253128365 |
| chr1-192555282-192555783 | 8 | 0.885000811454886 |
| chr7-30474774-30475275 | 8 | 0.845589980446488 |
| chr1-256291907-256292408 | 8 | 0.993042568138649 |
| chr10-14614316-14614817 | 8 | 0.977437978081699 |
| chr8-89916317-89916818 | 3 | 0.994529425938972 |
| chr6-35294996-35295497 | 8 | 0.894519800348898 |
| chr20-21942816-21943317 | 8 | 0.914139666839507 |
| chr9-62052958-62053459 | 5 | 0.985285513271726 |
| chr5-86575106-86575607 | 8 | 0.969117115946146 |
| chr2-50272332-50272833 | 8 | 0.818871336814979 |
| chr7-96907244-96907745 | 3 | 0.984309330980854 |
| chr12-23939268-23939769 | 4 | 0.877893200964666 |
| chr8-62171393-62171894 | 4 | 0.82433720550739 |
| chr15-41896269-41896770 | 5 | 0.987160464758214 |
| chr16-55785569-55786070 | 8 | 0.866452351945687 |
| chr5-140275682-140276183 | 5 | 0.989036919019091 |
| chr5-40598070-40598571 | 8 | 0.994987067458621 |
| chr9-17969012-17969513 | 3 | 0.996837546832634 |
| chr4-25871884-25872385 | 3 | 0.995236800293273 |
| chr7-59103594-59104095 | 8 | 0.985851376261946 |

| | | |
|---|---|---|
| chr17-58510473-58510974 | 8 | 0.920649070708472 |
| chr5-47273349-47273850 | 8 | 0.927889674633807 |
| chr11-10920722-10921223 | 5 | 0.89446970423549 |
| chr13-73953855-73954356 | 8 | 0.990873314890676 |
| chr14-41830550-41831051 | 3 | 0.846142323106654 |
| chr14-75642427-75642928 | 5 | 0.937461345415426 |
| chr1-131738344-131738845 | 6 | 0.993209707707936 |
| chr13-53642772-53643273 | 8 | 0.830130436466234 |
| chr4-12988817-12989318 | 6 | 0.993124779793147 |
| chr2-117437933-117438434 | 2 | 0.886421701610665 |
| chr20-41833364-41833865 | 8 | 0.861451479503303 |
| chr10-60206548-60207049 | 8 | 0.825314557616972 |
| chr10-33485070-33485571 | 6 | 0.935873085039983 |
| chr2-203639593-203640094 | 8 | 0.886519818837126 |
| chr17-54344014-54344515 | 8 | 0.913828692720139 |
| chr5-158672166-158672667 | 8 | 0.831389397427281 |
| chr2-192210304-192210805 | 8 | 0.841277893095237 |
| chr18-74985021-74985522 | 8 | 0.926372846438783 |
| chr13-68833482-68833983 | 4 | 0.939382807873292 |
| chr14-68722475-68722976 | 6 | 0.992775172226213 |
| chr18-12169179-12169680 | 8 | 0.892499851219279 |
| chr11-73184441-73184942 | 4 | 0.842193373283205 |
| chr1-228359938-228360439 | 3 | 0.99593553545041 |
| chr20-12247649-12248150 | 8 | 0.926834696480619 |
| chr7-52764102-52764603 | 8 | 0.949379702758036 |
| chr14-3047613-3048114 | 8 | 0.954865669901581 |
| chr17-54479236-54479737 | 8 | 0.992760394639137 |
| chr14-53845242-53845743 | 3 | 0.99593553545041 |
| chr19-38508097-38508598 | 8 | 0.993493681128311 |
| chrX-59558181-59558682 | 8 | 0.886834771717421 |
| chr2-82099248-82099749 | 3 | 0.99303735311034 |
| chr7-52580481-52580982 | 4 | 0.94376385277153 |
| chr3-131291785-131292286 | 8 | 0.965723492153487 |
| chr4-41585442-41585943 | 3 | 0.996813335382211 |
| chr7-78677678-78678179 | 8 | 0.842056499218522 |
| chr9-69629511-69630012 | 5 | 0.855631315944658 |
| chr18-13974051-13974552 | 3 | 0.996434843480982 |
| chr9-52398380-52398881 | 6 | 0.863862938098118 |
| chr9-87481481-87481982 | 4 | 0.972899314973445 |
| chr11-85868108-85868609 | 8 | 0.993493681128311 |
| chr19-12413555-12414056 | 7 | 0.918163413319342 |
| chr6-25644547-25645048 | 8 | 0.921339400831225 |
| chr8-99187013-99187514 | 4 | 0.965688414859016 |

| | | |
|---|---|---|
| chr3-94026909-94027410 | 8 | 0.894823680924408 |
| chr13-68940791-68941292 | 8 | 0.995131658671286 |
| chr2-23324018-23324519 | 3 | 0.817297909389018 |
| chr1-191166168-191166669 | 8 | 0.964370449380955 |
| chr3-165597716-165598217 | 2 | 0.821442117920053 |
| chr12-43454316-43454817 | 5 | 0.892733208011164 |
| chr8-106828888-106829389 | 8 | 0.981516864074801 |
| chr4-79304101-79304602 | 8 | 0.917223824878811 |
| chr8-12733597-12734098 | 8 | 0.992372510593451 |
| chr11-20791547-20792048 | 3 | 0.931512212132171 |
| chr7-83083992-83084493 | 8 | 0.889860928461127 |
| chr7-128820390-128820891 | 3 | 0.975048751831748 |
| chr15-80682346-80682847 | 5 | 0.964126061015581 |
| chr7-111537976-111538477 | 8 | 0.981971294696954 |
| chr3-132603623-132604124 | 8 | 0.86589339177396 |
| chr5-107922924-107923425 | 3 | 0.996884337828258 |
| chr8-7810289-7810790 | 3 | 0.893847155127086 |
| chr6-45634965-45635466 | 5 | 0.967475107781517 |
| chr10-69636810-69637311 | 2 | 0.857308561770735 |
| chr17-4570097-4570598 | 8 | 0.993115924208685 |
| chr1-16948146-16948647 | 8 | 0.984102602928831 |
| chr2-208990026-208990527 | 2 | 0.939705356285553 |
| chr4-17179163-17179664 | 4 | 0.896015273165546 |
| chr20-791313-791814 | 8 | 0.966277872982925 |
| chr4-107971164-107971665 | 3 | 0.99593553545041 |
| chr2-27873017-27873518 | 5 | 0.983919978840557 |
| chr6-97601807-97602308 | 3 | 0.99593553545041 |
| chr5-70420722-70421223 | 3 | 0.82544612565298 |
| chr6-122767666-122768167 | 8 | 0.976013002765456 |
| chr18-70042547-70043048 | 5 | 0.846811510831816 |
| chr1-178959778-178960279 | 8 | 0.992769043250022 |
| chr16-1318576-1319077 | 8 | 0.993933761261778 |
| chr5-118867817-118868318 | 3 | 0.997581431100579 |
| chr2-228772954-228773455 | 8 | 0.878325005533019 |
| chr2-207556840-207557341 | 6 | 0.962848801349788 |
| chr17-51552611-51553112 | 2 | 0.945155426568596 |
| chr4-70204621-70205122 | 8 | 0.871671719201235 |
| chr10-106423419-106423920 | 8 | 0.953727425759612 |
| chr15-50838259-50838760 | 2 | 0.914095106998551 |
| chr1-131942385-131942886 | 8 | 0.938807412122011 |
| chr12-26304699-26305200 | 4 | 0.955495166162121 |
| chr11-11111458-11111959 | 8 | 0.95998780339734 |
| chr14-12688346-12688847 | 8 | 0.887289370480919 |

| | | |
|---|---|---|
| chr2-68156606-68157107 | 4 | 0.944673660695072 |
| chr11-22967910-22968411 | 3 | 0.994679711638003 |
| chr10-10272708-10273209 | 1 | 0.848218387408459 |
| chr4-31998228-31998729 | 8 | 0.94069673251728 |
| chr16-60602582-60603083 | 8 | 0.983954109242465 |
| chr13-451497-451998 | 5 | 0.907096765411512 |
| chr5-138744232-138744733 | 8 | 0.859842333940704 |
| chr1-134501418-134501919 | 8 | 0.857161276256807 |
| chr1-173535230-173535731 | 8 | 0.993493681128311 |
| chr4-153225144-153225645 | 8 | 0.936523828242723 |
| chr15-34757441-34757942 | 8 | 0.984444892535828 |
| chr1-249128120-249128621 | 2 | 0.949438175264156 |
| chr2-146678761-146679262 | 3 | 0.99593553545041 |
| chr10-38575170-38575671 | 8 | 0.961935101046623 |
| chr7-106716788-106717289 | 8 | 0.992301295089048 |
| chr1-118670458-118670959 | 7 | 0.815235370067569 |
| chr5-89836166-89836667 | 8 | 0.993493681128311 |
| chr3-142778306-142778807 | 8 | 0.993665155734044 |
| chr20-11669154-11669655 | 4 | 0.899795237737357 |
| chr3-35848111-35848612 | 6 | 0.984218424713879 |
| chr14-2496382-2496883 | 8 | 0.830022757785246 |
| chr11-25001058-25001559 | 8 | 0.880982334539722 |
| chr7-105941848-105942349 | 8 | 0.993493681128311 |
| chr5-35674424-35674925 | 6 | 0.993187938156903 |
| chr6-54392926-54393427 | 8 | 0.860992390028404 |
| chr5-6285031-6285532 | 8 | 0.981181736919399 |
| chr16-49591910-49592411 | 3 | 0.995406033668234 |
| chr14-3921424-3921925 | 8 | 0.922131294564645 |
| chr14-40069215-40069716 | 8 | 0.87228914939855 |
| chr3-55812593-55813094 | 2 | 0.844557 |
| chr1-19439604-19440105 | 7 | 0.969314228411551 |
| chr2-148302502-148303003 | 8 | 0.921851985345098 |
| chr1-30964568-30965069 | 8 | 0.891954690934221 |
| chr8-106008528-106009029 | 8 | 0.962504555743949 |
| chr4-37157466-37157967 | 1 | 0.887144263966975 |
| chr15-90398775-90399276 | 8 | 0.934638403121259 |
| chr20-35903952-35904453 | 2 | 0.894838605375151 |
| chr8-36603613-36604114 | 2 | 0.867145508514133 |
| chr2-56428698-56429199 | 8 | 0.987953112535766 |
| chrX-8860760-8861261 | 3 | 0.99593553545041 |
| chr15-93595921-93596422 | 8 | 0.993493681128311 |
| chr8-82413614-82414115 | 3 | 0.988281309893553 |
| chr10-21430587-21431088 | 1 | 0.976559384859315 |

| | | |
|---|---|---|
| chr4-107623395-107623896 | 3 | 0.996943386121048 |
| chr10-35703063-35703564 | 8 | 0.846169470353004 |
| chr5-100667452-100667953 | 8 | 0.993493681128311 |
| chrX-111105696-111106197 | 7 | 0.939479431978012 |
| chr19-50261718-50262219 | 3 | 0.996584625651969 |
| chr2-194994529-194995030 | 4 | 0.944436728603579 |
| chr7-12258504-12259005 | 8 | 0.993493681128311 |
| chr7-28433404-28433905 | 8 | 0.924216107422009 |
| chr20-44091787-44092288 | 8 | 0.892508519897737 |
| chr5-149580173-149580674 | 8 | 0.897260689175407 |
| chr3-5812405-5812906 | 5 | 0.929921738898632 |
| chr14-32442311-32442812 | 8 | 0.894121282029023 |
| chr2-141956111-141956612 | 4 | 0.973447149806259 |
| chr4-141713298-141713799 | 3 | 0.995497227681099 |
| chr12-6193793-6194294 | 4 | 0.878959728218158 |
| chr9-69101239-69101740 | 1 | 0.929829884153372 |
| chr6-110774477-110774978 | 5 | 0.84061465928441 |
| chr10-95677966-95678467 | 8 | 0.974731458949562 |
| chr17-76921966-76922467 | 8 | 0.851505716593842 |
| chr16-77831659-77832160 | 8 | 0.932552311606501 |
| chr3-167131480-167131981 | 8 | 0.990538125396875 |
| chr3-8230265-8230766 | 3 | 0.9802643750653 |
| chr8-56633173-56633674 | 2 | 0.863831004753834 |
| chr5-102761727-102762228 | 8 | 0.993871078648758 |
| chr6-52785313-52785814 | 8 | 0.893913268156181 |
| chr6-76718779-76719280 | 8 | 0.990673814131895 |
| chr18-44390330-44390831 | 8 | 0.989887961750859 |
| chr5-97864820-97865321 | 8 | 0.964494925129622 |
| chr3-95902511-95903012 | 8 | 0.823969840224213 |
| chr16-29608736-29609237 | 6 | 0.996675133058718 |
| chr13-102014450-102014951 | 3 | 0.98562041325329 |
| chr8-34449263-34449764 | 6 | 0.960665170604316 |
| chr14-67604711-67605212 | 4 | 0.904944054872065 |
| chr16-52754731-52755232 | 3 | 0.99593553545041 |
| chr3-31494094-31494595 | 8 | 0.995119702271244 |
| chr6-125878766-125879267 | 8 | 0.936995195557881 |
| chr17-37331457-37331958 | 6 | 0.936778398375041 |
| chr14-59518249-59518750 | 8 | 0.982099565323434 |
| chr3-1612990-1613491 | 2 | 0.968775881877449 |
| chrX-20279470-20279971 | 8 | 0.914823823443418 |
| chr2-235519930-235520431 | 5 | 0.962170240754501 |
| chr8-75010122-75010623 | 4 | 0.904267648700211 |
| chr17-1059658-1060159 | 8 | 0.932477311924425 |

| | | |
|---|---|---|
| chr4-34482202-34482703 | 3 | 0.995712555119044 |
| chr9-27438692-27439193 | 3 | 0.997015006661942 |
| chr3-86169881-86170382 | 3 | 0.995909427885616 |
| chr18-72750217-72750718 | 8 | 0.968834156160438 |
| chr6-69597678-69598179 | 2 | 0.872003424293496 |
| chr20-45048156-45048657 | 5 | 0.991553562967622 |
| chr13-90102252-90102753 | 4 | 0.848504628817782 |
| chr2-22124771-22125272 | 8 | 0.922831465967122 |
| chr9-83535789-83536290 | 8 | 0.89523276868947 |
| chr9-85229981-85230482 | 3 | 0.99539721792479 |
| chr8-3445563-3446064 | 8 | 0.974477070557981 |
| chr15-420810-421311 | 8 | 0.995183046259004 |
| chr5-1307808-1308309 | 3 | 0.837944720661436 |
| chr3-116908510-116909011 | 2 | 0.929053379482433 |
| chr8-24968004-24968505 | 8 | 0.885007264272315 |
| chr7-32790363-32790864 | 8 | 0.818882209248107 |
| chr17-78941600-78942101 | 3 | 0.99593553545041 |
| chr2-134902435-134902936 | 4 | 0.865876949744782 |
| chr4-70555-71056 | 5 | 0.960216925751938 |
| chr1-6306655-6307156 | 3 | 0.995888232434425 |
| chr1-99855476-99855977 | 8 | 0.979423467408372 |
| chr4-45930472-45930973 | 1 | 0.826673368952453 |
| chr13-40686918-40687419 | 8 | 0.954875343553898 |
| chr9-32798537-32799038 | 8 | 0.993493681128311 |
| chr18-50925696-50926197 | 2 | 0.952007301311143 |
| chr2-113438691-113439192 | 5 | 0.844854382938149 |
| chr7-33382548-33383049 | 8 | 0.854459314536498 |
| chr16-52217996-52218497 | 8 | 0.849953956890896 |
| chr8-78507323-78507824 | 8 | 0.976972264215283 |
| chr1-234735761-234736262 | 8 | 0.994202115577943 |
| chr15-31783924-31784425 | 4 | 0.927706777373632 |
| chr3-144100569-144101070 | 8 | 0.863302716958428 |
| chr4-122080565-122081066 | 8 | 0.88000556015998 |
| chr2-87942413-87942914 | 8 | 0.995178629537486 |
| chr9-4853902-4854403 | 3 | 0.958433412573605 |
| chr8-80032497-80032998 | 8 | 0.886817015993442 |
| chr17-23958857-23959358 | 8 | 0.955976678014836 |
| chr8-117731741-117732242 | 4 | 0.819075064985261 |
| chr19-21147982-21148483 | 6 | 0.824498642579051 |
| chr5-33602648-33603149 | 8 | 0.967880582275327 |
| chr14-58755551-58756052 | 5 | 0.983919978840557 |
| chr1-221902737-221903238 | 2 | 0.90614258978986 |
| chr16-76952903-76953404 | 8 | 0.993493681128311 |

| | | |
|---|---|---|
| chr1-156229531-156230032 | 8 | 0.940737646232077 |
| chr4-105342750-105343251 | 8 | 0.968796244103088 |
| chr16-81092969-81093470 | 3 | 0.970427333499962 |
| chr16-37730251-37730752 | 3 | 0.98676923089446 |
| chr3-99866985-99867486 | 8 | 0.994984774749418 |
| chr11-76750481-76750982 | 8 | 0.963210232928788 |
| chr2-72415797-72416298 | 3 | 0.995916150169789 |
| chr3-149728048-149728549 | 8 | 0.819636783708409 |
| chr15-62117046-62117547 | 5 | 0.953656350967572 |
| chr7-67980053-67980554 | 8 | 0.981771396644968 |
| chr18-77019682-77020183 | 6 | 0.992874061343942 |
| chr9-76606393-76606894 | 8 | 0.878798782124596 |
| chrX-12166868-12167369 | 8 | 0.972065147151949 |
| chr3-125271908-125272409 | 2 | 0.907855295322775 |
| chrX-9835423-9835924 | 8 | 0.995219580918123 |
| chr1-43696203-43696704 | 3 | 0.853394414339003 |
| chr14-92097730-92098231 | 8 | 0.952583682493275 |
| chr9-42456158-42456659 | 3 | 0.940826155312022 |
| chr19-21426191-21426692 | 8 | 0.940954785900604 |
| chr6-101797931-101798432 | 8 | 0.987028030369408 |
| chr20-46138332-46138833 | 8 | 0.836497158107766 |
| chr12-13323534-13324035 | 8 | 0.986674550830588 |
| chr8-21405441-21405942 | 5 | 0.924908147172567 |
| chr9-22204133-22204634 | 8 | 0.906505331709774 |
| chr19-46575310-46575811 | 8 | 0.831333572777075 |
| chr9-5534134-5534635 | 5 | 0.97677194985742 |
| chr8-54423447-54423948 | 4 | 0.951397052049363 |
| chr3-57199629-57200130 | 8 | 0.914853358649472 |
| chr6-139471909-139472410 | 8 | 0.862057189537066 |
| chr15-20691388-20691889 | 3 | 0.855594963303656 |
| chr10-88477373-88477874 | 2 | 0.964721469643017 |
| chr10-104846992-104847493 | 4 | 0.946648151702213 |
| chr2-145559319-145559820 | 8 | 0.989774792279511 |
| chr1-181447562-181448063 | 8 | 0.992011104823382 |
| chr10-12643877-12644378 | 8 | 0.973831554193062 |
| chr3-125389878-125390379 | 1 | 0.960539128549873 |
| chr7-23417449-23417950 | 8 | 0.939232338061884 |
| chr13-3912853-3913354 | 8 | 0.956385058392412 |
| chr10-85970726-85971227 | 5 | 0.977225110126551 |
| chr16-62386435-62386936 | 2 | 0.931942928624724 |
| chr14-74728080-74728581 | 8 | 0.88735774954801 |
| chr1-82340286-82340787 | 3 | 0.970024046708142 |
| chr6-28554723-28555224 | 8 | 0.901245275440265 |

| | | |
|---|---|---|
| chr12-35831818-35832319 | 8 | 0.956918824455432 |
| chr2-126406063-126406564 | 7 | 0.923615008819642 |
| chr8-23177627-23178128 | 3 | 0.92756159546533 |
| chr5-150475054-150475555 | 4 | 0.895085141053783 |
| chr6-25037707-25038208 | 4 | 0.885544918435235 |
| chr12-13797866-13798367 | 8 | 0.994358566280896 |
| chr18-23338011-23338512 | 8 | 0.86903128178108 |
| chr20-24624875-24625376 | 8 | 0.952971844166977 |
| chr9-112171707-112172208 | 4 | 0.860701852467433 |
| chr14-31190203-31190704 | 8 | 0.955442809482693 |
| chr5-66974931-66975432 | 8 | 0.993493681128311 |
| chr5-48912914-48913415 | 8 | 0.969069409177957 |
| chr16-74813163-74813664 | 4 | 0.825037014690545 |
| chr8-52987586-52988087 | 8 | 0.847052907257275 |
| chr16-76104069-76104570 | 3 | 0.965863838598619 |
| chr17-68778074-68778575 | 8 | 0.993493681128311 |
| chr5-63524908-63525409 | 2 | 0.956030055332573 |
| chr16-22288187-22288688 | 8 | 0.88057192350964 |
| chr1-44825857-44826358 | 8 | 0.993493681128311 |
| chrX-71856274-71856775 | 8 | 0.853544644309419 |
| chrX-146738837-146739338 | 6 | 0.991910732343381 |
| chr18-74201079-74201580 | 5 | 0.96200603954612 |
| chr2-151952286-151952787 | 6 | 0.960818125160498 |
| chr3-128234078-128234579 | 8 | 0.90261489529505 |
| chr1-44968379-44968880 | 8 | 0.99423421058019 |
| chr6-139222202-139222703 | 8 | 0.989908042448463 |
| chr7-100189240-100189741 | 3 | 0.887689402519156 |
| chr6-70154770-70155271 | 8 | 0.951752223915151 |
| chr16-41471564-41472065 | 3 | 0.986349388078717 |
| chr14-101923877-101924378 | 6 | 0.902080766384035 |
| chr12-9146362-9146863 | 1 | 0.875347440326739 |
| chr10-98729898-98730399 | 7 | 0.904575797540696 |
| chr2-123396756-123397257 | 8 | 0.976851266108553 |
| chr3-45671018-45671519 | 4 | 0.924921582563135 |
| chr8-121168668-121169169 | 8 | 0.942168336162246 |
| chr2-166442464-166442965 | 8 | 0.931376319361098 |
| chr8-1831659-1832160 | 3 | 0.989648362141776 |
| chr7-116790867-116791368 | 8 | 0.957236750783774 |
| chr17-64087577-64088078 | 4 | 0.891271341780058 |
| chr8-50528423-50528924 | 8 | 0.876077578627966 |
| chr1-137139459-137139960 | 4 | 0.903146207808287 |
| chrX-101037574-101038075 | 6 | 0.991206925481043 |
| chr16-371386-371887 | 8 | 0.995202992231874 |

| | | |
|---|---|---|
| chr15-11742546-11743047 | 8 | 0.993493681128311 |
| chr4-90626162-90626663 | 8 | 0.899001652890025 |
| chr16-689760-690261 | 2 | 0.851704202065732 |
| chr14-43583241-43583742 | 8 | 0.989511244543474 |
| chr6-138433034-138433535 | 5 | 0.954745732953186 |
| chr18-61184143-61184644 | 8 | 0.873005203905064 |
| chr3-112811307-112811808 | 8 | 0.974574371058335 |
| chr1-45588580-45589081 | 8 | 0.852207312196261 |
| chr2-38923168-38923669 | 3 | 0.962494292129165 |
| chr10-41689485-41689986 | 4 | 0.847624522403144 |
| chr13-37574093-37574594 | 3 | 0.968309817278325 |
| chr16-1531644-1532145 | 4 | 0.954660517920697 |
| chr1-79441196-79441697 | 6 | 0.989057789397719 |
| chr20-49537318-49537819 | 8 | 0.978030095323696 |
| chr3-10937789-10938290 | 8 | 0.987624460077484 |
| chr3-146667928-146668429 | 5 | 0.826468827871127 |
| chr2-195798482-195798983 | 8 | 0.970979036900069 |
| chr5-114771940-114772441 | 2 | 0.957611824120021 |
| chr11-50754786-50755287 | 8 | 0.995108609752343 |
| chr3-8518258-8518759 | 5 | 0.993426196478867 |
| chr1-179933398-179933899 | 8 | 0.816116623279226 |
| chr2-95042215-95042716 | 5 | 0.959078361730927 |
| chr6-102811732-102812233 | 4 | 0.82957536275897 |
| chr5-103761366-103761867 | 8 | 0.835655667335068 |
| chr1-134384746-134385247 | 8 | 0.8827771252816 |
| chr18-13443213-13443714 | 8 | 0.905752838459398 |
| chr1-118332284-118332785 | 6 | 0.977216838435538 |
| chr2-78191465-78191966 | 8 | 0.846026168788476 |
| chr10-1342132-1342633 | 6 | 0.991016311444948 |
| chr4-171476425-171476926 | 8 | 0.989484782813041 |
| chr1-214372051-214372552 | 3 | 0.991244731582123 |
| chr18-75326999-75327500 | 8 | 0.847974005200148 |
| chr4-66421826-66422327 | 6 | 0.954614931785484 |
| chr1-172768253-172768754 | 8 | 0.988818416747586 |
| chr4-180029495-180029996 | 6 | 0.874133052114265 |
| chr18-37130126-37130627 | 4 | 0.933132976360223 |
| chr14-9478790-9479291 | 8 | 0.99242007879193 |
| chr5-59050810-59051311 | 5 | 0.988722322664309 |
| chr1-170452549-170453050 | 2 | 0.878437904455003 |
| chr2-237206397-237206898 | 8 | 0.852462007970282 |
| chr7-116140652-116141153 | 8 | 0.993493681128311 |
| chr1-228453490-228453991 | 8 | 0.952533476491339 |
| chr15-9474735-9475236 | 8 | 0.933208250147223 |

| | | |
|---|---|---|
| chr1-96814946-96815447 | 8 | 0.867716936419145 |
| chrX-2878434-2878935 | 1 | 0.878766445977872 |
| chr1-106600873-106601374 | 8 | 0.950383448365123 |
| chr4-126840779-126841280 | 8 | 0.840009507800485 |
| chr19-42527220-42527721 | 5 | 0.985113152002095 |
| chr6-27601464-27601965 | 8 | 0.993493681128311 |
| chr9-86857378-86857879 | 8 | 0.971779317880617 |
| chr15-15355402-15355903 | 8 | 0.898847228435145 |
| chrX-78907562-78908063 | 8 | 0.993493681128311 |
| chr12-41403448-41403949 | 5 | 0.984200062030051 |
| chr1-86872599-86873100 | 8 | 0.850551963068947 |
| chr1-47079123-47079624 | 8 | 0.942566244406527 |
| chr5-58305488-58305989 | 8 | 0.837429332590447 |
| chr5-23988382-23988883 | 8 | 0.98685210623688 |
| chr4-6411112-6411613 | 6 | 0.991517273792013 |
| chr4-36416250-36416751 | 7 | 0.903284845590226 |
| chr1-107071960-107072461 | 3 | 0.995770209791711 |
| chr10-10090117-10090618 | 8 | 0.989049741835318 |
| chr19-41509215-41509716 | 2 | 0.957671378769363 |
| chr2-176270918-176271419 | 8 | 0.872147739352027 |
| chr16-69793727-69794228 | 5 | 0.865818215744656 |
| chr15-14532950-14533451 | 3 | 0.981343086156129 |
| chr14-2062964-2063465 | 4 | 0.973565278337902 |
| chr1-193186520-193187021 | 3 | 0.864724442390483 |
| chr16-79526752-79527253 | 4 | 0.901178606583709 |
| chr3-59828344-59828845 | 5 | 0.957648370836396 |
| chr1-30165560-30166061 | 4 | 0.935078488627716 |
| chr15-62113038-62113539 | 8 | 0.991318912026429 |
| chr6-56113222-56113723 | 2 | 0.874906246314391 |
| chr18-79850031-79850532 | 8 | 0.993493681128311 |
| chr5-84016008-84016509 | 8 | 0.822368637589494 |
| chr3-145085464-145085965 | 8 | 0.884802764608642 |
| chr10-72134481-72134982 | 4 | 0.936084725309355 |
| chr2-152747661-152748162 | 3 | 0.939160059783766 |
| chr6-14412171-14412672 | 8 | 0.980262965851332 |
| chr9-80450035-80450536 | 3 | 0.993039571464162 |
| chr8-13341553-13342054 | 5 | 0.916166842149405 |
| chr1-41988232-41988733 | 2 | 0.815792899919994 |
| chr15-21874726-21875227 | 7 | 0.853905380004021 |
| chr14-8202252-8202753 | 5 | 0.990390054373089 |
| chr2-170489103-170489604 | 2 | 0.84883298531056 |
| chr12-30759563-30760064 | 6 | 0.984314220966776 |
| chr19-46335311-46335812 | 8 | 0.994229164154809 |

| | | |
|---|---|---|
| chr11-69853318-69853819 | 6 | 0.865227330791394 |
| chr8-93972201-93972702 | 3 | 0.991349271156984 |
| chr10-62407183-62407684 | 4 | 0.93456531713165 |
| chr10-14072990-14073491 | 8 | 0.967474723746857 |
| chr5-112059395-112059896 | 8 | 0.833032864129595 |
| chr11-76621979-76622480 | 8 | 0.864334386849007 |
| chrX-104593789-104594290 | 5 | 0.828483562986921 |
| chr14-40239465-40239966 | 8 | 0.899083167052521 |
| chr17-20524295-20524796 | 5 | 0.968364868343706 |
| chrX-65888406-65888907 | 2 | 0.816801062585024 |
| chr11-74446043-74446544 | 3 | 0.976724473128102 |
| chr9-111270001-111270502 | 8 | 0.981355892733916 |
| chr10-61990068-61990569 | 8 | 0.989663556472284 |
| chr14-96730581-96731082 | 8 | 0.961236311880311 |
| chr6-87199922-87200423 | 3 | 0.99593553545041 |
| chr10-78239970-78240471 | 3 | 0.99593553545041 |
| chr4-130536090-130536591 | 8 | 0.946518476395505 |
| chr1-153184862-153185363 | 8 | 0.992812928705883 |
| chr2-244234145-244234646 | 8 | 0.944802668133465 |
| chr1-61894967-61895468 | 3 | 0.996789008428348 |
| chr17-47218028-47218529 | 8 | 0.883177211804961 |
| chr17-27795282-27795783 | 8 | 0.925966689383416 |
| chr13-71327761-71328262 | 3 | 0.995764232198329 |
| chr7-39113930-39114431 | 8 | 0.994825939579589 |
| chr3-142666478-142666979 | 8 | 0.982115039832878 |
| chr3-109081340-109081841 | 5 | 0.867560060204978 |
| chr13-68567209-68567710 | 5 | 0.910180209794913 |
| chr7-71644862-71645363 | 5 | 0.830836852707844 |
| chr4-173702035-173702536 | 4 | 0.904917859988698 |
| chr16-51059527-51060028 | 6 | 0.986967258741252 |
| chr16-5020172-5020673 | 8 | 0.993856833500248 |
| chr9-6891233-6891734 | 2 | 0.926640185655787 |
| chr20-47423750-47424251 | 5 | 0.917800664321236 |
| chr12-24656318-24656819 | 8 | 0.891305749296109 |
| chr5-125920711-125921212 | 3 | 0.976570442636557 |
| chr13-14636808-14637309 | 5 | 0.958264081823656 |
| chr17-48712992-48713493 | 2 | 0.869664819313722 |
| chr6-71556894-71557395 | 8 | 0.980410134078293 |
| chr1-256015109-256015610 | 8 | 0.985828332594248 |
| chr4-157241443-157241944 | 8 | 0.973789943148092 |
| chr17-39557638-39558139 | 3 | 0.99593553545041 |
| chr18-62507569-62508070 | 5 | 0.983919978840557 |
| chr12-31058134-31058635 | 8 | 0.976565602508779 |

| | | |
|---|---|---|
| chr9-58511073-58511574 | 2 | 0.82143464866205 |
| chr17-16904614-16905115 | 3 | 0.99689847678666 |
| chr1-96718620-96719121 | 8 | 0.939563318459525 |
| chr13-8094471-8094972 | 3 | 0.9972466410393 |
| chr10-47467013-47467514 | 4 | 0.817309801300336 |
| chr6-96999782-97000283 | 4 | 0.906669472926994 |
| chr1-136093881-136094382 | 2 | 0.967593774755178 |
| chr18-81993621-81994122 | 3 | 0.946843917627425 |
| chr2-217129031-217129532 | 3 | 0.99324119889726 |
| chr17-63194438-63194939 | 6 | 0.983262171091657 |
| chr5-81065732-81066233 | 3 | 0.976228809677638 |
| chr2-120041515-120042016 | 8 | 0.963083699427182 |
| chr2-115846119-115846620 | 7 | 0.868467738964448 |
| chr6-52725211-52725712 | 3 | 0.977561054832901 |
| chr16-7961720-7962221 | 8 | 0.987827126020955 |
| chr13-91843482-91843983 | 8 | 0.97068283710734 |
| chr12-40725718-40726219 | 8 | 0.818159390712741 |
| chr18-20239195-20239696 | 8 | 0.993493681128311 |
| chr13-93067086-93067587 | 8 | 0.913116386302564 |
| chr12-23168745-23169246 | 8 | 0.986982370067312 |
| chr3-133115263-133115764 | 5 | 0.864689499278115 |
| chr2-33745127-33745628 | 8 | 0.938844703338789 |
| chr1-258517671-258518172 | 8 | 0.991175399513473 |
| chr9-64613946-64614447 | 5 | 0.86640700720251 |
| chr5-25236936-25237437 | 6 | 0.991206925481043 |
| chr18-46651930-46652431 | 8 | 0.989635608533839 |
| chr13-87604139-87604640 | 4 | 0.911367370482038 |
| chr9-61586762-61587263 | 8 | 0.919291682632691 |
| chr1-84059209-84059710 | 1 | 0.857288093840887 |
| chr18-41111948-41112449 | 5 | 0.920009020092496 |
| chr3-28729288-28729789 | 8 | 0.98726972440909 |
| chr2-239336491-239336992 | 4 | 0.880690320225164 |
| chr15-36239595-36240096 | 8 | 0.992141278506622 |
| chr20-13939111-13939612 | 3 | 0.888101751133585 |
| chr19-48969585-48970086 | 5 | 0.967088304101971 |
| chr11-37871865-37872366 | 3 | 0.995063142496304 |
| chr7-87772037-87772538 | 8 | 0.966512339961041 |
| chr20-50015492-50015993 | 2 | 0.922374112277084 |
| chr4-176638431-176638932 | 8 | 0.968017759144801 |
| chr5-13993905-13994406 | 5 | 0.928981639366681 |
| chr20-27188528-27189029 | 8 | 0.996143319895167 |
| chr7-108929259-108929760 | 8 | 0.929232945444794 |
| chr17-8312059-8312560 | 8 | 0.993493681128311 |

| | | |
|---|---|---|
| chr4-45163491-45163992 | 8 | 0.991145741847249 |
| chr1-191495139-191495640 | 8 | 0.827618422599563 |
| chr11-57832830-57833331 | 2 | 0.866665314264955 |
| chr7-73790065-73790566 | 8 | 0.860127807576719 |
| chr4-51032069-51032570 | 8 | 0.845667564755703 |
| chr2-108348452-108348953 | 3 | 0.9894807904912 |
| chr10-27545220-27545721 | 3 | 0.88253998422656 |
| chr3-144752198-144752699 | 8 | 0.988481805551286 |
| chr6-27384053-27384554 | 4 | 0.947026426089266 |
| chr8-101613527-101614028 | 4 | 0.891879037565868 |
| chr1-125660158-125660659 | 3 | 0.997116731242404 |
| chr5-25850206-25850707 | 5 | 0.975499943189572 |
| chr1-18189329-18189830 | 8 | 0.985028682635524 |
| chr6-97346296-97346797 | 3 | 0.99593553545041 |
| chr18-13963337-13963838 | 2 | 0.848179211737768 |
| chr19-1884869-1885370 | 8 | 0.970891178303595 |
| chr7-116932299-116932800 | 8 | 0.827007726582638 |
| chr20-48039216-48039717 | 8 | 0.993493681128311 |
| chr15-26744339-26744840 | 8 | 0.955628299277914 |
| chr17-22355294-22355795 | 5 | 0.991687644973901 |
| chr11-30293182-30293683 | 8 | 0.977687437065161 |
| chr1-7427433-7427934 | 8 | 0.865960072598242 |
| chr7-75941217-75941718 | 6 | 0.991206925481043 |
| chr5-80171600-80172101 | 3 | 0.99593553545041 |
| chr15-80234487-80234988 | 8 | 0.968472648376801 |
| chr19-45944914-45945415 | 8 | 0.959216048225032 |
| chr5-71483481-71483982 | 8 | 0.995248618730591 |
| chr1-216261348-216261849 | 8 | 0.995214911895729 |
| chr1-160575030-160575531 | 3 | 0.870892374250077 |
| chr18-68067784-68068285 | 8 | 0.970872748733962 |
| chr8-50678002-50678503 | 8 | 0.878515804695444 |
| chr2-228659005-228659506 | 8 | 0.987459390792809 |
| chr1-177828992-177829493 | 8 | 0.979646902994953 |
| chr19-40743946-40744447 | 8 | 0.987489701656936 |
| chr17-62874026-62874527 | 2 | 0.885905727605645 |
| chr3-59138352-59138853 | 4 | 0.968409060207112 |
| chr19-20766913-20767414 | 8 | 0.991780231276768 |
| chr10-36883076-36883577 | 3 | 0.998157532951077 |
| chr1-246746688-246747189 | 8 | 0.909899360881093 |
| chr1-102292949-102293450 | 3 | 0.991869996427527 |
| chr5-129641895-129642396 | 8 | 0.977516701133825 |
| chr5-151487004-151487505 | 2 | 0.817360579797996 |
| chr17-53973719-53974220 | 6 | 0.978355323023694 |

| | | |
|---|---|---|
| chr10-43140425-43140926 | 3 | 0.993978162918737 |
| chr19-55672473-55672974 | 5 | 0.817255666300359 |
| chr15-34963132-34963633 | 6 | 0.97500848566338 |
| chr8-70349567-70350068 | 8 | 0.928727872886835 |
| chr1-72837004-72837505 | 3 | 0.99593553545041 |
| chr17-40910140-40910641 | 8 | 0.974019444417135 |
| chr7-114618537-114619038 | 2 | 0.91217804290977 |
| chr11-39436924-39437425 | 6 | 0.993134475666426 |
| chr18-72394688-72395189 | 8 | 0.994986338912894 |
| chr15-55515062-55515563 | 8 | 0.981047929537273 |
| chr7-65315858-65316359 | 8 | 0.912422292559225 |
| chr2-59204649-59205150 | 5 | 0.929069135427944 |
| chr13-48061825-48062326 | 4 | 0.96861106901096 |
| chr6-38197970-38198471 | 8 | 0.821508053386974 |
| chr14-66963907-66964408 | 8 | 0.96959814864349 |
| chr4-75495752-75496253 | 3 | 0.996898464235776 |
| chr20-29494961-29495462 | 8 | 0.993493681128311 |
| chr6-102034205-102034706 | 2 | 0.84260170783877 |
| chr1-218902634-218903135 | 8 | 0.982704319924436 |
| chr2-23687259-23687760 | 6 | 0.924645144394535 |
| chr6-36861296-36861797 | 5 | 0.914399049400505 |
| chr2-102077315-102077816 | 8 | 0.953871630656732 |
| chr1-218241689-218242190 | 2 | 0.940262261220098 |
| chr3-72643626-72644127 | 3 | 0.985522894297961 |
| chr6-120949235-120949736 | 2 | 0.852284541932229 |
| chr13-42043515-42044016 | 3 | 0.996447766748777 |
| chr7-99996713-99997214 | 4 | 0.892160899584482 |
| chr4-117145862-117146363 | 8 | 0.972963243074343 |
| chr16-70130206-70130707 | 8 | 0.993493681128311 |
| chr16-51943791-51944292 | 8 | 0.936893404104736 |
| chr7-129367481-129367982 | 4 | 0.920374909084111 |
| chr3-95322052-95322553 | 2 | 0.897181075291074 |
| chrX-51481354-51481855 | 8 | 0.965988918037053 |
| chr6-129983736-129984237 | 8 | 0.838992273742566 |
| chr6-28903779-28904280 | 8 | 0.995208038573293 |
| chr9-112215289-112215790 | 5 | 0.948102945184286 |
| chr16-75956477-75956978 | 7 | 0.817833497741862 |
| chr14-66856291-66856792 | 8 | 0.986492162838503 |
| chr2-217938716-217939217 | 6 | 0.993417604363711 |
| chr18-36924091-36924592 | 4 | 0.879651044261949 |
| chr2-19775087-19775588 | 3 | 0.99593553545041 |
| chr18-14827822-14828323 | 3 | 0.996894879168387 |
| chr7-54731877-54732378 | 6 | 0.987196408579549 |

| | | |
|---|---|---|
| chr16-62494790-62495291 | 4 | 0.959823321317224 |
| chr6-75703420-75703921 | 6 | 0.986022827934464 |
| chr13-81740967-81741468 | 2 | 0.919128057862553 |
| chrX-67119545-67120046 | 8 | 0.945145348345315 |
| chr6-46601085-46601586 | 1 | 0.913336178837231 |
| chr6-46450006-46450507 | 8 | 0.957030541704252 |
| chr16-17360737-17361238 | 5 | 0.849014227813885 |
| chr19-20173885-20174386 | 6 | 0.861543450140253 |
| chr17-18872783-18873284 | 1 | 0.964595970883573 |
| chr19-54673649-54674150 | 8 | 0.979666126081061 |
| chr11-61347966-61348467 | 6 | 0.977641162155017 |
| chr3-70049166-70049667 | 2 | 0.903511455807782 |
| chrX-71128344-71128845 | 8 | 0.993493681128311 |
| chr3-133034646-133035147 | 6 | 0.980378293359617 |
| chr18-24594909-24595410 | 8 | 0.944160574258672 |
| chr20-12529215-12529716 | 6 | 0.947030135644942 |
| chr9-74739086-74739587 | 5 | 0.856566198781764 |
| chr2-62061894-62062395 | 8 | 0.864518662281099 |
| chr5-140445168-140445669 | 5 | 0.957680926036942 |
| chrX-115567750-115568251 | 5 | 0.818113101419983 |
| chr14-25167793-25168294 | 8 | 0.978010011021604 |
| chr7-23639688-23640189 | 8 | 0.953077806122738 |
| chr14-12378871-12379372 | 8 | 0.983053828719092 |
| chr4-40157728-40158229 | 3 | 0.926884877137249 |
| chr13-1004101-1004602 | 1 | 0.886397339840965 |
| chr1-183487164-183487665 | 5 | 0.990697560025163 |
| chr5-48212766-48213267 | 5 | 0.983919978840557 |
| chr4-59994854-59995355 | 8 | 0.94294245518538 |
| chr6-73237114-73237615 | 3 | 0.99489884251131 |
| chr13-68174685-68175186 | 4 | 0.84255385407156 |
| chr1-67426387-67426888 | 8 | 0.975455871678531 |
| chr2-223696066-223696567 | 6 | 0.975783597529751 |
| chr17-82538769-82539270 | 3 | 0.99593553545041 |
| chr2-111142405-111142906 | 8 | 0.938651425305976 |
| chr10-40909902-40910403 | 2 | 0.935408359896337 |
| chr17-66103179-66103680 | 4 | 0.971597056361243 |
| chr8-70377313-70377814 | 8 | 0.995210978082595 |
| chr1-117343785-117344286 | 8 | 0.912917908940124 |
| chr6-84077113-84077614 | 4 | 0.929505609166686 |
| chr2-211827091-211827592 | 8 | 0.988978426772579 |
| chr5-166081609-166082110 | 8 | 0.960779443209152 |
| chr18-12498770-12499271 | 8 | 0.908308382272893 |
| chr5-88214195-88214696 | 4 | 0.81960520133494 |

| | | |
|---|---|---|
| chr2-123110049-123110550 | 2 | 0.919647129561746 |
| chr13-86516283-86516784 | 8 | 0.818491686131178 |
| chr1-80424197-80424698 | 8 | 0.930385781259653 |
| chr13-104713881-104714382 | 8 | 0.993493681128311 |
| chr7-6446195-6446696 | 6 | 0.990177215955679 |
| chr1-1035218-1035719 | 2 | 0.903419799633278 |
| chr1-259119376-259119877 | 8 | 0.978435548767828 |
| chr5-113727657-113728158 | 8 | 0.994928650278706 |
| chr17-33767576-33768077 | 6 | 0.992845107306875 |
| chr10-92194935-92195436 | 8 | 0.850346765700317 |
| chr4-57422645-57423146 | 8 | 0.929076912315809 |
| chr10-43877615-43878116 | 8 | 0.842640529449309 |
| chr5-71721805-71722306 | 8 | 0.995249047383649 |
| chr1-221541431-221541932 | 8 | 0.955562291463887 |
| chr17-23603563-23604064 | 4 | 0.95399771337391 |
| chr14-75614644-75615145 | 5 | 0.983919978840557 |
| chr4-160157991-160158492 | 8 | 0.855079846114478 |
| chr11-8217914-8218415 | 8 | 0.993493681128311 |
| chrX-6504519-6505020 | 8 | 0.858684884018521 |
| chr7-31647324-31647825 | 8 | 0.854159885431753 |
| chr4-40883632-40884133 | 2 | 0.95493513272187 |
| chr4-12900768-12901269 | 3 | 0.99495098374011 |
| chr11-71802077-71802578 | 2 | 0.947578412802407 |
| chr7-77182871-77183372 | 6 | 0.978553202196809 |
| chr10-10179220-10179721 | 8 | 0.827616560012828 |
| chr7-868969-869470 | 8 | 0.852157521342669 |
| chr6-46879063-46879564 | 7 | 0.887798828650065 |
| chr13-91834755-91835256 | 1 | 0.864509768023234 |
| chr15-55409700-55410201 | 1 | 0.855438264121173 |
| chr12-36267741-36268242 | 8 | 0.937510061770309 |
| chr7-20446802-20447303 | 8 | 0.951807654427374 |
| chr15-48584977-48585478 | 5 | 0.894643771478801 |
| chr6-95834509-95835010 | 8 | 0.921232152676037 |
| chr18-72603985-72604486 | 2 | 0.889543264419882 |
| chr9-78598475-78598976 | 1 | 0.958011460885045 |
| chr15-48314191-48314692 | 8 | 0.948570756486619 |
| chr12-29286059-29286560 | 7 | 0.822616175946809 |
| chr4-70532883-70533384 | 8 | 0.922377149733083 |
| chr17-29522441-29522942 | 4 | 0.972570768863994 |
| chr10-61161779-61162280 | 3 | 0.945684761531117 |
| chr16-69778414-69778915 | 8 | 0.884453217964114 |
| chr1-2081169-2081670 | 7 | 0.901968053799556 |
| chrX-100439357-100439858 | 6 | 0.968513134588093 |

| | | |
|---|---|---|
| chr9-107366737-107367238 | 6 | 0.832998780795108 |
| chr3-53890294-53890795 | 5 | 0.986973448188932 |
| chr1-184976205-184976706 | 5 | 0.968594771986726 |
| chr5-146508104-146508605 | 3 | 0.930712924657789 |
| chr2-10403326-10403827 | 6 | 0.896041980355138 |
| chr10-18753241-18753742 | 4 | 0.925275895052288 |
| chr2-144226196-144226697 | 8 | 0.891114160514591 |
| chr15-32649928-32650429 | 4 | 0.955471065663851 |
| chr15-27383290-27383791 | 8 | 0.832704969649627 |
| chr2-33233334-33233835 | 8 | 0.993493681128311 |
| chr20-4086464-4086965 | 3 | 0.937545122128941 |
| chr1-191491042-191491543 | 8 | 0.953754916279439 |
| chr6-22563735-22564236 | 4 | 0.945714662618247 |
| chr4-161648559-161649060 | 8 | 0.989145470719426 |
| chr6-47313076-47313577 | 2 | 0.950036796433716 |
| chr7-85657399-85657900 | 2 | 0.86382796071967 |
| chr7-97706254-97706755 | 2 | 0.943133522508114 |
| chr19-46875679-46876180 | 5 | 0.961486449054818 |
| chr1-52543283-52543784 | 8 | 0.962903098860007 |
| chr1-91032837-91033338 | 5 | 0.991271298988764 |
| chr2-65646630-65647131 | 8 | 0.993493681128311 |
| chr11-46489905-46490406 | 3 | 0.995636584923328 |
| chr19-39543968-39544469 | 8 | 0.911345233796434 |
| chr2-238607779-238608280 | 8 | 0.854335277454328 |
| chr19-19443278-19443779 | 8 | 0.8958553762844 |
| chr10-51160983-51161484 | 6 | 0.978007245203924 |
| chr17-64069733-64070234 | 6 | 0.830804818990052 |
| chr19-43527870-43528371 | 8 | 0.988692783485726 |
| chr11-33437023-33437524 | 8 | 0.97744143618792 |
| chr5-6623846-6624347 | 5 | 0.984074992147598 |
| chr3-61496595-61497096 | 5 | 0.955391291782989 |
| chr5-59250087-59250588 | 8 | 0.839251314103432 |
| chr4-21535719-21536220 | 8 | 0.973429698243408 |
| chr13-64448116-64448617 | 8 | 0.96316351782653 |
| chr4-131445566-131446067 | 2 | 0.928093295642362 |
| chr6-53365145-53365646 | 7 | 0.890999730680586 |
| chr5-101561581-101562082 | 4 | 0.924887525459439 |
| chr5-58452382-58452883 | 8 | 0.934190580568623 |
| chr2-43728948-43729449 | 8 | 0.830481359068965 |
| chr13-101751544-101752045 | 6 | 0.942698451410164 |
| chr15-74325359-74325860 | 7 | 0.819131758966831 |
| chr3-113858028-113858529 | 8 | 0.956824022741411 |
| chr5-135253512-135254013 | 8 | 0.844475432477406 |

| | | |
|---|---|---|
| chr3-114169177-114169678 | 8 | 0.995068762889216 |
| chr1-246078377-246078878 | 8 | 0.837800881453215 |
| chr3-48125315-48125816 | 3 | 0.933985441399141 |
| chr17-5704424-5704925 | 2 | 0.957809952673182 |
| chr1-255685940-255686441 | 6 | 0.995390954290043 |
| chr3-64827562-64828063 | 8 | 0.993493681128311 |
| chr5-118626690-118627191 | 6 | 0.973961760297324 |
| chrX-76613996-76614497 | 3 | 0.993462382519535 |
| chr3-153351789-153352290 | 4 | 0.954531488970393 |
| chr4-172493129-172493630 | 8 | 0.993192319365391 |
| chr20-41596984-41597485 | 3 | 0.99593553545041 |
| chr9-40031750-40032251 | 5 | 0.932068070001305 |
| chr2-221183684-221184185 | 8 | 0.985256856369237 |
| chr1-91160632-91161133 | 8 | 0.93166621773701 |
| chr1-230734933-230735434 | 8 | 0.992373826810876 |
| chrX-63420191-63420692 | 8 | 0.862048316869685 |
| chr5-106609914-106610415 | 6 | 0.983490663130606 |
| chr9-84688755-84689256 | 1 | 0.885069759852533 |
| chr10-10022684-10023185 | 8 | 0.994934230377201 |
| chr11-3549965-3550466 | 3 | 0.996813811608627 |
| chrX-127378314-127378815 | 8 | 0.849814264695056 |
| chr19-48951601-48952102 | 5 | 0.852627155780455 |
| chr2-219067220-219067721 | 5 | 0.912059266852995 |
| chr1-164521075-164521576 | 8 | 0.879439879566283 |
| chr4-67597371-67597872 | 8 | 0.898095370714446 |
| chr8-98351156-98351657 | 8 | 0.94055508559287 |
| chr4-75973593-75974094 | 3 | 0.996933620104904 |
| chr12-22873774-22874275 | 8 | 0.854932250337385 |
| chrX-58076375-58076876 | 6 | 0.991206925481043 |
| chr12-31862145-31862646 | 4 | 0.882614188556833 |
| chr19-40655373-40655874 | 5 | 0.983356070312986 |
| chr1-220920970-220921471 | 7 | 0.8392847649807 |
| chr9-85970861-85971362 | 5 | 0.966887374767078 |
| chr13-77238160-77238661 | 8 | 0.955078733545923 |
| chr17-22632358-22632859 | 5 | 0.987156493582172 |
| chrX-13461517-13462018 | 5 | 0.976251952306158 |
| chr14-77838604-77839105 | 8 | 0.821115505873039 |
| chr7-44120678-44121179 | 3 | 0.997080009094888 |
| chr15-2080077-2080578 | 5 | 0.922312318200169 |
| chr11-73764101-73764602 | 3 | 0.99593553545041 |
| chr15-94657905-94658406 | 8 | 0.951348331200318 |
| chr17-31292568-31293069 | 8 | 0.961196862294352 |
| chrX-139542718-139543219 | 3 | 0.994376778907895 |

| | | |
|---|---|---|
| chr19-16877277-16877778 | 8 | 0.888881154227099 |
| chr11-27645127-27645628 | 4 | 0.864703109274162 |
| chr8-81682858-81683359 | 8 | 0.993493681128311 |
| chr11-35936600-35937101 | 3 | 0.984318599630584 |
| chr14-64792920-64793421 | 5 | 0.974863741799867 |
| chr1-44008048-44008549 | 8 | 0.993493681128311 |
| chr4-136412275-136412776 | 8 | 0.993493681128311 |
| chr14-94101694-94102195 | 2 | 0.863026333131835 |
| chr15-80128600-80129101 | 4 | 0.821855646335006 |
| chr9-109667082-109667583 | 2 | 0.911335688405461 |
| chr7-74830269-74830770 | 3 | 0.995129698747335 |
| chr10-96984724-96985225 | 8 | 0.984219468836641 |
| chr8-119483465-119483966 | 5 | 0.983919978840557 |
| chr13-55757406-55757907 | 8 | 0.969951036200719 |
| chr16-29538521-29539022 | 5 | 0.93967724971496 |
| chr8-33769422-33769923 | 8 | 0.868463071924816 |
| chr1-54198404-54198905 | 3 | 0.996446931262112 |
| chr9-38577449-38577950 | 8 | 0.916711699786175 |
| chr2-187581920-187582421 | 4 | 0.967782426332037 |
| chr12-10030741-10031242 | 8 | 0.994039969921281 |
| chr15-33889355-33889856 | 8 | 0.987107907143404 |
| chr1-218464445-218464946 | 4 | 0.837162837579627 |
| chr14-75680982-75681483 | 8 | 0.814995180967418 |
| chr1-194455815-194456316 | 6 | 0.994387035222024 |
| chr16-29534317-29534818 | 8 | 0.911215087031679 |
| chr6-6611781-6612282 | 8 | 0.818045535732102 |
| chrX-141885094-141885595 | 6 | 0.991206925481043 |
| chr18-32191660-32192161 | 8 | 0.984257708487394 |
| chr9-96599001-96599502 | 3 | 0.982097830742314 |
| chr16-72550374-72550875 | 3 | 0.997331805233606 |
| chr6-86993519-86994020 | 8 | 0.977839776155169 |
| chr18-3086445-3086946 | 4 | 0.925281182194083 |
| chr15-52372487-52372988 | 5 | 0.893797279647307 |
| chr7-40521211-40521712 | 8 | 0.864071332783019 |
| chr1-41991198-41991699 | 8 | 0.913069645125132 |
| chr1-201964379-201964880 | 8 | 0.989452833326451 |
| chr2-225666215-225666716 | 3 | 0.996836448549842 |
| chr16-23110737-23111238 | 8 | 0.948392016423249 |
| chr2-124001520-124002021 | 2 | 0.876542153572308 |
| chr8-9089785-9090286 | 6 | 0.947337570474416 |
| chr2-43867828-43868329 | 4 | 0.953241643618347 |
| chr3-128959997-128960498 | 8 | 0.993493681128311 |
| chr17-36102231-36102732 | 8 | 0.985843410569588 |

| | | |
|---|---|---|
| chr17-4107325-4107826 | 8 | 0.928247379049544 |
| chr4-141602014-141602515 | 1 | 0.879066446927316 |
| chr1-200887098-200887599 | 8 | 0.980724586026443 |
| chr18-77812626-77813127 | 5 | 0.971137631603887 |
| chr3-51766337-51766838 | 3 | 0.91908057329217 |
| chr6-57151799-57152300 | 8 | 0.839786305963326 |
| chr16-7272049-7272550 | 8 | 0.907562816228767 |
| chr20-39677281-39677782 | 3 | 0.996852868929874 |
| chr18-32768220-32768721 | 3 | 0.996447681127541 |
| chr3-165207891-165208392 | 8 | 0.977513278493746 |
| chr8-101145499-101146000 | 6 | 0.992565510414297 |
| chr14-40683939-40684440 | 5 | 0.989688475786812 |
| chr18-52046032-52046533 | 8 | 0.865838162386532 |
| chr1-185469114-185469615 | 8 | 0.987741986199935 |
| chr14-95182763-95183264 | 5 | 0.982236834607464 |
| chr4-37433572-37434073 | 2 | 0.951970845904124 |
| chr9-26812208-26812709 | 8 | 0.958310951288923 |
| chr5-66133676-66134177 | 3 | 0.946350526838321 |
| chr6-46442432-46442933 | 8 | 0.995238286029476 |
| chr7-84062505-84063006 | 8 | 0.988254037497999 |
| chr1-253186177-253186678 | 2 | 0.866203578736442 |
| chr1-15264820-15265321 | 5 | 0.978802885883043 |
| chr7-55674306-55674807 | 8 | 0.910332103674534 |
| chr16-9024452-9024953 | 7 | 0.96423588254163 |
| chr16-49421823-49422324 | 3 | 0.995292006098356 |
| chr19-18037962-18038463 | 8 | 0.905652262375405 |
| chr2-44526453-44526954 | 8 | 0.980511314261013 |
| chr20-39753496-39753997 | 6 | 0.882101177233993 |
| chr16-57987195-57987696 | 8 | 0.993493681128311 |
| chr16-49112808-49113309 | 3 | 0.989615380759939 |
| chr12-41995110-41995611 | 8 | 0.88130761980759 |
| chr15-69186882-69187383 | 2 | 0.940258847907091 |
| chr6-113448935-113449436 | 3 | 0.99593553545041 |
| chr15-73001182-73001683 | 3 | 0.966791820604251 |
| chr18-64116997-64117498 | 4 | 0.829644931861542 |
| chr2-214002778-214003279 | 5 | 0.941660882612777 |
| chr9-102725514-102726015 | 8 | 0.829729104964556 |
| chr20-30116211-30116712 | 4 | 0.901393920335357 |
| chr2-31566526-31567027 | 8 | 0.985553454029797 |
| chr5-24515933-24516434 | 8 | 0.857065260602078 |
| chr4-182386514-182387015 | 4 | 0.859656958283902 |
| chr1-163971132-163971633 | 8 | 0.852460728908231 |
| chr5-44363771-44364272 | 6 | 0.982581638245659 |

| | | |
|---|---|---|
| chr14-85842976-85843477 | 6 | 0.98155764236452 |
| chr3-103932150-103932651 | 8 | 0.947758859294089 |
| chr3-45306181-45306682 | 4 | 0.906975277918247 |
| chr20-33731064-33731565 | 3 | 0.91204268128248 |
| chr4-65020222-65020723 | 4 | 0.834055006388269 |
| chr10-69825865-69826366 | 6 | 0.91300359172963 |
| chr11-30004051-30004552 | 4 | 0.88339733350376 |
| chr2-61254657-61255158 | 5 | 0.875540375403325 |
| chr5-129804541-129805042 | 8 | 0.864450553153866 |
| chr2-17424094-17424595 | 3 | 0.99591682677401 |
| chr13-43545605-43546106 | 4 | 0.945329291488354 |
| chr20-28913360-28913861 | 5 | 0.955726821544548 |
| chr16-21764585-21765086 | 8 | 0.936672758590099 |
| chr4-97432570-97433071 | 3 | 0.99652012313321 |
| chr2-175035268-175035769 | 7 | 0.890985992319198 |
| chr16-38912377-38912878 | 8 | 0.992934028462259 |
| chr17-39759324-39759825 | 8 | 0.950693844901488 |
| chr11-74434855-74435356 | 4 | 0.86853442000281 |
| chr2-237417193-237417694 | 5 | 0.956706047727971 |
| chr1-161404843-161405344 | 3 | 0.99593553545041 |
| chr3-139568081-139568582 | 8 | 0.929696515987211 |
| chr12-46312660-46313161 | 8 | 0.994981269771822 |
| chr14-52784561-52785062 | 8 | 0.993493681128311 |
| chr17-19743962-19744463 | 8 | 0.98973490052286 |
| chr3-118892311-118892812 | 8 | 0.990317280745326 |
| chr9-74120772-74121273 | 8 | 0.982580824400392 |
| chr5-137443381-137443882 | 8 | 0.988304265214674 |
| chr11-82042362-82042863 | 5 | 0.97357619549471 |
| chr16-1916814-1917315 | 4 | 0.831183378711769 |
| chr11-64083850-64084351 | 8 | 0.867567568773811 |
| chr5-103947515-103948016 | 1 | 0.927984406226596 |
| chr15-14024799-14025300 | 8 | 0.836267570971862 |
| chr9-36109119-36109620 | 8 | 0.993493681128311 |
| chr4-141838678-141839179 | 8 | 0.89421952735716 |
| chr1-12475043-12475544 | 8 | 0.923794703359074 |
| chr9-27971181-27971682 | 8 | 0.993493681128311 |
| chr2-235885049-235885550 | 2 | 0.932531412971934 |
| chr16-7709671-7710172 | 8 | 0.988886562200025 |
| chr11-36712997-36713498 | 3 | 0.874428716404002 |
| chr2-205096606-205097107 | 8 | 0.818479319936104 |
| chr2-130441200-130441701 | 3 | 0.99593553545041 |
| chr5-149128045-149128546 | 5 | 0.9761417921985 |
| chr16-24155432-24155933 | 3 | 0.985953504394171 |

| | | |
|---|---|---|
| chr11-57577454-57577955 | 8 | 0.976098450979803 |
| chr16-46016723-46017224 | 4 | 0.922441930044437 |
| chr7-4204961-4205462 | 3 | 0.996886284397733 |
| chr4-50408064-50408565 | 8 | 0.837657410900763 |
| chr6-42089471-42089972 | 8 | 0.992183983120653 |
| chr20-9262579-9263080 | 8 | 0.954486126841276 |
| chr18-13804378-13804879 | 8 | 0.857047067749556 |
| chr14-74332660-74333161 | 8 | 0.987761459482756 |
| chr5-58588070-58588571 | 4 | 0.85916912954879 |
| chr12-38465183-38465684 | 4 | 0.959251585289082 |
| chr1-134586097-134586598 | 8 | 0.8645208354171 |
| chr17-52386208-52386709 | 4 | 0.821347515623078 |
| chr16-8454617-8455118 | 7 | 0.955396702222365 |
| chr14-65723382-65723883 | 4 | 0.919610220415759 |
| chrX-2317416-2317917 | 5 | 0.93518559806847 |
| chrX-57167274-57167775 | 3 | 0.99593553545041 |
| chr2-96675558-96676059 | 6 | 0.992053135346732 |
| chr2-189196383-189196884 | 2 | 0.9072804789803 |
| chr17-32533470-32533971 | 5 | 0.825556174292083 |
| chr15-69553590-69554091 | 8 | 0.984248712580069 |
| chr2-138487709-138488210 | 5 | 0.982965134454085 |
| chr7-64358875-64359376 | 3 | 0.988249065595898 |
| chr5-40158761-40159262 | 5 | 0.992710506156951 |
| chr5-30296484-30296985 | 3 | 0.992219167943066 |
| chr14-26366814-26367315 | 6 | 0.987939068261608 |
| chr8-73053567-73054068 | 8 | 0.906185895232714 |
| chr18-46924661-46925162 | 3 | 0.996350712481501 |
| chr11-10633638-10634139 | 8 | 0.962959972279498 |
| chr5-149670075-149670576 | 8 | 0.857969062496365 |
| chr18-77561388-77561889 | 8 | 0.995249046926399 |
| chr7-22100374-22100875 | 8 | 0.910147466144504 |
| chr18-73603146-73603647 | 8 | 0.917292742882452 |
| chr1-178979579-178980080 | 6 | 0.991206925481043 |
| chr5-48982015-48982516 | 3 | 0.994353086031747 |
| chr4-132748453-132748954 | 8 | 0.951427130165887 |
| chr18-30802067-30802568 | 5 | 0.975763288932635 |
| chr5-99394919-99395420 | 3 | 0.988561719389599 |
| chr7-45022116-45022617 | 3 | 0.997245636080483 |
| chr3-12146524-12147025 | 8 | 0.974695202566168 |
| chr9-26691544-26692045 | 5 | 0.979090798905672 |
| chr5-5516443-5516944 | 5 | 0.987139396972157 |
| chr5-120800888-120801389 | 6 | 0.963134055536708 |
| chr3-117347621-117348122 | 8 | 0.993493681128311 |

| | | |
|---|---|---|
| chr1-56355843-56356344 | 8 | 0.891256733430285 |
| chr6-132335488-132335989 | 8 | 0.907323378827231 |
| chr3-33266311-33266812 | 8 | 0.830433636317777 |
| chr14-60373924-60374425 | 3 | 0.99593553545041 |
| chr8-94619093-94619594 | 4 | 0.957475834360396 |
| chr1-127191983-127192484 | 3 | 0.996828733694509 |
| chr1-192909786-192910287 | 6 | 0.974552537415338 |
| chr6-109270536-109271037 | 3 | 0.871363532332789 |
| chrX-138823432-138823933 | 8 | 0.973942919113303 |
| chr14-35721492-35721993 | 8 | 0.971891665645531 |
| chr1-131683907-131684408 | 3 | 0.983737941286447 |
| chr3-83316702-83317203 | 5 | 0.915457331918206 |
| chr6-103972068-103972569 | 2 | 0.941069379534294 |
| chr7-24156324-24156825 | 3 | 0.988113520692144 |
| chr7-21273125-21273626 | 8 | 0.95974544596359 |
| chrX-69483178-69483679 | 7 | 0.829159863606048 |
| chr2-138508925-138509426 | 4 | 0.87513104193365 |
| chrX-97759791-97760292 | 8 | 0.987678403015074 |
| chr10-17021760-17022261 | 8 | 0.880471513128512 |
| chr10-7134767-7135268 | 7 | 0.916847506459611 |
| chrX-85185118-85185619 | 6 | 0.991206925481043 |
| chr8-119239231-119239732 | 3 | 0.996890294569399 |
| chr20-33420780-33421281 | 5 | 0.92520586598987 |
| chr15-33291505-33292006 | 8 | 0.909174354664312 |
| chr15-26748925-26749426 | 8 | 0.917764264368961 |
| chr4-45045229-45045730 | 8 | 0.915193247885769 |
| chr20-21818150-21818651 | 4 | 0.948870412143412 |
| chr3-48331093-48331594 | 6 | 0.967889998214576 |
| chr8-4328317-4328818 | 7 | 0.920422758672207 |
| chr12-43710637-43711138 | 8 | 0.889419256098898 |
| chr6-76175818-76176319 | 6 | 0.923782938546736 |
| chr1-193968530-193969031 | 8 | 0.994700006866399 |
| chr3-125233764-125234265 | 3 | 0.973645688063222 |
| chrX-132752945-132753446 | 8 | 0.936070280402866 |
| chr2-202026622-202027123 | 5 | 0.983919978840557 |
| chr2-137046763-137047264 | 6 | 0.988328570297545 |
| chr16-5191798-5192299 | 6 | 0.993077488252282 |
| chr2-127468963-127469464 | 3 | 0.99593553545041 |
| chr16-26711221-26711722 | 7 | 0.896683609063415 |
| chr8-122139185-122139686 | 5 | 0.949411643206533 |
| chr16-2718700-2719201 | 8 | 0.929715782699452 |
| chr12-743751-744252 | 7 | 0.820571120882416 |
| chr16-53198922-53199423 | 8 | 0.883899348194914 |

| | | |
|---|---|---|
| chr10-65006737-65007238 | 8 | 0.986657748536422 |
| chr17-38367454-38367955 | 4 | 0.947998701781981 |
| chr20-33889305-33889806 | 8 | 0.964652259820515 |
| chr13-5861192-5861693 | 8 | 0.978559397883091 |
| chr6-140561333-140561834 | 3 | 0.987152702643473 |
| chr16-31758391-31758892 | 3 | 0.99593553545041 |
| chr16-47152653-47153154 | 3 | 0.996713457637697 |
| chr2-237181289-237181790 | 2 | 0.904880642815262 |
| chr16-45039207-45039708 | 8 | 0.960709039000122 |
| chr19-48240159-48240660 | 3 | 0.837455767976305 |
| chr2-68842887-68843388 | 3 | 0.983012312452351 |
| chr11-82273726-82274227 | 7 | 0.926586945399917 |
| chr7-53151710-53152211 | 8 | 0.959017267106285 |
| chr6-45018961-45019462 | 8 | 0.837829945314989 |
| chr3-52558859-52559360 | 5 | 0.948993016099177 |
| chr11-6459550-6460051 | 6 | 0.991206925481043 |
| chr1-193710912-193711413 | 2 | 0.933515687605528 |
| chr3-87786431-87786932 | 5 | 0.959558314472832 |
| chr19-42391668-42392169 | 8 | 0.922576751906363 |
| chr3-82158363-82158864 | 3 | 0.951406641240291 |
| chr2-140105864-140106365 | 8 | 0.985031740853206 |
| chr4-137425843-137426344 | 3 | 0.996250261750741 |
| chr16-16292771-16293272 | 8 | 0.972838208668837 |
| chr18-22955328-22955829 | 4 | 0.96413433597075 |
| chr6-37556310-37556811 | 3 | 0.99593553545041 |
| chr8-27191634-27192135 | 8 | 0.932216726587886 |
| chr1-165052744-165053245 | 7 | 0.876908688216971 |
| chr3-57526974-57527475 | 8 | 0.907680994993659 |
| chr18-80023824-80024325 | 3 | 0.961265376274496 |
| chr8-99656237-99656738 | 8 | 0.977203496216434 |
| chr11-23850800-23851301 | 3 | 0.83643146821663 |
| chr6-124112210-124112711 | 4 | 0.822699046933944 |
| chr9-36537204-36537705 | 6 | 0.980891627286461 |
| chr3-102837163-102837664 | 8 | 0.99223637605502 |
| chr1-196882125-196882626 | 8 | 0.919043546899504 |
| chr5-58302927-58303428 | 8 | 0.907409020603374 |
| chr14-58303092-58303593 | 8 | 0.882223403423678 |
| chr10-46494055-46494556 | 8 | 0.873720399414952 |
| chr1-104852723-104853224 | 6 | 0.991206925481043 |
| chrX-133144218-133144719 | 6 | 0.992969138093695 |
| chr8-60847122-60847623 | 8 | 0.932152762729838 |
| chr12-11149771-11150272 | 8 | 0.955270028282941 |
| chr8-70321730-70322231 | 8 | 0.985604571545639 |

| | | |
|---|---|---|
| chr18-43882363-43882864 | 8 | 0.993493681128311 |
| chr3-140079724-140080225 | 6 | 0.988613855409543 |
| chr16-51287244-51287745 | 4 | 0.876639311040886 |
| chr17-23068348-23068849 | 4 | 0.876990073906887 |
| chr18-68927587-68928088 | 5 | 0.986281128285548 |
| chr15-18656013-18656514 | 8 | 0.975633491311248 |
| chr3-167642589-167643090 | 8 | 0.934853717078821 |
| chr15-28045069-28045570 | 8 | 0.819902827413338 |
| chrX-51911639-51912140 | 3 | 0.99686139990231 |
| chrX-11260504-11261005 | 8 | 0.873370410982571 |
| chr17-60110186-60110687 | 5 | 0.908066986186227 |
| chr11-30127331-30127832 | 2 | 0.903678084447719 |
| chr2-48270650-48271151 | 8 | 0.909206431615573 |
| chr2-83695199-83695700 | 8 | 0.820030771474953 |
| chr7-97535636-97536137 | 8 | 0.972898182987988 |
| chr10-92478959-92479460 | 8 | 0.851367546432571 |
| chr11-43478407-43478908 | 5 | 0.986740361238434 |
| chr5-118380630-118381131 | 3 | 0.990996834006645 |
| chr9-83616457-83616958 | 8 | 0.99466029816908 |
| chr10-104419335-104419836 | 5 | 0.995384387332191 |
| chr19-47302519-47303020 | 8 | 0.900992852551483 |
| chr17-1799785-1800286 | 8 | 0.964808387766829 |
| chr3-59037014-59037515 | 8 | 0.945216268824175 |
| chr3-106498864-106499365 | 8 | 0.876761387743116 |
| chr17-70424800-70425301 | 3 | 0.92328209738944 |
| chr13-91797281-91797782 | 3 | 0.996558941082292 |
| chr15-16873025-16873526 | 8 | 0.848793926751074 |
| chr5-160799735-160800236 | 4 | 0.938554017141219 |
| chr4-40356870-40357371 | 3 | 0.998287969792972 |
| chr16-17545216-17545717 | 5 | 0.961795060559604 |
| chr2-183794766-183795267 | 8 | 0.933481360388836 |
| chr4-107136848-107137349 | 3 | 0.920769067006032 |
| chr4-75058922-75059423 | 8 | 0.85593420615363 |
| chr12-40634455-40634956 | 5 | 0.954502768101915 |
| chr20-18239108-18239609 | 5 | 0.988292144100791 |
| chr4-36688207-36688708 | 8 | 0.928101243148491 |
| chr14-95658086-95658587 | 8 | 0.929370777772426 |
| chr5-124244729-124245230 | 8 | 0.983571143752193 |
| chr3-156612156-156612657 | 7 | 0.94396131339373 |
| chr15-40706719-40707220 | 6 | 0.99118790864789 |
| chr17-69942039-69942540 | 8 | 0.994941947722788 |
| chr3-163432065-163432566 | 3 | 0.930268532580402 |
| chr1-116403340-116403841 | 5 | 0.940295758069698 |

| | | |
|---|---|---|
| chr18-20032036-20032537 | 3 | 0.99593553545041 |
| chr6-37650428-37650929 | 5 | 0.983919978840557 |
| chr4-18525167-18525668 | 8 | 0.817616510077228 |
| chr4-58471970-58472471 | 8 | 0.993493681128311 |
| chr9-54848381-54848882 | 4 | 0.865636016348199 |
| chr20-12653163-12653664 | 8 | 0.994849419015346 |
| chr14-70949705-70950206 | 5 | 0.988694559775558 |
| chr6-79937447-79937948 | 8 | 0.984570081872741 |
| chr6-117952146-117952647 | 6 | 0.940814511396106 |
| chr1-203552973-203553474 | 8 | 0.896559244228572 |
| chr5-18768171-18768672 | 8 | 0.972379833010108 |
| chr8-110521190-110521691 | 8 | 0.956565360236512 |
| chr7-55065616-55066117 | 5 | 0.983006570365271 |
| chr4-101529153-101529654 | 3 | 0.99071621577461 |
| chr20-7816623-7817124 | 8 | 0.897347296672847 |
| chr20-9105808-9106309 | 8 | 0.862788953627554 |
| chr6-31887238-31887739 | 3 | 0.996832088469037 |
| chr5-120211951-120212452 | 1 | 0.934032671826641 |
| chr14-41771835-41772336 | 5 | 0.989028092883555 |
| chr4-178433528-178434029 | 5 | 0.916286492081471 |
| chr9-103720183-103720684 | 7 | 0.888377518781435 |
| chr16-76926872-76927373 | 6 | 0.970455307657824 |
| chr15-101477038-101477539 | 8 | 0.945938636982371 |
| chr7-110210792-110211293 | 2 | 0.841768116395068 |
| chr16-16870634-16871135 | 8 | 0.830278496191403 |
| chr1-118063735-118064236 | 4 | 0.953921075866967 |
| chr13-39069765-39070266 | 8 | 0.992352008640853 |
| chr2-129157669-129158170 | 4 | 0.930184503628412 |
| chr5-73697499-73698000 | 4 | 0.874281533601119 |
| chr1-247061787-247062288 | 8 | 0.914854745699449 |
| chr5-31240160-31240661 | 3 | 0.948003882463981 |
| chr14-31955762-31956263 | 8 | 0.860380678248283 |
| chr5-165005847-165006348 | 4 | 0.882753045562983 |
| chr14-81553118-81553619 | 5 | 0.990392300852183 |
| chr18-52984215-52984716 | 8 | 0.993493681128311 |
| chr9-25227016-25227517 | 3 | 0.99593553545041 |
| chr10-18496768-18497269 | 8 | 0.910842521074284 |
| chr15-86479518-86480019 | 3 | 0.913172919105263 |
| chr15-16877295-16877796 | 8 | 0.948123188011515 |
| chr11-75527158-75527659 | 5 | 0.859766291280272 |
| chr6-91602561-91603062 | 3 | 0.974846827294339 |
| chr8-96096915-96097416 | 8 | 0.941065079653588 |
| chr5-109613753-109614254 | 3 | 0.992687117976716 |

| | | |
|---|---|---|
| chr11-55163966-55164467 | 8 | 0.986389449615401 |
| chr4-95692406-95692907 | 5 | 0.991746238499823 |
| chr2-33603938-33604439 | 8 | 0.99479157491579 |
| chr5-32447188-32447689 | 8 | 0.927786704797029 |
| chr1-95135940-95136441 | 8 | 0.915304799139812 |
| chr9-33226718-33227219 | 5 | 0.947541204716694 |
| chr7-102035708-102036209 | 8 | 0.924426823976929 |
| chr3-156577032-156577533 | 8 | 0.857489072713023 |
| chr3-32208867-32209368 | 3 | 0.99593553545041 |
| chr8-20128779-20129280 | 4 | 0.959685877297921 |
| chr15-93394868-93395369 | 3 | 0.99593553545041 |
| chr18-52568622-52569123 | 5 | 0.862270508895369 |
| chr20-33928353-33928854 | 6 | 0.991206925481043 |
| chr1-185848764-185849265 | 6 | 0.907439133032986 |
| chrX-74243638-74244139 | 3 | 0.977652665599766 |
| chr3-50609258-50609759 | 8 | 0.94465372167849 |
| chr2-92416969-92417470 | 8 | 0.992068675006815 |
| chr1-27937448-27937949 | 8 | 0.873618526616671 |
| chr9-92519295-92519796 | 8 | 0.958096888593365 |
| chr13-45720520-45721021 | 8 | 0.921128607409675 |
| chr9-11111462-11111963 | 8 | 0.923777753133109 |
| chr6-138174414-138174915 | 8 | 0.97472935390112 |
| chr2-197182751-197183252 | 8 | 0.933884735285797 |
| chr2-149320780-149321281 | 8 | 0.994168124774611 |
| chr8-90699497-90699998 | 1 | 0.827338493603143 |
| chr7-120530465-120530966 | 8 | 0.933821987463447 |
| chr3-70262789-70263290 | 1 | 0.862125764217514 |
| chr2-59550596-59551097 | 5 | 0.99135775490503 |
| chr15-2892097-2892598 | 8 | 0.820044885392469 |
| chr2-132012300-132012801 | 8 | 0.993493681128311 |
| chr12-36095165-36095666 | 4 | 0.849553675467917 |
| chr14-21820855-21821356 | 7 | 0.822650863086994 |
| chr7-73043871-73044372 | 3 | 0.99812326093068 |
| chr1-87498295-87498796 | 8 | 0.973042706810397 |
| chr9-27733071-27733572 | 3 | 0.99593553545041 |
| chr1-52949311-52949812 | 5 | 0.984457400038231 |
| chr2-174908703-174909204 | 7 | 0.864780148548882 |
| chr8-6498075-6498576 | 4 | 0.948411139814452 |
| chr8-38731894-38732395 | 3 | 0.99593553545041 |
| chr18-25740125-25740626 | 2 | 0.91929247168285 |
| chr20-282271-282772 | 3 | 0.993773302509095 |
| chr14-61218482-61218983 | 6 | 0.991206925481043 |
| chr3-50189854-50190355 | 7 | 0.871976339093552 |

| | | |
|---|---|---|
| chr8-62552988-62553489 | 2 | 0.947354503987307 |
| chr1-8115079-8115580 | 5 | 0.966682458859881 |
| chr17-79460991-79461492 | 8 | 0.853520700576502 |
| chrX-72913990-72914491 | 5 | 0.93265255944256 |
| chr1-179378307-179378808 | 4 | 0.827478092034125 |
| chr1-6905900-6906401 | 8 | 0.909600836703377 |
| chr2-243763396-243763897 | 8 | 0.841285489400927 |
| chr3-151731575-151732076 | 7 | 0.852945979973116 |
| chr13-98308571-98309072 | 3 | 0.972420630567431 |
| chr16-1212088-1212589 | 4 | 0.881410825374023 |
| chr7-122711145-122711646 | 6 | 0.927931014369748 |
| chr2-41096128-41096629 | 8 | 0.879341396057688 |
| chr2-210775757-210776258 | 1 | 0.84921211198276 |
| chr2-14059824-14060325 | 3 | 0.873049648190567 |
| chr7-72869987-72870488 | 8 | 0.961013655672567 |
| chr3-147074046-147074547 | 8 | 0.993135360381209 |
| chr3-119919419-119919920 | 1 | 0.825763781196562 |
| chr5-121988504-121989005 | 8 | 0.877183363677082 |
| chr4-56047473-56047974 | 2 | 0.817547505425875 |
| chr18-36297401-36297902 | 3 | 0.989873469830769 |
| chr14-1523125-1523626 | 6 | 0.989536832523208 |
| chr10-41161796-41162297 | 8 | 0.993493681128311 |
| chr6-38513238-38513739 | 8 | 0.951414109852364 |
| chr1-77242212-77242713 | 8 | 0.938800713220375 |
| chr1-122021418-122021919 | 8 | 0.980967716668717 |
| chr6-23821272-23821773 | 8 | 0.993493681128311 |
| chr8-26157006-26157507 | 8 | 0.891856174078644 |
| chr6-59987751-59988252 | 6 | 0.977301018096882 |
| chr2-65029787-65030288 | 8 | 0.88568064075941 |
| chr2-236175661-236176162 | 8 | 0.823395124236805 |
| chr10-53245541-53246042 | 5 | 0.991567363026928 |
| chr5-110052317-110052818 | 5 | 0.981444095259985 |
| chr17-28470153-28470654 | 8 | 0.848211177021845 |
| chr14-35729831-35730332 | 4 | 0.867056972422413 |
| chr6-44510686-44511187 | 8 | 0.943225921882065 |
| chr16-50999155-50999656 | 8 | 0.995038115970088 |
| chr17-37276892-37277393 | 3 | 0.986852112243874 |
| chr14-32942857-32943358 | 8 | 0.920193406597798 |
| chr3-158332343-158332844 | 8 | 0.913535371289881 |
| chr19-53003952-53004453 | 8 | 0.831294320268914 |
| chr4-41794317-41794818 | 3 | 0.99593553545041 |
| chr3-14383358-14383859 | 8 | 0.993323811497312 |
| chr11-9866989-9867490 | 3 | 0.891225370564355 |

| | | |
|---|---|---|
| chr1-147796432-147796933 | 1 | 0.929659393199364 |
| chr2-46873266-46873767 | 6 | 0.990251214349038 |
| chr14-10432349-10432850 | 4 | 0.835401836860754 |
| chr1-26487000-26487501 | 6 | 0.983134436670358 |
| chr3-17232180-17232681 | 8 | 0.894396371126922 |
| chr6-25931674-25932175 | 5 | 0.988306536799975 |
| chr9-65407375-65407876 | 4 | 0.946679064329285 |
| chr8-50697393-50697894 | 8 | 0.842831250217405 |
| chr7-28636732-28637233 | 1 | 0.940336770196598 |
| chr8-15038321-15038822 | 3 | 0.996879128488781 |
| chr4-37856295-37856796 | 3 | 0.865800732331592 |
| chr11-77908715-77909216 | 3 | 0.996059911648099 |
| chr1-201938529-201939030 | 6 | 0.965697393173344 |
| chr6-43996651-43997152 | 7 | 0.921030723862081 |
| chr14-10622925-10623426 | 8 | 0.99513748592083 |
| chr20-24646356-24646857 | 2 | 0.88976975789241 |
| chr8-29911095-29911596 | 3 | 0.8273851050907 |
| chr5-119566481-119566982 | 8 | 0.874625606488514 |
| chr6-124059244-124059745 | 2 | 0.887254306891931 |
| chr5-99596282-99596783 | 3 | 0.996582925561868 |
| chr10-80439570-80440071 | 3 | 0.843400163192519 |
| chr2-169482713-169483214 | 5 | 0.895007056127164 |
| chr1-24167784-24168285 | 2 | 0.892286132843683 |
| chr1-189347137-189347638 | 4 | 0.971635246298473 |
| chr7-115150449-115150950 | 8 | 0.942168342688358 |
| chr19-46404475-46404976 | 6 | 0.89021619211323 |
| chr17-52881688-52882189 | 8 | 0.897674263424578 |
| chr5-138981719-138982220 | 8 | 0.836945400362171 |
| chr6-32141117-32141618 | 8 | 0.91800684280552 |
| chr15-44864217-44864718 | 8 | 0.944041210030376 |
| chr18-19046028-19046529 | 3 | 0.907381458078135 |
| chr5-165943538-165944039 | 4 | 0.901479077352047 |
| chr2-106940239-106940740 | 3 | 0.995671297566739 |
| chrX-103278485-103278986 | 4 | 0.967301363652442 |
| chr8-98880101-98880602 | 6 | 0.991206925481043 |
| chr20-33581097-33581598 | 8 | 0.993493681128311 |
| chr4-52284128-52284629 | 1 | 0.957078671249875 |
| chr19-18316028-18316529 | 5 | 0.981441526824141 |
| chr6-41992602-41993103 | 4 | 0.831863695436486 |
| chr3-8489690-8490191 | 8 | 0.930759825654741 |
| chr14-38616562-38617063 | 8 | 0.985890373236851 |
| chr3-89607560-89608061 | 3 | 0.997012207149577 |
| chr4-161156414-161156915 | 8 | 0.907528287015923 |

| | | |
|---|---|---|
| chr11-64853709-64854210 | 2 | 0.836046111582324 |
| chr15-31813034-31813535 | 8 | 0.896575446997129 |
| chrX-61014353-61014854 | 8 | 0.831620708864344 |
| chr19-5023621-5024122 | 8 | 0.940222669083681 |
| chr9-100905470-100905971 | 5 | 0.974733702029973 |
| chr15-1232312-1232813 | 1 | 0.843626846391155 |
| chr5-23404994-23405495 | 6 | 0.991206925481043 |
| chr6-30753336-30753837 | 2 | 0.818924342297744 |
| chr2-234585761-234586262 | 8 | 0.943646273605333 |
| chr8-10462091-10462592 | 8 | 0.904264633008113 |
| chr4-142190241-142190742 | 8 | 0.912417778803271 |
| chr17-16301674-16302175 | 2 | 0.928835553514974 |
| chr13-92276886-92277387 | 5 | 0.975653508132884 |
| chr9-57362874-57363375 | 8 | 0.94975657156437 |
| chr1-49372184-49372685 | 2 | 0.899100754227071 |
| chr16-7930863-7931364 | 8 | 0.953493004827547 |
| chr20-11956831-11957332 | 8 | 0.995070484238847 |
| chr11-44198544-44199045 | 7 | 0.865218627973302 |
| chr7-33863569-33864070 | 8 | 0.874816375427602 |
| chr7-129386376-129386877 | 8 | 0.993493681128311 |
| chrX-63164944-63165445 | 8 | 0.987914076026865 |
| chr11-48139589-48140090 | 1 | 0.862324326181048 |
| chr13-91476060-91476561 | 5 | 0.839365369210453 |
| chr2-248537519-248538020 | 8 | 0.995062397443016 |
| chr20-29917590-29918091 | 8 | 0.907346756401093 |
| chr7-126150277-126150778 | 7 | 0.895224000509615 |
| chr13-57993781-57994282 | 3 | 0.99593553545041 |
| chr4-96988594-96989095 | 6 | 0.993186017195353 |
| chr17-65270604-65271105 | 5 | 0.985213124671678 |
| chr3-22382530-22383031 | 8 | 0.940121095669279 |
| chr3-80287000-80287501 | 3 | 0.93278639518364 |
| chr14-99352927-99353428 | 8 | 0.9042446569522 |
| chr5-42033411-42033912 | 8 | 0.993493681128311 |
| chr7-120597862-120598363 | 3 | 0.948124360270946 |
| chr10-92846956-92847457 | 4 | 0.958994177005529 |
| chr5-44131118-44131619 | 8 | 0.910264370689343 |
| chr18-69318443-69318944 | 5 | 0.974369188322778 |
| chr6-67037168-67037669 | 8 | 0.987683450620483 |
| chr1-249392590-249393091 | 8 | 0.993493681128311 |
| chr5-124560262-124560763 | 8 | 0.947124572612691 |
| chrX-33711869-33712370 | 3 | 0.996557098311744 |
| chr1-191051249-191051750 | 4 | 0.962729937203316 |
| chr20-49112333-49112834 | 5 | 0.99183283586321 |

| | | |
|---|---|---|
| chr7-9900318-9900819 | 7 | 0.910403362076406 |
| chr2-169851986-169852487 | 4 | 0.968289247998047 |
| chr14-65236318-65236819 | 3 | 0.996845076535744 |
| chr2-246035348-246035849 | 6 | 0.970884415055294 |
| chr1-49377569-49378070 | 2 | 0.827053545101903 |
| chr2-52150126-52150627 | 4 | 0.938306341086592 |
| chr4-96392162-96392663 | 8 | 0.973034625114879 |
| chr2-248753323-248753824 | 6 | 0.989860842817247 |
| chr15-9847917-9848418 | 8 | 0.993957783263053 |
| chr14-102592195-102592696 | 4 | 0.967202988857924 |
| chr2-142639416-142639917 | 7 | 0.893716283398094 |
| chrX-30976339-30976840 | 8 | 0.990710103397398 |
| chr3-87940807-87941308 | 2 | 0.926728270466012 |
| chr3-142367693-142368194 | 3 | 0.996816168302729 |
| chr3-150246084-150246585 | 4 | 0.936681258458994 |
| chr4-71587375-71587876 | 4 | 0.92062926888985 |
| chr9-61656058-61656559 | 8 | 0.94651296239586 |
| chr13-98558187-98558688 | 8 | 0.994852890380179 |
| chr15-75110593-75111094 | 8 | 0.964699858520075 |
| chr7-23143450-23143951 | 8 | 0.857244367900522 |
| chr4-8020279-8020780 | 4 | 0.960882347558027 |
| chr5-128669522-128670023 | 3 | 0.993033690136579 |
| chr7-23507292-23507793 | 4 | 0.905774101435532 |
| chr7-55584793-55585294 | 4 | 0.972981878045407 |
| chr8-28672113-28672614 | 8 | 0.947348413880845 |
| chr15-7761568-7762069 | 6 | 0.920589285842757 |
| chr4-124919017-124919518 | 8 | 0.993493681128311 |
| chrX-61023511-61024012 | 2 | 0.899765987141031 |
| chr9-57889871-57890372 | 8 | 0.881094492441991 |
| chr12-27410021-27410522 | 8 | 0.959237388431107 |
| chr13-82644040-82644541 | 5 | 0.983919978840557 |
| chr3-95752640-95753141 | 7 | 0.852529426410657 |
| chr1-136584802-136585303 | 3 | 0.995346766097126 |
| chr13-54933298-54933799 | 3 | 0.992391923101182 |
| chr3-13913952-13914453 | 8 | 0.932263244623199 |
| chr12-28166377-28166878 | 8 | 0.880938224292855 |
| chr6-3775492-3775993 | 3 | 0.984501849301874 |
| chr9-76017788-76018289 | 8 | 0.875890113899788 |
| chr9-77063463-77063964 | 8 | 0.95693245620987 |
| chr5-120167872-120168373 | 8 | 0.906094370051112 |
| chr6-4995997-4996498 | 6 | 0.991206925481043 |
| chr7-11530175-11530676 | 5 | 0.9712641432793 |
| chr3-148166736-148167237 | 7 | 0.849400286662625 |

| | | |
|---|---|---|
| chr15-28992688-28993189 | 8 | 0.85412214470623 |
| chr7-72244769-72245270 | 5 | 0.991386003748959 |
| chr14-36860812-36861313 | 8 | 0.822643059144434 |
| chr1-228618155-228618656 | 8 | 0.993493681128311 |
| chr9-57664482-57664983 | 8 | 0.954405151463439 |
| chr18-82510632-82511133 | 3 | 0.99593553545041 |
| chr1-233761183-233761684 | 8 | 0.993558755669822 |
| chr16-61378803-61379304 | 2 | 0.948432942394546 |
| chr9-61984311-61984812 | 8 | 0.904382029032533 |
| chr17-68307867-68308368 | 6 | 0.977451795993614 |
| chr2-46411959-46412460 | 8 | 0.933020180455825 |
| chr18-3131010-3131511 | 8 | 0.875735366394361 |
| chr18-14852046-14852547 | 6 | 0.993223569601462 |
| chr10-3854005-3854506 | 3 | 0.998468550406401 |
| chr5-143379224-143379725 | 8 | 0.916913325889271 |
| chr15-14602378-14602879 | 3 | 0.961842589012692 |
| chr7-25067691-25068192 | 4 | 0.965277877670153 |
| chr20-41376689-41377190 | 5 | 0.965762081098111 |
| chr4-136951922-136952423 | 8 | 0.993493681128311 |
| chr4-145598374-145598875 | 8 | 0.889917595519454 |
| chr17-73070292-73070793 | 8 | 0.973478379816946 |
| chr2-8293701-8294202 | 1 | 0.905784780670079 |
| chr19-418004-418505 | 4 | 0.91917731302154 |
| chr3-159271408-159271909 | 4 | 0.971069582511955 |
| chr15-31861193-31861694 | 5 | 0.8500016214722 |
| chr18-53439242-53439743 | 6 | 0.992778478075623 |
| chr8-78189575-78190076 | 5 | 0.900944225792618 |
| chr13-16072704-16073205 | 8 | 0.993493681128311 |
| chr14-26839006-26839507 | 8 | 0.992586967507829 |
| chr4-31902206-31902707 | 7 | 0.835578952564919 |
| chr1-97069608-97070109 | 8 | 0.913537026623137 |
| chrX-108648731-108649232 | 1 | 0.930135107774838 |
| chr9-67638780-67639281 | 8 | 0.961535777183622 |
| chr3-60142712-60143213 | 4 | 0.912426791831454 |
| chr4-53988238-53988739 | 6 | 0.965084911203491 |
| chr8-109163834-109164335 | 5 | 0.993942824800524 |
| chr17-10614027-10614528 | 8 | 0.99483731609714 |
| chr2-56985514-56986015 | 8 | 0.973080825282728 |
| chr19-46037334-46037835 | 2 | 0.930401363063004 |
| chr18-2889140-2889641 | 2 | 0.859702930798066 |
| chr6-53462449-53462950 | 3 | 0.996570425402405 |
| chr5-109889302-109889803 | 2 | 0.899528218700857 |
| chr10-70448523-70449024 | 8 | 0.824661538541119 |

| | | |
|---|---|---|
| chr2-110662057-110662558 | 1 | 0.862719969537063 |
| chr12-11001731-11002232 | 4 | 0.959266810665326 |
| chr4-168342439-168342940 | 8 | 0.953518629409929 |
| chr7-112312849-112313350 | 4 | 0.880070231254505 |
| chr17-14687236-14687737 | 6 | 0.956289647373373 |
| chrX-6326149-6326650 | 6 | 0.988467896547754 |
| chr16-2667130-2667631 | 3 | 0.958067226432586 |
| chr3-122360965-122361466 | 3 | 0.99723966174989 |
| chr18-34602318-34602819 | 5 | 0.982132848594148 |
| chr7-121744155-121744656 | 8 | 0.980881899661956 |
| chr9-26995791-26996292 | 3 | 0.997045719306647 |
| chr18-18183054-18183555 | 5 | 0.985693565569029 |
| chrX-41622310-41622811 | 3 | 0.831875376413066 |
| chr8-120613699-120614200 | 8 | 0.819356141424333 |
| chr6-122998524-122999025 | 8 | 0.894544027412774 |
| chr5-36302723-36303224 | 3 | 0.99593553545041 |
| chr14-83199430-83199931 | 5 | 0.995819187978543 |
| chr15-28251898-28252399 | 8 | 0.956858962328155 |
| chr3-71694097-71694598 | 6 | 0.990945227186767 |
| chr1-19478453-19478954 | 8 | 0.980271277052006 |
| chr6-35979328-35979829 | 7 | 0.81984705433914 |
| chr3-152539343-152539844 | 5 | 0.991274232117738 |
| chr9-61898694-61899195 | 4 | 0.858186249794642 |
| chr16-32703486-32703987 | 8 | 0.964697439574003 |
| chr1-16654416-16654917 | 3 | 0.996483875848218 |
| chr9-107044889-107045390 | 4 | 0.843114994162807 |
| chr14-9347699-9348200 | 8 | 0.970247885156693 |
| chr19-29553766-29554267 | 8 | 0.835247394639012 |
| chr16-58419429-58419930 | 8 | 0.929627273375737 |
| chr4-114890929-114891430 | 1 | 0.918456500186301 |
| chr10-30309049-30309550 | 8 | 0.89618802560849 |
| chr2-40468536-40469037 | 3 | 0.993590401049137 |
| chrX-106196604-106197105 | 3 | 0.99593553545041 |
| chr3-8202985-8203486 | 8 | 0.980787792746756 |
| chr4-116132491-116132992 | 8 | 0.986665540380364 |
| chr1-256634625-256635126 | 4 | 0.88430198396587 |
| chr18-1360517-1361018 | 3 | 0.99593553545041 |
| chr15-55147819-55148320 | 5 | 0.899973546930124 |
| chr9-100576327-100576828 | 3 | 0.98928504114303 |
| chr9-80034064-80034565 | 4 | 0.853965516473323 |
| chr15-90393316-90393817 | 3 | 0.90794720091631 |
| chr9-29995057-29995558 | 3 | 0.996898584393495 |
| chr15-75732193-75732694 | 8 | 0.994568292580791 |

| | | |
|---|---|---|
| chr1-109372443-109372944 | 8 | 0.99523425612128 |
| chr4-123336891-123337392 | 8 | 0.90688652054881 |
| chr7-127895249-127895750 | 7 | 0.930557063496324 |
| chr14-99766836-99767337 | 8 | 0.859958908718618 |
| chr8-57604853-57605354 | 8 | 0.895878552559062 |
| chr1-123959492-123959993 | 7 | 0.839218149660881 |
| chr2-189315550-189316051 | 7 | 0.835055616407827 |
| chr18-26574860-26575361 | 2 | 0.880338968889618 |
| chr17-83778626-83779127 | 8 | 0.973888231897496 |
| chr7-83722980-83723481 | 8 | 0.993493681128311 |
| chr8-110541741-110542242 | 8 | 0.989856928535332 |
| chr6-135523275-135523776 | 3 | 0.996603538744077 |
| chr7-28072302-28072803 | 7 | 0.94686395378111 |
| chr1-145895533-145896034 | 3 | 0.99593553545041 |
| chr20-9325355-9325856 | 8 | 0.96227202637307 |
| chr14-17693024-17693525 | 8 | 0.907055298517312 |
| chr1-118511232-118511733 | 8 | 0.875902393883901 |
| chr8-103215383-103215884 | 8 | 0.882783656723247 |
| chr18-73383391-73383892 | 3 | 0.996873013089019 |
| chrX-100218131-100218632 | 8 | 0.853470682314959 |
| chr13-67669286-67669787 | 1 | 0.939636092901232 |
| chr10-13292725-13293226 | 8 | 0.902009695138439 |
| chr17-60524764-60525265 | 5 | 0.98800549583856 |
| chr1-30431342-30431843 | 8 | 0.993493681128311 |
| chr17-34465826-34466327 | 8 | 0.994925890321631 |
| chr4-66914139-66914640 | 8 | 0.892942055260565 |
| chr4-332839-333340 | 8 | 0.991116312209962 |
| chr5-132125942-132126443 | 8 | 0.991727242554605 |
| chr1-133362276-133362777 | 7 | 0.941462003648907 |
| chr15-14091499-14092000 | 7 | 0.868310688892075 |
| chr4-48665929-48666430 | 3 | 0.847562592188738 |
| chr6-94865438-94865939 | 1 | 0.955731770675914 |
| chr1-158890843-158891344 | 2 | 0.939654412197519 |
| chr4-152392081-152392582 | 8 | 0.977673048404812 |
| chr18-69237933-69238434 | 8 | 0.948297832412326 |
| chr10-98854918-98855419 | 5 | 0.990956373734286 |
| chr2-79054517-79055018 | 3 | 0.996085638737147 |
| chr2-108191304-108191805 | 3 | 0.996993892205929 |
| chr5-130366923-130367424 | 8 | 0.962023227001702 |
| chr4-8799405-8799906 | 6 | 0.992272278999421 |
| chr10-23560372-23560873 | 3 | 0.99593553545041 |
| chr4-40889270-40889771 | 8 | 0.994395846760472 |
| chr5-55271378-55271879 | 4 | 0.911329502277348 |

| | | |
|---|---|---|
| chrX-104137287-104137788 | 8 | 0.898406406218939 |
| chr10-27619432-27619933 | 2 | 0.915156079232068 |
| chr2-44278565-44279066 | 8 | 0.884473031472681 |
| chr16-81408295-81408796 | 8 | 0.988781790840773 |
| chr2-53925417-53925918 | 6 | 0.993216492591907 |
| chr13-71345119-71345620 | 3 | 0.907780899514584 |
| chr5-76235227-76235728 | 4 | 0.841806813914404 |
| chr5-39229323-39229824 | 6 | 0.991206925481043 |
| chr1-163798456-163798957 | 5 | 0.989625515817674 |
| chr4-40996510-40997011 | 1 | 0.887726727854531 |
| chr1-199198968-199199469 | 8 | 0.916849264086704 |
| chr2-234227349-234227850 | 8 | 0.983206462490764 |
| chr14-73187579-73188080 | 8 | 0.983088373313419 |
| chr1-176427361-176427862 | 2 | 0.942004700585397 |
| chr1-13752462-13752963 | 8 | 0.993387941083305 |
| chr10-8645890-8646391 | 8 | 0.993493681128311 |
| chr2-111560064-111560565 | 8 | 0.911387299249904 |
| chr11-48045326-48045827 | 3 | 0.970154375355568 |
| chr4-665987-666488 | 5 | 0.965273960390411 |
| chr10-39740730-39741231 | 3 | 0.996758971898571 |
| chr13-30562540-30563041 | 2 | 0.898708709041996 |
| chr13-80450889-80451390 | 8 | 0.885157571677406 |
| chr2-166885719-166886220 | 5 | 0.929467594277804 |
| chr1-228883388-228883889 | 4 | 0.845913381671485 |
| chr18-46491929-46492430 | 4 | 0.890790542385716 |
| chr18-38382139-38382640 | 5 | 0.986037191802367 |
| chr19-40149949-40150450 | 3 | 0.994721376872301 |
| chr8-56508610-56509111 | 8 | 0.964523477390188 |
| chr5-121486969-121487470 | 8 | 0.993521598722674 |
| chr10-95525195-95525696 | 1 | 0.87026017495611 |
| chr19-42172744-42173245 | 5 | 0.975466605791725 |
| chr16-81872562-81873063 | 8 | 0.908765363325669 |
| chr2-27161660-27162161 | 8 | 0.994344551106111 |
| chr1-135435867-135436368 | 8 | 0.855648199586505 |
| chr2-63376753-63377254 | 5 | 0.983919978840557 |
| chr6-126232437-126232938 | 2 | 0.959890405669343 |
| chr3-7448159-7448660 | 7 | 0.940998325629476 |
| chr1-17650691-17651192 | 3 | 0.934478943114012 |
| chr6-25915294-25915795 | 5 | 0.981044807173912 |
| chr9-64412337-64412838 | 8 | 0.995243082743842 |
| chr2-229514358-229514859 | 3 | 0.995021991321587 |
| chr8-113212101-113212602 | 6 | 0.989659049581836 |
| chr13-105368810-105369311 | 8 | 0.857687310473071 |

| | | |
|---|---|---|
| chr11-63720642-63721143 | 8 | 0.973412487187412 |
| chr1-94789863-94790364 | 8 | 0.993493681128311 |
| chr1-125444638-125445139 | 3 | 0.973853241187425 |
| chr8-58311429-58311930 | 7 | 0.872971398723827 |
| chr20-9239529-9240030 | 8 | 0.959534520382016 |
| chr4-57597835-57598336 | 8 | 0.968625223379036 |
| chr6-25642283-25642784 | 8 | 0.816005838730365 |
| chr17-28424791-28425292 | 6 | 0.982835817109148 |
| chr2-227864414-227864915 | 4 | 0.971366887255015 |
| chr2-29464542-29465043 | 5 | 0.898079385682532 |
| chr18-74239446-74239947 | 8 | 0.992287162627034 |
| chr12-41756002-41756503 | 8 | 0.995191464593294 |
| chr11-75258346-75258847 | 8 | 0.955491236775218 |
| chr1-241666833-241667334 | 4 | 0.895323696906527 |
| chr4-8591149-8591650 | 3 | 0.996846785772066 |
| chrX-74235489-74235990 | 4 | 0.917385779797837 |
| chr13-68821230-68821731 | 8 | 0.978716060594304 |
| chr9-55186142-55186643 | 8 | 0.949582687575002 |
| chr4-182448593-182449094 | 8 | 0.973697745365607 |
| chr4-172348400-172348901 | 8 | 0.925539730664012 |
| chr6-89572232-89572733 | 8 | 0.956719883164114 |
| chr7-56717206-56717707 | 4 | 0.944603590564064 |
| chr9-62794296-62794797 | 8 | 0.875857564625789 |
| chr8-53500682-53501183 | 6 | 0.993228060127262 |
| chr3-130186152-130186653 | 6 | 0.893901967626943 |
| chr8-97785948-97786449 | 8 | 0.994998743308501 |
| chr6-129323283-129323784 | 3 | 0.99593553545041 |
| chr14-32192697-32193198 | 7 | 0.854181298251184 |
| chr9-107132334-107132835 | 5 | 0.829695457301044 |
| chr7-93850715-93851216 | 8 | 0.990789224892692 |
| chr18-12985192-12985693 | 8 | 0.986085648877162 |
| chrX-106768874-106769375 | 6 | 0.962991681410945 |
| chr7-55866870-55867371 | 4 | 0.818608786383613 |
| chr12-12708074-12708575 | 2 | 0.867493419880473 |
| chr2-116199669-116200170 | 8 | 0.835374080542393 |
| chr4-27625298-27625799 | 6 | 0.926035416947087 |
| chr14-36686157-36686658 | 4 | 0.921377671907729 |
| chr17-51095437-51095938 | 3 | 0.995020171768342 |
| chr14-103070834-103071335 | 6 | 0.991206925481043 |
| chr2-173345351-173345852 | 3 | 0.992275112442168 |
| chr5-88789291-88789792 | 8 | 0.993493681128311 |
| chr9-28797364-28797865 | 8 | 0.993842429268704 |
| chr12-17362234-17362735 | 3 | 0.996549222714583 |

| | | |
|---|---|---|
| chr4-130435910-130436411 | 4 | 0.881722632691262 |
| chr5-63050914-63051415 | 4 | 0.900077062413416 |
| chr3-89027873-89028374 | 4 | 0.957446573266253 |
| chr3-138194145-138194646 | 5 | 0.972733516574498 |
| chr15-96200079-96200580 | 3 | 0.990673220895094 |
| chr7-69447446-69447947 | 8 | 0.837340667646357 |
| chr1-183792216-183792717 | 8 | 0.973949257155128 |
| chr1-254070441-254070942 | 2 | 0.953674867796328 |
| chrX-127221480-127221981 | 7 | 0.831278302477515 |
| chr11-49843137-49843638 | 8 | 0.972714122041475 |
| chr5-164310538-164311039 | 4 | 0.882901594380879 |
| chr16-43489651-43490152 | 8 | 0.939701742201549 |
| chr14-32507552-32508053 | 8 | 0.983317518571253 |
| chrX-6080996-6081497 | 8 | 0.918382012266037 |
| chr17-83266938-83267439 | 8 | 0.977269534498618 |
| chr1-174727652-174728153 | 5 | 0.817240772914364 |
| chr4-45272112-45272613 | 6 | 0.98802157065871 |
| chr14-81191239-81191740 | 1 | 0.935241477456877 |
| chr6-23159808-23160309 | 8 | 0.900447702165612 |
| chr8-122102079-122102580 | 8 | 0.964752640462731 |
| chr15-54649400-54649901 | 5 | 0.983300118190738 |
| chr3-121831230-121831731 | 8 | 0.992711420023718 |
| chrX-93344799-93345300 | 8 | 0.980871546760012 |
| chr2-161270722-161271223 | 3 | 0.99593553545041 |
| chr5-114990778-114991279 | 2 | 0.911191304116915 |
| chr2-62506592-62507093 | 8 | 0.832535253764677 |
| chr4-7827464-7827965 | 6 | 0.892044506579894 |
| chr3-146604826-146605327 | 4 | 0.872334724698214 |
| chr9-50057947-50058448 | 4 | 0.870798725794437 |
| chr2-164757309-164757810 | 7 | 0.879347212165494 |
| chr1-29883701-29884202 | 8 | 0.950562676770425 |
| chr4-171517533-171518034 | 3 | 0.988320869752487 |
| chr15-39209774-39210275 | 8 | 0.883144101044291 |
| chr4-116043593-116044094 | 8 | 0.992159050448197 |
| chr10-81682095-81682596 | 4 | 0.9479820208836 |
| chr14-18886578-18887079 | 8 | 0.854552227969303 |
| chr4-137229056-137229557 | 3 | 0.995981962345413 |
| chr5-28244696-28245197 | 8 | 0.967219470238382 |
| chr2-74261621-74262122 | 3 | 0.984090568537341 |
| chr5-154066413-154066914 | 8 | 0.950780683154835 |
| chr5-139362139-139362640 | 8 | 0.911649786974527 |
| chr1-233948024-233948525 | 7 | 0.880783093884037 |
| chr1-218969753-218970254 | 3 | 0.996704124978949 |

| | | |
|---|---|---|
| chr8-64362264-64362765 | 5 | 0.962278288994962 |
| chr20-28361804-28362305 | 8 | 0.893440112143955 |
| chr8-112902119-112902620 | 8 | 0.993493681128311 |
| chr3-462672-463173 | 2 | 0.818198361922129 |
| chr4-8115750-8116251 | 7 | 0.856868798321041 |
| chr11-11397881-11398382 | 8 | 0.994702242365837 |
| chr1-17951564-17952065 | 2 | 0.954680074489412 |
| chrX-36843553-36844054 | 8 | 0.993493681128311 |
| chr18-46829305-46829806 | 8 | 0.925763728343558 |
| chr4-60778024-60778525 | 8 | 0.897623881426534 |
| chr1-258810849-258811350 | 5 | 0.930560737191212 |
| chr2-49022589-49023090 | 3 | 0.987605780652121 |
| chr1-238640870-238641371 | 6 | 0.991206925481043 |
| chr4-139667468-139667969 | 8 | 0.855616578992447 |
| chr14-66351761-66352262 | 3 | 0.986950726305105 |
| chr5-113768615-113769116 | 5 | 0.973595417655019 |
| chr2-127207447-127207948 | 3 | 0.99593553545041 |
| chr17-55576923-55577424 | 8 | 0.954524247954907 |
| chr7-16517034-16517535 | 8 | 0.934971843553907 |
| chr9-109324790-109325291 | 2 | 0.967927408520686 |
| chr4-116469603-116470104 | 8 | 0.951841916653787 |
| chr17-58648200-58648701 | 3 | 0.9474447657127 |
| chr4-113075027-113075528 | 6 | 0.991839734135621 |
| chr20-20222977-20223478 | 8 | 0.856412784083684 |
| chr5-97766454-97766955 | 1 | 0.921788062785506 |
| chr4-137529105-137529606 | 3 | 0.996797982253014 |
| chrX-62940189-62940690 | 8 | 0.993493681128311 |
| chr18-79625426-79625927 | 8 | 0.952210716876258 |
| chr3-33235091-33235592 | 5 | 0.978960731243984 |
| chr7-69310474-69310975 | 8 | 0.850791439177799 |
| chr2-196828630-196829131 | 5 | 0.933473252136487 |
| chr11-70452239-70452740 | 6 | 0.985528143709898 |
| chr2-160664589-160665090 | 3 | 0.99581967375161 |
| chr1-185994760-185995261 | 3 | 0.995787728384418 |
| chrX-76698238-76698739 | 8 | 0.993053663855348 |
| chr2-176534811-176535312 | 3 | 0.996330933439854 |
| chr7-70661648-70662149 | 3 | 0.959447848767355 |
| chr19-48694588-48695089 | 2 | 0.923647190052354 |
| chr16-62734707-62735208 | 8 | 0.99500160002863 |
| chr4-120696726-120697227 | 8 | 0.840406497178475 |
| chr9-49616532-49617033 | 8 | 0.993493681128311 |
| chr14-65374180-65374681 | 3 | 0.993734401448598 |
| chr8-61105113-61105614 | 8 | 0.852641424608294 |

| | | |
|---|---|---|
| chr19-22639451-22639952 | 3 | 0.996896406369936 |
| chr6-139304528-139305029 | 3 | 0.912782070371678 |
| chr10-53038078-53038579 | 2 | 0.856690629749499 |
| chr3-94711935-94712436 | 1 | 0.951370952196338 |
| chr4-76423351-76423852 | 8 | 0.978080705559932 |
| chr19-3233567-3234068 | 3 | 0.99593553545041 |
| chr4-44222976-44223477 | 8 | 0.871316794880588 |
| chr14-56488742-56489243 | 8 | 0.995192510596935 |
| chr13-87154731-87155232 | 6 | 0.985150328446356 |
| chrX-76504808-76505309 | 8 | 0.971424802107004 |
| chr7-86629459-86629960 | 8 | 0.832233643589069 |
| chr8-103584712-103585213 | 4 | 0.851170192124298 |
| chr3-52004694-52005195 | 8 | 0.984424567908841 |
| chr17-70590824-70591325 | 3 | 0.960647280563893 |
| chrX-91635316-91635817 | 3 | 0.99590488775407 |
| chr2-269218-269719 | 3 | 0.99593553545041 |
| chr6-113539761-113540262 | 5 | 0.955679184080932 |
| chr7-108189822-108190323 | 5 | 0.989043937713608 |
| chr3-115455133-115455634 | 7 | 0.843882071397689 |
| chr1-52780823-52781324 | 5 | 0.931219341719886 |
| chr3-19246208-19246709 | 4 | 0.860618012207556 |
| chr13-3728776-3729277 | 6 | 0.994479209278336 |
| chr1-47247881-47248382 | 8 | 0.993493681128311 |
| chrX-87706685-87707186 | 6 | 0.991206925481043 |
| chr18-3776844-3777345 | 4 | 0.912219362456975 |
| chr15-4402644-4403145 | 8 | 0.824331064267817 |
| chrX-106116195-106116696 | 5 | 0.875417409520004 |
| chr17-67305756-67306257 | 5 | 0.977994103792607 |
| chr1-138210675-138211176 | 5 | 0.986717455562197 |
| chr2-38523712-38524213 | 8 | 0.938537944778306 |
| chr5-67182013-67182514 | 6 | 0.965139868877505 |
| chr2-123444813-123445314 | 5 | 0.977678275827285 |
| chr10-37145386-37145887 | 8 | 0.930040994051165 |
| chr10-41784058-41784559 | 8 | 0.845852919384964 |
| chr19-27459510-27460011 | 8 | 0.988945542182518 |
| chr1-147767801-147768302 | 6 | 0.991206925481043 |
| chr5-152079352-152079853 | 8 | 0.957182606663301 |
| chr19-13524825-13525326 | 8 | 0.937516276918483 |
| chr13-10417618-10418119 | 6 | 0.990775010426069 |
| chr3-112038829-112039330 | 8 | 0.895188256227861 |
| chr15-52494068-52494569 | 8 | 0.988366754302472 |
| chr2-16158225-16158726 | 3 | 0.918819988693195 |
| chr3-7846282-7846783 | 5 | 0.976851329711539 |

| | | |
|---|---|---|
| chr1-211788641-211789142 | 5 | 0.993168618330394 |
| chr19-19189346-19189847 | 5 | 0.988601457238039 |
| chr7-72883117-72883618 | 8 | 0.936932127168841 |
| chr2-241250711-241251212 | 8 | 0.994764593531304 |
| chr4-1545430-1545931 | 2 | 0.861126021282148 |
| chr3-129692079-129692580 | 8 | 0.954928213140278 |
| chr20-45950811-45951312 | 4 | 0.961682083688532 |
| chr16-71739101-71739602 | 3 | 0.996872242279685 |
| chr13-92219731-92220232 | 8 | 0.973892762118541 |
| chr8-100810610-100811111 | 4 | 0.862102762242211 |
| chr9-86995242-86995743 | 8 | 0.872697692285297 |
| chr11-70463370-70463871 | 5 | 0.988438965552113 |
| chr2-9141051-9141552 | 3 | 0.997014687738812 |
| chr4-62013843-62014344 | 8 | 0.956991428266373 |
| chr9-56830183-56830684 | 3 | 0.996626588650549 |
| chr2-160864027-160864528 | 6 | 0.987082054949776 |
| chr10-63573132-63573633 | 5 | 0.981861924087719 |
| chr2-234193094-234193595 | 5 | 0.982261533226046 |
| chr10-88979857-88980358 | 6 | 0.992994286968496 |
| chr3-70360256-70360757 | 1 | 0.866837552518574 |
| chr10-8707632-8708133 | 8 | 0.844324114857505 |
| chr7-92814617-92815118 | 8 | 0.971142431291409 |
| chr1-243075750-243076251 | 8 | 0.983813584712179 |
| chr1-51034805-51035306 | 8 | 0.966307772132505 |
| chr5-40604113-40604614 | 8 | 0.989577547700898 |
| chr2-205118910-205119411 | 8 | 0.898568716459139 |
| chr4-50418698-50419199 | 7 | 0.928944418044159 |
| chr14-42724819-42725320 | 5 | 0.991472622361748 |
| chr9-96488441-96488942 | 6 | 0.991206925481043 |
| chr2-239876297-239876798 | 3 | 0.995935535450401 |
| chr2-75565808-75566309 | 4 | 0.925947956547502 |
| chr5-5993905-5994406 | 8 | 0.995136916353795 |
| chr18-77255702-77256203 | 1 | 0.922922009139987 |
| chr13-103079551-103080052 | 3 | 0.975285666548954 |
| chr14-8789617-8790118 | 8 | 0.823256952257127 |
| chr6-30797143-30797644 | 1 | 0.914351839549245 |
| chr2-160394235-160394736 | 8 | 0.904590544498333 |
| chr6-8484478-8484979 | 4 | 0.969714550646068 |
| chr1-225642769-225643270 | 3 | 0.969521059626422 |
| chr19-16000172-16000673 | 8 | 0.962426202887886 |
| chr2-23212210-23212711 | 3 | 0.855397566806865 |
| chr11-17749527-17750028 | 3 | 0.995518608953621 |
| chr18-19384399-19384900 | 3 | 0.99593553545041 |

| | | |
|---|---|---|
| chr6-68551206-68551707 | 5 | 0.974616286477247 |
| chr3-154822876-154823377 | 8 | 0.922089133556354 |
| chr9-63894579-63895080 | 4 | 0.971416362694901 |
| chr3-18714938-18715439 | 8 | 0.88155081196857 |
| chr12-46137228-46137729 | 4 | 0.957971578270433 |
| chr1-239416025-239416526 | 5 | 0.98761110047377 |
| chr4-76596260-76596761 | 7 | 0.938236843708486 |
| chr2-171228671-171229172 | 7 | 0.877041027627406 |
| chr18-31172651-31173152 | 1 | 0.958518391485836 |
| chr4-50406026-50406527 | 8 | 0.994787341165884 |
| chr20-33197849-33198350 | 2 | 0.876720508493085 |
| chr8-87876935-87877436 | 4 | 0.876701025767291 |
| chr1-176523310-176523811 | 1 | 0.893832539978687 |
| chr8-48542919-48543420 | 6 | 0.949467349474506 |
| chr15-80843699-80844200 | 5 | 0.982398899028894 |
| chr5-128385891-128386392 | 3 | 0.966471370564569 |
| chr5-87609274-87609775 | 8 | 0.961907851349537 |
| chr2-153596769-153597270 | 5 | 0.980194673079863 |
| chr3-142828682-142829183 | 4 | 0.913042735499624 |
| chr13-37526081-37526582 | 8 | 0.963340866290121 |
| chr7-109883412-109883913 | 3 | 0.916363014609393 |
| chr2-68670102-68670603 | 3 | 0.99593553545041 |
| chr11-49462569-49463070 | 6 | 0.991206925481043 |
| chr4-138393144-138393645 | 6 | 0.992948883949602 |
| chr9-77880127-77880628 | 6 | 0.97682691918039 |
| chr7-37359615-37360116 | 8 | 0.974822702240826 |
| chr15-80058494-80058995 | 8 | 0.994547679909024 |
| chr14-87506026-87506527 | 6 | 0.9932149252023 |
| chr2-42715922-42716423 | 3 | 0.99593553545041 |
| chr4-79257324-79257825 | 5 | 0.982514188994461 |
| chr6-100577360-100577861 | 8 | 0.914710455443835 |
| chr2-121489334-121489835 | 8 | 0.879601475012394 |
| chr5-164887328-164887829 | 8 | 0.957698141305502 |
| chr5-21439276-21439777 | 8 | 0.937803554526913 |
| chr3-142900433-142900934 | 8 | 0.993493681128311 |
| chr4-50915105-50915606 | 8 | 0.990701960035188 |
| chr15-6568501-6569002 | 8 | 0.975524718608634 |
| chr9-82823242-82823743 | 6 | 0.953682077185612 |
| chr5-136748023-136748524 | 8 | 0.989499127226998 |
| chr14-87141762-87142263 | 4 | 0.898126598787843 |
| chr15-49104468-49104969 | 1 | 0.838154092060727 |
| chr3-119964025-119964526 | 8 | 0.90002532380047 |
| chr11-84659330-84659831 | 8 | 0.866741671843338 |

| | | |
|---|---|---|
| chr6-59068601-59069102 | 3 | 0.995935535545041 |
| chr8-106504381-106504882 | 8 | 0.861708176159622 |
| chr2-187470563-187471064 | 4 | 0.832406422925491 |
| chr13-104678172-104678673 | 8 | 0.913709059052859 |
| chr6-123316435-123316936 | 3 | 0.984933768239185 |
| chr3-84772531-84773032 | 8 | 0.888528804244262 |
| chr4-60772155-60772656 | 8 | 0.883914035819017 |
| chr3-125538689-125539190 | 1 | 0.921541049496702 |
| chr17-80408157-80408658 | 2 | 0.878003191556254 |
| chr2-46841048-46841549 | 8 | 0.884858178422124 |
| chr6-27312744-27313245 | 8 | 0.829534392704206 |
| chr7-119308273-119308774 | 2 | 0.937711327916121 |
| chr7-99829944-99830445 | 8 | 0.993493681128311 |
| chr18-72531859-72532360 | 8 | 0.995158235921521 |
| chr18-37533679-37534180 | 6 | 0.988485077347201 |
| chr7-76124413-76124914 | 3 | 0.966243619860232 |
| chr8-80518368-80518869 | 4 | 0.825635388455306 |
| chr16-77849036-77849537 | 8 | 0.97842259651426 |
| chr17-31456857-31457358 | 3 | 0.993697283156874 |
| chr7-34265289-34265790 | 1 | 0.946683631813927 |
| chr17-941806-942307 | 8 | 0.989681699261413 |
| chr16-64977672-64978173 | 8 | 0.983716884291196 |
| chr18-3058911-3059412 | 7 | 0.885291387426277 |
| chr6-4987138-4987639 | 8 | 0.970924523727977 |
| chr2-245368480-245368981 | 8 | 0.93446185810257 |
| chr8-62816008-62816509 | 8 | 0.875489581407371 |
| chr5-58584714-58585215 | 6 | 0.991755770490446 |
| chr20-9258243-9258744 | 4 | 0.972430183924583 |
| chr5-147641980-147642481 | 6 | 0.988700006962043 |
| chr18-14669168-14669669 | 5 | 0.984089332499984 |
| chr16-9063016-9063517 | 5 | 0.986553284274067 |
| chr15-75653237-75653738 | 5 | 0.81850391967159 |
| chr6-96622345-96622846 | 8 | 0.991341455457533 |
| chrX-131100075-131100576 | 5 | 0.983919978840557 |
| chr18-71541077-71541578 | 1 | 0.914579379693939 |
| chr3-152011211-152011712 | 8 | 0.98711379098194 |
| chr11-56723217-56723718 | 8 | 0.944627739136313 |
| chr18-2963754-2964255 | 8 | 0.824576160345094 |
| chr6-71122599-71123100 | 5 | 0.931232358034891 |
| chrX-6433239-6433740 | 5 | 0.991235418787437 |
| chr5-22209841-22210342 | 5 | 0.976491106195689 |
| chr2-30636361-30636862 | 7 | 0.851864839231836 |
| chr2-91204805-91205306 | 3 | 0.994245463682062 |

| | | |
|---|---|---|
| chr8-30571046-30571547 | 8 | 0.990942041107542 |
| chr2-168672382-168672883 | 8 | 0.995061793398975 |
| chr7-34106308-34106809 | 5 | 0.915589857098158 |
| chr20-24722900-24723401 | 8 | 0.993493681128311 |
| chr14-83668307-83668808 | 2 | 0.902276682066368 |
| chr4-14827819-14828320 | 7 | 0.904192512041792 |
| chr1-24691199-24691700 | 5 | 0.972931739409943 |
| chr1-6842170-6842671 | 8 | 0.892671496089969 |
| chr3-79541322-79541823 | 8 | 0.966190491702261 |
| chr7-110691524-110692025 | 8 | 0.848994917350004 |
| chr3-56458870-56459371 | 5 | 0.890149373414634 |
| chr18-19003119-19003620 | 3 | 0.996893320525359 |
| chr16-19783192-19783693 | 6 | 0.991437086523326 |
| chr5-150212556-150213057 | 4 | 0.917947302832633 |
| chr4-166250081-166250582 | 8 | 0.914260973780166 |
| chr3-167737160-167737661 | 5 | 0.985589393361481 |
| chr2-219125720-219126221 | 8 | 0.910384086541631 |
| chr13-95099926-95100427 | 8 | 0.984293710972551 |
| chr4-158742194-158742695 | 8 | 0.932751520801772 |
| chr5-51398159-51398660 | 2 | 0.854899030078036 |
| chr20-45383339-45383840 | 8 | 0.876115883294308 |
| chr9-74059134-74059635 | 4 | 0.956997161273811 |
| chr5-20369127-20369628 | 8 | 0.868759506854057 |
| chr2-240510360-240510861 | 7 | 0.900063937452316 |
| chr5-48640548-48641049 | 3 | 0.99593553545041 |
| chrX-75728737-75729238 | 8 | 0.993493681128311 |
| chr4-64840374-64840875 | 8 | 0.842471569248162 |
| chrX-109197401-109197902 | 8 | 0.993493681128311 |
| chr2-164972011-164972512 | 8 | 0.860510230747712 |
| chr4-131736429-131736930 | 4 | 0.942587509306476 |
| chr9-74750956-74751457 | 8 | 0.973603358929422 |
| chr5-76862547-76863048 | 6 | 0.992258680603813 |
| chr11-46708765-46709266 | 3 | 0.992681051188163 |
| chr4-9799239-9799740 | 5 | 0.883183662053226 |
| chr18-75616353-75616854 | 3 | 0.99593553545041 |
| chr15-80691998-80692499 | 8 | 0.840484021817601 |
| chr5-8171258-8171759 | 3 | 0.853350551978454 |
| chr9-12197786-12198287 | 5 | 0.827518781436004 |
| chr7-53697983-53698484 | 5 | 0.986894174641957 |
| chr7-108966884-108967385 | 8 | 0.932413324757799 |
| chr13-39231299-39231800 | 5 | 0.992321633121354 |
| chrX-101698396-101698897 | 3 | 0.99593553545041 |
| chr18-14874142-14874643 | 8 | 0.977773557341562 |

| | | |
|---|---|---|
| chr15-55097618-55098119 | 8 | 0.889683877306931 |
| chr15-4747441-4747942 | 2 | 0.946896364033507 |
| chr7-16675357-16675858 | 3 | 0.817390020702953 |
| chr5-81954266-81954767 | 8 | 0.987882301922486 |
| chrX-131669776-131670277 | 8 | 0.993493681128311 |
| chr17-24585763-24586264 | 3 | 0.995940369389171 |
| chr10-29943198-29943699 | 8 | 0.852352290207739 |
| chr5-165253907-165254408 | 5 | 0.988801739121309 |
| chr2-106762164-106762665 | 3 | 0.995805906688777 |
| chr16-6938865-6939366 | 8 | 0.941306848476946 |
| chr12-6279291-6279792 | 3 | 0.996336175113523 |
| chr5-136109549-136110050 | 4 | 0.921693023896605 |
| chrX-75361856-75362357 | 8 | 0.956268773680007 |
| chr7-26411065-26411566 | 5 | 0.989319883559888 |
| chr3-88210986-88211487 | 8 | 0.982171971690688 |
| chr13-92034617-92035118 | 5 | 0.973646837457843 |
| chr3-165926032-165926533 | 8 | 0.938923222165628 |
| chr6-130409066-130409567 | 8 | 0.937353777037479 |
| chr7-46563967-46564468 | 8 | 0.898859279320276 |
| chr17-75232273-75232774 | 8 | 0.931696456669503 |
| chr17-23777058-23777559 | 5 | 0.935803098164629 |
| chr9-24025221-24025722 | 3 | 0.996822374675157 |
| chr16-30637211-30637712 | 8 | 0.967479048781636 |
| chr1-172663659-172664160 | 5 | 0.894580975735203 |
| chrX-120309396-120309897 | 8 | 0.933776116426281 |
| chr3-146651016-146651517 | 8 | 0.986269722901172 |
| chr15-65552938-65553439 | 3 | 0.99593553545041 |
| chr7-85515328-85515829 | 8 | 0.924052546325364 |
| chr6-49192348-49192849 | 8 | 0.988561835548205 |
| chrX-149327095-149327596 | 7 | 0.946486400437644 |
| chr2-97899222-97899723 | 8 | 0.983955163366997 |
| chr9-107573510-107574011 | 6 | 0.997319352216967 |
| chr7-122784929-122785430 | 8 | 0.995075046896985 |
| chr15-75295397-75295898 | 3 | 0.996064978920777 |
| chr4-60923920-60924421 | 8 | 0.860095937282123 |
| chr1-42787088-42787589 | 6 | 0.991456792944673 |
| chr6-129979121-129979622 | 4 | 0.948631450733147 |
| chr14-49725228-49725729 | 6 | 0.989677817593526 |
| chr1-227175823-227176324 | 6 | 0.830151183918733 |
| chr17-68891491-68891992 | 3 | 0.967946532991927 |
| chr10-19543170-19543671 | 8 | 0.995249014518916 |
| chr11-79309977-79310478 | 8 | 0.993493681128311 |
| chr6-51850868-51851369 | 8 | 0.993493681128311 |

| | | |
|---|---|---|
| chr3-37225873-37226374 | 8 | 0.853642495615997 |
| chr7-59042905-59043406 | 5 | 0.83120900352725 |
| chr17-26636062-26636563 | 8 | 0.994713153725055 |
| chr19-21441892-21442393 | 3 | 0.996425906002666 |
| chr5-120909933-120910434 | 4 | 0.966212499988072 |
| chr2-154379796-154380297 | 3 | 0.99593553545041 |
| chr6-72248035-72248536 | 8 | 0.922438499394997 |
| chr10-49275690-49276191 | 8 | 0.968983361003641 |
| chr9-106586868-106587369 | 8 | 0.824424323184817 |
| chr14-7641634-7642135 | 8 | 0.860489421334931 |
| chr2-196125340-196125841 | 8 | 0.956539516949044 |
| chr11-16503778-16504279 | 8 | 0.920522342042344 |
| chr1-151574128-151574629 | 8 | 0.939848095904729 |
| chr18-33620775-33621276 | 3 | 0.99593553545041 |
| chr13-27417970-27418471 | 3 | 0.99593553545041 |
| chr5-60797080-60797581 | 8 | 0.941646419982432 |
| chr18-8891243-8891744 | 3 | 0.99593553545041 |
| chr5-156757045-156757546 | 2 | 0.8342348641903 |
| chr1-44148373-44148874 | 8 | 0.982590933219926 |
| chr1-150692085-150692586 | 4 | 0.936745767591723 |
| chr1-229209371-229209872 | 2 | 0.857208975992551 |
| chr16-43327116-43327617 | 5 | 0.983919978840557 |
| chr2-137583741-137584242 | 8 | 0.95936080966853 |
| chr4-45416554-45417055 | 3 | 0.99593553545041 |
| chr17-69047617-69048118 | 1 | 0.936631624891429 |
| chr7-120582648-120583149 | 3 | 0.888125218405635 |
| chr8-81756035-81756536 | 3 | 0.996895493162019 |
| chr5-46858044-46858545 | 3 | 0.936017888732831 |
| chr8-94512341-94512842 | 3 | 0.996885085403934 |
| chr1-105528469-105528970 | 8 | 0.985287026793208 |
| chr14-17508684-17509185 | 4 | 0.944309247672055 |
| chr15-46121105-46121606 | 8 | 0.96552768191316 |
| chr7-117492926-117493427 | 8 | 0.981892373129336 |
| chr20-7810506-7811007 | 8 | 0.994884307313204 |
| chr2-52101275-52101776 | 5 | 0.98907434483445 |
| chr10-60909586-60910087 | 3 | 0.945089462321788 |
| chr9-69038640-69039141 | 8 | 0.991320881053123 |
| chr2-8983714-8984215 | 3 | 0.99593553545041 |
| chr9-39508137-39508638 | 1 | 0.956495717118878 |
| chr18-20126594-20127095 | 5 | 0.978709147832825 |
| chr17-81301821-81302322 | 8 | 0.958220089639129 |
| chr3-134039065-134039566 | 7 | 0.907737337247172 |
| chr9-23280862-23281363 | 4 | 0.95112834143443 |

| | | |
|---|---|---|
| chr13-80519127-80519628 | 4 | 0.874601127192119 |
| chr13-65368992-65369493 | 8 | 0.858774520831572 |
| chr6-32336066-32336567 | 3 | 0.99593553545041 |
| chr20-15019611-15020112 | 8 | 0.943887994581343 |
| chr14-3638877-3639378 | 6 | 0.991748139593147 |
| chr6-125824268-125824769 | 8 | 0.926753950431647 |
| chr2-237622959-237623460 | 5 | 0.988622669614866 |
| chr15-49460338-49460839 | 3 | 0.996014196425285 |
| chr18-53142429-53142930 | 8 | 0.98000169055757 |
| chr3-111944250-111944751 | 3 | 0.947558439926702 |
| chr1-119495116-119495617 | 8 | 0.995198613229139 |
| chr9-22537836-22538337 | 8 | 0.921419931846859 |
| chr1-94794485-94794986 | 8 | 0.972901163921151 |
| chr10-95300403-95300904 | 8 | 0.959846491736435 |
| chr3-149909640-149910141 | 6 | 0.995721308336353 |
| chr2-216075202-216075703 | 8 | 0.869972314268389 |
| chr1-191552542-191553043 | 2 | 0.950886175431604 |
| chr3-90310486-90310987 | 3 | 0.988045010629544 |
| chr19-42984767-42985268 | 7 | 0.901546275416583 |
| chr2-42254664-42255165 | 3 | 0.981430331696364 |
| chr10-65680856-65681357 | 4 | 0.820433649899093 |
| chr3-48313746-48314247 | 3 | 0.994618784095937 |
| chr5-68636129-68636630 | 8 | 0.99435207791589 |
| chr9-60608841-60609342 | 2 | 0.873932439947519 |
| chr8-96013830-96014331 | 5 | 0.957595831352022 |
| chr19-27805097-27805598 | 8 | 0.994965413078843 |
| chr10-74312378-74312879 | 8 | 0.995240854146794 |
| chr1-194638690-194639191 | 8 | 0.910438682847639 |
| chr5-36863266-36863767 | 3 | 0.991205122998265 |
| chr20-43278000-43278501 | 8 | 0.931036067104435 |
| chr7-69435901-69436402 | 2 | 0.89167558140636 |
| chr6-92522200-92522701 | 8 | 0.892728470133174 |
| chr15-45161863-45162364 | 4 | 0.960741375720067 |
| chr11-16167802-16168303 | 7 | 0.856959913613802 |
| chr17-80772822-80773323 | 8 | 0.986605274468109 |
| chr2-150900364-150900865 | 8 | 0.87875932227892 |
| chr11-11606423-11606924 | 8 | 0.984661239461665 |
| chr2-8577934-8578435 | 4 | 0.905868153759368 |
| chr20-27269015-27269516 | 4 | 0.870798079417683 |
| chr4-180606381-180606882 | 4 | 0.958897638763733 |
| chr10-62212159-62212660 | 8 | 0.99520841481028 |
| chr2-178059222-178059723 | 5 | 0.986840535690406 |
| chr6-129326796-129327297 | 8 | 0.993744699156913 |

| | | |
|---|---|---|
| chr10-31807143-31807644 | 8 | 0.979474424011055 |
| chr8-68094068-68094569 | 3 | 0.985389860540397 |
| chr5-70514054-70514555 | 4 | 0.932329869185558 |
| chr8-31557402-31557903 | 8 | 0.968631146066424 |
| chr4-71516347-71516848 | 5 | 0.949194018500679 |
| chr8-82023543-82024044 | 8 | 0.900104844598385 |
| chr2-188932380-188932881 | 6 | 0.912308890925084 |
| chr10-15326653-15327154 | 8 | 0.947366132426231 |
| chr4-16669245-16669746 | 3 | 0.996582583656151 |
| chr5-114100402-114100903 | 8 | 0.918737605723074 |
| chr18-14807748-14808249 | 8 | 0.859291827401276 |
| chr2-216595097-216595598 | 6 | 0.994264890016363 |
| chr3-62285715-62286216 | 4 | 0.950574368305137 |
| chr4-18071665-18072166 | 8 | 0.918422069615471 |
| chr5-129820697-129821198 | 8 | 0.933961493293406 |
| chr12-6410240-6410741 | 8 | 0.932785188239042 |
| chr7-84138660-84139161 | 3 | 0.996561448107095 |
| chr8-31008580-31009081 | 4 | 0.918710967085303 |
| chr8-470177-470678 | 6 | 0.989549107092264 |
| chr20-25153129-25153630 | 8 | 0.845595363887306 |
| chr3-133066364-133066865 | 4 | 0.860807809935453 |
| chr5-120933529-120934030 | 8 | 0.993493681128311 |
| chr8-33365655-33366156 | 3 | 0.986096633150763 |
| chr8-66463355-66463856 | 8 | 0.924094993639865 |
| chr10-59805638-59806139 | 8 | 0.891882187259861 |
| chr4-141071536-141072037 | 2 | 0.952089475690067 |
| chr5-113683143-113683644 | 8 | 0.990737778259704 |
| chr17-41905472-41905973 | 3 | 0.892705856830895 |
| chr17-7755749-7756250 | 8 | 0.843108545118346 |
| chr13-98952169-98952670 | 7 | 0.941846029694049 |
| chr16-44139704-44140205 | 6 | 0.993169590443575 |
| chr5-42013566-42014067 | 3 | 0.994881391600792 |
| chr3-63983736-63984237 | 6 | 0.993227437711852 |
| chr4-158746309-158746810 | 6 | 0.991206925481043 |
| chr4-51250341-51250842 | 7 | 0.822908836374693 |
| chr14-96581950-96582451 | 8 | 0.836109459757243 |
| chr3-45592267-45592768 | 5 | 0.930548539516035 |
| chr1-84843821-84844322 | 5 | 0.915957314736288 |
| chr3-15294772-15295273 | 8 | 0.930551980711137 |
| chr5-34409892-34410393 | 2 | 0.958203700325914 |
| chr2-144817081-144817582 | 5 | 0.957188611304062 |
| chr9-87062583-87063084 | 8 | 0.98512936095591 |
| chr13-61926084-61926585 | 6 | 0.921739676722055 |

| | | |
|---|---|---|
| chr19-13838364-13838865 | 8 | 0.989826452493315 |
| chr7-54834669-54835170 | 8 | 0.874057317749866 |
| chr5-38783906-38784407 | 4 | 0.888620969576488 |
| chr4-125481529-125482030 | 1 | 0.874547613632169 |
| chr16-38128382-38128883 | 8 | 0.966124783328968 |
| chr8-117460427-117460928 | 3 | 0.997219011198082 |
| chr7-126607995-126608496 | 8 | 0.984553526067743 |
| chrX-2911289-2911790 | 5 | 0.962522993302938 |
| chr4-5291759-5292260 | 3 | 0.973863134861298 |
| chr11-57065585-57066086 | 8 | 0.993493681128311 |
| chr7-65458799-65459300 | 3 | 0.990657439038619 |
| chr13-86709417-86709918 | 8 | 0.992085392281718 |
| chrX-97222693-97223194 | 8 | 0.972941941433696 |
| chr10-34395904-34396405 | 8 | 0.923660131146887 |
| chr9-4985440-4985941 | 6 | 0.991206925481043 |
| chr4-151448171-151448672 | 8 | 0.990952129805584 |
| chr2-48985623-48986124 | 3 | 0.996710661384169 |
| chr9-61420212-61420713 | 8 | 0.951252016122593 |
| chrX-127433891-127434392 | 8 | 0.98349736815108 |
| chr9-6728549-6729050 | 2 | 0.872221169219229 |
| chr10-45417339-45417840 | 5 | 0.941332101648967 |
| chr8-54349368-54349869 | 8 | 0.920345775235663 |
| chr10-92848470-92848971 | 8 | 0.870614416568684 |
| chr2-220433305-220433806 | 5 | 0.978588343221154 |
| chr18-74935658-74936159 | 7 | 0.859747046229397 |
| chr7-100564450-100564951 | 6 | 0.993219851235241 |
| chr1-133366328-133366829 | 8 | 0.990962524822391 |
| chr3-10656074-10656575 | 8 | 0.988896816682504 |
| chr4-58717261-58717762 | 5 | 0.9830228154037 |
| chr3-31449925-31450426 | 2 | 0.823680582214686 |
| chr16-16928204-16928705 | 7 | 0.896920911027687 |
| chr3-79204906-79205407 | 8 | 0.965890790369581 |
| chr2-35420523-35421024 | 4 | 0.864939726186971 |
| chr14-90378160-90378661 | 3 | 0.996404360434705 |
| chr9-64794337-64794838 | 6 | 0.951703907407353 |
| chr9-90616744-90617245 | 8 | 0.9676240521962 |
| chrX-7948598-7949099 | 6 | 0.855369076269661 |
| chr12-20613610-20614111 | 3 | 0.842409620188395 |
| chr3-9419170-9419671 | 3 | 0.819172466055717 |
| chr20-52555782-52556283 | 6 | 0.993186590612345 |
| chr20-26512900-26513401 | 4 | 0.936682060222668 |
| chr3-159234021-159234522 | 6 | 0.853625638562121 |
| chr10-51823126-51823627 | 8 | 0.974734371851606 |

| | | |
|---|---|---|
| chr19-49792230-49792731 | 8 | 0.900665715362744 |
| chr1-108881610-108882111 | 8 | 0.986569348106224 |
| chr14-5239724-5240225 | 8 | 0.882466294264595 |
| chr17-1807586-1808087 | 5 | 0.980660410830711 |
| chr11-29753675-29754176 | 1 | 0.918343243762121 |
| chr3-38067944-38068445 | 4 | 0.94540362383289 |
| chr17-16257337-16257838 | 5 | 0.847656476077975 |
| chr2-216610541-216611042 | 4 | 0.907546022910172 |
| chr10-849565-850066 | 8 | 0.977934994579873 |
| chr5-138311129-138311630 | 8 | 0.993493681128311 |
| chr9-6900527-6901028 | 2 | 0.936354955288992 |
| chr5-114177901-114178402 | 8 | 0.860853595730414 |
| chr9-52472362-52472863 | 6 | 0.974428938383992 |
| chr13-49832357-49832858 | 5 | 0.980982517426203 |
| chr15-30176473-30176974 | 4 | 0.838314908707136 |
| chr14-15530855-15531356 | 3 | 0.88181150164071 |
| chr13-40509639-40510140 | 6 | 0.971658515185822 |
| chr10-88973556-88974057 | 8 | 0.948393631446475 |
| chr1-7444129-7444630 | 7 | 0.903379433838071 |
| chr8-100109844-100110345 | 8 | 0.92483570646757 |
| chrX-16162930-16163431 | 1 | 0.842008896916345 |
| chr2-113536137-113536638 | 8 | 0.987207315278715 |
| chr7-71047838-71048339 | 4 | 0.88724233087618 |
| chr19-23720366-23720867 | 5 | 0.899513286266922 |
| chr1-211917641-211918142 | 8 | 0.993561696490213 |
| chr5-38413629-38414130 | 6 | 0.996810559284194 |
| chr13-86053110-86053611 | 8 | 0.845078859793127 |
| chr4-50019363-50019864 | 8 | 0.959716056736134 |
| chr1-230727137-230727638 | 8 | 0.966336449981107 |
| chr1-163164322-163164823 | 3 | 0.858804442949449 |
| chr14-35278411-35278912 | 5 | 0.823805007491717 |
| chr16-7865384-7865885 | 4 | 0.887620529518108 |
| chr13-35130701-35131202 | 6 | 0.991206925481043 |
| chr10-13078096-13078597 | 8 | 0.950417197425978 |
| chr9-108600172-108600673 | 2 | 0.873419540937245 |
| chr7-18349697-18350198 | 6 | 0.987243221970331 |
| chr2-34749337-34749838 | 8 | 0.995109432947253 |
| chr12-29683271-29683772 | 4 | 0.942542851162992 |
| chr3-94368010-94368511 | 8 | 0.986568708591862 |
| chr11-38285362-38285863 | 5 | 0.828577314808886 |
| chr15-62839085-62839586 | 6 | 0.980878891466006 |
| chr3-160249359-160249860 | 6 | 0.988585915272453 |
| chr3-160489516-160490017 | 4 | 0.884725449198033 |

| | | |
|---|---|---|
| chr1-66885784-66886285 | 8 | 0.951539135176184 |
| chr4-16245543-16246044 | 3 | 0.996754637034844 |
| chr4-174510272-174510773 | 4 | 0.828129002129763 |
| chr6-84919885-84920386 | 3 | 0.818343053281414 |
| chr8-118144928-118145429 | 8 | 0.995004916942041 |
| chr9-56679017-56679518 | 7 | 0.897240876885302 |
| chr8-68878048-68878549 | 3 | 0.91265613067734 |
| chr9-46399606-46400107 | 8 | 0.971247313148793 |
| chr12-22122703-22123204 | 4 | 0.918931071983287 |
| chr3-61721780-61722281 | 8 | 0.833727949734183 |
| chr8-25393590-25394091 | 4 | 0.912240575799603 |
| chr8-48153170-48153671 | 8 | 0.915774437211667 |
| chr14-12009644-12010145 | 8 | 0.993493681128311 |
| chr4-113847949-113848450 | 8 | 0.993493681128311 |
| chr2-135032966-135033467 | 3 | 0.98712551545429 |
| chr11-54075933-54076434 | 8 | 0.975678980794063 |
| chr15-66429888-66430389 | 3 | 0.993992844418441 |
| chr7-55730306-55730807 | 4 | 0.910792225143269 |
| chr6-128036052-128036553 | 4 | 0.927294542784928 |
| chr18-21841436-21841937 | 8 | 0.858453728179884 |
| chr2-107664330-107664831 | 4 | 0.917915732232889 |
| chr2-34083884-34084385 | 8 | 0.857270430434912 |
| chr8-105529894-105530395 | 3 | 0.99593553545041 |
| chr7-116825676-116826177 | 8 | 0.994862258166993 |
| chr5-10540747-10541248 | 3 | 0.995107963045324 |
| chr7-56121194-56121695 | 8 | 0.954435540501866 |
| chr16-74098199-74098700 | 1 | 0.970696962943314 |
| chr5-56599513-56600014 | 5 | 0.974228799489207 |
| chr6-126788422-126788923 | 8 | 0.995062000409868 |
| chr5-155051369-155051870 | 8 | 0.959974687803115 |
| chr4-169504232-169504733 | 8 | 0.857667971288732 |
| chr5-128597330-128597831 | 8 | 0.921449426663672 |
| chr3-157684476-157684977 | 8 | 0.964633619758494 |
| chr1-173830810-173831311 | 4 | 0.899750166847714 |
| chr1-81971217-81971718 | 3 | 0.994795683447943 |
| chr12-26275283-26275784 | 8 | 0.928115524934964 |
| chr2-224948571-224949072 | 8 | 0.987823355180285 |
| chr2-121048204-121048705 | 8 | 0.995135099473691 |
| chr15-46231809-46232310 | 8 | 0.894966154431436 |
| chr6-35394197-35394698 | 5 | 0.937721567886918 |
| chr7-53841008-53841509 | 5 | 0.848423549753323 |
| chr16-5714804-5715305 | 8 | 0.954930068221544 |
| chr1-48754407-48754908 | 5 | 0.866850227973538 |

| | | |
|---|---|---|
| chr3-160047062-160047563 | 8 | 0.856589858891121 |
| chr8-50685277-50685778 | 4 | 0.893933219826821 |
| chr20-39412557-39413058 | 3 | 0.996883841256492 |
| chr9-42957782-42958283 | 8 | 0.972288846974328 |
| chr8-101505286-101505787 | 5 | 0.965359213645361 |
| chr3-33558317-33558818 | 5 | 0.986183305903212 |
| chr1-80272868-80273369 | 8 | 0.945507257628011 |
| chr8-55259084-55259585 | 4 | 0.834810440916329 |
| chr9-37761388-37761889 | 2 | 0.933197865029348 |
| chr4-111033399-111033900 | 3 | 0.953715360344666 |
| chr6-17409028-17409529 | 8 | 0.903608 |
| chr6-26949014-26949515 | 4 | 0.92937976921884 |
| chr16-3366724-3367225 | 8 | 0.963644411708579 |
| chr1-76611313-76611814 | 4 | 0.857791867796436 |
| chr16-20981431-20981932 | 7 | 0.933699473757275 |
| chr8-64869739-64870240 | 5 | 0.850469290502785 |
| chr4-88546245-88546746 | 6 | 0.974886844956075 |
| chr1-257434003-257434504 | 2 | 0.893289890443875 |
| chr14-67130593-67131094 | 2 | 0.933928696137728 |
| chr4-70538951-70539452 | 5 | 0.980730148513128 |
| chr8-116432000-116432501 | 5 | 0.971822056128953 |
| chr19-23717214-23717715 | 8 | 0.861816968924304 |
| chr8-14671624-14672125 | 8 | 0.901501786069173 |
| chr2-70252691-70253192 | 6 | 0.991346351218995 |
| chr5-17274892-17275393 | 4 | 0.89488537683665 |
| chr17-75669750-75670251 | 6 | 0.969389401350555 |
| chr2-194434540-194435041 | 3 | 0.995856179685057 |
| chr2-155553620-155554121 | 3 | 0.992702870972927 |
| chr13-39486941-39487442 | 4 | 0.868055328333318 |
| chr4-181741331-181741832 | 8 | 0.824661672199084 |
| chr5-58556200-58556701 | 8 | 0.941485733710212 |
| chr11-17329758-17330259 | 3 | 0.99593553545041 |
| chr16-45310005-45310506 | 5 | 0.86588595565226 |
| chr4-147909443-147909944 | 8 | 0.977456794982888 |
| chr2-182286359-182286860 | 2 | 0.941183400321471 |
| chr14-32715061-32715562 | 4 | 0.973568512483877 |
| chr20-17869486-17869987 | 3 | 0.987727550236744 |
| chr12-38693473-38693974 | 6 | 0.991653192286296 |
| chr2-193335866-193336367 | 3 | 0.920894303876011 |
| chr17-48267447-48267948 | 3 | 0.996988744267277 |
| chr6-107778399-107778900 | 8 | 0.909560065207344 |
| chr11-28482951-28483452 | 6 | 0.991823967710858 |
| chr5-76007248-76007749 | 8 | 0.817695378145353 |

| | | |
|---|---|---|
| chr17-12254553-12255054 | 8 | 0.832282262335387 |
| chr6-114839266-114839767 | 6 | 0.915967174777835 |
| chr10-101915858-101916359 | 8 | 0.963310654314072 |
| chr10-101061826-101062327 | 5 | 0.954040581307327 |
| chr9-50926574-50927075 | 7 | 0.861381030876639 |
| chr8-78927899-78928400 | 8 | 0.992438512360104 |
| chr10-24112434-24112935 | 8 | 0.990704049612876 |
| chr4-117844272-117844773 | 8 | 0.982964626920624 |
| chr8-113313256-113313757 | 6 | 0.991206925481043 |
| chr12-39984953-39985454 | 8 | 0.973277667246845 |
| chr1-85272923-85273424 | 6 | 0.992640662137898 |
| chr20-53898644-53899145 | 8 | 0.920131568794754 |
| chr1-46353488-46353989 | 3 | 0.986645091166631 |
| chr11-71759168-71759669 | 4 | 0.924662136528964 |
| chr4-181729035-181729536 | 8 | 0.993002908402472 |
| chr10-95070530-95071031 | 2 | 0.954848374278672 |
| chr16-57554046-57554547 | 6 | 0.93179282375699 |
| chr1-192618701-192619202 | 6 | 0.982800560677682 |
| chr14-95808737-95809238 | 8 | 0.993493681128311 |
| chr2-82421489-82421990 | 5 | 0.973198163254947 |
| chr3-32526160-32526661 | 8 | 0.822384212841885 |
| chr5-157806874-157807375 | 8 | 0.875616613713317 |
| chr5-73230986-73231487 | 8 | 0.893367131383859 |
| chr5-9077594-9078095 | 8 | 0.93560363884242 |
| chr6-5901294-5901795 | 7 | 0.911398996891459 |
| chr3-125917103-125917604 | 4 | 0.823182916970737 |
| chr8-113541862-113542363 | 5 | 0.987406168969971 |
| chr2-219874740-219875241 | 8 | 0.978808132298265 |
| chr8-81203768-81204269 | 8 | 0.978782241101284 |
| chr6-60905133-60905634 | 5 | 0.88611895523954 |
| chr7-19220372-19220873 | 4 | 0.878746586508464 |
| chr5-25339490-25339991 | 8 | 0.933733912347363 |
| chr6-75902260-75902761 | 2 | 0.840631778015928 |
| chr10-21015759-21016260 | 3 | 0.994446964784179 |
| chr4-117596783-117597284 | 3 | 0.997212496557648 |
| chr9-14592461-14592962 | 4 | 0.900587315903231 |
| chr3-147656296-147656797 | 8 | 0.993493681128311 |
| chr17-401138-401639 | 8 | 0.848394096692362 |
| chr5-88851855-88852356 | 3 | 0.969934080605834 |
| chr17-23212446-23212947 | 5 | 0.983919978840557 |
| chr2-216640901-216641402 | 5 | 0.847877941743505 |
| chr2-58004043-58004544 | 8 | 0.901391138378052 |
| chr9-74067074-74067575 | 6 | 0.990797172824179 |

| | | |
|---|---|---|
| chr4-38932141-38932642 | 3 | 0.99593553545041 |
| chr19-11234168-11234669 | 8 | 0.885054420804962 |
| chr7-41544500-41545001 | 3 | 0.99593553545041 |
| chrX-51007667-51008168 | 8 | 0.995223925499951 |
| chr7-1439789-1440290 | 2 | 0.894617537092371 |
| chr20-5558721-5559222 | 5 | 0.933495171472348 |
| chr11-18636865-18637366 | 8 | 0.99521385704227 |
| chr16-58972897-58973398 | 1 | 0.944069264986161 |
| chr7-1436052-1436553 | 3 | 0.996162499101901 |
| chr20-14377699-14378200 | 5 | 0.847439066289574 |
| chr7-18351635-18352136 | 1 | 0.864713715025775 |
| chr15-52665764-52666265 | 8 | 0.985627679970468 |
| chr4-170350551-170351052 | 8 | 0.956546314886767 |
| chr4-75802288-75802789 | 3 | 0.994420029040988 |
| chr1-54057000-54057501 | 6 | 0.997400128096088 |
| chrX-115125923-115126424 | 6 | 0.991206925481043 |
| chr20-36280338-36280839 | 8 | 0.946256172951676 |
| chr5-45652924-45653425 | 6 | 0.99239233066717 |
| chrX-24062935-24063436 | 8 | 0.993493681128311 |
| chr2-140545509-140546010 | 8 | 0.932011244998653 |
| chr8-72298504-72299005 | 5 | 0.851780205211437 |
| chr9-69975227-69975728 | 6 | 0.991206925481043 |
| chr3-93863417-93863918 | 5 | 0.986856455106576 |
| chr5-113393281-113393782 | 8 | 0.820565872945144 |
| chr2-97763139-97763640 | 3 | 0.997203808671902 |
| chr16-29023380-29023881 | 8 | 0.881146263421387 |
| chr7-111962379-111962880 | 1 | 0.930013490096086 |
| chr15-12983509-12984010 | 1 | 0.866072320096968 |
| chrX-135968701-135969202 | 6 | 0.991206925481043 |
| chr3-62487153-62487654 | 8 | 0.89024414337556 |
| chr14-80241771-80242272 | 8 | 0.895059982991332 |
| chr3-47232276-47232777 | 8 | 0.977317575917835 |
| chr1-258745581-258746082 | 2 | 0.881415102653762 |
| chr15-92419502-92420003 | 6 | 0.992985838680219 |
| chr19-37422227-37422728 | 4 | 0.908905192132494 |
| chr15-27749127-27749628 | 3 | 0.987695857269292 |
| chr1-27015647-27016148 | 6 | 0.92426092802986 |
| chr2-184350462-184350963 | 4 | 0.947820605508243 |
| chr10-42390295-42390796 | 2 | 0.841629699248887 |
| chr16-77516718-77517219 | 8 | 0.983988318535908 |
| chr5-95937719-95938220 | 4 | 0.900320575695841 |
| chr11-49069793-49070294 | 8 | 0.913940962832438 |
| chr8-52096963-52097464 | 8 | 0.968360819025488 |

| | | |
|---|---|---|
| chr9-104445281-104445782 | 7 | 0.921843820843932 |
| chr1-169994129-169994630 | 8 | 0.958178073206301 |
| chr2-61947323-61947824 | 8 | 0.993219682705801 |
| chr13-92844527-92845028 | 8 | 0.971078959068041 |
| chr12-38940654-38941155 | 6 | 0.971239259758363 |
| chr8-53329662-53330163 | 5 | 0.983919978840557 |
| chr14-39702674-39703175 | 3 | 0.99593553545041 |
| chr7-117034452-117034953 | 5 | 0.949715395847158 |
| chr1-246923953-246924454 | 8 | 0.943085144944515 |
| chr3-86409417-86409918 | 6 | 0.967322369327143 |
| chr4-117641689-117642190 | 8 | 0.992711241309882 |
| chr1-236069487-236069988 | 8 | 0.952076232021579 |
| chr18-47181286-47181787 | 8 | 0.927783122632009 |
| chr1-29246527-29247028 | 8 | 0.947593265265567 |
| chr7-11602047-11602548 | 1 | 0.974833222356135 |
| chr19-27986140-27986641 | 8 | 0.888461023474419 |
| chr1-87020609-87021110 | 3 | 0.863290643525705 |
| chr9-83147266-83147767 | 3 | 0.996587947940411 |
| chr9-83992512-83993013 | 6 | 0.971115613618299 |
| chr17-16908380-16908881 | 3 | 0.996556862894973 |
| chr13-40738355-40738856 | 4 | 0.904670157255616 |
| chr5-70443384-70443885 | 6 | 0.926112195496701 |
| chr2-230025572-230026073 | 3 | 0.996299126317259 |
| chr20-41830089-41830590 | 8 | 0.957004953482125 |
| chr13-14046605-14047106 | 3 | 0.986687577737362 |
| chr2-31744935-31745436 | 8 | 0.94194605844551 |
| chr9-37297990-37298491 | 6 | 0.935653676567604 |
| chr13-40122316-40122817 | 4 | 0.952870310475293 |
| chr15-38238667-38239168 | 3 | 0.868110304263213 |
| chr1-191932142-191932643 | 7 | 0.960178955983443 |
| chr2-77612454-77612955 | 5 | 0.908416896233685 |
| chr7-57322011-57322512 | 8 | 0.957103572083359 |
| chr2-191978305-191978806 | 5 | 0.983919978840557 |
| chr4-23764539-23765040 | 8 | 0.993493681128311 |
| chr1-13631917-13632418 | 8 | 0.896009844710787 |
| chr9-15848919-15849420 | 8 | 0.993563815417924 |
| chr18-17264549-17265050 | 3 | 0.996563717944791 |
| chr15-88855264-88855765 | 6 | 0.990002416941259 |
| chr5-89541713-89542214 | 8 | 0.8638625563141 |
| chr9-73914965-73915466 | 5 | 0.988774934462398 |
| chr17-15375369-15375870 | 5 | 0.949731322541089 |
| chr20-28374294-28374795 | 4 | 0.918952841168889 |
| chr18-59896499-59897000 | 5 | 0.975660198936747 |

| | | |
|---|---|---|
| chr8-9483621-9484122 | 3 | 0.897719788721722 |
| chr9-18129635-18130136 | 8 | 0.822335369075541 |
| chr3-41653688-41654189 | 8 | 0.99480362787823 |
| chr3-153446235-153446736 | 2 | 0.952239430538301 |
| chrX-25443032-25443533 | 6 | 0.991206925481043 |
| chr6-103380527-103381028 | 4 | 0.821376235858774 |
| chr19-17675552-17676053 | 5 | 0.989384079820651 |
| chr14-74958917-74959418 | 5 | 0.992548623882893 |
| chr3-79556435-79556936 | 8 | 0.938991884225371 |
| chr2-155978913-155979414 | 3 | 0.997092666047395 |
| chr10-72031883-72032384 | 8 | 0.90683363624032 |
| chr8-118041828-118042329 | 8 | 0.862636013235678 |
| chr4-55580170-55580671 | 8 | 0.993505947242278 |
| chr3-167641343-167641844 | 8 | 0.829910485152531 |
| chr4-176708829-176709330 | 5 | 0.968944052280864 |
| chr14-59623712-59624213 | 6 | 0.988165694483295 |
| chr18-1721647-1722148 | 6 | 0.990872035133228 |
| chr6-126812691-126813192 | 8 | 0.995209218802043 |
| chr14-82221388-82221889 | 5 | 0.990265932535206 |
| chr5-95981033-95981534 | 8 | 0.886882508101612 |
| chr6-131726135-131726636 | 8 | 0.909830653682199 |
| chr9-98413387-98413888 | 5 | 0.980144937613499 |
| chr17-6594130-6594631 | 3 | 0.997582307797457 |
| chr11-156298-156799 | 3 | 0.996294532256117 |
| chr18-69211493-69211994 | 7 | 0.945283191968694 |
| chr10-57465751-57466252 | 3 | 0.997115058607073 |
| chr10-60213038-60213539 | 8 | 0.920744789919159 |
| chr18-68144395-68144896 | 3 | 0.97512529054131 |
| chr2-213626987-213627488 | 3 | 0.981354581339185 |
| chr6-42258733-42259234 | 3 | 0.992910750754354 |
| chr19-40977849-40978350 | 3 | 0.99593553545041 |
| chr1-122411837-122412338 | 3 | 0.818646723608094 |
| chr14-15158654-15159155 | 3 | 0.910718750940112 |
| chr10-85042402-85042903 | 8 | 0.928168065798025 |
| chr8-122339911-122340412 | 5 | 0.948052204499346 |
| chr15-33578701-33579202 | 8 | 0.945790109091607 |
| chr11-29558942-29559443 | 2 | 0.897829551496906 |
| chr5-68414133-68414634 | 5 | 0.891071472585887 |
| chr20-14554648-14555149 | 6 | 0.842970111685849 |
| chrX-116149909-116150410 | 5 | 0.985890780683572 |
| chr18-69363016-69363517 | 5 | 0.987280155071826 |
| chr19-41537496-41537997 | 8 | 0.995166627017976 |
| chr2-122463588-122464089 | 5 | 0.991199437024476 |

| | | |
|---|---|---|
| chrX-48088606-48089107 | 2 | 0.823341270415452 |
| chr7-101439371-101439872 | 8 | 0.835491938738669 |
| chr10-49420975-49421476 | 8 | 0.980203600187233 |
| chr12-15838353-15838854 | 5 | 0.989542612108696 |
| chr14-32579406-32579907 | 5 | 0.986894767664926 |
| chr9-99493816-99494317 | 1 | 0.827279920040983 |
| chr17-82782913-82783414 | 8 | 0.89266135816987 |
| chr4-126636973-126637474 | 8 | 0.976087655899192 |
| chr5-34456384-34456885 | 6 | 0.992931162021524 |
| chr7-121993896-121994397 | 6 | 0.971712287045292 |
| chr14-63807166-63807667 | 8 | 0.985506180374004 |
| chr1-229825252-229825753 | 3 | 0.992971784648435 |
| chr9-38592492-38592993 | 4 | 0.931044436824874 |
| chr14-36350463-36350964 | 3 | 0.99593553545041 |
| chr12-33256421-33256922 | 8 | 0.988841760888876 |
| chr7-70804509-70805010 | 1 | 0.912122779583489 |
| chr17-80523100-80523601 | 8 | 0.891621003143025 |
| chr11-12512940-12513441 | 8 | 0.994903905854888 |
| chr8-111774403-111774904 | 3 | 0.995226417337936 |
| chr8-96654094-96654595 | 8 | 0.993493681128311 |
| chr19-2230641-2231142 | 8 | 0.876791434327995 |
| chr7-30939302-30939803 | 8 | 0.876813747002002 |
| chr5-142494249-142494750 | 8 | 0.859967222207055 |
| chr7-25914445-25914946 | 4 | 0.96793081254683 |
| chr14-90713374-90713875 | 6 | 0.877278333421314 |
| chr14-25440519-25441020 | 8 | 0.985356582955736 |
| chr18-50853003-50853504 | 6 | 0.868783351391248 |
| chr13-67511073-67511574 | 4 | 0.912456509031842 |
| chr13-32518994-32519495 | 4 | 0.96380460362654 |
| chr4-119079915-119080416 | 8 | 0.88494201914427 |
| chr8-23830104-23830605 | 2 | 0.915325450069051 |
| chr2-109145982-109146483 | 6 | 0.863470224055576 |
| chr16-74490930-74491431 | 2 | 0.823165397906597 |
| chr8-101067342-101067843 | 8 | 0.985451650085245 |
| chr18-30881560-30882061 | 5 | 0.98135039862121 |
| chr15-74700866-74701367 | 1 | 0.92338572149006 |
| chr6-89192547-89193048 | 8 | 0.881437222793518 |
| chr5-28841166-28841667 | 8 | 0.971088708813011 |
| chr16-76295212-76295713 | 8 | 0.942436009540544 |
| chr10-71959605-71960106 | 8 | 0.992389360257183 |
| chr17-18442720-18443221 | 8 | 0.993012988059606 |
| chr1-50340049-50340550 | 8 | 0.825146128058305 |
| chr18-20445017-20445518 | 8 | 0.993493681128311 |

| | | |
|---|---|---|
| chr15-95590998-95591499 | 4 | 0.96260374140028 |
| chr11-25482041-25482542 | 8 | 0.955640896237864 |
| chr5-147894688-147895189 | 8 | 0.962012353470779 |
| chr8-31948569-31949070 | 5 | 0.891743318217388 |
| chr10-101705257-101705758 | 2 | 0.874440365324486 |
| chr19-5366018-5366519 | 5 | 0.969628521984828 |
| chrX-139149763-139150264 | 1 | 0.952479112746708 |
| chr9-38329441-38329942 | 8 | 0.994146412195164 |
| chr2-234360385-234360886 | 8 | 0.899919080877061 |
| chr4-108425714-108426215 | 8 | 0.986276464210746 |
| chr19-39085950-39086451 | 8 | 0.891788351499142 |
| chr8-22787818-22788319 | 3 | 0.99593553545041 |
| chr2-60257892-60258393 | 8 | 0.992267497790316 |
| chr19-46398985-46399486 | 6 | 0.900955930071232 |
| chr16-81243020-81243521 | 8 | 0.969851416783461 |
| chr1-212677663-212678164 | 5 | 0.880330633577739 |
| chr2-62295598-62296099 | 5 | 0.921317927235324 |
| chr16-46616341-46616842 | 2 | 0.889625279056076 |
| chr20-48641928-48642429 | 8 | 0.910963598594349 |
| chr2-81660740-81661241 | 6 | 0.938433758891243 |
| chr7-119896445-119896946 | 5 | 0.989533058816118 |
| chr17-23111631-23112132 | 3 | 0.896834541844614 |
| chr9-101481768-101482269 | 8 | 0.918702797069759 |
| chr1-211136551-211137052 | 8 | 0.981525922961126 |
| chr16-68998252-68998753 | 8 | 0.869662410898878 |
| chr10-32291541-32292042 | 3 | 0.99597113823886 |
| chr13-85233870-85234371 | 5 | 0.988861328822785 |
| chr11-61942951-61943452 | 8 | 0.995241923083766 |
| chr16-65576742-65577243 | 7 | 0.929087249276444 |
| chr4-104128238-104128739 | 5 | 0.902190217899163 |
| chr10-60838290-60838791 | 8 | 0.930513835107605 |
| chr13-76727285-76727786 | 5 | 0.971855872713335 |
| chr19-54789802-54790303 | 3 | 0.818386699140536 |
| chr20-22059834-22060335 | 8 | 0.954748813432689 |
| chr17-7294734-7295235 | 8 | 0.992218629437173 |
| chr7-87917189-87917690 | 8 | 0.890353061285274 |
| chr1-229181174-229181675 | 8 | 0.884232802341725 |
| chr17-26296576-26297077 | 8 | 0.928880591445882 |
| chr18-39084426-39084927 | 4 | 0.890715025694449 |
| chr7-32020148-32020649 | 8 | 0.982060998311782 |
| chr15-16424490-16424991 | 6 | 0.992802888490099 |
| chr8-49465256-49465757 | 6 | 0.967350814394991 |
| chr2-32704870-32705371 | 5 | 0.900833524230565 |

| | | |
|---|---|---|
| chr2-185124648-185125149 | 8 | 0.993154047980838 |
| chr11-58278441-58278942 | 2 | 0.872547186578272 |
| chr9-78383854-78384355 | 8 | 0.890022519483175 |
| chr4-12298945-12299446 | 8 | 0.993493681128311 |
| chr9-3975583-3976084 | 3 | 0.934890831746495 |
| chr14-89455005-89455506 | 3 | 0.994462238719683 |
| chr7-121322847-121323348 | 8 | 0.825916806072881 |
| chrX-25948133-25948634 | 4 | 0.859853657622523 |
| chr6-137357743-137358244 | 3 | 0.990771814913891 |
| chr15-919773-920274 | 2 | 0.932529329931921 |
| chr5-133035806-133036307 | 3 | 0.868235680250667 |
| chrX-90981934-90982435 | 5 | 0.983919978840557 |
| chr8-89758280-89758781 | 3 | 0.997229176268692 |
| chr14-43387568-43388069 | 5 | 0.987402731982019 |
| chr5-152571268-152571769 | 8 | 0.817824561305123 |
| chr2-81713867-81714368 | 8 | 0.990880867085562 |
| chr1-247570386-247570887 | 8 | 0.901445822273188 |
| chr5-91847917-91848418 | 8 | 0.840133620625257 |
| chr1-10927181-10927682 | 6 | 0.991617375205002 |
| chr8-86876685-86877186 | 8 | 0.995241494867986 |
| chr7-71034704-71035205 | 8 | 0.989685888622127 |
| chr19-53352-53853 | 8 | 0.993493681128311 |
| chr18-34072277-34072778 | 8 | 0.97706615644807 |
| chr8-108545972-108546473 | 4 | 0.936122668918972 |
| chr4-113972982-113973483 | 8 | 0.945261977492857 |
| chr2-32761991-32762492 | 4 | 0.915478552492443 |
| chr1-12412928-12413429 | 5 | 0.834541148682665 |
| chr2-177336487-177336988 | 3 | 0.990292791481434 |
| chr15-52263454-52263955 | 5 | 0.966402214454731 |
| chr15-10541972-10542473 | 4 | 0.855029201777402 |
| chr10-51920930-51921431 | 8 | 0.846849953730016 |
| chr1-216710912-216711413 | 8 | 0.995103103768179 |
| chr15-9854706-9855207 | 5 | 0.987528973718467 |
| chr20-19937185-19937686 | 8 | 0.889744159911299 |
| chr15-84821613-84822114 | 8 | 0.993396132828487 |
| chr1-56078893-56079394 | 4 | 0.874742351249795 |
| chr12-9761974-9762475 | 8 | 0.976586963384959 |
| chr8-4092362-4092863 | 8 | 0.933741418574814 |
| chr5-109134688-109135189 | 6 | 0.991206925481043 |
| chr11-12037828-12038329 | 1 | 0.883681750166278 |
| chr15-67327840-67328341 | 3 | 0.995676000926024 |
| chr18-21599422-21599923 | 7 | 0.829888457963689 |
| chrX-29431569-29432070 | 6 | 0.836550894201376 |

| | | |
|---|---|---|
| chr2-58434710-58435211 | 6 | 0.996370463454665 |
| chrX-14219931-14220432 | 6 | 0.835995144978443 |
| chr11-36756027-36756528 | 8 | 0.970228753286271 |
| chr6-67767305-67767806 | 6 | 0.991113611001646 |
| chr7-10973840-10974341 | 4 | 0.849255272502609 |
| chr5-137564718-137565219 | 5 | 0.946322734738876 |
| chr1-107084375-107084876 | 8 | 0.939064610400459 |
| chr3-139177455-139177956 | 4 | 0.885467410245488 |
| chr9-41255856-41256357 | 8 | 0.958778838393893 |
| chr19-11360264-11360765 | 8 | 0.965941251844296 |
| chr3-33284933-33285434 | 8 | 0.971353328914777 |
| chr7-40575590-40576091 | 6 | 0.993710406385503 |
| chr6-118561904-118562405 | 5 | 0.867912349833464 |
| chr9-18208612-18209113 | 8 | 0.817071821013524 |
| chr1-224198942-224199443 | 3 | 0.995021292129093 |
| chr4-152200028-152200529 | 6 | 0.993228295527555 |
| chr18-75224374-75224875 | 8 | 0.970912165232325 |
| chr9-72230787-72231288 | 2 | 0.889316390130849 |
| chr5-116475049-116475550 | 3 | 0.992161020069214 |
| chr10-92798644-92799145 | 4 | 0.971581583388646 |
| chr13-43266195-43266696 | 7 | 0.863113247997551 |
| chr16-55818641-55819142 | 8 | 0.898206346550139 |
| chr16-52016157-52016658 | 8 | 0.95329294433495 |
| chr3-36964122-36964623 | 8 | 0.922306525605681 |
| chr18-73306204-73306705 | 8 | 0.988961988911973 |
| chr5-99331997-99332498 | 3 | 0.983856033298935 |
| chr11-35640605-35641106 | 8 | 0.896598621684576 |
| chr6-128812388-128812889 | 6 | 0.997480133631714 |
| chr10-91220747-91221248 | 8 | 0.908183391854128 |
| chr7-102268278-102268779 | 6 | 0.989411717225865 |
| chr1-104542033-104542534 | 4 | 0.959969611986886 |
| chr8-33855724-33856225 | 5 | 0.991680386895847 |
| chr7-23868941-23869442 | 5 | 0.980247318978162 |
| chr2-58908126-58908627 | 3 | 0.99593553545041 |
| chr1-189977474-189977975 | 4 | 0.923029261878718 |
| chr5-73096813-73097314 | 6 | 0.991856852544109 |
| chr8-29858518-29859019 | 2 | 0.836733577726761 |
| chr1-176981737-176982238 | 3 | 0.983396079280663 |
| chr1-105864618-105865119 | 3 | 0.995609536353085 |
| chr18-74692813-74693314 | 5 | 0.987303065906607 |
| chr15-12709546-12710047 | 8 | 0.968761478380184 |
| chr6-30132254-30132755 | 8 | 0.994780950715529 |
| chr15-82676747-82677248 | 5 | 0.979929536541819 |

| | | |
|---|---|---|
| chr4-133756029-133756530 | 4 | 0.904279432267671 |
| chr8-122282458-122282959 | 1 | 0.867490853848869 |
| chr11-29396962-29397463 | 8 | 0.919778090513051 |
| chr4-27045910-27046411 | 3 | 0.996491515561413 |
| chr15-34505137-34505638 | 8 | 0.984233753039155 |
| chr4-84215540-84216041 | 4 | 0.955401614293317 |
| chr17-12218039-12218540 | 5 | 0.987859371840808 |
| chr1-197654295-197654796 | 5 | 0.849086100903754 |
| chr10-68359402-68359903 | 8 | 0.992129063656169 |
| chr20-51634445-51634946 | 3 | 0.896704400190989 |
| chr1-207801773-207802274 | 4 | 0.970751311828324 |
| chr3-92230179-92230680 | 3 | 0.846652345928485 |
| chr17-62142553-62143054 | 3 | 0.984242631125474 |
| chr14-21962297-21962798 | 4 | 0.95006261625126 |
| chr4-24937311-24937812 | 8 | 0.839124749920178 |
| chr7-57286305-57286806 | 8 | 0.989020326245173 |
| chr15-46043125-46043626 | 4 | 0.896544904157942 |
| chr2-204407677-204408178 | 8 | 0.988617284054481 |
| chr15-78739907-78740408 | 8 | 0.816910871250303 |
| chr1-249608425-249608926 | 8 | 0.962298210447375 |
| chr6-50764481-50764982 | 2 | 0.939832318978678 |
| chr13-46713596-46714097 | 8 | 0.904663053081573 |
| chrX-1305419-1305920 | 7 | 0.832683036773206 |
| chr2-140856335-140856836 | 8 | 0.97827413486456 |
| chr6-119957041-119957542 | 8 | 0.928566145764233 |
| chr1-237042880-237043381 | 6 | 0.993051772932807 |
| chr10-42462421-42462922 | 8 | 0.95431039302722 |
| chr4-116688190-116688691 | 8 | 0.924040997112735 |
| chr17-1078200-1078701 | 5 | 0.98366192891458 |
| chr5-17059913-17060414 | 8 | 0.979958561131583 |
| chr7-83620159-83620660 | 8 | 0.993493681128311 |
| chr2-195169572-195170073 | 5 | 0.98011519707074 |
| chr18-58284111-58284612 | 1 | 0.972941657362974 |
| chr8-4232504-4233005 | 3 | 0.99001509922602 |
| chr7-53888921-53889422 | 7 | 0.946297693586323 |
| chr9-11779336-11779837 | 3 | 0.98422564467906 |
| chr15-24653658-24654159 | 8 | 0.995076729247021 |
| chr10-71160943-71161444 | 5 | 0.984619214441673 |
| chr6-72365008-72365509 | 8 | 0.99507196553308 |
| chr6-85140765-85141266 | 5 | 0.938835978942846 |
| chr14-82142441-82142942 | 1 | 0.872509508281739 |
| chr6-37925500-37926001 | 3 | 0.97518749772435 |
| chr2-248105031-248105532 | 8 | 0.871909452974201 |

| | | |
|---|---|---|
| chr2-211811582-211812083 | 4 | 0.853893399047068 |
| chr3-93166443-93166944 | 8 | 0.993240615344875 |
| chr14-64621291-64621792 | 8 | 0.989696158803897 |
| chr6-14843717-14844218 | 8 | 0.895822578863006 |
| chr11-49524734-49525235 | 3 | 0.957800007836866 |
| chr5-140398319-140398820 | 2 | 0.854078632496726 |
| chr9-99907943-99908444 | 3 | 0.993284051908127 |
| chr6-41639149-41639650 | 8 | 0.895249202256101 |
| chr17-68179150-68179651 | 8 | 0.832454188944926 |
| chr18-69742330-69742831 | 8 | 0.980520489308213 |
| chr5-70109067-70109568 | 8 | 0.915549813609707 |
| chr7-107668804-107669305 | 8 | 0.993493681128311 |
| chr2-138997764-138998265 | 8 | 0.836477079963094 |
| chr14-44554612-44555113 | 1 | 0.960827198476578 |
| chr8-43160375-43160876 | 7 | 0.907271502223741 |
| chr15-49242741-49243242 | 3 | 0.995403068315211 |
| chr17-45988370-45988871 | 8 | 0.981666820242165 |
| chr7-22345042-22345543 | 2 | 0.935222756269776 |
| chr5-125930924-125931425 | 8 | 0.875504265287059 |
| chr14-30199335-30199836 | 4 | 0.883087553041076 |
| chr17-44455431-44455932 | 7 | 0.902945737565474 |
| chr14-22865591-22866092 | 3 | 0.996889659618283 |
| chr3-46034609-46035110 | 8 | 0.94695987798215 |
| chr15-91312990-91313491 | 3 | 0.995222907948518 |
| chr2-41166461-41166962 | 8 | 0.956093960716557 |
| chr1-109135659-109136160 | 8 | 0.95579086477943 |
| chr3-41652517-41653018 | 8 | 0.848544495619119 |
| chr7-66274324-66274825 | 8 | 0.989223421808534 |
| chr7-45902150-45902651 | 5 | 0.872677711214763 |
| chr16-46113845-46114346 | 3 | 0.903540225216306 |
| chr15-99441791-99442292 | 4 | 0.860846486005035 |
| chr8-21340978-21341479 | 8 | 0.994749335658716 |
| chr1-172739938-172740439 | 8 | 0.939901685015074 |
| chr8-36614121-36614622 | 4 | 0.957729166512379 |
| chr2-233538043-233538544 | 5 | 0.939055747404612 |
| chr11-33125094-33125595 | 3 | 0.941461661936046 |
| chr18-47991500-47992001 | 8 | 0.993493681128311 |
| chr18-76385103-76385604 | 8 | 0.903277583402483 |
| chr17-83784456-83784957 | 2 | 0.846141693478039 |
| chr17-19555970-19556471 | 4 | 0.961920007865936 |
| chr11-14311825-14312326 | 6 | 0.984952253054664 |
| chr13-19756297-19756798 | 3 | 0.99593553545041 |
| chr20-24410707-24411208 | 8 | 0.910163886202669 |

| | | |
|---|---|---|
| chr17-19193084-19193585 | 6 | 0.951374082187249 |
| chr3-166953245-166953746 | 7 | 0.967902646004791 |
| chr14-75925388-75925889 | 1 | 0.905111004696302 |
| chr9-1225882-1226383 | 8 | 0.99506043622149 |
| chr20-10814248-10814749 | 8 | 0.986269541343177 |
| chr8-54850743-54851244 | 8 | 0.92753545760672 |
| chr6-53085460-53085961 | 8 | 0.994628681404951 |
| chr8-78312403-78312904 | 4 | 0.840733876342015 |
| chr14-9655251-9655752 | 4 | 0.968010516378556 |
| chr19-32992341-32992842 | 8 | 0.891103584190747 |
| chr10-64442322-64442823 | 8 | 0.981732213696297 |
| chr7-40539391-40539892 | 8 | 0.994966912587968 |
| chr14-45079583-45080084 | 8 | 0.943996555389745 |
| chr9-94949530-94950031 | 8 | 0.858924738977913 |
| chr17-28884918-28885419 | 5 | 0.98273162533797 |
| chr1-99125873-99126374 | 1 | 0.814979369815327 |
| chr12-34941318-34941819 | 4 | 0.971511201928662 |
| chrX-86716579-86717080 | 6 | 0.846026947114421 |
| chr10-88650576-88651077 | 8 | 0.92237618442639 |
| chr7-55581608-55582109 | 5 | 0.957478738965539 |
| chr6-22748897-22749398 | 1 | 0.816100238094451 |
| chrX-33320301-33320802 | 8 | 0.990939100576315 |
| chr2-78880429-78880930 | 6 | 0.992251539153639 |
| chr7-50959926-50960427 | 2 | 0.919172689036789 |
| chr6-40201252-40201753 | 6 | 0.941801290315027 |
| chr8-74689365-74689866 | 7 | 0.897831607235327 |
| chr3-43759603-43760104 | 4 | 0.860555706635751 |
| chr3-142386598-142387099 | 8 | 0.948880395740946 |
| chr3-120649482-120649983 | 8 | 0.962155932027906 |
| chr15-45762865-45763366 | 8 | 0.989576196037054 |
| chr13-14318403-14318904 | 8 | 0.878608312450387 |
| chr1-233461079-233461580 | 8 | 0.990971115476687 |
| chr1-105569760-105570261 | 3 | 0.977899222813375 |
| chr14-72109754-72110255 | 6 | 0.992640867869645 |
| chr5-13022900-13023401 | 3 | 0.990067108995023 |
| chr6-13200520-13201021 | 8 | 0.98517396480388 |
| chr7-124413481-124413982 | 8 | 0.92403746618912 |
| chr10-55452597-55453098 | 2 | 0.94540703059882 |
| chr13-48961731-48962232 | 5 | 0.960652475079396 |
| chr19-14733700-14734201 | 4 | 0.842316064241031 |
| chr6-46257720-46258221 | 8 | 0.986859853934944 |
| chr14-85123481-85123982 | 5 | 0.954809031048012 |
| chr2-191330829-191331330 | 8 | 0.970148054450031 |

| | | |
|---|---|---|
| chr13-35772909-35773410 | 5 | 0.988832280836386 |
| chr14-58675593-58676094 | 8 | 0.995173125917846 |
| chr18-61223703-61224204 | 8 | 0.972911178148287 |
| chr10-10213644-10214145 | 4 | 0.849574844505694 |
| chr3-141565963-141566464 | 8 | 0.994043141143163 |
| chr4-179077240-179077741 | 8 | 0.98277049595143 |
| chr8-84305843-84306344 | 3 | 0.996758589280804 |
| chr5-68951736-68952237 | 8 | 0.980945905725839 |
| chr16-59554835-59555336 | 3 | 0.997169119127373 |
| chr12-22850222-22850723 | 5 | 0.987654525972938 |
| chr14-90443706-90444207 | 3 | 0.964556968783981 |
| chr1-175713487-175713988 | 8 | 0.973717310170913 |
| chr3-142368413-142368914 | 8 | 0.965669201050515 |
| chr4-36068522-36069023 | 8 | 0.96032199522015 |
| chr15-72453028-72453529 | 8 | 0.887008698560505 |
| chr7-83817417-83817918 | 8 | 0.851284046395584 |
| chr5-61451080-61451581 | 2 | 0.867807142311204 |
| chr10-24533094-24533595 | 3 | 0.99593553545041 |
| chr5-149437711-149438212 | 4 | 0.893156928536409 |
| chr2-82683892-82684393 | 2 | 0.817977709437109 |
| chr14-20057781-20058282 | 3 | 0.863677230160499 |
| chr18-32217143-32217644 | 2 | 0.854485024824082 |
| chr11-2194631-2195132 | 3 | 0.99089004607084 |
| chr20-20555276-20555777 | 1 | 0.818198115103663 |
| chr5-57806854-57807355 | 4 | 0.882099289161896 |
| chr7-63674008-63674509 | 6 | 0.991206925481043 |
| chrX-14440516-14441017 | 6 | 0.996802531945538 |
| chr13-31879066-31879567 | 8 | 0.985390773967086 |
| chr1-58320033-58320534 | 8 | 0.948085465760393 |
| chr10-105966411-105966912 | 8 | 0.943589481639184 |
| chr1-165840720-165841221 | 3 | 0.991651118262396 |
| chr14-82015043-82015544 | 8 | 0.926590346236245 |
| chr13-55134185-55134686 | 2 | 0.826816180753593 |
| chr20-4596270-4596771 | 8 | 0.99123447028764 |
| chr7-27906768-27907269 | 7 | 0.879624193196952 |
| chr15-2010456-2010957 | 7 | 0.874078982076964 |
| chr2-210777355-210777856 | 8 | 0.99524893685131 |
| chr10-52755536-52756037 | 8 | 0.889885308928885 |
| chr1-246368798-246369299 | 8 | 0.917113261614234 |
| chr15-28871208-28871709 | 3 | 0.99593553545041 |
| chr13-24587082-24587583 | 1 | 0.937604381230491 |
| chr1-227144879-227145380 | 8 | 0.972154536064326 |
| chr15-33089737-33090238 | 5 | 0.848812702427098 |

| | | |
|---|---|---|
| chr4-58820333-58820834 | 5 | 0.991746774412507 |
| chr5-141285880-141286381 | 4 | 0.917651669636589 |
| chr10-99363536-99364037 | 4 | 0.897939446517479 |
| chr8-26269342-26269843 | 6 | 0.981028286415041 |
| chr1-255416751-255417252 | 8 | 0.938182482085936 |
| chr2-187662904-187663405 | 8 | 0.994985934986239 |
| chr18-70807000-70807501 | 3 | 0.892160662539608 |
| chrX-113792119-113792620 | 3 | 0.99593553545041 |
| chr6-23842586-23843087 | 8 | 0.84924514065068 |
| chr7-103928812-103929313 | 5 | 0.986541198905986 |
| chr5-49285648-49286149 | 8 | 0.995090280454927 |
| chr6-19126276-19126777 | 3 | 0.907078240920325 |
| chr5-132035012-132035513 | 8 | 0.995165673580393 |
| chr6-52927122-52927623 | 8 | 0.994314584081321 |
| chr19-4364183-4364684 | 8 | 0.971328742633202 |
| chr9-4010084-4010585 | 5 | 0.983919978840557 |
| chr17-55942484-55942985 | 8 | 0.995178637149639 |
| chr16-23870896-23871397 | 6 | 0.995592262390051 |
| chr1-88604059-88604560 | 8 | 0.893247336308196 |
| chr1-250660643-250661144 | 8 | 0.899414192521004 |
| chr9-31486606-31487107 | 8 | 0.905159977314605 |
| chr7-62262736-62263237 | 8 | 0.991177413512188 |
| chr13-91517660-91518161 | 8 | 0.897954891691522 |
| chr3-26888015-26888516 | 8 | 0.952857727956748 |
| chr1-259942838-259943339 | 5 | 0.99319356500838 |
| chr2-36078168-36078669 | 8 | 0.861274559431217 |
| chr8-102733523-102734024 | 3 | 0.997234042548622 |
| chr1-48751324-48751825 | 2 | 0.906533265936313 |
| chr17-50323972-50324473 | 3 | 0.96993600028298 |
| chr7-10009857-10010358 | 3 | 0.994500864784128 |
| chrX-116469308-116469809 | 5 | 0.974062466218128 |
| chr9-38598800-38599301 | 4 | 0.877905357326143 |
| chr7-46226053-46226554 | 8 | 0.873922674481343 |
| chr2-49567081-49567582 | 8 | 0.929691365104499 |
| chr4-133962956-133963457 | 7 | 0.854737549356872 |
| chrY-9829965-9830466 | 3 | 0.990002896833823 |
| chr17-47444990-47445491 | 8 | 0.877307553856892 |
| chr11-69768306-69768807 | 7 | 0.913131198514741 |
| chr6-130091318-130091819 | 8 | 0.840645217082798 |
| chr2-33218652-33219153 | 4 | 0.969240092197458 |
| chr6-120205870-120206371 | 8 | 0.845888215905634 |
| chr3-69940105-69940606 | 4 | 0.94315029063396 |
| chr6-40848435-40848936 | 8 | 0.87266533994053 |

| | | |
|---|---|---|
| chr5-162865765-162866266 | 8 | 0.975354170279469 |
| chr1-257187027-257187528 | 7 | 0.939362681195033 |
| chr8-29808057-29808558 | 4 | 0.87783693314702 |
| chr9-37131845-37132346 | 1 | 0.881383555643444 |
| chr18-19721666-19722167 | 8 | 0.880312948642621 |
| chr20-10722402-10722903 | 8 | 0.904575244547281 |
| chr7-35366385-35366886 | 8 | 0.993493681128311 |
| chr5-37354591-37355092 | 3 | 0.912883852065539 |
| chr5-9022516-9023017 | 5 | 0.973853698612542 |
| chr18-71699703-71700204 | 8 | 0.891148956537988 |
| chr4-105155458-105155959 | 8 | 0.982906585002943 |
| chr5-5661254-5661755 | 3 | 0.992282577814975 |
| chr10-64864077-64864578 | 8 | 0.993493681128311 |
| chr1-150777013-150777514 | 8 | 0.995209802647447 |
| chr3-25190231-25190732 | 6 | 0.991206925481043 |
| chr5-54205392-54205893 | 3 | 0.996208544337453 |
| chr5-45548042-45548543 | 7 | 0.860066866268389 |
| chr19-19296274-19296775 | 8 | 0.993493681128311 |
| chr1-18809335-18809836 | 8 | 0.974052954099261 |
| chr14-61103713-61104214 | 5 | 0.988411070131831 |
| chr20-45118245-45118746 | 2 | 0.940393270549469 |
| chr17-21005917-21006418 | 8 | 0.911814091673944 |
| chr9-7934404-7934905 | 6 | 0.992437810902631 |
| chr6-129001609-129002110 | 8 | 0.991687578096659 |
| chr9-85797524-85798025 | 8 | 0.924533572965495 |
| chr17-86447294-86447795 | 8 | 0.840122278261695 |
| chr2-135095519-135096020 | 8 | 0.993493681128311 |
| chr9-107389705-107390206 | 7 | 0.938299983149289 |
| chr8-57376928-57377429 | 8 | 0.982344624100442 |
| chr13-32072062-32072563 | 8 | 0.970831695937099 |
| chr1-123575827-123576328 | 6 | 0.959776886382773 |
| chr1-229596178-229596679 | 8 | 0.952085047574557 |
| chr12-45709585-45710086 | 8 | 0.976859916812232 |
| chr5-110480895-110481396 | 3 | 0.997721625774242 |
| chr14-36926375-36926876 | 3 | 0.996179840004265 |
| chr14-88738746-88739247 | 8 | 0.987789715720316 |
| chr4-172046711-172047212 | 3 | 0.996820713659207 |
| chr2-241960920-241961421 | 7 | 0.823178312992766 |
| chr17-74946763-74947264 | 4 | 0.852081704707365 |
| chr19-29168183-29168684 | 8 | 0.816452540270209 |
| chr8-34273183-34273684 | 3 | 0.988933947371583 |
| chr3-93687692-93688193 | 8 | 0.979618841277836 |
| chr18-51923872-51924373 | 8 | 0.993612594241317 |

| | | |
|---|---|---|
| chr20-43223394-43223895 | 8 | 0.966915092745696 |
| chr14-31962933-31963434 | 8 | 0.988671343166704 |
| chr15-51576390-51576891 | 8 | 0.924349429301028 |
| chr15-82404586-82405087 | 4 | 0.877555203272099 |
| chr18-47600902-47601403 | 8 | 0.992596841877379 |
| chr9-25044912-25045413 | 4 | 0.968887921713936 |
| chr5-119112293-119112794 | 7 | 0.920082902490678 |
| chr1-229256756-229257257 | 8 | 0.954150755368457 |
| chr10-34827267-34827768 | 8 | 0.83417147288529 |
| chr16-72518702-72519203 | 6 | 0.995689323112258 |
| chr1-127671789-127672290 | 3 | 0.99593553545041 |
| chr16-42504087-42504588 | 3 | 0.99593553545041 |
| chr7-5582176-5582677 | 3 | 0.997727574011814 |
| chr8-103560919-103561420 | 3 | 0.939267906292026 |
| chr14-84078310-84078811 | 3 | 0.99593553545041 |
| chr1-121536167-121536668 | 8 | 0.91994482513434 |
| chr4-160903526-160904027 | 8 | 0.979338624251982 |
| chr8-102793647-102794148 | 3 | 0.997592102779378 |
| chr3-117149378-117149879 | 8 | 0.993493681128311 |
| chrX-127562193-127562694 | 8 | 0.920304226518822 |
| chr15-1203780-1204281 | 4 | 0.861451803017381 |
| chrX-3341635-3342136 | 8 | 0.847490511215106 |
| chr4-91483442-91483943 | 3 | 0.993530461229751 |
| chr2-137513382-137513883 | 5 | 0.87995754009626 |
| chr20-6899936-6900437 | 8 | 0.915284189841314 |
| chr4-3669979-3670480 | 6 | 0.986180270603477 |
| chrX-115115288-115115789 | 2 | 0.835083243973302 |
| chr8-100146245-100146746 | 8 | 0.99472630726627 |
| chr13-50030857-50031358 | 8 | 0.993493681128311 |
| chr2-116129811-116130312 | 6 | 0.981744459788645 |
| chr14-66060739-66061240 | 8 | 0.993493681128311 |
| chr2-59927203-59927704 | 8 | 0.914759377556564 |
| chr11-50702335-50702836 | 8 | 0.993493681128311 |
| chr14-75627827-75628328 | 5 | 0.915596613089096 |
| chrX-51679226-51679727 | 6 | 0.992403689129333 |
| chrX-75835441-75835942 | 3 | 0.99593553545041 |
| chr1-189612016-189612517 | 3 | 0.817274333024555 |
| chr9-36960694-36961195 | 8 | 0.902049075101668 |
| chr4-118095973-118096474 | 3 | 0.998305705610864 |
| chr20-43574879-43575380 | 8 | 0.988516647906191 |
| chr6-52899545-52900046 | 8 | 0.819623243087742 |
| chr1-236773877-236774378 | 3 | 0.872234416947887 |
| chr1-148517256-148517757 | 8 | 0.994638060596724 |

| | | |
|---|---|---|
| chr6-117920114-117920615 | 2 | 0.93782213659568 |
| chr14-52671637-52672138 | 8 | 0.995175134294504 |
| chr1-150778612-150779113 | 8 | 0.962202567250932 |
| chr13-42567922-42568423 | 8 | 0.969543377729928 |
| chr16-57930683-57931184 | 8 | 0.892232099251012 |
| chr13-30831374-30831875 | 3 | 0.988312710223522 |
| chr11-33210567-33211068 | 7 | 0.964599746218132 |
| chr7-35878377-35878878 | 3 | 0.996613453281408 |
| chr12-34766276-34766777 | 8 | 0.895807571711223 |
| chrX-88226920-88227421 | 6 | 0.971356449128586 |
| chr4-63985201-63985702 | 8 | 0.959194609612719 |
| chr7-1753140-1753641 | 3 | 0.99593553545041 |
| chr4-93094572-93095073 | 3 | 0.996854246887445 |
| chr19-30931263-30931764 | 8 | 0.904484318976609 |
| chr1-51089008-51089509 | 3 | 0.99593553545041 |
| chr12-46322955-46323456 | 3 | 0.998316067264349 |
| chr1-14716054-14716555 | 2 | 0.841736734482323 |
| chr10-41354438-41354939 | 8 | 0.995182007550015 |
| chr2-130809002-130809503 | 4 | 0.972873907861843 |
| chr6-48675746-48676247 | 8 | 0.944248103468002 |
| chr5-139966735-139967236 | 8 | 0.833880240683251 |
| chr5-1918644-1919145 | 8 | 0.982018590111042 |
| chr7-108310677-108311178 | 8 | 0.878725481868167 |
| chr10-22994044-22994545 | 6 | 0.986809674511101 |
| chr11-56260730-56261231 | 3 | 0.997528624650192 |
| chr1-2255420-2255921 | 8 | 0.927592993908799 |
| chr1-120338832-120339333 | 5 | 0.983919978840557 |
| chr7-97858251-97858752 | 3 | 0.99662638484643 |
| chr14-99441435-99441936 | 8 | 0.935869432233022 |
| chr7-31712842-31713343 | 6 | 0.983317006025103 |
| chr4-52752788-52753289 | 6 | 0.989451465130694 |
| chr11-82253758-82254259 | 6 | 0.959030810507356 |
| chr19-29050875-29051376 | 8 | 0.97188248521919 |
| chr14-60070444-60070945 | 3 | 0.977545131178352 |
| chr19-13434893-13435394 | 8 | 0.937577925937076 |
| chr1-185973322-185973823 | 3 | 0.974117528750911 |
| chr10-74765880-74766381 | 5 | 0.876616914298627 |
| chr2-175555360-175555861 | 2 | 0.870486387802123 |
| chr6-6256555-6257056 | 4 | 0.91488282666685 |
| chr9-45794247-45794748 | 1 | 0.932392382859693 |
| chr8-111289937-111290438 | 8 | 0.899118703025963 |
| chr1-156242456-156242957 | 8 | 0.962763986325486 |
| chr5-49949650-49950151 | 8 | 0.931772265911168 |

| | | |
|---|---|---|
| chr8-64822126-64822627 | 5 | 0.974872991861749 |
| chr13-96793180-96793681 | 6 | 0.984174561405746 |
| chr3-141931387-141931888 | 8 | 0.899137654832986 |
| chr9-37990670-37991171 | 5 | 0.931635002068415 |
| chr4-91271097-91271598 | 3 | 0.922625037713335 |
| chr3-148240743-148241244 | 2 | 0.861950603153728 |
| chr13-78126475-78126976 | 5 | 0.969971497178053 |
| chr4-169326490-169326991 | 8 | 0.984720135046261 |
| chr7-83773377-83773878 | 3 | 0.995650113470664 |
| chr16-43732244-43732745 | 8 | 0.994870423980662 |
| chr12-11273245-11273746 | 8 | 0.935987184005892 |
| chr15-62367346-62367847 | 8 | 0.984348184561296 |
| chr4-181635643-181636144 | 3 | 0.996750594495595 |
| chr13-93623895-93624396 | 5 | 0.97487545379448 |
| chr3-112422278-112422779 | 2 | 0.926507623972299 |
| chr11-7552353-7552854 | 3 | 0.99593553545041 |
| chr14-70251255-70251756 | 8 | 0.852903814011955 |
| chr13-100536369-100536870 | 8 | 0.986253720134407 |
| chr4-176516335-176516836 | 3 | 0.996798493274048 |
| chr7-126864266-126864767 | 8 | 0.905177881843888 |
| chr5-83765018-83765519 | 3 | 0.953728737496609 |
| chr5-18477683-18478184 | 8 | 0.992811055479646 |
| chr8-66472122-66472623 | 8 | 0.971485889203936 |
| chr18-41044694-41045195 | 8 | 0.892602269686005 |
| chr1-143403405-143403906 | 4 | 0.948379559670944 |
| chr8-96948661-96949162 | 5 | 0.892409835460451 |
| chr3-21360454-21360955 | 4 | 0.942478755760754 |
| chr9-55801457-55801958 | 8 | 0.902700330414486 |
| chr6-738148-738649 | 3 | 0.99593553545041 |
| chr5-28509851-28510352 | 3 | 0.991781353051963 |
| chr9-97781565-97782066 | 5 | 0.932241689836963 |
| chr6-87392177-87392678 | 4 | 0.942569239379316 |
| chr3-105156940-105157441 | 2 | 0.938363850644714 |
| chr20-40773873-40774374 | 8 | 0.980116125968824 |
| chr6-83523587-83524088 | 3 | 0.993575526594232 |
| chr14-82586697-82587198 | 8 | 0.991359877957344 |
| chr5-110050997-110051498 | 6 | 0.996015948800712 |
| chr5-61832797-61833298 | 2 | 0.950191342178067 |
| chr5-91689691-91690192 | 8 | 0.966593899152791 |
| chr2-153338196-153338697 | 8 | 0.971389 |
| chr16-23593297-23593798 | 3 | 0.997389767637381 |
| chr1-92468378-92468879 | 2 | 0.897193053465481 |
| chr2-51616575-51617076 | 4 | 0.838152809671687 |

| | | |
|---|---|---|
| chr2-120017440-120017941 | 8 | 0.923364330281202 |
| chrX-148553429-148553930 | 3 | 0.991799824755738 |
| chr2-43380962-43381463 | 3 | 0.964676475137801 |
| chr5-54920804-54921305 | 3 | 0.978592968606984 |
| chr15-15918074-15918575 | 1 | 0.849314137261211 |
| chr2-31283445-31283946 | 8 | 0.917416388137247 |
| chr4-136633001-136633502 | 5 | 0.989916398570028 |
| chr1-177144879-177145380 | 8 | 0.951085120506144 |
| chr17-57374562-57375063 | 8 | 0.938412349271804 |
| chr19-47642540-47643041 | 5 | 0.97759257765303 |
| chr5-143686397-143686898 | 8 | 0.994631687646404 |
| chr6-84878409-84878910 | 8 | 0.93514400914118 |
| chr7-80731525-80732026 | 8 | 0.873266451936634 |
| chr10-43097354-43097855 | 6 | 0.991206925481043 |
| chr4-44031384-44031885 | 4 | 0.953100191315248 |
| chr11-58455025-58455526 | 8 | 0.988195879014477 |
| chr7-115047229-115047730 | 4 | 0.869299125046323 |
| chr10-84790535-84791036 | 8 | 0.952802643035133 |
| chr3-39317978-39318479 | 4 | 0.922382127094833 |
| chr3-44074048-44074549 | 4 | 0.84827215461163 |
| chr7-49280826-49281327 | 6 | 0.976058858498865 |
| chr17-85269414-85269915 | 5 | 0.991092112643062 |
| chr1-144556292-144556793 | 4 | 0.886346304866021 |
| chr5-20512852-20513353 | 1 | 0.828233542079733 |
| chr8-79707436-79707937 | 1 | 0.859151630476659 |
| chr4-122832986-122833487 | 8 | 0.844125865062122 |
| chr6-110150175-110150676 | 3 | 0.920709901298866 |
| chr1-191795770-191796271 | 8 | 0.934306763522995 |
| chr4-125107072-125107573 | 8 | 0.898088225821548 |
| chr2-19858286-19858787 | 3 | 0.99593553545041 |
| chr10-60498092-60498593 | 3 | 0.950297135310651 |
| chr11-34978014-34978515 | 3 | 0.982057014622362 |
| chr16-16874126-16874627 | 7 | 0.862638455944218 |
| chr11-47609128-47609629 | 2 | 0.93233945014526 |
| chr4-127024193-127024694 | 8 | 0.968499632563695 |
| chr2-83770326-83770827 | 8 | 0.993493681128311 |
| chr6-58348772-58349273 | 2 | 0.904152238355222 |
| chr5-140350125-140350626 | 6 | 0.94223625799605 |
| chr17-66192909-66193410 | 2 | 0.848306734703072 |
| chr10-53418108-53418609 | 7 | 0.969724146466636 |
| chr14-53612348-53612849 | 3 | 0.99593553545041 |
| chr7-114099636-114100137 | 8 | 0.961814064548332 |
| chr6-22174769-22175270 | 7 | 0.949689420286583 |

| | | |
|---|---|---|
| chr6-92040238-92040739 | 8 | 0.925801566582007 |
| chr14-83189797-83190298 | 5 | 0.956639444507985 |
| chr4-176486748-176487249 | 8 | 0.99513603045372 |
| chr1-134467481-134467982 | 5 | 0.985778175395781 |
| chr3-134201112-134201613 | 8 | 0.8418191852494 |
| chr7-105836202-105836703 | 1 | 0.881325706395845 |
| chr1-105516969-105517470 | 6 | 0.991206925481043 |
| chr17-85124710-85125211 | 2 | 0.872327163910893 |
| chr19-43368249-43368750 | 3 | 0.994818520046311 |
| chr19-13966990-13967491 | 8 | 0.816237320154314 |
| chr9-90407821-90408322 | 5 | 0.993135651031457 |
| chr11-48365668-48366169 | 3 | 0.993105592369514 |
| chr4-58479495-58479996 | 8 | 0.913089181155443 |
| chr11-75323495-75323996 | 8 | 0.951197861738178 |
| chr14-59072013-59072514 | 4 | 0.949044018137223 |
| chr20-49152940-49153441 | 6 | 0.979862189505702 |
| chr4-89902967-89903468 | 7 | 0.849508222143246 |
| chr6-26364641-26365142 | 8 | 0.951091956109309 |
| chr1-249510553-249511054 | 3 | 0.994305230876888 |
| chr1-219528819-219529320 | 8 | 0.869847057726491 |
| chr1-49714398-49714899 | 6 | 0.963063340235358 |
| chr17-42785598-42786099 | 3 | 0.912024244237906 |
| chr9-84056731-84057232 | 5 | 0.877096598054853 |
| chr1-69401022-69401523 | 8 | 0.929862693414071 |
| chr1-142191827-142192328 | 8 | 0.972484226464934 |
| chr12-31661508-31662009 | 8 | 0.957838701876485 |
| chr7-44250696-44251197 | 8 | 0.893714487924929 |
| chr3-159978385-159978886 | 7 | 0.918068337409502 |
| chr17-2574158-2574659 | 8 | 0.993493681128311 |
| chr4-23976993-23977494 | 3 | 0.993183237713101 |
| chr1-57304413-57304914 | 3 | 0.99593553545041 |
| chr6-54316297-54316798 | 8 | 0.875992122632417 |
| chr16-73018629-73019130 | 3 | 0.995344302823838 |
| chr4-129643710-129644211 | 5 | 0.996111827835971 |
| chr5-163406121-163406622 | 4 | 0.975128367116888 |
| chr6-113223360-113223861 | 5 | 0.929336357744576 |
| chr12-5220215-5220716 | 2 | 0.913726467348443 |
| chr12-12083452-12083953 | 5 | 0.95624318848748 |
| chr2-120588031-120588532 | 3 | 0.994922405565383 |
| chr1-143523234-143523735 | 6 | 0.992286821621332 |
| chr3-106434582-106435083 | 2 | 0.953380633922189 |
| chr2-57180668-57181169 | 2 | 0.945577117030331 |
| chr1-257973667-257974168 | 8 | 0.990380127727922 |

| | | |
|---|---|---|
| chr19-3261958-3262459 | 8 | 0.995040373203682 |
| chr11-60601030-60601531 | 8 | 0.925997609030091 |
| chr2-159976145-159976646 | 4 | 0.85137138663431 |
| chr2-215311439-215311940 | 8 | 0.993493681128311 |
| chr14-2944713-2945214 | 8 | 0.970507348547178 |
| chr3-119546151-119546652 | 8 | 0.928873274583807 |
| chr14-84184822-84185323 | 8 | 0.988970945549236 |
| chr13-33352955-33353456 | 3 | 0.99815516934407 |
| chr9-1037971-1038472 | 7 | 0.922596424176107 |
| chr19-54289446-54289947 | 5 | 0.970277415619737 |
| chr20-28495672-28496173 | 5 | 0.843739141651184 |
| chr20-32558591-32559092 | 8 | 0.993493681128311 |
| chr5-99307083-99307584 | 2 | 0.819761882800142 |
| chr13-81734576-81735077 | 8 | 0.994481293832383 |
| chr17-23268153-23268654 | 8 | 0.9883787697745 |
| chr11-83753568-83754069 | 8 | 0.958573333388932 |
| chrX-115463036-115463537 | 6 | 0.836341709286151 |
| chr17-78006652-78007153 | 1 | 0.902186852509171 |
| chr8-22820097-22820598 | 6 | 0.989295283789051 |
| chr9-52252456-52252957 | 3 | 0.934119140953401 |
| chr17-71061700-71062201 | 8 | 0.993493681128311 |
| chr12-46368293-46368794 | 3 | 0.98895012049733 |
| chr19-42502816-42503317 | 8 | 0.825976833031952 |
| chrX-35869773-35870274 | 6 | 0.988256792725008 |
| chr10-79284984-79285485 | 8 | 0.97261560548784 |
| chr19-36342503-36343004 | 8 | 0.988201667957702 |
| chr16-17571696-17572197 | 3 | 0.939268389395463 |
| chr7-38810527-38811028 | 3 | 0.995935535545041 |
| chr4-63103195-63103696 | 8 | 0.940522529315638 |
| chr14-44914925-44915426 | 3 | 0.998021770396126 |
| chr4-154829720-154830221 | 3 | 0.97528049530553 |
| chr6-5906216-5906717 | 5 | 0.920785150557163 |
| chr3-22024247-22024748 | 8 | 0.934377197874387 |
| chr1-167627074-167627575 | 8 | 0.90237059446779 |
| chr7-120315048-120315549 | 8 | 0.985772928747312 |
| chr4-100800919-100801420 | 3 | 0.995935535545041 |
| chr2-29337254-29337755 | 8 | 0.960634371641987 |
| chr17-80913426-80913927 | 2 | 0.905121920121911 |
| chr9-81285495-81285996 | 6 | 0.882420041140427 |
| chr16-63008304-63008805 | 8 | 0.990774768783042 |
| chr5-28497953-28498454 | 3 | 0.995935535545041 |
| chr3-7269122-7269623 | 2 | 0.95696540744469 |
| chr3-149222169-149222670 | 4 | 0.841689004338205 |

| | | |
|---|---|---|
| chr13-33043782-33044283 | 8 | 0.993493681128311 |
| chr3-34145513-34146014 | 8 | 0.977444482912384 |
| chr5-119829187-119829688 | 8 | 0.995218059353945 |
| chr2-153616639-153617140 | 8 | 0.984023348797939 |
| chr2-23111415-23111916 | 8 | 0.833979975176185 |
| chr3-45602153-45602654 | 8 | 0.913076675627006 |
| chr4-146665843-146666344 | 5 | 0.933352756236274 |
| chr11-56434999-56435500 | 4 | 0.974927190689243 |
| chr15-42410446-42410947 | 5 | 0.973563375639288 |
| chr19-20817786-20818287 | 8 | 0.966748116178076 |
| chr1-208091302-208091803 | 3 | 0.900875604162854 |
| chr3-126221266-126221767 | 8 | 0.993493681128311 |
| chr2-117779209-117779710 | 4 | 0.911567838070509 |
| chr2-219786910-219787411 | 8 | 0.99193401709521 |
| chr8-27665080-27665581 | 8 | 0.863649308800123 |
| chr17-25459844-25460345 | 3 | 0.99593553545041 |
| chr20-18658226-18658727 | 5 | 0.992519880135790 |
| chr17-58999341-58999842 | 3 | 0.86148016271581 |
| chr14-43137400-43137901 | 8 | 0.97152641397586 |
| chr7-68310532-68311033 | 8 | 0.985844232841029 |
| chr5-88690783-88691284 | 6 | 0.975714886138712 |
| chr1-205168177-205168678 | 7 | 0.949809003489688 |
| chr2-140540338-140540839 | 8 | 0.875696976847773 |
| chr5-48719083-48719584 | 8 | 0.905415599755058 |
| chr5-107857583-107858084 | 6 | 0.955018284360771 |
| chr10-66464983-66465484 | 7 | 0.926753364792667 |
| chr4-24285777-24286278 | 3 | 0.99593553545041 |
| chr10-16974616-16975117 | 8 | 0.978738090307365 |
| chr4-109872145-109872646 | 3 | 0.99593553545041 |
| chr20-19325712-19326213 | 8 | 0.943491114719767 |
| chr6-74804215-74804716 | 5 | 0.870323671080692 |
| chr17-44517553-44518054 | 5 | 0.983791549229828 |
| chr3-159637665-159638166 | 6 | 0.992586972870175 |
| chr20-15552818-15553319 | 3 | 0.980178858578278 |
| chr3-65052124-65052625 | 6 | 0.991985380425236 |
| chr2-82385910-82386411 | 8 | 0.989358710009268 |
| chr14-26514562-26515063 | 2 | 0.966590890969078 |
| chr1-175150297-175150798 | 4 | 0.878164072395399 |
| chr20-28797284-28797785 | 5 | 0.979905373963426 |
| chrX-63848348-63848849 | 8 | 0.863607492804242 |
| chr5-166601670-166602171 | 8 | 0.959902816367004 |
| chrX-31109229-31109730 | 8 | 0.993493681128311 |
| chr19-15872252-15872753 | 3 | 0.946615565954662 |

| | | |
|---|---|---|
| chr12-41777242-41777743 | 5 | 0.978716388037577 |
| chr18-46306116-46306617 | 8 | 0.918689831881818 |
| chr12-40648360-40648861 | 8 | 0.985549005384958 |
| chr3-126301586-126302087 | 8 | 0.94432765451685 |
| chr1-144584881-144585382 | 2 | 0.946101744761925 |
| chr8-33566875-33567376 | 8 | 0.945155850539477 |
| chr13-37052456-37052957 | 8 | 0.993493681128311 |
| chr5-103708391-103708892 | 6 | 0.991514065960296 |
| chr15-95604761-95605262 | 5 | 0.882609543368393 |
| chr1-149585623-149586124 | 8 | 0.976501893417225 |
| chr3-32335261-32335762 | 4 | 0.948940453988398 |
| chr17-37429601-37430102 | 5 | 0.887986863050255 |
| chr2-55491060-55491561 | 2 | 0.948972112067708 |
| chrX-131766996-131767497 | 5 | 0.989714845114254 |
| chr13-70962383-70962884 | 8 | 0.820804888452062 |
| chrX-128138982-128139483 | 6 | 0.898946198349534 |
| chr18-66591977-66592478 | 3 | 0.996826412221114 |
| chr4-45954231-45954732 | 4 | 0.830107988835591 |
| chr13-22922067-22922568 | 8 | 0.927370730955812 |
| chr5-111330883-111331384 | 3 | 0.993762085075582 |
| chr4-58499344-58499845 | 8 | 0.993493681128311 |
| chr16-74976412-74976913 | 8 | 0.933033699861084 |
| chr2-37270020-37270521 | 4 | 0.943578157957665 |
| chr8-67752168-67752669 | 8 | 0.955549581752173 |
| chr4-151287914-151288415 | 8 | 0.995239750392728 |
| chr19-25979427-25979928 | 8 | 0.958141199053776 |
| chr8-94507063-94507564 | 3 | 0.996892927205352 |
| chr2-242038336-242038837 | 4 | 0.901461866863128 |
| chr6-132635477-132635978 | 3 | 0.99593553545041 |
| chr11-547612-548113 | 5 | 0.868421607337303 |
| chr2-74648099-74648600 | 3 | 0.996768060346266 |
| chr8-59828775-59829276 | 5 | 0.965756891979294 |
| chr5-81621481-81621982 | 3 | 0.996867368066798 |
| chr10-99645883-99646384 | 8 | 0.890919339055551 |
| chr10-50383131-50383632 | 6 | 0.945139909404771 |
| chr11-36453254-36453755 | 3 | 0.971076702029771 |
| chr5-141915525-141916026 | 3 | 0.883705061352331 |
| chr7-106009059-106009560 | 8 | 0.990777757621015 |
| chr6-112385894-112386395 | 5 | 0.877464438032324 |
| chr14-59267792-59268293 | 8 | 0.983915135555041 |
| chr4-164352644-164353145 | 3 | 0.996845412217941 |
| chr5-20362217-20362718 | 8 | 0.924472669372339 |
| chr20-9386744-9387245 | 8 | 0.950365327198718 |

| | | |
|---|---|---|
| chr9-48091688-48092189 | 5 | 0.983111891131011 |
| chr18-14969176-14969677 | 5 | 0.983919978840557 |
| chr13-65145132-65145633 | 8 | 0.882877630978176 |
| chr5-6208988-6209489 | 3 | 0.974904529503569 |
| chr4-9069918-9070419 | 4 | 0.878057458594812 |
| chr2-173577849-173578350 | 8 | 0.970007600034136 |
| chr16-25477500-25478001 | 2 | 0.873356727772429 |
| chr5-23614108-23614609 | 8 | 0.910967675668493 |
| chr3-134245357-134245858 | 5 | 0.983919978840557 |
| chr15-9273589-9274090 | 3 | 0.966366115835657 |
| chr5-9727735-9728236 | 8 | 0.994759672830306 |
| chr11-20354679-20355180 | 7 | 0.825007469709441 |
| chr3-168903696-168904197 | 3 | 0.866368837002977 |
| chr1-8963563-8964064 | 4 | 0.954051311562055 |
| chr19-43683014-43683515 | 5 | 0.860101686691051 |
| chr6-13829866-13830367 | 8 | 0.993493681128311 |
| chr3-157641105-157641606 | 8 | 0.851394292040706 |
| chr17-43878034-43878535 | 2 | 0.950166202256989 |
| chr15-93546333-93546834 | 6 | 0.9912615069838 |
| chr6-58112561-58113062 | 6 | 0.993153211128866 |
| chr17-81993314-81993815 | 8 | 0.949373188859891 |
| chr1-171217086-171217587 | 1 | 0.822049417848157 |
| chr1-124145820-124146321 | 2 | 0.936468025776039 |
| chr4-164678555-164679056 | 3 | 0.996796897218897 |
| chr14-93319094-93319595 | 8 | 0.988248777927572 |
| chr17-17334846-17335347 | 5 | 0.989742261725034 |
| chr6-91131661-91132162 | 5 | 0.991725287752456 |
| chr3-38567682-38568183 | 8 | 0.954801436362133 |
| chr14-1915867-1916368 | 2 | 0.946339849671412 |
| chr19-33120453-33120954 | 4 | 0.849657283168177 |
| chr1-103952591-103953092 | 3 | 0.99593553545041 |
| chr20-18250389-18250890 | 8 | 0.937104170164926 |
| chr1-202889282-202889783 | 8 | 0.881584828068694 |
| chr1-54957406-54957907 | 8 | 0.888298077223646 |
| chr19-13219751-13220252 | 8 | 0.928903629222809 |
| chr15-14158399-14158900 | 5 | 0.985018781549612 |
| chr8-98947494-98947995 | 6 | 0.995976407310364 |
| chr1-258237021-258237522 | 8 | 0.87567209335085 |
| chr3-154303995-154304496 | 2 | 0.948643329611137 |
| chr4-101794620-101795121 | 3 | 0.99593553545041 |
| chrX-11147705-11148206 | 8 | 0.835129705361854 |
| chr2-140115805-140116306 | 3 | 0.968149292717095 |
| chr11-42017098-42017599 | 6 | 0.838709576034509 |

| | | |
|---|---|---|
| chr17-42609143-42609644 | 4 | 0.971506588144564 |
| chr14-75975504-75976005 | 5 | 0.985699431432156 |
| chr18-13040213-13040714 | 3 | 0.99593553545041 |
| chr17-32105098-32105599 | 8 | 0.921458316423025 |
| chr5-117190170-117190671 | 4 | 0.880647246047561 |
| chr8-19582578-19583079 | 3 | 0.874051889228427 |
| chr6-44684104-44684605 | 8 | 0.977399710437044 |
| chr15-20844120-20844621 | 8 | 0.993493681128311 |
| chr1-119223834-119224335 | 8 | 0.989524480380225 |
| chr6-139346617-139347118 | 4 | 0.962286743307133 |
| chr20-44152786-44153287 | 8 | 0.862004830298006 |
| chr1-249514251-249514752 | 8 | 0.965467204237073 |
| chr15-39873713-39874214 | 8 | 0.987889591651679 |
| chr7-95176100-95176601 | 8 | 0.870441796669454 |
| chr5-7725622-7726123 | 6 | 0.994360253583701 |
| chr16-5694489-5694990 | 8 | 0.855384366583569 |
| chr8-40961702-40962203 | 8 | 0.861108696573451 |
| chr3-117775046-117775547 | 4 | 0.914347204747012 |
| chr2-46512309-46512810 | 6 | 0.992236001810622 |
| chr9-86016608-86017109 | 8 | 0.887059647783287 |
| chr2-135833307-135833808 | 8 | 0.993493681128311 |
| chr6-27308692-27309193 | 8 | 0.993493681128311 |
| chr10-66652610-66653111 | 3 | 0.954913403126732 |
| chr7-27390085-27390586 | 8 | 0.985641799309007 |
| chr2-20845402-20845903 | 8 | 0.960508309745505 |
| chr2-54374270-54374771 | 8 | 0.991175785558071 |
| chr1-209363911-209364412 | 8 | 0.99397549589471 |
| chr2-151609865-151610366 | 8 | 0.919381833863723 |
| chr8-14014363-14014864 | 3 | 0.931356372705706 |
| chr6-115535820-115536321 | 3 | 0.986549766188483 |
| chr15-1537657-1538158 | 8 | 0.991948855342964 |
| chr2-10371120-10371621 | 6 | 0.983235163367558 |
| chr3-85676526-85677027 | 3 | 0.986740926690971 |
| chr6-43684210-43684711 | 5 | 0.917343638609745 |
| chr19-48558649-48559150 | 5 | 0.970645458955601 |
| chr10-21375280-21375781 | 1 | 0.944867160947875 |
| chr1-7156906-7157407 | 5 | 0.958024553626637 |
| chr1-70283437-70283938 | 3 | 0.993407678769924 |
| chr13-3696081-3696582 | 6 | 0.920030182523245 |
| chrY-13969454-13969955 | 3 | 0.994957495557718 |
| chr20-8135894-8136395 | 5 | 0.963154382909108 |
| chr5-61623448-61623949 | 8 | 0.930758635917954 |
| chr7-29019070-29019571 | 8 | 0.961554092457413 |

| | | |
|---|---|---|
| chr13-85115510-85116011 | 3 | 0.986184008633205 |
| chr13-97507455-97507956 | 4 | 0.975178744076571 |
| chr19-37439584-37440085 | 2 | 0.894579827344524 |
| chr1-219242873-219243374 | 8 | 0.873981452194564 |
| chr8-117850588-117851089 | 8 | 0.878415795849943 |
| chrX-24355635-24356136 | 2 | 0.880288722286736 |
| chr16-9354313-9354814 | 6 | 0.989761969552554 |
| chr2-112036251-112036752 | 4 | 0.956719140107321 |
| chr6-14687618-14688119 | 8 | 0.993493681128311 |
| chr7-113822950-113823451 | 8 | 0.878731577739782 |
| chr9-110594221-110594722 | 8 | 0.917832604657502 |
| chr11-35219969-35220470 | 8 | 0.954812202194475 |
| chr5-48718175-48718676 | 5 | 0.96703575319267 |
| chr10-37734749-37735250 | 8 | 0.824656275478903 |
| chr3-99837396-99837897 | 5 | 0.947777198026278 |
| chr1-200110504-200111005 | 8 | 0.938296381662895 |
| chr4-141637265-141637766 | 4 | 0.85278196730008 |
| chr16-79500874-79501375 | 8 | 0.916489120375144 |
| chr10-103262728-103263229 | 1 | 0.896492838048912 |
| chr12-29125929-29126430 | 3 | 0.943328495692177 |
| chr3-91810481-91810982 | 7 | 0.892234460902409 |
| chr5-142665412-142665913 | 5 | 0.984565594444946 |
| chr7-83909522-83910023 | 3 | 0.99593553545041 |
| chr10-26302409-26302910 | 8 | 0.916402313500905 |
| chr16-59005010-59005511 | 1 | 0.941243054695366 |
| chr9-54362087-54362588 | 8 | 0.82634091741934 |
| chr10-34049373-34049874 | 3 | 0.898690150481189 |
| chr15-46965475-46965976 | 8 | 0.92165008487129 |
| chr6-111559603-111560104 | 3 | 0.995445702245325 |
| chr8-27218728-27219229 | 3 | 0.995883475396051 |
| chr1-227517471-227517972 | 4 | 0.916626897245281 |
| chrX-111884969-111885470 | 8 | 0.993493681128311 |
| chr8-56175762-56176263 | 5 | 0.989458588425313 |
| chr9-70609955-70610456 | 3 | 0.953062332709917 |
| chr2-108834533-108835034 | 5 | 0.891020904670772 |
| chr7-65802202-65802703 | 2 | 0.917645726455962 |
| chr20-36402674-36403175 | 3 | 0.994648445337384 |
| chr13-70731851-70732352 | 4 | 0.8448935195764 |
| chr1-7507536-7508037 | 7 | 0.927382757728933 |
| chr7-98602989-98603490 | 4 | 0.973123798862735 |
| chr13-80113678-80114179 | 8 | 0.958938861505975 |
| chr13-57172001-57172502 | 8 | 0.99496973099361 |
| chr2-62109782-62110283 | 8 | 0.974563181166619 |

| | | |
|---|---|---|
| chr1-45447647-45448148 | 8 | 0.967245456908459 |
| chr10-18467169-18467670 | 6 | 0.987992542248998 |
| chr13-74879516-74880017 | 8 | 0.985799783018375 |
| chr20-19626797-19627298 | 8 | 0.986951272816455 |
| chr16-21353016-21353517 | 5 | 0.897825554823317 |
| chr1-118625735-118626236 | 6 | 0.990980740705316 |
| chr11-53534868-53535369 | 6 | 0.991206925481043 |
| chr5-35015372-35015873 | 6 | 0.867125431394257 |
| chr11-49272813-49273314 | 3 | 0.99593553545041 |
| chr15-20201686-20202187 | 5 | 0.938207645870034 |
| chr13-68569087-68569588 | 5 | 0.912558783283911 |
| chr7-33005784-33006285 | 8 | 0.83020174133677 |
| chr17-39663551-39664052 | 8 | 0.984376516020271 |
| chr10-71280560-71281061 | 8 | 0.993493681128311 |
| chr19-29611187-29611688 | 1 | 0.884093594601305 |
| chrX-128142130-128142631 | 4 | 0.92111284357392 |
| chr1-256079399-256079900 | 3 | 0.983475976351745 |
| chr18-60673749-60674250 | 8 | 0.993493681128311 |
| chr18-74180814-74181315 | 8 | 0.980497755231982 |
| chr16-77286719-77287220 | 8 | 0.993493681128311 |
| chrX-108630208-108630709 | 5 | 0.983919978840557 |
| chrX-106982716-106983217 | 3 | 0.984885256412346 |
| chr4-82482043-82482544 | 8 | 0.914020734678098 |
| chr9-92592822-92593323 | 4 | 0.951578832233771 |
| chr7-123657772-123658273 | 8 | 0.819887349487179 |
| chr6-140904936-140905437 | 8 | 0.995107883195904 |
| chr4-69715813-69716314 | 3 | 0.98626328286565 |
| chr15-43757233-43757734 | 2 | 0.957952710394278 |
| chr16-58055621-58056122 | 8 | 0.993493681128311 |
| chr14-103501734-103502235 | 8 | 0.96265147321898 |
| chr1-94312648-94313149 | 8 | 0.934150025274403 |
| chr6-129049961-129050462 | 8 | 0.993493681128311 |
| chr5-126327400-126327901 | 8 | 0.837655323456813 |
| chr6-18632019-18632520 | 8 | 0.891678956750728 |
| chr9-61768716-61769217 | 8 | 0.894444340200759 |
| chr11-2669548-2670049 | 1 | 0.906622915524458 |
| chr16-31162850-31163351 | 2 | 0.954171771364123 |
| chr8-43026230-43026731 | 8 | 0.993493681128311 |
| chr1-233371962-233372463 | 2 | 0.956191656948405 |
| chr3-41049370-41049871 | 3 | 0.991044573329043 |
| chr1-88358162-88358663 | 4 | 0.899476452765366 |
| chr1-164952523-164953024 | 8 | 0.960214679612158 |
| chr20-21147118-21147619 | 6 | 0.992638546640221 |

| | | |
|---|---|---|
| chr2-197991813-197992314 | 6 | 0.946249589441456 |
| chr2-93402723-93403224 | 5 | 0.95121433650134 |
| chr15-100049407-100049908 | 8 | 0.982461249756856 |
| chr19-50399835-50400336 | 8 | 0.938275124658361 |
| chr1-164347326-164347827 | 8 | 0.902315019398293 |
| chr17-77774263-77774764 | 8 | 0.990488961626945 |
| chr14-46428196-46428697 | 3 | 0.99593553545041 |
| chr9-38246302-38246803 | 3 | 0.996871674657255 |
| chr16-42912342-42912843 | 1 | 0.890479082205539 |
| chr1-18096149-18096650 | 8 | 0.956860504345036 |
| chr4-139840411-139840912 | 8 | 0.936086590969223 |
| chr8-99678102-99678603 | 8 | 0.841091886115438 |
| chr5-135062199-135062700 | 5 | 0.927901284401768 |
| chr5-133966114-133966615 | 4 | 0.855574310534113 |
| chr3-165873890-165874391 | 8 | 0.99015866091047 |
| chr2-21915444-21915945 | 6 | 0.961036160074254 |
| chr5-99161923-99162424 | 2 | 0.941712242123869 |
| chr3-101622237-101622738 | 6 | 0.974309777761852 |
| chr14-1426997-1427498 | 6 | 0.991823822561454 |
| chr16-74410007-74410508 | 3 | 0.997582740104186 |
| chr17-64327129-64327630 | 7 | 0.943987153664993 |
| chrX-122118838-122119339 | 2 | 0.868442280981622 |
| chr4-182325120-182325621 | 8 | 0.961451398860075 |
| chr16-23808787-23809288 | 3 | 0.903984619105343 |
| chr3-110534542-110535043 | 2 | 0.957600804921068 |
| chr18-61047433-61047934 | 8 | 0.920998375493262 |
| chr4-164281051-164281552 | 3 | 0.995795824730845 |
| chr12-11488102-11488603 | 2 | 0.940935431028049 |
| chr17-23690451-23690952 | 8 | 0.94393579454026 |
| chr4-71478747-71479248 | 7 | 0.855063898145054 |
| chr1-191906703-191907204 | 6 | 0.912501333152739 |
| chr11-74923388-74923889 | 4 | 0.940456782641671 |
| chr4-67315556-67316057 | 8 | 0.981361911654756 |
| chr17-80157528-80158029 | 5 | 0.844632383648066 |
| chr2-219177503-219178004 | 6 | 0.995014208257186 |
| chr20-38176626-38177127 | 6 | 0.965206349234653 |
| chr7-107299373-107299874 | 8 | 0.971483277040289 |
| chr8-55770023-55770524 | 5 | 0.994896570294365 |
| chr8-118166360-118166861 | 8 | 0.955223329198472 |
| chr15-31758405-31758906 | 8 | 0.853082516056401 |
| chr4-72248757-72249258 | 3 | 0.928834646291758 |
| chr15-49019433-49019934 | 8 | 0.950751429504288 |
| chr16-65737665-65738166 | 4 | 0.920685930249737 |

| | | |
|---|---|---|
| chr1-244017313-244017814 | 4 | 0.88286746060368 |
| chr14-60548962-60549463 | 6 | 0.953835632552801 |
| chr8-101650467-101650968 | 8 | 0.887061157239165 |
| chr14-83037106-83037607 | 1 | 0.930002596614739 |
| chr15-35133229-35133730 | 8 | 0.967791574826984 |
| chr11-44941556-44942057 | 8 | 0.875159450125619 |
| chr10-57239055-57239556 | 4 | 0.963269408763323 |
| chr8-27044885-27045386 | 6 | 0.943328374491521 |
| chr10-29235490-29235991 | 8 | 0.82786304016752 |
| chr11-69292708-69293209 | 7 | 0.892022551339259 |
| chr10-26866946-26867447 | 3 | 0.950745166538061 |
| chr13-6773952-6774453 | 3 | 0.99593553545041 |
| chr3-66857381-66857882 | 3 | 0.995343582645806 |
| chr18-73006590-73007091 | 8 | 0.931108968771309 |
| chr11-75604312-75604813 | 3 | 0.996831865635708 |
| chr13-5247994-5248495 | 3 | 0.995569029400898 |
| chr6-91312964-91313465 | 8 | 0.858275175728463 |
| chr9-4442539-4443040 | 8 | 0.912555398461278 |
| chr9-88788328-88788829 | 8 | 0.90500064766118 |
| chr11-5398524-5399025 | 6 | 0.993227812231235 |
| chr1-173843951-173844452 | 6 | 0.991206925481043 |
| chr13-70603488-70603989 | 8 | 0.993493681128311 |
| chr18-30765647-30766148 | 4 | 0.973637418506067 |
| chr14-33171931-33172432 | 2 | 0.87815679516264 |
| chr1-4327386-4327887 | 8 | 0.87771687400123 |
| chr5-60863566-60864067 | 5 | 0.993825481479112 |
| chr13-71566072-71566573 | 3 | 0.970066531763238 |
| chr2-96169727-96170228 | 3 | 0.99593553545041 |
| chr9-98891345-98891846 | 8 | 0.990354330143148 |
| chr18-32063353-32063854 | 8 | 0.960466344230041 |
| chr17-68513779-68514280 | 4 | 0.824092340610845 |
| chr3-110969923-110970424 | 6 | 0.991206925481043 |
| chr11-2737391-2737892 | 1 | 0.913450118388001 |
| chr9-18231982-18232483 | 8 | 0.994515303671343 |
| chr15-67418714-67419215 | 5 | 0.983526260070629 |
| chr20-43866154-43866655 | 4 | 0.866737506766527 |
| chr11-50945717-50946218 | 3 | 0.819356854517904 |
| chr3-102718434-102718935 | 8 | 0.914265022203164 |
| chr15-88208895-88209396 | 8 | 0.819783444448312 |
| chr4-128968797-128969298 | 3 | 0.995557592952688 |
| chr18-42815848-42816349 | 3 | 0.996591730891912 |
| chr2-219713098-219713599 | 5 | 0.83686188734727 |
| chr5-55314519-55315020 | 8 | 0.856731340350084 |

| | | |
|---|---|---|
| chr3-123174596-123175097 | 8 | 0.817971468962052 |
| chr18-15201656-15202157 | 8 | 0.994761147983332 |
| chr5-107974905-107975406 | 7 | 0.81936030558581 |
| chr19-4145259-4145760 | 5 | 0.987322638636864 |
| chr3-78268257-78268758 | 4 | 0.947473519173395 |
| chr1-6668733-6669234 | 4 | 0.955501126533583 |
| chrX-71211623-71212124 | 8 | 0.962976268305313 |
| chr2-95480435-95480936 | 8 | 0.994907782078608 |
| chr17-46585311-46585812 | 4 | 0.898905680422942 |
| chr1-45318588-45319089 | 5 | 0.892770234367184 |
| chr16-54579333-54579834 | 3 | 0.955974136871355 |
| chr7-124362027-124362528 | 1 | 0.932519624513477 |
| chr10-4437961-4438462 | 8 | 0.968198877510866 |
| chr2-58909417-58909918 | 3 | 0.996865726641406 |
| chr6-95849192-95849693 | 4 | 0.867960222891277 |
| chr3-156179268-156179769 | 5 | 0.991414469016434 |
| chr5-56964633-56965134 | 8 | 0.97452463478809 |
| chr6-51018711-51019212 | 8 | 0.995061397917813 |
| chr9-110255593-110256094 | 8 | 0.840172274578447 |
| chr15-10181727-10182228 | 8 | 0.993493681128311 |
| chr1-238609605-238610106 | 3 | 0.989726927254075 |
| chr3-147731906-147732407 | 3 | 0.99593553545041 |
| chr15-95352078-95352579 | 8 | 0.822604698600087 |
| chr17-27573460-27573961 | 4 | 0.94509230511995 |
| chr1-178963073-178963574 | 8 | 0.854214345726156 |
| chr2-62545002-62545503 | 8 | 0.936246660848877 |
| chr1-232119210-232119711 | 4 | 0.895738424914086 |
| chr5-128670273-128670774 | 3 | 0.997766248416143 |
| chr10-76324285-76324786 | 4 | 0.926490619581891 |
| chr14-57315399-57315900 | 3 | 0.834718053919528 |
| chr12-32346172-32346673 | 5 | 0.966724731084343 |
| chr10-44491451-44491952 | 8 | 0.961703713331127 |
| chr3-165519126-165519627 | 8 | 0.851894626293916 |
| chr8-114357379-114357880 | 3 | 0.997030136463289 |
| chr10-100180202-100180703 | 5 | 0.988788495091984 |
| chr4-177231940-177232441 | 7 | 0.897064705197928 |
| chr16-45359302-45359803 | 8 | 0.990010089471694 |
| chr4-169701419-169701920 | 8 | 0.913898980323728 |
| chr9-68706411-68706912 | 8 | 0.969499182607831 |
| chr13-70340947-70341448 | 5 | 0.946637106583585 |
| chr1-229472220-229472721 | 2 | 0.918889451518131 |
| chr9-79405310-79405811 | 8 | 0.995166801745748 |
| chrX-110111750-110112251 | 6 | 0.991206925481043 |

| | | |
|---|---|---|
| chr13-33347705-33348206 | 8 | 0.983647482044763 |
| chr1-98629123-98629624 | 7 | 0.967154336955723 |
| chr3-110832864-110833365 | 3 | 0.996048144414756 |
| chr1-22254663-22255164 | 6 | 0.967398611547904 |
| chr1-192431007-192431508 | 5 | 0.90461472209126 |
| chr2-142204374-142204875 | 8 | 0.950587474349039 |
| chr7-21880166-21880667 | 8 | 0.957228660389241 |
| chr11-76799564-76800065 | 5 | 0.935694359945105 |
| chr2-165386147-165386648 | 6 | 0.991206925481043 |
| chr4-141715464-141715965 | 4 | 0.966658738533666 |
| chrX-83230091-83230592 | 6 | 0.945934452668366 |
| chr5-99743174-99743675 | 3 | 0.985655423424945 |
| chr6-21947742-21948243 | 4 | 0.87098799645405 |
| chr1-199889710-199890211 | 4 | 0.885673542317076 |
| chr3-151663028-151663529 | 8 | 0.974652572292531 |
| chr2-54426790-54427291 | 1 | 0.871274287904229 |
| chr1-179995493-179995994 | 8 | 0.982694534568612 |
| chr5-68796151-68796652 | 5 | 0.958579202954809 |
| chr1-33131071-33131572 | 8 | 0.977008904478019 |
| chr2-138515367-138515868 | 4 | 0.970350771598634 |
| chr1-179486529-179487030 | 7 | 0.913281327215578 |
| chr13-33799673-33800174 | 3 | 0.99593553545041 |
| chr1-151341938-151342439 | 7 | 0.845384247223015 |
| chr13-11997040-11997541 | 8 | 0.993493681128311 |
| chr9-58218052-58218553 | 3 | 0.982513882372527 |
| chr6-7917163-7917664 | 3 | 0.988727135994957 |
| chr6-71992085-71992586 | 4 | 0.946579001148112 |
| chr13-106571324-106571825 | 8 | 0.957474733332047 |
| chr15-66557168-66557669 | 8 | 0.993493681128311 |
| chr18-70123110-70123611 | 4 | 0.865794980134203 |
| chr10-106022944-106023445 | 5 | 0.978696512043188 |
| chr4-150570624-150571125 | 1 | 0.925592325421913 |
| chr1-97458584-97459085 | 8 | 0.926753231526912 |
| chrX-40184653-40185154 | 6 | 0.969283456317594 |
| chr15-56495679-56496180 | 3 | 0.990858811948225 |
| chr7-51935042-51935543 | 5 | 0.913388519866206 |
| chr10-41754335-41754836 | 3 | 0.992436193702226 |
| chr20-34796475-34796976 | 4 | 0.945517093166306 |
| chr16-6093101-6093602 | 4 | 0.85126102223817 |
| chr9-4735101-4735602 | 2 | 0.855288692230253 |
| chr18-33493211-33493712 | 6 | 0.970309359518372 |
| chr13-67136189-67136690 | 8 | 0.993402142840724 |
| chr17-65746285-65746786 | 5 | 0.987725245388636 |

| | | |
|---|---|---|
| chr5-67609300-67609801 | 5 | 0.971726865366871 |
| chr14-68251518-68252019 | 8 | 0.968383566195616 |
| chr10-5867970-5868471 | 3 | 0.823405005795595 |
| chr1-2875215-2875716 | 1 | 0.826108837041461 |
| chr14-78068443-78068944 | 8 | 0.959124505514387 |
| chr15-100256239-100256740 | 8 | 0.864454370639738 |
| chr6-51130250-51130751 | 5 | 0.971735060705266 |
| chr16-77940-78441 | 2 | 0.932113996575259 |
| chr12-21808737-21809238 | 3 | 0.955301277549145 |
| chr7-23965179-23965680 | 8 | 0.993493681128311 |
| chr1-122533313-122533814 | 2 | 0.899742808886891 |
| chrX-98235052-98235553 | 8 | 0.98242553146473 |
| chr2-39558661-39559162 | 3 | 0.996796965992256 |
| chr2-171518415-171518916 | 8 | 0.959373426771584 |
| chr14-2327408-2327909 | 8 | 0.977129475184771 |
| chr8-60843670-60844171 | 8 | 0.91517290050211 |
| chr3-1108802-1109303 | 2 | 0.855807932750678 |
| chr16-20093391-20093892 | 2 | 0.942962383179382 |
| chr10-6665096-6665597 | 8 | 0.88875875542999 |
| chr10-86201272-86201773 | 5 | 0.984494559432575 |
| chr2-195479283-195479784 | 8 | 0.99805158658158 |
| chr14-43895666-43896167 | 3 | 0.991092764708295 |
| chr2-133914861-133915362 | 8 | 0.995230335984126 |
| chr8-54873993-54874494 | 8 | 0.944491886351464 |
| chr8-87477871-87478372 | 8 | 0.906393199760146 |
| chr13-12488875-12489376 | 3 | 0.869990051281619 |
| chr10-59980401-59980902 | 8 | 0.955943809372004 |
| chr10-8732218-8732719 | 1 | 0.872444771439234 |
| chr4-62183827-62184328 | 3 | 0.993543777965864 |
| chr2-163736484-163736985 | 2 | 0.861837061406266 |
| chrX-71761541-71762042 | 6 | 0.992375874254608 |
| chr6-10037026-10037527 | 4 | 0.829465877897553 |
| chr4-90094701-90095202 | 2 | 0.848981006760031 |
| chr4-109929333-109929834 | 6 | 0.885400599005611 |
| chr16-133066-133567 | 8 | 0.993493681128311 |
| chr5-57507173-57507674 | 8 | 0.977751140631537 |
| chr1-156892112-156892613 | 5 | 0.98144455427416 |
| chr19-27440262-27440763 | 8 | 0.888312121321391 |
| chr3-143850225-143850726 | 4 | 0.834420496061012 |
| chr5-143463093-143463594 | 8 | 0.993493681128311 |
| chr14-94705611-94706112 | 1 | 0.908778353364678 |
| chr7-19405315-19405816 | 8 | 0.995210684503526 |
| chr6-46701299-46701800 | 8 | 0.974480134318754 |

| | | |
|---|---|---|
| chr2-149872290-149872791 | 8 | 0.993493681128311 |
| chr1-150766459-150766960 | 8 | 0.965140843186118 |
| chr11-23839193-23839694 | 3 | 0.89000811593179 |
| chr9-89523331-89523832 | 6 | 0.991432246509226 |
| chr16-25502097-25502598 | 1 | 0.954511300560314 |
| chr16-37329757-37330258 | 3 | 0.99593553545041 |
| chr1-254121216-254121717 | 3 | 0.990881679305794 |
| chr6-14863486-14863987 | 8 | 0.860021216400864 |
| chr14-3762250-3762751 | 7 | 0.891579274970803 |
| chr2-71811336-71811837 | 3 | 0.988133923301963 |
| chr2-30363171-30363672 | 3 | 0.964567800506427 |
| chr15-78935341-78935842 | 6 | 0.993090155588391 |
| chr10-67362156-67362657 | 8 | 0.935452421933081 |
| chr6-3659573-3660074 | 3 | 0.988791733856131 |
| chr17-6526016-6526517 | 3 | 0.923151359515573 |
| chr1-22441329-22441830 | 4 | 0.950671774633594 |
| chr12-23245171-23245672 | 8 | 0.98620399524104 |
| chr9-99140247-99140748 | 8 | 0.99496285149544 |
| chr6-22273282-22273783 | 4 | 0.951726784868416 |
| chr8-86320318-86320819 | 6 | 0.974547064142819 |
| chr20-26035664-26036165 | 3 | 0.946236710924446 |
| chr2-22312184-22312685 | 8 | 0.859999166688695 |
| chr2-163407076-163407577 | 8 | 0.990296816888342 |
| chr10-68175613-68176114 | 8 | 0.973325753497726 |
| chr7-20978708-20979209 | 4 | 0.951025440334651 |
| chr2-173751525-173752026 | 8 | 0.966895324564967 |
| chr16-74236462-74236963 | 8 | 0.995175826035838 |
| chr4-110133904-110134405 | 3 | 0.921396713675932 |
| chr6-124498235-124498736 | 8 | 0.931148121508147 |
| chr3-121563911-121564412 | 1 | 0.847242651104827 |
| chr2-153321127-153321628 | 8 | 0.822934413721676 |
| chr13-1890043-1890544 | 3 | 0.99593553545041 |
| chr15-86887093-86887594 | 8 | 0.874623443775701 |
| chr18-74829179-74829680 | 7 | 0.93531019893931 |
| chr6-36058275-36058776 | 5 | 0.947755956982872 |
| chr3-108490134-108490635 | 8 | 0.995034636685775 |
| chr10-18048623-18049124 | 7 | 0.938388228170621 |
| chr15-61516847-61517348 | 3 | 0.979632316332613 |
| chr9-98687270-98687771 | 8 | 0.994009687288649 |
| chr3-59886725-59887226 | 7 | 0.913304248122462 |
| chr17-43803481-43803982 | 5 | 0.984309092820128 |
| chr5-143103017-143103518 | 3 | 0.997091612785809 |
| chr14-66732577-66733078 | 5 | 0.946076083135592 |

| | | |
|---|---|---|
| chr19-44699257-44699758 | 5 | 0.943951088491236 |
| chr19-28963140-28963641 | 4 | 0.974834217097081 |
| chr1-40325158-40325659 | 2 | 0.86723566830814 |
| chr16-4600262-4600763 | 5 | 0.885442987030226 |
| chr4-172812963-172813464 | 2 | 0.84255789653159 |
| chr14-3208133-3208634 | 8 | 0.877744503084654 |
| chr6-129256375-129256876 | 4 | 0.931728513029126 |
| chr8-25244505-25245006 | 3 | 0.995943497072293 |
| chr19-46415146-46415647 | 1 | 0.823019172620146 |
| chr4-92859030-92859531 | 8 | 0.994956795796151 |
| chr4-47370426-47370927 | 3 | 0.99593553545041 |
| chr2-110689662-110690163 | 8 | 0.980140293713799 |
| chr10-71187244-71187745 | 8 | 0.889256464875579 |
| chr20-49832360-49832861 | 3 | 0.99593553545041 |
| chr9-67834613-67835114 | 8 | 0.968252244875316 |
| chr10-9585228-9585729 | 8 | 0.864208406423823 |
| chr7-86447702-86448203 | 8 | 0.946015720429075 |
| chr3-154547642-154548143 | 8 | 0.99092891306373 |
| chr10-8950281-8950782 | 1 | 0.919096165353809 |
| chr1-257966776-257967277 | 4 | 0.8814177555358 |
| chr8-67440144-67440645 | 2 | 0.845837724340815 |
| chr18-73313389-73313890 | 8 | 0.966265793190274 |
| chr16-7621545-7622046 | 3 | 0.995727673066953 |
| chr4-116971272-116971773 | 8 | 0.975853374302242 |
| chr1-193400680-193401181 | 4 | 0.974367122645611 |
| chr6-52321693-52322194 | 8 | 0.993493681128311 |
| chr4-65478180-65478681 | 3 | 0.914942070828372 |
| chr11-46509496-46509997 | 4 | 0.911194821788698 |
| chr15-83438924-83439425 | 7 | 0.858688654416576 |
| chr2-242118395-242118896 | 4 | 0.875062737793033 |
| chrX-71248210-71248711 | 8 | 0.913712986612822 |
| chr8-23008695-23009196 | 8 | 0.993493681128311 |
| chr8-85196662-85197163 | 8 | 0.993493681128311 |
| chr7-95628394-95628895 | 8 | 0.83251866808912 |
| chr3-22351125-22351626 | 8 | 0.974215243018511 |
| chr4-159611609-159612110 | 8 | 0.979817770196919 |
| chr15-93418364-93418865 | 6 | 0.88577486900175 |
| chr2-235119254-235119755 | 8 | 0.830128118414439 |
| chr6-137486848-137487349 | 3 | 0.961342913578137 |
| chr5-137482518-137483019 | 8 | 0.855862363201582 |
| chr5-57359594-57360095 | 3 | 0.996922288339169 |
| chr6-14202881-14203382 | 8 | 0.951704776117362 |
| chr5-97690836-97691337 | 8 | 0.949289732784389 |

| | | |
|---|---|---|
| chr4-18195226-18195727 | 8 | 0.992953234103745 |
| chr16-66207369-66207870 | 8 | 0.993493681128311 |
| chr2-206916994-206917495 | 3 | 0.995510883790946 |
| chr1-91450791-91451292 | 3 | 0.933387967082256 |
| chr6-3324252-3324753 | 4 | 0.951820412605953 |
| chr5-39242812-39243313 | 3 | 0.976392146532891 |
| chr20-14140511-14141012 | 4 | 0.959934217736185 |
| chr12-15709974-15710475 | 5 | 0.951949886922021 |
| chr6-65895002-65895503 | 8 | 0.924615058641597 |
| chr10-87405800-87406301 | 8 | 0.991318610719527 |
| chr9-33782216-33782717 | 3 | 0.99593553545041 |
| chr4-104611116-104611617 | 8 | 0.979739755715088 |
| chr16-65223892-65224393 | 8 | 0.993493681128311 |
| chr3-103201776-103202277 | 3 | 0.996704215588976 |
| chr15-87354768-87355269 | 6 | 0.898813217102652 |
| chr3-17804059-17804560 | 8 | 0.994661622959023 |
| chr2-185183065-185183566 | 8 | 0.976884084835797 |
| chr6-21133732-21134233 | 8 | 0.832183228071789 |
| chr1-24721586-24722087 | 8 | 0.968754448297362 |
| chr1-226400944-226401445 | 8 | 0.880657553811163 |
| chr14-52705510-52706011 | 6 | 0.965629185388743 |
| chr3-18888237-18888738 | 8 | 0.835779567283923 |
| chrX-71711673-71712174 | 3 | 0.995648953226912 |
| chr1-189945183-189945684 | 4 | 0.973091925551148 |
| chr14-58467330-58467831 | 4 | 0.914662811529206 |
| chr10-47544695-47545196 | 8 | 0.957398182974581 |
| chr1-55653931-55654432 | 8 | 0.89585167552664 |
| chr9-8285493-8285994 | 3 | 0.994539451330046 |
| chr7-66220923-66221424 | 8 | 0.980210670213305 |
| chr2-87089209-87089710 | 3 | 0.992693910859628 |
| chr15-76231156-76231657 | 6 | 0.991055199891481 |
| chr4-59212925-59213426 | 8 | 0.854033990614708 |
| chr4-181561706-181562207 | 2 | 0.871336182310358 |
| chr4-43072394-43072895 | 8 | 0.884034335650142 |
| chr2-193948698-193949199 | 6 | 0.992895175265721 |
| chr7-75305176-75305677 | 3 | 0.9722836755921 |
| chr17-31408701-31409202 | 3 | 0.990801389229321 |
| chr10-4390414-4390915 | 8 | 0.926654901072173 |
| chr2-91900558-91901059 | 8 | 0.993493681128311 |
| chr2-28077337-28077838 | 6 | 0.986909408540346 |
| chr1-135121026-135121527 | 4 | 0.97109122958868 |
| chrX-68298585-68299086 | 2 | 0.918799325862769 |
| chr4-11177083-11177584 | 8 | 0.932414838422222 |

| | | |
|---|---|---|
| chr17-8746963-8747464 | 3 | 0.995935535545041 |
| chr4-96233943-96234444 | 3 | 0.930601058345971 |
| chr14-3063826-3064327 | 8 | 0.994891915155051 |
| chr6-126038097-126038598 | 8 | 0.989881179890593 |
| chr10-55974303-55974804 | 4 | 0.930579464661376 |
| chr1-223225970-223226471 | 8 | 0.991930409937198 |
| chr5-73889839-73890340 | 8 | 0.993493681128311 |
| chr19-729211-729712 | 2 | 0.82695688724847 |
| chr8-117223475-117223976 | 8 | 0.984763148012734 |
| chr6-5550860-5551361 | 4 | 0.957488434397787 |
| chr1-115560661-115561162 | 3 | 0.972746958730194 |
| chr12-34897306-34897807 | 8 | 0.965100670757381 |
| chr8-43881802-43882303 | 7 | 0.963757604502649 |
| chr5-12613185-12613686 | 8 | 0.98579814520045 |
| chr5-82826613-82827114 | 3 | 0.995935535545041 |
| chr4-113992863-113993364 | 3 | 0.995987402952235 |
| chr20-41810316-41810817 | 8 | 0.993077654322549 |
| chr14-28908569-28909070 | 3 | 0.8875093250367 |
| chr15-10194349-10194850 | 3 | 0.994262320067466 |
| chr15-53144275-53144776 | 8 | 0.959482060953984 |
| chr7-31436802-31437303 | 8 | 0.988291595429858 |
| chr14-97396916-97397417 | 8 | 0.98309765184785 |
| chrX-30149505-30150006 | 6 | 0.993186389538233 |
| chr2-115905087-115905588 | 6 | 0.983070503106323 |
| chr1-188215998-188216499 | 8 | 0.949992402937294 |
| chr3-33811272-33811773 | 6 | 0.980323150613748 |
| chr1-25607382-25607883 | 5 | 0.989655404552859 |
| chr4-33586004-33586505 | 3 | 0.856130912732897 |
| chr17-65197237-65197738 | 5 | 0.990765005425504 |
| chr2-77245962-77246463 | 5 | 0.890513491457804 |
| chr15-2124263-2124764 | 3 | 0.996867529630684 |
| chr6-9291032-9291533 | 8 | 0.955206555489478 |
| chr8-66607581-66608082 | 1 | 0.862884427303956 |
| chr5-23864026-23864527 | 8 | 0.939454659153985 |
| chr4-153628111-153628612 | 8 | 0.915872496763546 |
| chr1-56480669-56481170 | 4 | 0.930759109607864 |
| chr6-45911553-45912054 | 8 | 0.994491488212076 |
| chr8-30067439-30067940 | 8 | 0.994479372782305 |
| chr3-2248418-2248919 | 6 | 0.991206925481043 |
| chr19-1462747-1463248 | 8 | 0.897471223296499 |
| chr2-223687088-223687589 | 2 | 0.951838456967612 |
| chr16-23041117-23041618 | 8 | 0.993471612339338 |
| chr17-53333958-53334459 | 6 | 0.942506967903599 |

| | | |
|---|---|---|
| chr5-71172761-71173262 | 6 | 0.825187554081109 |
| chr9-61793533-61794034 | 3 | 0.996853298180617 |
| chr18-69167461-69167962 | 8 | 0.945569414967921 |
| chr8-52608497-52608998 | 6 | 0.99021403961603 |
| chr15-56710964-56711465 | 6 | 0.991206925481043 |
| chr11-44082808-44083309 | 6 | 0.990782405508966 |
| chr14-82787757-82788258 | 2 | 0.891327830904313 |
| chr1-180409115-180409616 | 3 | 0.884276206993514 |
| chr15-46926857-46927358 | 3 | 0.849873380409685 |
| chr19-53647250-53647751 | 8 | 0.990644724420188 |
| chr13-23336817-23337318 | 5 | 0.950154793759032 |
| chr9-97130163-97130664 | 8 | 0.990523709720869 |
| chr1-86514142-86514643 | 4 | 0.8685826724307 |
| chr2-242864709-242865210 | 8 | 0.911062137822779 |
| chr8-121473243-121473744 | 8 | 0.962676346195234 |
| chr5-47526817-47527318 | 4 | 0.819944837892071 |
| chr20-31301958-31302459 | 8 | 0.968310493205179 |
| chr15-62730269-62730770 | 8 | 0.816650391892204 |
| chr3-46821615-46822116 | 5 | 0.983919978840557 |
| chr5-10594852-10595353 | 8 | 0.878064800360596 |
| chr16-42115054-42115555 | 8 | 0.956884891804396 |
| chr17-77205120-77205621 | 8 | 0.993493681128311 |
| chr19-2326722-2327223 | 5 | 0.919796658747213 |
| chrX-83243798-83244299 | 6 | 0.991206925481043 |
| chr2-22715539-22716040 | 5 | 0.863664307733575 |
| chr11-51669048-51669549 | 3 | 0.952606266324842 |
| chr19-31761402-31761903 | 3 | 0.99490558475601 |
| chr8-105063231-105063732 | 8 | 0.995234229711778 |
| chr15-62836675-62837176 | 8 | 0.923032205285643 |
| chr1-194898724-194899225 | 8 | 0.982635956318607 |
| chr8-78769450-78769951 | 8 | 0.909474700843782 |
| chr19-8017922-8018423 | 3 | 0.992244100454722 |
| chr1-142744891-142745392 | 1 | 0.937843051110163 |
| chr8-8365815-8366316 | 3 | 0.996341548906442 |
| chr8-38659193-38659694 | 3 | 0.99593553545041 |
| chr9-88782843-88783344 | 8 | 0.845585289755391 |
| chr2-35211806-35212307 | 4 | 0.867992336998212 |
| chr10-43728306-43728807 | 8 | 0.995163770865393 |
| chr6-88644578-88645079 | 3 | 0.981160551467928 |
| chr6-124968314-124968815 | 6 | 0.984448445703706 |
| chr6-70359457-70359958 | 1 | 0.890872294898688 |
| chr10-95174777-95175278 | 8 | 0.993493681128311 |
| chr13-63117136-63117637 | 2 | 0.904958888088977 |

| | | |
|---|---|---|
| chr2-119514741-119515242 | 5 | 0.9760792599328 |
| chr9-101715016-101715517 | 6 | 0.989605513362158 |
| chr3-8477334-8477835 | 8 | 0.986347505719807 |
| chr3-50245005-50245506 | 7 | 0.883955660577533 |
| chr8-95262267-95262768 | 5 | 0.984508769610119 |
| chr15-37116050-37116551 | 8 | 0.930536887567952 |
| chr14-86883619-86884120 | 6 | 0.990034039029389 |
| chr2-834925-835426 | 4 | 0.838201555327702 |
| chr13-91505149-91505650 | 4 | 0.927105707742472 |
| chr12-12284267-12284768 | 8 | 0.862923205303186 |
| chr4-173787144-173787645 | 5 | 0.941528490429264 |
| chr3-88102739-88103240 | 8 | 0.923330655574993 |
| chr3-88962012-88962513 | 7 | 0.847795197309427 |
| chr14-12680789-12681290 | 8 | 0.991608603930658 |
| chr18-6466806-6467307 | 4 | 0.865164191791134 |
| chrX-42639176-42639677 | 8 | 0.99491056144807 |
| chr2-121640018-121640519 | 6 | 0.92109430514335 |
| chr16-6202825-6203326 | 8 | 0.956653474417542 |
| chr5-163533615-163534116 | 8 | 0.995043853381837 |
| chr18-26834170-26834671 | 4 | 0.836554893041947 |
| chr11-11314148-11314649 | 8 | 0.995204293906726 |
| chr1-141189664-141190165 | 3 | 0.847827370280887 |
| chr1-172446939-172447440 | 8 | 0.980711705717898 |
| chr3-52842722-52843223 | 8 | 0.988344746679323 |
| chr2-7683378-7683879 | 5 | 0.991213435300111 |
| chr18-2580254-2580755 | 8 | 0.928172950063971 |
| chr5-25206692-25207193 | 4 | 0.891523963463975 |
| chr11-16599119-16599620 | 3 | 0.986917329160385 |
| chr9-81235958-81236459 | 4 | 0.840610107149553 |
| chr15-11509329-11509830 | 8 | 0.873418356448317 |
| chr5-78675073-78675574 | 8 | 0.96833278573129 |
| chr3-127310469-127310970 | 8 | 0.872086173793692 |
| chr2-210903912-210904413 | 5 | 0.995513578754499 |
| chr6-98009902-98010403 | 8 | 0.948009230341207 |
| chr1-125033017-125033518 | 3 | 0.997220973740355 |
| chr4-84233845-84234346 | 3 | 0.83386160464576 |
| chr5-6059606-6060107 | 8 | 0.831765880481794 |
| chr2-152846536-152847037 | 8 | 0.958322033657431 |
| chr5-8161537-8162038 | 3 | 0.830679022740117 |
| chr5-77652226-77652727 | 6 | 0.954368510456647 |
| chr5-45084546-45085047 | 8 | 0.971860621464501 |
| chr6-5250619-5251120 | 3 | 0.981945747349821 |
| chr5-26629775-26630276 | 6 | 0.991206925481043 |

| | | |
|---|---|---|
| chr4-85570412-85570913 | 6 | 0.93533611493906 |
| chr17-23667511-23668012 | 8 | 0.993493681128311 |
| chr13-75874117-75874618 | 8 | 0.818860038738579 |
| chr10-78399633-78400134 | 6 | 0.993958311561746 |
| chr7-116857263-116857764 | 1 | 0.977651057508749 |
| chr16-50107580-50108081 | 3 | 0.878819190141328 |
| chr2-223012989-223013490 | 6 | 0.839146560247264 |
| chr7-88025568-88026069 | 8 | 0.979401793830385 |
| chr2-37049559-37050060 | 5 | 0.823185249076186 |
| chr7-11369036-11369537 | 4 | 0.831258352131658 |
| chr13-86984657-86985158 | 8 | 0.857282261329719 |
| chr14-87054899-87055400 | 8 | 0.993493681128311 |
| chrX-66435667-66436168 | 3 | 0.990371215149255 |
| chr1-236057583-236058084 | 4 | 0.930425678291854 |
| chr2-244697067-244697568 | 3 | 0.97274543499458 |
| chr2-73307752-73308253 | 8 | 0.925145962201218 |
| chr6-109890190-109890691 | 3 | 0.992708381557872 |
| chr1-84759709-84760210 | 5 | 0.837209585306282 |
| chr1-191425523-191426024 | 8 | 0.993493681128311 |
| chr14-72491469-72491970 | 6 | 0.992983863468712 |
| chr2-162763895-162764396 | 8 | 0.844021029594473 |
| chr1-109654151-109654652 | 8 | 0.94832460789081 |
| chr9-91766203-91766704 | 8 | 0.98478640220626 |
| chr5-132674246-132674747 | 4 | 0.869061310682502 |
| chr4-134039501-134040002 | 8 | 0.963161961129857 |
| chr2-194941737-194942238 | 2 | 0.919486754717326 |
| chr14-100044347-100044848 | 8 | 0.994771347336685 |
| chr9-40018841-40019342 | 8 | 0.994111805170119 |
| chr5-1237116-1237617 | 6 | 0.822183170326065 |
| chr4-40535825-40536326 | 8 | 0.814759244965479 |
| chr1-14987028-14987529 | 8 | 0.898034129153566 |
| chr3-114951128-114951629 | 3 | 0.956291497628401 |
| chr8-105512129-105512630 | 6 | 0.991206925481043 |
| chr4-84415885-84416386 | 2 | 0.897843756814573 |
| chr12-12909591-12910092 | 8 | 0.988252803824617 |
| chr9-3953569-3954070 | 8 | 0.968576884866033 |
| chr2-235758937-235759438 | 3 | 0.84938182502383 |
| chr9-70690647-70691148 | 8 | 0.95213511077616 |
| chr4-119187421-119187922 | 3 | 0.850232221347427 |
| chr17-59865288-59865789 | 3 | 0.871110159299202 |
| chr11-29561984-29562485 | 8 | 0.89085412088395 |
| chr19-40590181-40590682 | 6 | 0.990924577090702 |
| chr5-60311620-60312121 | 2 | 0.929813129247681 |

| | | |
|---|---|---|
| chr19-42169234-42169735 | 3 | 0.996744845615851 |
| chr5-76147044-76147545 | 3 | 0.821206321105969 |
| chr2-112532952-112533453 | 6 | 0.983396470028521 |
| chr4-113251126-113251627 | 3 | 0.996603669476879 |
| chr13-94946962-94947463 | 8 | 0.995244 |
| chr12-1413776-1414277 | 6 | 0.991206925481043 |
| chr13-96114092-96114593 | 2 | 0.875280935947978 |
| chr1-130366951-130367452 | 6 | 0.997336283477119 |
| chr3-153819368-153819869 | 7 | 0.920297897820044 |
| chr14-41485034-41485535 | 5 | 0.982429710134031 |
| chr17-10859131-10859632 | 8 | 0.974873661138759 |
| chr4-135897943-135898444 | 6 | 0.992675870299369 |
| chr2-13561577-13562078 | 2 | 0.837761669885398 |
| chr3-14644886-14645387 | 8 | 0.82349933996481 |
| chr13-95072375-95072876 | 1 | 0.947280203847471 |
| chr5-12933571-12934072 | 8 | 0.882672713886998 |
| chr3-163401378-163401879 | 7 | 0.839942314801067 |
| chr3-24010079-24010580 | 6 | 0.931990341696917 |
| chr14-85798409-85798910 | 6 | 0.996339428680782 |
| chr13-104716163-104716664 | 8 | 0.993493681128311 |
| chr8-65876804-65877305 | 8 | 0.957910741650864 |
| chr18-72082058-72082559 | 8 | 0.992133796446922 |
| chr16-10422557-10423058 | 8 | 0.881635023107117 |
| chr6-17043714-17044215 | 1 | 0.943022540275202 |
| chr14-64609876-64610377 | 8 | 0.913603854612354 |
| chr9-96492600-96493101 | 6 | 0.989432131883435 |
| chr20-13447189-13447690 | 5 | 0.985438349704936 |
| chr4-179453017-179453518 | 4 | 0.861664507849651 |
| chr2-128737058-128737559 | 4 | 0.94900347327216 |
| chr1-172882613-172883114 | 7 | 0.827321097032243 |
| chr10-69416452-69416953 | 4 | 0.904219414093182 |
| chr16-46200846-46201347 | 3 | 0.970178053735848 |
| chr4-25525620-25526121 | 5 | 0.820492852421039 |
| chr17-614505-615006 | 3 | 0.995926215555887 |
| chrX-15301769-15302270 | 8 | 0.993493681128311 |
| chr1-190725225-190725726 | 8 | 0.975274532757292 |
| chr15-38040605-38041106 | 5 | 0.991305489051261 |
| chr14-67027135-67027636 | 5 | 0.918341657487569 |
| chrX-99310169-99310670 | 8 | 0.960996243929046 |
| chr13-48345445-48345946 | 1 | 0.898430414331128 |
| chr2-79962788-79963289 | 8 | 0.96957492447643 |
| chr13-38201069-38201570 | 5 | 0.972683683244801 |
| chr17-77749640-77750141 | 2 | 0.838917230198925 |

| | | |
|---|---|---|
| chr10-66589854-66590355 | 8 | 0.938550687383188 |
| chr6-131824237-131824738 | 8 | 0.978620474705918 |
| chr7-14540224-14540725 | 8 | 0.992417647705264 |
| chr3-123169554-123170055 | 1 | 0.91758227159864 |
| chr4-5604742-5605243 | 8 | 0.979800328683014 |
| chr8-87293402-87293903 | 8 | 0.920730150892475 |
| chr2-194502761-194503262 | 7 | 0.821244515860554 |
| chr10-9625094-9625595 | 8 | 0.892593858583674 |
| chr7-29199745-29200246 | 5 | 0.986510538573507 |
| chr20-22164072-22164573 | 6 | 0.976345077723991 |
| chr7-84025837-84026338 | 2 | 0.852326308391653 |
| chr1-12326093-12326594 | 4 | 0.865953589236253 |
| chr4-136863535-136864036 | 3 | 0.969830615116071 |
| chr5-71480098-71480599 | 8 | 0.961444084695341 |
| chr3-60846228-60846729 | 5 | 0.8637784019425 |
| chr18-1528364-1528865 | 8 | 0.886705319513574 |
| chr7-31456938-31457439 | 3 | 0.995571144325591 |
| chr2-50854769-50855270 | 6 | 0.907579172376097 |
| chr1-215599912-215600413 | 3 | 0.995073831306338 |
| chr11-75316517-75317018 | 8 | 0.949076831497151 |
| chr19-42663493-42663994 | 5 | 0.99203305246032 |
| chr17-31231984-31232485 | 6 | 0.945393962724597 |
| chr1-86300592-86301093 | 8 | 0.865307918820707 |
| chr13-50535832-50536333 | 8 | 0.97811078483433 |
| chr4-57358579-57359080 | 4 | 0.873693219280909 |
| chr10-48404537-48405038 | 3 | 0.996348566729756 |
| chr3-62929894-62930395 | 3 | 0.995455849403848 |
| chr8-55511930-55512431 | 3 | 0.974368617291433 |
| chr2-35339393-35339894 | 8 | 0.979310609330573 |
| chr3-31150338-31150839 | 5 | 0.964108520939574 |
| chr16-36962901-36963402 | 3 | 0.99593553545041 |
| chr1-234011356-234011857 | 5 | 0.897997448197282 |
| chr15-39302854-39303355 | 8 | 0.992219815844232 |
| chr10-23927021-23927522 | 8 | 0.995053181408562 |
| chr3-81057355-81057856 | 8 | 0.985466989176177 |
| chr19-30895923-30896424 | 2 | 0.870571516819478 |
| chr2-121304120-121304621 | 7 | 0.92838327673684 |
| chr5-86753076-86753577 | 8 | 0.972985689397166 |
| chr6-44453423-44453924 | 3 | 0.996714862599703 |
| chr3-96821847-96822348 | 3 | 0.997038669087664 |
| chr5-119339077-119339578 | 8 | 0.981753728826486 |
| chr5-51835196-51835697 | 3 | 0.971790434506697 |
| chr8-105787364-105787865 | 8 | 0.892802054134844 |

| | | |
|---|---|---|
| chr8-6519872-6520373 | 8 | 0.815629656116876 |
| chr19-3088920-3089421 | 6 | 0.966937170036774 |
| chr16-46335042-46335543 | 8 | 0.967291927532276 |
| chr2-150321681-150322182 | 8 | 0.944025719673639 |
| chr7-48495808-48496309 | 3 | 0.99593553545041 |
| chr6-92167038-92167539 | 4 | 0.901616602445917 |
| chr6-119637787-119638288 | 5 | 0.980909612884822 |
| chr12-25337384-25337885 | 6 | 0.957976403330951 |
| chr9-15960279-15960780 | 8 | 0.882357292420889 |
| chr1-206228843-206229344 | 8 | 0.850306345047025 |
| chr3-33748462-33748963 | 6 | 0.929707632438025 |
| chr10-94547303-94547804 | 4 | 0.837282471071029 |
| chr1-124182799-124183300 | 7 | 0.849864271168378 |
| chr1-88510972-88511473 | 2 | 0.819810326370168 |
| chr15-53006366-53006867 | 2 | 0.92185223970541 |
| chr8-33428221-33428722 | 2 | 0.867750506344077 |
| chr20-6085302-6085803 | 3 | 0.889877699998417 |
| chr2-28907569-28908070 | 5 | 0.972814192243255 |
| chr1-164316589-164317090 | 5 | 0.922452935116286 |
| chr16-2125708-2126209 | 8 | 0.819420974412691 |
| chr14-85692126-85692627 | 6 | 0.986418811226527 |
| chr9-4903872-4904373 | 8 | 0.981142515486303 |
| chr15-1776849-1777350 | 8 | 0.99348330163782 |
| chr12-36688363-36688864 | 8 | 0.922223294510903 |
| chr6-7135591-7136092 | 8 | 0.954985154618426 |
| chr1-139416765-139417266 | 6 | 0.97381257047088 |
| chr7-90778787-90779288 | 8 | 0.895363316479176 |
| chr17-40663186-40663687 | 8 | 0.963344557132475 |
| chr8-59476210-59476711 | 8 | 0.994869793623336 |
| chr9-44886779-44887280 | 3 | 0.992338974664563 |
| chr12-44547211-44547712 | 5 | 0.833747854757047 |
| chr19-38091391-38091892 | 5 | 0.97996216684369 |
| chr16-58975040-58975541 | 1 | 0.90396144318634 |
| chr18-27372630-27373131 | 4 | 0.904707922098136 |
| chr6-71065687-71066188 | 7 | 0.891593515979136 |
| chr11-47368837-47369338 | 5 | 0.964968770115253 |
| chr19-51355407-51355908 | 5 | 0.850969301764829 |
| chr12-35884047-35884548 | 8 | 0.984420561760903 |
| chr1-156069200-156069701 | 1 | 0.927892954538012 |
| chr1-216595354-216595855 | 8 | 0.978609013272416 |
| chr13-44653541-44654042 | 4 | 0.916702048452512 |
| chrX-139928960-139929461 | 3 | 0.980215194766972 |
| chr10-15184622-15185123 | 8 | 0.988259045737347 |

| | | |
|---|---|---|
| chr1-233196886-233197387 | 3 | 0.989632628955175 |
| chr4-176790475-176790976 | 8 | 0.836500295173405 |
| chr12-22094221-22094722 | 1 | 0.966007590751134 |
| chr17-34935015-34935516 | 8 | 0.922109466067546 |
| chr4-158951025-158951526 | 3 | 0.998002814373725 |
| chr3-129126596-129127097 | 8 | 0.994524207053016 |
| chr3-142476015-142476516 | 6 | 0.948898754276407 |
| chr1-164127847-164128348 | 8 | 0.992515448635031 |
| chr17-39873476-39873977 | 7 | 0.816752776358263 |
| chr8-120120199-120120700 | 8 | 0.98188280683341 |
| chr18-57177333-57177834 | 7 | 0.828459211917319 |
| chr12-10963682-10964183 | 5 | 0.900443206227359 |
| chr1-78510239-78510740 | 8 | 0.995197436600237 |
| chr5-102614846-102615347 | 3 | 0.996897669285024 |
| chr17-26557299-26557800 | 6 | 0.997073656394841 |
| chr1-14557231-14557732 | 3 | 0.961345905623584 |
| chr18-43610821-43611322 | 3 | 0.99652668405983 |
| chr18-78145476-78145977 | 2 | 0.852399723525571 |
| chr18-56100075-56100576 | 3 | 0.999146007261213 |
| chr4-90039665-90040166 | 8 | 0.887009701559411 |
| chr10-88643172-88643673 | 3 | 0.82965597745088 |
| chr17-70713685-70714186 | 6 | 0.991206925481043 |
| chr5-105172590-105173091 | 3 | 0.993611542406818 |
| chr3-43729491-43729992 | 6 | 0.944889021351365 |
| chr17-42360731-42361232 | 2 | 0.881644878319939 |
| chr13-45606755-45607256 | 8 | 0.922412043751873 |
| chr17-67016148-67016649 | 8 | 0.932267639063259 |
| chr1-23230644-23231145 | 4 | 0.973726670181469 |
| chr2-150622006-150622507 | 8 | 0.993493681128311 |
| chr5-32801743-32802244 | 5 | 0.902024503910351 |
| chr10-75711024-75711525 | 3 | 0.974193879784105 |
| chr12-7646565-7647066 | 5 | 0.939243406971174 |
| chr17-48186576-48187077 | 2 | 0.896828201610237 |
| chr2-60958786-60959287 | 8 | 0.995193311868286 |
| chr2-201013222-201013723 | 3 | 0.989483363996485 |
| chr17-60580477-60580978 | 8 | 0.858355082070864 |
| chr4-80773357-80773858 | 2 | 0.874939110194043 |
| chr13-55815536-55816037 | 5 | 0.973741115495802 |
| chr1-250434473-250434974 | 6 | 0.974403522724788 |
| chrX-61046956-61047457 | 8 | 0.889492921293271 |
| chr14-72526584-72527085 | 6 | 0.840636518305952 |
| chr7-115134388-115134889 | 8 | 0.915587882595627 |
| chr2-171759453-171759954 | 8 | 0.983528396190102 |

| | | |
|---|---|---|
| chr5-111930088-111930589 | 8 | 0.99246155260815 |
| chr17-589951-590452 | 8 | 0.993493681128311 |
| chr2-108727707-108728208 | 6 | 0.977690887464001 |
| chr19-30787719-30788220 | 8 | 0.904760111176485 |
| chr9-85653252-85653753 | 8 | 0.936595045984706 |
| chr4-69968313-69968814 | 6 | 0.991206925481043 |
| chr8-123001016-123001517 | 5 | 0.975361470842746 |
| chr17-24865918-24866419 | 2 | 0.875625034341716 |
| chr16-57299218-57299719 | 8 | 0.905674599194844 |
| chr11-50084709-50085210 | 3 | 0.996887678610735 |
| chr8-12763725-12764226 | 8 | 0.88208085096594 |
| chr7-17492819-17493320 | 1 | 0.858581902113957 |
| chr1-100696636-100697137 | 8 | 0.946748133460375 |
| chr6-119359760-119360261 | 8 | 0.909778111812644 |
| chr2-101081383-101081884 | 4 | 0.874976148051897 |
| chr16-51341890-51342391 | 8 | 0.995729161269532 |
| chr1-119509846-119510347 | 8 | 0.974001098543858 |
| chr19-50554973-50555474 | 5 | 0.933989047170867 |
| chr4-9306972-9307473 | 5 | 0.96189248014176 |
| chr1-202603435-202603936 | 4 | 0.945658187588986 |
| chr12-27917419-27917920 | 2 | 0.892952384618974 |
| chr14-31166542-31167043 | 4 | 0.860397005411782 |
| chr7-111066920-111067421 | 4 | 0.915968575684896 |
| chr11-55202595-55203096 | 8 | 0.983110535560889 |
| chr17-42351061-42351562 | 2 | 0.862856037607599 |
| chr15-65130080-65130581 | 5 | 0.993530962702812 |
| chr2-242966456-242966957 | 8 | 0.979517360469485 |
| chr1-44567575-44568076 | 8 | 0.984143038575926 |
| chr2-35574746-35575247 | 2 | 0.926234357633049 |
| chr1-45803123-45803624 | 5 | 0.989417051473667 |
| chr12-26630003-26630504 | 8 | 0.956146686355906 |
| chr15-85148382-85148883 | 3 | 0.99593553545041 |
| chr15-39875218-39875719 | 8 | 0.886949169580508 |
| chr1-22890257-22890758 | 5 | 0.920637059876872 |
| chr2-161609485-161609986 | 6 | 0.99309732987552 |
| chr8-115340641-115341142 | 8 | 0.993493681128311 |
| chr10-95532977-95533478 | 7 | 0.906141142934086 |
| chr16-63546167-63546668 | 5 | 0.973787189578993 |
| chr3-145136318-145136819 | 8 | 0.995202897960661 |
| chr16-80166954-80167455 | 8 | 0.996481322476569 |
| chr5-152695691-152696192 | 6 | 0.991206925481043 |
| chr5-35496008-35496509 | 8 | 0.853941047833967 |
| chr4-157380403-157380904 | 8 | 0.822215684502485 |

| | | |
|---|---|---|
| chr15-39864740-39865241 | 2 | 0.952238697974421 |
| chr1-53926494-53926995 | 1 | 0.932717096079614 |
| chrX-3008228-3008729 | 3 | 0.99593553545041 |
| chr2-246642827-246643328 | 8 | 0.961814893499491 |
| chr10-49895574-49896075 | 6 | 0.972391601116976 |
| chr4-180826507-180827008 | 3 | 0.966355284470738 |
| chr1-215972146-215972647 | 4 | 0.818102856639061 |
| chr15-39005052-39005553 | 8 | 0.850957481612897 |
| chr5-45904403-45904904 | 3 | 0.996646097629639 |
| chr1-144221665-144222166 | 8 | 0.865124653556269 |
| chr11-57500111-57500612 | 4 | 0.933212174226919 |
| chr5-126392156-126392657 | 8 | 0.937327597891219 |
| chr13-105346564-105347065 | 7 | 0.844491645703085 |
| chr8-111010043-111010544 | 6 | 0.987305174859642 |
| chr13-98880172-98880673 | 7 | 0.921541358619628 |
| chr2-44595570-44596071 | 2 | 0.913886635133107 |
| chr16-1609618-1610119 | 8 | 0.948679403096563 |
| chr5-82518975-82519476 | 6 | 0.991206925481043 |
| chr9-35027876-35028377 | 6 | 0.991206925481043 |
| chr14-35410386-35410887 | 5 | 0.933161007467352 |
| chr5-130537360-130537861 | 8 | 0.955655565553998 |
| chr9-73003955-73004456 | 8 | 0.988601960334747 |
| chr5-11199783-11200284 | 5 | 0.982967866530617 |
| chr1-29161910-29162411 | 5 | 0.99112414717585 |
| chr5-3607837-3608338 | 8 | 0.831138026363577 |
| chr9-8847160-8847661 | 3 | 0.99593553545041 |
| chr14-88713065-88713566 | 4 | 0.971438453673808 |
| chr3-125147278-125147779 | 8 | 0.990141814285239 |
| chrX-73911599-73912100 | 3 | 0.99593553545041 |
| chr14-89531438-89531939 | 3 | 0.995814890010581 |
| chr5-126235181-126235682 | 3 | 0.886760844014749 |
| chr20-32229237-32229738 | 4 | 0.892670249810906 |
| chr13-46952436-46952937 | 5 | 0.990492806978931 |
| chr17-3193425-3193926 | 3 | 0.995069911857502 |
| chr2-247463739-247464240 | 7 | 0.856876238885853 |
| chr14-32262077-32262578 | 5 | 0.82088481574062 |
| chr11-39736934-39737435 | 3 | 0.957021189103856 |
| chr17-690318-690819 | 8 | 0.925586142359824 |
| chr8-89370115-89370616 | 2 | 0.820090343764255 |
| chr4-64383046-64383547 | 8 | 0.842944 |
| chr15-46352343-46352844 | 8 | 0.896409499647566 |
| chr17-25102474-25102975 | 3 | 0.99593553545041 |
| chr18-80233331-80233832 | 8 | 0.834902818974981 |

| | | |
|---|---|---|
| chr1-215846920-215847421 | 3 | 0.890384080370158 |
| chr16-26513-27014 | 4 | 0.88090809633648 |
| chr4-77350626-77351127 | 8 | 0.890727202551623 |
| chr14-27460674-27461175 | 3 | 0.996060409585683 |
| chr14-62804794-62805295 | 7 | 0.844329382364942 |
| chr13-33006780-33007281 | 8 | 0.985187898352153 |
| chr8-57708904-57709405 | 8 | 0.822867357794603 |
| chr1-50111703-50112204 | 8 | 0.964693065097217 |
| chr16-48713159-48713660 | 3 | 0.988855388925902 |
| chr2-41019587-41020088 | 2 | 0.936045750934605 |
| chr1-208032501-208033002 | 3 | 0.99593553545041 |
| chr9-12122513-12123014 | 7 | 0.879656072911428 |
| chr7-127494996-127495497 | 8 | 0.943027074619172 |
| chr3-62617140-62617641 | 8 | 0.983124638059788 |
| chr5-89018714-89019215 | 3 | 0.994915213462049 |
| chr3-143988773-143989274 | 6 | 0.968260091883899 |
| chr6-65077225-65077726 | 3 | 0.997887815547146 |
| chr14-42687904-42688405 | 8 | 0.816660255267824 |
| chrX-24344630-24345131 | 8 | 0.816115777137235 |
| chr11-54195739-54196240 | 3 | 0.996510975761972 |
| chr7-87205198-87205699 | 4 | 0.925333222276857 |
| chr16-25411229-25411730 | 8 | 0.917476803267772 |
| chr5-87705926-87706427 | 8 | 0.94876493834167 |
| chr14-16850602-16851103 | 8 | 0.859636811306347 |
| chr18-44586387-44586888 | 8 | 0.97067602990382 |
| chr8-98588433-98588934 | 4 | 0.910655735207689 |
| chrX-16082806-16083307 | 5 | 0.880361354713359 |
| chr3-120806445-120806946 | 2 | 0.869124883608171 |
| chr2-23297878-23298379 | 1 | 0.834178108409838 |
| chr2-216583238-216583739 | 3 | 0.988538136386109 |
| chr15-84016673-84017174 | 3 | 0.99593553545041 |
| chr20-13448414-13448915 | 5 | 0.967966478028449 |
| chr18-35369527-35370028 | 6 | 0.991767565469566 |
| chr3-43067009-43067510 | 5 | 0.906967864877279 |
| chr8-101593810-101594311 | 8 | 0.993493681128311 |
| chr5-151020868-151021369 | 8 | 0.993493681128311 |
| chr8-47923185-47923686 | 8 | 0.969700441703556 |
| chr4-180176548-180177049 | 8 | 0.965650460653345 |
| chr13-95796853-95797354 | 8 | 0.969743460103684 |
| chr14-34945668-34946169 | 8 | 0.993493681128311 |
| chrX-105055302-105055803 | 3 | 0.953415603684044 |
| chr13-86390540-86391041 | 8 | 0.933107565341857 |
| chr2-113815371-113815872 | 4 | 0.895915445543234 |

| | | |
|---|---|---|
| chr9-54759175-54759676 | 8 | 0.910388358238851 |
| chr19-14678374-14678875 | 8 | 0.94200264725111 |
| chr17-57467850-57468351 | 2 | 0.931149388681455 |
| chr4-50840115-50840616 | 8 | 0.993493681128311 |
| chr6-24592846-24593347 | 8 | 0.99337915137491 |
| chr1-29138950-29139451 | 8 | 0.969880403831804 |
| chr1-173866489-173866990 | 8 | 0.881515090013429 |
| chr18-40945203-40945704 | 1 | 0.93882320184518 |
| chr1-199054852-199055353 | 8 | 0.992391218881192 |
| chr13-22990346-22990847 | 8 | 0.988713468300707 |
| chr20-8569551-8570052 | 5 | 0.881145061125583 |
| chr10-13940498-13940999 | 8 | 0.968550844747143 |
| chr1-135272881-135273382 | 4 | 0.819850809094477 |
| chr15-49864577-49865078 | 1 | 0.89066139928747 |
| chr5-163803390-163803891 | 6 | 0.94090361184336 |
| chr5-45533095-45533596 | 8 | 0.98896138294571 |
| chr1-22041783-22042284 | 8 | 0.98246292699651 |
| chr4-136437642-136438143 | 8 | 0.965364064742348 |
| chr7-75610508-75611009 | 2 | 0.867577516591571 |
| chr9-16431547-16432048 | 8 | 0.993493681128311 |
| chr4-129753460-129753961 | 5 | 0.933060818317792 |
| chr5-108780819-108781320 | 3 | 0.987460608300338 |
| chr6-31041625-31042126 | 8 | 0.993563337851252 |
| chr14-13957054-13957555 | 3 | 0.987135561038095 |
| chr6-97743048-97743549 | 6 | 0.989412798000674 |
| chr4-150758361-150758862 | 1 | 0.869761261860686 |
| chr4-175355126-175355627 | 3 | 0.996215260964126 |
| chr10-1151812-1152313 | 3 | 0.906940250194867 |
| chr20-14804271-14804772 | 6 | 0.986174706065736 |
| chr14-70451762-70452263 | 3 | 0.815296901321348 |
| chr8-103055126-103055627 | 8 | 0.839651799966973 |
| chr8-11209326-11209827 | 8 | 0.977203504237778 |
| chr9-60068431-60068932 | 8 | 0.950023709265947 |
| chr4-39400390-39400891 | 8 | 0.815088795025796 |
| chr2-161666720-161667221 | 6 | 0.978633159458853 |
| chr20-13644804-13645305 | 8 | 0.968854863582188 |
| chr14-84103866-84104367 | 6 | 0.991206925481043 |
| chr1-136351646-136352147 | 1 | 0.912230613970737 |
| chr10-102221084-102221585 | 8 | 0.962104163262891 |
| chr4-127400169-127400670 | 4 | 0.930253427984047 |
| chr3-128287235-128287736 | 3 | 0.962567975366466 |
| chr8-107696233-107696734 | 5 | 0.983919978840557 |
| chr17-15826263-15826764 | 5 | 0.901303254727492 |

| | | |
|---|---|---|
| chr3-168745966-168746467 | 4 | 0.888333186522336 |
| chr11-72385849-72386350 | 5 | 0.982763105921473 |
| chr4-161495026-161495527 | 8 | 0.941872871851117 |
| chr4-63498080-63498581 | 3 | 0.995263531827676 |
| chr6-100749110-100749611 | 3 | 0.994500986477376 |
| chr8-116106166-116106667 | 5 | 0.913753040627348 |
| chr2-215414539-215415040 | 6 | 0.983889454880938 |
| chr6-3818107-3818608 | 5 | 0.956704072429278 |
| chr1-222089510-222090011 | 6 | 0.917029548437863 |
| chr2-134626537-134627038 | 5 | 0.961084763663175 |
| chr7-19110193-19110694 | 5 | 0.911385852182038 |
| chr8-67870674-67871175 | 5 | 0.973017647597863 |
| chr4-77994672-77995173 | 6 | 0.996330863733551 |
| chr7-110668318-110668819 | 7 | 0.832259527702977 |
| chr6-60553415-60553916 | 3 | 0.93395541627891 |
| chr20-11157122-11157623 | 2 | 0.819585232875676 |
| chr4-155967326-155967827 | 1 | 0.862906449889038 |
| chr6-30893345-30893846 | 2 | 0.862369337054718 |
| chr17-35763985-35764486 | 8 | 0.984499085198888 |
| chr7-92589904-92590405 | 6 | 0.990175614333467 |
| chr4-121151246-121151747 | 8 | 0.987403298836433 |
| chr5-44113318-44113819 | 8 | 0.893774175742643 |
| chr8-111986296-111986797 | 3 | 0.909893462776071 |
| chr5-48390036-48390537 | 2 | 0.950679498259472 |
| chr6-83824218-83824719 | 5 | 0.91169319892446 |
| chr5-33904049-33904550 | 6 | 0.99022570807672 |
| chr6-138067210-138067711 | 8 | 0.945569547300351 |
| chr2-188868519-188869020 | 8 | 0.992796706976959 |
| chr17-76300400-76300901 | 4 | 0.920381472836067 |
| chr9-61172699-61173200 | 4 | 0.920808806662187 |
| chr13-93390740-93391241 | 8 | 0.964527859770144 |
| chr7-15802421-15802922 | 2 | 0.900823755002024 |
| chr2-191748364-191748865 | 7 | 0.921062822122665 |
| chr20-11438350-11438851 | 6 | 0.974475476474296 |
| chr3-112075987-112076488 | 5 | 0.856849885552653 |
| chr8-37101869-37102370 | 6 | 0.99137744551631 |
| chr3-76907973-76908474 | 8 | 0.953392033411097 |
| chr9-81686306-81686807 | 3 | 0.996710831377659 |
| chr1-120257339-120257840 | 5 | 0.99166077970967 |
| chr13-58456185-58456686 | 3 | 0.99593553545041 |
| chr14-98973828-98974329 | 5 | 0.960532393584386 |
| chr6-83747484-83747985 | 2 | 0.921487337602171 |
| chr2-26501589-26502090 | 3 | 0.986197674642573 |

| | | |
|---|---|---|
| chr5-150256900-150257401 | 8 | 0.837766482110514 |
| chr11-69870071-69870572 | 7 | 0.859490184923228 |
| chr13-39064821-39065322 | 8 | 0.960308509540759 |
| chr10-60274178-60274679 | 8 | 0.931791519070525 |
| chr8-41776579-41777080 | 8 | 0.828955659717013 |
| chr15-44371726-44372227 | 8 | 0.881589637565909 |
| chr8-111400440-111400941 | 8 | 0.938945232593068 |
| chr11-6011952-6012453 | 5 | 0.988856809720223 |
| chrX-11636611-11637112 | 8 | 0.995014151360691 |
| chr15-49411904-49412405 | 1 | 0.855009307712643 |
| chr10-12903611-12904112 | 8 | 0.855510886992012 |
| chr17-80401111-80401612 | 8 | 0.903790526908776 |
| chr8-81137998-81138499 | 5 | 0.890370046976653 |
| chr2-229587837-229588338 | 3 | 0.99593553545041 |
| chr9-91830100-91830601 | 8 | 0.993493681128311 |
| chr9-103172819-103173320 | 1 | 0.898937136989929 |
| chr6-14480769-14481270 | 8 | 0.858141401459728 |
| chr13-10649100-10649601 | 3 | 0.894727086175543 |
| chr11-31402814-31403315 | 8 | 0.856950564537066 |
| chr6-17494424-17494925 | 3 | 0.982433607000375 |
| chr6-69800405-69800906 | 7 | 0.947539402344369 |
| chr6-64370375-64370876 | 3 | 0.987081624824659 |
| chr4-96018593-96019094 | 8 | 0.947981789254389 |
| chr8-56871188-56871689 | 1 | 0.921674552513984 |
| chr11-65513915-65514416 | 8 | 0.933246855408251 |
| chrX-32741020-32741521 | 5 | 0.991527454886902 |
| chr7-105428686-105429187 | 8 | 0.925306282078303 |
| chr13-65812250-65812751 | 7 | 0.822045419249417 |
| chr2-69136836-69137337 | 3 | 0.93316533924331 |
| chr19-23026992-23027493 | 4 | 0.919255084660906 |
| chr12-31569835-31570336 | 7 | 0.911096067571906 |
| chr14-104384335-104384836 | 1 | 0.826316131265811 |
| chr2-234536612-234537113 | 3 | 0.944154110027139 |
| chr19-37866830-37867331 | 8 | 0.993493681128311 |
| chr4-88673122-88673623 | 8 | 0.827205869521609 |
| chr14-72557551-72558052 | 8 | 0.983942625632151 |
| chr11-42612457-42612958 | 3 | 0.86842168165242 |
| chr14-46451453-46451954 | 8 | 0.995249000224298 |
| chr15-73661834-73662335 | 6 | 0.991206925481043 |
| chr11-41519376-41519877 | 3 | 0.988308749304143 |
| chr2-39178335-39178836 | 6 | 0.996337142365213 |
| chr16-50972966-50973467 | 5 | 0.871866528560033 |
| chr19-53489006-53489507 | 7 | 0.887885331864316 |

| | | |
|---|---|---|
| chrX-152295315-152295816 | 4 | 0.853027607072072 |
| chr16-505035-505536 | 4 | 0.852838184928882 |
| chr4-121040761-121041262 | 8 | 0.819387804143406 |
| chr20-52558630-52559131 | 3 | 0.99593553545041 |
| chr13-31351303-31351804 | 6 | 0.985340506589382 |
| chr2-216144183-216144684 | 8 | 0.837042576103271 |
| chr3-100623530-100624031 | 8 | 0.855716809765389 |
| chr14-9373377-9373878 | 8 | 0.991246211282547 |
| chr4-9255203-9255704 | 3 | 0.927172269021023 |
| chr1-32668250-32668751 | 6 | 0.939637203681475 |
| chr1-127825566-127826067 | 8 | 0.936931211221281 |
| chr3-48705191-48705692 | 6 | 0.828290722220431 |
| chr9-93731451-93731952 | 8 | 0.995043870509517 |
| chr14-77326715-77327216 | 4 | 0.936468671177608 |
| chr15-21182826-21183327 | 4 | 0.934151888069704 |
| chr9-47628534-47629035 | 7 | 0.946296164558397 |
| chr1-186379322-186379823 | 8 | 0.993493681128311 |
| chr15-92549107-92549608 | 3 | 0.994562573944604 |
| chr2-216917688-216918189 | 8 | 0.920985530667339 |
| chr3-107655097-107655598 | 4 | 0.817384912008822 |
| chr13-77284547-77285048 | 6 | 0.992672139683928 |
| chr15-31356055-31356556 | 8 | 0.920426661904164 |
| chr17-26459503-26460004 | 8 | 0.979285259376247 |
| chr7-96075442-96075943 | 8 | 0.96088710329243 |
| chr4-83251841-83252342 | 8 | 0.995127405170235 |
| chr1-260160875-260161376 | 6 | 0.996319445787319 |
| chr19-34210951-34211452 | 8 | 0.9619288471817 |
| chr1-142552856-142553357 | 1 | 0.887928176890446 |
| chr10-57530729-57531230 | 3 | 0.998818451354097 |
| chr5-124951178-124951679 | 2 | 0.95099797647012 |
| chr5-96126951-96127452 | 8 | 0.880036542579009 |
| chr1-182271900-182272401 | 8 | 0.932742203933483 |
| chr13-20267924-20268425 | 3 | 0.99593553545041 |
| chr20-15067798-15068299 | 3 | 0.99593553545041 |
| chr3-38914940-38915441 | 5 | 0.882654781771743 |
| chr18-68350543-68351044 | 6 | 0.994888074588938 |
| chr2-13966458-13966959 | 8 | 0.935056476743373 |
| chr11-78445021-78445522 | 5 | 0.967086614865653 |
| chr1-108714573-108715074 | 8 | 0.911471552584084 |
| chr7-43184539-43185040 | 3 | 0.99593553545041 |
| chr11-51548133-51548634 | 3 | 0.98898698001194 |
| chr7-18870285-18870786 | 8 | 0.910604230612418 |
| chr10-105965899-105966400 | 8 | 0.887658412484858 |

| | | |
|---|---|---|
| chr5-10494688-10495189 | 6 | 0.988198831294333 |
| chr10-6006155-6006656 | 8 | 0.833750410520389 |
| chr2-44979322-44979823 | 8 | 0.829540551920319 |
| chr1-10973172-10973673 | 1 | 0.910451022802323 |
| chr5-166320572-166321073 | 4 | 0.920638117437901 |
| chr2-112627545-112628046 | 3 | 0.993792343896668 |
| chr1-138507869-138508370 | 3 | 0.962512639256681 |
| chr1-28636977-28637478 | 8 | 0.876913113556486 |
| chr15-31318260-31318761 | 5 | 0.99116011950599 |
| chr18-81454328-81454829 | 6 | 0.992548560616391 |
| chr5-118995664-118996165 | 8 | 0.956437275934314 |
| chr20-17210693-17211194 | 5 | 0.843597523336885 |
| chr14-74579917-74580418 | 8 | 0.958477256585569 |
| chr18-56792199-56792700 | 5 | 0.952596224295758 |
| chr4-75331320-75331821 | 3 | 0.994440740908229 |
| chr6-20048330-20048831 | 8 | 0.872177375248415 |
| chr2-184249153-184249654 | 4 | 0.951032447619158 |
| chr20-8887366-8887867 | 5 | 0.865797855259252 |
| chr13-83473103-83473604 | 6 | 0.991930389319293 |
| chr13-13574894-13575395 | 8 | 0.934643195960212 |
| chr14-102296888-102297389 | 1 | 0.898462316325352 |
| chr17-6549822-6550323 | 2 | 0.927419111477223 |
| chr7-55455307-55455808 | 6 | 0.984963266783115 |
| chr17-5406613-5407114 | 3 | 0.997761930083336 |
| chr16-24341739-24342240 | 3 | 0.992668510367237 |
| chr15-45870231-45870732 | 2 | 0.95633343897054 |
| chr4-7365348-7365849 | 8 | 0.976213609082578 |
| chr4-170174171-170174672 | 8 | 0.994917653790396 |
| chr4-57910145-57910646 | 8 | 0.995228837685494 |
| chr9-67528481-67528982 | 5 | 0.940354037413768 |
| chrY-8655290-8655791 | 6 | 0.981048045757954 |
| chr13-32311324-32311825 | 5 | 0.854238604014078 |
| chr18-21930303-21930804 | 8 | 0.960512387489447 |
| chr3-89799185-89799686 | 2 | 0.886846615308695 |
| chrX-55146015-55146516 | 6 | 0.991206925481043 |
| chr12-22233501-22234002 | 8 | 0.905674759887435 |
| chr7-94441067-94441568 | 3 | 0.99593553545041 |
| chr7-111485098-111485599 | 8 | 0.933043105565899 |
| chr4-53330193-53330694 | 5 | 0.986716013519756 |
| chr2-73521151-73521652 | 6 | 0.991206925481043 |
| chr3-168164994-168165495 | 4 | 0.846863401848356 |
| chr7-40510393-40510894 | 1 | 0.918115946681984 |
| chr1-240863006-240863507 | 8 | 0.979995720002132 |

| | | |
|---|---|---|
| chr4-97556063-97556564 | 3 | 0.932878444627028 |
| chr19-9311918-9312419 | 5 | 0.93814564691164 |
| chr10-37779646-37780147 | 8 | 0.860951117768033 |
| chr18-56222520-56223021 | 5 | 0.94880759250941 |
| chr10-27618122-27618623 | 3 | 0.912995676872721 |
| chr14-34389409-34389910 | 8 | 0.988543087678444 |
| chr2-23295399-23295900 | 1 | 0.879706741767592 |
| chr8-42569910-42570411 | 8 | 0.848299502600563 |
| chr10-31148393-31148894 | 2 | 0.891397221999095 |
| chr19-906896-907397 | 8 | 0.970769268412211 |
| chr3-161109922-161110423 | 8 | 0.932487626291786 |
| chr4-144034366-144034867 | 6 | 0.990171277505846 |
| chr18-59771850-59772351 | 2 | 0.919227235332621 |
| chr3-155551067-155551568 | 8 | 0.958056212293321 |
| chr1-181526961-181527462 | 8 | 0.948345054645983 |
| chr9-25686007-25686508 | 3 | 0.989278874152851 |
| chr13-4286562-4287063 | 6 | 0.991206925481043 |
| chr17-14998690-14999191 | 8 | 0.964818528879928 |
| chr6-94080271-94080772 | 1 | 0.912930326478307 |
| chr1-152083039-152083540 | 8 | 0.993493681128311 |
| chr15-14547036-14547537 | 5 | 0.990839802779756 |
| chr2-35839027-35839528 | 4 | 0.85477845913497 |
| chr2-164606925-164607426 | 8 | 0.986498604646493 |
| chr16-6177889-6178390 | 8 | 0.93605386060258 |
| chr6-20864994-20865495 | 8 | 0.864740526457785 |
| chr9-5004165-5004666 | 2 | 0.915781369531348 |
| chr19-45573279-45573780 | 5 | 0.985594574960164 |
| chr18-9332469-9332970 | 6 | 0.990192745777369 |
| chr2-78631451-78631952 | 7 | 0.81826078494681 |
| chr1-179488610-179489111 | 7 | 0.882709313776719 |
| chr2-45373268-45373769 | 8 | 0.817486168962466 |
| chr13-75057333-75057834 | 8 | 0.839125569804956 |
| chr1-207607455-207607956 | 4 | 0.86725841284943 |
| chr1-96868153-96868654 | 3 | 0.99593553545041 |
| chr8-94428278-94428779 | 6 | 0.98599799882795 |
| chr2-241588953-241589454 | 8 | 0.838127141514918 |
| chr14-7332560-7333061 | 3 | 0.997239747082839 |
| chr13-41910050-41910551 | 3 | 0.99593553545041 |
| chr2-220519737-220520238 | 7 | 0.923047326737225 |
| chr10-9418884-9419385 | 3 | 0.99593553545041 |
| chr7-122570765-122571266 | 6 | 0.989748376001819 |
| chr20-25450656-25451157 | 6 | 0.941607885402903 |
| chr3-54726655-54727156 | 3 | 0.996794910505938 |

| | | |
|---|---|---|
| chr12-23765322-23765823 | 3 | 0.994831973199122 |
| chr6-7221207-7221708 | 3 | 0.997236964726726 |
| chr1-172474330-172474831 | 4 | 0.861663728161586 |
| chr12-37958687-37959188 | 8 | 0.989284702537235 |
| chr16-47787078-47787579 | 8 | 0.911540907355711 |
| chr6-101573828-101574329 | 5 | 0.827992037916861 |
| chr16-67069428-67069929 | 3 | 0.950174225047732 |
| chr8-64800579-64801080 | 3 | 0.995699716297182 |
| chr9-32280581-32281082 | 6 | 0.991946349872771 |
| chr1-176996426-176996927 | 3 | 0.998089922563146 |
| chr4-9132732-9133233 | 3 | 0.990971155700942 |
| chr12-32491524-32492025 | 5 | 0.949235498629815 |
| chr5-165605796-165606297 | 8 | 0.945688420640893 |
| chr16-69686096-69686597 | 5 | 0.899650513017784 |
| chr9-95882750-95883251 | 8 | 0.936897306093641 |
| chr14-41321207-41321708 | 6 | 0.944425379901259 |
| chr11-43514654-43515155 | 8 | 0.857508433329388 |
| chr20-41342351-41342852 | 8 | 0.93639726374587 |
| chr11-14796288-14796789 | 4 | 0.967996067969413 |
| chr2-167989262-167989763 | 8 | 0.869413231428845 |
| chr7-125846046-125846547 | 5 | 0.990179482849814 |
| chr7-69170984-69171485 | 2 | 0.89980761942735 |
| chr4-37805475-37805976 | 3 | 0.996115669236296 |
| chr13-28623853-28624354 | 3 | 0.983573731480347 |
| chr10-84636424-84636925 | 2 | 0.832672824230663 |
| chr19-46618476-46618977 | 1 | 0.890421155659589 |
| chr16-3113709-3114210 | 8 | 0.993493681128311 |
| chr7-76284201-76284702 | 5 | 0.942605299956544 |
| chr6-20285907-20286408 | 8 | 0.981621075407669 |
| chr10-91522387-91522888 | 8 | 0.879993511124513 |
| chr8-106693305-106693806 | 8 | 0.993493681128311 |
| chr6-34658465-34658966 | 8 | 0.984052407291257 |
| chr11-80657400-80657901 | 8 | 0.816403456872672 |
| chr2-13548445-13548946 | 6 | 0.991206925481043 |
| chr2-31139086-31139587 | 8 | 0.842766070917189 |
| chr3-152130540-152131041 | 8 | 0.987242152395463 |
| chr12-5256546-5257047 | 8 | 0.989668076508519 |
| chr1-121514959-121515460 | 8 | 0.990169646439054 |
| chr1-145305378-145305879 | 5 | 0.95042928115202 |
| chr10-87941380-87941881 | 8 | 0.979428125362937 |
| chr10-57377244-57377745 | 8 | 0.849577923504526 |
| chr14-1632825-1633326 | 8 | 0.893627685788025 |
| chrX-2655126-2655627 | 8 | 0.987200004598468 |

| | | |
|---|---|---|
| chr6-46718190-46718691 | 8 | 0.815628984993655 |
| chr14-3746155-3746656 | 6 | 0.991206925481043 |
| chr2-41009936-41010437 | 8 | 0.878760867387912 |
| chr2-45199005-45199506 | 8 | 0.964015345623687 |
| chr13-39695379-39695880 | 3 | 0.993889749337108 |
| chr1-217023948-217024449 | 5 | 0.978980394497587 |
| chr5-92136236-92136737 | 4 | 0.830138355877701 |
| chr1-225230220-225230721 | 8 | 0.993272515440937 |
| chr6-60218779-60219280 | 6 | 0.862292619135473 |
| chr1-122310817-122311318 | 3 | 0.981733669445948 |
| chrX-89230067-89230568 | 8 | 0.989625186102707 |
| chr13-17226838-17227339 | 4 | 0.951502022783725 |
| chr15-32646594-32647095 | 3 | 0.994993142420906 |
| chr2-53748486-53748987 | 8 | 0.8336949798019 |
| chr1-49650160-49650661 | 8 | 0.982581082113992 |
| chr12-40548142-40548643 | 5 | 0.91650457433096 |
| chr2-71629995-71630496 | 8 | 0.931859956742874 |
| chr5-139851542-139852043 | 8 | 0.920297426868637 |
| chr9-6747077-6747578 | 2 | 0.899681202922302 |
| chr13-43466998-43467499 | 5 | 0.986207083709542 |
| chr14-31296487-31296988 | 8 | 0.892349775792988 |
| chr9-57749917-57750418 | 8 | 0.97522828611875 |
| chr9-36764834-36765335 | 8 | 0.917768770030677 |
| chr13-69049689-69050190 | 8 | 0.917033766141273 |
| chr17-16229216-16229717 | 8 | 0.894436200071979 |
| chr4-115094525-115095026 | 1 | 0.900174975965092 |
| chr2-224819986-224820487 | 5 | 0.989889385943677 |
| chr11-71123774-71124275 | 8 | 0.822634172101068 |
| chr4-46674154-46674655 | 8 | 0.993493681128311 |
| chr8-88264791-88265292 | 4 | 0.862667442542278 |
| chr13-42567414-42567915 | 8 | 0.988212117308343 |
| chr15-68101848-68102349 | 3 | 0.957719310510451 |
| chr1-42398826-42399327 | 8 | 0.911396799902815 |
| chr8-15313323-15313824 | 5 | 0.852326510784187 |
| chr1-188859565-188860066 | 8 | 0.83969911234377 |
| chr7-106659708-106660209 | 6 | 0.995114130727675 |
| chr1-187542666-187543167 | 4 | 0.867075419503532 |
| chr17-85764932-85765433 | 3 | 0.988404241089061 |
| chr15-70817107-70817608 | 3 | 0.99593553545041 |
| chr9-47845223-47845724 | 3 | 0.99593553545041 |
| chr3-70293819-70294320 | 8 | 0.872362261325825 |
| chr1-252892117-252892618 | 4 | 0.967752022774567 |
| chr8-14010879-14011380 | 8 | 0.973739444351804 |

| | | |
|---|---|---|
| chr15-97932895-97933396 | 8 | 0.93701672988977 |
| chr9-35997793-35998294 | 8 | 0.920247970846283 |
| chrX-13206711-13207212 | 3 | 0.995593252507411 |
| chr14-71079800-71080301 | 8 | 0.993493681128311 |
| chr6-93947863-93948364 | 8 | 0.966815754974677 |
| chr14-43818222-43818723 | 5 | 0.970662658960266 |
| chr10-74961361-74961862 | 6 | 0.992787117747926 |
| chr8-122609784-122610285 | 8 | 0.913186678156369 |
| chr16-2115371-2115872 | 8 | 0.911104274999032 |
| chr19-1925783-1926284 | 3 | 0.922506979490305 |
| chr4-11179384-11179885 | 8 | 0.995204481378256 |
| chr5-151315933-151316434 | 8 | 0.9526853677333 |
| chr19-53981826-53982327 | 3 | 0.998392150222913 |
| chr1-181310319-181310820 | 8 | 0.933039650288755 |
| chr6-89896042-89896543 | 8 | 0.84557411173477 |
| chr9-107172705-107173206 | 5 | 0.957085012011397 |
| chr11-45912067-45912568 | 3 | 0.974367596108151 |
| chr9-51198673-51199174 | 8 | 0.993493681128311 |
| chr6-52442258-52442759 | 8 | 0.99221430179574 |
| chr2-127775393-127775894 | 6 | 0.830014439722252 |
| chrX-28063985-28064486 | 3 | 0.996155341520084 |
| chr3-139632930-139633431 | 8 | 0.973526486116083 |
| chr17-7484721-7485222 | 6 | 0.985980673297415 |
| chr13-37677153-37677654 | 8 | 0.991697314237118 |
| chr1-226743690-226744191 | 8 | 0.993160480928308 |
| chr10-39732571-39733072 | 8 | 0.894547009593897 |
| chr8-51608477-51608978 | 8 | 0.821337913265892 |
| chr10-79946293-79946794 | 2 | 0.861322876087839 |
| chr2-246723852-246724353 | 8 | 0.940412903748087 |
| chr1-243504684-243505185 | 4 | 0.958057884740778 |
| chr9-26954239-26954740 | 8 | 0.990671523168571 |
| chr14-86054590-86055091 | 8 | 0.994244561475692 |
| chr3-11233008-11233509 | 8 | 0.951959073429498 |
| chr16-61219778-61220279 | 8 | 0.991632676231729 |
| chr2-182711347-182711848 | 8 | 0.907620872435535 |
| chr10-7910794-7911295 | 3 | 0.99593553545041 |
| chr2-190393119-190393620 | 3 | 0.98284511093377 |
| chr17-68897063-68897564 | 8 | 0.840046384105774 |
| chr5-123867714-123868215 | 8 | 0.86673222803364 |
| chr1-171624963-171625464 | 6 | 0.982937277371897 |
| chr1-180123178-180123679 | 2 | 0.839065084337736 |
| chr7-91507529-91508030 | 8 | 0.892346912584752 |
| chr9-38837936-38838437 | 4 | 0.932495734304051 |

| | | |
|---|---|---|
| chr7-40348812-40349313 | 5 | 0.897948278184355 |
| chr7-76103812-76104313 | 3 | 0.967322500903879 |
| chr1-176757916-176758417 | 3 | 0.935182735691962 |
| chr2-117765408-117765909 | 8 | 0.848380606911352 |
| chr19-48181555-48182056 | 3 | 0.890844759049371 |
| chr14-50488226-50488727 | 3 | 0.931358982558854 |
| chr6-18435082-18435583 | 2 | 0.941551525868542 |
| chr3-65262369-65262870 | 8 | 0.972877842374481 |
| chr9-91015259-91015760 | 3 | 0.818561329195017 |
| chr8-79225819-79226320 | 8 | 0.985387398576634 |
| chr5-52549852-52550353 | 3 | 0.996806509369414 |
| chr6-2105148-2105649 | 3 | 0.99593553545041 |
| chr4-113820484-113820985 | 4 | 0.966474950238444 |
| chr3-41486247-41486748 | 5 | 0.983931979478917 |
| chr15-32515071-32515572 | 6 | 0.993092942266012 |
| chr1-173289703-173290204 | 8 | 0.849741606713939 |
| chrX-102670412-102670913 | 6 | 0.991206925481043 |
| chr15-53527970-53528471 | 8 | 0.948484228883906 |
| chr18-49352527-49353028 | 5 | 0.983534625869341 |
| chr13-36137991-36138492 | 2 | 0.875930099057894 |
| chr3-121360934-121361435 | 8 | 0.970968035589652 |
| chr5-71727184-71727685 | 5 | 0.997129347615888 |
| chr9-27796589-27797090 | 8 | 0.867673582420084 |
| chr17-33468133-33468634 | 2 | 0.959983564231473 |
| chr14-40227731-40228232 | 6 | 0.942484731463397 |
| chr7-55682203-55682704 | 8 | 0.845004363972267 |
| chr3-151449946-151450447 | 8 | 0.993493681128311 |
| chr1-90518548-90519049 | 8 | 0.96743686161987 |
| chr8-106254302-106254803 | 8 | 0.989696289380255 |
| chr5-165152317-165152818 | 2 | 0.881825434239602 |
| chr1-98591626-98592127 | 7 | 0.930662243391097 |
| chr20-30163957-30164458 | 1 | 0.970955648534439 |
| chr7-125658555-125659056 | 4 | 0.95969987232216 |
| chr6-31155172-31155673 | 8 | 0.881277155937682 |
| chr6-13557107-13557608 | 6 | 0.991206925481043 |
| chr17-81061905-81062406 | 5 | 0.875553544658366 |
| chr4-128171402-128171903 | 8 | 0.838580393031868 |
| chr12-39920552-39921053 | 8 | 0.987445467895592 |
| chr7-43739771-43740272 | 3 | 0.876565262657387 |
| chr17-23733311-23733812 | 8 | 0.992222753158444 |
| chr8-65509996-65510497 | 5 | 0.990496339061778 |
| chr14-13762046-13762547 | 3 | 0.88593763158935 |
| chr1-212780519-212781020 | 5 | 0.885651561621525 |

| | | |
|---|---|---|
| chr8-104518462-104518963 | 5 | 0.983919978840557 |
| chr5-117710619-117711120 | 5 | 0.979400312559706 |
| chr7-73341907-73342408 | 6 | 0.991278176569127 |
| chr12-33889616-33890117 | 2 | 0.853844534694591 |
| chrY-10768204-10768705 | 8 | 0.939359068462225 |
| chr2-209049003-209049504 | 8 | 0.963293901533691 |
| chr20-21705774-21706275 | 8 | 0.902670074248116 |
| chr13-85909554-85910055 | 8 | 0.993447946482345 |
| chr1-98400940-98401441 | 3 | 0.950626607860625 |
| chrX-25630330-25630831 | 8 | 0.98287331337078 |
| chr19-39739001-39739502 | 2 | 0.889880241642268 |
| chr2-69311771-69312272 | 3 | 0.996016579976726 |
| chr5-138930564-138931065 | 8 | 0.993493681128311 |
| chr19-36548684-36549185 | 8 | 0.835760953628675 |
| chr1-143681271-143681772 | 8 | 0.993493681128311 |
| chr5-58652242-58652743 | 8 | 0.990077895362341 |
| chr10-50380376-50380877 | 3 | 0.973909950103982 |
| chr17-66110658-66111159 | 8 | 0.993493681128311 |
| chr7-14485048-14485549 | 3 | 0.973745469178426 |
| chr3-113052959-113053460 | 3 | 0.998185601248305 |
| chr8-93881663-93882164 | 3 | 0.996682375475091 |
| chr10-46697101-46697602 | 4 | 0.941478711309699 |
| chr8-53121351-53121852 | 4 | 0.946862413781633 |
| chr4-120250226-120250727 | 8 | 0.895131327704276 |
| chr15-55704434-55704935 | 4 | 0.91965615145822 |
| chr12-29159520-29160021 | 5 | 0.989902521137397 |
| chr8-89887405-89887906 | 3 | 0.996184574464081 |
| chr6-36076868-36077369 | 4 | 0.892924033372723 |
| chr2-46686844-46687345 | 8 | 0.912490297481831 |
| chr19-41117009-41117510 | 6 | 0.969891356516054 |
| chr20-48550029-48550530 | 8 | 0.900606674716075 |
| chr1-80510843-80511344 | 4 | 0.84158455241343 |
| chr18-14463989-14464490 | 3 | 0.99593553545041 |
| chr1-39739982-39740483 | 4 | 0.956706075843022 |
| chr16-82032847-82033348 | 4 | 0.948359160937291 |
| chr12-18605875-18606376 | 8 | 0.964854767668189 |
| chr4-24051127-24051628 | 3 | 0.992977994112625 |
| chr7-28468984-28469485 | 6 | 0.990285535326096 |
| chr17-3804409-3804910 | 7 | 0.897015405922203 |
| chr8-74574014-74574515 | 2 | 0.946657377419299 |
| chr2-205745415-205745916 | 3 | 0.84332215610732 |
| chr2-124148651-124149152 | 6 | 0.985882821875658 |
| chr1-176743758-176744259 | 3 | 0.949895686281085 |

| | | |
|---|---|---|
| chr10-57881489-57881990 | 8 | 0.928864994807358 |
| chr5-125663007-125663508 | 3 | 0.99593553545041 |
| chr2-103523846-103524347 | 8 | 0.987576549838569 |
| chr12-19101288-19101789 | 8 | 0.817890696680863 |
| chr11-36739788-36740289 | 3 | 0.893007552749269 |
| chr12-39360003-39360504 | 5 | 0.990780133032782 |
| chr1-232586446-232586947 | 3 | 0.917075215537722 |
| chr20-48672185-48672686 | 6 | 0.97059813100746 |
| chr7-27953192-27953693 | 5 | 0.967633082876001 |
| chr5-166383090-166383591 | 3 | 0.980677383878438 |
| chr5-21958916-21959417 | 5 | 0.980165975540377 |
| chr10-93578420-93578921 | 7 | 0.938640807310853 |
| chr13-38837384-38837885 | 5 | 0.983047993381758 |
| chr1-31062804-31063305 | 6 | 0.857157761578292 |
| chr2-192386979-192387480 | 8 | 0.979118965094651 |
| chr18-38495649-38496150 | 3 | 0.995730101346191 |
| chr10-10629591-10630092 | 8 | 0.958765515619669 |
| chr4-7242444-7242945 | 8 | 0.841959322244636 |
| chr3-135702044-135702545 | 1 | 0.960047914325493 |
| chr8-106018493-106018994 | 4 | 0.829194591440547 |
| chr8-85008408-85008909 | 6 | 0.931308298801228 |
| chr14-31222128-31222629 | 8 | 0.851379715384569 |
| chr14-41348782-41349283 | 8 | 0.983944949366016 |
| chr16-1513797-1514298 | 8 | 0.987804544089799 |
| chr1-165010221-165010722 | 7 | 0.836578474658886 |
| chr6-111509211-111509712 | 4 | 0.936268138665734 |
| chr7-86635533-86636034 | 3 | 0.87717158378162 |
| chr9-81876842-81877343 | 3 | 0.99665347489431 |
| chrX-127542737-127543238 | 5 | 0.952674010214779 |
| chr13-64864449-64864950 | 8 | 0.95142289127503 |
| chr15-78348733-78349234 | 7 | 0.844382650881717 |
| chr15-51142480-51142981 | 8 | 0.916580464537279 |
| chrX-102402388-102402889 | 3 | 0.99473073244357 |
| chr17-10676598-10677099 | 2 | 0.840085826016338 |
| chr13-29523862-29524363 | 3 | 0.99000028888497 |
| chr3-52280472-52280973 | 3 | 0.99593553545041 |
| chr5-25865958-25866459 | 5 | 0.986771253925588 |
| chr4-128088172-128088673 | 8 | 0.844983237849115 |
| chr9-110574492-110574993 | 8 | 0.9569322964263 |
| chr18-24562192-24562693 | 3 | 0.907256634608734 |
| chr18-68433224-68433725 | 8 | 0.876595439197224 |
| chr3-147254291-147254792 | 8 | 0.887185760916555 |
| chr14-6206054-6206555 | 2 | 0.903000217991882 |

| | | |
|---|---|---|
| chr1-172200365-172200866 | 5 | 0.937074639114121 |
| chr7-106541699-106542200 | 7 | 0.880898640460998 |
| chr4-134926019-134926520 | 3 | 0.99593553545041 |
| chr8-59394410-59394911 | 8 | 0.943489279127872 |
| chr15-81868302-81868803 | 8 | 0.901042822089066 |
| chr13-68952822-68953323 | 4 | 0.880542924973467 |
| chr7-19902588-19903089 | 5 | 0.989479757616336 |
| chr5-79078470-79078971 | 8 | 0.92553940983474 |
| chr17-71784775-71785276 | 8 | 0.9786577191148 |
| chr4-12336422-12336923 | 1 | 0.926979955703767 |
| chr1-185263983-185264484 | 3 | 0.831927411116549 |
| chrX-20851345-20851846 | 6 | 0.88087314991115 |
| chr4-180413027-180413528 | 8 | 0.962409716446537 |
| chr19-47456981-47457482 | 7 | 0.884415950404875 |
| chr7-71991722-71992223 | 8 | 0.882235052596403 |
| chr19-35553894-35554395 | 4 | 0.863981205503549 |
| chr11-42895843-42896344 | 7 | 0.823695201764976 |
| chr8-119871672-119872173 | 8 | 0.848651116814579 |
| chr13-104361456-104361957 | 8 | 0.91607820027168 |
| chr1-240077545-240078046 | 8 | 0.923871027777213 |
| chr11-35176770-35177271 | 3 | 0.960480126094165 |
| chr8-76035908-76036409 | 5 | 0.969639565926293 |
| chr16-33168462-33168963 | 8 | 0.993493681128311 |
| chr5-63856166-63856667 | 2 | 0.876554619817351 |
| chr19-2745688-2746189 | 3 | 0.996897532724487 |
| chr6-57873713-57874214 | 8 | 0.990636256734665 |
| chr5-43083755-43084256 | 4 | 0.90612124854141 |
| chr6-71133908-71134409 | 8 | 0.945732791723547 |
| chr3-150782313-150782814 | 6 | 0.992240590998112 |
| chrX-49656571-49657072 | 8 | 0.988656403597106 |
| chr4-174466214-174466715 | 5 | 0.965068364378807 |
| chr6-20036379-20036880 | 8 | 0.989379173196804 |
| chr1-186392133-186392634 | 8 | 0.883655919731005 |
| chr17-69692481-69692982 | 3 | 0.981626211242577 |
| chrX-10703525-10704026 | 5 | 0.82973020321366 |
| chrX-81633807-81634308 | 3 | 0.925574184759578 |
| chr18-70820163-70820664 | 8 | 0.829031332941832 |
| chr2-183393811-183394312 | 3 | 0.996075728404651 |
| chr13-56253510-56254011 | 3 | 0.996024784503694 |
| chr7-107069405-107069906 | 8 | 0.853351423158838 |
| chr17-23371479-23371980 | 8 | 0.993493681128311 |
| chr2-109909832-109910333 | 3 | 0.99593553545041 |
| chr6-34078933-34079434 | 8 | 0.988884728261482 |

| | | |
|---|---|---|
| chr4-166909336-166909837 | 8 | 0.995226166137308 |
| chr13-64385693-64386194 | 8 | 0.902594168218213 |
| chr8-12319508-12320009 | 2 | 0.830944665634603 |
| chr19-41696978-41697479 | 6 | 0.83231045199843 |
| chr12-29796212-29796713 | 3 | 0.888502411340429 |
| chr12-10108606-10109107 | 8 | 0.995248532809436 |
| chr4-157978192-157978693 | 8 | 0.912865362785185 |
| chr1-136520206-136520707 | 2 | 0.917101457516642 |
| chr4-72713603-72714104 | 3 | 0.995608438370435 |
| chr10-70910360-70910861 | 8 | 0.904241447735984 |
| chr17-4193909-4194410 | 7 | 0.929770440510906 |
| chr8-49564484-49564985 | 8 | 0.994201870151637 |
| chr4-96215570-96216071 | 1 | 0.922584850316597 |
| chr17-5630240-5630741 | 7 | 0.841506796936524 |
| chr15-100171627-100172128 | 8 | 0.942964957314355 |
| chr7-126306962-126307463 | 6 | 0.953381666558087 |
| chr5-89963892-89964393 | 5 | 0.949546863872598 |
| chr1-45416585-45417086 | 8 | 0.878844147433832 |
| chr13-49366777-49367278 | 8 | 0.88739683626545 |
| chr8-12034666-12035167 | 3 | 0.844890111670462 |
| chr1-199093089-199093590 | 3 | 0.829269065378283 |
| chr5-58516429-58516930 | 5 | 0.978320682921652 |
| chr5-97398853-97399354 | 3 | 0.985891962668824 |
| chr15-20408787-20409288 | 4 | 0.846158640675039 |
| chr9-37385277-37385778 | 3 | 0.997293964871764 |
| chr10-42399232-42399733 | 3 | 0.996196088458596 |
| chr20-44590727-44591228 | 8 | 0.851422150681258 |
| chr9-74482346-74482847 | 6 | 0.993793334327063 |
| chr20-20702640-20703141 | 8 | 0.821535381241231 |
| chr3-131460339-131460840 | 8 | 0.986838723282227 |
| chr17-12257377-12257878 | 8 | 0.825231709760826 |
| chr18-69951830-69952331 | 6 | 0.923793231375164 |
| chrX-139590258-139590759 | 3 | 0.99593553545041 |
| chr1-4978303-4978804 | 3 | 0.99593553545041 |
| chr3-107168955-107169456 | 4 | 0.97486623839935 |
| chr14-77620126-77620627 | 8 | 0.970870692286231 |
| chr5-69932102-69932603 | 8 | 0.958554844187995 |
| chr2-84609639-84610140 | 3 | 0.996488464743796 |
| chr2-42777679-42778180 | 8 | 0.969201978921146 |
| chr14-28192437-28192938 | 3 | 0.936402244038762 |
| chr1-247454713-247455214 | 8 | 0.864556475277852 |
| chr2-119480246-119480747 | 8 | 0.849584600987149 |
| chr7-127666212-127666713 | 7 | 0.893894332379857 |

| | | |
|---|---|---|
| chr13-102335630-102336131 | 8 | 0.907700624718027 |
| chr9-63597592-63598093 | 7 | 0.955647952529898 |
| chr1-170440510-170441011 | 8 | 0.827837848787437 |
| chrX-106306307-106306808 | 8 | 0.823805414734978 |
| chr1-90883133-90883634 | 8 | 0.975688789013332 |
| chr6-39606772-39607273 | 8 | 0.993167762022506 |
| chr1-98883643-98884144 | 1 | 0.945821850635688 |
| chr4-95626378-95626879 | 3 | 0.953580340304932 |
| chr5-99815031-99815532 | 4 | 0.84266517050452 |
| chr9-57778058-57778559 | 5 | 0.98030911156561 |
| chr5-119277176-119277677 | 7 | 0.94257656810156 |
| chr2-188702785-188703286 | 5 | 0.983919978840557 |
| chr17-59674667-59675168 | 3 | 0.996368886984596 |
| chr8-67628542-67629043 | 8 | 0.919467254624268 |
| chr10-99432999-99433500 | 3 | 0.99617289233125 |
| chr6-47976489-47976990 | 6 | 0.997122873676964 |
| chr12-32693892-32694393 | 6 | 0.996228934030301 |
| chr6-31006680-31007181 | 8 | 0.899113997461355 |
| chr10-104937887-104938388 | 8 | 0.970930332228073 |
| chr2-139485850-139486351 | 8 | 0.99189730968073 |
| chr9-108371380-108371881 | 6 | 0.905485831206088 |
| chr4-163080488-163080989 | 3 | 0.996035653235031 |
| chr1-225081078-225081579 | 1 | 0.853798663840512 |
| chr1-241177444-241177945 | 8 | 0.927618107217538 |
| chr6-53720783-53721284 | 3 | 0.986626891382922 |
| chr19-23983933-23984434 | 8 | 0.944591890419288 |
| chr1-118170629-118171130 | 8 | 0.958129787352367 |
| chr14-95018836-95019337 | 3 | 0.856854407615072 |
| chrX-80952365-80952866 | 3 | 0.986860021383769 |
| chr13-90953448-90953949 | 8 | 0.885259111042183 |
| chr2-185155216-185155717 | 5 | 0.980932680260484 |
| chr2-86518673-86519174 | 6 | 0.993102606809521 |
| chr11-1250012-1250513 | 7 | 0.815371990594205 |
| chr18-64020135-64020636 | 8 | 0.957786846189813 |
| chr1-209187461-209187962 | 6 | 0.88631954556948 |
| chr17-17307806-17308307 | 3 | 0.996770152175886 |
| chr1-172391076-172391577 | 8 | 0.897084631065995 |
| chr3-155767752-155768253 | 8 | 0.974932100422805 |
| chr5-85305040-85305541 | 6 | 0.983568858387113 |
| chr14-100453087-100453588 | 4 | 0.836208872824819 |
| chr8-61240599-61241100 | 8 | 0.923695929174364 |
| chr3-147710013-147710514 | 3 | 0.91890873205231 |
| chr20-45036787-45037288 | 4 | 0.816031580794471 |

| | | |
|---|---|---|
| chr1-151903604-151904105 | 8 | 0.994923438605759 |
| chr5-25074326-25074827 | 1 | 0.949751234021723 |
| chr5-70205365-70205866 | 3 | 0.996895448726267 |
| chr6-84537644-84538145 | 3 | 0.991548370700568 |
| chrX-132180958-132181459 | 8 | 0.934157663062962 |
| chr20-28590710-28591211 | 4 | 0.906499442341639 |
| chr5-77077144-77077645 | 5 | 0.993229244567149 |
| chr13-100610789-100611290 | 3 | 0.99593553545041 |
| chr6-9603743-9604244 | 4 | 0.904172591721564 |
| chr8-67576098-67576599 | 8 | 0.943973149503023 |
| chr6-52734007-52734508 | 8 | 0.875879175817828 |
| chr14-72492050-72492551 | 6 | 0.992816758412934 |
| chr10-95495718-95496219 | 6 | 0.844895795385242 |
| chr19-37543417-37543918 | 5 | 0.957554223198429 |
| chr1-14680444-14680945 | 5 | 0.967239013232468 |
| chr10-17387656-17388157 | 4 | 0.930175940948668 |
| chr12-27875255-27875756 | 8 | 0.977870971647882 |
| chr16-75586619-75587120 | 8 | 0.991666332175796 |
| chr10-59657395-59657896 | 8 | 0.947784897422671 |
| chr6-9924879-9925380 | 2 | 0.920437409050874 |
| chr10-57915179-57915680 | 7 | 0.841958856187893 |
| chr5-50551839-50552340 | 8 | 0.870484317895262 |
| chr10-59962759-59963260 | 5 | 0.978413498664463 |
| chrX-78238459-78238960 | 6 | 0.989554488179095 |
| chr4-18251003-18251504 | 8 | 0.993493681128311 |
| chr7-58980475-58980976 | 6 | 0.992207948879996 |
| chr5-135344012-135344513 | 8 | 0.949604515569723 |
| chr1-96877100-96877601 | 3 | 0.99593553545041 |
| chr10-94486711-94487212 | 4 | 0.914938084877818 |
| chr7-125512950-125513451 | 6 | 0.995144314083709 |
| chr5-39065524-39066025 | 5 | 0.939996901592625 |
| chr11-54399208-54399709 | 4 | 0.893421091860764 |
| chr19-44000119-44000620 | 4 | 0.825432144594566 |
| chr2-219806068-219806569 | 5 | 0.98288684210311 |
| chr1-83539678-83540179 | 8 | 0.992928621756582 |
| chr6-15013933-15014434 | 5 | 0.885152108439234 |
| chr11-55550460-55550961 | 3 | 0.996041365453935 |
| chr12-12499546-12500047 | 8 | 0.993493681128311 |
| chr2-137799685-137800186 | 5 | 0.850712722204052 |
| chr7-93103003-93103504 | 8 | 0.875333474327058 |
| chr5-101076619-101077120 | 8 | 0.936090082157259 |
| chr6-124441271-124441772 | 1 | 0.878359831623158 |
| chr6-113034108-113034609 | 6 | 0.996522309823268 |

| | | |
|---|---|---|
| chr1-107960368-107960869 | 3 | 0.995815878076725 |
| chr14-95669428-95669929 | 2 | 0.910320477481226 |
| chr4-117710762-117711263 | 3 | 0.99655956927036 |
| chr7-87772662-87773163 | 2 | 0.819257004209548 |
| chr16-55371700-55372201 | 4 | 0.877336788925641 |
| chr6-49646361-49646862 | 8 | 0.983254149264059 |
| chr9-40304603-40305104 | 5 | 0.849918760217929 |
| chr8-2355957-2356458 | 6 | 0.909332270751482 |
| chr5-81409609-81410110 | 3 | 0.996778471232105 |
| chr14-24713816-24714317 | 8 | 0.857456451415222 |
| chr10-100022806-100023307 | 4 | 0.851695184209459 |
| chr15-19199578-19200079 | 8 | 0.976275200709388 |
| chr18-52276653-52277154 | 8 | 0.994505768098035 |
| chr2-24721323-24721824 | 8 | 0.855901580446855 |
| chr1-226605881-226606382 | 7 | 0.831636719863538 |
| chr11-85248037-85248538 | 8 | 0.973196748368879 |
| chr1-7741349-7741850 | 3 | 0.929296995203632 |
| chr4-7179305-7179806 | 5 | 0.988717827859362 |
| chr13-48264985-48265486 | 1 | 0.852739152185864 |
| chr5-166376930-166377431 | 8 | 0.909336744007618 |
| chr15-22087049-22087550 | 4 | 0.909917064940818 |
| chr16-49108677-49109178 | 2 | 0.882287526020696 |
| chr2-31897320-31897821 | 2 | 0.9499356240041 |
| chr2-226245074-226245575 | 6 | 0.982486946372097 |
| chr5-2146782-2147283 | 6 | 0.970380004387678 |
| chr17-7867556-7868057 | 8 | 0.995222180201727 |
| chr17-6563109-6563610 | 5 | 0.9856495027359 |
| chr5-50192244-50192745 | 3 | 0.99610823071023 |
| chr2-113188895-113189396 | 1 | 0.870573097610267 |
| chr19-16283706-16284207 | 6 | 0.949911665382645 |
| chr2-6839089-6839590 | 6 | 0.991206925481043 |
| chr16-41767638-41768139 | 8 | 0.888134110981514 |
| chr16-36285810-36286311 | 7 | 0.909198712973059 |
| chr9-50477638-50478139 | 6 | 0.993219783046378 |
| chr8-100493843-100494344 | 6 | 0.984276975604174 |
| chr5-16249914-16250415 | 3 | 0.997223780527493 |
| chr13-82157016-82157517 | 8 | 0.994972524074601 |
| chr10-53054699-53055200 | 8 | 0.843048440426911 |
| chrX-35929366-35929867 | 5 | 0.987681041860473 |
| chr2-137689779-137690280 | 4 | 0.881566032958975 |
| chr2-176195868-176196369 | 8 | 0.970387682835006 |
| chr7-108270749-108271250 | 2 | 0.897074842909927 |
| chr5-12371822-12372323 | 6 | 0.967116916048551 |

| | | |
|---|---|---|
| chr4-5915860-5916361 | 3 | 0.996883980006107 |
| chr20-11017311-11017812 | 8 | 0.834482388855807 |
| chr10-49558791-49559292 | 8 | 0.968991039910481 |
| chr1-171153978-171154479 | 6 | 0.991206925481043 |
| chr1-255345944-255346445 | 8 | 0.995578861674573 |
| chr3-123193833-123194334 | 3 | 0.996829805559368 |
| chr16-27765918-27766419 | 2 | 0.922324136703221 |
| chr3-76275565-76276066 | 3 | 0.996898085830519 |
| chr1-18465844-18466345 | 6 | 0.91662279608946 |
| chr20-12789632-12790133 | 1 | 0.944031573089681 |
| chr8-26676524-26677025 | 6 | 0.986606075928053 |
| chr15-14605297-14605798 | 5 | 0.849415191197269 |
| chr7-49621935-49622436 | 6 | 0.991665392023823 |
| chr10-85289381-85289882 | 5 | 0.875435719704132 |
| chr7-86027340-86027841 | 8 | 0.894305501804948 |
| chr11-50087570-50088071 | 8 | 0.966995044800624 |
| chr16-10306320-10306821 | 4 | 0.873237513408094 |
| chr11-55493844-55494345 | 3 | 0.99248983326709 |
| chr18-14608077-14608578 | 5 | 0.878542323020522 |
| chr17-72958760-72959261 | 8 | 0.956512579871459 |
| chr14-58583933-58584434 | 6 | 0.978666972266883 |
| chr12-23047962-23048463 | 5 | 0.983919978840557 |
| chr20-33746726-33747227 | 3 | 0.980760413502972 |
| chr1-128239183-128239684 | 1 | 0.819419852608577 |
| chr14-57331868-57332369 | 8 | 0.94201498552669 |
| chr8-37227874-37228375 | 3 | 0.979501793265657 |
| chr7-118110632-118111133 | 3 | 0.996423192982355 |
| chr5-105859746-105860247 | 8 | 0.848359517515509 |
| chr18-54422803-54423304 | 8 | 0.976577891839885 |
| chr3-117054221-117054722 | 4 | 0.921024775801428 |
| chr1-141887753-141888254 | 8 | 0.89504934232414 |
| chr17-23200499-23201000 | 5 | 0.958854456512373 |
| chrX-44979694-44980195 | 3 | 0.99593553545041 |
| chr18-62158057-62158558 | 3 | 0.899459910092122 |
| chr17-64300801-64301302 | 8 | 0.886955120944065 |
| chr1-132301615-132302116 | 2 | 0.954801316030335 |
| chr9-6908146-6908647 | 2 | 0.971620795041815 |
| chr9-77734312-77734813 | 8 | 0.919472007748199 |
| chr5-60525789-60526290 | 5 | 0.8292738425534 |
| chr11-81734993-81735494 | 3 | 0.964154482483955 |
| chr4-98644410-98644911 | 6 | 0.935895678259066 |
| chr14-57097591-57098092 | 5 | 0.880321503144474 |
| chr16-8739379-8739880 | 5 | 0.989023041829733 |

| | | |
|---|---|---|
| chr10-59801601-59802102 | 8 | 0.953718501743294 |
| chr8-58723363-58723864 | 8 | 0.840071971703756 |
| chr1-244683698-244684199 | 8 | 0.993493681128311 |
| chr16-3410029-3410530 | 3 | 0.996822300914249 |
| chr12-40280308-40280809 | 4 | 0.95396364652995 |
| chr9-46073734-46074235 | 8 | 0.889892238436868 |
| chr3-125044593-125045094 | 1 | 0.879104679407443 |
| chr3-141990811-141991312 | 8 | 0.983729908546286 |
| chr15-57484165-57484666 | 3 | 0.984430610318175 |
| chr17-26192107-26192608 | 2 | 0.957189476090916 |
| chr4-96648066-96648567 | 3 | 0.996897566881258 |
| chr15-81153436-81153937 | 3 | 0.996163285607072 |
| chr7-98928561-98929062 | 8 | 0.994876172671872 |
| chr14-97159802-97160303 | 8 | 0.925989553684017 |
| chr17-20532155-20532656 | 5 | 0.983548335635071 |
| chr1-165874483-165874984 | 7 | 0.956689956892717 |
| chr7-29015995-29016496 | 8 | 0.947470692685687 |
| chr14-32309145-32309646 | 5 | 0.940204124713214 |
| chr18-8423764-8424265 | 8 | 0.993493681128311 |
| chr5-150890034-150890535 | 6 | 0.992222166342784 |
| chr6-100183680-100184181 | 4 | 0.913935171103495 |
| chr4-51163016-51163517 | 2 | 0.945575421256468 |
| chr2-149429344-149429845 | 8 | 0.960757096448901 |
| chr15-82510601-82511102 | 2 | 0.887185146076579 |
| chr4-155116996-155117497 | 3 | 0.917351366705012 |
| chr13-91416341-91416842 | 8 | 0.851070649626962 |
| chr3-5747971-5748472 | 3 | 0.989050993862611 |
| chr11-38744408-38744909 | 3 | 0.877600808334058 |
| chr7-68982119-68982620 | 8 | 0.922792338215229 |
| chr9-34753052-34753553 | 2 | 0.957148508310848 |
| chr5-40908834-40909335 | 1 | 0.953929448646409 |
| chr3-61947336-61947837 | 8 | 0.862010563740406 |
| chr8-94612866-94613367 | 3 | 0.982403772862563 |
| chr1-132098214-132098715 | 6 | 0.991284614275097 |
| chr14-9969125-9969626 | 2 | 0.960480442181295 |
| chr12-19266121-19266622 | 5 | 0.983919978840557 |
| chr3-39726564-39727065 | 6 | 0.99322052248377 |
| chr13-104110539-104111040 | 1 | 0.829918018382957 |
| chr5-144843185-144843686 | 6 | 0.99125747181061 |
| chr11-25840066-25840567 | 1 | 0.905611820515872 |
| chr2-122926100-122926601 | 8 | 0.899548046160253 |
| chr6-71257401-71257902 | 5 | 0.960137620572756 |
| chr5-113024518-113025019 | 8 | 0.891414907079949 |

| | | |
|---|---|---|
| chr16-7374939-7375440 | 2 | 0.862227311849665 |
| chr2-61119569-61120070 | 4 | 0.939682045866071 |
| chr5-101433188-101433689 | 8 | 0.944871571576608 |
| chr5-150992165-150992666 | 8 | 0.986178467034945 |
| chr16-54890021-54890522 | 8 | 0.976609334283199 |
| chr2-49870085-49870586 | 5 | 0.868460319548246 |
| chr2-41026830-41027331 | 8 | 0.865467919062493 |
| chr11-10347071-10347572 | 2 | 0.862108056248202 |
| chr4-118169700-118170201 | 3 | 0.996372398751631 |
| chr19-9744364-9744865 | 2 | 0.884522381392861 |
| chr7-52484575-52485076 | 7 | 0.899340111770982 |
| chr19-24861733-24862234 | 8 | 0.895626108062309 |
| chr2-216859593-216860094 | 5 | 0.8225248226468 |
| chr2-77341174-77341675 | 2 | 0.910251491745192 |
| chr2-236790916-236791417 | 3 | 0.998304286837797 |
| chr12-15996909-15997410 | 3 | 0.99593553545041 |
| chr9-88778965-88779466 | 8 | 0.814811995976051 |
| chr7-15870865-15871366 | 2 | 0.907452936972599 |
| chr7-92435564-92436065 | 3 | 0.99570514172483 |
| chr6-9171484-9171985 | 4 | 0.971958582758704 |
| chr1-173328171-173328672 | 8 | 0.836016426608871 |
| chr10-76062642-76063143 | 2 | 0.938705648104873 |
| chr5-31467821-31468322 | 8 | 0.995026326919562 |
| chr9-40073692-40074193 | 3 | 0.997137179199004 |
| chr6-36859729-36860230 | 8 | 0.967492011685669 |
| chr17-42361296-42361797 | 2 | 0.933025434991937 |
| chr11-12786381-12786882 | 3 | 0.996706363878291 |
| chr20-29946329-29946830 | 8 | 0.975990315080907 |
| chr9-111262851-111263352 | 4 | 0.948045871734107 |
| chr15-58160959-58161460 | 3 | 0.954841271319205 |
| chr3-80878632-80879133 | 8 | 0.925592315267658 |
| chr2-138688184-138688685 | 8 | 0.983957205147011 |
| chr5-114258812-114259313 | 4 | 0.932133779767099 |
| chr8-44036102-44036603 | 4 | 0.917758304297288 |
| chr6-110212639-110213140 | 7 | 0.81740729686761 |
| chr13-94357859-94358360 | 3 | 0.985982987709795 |
| chr4-90235077-90235578 | 3 | 0.970911626943864 |
| chr1-156129022-156129523 | 8 | 0.909330188349501 |
| chr3-125541990-125542491 | 7 | 0.898196016131609 |
| chr19-9031484-9031985 | 3 | 0.995968890682665 |
| chr15-54817144-54817645 | 3 | 0.99593553545041 |
| chr15-34974689-34975190 | 4 | 0.906573466827176 |
| chr1-207576549-207577050 | 8 | 0.960364751844095 |

| | | |
|---|---|---|
| chr3-129852962-129853463 | 8 | 0.993493681128311 |
| chr6-96430481-96430982 | 8 | 0.993493681128311 |
| chr5-21123650-21124151 | 8 | 0.950826094377714 |
| chr10-76507156-76507657 | 5 | 0.979131733162012 |
| chr2-33527282-33527783 | 6 | 0.978815760767658 |
| chr3-23996279-23996780 | 6 | 0.991206925481043 |
| chr9-9501136-9501637 | 3 | 0.979923329662182 |
| chr9-110346692-110347193 | 8 | 0.926035084689534 |
| chr15-29956688-29957189 | 6 | 0.961791789205268 |
| chr9-48699369-48699870 | 8 | 0.925952410278375 |
| chr6-22068374-22068875 | 3 | 0.948580778975836 |
| chr2-148688215-148688716 | 3 | 0.99593553545041 |
| chr9-9717050-9717551 | 3 | 0.992945790095019 |
| chr9-90336367-90336868 | 7 | 0.822318805108501 |
| chr6-105585994-105586495 | 8 | 0.993636064227565 |
| chr7-97945058-97945559 | 8 | 0.881470726745634 |
| chr6-36172268-36172769 | 2 | 0.901767666346807 |
| chrX-65682396-65682897 | 6 | 0.823230726227001 |
| chr3-52983645-52984146 | 8 | 0.921201135783672 |
| chr9-74973168-74973669 | 5 | 0.979437000249609 |
| chr3-142494223-142494724 | 8 | 0.98387541716302 |
| chr5-45579176-45579677 | 3 | 0.99593553545041 |
| chr16-57364084-57364585 | 8 | 0.930439616205768 |
| chr8-99182250-99182751 | 5 | 0.958191490107793 |
| chr20-9995128-9995629 | 8 | 0.910643735778759 |
| chr1-126527430-126527931 | 8 | 0.993493681128311 |
| chr2-142016341-142016842 | 8 | 0.841100033622086 |
| chr5-101666266-101666767 | 4 | 0.87185616173929 |
| chr15-32967729-32968230 | 8 | 0.967802188202022 |
| chr3-28804687-28805188 | 5 | 0.900915385099038 |
| chr2-153774593-153775094 | 8 | 0.948519292615452 |
| chr18-25071533-25072034 | 4 | 0.872665370829304 |
| chr5-145206019-145206520 | 8 | 0.898937786838167 |
| chr13-3555627-3556128 | 6 | 0.983697830113674 |
| chr1-155959248-155959749 | 8 | 0.947593523800868 |
| chr4-136119720-136120221 | 2 | 0.940878275815785 |
| chr13-45890588-45891089 | 5 | 0.98488110227207 |
| chr13-89635430-89635931 | 2 | 0.951065183145121 |
| chr14-46033528-46034029 | 3 | 0.893658942418642 |
| chr1-7932125-7932626 | 4 | 0.913701410411511 |
| chr14-5600025-5600526 | 7 | 0.928665576501306 |
| chr14-57181711-57182212 | 8 | 0.898660090597154 |
| chr6-38947325-38947826 | 8 | 0.863159924775638 |

| | | |
|---|---|---|
| chr10-75152240-75152741 | 8 | 0.835664193822777 |
| chr19-21334273-21334774 | 8 | 0.955297475063808 |
| chrX-58192069-58192570 | 8 | 0.975662545667974 |
| chr11-4591012-4591513 | 3 | 0.99178846279883 |
| chr11-52402985-52403486 | 1 | 0.920890891225896 |
| chr1-121283380-121283881 | 8 | 0.943179346055703 |
| chr17-22904230-22904731 | 8 | 0.950476940227883 |
| chr5-73114882-73115383 | 4 | 0.933814898897464 |
| chr11-33350622-33351123 | 5 | 0.988598898659505 |
| chr4-93232349-93232850 | 6 | 0.992846695860325 |
| chr6-132143848-132144349 | 8 | 0.995076765489059 |
| chr14-63687922-63688423 | 8 | 0.867815048988807 |
| chr20-30547164-30547665 | 8 | 0.932922624393224 |
| chr13-44157808-44158309 | 2 | 0.840373727931665 |
| chr1-258285803-258286304 | 7 | 0.86564017998895 |
| chr9-38434414-38434915 | 2 | 0.955269033353368 |
| chrX-127381086-127381587 | 8 | 0.93736486208081 |
| chr16-78119741-78120242 | 7 | 0.816286725377189 |
| chr16-22088083-22088584 | 3 | 0.99593553545041 |
| chr3-51434456-51434957 | 8 | 0.947492222169682 |
| chr15-70719852-70720353 | 4 | 0.816567509926068 |
| chr2-226793059-226793560 | 6 | 0.993828747926915 |
| chr1-200928550-200929051 | 8 | 0.974712667830542 |
| chr13-52526333-52526834 | 8 | 0.901567274770534 |
| chr4-78447140-78447641 | 3 | 0.98707767860141 |
| chr6-16847239-16847740 | 3 | 0.890945561995148 |
| chr7-62918329-62918830 | 8 | 0.861457785276994 |
| chr8-75657584-75658085 | 8 | 0.978617857659222 |
| chr14-34593981-34594482 | 8 | 0.995183745820273 |
| chr3-34965418-34965919 | 8 | 0.9817276346849 |
| chr2-136921555-136922056 | 2 | 0.94961768892899 |
| chr5-140533540-140534041 | 8 | 0.993493681128311 |
| chr20-30562265-30562766 | 8 | 0.978169864438123 |
| chr7-125070238-125070739 | 5 | 0.991650519823049 |
| chr19-4928176-4928677 | 6 | 0.979747802827012 |
| chr6-14218976-14219477 | 8 | 0.968392392470383 |
| chr13-34964168-34964669 | 6 | 0.984273274367617 |
| chr1-219450585-219451086 | 1 | 0.817597797668111 |
| chr6-84969641-84970142 | 3 | 0.883576866712115 |
| chr14-58722341-58722842 | 3 | 0.99593553545041 |
| chr13-54299507-54300008 | 2 | 0.891718783708752 |
| chr19-52315573-52316074 | 6 | 0.987086588101339 |
| chr7-27348675-27349176 | 5 | 0.971907330891693 |

| | | |
|---|---|---|
| chr4-108594297-108594798 | 3 | 0.992737259981285 |
| chr3-44768829-44769330 | 8 | 0.930354414262004 |
| chr3-122357127-122357628 | 1 | 0.837881084761622 |
| chr13-79605686-79606187 | 8 | 0.916736939590837 |
| chr5-51489452-51489953 | 6 | 0.940095264890219 |
| chr7-11099427-11099928 | 3 | 0.996746221802541 |
| chr11-18438040-18438541 | 3 | 0.956469590017982 |
| chr1-221757017-221757518 | 4 | 0.819012096553617 |
| chr15-95874732-95875233 | 8 | 0.839826558797722 |
| chr6-48570434-48570935 | 6 | 0.986990660099803 |
| chr4-150039570-150040071 | 8 | 0.888787992658806 |
| chr4-37429058-37429559 | 1 | 0.936364768927347 |
| chr4-59839201-59839702 | 1 | 0.831224539601585 |
| chr20-47661605-47662106 | 7 | 0.906914597355071 |
| chr14-64074469-64074970 | 4 | 0.897367291738622 |
| chr3-63044412-63044913 | 2 | 0.951412423117401 |
| chrX-139396790-139397291 | 6 | 0.873446251474083 |
| chr5-78262078-78262579 | 6 | 0.993227411062818 |
| chr17-17800195-17800696 | 3 | 0.988860392167018 |
| chr4-93117461-93117962 | 4 | 0.892512177140622 |
| chr14-52481800-52482301 | 3 | 0.972526581135955 |
| chr11-56601314-56601815 | 8 | 0.958537958912539 |
| chr8-12749138-12749639 | 8 | 0.965225024450881 |
| chr14-38705056-38705557 | 6 | 0.987517212543133 |
| chr13-96111297-96111798 | 8 | 0.854677875335633 |
| chr5-12617806-12618307 | 3 | 0.893324682087745 |
| chr14-7433953-7434454 | 2 | 0.886199344450989 |
| chr15-37456718-37457219 | 4 | 0.830983736944116 |
| chr20-28961272-28961773 | 5 | 0.947755852499077 |
| chr11-5899924-5900425 | 3 | 0.856907391251296 |
| chr1-90940195-90940696 | 8 | 0.860992862502215 |
| chr15-80981124-80981625 | 7 | 0.88866032018098 |
| chr10-89398096-89398597 | 8 | 0.911673650836219 |
| chr4-14937221-14937722 | 4 | 0.847276713265689 |
| chrX-17836144-17836645 | 3 | 0.987151498923782 |
| chr1-86907302-86907803 | 3 | 0.953584533985568 |
| chr16-44947444-44947945 | 8 | 0.865531067407092 |
| chr17-56424379-56424880 | 7 | 0.866194311668982 |
| chr9-38317865-38318366 | 4 | 0.88834809821666 |
| chr8-62209898-62210399 | 4 | 0.926430215116359 |
| chr14-16196960-16197461 | 5 | 0.942736176444065 |
| chr6-106658844-106659345 | 8 | 0.905632058179238 |
| chr3-63081804-63082305 | 3 | 0.98560071852061 |

| | | |
|---|---|---|
| chr8-111389114-111389615 | 3 | 0.99593553545041 |
| chr19-47934425-47934926 | 5 | 0.988776450370087 |
| chr3-160132926-160133427 | 5 | 0.970410654464261 |
| chr13-44250546-44251047 | 4 | 0.965273350911401 |
| chr7-53857912-53858413 | 3 | 0.996454166174016 |
| chr9-6681084-6681585 | 8 | 0.986390151064129 |
| chr3-147959747-147960248 | 3 | 0.994623307118666 |
| chr2-191060570-191061071 | 8 | 0.97117487775977 |
| chr1-174660170-174660671 | 1 | 0.957422231061468 |
| chr2-103142321-103142822 | 8 | 0.827447648962807 |
| chr9-13838892-13839393 | 8 | 0.980503647129182 |
| chr10-96702605-96703106 | 4 | 0.881201931364418 |
| chr5-4812825-4813326 | 8 | 0.919414617801211 |
| chr3-9339135-9339636 | 4 | 0.886697410150423 |
| chr18-43248582-43249083 | 4 | 0.960915684563521 |
| chr1-223469428-223469929 | 8 | 0.988918269090364 |
| chr11-50887107-50887608 | 6 | 0.976970243879428 |
| chr7-37256190-37256691 | 8 | 0.995092971648716 |
| chr19-27476892-27477393 | 3 | 0.994975670106234 |
| chr19-25243331-25243832 | 8 | 0.940827597735275 |
| chr18-66230396-66230897 | 8 | 0.989430889751492 |
| chr2-41513262-41513763 | 2 | 0.929556987609842 |
| chr5-46505554-46506055 | 4 | 0.963120830164793 |
| chr8-66866339-66866840 | 3 | 0.95177699355563 |
| chr5-54206638-54207139 | 3 | 0.996715453791441 |
| chr18-22459615-22460116 | 5 | 0.984047249215827 |
| chr4-159358667-159359168 | 5 | 0.986909954837459 |
| chr5-120729399-120729900 | 3 | 0.903042231342903 |
| chr5-47084404-47084905 | 6 | 0.987076661417548 |
| chr2-41579435-41579936 | 8 | 0.993493681128311 |
| chr9-33230733-33231234 | 8 | 0.958837831147182 |
| chr10-67161072-67161573 | 4 | 0.832284069717264 |
| chr5-19919931-19920432 | 6 | 0.98982469665573 |
| chr13-85889467-85889968 | 3 | 0.99593553545041 |
| chr18-50157035-50157536 | 5 | 0.870246035360078 |
| chr7-20169263-20169764 | 8 | 0.90409260514068 |
| chr15-83241906-83242407 | 6 | 0.992412942750883 |
| chr19-55291598-55292099 | 8 | 0.993493681128311 |
| chrX-114283140-114283641 | 2 | 0.914084678210304 |
| chr1-254470876-254471377 | 8 | 0.995194362769321 |
| chr3-28744424-28744925 | 3 | 0.996303221112423 |
| chr3-137835807-137836308 | 3 | 0.996342784619027 |
| chr2-2733015-2733516 | 8 | 0.995228566937176 |

| | | |
|---|---|---|
| chr3-125182440-125182941 | 8 | 0.941964893510429 |
| chr11-14409337-14409838 | 8 | 0.990355171702628 |
| chr19-737479-737980 | 8 | 0.88197369071843 |
| chr10-80439052-80439553 | 3 | 0.907066543621851 |
| chr15-91720114-91720615 | 8 | 0.870851414568552 |
| chr13-105783529-105784030 | 3 | 0.996840128923007 |
| chr17-22320448-22320949 | 6 | 0.991698240220554 |
| chr9-82912243-82912744 | 8 | 0.993493681128311 |
| chr15-621123-621624 | 3 | 0.996869545083467 |
| chr6-129396540-129397041 | 5 | 0.983760760820639 |
| chr18-13705168-13705669 | 6 | 0.972439370380146 |
| chr10-60691876-60692377 | 8 | 0.844411331818381 |
| chr1-255227311-255227812 | 2 | 0.88485803066342 |
| chr14-59317109-59317610 | 5 | 0.970735938731879 |
| chr11-76397750-76398251 | 3 | 0.820911403823185 |
| chr13-6064333-6064834 | 3 | 0.965132819931063 |
| chr15-78161037-78161538 | 8 | 0.993430203457527 |
| chr6-88416200-88416701 | 6 | 0.991206925481043 |
| chr5-137851944-137852445 | 6 | 0.898026622415666 |
| chr10-46680549-46681050 | 4 | 0.869110576444084 |
| chr19-39458486-39458987 | 8 | 0.947608326200666 |
| chr18-24160986-24161487 | 4 | 0.938668879685991 |
| chr4-130396280-130396781 | 8 | 0.874707537880271 |
| chr2-117250283-117250784 | 8 | 0.897745648101029 |
| chr11-35847634-35848135 | 2 | 0.956955222207226 |
| chr1-180898761-180899262 | 8 | 0.977056054521173 |
| chr8-4873493-4873994 | 5 | 0.823818747768795 |
| chr5-38540027-38540528 | 3 | 0.997127707085811 |
| chr10-43785120-43785621 | 3 | 0.990692808411667 |
| chr17-39176758-39177259 | 3 | 0.925415213363492 |
| chr7-42098780-42099281 | 4 | 0.909729435483161 |
| chr2-194598008-194598509 | 8 | 0.880897135264582 |
| chr6-95408064-95408565 | 8 | 0.952816523486839 |
| chr17-41236493-41236994 | 2 | 0.931060685761869 |
| chr1-84302070-84302571 | 6 | 0.992449633272559 |
| chr8-118711774-118712275 | 4 | 0.961492987315662 |
| chr2-111919698-111920199 | 5 | 0.981558979520085 |
| chr9-38591409-38591910 | 8 | 0.839526679842249 |
| chr1-223208045-223208546 | 4 | 0.906443286508524 |
| chr19-39987201-39987702 | 5 | 0.914982144161058 |
| chr6-30613625-30614126 | 6 | 0.991206925481043 |
| chr2-228464693-228465194 | 3 | 0.986745664440927 |
| chr5-122751459-122751960 | 5 | 0.845456841934801 |

| | | |
|---|---|---|
| chr6-88133296-88133797 | 6 | 0.996171180806652 |
| chr4-170304320-170304821 | 8 | 0.900206805054718 |
| chr6-8552675-8553176 | 8 | 0.961932777938224 |
| chr13-23080075-23080576 | 8 | 0.973226769732139 |
| chr3-91719634-91720135 | 8 | 0.990985677478549 |
| chr19-16081343-16081844 | 8 | 0.993493681128311 |
| chr2-41259619-41260120 | 5 | 0.888098864709738 |
| chr14-88429765-88430266 | 6 | 0.917946399894752 |
| chr7-117186899-117187400 | 2 | 0.964241764873765 |
| chr4-141237375-141237876 | 6 | 0.978279896759453 |
| chr4-166700944-166701445 | 2 | 0.819830062268617 |
| chr18-40954438-40954939 | 8 | 0.917685340267828 |
| chr14-50738275-50738776 | 5 | 0.979905406571005 |
| chr4-75580538-75581039 | 5 | 0.981626678842736 |
| chr8-49158714-49159215 | 8 | 0.918814548283071 |
| chr2-195978768-195979269 | 3 | 0.941405735514823 |
| chr10-76554870-76555371 | 6 | 0.962675587603217 |
| chr3-9050602-9051103 | 4 | 0.820266083043931 |
| chr19-38507144-38507645 | 8 | 0.908359678178229 |
| chr14-10441974-10442475 | 8 | 0.993493681128311 |
| chr2-91859564-91860065 | 8 | 0.9327430313272 |
| chr15-80659958-80660459 | 6 | 0.995848799484165 |
| chr18-29787996-29788497 | 3 | 0.846011261699203 |
| chr11-44708276-44708777 | 7 | 0.869173150922018 |
| chr3-79951801-79952302 | 3 | 0.822028007449002 |
| chr20-14173845-14174346 | 7 | 0.84852145423716 |
| chr3-167351492-167351993 | 7 | 0.876912098763271 |
| chr13-104845589-104846090 | 8 | 0.895494114576707 |
| chr3-166820016-166820517 | 8 | 0.820875163839736 |
| chr3-122783708-122784209 | 1 | 0.842143100441906 |
| chr8-9339533-9340034 | 2 | 0.891045104243088 |
| chr4-91053442-91053943 | 5 | 0.819787654152147 |
| chr16-40654880-40655381 | 3 | 0.990162834420557 |
| chr16-83492001-83492502 | 3 | 0.996741400373135 |
| chr7-105590670-105591171 | 5 | 0.990447559538912 |
| chrX-74176333-74176834 | 8 | 0.994651016671484 |
| chr13-40726588-40727089 | 6 | 0.986527644050236 |
| chr6-65687442-65687943 | 3 | 0.99593553545041 |
| chr6-24499471-24499972 | 8 | 0.820254890005412 |
| chr3-135809711-135810212 | 2 | 0.851582659212714 |
| chr10-41257337-41257838 | 5 | 0.92316696625695 |
| chr1-155604908-155605409 | 2 | 0.956937120301164 |
| chr9-81514709-81515210 | 1 | 0.922688058127437 |

| | | |
|---|---|---|
| chrX-31849465-31849966 | 6 | 0.985631130155425 |
| chr18-83374040-83374541 | 6 | 0.958851770119484 |
| chr7-59189950-59190451 | 3 | 0.995943829062555 |
| chr9-96108038-96108539 | 2 | 0.893091422650667 |
| chr11-71164937-71165438 | 8 | 0.952854087899002 |
| chr1-228455834-228456335 | 8 | 0.819450278794113 |
| chr5-73734724-73735225 | 2 | 0.936216885196097 |
| chr2-87026792-87027293 | 5 | 0.963071296525387 |
| chr19-29547904-29548405 | 2 | 0.903147117324518 |
| chr11-71269695-71270196 | 8 | 0.894201760185534 |
| chr17-67364312-67364813 | 8 | 0.965507213091015 |
| chr10-9969959-9970460 | 8 | 0.856977571881207 |
| chr5-160776245-160776746 | 8 | 0.860012647152626 |
| chr4-114934891-114935392 | 1 | 0.860913499091819 |
| chr15-32849743-32850244 | 3 | 0.99593553545041 |
| chr2-46251703-46252204 | 8 | 0.901539950524598 |
| chr16-21583350-21583851 | 2 | 0.8355930249503 |
| chr6-61996291-61996792 | 2 | 0.919265998353953 |
| chr17-79673987-79674488 | 2 | 0.912047085377002 |
| chr20-3237362-3237863 | 8 | 0.993493681128311 |
| chr8-60340279-60340780 | 8 | 0.921236553792329 |
| chr7-29221889-29222390 | 8 | 0.855509190390123 |
| chr4-40833146-40833647 | 6 | 0.991206925481043 |
| chr10-28016998-28017499 | 6 | 0.869776044795146 |
| chr10-57208758-57209259 | 3 | 0.996694775001419 |
| chr7-71403362-71403863 | 5 | 0.983919978840557 |
| chr1-227166001-227166502 | 6 | 0.984617827883341 |
| chr16-31398298-31398799 | 4 | 0.920992899953364 |
| chr18-52032578-52033079 | 8 | 0.891025060080908 |
| chr7-94566225-94566726 | 8 | 0.892271159473177 |
| chr6-52665401-52665902 | 8 | 0.939628939030541 |
| chr17-62489443-62489944 | 3 | 0.99593553545041 |
| chr1-56407016-56407517 | 3 | 0.996882963505963 |
| chr2-134738898-134739399 | 3 | 0.99593553545041 |
| chr18-63789726-63790227 | 4 | 0.904003079842752 |
| chr14-12511496-12511997 | 8 | 0.8151506037921 |
| chr14-33634798-33635299 | 4 | 0.910007068997444 |
| chr7-29913187-29913688 | 4 | 0.944802736030865 |
| chr19-50133333-50133834 | 5 | 0.916764296571127 |
| chr11-81136096-81136597 | 8 | 0.994515968197285 |
| chr12-28014139-28014640 | 8 | 0.930881970059013 |
| chr1-44131947-44132448 | 4 | 0.850147910620854 |
| chr10-19557337-19557838 | 8 | 0.818923434185774 |

| | | |
|---|---|---|
| chr6-10403896-10404397 | 8 | 0.990078543195711 |
| chr10-69412750-69413251 | 8 | 0.914924776238711 |
| chr4-128275779-128276280 | 3 | 0.998285365055408 |
| chr12-22200173-22200674 | 8 | 0.963314667370865 |
| chr15-91423567-91424068 | 3 | 0.996871869827778 |
| chr1-8079292-8079793 | 8 | 0.963476880120105 |
| chr2-104310914-104311415 | 8 | 0.930956743982395 |
| chr19-38640095-38640596 | 5 | 0.942095768400419 |
| chr8-84128701-84129202 | 8 | 0.993074118526785 |
| chr6-131161309-131161810 | 8 | 0.991419054733034 |
| chr4-117772754-117773255 | 8 | 0.995097646812465 |
| chr3-42792319-42792820 | 3 | 0.997366551497976 |
| chr1-249606214-249606715 | 8 | 0.886430669341536 |
| chr4-126446297-126446798 | 8 | 0.869818564466856 |
| chr12-28447835-28448336 | 6 | 0.99189772873004 |
| chr9-63410046-63410547 | 7 | 0.945055821762391 |
| chr1-54081585-54082086 | 2 | 0.824492591110497 |
| chr4-65419095-65419596 | 3 | 0.960158494601362 |
| chrX-13502924-13503425 | 5 | 0.857401965474477 |
| chr12-33716307-33716808 | 4 | 0.94943990486077 |
| chr18-78229439-78229940 | 4 | 0.967569278916578 |
| chr4-115516759-115517260 | 8 | 0.935522907638001 |
| chr14-101013318-101013819 | 8 | 0.858766721927571 |
| chr5-45773638-45774139 | 7 | 0.901799771377597 |
| chrX-16168823-16169324 | 3 | 0.873780233862627 |
| chr4-175841613-175842114 | 5 | 0.861063230699203 |
| chr11-4416824-4417325 | 8 | 0.982104897971635 |
| chr20-51417754-51418255 | 3 | 0.99593553545041 |
| chr8-76118067-76118568 | 4 | 0.914761573460971 |
| chr10-59126581-59127082 | 3 | 0.996031578380205 |
| chr20-22966371-22966872 | 8 | 0.961468619550836 |
| chr5-121091184-121091685 | 4 | 0.816242017293351 |
| chr8-26302101-26302602 | 3 | 0.993737872562967 |
| chr8-75513679-75514180 | 6 | 0.989935270353471 |
| chr2-166497777-166498278 | 3 | 0.995621071471796 |
| chrX-94859554-94860055 | 6 | 0.990283243943055 |
| chr15-21252176-21252677 | 8 | 0.854945758531632 |
| chr13-82330272-82330773 | 4 | 0.917581938693309 |
| chr18-3860990-3861491 | 7 | 0.908998644514309 |
| chr7-27873782-27874283 | 1 | 0.975896873560983 |
| chr5-20462373-20462874 | 1 | 0.830399138026086 |
| chr2-61401500-61402001 | 8 | 0.919340087483374 |
| chr19-15906161-15906662 | 6 | 0.991544365477866 |

| | | |
|---|---|---|
| chr3-76091972-76092473 | 3 | 0.924408527703355 |
| chr9-30582063-30582564 | 3 | 0.984477606541983 |
| chr8-104489719-104490220 | 6 | 0.978746103847445 |
| chr10-105584864-105585365 | 5 | 0.996576381084485 |
| chr18-39570454-39570955 | 8 | 0.989797691538155 |
| chr16-38620703-38621204 | 6 | 0.988453358907536 |
| chr10-64462883-64463384 | 8 | 0.990974181818075 |
| chr8-23217669-23218170 | 5 | 0.973725777849207 |
| chr11-8774786-8775287 | 2 | 0.85299659137152 |
| chr6-3058906-3059407 | 6 | 0.982817979448901 |
| chr3-90520166-90520667 | 8 | 0.885982779592925 |
| chr20-25148386-25148887 | 3 | 0.871440073827718 |
| chr3-26639023-26639524 | 8 | 0.95525597076737 |
| chr10-87934466-87934967 | 8 | 0.98657966539428 |
| chr2-125136105-125136606 | 8 | 0.91875360102071 |
| chr1-251723238-251723739 | 1 | 0.949742527057987 |
| chr20-53849296-53849797 | 4 | 0.853417883469609 |
| chrX-22021422-22021923 | 7 | 0.960235812340116 |
| chr3-108107713-108108214 | 8 | 0.967876914318511 |
| chr10-62221520-62222021 | 8 | 0.98487603795137 |
| chr1-47299254-47299755 | 5 | 0.965183328421706 |
| chr14-78964931-78965432 | 8 | 0.819208859865944 |
| chr2-175259104-175259605 | 2 | 0.972876728587985 |
| chr15-19137257-19137758 | 2 | 0.950731626576375 |
| chr1-130813545-130814046 | 3 | 0.86074716303293 |
| chr5-60292506-60293007 | 8 | 0.964378084240513 |
| chr4-94724855-94725356 | 2 | 0.869755225450118 |
| chr4-105087506-105088007 | 2 | 0.854167960771094 |
| chr19-15933551-15934052 | 5 | 0.981999543066056 |
| chr18-7946272-7946773 | 6 | 0.921179420737455 |
| chr8-75286331-75286832 | 8 | 0.86066494651011 |
| chr10-40071750-40072251 | 8 | 0.983074949715127 |
| chr13-1584190-1584691 | 8 | 0.879741176728092 |
| chr2-136465778-136466279 | 8 | 0.979344946713488 |
| chr1-164497762-164498263 | 5 | 0.986747830317764 |
| chr1-189300191-189300692 | 4 | 0.950878481079921 |
| chr7-41536047-41536548 | 2 | 0.942636115863784 |
| chr20-50575466-50575967 | 3 | 0.995034678856006 |
| chr13-97008535-97009036 | 8 | 0.995219050663703 |
| chr14-79925571-79926072 | 8 | 0.890290510209716 |
| chr2-101506276-101506777 | 6 | 0.983931723605695 |
| chr8-47572771-47573272 | 5 | 0.935157549217605 |
| chr9-79217037-79217538 | 1 | 0.855313355794525 |

| | | |
|---|---|---|
| chr2-125248143-125248644 | 8 | 0.98033829250875 |
| chrX-47768957-47769458 | 4 | 0.826562525646756 |
| chr1-55769729-55770230 | 6 | 0.99246364086225 |
| chr11-20043430-20043931 | 8 | 0.993493681128311 |
| chr2-60655326-60655827 | 3 | 0.99593553545041 |
| chr3-9274459-9274960 | 2 | 0.852761450550064 |
| chr3-89651840-89652341 | 2 | 0.865266500293717 |
| chr11-11592913-11593414 | 3 | 0.99686641690099 |
| chr7-20077630-20078131 | 5 | 0.989718881133111 |
| chr5-21583786-21584287 | 5 | 0.951554704161067 |
| chr19-15994566-15995067 | 5 | 0.898665818442597 |
| chr1-133043973-133044474 | 2 | 0.836696589264493 |
| chr4-158392344-158392845 | 2 | 0.923341460245207 |
| chr1-234598729-234599230 | 5 | 0.897944831628718 |
| chr6-120189704-120190205 | 6 | 0.987639905223378 |
| chr13-78556556-78557057 | 3 | 0.996270586874103 |
| chr17-83088983-83089484 | 4 | 0.91222410075707 |
| chr4-57154561-57155062 | 5 | 0.825041138747491 |
| chr7-60648009-60648510 | 3 | 0.99593553545041 |
| chr1-190431526-190432027 | 6 | 0.911071452654699 |
| chr4-57218278-57218779 | 6 | 0.993221807934325 |
| chr3-13621609-13622110 | 8 | 0.96913811524025 |
| chr2-7900826-7901327 | 5 | 0.982857577587536 |
| chr7-51956704-51957205 | 3 | 0.986551978284141 |
| chr5-103534907-103535408 | 3 | 0.973490975553162 |
| chr9-74405226-74405727 | 8 | 0.980846514016873 |
| chr6-134648419-134648920 | 7 | 0.884561348908504 |
| chr1-153682383-153682884 | 7 | 0.932942823989645 |
| chr1-15660369-15660870 | 4 | 0.959658119951112 |
| chr15-49061691-49062192 | 8 | 0.991120618002126 |
| chr15-30452945-30453446 | 6 | 0.96845250740431 |
| chr4-48170981-48171482 | 3 | 0.994325038377665 |
| chr4-168935272-168935773 | 4 | 0.891680227817267 |
| chr15-40381666-40382167 | 8 | 0.84505649459886 |
| chr11-51272676-51273177 | 6 | 0.992763696056331 |
| chr10-33922299-33922800 | 7 | 0.89916281587944 |
| chr16-52581636-52582137 | 8 | 0.861802743309125 |
| chr13-36530910-36531411 | 8 | 0.993493681128311 |
| chr7-14273367-14273868 | 8 | 0.894848589421089 |
| chr7-113054373-113054874 | 5 | 0.902630232131875 |
| chr15-80573818-80574319 | 1 | 0.969249165027251 |
| chr6-71783192-71783693 | 1 | 0.879088541363728 |
| chr19-16055533-16056034 | 5 | 0.984462148053118 |

| | | |
|---|---|---|
| chr15-68811015-68811516 | 3 | 0.995935535545041 |
| chr19-53379136-53379637 | 5 | 0.962264553572301 |
| chr6-2169663-2170164 | 4 | 0.876825835911018 |
| chr12-155846-156347 | 8 | 0.995124781211608 |
| chr14-22256996-22257497 | 3 | 0.9903889327219 |
| chr14-11806431-11806932 | 8 | 0.859586818891976 |
| chr2-184327531-184328032 | 4 | 0.956433797798175 |
| chr20-18049302-18049803 | 5 | 0.868140392045972 |
| chr18-8011430-8011931 | 8 | 0.988812586270484 |
| chr3-145857896-145858397 | 5 | 0.939987451258792 |
| chr16-72919614-72920115 | 6 | 0.991206925481043 |
| chr4-121405721-121406222 | 5 | 0.864244100050065 |
| chr6-38561735-38562236 | 8 | 0.975059235114025 |
| chr13-39332586-39333087 | 3 | 0.990594048652778 |
| chr4-45628355-45628856 | 6 | 0.867031821272917 |
| chr16-66618384-66618885 | 5 | 0.992214578365133 |
| chr7-17885984-17886485 | 8 | 0.887120794153428 |
| chr1-254228116-254228617 | 8 | 0.993493681128311 |
| chr13-39432936-39433437 | 8 | 0.876102293614601 |
| chr19-11005363-11005864 | 4 | 0.876218032418701 |
| chr4-71078113-71078614 | 8 | 0.919008360460083 |
| chr13-86726228-86726729 | 8 | 0.89392562654009 |
| chr9-91792003-91792504 | 4 | 0.867076995081456 |
| chr7-104989411-104989912 | 8 | 0.991991725159938 |
| chr2-43425853-43426354 | 3 | 0.989817215173353 |
| chr20-13397362-13397863 | 7 | 0.899508417104251 |
| chr2-44787685-44788186 | 6 | 0.991260942200011 |
| chr1-153800373-153800874 | 8 | 0.960242451415366 |
| chr17-41028263-41028764 | 4 | 0.921116538670516 |
| chr11-66625088-66625589 | 4 | 0.868701600529354 |
| chr3-166411059-166411560 | 8 | 0.993493681128311 |
| chr3-142840612-142841113 | 5 | 0.989502657170865 |
| chr3-65282366-65282867 | 4 | 0.930553406278818 |
| chr3-1479464-1479965 | 2 | 0.909827744583116 |
| chr13-46289799-46290300 | 5 | 0.978295149590994 |
| chr1-173138617-173139118 | 6 | 0.898035152749252 |
| chr15-1511920-1512421 | 4 | 0.973054721615879 |
| chr9-107410895-107411396 | 5 | 0.98900388839453 |
| chr1-55772287-55772788 | 6 | 0.989567485256928 |
| chr17-51104446-51104947 | 5 | 0.869338763280963 |
| chr11-40417636-40418137 | 3 | 0.978107133274072 |
| chr1-212055216-212055717 | 6 | 0.957396826599093 |
| chr12-13990244-13990745 | 8 | 0.839473095211823 |

| | | |
|---|---|---|
| chr9-65563595-65564096 | 2 | 0.967144310586227 |
| chr16-34187420-34187921 | 6 | 0.870332113683559 |
| chr1-60267629-60268130 | 3 | 0.952120726314015 |
| chr7-111563013-111563514 | 5 | 0.946692807529131 |
| chr20-27117693-27118194 | 6 | 0.97009568544115 |
| chr16-42752831-42753332 | 3 | 0.99593553545041 |
| chr4-108494174-108494675 | 2 | 0.879627339697587 |
| chr16-13818798-13819299 | 6 | 0.972434140373252 |
| chr6-104284619-104285120 | 8 | 0.93503986582372 |
| chr2-224731418-224731919 | 5 | 0.869721812991647 |
| chr18-62129291-62129792 | 1 | 0.938846984099003 |
| chr19-54401514-54402015 | 5 | 0.994286234091894 |
| chr8-12737422-12737923 | 8 | 0.823083075940269 |
| chr2-123378002-123378503 | 4 | 0.915267052637378 |
| chr14-18459558-18460059 | 8 | 0.971572819490881 |
| chr11-17426443-17426944 | 4 | 0.954148973029963 |
| chr18-32174672-32175173 | 1 | 0.91138650864361 |
| chr3-6757949-6758450 | 6 | 0.843369691133198 |
| chr15-23812781-23813282 | 3 | 0.915473745748007 |
| chr1-44954370-44954871 | 8 | 0.991023793463436 |
| chr8-12274779-12275280 | 1 | 0.852297210566831 |
| chr2-231401513-231402014 | 3 | 0.996892637168296 |
| chr13-70975278-70975779 | 8 | 0.915666878827244 |
| chr15-76846883-76847384 | 7 | 0.862633088482085 |
| chr16-16006757-16007258 | 3 | 0.99593553545041 |
| chr8-66843146-66843647 | 8 | 0.864072311227155 |
| chr3-32848036-32848537 | 7 | 0.828718894983622 |
| chr6-124217554-124218055 | 8 | 0.825624784192106 |
| chrX-65889227-65889728 | 3 | 0.949294896147836 |
| chr6-131255237-131255738 | 3 | 0.88206779839883 |
| chr2-81849099-81849600 | 8 | 0.914283088942529 |
| chr1-110223589-110224090 | 3 | 0.996633542416283 |
| chr9-113319713-113320214 | 3 | 0.996054287676872 |
| chr18-1748858-1749359 | 3 | 0.916646009970009 |
| chr1-179118411-179118912 | 8 | 0.939861197670428 |
| chr15-11971602-11972103 | 5 | 0.981899455773227 |
| chr2-210323508-210324009 | 8 | 0.987713205507018 |
| chr14-2387343-2387844 | 2 | 0.954585875645695 |
| chr16-16708544-16709045 | 6 | 0.995948401889468 |
| chrX-147954807-147955308 | 8 | 0.993493681128311 |
| chr1-235245417-235245918 | 7 | 0.818570433902864 |
| chr8-50418993-50419494 | 3 | 0.990016105139205 |
| chr1-122039446-122039947 | 6 | 0.918693678603394 |

| | | |
|---|---|---|
| chr12-12052905-12053406 | 8 | 0.930386152007407 |
| chr15-48451657-48452158 | 8 | 0.992894726138898 |
| chr4-40929595-40930096 | 7 | 0.962677706241405 |
| chr18-24642591-24643092 | 6 | 0.949877568118708 |
| chr15-8233606-8234107 | 8 | 0.953995502974095 |
| chr14-13506743-13507244 | 2 | 0.925497720659152 |
| chr6-99042055-99042556 | 3 | 0.966202660369689 |
| chr17-81642168-81642669 | 3 | 0.990469072638263 |
| chr4-59317222-59317723 | 7 | 0.908736166847706 |
| chr14-93084662-93085163 | 6 | 0.99663128700614 |
| chr7-131791731-131792232 | 4 | 0.928055563704879 |
| chr11-55434831-55435332 | 4 | 0.861395952996685 |
| chrX-104342889-104343390 | 6 | 0.979845831042333 |
| chr7-114214144-114214645 | 7 | 0.932372884053451 |
| chr10-8348521-8349022 | 3 | 0.963934590243777 |
| chr9-21082566-21083067 | 4 | 0.861148164590802 |
| chr9-15691976-15692477 | 8 | 0.923260944088147 |
| chr11-24660022-24660523 | 3 | 0.916904133780168 |
| chr8-31582359-31582860 | 8 | 0.879114699160329 |
| chr8-102981361-102981862 | 3 | 0.996191228330091 |
| chr12-12078981-12079482 | 8 | 0.929971861125401 |
| chr20-2812731-2813232 | 5 | 0.992201841932777 |
| chr11-44154868-44155369 | 2 | 0.91874312284672 |
| chr17-82032051-82032552 | 8 | 0.993493681128311 |
| chr7-33963084-33963585 | 8 | 0.92596826276959 |
| chr3-78628831-78629332 | 8 | 0.889161498237186 |
| chr17-22703587-22704088 | 8 | 0.830510987283023 |
| chr8-28838188-28838689 | 8 | 0.865849893071202 |
| chr13-74111457-74111958 | 8 | 0.988526752346195 |
| chr13-82446612-82447113 | 5 | 0.82351612567166 |
| chr10-89162158-89162659 | 4 | 0.844047086210803 |
| chr1-120095331-120095832 | 8 | 0.989097478986817 |
| chr19-42944973-42945474 | 8 | 0.90122799596632 |
| chr5-110089128-110089629 | 6 | 0.993338363814715 |
| chr14-92114738-92115239 | 3 | 0.996855996153142 |
| chr2-120346405-120346906 | 8 | 0.993493681128311 |
| chr1-30343965-30344466 | 3 | 0.902075177418304 |
| chr17-39525029-39525530 | 6 | 0.989576413853067 |
| chr8-104750649-104751150 | 5 | 0.977391337903346 |
| chr2-240406039-240406540 | 1 | 0.885856363092775 |
| chr15-34086122-34086623 | 6 | 0.817913206184214 |
| chr12-5478560-5479061 | 8 | 0.90377079780038 |
| chr15-4184569-4185070 | 7 | 0.958450601106406 |

| | | |
|---|---|---|
| chrY-14117522-14118023 | 3 | 0.935059252386903 |
| chr17-80515206-80515707 | 5 | 0.978861840549482 |
| chr14-103021837-103022338 | 6 | 0.975853749010095 |
| chr16-73892268-73892769 | 6 | 0.993214152088585 |
| chr9-109616485-109616986 | 8 | 0.945413135660372 |
| chr3-157870593-157871094 | 8 | 0.981074975279802 |
| chr1-134171956-134172457 | 8 | 0.983677642536834 |
| chr8-77384509-77385010 | 6 | 0.991942647182129 |
| chr5-8411070-8411571 | 3 | 0.988536062287362 |
| chr8-114013427-114013928 | 8 | 0.851816139279638 |
| chr7-80912460-80912961 | 3 | 0.99593553545041 |
| chr1-34872853-34873354 | 6 | 0.992950680175248 |
| chr3-119545018-119545519 | 3 | 0.993101914060713 |
| chr9-98518118-98518619 | 2 | 0.877920578991674 |
| chr8-92673756-92674257 | 8 | 0.881255289206765 |
| chr17-5312465-5312966 | 8 | 0.922952059014854 |
| chr1-117829331-117829832 | 8 | 0.99477988177044 |
| chr5-60135668-60136169 | 5 | 0.957043933711373 |
| chr3-79549880-79550381 | 2 | 0.94588905000073 |
| chr11-71093340-71093841 | 8 | 0.952161851381489 |
| chr9-14563434-14563935 | 8 | 0.891154387744514 |
| chr9-82780232-82780733 | 8 | 0.995145039022299 |
| chr8-95648341-95648842 | 2 | 0.834218221157026 |
| chr4-33463642-33464143 | 2 | 0.860790910520593 |
| chr5-116081469-116081970 | 4 | 0.972910924199089 |
| chr18-44512622-44513123 | 6 | 0.987911061595736 |
| chr7-127227010-127227511 | 4 | 0.932364085987641 |
| chr10-15552012-15552513 | 7 | 0.921034277648718 |
| chr1-167622916-167623417 | 1 | 0.915698881992792 |
| chr5-83660825-83661326 | 4 | 0.951365145639644 |
| chr18-67151796-67152297 | 1 | 0.928860015897061 |
| chr20-46851039-46851540 | 5 | 0.993967023421824 |
| chr20-44932561-44933062 | 3 | 0.921685777620217 |
| chr17-7915809-7916310 | 3 | 0.828774701735015 |
| chr5-55883065-55883566 | 8 | 0.867271845702672 |
| chr6-19621650-19622151 | 8 | 0.832722519947792 |
| chr15-14786019-14786520 | 8 | 0.895478602068819 |
| chr1-192517480-192517981 | 8 | 0.878257010131787 |
| chr13-85692116-85692617 | 6 | 0.864779356325992 |
| chr19-41883812-41884313 | 8 | 0.928096514146776 |
| chr3-78600865-78601366 | 2 | 0.869762847019113 |
| chr2-33460127-33460628 | 8 | 0.993493681128311 |
| chr13-102012683-102013184 | 5 | 0.983425802318927 |

| | | |
|---|---|---|
| chr10-1314170-1314671 | 8 | 0.902267095636397 |
| chr10-94162466-94162967 | 4 | 0.948446204824266 |
| chr4-117991292-117991793 | 2 | 0.865703670263938 |
| chr8-57467875-57468376 | 8 | 0.964936590827269 |
| chr8-53733548-53734049 | 8 | 0.98455103056203 |
| chr1-31012634-31013135 | 8 | 0.876353580882893 |
| chr1-6379560-6380061 | 8 | 0.994985080610156 |
| chr4-36318162-36318663 | 8 | 0.974281128951541 |
| chr14-14038824-14039325 | 3 | 0.992112646533092 |
| chr10-41629328-41629829 | 7 | 0.882847403837905 |
| chr10-39683673-39684174 | 4 | 0.85105158909143 |
| chr1-4600937-4601438 | 5 | 0.914753073841265 |
| chr7-14568224-14568725 | 6 | 0.985801621027224 |
| chr17-70156105-70156606 | 2 | 0.962126065840543 |
| chr8-47261024-47261525 | 8 | 0.994666190910751 |
| chr9-59086303-59086804 | 5 | 0.986579374936707 |
| chr3-99220263-99220764 | 3 | 0.99321156170698 |
| chr18-55452111-55452612 | 2 | 0.926818456107688 |
| chr6-85407812-85408313 | 5 | 0.991733153331352 |
| chr4-18917026-18917527 | 6 | 0.991604992499129 |
| chr6-21572381-21572882 | 3 | 0.989999434895309 |
| chr8-29746490-29746991 | 8 | 0.968243451985951 |
| chr1-29676809-29677310 | 5 | 0.978262199562449 |
| chr16-36653998-36654499 | 6 | 0.855980836928336 |
| chr16-4529495-4529996 | 8 | 0.851472276907342 |
| chr4-76294655-76295156 | 3 | 0.996678488748895 |
| chr17-3204247-3204748 | 3 | 0.99593553545041 |
| chr2-141916255-141916756 | 3 | 0.994867313264439 |
| chr5-58061226-58061727 | 2 | 0.950509866104418 |
| chr4-13822172-13822673 | 8 | 0.990454005682935 |
| chr7-55839555-55840056 | 5 | 0.965028357992631 |
| chr19-3131678-3132179 | 3 | 0.9966470254186 |
| chr13-41538171-41538672 | 8 | 0.969198868870545 |
| chr4-9007207-9007708 | 6 | 0.94371290757631 |
| chr4-125758476-125758977 | 1 | 0.904427872559977 |
| chr6-88442025-88442526 | 6 | 0.985246028483527 |
| chr4-62431651-62432152 | 3 | 0.853832686181085 |
| chr3-22376074-22376575 | 8 | 0.910368176870827 |
| chr8-95728205-95728706 | 1 | 0.89972143400334 |
| chr2-219222438-219222939 | 8 | 0.984671217200747 |
| chr3-140129014-140129515 | 8 | 0.821223608641827 |
| chr12-7272180-7272681 | 5 | 0.837002954015163 |
| chr4-133806752-133807253 | 8 | 0.948940037577856 |

| | | |
|---|---|---|
| chr18-15438791-15439292 | 8 | 0.904706253765955 |
| chr11-34170063-34170564 | 4 | 0.957399248670789 |
| chr3-165516950-165517451 | 8 | 0.982898207971099 |
| chr5-106843348-106843849 | 4 | 0.843720558420071 |
| chr15-96048173-96048674 | 8 | 0.975694184461488 |
| chr13-56786403-56786904 | 3 | 0.993970449640953 |
| chr5-33369171-33369672 | 3 | 0.995719472103418 |
| chr14-57039798-57040299 | 1 | 0.843154029813067 |
| chr6-71240286-71240787 | 8 | 0.866247732513017 |
| chr6-8442334-8442835 | 5 | 0.969273879498112 |
| chr1-193482490-193482991 | 3 | 0.99593553545041 |
| chr4-88614765-88615266 | 3 | 0.997213524704538 |
| chr9-761472-761973 | 3 | 0.994986657140126 |
| chr1-30524226-30524727 | 8 | 0.858194403395488 |
| chr1-233136725-233137226 | 4 | 0.95520785840816 |
| chr11-84782631-84783132 | 8 | 0.974783704509271 |
| chr2-40647513-40648014 | 7 | 0.925816106682375 |
| chr14-102027329-102027830 | 6 | 0.96715205162991 |
| chr2-118856255-118856756 | 8 | 0.956888038105375 |
| chr1-77252147-77252648 | 1 | 0.937724678051087 |
| chr1-163416053-163416554 | 8 | 0.992659495367506 |
| chr6-97286241-97286742 | 8 | 0.842806028939707 |
| chr3-122855275-122855776 | 3 | 0.927962148426953 |
| chr15-92142373-92142874 | 7 | 0.91710837871614 |
| chr6-42635734-42636235 | 6 | 0.897364730597004 |
| chr20-24314393-24314894 | 3 | 0.894729539901099 |
| chr13-75882059-75882560 | 8 | 0.993493681128311 |
| chr7-75885583-75886084 | 1 | 0.910455818496809 |
| chr5-37880553-37881054 | 8 | 0.941866079887528 |
| chr14-83138845-83139346 | 5 | 0.840261636059002 |
| chr7-59244464-59244965 | 5 | 0.862547488997356 |
| chr6-130726038-130726539 | 8 | 0.916829736051384 |
| chr6-66051014-66051515 | 3 | 0.994692471711501 |
| chr5-170164-170665 | 3 | 0.979283874975892 |
| chr9-86056247-86056748 | 6 | 0.979421407538157 |
| chr5-37321065-37321566 | 8 | 0.902517869588449 |
| chr7-84282090-84282591 | 2 | 0.929827230213962 |
| chr10-44972719-44973220 | 8 | 0.952877793602173 |
| chr16-45970558-45971059 | 8 | 0.97330805960441 |
| chr2-25854985-25855486 | 5 | 0.843978816707856 |
| chr4-79913917-79914418 | 2 | 0.940417108691721 |
| chr16-64846184-64846685 | 8 | 0.951652389305956 |
| chrX-138460249-138460750 | 6 | 0.990187259795704 |

| | | |
|---|---|---|
| chr7-66598474-66598975 | 8 | 0.91543623163189 |
| chr1-196883630-196884131 | 8 | 0.982774120501658 |
| chr11-50834922-50835423 | 1 | 0.981660086188443 |
| chr15-95187561-95188062 | 5 | 0.950143018398277 |
| chrX-72079034-72079535 | 2 | 0.880674595014058 |
| chr4-18569439-18569940 | 8 | 0.98968885166961 |
| chr7-115021977-115022478 | 3 | 0.99708679447523 |
| chr19-22506494-22506995 | 3 | 0.988282371518855 |
| chr15-9333193-9333694 | 3 | 0.99593553545041 |
| chr16-79599723-79600224 | 6 | 0.923835046827014 |
| chrX-67770730-67771231 | 6 | 0.890464622731565 |
| chr18-6186540-6187041 | 8 | 0.943196528290201 |
| chr17-46921620-46922121 | 5 | 0.861326558027856 |
| chr2-246295759-246296260 | 3 | 0.996850060659576 |
| chrX-12187365-12187866 | 8 | 0.993381746361166 |
| chr9-40303602-40304103 | 7 | 0.955018510856572 |
| chr2-51329169-51329670 | 8 | 0.94704580349965 |
| chr6-51275793-51276294 | 8 | 0.903596428818029 |
| chr9-41002943-41003444 | 8 | 0.908638247698924 |
| chr2-82637176-82637677 | 4 | 0.855538036743124 |
| chr14-93280057-93280558 | 5 | 0.984150524704159 |
| chr15-92327296-92327797 | 6 | 0.93625916713532 |
| chrX-93254576-93255077 | 5 | 0.983919978840557 |
| chr4-97485734-97486235 | 3 | 0.99199586779674 |
| chr2-143548037-143548538 | 3 | 0.994760679084958 |
| chr6-89905283-89905784 | 8 | 0.838987444892305 |
| chr2-173159398-173159899 | 2 | 0.910805999240764 |
| chr3-112274460-112274961 | 5 | 0.990314775340657 |
| chr17-6715741-6716242 | 4 | 0.972436163725079 |
| chr16-44998169-44998670 | 8 | 0.865297061378665 |
| chr10-92988051-92988552 | 8 | 0.822516589155296 |
| chr14-104720257-104720758 | 3 | 0.99593553545041 |
| chr11-79992090-79992591 | 8 | 0.948999822781344 |
| chrX-125007790-125008291 | 6 | 0.982526620499194 |
| chr9-85428713-85429214 | 8 | 0.995465933111868 |
| chr14-53219938-53220439 | 3 | 0.998322087973705 |
| chr6-77532945-77533446 | 5 | 0.884257319041539 |
| chr7-109225803-109226304 | 3 | 0.930798719751482 |
| chr12-6275609-6276110 | 3 | 0.989478289573968 |
| chr5-57890128-57890629 | 4 | 0.868368574535663 |
| chr16-62694746-62695247 | 6 | 0.993156490444852 |
| chr15-32427054-32427555 | 3 | 0.997224872208587 |
| chr18-74229212-74229713 | 8 | 0.908566096137741 |

| | | |
|---|---|---|
| chr6-45507159-45507660 | 8 | 0.926327337421749 |
| chr4-173238007-173238508 | 8 | 0.993493681128311 |
| chr8-121451597-121452098 | 8 | 0.971824352126964 |
| chr15-80952400-80952901 | 3 | 0.995674404898769 |
| chr3-46274925-46275426 | 4 | 0.880305033935329 |
| chr2-103987824-103988325 | 3 | 0.81755689851358 |
| chr5-161713243-161713744 | 4 | 0.898344205244065 |
| chr7-105346555-105347056 | 6 | 0.92539240788289 |
| chr17-34777848-34778349 | 8 | 0.837690121442716 |
| chr1-214555440-214555941 | 1 | 0.820254778192202 |
| chr10-48701616-48702117 | 5 | 0.886950410665326 |
| chr2-30726615-30727116 | 6 | 0.988458739509076 |
| chr6-92271594-92272095 | 8 | 0.969308 |
| chr4-83639121-83639622 | 5 | 0.986150864736088 |
| chr9-33228004-33228505 | 5 | 0.847099927578533 |
| chr11-55543124-55543625 | 8 | 0.852031450606427 |
| chrX-25040307-25040808 | 8 | 0.993103252173188 |
| chr2-95993602-95994103 | 3 | 0.99593553545041 |
| chr2-48310171-48310672 | 8 | 0.845127807182848 |
| chr10-19156139-19156640 | 3 | 0.99593553545041 |
| chr18-12216708-12217209 | 2 | 0.858071616960734 |
| chr3-16444697-16445198 | 8 | 0.994490570953179 |
| chr3-7957865-7958366 | 7 | 0.834018900908553 |
| chr5-148854203-148854704 | 4 | 0.823502486523847 |
| chr8-96952797-96953298 | 1 | 0.820978932943141 |
| chr19-48386830-48387331 | 8 | 0.86712410801424 |
| chr17-56034196-56034697 | 3 | 0.995812438201949 |
| chr6-6386300-6386801 | 6 | 0.986800042552231 |
| chr6-86859959-86860460 | 3 | 0.996326572760017 |
| chr6-29829981-29830482 | 6 | 0.97952698884729 |
| chr2-134625113-134625614 | 5 | 0.863656998633966 |
| chr10-4011678-4012179 | 8 | 0.943462392855214 |
| chr8-62531739-62532240 | 8 | 0.925327945819004 |
| chr3-705952-706453 | 4 | 0.89179145837541 |
| chr15-75157097-75157598 | 2 | 0.966683006937966 |
| chr20-11958122-11958623 | 8 | 0.97996207149489 |
| chr11-80510246-80510747 | 8 | 0.987004436400685 |
| chr7-98961040-98961541 | 5 | 0.990200548884499 |
| chr4-155956545-155957046 | 8 | 0.923477491657334 |
| chr20-19709814-19710315 | 8 | 0.826028584263336 |
| chr10-107077852-107078353 | 8 | 0.965977348675453 |
| chr4-159105765-159106266 | 3 | 0.995025624475677 |
| chr11-33644678-33645179 | 5 | 0.835641559186892 |

| | | |
|---|---|---|
| chr9-62668164-62668665 | 2 | 0.884647421685563 |
| chr4-80633371-80633872 | 8 | 0.875924042260775 |
| chr4-164431865-164432366 | 3 | 0.995920812742515 |
| chr17-50165397-50165898 | 3 | 0.99593553545041 |
| chr14-41288959-41289460 | 7 | 0.887480457861797 |
| chr5-132873349-132873850 | 8 | 0.819151154422549 |
| chr2-120149361-120149862 | 8 | 0.895634901317304 |
| chr15-41040924-41041425 | 2 | 0.868528629080986 |
| chr11-55534742-55535243 | 8 | 0.890153258036728 |
| chr3-111737911-111738412 | 8 | 0.98775500589132 |
| chr13-66192749-66193250 | 8 | 0.924418228863833 |
| chr19-4847434-4847935 | 3 | 0.976178012784523 |
| chr20-49401698-49402199 | 3 | 0.99593553545041 |
| chr9-6913123-6913624 | 8 | 0.841214269015949 |
| chr20-4343546-4344047 | 8 | 0.966162851896389 |
| chr19-43874651-43875152 | 4 | 0.947036773739731 |
| chr11-16365816-16366317 | 5 | 0.920237201940669 |
| chr2-37420462-37420963 | 8 | 0.851003898101351 |
| chr9-107724128-107724629 | 8 | 0.819534377386284 |
| chr18-72051445-72051946 | 3 | 0.947667650800982 |
| chr8-92610870-92611371 | 7 | 0.909778167122747 |
| chr1-76587877-76588378 | 3 | 0.988024693176818 |
| chr20-51598833-51599334 | 3 | 0.978358919021395 |
| chr3-150839780-150840281 | 8 | 0.942143010003739 |
| chr1-26867733-26868234 | 8 | 0.968297251846551 |
| chr2-217074231-217074732 | 4 | 0.97126833501892 |
| chr14-32907966-32908467 | 5 | 0.980423388160036 |
| chr16-59885411-59885912 | 7 | 0.944578161496627 |
| chr5-18665226-18665727 | 2 | 0.956996889715028 |
| chr4-80875229-80875730 | 4 | 0.974687489886794 |
| chr10-19164152-19164653 | 3 | 0.99593553545041 |
| chr3-56512011-56512512 | 8 | 0.875385212982554 |
| chr14-81366462-81366963 | 8 | 0.986447295632543 |
| chr17-32602763-32603264 | 8 | 0.822975972178232 |
| chrX-135211715-135212216 | 6 | 0.991206925481043 |
| chr13-92998468-92998969 | 8 | 0.946078908234773 |
| chr7-27704179-27704680 | 8 | 0.856413297393517 |
| chr13-89329395-89329896 | 6 | 0.997195958950105 |
| chr20-25453759-25454260 | 3 | 0.991570535720291 |
| chr3-49893126-49893627 | 8 | 0.844558719820813 |
| chr4-154097297-154097798 | 2 | 0.858394919327686 |
| chr16-67457749-67458250 | 6 | 0.993329500844071 |
| chr18-70295552-70296053 | 4 | 0.888519661658738 |

| | | |
|---|---|---|
| chr11-51778817-51779318 | 3 | 0.99593553545041 |
| chr1-82413353-82413854 | 8 | 0.833797144007412 |
| chr9-17608343-17608844 | 3 | 0.834637343521901 |
| chr8-68829850-68830351 | 8 | 0.879736008073292 |
| chr10-79036102-79036603 | 2 | 0.96493424517026 |
| chr1-60755885-60756386 | 6 | 0.993163534620026 |
| chr1-182220899-182221400 | 8 | 0.909680898719492 |
| chr3-92618482-92618983 | 3 | 0.986996209479831 |
| chr8-45824626-45825127 | 8 | 0.894758238343397 |
| chr16-66666199-66666700 | 8 | 0.915211208005835 |
| chr16-4794296-4794797 | 4 | 0.935288145847828 |
| chr15-786283-786784 | 8 | 0.964398077310214 |
| chrX-40005537-40006038 | 6 | 0.991206925481043 |
| chr19-19589874-19590375 | 3 | 0.992272671405356 |
| chr19-29397405-29397906 | 8 | 0.988864304831906 |
| chr5-146720399-146720900 | 8 | 0.938634237313809 |
| chr9-43517420-43517921 | 4 | 0.905610169447826 |
| chr2-58406530-58407031 | 8 | 0.993493681128311 |
| chr4-142056099-142056600 | 8 | 0.993493681128311 |
| chr14-58346387-58346888 | 4 | 0.904194198743264 |
| chr7-28751247-28751748 | 8 | 0.986712275465418 |
| chr14-58224084-58224585 | 3 | 0.991729547241679 |
| chr4-5610099-5610600 | 8 | 0.978526770970116 |
| chr3-32512464-32512965 | 2 | 0.815666690092812 |
| chr17-14012315-14012816 | 5 | 0.990050227995684 |
| chr6-100773756-100774257 | 2 | 0.945048369003754 |
| chr14-56641260-56641761 | 8 | 0.993493681128311 |
| chr17-32240446-32240947 | 8 | 0.962891265105242 |
| chr6-109632803-109633304 | 3 | 0.995266484888213 |
| chr3-108182170-108182671 | 2 | 0.929420915085635 |
| chr4-174440044-174440545 | 7 | 0.970984820553646 |
| chr3-66496114-66496615 | 3 | 0.996762183194853 |
| chr8-55760547-55761048 | 8 | 0.96321590841965 |
| chr18-43143760-43144261 | 2 | 0.877591640848189 |
| chr9-96041782-96042283 | 1 | 0.887589583796505 |
| chr1-128983559-128984060 | 8 | 0.942518331555107 |
| chr4-13017938-13018439 | 8 | 0.993895306728392 |
| chr10-106820116-106820617 | 2 | 0.937062313113265 |
| chr12-23769283-23769784 | 8 | 0.974318035028958 |
| chr6-117863805-117864306 | 8 | 0.993493681128311 |
| chr8-24192469-24192970 | 3 | 0.994707637836735 |
| chr10-92566430-92566931 | 8 | 0.98595640628088 |
| chr5-4703789-4704290 | 6 | 0.991805754014272 |

| | | |
|---|---|---|
| chr15-1612716-1613217 | 8 | 0.995142633642585 |
| chr10-97527078-97527579 | 6 | 0.874079475926103 |
| chr8-86875422-86875923 | 8 | 0.860888905042349 |
| chr13-96473912-96474413 | 8 | 0.912784595423582 |
| chr5-109835528-109836029 | 8 | 0.966789023701419 |
| chr1-100392595-100393096 | 4 | 0.970651289522392 |
| chr2-155459446-155459947 | 3 | 0.964864557346121 |
| chr13-90292822-90293323 | 6 | 0.962192429634533 |
| chr6-44286509-44287010 | 4 | 0.829199358548778 |
| chr3-108477226-108477727 | 8 | 0.99283826974776 |
| chr9-65298976-65299477 | 2 | 0.900496719853632 |
| chr4-137689564-137690065 | 4 | 0.86061821188798 |
| chr9-38653178-38653679 | 8 | 0.948871807638006 |
| chr3-116923467-116923968 | 3 | 0.993745258573999 |
| chr9-11043862-11044363 | 7 | 0.853682483949814 |
| chr1-240535709-240536210 | 8 | 0.817549490819253 |
| chr15-76812666-76813167 | 3 | 0.95260686726094 |
| chr7-44114126-44114627 | 2 | 0.831874455414634 |
| chr2-240743168-240743669 | 5 | 0.988317108525239 |
| chr1-95006950-95007451 | 6 | 0.996266635448841 |
| chr4-117980519-117981020 | 6 | 0.840747309687903 |
| chr1-176730470-176730971 | 8 | 0.957682223758156 |
| chr2-129014870-129015371 | 3 | 0.97442753141364 |
| chr15-73533214-73533715 | 8 | 0.990282003215943 |
| chr6-49831136-49831637 | 8 | 0.908011841809726 |
| chr5-74360660-74361161 | 8 | 0.817332828753923 |
| chr15-3742048-3742549 | 5 | 0.98626954280233 |
| chr17-66283857-66284358 | 3 | 0.993524083943392 |
| chr10-65234066-65234567 | 2 | 0.821969020437924 |
| chr10-62639876-62640377 | 8 | 0.965016468463316 |
| chr2-82119714-82120215 | 5 | 0.822860919185433 |
| chr15-15741077-15741578 | 5 | 0.97751451842685 |
| chr4-169143352-169143853 | 5 | 0.947615246443736 |
| chr15-32094233-32094734 | 8 | 0.985900110466702 |
| chr3-58650947-58651448 | 8 | 0.88658389223344 |
| chr7-113603980-113604481 | 8 | 0.993111265173513 |
| chr13-63073471-63073972 | 8 | 0.976948529293467 |
| chr12-39836506-39837007 | 5 | 0.980200942544801 |
| chr17-78273622-78274123 | 2 | 0.945627837913685 |
| chr9-103962228-103962729 | 7 | 0.832654483013669 |
| chr6-23841119-23841620 | 2 | 0.843585815500645 |
| chr3-50593682-50594183 | 5 | 0.989312370254548 |
| chr10-37326168-37326669 | 8 | 0.902890300912356 |

| | | |
|---|---|---|
| chr13-5592582-5593083 | 3 | 0.99593553545041 |
| chr4-66392265-66392766 | 5 | 0.899309673370407 |
| chr3-19406366-19406867 | 1 | 0.937581959316105 |
| chr2-113028204-113028705 | 3 | 0.843223119721419 |
| chr20-19939839-19940340 | 8 | 0.983155354738596 |
| chr7-2097440-2097941 | 3 | 0.967221678204836 |
| chr20-9298618-9299119 | 8 | 0.858086949649737 |
| chr9-35401336-35401837 | 3 | 0.997671124914817 |
| chr6-67253236-67253737 | 3 | 0.956857316850102 |
| chr10-48929532-48930033 | 4 | 0.949351803746018 |
| chr9-83853256-83853757 | 4 | 0.964257014790037 |
| chr10-37063915-37064416 | 8 | 0.848176766102812 |
| chr6-106982071-106982572 | 3 | 0.996260702076582 |
| chr1-217369294-217369795 | 8 | 0.819173768464755 |
| chr13-48529453-48529954 | 8 | 0.995214546730281 |
| chr1-2067631-2068132 | 6 | 0.931665784469459 |
| chr11-29367169-29367670 | 4 | 0.950658817112221 |
| chr16-28545969-28546470 | 8 | 0.977752007789901 |
| chr1-42509622-42510123 | 8 | 0.990440443534031 |
| chr2-171542213-171542714 | 8 | 0.934129479628387 |
| chr10-93074266-93074767 | 4 | 0.852323648426432 |
| chr10-26380370-26380871 | 6 | 0.991206925481043 |
| chr5-113695248-113695749 | 8 | 0.994949930624093 |
| chr18-67742177-67742678 | 5 | 0.970821476764646 |
| chr13-82060102-82060603 | 8 | 0.878181024791738 |
| chr19-54184226-54184727 | 5 | 0.992534989523625 |
| chr6-41403203-41403704 | 4 | 0.903239021655959 |
| chr14-9506457-9506958 | 5 | 0.985612623212067 |
| chr4-137988756-137989257 | 2 | 0.816841996255319 |
| chr1-260025347-260025848 | 5 | 0.903630642711591 |
| chr7-66421692-66422193 | 8 | 0.954074072597433 |
| chr3-21924159-21924660 | 8 | 0.913802050780581 |
| chr2-221373101-221373602 | 3 | 0.899179661275066 |
| chr10-101777642-101778143 | 8 | 0.994556441426032 |
| chr18-15706726-15707227 | 3 | 0.985550934033111 |
| chr17-31180947-31181448 | 8 | 0.943311537481264 |
| chr6-106241382-106241883 | 8 | 0.971466314554786 |
| chr4-72106833-72107334 | 2 | 0.924230294452341 |
| chr15-40304800-40305301 | 8 | 0.857702576367824 |
| chr17-43198962-43199463 | 2 | 0.910695059631875 |
| chr3-166862235-166862736 | 8 | 0.98107912257875 |
| chr3-123722295-123722796 | 8 | 0.905007096345778 |
| chr18-51364768-51365269 | 8 | 0.890694756364438 |

| | | |
|---|---|---|
| chr1-215037373-215037874 | 8 | 0.923738725403357 |
| chr9-36706428-36706929 | 6 | 0.879326682004177 |
| chr1-57257713-57258214 | 8 | 0.989439373153604 |
| chr15-80595748-80596249 | 5 | 0.981021399480913 |
| chr17-28830131-28830632 | 8 | 0.971421399512828 |
| chr4-123365422-123365923 | 4 | 0.930693632714799 |
| chr18-48045926-48046427 | 6 | 0.949725702304139 |
| chr14-79421821-79422322 | 5 | 0.987405422991878 |
| chr5-17907569-17908070 | 6 | 0.987975589776673 |
| chr7-119388912-119389413 | 3 | 0.9967928410371 |
| chr10-33942732-33943233 | 8 | 0.849480279933662 |
| chr18-3586305-3586806 | 3 | 0.996827509729096 |
| chr15-46257078-46257579 | 6 | 0.940171753953691 |
| chr4-124973675-124974176 | 5 | 0.967524055464267 |
| chr13-69235567-69236068 | 8 | 0.977343444293909 |
| chr6-111822471-111822972 | 3 | 0.996812429855763 |
| chr4-148417877-148418378 | 4 | 0.892211367047311 |
| chr4-148134502-148135003 | 8 | 0.961693817531609 |
| chr14-74638923-74639424 | 5 | 0.982334093937637 |
| chr1-189424268-189424769 | 6 | 0.945757388067513 |
| chr4-93772981-93773482 | 8 | 0.853595439436259 |
| chr7-12050848-12051349 | 6 | 0.990058101990029 |
| chr2-127831088-127831589 | 8 | 0.993493681128311 |
| chr19-10281632-10282133 | 6 | 0.991975844055074 |
| chr4-159068561-159069062 | 6 | 0.84765193626708 |
| chr2-30584438-30584939 | 5 | 0.972746250115407 |
| chr2-133966510-133967011 | 4 | 0.898581830236503 |
| chr11-52240182-52240683 | 3 | 0.996768133280268 |
| chr9-6698701-6699202 | 2 | 0.871872236975833 |
| chr2-179078040-179078541 | 5 | 0.987228964843614 |
| chr1-179664420-179664921 | 8 | 0.941687258521543 |
| chr2-191463899-191464400 | 3 | 0.995278107005591 |
| chr3-148724393-148724894 | 8 | 0.910061161567956 |
| chr4-49169343-49169844 | 3 | 0.99593553545041 |
| chr7-91726332-91726833 | 2 | 0.939445803014906 |
| chr1-256218779-256219280 | 8 | 0.848741553112758 |
| chr2-145010060-145010561 | 8 | 0.994884364871509 |
| chr11-34546066-34546567 | 2 | 0.955381000238406 |
| chr3-123534560-123535061 | 7 | 0.894847341323709 |
| chr8-14508034-14508535 | 3 | 0.996585991555055 |
| chr17-59616105-59616606 | 3 | 0.995892597261526 |
| chr14-12782274-12782775 | 3 | 0.964892347436169 |
| chr13-91748260-91748761 | 4 | 0.83719544977383 |

| | | |
|---|---|---|
| chr13-76818601-76819102 | 2 | 0.877893009967011 |
| chr11-238550-239051 | 8 | 0.85821828117417 |
| chr15-38027514-38028015 | 5 | 0.906250053894709 |
| chr20-50423449-50423950 | 3 | 0.938659774001725 |
| chr16-10877912-10878413 | 8 | 0.995008190798393 |
| chr4-124506333-124506834 | 2 | 0.826105271006316 |
| chr13-33705237-33705738 | 3 | 0.89068789146904 |
| chr1-183943205-183943706 | 8 | 0.832332519784335 |
| chr16-56964576-56965077 | 2 | 0.854078182970453 |
| chr17-67022894-67023395 | 1 | 0.97770273266536 |
| chr5-132696011-132696512 | 8 | 0.964261076507302 |
| chr17-60724630-60725131 | 6 | 0.961978230453921 |
| chr15-82149893-82150394 | 1 | 0.81650778606391 |
| chr8-29836926-29837427 | 7 | 0.867463947542518 |
| chr1-169157782-169158283 | 8 | 0.988242274353192 |
| chr8-22380930-22381431 | 4 | 0.829670115523686 |
| chr4-133390597-133391098 | 8 | 0.976732545428403 |
| chr6-124004094-124004595 | 8 | 0.838100477820598 |
| chr13-71102334-71102835 | 8 | 0.995071445632928 |
| chr4-51120591-51121092 | 8 | 0.821135841735809 |
| chr14-29311355-29311856 | 2 | 0.907034188477221 |
| chr4-177084895-177085396 | 8 | 0.88532279755403 |
| chr1-143546744-143547245 | 8 | 0.878187394234694 |
| chr2-10682598-10683099 | 3 | 0.99593553545041 |
| chr14-64701212-64701713 | 3 | 0.988467246223218 |
| chr8-86969506-86970007 | 8 | 0.913161409338029 |
| chr2-238474369-238474870 | 3 | 0.958298786263517 |
| chr8-60998253-60998754 | 8 | 0.941423391332232 |
| chr5-166753176-166753677 | 8 | 0.971942401030485 |
| chr15-83864795-83865296 | 8 | 0.972223839090759 |
| chr8-14476103-14476604 | 3 | 0.995970437207271 |
| chr13-49485292-49485793 | 5 | 0.927358092842259 |
| chr3-45698232-45698733 | 7 | 0.964456177419394 |
| chr16-51149766-51150267 | 4 | 0.934362311894723 |
| chr6-46492075-46492576 | 4 | 0.82199108242908 |
| chr17-70442959-70443460 | 5 | 0.989873562097984 |
| chr20-42950648-42951149 | 6 | 0.949081890593085 |
| chr8-80408766-80409267 | 8 | 0.985300613682515 |
| chr18-47568710-47569211 | 3 | 0.998663517933723 |
| chr12-29001746-29002247 | 5 | 0.879700207772776 |
| chr4-157258819-157259320 | 3 | 0.996733236715367 |
| chr2-204672303-204672804 | 6 | 0.911919239733339 |
| chr11-64404363-64404864 | 4 | 0.816312851671639 |

| | | |
|---|---|---|
| chr3-149805594-149806095 | 8 | 0.90979145707035 |
| chr16-10645709-10646210 | 3 | 0.995099335843996 |
| chr17-26654516-26655017 | 8 | 0.85761391261118 |
| chr14-81142769-81143270 | 8 | 0.925278847957669 |
| chr4-84469706-84470207 | 3 | 0.972986477668524 |
| chr17-26869382-26869883 | 8 | 0.957330934421418 |
| chr5-148473330-148473831 | 2 | 0.816106132863384 |
| chr17-28709041-28709542 | 2 | 0.885586731696899 |
| chr16-23834455-23834956 | 6 | 0.984488080940628 |
| chr5-67499050-67499551 | 2 | 0.890940045751718 |
| chr1-21339741-21340242 | 5 | 0.986383627370342 |
| chr2-195388711-195389212 | 8 | 0.900074082912206 |
| chr3-102218456-102218957 | 8 | 0.993964672251086 |
| chr19-19884836-19885337 | 6 | 0.992737757460866 |
| chr13-12013449-12013950 | 3 | 0.975744328908357 |
| chr13-3074618-3075119 | 5 | 0.983919978840557 |
| chr8-28477673-28478174 | 2 | 0.877968141769439 |
| chr2-221970772-221971273 | 5 | 0.988538430444731 |
| chr2-78628744-78629245 | 7 | 0.969484503259656 |
| chr3-52014902-52015403 | 8 | 0.993493681128311 |
| chr4-106320331-106320832 | 3 | 0.991225268467353 |
| chr1-195360754-195361255 | 6 | 0.993183362470039 |
| chr11-34772880-34773381 | 3 | 0.960693969856017 |
| chr1-171705563-171706064 | 4 | 0.872908237796984 |
| chrX-83223068-83223569 | 3 | 0.990224315562538 |
| chr7-131562633-131563134 | 8 | 0.856176528623454 |
| chr6-14574460-14574961 | 5 | 0.988758438532114 |
| chr11-65541102-65541603 | 8 | 0.986519282271428 |
| chr18-31417615-31418116 | 3 | 0.989882818678652 |
| chr2-94443119-94443620 | 8 | 0.875449226227587 |
| chr1-186807587-186808088 | 8 | 0.883978328894881 |
| chr4-9075775-9076276 | 8 | 0.993493681128311 |
| chr8-76340454-76340955 | 8 | 0.913982241263186 |
| chrX-103605215-103605716 | 8 | 0.935891817961648 |
| chr18-15821604-15822105 | 2 | 0.834780162298243 |
| chr14-26562059-26562560 | 3 | 0.996674801929198 |
| chr6-6823851-6824352 | 8 | 0.977875182782689 |
| chr12-35873161-35873662 | 6 | 0.989190539008072 |
| chr2-116702013-116702514 | 8 | 0.934684434259968 |
| chr16-73003733-73004234 | 6 | 0.992314744798324 |
| chr13-82442573-82443074 | 8 | 0.966684878236623 |
| chr11-22517980-22518481 | 4 | 0.833144810045333 |
| chr13-2763095-2763596 | 3 | 0.99593553545041 |

| | | |
|---|---|---|
| chr2-52868796-52869297 | 6 | 0.994331547669686 |
| chr19-24627433-24627934 | 4 | 0.843492307647159 |
| chr14-18669460-18669961 | 3 | 0.99593553545041 |
| chrX-30638233-30638734 | 8 | 0.991168373552421 |
| chr1-154800142-154800643 | 8 | 0.972159043198922 |
| chr14-27546328-27546829 | 3 | 0.9968822138388 |
| chr7-37725473-37725974 | 8 | 0.972459137379764 |
| chr2-179838693-179839194 | 3 | 0.99593553545041 |
| chr6-38419901-38420402 | 5 | 0.940862501795483 |
| chr10-80911061-80911562 | 8 | 0.941695266799231 |
| chr19-46255557-46256058 | 5 | 0.984285134152873 |
| chr20-37006055-37006556 | 8 | 0.954680430088613 |
| chr5-135587043-135587544 | 8 | 0.935959638086136 |
| chr2-168875065-168875566 | 8 | 0.969163334764912 |
| chr6-109039044-109039545 | 3 | 0.99652777084639 |
| chr1-164941414-164941915 | 8 | 0.969779818990708 |
| chr14-82996463-82996964 | 4 | 0.972116296842901 |
| chr11-82241054-82241555 | 6 | 0.912313645268382 |
| chr13-41692453-41692954 | 8 | 0.988156938122633 |
| chr3-152618900-152619401 | 8 | 0.992319494024351 |
| chr10-97929564-97930065 | 8 | 0.857589367251087 |
| chr4-171122868-171123369 | 5 | 0.986890620238365 |
| chr4-45774792-45775293 | 8 | 0.978056431619351 |
| chr1-41099211-41099712 | 5 | 0.974023043169666 |
| chr5-94856936-94857437 | 6 | 0.988742014110839 |
| chr2-42733876-42734377 | 1 | 0.82177226291208 |
| chr6-7934557-7935058 | 7 | 0.890188671258969 |
| chr12-44647566-44648067 | 8 | 0.993434248355977 |
| chr10-95173481-95173982 | 8 | 0.942876617953413 |
| chr7-58418125-58418626 | 5 | 0.844035540458717 |
| chr1-129852378-129852879 | 8 | 0.866522569140492 |
| chr8-62990671-62991172 | 8 | 0.933321848874365 |
| chr9-95949573-95950074 | 3 | 0.951541289705157 |
| chr18-78464835-78465336 | 1 | 0.917786627207718 |
| chr17-48502170-48502671 | 2 | 0.929846797758224 |
| chr1-199527725-199528226 | 3 | 0.923519811782584 |
| chr9-93172944-93173445 | 6 | 0.995293343466177 |
| chr18-57268052-57268553 | 8 | 0.817292965467826 |
| chr3-19522239-19522740 | 8 | 0.961448799692018 |
| chr2-114425537-114426038 | 6 | 0.978936466041311 |
| chr12-33231684-33232185 | 8 | 0.993869070007562 |
| chr7-18230731-18231232 | 6 | 0.966863331889409 |
| chr4-62361795-62362296 | 3 | 0.967689608137056 |

| | | |
|---|---|---|
| chr14-6419844-6420345 | 3 | 0.994139195329967 |
| chr4-13708439-13708940 | 3 | 0.852346801221981 |
| chr6-22478788-22479289 | 1 | 0.879930627215435 |
| chr1-21718867-21719368 | 4 | 0.860438211286473 |
| chr15-27601046-27601547 | 8 | 0.920491061286935 |
| chr17-35890379-35890880 | 7 | 0.912293732671934 |
| chr6-122653656-122654157 | 6 | 0.99322715434238 |
| chr8-21901648-21902149 | 8 | 0.919598091743319 |
| chr1-141903691-141904192 | 2 | 0.881128404867834 |
| chr1-197464811-197465312 | 5 | 0.933849351966984 |
| chr14-37376841-37377342 | 3 | 0.972366379860099 |
| chr3-111890709-111891210 | 5 | 0.940106581359365 |
| chr9-56409958-56410459 | 8 | 0.912280866086382 |
| chr2-235760033-235760534 | 1 | 0.931725946859972 |
| chr1-5058307-5058808 | 2 | 0.944486818502775 |
| chr15-78763997-78764498 | 5 | 0.969765072751145 |
| chr1-179719863-179720364 | 3 | 0.990205815145553 |
| chr4-150011915-150012416 | 8 | 0.994947591556325 |
| chr6-135212289-135212790 | 3 | 0.948266598250996 |
| chr2-226532979-226533480 | 7 | 0.831835228037129 |
| chr1-215358741-215359242 | 3 | 0.97105499685899 |
| chr18-27034967-27035468 | 5 | 0.990247302822035 |
| chr17-45709554-45710055 | 2 | 0.917131779483262 |
| chr2-76936131-76936632 | 6 | 0.856252308071365 |
| chr4-160190185-160190686 | 8 | 0.988264692079785 |
| chr11-25523735-25524236 | 3 | 0.877077877235377 |
| chr4-161236060-161236561 | 8 | 0.817850506143387 |
| chr4-168844555-168845056 | 5 | 0.903517581767785 |
| chr8-532054-532555 | 6 | 0.992401124645678 |
| chr3-148909027-148909528 | 8 | 0.842135328521299 |
| chr2-211064297-211064798 | 3 | 0.973840358322025 |
| chr13-84733513-84734014 | 3 | 0.997525623307797 |
| chr15-11536834-11537335 | 6 | 0.867611528884445 |
| chr3-46379902-46380403 | 8 | 0.839150750891535 |
| chr5-31051820-31052321 | 5 | 0.969889710921996 |
| chr6-44080138-44080639 | 3 | 0.99428351001345 |
| chr8-120318056-120318557 | 4 | 0.842801326608462 |
| chr9-39056821-39057322 | 5 | 0.988600720135487 |
| chr13-86695706-86696207 | 5 | 0.96109026120874 |
| chr20-10909271-10909772 | 8 | 0.993493681128311 |
| chr6-42153562-42154063 | 8 | 0.92349107456813 |
| chr13-38607504-38608005 | 8 | 0.820520718409969 |
| chr3-96696124-96696625 | 8 | 0.981728112476511 |

| | | |
|---|---|---|
| chr13-55881473-55881974 | 5 | 0.958858119517232 |
| chr10-34730598-34731099 | 3 | 0.994359327951079 |
| chr3-60497704-60498205 | 6 | 0.82998505519722 |
| chr11-75032120-75032621 | 8 | 0.95392613716513 |
| chr10-92910654-92911155 | 2 | 0.851824999000314 |
| chr2-190802120-190802621 | 3 | 0.996393415481454 |
| chr7-41153324-41153825 | 5 | 0.987433256280139 |
| chr7-108915923-108916424 | 1 | 0.977917533272068 |
| chr4-148512278-148512779 | 8 | 0.992449101032812 |
| chr11-28142457-28142958 | 5 | 0.886593010493011 |
| chr12-13970720-13971221 | 7 | 0.933632434202725 |
| chr19-33923393-33923894 | 7 | 0.908099443904265 |
| chr15-98796495-98796996 | 7 | 0.857774342508279 |
| chr1-122664148-122664649 | 6 | 0.991206925481043 |
| chr8-19863186-19863687 | 8 | 0.993156135715065 |
| chr3-108063062-108063563 | 8 | 0.985582413263609 |
| chr3-43936376-43936877 | 4 | 0.865797035197933 |
| chr11-34001102-34001603 | 5 | 0.925141439082443 |
| chr17-79731752-79732253 | 4 | 0.96540655295438 |
| chr1-247538309-247538810 | 3 | 0.989838585808393 |
| chr3-116070066-116070567 | 3 | 0.948498013618857 |
| chr15-101319626-101320127 | 8 | 0.995104666317591 |
| chr8-84906161-84906662 | 8 | 0.837818883866183 |
| chr13-76426827-76427328 | 6 | 0.988909854716708 |
| chr4-131356981-131357482 | 5 | 0.943116232101891 |
| chr6-95179598-95180099 | 2 | 0.911393044970703 |
| chr5-13208327-13208828 | 8 | 0.838744342195562 |
| chr12-34937145-34937646 | 4 | 0.912657525437569 |
| chr19-31388631-31389132 | 5 | 0.95280271505992 |
| chr2-169480314-169480815 | 5 | 0.985851629790734 |
| chr4-121286790-121287291 | 5 | 0.828686731072886 |
| chr16-75412620-75413121 | 8 | 0.869472450267987 |
| chrX-39015574-39016075 | 3 | 0.985192717134987 |
| chr16-12713484-12713985 | 5 | 0.893701576616922 |
| chr17-71846453-71846954 | 5 | 0.829591211903324 |
| chr5-138749483-138749984 | 4 | 0.828024866820864 |
| chr2-140839551-140840052 | 6 | 0.991707965526091 |
| chr14-34551243-34551744 | 8 | 0.965581360564306 |
| chr4-65493591-65494092 | 6 | 0.831219865371241 |
| chr10-68984976-68985477 | 8 | 0.993493681128311 |
| chr2-128501512-128502013 | 8 | 0.993493681128311 |
| chr5-115435084-115435585 | 5 | 0.932976454182307 |
| chr4-95212469-95212970 | 5 | 0.983919978840557 |

| | | |
|---|---|---|
| chr1-184298659-184299160 | 5 | 0.97217696773474 |
| chr2-219037992-219038493 | 8 | 0.98234788983046 |
| chr5-136075178-136075679 | 1 | 0.870241479388609 |
| chr1-247306020-247306521 | 8 | 0.946115216812754 |
| chr1-63575678-63576179 | 3 | 0.987744898023284 |
| chr9-61387182-61387683 | 8 | 0.948312277396315 |
| chr1-21256507-21257008 | 3 | 0.99689578755327 |
| chr2-107483937-107484438 | 3 | 0.985785006865021 |
| chr10-95901852-95902353 | 5 | 0.978694615039021 |
| chr5-57429601-57430102 | 5 | 0.839555307300412 |
| chr18-24312023-24312524 | 3 | 0.826152785890942 |
| chr1-48311075-48311576 | 8 | 0.993493681128311 |
| chr3-93389890-93390391 | 2 | 0.924629487523069 |
| chr1-155844424-155844925 | 8 | 0.981634136798012 |
| chr13-79483584-79484085 | 5 | 0.990886239733335 |
| chr13-75742550-75743051 | 4 | 0.924972851231146 |
| chr2-29870964-29871465 | 4 | 0.92381336397894 |
| chr18-77996351-77996852 | 8 | 0.995160930405016 |
| chr14-61555604-61556105 | 5 | 0.984935598459337 |
| chr1-98157825-98158326 | 3 | 0.996895515637706 |
| chr14-103842199-103842700 | 8 | 0.937773007701039 |
| chr20-48197301-48197802 | 7 | 0.855423044168751 |
| chr17-33302977-33303478 | 5 | 0.944218609380808 |
| chr5-48362343-48362844 | 3 | 0.997496346202293 |
| chr20-40841624-40842125 | 8 | 0.861909209661902 |
| chr18-36289878-36290379 | 8 | 0.906060661339147 |
| chr20-27417529-27418030 | 7 | 0.954874115004696 |
| chr6-12112703-12113204 | 3 | 0.993688003566767 |
| chr3-55743039-55743540 | 8 | 0.960872902394187 |
| chr3-144052659-144053160 | 5 | 0.931205132487714 |
| chr16-44146814-44147315 | 6 | 0.987957571771903 |
| chr17-56185874-56186375 | 3 | 0.975449727495067 |
| chr17-32716240-32716741 | 8 | 0.960601069723379 |
| chr9-39740718-39741219 | 8 | 0.892169742955855 |
| chr7-42369971-42370472 | 3 | 0.8724794013243 |
| chr15-28448292-28448793 | 8 | 0.885348980514762 |
| chr1-7799192-7799693 | 3 | 0.996898561280417 |
| chr16-78864806-78865307 | 8 | 0.886016378038771 |
| chr1-197458194-197458695 | 8 | 0.914247385425331 |
| chr2-125229168-125229669 | 8 | 0.9952183083928 |
| chr2-195563140-195563641 | 5 | 0.827355669837822 |
| chr7-46015718-46016219 | 6 | 0.815553729188408 |
| chr7-53391033-53391534 | 4 | 0.887709020772528 |

| | | |
|---|---|---|
| chr1-48622819-48623320 | 8 | 0.950960398731481 |
| chr9-14399691-14400192 | 8 | 0.991571970032269 |
| chr20-32821564-32822065 | 2 | 0.928554169045845 |
| chr2-48236860-48237361 | 3 | 0.995666031511903 |
| chr1-42285573-42286074 | 8 | 0.932263056736488 |
| chr2-45299477-45299978 | 8 | 0.84738609506993 |
| chr3-55693259-55693760 | 8 | 0.899957246811104 |
| chr13-32197086-32197587 | 3 | 0.996514778878781 |
| chr6-129728907-129729408 | 8 | 0.993493681128311 |
| chr5-101470316-101470817 | 8 | 0.863938104107102 |
| chr7-24290221-24290722 | 8 | 0.993493681128311 |
| chr11-16059824-16060325 | 3 | 0.99593553545041 |
| chr2-174443986-174444487 | 8 | 0.951284868170145 |
| chr13-89955184-89955685 | 2 | 0.873393213284362 |
| chr14-95095133-95095634 | 3 | 0.950599235874246 |
| chr4-40533367-40533868 | 8 | 0.934269263815953 |
| chr3-26457577-26458078 | 3 | 0.947524348401938 |
| chr5-46400503-46401004 | 8 | 0.886950974027024 |
| chr1-213596605-213597106 | 8 | 0.879804573895351 |
| chr4-136936363-136936864 | 5 | 0.943001325518924 |
| chr1-199909309-199909810 | 8 | 0.909850809661942 |
| chr4-33223471-33223972 | 5 | 0.988505366354128 |
| chr8-70768043-70768544 | 6 | 0.989845086120882 |
| chr7-13825789-13826290 | 6 | 0.991206925481043 |
| chr11-54418573-54419074 | 8 | 0.974406784907429 |
| chr14-34702157-34702658 | 4 | 0.903266862383893 |
| chr10-68405642-68406143 | 2 | 0.916846966816944 |
| chr15-58862607-58863108 | 2 | 0.949807410106612 |
| chr19-39180815-39181316 | 2 | 0.862705371252284 |
| chr10-78865310-78865811 | 8 | 0.861400091561138 |
| chr10-73272855-73273356 | 4 | 0.944951550522835 |
| chr7-128854125-128854626 | 3 | 0.858024429284459 |
| chr7-27996164-27996665 | 8 | 0.821564508081797 |
| chr16-19956151-19956652 | 3 | 0.998738919658654 |
| chr19-28759843-28760344 | 5 | 0.981760870896903 |
| chr4-39905322-39905823 | 8 | 0.993219949697004 |
| chr15-99979719-99980220 | 5 | 0.953911321309424 |
| chr4-57595933-57596434 | 4 | 0.926426789429402 |
| chr4-22279319-22279820 | 8 | 0.954169002665358 |
| chr17-57422213-57422714 | 6 | 0.9859960321206 |
| chr3-27682563-27683064 | 2 | 0.943858356456128 |
| chr4-89785025-89785526 | 8 | 0.831529589961477 |
| chr3-168604948-168605449 | 4 | 0.878855540546074 |

| | | |
|---|---|---|
| chr1-23489743-23490244 | 5 | 0.952662119958099 |
| chr7-103657332-103657833 | 3 | 0.99593553545041 |
| chr6-89039691-89040192 | 4 | 0.935299876117616 |
| chr14-71432885-71433386 | 3 | 0.882732615619038 |
| chr5-151820278-151820779 | 8 | 0.847850684199932 |
| chr14-67197811-67198312 | 8 | 0.970494863814622 |
| chr5-143842425-143842926 | 3 | 0.996632698526141 |
| chr7-355961-356462 | 8 | 0.913704533304957 |
| chr10-63538614-63539115 | 2 | 0.953411264788595 |
| chr18-49691310-49691811 | 2 | 0.842618997419355 |
| chr6-30410239-30410740 | 1 | 0.874899687150458 |
| chr10-30634583-30635084 | 8 | 0.864251295864885 |
| chr15-64541646-64542147 | 8 | 0.986567924850793 |
| chr4-123444102-123444603 | 4 | 0.825413031205121 |
| chr6-22627907-22628408 | 8 | 0.975042955515873 |
| chr5-143675919-143676420 | 3 | 0.97809236227615 |
| chr4-19146922-19147423 | 3 | 0.996888761606514 |
| chr2-220878818-220879319 | 2 | 0.878041644076807 |
| chr7-130873616-130874117 | 4 | 0.936456375928137 |
| chr4-180028844-180029345 | 6 | 0.988077650148012 |
| chr2-225731504-225732005 | 3 | 0.996886942933905 |
| chr19-41628591-41629092 | 5 | 0.857650497233663 |
| chr12-9593829-9594330 | 6 | 0.991715865300873 |
| chr4-52487762-52488263 | 2 | 0.950735669506442 |
| chr6-90026786-90027287 | 5 | 0.96109205786575 |
| chr4-40117378-40117879 | 6 | 0.945080993499073 |
| chr4-58519452-58519953 | 8 | 0.970348706240279 |
| chr11-70284324-70284825 | 8 | 0.864042698496631 |
| chr8-122958651-122959152 | 8 | 0.982957472170408 |
| chr14-94170725-94171226 | 5 | 0.992419347258035 |
| chr13-76748240-76748741 | 3 | 0.964653044502504 |
| chr9-104081232-104081733 | 7 | 0.828879261185057 |
| chr17-84163798-84164299 | 6 | 0.989563639495005 |
| chr4-175362132-175362633 | 8 | 0.979184992996747 |
| chr5-61897194-61897695 | 4 | 0.925016528931506 |
| chr9-1273878-1274379 | 8 | 0.951855471920763 |
| chr11-3278622-3279123 | 6 | 0.972630122406167 |
| chr6-48032121-48032622 | 8 | 0.973913360126379 |
| chr8-65988759-65989260 | 8 | 0.967681143622566 |
| chr1-153252041-153252542 | 7 | 0.899736344363907 |
| chr3-137831874-137832375 | 3 | 0.975052231685203 |
| chr8-81623800-81624301 | 3 | 0.996352475852865 |
| chr1-157318078-157318579 | 3 | 0.98386457544126 |

| | | |
|---|---|---|
| chr5-80388986-80389487 | 5 | 0.95078275988035 |
| chr6-139079709-139080210 | 8 | 0.824199503622196 |
| chr6-51124254-51124755 | 8 | 0.997302227034509 |
| chr4-85100354-85100855 | 3 | 0.996523048838181 |
| chr13-4003215-4003716 | 3 | 0.99593553545041 |
| chr11-74270801-74271302 | 3 | 0.993952246178517 |
| chr20-27803791-27804292 | 4 | 0.922408652854774 |
| chr1-134540833-134541334 | 8 | 0.981064222554013 |
| chr4-169509628-169510129 | 5 | 0.843806815894771 |
| chr17-16246535-16247036 | 5 | 0.983919978840557 |
| chr18-73034234-73034735 | 6 | 0.986019538406521 |
| chr4-170176583-170177084 | 8 | 0.976943092205511 |
| chr6-91952799-91953300 | 5 | 0.961255806230318 |
| chr4-161030482-161030983 | 8 | 0.821991481845091 |
| chr11-69698964-69699465 | 8 | 0.987133361582493 |
| chr14-70855697-70856198 | 8 | 0.933779335650911 |
| chr6-24589389-24589890 | 4 | 0.944844276992973 |
| chr3-11813435-11813936 | 8 | 0.889956207325705 |
| chr18-46745124-46745625 | 8 | 0.953073822753152 |
| chr4-180772189-180772690 | 8 | 0.838838067422036 |
| chr15-98786560-98787061 | 1 | 0.819841881410266 |
| chr15-28941970-28942471 | 8 | 0.867886773827684 |
| chr20-7946932-7947433 | 8 | 0.995246359287424 |
| chr18-65600342-65600843 | 6 | 0.967878698933833 |
| chr6-77641415-77641916 | 3 | 0.892122978967072 |
| chr14-42651525-42652026 | 8 | 0.836577922983299 |
| chr18-78427087-78427588 | 3 | 0.967725359392957 |
| chr15-56757602-56758103 | 6 | 0.99067005005113 |
| chr3-47822811-47823312 | 6 | 0.990122459334646 |
| chr1-221747696-221748197 | 7 | 0.856564603959667 |
| chr19-46512216-46512717 | 2 | 0.823373600000609 |
| chr12-819554-820055 | 5 | 0.985347740547498 |
| chr2-13626951-13627452 | 3 | 0.896006548170374 |
| chr11-60544305-60544806 | 6 | 0.875767910798992 |
| chr1-21843716-21844217 | 8 | 0.977450898686268 |
| chr5-22344366-22344867 | 1 | 0.91099648627891 |
| chr20-3922103-3922604 | 7 | 0.950253738378109 |
| chr2-191682915-191683416 | 8 | 0.886409419995998 |
| chr20-12871516-12872017 | 2 | 0.906883921369184 |
| chr4-49484016-49484517 | 3 | 0.941373134893567 |
| chr3-44898090-44898591 | 2 | 0.959394168562028 |
| chr18-71164308-71164809 | 8 | 0.957688912201134 |
| chr15-9088038-9088539 | 8 | 0.912595929794022 |

| | | |
|---|---|---|
| chr5-161228512-161229013 | 4 | 0.940641157243298 |
| chr8-22259717-22260218 | 6 | 0.920393877446139 |
| chr13-73594565-73595066 | 4 | 0.834634427073233 |
| chr15-12824097-12824598 | 5 | 0.887732443749138 |
| chr7-95653087-95653588 | 8 | 0.907040585194369 |
| chr12-35895304-35895805 | 8 | 0.924113304978075 |
| chr8-38549931-38550432 | 6 | 0.991206925481043 |
| chr13-89813435-89813936 | 8 | 0.92170487480651 |
| chr2-140130369-140130870 | 8 | 0.952529866567463 |
| chr10-90984227-90984728 | 7 | 0.944325076707113 |
| chr15-27300163-27300664 | 5 | 0.953171212245461 |
| chr3-31367614-31368115 | 8 | 0.90432207233634 |
| chr9-51284511-51285012 | 3 | 0.996854105636614 |
| chr3-89420351-89420852 | 8 | 0.949410879110754 |
| chr7-103224139-103224640 | 3 | 0.996904088845888 |
| chr6-12012682-12013183 | 2 | 0.945027057330475 |
| chr15-34743752-34744253 | 3 | 0.900712470242057 |
| chr16-6271087-6271588 | 8 | 0.860869085627138 |
| chr1-152677680-152678181 | 8 | 0.891448545380675 |
| chr13-37513750-37514251 | 5 | 0.898239347165787 |
| chr17-49414200-49414701 | 5 | 0.926972697718174 |
| chr11-35979579-35980080 | 8 | 0.824247608866102 |
| chr5-71630761-71631262 | 6 | 0.991468244472202 |
| chr14-19492303-19492804 | 8 | 0.884380208229989 |
| chr14-40325381-40325882 | 6 | 0.951100417214088 |
| chr7-19931086-19931587 | 3 | 0.867782926479332 |
| chr8-28116198-28116699 | 3 | 0.994966087230895 |
| chr8-54994111-54994612 | 8 | 0.941385907975802 |
| chr2-153934308-153934809 | 3 | 0.837823381396743 |
| chr18-45259259-45259760 | 5 | 0.986513250235178 |
| chr9-3582544-3583045 | 8 | 0.960590778663984 |
| chr2-62094335-62094836 | 7 | 0.968601423210697 |
| chr10-36811037-36811538 | 6 | 0.994558608031975 |
| chr9-14238176-14238677 | 6 | 0.890164855400644 |
| chr11-25717405-25717906 | 8 | 0.988089072110898 |
| chr13-40146416-40146917 | 8 | 0.881261622408028 |
| chr17-19983049-19983550 | 4 | 0.841773866962704 |
| chr19-7973309-7973810 | 1 | 0.890479082205539 |
| chr4-117343308-117343809 | 4 | 0.819342917882697 |
| chr20-31186609-31187110 | 8 | 0.861203057395878 |
| chr2-175003073-175003574 | 7 | 0.938762353680481 |
| chr2-190933025-190933526 | 8 | 0.993895316024613 |
| chr16-24435700-24436201 | 6 | 0.993205216308377 |

| | | |
|---|---|---|
| chr15-96376486-96376987 | 4 | 0.930763894953406 |
| chr4-128614067-128614568 | 8 | 0.98587545221541 |
| chr5-68575873-68576374 | 4 | 0.825245837752615 |
| chr8-40844098-40844599 | 3 | 0.995448451443201 |
| chr16-42601917-42602418 | 3 | 0.996897898741737 |
| chr1-44360876-44361377 | 1 | 0.8356416789196 |
| chr3-10550260-10550761 | 5 | 0.943101035738044 |
| chr14-59115502-59116003 | 8 | 0.978221965007024 |
| chr2-74770678-74771179 | 3 | 0.99593553545041 |
| chr4-63487219-63487720 | 8 | 0.884355951930932 |
| chr15-69807249-69807750 | 2 | 0.959293122385596 |
| chr10-35885732-35886233 | 5 | 0.952355311298157 |
| chr14-3079062-3079563 | 5 | 0.877461653562515 |
| chr6-36230051-36230552 | 6 | 0.926402102205818 |
| chr1-177879639-177880140 | 8 | 0.965635006446224 |
| chr3-57248161-57248662 | 8 | 0.9258405331789 |
| chr17-29964794-29965295 | 6 | 0.989308843752634 |
| chr3-14872133-14872634 | 6 | 0.953539866175207 |
| chr6-117981433-117981934 | 5 | 0.987550298437455 |
| chr14-62604421-62604922 | 5 | 0.989943733944986 |
| chr1-32576767-32577268 | 8 | 0.992240074135667 |
| chr7-83550375-83550876 | 3 | 0.917046451751034 |
| chr1-77791093-77791594 | 8 | 0.991007976706948 |
| chr1-70801523-70802024 | 3 | 0.989776712479688 |
| chr5-89359564-89360065 | 6 | 0.991206925481043 |
| chr15-4927432-4927933 | 8 | 0.846721188176208 |
| chr8-106411436-106411937 | 6 | 0.990375734172156 |
| chr6-125087302-125087803 | 5 | 0.978130592359317 |
| chr16-11085546-11086047 | 3 | 0.989743498951621 |
| chr5-28133700-28134201 | 3 | 0.996778443188287 |
| chr5-116397803-116398304 | 1 | 0.891304362950758 |
| chr10-37325601-37326102 | 8 | 0.919517907105042 |
| chr7-67858546-67859047 | 5 | 0.978986539794649 |
| chr16-79824549-79825050 | 4 | 0.835263652180253 |
| chrX-115370790-115371291 | 4 | 0.817558157424666 |
| chr4-83968923-83969424 | 4 | 0.869560499025655 |
| chr7-122741791-122742292 | 2 | 0.867838519611811 |
| chr4-121666146-121666647 | 4 | 0.958650178899304 |
| chr15-29237481-29237982 | 7 | 0.856462793630797 |
| chr1-143659224-143659725 | 6 | 0.865313567581609 |
| chr13-90411941-90412442 | 8 | 0.935201408099083 |
| chrX-151318139-151318640 | 5 | 0.967594953874041 |
| chr8-8472968-8473469 | 8 | 0.993493681128311 |

| | | |
|---|---|---|
| chr11-60167302-60167803 | 7 | 0.920546875941229 |
| chr7-35202326-35202827 | 3 | 0.932655111624519 |
| chr8-25073912-25074413 | 8 | 0.993493681128311 |
| chr15-8582878-8583379 | 8 | 0.981405004366998 |
| chr1-173230751-173231252 | 6 | 0.989033535340754 |
| chr20-5510628-5511129 | 7 | 0.953595448422434 |
| chr1-18532958-18533459 | 5 | 0.975843774058046 |
| chr2-236711192-236711693 | 8 | 0.990892118616676 |
| chr15-58817864-58818365 | 3 | 0.99593553545041 |
| chr18-39940201-39940702 | 1 | 0.850671509112507 |
| chr4-128189975-128190476 | 3 | 0.904146846224811 |
| chr12-9820047-9820548 | 4 | 0.957392629096866 |
| chr2-139494730-139495231 | 4 | 0.891981797092932 |
| chr13-23925491-23925992 | 4 | 0.830120951769547 |
| chr20-28827998-28828499 | 3 | 0.997710904655319 |
| chr1-166423113-166423614 | 7 | 0.927012865661199 |
| chr15-47157798-47158299 | 5 | 0.989442769400044 |
| chr3-48113364-48113865 | 8 | 0.992889806070632 |
| chr6-117010010-117010511 | 6 | 0.991206925481043 |
| chr10-107076615-107077116 | 3 | 0.996896094500099 |
| chr6-71303645-71304146 | 8 | 0.980895801161126 |
| chr9-24997181-24997682 | 3 | 0.989893008843715 |
| chr4-148056741-148057242 | 1 | 0.854127045930289 |
| chr3-61072268-61072769 | 8 | 0.922283489781357 |
| chr9-61377390-61377891 | 4 | 0.81712237914655 |
| chr8-74658950-74659451 | 6 | 0.991206925481043 |
| chr4-25156232-25156733 | 2 | 0.914061938520923 |
| chr4-170328992-170329493 | 8 | 0.95647010766417 |
| chr8-89336318-89336819 | 8 | 0.898249610323624 |
| chr1-96519374-96519875 | 8 | 0.865018925355215 |
| chr15-76450166-76450667 | 8 | 0.968760101192368 |
| chr3-135973844-135974345 | 3 | 0.864074554873478 |
| chr8-29674060-29674561 | 8 | 0.881730539344675 |
| chr20-43310715-43311216 | 4 | 0.830232505821577 |
| chr18-51304102-51304603 | 1 | 0.964904016288974 |
| chr1-220545647-220546148 | 3 | 0.996438236829396 |
| chr17-70845224-70845725 | 3 | 0.990981449785923 |
| chr9-11777442-11777943 | 6 | 0.986804277661512 |
| chr6-76146944-76147445 | 6 | 0.991206925481043 |
| chr18-72472367-72472868 | 8 | 0.887466825177699 |
| chr1-136202255-136202756 | 5 | 0.825937358401205 |
| chr3-166829617-166830118 | 4 | 0.963678568712015 |
| chr9-74033119-74033620 | 8 | 0.991209500902028 |

| | | |
|---|---|---|
| chr3-124748224-124748725 | 6 | 0.822174201135421 |
| chr18-46614146-46614647 | 2 | 0.967981466663869 |
| chr15-2492380-2492881 | 8 | 0.907511827477045 |
| chr13-91604534-91605035 | 5 | 0.833892324698232 |
| chr20-46045736-46046237 | 8 | 0.968431512240545 |
| chr1-43697833-43698334 | 3 | 0.833728882829908 |
| chr4-123172886-123173387 | 7 | 0.862847832400193 |
| chr17-16313601-16314102 | 8 | 0.883917315176331 |
| chr1-69330362-69330863 | 3 | 0.983004003221614 |
| chr17-74065494-74065995 | 6 | 0.995453566340117 |
| chr10-7877748-7878249 | 6 | 0.99011229143968 |
| chr10-5643081-5643582 | 5 | 0.995156880948824 |
| chr8-114103655-114104156 | 8 | 0.964453306510332 |
| chr2-16932467-16932968 | 2 | 0.894658835601243 |
| chr4-87431841-87432342 | 4 | 0.955927241760607 |
| chr2-184863992-184864493 | 7 | 0.917266746066874 |
| chr15-22104064-22104565 | 8 | 0.993493681128311 |
| chr3-131194163-131194664 | 5 | 0.934212301622111 |
| chr19-11638475-11638976 | 6 | 0.991206925481043 |
| chr4-152369443-152369944 | 5 | 0.965629632371142 |
| chr9-70941547-70942048 | 6 | 0.965621416236656 |
| chr20-32950173-32950674 | 4 | 0.896699621870532 |
| chr5-31373822-31374323 | 8 | 0.993493681128311 |
| chr16-55615927-55616428 | 8 | 0.890122418264431 |
| chr8-26940183-26940684 | 8 | 0.865105931543169 |
| chr12-42323699-42324200 | 8 | 0.984468482189156 |
| chr19-34810342-34810843 | 8 | 0.919625019708274 |
| chr3-149793135-149793636 | 8 | 0.945810965856011 |
| chr9-92597590-92598091 | 8 | 0.995137589261461 |
| chr17-558954-559455 | 3 | 0.993524509001608 |
| chr5-77819630-77820131 | 8 | 0.82440907494061 |
| chr15-7618661-7619162 | 5 | 0.971310009044895 |
| chr8-122422490-122422991 | 6 | 0.926882060032304 |
| chr6-126625334-126625835 | 8 | 0.920986701837393 |
| chr14-72750606-72751107 | 8 | 0.840380139914685 |
| chr2-142842721-142843222 | 6 | 0.96003530125653 |
| chr12-41909006-41909507 | 8 | 0.971774360753593 |
| chr15-26628959-26629460 | 7 | 0.920019766650464 |
| chr15-6610935-6611436 | 8 | 0.993493681128311 |
| chr9-24073922-24074423 | 3 | 0.99593553545041 |
| chrX-7876498-7876999 | 3 | 0.976449781340945 |
| chr14-3050785-3051286 | 8 | 0.989723866681686 |
| chr7-52583140-52583641 | 2 | 0.956814717493453 |

| | | |
|---|---|---|
| chr7-115225590-115226091 | 4 | 0.876898502034207 |
| chr14-31058350-31058851 | 8 | 0.896446897937506 |
| chr1-108117519-108118020 | 3 | 0.99629648521096 |
| chr18-15256560-15257061 | 4 | 0.8157225918528 |
| chr10-3600100-3600601 | 1 | 0.88101160085124 |
| chr16-84523389-84523890 | 6 | 0.940124187089292 |
| chr3-57242311-57242812 | 4 | 0.935239726796041 |
| chr19-37348546-37349047 | 5 | 0.814797792887847 |
| chr10-102767324-102767825 | 3 | 0.873888464683567 |
| chr6-94116711-94117212 | 1 | 0.892219975299265 |
| chr8-96139640-96140141 | 5 | 0.915822546652504 |
| chr5-39031182-39031683 | 3 | 0.996717828429176 |
| chr1-20894483-20894984 | 8 | 0.988514181829739 |
| chrX-118326956-118327457 | 6 | 0.991206925481043 |
| chr8-62065477-62065978 | 3 | 0.970628380789511 |
| chr5-146628044-146628545 | 4 | 0.947123410116401 |
| chr15-37806033-37806534 | 4 | 0.829269046590118 |
| chr1-179483837-179484338 | 7 | 0.822944724519241 |
| chr9-20808726-20809227 | 3 | 0.931244440653619 |
| chr4-88204927-88205428 | 5 | 0.847521166536102 |
| chr15-74799175-74799676 | 3 | 0.985766159589048 |
| chr20-41054280-41054781 | 3 | 0.998426287032573 |
| chr7-76014777-76015278 | 1 | 0.866381204795672 |
| chr9-26105629-26106130 | 8 | 0.849896595913097 |
| chr6-87551180-87551681 | 8 | 0.993493681128311 |
| chrX-150702524-150703025 | 4 | 0.956930265279429 |
| chr3-11356840-11357341 | 8 | 0.991709778434335 |
| chr2-178118830-178119331 | 8 | 0.99314443647018 |
| chr2-218573223-218573724 | 3 | 0.993093294327543 |
| chr7-34367992-34368493 | 3 | 0.954324175029231 |
| chr19-35664777-35665278 | 8 | 0.893803651388656 |
| chr10-5466622-5467123 | 8 | 0.918743372369187 |
| chr2-231683687-231684188 | 8 | 0.896183110575406 |
| chr8-100411020-100411521 | 8 | 0.972844213763114 |
| chr13-38898812-38899313 | 3 | 0.99593553545041 |
| chr2-132736991-132737492 | 3 | 0.992685348007244 |
| chr1-242346536-242347037 | 8 | 0.879347648385035 |
| chr5-69320250-69320751 | 8 | 0.993493681128311 |
| chr17-79580034-79580535 | 6 | 0.992712783301745 |
| chr7-54091768-54092269 | 4 | 0.855583496357034 |
| chr2-47957428-47957929 | 8 | 0.944156034467732 |
| chr20-34008625-34009126 | 4 | 0.898251950700807 |
| chr14-12168140-12168641 | 1 | 0.858635106502732 |

| | | |
|---|---|---|
| chr7-27856862-27857363 | 3 | 0.973650720499788 |
| chr16-38589737-38590238 | 6 | 0.992560160282541 |
| chr1-213577877-213578378 | 4 | 0.842392441717136 |
| chr2-210450248-210450749 | 3 | 0.997418203204799 |
| chr10-38386008-38386509 | 1 | 0.951875165244596 |
| chr2-45504775-45505276 | 5 | 0.846910003188657 |
| chr6-89916513-89917014 | 8 | 0.969339124532679 |
| chr11-85375494-85375995 | 4 | 0.96927025986871 |
| chr2-172046219-172046720 | 3 | 0.963589216130854 |
| chr18-82364865-82365366 | 6 | 0.980630662178241 |
| chr8-67755495-67755996 | 8 | 0.988164977481767 |
| chr13-44984686-44985187 | 8 | 0.979411094518339 |
| chr3-50986363-50986864 | 8 | 0.959843262217067 |
| chr6-106224203-106224704 | 8 | 0.909763425950393 |
| chr1-189704475-189704976 | 4 | 0.934959759308536 |
| chr2-152624809-152625310 | 3 | 0.955102354175785 |
| chr1-83930290-83930791 | 1 | 0.905586129538326 |
| chr5-122168226-122168727 | 4 | 0.898995752881266 |
| chr11-70978502-70979003 | 3 | 0.83688778629849 |
| chr7-29799038-29799539 | 5 | 0.957211210602067 |
| chr16-8331064-8331565 | 6 | 0.992364975232757 |
| chr14-6979962-6980463 | 3 | 0.917492102754494 |
| chr7-53103350-53103851 | 8 | 0.992681453212056 |
| chr14-4209006-4209507 | 8 | 0.94948440446375 |
| chr8-85476014-85476515 | 4 | 0.971023206557965 |
| chr20-19545599-19546100 | 2 | 0.914699772467232 |
| chr2-244578815-244579316 | 3 | 0.99593553545041 |
| chr16-76777268-76777769 | 1 | 0.86419051814064 |
| chr5-5262926-5263427 | 6 | 0.878787436828094 |
| chr15-42610795-42611296 | 2 | 0.945125744597588 |
| chr8-116776400-116776901 | 8 | 0.891096569248591 |
| chr6-21615329-21615830 | 1 | 0.898051339450467 |
| chr18-32084190-32084691 | 4 | 0.937040418226114 |
| chr2-28837735-28838236 | 7 | 0.956181466558183 |
| chr4-91134494-91134995 | 3 | 0.96396609261947 |
| chr10-83842826-83843327 | 8 | 0.896449260311103 |
| chr9-42071256-42071757 | 8 | 0.896492576270568 |
| chr5-9106556-9107057 | 6 | 0.839877902951529 |
| chr11-84626061-84626562 | 6 | 0.988300150873811 |
| chr7-120933390-120933891 | 2 | 0.956098629773992 |
| chrX-152120988-152121489 | 3 | 0.97897880762728 |
| chr9-955747-956248 | 3 | 0.856127005401786 |
| chr4-61088304-61088805 | 1 | 0.888594519222521 |

| | | |
|---|---|---|
| chr17-54166571-54167072 | 4 | 0.970156438018595 |
| chr10-54537151-54537652 | 2 | 0.861903044258876 |
| chr1-134264757-134265258 | 1 | 0.877819428452905 |
| chr12-37789258-37789759 | 8 | 0.9754507624097 |
| chr11-62399811-62400312 | 8 | 0.993493681128311 |
| chr14-76671820-76672321 | 4 | 0.853863430696638 |
| chr9-70930215-70930716 | 8 | 0.943458624087905 |
| chr8-73451394-73451895 | 4 | 0.960603105590829 |
| chr2-165140095-165140596 | 7 | 0.896117052216371 |
| chrX-12010909-12011410 | 8 | 0.81768487505657 |
| chrX-130292580-130293081 | 4 | 0.948628921023735 |
| chr13-93426381-93426882 | 2 | 0.927462638734873 |
| chr14-81963462-81963963 | 4 | 0.87729262898141 |
| chr7-110038972-110039473 | 8 | 0.995002811775588 |
| chr7-29865869-29866370 | 1 | 0.83076165917741 |
| chr3-32433327-32433828 | 3 | 0.987853985758984 |
| chr6-72617677-72618178 | 8 | 0.993493681128311 |
| chr9-57142535-57143036 | 8 | 0.833095208336352 |
| chr1-56283074-56283575 | 7 | 0.849637789153316 |
| chr1-41715047-41715548 | 6 | 0.940306326324444 |
| chr4-10956485-10956986 | 3 | 0.851028907980504 |
| chr4-65240313-65240814 | 8 | 0.946900835171308 |
| chr5-143209835-143210336 | 5 | 0.977862695522349 |
| chr9-71246183-71246684 | 6 | 0.991206925481043 |
| chr3-128204510-128205011 | 7 | 0.933738880356252 |
| chr12-23169844-23170345 | 8 | 0.983813266652014 |
| chr9-68306008-68306509 | 2 | 0.911722865270819 |
| chrX-71603439-71603940 | 6 | 0.991206925481043 |
| chr10-14582408-14582909 | 8 | 0.968411866991947 |
| chr11-59890722-59891223 | 5 | 0.990157164022425 |
| chr6-72979937-72980438 | 6 | 0.844132636374985 |
| chr10-7460167-7460668 | 3 | 0.815587270422297 |
| chr12-11074485-11074986 | 6 | 0.92373226781106 |
| chr10-62669977-62670478 | 7 | 0.907790178689817 |
| chr17-80890493-80890994 | 8 | 0.911838579506191 |
| chr1-63124141-63124642 | 6 | 0.965927062862748 |
| chr1-117866677-117867178 | 1 | 0.866001529907166 |
| chr3-147322054-147322555 | 7 | 0.822224905312617 |
| chr11-29042044-29042545 | 8 | 0.843892899132217 |
| chr10-30933465-30933966 | 8 | 0.971629733451948 |
| chr1-11161213-11161714 | 6 | 0.991206925481043 |
| chr14-6332131-6332632 | 7 | 0.82200813709745 |
| chr15-53544938-53545439 | 8 | 0.953529484433813 |

| | | |
|---|---|---|
| chr11-54421606-54422107 | 1 | 0.952358865063383 |
| chr1-164147621-164148122 | 4 | 0.88478665876494 |
| chr5-72037526-72038027 | 6 | 0.965323674853934 |
| chr16-76152515-76153016 | 7 | 0.816230156080532 |
| chr3-44781466-44781967 | 5 | 0.932659667389211 |
| chr20-46734539-46735040 | 8 | 0.952557551856798 |
| chr10-34888073-34888574 | 2 | 0.933851741261891 |
| chr14-51053698-51054199 | 8 | 0.958567373574022 |
| chr14-25333703-25334204 | 4 | 0.910430502203725 |
| chr5-109183251-109183752 | 8 | 0.889865591506547 |
| chr10-22335134-22335635 | 8 | 0.995237183050753 |
| chr4-87849381-87849882 | 3 | 0.858652707459457 |
| chr3-147599995-147600496 | 6 | 0.992617771312633 |
| chr6-70035669-70036170 | 2 | 0.953662632615838 |
| chr8-83718751-83719252 | 3 | 0.944297688671131 |
| chr3-6923787-6924288 | 8 | 0.981583577849844 |
| chr1-28086775-28087276 | 4 | 0.906988765896632 |
| chr13-43544221-43544722 | 6 | 0.946887674505056 |
| chr2-218121025-218121526 | 7 | 0.857668433230753 |
| chr6-10572813-10573314 | 8 | 0.93613646676553 |
| chr10-33989954-33990455 | 8 | 0.971884162723112 |
| chr3-28971306-28971807 | 3 | 0.859997875575645 |
| chr3-93252114-93252615 | 8 | 0.817969744118412 |
| chr11-65463888-65464389 | 8 | 0.990023176435977 |
| chr8-88258039-88258540 | 8 | 0.988715954072259 |
| chr6-33932410-33932911 | 8 | 0.995247911177653 |
| chr5-78048640-78049141 | 8 | 0.904221231201454 |
| chr8-32603401-32603902 | 8 | 0.968906399453437 |
| chr9-15821063-15821564 | 8 | 0.98047905613433 |
| chr7-5776317-5776818 | 6 | 0.979517048741447 |
| chr5-4923362-4923863 | 8 | 0.985546667146257 |
| chr13-62942976-62943477 | 4 | 0.905185889708456 |
| chr10-56072845-56073346 | 8 | 0.89996390557549 |
| chr11-65501162-65501663 | 8 | 0.884146952975279 |
| chr1-255618429-255618930 | 4 | 0.896355282422192 |
| chr10-1507216-1507717 | 3 | 0.99593553545041 |
| chr10-12082245-12082746 | 3 | 0.995569456547676 |
| chr13-21166982-21167483 | 6 | 0.947987070304228 |
| chr6-118567790-118568291 | 8 | 0.978368072823828 |
| chr3-50387158-50387659 | 2 | 0.953112962354912 |
| chr2-24398756-24399257 | 3 | 0.910095416598002 |
| chr17-69820772-69821273 | 5 | 0.982489895889728 |
| chr12-35536064-35536565 | 6 | 0.99728374774239 |

| | | |
|---|---|---|
| chr20-19454226-19454727 | 4 | 0.837089558233483 |
| chr8-99276550-99277051 | 8 | 0.844245511045026 |
| chr5-103197372-103197873 | 3 | 0.972316861643745 |
| chr20-30471938-30472439 | 5 | 0.887453134450547 |
| chr13-99731686-99732187 | 3 | 0.911265911504896 |
| chr9-73283005-73283506 | 3 | 0.984587577575262 |
| chr9-16772303-16772804 | 1 | 0.969875666331038 |
| chr15-75572409-75572910 | 8 | 0.905332394952384 |
| chr18-1879015-1879516 | 8 | 0.919048602620049 |
| chr5-96300930-96301431 | 8 | 0.949890875494009 |
| chr20-27635745-27636246 | 6 | 0.99283241951558 |
| chr18-27656514-27657015 | 4 | 0.95512196301164 |
| chr13-41914661-41915162 | 8 | 0.942430799979497 |
| chr5-117988257-117988758 | 8 | 0.988153965805935 |
| chr14-36221490-36221991 | 5 | 0.946340028821805 |
| chr7-109012525-109013026 | 4 | 0.925547317510331 |
| chr17-73522174-73522675 | 4 | 0.967298882406869 |
| chr10-2767299-2767800 | 8 | 0.966911733029499 |
| chr14-1816657-1817158 | 3 | 0.996578966527631 |
| chr5-19250298-19250799 | 8 | 0.984509780367977 |
| chr18-12070656-12071157 | 7 | 0.87113516671001 |
| chr10-64849579-64850080 | 8 | 0.828322785412665 |
| chr8-44833882-44834383 | 4 | 0.929044880512534 |
| chr8-106566766-106567267 | 6 | 0.888600671089941 |
| chr10-74942802-74943303 | 5 | 0.942523983402761 |
| chr17-35770907-35771408 | 8 | 0.949757271600821 |
| chr9-84222191-84222692 | 5 | 0.967950836071123 |
| chr5-134328373-134328874 | 4 | 0.944281061523937 |
| chr8-109286521-109287022 | 5 | 0.858513120947264 |
| chrX-58291084-58291585 | 8 | 0.861069203293247 |
| chr1-42011414-42011915 | 8 | 0.828861244834443 |
| chr10-34190870-34191371 | 2 | 0.900849497481757 |
| chr9-22784421-22784922 | 6 | 0.995576701985923 |
| chr9-71101428-71101929 | 8 | 0.93492398155207 |
| chr2-37897518-37898019 | 1 | 0.945102965492357 |
| chr6-74106807-74107308 | 8 | 0.97840609695767 |
| chrX-49263443-49263944 | 8 | 0.993493681128311 |
| chr13-23783834-23784335 | 6 | 0.972241411083417 |
| chr10-54094578-54095079 | 4 | 0.969807891341833 |
| chr17-47190055-47190556 | 3 | 0.908521942313105 |
| chr10-16077092-16077593 | 3 | 0.994467196335241 |
| chr6-136187920-136188421 | 3 | 0.934938489706569 |
| chr8-88329141-88329642 | 8 | 0.954463821468736 |

| | | |
|---|---|---|
| chr3-35416209-35416710 | 2 | 0.952710700129486 |
| chr1-179588255-179588756 | 6 | 0.984744884492056 |
| chr3-54517269-54517770 | 5 | 0.96027965289129 |
| chr1-41571464-41571965 | 8 | 0.936668374806301 |
| chr18-24411854-24412355 | 1 | 0.889467812545941 |
| chr4-129540772-129541273 | 6 | 0.979745334886582 |
| chr6-20882661-20883162 | 5 | 0.96441271115569 |
| chr17-23612530-23613031 | 8 | 0.867212796841038 |
| chr15-95768702-95769203 | 8 | 0.933745133758544 |
| chr3-142638646-142639147 | 3 | 0.985824913827784 |
| chr15-31130247-31130748 | 6 | 0.991206925481043 |
| chr18-21962874-21963375 | 3 | 0.995741237937932 |
| chr20-31172412-31172913 | 5 | 0.98972078603459 |
| chr14-82798303-82798804 | 1 | 0.828039811528764 |
| chr3-152548185-152548686 | 4 | 0.890449524559126 |
| chr3-35886510-35887011 | 8 | 0.878932232605787 |
| chr3-126027372-126027873 | 6 | 0.991206925481043 |
| chr13-65689437-65689938 | 5 | 0.837078570120611 |
| chr4-140536252-140536753 | 5 | 0.917195431001808 |
| chr1-120401175-120401676 | 8 | 0.826367954133572 |
| chr2-136396102-136396603 | 8 | 0.905427620071676 |
| chr20-28734280-28734781 | 7 | 0.900510508448675 |
| chrX-132910444-132910945 | 8 | 0.988290190581453 |
| chr11-29097185-29097686 | 2 | 0.894926281280788 |
| chr4-22338065-22338566 | 2 | 0.831202340311383 |
| chr1-174423874-174424375 | 8 | 0.971450638647559 |
| chr6-125790569-125791070 | 7 | 0.952050971399127 |
| chr10-3626436-3626937 | 4 | 0.892012229594399 |
| chr3-56557328-56557829 | 3 | 0.864718294457305 |
| chr5-3077201-3077702 | 5 | 0.989617485419192 |
| chr13-91335049-91335550 | 2 | 0.931596565813637 |
| chr7-28169484-28169985 | 8 | 0.978465045636995 |
| chr1-218836521-218837022 | 3 | 0.982999122621224 |
| chr4-126765727-126766228 | 8 | 0.994963426460297 |
| chr20-44753665-44754166 | 8 | 0.993493681128311 |
| chr7-77077563-77078064 | 3 | 0.982153277777311 |
| chr3-132178116-132178617 | 5 | 0.989528469836083 |
| chr2-23097787-23098288 | 1 | 0.868594857341308 |
| chr3-146564306-146564807 | 3 | 0.996195026260868 |
| chr17-49673363-49673864 | 8 | 0.820434375047982 |
| chr1-48928286-48928787 | 8 | 0.952947357606621 |
| chr10-80598146-80598647 | 8 | 0.923931137217238 |
| chr2-121600467-121600968 | 5 | 0.935058674478792 |

| | | |
|---|---|---|
| chr14-79536646-79537147 | 5 | 0.892615877899327 |
| chr2-96705488-96705989 | 3 | 0.99593553545041 |
| chr2-103393607-103394108 | 3 | 0.99593553545041 |
| chr6-126490386-126490887 | 8 | 0.964593317595047 |
| chr13-48815095-48815596 | 7 | 0.920794342592222 |
| chr7-60733207-60733708 | 8 | 0.927784022499315 |
| chr3-10402429-10402930 | 8 | 0.962173797525967 |
| chr1-88111052-88111553 | 5 | 0.987925074165096 |
| chr3-136872643-136873144 | 6 | 0.84872874317781 |
| chr6-137258752-137259253 | 8 | 0.852376986854143 |
| chr3-104183169-104183670 | 8 | 0.938858786387181 |
| chr9-39311302-39311803 | 8 | 0.880402377300226 |
| chr2-34984496-34984997 | 8 | 0.917967923956069 |
| chr8-78163493-78163994 | 7 | 0.925217424927114 |
| chr1-143428493-143428994 | 4 | 0.928048545761006 |
| chr7-105523027-105523528 | 4 | 0.899621882728826 |
| chr17-46754348-46754849 | 6 | 0.993146873506981 |
| chr5-155417453-155417954 | 8 | 0.840450908237717 |
| chr1-176439372-176439873 | 3 | 0.942269545838667 |
| chr14-63106036-63106537 | 6 | 0.991206925481043 |
| chr4-11863771-11864272 | 5 | 0.89152777189893 |
| chr2-141894078-141894579 | 4 | 0.952165631505631 |
| chr16-80362541-80363042 | 5 | 0.963669272465371 |
| chr7-130764215-130764716 | 7 | 0.88342025148921 |
| chr4-52725413-52725914 | 8 | 0.930957413237032 |
| chr3-100082903-100083404 | 6 | 0.996375417465081 |
| chr12-15454708-15455209 | 8 | 0.864308628220451 |
| chr1-50461188-50461689 | 5 | 0.984023325393471 |
| chr6-25245739-25246240 | 8 | 0.914886623136692 |
| chr6-47798602-47799103 | 8 | 0.995189935399571 |
| chr6-88972751-88973252 | 8 | 0.993493681128311 |
| chr9-101963544-101964045 | 8 | 0.97979083784188 |
| chr5-139895697-139896198 | 6 | 0.991206925481043 |
| chr6-104636271-104636772 | 5 | 0.985978978726782 |
| chr4-91048844-91049345 | 4 | 0.96895146502314 |
| chr4-152971934-152972435 | 8 | 0.975056887935106 |
| chr19-27458220-27458721 | 8 | 0.877855335939111 |
| chr7-116845410-116845911 | 8 | 0.931384078414547 |
| chr7-69723955-69724456 | 3 | 0.996639860331809 |
| chr14-32000203-32000704 | 4 | 0.890388702850802 |
| chr2-184603573-184604074 | 3 | 0.913036043062793 |
| chr1-212483585-212484086 | 5 | 0.851485531888349 |
| chr19-23007153-23007654 | 2 | 0.869153856393181 |

| | | |
|---|---|---|
| chr5-36974080-36974581 | 3 | 0.995935535545041 |
| chr8-75872313-75872814 | 7 | 0.8273752154601 |
| chr4-28603660-28604161 | 3 | 0.887837192879767 |
| chr10-98785077-98785578 | 3 | 0.992818417247188 |
| chr3-41667915-41668416 | 8 | 0.993493681128311 |
| chr3-7657210-7657711 | 1 | 0.929864638201437 |
| chr16-51142010-51142511 | 8 | 0.874780607268621 |
| chr4-148626991-148627492 | 8 | 0.966266023684722 |
| chr4-19002971-19003472 | 5 | 0.944843868059729 |
| chr8-120218299-120218800 | 8 | 0.990272951706468 |
| chr1-235884804-235885305 | 2 | 0.869618721629371 |
| chr11-17016015-17016516 | 5 | 0.958900610236016 |
| chr3-57197983-57198484 | 8 | 0.89138267109697 |
| chr4-124737047-124737548 | 2 | 0.88817072472274 |
| chr15-99553177-99553678 | 8 | 0.887120917756461 |
| chr13-68452638-68453139 | 8 | 0.885690544353584 |
| chr7-116038470-116038971 | 2 | 0.915262854470227 |
| chr5-24150067-24150568 | 5 | 0.901274438494238 |
| chr3-55012809-55013310 | 8 | 0.971812206714399 |
| chr5-82778105-82778606 | 3 | 0.945925206156368 |
| chr6-58111129-58111630 | 2 | 0.827348140087022 |
| chr8-58018687-58019188 | 4 | 0.971537913946786 |
| chr13-72392399-72392900 | 8 | 0.913020729259093 |
| chr4-36483494-36483995 | 8 | 0.993493681128311 |
| chr14-92154894-92155395 | 3 | 0.996498783300807 |
| chr1-25942940-25943441 | 8 | 0.972151934409802 |
| chr2-173155582-173156083 | 8 | 0.993925631743961 |
| chr7-56568095-56568596 | 6 | 0.840240429942268 |
| chr3-64510990-64511491 | 7 | 0.844098918245266 |
| chr2-187868512-187869013 | 8 | 0.830817720674327 |
| chr8-107382358-107382859 | 3 | 0.99572062335703 |
| chr7-2263168-2263669 | 3 | 0.997900720571403 |
| chr17-83344499-83345000 | 3 | 0.995160559367578 |
| chr16-60084864-60085365 | 3 | 0.924124979409758 |
| chr6-67725539-67726040 | 8 | 0.993603106699727 |
| chr14-76041292-76041793 | 4 | 0.850255572049547 |
| chr7-127729722-127730223 | 5 | 0.979391789514322 |
| chr3-64893835-64894336 | 5 | 0.976585957100725 |
| chr14-75188037-75188538 | 8 | 0.873989000671358 |
| chr11-59064980-59065481 | 6 | 0.948582307977824 |
| chr1-234136418-234136919 | 2 | 0.944077444282493 |
| chr9-77974276-77974777 | 8 | 0.983704641442911 |
| chr6-119884325-119884826 | 5 | 0.990380472872057 |

| | | |
|---|---|---|
| chr6-119057766-119058267 | 8 | 0.955309840634537 |
| chr1-139780109-139780610 | 6 | 0.884510899209635 |
| chr18-40264914-40265415 | 1 | 0.924876167931856 |
| chr17-40392873-40393374 | 8 | 0.971881855499496 |
| chr11-27626968-27627469 | 8 | 0.958561990934962 |
| chr1-97178970-97179471 | 8 | 0.97402104050638 |
| chr3-78696347-78696848 | 8 | 0.844948609182608 |
| chr3-165327556-165328057 | 3 | 0.983013727174295 |
| chr7-109869797-109870298 | 6 | 0.875858475801895 |
| chr7-26977891-26978392 | 8 | 0.976755525664783 |
| chr3-108126959-108127460 | 1 | 0.965385758742995 |
| chrX-58115942-58116443 | 6 | 0.836295662760918 |
| chrX-138687699-138688200 | 3 | 0.918711568125055 |
| chr1-155825117-155825618 | 5 | 0.861555911060757 |
| chr17-54001015-54001516 | 5 | 0.846617762954435 |
| chr11-37868709-37869210 | 6 | 0.991187537489719 |
| chr7-115354631-115355132 | 8 | 0.904649928835132 |
| chr2-44805765-44806266 | 6 | 0.958395217417051 |
| chr15-75509629-75510130 | 3 | 0.991277177720554 |
| chr4-159019651-159020152 | 4 | 0.892529616412771 |
| chr14-27395385-27395886 | 6 | 0.96280875843103 |
| chr20-33316110-33316611 | 7 | 0.846653012072181 |
| chr8-29671070-29671571 | 7 | 0.825300901167587 |
| chr11-3781639-3782140 | 3 | 0.863031110941063 |
| chr1-242747635-242748136 | 1 | 0.953682858462138 |
| chr8-111737555-111738056 | 6 | 0.993185219985303 |
| chr4-162701029-162701530 | 4 | 0.966847760808149 |
| chr13-103244462-103244963 | 2 | 0.956812062106853 |
| chr6-113065873-113066374 | 5 | 0.970334553710809 |
| chrX-58114932-58115433 | 8 | 0.995189189258915 |
| chr1-249464333-249464834 | 6 | 0.991206925481043 |
| chr16-78960500-78961001 | 3 | 0.9976056837072 |
| chr20-35262398-35262899 | 6 | 0.95109384869132 |
| chr9-109177897-109178398 | 8 | 0.990278783299666 |
| chr4-169592117-169592618 | 6 | 0.876099716780297 |
| chr20-41162252-41162753 | 5 | 0.936143726181303 |
| chr2-136542408-136542909 | 8 | 0.834291572319334 |
| chr5-164312929-164313430 | 6 | 0.981290725010887 |
| chr3-95953037-95953538 | 3 | 0.921263499784847 |
| chr1-221214544-221215045 | 6 | 0.993030949886154 |
| chr2-178933052-178933553 | 3 | 0.99593553545041 |
| chr2-187841403-187841904 | 8 | 0.978269161764599 |
| chr10-90001185-90001686 | 4 | 0.815280676619734 |

| | | |
|---|---|---|
| chr3-19559822-19560323 | 8 | 0.904629431147796 |
| chr17-34849026-34849527 | 1 | 0.840052774465406 |
| chr14-1229333-1229834 | 4 | 0.934037055698976 |
| chr3-25658681-25659182 | 3 | 0.99654889319109 |
| chr5-60612014-60612515 | 3 | 0.995882684063526 |
| chr1-13304868-13305369 | 3 | 0.995561748587596 |
| chr17-45985460-45985961 | 3 | 0.97255404809779 |
| chr9-39243472-39243973 | 8 | 0.964592560907712 |
| chr9-65380311-65380812 | 3 | 0.995585730515071 |
| chr1-4321995-4322496 | 8 | 0.982725219949402 |
| chr14-76034656-76035157 | 5 | 0.82291343556427 |
| chr20-46621038-46621539 | 8 | 0.948563740427273 |
| chr16-22149612-22150113 | 3 | 0.99593553545041 |
| chr7-11309923-11310424 | 8 | 0.84146592294509 |
| chr7-101410008-101410509 | 4 | 0.931069786542846 |
| chr18-44335020-44335521 | 3 | 0.994532676714737 |
| chr6-132494420-132494921 | 3 | 0.997217186056032 |
| chr6-106681053-106681554 | 2 | 0.920408885046424 |
| chr1-172201600-172202101 | 8 | 0.956646868075726 |
| chr17-23885148-23885649 | 4 | 0.93117761454082 |
| chr12-31963639-31964140 | 5 | 0.989319594787223 |
| chr10-33945931-33946432 | 8 | 0.888793506847022 |
| chr5-110673509-110674010 | 3 | 0.990170536744366 |
| chr7-62435726-62436227 | 8 | 0.888296871770765 |
| chr1-39832495-39832996 | 8 | 0.938615833671814 |
| chr1-243010795-243011296 | 4 | 0.818773844938866 |
| chr17-45399353-45399854 | 4 | 0.820648628583494 |
| chr14-65453186-65453687 | 4 | 0.843021641121125 |
| chr8-29621631-29622132 | 3 | 0.990926244336402 |
| chr1-149724300-149724801 | 8 | 0.988167813183487 |
| chr7-113804051-113804552 | 8 | 0.93153934790451 |
| chr4-181472115-181472616 | 8 | 0.960851651504512 |
| chr3-123931491-123931992 | 6 | 0.993063726933853 |
| chr20-12461489-12461990 | 3 | 0.987576976517996 |
| chr2-241976851-241977352 | 8 | 0.94570776582424 |
| chr2-165675754-165676255 | 3 | 0.99593553545041 |
| chr2-157590740-157591241 | 8 | 0.990255907806572 |
| chr9-88909157-88909658 | 4 | 0.835297737540725 |
| chr16-43739023-43739524 | 8 | 0.993177249272971 |
| chr6-17007341-17007842 | 4 | 0.886933452837732 |
| chr14-35475092-35475593 | 8 | 0.993493681128311 |
| chr4-141073787-141074288 | 8 | 0.939548153148504 |
| chr6-75581596-75582097 | 8 | 0.955311325842304 |

| | | |
|---|---|---|
| chrX-83532252-83532753 | 6 | 0.931170588060549 |
| chr10-9005695-9006196 | 3 | 0.995938867728848 |
| chr16-17574587-17575088 | 3 | 0.943651582156581 |
| chr18-76490930-76491431 | 8 | 0.896140324665222 |
| chr19-48888066-48888567 | 8 | 0.945481100683129 |
| chr6-112208019-112208520 | 3 | 0.976889017669273 |
| chr8-25729737-25730238 | 2 | 0.82033772489509 |
| chr13-92101239-92101740 | 3 | 0.89131010226202 |
| chr14-35640641-35641142 | 5 | 0.860417279241892 |
| chr8-57055197-57055698 | 8 | 0.879071897607876 |
| chr1-122254189-122254690 | 5 | 0.983919978840557 |
| chr5-49846736-49847237 | 5 | 0.983919978840557 |
| chr1-255791435-255791936 | 8 | 0.985038629340082 |
| chr9-68258194-68258695 | 3 | 0.971781933657309 |
| chr5-142714252-142714753 | 4 | 0.961814907336904 |
| chr14-13487736-13488237 | 8 | 0.993379752922424 |
| chr9-74168130-74168631 | 2 | 0.970782231499784 |
| chr12-45358290-45358791 | 5 | 0.983314837226859 |
| chr5-71602671-71603172 | 6 | 0.986738217333259 |
| chr18-77571806-77572307 | 1 | 0.954822741013824 |
| chr8-80928214-80928715 | 8 | 0.96770410265327 |
| chr4-124456889-124457390 | 8 | 0.832303110478002 |
| chr4-177153982-177154483 | 8 | 0.981525940176275 |
| chr17-1672000-1672501 | 8 | 0.973187720639243 |
| chr15-51090114-51090615 | 8 | 0.842049297347958 |
| chr5-66147121-66147622 | 4 | 0.891191919028218 |
| chr5-165185637-165186138 | 8 | 0.882217717099018 |
| chr19-38620905-38621406 | 8 | 0.829153355673474 |
| chr4-140067900-140068401 | 3 | 0.996898542337441 |
| chr17-53300124-53300625 | 2 | 0.890185809564705 |
| chr5-142151251-142151752 | 3 | 0.963367342036804 |
| chr3-140124663-140125164 | 6 | 0.989199388233201 |
| chr15-62479423-62479924 | 8 | 0.975461165805654 |
| chr11-10865972-10866473 | 2 | 0.980253940737516 |
| chr13-48787256-48787757 | 5 | 0.988204601825638 |
| chr1-191611407-191611908 | 7 | 0.939018387483283 |
| chr12-25084869-25085370 | 8 | 0.927041137686798 |
| chr8-123069146-123069647 | 8 | 0.936004311100158 |
| chr1-258385268-258385769 | 2 | 0.860523798758398 |
| chr14-72601874-72602375 | 8 | 0.881280467020479 |
| chr9-94089752-94090253 | 2 | 0.954154371415482 |
| chr14-35214217-35214718 | 6 | 0.867198188713105 |
| chr12-46557272-46557773 | 8 | 0.993493681128311 |

| | | |
|---|---|---|
| chr1-114293631-114294132 | 3 | 0.994892262613339 |
| chr18-26607471-26607972 | 6 | 0.890067033824954 |
| chr4-21233066-21233567 | 8 | 0.993493681128311 |
| chr17-40555744-40556245 | 6 | 0.943343893508553 |
| chr3-116892068-116892569 | 8 | 0.974674515904274 |
| chr10-42659286-42659787 | 3 | 0.998098877256755 |
| chr1-173134554-173135055 | 4 | 0.89919125789848 |
| chr6-23852727-23853228 | 3 | 0.99593553545041 |
| chr1-164686165-164686666 | 8 | 0.834904 |
| chr2-76233140-76233641 | 7 | 0.901264116245953 |
| chr3-100287998-100288499 | 4 | 0.973129273479518 |
| chr8-74275618-74276119 | 8 | 0.825083454315473 |
| chr18-67000799-67001300 | 7 | 0.929062122813883 |
| chr4-84477433-84477934 | 4 | 0.819334571625804 |
| chr4-125821774-125822275 | 3 | 0.99593553545041 |
| chr15-64139358-64139859 | 6 | 0.942428336440842 |
| chr1-79630730-79631231 | 4 | 0.939075713773399 |
| chr11-4342808-4343309 | 6 | 0.865663974308891 |
| chr10-33554131-33554632 | 3 | 0.987645069491408 |
| chr20-50238681-50239182 | 6 | 0.991206925481043 |
| chr13-72278612-72279113 | 1 | 0.931249263739096 |
| chr7-59029440-59029941 | 8 | 0.943824076675382 |
| chr8-27841096-27841597 | 5 | 0.988562720221849 |
| chr11-59896160-59896661 | 5 | 0.973310408963638 |
| chr2-159538812-159539313 | 6 | 0.822525472284833 |
| chr7-24496117-24496618 | 7 | 0.947826151556191 |
| chr14-96372282-96372783 | 8 | 0.995025902378355 |
| chr10-84564340-84564841 | 8 | 0.995242755555335 |
| chr4-103570985-103571486 | 5 | 0.980699046077138 |
| chr3-7018616-7019117 | 6 | 0.975198038070117 |
| chr13-102709612-102710113 | 2 | 0.854834634687554 |
| chr12-43340154-43340655 | 8 | 0.970511612803339 |
| chr1-179592812-179593313 | 8 | 0.968684725066591 |
| chr3-43609253-43609754 | 6 | 0.918676937665403 |
| chr5-4725152-4725653 | 6 | 0.842610643617533 |
| chr3-80015893-80016394 | 2 | 0.864528104662273 |
| chr2-81521788-81522289 | 8 | 0.989362937894576 |
| chr5-114889967-114890468 | 8 | 0.932847328446307 |
| chr5-108204167-108204668 | 3 | 0.99593553545041 |
| chr15-94647707-94648208 | 6 | 0.981219687960823 |
| chr3-52932021-52932522 | 3 | 0.987098450779483 |
| chr1-221161918-221162419 | 3 | 0.985811056387787 |
| chr7-106933680-106934181 | 2 | 0.952941332809356 |

| | | |
|---|---|---|
| chr7-30225385-30225886 | 3 | 0.996764268534007 |
| chr15-17323858-17324359 | 6 | 0.989168402104167 |
| chr4-123340519-123341020 | 8 | 0.966741623605514 |
| chr1-54856591-54857092 | 3 | 0.99593553545041 |
| chr10-104805509-104806010 | 3 | 0.996725033120008 |
| chr4-60350703-60351204 | 5 | 0.897536541556149 |
| chr10-27264668-27265169 | 6 | 0.964981097361717 |
| chr7-49002550-49003051 | 6 | 0.977626445321926 |
| chr13-70195390-70195891 | 8 | 0.877748934218408 |
| chr2-210108448-210108949 | 1 | 0.943873063716057 |
| chr11-57495034-57495535 | 5 | 0.97802629472064 |
| chr17-74713859-74714360 | 8 | 0.928949951116286 |
| chr18-36675642-36676143 | 4 | 0.914239125766497 |
| chr6-56905929-56906430 | 3 | 0.99593553545041 |
| chr6-46793489-46793990 | 8 | 0.870144687368945 |
| chr5-129080781-129081282 | 6 | 0.991789764178166 |
| chr15-37821625-37822126 | 8 | 0.909569229231866 |
| chr3-11614823-11615324 | 4 | 0.959230243157655 |
| chr16-33318626-33319127 | 8 | 0.958451616952749 |
| chr4-30789617-30790118 | 8 | 0.906871836421652 |
| chr19-18030140-18030641 | 4 | 0.972424114656692 |
| chr11-43644392-43644893 | 3 | 0.967505195220091 |
| chr11-63069430-63069931 | 4 | 0.892512160411634 |
| chr4-141986023-141986524 | 6 | 0.977197348724259 |
| chr7-31698492-31698993 | 8 | 0.910765893604045 |
| chr15-21750766-21751267 | 6 | 0.939786370494039 |
| chr7-29167568-29168069 | 7 | 0.928282518208004 |
| chr2-112615706-112616207 | 2 | 0.831333973629894 |
| chr13-50341166-50341667 | 3 | 0.968408698116157 |
| chr1-32227574-32228075 | 6 | 0.991206925481043 |
| chr14-83768872-83769373 | 3 | 0.862027480861691 |
| chr2-10953964-10954465 | 8 | 0.993493681128311 |
| chr19-12002907-12003408 | 8 | 0.964595866236607 |
| chr7-126524196-126524697 | 3 | 0.99231489434187 |
| chr4-102280648-102281149 | 3 | 0.840996350025199 |
| chr6-8933914-8934415 | 8 | 0.975275106523284 |
| chr7-22369630-22370131 | 3 | 0.996057811158863 |
| chr6-95243363-95243864 | 3 | 0.975812542968088 |
| chr16-61814906-61815407 | 3 | 0.984390457737135 |
| chr19-136229-136730 | 4 | 0.860508046457377 |
| chr14-34530906-34531407 | 2 | 0.886443221502221 |
| chr13-16129888-16130389 | 8 | 0.97188294866849 |
| chrX-151233569-151234070 | 5 | 0.938982190708354 |

| | | |
|---|---|---|
| chr10-75089669-75090170 | 8 | 0.908748306793406 |
| chr9-111062316-111062817 | 8 | 0.966669104376727 |
| chr13-67112203-67112704 | 8 | 0.978430624896684 |
| chr4-74717609-74718110 | 3 | 0.99593553545041 |
| chr4-146988719-146989220 | 3 | 0.967892432060966 |
| chr5-9235806-9236307 | 8 | 0.991216864382594 |
| chr18-59343087-59343588 | 8 | 0.991513915800637 |
| chr2-151547429-151547930 | 5 | 0.946281801719312 |
| chr16-44018272-44018773 | 5 | 0.913337461579965 |
| chr20-42419754-42420255 | 7 | 0.912355409436322 |
| chr4-102241626-102242127 | 8 | 0.922088779073622 |
| chr2-240561907-240562408 | 6 | 0.989176547286718 |
| chr5-137492902-137493403 | 8 | 0.993493681128311 |
| chr13-48029728-48030229 | 2 | 0.931947601463849 |
| chr13-37841204-37841705 | 3 | 0.99593553545041 |
| chr3-114873215-114873716 | 8 | 0.986425771992889 |
| chr12-28010377-28010878 | 8 | 0.983676284557418 |
| chr17-8778760-8779261 | 5 | 0.985836706142151 |
| chr7-40555557-40556058 | 6 | 0.9670538161949 |
| chr14-77128228-77128729 | 8 | 0.97695674724735 |
| chr15-40823331-40823832 | 8 | 0.939013987328525 |
| chr3-78986772-78987273 | 6 | 0.873487658206657 |
| chr3-36852221-36852722 | 4 | 0.857452371258842 |
| chr10-104766954-104767455 | 4 | 0.820307844059532 |
| chr3-26244084-26244585 | 6 | 0.994211830739455 |
| chr16-73707060-73707561 | 3 | 0.954005107747979 |
| chr14-64506349-64506850 | 3 | 0.928580141259335 |
| chr20-15592087-15592588 | 6 | 0.991967081097411 |
| chr5-131774299-131774800 | 7 | 0.825643340846191 |
| chr5-62861315-62861816 | 8 | 0.910033845318313 |
| chr9-64888987-64889488 | 8 | 0.965355057576191 |
| chr7-69693624-69694125 | 8 | 0.938759775318672 |
| chr16-77849946-77850447 | 3 | 0.901366815341638 |
| chr5-24357542-24358043 | 5 | 0.983919978840557 |
| chr8-41028206-41028707 | 8 | 0.994654903899323 |
| chr9-105199507-105200008 | 4 | 0.870834695626234 |
| chr9-24669744-24670245 | 8 | 0.992182792411678 |
| chr5-135054880-135055381 | 5 | 0.989137558831664 |
| chr17-63425381-63425882 | 5 | 0.983919978840557 |
| chr7-108817925-108818426 | 8 | 0.92000421133097 |
| chr2-246717786-246718287 | 8 | 0.972650826332159 |
| chr7-23942024-23942525 | 4 | 0.854951494752092 |
| chr11-52932088-52932589 | 8 | 0.990989839507847 |

| | | |
|---|---|---|
| chr9-89480010-89480511 | 8 | 0.822566232300548 |
| chr6-139506343-139506844 | 8 | 0.814746756471836 |
| chr1-44434263-44434764 | 4 | 0.825524207736577 |
| chr1-202080037-202080538 | 8 | 0.835292601278795 |
| chr3-44823199-44823700 | 5 | 0.993131177854135 |
| chr4-7168912-7169413 | 5 | 0.950862490084422 |
| chr1-49282616-49283117 | 3 | 0.99593553545041 |
| chr2-157010718-157011219 | 3 | 0.996530804082791 |
| chr12-37350918-37351419 | 8 | 0.967582695171725 |
| chr17-84628375-84628876 | 6 | 0.997395485916031 |
| chr1-84867231-84867732 | 3 | 0.967747555946142 |
| chr3-93186976-93187477 | 8 | 0.972515814960066 |
| chr4-56505840-56506341 | 8 | 0.900748164933302 |
| chr10-34329736-34330237 | 3 | 0.99593553545041 |
| chr2-149842366-149842867 | 2 | 0.927711983469644 |
| chr11-67296774-67297275 | 8 | 0.832288553584912 |
| chr1-213714205-213714706 | 1 | 0.866988962473118 |
| chr12-13192402-13192903 | 8 | 0.899258451954816 |
| chr13-48370077-48370578 | 3 | 0.891452998626166 |
| chr20-26409360-26409861 | 6 | 0.918606741164188 |
| chr16-47371356-47371857 | 8 | 0.961129314060522 |
| chr3-142642722-142643223 | 3 | 0.997142031304554 |
| chr12-7515143-7515644 | 8 | 0.922868521403394 |
| chr3-165404272-165404773 | 3 | 0.997838988345321 |
| chr7-100415204-100415705 | 8 | 0.957834974876871 |
| chr2-207895312-207895813 | 8 | 0.993493681128311 |
| chr9-98555069-98555570 | 8 | 0.854907629035767 |
| chr17-52359328-52359829 | 5 | 0.984151869537007 |
| chr15-99163364-99163865 | 8 | 0.829481879834258 |
| chrX-21969860-21970361 | 3 | 0.818800176961901 |
| chr4-115030708-115031209 | 2 | 0.969928935739881 |
| chr7-28100020-28100521 | 4 | 0.883825060454567 |
| chr13-91128179-91128680 | 6 | 0.947561383518795 |
| chr1-255904676-255905177 | 8 | 0.884883645089331 |
| chr1-192215938-192216439 | 8 | 0.945796547469823 |
| chr9-50316069-50316570 | 8 | 0.85812257064483 |
| chr9-13843198-13843699 | 2 | 0.929391633529218 |
| chr17-32927381-32927882 | 3 | 0.95540145921692 |
| chr10-17682385-17682886 | 5 | 0.973471425932334 |
| chr2-206001000-206001501 | 6 | 0.991206925481043 |
| chr2-216900937-216901438 | 6 | 0.831867175326496 |
| chr1-198602694-198603195 | 5 | 0.835720096104101 |
| chr7-65038181-65038682 | 8 | 0.993493681128311 |

| | | |
|---|---|---|
| chr19-40179752-40180253 | 3 | 0.989942282057779 |
| chr3-77529508-77530009 | 4 | 0.892910941936716 |
| chr11-59366574-59367075 | 8 | 0.93396316925438 |
| chr1-78455550-78456051 | 3 | 0.983055014916196 |
| chr14-94198752-94199253 | 5 | 0.992450002181107 |
| chr7-110964866-110965367 | 5 | 0.958417391646968 |
| chr2-198288073-198288574 | 3 | 0.992025758129544 |
| chr5-119241018-119241519 | 8 | 0.995248741195107 |
| chr16-19725222-19725723 | 5 | 0.958245352890176 |
| chr11-54486392-54486893 | 5 | 0.884489892208203 |
| chr5-133620981-133621482 | 5 | 0.874948531095612 |
| chr15-15575709-15576210 | 6 | 0.993207206135076 |
| chr12-12904042-12904543 | 6 | 0.993201612459791 |
| chr4-126299274-126299775 | 8 | 0.889296312425967 |
| chr4-153650294-153650795 | 4 | 0.963274363126895 |
| chr14-15878684-15879185 | 2 | 0.872793290892721 |
| chr1-254823244-254823745 | 8 | 0.960214691427428 |
| chr9-111895115-111895616 | 6 | 0.880965421763409 |
| chr4-15531382-15531883 | 6 | 0.90271680396468 |
| chr5-151737461-151737962 | 8 | 0.953312787530892 |
| chr3-132978955-132979456 | 8 | 0.993820346326267 |
| chr2-116967370-116967871 | 8 | 0.993493681128311 |
| chr4-94189714-94190215 | 8 | 0.993493681128311 |
| chr7-52517681-52518182 | 8 | 0.842682490971149 |
| chr9-100024928-100025429 | 6 | 0.99283156371011 |
| chrX-3363302-3363803 | 3 | 0.954906160808171 |
| chr12-29416385-29416886 | 3 | 0.996880910700886 |
| chr15-85682419-85682920 | 4 | 0.942344268095434 |
| chr10-43014962-43015463 | 6 | 0.990563221835028 |
| chr3-130394031-130394532 | 3 | 0.996328443495776 |
| chr13-81569759-81570260 | 3 | 0.857339477042407 |
| chr14-18967455-18967956 | 5 | 0.993411993829884 |
| chr2-33391588-33392089 | 7 | 0.929450660550709 |
| chr1-46584967-46585468 | 8 | 0.946012968808814 |
| chr2-192686449-192686950 | 8 | 0.932434417072654 |
| chr8-115610126-115610627 | 8 | 0.874271353133208 |
| chr9-12634751-12635252 | 2 | 0.929472104251184 |
| chr10-73625653-73626154 | 1 | 0.840781823656798 |
| chr1-19293558-19294059 | 1 | 0.922142590579484 |
| chr18-83326585-83327086 | 3 | 0.99593553545041 |
| chr1-7690511-7691012 | 5 | 0.879142581477179 |
| chr3-112527874-112528375 | 8 | 0.894432524888255 |
| chr17-64498321-64498822 | 1 | 0.945788116356725 |

| | | |
|---|---|---|
| chrX-126822070-126822571 | 5 | 0.941158521050047 |
| chr5-144821287-144821788 | 8 | 0.952087532138997 |
| chr4-72703240-72703741 | 8 | 0.980939085812107 |
| chr3-112504955-112505456 | 8 | 0.993346865058617 |
| chr7-46798480-46798981 | 4 | 0.972591354160969 |
| chr3-18660374-18660875 | 8 | 0.889343134622326 |
| chr6-47905472-47905973 | 2 | 0.924208263988828 |
| chr5-62208487-62208988 | 1 | 0.960165988309335 |
| chr9-49031803-49032304 | 3 | 0.946189744291746 |
| chr8-61362277-61362778 | 3 | 0.996420894665031 |
| chr3-135792041-135792542 | 1 | 0.927236853530569 |
| chr2-164909890-164910391 | 3 | 0.99387445873643 |
| chr10-33930742-33931243 | 6 | 0.951980210414729 |
| chr1-226473487-226473988 | 8 | 0.983813297798762 |
| chr2-198060907-198061408 | 3 | 0.995094180703403 |
| chr20-21937111-21937612 | 8 | 0.979379496458008 |
| chrX-113772765-113773266 | 3 | 0.99593553545041 |
| chr20-26869451-26869952 | 5 | 0.966418071382306 |
| chr20-52417101-52417602 | 4 | 0.970624015846191 |
| chr1-20395310-20395811 | 4 | 0.919603992089752 |
| chr9-92887587-92888088 | 8 | 0.931915810981369 |
| chr10-69019103-69019604 | 3 | 0.993832797987757 |
| chr5-82693190-82693691 | 6 | 0.991206925481043 |
| chr7-62568414-62568915 | 3 | 0.99593553545041 |
| chr17-81041468-81041969 | 8 | 0.970811876430574 |
| chr17-62628389-62628890 | 6 | 0.99638836466507 |
| chr3-60474402-60474903 | 1 | 0.891993166827653 |
| chr4-29071107-29071608 | 4 | 0.879780305488875 |
| chr15-60209475-60209976 | 6 | 0.991206925481043 |
| chr6-92568161-92568662 | 6 | 0.983702507274721 |
| chr3-167368917-167369418 | 8 | 0.951579988513854 |
| chr7-115024441-115024942 | 3 | 0.995951796341764 |
| chr13-101504250-101504751 | 8 | 0.993792561356577 |
| chr8-46842685-46843186 | 8 | 0.949387127814022 |
| chr4-110493622-110494123 | 6 | 0.991427686799978 |
| chr15-31979217-31979718 | 2 | 0.909838751069905 |
| chr1-253926614-253927115 | 3 | 0.996875742055646 |
| chr2-117763924-117764425 | 5 | 0.989670931557546 |
| chr4-103639697-103640198 | 5 | 0.881259730414794 |
| chr1-171704150-171704651 | 2 | 0.868622691309784 |
| chr14-98300491-98300992 | 5 | 0.970768370227948 |
| chr1-232252614-232253115 | 8 | 0.996183895941798 |
| chr13-10665278-10665779 | 3 | 0.959949880906965 |

| | | |
|---|---|---|
| chr4-110006162-110006663 | 2 | 0.942004551405862 |
| chr4-173700121-173700622 | 4 | 0.933754463364561 |
| chr1-234393637-234394138 | 8 | 0.995226168836144 |
| chr3-72050311-72050812 | 6 | 0.89346442147869 |
| chr13-98915824-98916325 | 8 | 0.917954492887203 |
| chr14-72403826-72404327 | 3 | 0.925296647167284 |
| chr2-152910969-152911470 | 4 | 0.858287389322552 |
| chr5-114273698-114274199 | 6 | 0.993132993838828 |
| chr15-82636119-82636620 | 5 | 0.881251230905553 |
| chr1-180741723-180742224 | 8 | 0.881140277085518 |
| chr19-54201792-54202293 | 8 | 0.956556162994583 |
| chr19-53941497-53941998 | 5 | 0.887599485120456 |
| chr9-58272588-58273089 | 3 | 0.997225870789829 |
| chr5-118558922-118559423 | 6 | 0.992241257050088 |
| chr5-116887723-116888224 | 5 | 0.870255797446729 |
| chr17-14965799-14966300 | 8 | 0.954904714849397 |
| chr3-145766009-145766510 | 5 | 0.916355535852155 |
| chr1-48777531-48778032 | 7 | 0.883667933707357 |
| chr5-31415848-31416349 | 3 | 0.99593553545041 |
| chr7-108716707-108717208 | 8 | 0.864383696508686 |
| chr7-83127180-83127681 | 8 | 0.947985999779025 |
| chr7-99038730-99039231 | 8 | 0.975143761637691 |
| chr15-40799101-40799602 | 3 | 0.871240583771724 |
| chr11-85401917-85402418 | 3 | 0.998922725492795 |
| chr17-7789399-7789900 | 4 | 0.873809727886881 |
| chr6-44760122-44760623 | 8 | 0.994868702702304 |
| chr4-35417653-35418154 | 3 | 0.948961033288849 |
| chr14-3397352-3397853 | 4 | 0.973130040819077 |
| chr19-50000735-50001236 | 5 | 0.956658047702085 |
| chr7-109890861-109891362 | 2 | 0.89154678727804 |
| chr2-139092418-139092919 | 4 | 0.972803814502132 |
| chr9-38290414-38290915 | 5 | 0.990508243761936 |
| chr1-155429719-155430220 | 5 | 0.969244060239526 |
| chr2-220507151-220507652 | 7 | 0.898481865675115 |
| chr16-40041042-40041543 | 3 | 0.986222037570618 |
| chr7-122121959-122122460 | 5 | 0.969479358791703 |
| chr16-55147072-55147573 | 3 | 0.855616248847579 |
| chr4-58057991-58058492 | 4 | 0.929076293083328 |
| chr7-81803438-81803939 | 4 | 0.932546797213151 |
| chr2-13967005-13967506 | 6 | 0.948066316089693 |
| chr1-95210648-95211149 | 4 | 0.924378060313194 |
| chr15-19644389-19644890 | 8 | 0.860119891051683 |
| chr13-35022962-35023463 | 6 | 0.991206925481043 |

| | | |
|---|---|---|
| chr9-104569075-104569576 | 8 | 0.994143412189408 |
| chr19-48696727-48697228 | 8 | 0.940929243208771 |
| chr6-45251128-45251629 | 8 | 0.868200015125831 |
| chr15-18158414-18158915 | 4 | 0.916571747575771 |
| chr8-105067798-105068299 | 8 | 0.993493681128311 |
| chr11-54298672-54299173 | 2 | 0.865228407054128 |
| chr16-24799386-24799887 | 6 | 0.991206925481043 |
| chr6-89389993-89390494 | 8 | 0.980641307928714 |
| chr3-162145351-162145852 | 8 | 0.954470807611118 |
| chr18-12521893-12522394 | 8 | 0.994927113989791 |
| chr6-91616494-91616995 | 8 | 0.995062468678045 |
| chr17-51308549-51309050 | 5 | 0.978518745268174 |
| chr10-1155402-1155903 | 3 | 0.968662797690413 |
| chr3-64408644-64409145 | 8 | 0.842999908606208 |
| chr14-93455110-93455611 | 8 | 0.889087880935013 |
| chr7-48337610-48338111 | 6 | 0.93036269680519 |
| chr1-138141806-138142307 | 2 | 0.883341394473889 |
| chr9-95991068-95991569 | 8 | 0.89360573275846 |
| chr8-96985944-96986445 | 4 | 0.923558095114883 |
| chr10-36740193-36740694 | 5 | 0.990118120508577 |
| chr10-93683520-93684021 | 8 | 0.972604668816793 |
| chr3-106444458-106444959 | 8 | 0.818975878919481 |
| chr7-114223422-114223923 | 7 | 0.834477728535106 |
| chr4-151409781-151410282 | 6 | 0.991206925481043 |
| chr5-160065542-160066043 | 8 | 0.884632448107965 |
| chr7-26378352-26378853 | 3 | 0.99593553545041 |
| chr20-19151463-19151964 | 8 | 0.973778800431811 |
| chr6-56597620-56598121 | 3 | 0.880104358259155 |
| chrX-42949931-42950432 | 3 | 0.919840913048293 |
| chr1-154281661-154282162 | 4 | 0.81905753122183 |
| chr13-67193676-67194177 | 8 | 0.969792655493147 |
| chr8-37982226-37982727 | 3 | 0.995903166523414 |
| chrX-80367926-80368427 | 8 | 0.993493681128311 |
| chr20-47323780-47324281 | 4 | 0.889855392796875 |
| chr5-141727148-141727649 | 3 | 0.872346670888276 |
| chr19-25364573-25365074 | 8 | 0.993493681128311 |
| chr17-82617108-82617609 | 8 | 0.993493681128311 |
| chr16-44549581-44550082 | 3 | 0.918984694005333 |
| chr4-154858133-154858634 | 5 | 0.885467529362917 |
| chr10-53633250-53633751 | 4 | 0.878698729161787 |
| chr5-25436822-25437323 | 8 | 0.994999804958734 |
| chr7-592856-593357 | 8 | 0.842340749638387 |
| chr17-29970322-29970823 | 4 | 0.866047481196123 |

| | | |
|---|---|---|
| chr6-71680740-71681241 | 8 | 0.993493681128311 |
| chr5-113672130-113672631 | 8 | 0.99518939178809 |
| chr15-13630874-13631375 | 6 | 0.993212814131786 |
| chr12-37146145-37146646 | 8 | 0.923836178668215 |
| chr3-117919612-117920113 | 8 | 0.963570139808583 |
| chr3-141415482-141415983 | 4 | 0.922835114872311 |
| chr5-150878541-150879042 | 7 | 0.851157906084157 |
| chr7-69418560-69419061 | 3 | 0.964235678267982 |
| chr10-44779756-44780257 | 7 | 0.942020146145419 |
| chr2-136990779-136991280 | 3 | 0.831550753468024 |
| chr17-55657380-55657881 | 8 | 0.993493681128311 |
| chr13-105293955-105294456 | 3 | 0.99593553545041 |
| chr5-98852444-98852945 | 4 | 0.888658637621183 |
| chr7-55143722-55144223 | 3 | 0.995986758621932 |
| chr5-135991690-135992191 | 5 | 0.975635391182239 |
| chr10-93771828-93772329 | 8 | 0.983879561315498 |
| chr1-260393626-260394127 | 3 | 0.995291490405118 |
| chr16-7968595-7969096 | 8 | 0.990826315484373 |
| chr15-93566089-93566590 | 3 | 0.975869183819569 |
| chr8-120672034-120672535 | 4 | 0.832911010277153 |
| chr20-29374386-29374887 | 5 | 0.983877222154974 |
| chr4-162273992-162274493 | 4 | 0.888242453397482 |
| chr2-61884086-61884587 | 7 | 0.961670285378287 |
| chr10-5887945-5888446 | 5 | 0.964430477014113 |
| chr14-11904533-11905034 | 8 | 0.887139476104502 |
| chr6-49724439-49724940 | 8 | 0.894107586611241 |
| chr15-28895369-28895870 | 8 | 0.941417582949894 |
| chr3-123305044-123305545 | 8 | 0.892468468967757 |
| chr14-101133630-101134131 | 3 | 0.964673771419438 |
| chr8-66328590-66329091 | 8 | 0.972492115140076 |
| chr1-130036980-130037481 | 3 | 0.997140030988367 |
| chr1-218802934-218803435 | 2 | 0.958720173990005 |
| chr19-704210-704711 | 8 | 0.993493681128311 |
| chr15-99345434-99345935 | 8 | 0.915635671022847 |
| chr17-23980685-23981186 | 4 | 0.954776012568669 |
| chr5-136790039-136790540 | 5 | 0.832785194438548 |
| chr1-69803520-69804021 | 8 | 0.993923040384983 |
| chr3-41878976-41879477 | 6 | 0.993774610613793 |
| chr13-55767049-55767550 | 8 | 0.915372068763032 |
| chr2-81965160-81965661 | 8 | 0.96687932671758 |
| chr1-41137206-41137707 | 8 | 0.993493681128311 |
| chr9-81564706-81565207 | 4 | 0.905783889552666 |
| chr2-33055860-33056361 | 6 | 0.917303491285628 |

| | | |
|---|---|---|
| chr20-25238016-25238517 | 5 | 0.978112878783037 |
| chr1-213197741-213198242 | 2 | 0.877629382750471 |
| chr9-62662766-62663267 | 4 | 0.951802328711424 |
| chr14-76590651-76591152 | 5 | 0.912959352591762 |
| chr15-84581483-84581984 | 3 | 0.99593553545041 |
| chr17-36100460-36100961 | 8 | 0.982662607526936 |
| chr8-100039407-100039908 | 2 | 0.833633612552664 |
| chr10-102899763-102900264 | 2 | 0.903495001512691 |
| chr8-25022544-25023045 | 6 | 0.990793452844653 |
| chr16-58983168-58983669 | 1 | 0.928068024844144 |
| chr9-86510689-86511190 | 8 | 0.985053054043863 |
| chr14-11929363-11929864 | 8 | 0.927309086161437 |
| chr7-73625936-73626437 | 3 | 0.9977431720341 |
| chr8-105357771-105358272 | 3 | 0.982069574244413 |
| chr13-42103516-42104017 | 6 | 0.992314594543223 |
| chr6-10169050-10169551 | 8 | 0.984919426989496 |
| chr2-133213766-133214267 | 3 | 0.99593553545041 |
| chr4-150761654-150762155 | 1 | 0.908666196702764 |
| chr10-82595650-82596151 | 1 | 0.953873547139783 |
| chr7-109591247-109591748 | 8 | 0.932350797408208 |
| chr4-80907722-80908223 | 8 | 0.960610743855653 |
| chr2-8415322-8415823 | 8 | 0.95777550086026 |
| chr5-133886583-133887084 | 3 | 0.998435883111036 |
| chr2-96280052-96280553 | 5 | 0.904793118348362 |
| chr16-58743156-58743657 | 3 | 0.996581367801233 |
| chr5-113803269-113803770 | 8 | 0.959149318456012 |
| chr3-156419160-156419661 | 8 | 0.874797343605167 |
| chr16-58958693-58959194 | 1 | 0.879344890512478 |
| chr1-91859827-91860328 | 8 | 0.916673952983507 |
| chr15-34705828-34706329 | 1 | 0.822820856233979 |
| chr3-52727187-52727688 | 5 | 0.961282015322314 |
| chr1-243268329-243268830 | 8 | 0.995201208781541 |
| chr1-90650347-90650848 | 2 | 0.97069583798217 |
| chr14-21390223-21390724 | 6 | 0.925936045898179 |
| chr14-66789969-66790470 | 6 | 0.978190344344417 |
| chr2-196316540-196317041 | 8 | 0.982804548890128 |
| chrX-69602348-69602849 | 7 | 0.853465349371732 |
| chr19-46802639-46803140 | 8 | 0.840855426238499 |
| chr3-134080020-134080521 | 3 | 0.99593553545041 |
| chr8-5872945-5873446 | 4 | 0.961750071178646 |
| chr8-69533111-69533612 | 5 | 0.81909078455579 |
| chr12-1587413-1587914 | 6 | 0.97796700198602 |
| chr13-76799999-76800500 | 8 | 0.864053290585473 |

| | | |
|---|---|---|
| chr5-65261463-65261964 | 6 | 0.98904736082871 |
| chr3-93358799-93359300 | 8 | 0.878264284652003 |
| chr14-17940332-17940833 | 5 | 0.979980115580396 |
| chr1-167177641-167178142 | 3 | 0.92724825966625 |
| chr5-40423910-40424411 | 7 | 0.964424731961184 |
| chr16-59925892-59926393 | 5 | 0.966605094630264 |
| chr17-53233396-53233897 | 1 | 0.851895120361856 |
| chrX-71347149-71347650 | 3 | 0.996702279562969 |
| chr1-255495960-255496461 | 8 | 0.942597354007354 |
| chrX-47512546-47513047 | 3 | 0.99593553545041 |
| chr3-88524407-88524908 | 3 | 0.996898035496883 |
| chr17-48452881-48453382 | 4 | 0.845490421331959 |
| chr2-189022002-189022503 | 3 | 0.897086930825462 |
| chr18-54082072-54082573 | 1 | 0.896064031474202 |
| chr10-93851834-93852335 | 8 | 0.931208029848363 |
| chr2-160608857-160609358 | 3 | 0.991234024619994 |
| chr3-18569192-18569693 | 4 | 0.900492348997741 |
| chr10-87005368-87005869 | 6 | 0.951198441492886 |
| chr9-53124147-53124648 | 3 | 0.996869166363192 |
| chr2-166885056-166885557 | 6 | 0.82219583154429 |
| chr4-93719655-93720156 | 6 | 0.990595827032828 |
| chr14-78144411-78144912 | 6 | 0.989759845512388 |
| chr19-28484075-28484576 | 8 | 0.982631198663929 |
| chr6-100333874-100334375 | 3 | 0.959220572142537 |
| chr6-82185568-82186069 | 3 | 0.99593553545041 |
| chr13-64464760-64465261 | 6 | 0.981629414472224 |
| chr11-42523439-42523940 | 8 | 0.81832781585108 |
| chr13-105233061-105233562 | 3 | 0.99679599987285 |
| chr2-44341725-44342226 | 8 | 0.87167 |
| chr7-57884109-57884610 | 2 | 0.915676404093272 |
| chr5-72489689-72490190 | 8 | 0.942265062234253 |
| chr9-98684262-98684763 | 8 | 0.955063646289455 |
| chr5-137797472-137797973 | 6 | 0.864716124702042 |
| chr7-70180049-70180550 | 6 | 0.975530373587872 |
| chr9-26462242-26462743 | 3 | 0.91084157939034 |
| chr6-107126045-107126546 | 6 | 0.991558720825738 |
| chr15-52200831-52201332 | 8 | 0.898363724651711 |
| chr7-114786323-114786824 | 6 | 0.991651925508046 |
| chr2-104255530-104256031 | 8 | 0.945895814529706 |
| chr8-61145965-61146466 | 4 | 0.903054335441641 |
| chr9-22942039-22942540 | 3 | 0.858475195081983 |
| chr3-101405315-101405816 | 1 | 0.871179612901306 |
| chr4-145885276-145885777 | 2 | 0.816834887036522 |

| | | |
|---|---|---|
| chr13-30980246-30980747 | 2 | 0.827243818841956 |
| chr20-10227345-10227846 | 4 | 0.858406896634838 |
| chr11-1930523-1931024 | 4 | 0.874817084169779 |
| chr7-48041053-48041554 | 6 | 0.93971881730174 |
| chr18-16262666-16263167 | 8 | 0.961479453799751 |
| chrX-70451495-70451996 | 6 | 0.958873066589008 |
| chr5-71033427-71033928 | 8 | 0.98961754681017 |
| chr13-44091316-44091817 | 8 | 0.994774762914746 |
| chr19-49863491-49863992 | 5 | 0.970276658685935 |
| chr4-11516019-11516520 | 2 | 0.947037171885825 |
| chr7-8737767-8738268 | 4 | 0.920501603096769 |
| chr7-18175776-18176277 | 6 | 0.992891508165551 |
| chr5-121941547-121942048 | 1 | 0.841136666515984 |
| chr20-26984571-26985072 | 4 | 0.83459198155916 |
| chr17-40552242-40552743 | 6 | 0.992187084630191 |
| chr14-78986309-78986810 | 8 | 0.820628682705233 |
| chr2-224406422-224406923 | 6 | 0.991206925481043 |
| chr10-86067492-86067993 | 3 | 0.838972639317263 |
| chr12-40661972-40662473 | 8 | 0.885286638535782 |
| chr8-54217638-54218139 | 4 | 0.861301585704436 |
| chr18-43846104-43846605 | 6 | 0.866289821601971 |
| chr6-44828979-44829480 | 3 | 0.965281178671686 |
| chr9-103748991-103749492 | 6 | 0.862679835286802 |
| chr16-29038926-29039427 | 8 | 0.892071593159105 |
| chr8-80931314-80931815 | 2 | 0.96400657962276 |
| chr1-91776156-91776657 | 3 | 0.995826755335997 |
| chr5-40689193-40689694 | 3 | 0.99593553545041 |
| chr1-30283578-30284079 | 8 | 0.994134624377324 |
| chr2-217609380-217609881 | 3 | 0.993886571050433 |
| chr15-14591068-14591569 | 5 | 0.973508787277361 |
| chr14-32677977-32678478 | 8 | 0.995104592802646 |
| chr6-79807263-79807764 | 6 | 0.842936230870311 |
| chr8-100661497-100661998 | 5 | 0.858914788421555 |
| chr19-43625432-43625933 | 3 | 0.998653384831517 |
| chr1-221846222-221846723 | 5 | 0.981329332789157 |
| chr19-38971866-38972367 | 5 | 0.965768844003827 |
| chr7-125478624-125479125 | 4 | 0.961214849438855 |
| chr18-39708765-39709266 | 5 | 0.977236392549118 |
| chr8-88382567-88383068 | 8 | 0.940258867369858 |
| chr6-74973065-74973566 | 3 | 0.991641991352647 |
| chr17-18096048-18096549 | 5 | 0.83222449578098 |
| chr15-52529243-52529744 | 8 | 0.99426380500675 |
| chr1-124537942-124538443 | 4 | 0.928936494902355 |

| | | |
|---|---|---|
| chr11-44733837-44734338 | 5 | 0.918959118449569 |
| chr10-98648405-98648906 | 6 | 0.986840189104527 |
| chr3-125048891-125049392 | 1 | 0.893992498477396 |
| chr2-31518607-31519108 | 4 | 0.927066975140135 |
| chr11-49643212-49643713 | 8 | 0.880278047123374 |
| chr7-133078221-133078722 | 8 | 0.912709211786727 |
| chr19-14021545-14022046 | 8 | 0.896654256574651 |
| chr8-79318084-79318585 | 2 | 0.962524892895887 |
| chr2-243099699-243100200 | 3 | 0.995868675120575 |
| chrX-74307576-74308077 | 5 | 0.954908465884428 |
| chr2-102213841-102214342 | 7 | 0.959269896260796 |
| chr4-33043800-33044301 | 5 | 0.991532498634628 |
| chr9-32967068-32967569 | 8 | 0.835040898885527 |
| chr16-15135339-15135840 | 3 | 0.975111918024787 |
| chr6-57024334-57024835 | 5 | 0.81465904973919 |
| chr1-183410367-183410868 | 8 | 0.943063098174119 |
| chr6-140484245-140484746 | 5 | 0.97517939591677 |
| chr18-35169544-35170045 | 3 | 0.996794459847098 |
| chr9-1969211-1969712 | 2 | 0.888152575747339 |
| chr5-27504662-27505163 | 4 | 0.827550714120918 |
| chr3-135922576-135923077 | 1 | 0.85992464520185 |
| chr18-46563952-46564453 | 4 | 0.920422365290137 |
| chr12-20305303-20305804 | 8 | 0.86642934961735 |
| chr5-138085610-138086111 | 8 | 0.938526352018234 |
| chr4-115307316-115307817 | 6 | 0.98984120235888 |
| chr4-5345963-5346464 | 5 | 0.955363874206984 |
| chr2-123362120-123362621 | 8 | 0.957927382032707 |
| chr17-56579344-56579845 | 8 | 0.819033841662325 |
| chr13-76227317-76227818 | 6 | 0.992885663627174 |
| chr8-53474994-53475495 | 3 | 0.995837258032525 |
| chr3-168150817-168151318 | 4 | 0.937408408659082 |
| chr13-44111239-44111740 | 8 | 0.943279129330965 |
| chr19-43590898-43591399 | 5 | 0.948617185209338 |
| chr4-51223645-51224146 | 8 | 0.843214023203124 |
| chr6-126786310-126786811 | 3 | 0.988913387446388 |
| chr5-79301619-79302120 | 8 | 0.866267309959987 |
| chr4-101857898-101858399 | 8 | 0.815130043848162 |
| chr10-3474929-3475430 | 3 | 0.987897532212811 |
| chr9-56463507-56464008 | 8 | 0.884782482933123 |
| chr15-37871551-37872052 | 6 | 0.986990865329594 |
| chr8-110617053-110617554 | 6 | 0.968691918623808 |
| chr10-66111828-66112329 | 7 | 0.837803991289048 |
| chr9-92681993-92682494 | 8 | 0.92162484233811 |

| | | |
|---|---|---|
| chr13-87151584-87152085 | 6 | 0.99638196715315 |
| chr1-156893628-156894129 | 8 | 0.954709072440635 |
| chr17-45640311-45640812 | 2 | 0.9113289091205 |
| chr6-125987860-125988361 | 3 | 0.970618309078551 |
| chr12-40918621-40919122 | 5 | 0.957462768006145 |
| chr17-79774219-79774720 | 3 | 0.996666637414586 |
| chr12-26794273-26794774 | 8 | 0.913678061360305 |
| chr2-183865246-183865747 | 8 | 0.923468205466468 |
| chr6-34972316-34972817 | 4 | 0.823977279798931 |
| chr4-94149067-94149568 | 5 | 0.972585784890275 |
| chr13-47021899-47022400 | 7 | 0.896919771092739 |
| chr13-89746135-89746636 | 8 | 0.914595820984219 |
| chr6-72285232-72285733 | 8 | 0.895619497853037 |
| chrX-137580524-137581025 | 8 | 0.931543735751347 |
| chr4-144914737-144915238 | 3 | 0.99593553545041 |
| chrX-28246661-28247162 | 6 | 0.976241445611364 |
| chr2-52540755-52541256 | 8 | 0.983837641373446 |
| chr3-1247903-1248404 | 3 | 0.997726689962805 |
| chr14-39865263-39865764 | 3 | 0.99628665228603 |
| chr14-84416462-84416963 | 6 | 0.985795488513594 |
| chr6-36589657-36590158 | 6 | 0.959149443338982 |
| chr3-128284070-128284571 | 6 | 0.877582958079052 |
| chr13-37152000-37152501 | 1 | 0.912034495600694 |
| chr10-56250376-56250877 | 5 | 0.928044804182707 |
| chr1-190425860-190426361 | 6 | 0.995103594350473 |
| chr11-29745888-29746389 | 3 | 0.996874503534431 |
| chr9-60458673-60459174 | 5 | 0.95991211985239 |
| chr1-51089531-51090032 | 3 | 0.99593553545041 |
| chr1-86118476-86118977 | 8 | 0.855645520997631 |
| chrX-128689915-128690416 | 6 | 0.980533021565667 |
| chr17-80418602-80419103 | 6 | 0.991206925481043 |
| chr2-21851530-21852031 | 1 | 0.827877766770768 |
| chr2-74664349-74664850 | 8 | 0.913983211447528 |
| chr5-152071285-152071786 | 4 | 0.939355816940798 |
| chr6-66224789-66225290 | 8 | 0.993493681128311 |
| chr7-8892973-8893474 | 8 | 0.966047732403729 |
| chr16-55362096-55362597 | 5 | 0.937565298093581 |
| chr4-152430931-152431432 | 5 | 0.932991924044917 |
| chr9-74347085-74347586 | 5 | 0.94463130352916 |
| chr8-98660460-98660961 | 8 | 0.871724852747858 |
| chr4-53051846-53052347 | 3 | 0.99526255473666 |
| chr3-111417472-111417973 | 8 | 0.993493681128311 |
| chr4-85043245-85043746 | 8 | 0.894131932394128 |

| | | |
|---|---|---|
| chr15-1051469-1051970 | 8 | 0.966418117356272 |
| chr4-34451627-34452128 | 3 | 0.99593553545041 |
| chr2-234357693-234358194 | 7 | 0.958558753196486 |
| chr1-43972111-43972612 | 7 | 0.831877296837365 |
| chr10-23664772-23665273 | 6 | 0.991206925481043 |
| chr5-151626518-151627019 | 4 | 0.932279518810101 |
| chr12-40139059-40139560 | 8 | 0.889190050764598 |
| chr5-36787691-36788192 | 2 | 0.839148080178284 |
| chr3-12393847-12394348 | 5 | 0.978529829015398 |
| chr16-62678863-62679364 | 6 | 0.824172545505891 |
| chr13-94490831-94491332 | 8 | 0.947282824430835 |
| chr2-30131583-30132084 | 3 | 0.984026186301508 |
| chr3-70430153-70430654 | 1 | 0.910552737362399 |
| chr2-47068116-47068617 | 5 | 0.883569663810251 |
| chr18-76267095-76267596 | 6 | 0.815685318372338 |
| chr13-72311093-72311594 | 1 | 0.895416891419006 |
| chr6-83898080-83898581 | 4 | 0.958302378078604 |
| chr19-39816287-39816788 | 8 | 0.887735302542394 |
| chr15-38112942-38113443 | 8 | 0.869385797264427 |
| chr17-79091602-79092103 | 3 | 0.872750840599065 |
| chr18-59148309-59148810 | 8 | 0.90860760729925 |
| chr8-14604167-14604668 | 3 | 0.996883962010668 |
| chrX-15411000-15411501 | 3 | 0.981800127981603 |
| chr5-35437891-35438392 | 8 | 0.90502250134749 |
| chr2-61768807-61769308 | 8 | 0.989361097211749 |
| chr11-57239480-57239981 | 8 | 0.863778877858418 |
| chr9-11991460-11991961 | 8 | 0.994640979724106 |
| chr18-71131021-71131522 | 5 | 0.984882587888028 |
| chr11-7504208-7504709 | 1 | 0.864559176197078 |
| chr7-80302732-80303233 | 6 | 0.993064316573543 |
| chr8-3848369-3848870 | 8 | 0.957075272635338 |
| chr4-136317405-136317906 | 6 | 0.986615111581569 |
| chr15-32899892-32900393 | 2 | 0.923253293645064 |
| chr16-42002778-42003279 | 8 | 0.993299373933394 |
| chr4-132948322-132948823 | 8 | 0.939833763691849 |
| chrX-20045976-20046477 | 8 | 0.988799670305722 |
| chr15-675619-676120 | 3 | 0.996051292009584 |
| chr18-14581747-14582248 | 3 | 0.996896544898946 |
| chrX-36736688-36737189 | 6 | 0.991206925481043 |
| chr5-144449945-144450446 | 8 | 0.830719472398973 |
| chr4-135174629-135175130 | 8 | 0.994206541544251 |
| chr13-92367859-92368360 | 5 | 0.867334147299895 |
| chr1-55303242-55303743 | 3 | 0.994844228670908 |

| | | |
|---|---|---|
| chr9-97167788-97168289 | 6 | 0.990542210029202 |
| chr3-79588919-79589420 | 6 | 0.985818638684277 |
| chr1-152105903-152106404 | 5 | 0.988755500501293 |
| chr6-126224841-126225342 | 7 | 0.914488478125437 |
| chr4-170071369-170071870 | 7 | 0.926746317998386 |
| chr6-97300869-97301370 | 2 | 0.881815100817188 |
| chr11-60848013-60848514 | 8 | 0.90613073419546 |
| chr15-95785963-95786464 | 8 | 0.887851130359473 |
| chr3-131649225-131649726 | 2 | 0.966128808080628 |
| chr16-45405676-45406177 | 8 | 0.995230218403634 |
| chrX-59650254-59650755 | 5 | 0.96395252464132 |
| chr13-103985557-103986058 | 8 | 0.988926787487936 |
| chr1-166680503-166681004 | 1 | 0.938959377514884 |
| chr3-91341496-91341997 | 8 | 0.993493681128311 |
| chr16-35541284-35541785 | 3 | 0.99593553545041 |
| chr2-146952779-146953280 | 6 | 0.993086183182878 |
| chr19-47582627-47583128 | 2 | 0.915217096627968 |
| chr14-62945026-62945527 | 8 | 0.929494012087286 |
| chr7-52324394-52324895 | 1 | 0.836115169946187 |
| chr13-64187284-64187785 | 5 | 0.957734947265857 |
| chr14-87049335-87049836 | 8 | 0.993493681128311 |
| chr9-80568749-80569250 | 1 | 0.880918549493941 |
| chr14-80905948-80906449 | 2 | 0.892562491654669 |
| chr8-23781007-23781508 | 2 | 0.915205943386364 |
| chr13-55884521-55885022 | 8 | 0.928147234640718 |
| chr2-213545322-213545823 | 3 | 0.980444993821869 |
| chr19-23043882-23044383 | 8 | 0.995345684743764 |
| chr17-38792645-38793146 | 3 | 0.993715411117507 |
| chr7-28311599-28312100 | 5 | 0.842505720430365 |
| chr7-1434363-1434864 | 2 | 0.918006136202112 |
| chr15-44316301-44316802 | 1 | 0.959594905660848 |
| chr1-4369784-4370285 | 3 | 0.996596784965579 |
| chr9-45068751-45069252 | 3 | 0.99912712826445 |
| chr11-77279488-77279989 | 2 | 0.830929826302681 |
| chr9-68036973-68037474 | 3 | 0.99593553545041 |
| chr2-169631888-169632389 | 8 | 0.938706333770881 |
| chr12-17065928-17066429 | 8 | 0.980341292323927 |
| chr4-60133155-60133656 | 2 | 0.965101790741002 |
| chr8-32446726-32447227 | 3 | 0.995368170209246 |
| chr14-24704944-24705445 | 3 | 0.984346786051193 |
| chr9-108192912-108193413 | 5 | 0.927420274530861 |
| chrX-100424612-100425113 | 3 | 0.99593553545041 |
| chr17-77473148-77473649 | 3 | 0.945121425922879 |

| | | |
|---|---|---|
| chr16-71009410-71009911 | 7 | 0.85568921198038 |
| chr18-30157877-30158378 | 8 | 0.995238981989093 |
| chr9-88705416-88705917 | 2 | 0.919307378788629 |
| chr3-113661447-113661948 | 8 | 0.988713291673294 |
| chr3-54137768-54138269 | 8 | 0.881920912080842 |
| chr19-50966903-50967404 | 8 | 0.880324973964243 |
| chr19-31027195-31027696 | 6 | 0.988721271334229 |
| chr3-127599119-127599620 | 8 | 0.938240092605977 |
| chr19-899379-899880 | 3 | 0.990434689097275 |
| chr1-9010544-9011045 | 6 | 0.993157401275851 |
| chr4-88234576-88235077 | 5 | 0.894338801106534 |
| chr15-101617896-101618397 | 3 | 0.895785849025426 |
| chr20-46587050-46587551 | 8 | 0.963509143421597 |
| chr5-23930592-23931093 | 8 | 0.962512080605226 |
| chr4-52781807-52782308 | 8 | 0.864952264449762 |
| chrX-30747063-30747564 | 6 | 0.958153698862741 |
| chr7-111488595-111489096 | 8 | 0.965533728580349 |
| chr1-198579559-198580060 | 8 | 0.908245523933801 |
| chr12-40731568-40732069 | 8 | 0.915356055698493 |
| chr11-68114985-68115486 | 1 | 0.882074477288636 |
| chr2-207275145-207275646 | 5 | 0.950930351749404 |
| chr6-8533489-8533990 | 5 | 0.836367435600904 |
| chr1-59068769-59069270 | 6 | 0.959410011440007 |
| chr17-36180654-36181155 | 1 | 0.86549865196178 |
| chr15-42658063-42658564 | 5 | 0.983919978840557 |
| chr2-84777858-84778359 | 4 | 0.84844112331181 |
| chr4-109056050-109056551 | 5 | 0.983786863996452 |
| chr7-47399176-47399677 | 8 | 0.964268947243803 |
| chr14-54677622-54678123 | 6 | 0.938476917485371 |
| chr16-28454858-28455359 | 6 | 0.990363115606696 |
| chr1-153056295-153056796 | 8 | 0.971331025397383 |
| chr15-7672959-7673460 | 4 | 0.816009812671477 |
| chr7-28253842-28254343 | 5 | 0.941558833346536 |
| chr4-15915212-15915713 | 3 | 0.926347697906305 |
| chr4-96126273-96126774 | 3 | 0.998086730068755 |
| chr2-140802930-140803431 | 6 | 0.865750144405932 |
| chr2-146153046-146153547 | 3 | 0.921417758649512 |
| chr20-36921514-36922015 | 8 | 0.983997910756415 |
| chr7-89793479-89793980 | 2 | 0.935013190056861 |
| chr19-39472325-39472826 | 4 | 0.876355888124484 |
| chr15-1981117-1981618 | 3 | 0.998076505355918 |
| chr10-23121469-23121970 | 5 | 0.91977528723226 |
| chr13-66439041-66439542 | 3 | 0.851159006665109 |

| | | |
|---|---|---|
| chr18-35293270-35293771 | 8 | 0.985063260118654 |
| chr13-94540870-94541371 | 8 | 0.892749779784092 |
| chr1-28138002-28138503 | 3 | 0.994603656352347 |
| chr6-33934136-33934637 | 2 | 0.952634709522793 |
| chr17-60903740-60904241 | 5 | 0.816658019355541 |
| chr18-55676698-55677199 | 4 | 0.830713055162362 |
| chr6-71648523-71649024 | 4 | 0.835951870421503 |
| chr9-99690544-99691045 | 8 | 0.86269724037106 |
| chr10-12059928-12060429 | 3 | 0.988726658325252 |
| chr18-81938456-81938957 | 3 | 0.99161577752418 |
| chr15-91446196-91446697 | 8 | 0.876495164060099 |
| chr13-48265712-48266213 | 7 | 0.853241438173754 |
| chr2-212998294-212998795 | 6 | 0.98953469711311 |
| chr18-68422266-68422767 | 5 | 0.984813919135148 |
| chr19-27781978-27782479 | 8 | 0.963999843701842 |
| chr1-251562540-251563041 | 3 | 0.992910227085675 |
| chrX-15960958-15961459 | 3 | 0.99593553545041 |
| chr7-93325624-93326125 | 7 | 0.838862756741464 |
| chr7-19090619-19091120 | 4 | 0.875586486093452 |
| chr7-10761408-10761909 | 3 | 0.99593553545041 |
| chr5-159089897-159090398 | 8 | 0.957084359574894 |
| chr15-47160181-47160682 | 8 | 0.861942673330693 |
| chr6-48216601-48217102 | 3 | 0.996870918214245 |
| chr14-16172224-16172725 | 4 | 0.933491479860165 |
| chr11-43383740-43384241 | 2 | 0.925572764224337 |
| chr11-50085646-50086147 | 3 | 0.99593553545041 |
| chr7-92722916-92723417 | 1 | 0.831192391141609 |
| chr12-39319385-39319886 | 5 | 0.991938957117927 |
| chr14-75516630-75517131 | 5 | 0.976140153562509 |
| chr8-109977871-109978372 | 8 | 0.935278123843685 |
| chr1-122446725-122447226 | 5 | 0.927061791803708 |
| chr17-1332908-1333409 | 2 | 0.940173409087317 |
| chr13-66285613-66286114 | 4 | 0.959249502314113 |
| chr8-57792859-57793360 | 4 | 0.861760496264875 |
| chr13-95840750-95841251 | 7 | 0.852362519221258 |
| chr2-33352381-33352882 | 8 | 0.846054578291022 |
| chr2-74401608-74402109 | 4 | 0.968528657130175 |
| chr18-28867042-28867543 | 3 | 0.99593553545041 |
| chr10-61725230-61725731 | 2 | 0.854611826338041 |
| chr9-21602519-21603020 | 3 | 0.994476903714136 |
| chr6-96325893-96326394 | 8 | 0.993493681128311 |
| chr8-118764293-118764794 | 5 | 0.985235094574135 |
| chr14-88393516-88394017 | 3 | 0.996901949661083 |

| | | |
|---|---|---|
| chr16-20847454-20847955 | 4 | 0.952106368269797 |
| chr2-53559218-53559719 | 8 | 0.901612847917611 |
| chr20-33854784-33855285 | 6 | 0.971575410464868 |
| chr11-73879091-73879592 | 6 | 0.990335621167109 |
| chr2-186383476-186383977 | 6 | 0.991206925481043 |
| chr7-46673365-46673866 | 8 | 0.978160734887496 |
| chr12-40603680-40604181 | 8 | 0.831894298122677 |
| chr9-22206727-22207228 | 8 | 0.994618809860366 |
| chr14-83149574-83150075 | 4 | 0.922043774748727 |
| chr5-9249553-9250054 | 8 | 0.881145051801027 |
| chr10-39315813-39316314 | 8 | 0.952358877243019 |
| chr1-123291177-123291678 | 3 | 0.99593553545041 |
| chr6-49966795-49967296 | 8 | 0.98917440956943 |
| chr15-60188458-60188959 | 3 | 0.99003941134571 |
| chr4-83840532-83841033 | 6 | 0.970557779057439 |
| chr14-58756078-58756579 | 5 | 0.988454963965859 |
| chr7-86997196-86997697 | 8 | 0.995247170519694 |
| chr3-45754414-45754915 | 8 | 0.955978943158956 |
| chr3-101084891-101085392 | 8 | 0.891238198535566 |
| chr8-63261162-63261663 | 8 | 0.958022144371218 |
| chr2-147346320-147346821 | 8 | 0.988215461403242 |
| chr3-160176636-160177137 | 8 | 0.894028505713382 |
| chr5-146131924-146132425 | 8 | 0.972425621406605 |
| chr18-20692565-20693066 | 3 | 0.996874130427153 |
| chr14-37008920-37009421 | 3 | 0.934993406012922 |
| chr7-123216600-123217101 | 5 | 0.829993743078172 |
| chr18-32320037-32320538 | 3 | 0.996515737160711 |
| chr1-56971049-56971550 | 8 | 0.99451121777372 |
| chr7-45192575-45193076 | 8 | 0.980890720718562 |
| chr6-99294403-99294904 | 8 | 0.967093435764569 |
| chr7-124626091-124626592 | 8 | 0.903965219526245 |
| chr4-40037408-40037909 | 3 | 0.99269497529728 |
| chr8-79392213-79392714 | 3 | 0.995734316199332 |
| chr7-121184620-121185121 | 3 | 0.99593553545041 |
| chr5-3525511-3526012 | 6 | 0.872618809354774 |
| chr8-75573518-75574019 | 8 | 0.896692167085123 |
| chr7-105501116-105501617 | 1 | 0.815419549189396 |
| chr6-129917519-129918020 | 8 | 0.883827748178999 |
| chr6-52955224-52955725 | 7 | 0.831728878866589 |
| chr2-224304044-224304545 | 5 | 0.916772061986187 |
| chr17-33231247-33231748 | 7 | 0.850791842733441 |
| chr4-30428278-30428779 | 8 | 0.885421043553819 |
| chr15-25545829-25546330 | 5 | 0.983919978840557 |

| | | |
|---|---|---|
| chr2-34702342-34702843 | 3 | 0.992109646768372 |
| chr5-134211835-134212336 | 6 | 0.970959273852389 |
| chr2-168550545-168551046 | 2 | 0.943004906792591 |
| chr8-45159927-45160428 | 4 | 0.969181020188046 |
| chr3-78103988-78104489 | 3 | 0.996722204288609 |
| chr9-105590838-105591339 | 5 | 0.987228113606421 |
| chr13-25333829-25334330 | 3 | 0.99593553545041 |
| chr1-184200222-184200723 | 3 | 0.98141071186734 |
| chr14-98012234-98012735 | 8 | 0.94497942743378 |
| chr8-29789038-29789539 | 8 | 0.993873415363321 |
| chr19-29807539-29808040 | 2 | 0.946095649054684 |
| chr6-23123386-23123887 | 7 | 0.927610281932023 |
| chr18-50840181-50840682 | 8 | 0.957641813016902 |
| chr2-121868555-121869056 | 8 | 0.9114684670228 |
| chr11-37738361-37738862 | 5 | 0.983919978840557 |
| chr1-15339343-15339844 | 4 | 0.962287220922307 |
| chr10-9564439-9564940 | 3 | 0.993887886894354 |
| chrX-126771739-126772240 | 5 | 0.983919978840557 |
| chr8-119298330-119298831 | 8 | 0.860740697685149 |
| chr1-209854246-209854747 | 8 | 0.878076889154796 |
| chr15-81449623-81450124 | 8 | 0.97612576551457 |
| chr2-50564870-50565371 | 6 | 0.991206925481043 |
| chr7-82789244-82789745 | 8 | 0.995069598027495 |
| chr6-139705140-139705641 | 1 | 0.94028585642955 |
| chr9-98281957-98282458 | 6 | 0.984069925030642 |
| chr2-189375735-189376236 | 8 | 0.946394626765062 |
| chr7-27912683-27913184 | 7 | 0.908467307407939 |
| chr13-91916722-91917223 | 8 | 0.845373279442294 |
| chr11-65756895-65757396 | 8 | 0.96066904575463 |
| chr10-33537635-33538136 | 8 | 0.937881885131993 |
| chr4-69672734-69673235 | 3 | 0.964306425602857 |
| chr6-83917523-83918024 | 3 | 0.970911946511071 |
| chr14-65244745-65245246 | 7 | 0.823008835026805 |
| chr13-65037475-65037976 | 8 | 0.89283615666234 |
| chr2-188276006-188276507 | 5 | 0.967419495886522 |
| chr7-20296796-20297297 | 8 | 0.889776369621508 |
| chr13-79262060-79262561 | 1 | 0.927453796952525 |
| chr18-32947132-32947633 | 5 | 0.906509023632724 |
| chr16-70608195-70608696 | 3 | 0.99593553545041 |
| chr10-47855350-47855851 | 3 | 0.996389180740731 |
| chr18-34393157-34393658 | 8 | 0.993493681128311 |
| chr6-125292498-125292999 | 8 | 0.871697621875993 |
| chr15-31954328-31954829 | 4 | 0.820235092480221 |

| | | |
|---|---|---|
| chr16-68945712-68946213 | 1 | 0.869875610684463 |
| chr19-46488515-46489016 | 1 | 0.885183413176144 |
| chr5-162584112-162584613 | 4 | 0.952718594547332 |
| chr5-59017775-59018276 | 1 | 0.953731358703435 |
| chr20-44347174-44347675 | 2 | 0.956845591567764 |
| chr14-22127959-22128460 | 8 | 0.852979591083988 |
| chr6-8572860-8573361 | 6 | 0.997225157535748 |
| chr12-38623673-38624174 | 8 | 0.908304112877581 |
| chr11-66681693-66682194 | 8 | 0.927412953537393 |
| chr18-83388050-83388551 | 6 | 0.991206925481043 |
| chr5-79652192-79652693 | 8 | 0.908867561370643 |
| chr10-48029999-48030500 | 8 | 0.994462288922286 |
| chr3-103233388-103233889 | 1 | 0.814680020099831 |
| chr11-66407028-66407529 | 2 | 0.925131083808533 |
| chrX-130930649-130931150 | 8 | 0.89095614999465 |
| chr10-52228780-52229281 | 8 | 0.98218372809308 |
| chr5-61567229-61567730 | 1 | 0.943781127173182 |
| chr13-99714978-99715479 | 3 | 0.99593553545041 |
| chr1-143539504-143540005 | 1 | 0.935907666487705 |
| chr17-33472748-33473249 | 8 | 0.958055032923879 |
| chr2-46596923-46597424 | 5 | 0.870286560374419 |
| chr15-98740434-98740935 | 5 | 0.97971569066541 |
| chr5-41870218-41870719 | 3 | 0.987712277955826 |
| chr20-13295158-13295659 | 5 | 0.92779965834632 |
| chr1-225901907-225902408 | 5 | 0.9041255567423 |
| chr13-41058017-41058518 | 8 | 0.993493681128311 |
| chr13-1526604-1527105 | 6 | 0.975217831161368 |
| chr1-222753919-222754420 | 8 | 0.940143403247396 |
| chr10-52314839-52315340 | 7 | 0.829803417921844 |
| chr1-69253871-69254372 | 3 | 0.996799423729359 |
| chr4-151984489-151984990 | 8 | 0.880977286923171 |
| chr5-133548026-133548527 | 8 | 0.988784623930826 |
| chr16-60249-60750 | 3 | 0.996055877284061 |
| chr8-69798108-69798609 | 8 | 0.914165814194408 |
| chr17-22657299-22657800 | 8 | 0.925626386907593 |
| chr3-29489774-29490275 | 8 | 0.93624968934816 |
| chr9-56947757-56948258 | 8 | 0.927509688554651 |
| chr3-69364230-69364731 | 1 | 0.892987728342709 |
| chr9-48331345-48331846 | 8 | 0.931804546416988 |
| chr9-26477001-26477502 | 3 | 0.995367172002261 |
| chr7-89556634-89557135 | 1 | 0.929212195415513 |
| chr18-76549834-76550335 | 6 | 0.897378562182009 |
| chr9-83154757-83155258 | 3 | 0.935368816295751 |

| | | |
|---|---|---|
| chr9-83686835-83687336 | 8 | 0.822126824799558 |
| chr12-32504681-32505182 | 5 | 0.986130651420782 |
| chr3-141491605-141492106 | 8 | 0.977311018829106 |
| chr2-4727117-4727618 | 3 | 0.966994645306599 |
| chr13-100313968-100314469 | 2 | 0.890541629973072 |
| chr1-31407150-31407651 | 8 | 0.896545519749861 |
| chr1-23576296-23576797 | 8 | 0.874860878180478 |
| chr7-128363902-128364403 | 3 | 0.992646994408225 |
| chr16-17724973-17725474 | 3 | 0.985047516757873 |
| chr5-32073602-32074103 | 3 | 0.983731627868125 |
| chr11-44971860-44972361 | 1 | 0.905418659908915 |
| chr5-130922645-130923146 | 8 | 0.962886466301744 |
| chr10-25031126-25031627 | 2 | 0.85990694931584 |
| chr15-22201072-22201573 | 7 | 0.903203339631572 |
| chr14-43232693-43233194 | 5 | 0.960806377710653 |
| chr9-26729911-26730412 | 8 | 0.930769945454201 |
| chr1-107429771-107430272 | 3 | 0.984676541540828 |
| chr13-31651118-31651619 | 8 | 0.85079566586319 |
| chr4-37686628-37687129 | 3 | 0.853209640336967 |
| chr5-158483801-158484302 | 6 | 0.984074353524426 |
| chr6-27343747-27344248 | 3 | 0.97094991794888 |
| chr3-123649459-123649960 | 8 | 0.980687765666815 |
| chr10-91073221-91073722 | 4 | 0.86948112487998 |
| chr5-147209528-147210029 | 7 | 0.93113480119645 |
| chr6-132687555-132688056 | 3 | 0.99593553545041 |
| chr3-99121844-99122345 | 3 | 0.977742615627349 |
| chr7-104501709-104502210 | 3 | 0.992237447755906 |
| chr14-74937094-74937595 | 8 | 0.871494447752188 |
| chr9-103095898-103096399 | 1 | 0.897048923928524 |
| chr3-148301671-148302172 | 8 | 0.993493681128311 |
| chr1-234622243-234622744 | 8 | 0.837680287348864 |
| chr6-40677324-40677825 | 7 | 0.828422960083916 |
| chr7-118560078-118560579 | 3 | 0.985437472931216 |
| chr17-31260763-31261264 | 3 | 0.963305582981169 |
| chr8-106093304-106093805 | 8 | 0.8968345347745 |
| chr11-4050647-4051148 | 8 | 0.919793226716836 |
| chr2-244988151-244988652 | 3 | 0.99593553545041 |
| chr20-25143218-25143719 | 8 | 0.978035660922918 |
| chr2-38540725-38541226 | 1 | 0.920815127320469 |
| chr7-67408216-67408717 | 8 | 0.819733697419761 |
| chr13-44015412-44015913 | 8 | 0.992583971805806 |
| chr13-45906467-45906968 | 4 | 0.834935233902988 |
| chr17-65042830-65043331 | 1 | 0.917921462655034 |

| | | |
|---|---|---|
| chr1-161348644-161349145 | 8 | 0.901758868280262 |
| chr15-6772066-6772567 | 6 | 0.992143913531083 |
| chr3-108031120-108031621 | 3 | 0.985931999374274 |
| chrX-63954086-63954587 | 3 | 0.99681624195534 |
| chr4-154679530-154680031 | 3 | 0.990003381704189 |
| chr14-69759983-69760484 | 8 | 0.983941901122751 |
| chr3-54680235-54680736 | 8 | 0.993493681128311 |
| chr7-90774046-90774547 | 8 | 0.824609757308435 |
| chr9-2187446-2187947 | 3 | 0.997213450946204 |
| chr2-110363954-110364455 | 8 | 0.903101332649866 |
| chr8-21843654-21844155 | 5 | 0.987713580657996 |
| chr1-15169351-15169852 | 4 | 0.819639326195535 |
| chr6-27081822-27082323 | 2 | 0.951990612461287 |
| chr20-4473846-4474347 | 8 | 0.970775624498632 |
| chr12-8471665-8472166 | 3 | 0.995642327356344 |
| chr3-11544750-11545251 | 8 | 0.993875942382002 |
| chr15-32217661-32218162 | 3 | 0.975655097722821 |
| chr8-12952319-12952820 | 3 | 0.930425941543969 |
| chr5-130416042-130416543 | 8 | 0.823340199303476 |
| chr2-22341347-22341848 | 5 | 0.988005462195933 |
| chr1-213264595-213265096 | 6 | 0.991889664478073 |
| chr1-44013014-44013515 | 8 | 0.961904442544512 |
| chr18-30595978-30596479 | 8 | 0.948687384799746 |
| chr10-35884931-35885432 | 5 | 0.931579111740944 |
| chr10-11603137-11603638 | 8 | 0.98651051504533 |
| chr13-89752977-89753478 | 8 | 0.918001638201071 |
| chr17-8133094-8133595 | 8 | 0.844750514362831 |
| chr16-76910906-76911407 | 7 | 0.964166041149797 |
| chr13-69251967-69252468 | 8 | 0.992059190198966 |
| chr9-10689762-10690263 | 8 | 0.974978843858713 |
| chr16-84013763-84014264 | 3 | 0.995813209262692 |
| chr1-163966709-163967210 | 8 | 0.864644124770508 |
| chr5-70392161-70392662 | 8 | 0.990428009290968 |
| chr13-82134735-82135236 | 5 | 0.977427004180296 |
| chr1-135119877-135120378 | 6 | 0.992981776339421 |
| chr19-27652768-27653269 | 8 | 0.908049875054119 |
| chr4-181216689-181217190 | 6 | 0.983756700559874 |
| chr13-104265508-104266009 | 3 | 0.995067032453406 |
| chr7-105487379-105487880 | 8 | 0.870743490199964 |
| chr2-219400659-219401160 | 7 | 0.818783174062586 |
| chr7-43008718-43009219 | 8 | 0.87001004767821 |
| chr7-115386566-115387067 | 3 | 0.992488174469376 |
| chr14-27666565-27667066 | 3 | 0.99593553545041 |

| | | |
|---|---|---|
| chr10-820808-821309 | 5 | 0.874798443785817 |
| chr8-58744340-58744841 | 8 | 0.986541818548428 |
| chr6-9725752-9726253 | 4 | 0.928585283286778 |
| chr18-25909431-25909932 | 2 | 0.85340770920094 |
| chr16-68103005-68103506 | 5 | 0.927470493800926 |
| chr7-55681450-55681951 | 8 | 0.993493681128311 |
| chr2-138434988-138435489 | 3 | 0.99593553545041 |
| chr9-12478404-12478905 | 8 | 0.899305786124885 |
| chr7-19464268-19464769 | 2 | 0.889636401248704 |
| chr14-66456132-66456633 | 1 | 0.905845767156541 |
| chr18-45839848-45840349 | 8 | 0.993493681128311 |
| chr7-122020847-122021348 | 6 | 0.993133072031873 |
| chr4-39982312-39982813 | 4 | 0.841592600804645 |
| chr6-29623892-29624393 | 6 | 0.980510383418924 |
| chr1-85351061-85351562 | 1 | 0.934494864156645 |
| chr3-114230066-114230567 | 8 | 0.992641570275353 |
| chr6-41507307-41507808 | 5 | 0.934407197567613 |
| chr14-84064986-84065487 | 8 | 0.968950466930109 |
| chr4-173727070-173727571 | 3 | 0.996852060459077 |
| chr16-59374903-59375404 | 6 | 0.838490406591797 |
| chr8-44111646-44112147 | 8 | 0.964508194885904 |
| chr5-58574144-58574645 | 8 | 0.877368153522031 |
| chr15-44989837-44990338 | 2 | 0.873858065573185 |
| chr13-65142647-65143148 | 8 | 0.906620302826238 |
| chr9-16291247-16291748 | 8 | 0.97734262382992 |
| chrX-139730423-139730924 | 6 | 0.971930618914801 |
| chr3-1481042-1481543 | 4 | 0.823197580704773 |
| chr14-48094654-48095155 | 6 | 0.944272104424662 |
| chr20-3461710-3462211 | 3 | 0.987735613562801 |
| chr3-111513365-111513866 | 8 | 0.974571491165352 |
| chr7-133763581-133764082 | 8 | 0.985330823548184 |
| chr13-205006-205507 | 3 | 0.99389772888381 |
| chr10-35961281-35961782 | 4 | 0.935247151507784 |
| chr7-100124732-100125233 | 8 | 0.98208346430115 |
| chr2-234002605-234003106 | 3 | 0.895703429635151 |
| chr1-33419589-33420090 | 8 | 0.83254484721097 |
| chr5-152925513-152926014 | 8 | 0.985912376840763 |
| chr1-124709974-124710475 | 5 | 0.978355477589891 |
| chr6-106047717-106048218 | 1 | 0.92123587261018 |
| chr1-181491060-181491561 | 5 | 0.862553698383308 |
| chr3-30509528-30510029 | 3 | 0.995240782087349 |
| chr15-99485087-99485588 | 8 | 0.932704979195708 |
| chr5-28742635-28743136 | 7 | 0.959621597303679 |

| | | |
|---|---|---|
| chr3-142039217-142039718 | 5 | 0.955495748689693 |
| chr10-50011491-50011992 | 8 | 0.993493681128311 |
| chr14-83718591-83719092 | 3 | 0.908990572526733 |
| chr8-80246577-80247078 | 8 | 0.98914115413711 |
| chr2-234999399-234999900 | 5 | 0.965304647353189 |
| chr19-42625295-42625796 | 5 | 0.983032979668418 |
| chr5-165367591-165368092 | 8 | 0.935811817490458 |
| chr10-95428463-95428964 | 5 | 0.863224957093669 |
| chr1-214320443-214320944 | 3 | 0.99539920495285 |
| chr1-107209142-107209643 | 6 | 0.974463414693927 |
| chr1-52286604-52287105 | 5 | 0.967869148045426 |
| chr1-178439711-178440212 | 6 | 0.991132373928974 |
| chr15-12643391-12643892 | 5 | 0.96413185112567 |
| chr20-9965104-9965605 | 3 | 0.906629245170629 |
| chr16-2979869-2980370 | 8 | 0.993493681128311 |
| chr1-11863199-11863700 | 3 | 0.993850182644165 |
| chr9-37462582-37463083 | 8 | 0.924342530466942 |
| chr5-35064736-35065237 | 6 | 0.888793931109723 |
| chr19-40501516-40502017 | 3 | 0.99593553545041 |
| chr6-125046884-125047385 | 4 | 0.928765078922322 |
| chr6-138407963-138408464 | 7 | 0.944749131471199 |
| chr9-65502428-65502929 | 2 | 0.896317072223991 |
| chr4-110027897-110028398 | 8 | 0.986448504848196 |
| chr2-20605631-20606132 | 3 | 0.881452213644868 |
| chr10-8343284-8343785 | 3 | 0.996530686878949 |
| chr3-72895057-72895558 | 6 | 0.991206925481043 |
| chr4-26670802-26671303 | 3 | 0.99593553545041 |
| chr14-7782180-7782681 | 7 | 0.937156697086769 |
| chr9-42549962-42550463 | 8 | 0.906783533308076 |
| chr16-80246185-80246686 | 4 | 0.832926434909546 |
| chr16-46141064-46141565 | 8 | 0.851107850183703 |
| chr7-55483893-55484394 | 7 | 0.930493636358998 |
| chr3-159017747-159018248 | 5 | 0.988603059858181 |
| chr4-176159086-176159587 | 4 | 0.94819106416891 |
| chr18-21920388-21920889 | 3 | 0.99593553545041 |
| chr3-116369916-116370417 | 3 | 0.979413369045181 |
| chr16-66665655-66666156 | 8 | 0.974576057206222 |
| chr1-107357033-107357534 | 8 | 0.988260174889352 |
| chr16-81818198-81818699 | 6 | 0.979758612496825 |
| chr20-20184965-20185466 | 8 | 0.866743951068785 |
| chr6-64447066-64447567 | 8 | 0.834530056928986 |
| chr2-189318283-189318784 | 8 | 0.958161513033618 |
| chr6-58577747-58578248 | 3 | 0.997674275132125 |

| | | |
|---|---|---|
| chr1-19503043-19503544 | 8 | 0.995202849486469 |
| chr17-1014794-1015295 | 5 | 0.986359601934013 |
| chr1-63623201-63623702 | 6 | 0.939879212000225 |
| chr1-126851720-126852221 | 5 | 0.876568474851223 |
| chr3-155034954-155035455 | 8 | 0.954410867797046 |
| chr1-124038808-124039309 | 5 | 0.990218007312326 |
| chr4-178363269-178363770 | 5 | 0.985847674938417 |
| chr20-19947954-19948455 | 6 | 0.99144896167999 |
| chr16-61095513-61096014 | 8 | 0.949717786972461 |
| chrX-17233847-17234348 | 6 | 0.984575027185221 |
| chr19-45432323-45432824 | 3 | 0.8187385567852 |
| chr5-124639111-124639612 | 8 | 0.832525105856515 |
| chr8-121790075-121790576 | 6 | 0.949445867986137 |
| chr2-217756093-217756594 | 6 | 0.992369219361493 |
| chr20-19943232-19943733 | 3 | 0.99593553545041 |
| chr9-106644126-106644627 | 8 | 0.882243265996872 |
| chr13-44935333-44935834 | 7 | 0.897746420452755 |
| chr9-23271821-23272322 | 5 | 0.986830401677125 |
| chr6-117941710-117942211 | 4 | 0.961608126400428 |
| chr7-120300916-120301417 | 2 | 0.94206768207741 |
| chr11-28852509-28853010 | 8 | 0.902605263820729 |
| chr15-34542595-34543096 | 6 | 0.991206925481043 |
| chr2-186126272-186126773 | 3 | 0.994880747680331 |
| chr7-63854084-63854585 | 8 | 0.942624790418819 |
| chr16-41093424-41093925 | 6 | 0.977177325594267 |
| chr6-123313093-123313594 | 1 | 0.818802740391022 |
| chr11-47219004-47219505 | 6 | 0.993228211943007 |
| chr5-20769751-20770252 | 6 | 0.987140524145356 |
| chr20-7040870-7041371 | 5 | 0.988578315053284 |
| chr14-43703176-43703677 | 8 | 0.967766920496858 |
| chr17-59771665-59772166 | 8 | 0.870964979872237 |
| chr13-94370044-94370545 | 2 | 0.959525405424327 |
| chr14-7943580-7944081 | 4 | 0.966480586185688 |
| chr14-74540027-74540528 | 8 | 0.973894786655065 |
| chr7-92606728-92607229 | 3 | 0.996309270057404 |
| chr7-112270806-112271307 | 8 | 0.894695386043686 |
| chr4-82422889-82423390 | 4 | 0.967478416126122 |
| chr6-127694770-127695271 | 6 | 0.976786314748949 |
| chr16-20743805-20744306 | 8 | 0.916063909149208 |
| chr9-37423673-37424174 | 8 | 0.92703673735892 |
| chr3-87707764-87708265 | 3 | 0.886143753862214 |
| chr18-26205160-26205661 | 7 | 0.901217106399162 |
| chr2-25573578-25574079 | 8 | 0.992615820450424 |

| | | |
|---|---|---|
| chr3-124234524-124235025 | 5 | 0.927340736954421 |
| chr6-104971037-104971538 | 8 | 0.838939292814569 |
| chr16-32283347-32283848 | 6 | 0.992831884410744 |
| chr2-11824646-11825147 | 2 | 0.856886696239541 |
| chr12-12489693-12490194 | 8 | 0.976171826279177 |
| chr8-20758204-20758705 | 6 | 0.992608833573697 |
| chr3-155553466-155553967 | 8 | 0.934990493016784 |
| chr5-142104937-142105438 | 4 | 0.833020204398999 |
| chr5-144453149-144453650 | 8 | 0.96566758739827 |
| chr19-4952575-4953076 | 3 | 0.994224344361744 |
| chr6-138467668-138468169 | 8 | 0.926568877064595 |
| chr10-41815904-41816405 | 6 | 0.92971056316036 |
| chr6-54359027-54359528 | 3 | 0.996873202415663 |
| chr15-29402617-29403118 | 8 | 0.913592291889823 |
| chr1-145286998-145287499 | 8 | 0.887096021948295 |
| chr2-221328528-221329029 | 7 | 0.887378150224281 |
| chr9-108761296-108761797 | 3 | 0.996049327272995 |
| chr6-99325498-99325999 | 8 | 0.944908528113066 |
| chr6-76724288-76724789 | 8 | 0.845432936614411 |
| chr1-225400623-225401124 | 2 | 0.832021191327104 |
| chr2-153300418-153300919 | 8 | 0.853139248494638 |
| chr1-177172694-177173195 | 6 | 0.986887501800423 |
| chr6-83299872-83300373 | 3 | 0.876617643760151 |
| chr14-30717413-30717914 | 8 | 0.993806219352565 |
| chr8-85032127-85032628 | 8 | 0.855890858721058 |
| chr3-149187038-149187539 | 6 | 0.977564584242684 |
| chr16-76520035-76520536 | 8 | 0.96891281457872 |
| chr2-183049706-183050207 | 6 | 0.920678787695997 |
| chr6-48354954-48355455 | 6 | 0.952443866725482 |
| chr16-64140256-64140757 | 8 | 0.995245916633714 |
| chr4-61048753-61049254 | 8 | 0.949579039343545 |
| chr14-26218233-26218734 | 6 | 0.840036690134396 |
| chr9-41097233-41097734 | 3 | 0.99593553545041 |
| chr18-37983021-37983522 | 3 | 0.99593553545041 |
| chr2-74900554-74901055 | 8 | 0.966147657326977 |
| chr14-100233247-100233748 | 3 | 0.985377958231181 |
| chr12-12964466-12964967 | 8 | 0.90927278826754 |
| chr5-22939973-22940474 | 5 | 0.965497931020096 |
| chr8-34065334-34065835 | 6 | 0.963534623639078 |
| chr6-58022738-58023239 | 3 | 0.956660219218504 |
| chr1-235146476-235146977 | 8 | 0.831471102491221 |
| chr13-380038-380539 | 3 | 0.98362834699671 |
| chr13-78153025-78153526 | 8 | 0.956826730919193 |

| | | |
|---|---|---|
| chr5-155889778-155890279 | 6 | 0.977348094730305 |
| chr8-88651329-88651830 | 8 | 0.993493681128311 |
| chr6-76237539-76238040 | 3 | 0.996644249616279 |
| chr20-44901498-44901999 | 5 | 0.966908732933029 |
| chr16-25560750-25561251 | 1 | 0.963426429596441 |
| chr1-121925263-121925764 | 8 | 0.891992771858842 |
| chr5-125299278-125299779 | 5 | 0.91544518888676 |
| chr6-36284234-36284735 | 3 | 0.996681576843158 |
| chr2-232119672-232120173 | 4 | 0.872937795911139 |
| chr15-10594980-10595481 | 3 | 0.996648312866657 |
| chr8-111417329-111417830 | 3 | 0.916173561088745 |
| chr3-57944982-57945483 | 8 | 0.973399971053941 |
| chr18-66132461-66132962 | 6 | 0.991206925481043 |
| chr15-10019878-10020379 | 4 | 0.888401863024329 |
| chr13-24308424-24308925 | 1 | 0.9610559278232 |
| chr1-47629163-47629664 | 2 | 0.843940591246914 |
| chr18-14781242-14781743 | 3 | 0.996282107486874 |
| chr7-122500908-122501409 | 8 | 0.969306467728788 |
| chr3-33347521-33348022 | 8 | 0.993493681128311 |
| chr4-181866388-181866889 | 6 | 0.993224058629944 |
| chr2-209233749-209234250 | 8 | 0.993823476629549 |
| chrX-10283218-10283719 | 5 | 0.854660681727789 |
| chr3-31647322-31647823 | 2 | 0.830697264420099 |
| chr2-133632171-133632672 | 5 | 0.977920184384518 |
| chr18-30591499-30592000 | 8 | 0.914862764707165 |
| chr7-32223803-32224304 | 6 | 0.986596811915084 |
| chr11-74493425-74493926 | 8 | 0.922846074346778 |
| chr7-79745037-79745538 | 3 | 0.991575780628372 |
| chr9-41800342-41800843 | 8 | 0.816409867356964 |
| chr1-175856740-175857241 | 8 | 0.906647949959444 |
| chr1-58396860-58397361 | 8 | 0.992631028320378 |
| chr5-88933217-88933718 | 8 | 0.854967329377706 |
| chr4-29936382-29936883 | 3 | 0.99593553545041 |
| chr7-122907248-122907749 | 7 | 0.85693013442193 |
| chr9-19387012-19387513 | 6 | 0.991206925481043 |
| chr17-18154935-18155436 | 8 | 0.996825737481537 |
| chr16-68653179-68653680 | 4 | 0.833955452846883 |
| chr12-29461821-29462322 | 3 | 0.985499284864491 |
| chr14-89601239-89601740 | 6 | 0.991206925481043 |
| chr8-101866102-101866603 | 8 | 0.83409890835582 |
| chr7-556643-557144 | 8 | 0.822075411183749 |
| chr13-78288059-78288560 | 8 | 0.882190976752812 |
| chr8-72392121-72392622 | 7 | 0.829374642283977 |

| | | |
|---|---|---|
| chr4-35410724-35411225 | 3 | 0.996702155846994 |
| chr4-115029744-115030245 | 1 | 0.960048317456435 |
| chr3-28507088-28507589 | 6 | 0.992207014673326 |
| chr8-49943919-49944420 | 8 | 0.993493681128311 |
| chr2-48235467-48235968 | 3 | 0.995876447092654 |
| chr1-256323811-256324312 | 3 | 0.840263413215812 |
| chr9-25072534-25073035 | 2 | 0.957194923851561 |
| chr11-68892272-68892773 | 8 | 0.869865911812525 |
| chr16-79133762-79134263 | 8 | 0.993493681128311 |
| chr17-48923515-48924016 | 4 | 0.886573867340275 |
| chr8-12265647-12266148 | 3 | 0.996181334565585 |
| chr18-31056048-31056549 | 8 | 0.953288233891771 |
| chr16-48784357-48784858 | 3 | 0.99593553545041 |
| chr8-24744207-24744708 | 3 | 0.996822318479957 |
| chr2-112125419-112125920 | 6 | 0.941103649331035 |
| chr8-78197403-78197904 | 5 | 0.934483669571555 |
| chr8-30221802-30222303 | 5 | 0.944011018877705 |
| chr2-40801891-40802392 | 5 | 0.83520234788705 |
| chr3-62975846-62976347 | 3 | 0.996895340283264 |
| chr4-115093772-115094273 | 1 | 0.851849064391955 |
| chr15-5207747-5208248 | 6 | 0.975800192152586 |
| chr8-8184084-8184585 | 6 | 0.852762798448559 |
| chr1-240645326-240645827 | 7 | 0.951205213119957 |
| chr14-73797164-73797665 | 6 | 0.991206925481043 |
| chr4-56919069-56919570 | 6 | 0.968254975653461 |
| chr3-143941968-143942469 | 4 | 0.893896153254488 |
| chr20-1463104-1463605 | 8 | 0.828836144105728 |
| chr19-13696751-13697252 | 8 | 0.818848960306392 |
| chrX-13059027-13059528 | 2 | 0.86557670365238 |
| chr4-23897527-23898028 | 4 | 0.937886686117915 |
| chr4-127958627-127959128 | 8 | 0.994390007873033 |
| chr6-34219995-34220496 | 3 | 0.99593553545041 |
| chr19-13192561-13193062 | 5 | 0.952931750638306 |
| chr16-56063420-56063921 | 1 | 0.920588522438523 |
| chr19-57322245-57322746 | 8 | 0.924880535854984 |
| chrX-63728980-63729481 | 3 | 0.996374611770024 |
| chr3-92560911-92561412 | 6 | 0.926407676123526 |
| chr18-44097936-44098437 | 8 | 0.995233334623945 |
| chr4-138500252-138500753 | 5 | 0.983919978840557 |
| chr13-64468265-64468766 | 6 | 0.982866725406179 |
| chr7-7146311-7146812 | 3 | 0.996385798644127 |
| chr8-99987958-99988459 | 2 | 0.816753680507757 |
| chr16-56251-56752 | 8 | 0.989647879755761 |

| | | |
|---|---|---|
| chr16-7308742-7309243 | 5 | 0.945825356473338 |
| chr9-44973958-44974459 | 2 | 0.936038929336236 |
| chr18-33641342-33641843 | 3 | 0.99593553545041 |
| chr4-95975789-95976290 | 2 | 0.843607776111483 |
| chr4-135701739-135702240 | 2 | 0.950560280729254 |
| chr1-165672398-165672899 | 3 | 0.995661620715337 |
| chr15-3852781-3853282 | 8 | 0.858795088312794 |
| chr2-48223603-48224104 | 2 | 0.913818468797258 |
| chr14-26743181-26743682 | 3 | 0.902940793026182 |
| chr6-26043507-26044008 | 5 | 0.973083015309622 |
| chr11-62014933-62015434 | 4 | 0.944837622380528 |
| chr4-51830028-51830529 | 4 | 0.958295043728667 |
| chr1-111524619-111525120 | 3 | 0.938484420099422 |
| chr1-208048669-208049170 | 6 | 0.992787794897802 |
| chr11-68092693-68093194 | 8 | 0.896217006006771 |
| chr17-75589262-75589763 | 5 | 0.826556071203859 |
| chr18-70715212-70715713 | 8 | 0.968466600217324 |
| chr8-84434075-84434576 | 3 | 0.983910013143506 |
| chr6-108179049-108179550 | 8 | 0.985666004474968 |
| chr2-95037475-95037976 | 5 | 0.983919978840557 |
| chr16-43488086-43488587 | 2 | 0.918063798800379 |
| chr1-253449373-253449874 | 8 | 0.993493681128311 |
| chr2-209347999-209348500 | 8 | 0.926747252813057 |
| chr7-72913020-72913521 | 8 | 0.886890028098798 |
| chr1-258129449-258129950 | 4 | 0.839593514870506 |
| chr18-67960827-67961328 | 2 | 0.942184358752333 |
| chr4-117959908-117960409 | 2 | 0.889004169892699 |
| chr8-25014267-25014768 | 6 | 0.99511231930262 |
| chr4-38585786-38586287 | 2 | 0.820178661850255 |
| chr13-63692180-63692681 | 8 | 0.993493681128311 |
| chr3-91101113-91101614 | 3 | 0.988282400819785 |
| chr2-164147509-164148010 | 6 | 0.991206925481043 |
| chr2-169407019-169407520 | 7 | 0.845244191106716 |
| chr11-66355089-66355590 | 5 | 0.967901412875878 |
| chr20-11351126-11351627 | 8 | 0.988374223276296 |
| chr2-243451100-243451601 | 5 | 0.985518510470041 |
| chr14-1838320-1838821 | 4 | 0.879549397895452 |
| chr3-11613441-11613942 | 8 | 0.964263545964785 |
| chr1-148104005-148104506 | 3 | 0.989445690705285 |
| chr15-29955894-29956395 | 4 | 0.88309315484654 |
| chr2-102591119-102591620 | 8 | 0.946964795942432 |
| chr4-106080646-106081147 | 3 | 0.885651302698149 |
| chr5-56088573-56089074 | 6 | 0.994869494827618 |

| | | |
|---|---|---|
| chr10-48242250-48242751 | 8 | 0.993493681128311 |
| chr10-62136142-62136643 | 4 | 0.947795485520683 |
| chr8-100023708-100024209 | 8 | 0.993493681128311 |
| chr10-97287378-97287879 | 4 | 0.924148897161212 |
| chr11-82984798-82985299 | 8 | 0.988494656628051 |
| chr2-51639922-51640423 | 4 | 0.935505864190015 |
| chr1-202544610-202545111 | 8 | 0.99459262864788 |
| chr11-19894021-19894522 | 8 | 0.827673396576491 |
| chr5-54834833-54835334 | 2 | 0.887325534887449 |
| chr1-135866325-135866826 | 8 | 0.894509106602196 |
| chr4-43570700-43571201 | 5 | 0.835163895740708 |
| chr7-113267048-113267549 | 4 | 0.867725452964756 |
| chr2-225301500-225302001 | 8 | 0.922623152309501 |
| chr6-103936910-103937411 | 6 | 0.992205248098334 |
| chr4-26253564-26254065 | 3 | 0.994468909929851 |
| chr15-70276309-70276810 | 3 | 0.99593553545041 |
| chr10-73416583-73417084 | 4 | 0.966680744687828 |
| chr17-52955284-52955785 | 8 | 0.967771747509374 |
| chr3-157794985-157795486 | 8 | 0.860901681823365 |
| chr10-39236848-39237349 | 8 | 0.923912404500616 |
| chr19-46494658-46495159 | 5 | 0.976480097170728 |
| chr19-34188749-34189250 | 1 | 0.956467290773653 |
| chr17-73462399-73462900 | 8 | 0.819393801605667 |
| chr2-193583844-193584345 | 5 | 0.846792124441491 |
| chr12-45915901-45916402 | 8 | 0.903507914141696 |
| chr2-122698129-122698630 | 4 | 0.961570960752292 |
| chr3-146357045-146357546 | 8 | 0.957319273220291 |
| chr15-12546724-12547225 | 6 | 0.98193938815218 |
| chr17-16709083-16709584 | 7 | 0.825117592808582 |
| chr15-22039472-22039973 | 6 | 0.96684313878213 |
| chr1-102014126-102014627 | 3 | 0.993319381736085 |
| chr9-22702859-22703360 | 5 | 0.839858826102869 |
| chr17-66320763-66321264 | 8 | 0.928072170311075 |
| chr5-103763946-103764447 | 1 | 0.929595336238961 |
| chr10-43544872-43545373 | 8 | 0.988511049412986 |
| chr10-15342420-15342921 | 8 | 0.963423967099695 |
| chr1-105469478-105469979 | 3 | 0.989823385876404 |
| chr3-56484336-56484837 | 6 | 0.841359880683042 |
| chr8-78514690-78515191 | 4 | 0.820275069124973 |
| chr1-56177378-56177879 | 8 | 0.927530791404354 |
| chr5-134568539-134569040 | 2 | 0.822138818231988 |
| chr13-73563528-73564029 | 8 | 0.935207362375462 |
| chr15-67754677-67755178 | 6 | 0.991206925481043 |

| | | |
|---|---|---|
| chr7-10386736-10387237 | 6 | 0.958115641198757 |
| chr5-160743894-160744395 | 3 | 0.885941390337048 |
| chr1-181949460-181949961 | 3 | 0.996893136913015 |
| chr5-103906561-103907062 | 8 | 0.818788064514631 |
| chr18-75262187-75262688 | 8 | 0.897234186817781 |
| chr2-81788501-81789002 | 3 | 0.996013345773066 |
| chr15-5443817-5444318 | 8 | 0.921389028181876 |
| chr15-39668712-39669213 | 8 | 0.983349732466945 |
| chr2-172149410-172149911 | 5 | 0.869586039647173 |
| chr4-26930847-26931348 | 5 | 0.985156777599357 |
| chr2-245822380-245822881 | 5 | 0.87834905529251 |
| chr4-180777845-180778346 | 8 | 0.978690656262127 |
| chr9-84208483-84208984 | 8 | 0.908894232627222 |
| chr9-720685-721186 | 8 | 0.87779519983718 |
| chr11-82418620-82419121 | 7 | 0.875052367779689 |
| chr18-1444546-1445047 | 8 | 0.990631477356904 |
| chr4-61848591-61849092 | 6 | 0.953094126675614 |
| chr7-106956657-106957158 | 8 | 0.846319277409508 |
| chr4-177637794-177638295 | 8 | 0.912936093022699 |
| chr12-7198660-7199161 | 1 | 0.850945867145227 |
| chr6-33923877-33924378 | 8 | 0.986187016068827 |
| chr6-26604530-26605031 | 4 | 0.933501403495869 |
| chr2-39545580-39546081 | 8 | 0.976863613333486 |
| chr7-78252285-78252786 | 3 | 0.970420879814843 |
| chr5-66168395-66168896 | 3 | 0.99601028775109 |
| chr6-72326458-72326959 | 8 | 0.929931802148266 |
| chr3-145046614-145047115 | 3 | 0.988984666274311 |
| chr5-135606512-135607013 | 8 | 0.916216477913902 |
| chr9-96453416-96453917 | 5 | 0.919100526505088 |
| chr2-47245303-47245804 | 3 | 0.99593553545041 |
| chr8-41340936-41341437 | 3 | 0.989085360487387 |
| chr18-69597797-69598298 | 5 | 0.9834776003012 |
| chr8-47408228-47408729 | 8 | 0.993625891006191 |
| chr19-33997874-33998375 | 8 | 0.953423110772346 |
| chr15-10269848-10270349 | 8 | 0.827178471544481 |
| chr8-112101947-112102448 | 8 | 0.96675797863167 |
| chr1-214859116-214859617 | 3 | 0.986985383402632 |
| chr8-117493544-117494045 | 4 | 0.899676560675129 |
| chr2-70240803-70241304 | 3 | 0.921055698171401 |
| chr4-111186511-111187012 | 3 | 0.976226715254847 |
| chr12-7771964-7772465 | 6 | 0.931017341194756 |
| chr20-47532298-47532799 | 4 | 0.9334062980013 |
| chr19-19937304-19937805 | 1 | 0.9304605867302 |

| | | |
|---|---|---|
| chr8-47475252-47475753 | 8 | 0.962349994533926 |
| chr4-35893735-35894236 | 8 | 0.835792331419292 |
| chr5-142864058-142864559 | 5 | 0.948670711318109 |
| chr20-14299713-14300214 | 8 | 0.993493681128311 |
| chr15-52727078-52727579 | 4 | 0.901468787195521 |
| chr18-72158977-72159478 | 3 | 0.996871671795114 |
| chr13-85139481-85139982 | 8 | 0.981140111734468 |
| chr5-48176802-48177303 | 4 | 0.879452936860905 |
| chr6-20774827-20775328 | 8 | 0.938910437282988 |
| chr1-138144662-138145163 | 3 | 0.990835851233029 |
| chr1-120693834-120694335 | 8 | 0.970456391872389 |
| chr9-93382122-93382623 | 4 | 0.971540908963371 |
| chr4-135682776-135683277 | 6 | 0.991206925481043 |
| chr2-49356606-49357107 | 3 | 0.99593553545041 |
| chr13-102946400-102946901 | 3 | 0.975989124941153 |
| chr2-213234525-213235026 | 3 | 0.996891926787176 |
| chr5-30238810-30239311 | 5 | 0.825123441673364 |
| chr2-50782809-50783310 | 3 | 0.923474471207576 |
| chr3-77806681-77807182 | 3 | 0.94016112501253 |
| chr20-10707228-10707729 | 8 | 0.847382578949827 |
| chr4-68899975-68900476 | 5 | 0.990697477632501 |
| chr9-66020875-66021376 | 5 | 0.991751877563292 |
| chr2-236690805-236691306 | 8 | 0.964914627059513 |
| chr17-66860390-66860891 | 8 | 0.883188615015639 |
| chr1-46311891-46312392 | 3 | 0.985374401281284 |
| chr5-108733260-108733761 | 3 | 0.968988656268091 |
| chr18-30853013-30853514 | 8 | 0.839812516559326 |
| chr11-50588581-50589082 | 7 | 0.851873184223184 |
| chr15-49120883-49121384 | 1 | 0.921767776949711 |
| chr3-78573521-78574022 | 5 | 0.989221690927555 |
| chr7-116800891-116801392 | 2 | 0.887500683667637 |
| chr11-30717413-30717914 | 4 | 0.874098001498627 |
| chr14-99940161-99940662 | 6 | 0.990302643927732 |
| chr7-129199099-129199600 | 8 | 0.822403409049464 |
| chr13-102582663-102583164 | 8 | 0.97073416969907 |
| chr6-86397980-86398481 | 3 | 0.99593553545041 |
| chr3-43006443-43006944 | 8 | 0.993493681128311 |
| chr17-13093514-13094015 | 8 | 0.842195420157346 |
| chr2-121296761-121297262 | 7 | 0.880166615659933 |
| chr7-71294727-71295228 | 8 | 0.97191627880562 |
| chrX-116083363-116083864 | 8 | 0.993493681128311 |
| chr14-83495143-83495644 | 8 | 0.956470966353436 |
| chr15-55389090-55389591 | 8 | 0.976118929704084 |

| | | |
|---|---|---|
| chr7-20089837-20090338 | 4 | 0.837916687510868 |
| chr7-115916402-115916903 | 8 | 0.930975474213084 |
| chr19-55539538-55540039 | 8 | 0.961147709406633 |
| chr5-135911828-135912329 | 8 | 0.891830743649939 |
| chr20-19723122-19723623 | 8 | 0.865818507733769 |
| chr5-103617968-103618469 | 8 | 0.940569780915715 |
| chr1-140980839-140981340 | 3 | 0.99593553545041 |
| chr2-87979091-87979592 | 8 | 0.973530021145424 |
| chr1-173734511-173735012 | 3 | 0.984438783993147 |
| chr1-28301724-28302225 | 6 | 0.990473919448065 |
| chr8-9302563-9303064 | 4 | 0.934444730917181 |
| chr7-90989819-90990320 | 5 | 0.893386743443368 |
| chr5-86714408-86714909 | 3 | 0.995013707022048 |
| chr6-104453721-104454222 | 1 | 0.847147321101607 |
| chr3-77402094-77402595 | 8 | 0.990889748985414 |
| chr9-74526294-74526795 | 4 | 0.947588192904901 |
| chr7-82190433-82190934 | 7 | 0.956637804698692 |
| chr19-24322057-24322558 | 7 | 0.851545857648389 |
| chr1-93679040-93679541 | 3 | 0.980358219927984 |
| chr5-123503664-123504165 | 5 | 0.92375017786249 |
| chr14-7903883-7904384 | 8 | 0.993493681128311 |
| chr12-46552218-46552719 | 3 | 0.970251009461337 |
| chr14-72984530-72985031 | 5 | 0.983589183956838 |
| chr1-168556942-168557443 | 8 | 0.926113513650966 |
| chr2-92851871-92852372 | 8 | 0.992319962909881 |
| chr3-16600207-16600708 | 7 | 0.912361335996487 |
| chr19-28213022-28213523 | 6 | 0.978773195405526 |
| chr18-21334276-21334777 | 3 | 0.99026860895629 |
| chr9-101250465-101250966 | 8 | 0.984495543443049 |
| chr7-129795604-129796105 | 6 | 0.983657761545211 |
| chr18-70330845-70331346 | 5 | 0.983920168976783 |
| chr15-95065057-95065558 | 8 | 0.982733208138574 |
| chr14-77856399-77856900 | 4 | 0.868132365687201 |
| chr8-71842794-71843295 | 8 | 0.993493681128311 |
| chr17-71167474-71167975 | 8 | 0.928778723539888 |
| chr1-45614279-45614780 | 4 | 0.928340231033117 |
| chr1-145826590-145827091 | 7 | 0.860472876160191 |
| chr1-239256866-239257367 | 2 | 0.838523505310303 |
| chr2-246875373-246875874 | 8 | 0.988061215982834 |
| chr19-34312953-34313454 | 3 | 0.988660546241261 |
| chr11-26074962-26075463 | 3 | 0.966790077919273 |
| chr18-56267781-56268282 | 8 | 0.854544026416124 |
| chr7-48689629-48690130 | 3 | 0.991339572179871 |

| | | |
|---|---|---|
| chr16-53599903-53600404 | 3 | 0.988155233037385 |
| chr15-38473043-38473544 | 5 | 0.927097030828788 |
| chr1-149453453-149453954 | 8 | 0.896891207598895 |
| chr5-129167858-129168359 | 3 | 0.990867904526153 |
| chr19-49795656-49796157 | 8 | 0.894849926446798 |
| chr5-38203861-38204362 | 8 | 0.861883514381506 |
| chr18-31972495-31972996 | 8 | 0.821897704044448 |
| chr6-100680032-100680533 | 2 | 0.929625001324912 |
| chr16-53467627-53468128 | 8 | 0.991074776642282 |
| chr13-64080071-64080572 | 3 | 0.848496313541025 |
| chrX-126243749-126244250 | 3 | 0.918453366181199 |
| chr6-138405951-138406452 | 6 | 0.997544061098176 |
| chr8-34063489-34063990 | 8 | 0.919859575623813 |
| chr19-56871740-56872241 | 3 | 0.994185258012372 |
| chr20-25358920-25359421 | 3 | 0.953564996744095 |
| chr2-28397475-28397976 | 2 | 0.875083177358961 |
| chr15-40109652-40110153 | 3 | 0.973781180190657 |
| chr1-256310412-256310913 | 3 | 0.982570118689954 |
| chr10-1177212-1177713 | 1 | 0.913208109182236 |
| chr13-80214452-80214953 | 2 | 0.849230374702641 |
| chr5-103944359-103944860 | 1 | 0.826029494147484 |
| chr18-33715167-33715668 | 3 | 0.99376678175589 |
| chr1-120800814-120801315 | 6 | 0.983500745355969 |
| chr2-31330922-31331423 | 6 | 0.952776475459812 |
| chr18-62639544-62640045 | 8 | 0.981343683590903 |
| chr6-103261716-103262217 | 8 | 0.945845445406466 |
| chr4-113700746-113701247 | 8 | 0.990206754831308 |
| chr10-3528999-3529500 | 5 | 0.906072469161828 |
| chr7-54204032-54204533 | 5 | 0.96158035428489 |
| chr18-20135555-20136056 | 3 | 0.995435821011597 |
| chr1-72212260-72212761 | 3 | 0.981484996512347 |
| chr1-230108188-230108689 | 1 | 0.837048684468271 |
| chr9-4269852-4270353 | 3 | 0.997018757605653 |
| chr6-68650017-68650518 | 3 | 0.994204009436983 |
| chr1-119387722-119388223 | 5 | 0.952295655174773 |
| chr7-94836444-94836945 | 3 | 0.886932456978513 |
| chr17-27694744-27695245 | 3 | 0.961697817823712 |
| chr5-132129714-132130215 | 8 | 0.91441667094124 |
| chr11-52373008-52373509 | 3 | 0.996765282887223 |
| chr19-36401310-36401811 | 8 | 0.980961770945645 |
| chr10-99060104-99060605 | 5 | 0.814721577407869 |
| chr1-207104196-207104697 | 3 | 0.996769606032763 |
| chr11-72289095-72289596 | 5 | 0.933565910217922 |

| | | |
|---|---|---|
| chr13-22487526-22488027 | 6 | 0.991249635048649 |
| chr17-25964757-25965258 | 5 | 0.977845440431194 |
| chr16-36964505-36965006 | 6 | 0.814704493174548 |
| chr7-49896087-49896588 | 3 | 0.99433573842773 |
| chr1-48264382-48264883 | 8 | 0.873246064084021 |
| chr5-66853686-66854187 | 6 | 0.967638375740138 |
| chr2-60551631-60552132 | 2 | 0.955375172398288 |
| chr12-16221218-16221719 | 3 | 0.99593553545041 |
| chr4-55889980-55890481 | 8 | 0.894553625694792 |
| chr2-184532160-184532661 | 8 | 0.962662524798453 |
| chr16-34006563-34007064 | 4 | 0.855167317646074 |
| chr1-196103812-196104313 | 2 | 0.953437644042566 |
| chr2-101860914-101861415 | 6 | 0.996281566115052 |
| chr1-212040931-212041432 | 6 | 0.988349344930207 |
| chr18-66428119-66428620 | 8 | 0.989517295159356 |
| chr2-202038853-202039354 | 8 | 0.914408474668935 |
| chr7-20346471-20346972 | 8 | 0.921224035282421 |
| chr18-47452716-47453217 | 5 | 0.947240396531148 |
| chr6-19779416-19779917 | 8 | 0.93726588841463 |
| chr8-42787533-42788034 | 8 | 0.919597188523461 |
| chr16-83375072-83375573 | 3 | 0.99593553545041 |
| chr1-102890302-102890803 | 8 | 0.978946929539575 |
| chr15-3719381-3719882 | 5 | 0.993654297304987 |
| chr4-55095702-55096203 | 5 | 0.879165192229998 |
| chr1-137803795-137804296 | 3 | 0.996080650315736 |
| chr16-22257261-22257762 | 3 | 0.996084847689616 |
| chr2-68113455-68113956 | 5 | 0.878307809650961 |
| chr8-101748539-101749040 | 3 | 0.973132179189159 |
| chr4-89678437-89678938 | 6 | 0.895417767455491 |
| chr9-38042752-38043253 | 3 | 0.996272024145496 |
| chr10-96892839-96893340 | 6 | 0.955077335498751 |
| chr18-18806501-18807002 | 5 | 0.988495019417891 |
| chr18-2921018-2921519 | 8 | 0.877339071000244 |
| chr18-69308363-69308864 | 7 | 0.921882048747197 |
| chr7-65313694-65314195 | 4 | 0.832024065351236 |
| chr4-116161309-116161810 | 5 | 0.954344714259618 |
| chr11-31029753-31030254 | 4 | 0.868948100113088 |
| chr2-22702335-22702836 | 6 | 0.909789993844034 |
| chr17-1518630-1519131 | 4 | 0.847196332394287 |
| chr8-50172479-50172980 | 8 | 0.973158115619234 |
| chr14-75866146-75866647 | 5 | 0.816095208709966 |
| chr17-81844204-81844705 | 3 | 0.996022657730317 |
| chr6-63734984-63735485 | 3 | 0.99593553545041 |

| | | |
|---|---|---|
| chr14-57193094-57193595 | 5 | 0.993542075757161 |
| chr19-11352218-11352719 | 3 | 0.99593553545041 |
| chr20-16408411-16408912 | 3 | 0.982549085464864 |
| chr1-30996647-30997148 | 2 | 0.915148265905261 |
| chr1-36483571-36484072 | 3 | 0.995308959043756 |
| chr10-72766263-72766764 | 8 | 0.993493681128311 |
| chr3-142411852-142412353 | 8 | 0.884778333477586 |
| chr2-241498464-241498965 | 7 | 0.892216783747899 |
| chr18-45224653-45225154 | 3 | 0.98989255007538 |
| chr8-122023025-122023526 | 5 | 0.979511906133991 |
| chr4-17190423-17190924 | 6 | 0.816847091556287 |
| chr11-16760876-16761377 | 3 | 0.996164289134162 |
| chr9-39421499-39422000 | 5 | 0.98243110506596 |
| chr14-52778537-52779038 | 6 | 0.955468460519548 |
| chr3-78177878-78178379 | 5 | 0.983429969262885 |
| chr15-42135496-42135997 | 8 | 0.831444336542266 |
| chr7-90746860-90747361 | 8 | 0.919389487119298 |
| chr18-37738634-37739135 | 8 | 0.981872700243719 |
| chr20-26294010-26294511 | 5 | 0.980164327006862 |
| chr5-100298065-100298566 | 3 | 0.997770634871527 |
| chr4-33222337-33222838 | 8 | 0.996085084261586 |
| chr2-101147102-101147603 | 3 | 0.99593553545041 |
| chr4-10124963-10125464 | 4 | 0.962186448364447 |
| chr1-60061471-60061972 | 3 | 0.99593553545041 |
| chr6-110687478-110687979 | 6 | 0.983687452533493 |
| chr8-102702595-102703096 | 3 | 0.993361687472077 |
| chr13-87157884-87158385 | 6 | 0.992684140291683 |
| chr5-2061098-2061599 | 7 | 0.881464069082854 |
| chr2-150731002-150731503 | 3 | 0.99593553545041 |
| chr1-98145695-98146196 | 3 | 0.996807273164004 |
| chr5-53708117-53708618 | 6 | 0.932302730993135 |
| chr15-46920079-46920580 | 4 | 0.886670579961246 |
| chr1-50131285-50131786 | 7 | 0.901221602221573 |
| chr13-1638887-1639388 | 3 | 0.946259492314375 |
| chr1-173125525-173126026 | 6 | 0.978039300317217 |
| chr2-144303395-144303896 | 8 | 0.958527643661531 |
| chr6-51015362-51015863 | 4 | 0.843684614423317 |
| chr1-158473512-158474013 | 6 | 0.988874472443196 |
| chr17-64495854-64496355 | 8 | 0.99515233226494 |
| chr10-48774961-48775462 | 5 | 0.853092982186901 |
| chr8-101068786-101069287 | 8 | 0.9928773865149 |
| chr5-110413295-110413796 | 8 | 0.989937189667022 |
| chr9-49885635-49886136 | 8 | 0.820921965453118 |

| | | |
|---|---|---|
| chrX-35045792-35046293 | 5 | 0.887609862422577 |
| chr9-20783405-20783906 | 3 | 0.995651762128785 |
| chr16-79549404-79549905 | 7 | 0.928538057055677 |
| chr17-27801418-27801919 | 8 | 0.889107406824634 |
| chr12-13334852-13335353 | 3 | 0.983981860521442 |
| chr3-135072483-135072984 | 6 | 0.99249975415989 |
| chr5-123471387-123471888 | 3 | 0.979228032764253 |
| chr16-80280978-80281479 | 6 | 0.991803993577903 |
| chr2-37690240-37690741 | 1 | 0.883560226127921 |
| chr1-171824453-171824954 | 8 | 0.986568438788106 |
| chr8-29960606-29961107 | 8 | 0.993493681128311 |
| chr2-23086729-23087230 | 6 | 0.987765074728663 |
| chr2-137267071-137267572 | 7 | 0.882624253906271 |
| chr6-72003534-72004035 | 3 | 0.985573685040978 |
| chr16-6428400-6428901 | 7 | 0.90934101191474 |
| chr5-146728451-146728952 | 8 | 0.939616696782719 |
| chr2-205520831-205521332 | 8 | 0.984654261392286 |
| chr9-12726867-12727368 | 6 | 0.992521704626841 |
| chr3-129160847-129161348 | 3 | 0.997662929318247 |
| chr9-72258279-72258780 | 3 | 0.860400019042801 |
| chr4-168967387-168967888 | 3 | 0.995361666739754 |
| chr1-78323972-78324473 | 1 | 0.85737815783084 |
| chr8-119487268-119487769 | 3 | 0.982862619701189 |
| chr1-178101115-178101616 | 8 | 0.994702640714364 |
| chr8-101769825-101770326 | 1 | 0.884694031296022 |
| chr9-45032159-45032660 | 8 | 0.914603711923742 |
| chr5-61640382-61640883 | 3 | 0.996835228779196 |
| chrX-30423204-30423705 | 6 | 0.991598049934165 |
| chr2-108404624-108405125 | 3 | 0.99665882911282 |
| chr9-59266591-59267092 | 8 | 0.93873875971042 |
| chr6-3719397-3719898 | 1 | 0.829131961817219 |
| chr16-1954018-1954519 | 8 | 0.990657376063323 |
| chr2-196767147-196767648 | 8 | 0.928069647495318 |
| chr5-119367897-119368398 | 6 | 0.994740498879575 |
| chr8-111446608-111447109 | 6 | 0.901274728212712 |
| chr5-2889164-2889665 | 3 | 0.995574667621554 |
| chr2-235447287-235447788 | 8 | 0.951603040847117 |
| chr14-64538710-64539211 | 6 | 0.993070730320936 |
| chr10-34072631-34073132 | 4 | 0.872407393072059 |
| chr9-88122740-88123241 | 8 | 0.980616191610055 |
| chr3-141361748-141362249 | 8 | 0.892217307473522 |
| chr4-23736543-23737044 | 3 | 0.985414938111329 |
| chr16-66706452-66706953 | 3 | 0.830895334650008 |

| | | |
|---|---|---|
| chr18-71341529-71342030 | 3 | 0.995788133461565 |
| chr2-143709150-143709651 | 8 | 0.95251616078648 |
| chr15-52482557-52483058 | 3 | 0.990184285243731 |
| chr2-5164187-5164688 | 2 | 0.893693162737138 |
| chr9-37766409-37766910 | 3 | 0.991288559990946 |
| chr3-167429745-167430246 | 5 | 0.989526036454242 |
| chr9-57070780-57071281 | 4 | 0.933630747855905 |
| chr5-120285341-120285842 | 2 | 0.912757325913238 |
| chr3-117577686-117578187 | 6 | 0.943769352035167 |
| chr19-26780602-26781103 | 4 | 0.828162112245263 |
| chr9-42444760-42445261 | 1 | 0.925186380219955 |
| chr2-188759994-188760495 | 7 | 0.834783988214067 |
| chr9-21543379-21543880 | 3 | 0.990162939654253 |
| chr3-147513608-147514109 | 6 | 0.9419382821166 |
| chr2-24513288-24513789 | 3 | 0.994373273868786 |
| chr5-119905541-119906042 | 8 | 0.998169962550069 |
| chr9-42672236-42672737 | 8 | 0.843126667532485 |
| chr9-58686797-58687298 | 3 | 0.98755506708969 |
| chr17-16201575-16202076 | 8 | 0.967244468054483 |
| chr1-216850090-216850591 | 3 | 0.996233423676031 |
| chr15-35028844-35029345 | 8 | 0.940075739989327 |
| chr12-10720101-10720602 | 5 | 0.986994992872723 |
| chr4-58135237-58135738 | 8 | 0.939815143017519 |
| chr2-188106941-188107442 | 5 | 0.990097870541145 |
| chr3-145846878-145847379 | 3 | 0.995196655607805 |
| chr5-137081140-137081641 | 8 | 0.945934110462412 |
| chr1-249869057-249869558 | 6 | 0.991206925481043 |
| chr13-18449256-18449757 | 3 | 0.99593553545041 |
| chr11-68377653-68378154 | 8 | 0.901420597299792 |
| chr1-259307803-259308304 | 8 | 0.931598762233506 |
| chr2-99965525-99966026 | 3 | 0.995206178253483 |
| chr4-33154424-33154925 | 5 | 0.877111624979076 |
| chr1-11985032-11985533 | 8 | 0.95663955472402 |
| chr4-133577693-133578194 | 2 | 0.865099376726939 |
| chr1-191448305-191448806 | 8 | 0.896109982185941 |
| chr2-46953711-46954212 | 5 | 0.97946983751548 |
| chr17-66438436-66438937 | 8 | 0.989363373124784 |
| chr1-117246643-117247144 | 3 | 0.949652378337432 |
| chr17-68856021-68856522 | 3 | 0.98164373114326 |
| chr15-6960432-6960933 | 6 | 0.989326307670765 |
| chr9-63823192-63823693 | 5 | 0.905334893932385 |
| chr1-67790053-67790554 | 8 | 0.995239673474855 |
| chr5-154697328-154697829 | 8 | 0.844186413900344 |

| | | |
|---|---|---|
| chrX-116221799-116222300 | 5 | 0.971008718939277 |
| chr2-183271693-183272194 | 2 | 0.867044362702551 |
| chr7-81069547-81070048 | 3 | 0.99593553545041 |
| chr3-78294716-78295217 | 8 | 0.850167424821073 |
| chr1-213241237-213241738 | 7 | 0.894829 |
| chr12-43002320-43002821 | 3 | 0.857864387479398 |
| chr3-145904309-145904810 | 6 | 0.989762832855026 |
| chr15-32398296-32398797 | 4 | 0.916240216438904 |
| chr19-12942368-12942869 | 5 | 0.96041644976013 |
| chr7-17284962-17285463 | 3 | 0.994379723098762 |
| chr14-54903493-54903994 | 3 | 0.982222043075846 |
| chr3-42548377-42548878 | 3 | 0.996862674119375 |
| chr5-74700833-74701334 | 8 | 0.971635718958247 |
| chr15-25779555-25780056 | 5 | 0.977909868105945 |
| chr13-76326905-76327406 | 8 | 0.971563342898172 |
| chr2-197966248-197966749 | 3 | 0.984526023806785 |
| chr13-99459385-99459886 | 3 | 0.973294127135665 |
| chr13-77179658-77180159 | 4 | 0.919622750214585 |
| chr8-48546617-48547118 | 1 | 0.874100817701954 |
| chr12-39368728-39369229 | 8 | 0.979314273955132 |
| chr7-22336466-22336967 | 2 | 0.946144874937486 |
| chr1-169231777-169232278 | 8 | 0.995173406049565 |
| chr2-190740861-190741362 | 8 | 0.993493681128311 |
| chr5-91221132-91221633 | 5 | 0.974949357228145 |
| chr14-81830209-81830710 | 3 | 0.995927856842658 |
| chrX-30047481-30047982 | 8 | 0.820168474568456 |
| chr7-58438392-58438893 | 5 | 0.852827076159195 |
| chr8-66586167-66586668 | 3 | 0.832746909814684 |
| chr6-40859776-40860277 | 8 | 0.966876294579102 |
| chr9-111343583-111344084 | 8 | 0.964221834449262 |
| chr15-89292021-89292522 | 3 | 0.850219412150736 |
| chr1-121747392-121747893 | 6 | 0.817466662418516 |
| chr16-82221270-82221771 | 6 | 0.8277321121875 |
| chr8-25930232-25930733 | 6 | 0.967542962936556 |
| chr4-67454586-67455087 | 8 | 0.954875664657928 |
| chr2-119733933-119734434 | 3 | 0.996883950621111 |
| chr4-80872094-80872595 | 3 | 0.99679107070141 |
| chr15-93057451-93057952 | 6 | 0.987102814885328 |
| chr13-86713480-86713981 | 8 | 0.884355653033269 |
| chr15-33284636-33285137 | 8 | 0.983587276572597 |
| chr19-14965723-14966224 | 6 | 0.917801157400926 |
| chrX-11911896-11912397 | 6 | 0.829894555944284 |
| chr9-59754563-59755064 | 3 | 0.99437354420506 |

| | | |
|---|---|---|
| chr6-51522927-51523428 | 1 | 0.838438517094273 |
| chr5-16582574-16583075 | 8 | 0.851239759573908 |
| chr17-28854739-28855240 | 4 | 0.933241452771631 |
| chr10-38022807-38023308 | 2 | 0.899978321707217 |
| chr5-61689492-61689993 | 3 | 0.997856564513066 |
| chr6-110187804-110188305 | 3 | 0.989293107663766 |
| chr11-14669790-14670291 | 3 | 0.99593553545041 |
| chr10-57822905-57823406 | 5 | 0.991593868605148 |
| chr2-234048233-234048734 | 8 | 0.931976715134812 |
| chr8-46602910-46603411 | 8 | 0.974688891868676 |
| chr2-110201294-110201795 | 3 | 0.818119772574765 |
| chr2-229356303-229356804 | 3 | 0.99241157486974 |
| chr2-207029539-207030040 | 6 | 0.966482402139168 |
| chr2-27641274-27641775 | 3 | 0.987358900344792 |
| chr15-35553290-35553791 | 4 | 0.951394942874892 |
| chr2-71209879-71210380 | 3 | 0.994783185509647 |
| chr14-52865318-52865819 | 8 | 0.910697214069296 |
| chr20-52452658-52453159 | 3 | 0.996729771489655 |
| chr1-187281955-187282456 | 8 | 0.933531378013666 |
| chr11-83404880-83405381 | 8 | 0.901896423781149 |
| chr18-55383274-55383775 | 3 | 0.962781418002175 |
| chr16-64735651-64736152 | 5 | 0.985370343016277 |
| chr16-35332590-35333091 | 6 | 0.951186061471785 |
| chr13-16087076-16087577 | 4 | 0.926392278093125 |
| chr8-111422548-111423049 | 3 | 0.957992671403857 |
| chr5-19913010-19913511 | 6 | 0.848328443248679 |
| chr16-6994991-6995492 | 2 | 0.894251484007404 |
| chr16-71423522-71424023 | 2 | 0.924612508395985 |
| chr6-86299164-86299665 | 5 | 0.991521993870562 |
| chr3-91884791-91885292 | 3 | 0.988503524696689 |
| chr4-51352771-51353272 | 8 | 0.839892943348725 |
| chr1-52304314-52304815 | 8 | 0.821062355824195 |
| chr3-162915324-162915825 | 5 | 0.931776889442451 |
| chr19-855084-855585 | 1 | 0.889816544015958 |
| chr2-94026060-94026561 | 4 | 0.842771813426673 |
| chr5-60449633-60450134 | 8 | 0.943495040372607 |
| chr3-156719882-156720383 | 7 | 0.818690871514872 |
| chr7-115589716-115590217 | 8 | 0.944091436376421 |
| chr9-47353340-47353841 | 8 | 0.879194871681441 |
| chr19-53848976-53849477 | 4 | 0.938855868693034 |
| chr8-118361775-118362276 | 8 | 0.837767370167247 |
| chr5-55956174-55956675 | 8 | 0.908148294757674 |
| chr3-107703616-107704117 | 8 | 0.972916368835791 |

| | | |
|---|---|---|
| chr11-84539622-84540123 | 6 | 0.991206925481043 |
| chr15-452484-452985 | 6 | 0.989071244094626 |
| chr1-161512810-161513311 | 8 | 0.993493681128311 |
| chr10-41409276-41409777 | 1 | 0.92736281443066 |
| chr16-29553052-29553553 | 4 | 0.959132416134295 |
| chr18-61307299-61307800 | 3 | 0.940740620882175 |
| chrX-46807956-46808457 | 3 | 0.89518242654817 |
| chr14-13562685-13563186 | 6 | 0.911727763754581 |
| chr4-85510613-85511114 | 1 | 0.903561177881908 |
| chr5-118394916-118395417 | 2 | 0.833993481217412 |
| chr9-1787357-1787858 | 6 | 0.990810728010163 |
| chr20-43992309-43992810 | 8 | 0.995200110549519 |
| chr2-138344258-138344759 | 3 | 0.985533830059371 |
| chr2-40662016-40662517 | 8 | 0.913756969031869 |
| chr2-189545918-189546419 | 8 | 0.840572391702911 |
| chr4-50580857-50581358 | 1 | 0.927604072035894 |
| chr5-113241923-113242424 | 3 | 0.996584577905947 |
| chr1-14718348-14718849 | 8 | 0.989042828067529 |
| chr20-9517422-9517923 | 3 | 0.983707919327671 |
| chr17-6363466-6363967 | 8 | 0.994997273549595 |
| chr9-42943328-42943829 | 6 | 0.933067241402514 |
| chr4-137681818-137682319 | 3 | 0.99593553545041 |
| chr5-41819065-41819566 | 6 | 0.98518135827704 |
| chr18-78708276-78708777 | 3 | 0.995823991788683 |
| chr17-68577143-68577644 | 8 | 0.982503745930165 |
| chr2-24299115-24299616 | 6 | 0.962642113481868 |
| chr10-78628832-78629333 | 3 | 0.996853714341045 |
| chr9-45231392-45231893 | 5 | 0.968053210081435 |
| chr15-14250422-14250923 | 6 | 0.861635733612748 |
| chr14-10379467-10379968 | 1 | 0.980565514811525 |
| chr16-69929860-69930361 | 1 | 0.923672253670683 |
| chr2-227403508-227404009 | 5 | 0.820565287165639 |
| chr13-54846762-54847263 | 1 | 0.955920143159584 |
| chr3-59866148-59866649 | 5 | 0.831506811732692 |
| chr12-15006072-15006573 | 8 | 0.815684028459289 |
| chrX-139850600-139851101 | 5 | 0.934702936494926 |
| chr15-99217256-99217757 | 4 | 0.917895176446479 |
| chr20-38282887-38283388 | 3 | 0.858229772273228 |
| chr16-69907501-69908002 | 3 | 0.965879173324053 |
| chr14-93804913-93805414 | 7 | 0.959123726353062 |
| chr7-116180960-116181461 | 8 | 0.896468629972947 |
| chr1-260109446-260109947 | 8 | 0.946020970381989 |
| chr8-26941336-26941837 | 4 | 0.951718485681677 |

| | | |
|---|---|---|
| chr16-36130802-36131303 | 8 | 0.898053473949122 |
| chr4-181774054-181774555 | 5 | 0.99130519299749 |
| chr7-28849788-28850289 | 3 | 0.996785449897633 |
| chr11-61754340-61754841 | 8 | 0.990477550361356 |
| chr1-216537711-216538212 | 2 | 0.829756184056312 |
| chr4-39621555-39622056 | 3 | 0.996787959960612 |
| chr10-25819989-25820490 | 8 | 0.901373664441528 |
| chr6-3439650-3440151 | 3 | 0.997930646873029 |
| chr14-55499353-55499854 | 8 | 0.993493681128311 |
| chr15-10415873-10416374 | 6 | 0.993214316467802 |
| chr16-17449820-17450321 | 8 | 0.849804750864519 |
| chr9-58500227-58500728 | 3 | 0.997175185530614 |
| chr2-28545489-28545990 | 7 | 0.932279938997871 |
| chr1-156679566-156680067 | 3 | 0.996033437488201 |
| chr15-94886706-94887207 | 4 | 0.837306835260203 |
| chr1-94989896-94990397 | 8 | 0.944774140404828 |
| chr8-63709781-63710282 | 1 | 0.954444619002682 |
| chr6-91458675-91459176 | 8 | 0.881914090935857 |
| chr19-36062550-36063051 | 8 | 0.997349886794605 |
| chr18-3022338-3022839 | 5 | 0.944663697302243 |
| chr15-37757134-37757635 | 6 | 0.986539625362955 |
| chr14-30446566-30447067 | 3 | 0.989737371617513 |
| chr15-75294676-75295177 | 3 | 0.99519653936785 |
| chr4-11238907-11239408 | 8 | 0.98566155825857 |
| chr17-20868973-20869474 | 8 | 0.97599148315476 |
| chr13-87898798-87899299 | 5 | 0.972302967168806 |
| chr4-80324746-80325247 | 3 | 0.872034919645522 |
| chr9-59296835-59297336 | 4 | 0.886401409515673 |
| chr4-147614366-147614867 | 3 | 0.989496528085492 |
| chr3-25097956-25098457 | 3 | 0.996781115131545 |
| chr1-136721964-136722465 | 8 | 0.873326137518571 |
| chr18-6610758-6611259 | 2 | 0.931480995552814 |
| chr16-12906426-12906927 | 6 | 0.989547771956942 |
| chr3-36794561-36795062 | 2 | 0.901537617475246 |
| chr15-82500663-82501164 | 8 | 0.986977517993786 |
| chr1-152288807-152289308 | 3 | 0.996528568860468 |
| chrX-38633352-38633853 | 3 | 0.99593553545041 |
| chr1-21585463-21585964 | 3 | 0.996738457809876 |
| chr2-79577427-79577928 | 8 | 0.920504119296691 |
| chr5-122337338-122337839 | 8 | 0.892634129892433 |
| chr19-33192795-33193296 | 8 | 0.889705343396788 |
| chr6-45356656-45357157 | 6 | 0.991819222401598 |
| chr19-47386651-47387152 | 6 | 0.931986491414273 |

| | | |
|---|---|---|
| chr7-24298211-24298712 | 6 | 0.990291597900478 |
| chr18-15638371-15638872 | 8 | 0.993493681128311 |
| chr18-52334774-52335275 | 5 | 0.959272875147344 |
| chr8-46934036-46934537 | 5 | 0.984869109373247 |
| chr18-40286896-40287397 | 4 | 0.919300589168551 |
| chr16-33151845-33152346 | 6 | 0.99612820892675 |
| chr4-104253944-104254445 | 5 | 0.980216807099712 |
| chr3-59827500-59828001 | 7 | 0.85541897841837 |
| chr3-18763591-18764092 | 8 | 0.818016260777405 |
| chrX-81618051-81618552 | 3 | 0.99593553545041 |
| chr16-72126868-72127369 | 3 | 0.886906160153557 |
| chr2-158685378-158685879 | 3 | 0.900699546355479 |
| chr1-107664995-107665496 | 7 | 0.920826585645134 |
| chr8-115273191-115273692 | 4 | 0.822738694010671 |
| chr16-18178877-18179378 | 5 | 0.97603062752312 |
| chr20-45258499-45259000 | 6 | 0.997148273292681 |
| chr4-164673150-164673651 | 3 | 0.993446777449637 |
| chr4-20145645-20146146 | 8 | 0.994866899473428 |
| chr1-31793096-31793597 | 3 | 0.996444560010941 |
| chr9-111808243-111808744 | 6 | 0.967416672023132 |
| chr1-80360074-80360575 | 8 | 0.899025683443877 |
| chr13-62966682-62967183 | 3 | 0.980814742296655 |
| chr15-79075228-79075729 | 8 | 0.983747061922277 |
| chr8-80177589-80178090 | 8 | 0.921852647966197 |
| chr10-26272171-26272672 | 6 | 0.989690622310852 |
| chr7-29053397-29053898 | 3 | 0.82448024739674 |
| chr10-101469825-101470326 | 5 | 0.868067802468253 |
| chr12-44163429-44163930 | 8 | 0.914865155528144 |
| chr9-108450093-108450594 | 3 | 0.976688023008166 |
| chr14-97037994-97038495 | 4 | 0.876717997701411 |
| chr5-50596852-50597353 | 8 | 0.881434748692063 |
| chr17-51774820-51775321 | 5 | 0.984940598794784 |
| chr11-13894131-13894632 | 8 | 0.986004527074238 |
| chr4-178100214-178100715 | 6 | 0.988936578293106 |
| chr6-48398007-48398508 | 3 | 0.996856911826977 |
| chr4-95317677-95318178 | 4 | 0.907990068460588 |
| chr2-102632126-102632627 | 3 | 0.988342813788185 |
| chr4-15342264-15342765 | 8 | 0.993493681128311 |
| chr6-139151473-139151974 | 3 | 0.94091283001361 |
| chr2-77187876-77188377 | 1 | 0.848231461977924 |
| chr6-50045122-50045623 | 8 | 0.993915920040576 |
| chrX-79034462-79034963 | 3 | 0.991649671403033 |
| chr6-14756761-14757262 | 2 | 0.971889201426676 |

| | | |
|---|---|---|
| chr5-7986040-7986541 | 6 | 0.926625674962354 |
| chr17-12956425-12956926 | 2 | 0.846403948353355 |
| chr3-52294804-52295305 | 8 | 0.977776136337669 |
| chr9-80651629-80652130 | 3 | 0.932632819930373 |
| chr1-6812320-6812821 | 5 | 0.983919978840557 |
| chr10-35751911-35752412 | 8 | 0.992643521666672 |
| chr3-110732658-110733159 | 1 | 0.922946788822707 |
| chr5-59566021-59566522 | 7 | 0.865264640554019 |
| chr5-54383106-54383607 | 7 | 0.850035280136388 |
| chr4-108350503-108351004 | 3 | 0.99593553545041 |
| chr9-112475295-112475796 | 3 | 0.996339321561447 |
| chr5-97244827-97245328 | 2 | 0.883000452589677 |
| chr8-62771749-62772250 | 8 | 0.958535428074057 |
| chr20-45781566-45782067 | 6 | 0.927007282327117 |
| chr7-90183447-90183948 | 8 | 0.81563683610947 |
| chr5-77737866-77738367 | 8 | 0.885406895758978 |
| chr6-6695441-6695942 | 2 | 0.966825472354166 |
| chrX-68971784-68972285 | 5 | 0.952008755934658 |
| chr20-7089266-7089767 | 7 | 0.820537346961012 |
| chr6-108320473-108320974 | 8 | 0.99521137587376 |
| chr2-214955847-214956348 | 5 | 0.978814847176666 |
| chr4-65107923-65108424 | 8 | 0.959108825234367 |
| chr16-81234825-81235326 | 1 | 0.916328944909764 |
| chr6-73571850-73572351 | 5 | 0.980129119993335 |
| chr19-33614977-33615478 | 5 | 0.988829036707359 |
| chr5-100091389-100091890 | 4 | 0.850774137356991 |
| chr10-75017525-75018026 | 3 | 0.996461347716699 |
| chr2-136739120-136739621 | 1 | 0.913498821279849 |
| chr10-11286333-11286834 | 4 | 0.909634901518828 |
| chr2-177572078-177572579 | 3 | 0.99047612582258 |
| chr20-9670116-9670617 | 1 | 0.819792041983034 |
| chr3-32332382-32332883 | 3 | 0.995595474605443 |
| chr5-138081086-138081587 | 8 | 0.819856449551955 |
| chr10-89387463-89387964 | 8 | 0.938526204196328 |
| chr3-168999800-169000301 | 4 | 0.905162138900184 |
| chr15-72337758-72338259 | 8 | 0.932363719020969 |
| chr17-21410375-21410876 | 5 | 0.989371066924074 |
| chr2-164852035-164852536 | 5 | 0.987511615481425 |
| chr6-13003433-13003934 | 4 | 0.971701476117084 |
| chr5-46421695-46422196 | 5 | 0.978338127306618 |
| chr15-51237226-51237727 | 4 | 0.884672028293672 |
| chr8-102136123-102136624 | 4 | 0.897610601009573 |
| chr2-166447827-166448328 | 5 | 0.976498868212392 |

| | | |
|---|---|---|
| chr17-75204899-75205400 | 7 | 0.821407444259472 |
| chr9-19058388-19058889 | 8 | 0.995169025235181 |
| chr5-121311532-121312033 | 8 | 0.823932020452621 |
| chr4-14550469-14550970 | 8 | 0.866536896865737 |
| chr3-38292250-38292751 | 6 | 0.9922742898958 |
| chr11-25791555-25792056 | 3 | 0.985892252972753 |
| chr6-104118687-104119188 | 8 | 0.935535537480613 |
| chr20-36404819-36405320 | 8 | 0.995056992043118 |
| chr14-57865116-57865617 | 4 | 0.955756072529305 |
| chr4-62852161-62852662 | 6 | 0.875886260474434 |
| chr4-61268803-61269304 | 8 | 0.972775879017443 |
| chr1-204079750-204080251 | 8 | 0.905778070258203 |
| chrX-17621151-17621652 | 2 | 0.889044759590379 |
| chr18-82804211-82804712 | 3 | 0.985380167008127 |
| chr2-110455505-110456006 | 8 | 0.884987182394637 |
| chr9-105407370-105407871 | 8 | 0.977896401088148 |
| chr1-221041366-221041867 | 3 | 0.99593553545041 |
| chr16-4748559-4749060 | 5 | 0.956883175381996 |
| chr4-29298316-29298817 | 2 | 0.831796798410195 |
| chr18-23252217-23252718 | 8 | 0.913549532775258 |
| chr2-175830894-175831395 | 8 | 0.831198238590632 |
| chrX-111456213-111456714 | 5 | 0.962534264581091 |
| chr15-62034603-62035104 | 8 | 0.814792041720439 |
| chr13-103256930-103257431 | 6 | 0.971249639845022 |
| chr13-36815894-36816395 | 5 | 0.843530625705763 |
| chr8-89699156-89699657 | 8 | 0.986549472247094 |
| chrY-6814517-6815018 | 3 | 0.990596827919659 |
| chr4-69775012-69775513 | 4 | 0.869775425472001 |
| chr14-79928927-79929428 | 8 | 0.986767730254833 |
| chr17-12352480-12352981 | 5 | 0.982405395335005 |
| chr6-117259816-117260317 | 2 | 0.953464004918759 |
| chr13-101595330-101595831 | 7 | 0.879010681156722 |
| chr3-117346618-117347119 | 8 | 0.86352515469602 |
| chr15-92195404-92195905 | 2 | 0.834891379728613 |
| chr11-47522795-47523296 | 5 | 0.956870992707019 |
| chr6-84058661-84059162 | 6 | 0.993186027050736 |
| chr17-1317337-1317838 | 3 | 0.99593553545041 |
| chr8-98245304-98245805 | 3 | 0.99593553545041 |
| chr13-98237559-98238060 | 7 | 0.88272098995403 |
| chr1-180870991-180871492 | 8 | 0.844915282853926 |
| chr9-106977793-106978294 | 8 | 0.931081590573719 |
| chr4-28536137-28536638 | 1 | 0.850826574774246 |
| chr6-121973469-121973970 | 8 | 0.890725356567598 |

| | | |
|---|---|---|
| chr13-47262652-47263153 | 8 | 0.894205705121029 |
| chr4-151012193-151012694 | 8 | 0.879440924374751 |
| chr8-77952506-77953007 | 8 | 0.97164450963523 |
| chr3-55629001-55629502 | 5 | 0.816515511277926 |
| chr5-138082004-138082505 | 2 | 0.85507506100352 |
| chr9-41582402-41582903 | 1 | 0.832556994827119 |
| chr9-80628850-80629351 | 2 | 0.931056713137803 |
| chr7-107305185-107305686 | 7 | 0.953296292242156 |
| chr10-75850391-75850892 | 6 | 0.917355161615008 |
| chr5-161665477-161665978 | 8 | 0.899783576091521 |
| chrX-133075456-133075957 | 1 | 0.850168477157195 |
| chr1-90329326-90329827 | 5 | 0.964693067161461 |
| chr7-68573536-68574037 | 5 | 0.976298550372357 |
| chr2-30982875-30983376 | 8 | 0.815143358650224 |
| chr18-76905978-76906479 | 8 | 0.964572587035592 |
| chr20-20103187-20103688 | 6 | 0.99497434619307 |
| chr4-80899788-80900289 | 3 | 0.988616904346355 |
| chr2-234021055-234021556 | 4 | 0.891143632405645 |
| chr2-111267934-111268435 | 2 | 0.947685104390663 |
| chr3-34012523-34013024 | 4 | 0.912984835222001 |
| chr3-34177861-34178362 | 7 | 0.907643808757694 |
| chr10-4404502-4405003 | 8 | 0.898699990995408 |
| chr3-107941382-107941883 | 6 | 0.992861233540875 |
| chr9-70730199-70730700 | 6 | 0.980660845815372 |
| chr2-126108837-126109338 | 3 | 0.99236739025359 |
| chr13-22576965-22577466 | 8 | 0.97969062170687 |
| chr13-93492832-93493333 | 8 | 0.921086272359416 |
| chr5-104358687-104359188 | 6 | 0.993200508190598 |
| chr2-95851177-95851678 | 3 | 0.99595853924407 |
| chr5-145522930-145523431 | 8 | 0.981097811429857 |
| chr15-47916892-47917393 | 8 | 0.935633086893808 |
| chr18-69545924-69546425 | 5 | 0.990354926155878 |
| chr17-42022789-42023290 | 3 | 0.954923151784022 |
| chr7-39779528-39780029 | 3 | 0.960866444379565 |
| chr1-151655498-151655999 | 7 | 0.922719711542211 |
| chr4-86817947-86818448 | 3 | 0.99769331504779 |
| chr13-67827860-67828361 | 5 | 0.925348059985929 |
| chr2-70702790-70703291 | 3 | 0.9963597202227 |
| chr12-27363519-27364020 | 8 | 0.994777293103272 |
| chr4-77355687-77356188 | 2 | 0.960274021446646 |
| chr5-66741051-66741552 | 6 | 0.993225565110815 |
| chr13-44434179-44434680 | 6 | 0.989750081460807 |
| chr3-108562176-108562677 | 5 | 0.971232421184897 |

| | | |
|---|---|---|
| chr1-176844484-176844985 | 6 | 0.984642952250453 |
| chr5-43288646-43289147 | 6 | 0.992287503716321 |
| chr3-22857001-22857502 | 4 | 0.945781511146598 |
| chr7-92873951-92874452 | 6 | 0.92805166238011 |
| chr7-19482647-19483148 | 3 | 0.989958565042543 |
| chr13-71725834-71726335 | 5 | 0.976476377992192 |
| chr19-22816733-22817234 | 5 | 0.987736616383114 |
| chr12-27661811-27662312 | 8 | 0.835310497679478 |
| chr5-107936011-107936512 | 6 | 0.938019474430184 |
| chr8-69711213-69711714 | 5 | 0.901226806775154 |
| chr14-58554701-58555202 | 8 | 0.910755219496596 |
| chr2-32543205-32543706 | 3 | 0.996240910461671 |
| chr5-116451556-116452057 | 6 | 0.991206925481043 |
| chr4-51432620-51433121 | 4 | 0.898583506664569 |
| chr3-8303581-8304082 | 4 | 0.903465968217285 |
| chr4-4991486-4991987 | 6 | 0.991206925481043 |
| chr10-67133451-67133952 | 5 | 0.853314349935117 |
| chr15-51766908-51767409 | 8 | 0.949507149261252 |
| chr16-47160088-47160589 | 3 | 0.996781153337878 |
| chr7-113875911-113876412 | 5 | 0.962081244085662 |
| chr16-69459133-69459634 | 7 | 0.919707041289748 |
| chr17-74625488-74625989 | 8 | 0.987252065211722 |
| chr13-100444520-100445021 | 5 | 0.987874286797518 |
| chr19-25857277-25857778 | 2 | 0.955886482931551 |
| chr7-66590509-66591010 | 3 | 0.995588573809693 |
| chr2-173093678-173094179 | 3 | 0.984224318438049 |
| chr4-138851181-138851682 | 8 | 0.937699793173059 |
| chr3-139758473-139758974 | 8 | 0.929422147551718 |
| chr2-149486580-149487081 | 7 | 0.945107266501984 |
| chr6-60892525-60893026 | 8 | 0.850013061687972 |
| chr5-121183060-121183561 | 4 | 0.83322947641694 |
| chr3-119138049-119138550 | 8 | 0.89575393747998 |
| chr16-42703998-42704499 | 5 | 0.972412874269378 |
| chr6-138783719-138784220 | 6 | 0.993208891079114 |
| chr14-24767801-24768302 | 3 | 0.97619171975314 |
| chr11-65881994-65882495 | 2 | 0.971351201616674 |
| chr7-112898514-112899015 | 8 | 0.934668024688541 |
| chr4-61101386-61101887 | 1 | 0.836079099729529 |
| chr6-42388869-42389370 | 4 | 0.883579190572533 |
| chr2-56182266-56182767 | 1 | 0.847024055306116 |
| chr10-65690634-65691135 | 8 | 0.856603428285659 |
| chr11-24655019-24655520 | 1 | 0.90232450323127 |
| chr9-71447742-71448243 | 2 | 0.931524643541087 |

| | | |
|---|---|---|
| chrX-133443702-133444203 | 5 | 0.94605930546835 |
| chr15-81535471-81535972 | 8 | 0.885616333571574 |
| chr1-224896920-224897421 | 6 | 0.992793451071027 |
| chr20-16659393-16659894 | 3 | 0.981038255521931 |
| chr12-44798226-44798727 | 5 | 0.977015722774707 |
| chr2-197526706-197527207 | 5 | 0.930985123972792 |
| chr5-166716377-166716878 | 8 | 0.870367159415136 |
| chr2-244913939-244914440 | 3 | 0.997717833845903 |
| chr10-40102570-40103071 | 3 | 0.992330925201346 |
| chr2-184018823-184019324 | 5 | 0.843911051135678 |
| chrX-56282340-56282841 | 5 | 0.950802106938555 |
| chr16-2929590-2930091 | 6 | 0.992877696061462 |
| chr17-76581827-76582328 | 3 | 0.831852774241139 |
| chr5-119933978-119934479 | 8 | 0.874798275784264 |
| chr4-98550458-98550959 | 3 | 0.996886624752824 |
| chr6-135209281-135209782 | 6 | 0.962594286651167 |
| chr9-38958817-38959318 | 2 | 0.937465093935004 |
| chr5-79323704-79324205 | 8 | 0.939610001849561 |
| chr3-57287136-57287637 | 8 | 0.952242830939079 |
| chr17-30322609-30323110 | 3 | 0.99779202960572 |
| chr13-8070067-8070568 | 5 | 0.881371690838569 |
| chr1-79201038-79201539 | 8 | 0.97065498821039 |
| chr2-221131220-221131721 | 5 | 0.978269861451436 |
| chr7-110650526-110651027 | 4 | 0.841634613978747 |
| chr4-53592245-53592746 | 5 | 0.986872808133299 |
| chr1-132208123-132208624 | 6 | 0.991206925481043 |
| chr16-57517848-57518349 | 8 | 0.901274564761519 |
| chr14-6794398-6794899 | 3 | 0.978770881681911 |
| chr15-37541331-37541832 | 8 | 0.936936895170635 |
| chr4-168511714-168512215 | 4 | 0.851723620984788 |
| chr5-164282872-164283373 | 8 | 0.991322790963401 |
| chr8-43003485-43003986 | 4 | 0.820380943104056 |
| chr17-8181553-8182054 | 8 | 0.990764693419537 |
| chr13-63712435-63712936 | 8 | 0.956367208113501 |
| chr16-9185342-9185843 | 8 | 0.833139274926382 |
| chr11-39784599-39785100 | 6 | 0.983731044222283 |
| chr1-179688619-179689120 | 8 | 0.861010238741091 |
| chr13-106151801-106152302 | 8 | 0.846095513486607 |
| chr9-110861519-110862020 | 8 | 0.972349141605752 |
| chr2-27846696-27847197 | 8 | 0.938219540605475 |
| chr8-78446097-78446598 | 6 | 0.995269188744307 |
| chr8-59340577-59341078 | 6 | 0.88144976279311 |
| chr17-24714300-24714801 | 5 | 0.983892405756546 |

| | | |
|---|---|---|
| chr9-22120553-22121054 | 3 | 0.984142853494004 |
| chr17-7527292-7527793 | 8 | 0.961305772598806 |
| chr4-3889037-3889538 | 8 | 0.985293638651764 |
| chr14-99477196-99477697 | 8 | 0.978712332218583 |
| chr9-92914639-92915140 | 3 | 0.996456099131366 |
| chr8-6776999-6777500 | 8 | 0.903294089578131 |
| chr5-39553923-39554424 | 3 | 0.992128483580648 |
| chr17-52488637-52489138 | 2 | 0.960369773313132 |
| chr3-103679509-103680010 | 3 | 0.981749440628758 |
| chr11-36090293-36090794 | 3 | 0.930483487107782 |
| chr6-58655016-58655517 | 3 | 0.996589601313221 |
| chr5-113270184-113270685 | 6 | 0.828974585768718 |
| chr16-26537327-26537828 | 5 | 0.981721081002819 |
| chr2-42982088-42982589 | 8 | 0.976464671599032 |
| chr11-70770294-70770795 | 8 | 0.993493681128311 |
| chr19-39854645-39855146 | 8 | 0.935564860105566 |
| chr5-47841481-47841982 | 8 | 0.99520821769422 |
| chr4-50096529-50097030 | 3 | 0.907985629164658 |
| chr4-26135678-26136179 | 8 | 0.985263889889432 |
| chrX-68425650-68426151 | 3 | 0.976263648390382 |
| chr1-199441665-199442166 | 5 | 0.970414375567071 |
| chr1-223725017-223725518 | 4 | 0.89360587646182 |
| chr12-35608999-35609500 | 5 | 0.975237148379913 |
| chr10-46736267-46736768 | 8 | 0.848298568867226 |
| chr14-73047824-73048325 | 3 | 0.942136302576398 |
| chrX-31518341-31518842 | 6 | 0.920559053793146 |
| chr3-121057640-121058141 | 5 | 0.948555517692537 |
| chr4-170691922-170692423 | 5 | 0.89366384423957 |
| chr11-40250085-40250586 | 5 | 0.956092483044921 |
| chr1-235909194-235909695 | 8 | 0.994201802219467 |
| chr5-158916498-158916999 | 2 | 0.953933557086916 |
| chr1-177224675-177225176 | 5 | 0.891773546749804 |
| chr17-21737962-21738463 | 3 | 0.836105030497619 |
| chr16-47369941-47370442 | 8 | 0.870943372832862 |
| chrX-31111050-31111551 | 8 | 0.993493681128311 |
| chr5-129100768-129101269 | 3 | 0.878759110762631 |
| chr8-94474412-94474913 | 8 | 0.99242979423574 |
| chr4-86132568-86133069 | 8 | 0.995244782978491 |
| chr14-83125890-83126391 | 5 | 0.978540066224443 |
| chr1-41622937-41623438 | 8 | 0.991164958723423 |
| chr13-95546567-95547068 | 8 | 0.880812654236794 |
| chr10-54032302-54032803 | 2 | 0.844911787382016 |
| chr9-83737275-83737776 | 8 | 0.959942201400442 |

| | | |
|---|---|---|
| chr3-78649587-78650088 | 7 | 0.831832582539105 |
| chr9-59716529-59717030 | 5 | 0.975329173709019 |
| chr4-140980605-140981106 | 2 | 0.935161583376243 |
| chr4-12766638-12767139 | 6 | 0.98524965674539 |
| chr18-69212707-69213208 | 6 | 0.817721286745685 |
| chr14-94006067-94006568 | 8 | 0.955772203000745 |
| chr4-116303249-116303750 | 8 | 0.836835016990249 |
| chr9-19817196-19817697 | 6 | 0.989710700815939 |
| chr8-37273444-37273945 | 5 | 0.980988529500668 |
| chr6-47463653-47464154 | 8 | 0.928747703424343 |
| chr8-45097341-45097842 | 8 | 0.993493681128311 |
| chr14-65804636-65805137 | 2 | 0.854220144976588 |
| chr3-121485991-121486492 | 3 | 0.997664227550862 |
| chr7-37477397-37477898 | 8 | 0.993493681128311 |
| chr20-42041420-42041921 | 7 | 0.912457403183189 |
| chr2-227702177-227702678 | 1 | 0.904995965993696 |
| chr4-56978199-56978700 | 6 | 0.988695906638745 |
| chr5-165471749-165472250 | 7 | 0.912296919452422 |
| chr10-33927479-33927980 | 7 | 0.913010647074667 |
| chr13-6316453-6316954 | 3 | 0.998901306102065 |
| chr19-19328961-19329462 | 5 | 0.98250023838507 |
| chr9-59747086-59747587 | 4 | 0.884244436330554 |
| chr1-151693304-151693805 | 8 | 0.929940927841883 |
| chr7-67149961-67150462 | 3 | 0.99593553545041 |
| chr3-85534713-85535214 | 6 | 0.980136946230959 |
| chr4-164182063-164182564 | 8 | 0.988210216715093 |
| chr2-207282097-207282598 | 4 | 0.938446698401951 |
| chr18-43198901-43199402 | 8 | 0.993493681128311 |
| chr9-92188560-92189061 | 8 | 0.918419497636821 |
| chr1-107559516-107560017 | 3 | 0.836720743812622 |
| chr5-36440292-36440793 | 3 | 0.99401063161011 |
| chr5-64007186-64007687 | 3 | 0.99705637889073 |
| chr20-27612627-27613128 | 8 | 0.913744709168621 |
| chr18-6755111-6755612 | 8 | 0.985579316174141 |
| chr2-13926429-13926930 | 5 | 0.983718221406844 |
| chr12-37148892-37149393 | 8 | 0.96734637201748 |
| chr8-7289517-7290018 | 8 | 0.924960479481055 |
| chr2-41488202-41488703 | 8 | 0.93133616522551 |
| chr13-89168281-89168782 | 5 | 0.850429593560908 |
| chr14-101811549-101812050 | 8 | 0.837304440977087 |
| chr11-27722756-27723257 | 8 | 0.979223534140656 |
| chr14-97656734-97657235 | 3 | 0.996991138826209 |
| chr4-13697520-13698021 | 2 | 0.862263801672541 |

| | | |
|---|---|---|
| chr19-19585969-19586470 | 8 | 0.961591597446176 |
| chr13-47242424-47242925 | 8 | 0.944758186645866 |
| chr14-8361207-8361708 | 6 | 0.961001721078816 |
| chr13-82812937-82813438 | 5 | 0.990373269042194 |
| chr8-97191861-97192362 | 8 | 0.858078212214107 |
| chr10-61019794-61020295 | 2 | 0.927611120179017 |
| chr1-152496852-152497353 | 8 | 0.99518526185438 |
| chr1-94790410-94790911 | 2 | 0.952116867475702 |
| chr16-60026537-60027038 | 3 | 0.984256415046441 |
| chr12-405785-406286 | 7 | 0.969118955596412 |
| chr17-79676687-79677188 | 6 | 0.964412940224746 |
| chr14-47287930-47288431 | 8 | 0.981332703411473 |
| chr3-152158385-152158886 | 8 | 0.991078280605019 |
| chr14-13664640-13665141 | 8 | 0.904565629490125 |
| chr2-62687611-62688112 | 6 | 0.987526745992313 |
| chr4-140070511-140071012 | 8 | 0.993493681128311 |
| chr13-102352136-102352637 | 6 | 0.991074666990715 |
| chr5-101417991-101418492 | 2 | 0.960585565423293 |
| chr10-86692352-86692853 | 6 | 0.992145906968967 |
| chr7-93552849-93553350 | 5 | 0.900795768617327 |
| chr1-34116745-34117246 | 8 | 0.825773613016658 |
| chr1-56755346-56755847 | 8 | 0.993493681128311 |
| chr1-151841174-151841675 | 8 | 0.900295664028436 |
| chr9-95000999-95001500 | 6 | 0.907379203072324 |
| chr4-105940023-105940524 | 3 | 0.996859836392847 |
| chr4-68020916-68021417 | 8 | 0.821455253264633 |
| chr2-233167425-233167926 | 3 | 0.890404070678626 |
| chr4-97002012-97002513 | 3 | 0.99593553545041 |
| chr18-64964088-64964589 | 6 | 0.989945442766698 |
| chr4-30421743-30422244 | 8 | 0.88161304408604 |
| chr11-7380374-7380875 | 3 | 0.991913739680437 |
| chr6-125180304-125180805 | 5 | 0.903156552334066 |
| chr15-81898454-81898955 | 6 | 0.989592427599414 |
| chr20-29296203-29296704 | 5 | 0.860077699117071 |
| chr6-118350012-118350513 | 3 | 0.906369242010349 |
| chr1-242257081-242257582 | 8 | 0.993493681128311 |
| chr1-82679473-82679974 | 8 | 0.825201036748058 |
| chr6-138421092-138421593 | 7 | 0.939665268034567 |
| chr1-253287194-253287695 | 8 | 0.977462969086308 |
| chr4-83659368-83659869 | 8 | 0.927878502828878 |
| chr1-123808915-123809416 | 8 | 0.993493681128311 |
| chr4-23012896-23013397 | 8 | 0.993493681128311 |
| chr4-28534139-28534640 | 3 | 0.995271137668413 |

| | | |
|---|---|---|
| chr19-49887120-49887621 | 8 | 0.988618184226006 |
| chr6-100300650-100301151 | 3 | 0.996328592261637 |
| chr1-167061112-167061613 | 3 | 0.971620869823306 |
| chr2-218282742-218283243 | 3 | 0.98049861809742 |
| chr18-30431056-30431557 | 1 | 0.869000080973554 |
| chr16-46524804-46525305 | 2 | 0.863787740906045 |
| chr11-77703772-77704273 | 3 | 0.845617969056777 |
| chr14-34847088-34847589 | 6 | 0.913485484768749 |
| chr18-52353308-52353809 | 3 | 0.964022659111666 |
| chr3-22594771-22595272 | 6 | 0.948788330866251 |
| chr17-61224804-61225305 | 8 | 0.914790653685366 |
| chr4-142310260-142310761 | 3 | 0.99593553545041 |
| chr1-90420786-90421287 | 8 | 0.992963903001503 |
| chr9-86349015-86349516 | 3 | 0.998575172634325 |
| chrX-25056900-25057401 | 5 | 0.889388840189785 |
| chr2-71168497-71168998 | 8 | 0.881270659480799 |
| chr12-39149301-39149802 | 5 | 0.972712285974684 |
| chr14-82336315-82336816 | 8 | 0.986156300591078 |
| chr14-75835057-75835558 | 6 | 0.897437937164601 |
| chr17-74933658-74934159 | 4 | 0.850269652180112 |
| chr10-86042052-86042553 | 8 | 0.963512946956433 |
| chr17-64977903-64978404 | 8 | 0.864821291482569 |
| chr4-116105309-116105810 | 8 | 0.914337953593745 |
| chr12-35869795-35870296 | 6 | 0.863689826180815 |
| chr10-13366270-13366771 | 2 | 0.968702451992612 |
| chr2-118036640-118037141 | 5 | 0.92759301585079 |
| chrX-9515499-9516000 | 8 | 0.865977286631938 |
| chr14-16631091-16631592 | 3 | 0.99593553545041 |
| chr4-56528670-56529171 | 3 | 0.99593553545041 |
| chr16-6628760-6629261 | 2 | 0.911457220134608 |
| chr2-235499976-235500477 | 8 | 0.933117953509613 |
| chr14-70728082-70728583 | 5 | 0.974490211078165 |
| chr7-43771971-43772472 | 4 | 0.824012815381021 |
| chr3-150771143-150771644 | 2 | 0.81590200000948 |
| chr9-14998776-14999277 | 8 | 0.962641791171441 |
| chr4-19122030-19122531 | 3 | 0.841151247134493 |
| chr3-122754964-122755465 | 1 | 0.853142669437081 |
| chr2-203558386-203558887 | 1 | 0.867639377885524 |
| chr18-74918120-74918621 | 8 | 0.982073101702648 |
| chr9-41281280-41281781 | 8 | 0.911538051166401 |
| chr2-142027730-142028231 | 6 | 0.891202025032542 |
| chr4-178523416-178523917 | 7 | 0.927107499194303 |
| chr14-58132203-58132704 | 3 | 0.987943849917384 |

| | | |
|---|---|---|
| chr4-175727970-175728471 | 6 | 0.993225962923485 |
| chr10-72940480-72940981 | 4 | 0.878529108226389 |
| chr18-71492411-71492912 | 8 | 0.84179364936356 |
| chr20-34985413-34985914 | 3 | 0.996869240692451 |
| chr17-58854034-58854535 | 3 | 0.976500675492917 |
| chr3-37251887-37252388 | 8 | 0.927543932212196 |
| chr4-58750004-58750505 | 6 | 0.825228870176599 |
| chr3-136243846-136244347 | 3 | 0.983475580028398 |
| chr12-8366716-8367217 | 8 | 0.952117072006503 |
| chr10-31641920-31642421 | 4 | 0.850115129254745 |
| chr16-62952871-62953372 | 6 | 0.924256966546836 |
| chr18-81928713-81929214 | 3 | 0.99593553545041 |
| chr11-50298645-50299146 | 6 | 0.879263154137809 |
| chr12-4536130-4536631 | 8 | 0.868540451223297 |
| chr3-132527312-132527813 | 8 | 0.992235378173467 |
| chr10-56900257-56900758 | 5 | 0.92486306528951 |
| chr3-22270589-22271090 | 4 | 0.974397680384562 |
| chr14-75829020-75829521 | 5 | 0.968867739054725 |
| chr1-134524457-134524958 | 1 | 0.880308001558652 |
| chr14-66396923-66397424 | 7 | 0.829833637253703 |
| chr10-75876541-75877042 | 3 | 0.98126011048011 |
| chr1-252934111-252934612 | 4 | 0.816519358771234 |
| chr11-60377465-60377966 | 8 | 0.894104622534037 |
| chr3-108809353-108809854 | 2 | 0.922431948758216 |
| chr1-163422483-163422984 | 4 | 0.922038760717033 |
| chr1-52823730-52824231 | 6 | 0.991206925481043 |
| chr4-115046027-115046528 | 2 | 0.949941224549928 |
| chr9-100123221-100123722 | 3 | 0.990415142852872 |
| chr9-43296107-43296608 | 4 | 0.833091438861108 |
| chr8-92666592-92667093 | 5 | 0.9495319085439 |
| chr2-24104916-24105417 | 8 | 0.993493681128311 |
| chr5-68521115-68521616 | 8 | 0.982917483858792 |
| chr5-129271893-129272394 | 2 | 0.919970740007596 |
| chr11-76534954-76535455 | 8 | 0.826658407410547 |
| chr4-128194167-128194668 | 1 | 0.93472032694512 |
| chr2-200886058-200886559 | 3 | 0.996878158827239 |
| chr8-95335306-95335807 | 3 | 0.848378553237498 |
| chr2-80387148-80387649 | 6 | 0.991206925481043 |
| chr2-117358244-117358745 | 2 | 0.886910073198554 |
| chr6-118338669-118339170 | 8 | 0.991554619999352 |
| chr15-80786275-80786776 | 5 | 0.906508940670788 |
| chr3-18447611-18448112 | 2 | 0.924794183772762 |
| chr1-189741844-189742345 | 2 | 0.915685302350691 |

| | | |
|---|---|---|
| chr7-48853320-48853821 | 3 | 0.996832954026632 |
| chr16-47822361-47822862 | 6 | 0.926401829535873 |
| chr2-144964236-144964737 | 3 | 0.996771771137919 |
| chr5-124913656-124914157 | 3 | 0.942408766740296 |
| chr3-107483888-107484389 | 8 | 0.916213941966608 |
| chr8-60188243-60188744 | 5 | 0.98141486234525 |
| chr8-55042129-55042630 | 5 | 0.925430435088137 |
| chr6-14511440-14511941 | 8 | 0.922007125758166 |
| chr12-42325421-42325922 | 8 | 0.892265749996574 |
| chr1-17272589-17273090 | 1 | 0.814987088473854 |
| chr4-134828631-134829132 | 3 | 0.996027781124953 |
| chr15-24974562-24975063 | 4 | 0.852860912888412 |
| chr2-76638493-76638994 | 3 | 0.99593553545041 |
| chr19-36339767-36340268 | 4 | 0.879327601674989 |
| chr1-92151598-92152099 | 3 | 0.97546696058699 |
| chr20-25436582-25437083 | 5 | 0.866770461664785 |
| chr3-104706655-104707156 | 8 | 0.967874741490527 |
| chr1-232324603-232325104 | 8 | 0.993493681128311 |
| chr5-165156119-165156620 | 3 | 0.956155011024844 |
| chr9-103973758-103974259 | 8 | 0.911174545166564 |
| chrX-118031740-118032241 | 8 | 0.956504074306875 |
| chr9-98830599-98831100 | 3 | 0.99593553545041 |
| chr5-141504363-141504864 | 7 | 0.938725751881294 |
| chr1-73576422-73576923 | 5 | 0.982392756212391 |
| chr1-234639479-234639980 | 8 | 0.951264210174657 |
| chr18-71419669-71420170 | 2 | 0.90976756691966 |
| chr19-40305314-40305815 | 5 | 0.990258229749083 |
| chr1-77410496-77410997 | 5 | 0.881368371863794 |
| chr13-65327008-65327509 | 2 | 0.844546006912153 |
| chr9-104199760-104200261 | 2 | 0.861937317787204 |
| chr6-83468971-83469472 | 5 | 0.963714649565262 |
| chr8-13547765-13548266 | 6 | 0.992409593253672 |
| chr13-35956102-35956603 | 6 | 0.987312708760269 |
| chr1-173827904-173828405 | 3 | 0.995279966849318 |
| chr5-32070915-32071416 | 8 | 0.943983884933061 |
| chr2-4073983-4074484 | 3 | 0.997328621774744 |
| chr1-89512911-89513412 | 8 | 0.99523383929113 |
| chr14-80840491-80840992 | 5 | 0.986570849657493 |
| chr2-61953723-61954224 | 8 | 0.925548523791243 |
| chr7-86982281-86982782 | 8 | 0.849937097924704 |
| chr2-102068994-102069495 | 2 | 0.951113013627785 |
| chr14-78017541-78018042 | 4 | 0.956507373584802 |
| chr6-56731348-56731849 | 1 | 0.873842431744383 |

| | | |
|---|---|---|
| chr7-28127996-28128497 | 3 | 0.995865250790078 |
| chr4-39842329-39842830 | 5 | 0.983919978840557 |
| chr14-19059334-19059835 | 8 | 0.988725640419776 |
| chr15-92115379-92115880 | 7 | 0.932424005909668 |
| chr12-33326863-33327364 | 8 | 0.933784778321583 |
| chr12-20266313-20266814 | 4 | 0.820656616168253 |
| chr3-76743832-76744333 | 4 | 0.857152593606624 |
| chr2-50456309-50456810 | 6 | 0.914499181225182 |
| chr8-27320643-27321144 | 3 | 0.996442689413338 |
| chr7-131914654-131915155 | 4 | 0.972813199474857 |
| chr2-225626130-225626631 | 3 | 0.9959767213503 |
| chr3-137694204-137694705 | 3 | 0.988012728491063 |
| chr14-20954102-20954603 | 3 | 0.995839425040498 |
| chr19-35810949-35811450 | 3 | 0.995245258268197 |
| chr6-33883495-33883996 | 8 | 0.989036236354329 |
| chr4-66630232-66630733 | 6 | 0.992913304024657 |
| chr3-132427841-132428342 | 6 | 0.831503914770833 |
| chr2-110927444-110927945 | 6 | 0.875227317570826 |
| chr8-113813423-113813924 | 6 | 0.992477429544254 |
| chr16-61456496-61456997 | 8 | 0.822146958147056 |
| chrX-48196820-48197321 | 8 | 0.985509828187044 |
| chr15-76507573-76508074 | 8 | 0.975824754212466 |
| chr10-80922894-80923395 | 2 | 0.869778959412877 |
| chr5-43321999-43322500 | 8 | 0.97088785852536 |
| chr6-111514687-111515188 | 3 | 0.99593553545041 |
| chrX-2324762-2325263 | 8 | 0.986273112077613 |
| chr6-140933404-140933905 | 5 | 0.990326284599113 |
| chr16-59942417-59942918 | 5 | 0.944800740350309 |
| chr7-94945865-94946366 | 4 | 0.934330026239665 |
| chr4-63998094-63998595 | 6 | 0.991488853289354 |
| chr1-101387101-101387602 | 8 | 0.994975625476894 |
| chr10-96628667-96629168 | 3 | 0.99593553545041 |
| chr5-69475522-69476023 | 1 | 0.946194638709091 |
| chr3-92585880-92586381 | 5 | 0.909806301471016 |
| chr18-69738630-69739131 | 3 | 0.99593553545041 |
| chr9-50238281-50238782 | 8 | 0.985054418272636 |
| chr15-41635085-41635586 | 6 | 0.993222331196379 |
| chr1-104340958-104341459 | 3 | 0.97382168207132 |
| chr16-7402818-7403319 | 8 | 0.961282658033518 |
| chr2-224838805-224839306 | 8 | 0.835063005640622 |
| chr1-29075338-29075839 | 3 | 0.996817530881845 |
| chr1-7140856-7141357 | 2 | 0.889009041749134 |
| chr16-2801511-2802012 | 2 | 0.84311870283448 |

| | | |
|---|---|---|
| chr15-11624351-11624852 | 3 | 0.99251594839107 |
| chr3-147516667-147517168 | 4 | 0.928864696966375 |
| chr9-41993665-41994166 | 7 | 0.904763669345744 |
| chr14-82842210-82842711 | 8 | 0.977538930235284 |
| chr13-34455692-34456193 | 8 | 0.967240147950808 |
| chr19-24607815-24608316 | 3 | 0.996708337174514 |
| chr13-63947351-63947852 | 8 | 0.958038757366002 |
| chr14-13756527-13757028 | 6 | 0.967007707572384 |
| chr8-5723785-5724286 | 3 | 0.967058 |
| chr9-64430907-64431408 | 5 | 0.974182441727273 |
| chr19-17433490-17433991 | 1 | 0.854393616959362 |
| chr18-12103411-12103912 | 7 | 0.911291319867377 |
| chr8-50795620-50796121 | 4 | 0.965936184367867 |
| chr8-5423099-5423600 | 3 | 0.996760226644761 |
| chr9-43210282-43210783 | 8 | 0.844612290982125 |
| chr2-185811811-185812312 | 8 | 0.983673087151712 |
| chr15-32151975-32152476 | 8 | 0.970794866150171 |
| chr7-23257682-23258183 | 8 | 0.978573096024082 |
| chr7-59044108-59044609 | 8 | 0.842705944698723 |
| chr6-111224770-111225271 | 3 | 0.991940367329203 |
| chr4-40391938-40392439 | 3 | 0.949280864825625 |
| chr20-34123900-34124401 | 3 | 0.996708369665338 |
| chr1-170914125-170914626 | 8 | 0.962681426870035 |
| chr1-177450022-177450523 | 3 | 0.99407137180882 |
| chr4-76019888-76020389 | 8 | 0.831446442547177 |
| chr2-167422879-167423380 | 6 | 0.98994394192052 |
| chr4-45224020-45224521 | 8 | 0.99515875626336 |
| chr2-76973138-76973639 | 3 | 0.984670720847708 |
| chr2-114514159-114514660 | 3 | 0.877215950665732 |
| chr10-106207162-106207663 | 8 | 0.962318529859108 |
| chr10-53545070-53545571 | 8 | 0.817203738385576 |
| chr1-204884915-204885416 | 6 | 0.991206925481043 |
| chr2-239758106-239758607 | 3 | 0.989663975161409 |
| chr5-115232369-115232870 | 6 | 0.993074742668329 |
| chr15-26655825-26656326 | 3 | 0.922714430221401 |
| chr14-24663306-24663807 | 3 | 0.99593553545041 |
| chr20-48090019-48090520 | 2 | 0.816020914333317 |
| chr2-210646679-210647180 | 7 | 0.835218297467132 |
| chr17-80494267-80494768 | 8 | 0.951997805413392 |
| chr2-45676795-45677296 | 5 | 0.989377435529891 |
| chr14-16359059-16359560 | 6 | 0.827645609333637 |
| chr14-8674210-8674711 | 3 | 0.992774028288213 |
| chr17-78833249-78833750 | 8 | 0.992463081251146 |

| | | |
|---|---|---|
| chr1-110110819-110111320 | 3 | 0.99611681710662 |
| chr9-28083095-28083596 | 4 | 0.959133663009545 |
| chr13-83909281-83909782 | 8 | 0.878078815190689 |
| chr10-74909322-74909823 | 5 | 0.983785020576424 |
| chr2-34757035-34757536 | 8 | 0.845630289461026 |
| chr4-33920531-33921032 | 8 | 0.850705798531257 |
| chr18-45165375-45165876 | 8 | 0.879051119150292 |
| chr18-34113080-34113581 | 2 | 0.866959638913679 |
| chr1-255852812-255853313 | 5 | 0.923622944249582 |
| chr13-70842517-70843018 | 6 | 0.985120730442661 |
| chr6-87464856-87465357 | 7 | 0.832162407412896 |
| chr18-66262618-66263119 | 3 | 0.98743371158213 |
| chr15-35396371-35396872 | 8 | 0.886263731634658 |
| chr10-5526093-5526594 | 8 | 0.988112217825314 |
| chrX-9126578-9127079 | 4 | 0.825071713244564 |
| chr17-42411275-42411776 | 6 | 0.989203483164015 |
| chrY-5557542-5558043 | 3 | 0.948256331244354 |
| chr3-93520712-93521213 | 4 | 0.965664354630387 |
| chr1-5595994-5596495 | 5 | 0.952482471413456 |
| chr2-224837714-224838215 | 3 | 0.985265542606799 |
| chr11-51096833-51097334 | 8 | 0.916357440843089 |
| chr14-6367656-6368157 | 3 | 0.993149481354719 |
| chr15-5108905-5109406 | 8 | 0.855155396676831 |
| chr3-153207380-153207881 | 8 | 0.839636990589502 |
| chr4-181460992-181461493 | 8 | 0.993493681128311 |
| chrX-131467092-131467593 | 6 | 0.945985491630311 |
| chr2-33385118-33385619 | 5 | 0.971865867251359 |
| chr5-105920850-105921351 | 8 | 0.989572174914858 |
| chr8-84992760-84993261 | 8 | 0.993831144794386 |
| chr17-76813785-76814286 | 8 | 0.992088962664559 |
| chr14-27388584-27389085 | 8 | 0.993493681128311 |
| chr9-57187264-57187765 | 8 | 0.879088199950357 |
| chr2-204592931-204593432 | 2 | 0.827432802728374 |
| chr14-81833405-81833906 | 2 | 0.900569389950279 |
| chr5-100025653-100026154 | 8 | 0.863685579467975 |
| chr14-80323841-80324342 | 8 | 0.994834852291933 |
| chr20-12281504-12282005 | 8 | 0.938810409589141 |
| chr3-108247909-108248410 | 2 | 0.872637502328872 |
| chr13-103644211-103644712 | 8 | 0.992992884059479 |
| chr4-127142892-127143393 | 8 | 0.963756168872909 |
| chr4-173402501-173403002 | 8 | 0.841296420545481 |
| chr4-14146028-14146529 | 7 | 0.961957522051897 |
| chr20-15556792-15557293 | 3 | 0.99593553545041 |

| | | |
|---|---|---|
| chr18-61166310-61166811 | 8 | 0.991649314460623 |
| chr8-44405661-44406162 | 4 | 0.950169042099472 |
| chr7-31720085-31720586 | 6 | 0.902867822124995 |
| chr3-107639044-107639545 | 3 | 0.97401819302982 |
| chr11-17527533-17528034 | 3 | 0.996249270879472 |
| chr1-147326334-147326835 | 6 | 0.993219070775656 |
| chr4-172457445-172457946 | 3 | 0.99593553545041 |
| chr7-31835180-31835681 | 8 | 0.993493681128311 |
| chr4-168451795-168452296 | 8 | 0.892389561544852 |
| chr5-129821324-129821825 | 4 | 0.95421728585015 |
| chr15-23518770-23519271 | 5 | 0.983919978840557 |
| chr5-33831008-33831509 | 6 | 0.973310436582248 |
| chr2-40066858-40067359 | 8 | 0.918665319823741 |
| chr3-142634693-142635194 | 3 | 0.931493542992963 |
| chr10-49884043-49884544 | 8 | 0.98416622990092 |
| chr2-73143986-73144487 | 3 | 0.983327469213813 |
| chr16-14804512-14805013 | 6 | 0.963996536030786 |
| chr14-89733146-89733647 | 6 | 0.858255340810361 |
| chr7-74338282-74338783 | 2 | 0.94107372815497 |
| chr9-100295733-100296234 | 3 | 0.957533459530723 |
| chr20-20284884-20285385 | 8 | 0.962335355767114 |
| chr10-42802710-42803211 | 3 | 0.99593553545041 |
| chr9-99111883-99112384 | 8 | 0.982771148048923 |
| chr13-103773598-103774099 | 8 | 0.978737547829736 |
| chr2-24286576-24287077 | 8 | 0.984838898884685 |
| chr5-118178689-118179190 | 8 | 0.936054716252683 |
| chr19-39641571-39642072 | 3 | 0.945881233129705 |
| chr1-244571505-244572006 | 2 | 0.835089996449832 |
| chr2-184426784-184427285 | 8 | 0.857610445723288 |
| chr10-94369470-94369971 | 4 | 0.926168797577005 |
| chr13-44037139-44037640 | 2 | 0.957501139983152 |
| chr13-86496622-86497123 | 8 | 0.950718893513354 |
| chr2-225269996-225270497 | 1 | 0.89324678378216 |
| chr10-19950189-19950690 | 8 | 0.990664961127536 |
| chr1-99367605-99368106 | 6 | 0.997398992854999 |
| chr17-28444453-28444954 | 4 | 0.88857284594955 |
| chr3-161490020-161490521 | 8 | 0.874086566089641 |
| chr17-63324540-63325041 | 3 | 0.987084602757637 |
| chr2-205740302-205740803 | 5 | 0.979915026896803 |
| chr5-10680552-10681053 | 3 | 0.99593553545041 |
| chr2-84512238-84512739 | 3 | 0.998189105034073 |
| chr5-148137789-148138290 | 8 | 0.858615341478777 |
| chr17-30191924-30192425 | 8 | 0.83016590526279 |

| | | |
|---|---|---|
| chr3-72559438-72559939 | 5 | 0.983919978840557 |
| chr6-7502423-7502924 | 3 | 0.919392218279432 |
| chr1-3975261-3975762 | 3 | 0.993523997660956 |
| chr1-223098593-223099094 | 8 | 0.91802272768964 |
| chr2-119634907-119635408 | 7 | 0.899837311722821 |
| chr8-50637358-50637859 | 3 | 0.967810915623842 |
| chr16-20748228-20748729 | 5 | 0.923095017249676 |
| chr7-73947945-73948446 | 2 | 0.837651963169032 |
| chr4-18412559-18413060 | 5 | 0.970344723506381 |
| chr8-106495345-106495846 | 6 | 0.976730275811268 |
| chr3-157553469-157553970 | 8 | 0.881421564981819 |
| chr8-68879375-68879876 | 1 | 0.911740561445621 |
| chr3-63400647-63401148 | 5 | 0.986603199837457 |
| chr13-92444684-92445185 | 8 | 0.948574525526678 |
| chr14-12926571-12927072 | 3 | 0.99593553545041 |
| chr4-92811113-92811614 | 3 | 0.967151025820075 |
| chr8-116318635-116319136 | 4 | 0.959059038493512 |
| chr19-53757719-53758220 | 5 | 0.995931744719597 |
| chr6-104451892-104452393 | 1 | 0.884834280528109 |
| chr20-10358317-10358818 | 5 | 0.986159204577693 |
| chr5-100698430-100698931 | 8 | 0.8370669659256 |
| chr20-26071609-26072110 | 8 | 0.821812354366322 |
| chr5-945730-946231 | 6 | 0.991206925481043 |
| chr20-36949503-36950004 | 2 | 0.899856221962646 |
| chr13-40233447-40233948 | 5 | 0.988441319770842 |
| chr2-161713892-161714393 | 8 | 0.943952448584214 |
| chr2-117229230-117229731 | 3 | 0.996872314529127 |
| chr2-117191794-117192295 | 2 | 0.947485057021522 |
| chr3-77114253-77114754 | 8 | 0.8756782695644 |
| chr6-22356399-22356900 | 3 | 0.988094531188916 |
| chr19-53759646-53760147 | 8 | 0.969617251945193 |
| chr1-193851196-193851697 | 6 | 0.935544751950963 |
| chr14-42729521-42730022 | 8 | 0.977560356959369 |
| chr17-8664721-8665222 | 5 | 0.91721651743085 |
| chr18-67599454-67599955 | 5 | 0.988367711693253 |
| chr4-109664879-109665380 | 2 | 0.95791608429324 |
| chr12-9960993-9961494 | 5 | 0.965386458987166 |
| chr2-58582997-58583498 | 3 | 0.936989725834216 |
| chrX-32617621-32618122 | 4 | 0.845727586611458 |
| chr19-46708386-46708887 | 6 | 0.977532726997371 |
| chr3-19274617-19275118 | 2 | 0.951689687022798 |
| chr1-142796344-142796845 | 8 | 0.820762147218797 |
| chr18-55252772-55253273 | 5 | 0.950152933949976 |

| | | |
|---|---|---|
| chr10-31642593-31643094 | 6 | 0.993165040143682 |
| chr7-97067391-97067892 | 3 | 0.996518820654679 |
| chr2-169476866-169477367 | 5 | 0.963466174329492 |
| chr3-57290889-57291390 | 4 | 0.856409850461739 |
| chr14-21195033-21195534 | 5 | 0.916756825075389 |
| chr17-21067254-21067755 | 5 | 0.985972909125294 |
| chr7-53752223-53752724 | 2 | 0.824991880399149 |
| chr13-64327426-64327927 | 8 | 0.937623627435629 |
| chr18-67069049-67069550 | 2 | 0.944040320247596 |
| chr14-86082714-86083215 | 5 | 0.991111064607434 |
| chr4-22914393-22914894 | 3 | 0.946637238946441 |
| chr10-8208524-8209025 | 2 | 0.835080806586502 |
| chr1-88929067-88929568 | 8 | 0.995154516316724 |
| chr18-23855555-23856056 | 8 | 0.912733130224423 |
| chr5-102390173-102390674 | 8 | 0.888171511544962 |
| chr4-35975286-35975787 | 8 | 0.983748208917374 |
| chrX-26618864-26619365 | 5 | 0.983919978840557 |
| chr18-77213489-77213990 | 8 | 0.847888814934631 |
| chr1-231052752-231053253 | 2 | 0.842025016191828 |
| chr12-28377147-28377648 | 5 | 0.885495751296863 |
| chr2-338969-339470 | 8 | 0.993493681128311 |
| chr20-21915782-21916283 | 3 | 0.956022870521772 |
| chr1-99964128-99964629 | 6 | 0.97117244508496 |
| chr3-56338434-56338935 | 8 | 0.993493681128311 |
| chr1-178030594-178031095 | 6 | 0.932636512133259 |
| chr1-99470810-99471311 | 1 | 0.921359595278687 |
| chr5-58460055-58460556 | 8 | 0.923647018129634 |
| chr7-37976810-37977311 | 5 | 0.953277809698845 |
| chr4-43523266-43523767 | 8 | 0.869325740068382 |
| chr9-16778409-16778910 | 6 | 0.991206925481043 |
| chr4-126461675-126462176 | 1 | 0.960859718582327 |
| chr11-65591807-65592308 | 8 | 0.979920632670985 |
| chr15-40110503-40111004 | 3 | 0.948384365268616 |
| chr19-49433924-49434425 | 6 | 0.914656989240298 |
| chr3-142260035-142260536 | 8 | 0.958334190356264 |
| chr14-36309351-36309852 | 3 | 0.973243623810665 |
| chr14-81641939-81642440 | 6 | 0.834506565035586 |
| chr10-4108895-4109396 | 8 | 0.851214945040913 |
| chr5-83855494-83855995 | 3 | 0.996876323793869 |
| chrX-131217204-131217705 | 8 | 0.860038616936757 |
| chr6-97744622-97745123 | 3 | 0.996626977613015 |
| chr13-43232454-43232955 | 6 | 0.942779328294671 |
| chr6-40239412-40239913 | 7 | 0.820506731543092 |

| | | |
|---|---|---|
| chr4-164864376-164864877 | 3 | 0.994043522719322 |
| chr1-8577416-8577917 | 7 | 0.93210878000797 |
| chrX-103440670-103441171 | 8 | 0.991960922617798 |
| chr5-72797665-72798166 | 8 | 0.986815577690293 |
| chr2-225095733-225096234 | 6 | 0.939149847763434 |
| chr9-61937647-61938148 | 8 | 0.977820042804581 |
| chr4-97306727-97307228 | 8 | 0.822142153082056 |
| chr3-87516501-87517002 | 8 | 0.857258606830686 |
| chr8-32159109-32159610 | 5 | 0.844215699411496 |
| chr18-66339888-66340389 | 8 | 0.993368869392013 |
| chr13-99820288-99820789 | 3 | 0.960373530058288 |
| chr17-86421258-86421759 | 3 | 0.99593553545041 |
| chr3-13269432-13269933 | 8 | 0.994982183889212 |
| chr19-22682108-22682609 | 3 | 0.99593553545041 |
| chr4-150114085-150114586 | 8 | 0.993493681128311 |
| chr1-127224184-127224685 | 8 | 0.918374339572459 |
| chr4-124503840-124504341 | 2 | 0.948886610444538 |
| chr3-29769591-29770092 | 8 | 0.968915988506368 |
| chr10-38943384-38943885 | 8 | 0.98643106634677 |
| chr14-1809605-1810106 | 2 | 0.909755851350624 |
| chrX-24767573-24768074 | 5 | 0.906484798329675 |
| chr5-26749588-26750089 | 3 | 0.996858681817304 |
| chr8-82568924-82569425 | 3 | 0.857489242109809 |
| chr12-26839995-26840496 | 8 | 0.907001186996301 |
| chr20-26878321-26878822 | 3 | 0.98039440084787 |
| chr19-40812324-40812825 | 6 | 0.947676267238078 |
| chrX-20079400-20079901 | 5 | 0.962957080183948 |
| chr15-27431817-27432318 | 8 | 0.986104937459192 |
| chr10-65083799-65084300 | 2 | 0.860831924505757 |
| chr10-24232367-24232868 | 8 | 0.867339012319824 |
| chr5-135307257-135307758 | 4 | 0.890206285661115 |
| chr19-11096439-11096940 | 4 | 0.853410351112881 |
| chr2-113636855-113637356 | 8 | 0.989846442660266 |
| chr2-193321089-193321590 | 3 | 0.913629803328287 |
| chr16-60929884-60930385 | 3 | 0.91484812608079 |
| chr5-123504844-123505345 | 5 | 0.917653862121308 |
| chr7-37786438-37786939 | 3 | 0.99593553545041 |
| chr8-116480908-116481409 | 5 | 0.895601069022131 |
| chr3-9047434-9047935 | 8 | 0.995177996696644 |
| chrX-101432139-101432640 | 3 | 0.99593553545041 |
| chr4-115374974-115375475 | 8 | 0.987563439565059 |
| chr6-132462097-132462598 | 4 | 0.9132483105301 |
| chr1-182395931-182396432 | 5 | 0.991626405278659 |

| | | |
|---|---|---|
| chr18-20166892-20167393 | 3 | 0.987164242270332 |
| chr2-56316560-56317061 | 4 | 0.965359979073788 |
| chr15-2470218-2470719 | 8 | 0.85519115511846 |
| chr6-43386809-43387310 | 6 | 0.98857327839645 |
| chr12-40860964-40861465 | 8 | 0.897121932248185 |
| chr11-76465769-76466270 | 8 | 0.948430826159201 |
| chr5-119326655-119327156 | 3 | 0.99593553545041 |
| chr19-31452436-31452937 | 3 | 0.978118182313188 |
| chr6-13376325-13376826 | 2 | 0.828710745940726 |
| chr8-72234930-72235431 | 4 | 0.926216043425344 |
| chr17-15412136-15412637 | 8 | 0.976342148542666 |
| chr9-14240953-14241454 | 4 | 0.974935866672013 |
| chr19-8275447-8275948 | 8 | 0.993493681128311 |
| chr16-9536262-9536763 | 8 | 0.867975850181182 |
| chr15-31387073-31387574 | 8 | 0.970972410714851 |
| chr16-70760291-70760792 | 6 | 0.997315827561035 |
| chrX-35289808-35290309 | 3 | 0.99593553545041 |
| chr2-209769281-209769782 | 8 | 0.969636731643452 |
| chr15-95096791-95097292 | 8 | 0.901720317141182 |
| chr1-221271496-221271997 | 3 | 0.995964692379161 |
| chr19-46408681-46409182 | 1 | 0.920502389523775 |
| chr17-19705582-19706083 | 8 | 0.943501144902767 |
| chr2-54888853-54889354 | 1 | 0.909514629978738 |
| chr17-9535006-9535507 | 8 | 0.839749686397254 |
| chr4-84998866-84999367 | 3 | 0.996031820377014 |
| chrX-107297471-107297972 | 3 | 0.989265570082102 |
| chr17-60111745-60112246 | 3 | 0.974153198412219 |
| chr11-68246799-68247300 | 5 | 0.992384691321151 |
| chr11-84677091-84677592 | 3 | 0.937059152035821 |
| chr1-174270838-174271339 | 3 | 0.996361525465909 |
| chr8-3444955-3445456 | 3 | 0.990351255213131 |
| chr2-117463069-117463570 | 1 | 0.97764783444183 |
| chr1-176946513-176947014 | 7 | 0.899935534763721 |
| chr16-73279235-73279736 | 8 | 0.958254188128147 |
| chr10-87232239-87232740 | 8 | 0.930313863994518 |
| chr2-48721740-48722241 | 3 | 0.996888186104297 |
| chr11-29820959-29821460 | 8 | 0.993493681128311 |
| chr9-73976376-73976877 | 8 | 0.849348115282345 |
| chr15-37870180-37870681 | 6 | 0.992802471416283 |
| chr8-4617942-4618443 | 6 | 0.958644046928498 |
| chr7-60223003-60223504 | 6 | 0.889856451151409 |
| chr4-30606589-30607090 | 5 | 0.968307901411297 |
| chr3-61878455-61878956 | 2 | 0.936323128640049 |

| | | |
|---|---|---|
| chr8-55910742-55911243 | 7 | 0.903475805291864 |
| chr11-55493239-55493740 | 3 | 0.996351321575937 |
| chr9-31629534-31630035 | 3 | 0.99593553545041 |
| chr15-62347461-62347962 | 6 | 0.851983435465259 |
| chr6-123315772-123316273 | 3 | 0.914594145144845 |
| chr12-38369939-38370440 | 3 | 0.868118088417698 |
| chr3-79312083-79312584 | 7 | 0.957182640393105 |
| chr10-102982633-102983134 | 8 | 0.910818481124912 |
| chr11-25907924-25908425 | 5 | 0.841176949303333 |
| chr4-152930456-152930957 | 6 | 0.992407347055655 |
| chr17-72383639-72384140 | 8 | 0.990007878801905 |
| chr18-6437405-6437906 | 8 | 0.829932776971319 |
| chr2-106963228-106963729 | 6 | 0.969390777512734 |
| chr2-216019236-216019737 | 8 | 0.953215576832158 |
| chr4-143250174-143250675 | 3 | 0.997343032841785 |
| chr1-144973608-144974109 | 8 | 0.894632015342835 |
| chr1-34489134-34489635 | 2 | 0.90416688267953 |
| chr15-33728255-33728756 | 8 | 0.921863587210846 |
| chr2-14648222-14648723 | 5 | 0.983919978840557 |
| chr3-22825469-22825970 | 4 | 0.882186063929432 |
| chr1-92023086-92023587 | 3 | 0.99593553545041 |
| chr19-51522408-51522909 | 8 | 0.935053234135147 |
| chr16-77851847-77852348 | 3 | 0.987569294922334 |
| chr1-145013088-145013589 | 5 | 0.988398375951614 |
| chr10-41407195-41407696 | 1 | 0.861081178978383 |
| chr4-54039414-54039915 | 5 | 0.988732339400237 |
| chr18-6993257-6993758 | 3 | 0.986955795863855 |
| chr15-73183942-73184443 | 8 | 0.987242064220893 |
| chr15-80039552-80040053 | 6 | 0.91195715948955 |
| chr3-123558898-123559399 | 6 | 0.985620846476362 |
| chr11-13161523-13162024 | 6 | 0.984255941372768 |
| chr9-4740338-4740839 | 6 | 0.991206925481043 |
| chr8-46001057-46001558 | 5 | 0.990292648883685 |
| chr18-20802641-20803142 | 8 | 0.944882107566012 |
| chr6-130712934-130713435 | 8 | 0.842584057853706 |
| chr12-44867248-44867749 | 8 | 0.929333242373305 |
| chr11-50698310-50698811 | 8 | 0.995167936239416 |
| chr4-136755375-136755876 | 8 | 0.941412805768866 |
| chr5-103566793-103567294 | 6 | 0.983550054465426 |
| chr15-78196844-78197345 | 7 | 0.917191429572459 |
| chr18-74340314-74340815 | 4 | 0.855132071099882 |
| chr10-78525242-78525743 | 2 | 0.848067227694387 |
| chr6-136008887-136009388 | 3 | 0.996569731174574 |

| | | |
|---|---|---|
| chr15-34156166-34156667 | 7 | 0.968245835883954 |
| chr8-117699749-117700250 | 8 | 0.913697430383822 |
| chr5-84560251-84560752 | 8 | 0.995068185291067 |
| chr2-152187641-152188142 | 8 | 0.993414839133152 |
| chr10-66268476-66268977 | 4 | 0.841602563472535 |
| chr7-102704655-102705156 | 6 | 0.981460885544679 |
| chr10-89285021-89285522 | 8 | 0.854841468245081 |
| chr8-82178667-82179168 | 3 | 0.989637447818006 |
| chr9-99676905-99677406 | 3 | 0.996766900902437 |
| chr10-40735959-40736460 | 8 | 0.888663703250597 |
| chr5-20393212-20393713 | 1 | 0.878441503903929 |
| chr8-101143657-101144158 | 6 | 0.992946846260817 |
| chr20-10985964-10986465 | 7 | 0.916165694310836 |
| chr17-60467483-60467984 | 8 | 0.994704117542318 |
| chr7-128184931-128185432 | 8 | 0.993493681128311 |
| chr4-179236167-179236668 | 8 | 0.995247910447258 |
| chr4-124453000-124453501 | 6 | 0.936686845531391 |
| chr4-82843244-82843745 | 8 | 0.991082747547233 |
| chr2-134961875-134962376 | 7 | 0.923496147938285 |
| chr17-6163649-6164150 | 8 | 0.994466496904208 |
| chr19-54276986-54277487 | 4 | 0.957723200298639 |
| chr7-58035816-58036317 | 3 | 0.996039340467025 |
| chr4-118863741-118864242 | 5 | 0.991280272673494 |
| chr20-20538196-20538697 | 5 | 0.93867809989156 |
| chr3-66143070-66143571 | 8 | 0.982948594831799 |
| chr4-170396317-170396818 | 5 | 0.94244713335786 |
| chr13-42272083-42272584 | 8 | 0.920457673718591 |
| chr11-46905703-46906204 | 1 | 0.928375921160443 |
| chr13-102345115-102345616 | 7 | 0.81772229859159 |
| chr10-37308049-37308550 | 8 | 0.963379616300607 |
| chr15-47164693-47165194 | 2 | 0.917039662314276 |
| chr3-55750827-55751328 | 8 | 0.946755350420586 |
| chr20-7745705-7746206 | 8 | 0.913705473362895 |
| chr4-118460300-118460801 | 6 | 0.913927880098556 |
| chr2-197330407-197330908 | 3 | 0.996860215652949 |
| chr2-190926568-190927069 | 2 | 0.890636620894108 |
| chr7-63529088-63529589 | 5 | 0.943195056666898 |
| chr19-4174128-4174629 | 8 | 0.993493681128311 |
| chr9-27702359-27702860 | 5 | 0.890068838031756 |
| chr20-31194590-31195091 | 6 | 0.991206925481043 |
| chr11-21313251-21313752 | 8 | 0.992841624407393 |
| chr17-54267592-54268093 | 5 | 0.891306278600508 |
| chr2-234018371-234018872 | 3 | 0.992401231667888 |

| | | |
|---|---|---|
| chr4-114660676-114661177 | 4 | 0.909444781691396 |
| chr13-85433927-85434428 | 1 | 0.910778860759258 |
| chr15-36601775-36602276 | 5 | 0.957694980475461 |
| chr4-136163034-136163535 | 8 | 0.956619836686992 |
| chr3-91102134-91102635 | 8 | 0.993493681128311 |
| chr2-137377384-137377885 | 7 | 0.956988602778352 |
| chr17-73503803-73504304 | 8 | 0.995096536365639 |
| chr13-101934140-101934641 | 5 | 0.982341434969248 |
| chr12-43910969-43911470 | 8 | 0.993493681128311 |
| chr12-1662139-1662640 | 8 | 0.840137634214595 |
| chr1-143906859-143907360 | 8 | 0.890116277972189 |
| chr3-12185362-12185863 | 8 | 0.975621761447533 |
| chr16-29494635-29495136 | 2 | 0.865652167131434 |
| chr1-180483487-180483988 | 2 | 0.953772444244331 |
| chr2-192695912-192696413 | 4 | 0.965958215610747 |
| chr6-5869185-5869686 | 6 | 0.997138193881752 |
| chr2-189731797-189732298 | 3 | 0.996898394955622 |
| chr17-23601986-23602487 | 6 | 0.956016782532541 |
| chr13-449552-450053 | 7 | 0.949321021703585 |
| chr6-31335589-31336090 | 8 | 0.919242851525447 |
| chr8-8784981-8785482 | 3 | 0.973645062257353 |
| chr1-236236924-236237425 | 8 | 0.993493681128311 |
| chr1-23687048-23687549 | 3 | 0.902710021694605 |
| chr4-84975913-84976414 | 3 | 0.896859501758916 |
| chr5-117050062-117050563 | 3 | 0.982372409516314 |
| chr18-36916475-36916976 | 2 | 0.896096120257217 |
| chr16-51784531-51785032 | 8 | 0.948158516572931 |
| chr1-205003073-205003574 | 3 | 0.99593553545041 |
| chr3-22225742-22226243 | 8 | 0.94358386821758 |
| chr13-104498729-104499230 | 3 | 0.99646857483134 |
| chr2-188103292-188103793 | 3 | 0.99593553545041 |
| chr3-158640466-158640967 | 4 | 0.893141081293059 |
| chr5-128382285-128382786 | 5 | 0.849023809974351 |
| chr4-63449660-63450161 | 8 | 0.978617308913901 |
| chr2-159340067-159340568 | 6 | 0.88425293024649 |
| chr7-104885018-104885519 | 5 | 0.978229185505236 |
| chr6-140521754-140522255 | 6 | 0.992756534654393 |
| chr7-65756230-65756731 | 3 | 0.996550872137766 |
| chr10-102842399-102842900 | 3 | 0.899610448467774 |
| chr8-31304111-31304612 | 3 | 0.996524657024323 |
| chr4-44620464-44620965 | 8 | 0.832347158894147 |
| chr1-202213184-202213685 | 4 | 0.945287399354966 |
| chr18-77037128-77037629 | 7 | 0.81469377627897 |

| | | |
|---|---|---|
| chr3-105873220-105873721 | 8 | 0.934644376837411 |
| chr10-96360926-96361427 | 7 | 0.84777471917185 |
| chr8-103754103-103754604 | 8 | 0.922151427428476 |
| chr2-105802124-105802625 | 3 | 0.996316754550631 |
| chr3-158030571-158031072 | 7 | 0.871893548104716 |
| chr6-130324026-130324527 | 8 | 0.968761039129048 |
| chr1-172096257-172096758 | 6 | 0.987373698149628 |
| chr1-116719170-116719671 | 6 | 0.991206925481043 |
| chr5-86977624-86978125 | 3 | 0.856592451619648 |
| chr6-8436031-8436532 | 2 | 0.931732001676883 |
| chr8-64809495-64809996 | 5 | 0.988209935018953 |
| chr1-218651844-218652345 | 3 | 0.993154571242079 |
| chr15-4971331-4971832 | 5 | 0.980382282378868 |
| chr13-81735267-81735768 | 8 | 0.993493681128311 |
| chr20-51373534-51374035 | 6 | 0.948947590806076 |
| chr14-75002335-75002836 | 3 | 0.998934181607667 |
| chr4-6402216-6402717 | 3 | 0.990887286505522 |
| chr15-92086838-92087339 | 7 | 0.849165386510068 |
| chr8-59543727-59544228 | 4 | 0.882089127463043 |
| chr11-55473840-55474341 | 1 | 0.878833149383388 |
| chrX-63618400-63618901 | 6 | 0.992776787170018 |
| chr1-108426019-108426520 | 3 | 0.95751828531279 |
| chr1-256738659-256739160 | 8 | 0.885909927580141 |
| chr8-95692947-95693448 | 5 | 0.883220868908761 |
| chr9-47473315-47473816 | 4 | 0.963598576741752 |
| chr4-32053184-32053685 | 3 | 0.995082 |
| chr6-54364191-54364692 | 1 | 0.884047506251288 |
| chr12-10790237-10790738 | 6 | 0.99166434824821 |
| chr13-42941160-42941661 | 5 | 0.982049215093606 |
| chr4-154224246-154224747 | 8 | 0.952061285599628 |
| chr1-45463192-45463693 | 8 | 0.865146944029165 |
| chr6-138542549-138543050 | 1 | 0.954472288172882 |
| chr8-65093245-65093746 | 3 | 0.916334715412759 |
| chr13-47407367-47407868 | 4 | 0.83121106532907 |
| chr6-129295800-129296301 | 3 | 0.996561681686143 |
| chr13-1803294-1803795 | 3 | 0.993828773069016 |
| chr15-31539340-31539841 | 8 | 0.929409851981086 |
| chr11-25134385-25134886 | 3 | 0.982730534885752 |
| chr10-82107483-82107984 | 8 | 0.847964464272831 |
| chr5-134114499-134115000 | 6 | 0.933905107716443 |
| chr11-30786698-30787199 | 4 | 0.930042096973102 |
| chr1-164403035-164403536 | 2 | 0.953059432784389 |
| chr1-7291575-7292076 | 6 | 0.992866818878576 |

| | | |
|---|---|---|
| chr11-45499751-45500252 | 3 | 0.996850259457504 |
| chr3-21496647-21497148 | 8 | 0.978637461977517 |
| chr11-1363966-1364467 | 6 | 0.992310969428881 |
| chrX-38869938-38870439 | 5 | 0.983919978840557 |
| chr4-22690821-22691322 | 8 | 0.995188014254399 |
| chr4-34243407-34243908 | 8 | 0.963463699325182 |
| chr2-4499638-4500139 | 8 | 0.975392432295924 |
| chr4-15376564-15377065 | 3 | 0.974054148727137 |
| chr15-94741635-94742136 | 8 | 0.91878229900519 |
| chr18-78172116-78172617 | 5 | 0.954438727753975 |
| chr8-45771894-45772395 | 8 | 0.967208720125551 |
| chr8-121546922-121547423 | 8 | 0.871029503884647 |
| chr4-169205187-169205688 | 8 | 0.995184538485205 |
| chr2-249027856-249028357 | 5 | 0.857662625582988 |
| chr10-33125466-33125967 | 8 | 0.993493681128311 |
| chr3-87687240-87687741 | 8 | 0.821603809862131 |
| chr11-12935107-12935608 | 2 | 0.958812369814636 |
| chr7-49587157-49587658 | 6 | 0.991206925481043 |
| chr19-8162247-8162748 | 6 | 0.893209187902841 |
| chr4-181989673-181990174 | 8 | 0.912652594448636 |
| chr11-53482139-53482640 | 3 | 0.99684655008674 |
| chr2-196350781-196351282 | 4 | 0.835426805171567 |
| chr3-158841900-158842401 | 8 | 0.958527383650977 |
| chr2-233863734-233864235 | 7 | 0.816607195112635 |
| chr17-1461240-1461741 | 8 | 0.993493681128311 |
| chr2-89875374-89875875 | 3 | 0.981245492153688 |
| chr9-26765608-26766109 | 8 | 0.941934811993555 |
| chr1-44134791-44135292 | 6 | 0.967495011276736 |
| chr2-194596744-194597245 | 3 | 0.977288854989358 |
| chr3-150233929-150234430 | 3 | 0.99593553545041 |
| chr2-94509459-94509960 | 8 | 0.956455285597688 |
| chr17-59009679-59010180 | 1 | 0.830880891826973 |
| chr15-98665575-98666076 | 1 | 0.907572014160732 |
| chr2-59350681-59351182 | 2 | 0.917755994170986 |
| chr11-77566198-77566699 | 1 | 0.886289653630439 |
| chr17-79888023-79888524 | 7 | 0.903304041249255 |
| chr7-48857095-48857596 | 3 | 0.934364463959507 |
| chr16-74950465-74950966 | 8 | 0.994403199927586 |
| chr5-123226346-123226847 | 2 | 0.82537695646414 |
| chr3-10608138-10608639 | 1 | 0.819514450649026 |
| chr14-36645466-36645967 | 8 | 0.967965170794563 |
| chr2-161816761-161817262 | 3 | 0.99593553545041 |
| chr6-2547538-2548039 | 8 | 0.994625471843157 |

| | | |
|---|---|---|
| chr4-10022318-10022819 | 7 | 0.907123485805377 |
| chr3-132085092-132085593 | 6 | 0.993202181072587 |
| chr9-71895276-71895777 | 3 | 0.994664619587616 |
| chr4-61606167-61606668 | 3 | 0.981852701342383 |
| chr11-62305696-62306197 | 6 | 0.992132091744698 |
| chr13-76904607-76905108 | 3 | 0.99447344063943 |
| chr8-113098163-113098664 | 3 | 0.851941081579154 |
| chr3-132787835-132788336 | 3 | 0.950959786550806 |
| chr3-168236970-168237471 | 8 | 0.861908266659996 |
| chr4-103823409-103823910 | 3 | 0.989421142937724 |
| chr11-34622016-34622517 | 3 | 0.819117641307972 |
| chr20-434440-434941 | 3 | 0.994979867578316 |
| chr2-25738240-25738741 | 8 | 0.995169568539088 |
| chr16-28330021-28330522 | 1 | 0.889842538872464 |
| chr6-126117730-126118231 | 3 | 0.997039585098376 |
| chr1-126295647-126296148 | 8 | 0.985016606242462 |
| chr19-36152343-36152844 | 3 | 0.995655836245281 |
| chr11-54842018-54842519 | 4 | 0.910994232424125 |
| chr4-11944584-11945085 | 8 | 0.8699324823975 |
| chr19-47368125-47368626 | 5 | 0.992630843359447 |
| chr5-66035005-66035506 | 4 | 0.969739937703207 |
| chrY-1064817-1065318 | 3 | 0.939127314528304 |
| chr10-66726543-66727044 | 3 | 0.987162890930284 |
| chr5-57122455-57122956 | 3 | 0.966663164175664 |
| chr17-8952607-8953108 | 3 | 0.995762965705532 |
| chr8-38232584-38233085 | 2 | 0.927275421992298 |
| chr19-37230634-37231135 | 8 | 0.991246543519347 |
| chr13-38204796-38205297 | 5 | 0.956662150606008 |
| chr2-182689797-182690298 | 8 | 0.966175749699636 |
| chr1-216243733-216244234 | 4 | 0.866872338276908 |
| chr15-40940781-40941282 | 8 | 0.928592188847704 |
| chr11-66507461-66507962 | 3 | 0.981491527736852 |
| chr2-92883536-92884037 | 5 | 0.885356112203976 |
| chr1-39572879-39573380 | 3 | 0.99593553545041 |
| chr2-118014113-118014614 | 6 | 0.98058858162436 |
| chr1-259700002-259700503 | 8 | 0.893047138792186 |
| chr1-154687026-154687527 | 2 | 0.922612863445399 |
| chr2-54320934-54321435 | 7 | 0.903369452294153 |
| chr6-70913225-70913726 | 3 | 0.957574264999765 |
| chr14-42468021-42468522 | 6 | 0.992908571833465 |
| chr14-41042532-41043033 | 3 | 0.99593553545041 |
| chr8-75198268-75198769 | 6 | 0.991206925481043 |
| chr3-42529070-42529571 | 5 | 0.983919978840557 |

| | | |
|---|---|---|
| chr10-81120698-81121199 | 6 | 0.990731667702015 |
| chr3-146047736-146048237 | 8 | 0.979847183520601 |
| chr8-32160328-32160829 | 3 | 0.99593553545041 |
| chr9-50056555-50057056 | 4 | 0.87356550318867 |
| chr16-63384877-63385378 | 8 | 0.992680575908086 |
| chr5-26724667-26725168 | 7 | 0.905927237102375 |
| chr2-68540493-68540994 | 5 | 0.95979491028771 |
| chr6-20518983-20519484 | 7 | 0.854674727134001 |
| chr18-4000872-4001373 | 5 | 0.861939368040471 |
| chr5-97359847-97360348 | 8 | 0.878339966085566 |
| chr8-116967999-116968500 | 8 | 0.987320523266943 |
| chr17-59230216-59230717 | 8 | 0.993493681128311 |
| chr14-81742520-81743021 | 5 | 0.986323762529136 |
| chr13-77081198-77081699 | 8 | 0.958922801706502 |
| chr15-73581094-73581595 | 8 | 0.943767335097027 |
| chr13-104262920-104263421 | 5 | 0.98204669434601 |
| chr5-9413309-9413810 | 7 | 0.831980534908748 |
| chr10-53590515-53591016 | 8 | 0.955951892150739 |
| chr14-103763582-103764083 | 8 | 0.982444265637192 |
| chr4-43005205-43005706 | 6 | 0.976673184930887 |
| chr9-78665367-78665868 | 3 | 0.995106382812854 |
| chr1-75565109-75565610 | 3 | 0.820492655556227 |
| chr1-193796273-193796774 | 3 | 0.993436066154043 |
| chr14-75974610-75975111 | 5 | 0.986206276603943 |
| chr10-37317556-37318057 | 4 | 0.895925264850729 |
| chr11-77468479-77468980 | 4 | 0.922898336118045 |
| chr7-92949140-92949641 | 1 | 0.910682724013726 |
| chr19-23719464-23719965 | 3 | 0.990315549360253 |
| chr12-41208524-41209025 | 8 | 0.941348628829064 |
| chr1-215309575-215310076 | 8 | 0.865989876309339 |
| chr14-91780859-91781360 | 3 | 0.997778872618577 |
| chr11-74435507-74436008 | 8 | 0.832799226797965 |
| chr18-61578172-61578673 | 4 | 0.866847110393877 |
| chr4-68018091-68018592 | 5 | 0.986045816544085 |
| chr1-244363024-244363525 | 6 | 0.978821850635015 |
| chr11-31074090-31074591 | 8 | 0.99177751435409 |
| chr5-159820967-159821468 | 7 | 0.849935927960836 |
| chr20-46078189-46078690 | 8 | 0.938598205526579 |
| chr14-79158611-79159112 | 4 | 0.852526219351768 |
| chr7-30958295-30958796 | 3 | 0.99593553545041 |
| chr3-112265212-112265713 | 8 | 0.978290152330842 |
| chr2-33065682-33066183 | 8 | 0.886964765612388 |
| chr15-78714340-78714841 | 6 | 0.984475631299361 |

| | | |
|---|---|---|
| chr14-66067987-66068488 | 6 | 0.90213686520283 |
| chr4-118959317-118959818 | 4 | 0.924787949790901 |
| chr11-66006330-66006831 | 5 | 0.983919978840557 |
| chr1-41695589-41696090 | 5 | 0.981509747563348 |
| chr8-52133141-52133642 | 5 | 0.983919978840557 |
| chr18-50372058-50372559 | 3 | 0.959516713200421 |
| chr4-63569250-63569751 | 8 | 0.958172190607702 |
| chr18-64269523-64270024 | 2 | 0.846382068635548 |
| chr16-68061759-68062260 | 8 | 0.992673413118251 |
| chr4-93135030-93135531 | 3 | 0.981860800489687 |
| chr5-6055200-6055701 | 8 | 0.843417001865428 |
| chr4-65685738-65686239 | 6 | 0.991631364557287 |
| chr19-27615729-27616230 | 5 | 0.983919978840557 |
| chr11-37823013-37823514 | 8 | 0.824969666248783 |
| chr9-81189140-81189641 | 5 | 0.956083255076676 |
| chr20-32874460-32874961 | 2 | 0.832388530914249 |
| chr5-136460258-136460759 | 7 | 0.865369584729623 |
| chr4-8409880-8410381 | 1 | 0.91759481069812 |
| chr9-30175211-30175712 | 5 | 0.98898102220192 |
| chr10-98806733-98807234 | 2 | 0.882278664922283 |
| chr16-81002265-81002766 | 2 | 0.901220346118016 |
| chr6-41445180-41445681 | 8 | 0.995141332966185 |
| chr9-77173765-77174266 | 6 | 0.990481017701781 |
| chr19-10672194-10672695 | 3 | 0.81486486762419 |
| chr18-1997078-1997579 | 6 | 0.993172921199411 |
| chr9-63919364-63919865 | 4 | 0.834646861329649 |
| chr16-15730828-15731329 | 7 | 0.890764739987742 |
| chr13-101140639-101141140 | 2 | 0.916240141484546 |
| chr1-194607465-194607966 | 8 | 0.874445422962535 |
| chr2-151333123-151333624 | 7 | 0.959922522893759 |
| chr5-148816436-148816937 | 8 | 0.911161311685335 |
| chr2-173663784-173664285 | 8 | 0.82520370135855 |
| chr14-85756993-85757494 | 6 | 0.991206925481043 |
| chr1-149705825-149706326 | 3 | 0.905749917648993 |
| chr14-64442394-64442895 | 8 | 0.995231794611931 |
| chr5-120186866-120187367 | 7 | 0.880339888191325 |
| chr9-57656791-57657292 | 6 | 0.851957171995564 |
| chr20-7269462-7269963 | 4 | 0.821010974263983 |
| chr7-89893168-89893669 | 6 | 0.991799351121162 |
| chr13-90809511-90810012 | 8 | 0.976930170815585 |
| chr19-56436106-56436607 | 5 | 0.930971078856378 |
| chr2-150839328-150839829 | 4 | 0.962898557159108 |
| chr1-226532592-226533093 | 1 | 0.890896933926234 |

| | | |
|---|---|---|
| chr2-167445923-167446424 | 8 | 0.955896079201381 |
| chr11-23695210-23695711 | 8 | 0.933115157022599 |
| chr10-43809884-43810385 | 8 | 0.832351349990321 |
| chr4-146351955-146352456 | 2 | 0.957078477861456 |
| chr16-113886-114387 | 3 | 0.996761285423325 |
| chr2-83458238-83458739 | 8 | 0.912323967650736 |
| chr7-31199767-31200268 | 3 | 0.99593553545041 |
| chr10-70718904-70719405 | 8 | 0.956749017214357 |
| chr2-104918672-104919173 | 3 | 0.970791409666823 |
| chr9-23741540-23742041 | 5 | 0.974596921224026 |
| chr3-48053237-48053738 | 3 | 0.99593553545041 |
| chr13-44007683-44008184 | 8 | 0.993493681128311 |
| chrX-68576765-68577266 | 5 | 0.843280525865606 |
| chr18-71499260-71499761 | 8 | 0.848703744510641 |
| chr1-204193529-204194030 | 5 | 0.980823025251593 |
| chr4-52711886-52712387 | 6 | 0.991206925481043 |
| chr16-44550167-44550668 | 3 | 0.918170878335968 |
| chr3-163420665-163421166 | 6 | 0.990184538464911 |
| chr1-67755023-67755524 | 1 | 0.821794191220484 |
| chrX-95994993-95995494 | 5 | 0.839544032942879 |
| chr3-82358559-82359060 | 3 | 0.99593553545041 |
| chr6-117569168-117569669 | 3 | 0.988836865675564 |
| chr1-185718132-185718633 | 3 | 0.914442709828393 |
| chr11-71162426-71162927 | 8 | 0.95218982587663 |
| chr14-33773834-33774335 | 2 | 0.950016863950454 |
| chr3-125909233-125909734 | 6 | 0.993739460794304 |
| chr8-60765579-60766080 | 8 | 0.969768323910737 |
| chr3-19083315-19083816 | 6 | 0.944442511220722 |
| chr15-60817464-60817965 | 6 | 0.988639748009796 |
| chr18-73734615-73735116 | 4 | 0.8986268561208 |
| chr15-61967174-61967675 | 1 | 0.963470880767755 |
| chr11-28558292-28558793 | 6 | 0.906574462494711 |
| chr1-153938667-153939168 | 8 | 0.896950623163086 |
| chr1-239733706-239734207 | 4 | 0.85020309932504 |
| chr16-32032711-32033212 | 5 | 0.973337227075162 |
| chr8-65854596-65855097 | 8 | 0.938966509922934 |
| chr1-45268061-45268562 | 8 | 0.993493681128311 |
| chr12-13812910-13813411 | 8 | 0.855214944002816 |
| chr4-120735613-120736114 | 5 | 0.992672557607459 |
| chr6-128323874-128324375 | 8 | 0.892424686498272 |
| chr9-87572578-87573079 | 4 | 0.929978833094843 |
| chr5-110310496-110310997 | 7 | 0.937324878888452 |
| chr13-106701584-106702085 | 5 | 0.983516754572625 |

| | | |
|---|---|---|
| chr3-7921907-7922408 | 7 | 0.908136787472196 |
| chr16-56039866-56040367 | 4 | 0.937702430776053 |
| chr4-11342301-11342802 | 4 | 0.867957511459299 |
| chr2-210183416-210183917 | 1 | 0.961076 |
| chr11-63219492-63219993 | 8 | 0.993593487556906 |
| chr3-8741549-8742050 | 8 | 0.993150480918962 |
| chr3-43351806-43352307 | 3 | 0.99593553545041 |
| chr1-185402841-185403342 | 5 | 0.945711961845778 |
| chr17-21901894-21902395 | 5 | 0.966407028804354 |
| chr6-28905261-28905762 | 8 | 0.993493681128311 |
| chr13-86265267-86265768 | 8 | 0.87640068356975 |
| chr6-9481541-9482042 | 3 | 0.99593553545041 |
| chr5-92374134-92374635 | 8 | 0.825927352374873 |
| chr2-24665473-24665974 | 7 | 0.967941051330686 |
| chr9-14171681-14172182 | 4 | 0.896351453572673 |
| chrX-103451820-103452321 | 8 | 0.963984135617879 |
| chr19-45024278-45024779 | 5 | 0.96763474777221 |
| chr9-60067845-60068346 | 8 | 0.993493681128311 |
| chr8-30409392-30409893 | 3 | 0.958183038068164 |
| chr5-124446154-124446655 | 8 | 0.974207979425143 |
| chr6-82301852-82302353 | 3 | 0.996751104578755 |
| chr9-35592915-35593416 | 3 | 0.991583628551509 |
| chr2-136746408-136746909 | 8 | 0.889771986272322 |
| chr7-94009501-94010002 | 8 | 0.991642151359196 |
| chr6-105775974-105776475 | 4 | 0.864024839852277 |
| chr9-45390811-45391312 | 4 | 0.939606056454526 |
| chr4-68537167-68537668 | 3 | 0.995427132094967 |
| chr17-19208523-19209024 | 5 | 0.885404178368548 |
| chr2-62044385-62044886 | 6 | 0.949771392178814 |
| chr8-76695816-76696317 | 1 | 0.95484101300226 |
| chr13-63321423-63321924 | 1 | 0.870547356608876 |
| chr13-52606011-52606512 | 5 | 0.857100038878245 |
| chr5-41818392-41818893 | 6 | 0.989622731112471 |
| chr2-130157347-130157848 | 8 | 0.967752621287755 |
| chr15-6626223-6626724 | 5 | 0.863728968929989 |
| chrX-35090923-35091424 | 5 | 0.990311370913154 |
| chr2-195977477-195977978 | 2 | 0.864506003710192 |
| chr14-86139943-86140444 | 5 | 0.837675732421225 |
| chr11-72049402-72049903 | 8 | 0.993493681128311 |
| chr3-88138480-88138981 | 6 | 0.860123458970235 |
| chr4-5759134-5759635 | 6 | 0.991206925481043 |
| chr14-92545112-92545613 | 8 | 0.824215994108884 |
| chr4-103893414-103893915 | 2 | 0.962415907170342 |

| | | |
|---|---|---|
| chr20-50586158-50586659 | 6 | 0.899336329556439 |
| chr7-46514449-46514950 | 8 | 0.942929602395596 |
| chr5-145723165-145723666 | 8 | 0.8666713103277 |
| chr9-56156109-56156610 | 5 | 0.88151307969889 |
| chr4-177499682-177500183 | 6 | 0.967550099257853 |
| chr8-11230479-11230980 | 2 | 0.822136798554346 |
| chr10-18383630-18384131 | 8 | 0.852538087144211 |
| chr4-163883999-163884500 | 3 | 0.988992689810603 |
| chr4-14508359-14508860 | 6 | 0.879125181694954 |
| chr2-47154731-47155232 | 3 | 0.951988671775242 |
| chr9-72828628-72829129 | 5 | 0.989772444642198 |
| chr11-2962299-2962800 | 5 | 0.919653061806848 |
| chr2-56906162-56906663 | 8 | 0.987810523947109 |
| chr5-142538439-142538940 | 6 | 0.871316616866718 |
| chr7-100083377-100083878 | 8 | 0.948820172995331 |
| chr3-117751805-117752306 | 8 | 0.841858110601745 |
| chr6-71730652-71731153 | 3 | 0.998848932273241 |
| chr8-74687573-74688074 | 6 | 0.861667638070019 |
| chr3-8728287-8728788 | 4 | 0.951373018639172 |
| chr8-95728772-95729273 | 1 | 0.90781936230688 |
| chr4-72727527-72728028 | 3 | 0.977534437032775 |
| chr18-33842473-33842974 | 3 | 0.888006020078813 |
| chr6-15593640-15594141 | 8 | 0.917303571093118 |
| chr8-33119251-33119752 | 5 | 0.988125361095638 |
| chr1-206903761-206904262 | 8 | 0.819889051895567 |
| chr18-4423514-4424015 | 3 | 0.996637049551616 |
| chr15-88022784-88023285 | 8 | 0.99479639811052 |
| chr15-80049821-80050322 | 8 | 0.839209245468322 |
| chr8-101974703-101975204 | 3 | 0.99593553545041 |
| chr16-18957968-18958469 | 8 | 0.977690490307554 |
| chr16-25394436-25394937 | 1 | 0.962640258912476 |
| chr17-20218272-20218773 | 4 | 0.855225412460844 |
| chr15-38557612-38558113 | 1 | 0.850310786726475 |
| chr3-144146270-144146771 | 4 | 0.881633768556946 |
| chr6-90170392-90170893 | 4 | 0.966827297431083 |
| chr2-44370382-44370883 | 8 | 0.893395796904929 |
| chr11-2534902-2535403 | 3 | 0.993320962655609 |
| chr18-1012725-1013226 | 5 | 0.97882268042844 |
| chr13-77336130-77336631 | 3 | 0.99593553545041 |
| chr16-54237612-54238113 | 6 | 0.991978083515439 |
| chr1-49752231-49752732 | 7 | 0.819874392266577 |
| chr4-82363420-82363921 | 8 | 0.926180906343853 |
| chr3-61417644-61418145 | 8 | 0.977857008939018 |

| | | |
|---|---|---|
| chr20-33205207-33205708 | 5 | 0.96623799361602 |
| chr16-42143815-42144316 | 6 | 0.991206925481043 |
| chr3-121599870-121600371 | 3 | 0.953203596834156 |
| chr19-46742426-46742927 | 3 | 0.998544666814185 |
| chrX-119460474-119460975 | 3 | 0.996422363032864 |
| chr1-132785368-132785869 | 4 | 0.840391654359331 |
| chr18-78366556-78367057 | 3 | 0.994830332618095 |
| chr4-68133982-68134483 | 5 | 0.990157774324612 |
| chr5-19501400-19501901 | 4 | 0.815963602564899 |
| chr13-103739953-103740454 | 6 | 0.994271736137241 |
| chr2-42474145-42474646 | 3 | 0.966695665458928 |
| chr14-76722418-76722919 | 8 | 0.95258895412737 |
| chr6-46593591-46594092 | 1 | 0.870417563901943 |
| chr3-106438007-106438508 | 5 | 0.973978070144304 |
| chr4-149438015-149438516 | 2 | 0.871943786547679 |
| chr18-357617-358118 | 6 | 0.872815452893491 |
| chr9-64914935-64915436 | 8 | 0.877195730286036 |
| chr8-907648-908149 | 6 | 0.991206925481043 |
| chr5-153529185-153529686 | 2 | 0.895177009455543 |
| chr1-218385937-218386438 | 8 | 0.993493681128311 |
| chrX-10526610-10527111 | 8 | 0.995169207708373 |
| chr7-103066001-103066502 | 6 | 0.971253469318272 |
| chr2-170022542-170023043 | 8 | 0.870829674676934 |
| chr1-9686820-9687321 | 4 | 0.871164280976042 |
| chr18-69694870-69695371 | 2 | 0.859637415298935 |
| chr13-24835260-24835761 | 6 | 0.917552358342899 |
| chr2-36834295-36834796 | 3 | 0.996869946862024 |
| chr1-21659003-21659504 | 1 | 0.868760360572483 |
| chr6-16503812-16504313 | 3 | 0.917538366975955 |
| chr20-24868125-24868626 | 4 | 0.915783362859681 |
| chr12-1279195-1279696 | 3 | 0.995213127219095 |
| chr2-139905096-139905597 | 5 | 0.966716843903406 |
| chr14-92380812-92381313 | 6 | 0.976549124330896 |
| chr16-10470511-10471012 | 7 | 0.933366512991694 |
| chr15-44544952-44545453 | 5 | 0.982114903854221 |
| chr18-31524674-31525175 | 8 | 0.993535212877223 |
| chr11-70177148-70177649 | 3 | 0.922031841133563 |
| chr7-56042592-56043093 | 8 | 0.894569528541365 |
| chr2-69826232-69826733 | 3 | 0.891274267591268 |
| chr17-26295253-26295754 | 3 | 0.993492291467515 |
| chr19-40319970-40320471 | 6 | 0.919621114398887 |
| chr5-23411687-23412188 | 4 | 0.916161692775233 |
| chr4-52140368-52140869 | 1 | 0.948072044837463 |

| | | |
|---|---|---|
| chr8-95005421-95005922 | 7 | 0.894971747198187 |
| chr3-18547477-18547978 | 8 | 0.906226540289599 |
| chr8-64008937-64009438 | 6 | 0.956127042163304 |
| chr10-95471453-95471954 | 8 | 0.868506615350207 |
| chr8-59006919-59007420 | 4 | 0.838515273034472 |
| chr5-144383224-144383725 | 8 | 0.980377790532972 |
| chr9-50468521-50469022 | 6 | 0.992062858918006 |
| chr18-16100411-16100912 | 8 | 0.857344337729116 |
| chr2-224211179-224211680 | 2 | 0.884187551964388 |
| chr7-87569365-87569866 | 5 | 0.926665250769389 |
| chrX-12722589-12723090 | 8 | 0.87655189553735 |
| chr4-30865328-30865829 | 8 | 0.9763730874898 |
| chr15-12769772-12770273 | 5 | 0.990691441369711 |
| chr17-6203154-6203655 | 5 | 0.883087211665675 |
| chr7-51336528-51337029 | 8 | 0.938582822811594 |
| chr2-39476180-39476681 | 6 | 0.98606940007988 |
| chr18-44388646-44389147 | 4 | 0.877388692948721 |
| chr1-219183894-219184395 | 1 | 0.892673998397363 |
| chr1-29889366-29889867 | 8 | 0.993493681128311 |
| chr3-159551043-159551544 | 8 | 0.846876836498127 |
| chr7-76022056-76022557 | 8 | 0.96935021090113 |
| chr3-122153751-122154252 | 1 | 0.92908210753935 |
| chr7-25362775-25363276 | 6 | 0.993022508286636 |
| chr16-79043869-79044370 | 8 | 0.992672713234939 |
| chr4-50767587-50768088 | 8 | 0.978607137446113 |
| chr17-25154246-25154747 | 6 | 0.992823139929147 |
| chr14-89602099-89602600 | 5 | 0.983919978840557 |
| chr4-84495136-84495637 | 3 | 0.985367209814985 |
| chr8-98323478-98323979 | 8 | 0.913727259656134 |
| chr1-257960553-257961054 | 3 | 0.997212240988247 |
| chr2-170555417-170555918 | 8 | 0.958998290891198 |
| chr6-70878279-70878780 | 8 | 0.874874848036543 |
| chr5-126175237-126175738 | 3 | 0.981015358577734 |
| chr10-15521164-15521665 | 8 | 0.849251037060485 |
| chr6-15495452-15495953 | 4 | 0.823234154321803 |
| chr1-180100137-180100638 | 2 | 0.95321488700671 |
| chr14-68204319-68204820 | 6 | 0.946918076343255 |
| chr3-108025090-108025591 | 4 | 0.864032018868637 |
| chr10-64426486-64426987 | 8 | 0.896239047146281 |
| chr17-67941467-67941968 | 3 | 0.995106274408176 |
| chr16-61887903-61888404 | 3 | 0.99593553545041 |
| chr2-188667406-188667907 | 8 | 0.814692621109821 |
| chr5-15302138-15302639 | 1 | 0.864306438352937 |

| | | |
|---|---|---|
| chr11-2661988-2662489 | 5 | 0.877226463111614 |
| chr19-8865152-8865653 | 4 | 0.869649118220709 |
| chr2-243185723-243186224 | 8 | 0.836981461671209 |
| chr14-25143533-25144034 | 3 | 0.993635659128688 |
| chr14-46408971-46409472 | 5 | 0.858309688693282 |
| chr2-101726018-101726519 | 8 | 0.868779701566459 |
| chr20-8114105-8114606 | 4 | 0.883282221011166 |
| chr6-44618835-44619336 | 8 | 0.966706210434797 |
| chr15-7620159-7620660 | 5 | 0.980946123686553 |
| chr16-70584043-70584544 | 5 | 0.898021 |
| chr7-75848669-75849170 | 1 | 0.945945389183251 |
| chr4-43178960-43179461 | 3 | 0.968688430450691 |
| chr2-7510239-7510740 | 6 | 0.979899504104703 |
| chr2-169362345-169362846 | 8 | 0.974890852370021 |
| chr15-92857108-92857609 | 8 | 0.977331421708112 |
| chr17-79792249-79792750 | 3 | 0.932083154115044 |
| chr2-26004521-26005022 | 3 | 0.983951826918071 |
| chr1-93901545-93902046 | 8 | 0.922286039630256 |
| chr12-26774078-26774579 | 8 | 0.993493681128311 |
| chr17-45915823-45916324 | 3 | 0.986058324088949 |
| chr18-52589134-52589635 | 6 | 0.99604674796319 |
| chr4-71439193-71439694 | 5 | 0.892975033169339 |
| chr6-19458159-19458660 | 1 | 0.89458796139615 |
| chr7-85162660-85163161 | 8 | 0.872673570273182 |
| chr8-122034351-122034852 | 6 | 0.94654956574851 |
| chr16-81842335-81842836 | 4 | 0.824218834684826 |
| chr4-17782579-17783080 | 5 | 0.984763343311764 |
| chr8-2418994-2419495 | 5 | 0.954607600747095 |
| chr4-133845478-133845979 | 7 | 0.894587076841821 |
| chr10-26878611-26879112 | 3 | 0.996813981408684 |
| chr5-162244405-162244906 | 5 | 0.987140534610105 |
| chr10-42475751-42476252 | 8 | 0.974122513997053 |
| chr12-32077266-32077767 | 8 | 0.927298532546954 |
| chr16-1487558-1488059 | 4 | 0.972548120767105 |
| chr18-30067204-30067705 | 2 | 0.941570359785178 |
| chr8-56205791-56206292 | 6 | 0.991206925481043 |
| chr18-649998-650499 | 3 | 0.949201488830753 |
| chr10-27485116-27485617 | 6 | 0.99229172785584 |
| chr13-44260328-44260829 | 3 | 0.981945919438092 |
| chr16-9528359-9528860 | 3 | 0.866520280034907 |
| chr2-104589236-104589737 | 6 | 0.987508083509351 |
| chr19-17451388-17451889 | 7 | 0.866955804928371 |
| chrY-16347833-16348334 | 3 | 0.99593553545041 |

| | | |
|---|---|---|
| chr3-66165412-66165913 | 5 | 0.848368739897356 |
| chrX-33478819-33479320 | 6 | 0.997361207451738 |
| chr2-231774460-231774961 | 5 | 0.946225144949829 |
| chr13-55558763-55559264 | 5 | 0.990615520394705 |
| chr1-74578703-74579204 | 3 | 0.99593553545041 |
| chr4-180221290-180221791 | 2 | 0.955005251685376 |
| chr14-74962657-74963158 | 6 | 0.882787509722502 |
| chr13-103601569-103602070 | 3 | 0.933701193888506 |
| chr3-12368878-12369379 | 8 | 0.869435373118771 |
| chr9-19132930-19133431 | 3 | 0.993308785348917 |
| chr2-34702896-34703397 | 8 | 0.992533571740032 |
| chr10-98588219-98588720 | 8 | 0.92301753393011 |
| chr17-30090601-30091102 | 5 | 0.97894769788217 |
| chr1-221889306-221889807 | 6 | 0.945058328543429 |
| chr12-17086867-17087368 | 5 | 0.987990042357378 |
| chr17-42461929-42462430 | 6 | 0.947880546735885 |
| chr13-90312712-90313213 | 2 | 0.969375788164346 |
| chr19-37312569-37313070 | 8 | 0.870941608896229 |
| chr3-119908822-119909323 | 8 | 0.962300893231961 |
| chr12-26752815-26753316 | 8 | 0.831571680221989 |
| chr4-148117706-148118207 | 8 | 0.817248924365227 |
| chr2-14794912-14795413 | 3 | 0.99593553545041 |
| chr2-43231494-43231995 | 8 | 0.993028328509668 |
| chr10-5915439-5915940 | 3 | 0.99593553545041 |
| chr6-90054693-90055194 | 4 | 0.970924325311068 |
| chr3-150824373-150824874 | 4 | 0.82810638740749 |
| chr1-52593418-52593919 | 3 | 0.99593553545041 |
| chr6-31949592-31950093 | 8 | 0.984448374513365 |
| chr5-91003344-91003845 | 8 | 0.992534563546638 |
| chr16-21290219-21290720 | 8 | 0.96144126173487 |
| chr7-115178512-115179013 | 6 | 0.982068516734074 |
| chr15-53112370-53112871 | 8 | 0.834678309647554 |
| chr8-52985267-52985768 | 8 | 0.889435181593094 |
| chr18-23319384-23319885 | 4 | 0.885447553801232 |
| chr6-45732601-45733102 | 4 | 0.966068882415221 |
| chr8-109045816-109046317 | 8 | 0.95467765804424 |
| chr14-42981194-42981695 | 4 | 0.830708027266995 |
| chr5-16107529-16108030 | 3 | 0.990099087344824 |
| chr2-113493123-113493624 | 1 | 0.903220168789818 |
| chr8-113901792-113902293 | 8 | 0.821308838992708 |
| chr9-50860503-50861004 | 3 | 0.996892495833392 |
| chr4-175853512-175854013 | 8 | 0.990116633663643 |
| chr13-93311816-93312317 | 4 | 0.865262964614262 |

| | | |
|---|---|---|
| chr12-44166827-44167328 | 8 | 0.889133355884351 |
| chr15-19742835-19743336 | 3 | 0.996372203894258 |
| chr18-41829078-41829579 | 3 | 0.844593756174097 |
| chr11-78462623-78463124 | 8 | 0.919441964719679 |
| chr5-20133650-20134151 | 6 | 0.990285552005993 |
| chr1-223678491-223678992 | 5 | 0.93998712975646 |
| chr10-74746646-74747147 | 8 | 0.818312554292378 |
| chr15-45044448-45044949 | 5 | 0.945626534937349 |
| chr1-99006601-99007102 | 3 | 0.946555492290409 |
| chr3-140123891-140124392 | 6 | 0.895647583191369 |
| chr15-64128189-64128690 | 5 | 0.831257926117256 |
| chr18-50649804-50650305 | 4 | 0.949099375884655 |
| chr4-80726152-80726653 | 8 | 0.917087590136503 |
| chr18-27250989-27251490 | 5 | 0.831130553812958 |
| chr2-35110576-35111077 | 4 | 0.945174769663999 |
| chr17-10705752-10706253 | 8 | 0.979301305469176 |
| chr16-57382796-57383297 | 7 | 0.910680796632359 |
| chr4-67841898-67842399 | 4 | 0.886444178426954 |
| chr3-42660890-42661391 | 3 | 0.937949897266771 |
| chr4-127972443-127972944 | 3 | 0.996854163093434 |
| chr13-77346538-77347039 | 5 | 0.9868442763344 |
| chr5-130434454-130434955 | 8 | 0.856669882452216 |
| chr5-41086141-41086642 | 3 | 0.995666599800384 |
| chr5-91794616-91795117 | 8 | 0.893662637742225 |
| chr4-52988063-52988564 | 8 | 0.842807924174053 |
| chr3-148012065-148012566 | 7 | 0.851121047351769 |
| chr15-41350328-41350829 | 8 | 0.917878507978168 |
| chr8-107254918-107255419 | 6 | 0.85961321444283 |
| chr4-72983735-72984236 | 6 | 0.957607306940818 |
| chr20-5429131-5429632 | 8 | 0.834216932333932 |
| chr5-82978739-82979240 | 2 | 0.886790018726367 |
| chr3-134175727-134176228 | 3 | 0.995953224783561 |
| chr3-162625205-162625706 | 5 | 0.97565579582977 |
| chr4-162386292-162386793 | 8 | 0.997731358211563 |
| chr4-118755456-118755957 | 2 | 0.946362814452236 |
| chr19-26987841-26988342 | 8 | 0.983340483993734 |
| chr2-91802053-91802554 | 2 | 0.955831905831136 |
| chr7-131685857-131686358 | 8 | 0.869191817472507 |
| chr18-26029921-26030422 | 8 | 0.905895153164944 |
| chr20-41607457-41607958 | 8 | 0.834157822368945 |
| chr19-50573903-50574404 | 3 | 0.881199583948614 |
| chr1-192061517-192062018 | 8 | 0.93907963684465 |
| chr15-21213697-21214198 | 7 | 0.905336615587407 |

| | | |
|---|---|---|
| chr19-20719426-20719927 | 6 | 0.985613484703734 |
| chr4-136538688-136539189 | 6 | 0.991206925481043 |
| chr6-3245809-3246310 | 6 | 0.969979032467039 |
| chr18-82351772-82352273 | 6 | 0.820972836940916 |
| chr17-38530676-38531177 | 8 | 0.923165105901579 |
| chr16-69638107-69638608 | 8 | 0.993493681128311 |
| chr8-73879771-73880272 | 8 | 0.994363540132764 |
| chr9-74663329-74663830 | 3 | 0.99710423822574 |
| chr17-81991649-81992150 | 3 | 0.980679911785082 |
| chr4-72262982-72263483 | 7 | 0.948926736711147 |
| chr19-53341737-53342238 | 3 | 0.947692951431334 |
| chr3-7266026-7266527 | 8 | 0.986377779347423 |
| chr6-106945420-106945921 | 4 | 0.884935876583552 |
| chr15-44140063-44140564 | 4 | 0.921720026939049 |
| chr2-46475237-46475738 | 6 | 0.833989985001203 |
| chr11-55458494-55458995 | 5 | 0.890475954876702 |
| chr6-109522581-109523082 | 6 | 0.989505593146196 |
| chr14-66938266-66938767 | 3 | 0.996765388867973 |
| chr2-183605298-183605799 | 3 | 0.982093549044534 |
| chr16-9845871-9846372 | 3 | 0.996362695132671 |
| chr9-86346106-86346607 | 3 | 0.991146771269865 |
| chr8-37247104-37247605 | 5 | 0.85967880000766 |
| chr1-16008059-16008560 | 2 | 0.893227596817787 |
| chr2-187851271-187851772 | 4 | 0.880955904045866 |
| chr11-52369343-52369844 | 1 | 0.919746959113406 |
| chr11-71548641-71549142 | 6 | 0.92760804701803 |
| chr6-54951455-54951956 | 3 | 0.978681251149223 |
| chr12-33319022-33319523 | 8 | 0.901186122722985 |
| chr10-37053338-37053839 | 4 | 0.906994731028541 |
| chr1-161793549-161794050 | 6 | 0.989383467536811 |
| chr1-125277931-125278432 | 6 | 0.925529481250743 |
| chr6-3994153-3994654 | 3 | 0.99593553545041 |
| chr7-127228509-127229010 | 1 | 0.940225294273371 |
| chr3-38866442-38866943 | 3 | 0.996795896493435 |
| chr13-93175202-93175703 | 4 | 0.928923703988755 |
| chr13-93974109-93974610 | 4 | 0.824491132364398 |
| chr5-959512-960013 | 3 | 0.988442776938195 |
| chr16-44287000-44287501 | 5 | 0.868229340920695 |
| chr1-243929299-243929800 | 4 | 0.838531289559261 |
| chr4-171999891-172000392 | 8 | 0.988723924452092 |
| chr10-14662000-14662501 | 5 | 0.98042881447041 |
| chr19-9552505-9553006 | 4 | 0.899032954351645 |
| chr20-19608568-19609069 | 2 | 0.957494444123375 |

| | | |
|---|---|---|
| chr14-33323546-33324047 | 8 | 0.993493681128311 |
| chr4-172239713-172240214 | 6 | 0.987275521252346 |
| chr15-53192723-53193224 | 5 | 0.942086997017476 |
| chr1-98882382-98882883 | 1 | 0.892305061455585 |
| chr9-98632294-98632795 | 2 | 0.859414041519964 |
| chr14-79523894-79524395 | 8 | 0.957021725626935 |
| chr1-243229796-243230297 | 4 | 0.96359584367927 |
| chr5-11147703-11148204 | 3 | 0.997001812610079 |
| chr13-70338308-70338809 | 5 | 0.858131288718228 |
| chr2-221329918-221330419 | 5 | 0.981370826603699 |
| chr9-38900803-38901304 | 5 | 0.941310188914193 |
| chrX-146686813-146687314 | 5 | 0.978179635525043 |
| chr2-62283445-62283946 | 3 | 0.950656982699278 |
| chr12-22372005-22372506 | 3 | 0.99593553545041 |
| chr14-12687492-12687993 | 8 | 0.986669510963414 |
| chr4-157351180-157351681 | 8 | 0.968088713238052 |
| chr5-109770324-109770825 | 8 | 0.903945611619539 |
| chr14-71611556-71612057 | 3 | 0.857334321833275 |
| chr17-6609550-6610051 | 3 | 0.883241322893504 |
| chrX-131032214-131032715 | 5 | 0.948560307617569 |
| chr11-41389653-41390154 | 5 | 0.98199596741887 |
| chr17-40463664-40464165 | 8 | 0.882584458942544 |
| chr16-14939322-14939823 | 3 | 0.996741839517006 |
| chr10-99327381-99327882 | 6 | 0.997062100170004 |
| chr8-47400080-47400581 | 6 | 0.967465209082495 |
| chr2-106767157-106767658 | 3 | 0.99053143589347 |
| chr14-86239182-86239683 | 3 | 0.996412772647029 |
| chr20-44709278-44709779 | 6 | 0.967421714210101 |
| chr1-201166151-201166652 | 8 | 0.979340944413938 |
| chr4-73583029-73583530 | 3 | 0.996804914131944 |
| chr6-43886822-43887323 | 3 | 0.995773207572645 |
| chr15-100990458-100990959 | 3 | 0.99593553545041 |
| chr2-51853017-51853518 | 4 | 0.923129577935617 |
| chr9-109187641-109188142 | 7 | 0.916364982878037 |
| chr2-172975076-172975577 | 8 | 0.894613325992837 |
| chr10-36281875-36282376 | 8 | 0.816620226941335 |
| chrX-24265622-24266123 | 8 | 0.932628719043136 |
| chr1-94512132-94512633 | 8 | 0.8674347937327 |
| chr4-168451188-168451689 | 8 | 0.951495241330752 |
| chr6-126386511-126387012 | 3 | 0.936269432680251 |
| chr19-6982225-6982726 | 3 | 0.99593553545041 |
| chr10-102281854-102282355 | 5 | 0.820296286778094 |
| chr19-46736209-46736710 | 8 | 0.8586621661827 |

| | | |
|---|---|---|
| chr5-70128653-70129154 | 4 | 0.86993014062063 |
| chr9-91577305-91577806 | 8 | 0.994848827457589 |
| chr16-68724830-68725331 | 8 | 0.93367873180213 |
| chr13-65978394-65978895 | 8 | 0.955429706169085 |
| chr12-5763353-5763854 | 8 | 0.968006715624568 |
| chr5-152101185-152101686 | 8 | 0.976532430665467 |
| chr3-148430412-148430913 | 3 | 0.996808467224028 |
| chr13-38848918-38849419 | 5 | 0.980892317013054 |
| chr15-94507776-94508277 | 5 | 0.904563496466545 |
| chr10-25481504-25482005 | 3 | 0.892008551887863 |
| chr12-7916786-7917287 | 8 | 0.843820158516282 |
| chr9-61410857-61411358 | 3 | 0.956410262694165 |
| chr16-17308495-17308996 | 2 | 0.84863084136273 |
| chr6-81934575-81935076 | 3 | 0.996791108218244 |
| chr19-48091914-48092415 | 8 | 0.992215381342621 |
| chr3-64319891-64320392 | 3 | 0.995235571385819 |
| chr18-69217375-69217876 | 8 | 0.892526800227842 |
| chr7-8596513-8597014 | 8 | 0.887381147293287 |
| chr18-11393080-11393581 | 6 | 0.983984090016715 |
| chr11-13051421-13051922 | 6 | 0.964630521693261 |
| chr10-35888137-35888638 | 5 | 0.941043426290086 |
| chr13-67488766-67489267 | 5 | 0.986729747089484 |
| chr18-60313956-60314457 | 1 | 0.862434156262529 |
| chr3-49627207-49627708 | 8 | 0.95363275750044 |
| chr1-233628964-233629465 | 6 | 0.991893050419162 |
| chr3-161545768-161546269 | 6 | 0.993072080044206 |
| chr20-18599542-18600043 | 4 | 0.940962445361097 |
| chr1-40484319-40484820 | 8 | 0.954787021911671 |
| chr4-84402500-84403001 | 3 | 0.99593553545041 |
| chr10-74594238-74594739 | 3 | 0.972652720523161 |
| chr5-124056318-124056819 | 2 | 0.92426827745052 |
| chr1-152817613-152818114 | 3 | 0.961866522918511 |
| chr5-85273680-85274181 | 8 | 0.961290660618194 |
| chr4-120896233-120896734 | 8 | 0.978054498289118 |
| chr6-39257675-39258176 | 8 | 0.923658377156192 |
| chr14-95903624-95904125 | 3 | 0.975499076639385 |
| chr13-93808231-93808732 | 8 | 0.883619526657272 |
| chr13-63659754-63660255 | 3 | 0.987040038632554 |
| chr16-56349446-56349947 | 5 | 0.9912339278223 |
| chr4-14855007-14855508 | 6 | 0.878506093051855 |
| chr13-64709690-64710191 | 1 | 0.910498108385133 |
| chr4-51046779-51047280 | 8 | 0.895809685513761 |
| chr11-80330755-80331256 | 8 | 0.995243142913875 |

| | | |
|---|---|---|
| chrX-19774201-19774702 | 5 | 0.872427156924691 |
| chr2-240820870-240821371 | 4 | 0.934969047868185 |
| chr3-34553924-34554425 | 6 | 0.9935928183376 |
| chr17-22644332-22644833 | 5 | 0.972860609830868 |
| chr16-7273197-7273698 | 8 | 0.82678036858821 |
| chr5-143240514-143241015 | 8 | 0.968644969539934 |
| chr8-111161895-111162396 | 4 | 0.889485316009627 |
| chr2-53195844-53196345 | 5 | 0.970648724069164 |
| chr4-81585420-81585921 | 7 | 0.939939655775954 |
| chr17-53823080-53823581 | 6 | 0.99042137756302 |
| chr2-212104745-212105246 | 2 | 0.92482022450868 |
| chr12-46376872-46377373 | 5 | 0.975421764667608 |
| chr2-213136414-213136915 | 3 | 0.99593553545041 |
| chr3-165930535-165931036 | 8 | 0.894458816491972 |
| chr3-76816428-76816929 | 6 | 0.98943124371163 |
| chr6-92351483-92351984 | 6 | 0.991118779454105 |
| chr2-112628213-112628714 | 3 | 0.950409260124716 |
| chr4-90796074-90796575 | 7 | 0.956215850454304 |
| chr5-36654195-36654696 | 8 | 0.932597316218472 |
| chr19-19139768-19140269 | 3 | 0.875549738697087 |
| chr12-43239914-43240415 | 5 | 0.993482052496622 |
| chr1-236480471-236480972 | 3 | 0.818246548410871 |
| chr9-92110685-92111186 | 8 | 0.994654879976391 |
| chr10-19910065-19910566 | 8 | 0.995016656525936 |
| chr12-342115-342616 | 1 | 0.933900347512742 |
| chr3-122133869-122134370 | 6 | 0.98232666614138 |
| chrX-130482902-130483403 | 8 | 0.949223302162738 |
| chr17-68219344-68219845 | 8 | 0.993290950986883 |
| chr1-152346780-152347281 | 5 | 0.952142345808266 |
| chr15-32402027-32402528 | 8 | 0.991196787932197 |
| chr7-119678856-119679357 | 7 | 0.837544764246661 |
| chr7-93994896-93995397 | 3 | 0.994121235519034 |
| chr1-181206819-181207320 | 4 | 0.849891911816611 |
| chr5-111693612-111694113 | 8 | 0.995249052783328 |
| chr9-39334041-39334542 | 8 | 0.993493681128311 |
| chr17-46195782-46196283 | 8 | 0.985648752006408 |
| chr15-27032734-27033235 | 3 | 0.996314425001734 |
| chr2-105691219-105691720 | 3 | 0.994307140285276 |
| chr5-4583188-4583689 | 8 | 0.991448795954173 |
| chr7-71365226-71365727 | 1 | 0.863608980810649 |
| chr1-29979633-29980134 | 8 | 0.823818958568879 |
| chr3-43593198-43593699 | 7 | 0.82109114061123 |
| chr12-46484126-46484627 | 6 | 0.989683212347944 |

| | | |
|---|---|---|
| chr16-83445924-83446425 | 3 | 0.994939697148759 |
| chr8-105921495-105921996 | 8 | 0.833752265860408 |
| chr14-29166207-29166708 | 6 | 0.951376841042623 |
| chr12-131832-132333 | 8 | 0.986736317128256 |
| chr1-178171545-178172046 | 6 | 0.928109550814679 |
| chr14-264727-265228 | 5 | 0.869497728761255 |
| chr20-13819629-13820130 | 6 | 0.967602548769566 |
| chr1-76536347-76536848 | 3 | 0.99593553545041 |
| chr17-13341765-13342266 | 3 | 0.838602757618894 |
| chr8-57107726-57108227 | 8 | 0.945284649905621 |
| chr6-49487988-49488489 | 1 | 0.913147461493766 |
| chr1-126616236-126616737 | 4 | 0.909544886020556 |
| chr2-77387553-77388054 | 1 | 0.847249990571014 |
| chr4-83668323-83668824 | 8 | 0.919922622126487 |
| chr13-69129811-69130312 | 6 | 0.837611619039401 |
| chr1-65946668-65947169 | 6 | 0.831441848976785 |
| chr4-118570707-118571208 | 6 | 0.964481764590424 |
| chr20-38108779-38109280 | 8 | 0.954535795504942 |
| chr5-97773983-97774484 | 3 | 0.973687213480431 |
| chr10-67746079-67746580 | 3 | 0.996766939926028 |
| chr3-15885593-15886094 | 6 | 0.849888581759355 |
| chr7-71401846-71402347 | 6 | 0.929147725230279 |
| chr4-163646468-163646969 | 4 | 0.816681050693876 |
| chrX-126010312-126010813 | 6 | 0.988059349382005 |
| chr5-125206473-125206974 | 8 | 0.892352806147722 |
| chr18-47753288-47753789 | 4 | 0.892541053277084 |
| chr11-74483726-74484227 | 8 | 0.979720318479345 |
| chr16-48128832-48129333 | 4 | 0.971558627455467 |
| chr9-76547110-76547611 | 5 | 0.951528219390022 |
| chr8-101749524-101750025 | 4 | 0.954087794564732 |
| chr19-16086283-16086784 | 5 | 0.97865981343244 |
| chr3-149467570-149468071 | 1 | 0.94280136657835 |
| chrX-39755163-39755664 | 5 | 0.982299489905808 |
| chr1-250114680-250115181 | 3 | 0.991470386531119 |
| chr7-106813096-106813597 | 3 | 0.99593553545041 |
| chr6-37111434-37111935 | 3 | 0.99688392666275 |
| chr12-29114724-29115225 | 5 | 0.982522619673715 |
| chr1-164899541-164900042 | 7 | 0.912511840963671 |
| chr6-138992196-138992697 | 8 | 0.833658326602283 |
| chr3-123017671-123018172 | 1 | 0.896821825284991 |
| chr17-18315249-18315750 | 3 | 0.958989991007967 |
| chr4-125414693-125415194 | 2 | 0.847025240535619 |
| chr9-41543243-41543744 | 6 | 0.986486909128116 |

| | | |
|---|---|---|
| chr3-8518938-8519439 | 5 | 0.99077067378998 |
| chr4-93247309-93247810 | 3 | 0.995467382841383 |
| chr17-32749029-32749530 | 8 | 0.99343796502999 |
| chr11-16210828-16211329 | 8 | 0.985128059485854 |
| chr6-5159385-5159886 | 8 | 0.993493681128311 |
| chrX-128767976-128768477 | 6 | 0.991206925481043 |
| chr4-71493677-71494178 | 5 | 0.842421694296968 |
| chr7-45007794-45008295 | 3 | 0.986806835935234 |
| chr16-76875629-76876130 | 5 | 0.989107547075731 |
| chr4-140591850-140592351 | 3 | 0.99686246439296 |
| chr5-138881403-138881904 | 8 | 0.96137608146315 |
| chr4-123397352-123397853 | 4 | 0.839150808641909 |
| chr18-80216375-80216876 | 8 | 0.876696734900585 |
| chr20-52765711-52766212 | 8 | 0.993493681128311 |
| chr9-77593282-77593783 | 8 | 0.977852202094554 |
| chr18-32290992-32291493 | 3 | 0.9936383125589 |
| chr3-7920466-7920967 | 7 | 0.927631491488398 |
| chr9-1755811-1756312 | 6 | 0.988371320336821 |
| chr4-54554249-54554750 | 3 | 0.926321967945054 |
| chr2-59594317-59594818 | 8 | 0.91097546190353 |
| chr18-38309545-38310046 | 6 | 0.892160649072393 |
| chr5-18879774-18880275 | 8 | 0.992871864219775 |
| chr9-18754439-18754940 | 6 | 0.983879498109512 |
| chr9-3917006-3917507 | 8 | 0.972981532525027 |
| chr1-118367515-118368016 | 4 | 0.921172808939869 |
| chr8-26542494-26542995 | 3 | 0.99593553545041 |
| chr7-94436937-94437438 | 8 | 0.957750383322929 |
| chr9-3227833-3228334 | 1 | 0.920067126228465 |
| chr1-195418966-195419467 | 3 | 0.979564582894966 |
| chr15-94791692-94792193 | 6 | 0.992277950330663 |
| chr10-32302725-32303226 | 3 | 0.815009051322576 |
| chr1-220034226-220034727 | 8 | 0.957898492016209 |
| chr3-37811495-37811996 | 5 | 0.98618834865241 |
| chr1-129121410-129121911 | 8 | 0.905339724880224 |
| chr2-222731295-222731796 | 3 | 0.990247608960949 |
| chr16-60173291-60173792 | 8 | 0.984184195595576 |
| chrX-67216043-67216544 | 8 | 0.976584336364265 |
| chr2-153607384-153607885 | 8 | 0.97052800898644 |
| chr5-49766337-49766838 | 3 | 0.99593553545041 |
| chr9-70891864-70892365 | 5 | 0.915828290244456 |
| chr17-64353124-64353625 | 5 | 0.985903164255205 |
| chr12-15951075-15951576 | 8 | 0.89871953133767 |
| chr1-122528715-122529216 | 2 | 0.934271914707678 |

| | | |
|---|---|---|
| chr2-159393668-159394169 | 3 | 0.99593553545041 |
| chr1-207144796-207145297 | 8 | 0.886184409348409 |
| chr10-99153684-99154185 | 8 | 0.905560223389614 |
| chr12-40090358-40090859 | 8 | 0.917295622056187 |
| chr15-20902888-20903389 | 8 | 0.816264146683736 |
| chr2-78498688-78499189 | 5 | 0.948749697100954 |
| chr4-107274542-107275043 | 3 | 0.991274658473959 |
| chr12-37804303-37804804 | 8 | 0.985503934313584 |
| chr9-3426922-3427423 | 8 | 0.848131561693578 |
| chr1-99374845-99375346 | 6 | 0.980749936373792 |
| chr4-128320970-128321471 | 8 | 0.988952782922706 |
| chr14-19159838-19160339 | 8 | 0.906366499306118 |
| chr4-9957186-9957687 | 6 | 0.946476575764483 |
| chr15-92346410-92346911 | 3 | 0.989954318297675 |
| chr4-82196483-82196984 | 2 | 0.924758223915411 |
| chr5-166138416-166138917 | 8 | 0.922354014284253 |
| chr16-39767268-39767769 | 7 | 0.870450763330307 |
| chr12-15898434-15898935 | 8 | 0.837831438282682 |
| chr4-79938808-79939309 | 5 | 0.991916305638433 |
| chr1-92846907-92847408 | 3 | 0.940352443678381 |
| chrX-104887640-104888141 | 6 | 0.909965298097436 |
| chr17-74007434-74007935 | 7 | 0.867101968760562 |
| chr13-95938418-95938919 | 8 | 0.816412270934591 |
| chr4-76090507-76091008 | 8 | 0.929664849689925 |
| chr7-26522164-26522665 | 6 | 0.990132632889215 |
| chr13-105191856-105192357 | 2 | 0.94774295384541 |
| chr2-226119512-226120013 | 8 | 0.995081136962402 |
| chr15-61965861-61966362 | 3 | 0.815992557680269 |
| chr10-52055872-52056373 | 8 | 0.993493681128311 |
| chrX-46303611-46304112 | 5 | 0.967373190671018 |
| chr6-44175341-44175842 | 2 | 0.830780427200397 |
| chr9-17855673-17856174 | 8 | 0.905024761037441 |
| chr18-62916559-62917060 | 8 | 0.900882299771804 |
| chr6-132006388-132006889 | 2 | 0.863762350824965 |
| chrX-121248506-121249007 | 3 | 0.99592822663741 |
| chr16-74461940-74462441 | 8 | 0.883262657470917 |
| chr17-68701340-68701841 | 6 | 0.989329065620784 |
| chr6-124726361-124726862 | 4 | 0.920777637800347 |
| chr5-101285353-101285854 | 8 | 0.851289207315448 |
| chr14-88765499-88766000 | 6 | 0.992022920571481 |
| chr20-5398646-5399147 | 7 | 0.823033600317083 |
| chr16-49398342-49398843 | 3 | 0.966089578460968 |
| chr15-39280583-39281084 | 7 | 0.836213535933712 |

| | | |
|---|---|---|
| chr13-22572701-22573202 | 5 | 0.858522661654098 |
| chr2-195994143-195994644 | 8 | 0.946140006217098 |
| chr2-175291744-175292245 | 8 | 0.878950780647195 |
| chr17-27265729-27266230 | 5 | 0.980702501284673 |
| chr7-28446147-28446648 | 3 | 0.99657388123021 |
| chr7-88985134-88985635 | 8 | 0.866889 |
| chr5-162823078-162823579 | 8 | 0.954647941788032 |
| chr18-11849628-11850129 | 8 | 0.880533373146074 |
| chr4-30540721-30541222 | 6 | 0.985000915561644 |
| chr3-166850630-166851131 | 4 | 0.972975481236495 |
| chr9-102482181-102482682 | 7 | 0.848330964490827 |
| chr20-46000910-46001411 | 4 | 0.849295431801203 |
| chr6-10683833-10684334 | 8 | 0.993493681128311 |
| chr13-14784847-14785348 | 6 | 0.973385531715858 |
| chr5-62807148-62807649 | 5 | 0.927355925225082 |
| chr20-30604368-30604869 | 8 | 0.854539898809102 |
| chr19-13522104-13522605 | 5 | 0.983919978840557 |
| chr15-13995886-13996387 | 7 | 0.830430806064344 |
| chr16-52135369-52135870 | 6 | 0.975483072077105 |
| chr3-112485098-112485599 | 8 | 0.967234263819676 |
| chr3-133252733-133253234 | 8 | 0.919087118352697 |
| chr1-30093286-30093787 | 8 | 0.995248837508557 |
| chr8-69979023-69979524 | 8 | 0.949820538623644 |
| chr2-124602426-124602927 | 6 | 0.990121201348237 |
| chr2-219408113-219408614 | 4 | 0.818988629407328 |
| chr2-219402556-219403057 | 4 | 0.911887099043009 |
| chr14-58485899-58486400 | 5 | 0.967379363821867 |
| chr20-30162125-30162626 | 1 | 0.837026786785189 |
| chr10-72487370-72487871 | 6 | 0.983152624644642 |
| chr1-131993736-131994237 | 3 | 0.961535464829591 |
| chr7-50145395-50145896 | 3 | 0.994887580629061 |
| chr9-56276027-56276528 | 3 | 0.978997011707826 |
| chr4-115918132-115918633 | 8 | 0.915159340730897 |
| chr4-131306128-131306629 | 6 | 0.990200887846823 |
| chr10-56137672-56138173 | 8 | 0.911316598813576 |
| chr1-45930497-45930998 | 8 | 0.992178224139345 |
| chr17-24562356-24562857 | 6 | 0.942072951316711 |
| chr7-118191123-118191624 | 8 | 0.888402723709189 |
| chr13-94448240-94448741 | 8 | 0.864876416141289 |
| chr7-63700985-63701486 | 3 | 0.963576881486014 |
| chr17-12703829-12704330 | 3 | 0.990013492048584 |
| chr20-36506813-36507314 | 6 | 0.983559464165036 |
| chr3-106939050-106939551 | 4 | 0.817617047681865 |

| | | |
|---|---|---|
| chr8-104761609-104762110 | 6 | 0.818159075363391 |
| chr1-42746385-42746886 | 8 | 0.992349223726756 |
| chr12-45093461-45093962 | 8 | 0.815059617000725 |
| chr13-98627152-98627653 | 4 | 0.9023962255871 |
| chr11-36221212-36221713 | 3 | 0.995620153138524 |
| chr6-69397774-69398275 | 5 | 0.957957576887187 |
| chr15-53775656-53776157 | 3 | 0.988013724258845 |
| chr14-82422166-82422667 | 6 | 0.972563888150864 |
| chr2-247352413-247352914 | 5 | 0.959110332012085 |
| chr3-40869605-40870106 | 5 | 0.92363639343092 |
| chr4-175065728-175066229 | 6 | 0.991206925481043 |
| chr13-30859730-30860231 | 4 | 0.830225219136456 |
| chr1-107769499-107770000 | 6 | 0.992504103471873 |
| chr6-30151396-30151897 | 3 | 0.992527866754862 |
| chr13-30787329-30787830 | 8 | 0.995067664035511 |
| chr2-198513423-198513924 | 3 | 0.99593553545041 |
| chr17-44623728-44624229 | 2 | 0.895858288268349 |
| chr4-15905270-15905771 | 3 | 0.949168176579027 |
| chr2-148013013-148013514 | 3 | 0.960894009453918 |
| chr10-104229491-104229992 | 6 | 0.959013755140471 |
| chr10-69631161-69631662 | 8 | 0.930454603493609 |
| chr3-124324842-124325343 | 8 | 0.952359060206323 |
| chr1-212317004-212317505 | 6 | 0.991206925481043 |
| chr1-220178827-220179328 | 4 | 0.817105436664691 |
| chr4-66160155-66160656 | 8 | 0.889463510134396 |
| chr8-102575365-102575866 | 8 | 0.827608587786861 |
| chr13-106521685-106522186 | 3 | 0.98797519865479 |
| chr2-3285135-3285636 | 3 | 0.973753900185531 |
| chr8-29006040-29006541 | 4 | 0.920431846808207 |
| chr13-36465250-36465751 | 3 | 0.972367090611746 |
| chr8-68905311-68905812 | 1 | 0.860001996691763 |
| chr1-12189185-12189686 | 2 | 0.950851290919703 |
| chr2-67497942-67498443 | 8 | 0.983416589766416 |
| chr6-69476907-69477408 | 8 | 0.923789446483016 |
| chr18-4235325-4235826 | 8 | 0.930691437821227 |
| chr12-38380542-38381043 | 5 | 0.865421557679708 |
| chr2-46321250-46321751 | 6 | 0.993183757411015 |
| chr17-38751211-38751712 | 5 | 0.983919978840557 |
| chr1-109395675-109396176 | 8 | 0.825899625709686 |
| chr4-143379410-143379911 | 3 | 0.992083187984098 |
| chr12-20090918-20091419 | 7 | 0.836546364367928 |
| chr8-55448200-55448701 | 8 | 0.929272976886326 |
| chr20-36953324-36953825 | 8 | 0.933900976944438 |

| | | |
|---|---|---|
| chr15-29881060-29881561 | 8 | 0.815720231754169 |
| chr8-106109784-106110285 | 8 | 0.826081885038455 |
| chr18-11904768-11905269 | 3 | 0.99593553545041 |
| chr16-59003384-59003885 | 1 | 0.873390053878288 |
| chr20-36806807-36807308 | 3 | 0.995940367101444 |
| chr11-63877218-63877719 | 6 | 0.970703663274145 |
| chr8-5822688-5823189 | 3 | 0.905962225271067 |
| chr5-137716047-137716548 | 6 | 0.994552293653906 |
| chr20-31699218-31699719 | 7 | 0.921220392772741 |
| chr12-44479124-44479625 | 6 | 0.878556536316849 |
| chr6-119818718-119819219 | 5 | 0.981701944254491 |
| chr8-55064722-55065223 | 5 | 0.973505278989607 |
| chr9-45046662-45047163 | 5 | 0.919979627469248 |
| chr14-79831979-79832480 | 8 | 0.96040570436612 |
| chr10-37485607-37486108 | 8 | 0.823059886414228 |
| chr1-55656440-55656941 | 2 | 0.874873314765022 |
| chr5-145774810-145775311 | 8 | 0.821677609459424 |
| chr7-71380147-71380648 | 5 | 0.852193605501278 |
| chr3-122121172-122121673 | 6 | 0.932288420222738 |
| chr4-109826973-109827474 | 2 | 0.931001046156882 |
| chr10-94755943-94756444 | 8 | 0.855262913091918 |
| chr1-76718982-76719483 | 8 | 0.988442206133164 |
| chr7-61196287-61196788 | 1 | 0.900547703928514 |
| chr14-102092603-102093104 | 3 | 0.996893226829662 |
| chr9-30222580-30223081 | 3 | 0.995114704191427 |
| chr16-43866176-43866677 | 5 | 0.855932179895307 |
| chr8-78506712-78507213 | 8 | 0.912097263897208 |
| chr2-24302978-24303479 | 6 | 0.997053579287151 |
| chr1-160670411-160670912 | 3 | 0.870733862061607 |
| chr1-81271873-81272374 | 5 | 0.901536488969406 |
| chr4-96351391-96351892 | 8 | 0.977463811112572 |
| chr19-29858939-29859440 | 6 | 0.995815136968356 |
| chr1-135910516-135911017 | 5 | 0.92573100617634 |
| chr15-96014681-96015182 | 3 | 0.996062237441088 |
| chr14-5433164-5433665 | 8 | 0.939109965308385 |
| chr12-35757998-35758499 | 2 | 0.878003563001385 |
| chr13-98348360-98348861 | 4 | 0.964025115920817 |
| chr17-68792756-68793257 | 3 | 0.99593553545041 |
| chr8-92390072-92390573 | 3 | 0.852430072769947 |
| chr4-124301820-124302321 | 5 | 0.916097816877132 |
| chr19-44108619-44109120 | 8 | 0.994962813048223 |
| chr1-258037997-258038498 | 8 | 0.826357606102725 |
| chr5-132651349-132651850 | 5 | 0.920749460723337 |

| | | |
|---|---|---|
| chr1-138384331-138384832 | 8 | 0.855540247485498 |
| chr4-43000475-43000976 | 3 | 0.995448276485524 |
| chr5-133599393-133599894 | 8 | 0.974015359669842 |
| chr9-70938863-70939364 | 8 | 0.993493681128311 |
| chr9-55182502-55183003 | 3 | 0.99593553545041 |
| chr8-95583365-95583866 | 8 | 0.87441141928579 |
| chr6-103111382-103111883 | 3 | 0.976777021424731 |
| chr4-9568771-9569272 | 3 | 0.970997904379163 |
| chr7-114427376-114427877 | 2 | 0.901033973335908 |
| chr7-44868490-44868991 | 2 | 0.914896299132065 |
| chr6-43224449-43224950 | 4 | 0.909933482837981 |
| chr7-17556932-17557433 | 3 | 0.954167110919503 |
| chr12-37933449-37933950 | 5 | 0.830289543482197 |
| chr10-102637369-102637870 | 8 | 0.920067142866686 |
| chr12-9600628-9601129 | 8 | 0.915644549241975 |
| chr16-43484551-43485052 | 3 | 0.996844549082093 |
| chr1-146392620-146393121 | 8 | 0.995180486496983 |
| chr12-20873037-20873538 | 8 | 0.98189196458416 |
| chr16-50645022-50645523 | 8 | 0.973788768098631 |
| chr9-83110628-83111129 | 8 | 0.982216499039502 |
| chr12-10631403-10631904 | 8 | 0.971404046870842 |
| chr4-75673812-75674313 | 3 | 0.995443758777384 |
| chr19-38446241-38446742 | 8 | 0.970085369689624 |
| chr4-75941787-75942288 | 6 | 0.890932496270535 |
| chr6-21190194-21190695 | 5 | 0.927398823310555 |
| chr20-12904602-12905103 | 8 | 0.992362370151869 |
| chr3-19144895-19145396 | 8 | 0.879858531207098 |
| chr10-67180738-67181239 | 8 | 0.840510712283355 |
| chr12-24071946-24072447 | 5 | 0.978932268471773 |
| chr5-8150240-8150741 | 2 | 0.902378810514337 |
| chr18-55377166-55377667 | 2 | 0.84659627198146 |
| chr13-44235661-44236162 | 4 | 0.93267986471002 |
| chr8-12022182-12022683 | 3 | 0.996896889538922 |
| chr9-22116784-22117285 | 6 | 0.992997532466427 |
| chr2-59311023-59311524 | 5 | 0.975520649836134 |
| chr6-102512766-102513267 | 8 | 0.82464252587963 |
| chr2-113251630-113252131 | 1 | 0.944481587055094 |
| chr13-106528282-106528783 | 2 | 0.882812505091298 |
| chr8-57448268-57448769 | 8 | 0.963105004506859 |
| chr11-52520554-52521055 | 2 | 0.925584339936247 |
| chr10-18313569-18314070 | 7 | 0.904069963830387 |
| chr2-35715154-35715655 | 8 | 0.81513919955834 |
| chr14-72529215-72529716 | 6 | 0.991206925481043 |

| | | |
|---|---|---|
| chr10-92056743-92057244 | 5 | 0.987950439104842 |
| chr5-14658472-14658973 | 4 | 0.840651916753398 |
| chr6-49488711-49489212 | 1 | 0.904693085552731 |
| chr10-60718335-60718836 | 1 | 0.97137074623205 |
| chr1-122312007-122312508 | 3 | 0.975413824336045 |
| chr4-17864504-17865005 | 3 | 0.995430764757834 |
| chr10-43563880-43564381 | 1 | 0.875603906042278 |
| chr7-70799401-70799902 | 2 | 0.916652281617203 |
| chr2-101123992-101124493 | 8 | 0.820695759204371 |
| chr2-104623400-104623901 | 8 | 0.990480291367518 |
| chr1-158388812-158389313 | 3 | 0.836621408303927 |
| chr4-148305233-148305734 | 2 | 0.969603759400855 |
| chr11-59729489-59729990 | 8 | 0.975026415053891 |
| chrX-57465038-57465539 | 5 | 0.983919978840557 |
| chr7-23922460-23922961 | 6 | 0.992594504164124 |
| chr11-81563134-81563635 | 6 | 0.839386725724589 |
| chrX-33269499-33270000 | 8 | 0.991739207320445 |
| chr8-119376203-119376704 | 8 | 0.905650639008778 |
| chr15-20561623-20562124 | 8 | 0.962852243908794 |
| chr1-31860198-31860699 | 8 | 0.973465084268706 |
| chr14-85618456-85618957 | 6 | 0.927404652813512 |
| chr1-97386558-97387059 | 2 | 0.885480099037157 |
| chr12-6484387-6484888 | 2 | 0.955093011955656 |
| chr13-47743401-47743902 | 8 | 0.975433389320927 |
| chr20-43627543-43628044 | 8 | 0.987903112532265 |
| chr3-138172464-138172965 | 6 | 0.989970133445094 |
| chr7-47766994-47767495 | 6 | 0.993304994523469 |
| chr4-170183529-170184030 | 8 | 0.992120412919965 |
| chr3-54253648-54254149 | 5 | 0.957220771441708 |
| chr3-61099529-61100030 | 5 | 0.990998659880451 |
| chr11-78457742-78458243 | 5 | 0.92700297227535 |
| chr14-60862017-60862518 | 8 | 0.984880132963347 |
| chr6-87225949-87226450 | 6 | 0.925669305852227 |
| chr3-149033424-149033925 | 3 | 0.905111699446642 |
| chr4-54015236-54015737 | 4 | 0.939559891835861 |
| chr14-94130216-94130717 | 3 | 0.997010763373947 |
| chr19-34529844-34530345 | 8 | 0.947337350850971 |
| chr15-98412348-98412849 | 4 | 0.959113675581024 |
| chr9-64305197-64305698 | 8 | 0.98157523778424 |
| chr2-22751431-22751932 | 8 | 0.995228050916509 |
| chr8-47779126-47779627 | 6 | 0.857552912942718 |
| chr7-122808683-122809184 | 6 | 0.995347013089166 |
| chr5-83184675-83185176 | 3 | 0.995455431061296 |

| | | |
|---|---|---|
| chr2-220645383-220645884 | 8 | 0.975786762341434 |
| chr1-219418857-219419358 | 1 | 0.8235540043776 |
| chrX-150909882-150910383 | 3 | 0.912567498378933 |
| chr14-98379666-98380167 | 4 | 0.932339724271835 |
| chr10-35334834-35335335 | 3 | 0.984471374213276 |
| chr16-46285326-46285827 | 4 | 0.958592484358292 |
| chr10-27986970-27987471 | 7 | 0.818774997010958 |
| chr1-32950476-32950977 | 4 | 0.8345702794628 |
| chr17-25155161-25155662 | 6 | 0.992780404457086 |
| chr4-83844275-83844776 | 8 | 0.914841979023097 |
| chr1-177850881-177851382 | 2 | 0.878295942442803 |
| chr9-40597798-40598299 | 8 | 0.894437285264895 |
| chr6-20028411-20028912 | 3 | 0.995069854968598 |
| chr9-102589828-102590329 | 8 | 0.983182337674562 |
| chr6-119667185-119667686 | 8 | 0.834029163286268 |
| chr7-123375408-123375909 | 8 | 0.969600338711184 |
| chr1-117514750-117515251 | 2 | 0.844297084069276 |
| chr17-81814669-81815170 | 6 | 0.991954750009834 |
| chr14-101172402-101172903 | 5 | 0.990892352430047 |
| chr18-54632143-54632644 | 4 | 0.869722372144031 |
| chr8-89348560-89349061 | 6 | 0.988685851034687 |
| chr2-187832360-187832861 | 6 | 0.987184600776259 |
| chr19-26394725-26395226 | 5 | 0.987834774441003 |
| chr10-53246098-53246599 | 7 | 0.93749040881339 |
| chr8-112596813-112597314 | 5 | 0.88403169742771 |
| chr3-104487813-104488314 | 5 | 0.973586542631572 |
| chr18-34655327-34655828 | 8 | 0.891541601748853 |
| chr14-97048860-97049361 | 2 | 0.918184835753959 |
| chr3-91497860-91498361 | 8 | 0.920072603376131 |
| chr2-154397219-154397720 | 3 | 0.885585749054929 |
| chr3-125668037-125668538 | 6 | 0.947601407684254 |
| chr2-184391082-184391583 | 8 | 0.954211579168382 |
| chr11-25146827-25147328 | 6 | 0.994831682646368 |
| chr6-27737434-27737935 | 6 | 0.97961961904057 |
| chr5-66211465-66211966 | 6 | 0.978682889684968 |
| chrX-64363828-64364329 | 7 | 0.882358702436407 |
| chr1-87182815-87183316 | 2 | 0.956678283548029 |
| chr19-45230601-45231102 | 5 | 0.837342468557073 |
| chr2-33694338-33694839 | 3 | 0.996703111572272 |
| chr9-30535145-30535646 | 3 | 0.99593553545041 |
| chr2-225238841-225239342 | 8 | 0.929305367874572 |
| chr18-51608421-51608922 | 1 | 0.94553451254943 |
| chr3-101016668-101017169 | 5 | 0.979512197111429 |

| | | |
|---|---|---|
| chr6-41775121-41775622 | 8 | 0.907242594494932 |
| chr2-79939363-79939864 | 3 | 0.996877730109922 |
| chr4-77136770-77137271 | 8 | 0.993493681128311 |
| chr3-13603826-13604327 | 8 | 0.936357177236945 |
| chr10-10019669-10020170 | 4 | 0.959173528813235 |
| chr7-17297208-17297709 | 8 | 0.980144107861631 |
| chr12-17273158-17273659 | 2 | 0.830388458332456 |
| chr9-49762830-49763331 | 7 | 0.854219895058629 |
| chr3-108343211-108343712 | 8 | 0.948746892566629 |
| chr18-47881229-47881730 | 8 | 0.994501415480041 |
| chr9-24415270-24415771 | 3 | 0.995718032319545 |
| chr2-167286527-167287028 | 6 | 0.969983864219663 |
| chr2-233147689-233148190 | 3 | 0.996347983854157 |
| chr17-7203199-7203700 | 6 | 0.997103751921028 |
| chr17-15842968-15843469 | 8 | 0.97362839265253 |
| chr1-242820235-242820736 | 6 | 0.944731781511035 |
| chr5-148755751-148756252 | 4 | 0.834397787163337 |
| chr3-95301313-95301814 | 4 | 0.959283344773625 |
| chr1-120360150-120360651 | 8 | 0.994533651481151 |
| chr1-12346368-12346869 | 4 | 0.884045263370145 |
| chrX-74104303-74104804 | 5 | 0.919947974247089 |
| chr3-35504224-35504725 | 3 | 0.985072721108869 |
| chr15-65881418-65881919 | 3 | 0.99593553545041 |
| chr2-57526014-57526515 | 8 | 0.860371099097104 |
| chr13-38216166-38216667 | 5 | 0.863783242702357 |
| chr16-19974269-19974770 | 3 | 0.997760472672375 |
| chr7-122989526-122990027 | 6 | 0.992941526645466 |
| chr2-29933840-29934341 | 4 | 0.957007854681196 |
| chr2-44880367-44880868 | 8 | 0.91052561941569 |
| chr14-34307445-34307946 | 4 | 0.840190214530987 |
| chr10-74934139-74934640 | 8 | 0.839585445392104 |
| chr16-63158780-63159281 | 8 | 0.995248293852869 |
| chr1-257771218-257771719 | 8 | 0.836160056738811 |
| chr1-121127949-121128450 | 8 | 0.906818464306836 |
| chr20-48441044-48441545 | 1 | 0.942707691716262 |
| chr19-29603947-29604448 | 8 | 0.970272726745612 |
| chr17-60371186-60371687 | 8 | 0.981098974263158 |
| chr16-5629564-5630065 | 8 | 0.995245014470283 |
| chr11-78248105-78248606 | 3 | 0.9970370518132 |
| chr11-69700275-69700776 | 8 | 0.956035102059588 |
| chr15-2107724-2108225 | 6 | 0.978489251678467 |
| chr1-207236843-207237344 | 8 | 0.966428577365746 |
| chr14-101174057-101174558 | 5 | 0.991544323684295 |

| | | |
|---|---|---|
| chr3-11736120-11736621 | 8 | 0.916787769464593 |
| chr1-154039038-154039539 | 3 | 0.991402571138727 |
| chr16-62547619-62548120 | 6 | 0.981256820278151 |
| chr6-59688335-59688836 | 3 | 0.99593553545041 |
| chr6-120203600-120204101 | 1 | 0.88555137702008 |
| chr10-89053221-89053722 | 3 | 0.99676946451432 |
| chr14-24768463-24768964 | 3 | 0.963456504098733 |
| chr1-89557473-89557974 | 6 | 0.962772933247433 |
| chr2-151011396-151011897 | 2 | 0.847588125640555 |
| chr7-121963933-121964434 | 8 | 0.984101776694949 |
| chr10-79945675-79946176 | 8 | 0.936130040308834 |
| chr3-148729814-148730315 | 8 | 0.817839171498938 |
| chr1-26349262-26349763 | 8 | 0.848367295714979 |
| chr9-50428980-50429481 | 8 | 0.847758848467201 |
| chr8-56756219-56756720 | 4 | 0.919565260830822 |
| chr4-72317154-72317655 | 6 | 0.879390278483934 |
| chr1-42539674-42540175 | 5 | 0.989119525256711 |
| chr2-170870654-170871155 | 8 | 0.98076014362548 |
| chr4-2599402-2599903 | 8 | 0.893090495715837 |
| chr13-46938112-46938613 | 1 | 0.842322179866255 |
| chr10-13154574-13155075 | 8 | 0.984638522744969 |
| chr4-63105443-63105944 | 8 | 0.874656071264724 |
| chr1-220057663-220058164 | 2 | 0.92428272426245 |
| chr16-79152417-79152918 | 1 | 0.873416362576396 |
| chr8-95211056-95211557 | 8 | 0.954378206360365 |
| chr14-41862099-41862600 | 5 | 0.994864442031338 |
| chr17-71940552-71941053 | 5 | 0.989807518833273 |
| chr5-162330807-162331308 | 4 | 0.873200559430878 |
| chr14-10803238-10803739 | 8 | 0.969344621116184 |
| chr14-30443695-30444196 | 3 | 0.892563462083556 |
| chr12-19373793-19374294 | 5 | 0.885863232033418 |
| chr2-240368540-240369041 | 3 | 0.939402284794333 |
| chr8-63794097-63794598 | 3 | 0.995524056476941 |
| chr10-56394303-56394804 | 8 | 0.928569409038674 |
| chr16-37788889-37789390 | 8 | 0.864338743518942 |
| chr16-69989115-69989616 | 5 | 0.896514263271994 |
| chr7-31806411-31806912 | 5 | 0.958544968788073 |
| chr15-40921086-40921587 | 8 | 0.895843433188973 |
| chr6-110272040-110272541 | 8 | 0.884977270977384 |
| chr2-191434069-191434570 | 8 | 0.974897205710985 |
| chr1-194019704-194020205 | 8 | 0.985511753796842 |
| chr14-41743887-41744388 | 5 | 0.957878842888395 |
| chr5-159916270-159916771 | 6 | 0.985660417382684 |

| | | |
|---|---|---|
| chr14-81999673-82000174 | 4 | 0.851353935012164 |
| chr10-46576409-46576910 | 4 | 0.954892454225252 |
| chr6-128297346-128297847 | 6 | 0.977700240395201 |
| chr15-10151346-10151847 | 3 | 0.99593553545041 |
| chr15-95881158-95881659 | 8 | 0.89542589705963 |
| chr1-255494274-255494775 | 8 | 0.893609139030902 |
| chr2-113834737-113835238 | 3 | 0.98640891564846 |
| chr13-69244197-69244698 | 8 | 0.875964617555216 |
| chr3-156107524-156108025 | 1 | 0.911506235756419 |
| chr3-108803602-108804103 | 8 | 0.993425005879457 |
| chr15-18136443-18136944 | 6 | 0.992258980624612 |
| chr3-142743574-142744075 | 5 | 0.968287373643827 |
| chr5-12286523-12287024 | 5 | 0.983565491365917 |
| chr10-75496298-75496799 | 5 | 0.983919978840557 |
| chr20-42601757-42602258 | 2 | 0.840212821246084 |
| chr20-53797887-53798388 | 8 | 0.974095144100722 |
| chr8-70933710-70934211 | 8 | 0.824615840492898 |
| chr1-198718517-198719018 | 4 | 0.892607563199447 |
| chrX-151447545-151448046 | 6 | 0.958786985484682 |
| chr8-46781870-46782371 | 3 | 0.976322341856074 |
| chr16-16930693-16931194 | 6 | 0.990590383164384 |
| chr14-75500782-75501283 | 8 | 0.982481206343748 |
| chr5-107506647-107507148 | 6 | 0.846326511764235 |
| chr13-34902627-34903128 | 3 | 0.936069105207486 |
| chr15-296166-296667 | 4 | 0.971889206063868 |
| chr3-168279281-168279782 | 1 | 0.854003784333661 |
| chr2-81685188-81685689 | 3 | 0.996007026251852 |
| chr2-236111197-236111698 | 8 | 0.932448470787304 |
| chr7-59672757-59673258 | 5 | 0.975829072177202 |
| chr13-66705292-66705793 | 6 | 0.989365610653954 |
| chr3-89718344-89718845 | 8 | 0.969523859815527 |
| chr4-121493281-121493782 | 5 | 0.97570617304009 |
| chr17-41158641-41159142 | 1 | 0.889119936887018 |
| chr1-256200408-256200909 | 3 | 0.996080124609011 |
| chr3-69141290-69141791 | 5 | 0.892784881666341 |
| chr4-81510782-81511283 | 8 | 0.995243997298478 |
| chr9-108701060-108701561 | 4 | 0.817043527273891 |
| chr16-57471472-57471973 | 8 | 0.94724580545816 |
| chr5-91668137-91668638 | 3 | 0.996483628196343 |
| chr7-51191249-51191750 | 2 | 0.95370920667954 |
| chr9-736291-736792 | 8 | 0.839450594660936 |
| chr3-100112271-100112772 | 3 | 0.997222008256667 |
| chr3-34256076-34256577 | 4 | 0.904112265981866 |

| | | |
|---|---|---|
| chr12-14431313-14431814 | 7 | 0.938599323288798 |
| chr10-97830519-97831020 | 3 | 0.99593553545041 |
| chr8-78253016-78253517 | 4 | 0.890734559948265 |
| chr9-35531314-35531815 | 3 | 0.996873902913546 |
| chr17-72283843-72284344 | 4 | 0.824843062374753 |
| chr7-60985711-60986212 | 3 | 0.938793073150059 |
| chr8-106230365-106230866 | 8 | 0.993493681128311 |
| chr16-24996090-24996591 | 2 | 0.960418209424315 |
| chr4-149476589-149477090 | 5 | 0.877740236765216 |
| chr10-97436331-97436832 | 3 | 0.996360262992817 |
| chr5-159320761-159321262 | 3 | 0.966201479247291 |
| chr1-202572809-202573310 | 8 | 0.848579412595375 |
| chr1-121431404-121431905 | 3 | 0.996586211458065 |
| chr2-210666233-210666734 | 8 | 0.907675171392779 |
| chr2-34140862-34141363 | 8 | 0.992449711394062 |
| chr2-35190795-35191296 | 8 | 0.860165133687595 |
| chr15-29437987-29438488 | 1 | 0.819908152027788 |
| chr11-12104593-12105094 | 4 | 0.927375405300566 |
| chr1-143590229-143590730 | 1 | 0.949289413980551 |
| chr12-28612379-28612880 | 8 | 0.95886688334373 |
| chr16-80799274-80799775 | 1 | 0.960851193047357 |
| chr4-173883281-173883782 | 5 | 0.922113112251202 |
| chr20-39568933-39569434 | 6 | 0.970616125332377 |
| chr15-14570634-14571135 | 3 | 0.99593553545041 |
| chr5-60808533-60809034 | 8 | 0.935087807330676 |
| chr2-87933570-87934071 | 3 | 0.99593553545041 |
| chr6-45455710-45456211 | 4 | 0.817399359044828 |
| chr17-931437-931938 | 8 | 0.878812021512004 |
| chr13-2585928-2586429 | 3 | 0.985116294226491 |
| chr16-72073310-72073811 | 5 | 0.979447822632635 |
| chr8-40275929-40276430 | 8 | 0.993493681128311 |
| chr11-66638390-66638891 | 5 | 0.937905858616566 |
| chr8-109323562-109324063 | 4 | 0.903882399131265 |
| chr12-37081631-37082132 | 8 | 0.878162951691263 |
| chr18-61725196-61725697 | 3 | 0.947309522137735 |
| chr12-9986507-9987008 | 8 | 0.993493681128311 |
| chr7-18214943-18215444 | 3 | 0.991808635302263 |
| chr14-35748820-35749321 | 2 | 0.951996411026712 |
| chr2-2805625-2806126 | 3 | 0.997081370540384 |
| chr19-23664884-23665385 | 8 | 0.962289233828731 |
| chr4-126501926-126502427 | 1 | 0.859951157643512 |
| chr4-18197804-18198305 | 8 | 0.963571129090214 |
| chr5-18872759-18873260 | 8 | 0.942857688605922 |

| | | |
|---|---|---|
| chr15-39287324-39287825 | 3 | 0.982084345126225 |
| chr11-45697742-45698243 | 8 | 0.984934778252571 |
| chr1-165219864-165220365 | 2 | 0.916377017648055 |
| chr9-89144299-89144800 | 4 | 0.839968246171932 |
| chr10-97883534-97884035 | 2 | 0.859070770616795 |
| chr2-221184414-221184915 | 6 | 0.915886738974416 |
| chr18-44747058-44747559 | 1 | 0.916684941734754 |
| chr1-172014681-172015182 | 1 | 0.875031725804446 |
| chr4-43049531-43050032 | 6 | 0.837229194818836 |
| chr10-65648707-65649208 | 6 | 0.994436946669374 |
| chr7-46342414-46342915 | 2 | 0.952431838162654 |
| chr3-138088390-138088891 | 6 | 0.98785857139308 |
| chr10-9854512-9855013 | 3 | 0.994548421062392 |
| chr8-39177946-39178447 | 6 | 0.992887340838313 |
| chr8-30981428-30981929 | 3 | 0.993312476496703 |
| chr13-42940614-42941115 | 5 | 0.991482990740515 |
| chr19-20463482-20463983 | 6 | 0.952707946295068 |
| chr12-35868116-35868617 | 6 | 0.961019111653904 |
| chr6-122518147-122518648 | 3 | 0.985099375186316 |
| chr6-97620444-97620945 | 8 | 0.957766031176639 |
| chr10-34404074-34404575 | 3 | 0.99593553545041 |
| chr4-48953070-48953571 | 6 | 0.880102819087116 |
| chr8-86501318-86501819 | 8 | 0.948056133578377 |
| chr9-99529712-99530213 | 8 | 0.866404035386132 |
| chr3-56235980-56236481 | 4 | 0.970936473518667 |
| chr2-111491736-111492237 | 8 | 0.990665453934231 |
| chr6-59404906-59405407 | 2 | 0.922556851386691 |
| chr15-93705025-93705526 | 5 | 0.989801877106322 |
| chr17-26573077-26573578 | 5 | 0.988438606429561 |
| chr18-58805146-58805647 | 4 | 0.892881677320097 |
| chr18-51720012-51720513 | 3 | 0.964967730758526 |
| chr17-57388869-57389370 | 2 | 0.950324634014666 |
| chr8-43259500-43260001 | 4 | 0.95177605877627 |
| chr7-111309378-111309879 | 3 | 0.998446434709396 |
| chr17-64241861-64242362 | 4 | 0.923872989140862 |
| chr1-169381826-169382327 | 8 | 0.970374300416302 |
| chr3-118525808-118526309 | 6 | 0.974840811254783 |
| chr5-122547287-122547788 | 5 | 0.96026826217531 |
| chr5-124797958-124798459 | 7 | 0.825017232050614 |
| chr3-129866563-129867064 | 8 | 0.993493681128311 |
| chr3-96845465-96845966 | 8 | 0.954174675749166 |
| chr7-14544493-14544994 | 8 | 0.968200565424709 |
| chr5-77127190-77127691 | 8 | 0.993130991512663 |

| | | |
|---|---|---|
| chr1-126528108-126528609 | 8 | 0.993493681128311 |
| chr10-73173895-73174396 | 8 | 0.891383124011011 |
| chr4-64854550-64855051 | 3 | 0.980061458564301 |
| chr9-58153522-58154023 | 3 | 0.99593553545041 |
| chr5-80500864-80501365 | 5 | 0.871880012728243 |
| chr13-20772838-20773339 | 5 | 0.952695364165939 |
| chr10-72081807-72082308 | 8 | 0.967300713415229 |
| chr4-37655136-37655637 | 1 | 0.95801496235297 |
| chr17-69807950-69808451 | 3 | 0.993638123052079 |
| chr14-78754966-78755467 | 4 | 0.960623344546824 |
| chr13-21090172-21090673 | 5 | 0.8291803304018 |
| chr5-53643034-53643535 | 6 | 0.991206925481043 |
| chr19-54232195-54232696 | 8 | 0.852702640824871 |
| chr18-62369082-62369583 | 2 | 0.88583508795705 |
| chr10-34642282-34642783 | 4 | 0.865195752404002 |
| chr1-10678905-10679406 | 3 | 0.99593553545041 |
| chr8-79267996-79268497 | 2 | 0.846813853451144 |
| chr12-9322917-9323418 | 5 | 0.862190075128319 |
| chr4-120171271-120171772 | 5 | 0.915441832577779 |
| chr17-51250465-51250966 | 8 | 0.980430324657562 |
| chr2-47582243-47582744 | 6 | 0.975711469609296 |
| chr13-102384422-102384923 | 5 | 0.834007198044538 |
| chr1-175246316-175246817 | 1 | 0.960866872469454 |
| chr9-70633844-70634345 | 8 | 0.993493681128311 |
| chr11-86055466-86055967 | 5 | 0.987236445300548 |
| chr5-72213478-72213979 | 5 | 0.983919978840557 |
| chr7-123283382-123283883 | 6 | 0.992880372787134 |
| chr9-69861210-69861711 | 4 | 0.97476899968332 |
| chr9-17630610-17631111 | 3 | 0.916061158203493 |
| chr17-6759608-6760109 | 8 | 0.936041959135224 |
| chr18-30885298-30885799 | 3 | 0.975135910101772 |
| chr15-48261692-48262193 | 6 | 0.984880990652919 |
| chr10-19053438-19053939 | 3 | 0.99357191179718 |
| chr3-2310660-2311161 | 3 | 0.98160518241208 |
| chr9-62688615-62689116 | 5 | 0.90857648526579 |
| chr12-15259827-15260328 | 3 | 0.893374511084688 |
| chr6-15875177-15875678 | 8 | 0.945272135929636 |
| chr5-15013061-15013562 | 3 | 0.909921252053481 |
| chr9-29831073-29831574 | 8 | 0.981621939558881 |
| chr4-67837404-67837905 | 6 | 0.993207006622967 |
| chr7-127126188-127126689 | 6 | 0.99242596610993 |
| chr1-172934854-172935355 | 5 | 0.977472607195906 |
| chr1-49918430-49918931 | 3 | 0.99628644413609 |

| | | |
|---|---|---|
| chr5-143554655-143555156 | 4 | 0.931242721109359 |
| chr4-71458297-71458798 | 1 | 0.909591851868803 |
| chr20-43196787-43197288 | 8 | 0.989209891125309 |
| chr4-123791836-123792337 | 8 | 0.971335084025729 |
| chr18-75551973-75552474 | 8 | 0.995243605301839 |
| chr3-153208965-153209466 | 8 | 0.907397387063963 |
| chr9-29382060-29382561 | 6 | 0.993214364471501 |
| chr10-48731742-48732243 | 8 | 0.994488107555508 |
| chr11-67037948-67038449 | 3 | 0.849431553478135 |
| chr3-102619420-102619921 | 8 | 0.952793366813173 |
| chr7-11284005-11284506 | 8 | 0.962783206073237 |
| chr3-93312813-93313314 | 7 | 0.888206052012457 |
| chr16-81635657-81636158 | 7 | 0.924653207158244 |
| chr2-218776116-218776617 | 5 | 0.991929576511051 |
| chr17-7469752-7470253 | 6 | 0.983333941848679 |
| chr10-45922022-45922523 | 2 | 0.938897415035518 |
| chr3-147262364-147262865 | 5 | 0.925533443899383 |
| chr2-129335921-129336422 | 6 | 0.991206925481043 |
| chr3-65003013-65003514 | 8 | 0.943088990291071 |
| chr4-121507642-121508143 | 8 | 0.921227476234802 |
| chr3-22277901-22278402 | 8 | 0.98667251033888 |
| chr1-185988817-185989318 | 3 | 0.995272527069535 |
| chr19-48230501-48231002 | 5 | 0.993215813683507 |
| chr4-13858214-13858715 | 7 | 0.949839533996394 |
| chr13-46418424-46418925 | 2 | 0.949729440974336 |
| chr5-59891807-59892308 | 5 | 0.979704775272941 |
| chr5-130640744-130641245 | 6 | 0.87182643081595 |
| chr15-31995995-31996496 | 6 | 0.986746490839237 |
| chr13-77370098-77370599 | 8 | 0.982965339520593 |
| chr1-192573211-192573712 | 2 | 0.961304724998958 |
| chr10-79698084-79698585 | 8 | 0.833182084994676 |
| chr8-102641503-102642004 | 8 | 0.973680722615934 |
| chr1-157102101-157102602 | 5 | 0.983070662306076 |
| chr19-31199862-31200363 | 3 | 0.99593553545041 |
| chr2-216983213-216983714 | 3 | 0.996897347404138 |
| chr10-44035614-44036115 | 8 | 0.819874533911602 |
| chr3-56461975-56462476 | 1 | 0.83963558144679 |
| chr20-10882655-10883156 | 6 | 0.991432535496177 |
| chr2-221541710-221542211 | 8 | 0.949264588045795 |
| chr5-58608728-58609229 | 8 | 0.974808941800421 |
| chr13-38843026-38843527 | 5 | 0.989853098992835 |
| chr16-74206779-74207280 | 8 | 0.86323172077251 |
| chr10-17206792-17207293 | 7 | 0.842038579733513 |

| | | |
|---|---|---|
| chr14-32157106-32157607 | 7 | 0.852029280127631 |
| chr9-65028131-65028632 | 6 | 0.972806156040258 |
| chr3-157843419-157843920 | 4 | 0.820285639338514 |
| chr7-128797867-128798368 | 8 | 0.866631240376486 |
| chr7-11161272-11161773 | 5 | 0.988919780239301 |
| chr13-68008574-68009075 | 1 | 0.89261884446938 |
| chr3-34098026-34098527 | 8 | 0.995248877971997 |
| chr11-48627858-48628359 | 5 | 0.931923965154614 |
| chr1-228017938-228018439 | 5 | 0.991133906490909 |
| chr8-10763293-10763794 | 8 | 0.992412695674001 |
| chr13-52538850-52539351 | 3 | 0.99561409512842 |
| chr7-59155334-59155835 | 8 | 0.95864473154366 |
| chr2-171057807-171058308 | 7 | 0.968412071780893 |
| chr4-160471230-160471731 | 4 | 0.958803626268144 |
| chr19-44684507-44685008 | 5 | 0.986062997913894 |
| chr7-90514671-90515172 | 5 | 0.991739003278754 |
| chr7-103384969-103385470 | 3 | 0.99593553545041 |
| chr4-65635505-65636006 | 3 | 0.996416485221169 |
| chr4-181211468-181211969 | 8 | 0.993493681128311 |
| chr6-101945620-101946121 | 8 | 0.882708021076732 |
| chr13-28140756-28141257 | 7 | 0.837001581378254 |
| chr15-5868644-5869145 | 6 | 0.98054 |
| chr18-31063832-31064333 | 2 | 0.950828858354029 |
| chr11-83599426-83599927 | 5 | 0.984315803011563 |
| chr7-62551631-62552132 | 2 | 0.931482461740605 |
| chr3-6395299-6395800 | 3 | 0.99630974520581 |
| chr17-28285091-28285592 | 3 | 0.996887814936809 |
| chr14-80849292-80849793 | 5 | 0.84523540238052 |
| chr1-178186059-178186560 | 6 | 0.992814131233246 |
| chr4-85631346-85631847 | 1 | 0.949843035415182 |
| chr1-117430940-117431441 | 3 | 0.83082188698837 |
| chr9-39304178-39304679 | 8 | 0.993493681128311 |
| chr20-43430762-43431263 | 3 | 0.870343982494983 |
| chr2-10929386-10929887 | 3 | 0.992675801153738 |
| chr2-233254399-233254900 | 8 | 0.932938387551323 |
| chr4-92441348-92441849 | 3 | 0.878939672591798 |
| chr3-41461724-41462225 | 3 | 0.99319236041579 |
| chr11-12966661-12967162 | 8 | 0.838928145651811 |
| chr1-188774854-188775355 | 8 | 0.968194408876689 |
| chr8-111936003-111936504 | 3 | 0.992081253022961 |
| chr7-89744026-89744527 | 4 | 0.904597813531694 |
| chrX-9583970-9584471 | 4 | 0.924457993940277 |
| chr17-23158776-23159277 | 1 | 0.818295159517976 |

| | | |
|---|---|---|
| chr15-76033355-76033856 | 8 | 0.876781715962774 |
| chr1-89320846-89321347 | 3 | 0.984545628854109 |
| chr4-41410396-41410897 | 6 | 0.994930193701681 |
| chr1-84876254-84876755 | 3 | 0.921182913616454 |
| chr4-174423060-174423561 | 5 | 0.992082802440933 |
| chr11-23802432-23802933 | 1 | 0.889484561049119 |
| chr3-68833856-68834357 | 1 | 0.93885092510239 |
| chr10-37783877-37784378 | 8 | 0.843110597223499 |
| chr1-43273404-43273905 | 7 | 0.848286348502568 |
| chr4-107056067-107056568 | 3 | 0.979605803883369 |
| chr1-206043376-206043877 | 5 | 0.929674120048956 |
| chr15-52884486-52884987 | 6 | 0.971284990552113 |
| chr2-10701959-10702460 | 3 | 0.992461896708067 |
| chr17-72945300-72945801 | 3 | 0.99794416464115 |
| chr2-33329482-33329983 | 5 | 0.983919978840557 |
| chr1-77290813-77291314 | 2 | 0.926805873899416 |
| chr18-74658703-74659204 | 7 | 0.946284756477139 |
| chr8-42183101-42183602 | 4 | 0.830992752998136 |
| chr6-101366837-101367338 | 6 | 0.910242099543997 |
| chr7-52387303-52387804 | 1 | 0.941374690608156 |
| chr7-110288662-110289163 | 4 | 0.942885314050382 |
| chr7-128935119-128935620 | 8 | 0.918015994996924 |
| chr20-45251926-45252427 | 2 | 0.903089367069413 |
| chr1-250778731-250779232 | 8 | 0.949322329462267 |
| chr18-4598341-4598842 | 8 | 0.990419705491589 |
| chr2-67043082-67043583 | 3 | 0.922801762395249 |
| chr19-16446631-16447132 | 8 | 0.909369434618705 |
| chr11-57742107-57742608 | 1 | 0.885590939216568 |
| chrX-102201797-102202298 | 3 | 0.958634602518384 |
| chr3-152770118-152770619 | 3 | 0.985629310764732 |
| chr5-122855969-122856470 | 4 | 0.867353767939146 |
| chr7-102720654-102721155 | 2 | 0.821656111975133 |
| chr9-6730342-6730843 | 2 | 0.902814338410309 |
| chr8-116496159-116496660 | 2 | 0.9513214507412 |
| chr3-100634396-100634897 | 6 | 0.992889319152502 |
| chr7-10251842-10252343 | 8 | 0.973995689598676 |
| chr8-88156416-88156917 | 5 | 0.892914359657093 |
| chr5-14883325-14883826 | 1 | 0.926262107809338 |
| chr2-232573472-232573973 | 6 | 0.988852836909566 |
| chr18-68622747-68623248 | 6 | 0.996850189874423 |
| chr10-45685650-45686151 | 7 | 0.942790742647976 |
| chr15-92307006-92307507 | 4 | 0.891825727031152 |
| chr1-130925491-130925992 | 8 | 0.90138882831071 |

| | | |
|---|---|---|
| chr5-13387998-13388499 | 2 | 0.93504715504402 |
| chr16-25585793-25586294 | 2 | 0.933451921367431 |
| chr7-29557404-29557905 | 8 | 0.987313564041556 |
| chr8-51307574-51308075 | 3 | 0.993371956051292 |
| chr7-50639745-50640246 | 3 | 0.996851304406847 |
| chr14-100263818-100264319 | 6 | 0.993220336965468 |
| chr5-3149331-3149832 | 1 | 0.844343574381482 |
| chr3-36546738-36547239 | 8 | 0.952607412753826 |
| chr18-59323050-59323551 | 1 | 0.887143050437786 |
| chr5-21548781-21549282 | 8 | 0.835757776739664 |
| chr5-49667634-49668135 | 3 | 0.987855986239592 |
| chr8-113899680-113900181 | 5 | 0.991092319078656 |
| chr16-55520678-55521179 | 5 | 0.981473276010259 |
| chr17-10391784-10392285 | 8 | 0.984838293772457 |
| chr10-49679082-49679583 | 6 | 0.99656317047143 |
| chr12-8499881-8500382 | 5 | 0.983919978840557 |
| chr16-25735903-25736404 | 3 | 0.995387310687829 |
| chr7-46003684-46004185 | 8 | 0.972113740989771 |
| chr19-114700-115201 | 8 | 0.932293588303734 |
| chr5-146764622-146765123 | 7 | 0.932957189350904 |
| chr2-184594783-184595284 | 3 | 0.962256218423682 |
| chr7-98978282-98978783 | 2 | 0.946623033604052 |
| chr17-18053173-18053674 | 8 | 0.848773006594699 |
| chr11-31538959-31539460 | 8 | 0.967034810974943 |
| chrX-145025675-145026176 | 3 | 0.99593553545041 |
| chr18-56800065-56800566 | 7 | 0.915695259836032 |
| chr11-43792544-43793045 | 6 | 0.993433115984242 |
| chr8-11018928-11019429 | 8 | 0.92681046354256 |
| chr3-88400391-88400892 | 8 | 0.817961076667351 |
| chr3-117193788-117194289 | 3 | 0.996279905245101 |
| chr4-109082072-109082573 | 8 | 0.986246585607901 |
| chr19-41967597-41968098 | 3 | 0.847489841058574 |
| chr3-37396743-37397244 | 4 | 0.971832020370852 |
| chr1-61103289-61103790 | 2 | 0.85135866732818 |
| chr16-59856515-59857016 | 3 | 0.992185655008656 |
| chr6-67630548-67631049 | 3 | 0.993256476724116 |
| chr8-59911204-59911705 | 2 | 0.919593020640207 |
| chr8-57948690-57949191 | 8 | 0.86998625881723 |
| chr5-41503123-41503624 | 3 | 0.996283568154913 |
| chr6-19591373-19591874 | 6 | 0.885694996712843 |
| chr10-104202518-104203019 | 4 | 0.912644848481531 |
| chr20-29942339-29942840 | 8 | 0.940327020224041 |
| chr20-39446236-39446737 | 3 | 0.994588945537182 |

| | | |
|---|---|---|
| chr17-83024405-83024906 | 1 | 0.959678560023341 |
| chr16-57940678-57941179 | 8 | 0.910097078526659 |
| chr13-51027748-51028249 | 6 | 0.979990041039635 |
| chr8-52157245-52157746 | 6 | 0.991206925481043 |
| chr8-63235083-63235584 | 8 | 0.930917458581106 |
| chr14-61440801-61441302 | 3 | 0.996318088801004 |
| chr8-103112900-103113401 | 4 | 0.926383192712954 |
| chr3-126663608-126664109 | 3 | 0.982652339115002 |
| chr8-19814561-19815062 | 8 | 0.925236412272953 |
| chr14-86746462-86746963 | 3 | 0.993419379445788 |
| chr1-16927402-16927903 | 7 | 0.940127486829979 |
| chr8-32192671-32193172 | 5 | 0.934920196247748 |
| chr8-107340681-107341182 | 8 | 0.912382588646092 |
| chr3-53758758-53759259 | 4 | 0.827786083357977 |
| chr13-100461320-100461821 | 3 | 0.997217348945263 |
| chr14-97083038-97083539 | 8 | 0.882965054719447 |
| chr10-36608610-36609111 | 2 | 0.931166133617466 |
| chr6-123519249-123519750 | 4 | 0.975270665235187 |
| chr4-9288166-9288667 | 8 | 0.980426682587228 |
| chr4-68881610-68882111 | 8 | 0.995143950747246 |
| chr11-9756515-9757016 | 8 | 0.943044711325474 |
| chr14-17894607-17895108 | 8 | 0.894975202315806 |
| chr20-43937797-43938298 | 3 | 0.996059076377776 |
| chr7-76128961-76129462 | 3 | 0.986796187264149 |
| chr19-7655327-7655828 | 6 | 0.877684239778547 |
| chr20-1318082-1318583 | 8 | 0.913028388304391 |
| chr13-96859267-96859768 | 8 | 0.936146077821958 |
| chr15-1006755-1007256 | 4 | 0.847195506876132 |
| chr3-16687788-16688289 | 5 | 0.943814869932954 |
| chr6-28698709-28699210 | 3 | 0.973154391671434 |
| chr17-53400818-53401319 | 2 | 0.818212050352169 |
| chr1-120318841-120319342 | 8 | 0.987997246158097 |
| chr1-124480228-124480729 | 8 | 0.986526720671404 |
| chr4-62180624-62181125 | 5 | 0.881408252975546 |
| chr7-116031887-116032388 | 8 | 0.852195424434466 |
| chr15-11786109-11786610 | 3 | 0.997142781520563 |
| chr3-38332864-38333365 | 3 | 0.905340572890826 |
| chr3-11429300-11429801 | 8 | 0.955567519476215 |
| chr17-307980-308481 | 3 | 0.985364978425154 |
| chr15-45030904-45031405 | 8 | 0.851103508721927 |
| chr3-161894591-161895092 | 8 | 0.980204522060699 |
| chr4-15169557-15170058 | 5 | 0.95317097479844 |
| chr13-78414793-78415294 | 5 | 0.968750891670852 |

| | | |
|---|---|---|
| chr4-157119504-157120005 | 6 | 0.996835687012685 |
| chr9-105854221-105854722 | 8 | 0.957451433115733 |
| chr15-6011395-6011896 | 3 | 0.949750137062036 |
| chr14-34375731-34376232 | 8 | 0.845411394661784 |
| chr1-145761575-145762076 | 5 | 0.910469688621025 |
| chr10-63892839-63893340 | 5 | 0.950773673110236 |
| chr10-72639703-72640204 | 6 | 0.936626288042979 |
| chr2-33889962-33890463 | 4 | 0.906264164013894 |
| chr7-112399395-112399896 | 5 | 0.985933952148483 |
| chr8-78868114-78868615 | 8 | 0.870107139393582 |
| chr10-50395238-50395739 | 8 | 0.853990644805881 |
| chr11-11808494-11808995 | 2 | 0.877447785237012 |
| chr2-165797507-165798008 | 6 | 0.982426188508592 |
| chr15-31400684-31401185 | 4 | 0.880876973723345 |
| chr9-96948841-96949342 | 3 | 0.955079220185299 |
| chr16-15071193-15071694 | 5 | 0.982935589342089 |
| chr8-87256707-87257208 | 8 | 0.994851912928974 |
| chr7-21454185-21454686 | 8 | 0.872515462373085 |
| chr12-42882373-42882874 | 5 | 0.982875988352253 |
| chr2-40288507-40289008 | 5 | 0.820171254000443 |
| chr6-42463067-42463568 | 8 | 0.911320638405527 |
| chr19-15653597-15654098 | 4 | 0.878773958328864 |
| chr3-123280919-123281420 | 6 | 0.823719778015297 |
| chr2-137182478-137182979 | 8 | 0.941003615643705 |
| chr2-102164923-102165424 | 4 | 0.909105983606454 |
| chr5-104647623-104648124 | 3 | 0.993212285260766 |
| chr7-34435099-34435600 | 4 | 0.926454709030038 |
| chr8-58540753-58541254 | 8 | 0.993493681128311 |
| chr15-53708424-53708925 | 5 | 0.968281709822606 |
| chr9-3567879-3568380 | 3 | 0.994498140661478 |
| chr16-19458197-19458698 | 7 | 0.930315741543176 |
| chr9-23664736-23665237 | 2 | 0.862967058020892 |
| chr9-64068793-64069294 | 8 | 0.944022517302467 |
| chr17-82215311-82215812 | 8 | 0.911087672866612 |
| chr1-78866653-78867154 | 2 | 0.875865099912967 |
| chr5-110457324-110457825 | 1 | 0.858065793713523 |
| chr2-55022189-55022690 | 3 | 0.96390384321672 |
| chr2-164070272-164070773 | 8 | 0.994667029752205 |
| chr3-107215735-107216236 | 8 | 0.993493681128311 |
| chr4-57244094-57244595 | 8 | 0.880021169784244 |
| chr14-47705655-47706156 | 3 | 0.996855407378503 |
| chr9-6940373-6940874 | 3 | 0.938134655809053 |
| chr9-49570157-49570658 | 3 | 0.996885776731633 |

| | | |
|---|---|---|
| chr16-73626213-73626714 | 6 | 0.991206925481043 |
| chr1-207077539-207078040 | 7 | 0.853714901379743 |
| chr1-48792608-48793109 | 5 | 0.958293576994459 |
| chr1-257761175-257761676 | 8 | 0.948459404049069 |
| chr7-22471874-22472375 | 4 | 0.882557391227374 |
| chr4-36603359-36603860 | 3 | 0.999040199869997 |
| chr4-89873484-89873985 | 6 | 0.992730054457345 |
| chr18-55332452-55332953 | 1 | 0.879706240237182 |
| chr15-70553478-70553979 | 8 | 0.98514346769475 |
| chr1-82534583-82535084 | 6 | 0.991517809414829 |
| chr3-77538125-77538626 | 4 | 0.846163124176935 |
| chr11-80067071-80067572 | 6 | 0.907308790673052 |
| chr7-99246196-99246697 | 6 | 0.970491451506436 |
| chr10-21480119-21480620 | 6 | 0.991206925481043 |
| chr11-59044894-59045395 | 3 | 0.99244760210055 |
| chr2-235845965-235846466 | 8 | 0.84542975973644 |
| chr18-27581209-27581710 | 4 | 0.841607235921436 |
| chr18-17596335-17596836 | 3 | 0.995624653355086 |
| chr4-56979939-56980440 | 1 | 0.917512002623803 |
| chr10-49405222-49405723 | 3 | 0.957003886059291 |
| chr4-40957570-40958071 | 3 | 0.868466204218644 |
| chr13-30609344-30609845 | 8 | 0.8706516434889 |
| chr2-210184727-210185228 | 3 | 0.879136992586526 |
| chr6-119930261-119930762 | 5 | 0.980161619692621 |
| chr3-9430318-9430819 | 3 | 0.977533194238896 |
| chrX-2284724-2285225 | 6 | 0.988728306081095 |
| chr15-1215506-1216007 | 6 | 0.993192878794881 |
| chr17-26279307-26279808 | 4 | 0.953928770074314 |
| chr17-18178730-18179231 | 8 | 0.993881987089913 |
| chr9-17611416-17611917 | 3 | 0.938719661881804 |
| chr18-65129397-65129898 | 7 | 0.943360050948457 |
| chr10-80630216-80630717 | 2 | 0.913488670262923 |
| chr14-31148935-31149436 | 8 | 0.988628355495835 |
| chr4-45591468-45591969 | 4 | 0.844852234548007 |
| chr6-9408224-9408725 | 8 | 0.980478384384595 |
| chr7-105048807-105049308 | 8 | 0.992091772650707 |
| chr2-78702548-78703049 | 6 | 0.926531246784536 |
| chr9-61556472-61556973 | 3 | 0.990272922640519 |
| chr7-131876654-131877155 | 6 | 0.989682488952228 |
| chr2-142620515-142621016 | 4 | 0.898805325138325 |
| chr7-51911088-51911589 | 5 | 0.898516627746471 |
| chr9-85316046-85316547 | 4 | 0.89061810596006 |
| chr11-65339894-65340395 | 6 | 0.97591370360914 |

| | | |
|---|---|---|
| chr2-242444366-242444867 | 2 | 0.936282774917366 |
| chr5-60253548-60254049 | 3 | 0.997838449965887 |
| chr7-47108468-47108969 | 6 | 0.991206925481043 |
| chr14-11025885-11026386 | 3 | 0.996737422475446 |
| chr13-78542007-78542508 | 5 | 0.862207456400574 |
| chr1-185259081-185259582 | 8 | 0.897057380114354 |
| chr4-182208385-182208886 | 5 | 0.954688072269805 |
| chr2-21083990-21084491 | 6 | 0.99318235285881 |
| chr7-37478651-37479152 | 5 | 0.954128175110684 |
| chr4-149792220-149792721 | 6 | 0.888036659918091 |
| chr1-59013202-59013703 | 6 | 0.991206925481043 |
| chr8-117517910-117518411 | 3 | 0.988465769623828 |
| chr6-128295151-128295652 | 6 | 0.991206925481043 |
| chr1-200701162-200701663 | 8 | 0.859818334018063 |
| chr2-33235595-33236096 | 3 | 0.968381782832547 |
| chr5-140340471-140340972 | 8 | 0.816576960241992 |
| chr2-109926630-109927131 | 3 | 0.990961114516587 |
| chr5-165959091-165959592 | 6 | 0.993587391914085 |
| chr10-20437634-20438135 | 8 | 0.817523127884904 |
| chr12-39910957-39911458 | 3 | 0.99593553545041 |
| chr13-21933655-21934156 | 6 | 0.922757144650859 |
| chr1-140440913-140441414 | 3 | 0.930699096492788 |
| chr1-256283657-256284158 | 3 | 0.907970539098365 |
| chr15-26735511-26736012 | 3 | 0.994416583266237 |
| chr14-83132920-83133421 | 3 | 0.996158384547741 |
| chr14-98139901-98140402 | 3 | 0.99682032655819 |
| chr4-23909-24410 | 6 | 0.991206925481043 |
| chr1-121493293-121493794 | 5 | 0.9453345850505 |
| chr13-48474919-48475420 | 6 | 0.87902533813781 |
| chr7-17669613-17670114 | 6 | 0.983884809362905 |
| chr1-53966295-53966796 | 2 | 0.821754161647752 |
| chr13-98678048-98678549 | 8 | 0.986672332441423 |
| chr5-82559350-82559851 | 3 | 0.990243180646188 |
| chr3-99567289-99567790 | 5 | 0.833581948737106 |
| chr5-123488528-123489029 | 3 | 0.99593553545041 |
| chr1-21992037-21992538 | 5 | 0.990361819447843 |
| chr3-119525067-119525568 | 5 | 0.954659350009512 |
| chr5-48440365-48440866 | 3 | 0.996296863068534 |
| chr3-60193517-60194018 | 6 | 0.902422961538669 |
| chr19-14284998-14285499 | 1 | 0.897948558476419 |
| chr17-29052835-29053336 | 1 | 0.934482878140115 |
| chr18-17591315-17591816 | 3 | 0.94595004361201 |
| chr9-108124563-108125064 | 7 | 0.936535946832898 |

| | | |
|---|---|---|
| chr13-31014359-31014860 | 7 | 0.845700898589992 |
| chr10-69313801-69314302 | 8 | 0.989292755764891 |
| chr7-83811549-83812050 | 8 | 0.961467832766542 |
| chr3-7258289-7258790 | 6 | 0.969224557673245 |
| chr1-32191380-32191881 | 4 | 0.929225234153942 |
| chr15-46730350-46730851 | 2 | 0.941907441052766 |
| chr3-104707737-104708238 | 2 | 0.923907123648859 |
| chr19-39146434-39146935 | 2 | 0.952089516370837 |
| chr6-101429316-101429817 | 5 | 0.951182118266351 |
| chr10-16477771-16478272 | 8 | 0.885569262421681 |
| chr8-14507426-14507927 | 3 | 0.996771376398237 |
| chr4-28115314-28115815 | 4 | 0.927805657512537 |
| chr18-66348379-66348880 | 6 | 0.973119413363725 |
| chr1-188433837-188434338 | 8 | 0.993525220546207 |
| chr1-50783328-50783829 | 5 | 0.96641008058231 |
| chr13-43371586-43372087 | 6 | 0.901617385892199 |
| chr10-81607224-81607725 | 4 | 0.931353761699317 |
| chr19-25519954-25520455 | 6 | 0.983867981277849 |
| chr10-18840715-18841216 | 6 | 0.993078760768967 |
| chr9-100820401-100820902 | 6 | 0.863242153163559 |
| chr7-121139652-121140153 | 8 | 0.906170364050508 |
| chr7-131528685-131529186 | 4 | 0.919575810770695 |
| chr10-60693940-60694441 | 5 | 0.960609495994165 |
| chr7-98436526-98437027 | 6 | 0.987084244417287 |
| chr9-57476556-57477057 | 8 | 0.979096283754129 |
| chr1-175345833-175346334 | 8 | 0.97788623267949 |
| chr4-23067547-23068048 | 5 | 0.990294764608162 |
| chr1-1311939-1312440 | 4 | 0.931031828531027 |
| chrX-111328709-111329210 | 8 | 0.993493681128311 |
| chr13-34847461-34847962 | 3 | 0.995609096309321 |
| chr18-13328990-13329491 | 5 | 0.924176951577337 |
| chr1-50557273-50557774 | 2 | 0.956936103517248 |
| chr12-26516824-26517325 | 4 | 0.835090984401436 |
| chr10-74119485-74119986 | 3 | 0.995921814690859 |
| chr18-61224747-61225248 | 8 | 0.932096199761734 |
| chr7-96351717-96352218 | 2 | 0.907376952146601 |
| chr6-137582256-137582757 | 5 | 0.909742752445601 |
| chr5-117526443-117526944 | 8 | 0.986850205140325 |
| chr8-45044680-45045181 | 7 | 0.916696117422435 |
| chr19-54990515-54991016 | 8 | 0.97681027251897 |
| chr2-149067036-149067537 | 8 | 0.909905543594843 |
| chr3-108658504-108659005 | 3 | 0.857278921983903 |
| chr14-78577795-78578296 | 3 | 0.989353849217135 |

| | | |
|---|---|---|
| chr2-245542910-245543411 | 3 | 0.99593553545041 |
| chr3-30383226-30383727 | 8 | 0.993493681128311 |
| chr8-119435168-119435669 | 2 | 0.971399652818757 |
| chr15-47749291-47749792 | 6 | 0.991206925481043 |
| chr1-29154703-29155204 | 8 | 0.990888896915094 |
| chr1-247307786-247308287 | 8 | 0.970069736993113 |
| chrX-135826450-135826951 | 5 | 0.985596280138798 |
| chr9-11301075-11301576 | 2 | 0.830897949213658 |
| chr8-71043714-71044215 | 1 | 0.919747660129577 |
| chr14-47337766-47338267 | 1 | 0.919309442044673 |
| chr3-46755472-46755973 | 3 | 0.99387895837924 |
| chr2-157402452-157402953 | 6 | 0.983637990233309 |
| chr4-58648832-58649333 | 8 | 0.984772997561739 |
| chrX-102103088-102103589 | 6 | 0.970870856235827 |
| chr16-79044871-79045372 | 8 | 0.994699514862584 |
| chr4-181140622-181141123 | 4 | 0.929614602293723 |
| chr1-231636715-231637216 | 5 | 0.902199310568744 |
| chr19-28671328-28671829 | 8 | 0.956761498340595 |
| chr15-4843917-4844418 | 6 | 0.988356375885268 |
| chr17-7462118-7462619 | 3 | 0.998325436480714 |
| chr5-137791089-137791590 | 2 | 0.946613366709631 |
| chr2-193761544-193762045 | 6 | 0.925269109798497 |
| chr2-133342602-133343103 | 8 | 0.866343756552895 |
| chr14-13663087-13663588 | 8 | 0.979628621797536 |
| chr8-74154828-74155329 | 6 | 0.991206925481043 |
| chr2-104728325-104728826 | 6 | 0.986067780151949 |
| chr3-138594789-138595290 | 3 | 0.869475943910701 |
| chr5-160577155-160577656 | 6 | 0.904795427124437 |
| chr3-152016681-152017182 | 5 | 0.990482103912263 |
| chr5-144328448-144328949 | 1 | 0.958391679450441 |
| chr8-101588651-101589152 | 3 | 0.999034804502132 |
| chr5-100494208-100494709 | 8 | 0.995005673596238 |
| chr14-17505652-17506153 | 3 | 0.920835016806795 |
| chr16-22945037-22945538 | 8 | 0.975839750005279 |
| chr3-114829007-114829508 | 3 | 0.996864629785686 |
| chr13-71556998-71557499 | 3 | 0.990637278931022 |
| chr19-39838977-39839478 | 7 | 0.883414952435643 |
| chr10-65624177-65624678 | 8 | 0.979512068660338 |
| chr20-46109417-46109918 | 5 | 0.934273987237006 |
| chr13-43671490-43671991 | 8 | 0.986967328675813 |
| chr7-24585091-24585592 | 3 | 0.99593553545041 |
| chr3-32463084-32463585 | 3 | 0.952657750255591 |
| chr20-13398402-13398903 | 7 | 0.856297879240948 |

| | | |
|---|---|---|
| chr2-108555024-108555525 | 3 | 0.892888090635594 |
| chr5-92334186-92334687 | 3 | 0.94632193612834 |
| chr8-66617237-66617738 | 8 | 0.950677378232059 |
| chr3-154588368-154588869 | 3 | 0.944274923744535 |
| chr2-189512069-189512570 | 5 | 0.934346658172656 |
| chr5-166647633-166648134 | 8 | 0.845898926201606 |
| chr7-131652862-131653363 | 8 | 0.856613502568788 |
| chr6-123104260-123104761 | 8 | 0.993493681128311 |
| chr1-53093724-53094225 | 3 | 0.988847814263583 |
| chr8-23904358-23904859 | 4 | 0.879318167062758 |
| chr13-91033173-91033674 | 3 | 0.821479049698075 |
| chr10-70976890-70977391 | 3 | 0.976985682709673 |
| chr3-67379263-67379764 | 3 | 0.912537996819616 |
| chr4-58753810-58754311 | 6 | 0.948858234579663 |
| chr2-243798345-243798846 | 5 | 0.991184957255477 |
| chr5-9938489-9938990 | 3 | 0.988494726197429 |
| chr1-85203632-85204133 | 8 | 0.91938025229659 |
| chr14-88764573-88765074 | 6 | 0.991206925481043 |
| chr3-16659799-16660300 | 8 | 0.880850292205405 |
| chr8-54445689-54446190 | 7 | 0.863802969033817 |
| chr9-63226103-63226604 | 3 | 0.99461974698998 |
| chr1-190605967-190606468 | 8 | 0.985238635917008 |
| chr19-23156212-23156713 | 8 | 0.99524680627883 |
| chr2-243063497-243063998 | 6 | 0.96362283520478 |
| chr2-30726051-30726552 | 6 | 0.993736169604234 |
| chr10-91713570-91714071 | 5 | 0.985267966277133 |
| chr11-30987386-30987887 | 5 | 0.91789761021817 |
| chr3-62861682-62862183 | 8 | 0.980595140006527 |
| chr14-8786226-8786727 | 3 | 0.99593553545041 |
| chr6-111120067-111120568 | 2 | 0.909296146029004 |
| chr12-26361927-26362428 | 1 | 0.878567848329721 |
| chr10-22729070-22729571 | 3 | 0.922723930424366 |
| chr8-115108255-115108756 | 3 | 0.992874558279181 |
| chr15-81969464-81969965 | 6 | 0.993960756099246 |
| chr10-57734365-57734866 | 8 | 0.934034902299775 |
| chr16-2958004-2958505 | 2 | 0.83010295489367 |
| chr2-232362917-232363418 | 4 | 0.852277808572243 |
| chr1-150606959-150607460 | 3 | 0.951117306454311 |
| chr7-78123141-78123642 | 6 | 0.984161114960353 |
| chr7-20622014-20622515 | 8 | 0.973519559074563 |
| chr1-137138851-137139352 | 8 | 0.977352185314889 |
| chr17-20658951-20659452 | 5 | 0.985887398887293 |
| chr2-66051817-66052318 | 3 | 0.955030129519966 |

| | | |
|---|---|---|
| chr19-427091-427592 | 7 | 0.885132721128192 |
| chr9-11690185-11690686 | 6 | 0.976590654343408 |
| chr2-83511002-83511503 | 5 | 0.928051837260018 |
| chr5-71974273-71974774 | 3 | 0.994822492177849 |
| chr16-3458515-3459016 | 8 | 0.994929718700582 |
| chr5-20423510-20424011 | 1 | 0.840200712380858 |
| chr20-45631215-45631716 | 2 | 0.958673554599314 |
| chr4-178152412-178152913 | 4 | 0.871846122873958 |
| chr16-33118878-33119379 | 3 | 0.996548211281203 |
| chr14-42425424-42425925 | 8 | 0.858876555936528 |
| chr14-7612736-7613237 | 5 | 0.945766438910441 |
| chr7-64207488-64207989 | 2 | 0.826573400283648 |
| chr1-245034087-245034588 | 8 | 0.964519861032215 |
| chr2-33979452-33979953 | 3 | 0.989830988422047 |
| chr3-129476369-129476870 | 6 | 0.992330375107394 |
| chr11-28296179-28296680 | 1 | 0.855068279835849 |
| chr2-226186648-226187149 | 8 | 0.87170838731263 |
| chr16-72140340-72140841 | 3 | 0.983206930513854 |
| chr6-21501510-21502011 | 8 | 0.859901381145553 |
| chr20-35922536-35923037 | 8 | 0.838539440529703 |
| chr1-108024773-108025274 | 4 | 0.830510881446508 |
| chr12-32109039-32109540 | 8 | 0.932133499478113 |
| chr13-103903375-103903876 | 7 | 0.928638659863344 |
| chr5-130295599-130296100 | 8 | 0.863884742261909 |
| chr8-65272052-65272553 | 1 | 0.950747974670771 |
| chr2-115879742-115880243 | 3 | 0.99593553545041 |
| chr5-122961275-122961776 | 3 | 0.99593553545041 |
| chr17-52690981-52691482 | 5 | 0.962943677335828 |
| chr9-88853720-88854221 | 8 | 0.827585963139839 |
| chr7-121352854-121353355 | 4 | 0.873432786847575 |
| chr20-34971051-34971552 | 6 | 0.99239629986111 |
| chr13-66272447-66272948 | 8 | 0.994124065528926 |
| chr17-52798200-52798701 | 5 | 0.987078720611311 |
| chr2-112673030-112673531 | 4 | 0.877926994557241 |
| chr15-64316938-64317439 | 3 | 0.918134804598389 |
| chrX-50247319-50247820 | 3 | 0.992972296662505 |
| chr11-15581394-15581895 | 6 | 0.972397162607536 |
| chr17-72166520-72167021 | 6 | 0.956370133084458 |
| chr4-50213279-50213780 | 3 | 0.988215574311551 |
| chr15-83635167-83635668 | 5 | 0.983919978840557 |
| chr17-68758348-68758849 | 3 | 0.99593553545041 |
| chr6-106926768-106927269 | 6 | 0.968796177123345 |
| chr4-60239556-60240057 | 1 | 0.901735527727429 |

| | | |
|---|---|---|
| chr18-73555913-73556414 | 8 | 0.89910475730874 |
| chr11-33813517-33814018 | 5 | 0.986491355131835 |
| chr12-23004037-23004538 | 6 | 0.983994477769524 |
| chr10-67162078-67162579 | 3 | 0.895561086760885 |
| chr1-71685987-71686488 | 3 | 0.99593553545041 |
| chr13-78292242-78292743 | 8 | 0.995207278526654 |
| chr17-73705229-73705730 | 5 | 0.960561783113019 |
| chr15-85356873-85357374 | 3 | 0.99593553545041 |
| chr3-131651422-131651923 | 5 | 0.991357974645237 |
| chr17-75259481-75259982 | 5 | 0.990746473290419 |
| chr4-60232239-60232740 | 1 | 0.940630749055045 |
| chr15-74316677-74317178 | 8 | 0.993360565400698 |
| chr10-65635690-65636191 | 6 | 0.991206925481043 |
| chr1-193918659-193919160 | 8 | 0.989876640865756 |
| chr2-152825241-152825742 | 8 | 0.990424147175861 |
| chr17-41254871-41255372 | 5 | 0.927185580609898 |
| chr3-114862976-114863477 | 8 | 0.88252399133885 |
| chr9-27978950-27979451 | 3 | 0.99593553545041 |
| chr6-18065354-18065855 | 3 | 0.981482307428783 |
| chr18-58302882-58303383 | 8 | 0.862030486531705 |
| chr6-44192769-44193270 | 3 | 0.99593553545041 |
| chr8-115547594-115548095 | 8 | 0.978925490539533 |
| chr3-53262978-53263479 | 8 | 0.993493681128311 |
| chr18-78747093-78747594 | 6 | 0.846577583044786 |
| chr3-41866614-41867115 | 3 | 0.997476786801526 |
| chr7-26771824-26772325 | 5 | 0.973416295709778 |
| chr2-211642989-211643490 | 6 | 0.960952238291284 |
| chr1-101396578-101397079 | 2 | 0.882325770842386 |
| chr2-167400641-167401142 | 8 | 0.946532384495291 |
| chr17-22785739-22786240 | 3 | 0.846004310313684 |
| chr1-42340263-42340764 | 8 | 0.917659331370215 |
| chr2-94560407-94560908 | 3 | 0.995846198532283 |
| chr13-104149915-104150416 | 8 | 0.930303622674617 |
| chr13-82963377-82963878 | 4 | 0.884030299179437 |
| chr8-68080087-68080588 | 8 | 0.990574273171622 |
| chr14-32531363-32531864 | 8 | 0.894977769114623 |
| chr5-145280930-145281431 | 4 | 0.873166751239185 |
| chr2-113335626-113336127 | 5 | 0.946817533945969 |
| chr5-16756866-16757367 | 6 | 0.989917642512723 |
| chr1-219451153-219451654 | 1 | 0.950325329747113 |
| chr2-143698138-143698639 | 4 | 0.906680094866216 |
| chr14-95143343-95143844 | 1 | 0.93796766761576 |
| chr10-18816061-18816562 | 8 | 0.95979671943082 |

| | | |
|---|---|---|
| chr5-84681468-84681969 | 3 | 0.995924534943556 |
| chr12-40969655-40970156 | 8 | 0.980448909449023 |
| chr18-20250095-20250596 | 5 | 0.983919978840557 |
| chr16-48800921-48801422 | 3 | 0.995552263510682 |
| chr5-76843699-76844200 | 6 | 0.97894787932159 |
| chr6-42187973-42188474 | 5 | 0.979878693890859 |
| chr11-56237935-56238436 | 8 | 0.901885112708267 |
| chr2-136598686-136599187 | 8 | 0.904166006781795 |
| chr3-65460734-65461235 | 4 | 0.875722893768056 |
| chr3-12876432-12876933 | 6 | 0.979875668824858 |
| chr10-63439842-63440343 | 4 | 0.83607489185493 |
| chr6-92757738-92758239 | 7 | 0.831324995012239 |
| chr16-80973094-80973595 | 8 | 0.938073454798473 |
| chr8-115556066-115556567 | 8 | 0.968775417632391 |
| chr1-124464026-124464527 | 2 | 0.947250222583754 |
| chr2-89658513-89659014 | 3 | 0.990381286058722 |
| chr4-7848474-7848975 | 2 | 0.843591317954021 |
| chr1-165671333-165671834 | 3 | 0.99822280627762 |
| chr10-54450116-54450617 | 8 | 0.94746157276084 |
| chr2-115603479-115603980 | 8 | 0.993493681128311 |
| chr6-65579378-65579879 | 3 | 0.99593553545041 |
| chr2-107591668-107592169 | 3 | 0.99593553545041 |
| chr8-28173586-28174087 | 8 | 0.816707041347159 |
| chr2-172143084-172143585 | 6 | 0.991206925481043 |
| chr1-216587415-216587916 | 8 | 0.906630178575008 |
| chr3-16230697-16231198 | 8 | 0.94592646232169 |
| chr4-149758587-149759088 | 4 | 0.872220478499314 |
| chr7-74923165-74923666 | 5 | 0.987405145827243 |
| chr2-237879298-237879799 | 3 | 0.9762927159429 |
| chr2-205099859-205100360 | 3 | 0.996707936269173 |
| chr1-198265379-198265880 | 5 | 0.891016104810415 |
| chr11-32342232-32342733 | 8 | 0.945199770495402 |
| chr16-46369883-46370384 | 3 | 0.995554726929847 |
| chr10-87894958-87895459 | 8 | 0.979531441725351 |
| chr9-109176939-109177440 | 8 | 0.893734756767378 |
| chr6-102250275-102250776 | 6 | 0.985876608939811 |
| chr14-74327368-74327869 | 5 | 0.824837397882298 |
| chr4-82642650-82643151 | 4 | 0.85890385309429 |
| chr4-117073231-117073732 | 4 | 0.92848226817868 |
| chr2-44324879-44325380 | 8 | 0.831190901181159 |
| chr4-104545228-104545729 | 8 | 0.831264797263003 |
| chr1-219028149-219028650 | 8 | 0.892175704535135 |
| chr11-50596134-50596635 | 3 | 0.997715491341992 |

| | | |
|---|---|---|
| chr2-77159804-77160305 | 3 | 0.914455451735959 |
| chrX-79022284-79022785 | 6 | 0.991206925481043 |
| chr4-9360463-9360964 | 3 | 0.98863672456975 |
| chr2-146286279-146286780 | 3 | 0.84056352444764 |
| chr4-68579987-68580488 | 7 | 0.947648514465947 |
| chr4-141654958-141655459 | 6 | 0.952875365756835 |
| chr6-137066358-137066859 | 6 | 0.995437664307759 |
| chr3-56744621-56745122 | 6 | 0.883810333037502 |
| chr16-56172534-56173035 | 6 | 0.957296089837754 |
| chr5-143342237-143342738 | 6 | 0.993005227069393 |
| chr4-84499167-84499668 | 3 | 0.989062534780992 |
| chr3-53051014-53051515 | 5 | 0.89479506001388 |
| chr20-20851479-20851980 | 5 | 0.828559084349167 |
| chr9-55378210-55378711 | 4 | 0.90639214725245 |
| chr7-47841486-47841987 | 6 | 0.915772000339588 |
| chr15-58467577-58468078 | 3 | 0.995875741121172 |
| chr5-64937305-64937806 | 3 | 0.995757022317915 |
| chr14-77188991-77189492 | 8 | 0.950507101333644 |
| chr6-43835927-43836428 | 3 | 0.996583164498458 |
| chr13-84046414-84046915 | 6 | 0.963471542684957 |
| chr5-71230709-71231210 | 8 | 0.968834852148849 |
| chr1-220202855-220203356 | 1 | 0.929524380155955 |
| chr5-10660242-10660743 | 8 | 0.990798842371327 |
| chr9-61902252-61902753 | 8 | 0.985610470795226 |
| chr3-114957425-114957926 | 3 | 0.959270028862933 |
| chr5-99317579-99318080 | 3 | 0.996719523431279 |
| chr1-229050009-229050510 | 3 | 0.994717080969882 |
| chr3-102603189-102603690 | 4 | 0.926661649320321 |
| chr4-67441942-67442443 | 8 | 0.936308104790294 |
| chr7-29876583-29877084 | 3 | 0.997666293197799 |
| chr9-83401106-83401607 | 3 | 0.995814604894927 |
| chr1-186337635-186338136 | 3 | 0.994090247939285 |
| chr1-3276354-3276855 | 3 | 0.972768468980521 |
| chr8-31414268-31414769 | 8 | 0.977518762301788 |
| chr3-60844186-60844687 | 8 | 0.850474824340717 |
| chr13-100131782-100132283 | 8 | 0.950877963351659 |
| chr15-21609016-21609517 | 8 | 0.915442146466854 |
| chr3-38598240-38598741 | 3 | 0.900025464324126 |
| chr9-111876038-111876539 | 3 | 0.930339862840483 |
| chr10-62996388-62996889 | 3 | 0.874980399452662 |
| chr9-106902724-106903225 | 3 | 0.922595910378899 |
| chr16-22711122-22711623 | 3 | 0.994646728528655 |
| chr6-30751548-30752049 | 6 | 0.993226760886017 |

| | | |
|---|---|---|
| chr3-94090258-94090759 | 3 | 0.834389748402137 |
| chr13-103912330-103912831 | 8 | 0.990548124971545 |
| chr9-50526424-50526925 | 7 | 0.857902827017393 |
| chr14-33528710-33529211 | 3 | 0.945508905260724 |
| chr14-56742546-56743047 | 6 | 0.995090810591101 |
| chr1-191498823-191499324 | 3 | 0.996828879447509 |
| chr4-61454933-61455434 | 8 | 0.836943237928347 |
| chr16-44881741-44882242 | 4 | 0.876013425263287 |
| chr13-105584086-105584587 | 8 | 0.953424723659739 |
| chr1-96473264-96473765 | 4 | 0.867911176382225 |
| chr13-86716717-86717218 | 8 | 0.981912575424024 |
| chr19-13438293-13438794 | 5 | 0.898201678686214 |
| chr2-188853080-188853581 | 8 | 0.834118019358734 |
| chr2-134928220-134928721 | 6 | 0.99310842798224 |
| chr4-114996317-114996818 | 1 | 0.874353278484944 |
| chr5-50726537-50727038 | 3 | 0.980653266470173 |
| chr14-9120749-9121250 | 8 | 0.986570341445266 |
| chr16-5322749-5323250 | 5 | 0.843931026310326 |
| chr3-59511629-59512130 | 6 | 0.991718554891207 |
| chr4-127373691-127374192 | 6 | 0.854547868634802 |
| chr4-67021842-67022343 | 6 | 0.953777717779865 |
| chr19-46724638-46725139 | 5 | 0.989189576747312 |
| chr16-8401872-8402373 | 6 | 0.885684145138511 |
| chr5-63658440-63658941 | 8 | 0.98872326664934 |
| chr16-37738593-37739094 | 3 | 0.908850927347836 |
| chr17-64984300-64984801 | 1 | 0.878162232040481 |
| chr9-104489767-104490268 | 8 | 0.966907278290483 |
| chr5-162015317-162015818 | 3 | 0.984700679039855 |
| chrX-96321146-96321647 | 1 | 0.818426300028893 |
| chr18-1443639-1444140 | 4 | 0.836067342309804 |
| chr10-100353631-100354132 | 8 | 0.848009505456617 |
| chr18-74831480-74831981 | 6 | 0.94562896975511 |
| chr7-25996962-25997463 | 3 | 0.995022982837103 |
| chr19-33617065-33617566 | 1 | 0.822812285440876 |
| chr15-76184073-76184574 | 3 | 0.994490375925329 |
| chr4-72113361-72113862 | 2 | 0.955079810686952 |
| chr10-85565713-85566214 | 6 | 0.991867949511896 |
| chr8-78286535-78287036 | 8 | 0.892613868577175 |
| chr9-38239062-38239563 | 8 | 0.993493681128311 |
| chr5-142646798-142647299 | 5 | 0.989991914796402 |
| chr15-41315344-41315845 | 8 | 0.932347584045371 |
| chr2-230667849-230668350 | 8 | 0.968918004191067 |
| chr3-13941697-13942198 | 8 | 0.927090317548215 |

| | | |
|---|---|---|
| chr14-56890803-56891304 | 2 | 0.938641265641878 |
| chr2-186632746-186633247 | 8 | 0.984948899006401 |
| chr20-35763878-35764379 | 8 | 0.903701570414838 |
| chr17-45753298-45753799 | 3 | 0.997339387890769 |
| chr17-79140634-79141135 | 8 | 0.991284943754411 |
| chr20-36207557-36208058 | 8 | 0.937426841078628 |
| chr5-42858191-42858692 | 6 | 0.892833086955213 |
| chr1-106195245-106195746 | 6 | 0.983745742388889 |
| chr8-76735893-76736394 | 1 | 0.944504370813467 |
| chr8-21237971-21238472 | 2 | 0.931084421850854 |
| chr2-55760587-55761088 | 5 | 0.991619407414557 |
| chr2-173688703-173689204 | 8 | 0.846277712177265 |
| chr13-95197922-95198423 | 3 | 0.996667058505036 |
| chr9-11347311-11347812 | 8 | 0.993493681128311 |
| chr8-107389891-107390392 | 3 | 0.99593553545041 |
| chr9-90728866-90729367 | 8 | 0.951170994273425 |
| chr16-6100203-6100704 | 8 | 0.991008416167834 |
| chr1-199437101-199437602 | 8 | 0.994984333610293 |
| chr4-148642006-148642507 | 8 | 0.952803841054735 |
| chr17-64448918-64449419 | 8 | 0.875747298629968 |
| chr8-100574460-100574961 | 6 | 0.963944503691193 |
| chr7-51073662-51074163 | 2 | 0.902185871260695 |
| chr15-1221917-1222418 | 1 | 0.838321047806993 |
| chr3-70180292-70180793 | 8 | 0.984207095190466 |
| chr5-155602750-155603251 | 6 | 0.94881286167257 |
| chr2-218149848-218150349 | 6 | 0.987997793581122 |
| chr14-49785328-49785829 | 3 | 0.949897090113253 |
| chr10-8317172-8317673 | 8 | 0.894177700560265 |
| chr5-29319084-29319585 | 5 | 0.988038790271037 |
| chr7-54176058-54176559 | 3 | 0.998004914840928 |
| chr16-66272692-66273193 | 8 | 0.995106675910537 |
| chr19-36705190-36705691 | 5 | 0.930200075635223 |
| chr2-25693940-25694441 | 8 | 0.918658306905501 |
| chr10-99040400-99040901 | 5 | 0.920855347254841 |
| chr8-117648352-117648853 | 6 | 0.987597881173947 |
| chr1-137162545-137163046 | 3 | 0.989096112472958 |
| chr3-128194018-128194519 | 5 | 0.944070606771317 |
| chr7-26232480-26232981 | 7 | 0.856968914416357 |
| chr1-149285865-149286366 | 6 | 0.99247715868385 |
| chr3-163468140-163468641 | 3 | 0.871999077190877 |
| chr1-144823711-144824212 | 8 | 0.979129626666377 |
| chr10-1026489-1026990 | 8 | 0.975319319685051 |
| chr9-62167035-62167536 | 8 | 0.927308346592729 |

| | | |
|---|---|---|
| chr1-168342992-168343493 | 8 | 0.825400572826531 |
| chr10-103874578-103875079 | 8 | 0.923555766413994 |
| chr7-69270521-69271022 | 8 | 0.995184176137661 |
| chr15-38377272-38377773 | 1 | 0.928802800759943 |
| chr1-259796455-259796956 | 8 | 0.89479990309194 |
| chr8-111496445-111496946 | 3 | 0.995935535450041 |
| chr5-71519231-71519732 | 4 | 0.969324336125022 |
| chr4-105302595-105303096 | 4 | 0.903087724174875 |
| chr15-99231763-99232264 | 4 | 0.892435152631703 |
| chr6-50286574-50287075 | 6 | 0.98350659776052 |
| chr1-39816167-39816668 | 6 | 0.886510210268062 |
| chr3-20344461-20344962 | 3 | 0.995975368491199 |
| chr9-6887739-6888240 | 2 | 0.910719861666149 |
| chr7-29573428-29573929 | 8 | 0.932641744024601 |
| chr1-219097952-219098453 | 2 | 0.966654296218827 |
| chr4-82132116-82132617 | 8 | 0.967951995658161 |
| chr8-47739996-47740497 | 5 | 0.987812098431893 |
| chr17-86013410-86013911 | 8 | 0.991100712496073 |
| chr10-42268751-42269252 | 3 | 0.982001712915839 |
| chr18-25966762-25967263 | 8 | 0.974292407342685 |
| chr1-229126098-229126599 | 8 | 0.905657661205597 |
| chr2-204699637-204700138 | 5 | 0.984344138401646 |
| chr4-88360627-88361128 | 3 | 0.994628003496879 |
| chr1-259148453-259148954 | 8 | 0.993493681128311 |
| chr2-190691399-190691900 | 8 | 0.978595240055365 |
| chr5-19498121-19498622 | 6 | 0.989070397512939 |
| chr14-18620388-18620889 | 4 | 0.972452095340463 |
| chr5-78312822-78313323 | 3 | 0.99593553545041 |
| chr6-118030806-118031307 | 4 | 0.82470540889327 |
| chr20-19344880-19345381 | 8 | 0.965351229213173 |
| chrX-133057532-133058033 | 3 | 0.99593553545041 |
| chr19-48632601-48633102 | 4 | 0.825708369640626 |
| chrX-10702290-10702791 | 4 | 0.937211922674044 |
| chr5-82599024-82599525 | 3 | 0.99593553545041 |
| chr7-46003015-46003516 | 8 | 0.9864944431108 |
| chr7-89343493-89343994 | 8 | 0.958005452479226 |
| chr15-47014409-47014910 | 6 | 0.989083892043489 |
| chr18-55935291-55935792 | 3 | 0.977305211111825 |
| chr3-120659689-120660190 | 3 | 0.977849800973773 |
| chr19-46721339-46721840 | 6 | 0.935983696376853 |
| chr16-7316303-7316804 | 8 | 0.947824846183532 |
| chr13-87399856-87400357 | 3 | 0.993705733262763 |
| chr1-137939952-137940453 | 4 | 0.926745616116053 |

| | | |
|---|---|---|
| chr2-50301669-50302170 | 6 | 0.853055782931624 |
| chr19-42351107-42351608 | 7 | 0.870631965476964 |
| chr16-57329484-57329985 | 6 | 0.996016255547304 |
| chr4-108803968-108804469 | 3 | 0.932676869862292 |
| chr6-123323862-123324363 | 1 | 0.86079768597454 |
| chr1-245486293-245486794 | 3 | 0.966934796454594 |
| chr8-61938479-61938980 | 1 | 0.819663997855808 |
| chr5-99787713-99788214 | 6 | 0.992566101472949 |
| chr1-10041890-10042391 | 3 | 0.870955692108651 |
| chr3-120069823-120070324 | 6 | 0.991206925481043 |
| chr10-48721722-48722223 | 8 | 0.814967810257943 |
| chr14-69196535-69197036 | 8 | 0.964942232090365 |
| chr18-13327968-13328469 | 6 | 0.981711114071626 |
| chr4-89766714-89767215 | 8 | 0.987026112250246 |
| chr1-222250001-222250502 | 8 | 0.995153004418284 |
| chr13-79294319-79294820 | 6 | 0.992722904314563 |
| chr1-158761885-158762386 | 6 | 0.940883658954042 |
| chr16-39784024-39784525 | 3 | 0.973887975470196 |
| chr7-101010280-101010781 | 6 | 0.992673163291303 |
| chr1-233608421-233608922 | 6 | 0.992212586164696 |
| chr1-222965673-222966174 | 4 | 0.828955508439191 |
| chr9-53703014-53703515 | 3 | 0.996667916347605 |
| chr6-7125879-7126380 | 8 | 0.993396263088517 |
| chr4-113738846-113739347 | 5 | 0.991279367457463 |
| chr13-65612660-65613161 | 5 | 0.95682728756915 |
| chr1-216220632-216221133 | 3 | 0.976508107440066 |
| chr6-121200303-121200804 | 3 | 0.996709201433337 |
| chr2-116354947-116355448 | 3 | 0.908259458691794 |
| chr1-185147241-185147742 | 2 | 0.956198123157754 |
| chr4-154932177-154932678 | 1 | 0.916295517706249 |
| chr8-64995207-64995708 | 4 | 0.898630213242696 |
| chr2-78886479-78886980 | 6 | 0.885873477808501 |
| chr20-1947777-1948278 | 6 | 0.965881516772226 |
| chr8-82839497-82839998 | 3 | 0.996542439612702 |
| chr4-116147603-116148104 | 8 | 0.83236376251562 |
| chr7-99495405-99495906 | 3 | 0.99593553545041 |
| chr18-2928398-2928899 | 2 | 0.924400880424717 |
| chr3-143730938-143731439 | 8 | 0.86043597944876 |
| chr6-8636622-8637123 | 8 | 0.94333269222264 |
| chr13-21044455-21044956 | 5 | 0.9841168719532 |
| chr5-109203204-109203705 | 3 | 0.992520402377033 |
| chr10-77122258-77122759 | 3 | 0.950409025092972 |
| chr16-56506605-56507106 | 8 | 0.948289950262993 |

| | | |
|---|---|---|
| chr5-129380519-129381020 | 7 | 0.83724163131233 |
| chr1-126943894-126944395 | 3 | 0.996566353929505 |
| chr6-90752411-90752912 | 7 | 0.965875060420093 |
| chr8-92188898-92189399 | 3 | 0.957349355728466 |
| chr10-17108175-17108676 | 4 | 0.945423359129123 |
| chr14-59903290-59903791 | 5 | 0.97942055441416 |
| chr14-19404148-19404649 | 2 | 0.830224642213134 |
| chr14-35904150-35904651 | 6 | 0.967665439463165 |
| chr17-16570213-16570714 | 5 | 0.983180737290894 |
| chr10-48865246-48865747 | 8 | 0.955125566245621 |
| chr5-87832958-87833459 | 3 | 0.996610248228935 |
| chr9-69032587-69033088 | 6 | 0.988971295156621 |
| chr14-79922737-79923238 | 8 | 0.947279136894053 |
| chr4-106862539-106863040 | 8 | 0.991977841818188 |
| chr11-9113793-9114294 | 3 | 0.99593553545041 |
| chr9-37849818-37850319 | 3 | 0.99593553545041 |
| chr11-61990164-61990665 | 2 | 0.960840186647126 |
| chr4-8793765-8794266 | 6 | 0.990733359279283 |
| chr6-72610005-72610506 | 8 | 0.940476837786647 |
| chr8-22422332-22422833 | 8 | 0.994559742841121 |
| chr6-20575598-20576099 | 3 | 0.884574138025342 |
| chr6-114639250-114639751 | 8 | 0.930479226725421 |
| chr2-167917337-167917838 | 3 | 0.99593553545041 |
| chr1-159499140-159499641 | 3 | 0.99593553545041 |
| chr7-99035877-99036378 | 8 | 0.908720286108824 |
| chr13-33221636-33222137 | 3 | 0.939778937209957 |
| chr17-64259301-64259802 | 5 | 0.846982199268887 |
| chr1-223375680-223376181 | 8 | 0.995232271402472 |
| chr6-31276054-31276555 | 3 | 0.90045045802862 |
| chr14-41390339-41390840 | 8 | 0.897626553303447 |
| chr8-97717211-97717712 | 8 | 0.985978901028341 |
| chr2-81282782-81283283 | 6 | 0.847085674351862 |
| chr10-27146065-27146566 | 3 | 0.873822025173216 |
| chr3-64220895-64221396 | 8 | 0.992782779155339 |
| chr20-10860585-10861086 | 8 | 0.825929512698147 |
| chr7-60085059-60085560 | 8 | 0.995246341630326 |
| chr2-186372354-186372855 | 4 | 0.968639547990024 |
| chr4-123570017-123570518 | 8 | 0.913757884835234 |
| chrX-106440390-106440891 | 7 | 0.856876646812543 |
| chr18-37070612-37071113 | 1 | 0.878309033250622 |
| chr9-1909223-1909724 | 4 | 0.901143590013405 |
| chr9-57503199-57503700 | 8 | 0.950091635081441 |
| chr11-70187935-70188436 | 2 | 0.957478892400803 |

| | | |
|---|---|---|
| chr5-33353070-33353571 | 3 | 0.904643920619509 |
| chr15-8098518-8099019 | 3 | 0.974924743729247 |
| chr4-104619531-104620032 | 5 | 0.986076432418345 |
| chr2-111270896-111271397 | 8 | 0.851207750195739 |
| chr8-57159967-57160468 | 8 | 0.950254798473787 |
| chrX-76663747-76664248 | 5 | 0.848391396838662 |
| chr3-33173746-33174247 | 8 | 0.886197045429999 |
| chr1-218483026-218483527 | 6 | 0.952206046312039 |
| chr5-91232426-91232927 | 8 | 0.981936529013587 |
| chr10-10631402-10631903 | 4 | 0.895947073347236 |
| chr7-60331435-60331936 | 3 | 0.99593553545041 |
| chr16-69141927-69142428 | 8 | 0.955763203185551 |
| chr6-97921621-97922122 | 8 | 0.880318321283323 |
| chr9-79910011-79910512 | 8 | 0.842663006300393 |
| chr1-207058468-207058969 | 1 | 0.932607209854103 |
| chr7-34781868-34782369 | 5 | 0.990485615476185 |
| chr7-59197496-59197997 | 3 | 0.972415765017991 |
| chr15-56305096-56305597 | 3 | 0.989671264800668 |
| chr8-90005374-90005875 | 8 | 0.993493681128311 |
| chr7-126517190-126517691 | 8 | 0.948288973256676 |
| chr13-22722531-22723032 | 6 | 0.863305102522622 |
| chr8-63343765-63344266 | 2 | 0.923647452700881 |
| chr14-77323650-77324151 | 4 | 0.917970604195101 |
| chr11-13333029-13333530 | 8 | 0.957977582822757 |
| chr2-123349095-123349596 | 4 | 0.934043321262994 |
| chr8-122115812-122116313 | 8 | 0.989100704754532 |
| chr3-54416031-54416532 | 4 | 0.880468545800135 |
| chr8-36824043-36824544 | 2 | 0.905710661028793 |
| chr16-31284517-31285018 | 3 | 0.981610537649359 |
| chr9-107316081-107316582 | 1 | 0.824683138874898 |
| chr2-140458641-140459142 | 8 | 0.964642185261253 |
| chr6-17450729-17451230 | 6 | 0.991206925481043 |
| chr15-18334014-18334515 | 1 | 0.94076479181528 |
| chr16-46078483-46078984 | 2 | 0.835264239008658 |
| chr14-80267524-80268025 | 2 | 0.91970016926947 |
| chr5-113131720-113132221 | 3 | 0.997163995135549 |
| chr1-121610168-121610669 | 8 | 0.897898162384234 |
| chrX-107198394-107198895 | 8 | 0.995226541933262 |
| chr20-31874431-31874932 | 8 | 0.95830416410406 |
| chr10-49657087-49657588 | 2 | 0.902093007890272 |
| chr5-55299817-55300318 | 8 | 0.993493681128311 |
| chr6-138318263-138318764 | 6 | 0.936874040593042 |
| chr2-139988825-139989326 | 5 | 0.866511946112116 |

| | | |
|---|---|---|
| chr10-11795613-11796114 | 8 | 0.994364960509474 |
| chr13-33614620-33615121 | 3 | 0.957809541562387 |
| chr7-118478555-118479056 | 3 | 0.995044545148374 |
| chr8-121069874-121070375 | 6 | 0.97323029211718 |
| chr1-252451000-252451501 | 3 | 0.959469806996889 |
| chr2-183934121-183934622 | 8 | 0.886431037642107 |
| chr7-27599920-27600421 | 3 | 0.98476646230379 |
| chr14-9385265-9385766 | 8 | 0.992055450525955 |
| chr5-74527151-74527652 | 7 | 0.86300471489206 |
| chr4-136211356-136211857 | 5 | 0.93782311504702 |
| chr16-26889954-26890455 | 3 | 0.960205073550857 |
| chr15-62388887-62389388 | 3 | 0.995687195676245 |
| chr7-66102744-66103245 | 8 | 0.90073481339022 |
| chr5-105988563-105989064 | 8 | 0.974406770545099 |
| chr18-55350299-55350800 | 8 | 0.851629996325454 |
| chr19-18827194-18827695 | 1 | 0.958502356876963 |
| chr15-13183689-13184190 | 8 | 0.883940655555277 |
| chr2-90884508-90885009 | 6 | 0.993154086598148 |
| chr11-49771292-49771793 | 3 | 0.996442388254158 |
| chr11-54454961-54455462 | 4 | 0.934395745708266 |
| chr11-72214729-72215230 | 6 | 0.993191120017266 |
| chr4-178280816-178281317 | 3 | 0.99593553545041 |
| chr2-238332030-238332531 | 6 | 0.944622967851572 |
| chr8-105933968-105934469 | 3 | 0.996110138046083 |
| chr18-73783853-73784354 | 3 | 0.972313695664913 |
| chr12-10759743-10760244 | 8 | 0.93427467871112 |
| chr4-15514946-15515447 | 6 | 0.87766519770172 |
| chr4-57605245-57605746 | 8 | 0.924428916975826 |
| chr2-33648092-33648593 | 8 | 0.943302319212795 |
| chr1-34190326-34190827 | 3 | 0.994964290531962 |
| chr7-122165693-122166194 | 3 | 0.996993592986769 |
| chr10-20501171-20501672 | 8 | 0.940693396719935 |
| chr17-25645938-25646439 | 8 | 0.981824900745555 |
| chr9-16283393-16283894 | 8 | 0.975673152929464 |
| chr9-37512435-37512936 | 3 | 0.995815571152109 |
| chr14-26986469-26986970 | 3 | 0.967660456245225 |
| chr10-106619776-106620277 | 5 | 0.890366256461618 |
| chr3-116454713-116455214 | 4 | 0.921078287502345 |
| chr7-24954099-24954600 | 2 | 0.831846919822867 |
| chr5-4563188-4563689 | 6 | 0.991206925481043 |
| chr1-54094116-54094617 | 3 | 0.995202787033579 |
| chr2-63473077-63473578 | 6 | 0.991926435001023 |
| chr16-52284389-52284890 | 3 | 0.992836852448801 |

| | | |
|---|---|---|
| chr17-56539615-56540116 | 6 | 0.98839974893394 |
| chr18-44542313-44542814 | 3 | 0.969680344631407 |
| chr15-28926431-28926932 | 8 | 0.894464993599505 |
| chr1-239919029-239919530 | 2 | 0.92952845213992 |
| chr17-17959306-17959807 | 4 | 0.970493683356963 |
| chr5-150739410-150739911 | 8 | 0.979074863719178 |
| chr17-62274903-62275404 | 8 | 0.902631669187745 |
| chr3-12391217-12391718 | 8 | 0.860638563323535 |
| chr15-35070877-35071378 | 6 | 0.987271641740978 |
| chr7-86633730-86634231 | 3 | 0.818388674312947 |
| chr4-66367143-66367644 | 2 | 0.905964286398482 |
| chr12-44075495-44075996 | 3 | 0.995826275508609 |
| chr1-215199943-215200444 | 2 | 0.947792482952639 |
| chr2-77628022-77628523 | 5 | 0.916494131253148 |
| chr6-126319397-126319898 | 3 | 0.998450418807014 |
| chr7-50359234-50359735 | 6 | 0.968710220374559 |
| chr9-86268780-86269281 | 1 | 0.866603981014963 |
| chr5-71163192-71163693 | 6 | 0.990494757659631 |
| chr12-17211583-17212084 | 3 | 0.8262331909089 |
| chr3-134369961-134370462 | 3 | 0.995311200741689 |
| chr14-76361388-76361889 | 8 | 0.889522477295899 |
| chr8-41825465-41825966 | 8 | 0.996848898185876 |
| chr2-147021233-147021734 | 3 | 0.99205037170602 |
| chr1-13206971-13207472 | 8 | 0.977863170475489 |
| chr4-105874615-105875116 | 3 | 0.993775710161671 |
| chr9-36992221-36992722 | 8 | 0.945726794987927 |
| chr10-63636273-63636774 | 6 | 0.98749978379207 |
| chr3-36981150-36981651 | 3 | 0.849206976869078 |
| chr10-21943836-21944337 | 8 | 0.991053483875848 |
| chr7-69872721-69873222 | 8 | 0.818393472795624 |
| chr7-17494620-17495121 | 8 | 0.916934112693595 |
| chr8-32967013-32967514 | 6 | 0.991206925481043 |
| chr8-61154512-61155013 | 5 | 0.966299888848664 |
| chr1-152690875-152691376 | 1 | 0.903147678082478 |
| chr4-43313527-43314028 | 5 | 0.994292171447612 |
| chr15-36334429-36334930 | 5 | 0.987952190265581 |
| chr14-87053830-87054331 | 2 | 0.891967911153604 |
| chr5-80055599-80056100 | 8 | 0.994403199927586 |
| chr2-219336188-219336689 | 8 | 0.995223049512792 |
| chr6-53911684-53912185 | 1 | 0.819405441711798 |
| chr19-47101990-47102491 | 3 | 0.994046906856526 |
| chr13-50141116-50141617 | 6 | 0.975718458744978 |
| chr17-31962842-31963343 | 8 | 0.953825541853651 |

| | | |
|---|---|---|
| chr13-65029489-65029990 | 5 | 0.973783017564678 |
| chr7-39114504-39115005 | 8 | 0.828823669604691 |
| chr15-16735173-16735674 | 8 | 0.990112838093362 |
| chr17-48820786-48821287 | 3 | 0.94434416577743 |
| chr7-109253885-109254386 | 6 | 0.850550352855147 |
| chr1-187666730-187667231 | 8 | 0.964532646523908 |
| chr4-122487831-122488332 | 3 | 0.92077695241113 |
| chr2-22734305-22734806 | 6 | 0.832959873658466 |
| chr16-15087267-15087768 | 3 | 0.996897467934083 |
| chr4-125767188-125767689 | 2 | 0.95801873825263 |
| chr3-67608356-67608857 | 8 | 0.94736825512985 |
| chr3-113865673-113866174 | 6 | 0.867127356512239 |
| chr7-103311426-103311927 | 6 | 0.908858971257047 |
| chr18-24634679-24635180 | 6 | 0.859423304559701 |
| chr8-51400874-51401375 | 3 | 0.980914658808508 |
| chr3-79122164-79122665 | 1 | 0.936091636440261 |
| chr8-50839539-50840040 | 6 | 0.935110106499568 |
| chr4-84448417-84448918 | 7 | 0.867030297636068 |
| chr1-213339205-213339706 | 6 | 0.972839686294957 |
| chr17-21513713-21514214 | 8 | 0.992034754836858 |
| chr16-45782779-45783280 | 7 | 0.943161062961004 |
| chr19-47731310-47731811 | 5 | 0.984959163770208 |
| chr5-137586196-137586697 | 2 | 0.843373375429845 |
| chr4-51024781-51025282 | 8 | 0.839323893540635 |
| chr15-95793393-95793894 | 3 | 0.982221209428539 |
| chr10-86278832-86279333 | 5 | 0.988386899288467 |
| chr17-85092072-85092573 | 8 | 0.986458991656395 |
| chr4-50502589-50503090 | 6 | 0.993017026766009 |
| chr9-12453378-12453879 | 4 | 0.826914268476105 |
| chr7-36310810-36311311 | 3 | 0.99593553545041 |
| chr9-91011762-91012263 | 3 | 0.908480457909944 |
| chr6-51167428-51167929 | 8 | 0.930796630306894 |
| chr19-37523792-37524293 | 8 | 0.909634181643856 |
| chr4-104223695-104224196 | 5 | 0.987730077253566 |
| chr5-158855934-158856435 | 1 | 0.93677197663618 |
| chr6-40205777-40206278 | 3 | 0.996045245542361 |
| chr20-30770179-30770680 | 2 | 0.883574790208368 |
| chr6-11975625-11976126 | 8 | 0.887331094196907 |
| chr3-82651562-82652063 | 3 | 0.99593553545041 |
| chr19-2194022-2194523 | 8 | 0.994153680037558 |
| chr14-102032455-102032956 | 8 | 0.994980944210236 |
| chr17-63468113-63468614 | 8 | 0.927958348594177 |
| chr9-42688509-42689010 | 8 | 0.916068063060558 |

| | | |
|---|---|---|
| chr10-56067552-56068053 | 8 | 0.968145184709039 |
| chr8-49357833-49358334 | 3 | 0.99478338826787 |
| chr9-94292097-94292598 | 3 | 0.996692671047639 |
| chr4-60725911-60726412 | 3 | 0.99593553545041 |
| chr8-96402418-96402919 | 5 | 0.958463443122067 |
| chrX-17244589-17245090 | 3 | 0.917728057749773 |
| chr11-36930402-36930903 | 3 | 0.995016287516848 |
| chr1-207115966-207116467 | 6 | 0.980337989280986 |
| chr1-12779330-12779831 | 5 | 0.985529474978867 |
| chr3-143633374-143633875 | 8 | 0.866380527676822 |
| chr16-3061199-3061700 | 8 | 0.892738139352657 |
| chr13-29705768-29706269 | 8 | 0.816406335603163 |
| chr9-26994791-26995292 | 3 | 0.994640527301336 |
| chr6-35395196-35395697 | 2 | 0.861468593197332 |
| chrX-37195791-37196292 | 8 | 0.829761387016635 |
| chr1-3197833-3198334 | 2 | 0.967088199225675 |
| chr2-38477009-38477510 | 5 | 0.970238514148418 |
| chr11-43772722-43773223 | 6 | 0.991206925481043 |
| chr10-41512080-41512581 | 3 | 0.902821718361398 |
| chr2-240405145-240405646 | 1 | 0.870927231007338 |
| chr9-17632217-17632718 | 5 | 0.990746976417433 |
| chr16-76880466-76880967 | 8 | 0.921562069959513 |
| chr2-162115004-162115505 | 8 | 0.995248859488942 |
| chr1-76271132-76271633 | 3 | 0.99593553545041 |
| chr2-22349492-22349993 | 4 | 0.86255058026029 |
| chr6-123944238-123944739 | 8 | 0.983146847254847 |
| chr3-17910123-17910624 | 4 | 0.883300844333624 |
| chr14-9081588-9082089 | 6 | 0.847377123903534 |
| chr2-194465538-194466039 | 3 | 0.832620251805178 |
| chr2-63114321-63114822 | 8 | 0.849950687998709 |
| chr1-9336260-9336761 | 1 | 0.911757189326417 |
| chr4-12835651-12836152 | 3 | 0.960492142564211 |
| chr5-149163771-149164272 | 5 | 0.986817809822641 |
| chr1-42032826-42033327 | 6 | 0.884158266644416 |
| chr11-62878863-62879364 | 7 | 0.885987486916031 |
| chr20-30036677-30037178 | 3 | 0.832087657302451 |
| chr18-16900620-16901121 | 4 | 0.879575353819111 |
| chr17-54355790-54356291 | 5 | 0.869367175287437 |
| chr6-99295133-99295634 | 8 | 0.965193296397442 |
| chr6-94782512-94783013 | 5 | 0.960733532274367 |
| chr11-36385471-36385972 | 3 | 0.996103773592435 |
| chr3-2294043-2294544 | 7 | 0.929609300713583 |
| chr16-51243073-51243574 | 3 | 0.998856709670896 |

| | | |
|---|---|---|
| chr20-18780614-18781115 | 8 | 0.902140536299978 |
| chr5-113951668-113952169 | 8 | 0.992295009526017 |
| chr2-175503948-175504449 | 8 | 0.850247776331661 |
| chr19-55668205-55668706 | 7 | 0.845205763682802 |
| chr5-20124148-20124649 | 1 | 0.968566836537958 |
| chr10-5431103-5431604 | 5 | 0.976888447795076 |
| chr6-43674395-43674896 | 5 | 0.972467590809782 |
| chr2-115191909-115192410 | 2 | 0.819285241917083 |
| chr15-63321282-63321783 | 5 | 0.893712590925126 |
| chr4-53944382-53944883 | 8 | 0.98725357158232 |
| chr5-105502509-105503010 | 3 | 0.996785280780476 |
| chr14-97638378-97638879 | 5 | 0.984311871286195 |
| chr11-49423843-49424344 | 4 | 0.883519090874843 |
| chr10-97420231-97420732 | 3 | 0.996064518998858 |
| chr16-60331715-60332216 | 6 | 0.992378717618084 |
| chr6-36980149-36980650 | 8 | 0.954979438388199 |
| chr17-29656686-29657187 | 1 | 0.846283279992168 |
| chr5-90830162-90830663 | 8 | 0.986735362741399 |
| chr20-39623763-39624264 | 3 | 0.99593553545041 |
| chr5-72791919-72792420 | 8 | 0.982353099436195 |
| chr19-38561591-38562092 | 6 | 0.994831902039436 |
| chr17-71607739-71608240 | 8 | 0.954714861000842 |
| chr1-12043784-12044285 | 6 | 0.988916148774189 |
| chr3-163001970-163002471 | 7 | 0.879579433364598 |
| chr2-75718171-75718672 | 3 | 0.977520544792785 |
| chr11-22452552-22453053 | 6 | 0.991206925481043 |
| chr20-29016138-29016639 | 5 | 0.960006954544167 |
| chr4-61407693-61408194 | 6 | 0.993063041173872 |
| chr6-37573784-37574285 | 3 | 0.996064713943643 |
| chr2-176136599-176137100 | 5 | 0.98757837948099 |
| chr7-28568291-28568792 | 3 | 0.882743453814069 |
| chr5-71802613-71803114 | 4 | 0.944051450463312 |
| chr2-144825331-144825832 | 5 | 0.81845500318154 |
| chr2-152597047-152597548 | 3 | 0.99593553545041 |
| chr5-121510752-121511253 | 6 | 0.9932378925482 |
| chr15-38367323-38367824 | 1 | 0.929507113859707 |
| chr13-99258543-99259044 | 3 | 0.994173093617091 |
| chr18-69319742-69320243 | 5 | 0.823196699351966 |
| chr13-29597979-29598480 | 8 | 0.933632054886891 |
| chr13-99574663-99575164 | 3 | 0.98385162741418 |
| chr18-11966758-11967259 | 8 | 0.969761479770924 |
| chr13-70684495-70684996 | 5 | 0.947457316128206 |
| chr1-245992948-245993449 | 3 | 0.904362335196027 |

| | | |
|---|---|---|
| chr9-70683378-70683879 | 5 | 0.975999200880075 |
| chr2-204186648-204187149 | 8 | 0.967456626914029 |
| chr10-11625317-11625818 | 8 | 0.865330355440124 |
| chr17-26423869-26424370 | 1 | 0.885101761268521 |
| chr2-211696974-211697475 | 8 | 0.96257482370271 |
| chr2-40317156-40317657 | 3 | 0.9870417955052 |
| chr13-50770231-50770732 | 6 | 0.989944450480595 |
| chr3-68678812-68679313 | 5 | 0.958821882248705 |
| chr8-71712152-71712653 | 6 | 0.99359618223639 |
| chr3-60726024-60726525 | 8 | 0.973772238719434 |
| chr3-133055372-133055873 | 8 | 0.984467592672194 |
| chr19-34988912-34989413 | 8 | 0.818440286776921 |
| chr1-196693270-196693771 | 6 | 0.899368639811308 |
| chr4-25378231-25378732 | 7 | 0.95539616656959 |
| chr6-35890002-35890503 | 1 | 0.860928093147633 |
| chr12-39295609-39296110 | 4 | 0.860033871741859 |
| chr13-63944497-63944998 | 6 | 0.950980188451224 |
| chr17-22665717-22666218 | 1 | 0.875492626769663 |
| chr13-105171958-105172459 | 3 | 0.9959881822212 |
| chr5-5759338-5759839 | 8 | 0.84241041196485 |
| chr14-67184393-67184894 | 4 | 0.935117975921206 |
| chr19-9166374-9166875 | 4 | 0.947113638610596 |
| chr7-49525416-49525917 | 3 | 0.99593553545041 |
| chr3-7257749-7258250 | 6 | 0.91935878431396 |
| chr3-28017969-28018470 | 8 | 0.834205097672234 |
| chr12-38457296-38457797 | 4 | 0.956340497027119 |
| chr2-23436817-23437318 | 1 | 0.908870650756474 |
| chr18-73838635-73839136 | 8 | 0.959769677766544 |
| chr5-166288214-166288715 | 8 | 0.880158669674408 |
| chr17-32037957-32038458 | 7 | 0.815500898695025 |
| chr6-132688387-132688888 | 3 | 0.946400783703193 |
| chr6-19946281-19946782 | 1 | 0.884734400872907 |
| chr1-121720861-121721362 | 4 | 0.822237454866721 |
| chr18-18263700-18264201 | 3 | 0.827595114055821 |
| chr1-12182755-12183256 | 5 | 0.988956487905615 |
| chr18-13994722-13995223 | 6 | 0.991206925481043 |
| chr10-73646102-73646603 | 2 | 0.901153542355495 |
| chr16-41423571-41424072 | 5 | 0.976528702829629 |
| chr3-166680523-166681024 | 2 | 0.86228085405078 |
| chr7-30777836-30778337 | 6 | 0.902293452157212 |
| chr10-100004319-100004820 | 3 | 0.997213688367294 |
| chr4-120008450-120008951 | 8 | 0.96942408887644 |
| chr1-169732765-169733266 | 8 | 0.919331370489976 |

| | | |
|---|---|---|
| chr6-105436836-105437337 | 8 | 0.898257647686736 |
| chr18-8050311-8050812 | 3 | 0.996030335890507 |
| chr4-21377256-21377757 | 6 | 0.987259512326222 |
| chr7-35044432-35044933 | 8 | 0.993493681128311 |
| chr7-56708529-56709030 | 8 | 0.961325819338905 |
| chr13-87152163-87152664 | 6 | 0.976704112056354 |
| chr14-66970218-66970719 | 8 | 0.956595280699038 |
| chr19-52544745-52545246 | 6 | 0.992885972978943 |
| chr7-123917321-123917822 | 5 | 0.856435203736555 |
| chr2-242870006-242870507 | 3 | 0.844293709173465 |
| chr1-256796904-256797405 | 3 | 0.989315847356981 |
| chr7-113338036-113338537 | 2 | 0.931869613947622 |
| chr17-64812818-64813319 | 6 | 0.967548037748511 |
| chr2-4218111-4218612 | 2 | 0.910612953699041 |
| chr15-10102540-10103041 | 5 | 0.880094847916822 |
| chr1-118473432-118473933 | 7 | 0.886275231801594 |
| chr3-88911341-88911842 | 3 | 0.997236681685013 |
| chr15-68562842-68563343 | 6 | 0.991206925481043 |
| chr17-78556831-78557332 | 3 | 0.848661632926371 |
| chr1-96764792-96765293 | 5 | 0.898428310899397 |
| chr2-216316828-216317329 | 6 | 0.971309372983181 |
| chr19-27908266-27908767 | 8 | 0.97455201620785 |
| chr5-21998154-21998655 | 5 | 0.918713447213192 |
| chr11-49809443-49809944 | 8 | 0.893214140497344 |
| chr3-97513835-97514336 | 3 | 0.98609398230243 |
| chr15-27010672-27011173 | 3 | 0.99674546723326 |
| chr2-121543040-121543541 | 8 | 0.84254616130651 |
| chr9-11320980-11321481 | 4 | 0.970776747473564 |
| chr15-35783647-35784148 | 8 | 0.85225101778588 |
| chr7-109287557-109288058 | 3 | 0.991675573275343 |
| chr4-137662700-137663201 | 3 | 0.99593553545041 |
| chr4-165572934-165573435 | 3 | 0.991591675341721 |
| chr7-121118022-121118523 | 3 | 0.996890803379835 |
| chr14-76389151-76389652 | 5 | 0.975197849278515 |
| chr2-53326085-53326586 | 8 | 0.9816106684395 |
| chr5-115307489-115307990 | 8 | 0.873216540328511 |
| chr8-32805977-32806478 | 8 | 0.964489900750402 |
| chr15-92570046-92570547 | 2 | 0.85397287581985 |
| chr3-10590874-10591375 | 1 | 0.950458772699935 |
| chr12-15796856-15797357 | 8 | 0.939889649746431 |
| chr15-42669703-42670204 | 3 | 0.995704234778989 |
| chr20-15558955-15559456 | 6 | 0.991206925481043 |
| chrX-26207812-26208313 | 6 | 0.991206925481043 |

| | | |
|---|---|---|
| chr16-32307041-32307542 | 3 | 0.95589179315599 |
| chrX-97739194-97739695 | 5 | 0.903242485694359 |
| chr2-170548166-170548667 | 3 | 0.883891710629238 |
| chr3-55435188-55435689 | 2 | 0.956532043768478 |
| chr19-26834927-26835428 | 4 | 0.85216945792898 |
| chr20-10020218-10020719 | 3 | 0.858522801522824 |
| chr16-69696382-69696883 | 8 | 0.881333046755865 |
| chr6-105552169-105552670 | 2 | 0.946607451665543 |
| chr1-21864431-21864932 | 3 | 0.99593553545041 |
| chr9-3205866-3206367 | 2 | 0.952095665361803 |
| chr17-66574405-66574906 | 3 | 0.997161870124846 |
| chr4-137324468-137324969 | 5 | 0.924748687054253 |
| chr6-91920588-91921089 | 5 | 0.903278926793844 |
| chr7-124899536-124900037 | 5 | 0.982704381163367 |
| chr1-163493882-163494383 | 8 | 0.859593723770876 |
| chr2-104566574-104567075 | 3 | 0.993447107478162 |
| chr3-54969205-54969706 | 1 | 0.83111943690199 |
| chr18-16293826-16294327 | 7 | 0.846729783050205 |
| chr20-8496441-8496942 | 4 | 0.860105379213457 |
| chr6-84359835-84360336 | 3 | 0.99593553545041 |
| chr10-52711505-52712006 | 6 | 0.970177809113545 |
| chr19-51835943-51836444 | 5 | 0.88945459731196 |
| chr8-20010916-20011417 | 4 | 0.86911011034188 |
| chr5-141996007-141996508 | 8 | 0.99076203678828 |
| chr19-8662810-8663311 | 5 | 0.990233838321712 |
| chr3-53498257-53498758 | 2 | 0.888807591376011 |
| chr19-28902104-28902605 | 6 | 0.995508266902458 |
| chr13-85251571-85252072 | 5 | 0.983919978840557 |
| chr8-96961378-96961879 | 3 | 0.918989158432561 |
| chr14-38716995-38717496 | 3 | 0.996352522315109 |
| chr11-51786002-51786503 | 4 | 0.926309546221374 |
| chr6-23221885-23222386 | 8 | 0.850593830638575 |
| chr2-141755564-141756065 | 1 | 0.876896357375539 |
| chr8-63651460-63651961 | 5 | 0.981622286711607 |
| chr2-107559675-107560176 | 3 | 0.99560933482721 |
| chr8-37372733-37373234 | 5 | 0.947427380780213 |
| chrX-73424355-73424856 | 3 | 0.99593553545041 |
| chr2-78217248-78217749 | 8 | 0.961194856745093 |
| chr17-16794883-16795384 | 8 | 0.990498680960834 |
| chr15-26549498-26549999 | 2 | 0.953932865645374 |
| chr11-52244272-52244773 | 5 | 0.980509218838984 |
| chr9-36710360-36710861 | 8 | 0.896948935431929 |
| chrX-126415059-126415560 | 3 | 0.99447714795145 |

| | | |
|---|---|---|
| chr19-19326583-19327084 | 5 | 0.907613590599914 |
| chr4-26288799-26289300 | 5 | 0.916088109351709 |
| chr18-24423440-24423941 | 1 | 0.830552796816118 |
| chr14-11467282-11467783 | 5 | 0.978199722175455 |
| chr12-31988099-31988600 | 1 | 0.894130739020971 |
| chr2-32481168-32481669 | 4 | 0.834622100585666 |
| chr14-34662360-34662861 | 3 | 0.99593553545041 |
| chr8-101207241-101207742 | 3 | 0.883700526814161 |
| chr19-16876382-16876883 | 5 | 0.896622820957016 |
| chr20-36051108-36051609 | 7 | 0.824573875490767 |
| chr17-30291561-30292062 | 5 | 0.962302760778693 |
| chr7-115877656-115878157 | 8 | 0.959699937861911 |
| chr3-57283166-57283667 | 2 | 0.968215976960357 |
| chr11-44597952-44598453 | 8 | 0.919612214269662 |
| chr18-56242182-56242683 | 3 | 0.878798144504984 |
| chr8-18900055-18900556 | 3 | 0.99559006396072 |
| chr18-52598355-52598856 | 3 | 0.920461934962208 |
| chr11-40717867-40718368 | 1 | 0.960399003292631 |
| chr13-87947894-87948395 | 8 | 0.97314706824671 |
| chr7-27644260-27644761 | 6 | 0.909836207411112 |
| chr9-14391804-14392305 | 8 | 0.872517182864274 |
| chr19-3085074-3085575 | 6 | 0.994016248448758 |
| chr10-41182970-41183471 | 8 | 0.938206170023661 |
| chr14-50785566-50786067 | 3 | 0.958455505007571 |
| chr3-150423410-150423911 | 8 | 0.988793327588371 |
| chr3-158113203-158113704 | 5 | 0.944189303756894 |
| chr2-223328114-223328615 | 3 | 0.996956479647444 |
| chr1-240873435-240873936 | 4 | 0.958578909946226 |
| chr16-23034718-23035219 | 3 | 0.995671044906099 |
| chr5-110665835-110666336 | 5 | 0.896565946116589 |
| chr13-82422423-82422924 | 3 | 0.996352703471457 |
| chr11-40810523-40811024 | 5 | 0.843628132658024 |
| chr8-30410484-30410985 | 8 | 0.877553519152638 |
| chr19-52353265-52353766 | 8 | 0.993493681128311 |
| chr11-19387486-19387987 | 3 | 0.996382235156185 |
| chr6-112557830-112558331 | 6 | 0.975722365443783 |
| chr18-69843690-69844191 | 8 | 0.925931339169953 |
| chr19-14356220-14356721 | 5 | 0.831585986413919 |
| chr8-81211628-81212129 | 3 | 0.840017541231212 |
| chr14-44014197-44014698 | 5 | 0.957006558327174 |
| chr4-127980162-127980663 | 3 | 0.969976515067553 |
| chr10-41866560-41867061 | 5 | 0.851471204843869 |
| chr15-10240366-10240867 | 8 | 0.923962904262548 |

| | | |
|---|---|---|
| chr3-109703663-109704164 | 5 | 0.826462757111907 |
| chr7-43443262-43443763 | 2 | 0.898480377368694 |
| chr14-22485665-22486166 | 3 | 0.99532905608205 |
| chr4-75804776-75805277 | 5 | 0.980784419012058 |
| chr1-176060010-176060511 | 8 | 0.853947871996732 |
| chr9-56436924-56437425 | 8 | 0.990945706857936 |
| chr10-102758233-102758734 | 6 | 0.924839031451699 |
| chr14-81239878-81240379 | 3 | 0.988857506560034 |
| chr8-43980487-43980988 | 8 | 0.923539391351882 |
| chr18-44208881-44209382 | 3 | 0.996853107508974 |
| chr1-116959469-116959970 | 4 | 0.868231438331269 |
| chr1-118964211-118964712 | 5 | 0.964138534146951 |
| chr3-47452526-47453027 | 8 | 0.993493681128311 |
| chr5-97263940-97264441 | 8 | 0.884214572367266 |
| chr13-51117413-51117914 | 3 | 0.994155964605852 |
| chr20-49410241-49410742 | 8 | 0.908737324342026 |
| chr3-35414986-35415487 | 3 | 0.995475352675392 |
| chr13-7691100-7691601 | 3 | 0.903766152388814 |
| chr8-55636165-55636666 | 6 | 0.900642656835387 |
| chr19-1651130-1651631 | 6 | 0.990823966381369 |
| chr3-133862233-133862734 | 8 | 0.913277303190818 |
| chr10-67580612-67581113 | 8 | 0.911412730921648 |
| chr15-65520744-65521245 | 6 | 0.979237049622411 |
| chr2-60563912-60564413 | 8 | 0.856376326469436 |
| chr6-20221738-20222239 | 4 | 0.963902270504486 |
| chr7-34107312-34107813 | 5 | 0.877048178591569 |
| chr7-33780779-33781280 | 8 | 0.995248391549973 |
| chr17-180292-180793 | 8 | 0.993493681128311 |
| chr2-38426509-38427010 | 1 | 0.817399672278833 |
| chr2-47524599-47525100 | 5 | 0.981590196038017 |
| chr13-97369726-97370227 | 8 | 0.993493681128311 |
| chr2-190698581-190699082 | 8 | 0.941756574695585 |
| chr1-173324789-173325290 | 5 | 0.957150672525898 |
| chr2-234798250-234798751 | 1 | 0.883211365222228 |
| chr19-41443713-41444214 | 5 | 0.929535883749044 |
| chr13-96971426-96971927 | 8 | 0.927474091280753 |
| chr12-19791294-19791795 | 4 | 0.918503271529987 |
| chr7-82571931-82572432 | 8 | 0.980800322140071 |
| chr10-3782855-3783356 | 6 | 0.958400834098534 |
| chr13-100791322-100791823 | 8 | 0.995241984852863 |
| chr1-152463365-152463866 | 8 | 0.994095912069985 |
| chr9-94224723-94225224 | 8 | 0.860788143375171 |
| chr3-32881248-32881749 | 3 | 0.99593553545041 |

| | | |
|---|---|---|
| chr3-124926959-124927460 | 8 | 0.987388152797503 |
| chr4-52401163-52401664 | 6 | 0.991206925481043 |
| chr2-224655474-224655975 | 7 | 0.863653987140301 |
| chr6-104369650-104370151 | 5 | 0.978796756362467 |
| chr18-24286394-24286895 | 7 | 0.924035228869087 |
| chr2-183020412-183020913 | 8 | 0.908190941518698 |
| chr2-225916516-225917017 | 1 | 0.824989197381361 |
| chr7-51245570-51246071 | 3 | 0.947864194012094 |
| chr17-7017681-7018182 | 6 | 0.985762408875011 |
| chr2-40639854-40640355 | 3 | 0.949686656879072 |
| chr14-69743280-69743781 | 4 | 0.930054580338361 |
| chr2-140318849-140319350 | 3 | 0.982010325163162 |
| chr7-117538678-117539179 | 3 | 0.997101028203166 |
| chr5-96124718-96125219 | 4 | 0.920580068357258 |
| chr16-16484377-16484878 | 3 | 0.975944549711343 |
| chr12-35918689-35919190 | 3 | 0.851714142104762 |
| chr1-146880344-146880845 | 8 | 0.886227731310395 |
| chr8-98876190-98876691 | 3 | 0.988896112365191 |
| chr3-46146423-46146924 | 2 | 0.870267287483697 |
| chr9-18041290-18041791 | 3 | 0.970863796375216 |
| chr2-226945022-226945523 | 3 | 0.981060786782794 |
| chr10-62751353-62751854 | 1 | 0.855618305962094 |
| chr9-4792226-4792727 | 7 | 0.956554045734813 |
| chr2-82389270-82389771 | 5 | 0.988799968716129 |
| chr2-20682123-20682624 | 8 | 0.993493681128311 |
| chr14-20390305-20390806 | 5 | 0.991493727548763 |
| chr10-50891108-50891609 | 3 | 0.940699682701965 |
| chr15-63991108-63991609 | 3 | 0.996745637977307 |
| chr1-195912207-195912708 | 4 | 0.949997397414972 |
| chr8-21482997-21483498 | 8 | 0.989310960375905 |
| chr11-55171860-55172361 | 5 | 0.971888424445493 |
| chr3-102404961-102405462 | 8 | 0.993493681128311 |
| chr1-5106822-5107323 | 3 | 0.997061507001211 |
| chr8-5770171-5770672 | 3 | 0.948638211725407 |
| chr14-52883691-52884192 | 8 | 0.858168379842093 |
| chr8-73701611-73702112 | 8 | 0.956433703170866 |
| chr5-24370841-24371342 | 4 | 0.925634405544452 |
| chr19-36340382-36340883 | 8 | 0.895915214399391 |
| chr5-138887036-138887537 | 8 | 0.973116280815416 |
| chr8-69183954-69184455 | 1 | 0.922772070508778 |
| chr2-162529984-162530485 | 6 | 0.991206925481043 |
| chr6-22146407-22146908 | 8 | 0.959458017026557 |
| chr5-89162368-89162869 | 3 | 0.99593553545041 |

| | | |
|---|---|---|
| chr12-9516677-9517178 | 8 | 0.853253286378554 |
| chr1-67776544-67777045 | 4 | 0.874670782760698 |
| chr17-79680742-79681243 | 4 | 0.891065677533935 |
| chr4-106578878-106579379 | 3 | 0.940981975115582 |
| chr16-67821479-67821980 | 3 | 0.921308830965555 |
| chr1-180937113-180937614 | 5 | 0.98417818753991 |
| chr8-40332003-40332504 | 3 | 0.996871297299881 |
| chr16-73157165-73157666 | 8 | 0.993493681128311 |
| chr5-97420710-97421211 | 8 | 0.964314511396098 |
| chr7-22206493-22206994 | 8 | 0.986914964567932 |
| chr12-44545055-44545556 | 6 | 0.870178435990656 |
| chr14-1513212-1513713 | 8 | 0.983521036209224 |
| chr1-156760532-156761033 | 8 | 0.989865485463721 |
| chr10-17566958-17567459 | 8 | 0.930713945620407 |
| chr13-63631323-63631824 | 5 | 0.978807296624394 |
| chr3-106908609-106909110 | 3 | 0.989669059690307 |
| chr5-6842337-6842838 | 8 | 0.928246658588994 |
| chr5-57441252-57441753 | 1 | 0.925706303873513 |
| chr11-67450098-67450599 | 3 | 0.99593553545041 |
| chr7-114552798-114553299 | 8 | 0.842907376592027 |
| chr3-155985623-155986124 | 3 | 0.938725174688406 |
| chr5-118575728-118576229 | 3 | 0.884939933067404 |
| chr2-235622176-235622677 | 3 | 0.993014636108877 |
| chr9-6180968-6181469 | 3 | 0.959995654257502 |
| chr4-158553373-158553874 | 8 | 0.977468450355265 |
| chr4-162227618-162228119 | 3 | 0.915104977540296 |
| chr13-100266715-100267216 | 3 | 0.996688090642187 |
| chr20-31606996-31607497 | 3 | 0.901680031653272 |
| chr11-17270087-17270588 | 3 | 0.988056121279129 |
| chr7-123993957-123994458 | 3 | 0.996232676419417 |
| chr18-11083844-11084345 | 5 | 0.908092388694338 |
| chrX-24146956-24147457 | 6 | 0.843849060927121 |
| chr14-71730778-71731279 | 3 | 0.946066941431386 |
| chr19-39108085-39108586 | 8 | 0.959797589424551 |
| chr15-14663369-14663870 | 7 | 0.860607153327214 |
| chr2-193041280-193041781 | 8 | 0.855266825060221 |
| chr6-26499871-26500372 | 6 | 0.992064616402287 |
| chr11-55881981-55882482 | 8 | 0.973780207144453 |
| chr3-137712262-137712763 | 3 | 0.996805997442978 |
| chr10-5615429-5615930 | 6 | 0.981247049849965 |
| chr4-76989974-76990475 | 5 | 0.990647185669938 |
| chr13-93202239-93202740 | 8 | 0.951380479773096 |
| chr5-86227102-86227603 | 3 | 0.867322143750279 |

| | | |
|---|---|---|
| chr4-92416687-92417188 | 8 | 0.825045621685792 |
| chr6-123237340-123237841 | 8 | 0.883911928870456 |
| chr5-121277522-121278023 | 8 | 0.834398075243152 |
| chr4-61790888-61791389 | 2 | 0.844935670093668 |
| chr15-34961609-34962110 | 4 | 0.874453177606776 |
| chr1-213450215-213450716 | 3 | 0.996649819184355 |
| chr5-127898913-127899414 | 8 | 0.905676296177498 |
| chr18-3741996-3742497 | 8 | 0.982305076683009 |
| chr1-246267272-246267773 | 8 | 0.828139491854749 |
| chr9-47446876-47447377 | 8 | 0.824347259575121 |
| chr6-91846005-91846506 | 3 | 0.989493714619499 |
| chr4-151479457-151479958 | 2 | 0.943210188312171 |
| chr15-37554151-37554652 | 8 | 0.820453341887842 |
| chr5-131142922-131143423 | 6 | 0.991786392415614 |
| chr8-57136532-57137033 | 2 | 0.903503764021731 |
| chr15-44446816-44447317 | 5 | 0.981373567062077 |
| chr10-25034515-25035016 | 8 | 0.984934467337153 |
| chr1-6681391-6681892 | 6 | 0.974029552370462 |
| chr4-67893415-67893916 | 5 | 0.977574851372476 |
| chr18-56902627-56903128 | 8 | 0.825218397959348 |
| chr11-70172623-70173124 | 6 | 0.858683664370486 |
| chr6-107996844-107997345 | 8 | 0.959446154763879 |
| chr20-19335960-19336461 | 4 | 0.852302636872302 |
| chr5-137004996-137005497 | 4 | 0.891687059516836 |
| chr11-75099500-75100001 | 2 | 0.892157712529387 |
| chr17-21937300-21937801 | 6 | 0.96885996310418 |
| chr15-1561344-1561845 | 6 | 0.991811392061077 |
| chr8-20547465-20547966 | 4 | 0.904194007669773 |
| chr9-21761025-21761526 | 3 | 0.915188544001564 |
| chr3-46032280-46032781 | 8 | 0.928146203484723 |
| chr3-103973446-103973947 | 7 | 0.860101738803886 |
| chr7-99238832-99239333 | 2 | 0.901196784095479 |
| chr14-79362574-79363075 | 4 | 0.950620362641962 |
| chrX-25551168-25551669 | 5 | 0.99165349760757 |
| chr3-49239834-49240335 | 8 | 0.918247541391394 |
| chr5-104069515-104070016 | 3 | 0.99383710578225 |
| chr10-75315174-75315675 | 5 | 0.857044901510972 |
| chr15-53174201-53174702 | 1 | 0.940390677511128 |
| chr6-101989876-101990377 | 8 | 0.833429567908101 |
| chr14-35525792-35526293 | 2 | 0.936572925951918 |
| chr7-89907426-89907927 | 4 | 0.924618957222918 |
| chr1-192060318-192060819 | 5 | 0.976027389865621 |
| chr6-44479967-44480468 | 4 | 0.841170277265985 |

| | | |
|---|---|---|
| chr1-31210650-31211151 | 3 | 0.996727161871002 |
| chr15-41600642-41601143 | 8 | 0.853988577461894 |
| chr15-27712216-27712717 | 3 | 0.850754906677795 |
| chr5-106112746-106113247 | 3 | 0.996805938757694 |
| chr14-11889345-11889846 | 6 | 0.942847991812791 |
| chr10-20440326-20440827 | 8 | 0.927346830123819 |
| chr2-126685555-126686056 | 4 | 0.936633229796057 |
| chr1-207089348-207089849 | 8 | 0.82083910465654 |
| chr20-9536892-9537393 | 3 | 0.957071498510491 |
| chr10-57419114-57419615 | 8 | 0.918881863711973 |
| chr18-38396954-38397455 | 6 | 0.960275409692481 |
| chr1-48411395-48411896 | 8 | 0.935580173239425 |
| chr8-55216066-55216567 | 8 | 0.904304593077882 |
| chr5-29191774-29192275 | 5 | 0.988410750236134 |
| chr5-14501816-14502317 | 7 | 0.947187803622909 |
| chr5-38890497-38890998 | 6 | 0.983701108672025 |
| chr13-37199975-37200476 | 5 | 0.837895898395945 |
| chr12-8345280-8345781 | 6 | 0.948020814835666 |
| chr7-17653404-17653905 | 8 | 0.99450907003268 |
| chr13-78171873-78172374 | 3 | 0.996722125977629 |
| chr17-27795873-27796374 | 3 | 0.875728586995099 |
| chr9-17423513-17424014 | 3 | 0.99633494039641 |
| chr16-22829941-22830442 | 8 | 0.984908185577987 |
| chr2-231053353-231053854 | 8 | 0.943377214269783 |
| chr4-97456278-97456779 | 1 | 0.890479082205539 |
| chr4-176131674-176132175 | 5 | 0.859149623765682 |
| chr6-122851415-122851916 | 6 | 0.991758739941446 |
| chr14-40742867-40743368 | 8 | 0.916608420637115 |
| chr10-84890388-84890889 | 3 | 0.997047358474424 |
| chr7-21443919-21444420 | 3 | 0.995688685850912 |
| chr14-9529726-9530227 | 5 | 0.987871451392268 |
| chr4-127024735-127025236 | 2 | 0.831436377494469 |
| chr2-31853390-31853891 | 5 | 0.98812407808238 |
| chr5-82184479-82184980 | 3 | 0.996228875012517 |
| chr6-116422174-116422675 | 6 | 0.992795188810831 |
| chr1-240505995-240506496 | 8 | 0.911503030431784 |
| chr13-105527406-105527907 | 3 | 0.940958257160161 |
| chr9-70521414-70521915 | 3 | 0.96145817669012 |
| chr8-81516039-81516540 | 7 | 0.886213160396446 |
| chr10-58317929-58318430 | 3 | 0.928923050562318 |
| chr15-25706450-25706951 | 8 | 0.836621469971768 |
| chr2-5656230-5656731 | 3 | 0.987052898044121 |
| chr5-143914956-143915457 | 6 | 0.996088871871137 |

| | | |
|---|---|---|
| chr13-24901027-24901528 | 5 | 0.900908977191138 |
| chr6-70357304-70357805 | 1 | 0.816810066930187 |
| chr1-95732145-95732646 | 3 | 0.996588751698973 |
| chr8-28915635-28916136 | 8 | 0.909686723456207 |
| chr9-63323581-63324082 | 4 | 0.867043436772396 |
| chr11-39373620-39374121 | 3 | 0.971670767970734 |
| chr5-135589876-135590377 | 8 | 0.949653521295313 |
| chr6-55604097-55604598 | 7 | 0.847483283288564 |
| chr18-14956338-14956839 | 6 | 0.992794454436234 |
| chr7-9202702-9203203 | 6 | 0.992998587141351 |
| chr7-90356686-90357187 | 5 | 0.971323683338898 |
| chr7-23597177-23597678 | 8 | 0.831974517545997 |
| chr2-177581822-177582323 | 6 | 0.985663733224509 |
| chr2-169710789-169711290 | 5 | 0.983919978840557 |
| chr1-13845695-13846196 | 8 | 0.976576120737881 |
| chr14-75324827-75325328 | 8 | 0.944896943513364 |
| chr14-77264059-77264560 | 6 | 0.9850399734024 |
| chr2-221806461-221806962 | 8 | 0.886778261649899 |
| chr5-131460165-131460666 | 8 | 0.994162277421313 |
| chr4-116481825-116482326 | 2 | 0.879956461494521 |
| chr4-84306480-84306981 | 1 | 0.975184613282796 |
| chr15-62529150-62529651 | 8 | 0.992340172353455 |
| chr18-57227947-57228448 | 3 | 0.922484456224866 |
| chr4-63619798-63620299 | 3 | 0.986193452436488 |
| chr4-7099292-7099793 | 3 | 0.983834804919932 |
| chr1-164031573-164032074 | 1 | 0.930999586338836 |
| chr1-221226167-221226668 | 3 | 0.99593553545041 |
| chr10-89732938-89733439 | 5 | 0.818550469056904 |
| chr18-22754368-22754869 | 6 | 0.859981088678877 |
| chr14-41505712-41506213 | 8 | 0.898106960338444 |
| chr1-107650863-107651364 | 3 | 0.957814915733555 |
| chr2-32468280-32468781 | 5 | 0.991489357272788 |
| chr5-119523066-119523567 | 3 | 0.995822557728283 |
| chrX-106344417-106344918 | 2 | 0.881003832322524 |
| chrX-60992014-60992515 | 3 | 0.976962245041949 |
| chr19-46650646-46651147 | 1 | 0.892264383170292 |
| chr9-28833873-28834374 | 3 | 0.856748381144871 |
| chr3-47156393-47156894 | 8 | 0.89120313674389 |
| chr2-211157117-211157618 | 1 | 0.83621181001791 |
| chr14-63152067-63152568 | 4 | 0.922100075052596 |
| chr16-63235863-63236364 | 8 | 0.96948301637079 |
| chr13-46560355-46560856 | 5 | 0.957935995291546 |
| chr8-76272698-76273199 | 2 | 0.954001949207667 |

| | | |
|---|---|---|
| chr1-220142411-220142912 | 6 | 0.992990919073515 |
| chr9-94615874-94616375 | 3 | 0.996537846399659 |
| chr2-103833424-103833925 | 8 | 0.955221665240566 |
| chr3-79953891-79954392 | 3 | 0.940544675893182 |
| chr3-168363112-168363613 | 3 | 0.996832592577172 |
| chr3-60109212-60109713 | 3 | 0.983152810871207 |
| chr3-105461716-105462217 | 5 | 0.81840104463986 |
| chr4-107081858-107082359 | 3 | 0.996888696033121 |
| chr11-28748281-28748782 | 3 | 0.992775297621559 |
| chr3-102147716-102148217 | 8 | 0.949863756407834 |
| chr7-123272706-123273207 | 2 | 0.951886155271278 |
| chr1-33523754-33524255 | 3 | 0.949101093529429 |
| chr16-55638459-55638960 | 3 | 0.847340256391384 |
| chr4-76279994-76280495 | 5 | 0.990163690561291 |
| chr1-245536639-245537140 | 3 | 0.890914719972348 |
| chr10-35928887-35929388 | 8 | 0.915734445253515 |
| chr11-1231005-1231506 | 4 | 0.947179049160125 |
| chr1-153688317-153688818 | 5 | 0.986626825350919 |
| chr3-16394242-16394743 | 4 | 0.893415964657485 |
| chr8-122459623-122460124 | 6 | 0.993008373142359 |
| chr12-26925996-26926497 | 2 | 0.896437094601247 |
| chr12-42117637-42118138 | 3 | 0.995950387451304 |
| chr19-5472623-5473124 | 3 | 0.979338257783769 |
| chr2-149465957-149466458 | 3 | 0.971599734662774 |
| chr1-164503462-164503963 | 2 | 0.847241640861514 |
| chr13-88214201-88214702 | 5 | 0.967461458119571 |
| chr6-73067965-73068466 | 4 | 0.95164420769682 |
| chr14-72905750-72906251 | 7 | 0.893453538497455 |
| chr1-172005463-172005964 | 6 | 0.993655748339365 |
| chr15-76566830-76567331 | 6 | 0.993066923955338 |
| chr6-89862185-89862686 | 8 | 0.86079588524941 |
| chr2-209880808-209881309 | 5 | 0.983932505587008 |
| chr16-3334156-3334657 | 8 | 0.940442353688817 |
| chr15-93238180-93238681 | 3 | 0.962777847280361 |
| chr11-39135795-39136296 | 5 | 0.828726133898602 |
| chr18-56298696-56299197 | 7 | 0.888268692040811 |
| chr8-6983648-6984149 | 8 | 0.98851000626342 |
| chr8-51147059-51147560 | 8 | 0.852939126557132 |
| chr13-99511602-99512103 | 3 | 0.99593553545041 |
| chr16-70301915-70302416 | 3 | 0.976919905328209 |
| chr10-100719271-100719772 | 5 | 0.918097677876677 |
| chr15-82103293-82103794 | 8 | 0.870128010785818 |
| chr13-73719005-73719506 | 2 | 0.911533247715922 |

| | | |
|---|---|---|
| chr8-64163455-64163956 | 8 | 0.911095781268492 |
| chr15-6509168-6509669 | 3 | 0.95931700466165 |
| chr5-77593832-77594333 | 8 | 0.995090100937469 |
| chr14-66860816-66861317 | 8 | 0.862818634043759 |
| chr4-52234084-52234585 | 1 | 0.956223067841829 |
| chr2-43100154-43100655 | 4 | 0.847042122728054 |
| chr14-47992773-47993274 | 3 | 0.982776200776766 |
| chr4-159326861-159327362 | 8 | 0.879244437505464 |
| chr14-4464998-4465499 | 3 | 0.977201323848812 |
| chr10-45171783-45172284 | 8 | 0.980322948325665 |
| chr11-45370482-45370983 | 8 | 0.982176947026403 |
| chr1-130635015-130635516 | 7 | 0.849052854027824 |
| chr10-95382469-95382970 | 5 | 0.872597070475756 |
| chr7-43051735-43052236 | 3 | 0.996747305008973 |
| chr1-169656305-169656806 | 1 | 0.906823585305894 |
| chr10-50737633-50738134 | 3 | 0.826236638460602 |
| chr1-176436059-176436560 | 3 | 0.906249547012047 |
| chr7-12290545-12291046 | 3 | 0.876671258600564 |
| chr2-60916626-60917127 | 4 | 0.857212698846799 |
| chr9-23603163-23603664 | 6 | 0.984989968123706 |
| chr18-77722654-77723155 | 5 | 0.972735062523435 |
| chr4-175885370-175885871 | 7 | 0.853874753533865 |
| chr8-103931685-103932186 | 8 | 0.933004315818261 |
| chr7-70811645-70812146 | 3 | 0.844560942318938 |
| chr6-120323196-120323697 | 8 | 0.830645908955244 |
| chr4-59831546-59832047 | 8 | 0.99052987696514 |
| chr3-129911379-129911880 | 8 | 0.993493681128311 |
| chr14-94102386-94102887 | 8 | 0.948594750826412 |
| chr2-215619402-215619903 | 8 | 0.967220404911234 |
| chr1-198084907-198085408 | 5 | 0.933648201454109 |
| chr1-194440041-194440542 | 8 | 0.892764189814827 |
| chr4-116630879-116631380 | 4 | 0.942864401040341 |
| chr11-57059358-57059859 | 2 | 0.929441922722474 |
| chr5-114095735-114096236 | 3 | 0.991998958269851 |
| chr14-66578585-66579086 | 8 | 0.904185975454567 |
| chr8-111153987-111154488 | 8 | 0.834581555695335 |
| chr2-195873711-195874212 | 3 | 0.996306490449243 |
| chr1-202689877-202690378 | 8 | 0.95825970834866 |
| chr8-96034557-96035058 | 7 | 0.947785294946563 |
| chr18-35417341-35417842 | 8 | 0.816969444091263 |
| chr6-102538655-102539156 | 6 | 0.913378744942233 |
| chr12-11108125-11108626 | 6 | 0.992028049541858 |
| chr15-76519359-76519860 | 6 | 0.92862864086746 |

| | | |
|---|---|---|
| chr1-36316884-36317385 | 3 | 0.99593553545041 |
| chr10-89565339-89565840 | 3 | 0.99593553545041 |
| chr8-44114836-44115337 | 8 | 0.888205482027925 |
| chr11-33285119-33285620 | 1 | 0.824891170143562 |
| chr19-34186261-34186762 | 5 | 0.920784678422311 |
| chr4-77624688-77625189 | 3 | 0.960371243372148 |
| chr19-38732649-38733150 | 8 | 0.972373217578345 |
| chr6-100875964-100876465 | 4 | 0.892696616993626 |
| chr3-142789297-142789798 | 8 | 0.926864961984217 |
| chr2-11063262-11063763 | 3 | 0.995997033697535 |
| chrX-100013025-100013526 | 6 | 0.974317888375527 |
| chr7-100815441-100815942 | 3 | 0.867500362873127 |
| chr6-103343803-103344304 | 6 | 0.981758700436665 |
| chr6-48499918-48500419 | 3 | 0.892403616405937 |
| chr2-60785393-60785894 | 6 | 0.991206925481043 |
| chr11-74246097-74246598 | 1 | 0.897260077793368 |
| chr18-24867252-24867753 | 7 | 0.948481075483925 |
| chr1-185182981-185183482 | 1 | 0.958021564551759 |
| chr11-75893611-75894112 | 5 | 0.987260681312237 |
| chr3-79581190-79581691 | 6 | 0.991206925481043 |
| chr3-22100630-22101131 | 8 | 0.849476596975546 |
| chr16-35632336-35632837 | 3 | 0.99593553545041 |
| chr16-77584068-77584569 | 4 | 0.820329681149229 |
| chr16-64734887-64735388 | 5 | 0.992744791552034 |
| chr14-19001793-19002294 | 3 | 0.99177007701126 |
| chr11-58022712-58023213 | 4 | 0.85517690860011 |
| chr2-118518505-118519006 | 8 | 0.986716884343143 |
| chr9-105190686-105191187 | 3 | 0.942159811261228 |
| chr6-119811986-119812487 | 7 | 0.88486710514848 |
| chrX-128268058-128268559 | 3 | 0.955940388607303 |
| chr2-170680324-170680825 | 8 | 0.953056809306581 |
| chr12-11442120-11442621 | 8 | 0.912317771539753 |
| chr3-89717261-89717762 | 8 | 0.980877817768993 |
| chr2-113797184-113797685 | 3 | 0.996731311350224 |
| chr15-69135729-69136230 | 4 | 0.885499705641588 |
| chr5-154719228-154719729 | 7 | 0.821331392570717 |
| chr2-4029508-4030009 | 3 | 0.961108566639508 |
| chr13-20640916-20641417 | 8 | 0.991800669829484 |
| chr4-34300090-34300591 | 5 | 0.94985942609078 |
| chr5-25497354-25497855 | 8 | 0.82032327881397 |
| chr9-108396039-108396540 | 5 | 0.835437570532175 |
| chr18-32940879-32941380 | 3 | 0.988968792270182 |
| chr17-18576928-18577429 | 3 | 0.99593553545041 |

| | | |
|---|---|---|
| chr15-8109493-8109994 | 8 | 0.951068412751425 |
| chr4-3888533-3889034 | 8 | 0.914165473139477 |
| chr6-123344672-123345173 | 8 | 0.837348439657616 |
| chr7-70678350-70678851 | 5 | 0.968424704426692 |
| chr15-76260925-76261426 | 4 | 0.930371170475037 |
| chr2-61990722-61991223 | 6 | 0.992716325582624 |
| chr2-213350104-213350605 | 6 | 0.992413150323096 |
| chr12-35962929-35963430 | 6 | 0.843814381390545 |
| chr15-36744308-36744809 | 5 | 0.983919978840557 |
| chr20-24531248-24531749 | 5 | 0.983625131588857 |
| chr8-29955153-29955654 | 2 | 0.836122916721081 |
| chr3-8744384-8744885 | 8 | 0.955284917971869 |
| chr16-57658793-57659294 | 3 | 0.941362265788453 |
| chr17-85781370-85781871 | 3 | 0.984905962280536 |
| chr3-136265032-136265533 | 3 | 0.991239851186375 |
| chr15-72479507-72480008 | 3 | 0.991860036228589 |
| chr14-81318650-81319151 | 2 | 0.868052014984466 |
| chr5-49259305-49259806 | 3 | 0.995449553135518 |
| chr2-242956708-242957209 | 8 | 0.988289877434572 |
| chr1-184115755-184116256 | 8 | 0.964362891598838 |
| chr1-90973846-90974347 | 8 | 0.815351802713358 |
| chr2-16161996-16162497 | 3 | 0.890597863731363 |
| chr2-138784698-138785199 | 8 | 0.995160239051892 |
| chr13-76910245-76910746 | 1 | 0.892577153804324 |
| chr11-84746990-84747491 | 8 | 0.88887 |
| chr1-188150225-188150726 | 1 | 0.918894680708066 |
| chr11-78297422-78297923 | 8 | 0.987227534621699 |
| chr1-50118704-50119205 | 7 | 0.855352307475731 |
| chr17-59101715-59102216 | 6 | 0.991206925481043 |
| chr4-121215679-121216180 | 1 | 0.816746216215839 |
| chrX-152287099-152287600 | 4 | 0.964618682704522 |
| chr15-98300927-98301428 | 2 | 0.949829822203438 |
| chr19-30556180-30556681 | 5 | 0.990817132335935 |
| chr1-136657290-136657791 | 3 | 0.99663282948166 |
| chr2-107641220-107641721 | 8 | 0.850627584340837 |
| chr11-77859242-77859743 | 6 | 0.991206925481043 |
| chr9-6199604-6200105 | 5 | 0.968512896685289 |
| chr10-72698185-72698686 | 6 | 0.992552023256053 |
| chr4-137175620-137176121 | 3 | 0.995825473982587 |
| chr13-55398877-55399378 | 2 | 0.833421036626739 |
| chr12-1423749-1424250 | 5 | 0.937222071892364 |
| chr6-57141269-57141770 | 3 | 0.910690231834575 |
| chr12-43737091-43737592 | 8 | 0.863131840444732 |

| | | |
|---|---|---|
| chr2-125732411-125732912 | 6 | 0.974840178602513 |
| chr17-37920848-37921349 | 3 | 0.991791928128403 |
| chr6-52806776-52807277 | 4 | 0.879661629671029 |
| chrX-67200593-67201094 | 3 | 0.930275146330359 |
| chr2-22953742-22954243 | 6 | 0.981410337281078 |
| chr20-14508985-14509486 | 3 | 0.99593553545041 |
| chr10-40174643-40175144 | 8 | 0.838799094697068 |
| chr1-161550923-161551424 | 8 | 0.991034854304957 |
| chr19-40759572-40760073 | 3 | 0.973580707564204 |
| chr17-67850332-67850833 | 1 | 0.881006635172288 |
| chr3-40485104-40485605 | 2 | 0.878197784076833 |
| chr9-33235614-33236115 | 4 | 0.869349385831244 |
| chr2-233676770-233677271 | 5 | 0.882536586037042 |
| chr5-139168646-139169147 | 1 | 0.940368554118086 |
| chr3-42961555-42962056 | 8 | 0.963532881296278 |
| chr10-51077294-51077795 | 3 | 0.984106657165537 |
| chr1-240016615-240017116 | 1 | 0.954444222515315 |
| chr10-87635635-87636136 | 5 | 0.989346985357881 |
| chr3-51073384-51073885 | 6 | 0.993183386633752 |
| chr8-97446514-97447015 | 4 | 0.925572800348165 |
| chr6-30625338-30625839 | 3 | 0.961539864229366 |
| chr10-71858994-71859495 | 8 | 0.937139468239671 |
| chr6-103842443-103842944 | 2 | 0.884446522030512 |
| chr2-106680486-106680987 | 3 | 0.974539622700298 |
| chr9-67818002-67818503 | 3 | 0.99593553545041 |
| chr2-113372866-113373367 | 1 | 0.90564353219494 |
| chr1-12916402-12916903 | 4 | 0.883332390608464 |
| chr2-33330188-33330689 | 5 | 0.846918583872217 |
| chr4-153141118-153141619 | 4 | 0.832375974059703 |
| chr7-11280012-11280513 | 8 | 0.9529202772098 |
| chr17-74695966-74696467 | 7 | 0.892858848180034 |
| chr4-131622621-131623122 | 4 | 0.946628607399142 |
| chr18-2463746-2464247 | 4 | 0.896748370395811 |
| chr17-12385191-12385692 | 8 | 0.994945621465006 |
| chr4-12612180-12612681 | 8 | 0.96975700251019 |
| chr11-67471935-67472436 | 2 | 0.900483651622952 |
| chr7-72128906-72129407 | 6 | 0.993105317734455 |
| chr14-67980792-67981293 | 7 | 0.96297195504115 |
| chr4-177787121-177787622 | 5 | 0.975291351842227 |
| chr17-48747753-48748254 | 6 | 0.913151999940065 |
| chr1-104305976-104306477 | 6 | 0.9928674922501 |
| chr14-33388371-33388872 | 8 | 0.991872411605247 |
| chr6-23341653-23342154 | 3 | 0.995957116372111 |

| | | |
|---|---|---|
| chr15-14235048-14235549 | 7 | 0.817018484302612 |
| chr4-82122514-82123015 | 6 | 0.836429802372169 |
| chr17-10549194-10549695 | 3 | 0.988935764467153 |
| chr1-79137168-79137669 | 5 | 0.992316404591661 |
| chr20-12513464-12513965 | 8 | 0.932660755413315 |
| chr11-11315381-11315882 | 5 | 0.940106575688647 |
| chr1-133909191-133909692 | 5 | 0.983770031856208 |
| chr1-117415338-117415839 | 4 | 0.874368872116265 |
| chr13-41636831-41637332 | 8 | 0.889225046755228 |
| chr17-20931461-20931962 | 8 | 0.878856798864102 |
| chr15-62822224-62822725 | 4 | 0.828313046960974 |
| chr17-6131443-6131944 | 3 | 0.997035489364825 |
| chr3-93402054-93402555 | 3 | 0.977419882653856 |
| chr1-51921102-51921603 | 5 | 0.978558447076817 |
| chr3-166769737-166770238 | 2 | 0.835819349773514 |
| chr13-76221090-76221591 | 6 | 0.993228316686902 |
| chr1-135213902-135214403 | 7 | 0.824161228360168 |
| chr7-63653648-63654149 | 4 | 0.948241010460401 |
| chr16-1459039-1459540 | 1 | 0.954222709488962 |
| chr13-41784724-41785225 | 5 | 0.910882963138843 |
| chr3-57516900-57517401 | 8 | 0.895949524764894 |
| chr4-66733657-66734158 | 3 | 0.99761279962836 |
| chr14-50134149-50134650 | 8 | 0.976507476761123 |
| chr3-16850572-16851073 | 8 | 0.995181983732015 |
| chr16-64704284-64704785 | 4 | 0.891485226985186 |
| chr12-4705034-4705535 | 2 | 0.905135378220355 |
| chr6-20898561-20899062 | 5 | 0.963684234329365 |
| chr11-54164171-54164672 | 8 | 0.953996342034152 |
| chr4-28723090-28723591 | 8 | 0.867973912652141 |
| chr2-245959253-245959754 | 6 | 0.983132580639028 |
| chr15-80911875-80912376 | 3 | 0.993321762633891 |
| chr1-195045182-195045683 | 8 | 0.871208206222072 |
| chr5-71435490-71435991 | 5 | 0.98801842987901 |
| chr1-214835169-214835670 | 2 | 0.945669400580585 |
| chr2-23432852-23433353 | 6 | 0.990349159902184 |
| chr5-21879145-21879646 | 7 | 0.842139026234623 |
| chr8-109789188-109789689 | 4 | 0.966123798488476 |
| chr4-28795451-28795952 | 4 | 0.848051789857357 |
| chr13-37628130-37628631 | 6 | 0.98058672658214 |
| chr6-58605488-58605989 | 3 | 0.978013172201084 |
| chr14-17822900-17823401 | 6 | 0.989142144948467 |
| chr4-149913748-149914249 | 1 | 0.884390504612024 |
| chr11-5409067-5409568 | 6 | 0.931693601071094 |

| | | |
|---|---|---|
| chr13-93439158-93439659 | 8 | 0.995069440976113 |
| chr14-55390063-55390564 | 6 | 0.96798862743366 |
| chr1-208927178-208927679 | 8 | 0.972972597056571 |
| chr5-68940667-68941168 | 3 | 0.99593553545041 |
| chr3-90833970-90834471 | 8 | 0.868108583884869 |
| chr10-76750082-76750583 | 8 | 0.969937932170353 |
| chr14-32265552-32266053 | 6 | 0.852060699227979 |
| chr1-36684701-36685202 | 1 | 0.849046098424177 |
| chr15-20115382-20115883 | 8 | 0.849759968028015 |
| chr6-72906246-72906747 | 3 | 0.97819718603282 |
| chr7-125921796-125922297 | 5 | 0.990858330119843 |
| chr7-58779607-58780108 | 4 | 0.90173856004343 |
| chr3-79870367-79870868 | 4 | 0.925713786337599 |
| chr2-233696562-233697063 | 4 | 0.935735662936926 |
| chr9-108699791-108700292 | 2 | 0.852874915032972 |
| chr2-116747881-116748382 | 6 | 0.892510474404401 |
| chr20-23611290-23611791 | 3 | 0.836047961585812 |
| chr18-1795078-1795579 | 3 | 0.978613725064332 |
| chr18-51785498-51785999 | 8 | 0.97219558507053 |
| chr15-75396414-75396915 | 2 | 0.945022322362589 |
| chr7-73221186-73221687 | 8 | 0.969736700139121 |
| chr16-73380862-73381363 | 8 | 0.990390868220918 |
| chr2-141877700-141878201 | 6 | 0.988414215054656 |
| chr2-229059206-229059707 | 8 | 0.970117867987065 |
| chr3-124944819-124945320 | 6 | 0.985394672230491 |
| chr2-113972609-113973110 | 3 | 0.959753005333797 |
| chr13-64737586-64738087 | 7 | 0.830221273291046 |
| chr5-157181952-157182453 | 8 | 0.993380317818848 |
| chrX-108077355-108077856 | 4 | 0.849283589199514 |
| chr10-67319991-67320492 | 7 | 0.823769025296504 |
| chr13-97019720-97020221 | 8 | 0.844231083263315 |
| chr13-41346118-41346619 | 8 | 0.985284590598188 |
| chr7-7924871-7925372 | 3 | 0.885766425913368 |
| chr13-104295856-104296357 | 6 | 0.943649707610575 |
| chr1-155129560-155130061 | 8 | 0.922147203251748 |
| chr3-54907555-54908056 | 1 | 0.88801003044325 |
| chr5-128170204-128170705 | 3 | 0.995735609772143 |
| chr5-120074949-120075450 | 3 | 0.983815760499316 |
| chr16-19369260-19369761 | 2 | 0.827296670268714 |
| chr2-198871285-198871786 | 3 | 0.937136598848331 |
| chr19-43471231-43471732 | 3 | 0.994686403711227 |
| chr17-23831234-23831735 | 8 | 0.849466318264473 |
| chr3-118127693-118128194 | 5 | 0.979087220176682 |

| | | |
|---|---|---|
| chr14-22504960-22505461 | 3 | 0.901105899259318 |
| chr3-83118291-83118792 | 3 | 0.99593553545041 |
| chr3-144652751-144653252 | 5 | 0.988770912564535 |
| chr6-43295066-43295567 | 2 | 0.883258028359315 |
| chr15-47170979-47171480 | 8 | 0.938702750405717 |
| chr16-60747789-60748290 | 4 | 0.859227975383635 |
| chr15-95637636-95638137 | 3 | 0.985111840819176 |
| chr1-19798143-19798644 | 8 | 0.906007446140777 |
| chr19-26410151-26410652 | 3 | 0.993910823927479 |
| chr13-93812115-93812616 | 3 | 0.90348885834254 |
| chr4-23272191-23272692 | 6 | 0.960168137431802 |
| chr7-12687477-12687978 | 5 | 0.890695416615565 |
| chr19-16929177-16929678 | 6 | 0.833872241972305 |
| chr2-205243548-205244049 | 1 | 0.931537275063198 |
| chr17-1523245-1523746 | 8 | 0.835244863761634 |
| chr1-152622423-152622924 | 5 | 0.989122971242473 |
| chr9-109984644-109985145 | 5 | 0.888868241848419 |
| chr2-91649044-91649545 | 8 | 0.827469035033708 |
| chr8-73517894-73518395 | 5 | 0.99085061468057 |
| chr8-51179931-51180432 | 7 | 0.941269063867234 |
| chr13-46488743-46489244 | 4 | 0.873104647378593 |
| chr16-11399060-11399561 | 6 | 0.991206925481043 |
| chr4-104162224-104162725 | 8 | 0.924063101929606 |
| chr20-31925252-31925753 | 3 | 0.99593553545041 |
| chr1-30523103-30523604 | 2 | 0.824340483400096 |
| chr6-126957860-126958361 | 2 | 0.954523427714695 |
| chr11-65221768-65222269 | 3 | 0.87471445047726 |
| chr3-45990778-45991279 | 6 | 0.990963542360934 |
| chr4-61433085-61433586 | 8 | 0.961904375154081 |
| chr5-149460740-149461241 | 6 | 0.923801099330946 |
| chr5-117929285-117929786 | 8 | 0.935438384473534 |
| chrX-64341487-64341988 | 2 | 0.930976629323021 |
| chr9-106033253-106033754 | 5 | 0.962881361624681 |
| chr7-21338079-21338580 | 3 | 0.992994408485222 |
| chr11-25156751-25157252 | 6 | 0.991206925481043 |
| chr8-116902554-116903055 | 2 | 0.864549868873347 |
| chr6-118861053-118861554 | 8 | 0.985183000912591 |
| chr5-14316957-14317458 | 3 | 0.816812391609053 |
| chr10-44415092-44415593 | 8 | 0.932224584611631 |
| chr10-18901053-18901554 | 8 | 0.941496033872185 |
| chr2-237126899-237127400 | 6 | 0.99293276700761 |
| chr2-80376515-80377016 | 6 | 0.993226869046765 |
| chr1-91323999-91324500 | 3 | 0.996365778240591 |

| | | |
|---|---|---|
| chr3-32407228-32407729 | 5 | 0.931305634660025 |
| chr5-57056547-57057048 | 3 | 0.96955638117889 |
| chr2-33867689-33868190 | 8 | 0.874612460208834 |
| chr11-79338159-79338660 | 8 | 0.838708833803653 |
| chr16-83969200-83969701 | 8 | 0.994907765504038 |
| chr1-56018922-56019423 | 3 | 0.995960105884224 |
| chr11-14465325-14465826 | 3 | 0.9916440437903 |
| chr17-32387502-32388003 | 4 | 0.850501020790549 |
| chr2-220549712-220550213 | 6 | 0.986255957874881 |
| chr14-65107753-65108254 | 7 | 0.897150019960263 |
| chr3-40795279-40795780 | 6 | 0.991206925481043 |
| chr9-73171576-73172077 | 8 | 0.934953310398722 |
| chr3-62287455-62287956 | 4 | 0.907019941325929 |
| chr3-152406941-152407442 | 3 | 0.979258559310043 |
| chr1-32038375-32038876 | 6 | 0.991206925481043 |
| chr10-43960413-43960914 | 8 | 0.8213300118745 |
| chr1-29583754-29584255 | 1 | 0.881375280449929 |
| chr9-29200350-29200851 | 8 | 0.967253596440151 |
| chr5-62209505-62210006 | 1 | 0.970889307046725 |
| chr1-31501786-31502287 | 5 | 0.961466685453333 |
| chr3-68789837-68790338 | 1 | 0.862620037242107 |
| chr15-101463374-101463875 | 3 | 0.865406757165348 |
| chr1-225793848-225794349 | 5 | 0.971775 |
| chr3-125608654-125609155 | 6 | 0.979034787104312 |
| chr4-161671600-161672101 | 3 | 0.991720008807184 |
| chr18-46627007-46627508 | 5 | 0.970725420241972 |
| chr20-32843218-32843719 | 3 | 0.945343988554649 |
| chr8-109777373-109777874 | 8 | 0.870664484494034 |
| chr7-91510950-91511451 | 5 | 0.831394735397476 |
| chr9-68245137-68245638 | 6 | 0.956032018471106 |
| chr18-59283671-59284172 | 4 | 0.833131991496237 |
| chr20-34525661-34526162 | 3 | 0.99593553545041 |
| chr6-123804116-123804617 | 6 | 0.986737697245501 |
| chr8-106493474-106493975 | 6 | 0.993134416967774 |
| chr19-1226632-1227133 | 3 | 0.827945107750171 |
| chr2-82133056-82133557 | 3 | 0.99593553545041 |
| chr6-68545087-68545588 | 8 | 0.840564328535953 |
| chr18-7919606-7920107 | 8 | 0.955734365467809 |
| chr17-36839063-36839564 | 2 | 0.912273146803534 |
| chr15-55526948-55527449 | 8 | 0.973839823094862 |
| chr7-65924727-65925228 | 1 | 0.937811745011499 |
| chr5-109502680-109503181 | 3 | 0.996891894509581 |
| chr13-77537706-77538207 | 3 | 0.993312750081796 |

| | | |
|---|---|---|
| chr6-67999409-67999910 | 2 | 0.947883806159316 |
| chr10-48179957-48180458 | 8 | 0.841160927937154 |
| chr3-162397237-162397738 | 3 | 0.983577473351225 |
| chr11-62002188-62002689 | 5 | 0.974910699515799 |
| chr14-4796409-4796910 | 6 | 0.994544647350829 |
| chr6-28413320-28413821 | 8 | 0.985769833275267 |
| chr14-75778662-75779163 | 2 | 0.822069401682496 |
| chr15-66019065-66019566 | 8 | 0.863495229066827 |
| chr5-162553190-162553691 | 2 | 0.948704031652424 |
| chr3-15667191-15667692 | 6 | 0.916151033786406 |
| chr7-67705627-67706128 | 3 | 0.872725474131588 |
| chr13-42937784-42938285 | 8 | 0.913780327060353 |
| chr2-4023115-4023616 | 3 | 0.98680630442556 |
| chr14-17837985-17838486 | 5 | 0.980848534166288 |
| chr2-42049308-42049809 | 4 | 0.896322121591067 |
| chr1-241915273-241915774 | 3 | 0.995823801966529 |
| chr15-64405535-64406036 | 3 | 0.99593553545041 |
| chr1-135575801-135576302 | 8 | 0.993493681128311 |
| chr8-69784035-69784536 | 3 | 0.99593553545041 |
| chr20-25595724-25596225 | 5 | 0.947608959567483 |
| chr2-198063036-198063537 | 3 | 0.996888159078554 |
| chr8-62362779-62363280 | 8 | 0.859937605940569 |
| chr3-56573119-56573620 | 5 | 0.989230884588909 |
| chr9-70685292-70685793 | 5 | 0.992456208344287 |
| chr3-79769668-79770169 | 7 | 0.957420916153873 |
| chr6-51948422-51948923 | 8 | 0.851370711845724 |
| chr5-104414813-104415314 | 3 | 0.856725733031467 |
| chr6-129069379-129069880 | 8 | 0.9868297832212 |
| chr7-43892453-43892954 | 3 | 0.98171641150996 |
| chr14-99610696-99611197 | 3 | 0.995578825850607 |
| chr13-38748352-38748853 | 8 | 0.873449707761335 |
| chr19-10357114-10357615 | 8 | 0.993107506000516 |
| chr10-86680487-86680988 | 8 | 0.959292062059821 |
| chr2-184918056-184918557 | 3 | 0.996896689926473 |
| chr14-58187723-58188224 | 5 | 0.820983830266399 |
| chr5-158219253-158219754 | 3 | 0.894926935849817 |
| chr10-44885511-44886012 | 4 | 0.924865513112585 |
| chr5-71271092-71271593 | 3 | 0.99593553545041 |
| chr6-65326527-65327028 | 3 | 0.990910647688105 |
| chr16-67997464-67997965 | 8 | 0.96278270686168 |
| chr8-68067908-68068409 | 8 | 0.838718028655509 |
| chr20-53740172-53740673 | 8 | 0.988808476443567 |
| chr9-48329432-48329933 | 6 | 0.870334407814264 |

| | | |
|---|---|---|
| chr4-7383520-7384021 | 3 | 0.990290280430965 |
| chr11-44365658-44366159 | 3 | 0.874283203072147 |
| chr15-15559850-15560351 | 6 | 0.932403840600557 |
| chr17-52381816-52382317 | 8 | 0.920510667966415 |
| chr2-86221028-86221529 | 8 | 0.971738563500609 |
| chr7-116846407-116846908 | 8 | 0.834206472742823 |
| chr14-17823561-17824062 | 6 | 0.98468173299478 |
| chr3-122268139-122268640 | 8 | 0.977678364025169 |
| chr3-28586211-28586712 | 8 | 0.984628770622273 |
| chr2-17731194-17731695 | 8 | 0.91849192075006 |
| chr14-94118604-94119105 | 8 | 0.854658748858751 |
| chr3-113334576-113335077 | 8 | 0.979661179852662 |
| chr1-223965417-223965918 | 3 | 0.971497451853858 |
| chr20-12151154-12151655 | 6 | 0.997217992352759 |
| chr2-223644037-223644538 | 1 | 0.957925920092624 |
| chr6-125966281-125966782 | 6 | 0.988951661767663 |
| chr7-33070683-33071184 | 6 | 0.965546442471794 |
| chr20-10072758-10073259 | 4 | 0.871294163240185 |
| chr19-46585271-46585772 | 1 | 0.830552206965508 |
| chr5-57538161-57538662 | 5 | 0.9006682146437 |
| chr11-82607232-82607733 | 4 | 0.928924615708572 |
| chrX-31512753-31513254 | 5 | 0.953831372118193 |
| chr3-35350400-35350901 | 8 | 0.943257764726447 |
| chr13-95243836-95244337 | 6 | 0.993025486282457 |
| chrX-40583442-40583943 | 3 | 0.99593553545041 |
| chr14-57660257-57660758 | 8 | 0.95415639774569 |
| chr18-71259291-71259792 | 5 | 0.977381817555229 |
| chr6-126224214-126224715 | 4 | 0.973643973222974 |
| chrX-105327996-105328497 | 6 | 0.992271208191485 |
| chr1-255980929-255981430 | 8 | 0.876439724145167 |
| chr9-89471195-89471696 | 8 | 0.995160239051892 |
| chr14-71939824-71940325 | 8 | 0.993493681128311 |
| chr7-104341136-104341637 | 8 | 0.9865642634219 |
| chr8-116156307-116156808 | 3 | 0.992708839550249 |
| chr17-20952440-20952941 | 6 | 0.987402218731968 |
| chr16-8532069-8532570 | 3 | 0.997443689175671 |
| chr6-8631381-8631882 | 6 | 0.99534553783543 |
| chr12-26335980-26336481 | 6 | 0.993180159286168 |
| chr14-81671486-81671987 | 5 | 0.989554287606606 |
| chr13-66055296-66055797 | 8 | 0.977258363593895 |
| chr5-71914206-71914707 | 5 | 0.988377548401991 |
| chr1-182344853-182345354 | 8 | 0.827523755976996 |
| chr4-40837614-40838115 | 1 | 0.916072252063111 |

| | | |
|---|---|---|
| chr2-203934637-203935138 | 8 | 0.974370950008889 |
| chr4-35223863-35224364 | 8 | 0.970680698093818 |
| chr9-68529011-68529512 | 6 | 0.849414898264814 |
| chr2-219003276-219003777 | 8 | 0.826287690118711 |
| chr1-56233856-56234357 | 1 | 0.868260243360313 |
| chr1-233609355-233609856 | 6 | 0.997558872160391 |
| chr10-70119925-70120426 | 6 | 0.991810075242147 |
| chr15-12003049-12003550 | 6 | 0.992222025426775 |
| chr7-29616311-29616812 | 8 | 0.92320190733933 |
| chr7-20441768-20442269 | 5 | 0.98750436841325 |
| chr4-95356379-95356880 | 6 | 0.987817953055755 |
| chr1-99650618-99651119 | 2 | 0.868606501268806 |
| chr14-66520117-66520618 | 4 | 0.869715707699634 |
| chr15-52827553-52828054 | 3 | 0.995965984901191 |
| chr2-48737767-48738268 | 5 | 0.824314429781956 |
| chr1-3332943-3333444 | 7 | 0.840607454931359 |
| chr18-60343533-60344034 | 8 | 0.949544323177106 |
| chr4-70539696-70540197 | 3 | 0.99607436113543 |
| chr13-77023089-77023590 | 4 | 0.846030517254772 |
| chr6-16664594-16665095 | 5 | 0.973683621499041 |
| chr16-6977057-6977558 | 4 | 0.859233081976908 |
| chr8-72699979-72700480 | 8 | 0.993493681128311 |
| chr5-7491816-7492317 | 5 | 0.955367468448348 |
| chr13-103229479-103229980 | 3 | 0.819630784225953 |
| chr8-56431100-56431601 | 3 | 0.99593553545041 |
| chr16-81436946-81437447 | 8 | 0.875260563345153 |
| chr1-91817127-91817628 | 3 | 0.99593553545041 |
| chr18-75767226-75767727 | 8 | 0.993493681128311 |
| chr20-18762530-18763031 | 8 | 0.850537690704263 |
| chr17-52034383-52034884 | 5 | 0.982750613684041 |
| chr9-100691303-100691804 | 3 | 0.99593553545041 |
| chr15-64441437-64441938 | 2 | 0.925087516688325 |
| chr18-53369748-53370249 | 8 | 0.829335447677946 |
| chr7-52044412-52044913 | 6 | 0.894469038592361 |
| chr3-102820859-102821360 | 2 | 0.941728006904332 |
| chr7-64324573-64325074 | 6 | 0.991206925481043 |
| chr5-51439660-51440161 | 2 | 0.93955044169553 |
| chr11-83325235-83325736 | 3 | 0.882956376466689 |
| chr5-22168513-22169014 | 8 | 0.975448803072319 |
| chr20-2901682-2902183 | 5 | 0.981999636990036 |
| chr14-22308973-22309474 | 5 | 0.982429362085923 |
| chr5-12556297-12556798 | 5 | 0.989400580943028 |
| chr4-94758793-94759294 | 3 | 0.841765548187705 |

| | | |
|---|---|---|
| chr7-26474962-26475463 | 3 | 0.996894645922101 |
| chrX-10197935-10198436 | 4 | 0.965445614470174 |
| chr5-20399908-20400409 | 8 | 0.905548457249927 |
| chr18-5078037-5078538 | 3 | 0.997009806063225 |
| chr4-178034159-178034660 | 5 | 0.983793481382367 |
| chr10-3367680-3368181 | 6 | 0.892067392967755 |
| chr14-41836517-41837018 | 8 | 0.90860026458806 |
| chr19-36939069-36939570 | 8 | 0.993493681128311 |
| chr2-221340261-221340762 | 8 | 0.850198850908147 |
| chr2-101238531-101239032 | 2 | 0.904817350304485 |
| chr1-219326404-219326905 | 4 | 0.885885777078264 |
| chr11-48474700-48475201 | 8 | 0.864315394766629 |
| chr18-42766422-42766923 | 3 | 0.886686687684039 |
| chr3-79117374-79117875 | 1 | 0.969415483091366 |
| chr11-63968726-63969227 | 8 | 0.934055983953227 |
| chr18-55561492-55561993 | 3 | 0.99807578289294 |
| chr3-160478748-160479249 | 8 | 0.889664656344722 |
| chr13-50572902-50573403 | 4 | 0.93217962963082 |
| chr5-62335943-62336444 | 3 | 0.996790518280796 |
| chr15-50277299-50277800 | 6 | 0.987242358840542 |
| chr6-106974941-106975442 | 3 | 0.991775522170385 |
| chr13-77970946-77971447 | 3 | 0.997842994268696 |
| chr15-11283523-11284024 | 3 | 0.99593553545041 |
| chr5-115924318-115924819 | 8 | 0.866036380897894 |
| chr8-32596512-32597013 | 6 | 0.938724077092084 |
| chr2-173224218-173224719 | 2 | 0.816683714900814 |
| chr5-128653800-128654301 | 5 | 0.91540897872675 |
| chr13-91477393-91477894 | 8 | 0.968953322980912 |
| chr9-105652386-105652887 | 8 | 0.94978549330327 |
| chr12-14549828-14550329 | 8 | 0.976182710149052 |
| chr8-52600860-52601361 | 3 | 0.980170362721279 |
| chr19-39317015-39317516 | 5 | 0.819891994762097 |
| chr3-140372309-140372810 | 8 | 0.991663606893249 |
| chr10-23420338-23420839 | 6 | 0.981199389152478 |
| chr6-110066125-110066626 | 5 | 0.944230903991495 |
| chr8-30052416-30052917 | 5 | 0.971096807877123 |
| chr13-82941896-82942397 | 5 | 0.99158103843501 |
| chr19-9485315-9485816 | 6 | 0.864281034554335 |
| chr6-46981243-46981744 | 3 | 0.995768997266088 |
| chr14-71954600-71955101 | 8 | 0.981161283649974 |
| chr15-17747896-17748397 | 4 | 0.823524474564755 |
| chr2-137531523-137532024 | 5 | 0.971431096976248 |
| chr3-125267238-125267739 | 3 | 0.983398654265866 |

| | | |
|---|---|---|
| chr2-13875865-13876366 | 3 | 0.996718654568811 |
| chr2-4627159-4627660 | 5 | 0.959627847613344 |
| chr10-72431552-72432053 | 8 | 0.823726115041487 |
| chr13-65549078-65549579 | 8 | 0.995246625390252 |
| chr2-142193492-142193993 | 8 | 0.978934430083235 |
| chr2-46652086-46652587 | 3 | 0.996877828312262 |
| chr11-15924923-15925424 | 7 | 0.864134887928649 |
| chr14-35407715-35408216 | 5 | 0.982862857459958 |
| chr6-93430222-93430723 | 6 | 0.991206925481043 |
| chr4-76666691-76667192 | 4 | 0.815771097960451 |
| chr2-41024069-41024570 | 2 | 0.95638876009745 |
| chr3-49956639-49957140 | 3 | 0.996699239363713 |
| chr1-198730263-198730764 | 2 | 0.862019355408333 |
| chr4-14676497-14676998 | 6 | 0.988301399720597 |
| chr6-33768070-33768571 | 6 | 0.932708064826135 |
| chr1-173676981-173677482 | 6 | 0.991206925481043 |
| chr2-241536798-241537299 | 6 | 0.893144962560394 |
| chr9-111465599-111466100 | 6 | 0.974937724448335 |
| chr10-3493576-3494077 | 3 | 0.988449358738361 |
| chrX-142573384-142573885 | 3 | 0.99593553545041 |
| chr1-189341008-189341509 | 7 | 0.937289327580901 |
| chr6-54308476-54308977 | 8 | 0.938580007738151 |
| chr3-87534205-87534706 | 3 | 0.990217999476869 |
| chr15-16420203-16420704 | 8 | 0.831820543288292 |
| chr20-23903482-23903983 | 4 | 0.903042856577225 |
| chr14-83265922-83266423 | 7 | 0.900419006184618 |
| chr18-9708145-9708646 | 3 | 0.993058468791105 |
| chr13-88097311-88097812 | 7 | 0.877215852124701 |
| chr9-40455583-40456084 | 8 | 0.884398450824859 |
| chr6-81431563-81432064 | 6 | 0.991206925481043 |
| chr3-114210434-114210935 | 1 | 0.927237297748269 |
| chr14-33969639-33970140 | 3 | 0.971782444717419 |
| chr6-5762958-5763459 | 6 | 0.884684057367165 |
| chr17-68979102-68979603 | 6 | 0.988287235479087 |
| chr20-24837482-24837983 | 8 | 0.939973397156721 |
| chr20-18644205-18644706 | 8 | 0.977752009014015 |
| chr7-128677608-128678109 | 5 | 0.98280437340091 |
| chrX-105990401-105990902 | 4 | 0.97561785725183 |
| chr1-207421176-207421677 | 8 | 0.982934919734043 |
| chr4-181390762-181391263 | 6 | 0.991206925481043 |
| chr14-10329285-10329786 | 8 | 0.863961213562134 |
| chr1-204761635-204762136 | 2 | 0.931700386556795 |
| chr6-45654298-45654799 | 6 | 0.81858478079202 |

| | | |
|---|---|---|
| chr9-45814336-45814837 | 8 | 0.897029424259867 |
| chr1-11835670-11836171 | 8 | 0.988289317703882 |
| chr6-14442812-14443313 | 4 | 0.974198654702303 |
| chr7-133921764-133922265 | 4 | 0.975752411495426 |
| chr9-98240661-98241162 | 8 | 0.849180734230741 |
| chr1-214549434-214549935 | 3 | 0.949830674578038 |
| chr10-72208341-72208842 | 6 | 0.988069137719631 |
| chr5-123400474-123400975 | 6 | 0.977876940406819 |
| chr6-75846996-75847497 | 6 | 0.990606050091032 |
| chr18-75141434-75141935 | 3 | 0.990243316971838 |
| chr1-243029636-243030137 | 8 | 0.984282397501767 |
| chr17-51222277-51222778 | 8 | 0.850070132534163 |
| chr11-31349673-31350174 | 5 | 0.961670104585999 |
| chr15-47232896-47233397 | 6 | 0.860033592986087 |
| chr6-70390877-70391378 | 2 | 0.936815310452037 |
| chr14-65636728-65637229 | 6 | 0.987215701386629 |
| chr8-108854181-108854682 | 4 | 0.867919640307905 |
| chr9-111131657-111132158 | 5 | 0.928306879973605 |
| chrX-151396326-151396827 | 2 | 0.885198772569686 |
| chr8-3933068-3933569 | 6 | 0.991206925481043 |
| chr1-226323157-226323658 | 8 | 0.988452920740747 |
| chr4-126398443-126398944 | 8 | 0.903354196856615 |
| chr4-81458735-81459236 | 5 | 0.884563428739044 |
| chr1-201032524-201033025 | 5 | 0.864199471529909 |
| chr8-83930464-83930965 | 3 | 0.996641751427291 |
| chr3-102696830-102697331 | 2 | 0.841865542781358 |
| chr2-33788098-33788599 | 3 | 0.994865557495073 |
| chr5-117314317-117314818 | 4 | 0.942105954046484 |
| chr17-68964054-68964555 | 6 | 0.987501725718177 |
| chrX-50026217-50026718 | 6 | 0.897955013799213 |
| chr1-31391985-31392486 | 3 | 0.99593553545041 |
| chr8-63275485-63275986 | 7 | 0.894893566203855 |
| chr2-41955304-41955805 | 8 | 0.824473727991301 |
| chr3-120058269-120058770 | 6 | 0.956019651604816 |
| chr4-8312736-8313237 | 8 | 0.946825822357052 |
| chr6-119426713-119427214 | 5 | 0.948435385351873 |
| chr1-151718742-151719243 | 4 | 0.855431540567888 |
| chr5-165884914-165885415 | 3 | 0.904427977409668 |
| chr2-174466368-174466869 | 8 | 0.9822225797294 |
| chr20-9688611-9689112 | 7 | 0.85914475093658 |
| chr17-24648303-24648804 | 6 | 0.976798044920234 |
| chr15-32670449-32670950 | 8 | 0.925376776753475 |
| chr9-90320854-90321355 | 8 | 0.920861537973144 |

| | | |
|---|---|---|
| chr14-66374305-66374806 | 8 | 0.930811091992802 |
| chr16-69677473-69677974 | 4 | 0.973691695978432 |
| chr3-62086062-62086563 | 8 | 0.827548641931189 |
| chr6-125890388-125890889 | 3 | 0.994699243948489 |
| chrX-90385613-90386114 | 6 | 0.992988237256966 |
| chr17-21552752-21553253 | 8 | 0.955935753536018 |
| chr16-45723542-45724043 | 8 | 0.871709558069161 |
| chr3-95630319-95630820 | 3 | 0.99593553545041 |
| chr6-85219144-85219645 | 6 | 0.965225280811528 |
| chr13-103147139-103147640 | 8 | 0.993094823723113 |
| chr19-49624162-49624663 | 3 | 0.898266664143276 |
| chr9-105448263-105448764 | 7 | 0.8896017514682 |
| chr10-98151680-98152181 | 8 | 0.958746572201038 |
| chrX-28436655-28437156 | 5 | 0.962521654908939 |
| chr2-133439539-133440040 | 3 | 0.930343092192623 |
| chr10-21711934-21712435 | 3 | 0.954235873411964 |
| chr1-34857319-34857820 | 6 | 0.990685681726066 |
| chr9-64729735-64730236 | 2 | 0.900838061511161 |
| chr13-58628941-58629442 | 3 | 0.988942289547123 |
| chr5-97109215-97109716 | 6 | 0.996997047559388 |
| chr6-8262855-8263356 | 8 | 0.97593480501444 |
| chr5-101447805-101448306 | 5 | 0.969508606015979 |
| chr10-92586038-92586539 | 4 | 0.874544783656342 |
| chr3-122602593-122603094 | 1 | 0.909487523868063 |
| chr6-15215051-15215552 | 5 | 0.876276885070782 |
| chr2-185859935-185860436 | 8 | 0.903808237575805 |
| chr10-66232822-66233323 | 6 | 0.996053411394282 |
| chr20-49094508-49095009 | 1 | 0.930556598982208 |
| chr10-105078724-105079225 | 8 | 0.992916460687806 |
| chr4-175004901-175005402 | 3 | 0.900414430101722 |
| chr1-234999397-234999898 | 8 | 0.992778896632744 |
| chr4-43447860-43448361 | 8 | 0.944575785989204 |
| chr5-43111240-43111741 | 3 | 0.879286894069774 |
| chr14-7631554-7632055 | 2 | 0.930785688515973 |
| chr19-53284492-53284993 | 1 | 0.832175426441685 |
| chr20-5008236-5008737 | 8 | 0.825888103302099 |
| chr6-13344403-13344904 | 8 | 0.993493681128311 |
| chr5-79303280-79303781 | 2 | 0.95770064046367 |
| chr6-16636503-16637004 | 8 | 0.905168454399712 |
| chr7-11267932-11268433 | 8 | 0.876236118184519 |
| chr8-78430379-78430880 | 6 | 0.924014861106678 |
| chr1-154589406-154589907 | 8 | 0.981877277593851 |
| chr10-62127220-62127721 | 3 | 0.996461471626729 |

| | | |
|---|---|---|
| chr20-5777196-5777697 | 8 | 0.995542155587138 |
| chr3-54547449-54547950 | 2 | 0.850330344473098 |
| chr13-50559257-50559758 | 8 | 0.970672136055718 |
| chr4-88908706-88909207 | 3 | 0.891380488675827 |
| chr14-4323635-4324136 | 8 | 0.853942046879136 |
| chr7-30376354-30376855 | 4 | 0.909165714477036 |
| chr5-122265620-122266121 | 6 | 0.996535175817113 |
| chr5-122307354-122307855 | 6 | 0.996528630566277 |
| chr7-130982622-130983123 | 8 | 0.894210096284264 |
| chr16-53539039-53539540 | 3 | 0.994405638104216 |
| chr5-164921756-164922257 | 3 | 0.820429786790711 |
| chr2-150836199-150836700 | 8 | 0.993493681128311 |
| chr1-127565014-127565515 | 6 | 0.884183051415678 |
| chr15-28746963-28747464 | 5 | 0.964330623823044 |
| chr5-78242427-78242928 | 4 | 0.890303075139207 |
| chr6-51674071-51674572 | 1 | 0.937842673715779 |
| chr19-19933221-19933722 | 1 | 0.871534646188016 |
| chr9-112714493-112714994 | 3 | 0.99593553545041 |
| chr4-168379273-168379774 | 8 | 0.993493681128311 |
| chr14-104423133-104423634 | 8 | 0.987859415458081 |
| chr2-235215254-235215755 | 8 | 0.865826229713982 |
| chr3-134266388-134266889 | 3 | 0.994054065310948 |
| chr2-45593453-45593954 | 6 | 0.997359657684954 |
| chr3-151003760-151004261 | 8 | 0.815146510581264 |
| chr11-30363760-30364261 | 8 | 0.988668717454696 |
| chr4-41314806-41315307 | 4 | 0.888888539769172 |
| chr8-112811566-112812067 | 6 | 0.936572819723477 |
| chr13-102005742-102006243 | 3 | 0.99682055579757 |
| chr9-62219355-62219856 | 4 | 0.883672898354712 |
| chr1-89983510-89984011 | 2 | 0.920028315309666 |
| chr1-47484030-47484531 | 1 | 0.95038429797362 |
| chr4-49924821-49925322 | 3 | 0.981289918375805 |
| chr1-254210931-254211432 | 8 | 0.900955059679851 |
| chr1-173605580-173606081 | 3 | 0.99382446410171 |
| chr3-50431482-50431983 | 6 | 0.994343499604771 |
| chr10-6297814-6298315 | 5 | 0.956057279684501 |
| chr11-26247489-26247990 | 6 | 0.991206925481043 |
| chr6-111569453-111569954 | 8 | 0.947475854235619 |
| chr9-17296580-17297081 | 6 | 0.991206925481043 |
| chr19-26857721-26858222 | 8 | 0.931950130156053 |
| chr15-70161465-70161966 | 3 | 0.982940951099769 |
| chr13-96050664-96051165 | 8 | 0.990056419316456 |
| chr14-99183879-99184380 | 6 | 0.992903737563989 |

| | | |
|---|---|---|
| chr4-118078498-118078999 | 8 | 0.964502674430325 |
| chr15-7621243-7621744 | 5 | 0.970699324727767 |
| chr15-19543794-19544295 | 1 | 0.870143499692803 |
| chr8-47553573-47554074 | 2 | 0.834204363740247 |
| chr8-85846304-85846805 | 3 | 0.974631528029034 |
| chr9-76862778-76863279 | 4 | 0.891011916606651 |
| chr14-88458463-88458964 | 8 | 0.908044773282533 |
| chr4-126930554-126931055 | 6 | 0.990553712069381 |
| chr6-51456307-51456808 | 6 | 0.892070764989921 |
| chr18-73591563-73592064 | 8 | 0.869389843776848 |
| chr9-34805728-34806229 | 6 | 0.991206925481043 |
| chr2-94571007-94571508 | 3 | 0.980191959552838 |
| chr18-35287239-35287740 | 8 | 0.995244688456174 |
| chr1-188873149-188873650 | 8 | 0.981152366338519 |
| chr10-80001178-80001679 | 8 | 0.825713574685057 |
| chr4-142593037-142593538 | 8 | 0.983241772715186 |
| chr10-59341460-59341961 | 3 | 0.995877129208933 |
| chr16-50517792-50518293 | 6 | 0.98397790505587 |
| chr14-73937716-73938217 | 8 | 0.827145387368663 |
| chr20-53761349-53761850 | 6 | 0.994678606198107 |
| chr8-47501210-47501711 | 3 | 0.99593553545041 |
| chr8-28835844-28836345 | 8 | 0.863100614562599 |
| chr8-69148667-69149168 | 1 | 0.816433383743844 |
| chr6-123379008-123379509 | 2 | 0.971201498109347 |
| chr14-79325862-79326363 | 5 | 0.981694937043236 |
| chr5-68659796-68660297 | 8 | 0.877954859498387 |
| chr13-37885787-37886288 | 8 | 0.829253343640901 |
| chr10-77013663-77014164 | 3 | 0.996896496378318 |
| chr12-40682253-40682754 | 8 | 0.989156086685695 |
| chr20-11580135-11580636 | 2 | 0.979114963976027 |
| chr15-66660460-66660961 | 3 | 0.99593553545041 |
| chr10-90987633-90988134 | 5 | 0.942720452355066 |
| chr16-5898316-5898817 | 8 | 0.82780227947485 |
| chr4-8204772-8205273 | 6 | 0.995583125430695 |
| chr2-145614761-145615262 | 3 | 0.996730464719458 |
| chr16-17172506-17173007 | 8 | 0.834548675995402 |
| chr18-30002288-30002789 | 7 | 0.880450185609833 |
| chr10-48602594-48603095 | 8 | 0.944366301307029 |
| chr10-105968909-105969410 | 4 | 0.899969504240266 |
| chr7-116703006-116703507 | 3 | 0.997910956067315 |
| chr13-101662157-101662658 | 6 | 0.907129535973376 |
| chr3-45200146-45200647 | 5 | 0.89968308854111 |
| chr8-9408368-9408869 | 3 | 0.996730397514012 |

| | | |
|---|---|---|
| chr9-111732427-111732928 | 3 | 0.99628935760528 |
| chr10-94838041-94838542 | 6 | 0.974591588134847 |
| chr7-56504761-56505262 | 8 | 0.979324793043075 |
| chr11-9914518-9915019 | 5 | 0.95871705531043 |
| chr14-4203197-4203698 | 3 | 0.97094742910265 |
| chr1-224307229-224307730 | 1 | 0.840654446846879 |
| chr6-110443831-110444332 | 8 | 0.91372476112902 |
| chr7-65710925-65711426 | 3 | 0.995424318823653 |
| chr14-3640041-3640542 | 6 | 0.989983705911407 |
| chr8-4869327-4869828 | 8 | 0.895255456447633 |
| chr17-18058493-18058994 | 8 | 0.973207609330527 |
| chr8-68418344-68418845 | 4 | 0.925069598488787 |
| chr11-65439654-65440155 | 8 | 0.842698027502777 |
| chr6-40934300-40934801 | 4 | 0.874455968805738 |
| chr1-146494630-146495131 | 8 | 0.976986710106099 |
| chr1-41933901-41934402 | 6 | 0.985151708203016 |
| chr15-54978378-54978879 | 3 | 0.972485680913378 |
| chr13-94995033-94995534 | 4 | 0.940244718144293 |
| chr7-31881141-31881642 | 3 | 0.996897968208821 |
| chr13-38929005-38929506 | 8 | 0.838721620925697 |
| chr10-7613615-7614116 | 6 | 0.989967239529868 |
| chr2-122718610-122719111 | 8 | 0.816284440113856 |
| chr7-19062576-19063077 | 8 | 0.976643553434207 |
| chr9-2225443-2225944 | 5 | 0.964619261121256 |
| chr1-190041158-190041659 | 3 | 0.993650122125856 |
| chr18-67382392-67382893 | 8 | 0.836185071650059 |
| chr10-36792698-36793199 | 8 | 0.905713389324213 |
| chr5-7342459-7342960 | 3 | 0.990430608199649 |
| chr12-9520463-9520964 | 1 | 0.898332245482668 |
| chr3-80800346-80800847 | 6 | 0.983495857222109 |
| chr8-107317130-107317631 | 8 | 0.905034082440155 |
| chr2-170736067-170736568 | 5 | 0.989697533322907 |
| chr18-76540300-76540801 | 8 | 0.992080897288885 |
| chr14-37951653-37952154 | 3 | 0.962706612079999 |
| chr1-19217220-19217721 | 8 | 0.982815514427298 |
| chr13-22140475-22140976 | 3 | 0.969124460857403 |
| chr7-89934479-89934980 | 1 | 0.89824416580917 |
| chr11-12273229-12273730 | 6 | 0.993221599223871 |
| chr2-124055282-124055783 | 3 | 0.993711700784146 |
| chr5-85421705-85422206 | 6 | 0.993159480421512 |
| chr7-126140608-126141109 | 5 | 0.964283573895657 |
| chr19-11559817-11560318 | 4 | 0.884447338248911 |
| chr4-175719557-175720058 | 7 | 0.886248408918038 |

| | | |
|---|---|---|
| chr8-36944429-36944930 | 8 | 0.875462893760429 |
| chr16-52104319-52104820 | 1 | 0.903063234858441 |
| chr1-101566681-101567182 | 7 | 0.925069575536125 |
| chr10-29087033-29087534 | 5 | 0.984320620593261 |
| chr13-44275221-44275722 | 8 | 0.863498395597668 |
| chr1-47401813-47402314 | 6 | 0.994352621386118 |
| chr20-44276584-44277085 | 2 | 0.939701633412841 |
| chr18-60466963-60467464 | 5 | 0.884847007350239 |
| chr13-88952886-88953387 | 4 | 0.839963522564347 |
| chr16-43958181-43958682 | 8 | 0.993493681128311 |
| chr9-92097436-92097937 | 7 | 0.856661679929273 |
| chr19-8261712-8262213 | 3 | 0.996849966128339 |
| chr6-55595552-55596053 | 6 | 0.991206925481043 |
| chr14-26447410-26447911 | 8 | 0.91905817525639 |
| chr3-39055504-39056005 | 3 | 0.831194662014537 |
| chr1-62449092-62449593 | 3 | 0.99593553545041 |
| chr20-30942325-30942826 | 6 | 0.991206925481043 |
| chr17-84800390-84800891 | 3 | 0.903216758845245 |
| chr3-444682-445183 | 2 | 0.859789331365917 |
| chr10-54220465-54220966 | 5 | 0.861636105259238 |
| chr16-16935735-16936236 | 5 | 0.918052476770868 |
| chr3-131308054-131308555 | 2 | 0.90981693928606 |
| chr4-140620855-140621356 | 6 | 0.986831926648019 |
| chr1-100386426-100386927 | 3 | 0.920391582177834 |
| chr3-163628483-163628984 | 3 | 0.99593553545041 |
| chr4-80941419-80941920 | 8 | 0.873895372729484 |
| chr8-53111645-53112146 | 3 | 0.991924631064603 |
| chr2-157724339-157724840 | 3 | 0.99593553545041 |
| chr18-63312937-63313438 | 1 | 0.857831267885934 |
| chr2-239048497-239048998 | 6 | 0.992888217922468 |
| chr7-105973181-105973682 | 3 | 0.983181746920879 |
| chr2-119496621-119497122 | 4 | 0.966938915865596 |
| chr1-89418311-89418812 | 8 | 0.969715477547038 |
| chr6-31329234-31329735 | 8 | 0.827131509065474 |
| chr6-37276232-37276733 | 6 | 0.992944914084811 |
| chr17-6658279-6658780 | 5 | 0.854667038904045 |
| chr17-23698607-23699108 | 5 | 0.989619300184635 |
| chr18-53917742-53918243 | 6 | 0.992453804202721 |
| chr1-194480020-194480521 | 2 | 0.872599532221531 |
| chr11-81807886-81808387 | 5 | 0.933279675290579 |
| chr10-82953092-82953593 | 8 | 0.993493681128311 |
| chr1-163123665-163124166 | 3 | 0.988982656865276 |
| chr3-166537492-166537993 | 8 | 0.957240234305917 |

| | | |
|---|---|---|
| chr12-39771843-39772344 | 5 | 0.993414961138274 |
| chr11-48734123-48734624 | 8 | 0.981732814148383 |
| chr3-143905114-143905615 | 4 | 0.948279583233748 |
| chr4-167123777-167124278 | 8 | 0.858717638368621 |
| chr2-240249983-240250484 | 6 | 0.929755699341977 |
| chr5-72148295-72148796 | 2 | 0.891494642762172 |
| chr18-50523354-50523855 | 5 | 0.976317745843577 |
| chr7-46538627-46539128 | 8 | 0.973239907544726 |
| chr10-20469843-20470344 | 6 | 0.990376170190373 |
| chr15-25656258-25656759 | 7 | 0.952042532027573 |
| chrX-65223829-65224330 | 5 | 0.970199474957913 |
| chr2-160360110-160360611 | 3 | 0.996410648817179 |
| chr3-80885459-80885960 | 3 | 0.995793090618271 |
| chr13-44846178-44846679 | 5 | 0.928379660867808 |
| chrX-58087373-58087874 | 6 | 0.979798849302838 |
| chr3-101388996-101389497 | 3 | 0.954936192830502 |
| chr11-51729900-51730401 | 6 | 0.991407482837903 |
| chr13-69810156-69810657 | 3 | 0.99593553545041 |
| chr14-67380896-67381397 | 4 | 0.955737203807356 |
| chr5-166471014-166471515 | 8 | 0.976738786451707 |
| chr18-12499783-12500284 | 3 | 0.996088552540347 |
| chr17-55519302-55519803 | 8 | 0.984297501808547 |
| chr6-27455663-27456164 | 8 | 0.951263589157878 |
| chr8-62976177-62976678 | 8 | 0.882526416861265 |
| chr8-82822354-82822855 | 2 | 0.947209711051289 |
| chr3-104753800-104754301 | 1 | 0.904342556137641 |
| chr8-37383440-37383941 | 8 | 0.965180115109781 |
| chr5-99824703-99825204 | 5 | 0.861245778399394 |
| chr6-61000209-61000710 | 3 | 0.996594090310781 |
| chr1-32925404-32925905 | 8 | 0.864174984989573 |
| chr15-5671792-5672293 | 8 | 0.92504301833475 |
| chr13-84463466-84463967 | 5 | 0.976863577703874 |
| chr19-37655758-37656259 | 8 | 0.85186985239345 |
| chr1-170119381-170119882 | 6 | 0.986579452308807 |
| chr19-50869296-50869797 | 8 | 0.929027342368892 |
| chr2-210963441-210963942 | 4 | 0.832802382718563 |
| chr7-64326051-64326552 | 6 | 0.991206925481043 |
| chr5-145275132-145275633 | 4 | 0.924157142475141 |
| chr8-78138705-78139206 | 8 | 0.98609461996581 |
| chr10-21091474-21091975 | 6 | 0.987294481475655 |
| chr17-51230684-51231185 | 8 | 0.917894145511538 |
| chr1-45499236-45499737 | 3 | 0.994059600462426 |
| chr1-182418139-182418640 | 8 | 0.907060544825911 |

| | | |
|---|---|---|
| chr8-52339851-52340352 | 2 | 0.90309712402696 |
| chr5-83741925-83742426 | 5 | 0.983919978840557 |
| chr8-70489042-70489543 | 7 | 0.862716614245986 |
| chr7-14450539-14451040 | 8 | 0.819878124418206 |
| chr17-53101933-53102434 | 5 | 0.857828621785478 |
| chr4-22018189-22018690 | 5 | 0.907228750290527 |
| chr7-98969053-98969554 | 5 | 0.990022129809314 |
| chr12-2799855-2800356 | 8 | 0.819597762789441 |
| chr19-29600455-29600956 | 8 | 0.972292488662755 |
| chr3-165696401-165696902 | 5 | 0.822424634050116 |
| chr14-44714949-44715450 | 1 | 0.901009242978542 |
| chr5-139231701-139232202 | 5 | 0.985154991200442 |
| chr9-91412991-91413492 | 6 | 0.963965519067461 |
| chr3-158333563-158334064 | 8 | 0.846970801326731 |
| chr7-416301-416802 | 3 | 0.99193081355542 |
| chr1-131888166-131888667 | 5 | 0.919891525156025 |
| chr6-13462328-13462829 | 6 | 0.992832839064043 |
| chr1-193555340-193555841 | 3 | 0.983782114390327 |
| chr6-24230223-24230724 | 2 | 0.944340276087842 |
| chr2-39737193-39737694 | 8 | 0.916254376619225 |
| chr1-86282886-86283387 | 5 | 0.979288839848978 |
| chr15-26796816-26797317 | 8 | 0.847243992261102 |
| chr6-52240673-52241174 | 8 | 0.946264298212829 |
| chr3-115472618-115473119 | 8 | 0.942418578989366 |
| chr1-49385990-49386491 | 8 | 0.852555445728277 |
| chr3-57252567-57253068 | 3 | 0.92094135915735 |
| chr1-164228550-164229051 | 6 | 0.990679521118535 |
| chr2-103750484-103750985 | 3 | 0.99593553545041 |
| chr2-44703354-44703855 | 3 | 0.946312406133147 |
| chr4-32283078-32283579 | 8 | 0.976292132558626 |
| chr1-218801260-218801761 | 6 | 0.973267980230938 |
| chr17-27908820-27909321 | 8 | 0.89776622416177 |
| chr9-71039593-71040094 | 5 | 0.983919978840557 |
| chr13-49820550-49821051 | 3 | 0.996779069358105 |
| chr3-158313872-158314373 | 5 | 0.990397620474143 |
| chr3-115562187-115562688 | 5 | 0.936977993527115 |
| chr6-64103279-64103780 | 3 | 0.99655253922721 |
| chr14-31583829-31584330 | 5 | 0.873512499041062 |
| chr6-13164499-13165000 | 6 | 0.993212168551819 |
| chr6-42945962-42946463 | 7 | 0.929899161913742 |
| chr15-52617081-52617582 | 8 | 0.929001168998743 |
| chr7-57403944-57404445 | 8 | 0.931864902522124 |
| chr13-46642047-46642548 | 6 | 0.989176412259215 |

| | | |
|---|---|---|
| chr16-67069945-67070446 | 3 | 0.871191885950434 |
| chr13-100867183-100867684 | 8 | 0.891961707724116 |
| chr17-51754170-51754671 | 3 | 0.970568525241885 |
| chr2-114855295-114855796 | 8 | 0.8914575892951 |
| chr6-46278494-46278995 | 8 | 0.989453946610411 |
| chr8-60521937-60522438 | 8 | 0.987540824617156 |
| chr20-26716313-26716814 | 8 | 0.979898576775918 |
| chr4-176180288-176180789 | 3 | 0.922413331739546 |
| chr7-74226890-74227391 | 3 | 0.847527532929341 |
| chr18-70922008-70922509 | 7 | 0.901431913055004 |
| chr10-102654083-102654584 | 5 | 0.902043496506498 |
| chr4-118719441-118719942 | 8 | 0.994779947339339 |
| chr2-184647585-184648086 | 2 | 0.86783327106924 |
| chr3-168954668-168955169 | 3 | 0.994326014241192 |
| chr8-104531173-104531674 | 8 | 0.869943456163691 |
| chr14-97907653-97908154 | 2 | 0.826801212215924 |
| chr9-60659762-60660263 | 8 | 0.984840470472219 |
| chr18-51454838-51455339 | 4 | 0.919857872419529 |
| chr16-9403767-9404268 | 8 | 0.891755437391847 |
| chr17-83064387-83064888 | 8 | 0.988889956202136 |
| chr19-11805300-11805801 | 7 | 0.820812101974337 |
| chr4-60360190-60360691 | 3 | 0.935860588663072 |
| chr9-57275769-57276270 | 8 | 0.882569475209946 |
| chr1-22823667-22824168 | 8 | 0.994690214503667 |
| chr7-99746989-99747490 | 6 | 0.977792173161147 |
| chr1-127586779-127587280 | 6 | 0.851718121735444 |
| chr5-117825950-117826451 | 8 | 0.837694211448347 |
| chr10-23040916-23041417 | 6 | 0.991206925481043 |
| chr6-56199302-56199803 | 2 | 0.891565537709273 |
| chr3-142323001-142323502 | 3 | 0.989761718689481 |
| chr4-90292149-90292650 | 3 | 0.989749821455503 |
| chr18-21473912-21474413 | 3 | 0.996894471679704 |
| chr5-49286696-49287197 | 5 | 0.970087344464102 |
| chr5-24830326-24830827 | 5 | 0.927052645781946 |
| chr18-69250790-69251291 | 7 | 0.875733264363691 |
| chr7-63075328-63075829 | 3 | 0.996697403150195 |
| chr14-84122966-84123467 | 6 | 0.91648430760712 |
| chr6-45744758-45745259 | 2 | 0.904892091420879 |
| chr1-254504627-254505128 | 3 | 0.996515915489637 |
| chr1-118835046-118835547 | 5 | 0.820345456992882 |
| chr4-90239340-90239841 | 7 | 0.943236008976239 |
| chr12-12298397-12298898 | 6 | 0.931222975911431 |
| chr1-48058214-48058715 | 4 | 0.895017281887141 |

| | | |
|---|---|---|
| chr7-120297213-120297714 | 8 | 0.882487744293956 |
| chr14-30714349-30714850 | 5 | 0.890996738349114 |
| chr18-4689290-4689791 | 6 | 0.993187436927845 |
| chr7-129423312-129423813 | 2 | 0.970109921368683 |
| chr7-119095348-119095849 | 8 | 0.995240061793169 |
| chr2-232396654-232397155 | 6 | 0.960766501101573 |
| chr7-112546267-112546768 | 8 | 0.956959551713027 |
| chr4-83739553-83740054 | 3 | 0.994335031227708 |
| chr8-19764613-19765114 | 5 | 0.98858796030953 |
| chr18-67638485-67638986 | 8 | 0.995151522153681 |
| chr2-182925182-182925683 | 8 | 0.993067452972919 |
| chr2-197879214-197879715 | 5 | 0.896995388158151 |
| chr3-69632887-69633388 | 8 | 0.89193136634518 |
| chr3-159620514-159621015 | 6 | 0.983239633183659 |
| chr9-23099038-23099539 | 3 | 0.99593553545041 |
| chr7-103654723-103655224 | 5 | 0.934316803008063 |
| chr4-134024883-134025384 | 6 | 0.99034244942314 |
| chr3-1252305-1252806 | 3 | 0.976275987810444 |
| chr15-1699423-1699924 | 8 | 0.954956359671194 |
| chr4-58110812-58111313 | 7 | 0.852962389259859 |
| chr3-145999253-145999754 | 7 | 0.883430802830721 |
| chr6-68449569-68450070 | 3 | 0.988479317384763 |
| chr3-145696817-145697318 | 5 | 0.993482622838122 |
| chr19-34760405-34760906 | 8 | 0.891488602692198 |
| chr18-45749462-45749963 | 8 | 0.99518320586467 |
| chr14-67741500-67742001 | 8 | 0.993493681128311 |
| chr9-18092563-18093064 | 3 | 0.861007407488055 |
| chr10-23335778-23336279 | 3 | 0.994771753495956 |
| chr2-206545937-206546438 | 3 | 0.945512453185334 |
| chr2-238189438-238189939 | 8 | 0.98671100585591 |
| chr19-53725040-53725541 | 6 | 0.823793050977603 |
| chr2-135145766-135146267 | 8 | 0.975356821139186 |
| chr18-30743046-30743547 | 4 | 0.816627250493826 |
| chr18-23553570-23554071 | 3 | 0.86872026802863 |
| chr6-48704059-48704560 | 6 | 0.982810315314175 |
| chr12-16988437-16988938 | 8 | 0.874653849489231 |
| chr15-92653287-92653788 | 5 | 0.844652664242462 |
| chr16-69202896-69203397 | 3 | 0.912532618432614 |
| chr14-79448103-79448604 | 7 | 0.839742493549037 |
| chr3-140246527-140247028 | 1 | 0.928107188391251 |
| chr3-91453083-91453584 | 8 | 0.83173228831274 |
| chr1-175225553-175226054 | 6 | 0.920263369153899 |
| chr14-4863132-4863633 | 3 | 0.997647272197524 |

| | | |
|---|---|---|
| chr6-55675391-55675892 | 7 | 0.917242124351984 |
| chrX-115363373-115363874 | 1 | 0.823775158695192 |
| chr13-74325408-74325909 | 6 | 0.963421088426723 |
| chr4-154670271-154670772 | 8 | 0.974148521032846 |
| chr9-84869229-84869730 | 4 | 0.974276689002774 |
| chr6-38945676-38946177 | 8 | 0.897853754738056 |
| chr1-222324718-222325219 | 6 | 0.885891413137449 |
| chr15-100773381-100773882 | 6 | 0.993226489483441 |
| chr3-134534335-134534836 | 3 | 0.995595140575314 |
| chr17-62125310-62125811 | 4 | 0.874514758741126 |
| chr2-172288828-172289329 | 8 | 0.822659330739162 |
| chr9-62971288-62971789 | 8 | 0.844006707823997 |
| chr5-2850646-2851147 | 3 | 0.987020639861689 |
| chr6-15149122-15149623 | 2 | 0.840146851576324 |
| chr6-140003458-140003959 | 5 | 0.819218084834641 |
| chr18-66419027-66419528 | 6 | 0.995544177003485 |
| chr1-7625746-7626247 | 5 | 0.972447134196977 |
| chr2-112475110-112475611 | 4 | 0.892675074409787 |
| chr1-165117208-165117709 | 3 | 0.996622864021207 |
| chrX-17254764-17255265 | 3 | 0.918658677755925 |
| chr7-93468374-93468875 | 3 | 0.827816641865597 |
| chr12-4874681-4875182 | 8 | 0.980829189413113 |
| chr4-61371962-61372463 | 8 | 0.992712418312491 |
| chr8-87035385-87035886 | 2 | 0.86034077357897 |
| chr8-59209054-59209555 | 8 | 0.959248491454901 |
| chr1-15735003-15735504 | 4 | 0.881496934594672 |
| chr15-44849010-44849511 | 5 | 0.835026523904839 |
| chr5-38851788-38852289 | 3 | 0.996797337086159 |
| chrX-12076357-12076858 | 3 | 0.99593553545041 |
| chr8-38315038-38315539 | 3 | 0.996880015486123 |
| chr8-40946202-40946703 | 6 | 0.991206925481043 |
| chr9-95107409-95107910 | 8 | 0.855698949206757 |
| chr2-137514283-137514784 | 8 | 0.908674001816662 |
| chr7-134707196-134707697 | 4 | 0.873989898756188 |
| chr8-30226372-30226873 | 3 | 0.989829335403468 |
| chr9-7416552-7417053 | 2 | 0.891594870799915 |
| chr20-12157400-12157901 | 6 | 0.833735131061582 |
| chr1-174598780-174599281 | 8 | 0.962594045862265 |
| chr4-165560301-165560802 | 6 | 0.878182932366689 |
| chr16-60811894-60812395 | 8 | 0.993493681128311 |
| chrX-14718301-14718802 | 8 | 0.898337638796999 |
| chr9-32030589-32031090 | 3 | 0.99593553545041 |
| chr8-83620814-83621315 | 8 | 0.975375803897654 |

| | | |
|---|---|---|
| chr2-185778875-185779376 | 8 | 0.99466478533934 |
| chr7-100534408-100534909 | 2 | 0.946704244976497 |
| chr10-12964805-12965306 | 3 | 0.947811367416139 |
| chr16-37651106-37651607 | 6 | 0.992127698027661 |
| chr1-217734967-217735468 | 3 | 0.99682536331862 |
| chr6-140261799-140262300 | 6 | 0.991206925481043 |
| chrX-117221149-117221650 | 5 | 0.992860122309037 |
| chrX-125795132-125795633 | 5 | 0.949624920614036 |
| chr14-94192404-94192905 | 1 | 0.898027884552763 |
| chr1-3127927-3128428 | 6 | 0.993034043813778 |
| chr20-26641340-26641841 | 5 | 0.989258827655682 |
| chr16-33952285-33952786 | 2 | 0.954816528764936 |
| chr14-19229663-19230164 | 8 | 0.837108865190459 |
| chr5-131004019-131004520 | 5 | 0.932621731261367 |
| chr1-132891921-132892422 | 8 | 0.862426351595828 |
| chr18-62263454-62263955 | 6 | 0.992563133362902 |
| chr2-112611907-112612408 | 4 | 0.875272299370816 |
| chr3-7921279-7921780 | 5 | 0.816362818121347 |
| chr10-86615540-86616041 | 3 | 0.930292806999697 |
| chr5-108882716-108883217 | 3 | 0.996743955096491 |
| chr6-118057221-118057722 | 5 | 0.970935445121998 |
| chr2-97757427-97757928 | 8 | 0.988938396082308 |
| chr14-41685769-41686270 | 2 | 0.873159478581326 |
| chr1-12432041-12432542 | 3 | 0.996701323010931 |
| chr9-94500774-94501275 | 6 | 0.985020967835858 |
| chr8-105434021-105434522 | 8 | 0.981465986974216 |
| chr7-122675597-122676098 | 8 | 0.898229822780873 |
| chr3-94488831-94489332 | 1 | 0.880807989353786 |
| chr15-63406200-63406701 | 5 | 0.987205218474872 |
| chr4-62269795-62270296 | 5 | 0.856722070501011 |
| chr3-21585165-21585666 | 6 | 0.963488770445976 |
| chr7-27309324-27309825 | 5 | 0.906748914154767 |
| chr11-74867069-74867570 | 8 | 0.825785518413204 |
| chr1-227451177-227451678 | 2 | 0.952900945684685 |
| chr9-70326039-70326540 | 3 | 0.988893624891112 |
| chr9-26071016-26071517 | 8 | 0.821455463372308 |
| chr3-83595892-83596393 | 3 | 0.99593553545041 |
| chr8-107134816-107135317 | 8 | 0.959909352204833 |
| chr1-83948811-83949312 | 8 | 0.814783131017205 |
| chr1-214123107-214123608 | 2 | 0.90655213851175 |
| chr19-20966820-20967321 | 3 | 0.996704915349598 |
| chr14-53438009-53438510 | 6 | 0.939152377424073 |
| chr6-21630148-21630649 | 1 | 0.878344132505239 |

| | | |
|---|---|---|
| chr14-41406945-41407446 | 2 | 0.85180423134877 |
| chr1-179926504-179927005 | 5 | 0.841722249828423 |
| chr3-128510467-128510968 | 8 | 0.844181 |
| chr3-123393644-123394145 | 6 | 0.985030952947783 |
| chr3-38302467-38302968 | 3 | 0.983477877316475 |
| chr6-52286338-52286839 | 8 | 0.974297008376509 |
| chr1-138853197-138853698 | 3 | 0.996558017959596 |
| chr20-25295190-25295691 | 6 | 0.991206925481043 |
| chr11-76603873-76604374 | 4 | 0.856347047150628 |
| chr17-13629477-13629978 | 3 | 0.996642688904772 |
| chr2-45706989-45707490 | 5 | 0.952445568502526 |
| chr1-146502967-146503468 | 2 | 0.876682042677253 |
| chr13-56311842-56312343 | 3 | 0.99593553545041 |
| chr7-52096495-52096996 | 5 | 0.985647999762904 |
| chr1-133797522-133798023 | 7 | 0.847509846098016 |
| chr10-15893075-15893576 | 8 | 0.985655568415206 |
| chr9-111013450-111013951 | 3 | 0.996744306092351 |
| chr6-38652487-38652988 | 8 | 0.921549465656204 |
| chr13-72087130-72087631 | 1 | 0.896442029556317 |
| chr17-7912681-7913182 | 3 | 0.958170421145251 |
| chr10-4376658-4377159 | 3 | 0.843750192358292 |
| chr5-96069966-96070467 | 8 | 0.958746965155389 |
| chr16-75147400-75147901 | 8 | 0.993493681128311 |
| chr6-125050665-125051166 | 4 | 0.845324810048307 |
| chr4-91070961-91071462 | 3 | 0.99593553545041 |
| chr8-80474256-80474757 | 3 | 0.996895109213653 |
| chr15-39757705-39758206 | 8 | 0.834219815601882 |
| chr18-16498439-16498940 | 3 | 0.989615629648154 |
| chr2-218369671-218370172 | 3 | 0.978645666300331 |
| chr3-55782206-55782707 | 3 | 0.996896883020966 |
| chr6-11056549-11057050 | 3 | 0.991226205098321 |
| chr14-34085897-34086398 | 4 | 0.960842214975684 |
| chr3-92880845-92881346 | 4 | 0.902140695168812 |
| chr3-50228638-50229139 | 6 | 0.991164919111583 |
| chr1-146287800-146288301 | 3 | 0.991553540115301 |
| chr12-37632794-37633295 | 6 | 0.933217366306083 |
| chr15-12562747-12563248 | 6 | 0.992788227893833 |
| chr1-47523753-47524254 | 3 | 0.992668652338774 |
| chr10-40669756-40670257 | 8 | 0.924703394212338 |
| chr6-127653230-127653731 | 8 | 0.995180102467765 |
| chr5-150991651-150992152 | 8 | 0.877893621505771 |
| chr2-91051637-91052138 | 8 | 0.972998538305048 |
| chr12-23021335-23021836 | 8 | 0.981452169810274 |

| | | |
|---|---|---|
| chrX-10286585-10287086 | 3 | 0.993733501899129 |
| chr3-167881325-167881826 | 6 | 0.977600357011101 |
| chr1-245466643-245467144 | 8 | 0.816993163114302 |
| chr6-123524018-123524519 | 3 | 0.996303649600303 |
| chr3-25426739-25427240 | 3 | 0.997467960466652 |
| chr8-1903992-1904493 | 4 | 0.821345380340532 |
| chr2-171253125-171253626 | 8 | 0.881845789253452 |
| chr6-36296798-36297299 | 3 | 0.962478926689549 |
| chr8-15492569-15493070 | 6 | 0.882521472920987 |
| chr13-70939407-70939908 | 3 | 0.99593553545041 |
| chr16-46440412-46440913 | 3 | 0.846978659328187 |
| chr2-191723372-191723873 | 7 | 0.823027078794154 |
| chr9-100053209-100053710 | 3 | 0.910006935074574 |
| chr2-107581516-107582017 | 3 | 0.970090145997043 |
| chr15-12699158-12699659 | 7 | 0.941673959918597 |
| chr8-62589093-62589594 | 8 | 0.932063508479936 |
| chr8-60263778-60264279 | 8 | 0.927573982310419 |
| chr10-81541702-81542203 | 8 | 0.860040353172868 |
| chrX-27931117-27931618 | 8 | 0.985092047826672 |
| chr7-58023240-58023741 | 2 | 0.897003749680316 |
| chr14-57889100-57889601 | 3 | 0.995918694716553 |
| chr18-44211452-44211953 | 6 | 0.991206925481043 |
| chr12-45769080-45769581 | 6 | 0.991206925481043 |
| chr16-75250168-75250669 | 8 | 0.949296370195726 |
| chr4-39622581-39623082 | 3 | 0.994016972856281 |
| chr20-30872754-30873255 | 6 | 0.994036754344662 |
| chr17-64797271-64797772 | 5 | 0.820381439451914 |
| chr7-38754012-38754513 | 3 | 0.997896144213226 |
| chr4-177504322-177504823 | 6 | 0.898962823348754 |
| chr12-43545045-43545546 | 1 | 0.948736096472324 |
| chr5-83992545-83993046 | 8 | 0.897302135348635 |
| chr7-44064183-44064684 | 6 | 0.991206925481043 |
| chr5-158805240-158805741 | 8 | 0.990871095072864 |
| chr13-63799843-63800344 | 1 | 0.867736381153422 |
| chr6-111653305-111653806 | 3 | 0.99593553545041 |
| chr1-5058851-5059352 | 2 | 0.830353263307202 |
| chr17-32422834-32423335 | 7 | 0.872418867237756 |
| chr10-42853654-42854155 | 3 | 0.993401348332182 |
| chr1-34302707-34303208 | 5 | 0.99166492979399 |
| chr6-53453463-53453964 | 3 | 0.995272287113877 |
| chr3-14631769-14632270 | 8 | 0.917585943810195 |
| chr6-115905586-115906087 | 4 | 0.925350746867526 |
| chr8-63374537-63375038 | 8 | 0.941600282875801 |

| | | |
|---|---|---|
| chr4-65096397-65096898 | 8 | 0.984658984171835 |
| chr3-11789221-11789722 | 6 | 0.990934130709948 |
| chr16-71816645-71817146 | 3 | 0.99593553545041 |
| chr16-566885-567386 | 4 | 0.872664715153613 |
| chr11-27116226-27116727 | 8 | 0.995034981941983 |
| chrX-56570902-56571403 | 3 | 0.985949336546827 |
| chr7-95013279-95013780 | 5 | 0.98895581994682 |
| chr10-19112047-19112548 | 3 | 0.955010179675178 |
| chr8-76712768-76713269 | 5 | 0.860840151608423 |
| chr6-137462884-137463385 | 2 | 0.836967682058954 |
| chr9-64998468-64998969 | 7 | 0.927229347627833 |
| chr7-24010799-24011300 | 5 | 0.928138542894872 |
| chr9-104338239-104338740 | 1 | 0.85425550213927 |
| chr1-243832934-243833435 | 8 | 0.923768749949054 |
| chr3-141372339-141372840 | 3 | 0.886159846974737 |
| chr17-40377608-40378109 | 4 | 0.967970546988431 |
| chr9-7334842-7335343 | 6 | 0.954840465821261 |
| chr10-78435675-78436176 | 8 | 0.993493681128311 |
| chr8-113510173-113510674 | 3 | 0.997776331861542 |
| chr2-57808599-57809100 | 5 | 0.983919978840557 |
| chr14-46500286-46500787 | 3 | 0.995595803869061 |
| chr1-233201858-233202359 | 6 | 0.930052468166583 |
| chr7-62480794-62481295 | 8 | 0.993493681128311 |
| chr1-178621902-178622403 | 3 | 0.843454258894892 |
| chr18-23930537-23931038 | 7 | 0.873724669845422 |
| chr16-50290158-50290659 | 3 | 0.996771925094162 |
| chr18-61389334-61389835 | 3 | 0.988548070828516 |
| chr13-1115578-1116079 | 8 | 0.934665721968691 |
| chr19-9545122-9545623 | 6 | 0.923801498396584 |
| chr18-81889911-81890412 | 8 | 0.899016764599203 |
| chr2-136381311-136381812 | 5 | 0.825108461172108 |
| chr16-49178287-49178788 | 4 | 0.875613093199463 |
| chr13-33195986-33196487 | 3 | 0.986577170145212 |
| chr5-148646631-148647132 | 5 | 0.984704281387296 |
| chr3-166974686-166975187 | 2 | 0.972572349575401 |
| chr2-113960384-113960885 | 3 | 0.995544466061681 |
| chr19-17771592-17772093 | 3 | 0.995694858250708 |
| chr19-21938810-21939311 | 2 | 0.830269089128911 |
| chr19-12803321-12803822 | 3 | 0.994572110619253 |
| chr5-47112305-47112806 | 8 | 0.880341979836906 |
| chr2-42184200-42184701 | 3 | 0.994645654966188 |
| chr9-84578973-84579474 | 2 | 0.953181722288917 |
| chr14-9432266-9432767 | 3 | 0.999228947704706 |

| | | |
|---|---|---|
| chr5-109827525-109828026 | 8 | 0.835030967125015 |
| chr19-56291200-56291701 | 6 | 0.98699381263821 |
| chr1-245576041-245576542 | 3 | 0.999071111019307 |
| chr5-153409855-153410356 | 4 | 0.915945675761851 |
| chr2-204136823-204137324 | 8 | 0.841151877229491 |
| chr8-69055362-69055863 | 3 | 0.996524002687949 |
| chr1-97279645-97280146 | 8 | 0.883431035676631 |
| chr8-53792367-53792868 | 1 | 0.927574261785646 |
| chr14-90759220-90759721 | 6 | 0.991206925481043 |
| chr5-133220668-133221169 | 4 | 0.968605338294872 |
| chr5-109653339-109653840 | 1 | 0.842267310891249 |
| chr16-61266081-61266582 | 3 | 0.995533500693784 |
| chr11-67463058-67463559 | 8 | 0.949077483051189 |
| chr6-114038605-114039106 | 3 | 0.996710063184718 |
| chr4-134089228-134089729 | 4 | 0.971087976644435 |
| chr9-109608700-109609201 | 8 | 0.867160089431543 |
| chr15-12081116-12081617 | 3 | 0.954389043751392 |
| chr19-42886649-42887150 | 4 | 0.950357025679519 |
| chr12-13722712-13723213 | 6 | 0.993141580142726 |
| chr11-3591757-3592258 | 4 | 0.926884151323152 |
| chr16-44403610-44404111 | 8 | 0.971440119198921 |
| chr17-14444216-14444717 | 2 | 0.937743213757169 |
| chr1-14786049-14786550 | 6 | 0.991858638122673 |
| chr2-114241543-114242044 | 6 | 0.972105884835964 |
| chr10-74692830-74693331 | 3 | 0.952321728413437 |
| chr6-27471241-27471742 | 2 | 0.817130499395741 |
| chr4-180246702-180247203 | 7 | 0.935335144602817 |
| chr2-25083232-25083733 | 6 | 0.817601964931082 |
| chr15-27825209-27825710 | 4 | 0.883131613906332 |
| chr2-21569716-21570217 | 6 | 0.95979022400667 |
| chr2-18611713-18612214 | 6 | 0.885956922989879 |
| chr15-36983786-36984287 | 6 | 0.880825073835339 |
| chr1-11682685-11683186 | 3 | 0.994266916363937 |
| chr6-4864277-4864778 | 8 | 0.939663923666922 |
| chr8-33361793-33362294 | 3 | 0.897139825943528 |
| chr6-5649188-5649689 | 6 | 0.991206925481043 |
| chr10-102902220-102902721 | 3 | 0.888203333753407 |
| chr5-143387181-143387682 | 4 | 0.882723878308728 |
| chr7-119893007-119893508 | 5 | 0.98345467632347 |
| chr2-76352053-76352554 | 5 | 0.983919978840557 |
| chr1-199501640-199502141 | 5 | 0.983919978840557 |
| chr2-34343601-34344102 | 8 | 0.897027134689613 |
| chr7-133465408-133465909 | 8 | 0.905163190620606 |

| | | |
|---|---|---|
| chr1-120255694-120256195 | 5 | 0.990763858463092 |
| chr3-148208140-148208641 | 3 | 0.99593553545041 |
| chr9-58779294-58779795 | 2 | 0.941529910437192 |
| chr2-12386211-12386712 | 3 | 0.995172866945922 |
| chr8-113909002-113909503 | 5 | 0.970147882545171 |
| chr6-140530828-140531329 | 3 | 0.993093014745885 |
| chr14-9696072-9696573 | 8 | 0.918378548813999 |
| chr7-61805680-61806181 | 3 | 0.989744108761911 |
| chr14-77368642-77369143 | 3 | 0.99593553545041 |
| chr7-68653389-68653890 | 3 | 0.996777445766287 |
| chr5-121766653-121767154 | 3 | 0.855202924287031 |
| chr8-78433647-78434148 | 8 | 0.990357688288325 |
| chr17-10475919-10476420 | 8 | 0.890844903037529 |
| chr2-186910586-186911087 | 8 | 0.98851207150841 |
| chr10-103502441-103502942 | 3 | 0.89426838773432 |
| chr6-114870257-114870758 | 8 | 0.87783716907871 |
| chr17-26247091-26247592 | 3 | 0.991766993245642 |
| chr10-30588447-30588948 | 8 | 0.947540790884072 |
| chr1-129379819-129380320 | 7 | 0.847666180108917 |
| chr14-82990309-82990810 | 1 | 0.93851587522281 |
| chr7-124056269-124056770 | 3 | 0.988425528898609 |
| chr1-222275810-222276311 | 5 | 0.98333709086421 |
| chr7-98091287-98091788 | 3 | 0.99593553545041 |
| chr2-155173265-155173766 | 3 | 0.99593553545041 |
| chr1-106800264-106800765 | 3 | 0.843056938603419 |
| chr8-10262795-10263296 | 8 | 0.94873241854959 |
| chr2-121250624-121251125 | 5 | 0.858837502315655 |
| chr8-84969363-84969864 | 4 | 0.941195121738968 |
| chr2-197295633-197296134 | 8 | 0.955890972845255 |
| chr5-112346511-112347012 | 8 | 0.952043495330088 |
| chr11-70647797-70648298 | 5 | 0.920643762028004 |
| chr6-126350532-126351033 | 3 | 0.97111567547504 |
| chr15-78074902-78075403 | 7 | 0.912055385900959 |
| chr12-26923256-26923757 | 5 | 0.932096433558957 |
| chr4-123431089-123431590 | 5 | 0.970886652589135 |
| chr1-248534480-248534981 | 5 | 0.84219562350291 |
| chr11-58329953-58330454 | 6 | 0.935696895037283 |
| chr15-97356077-97356578 | 3 | 0.996979040468017 |
| chr4-7717016-7717517 | 3 | 0.998992451104953 |
| chrX-72760570-72761071 | 8 | 0.834438077267551 |
| chr16-22309956-22310457 | 3 | 0.971201593089019 |
| chr8-122128125-122128626 | 3 | 0.996893376075211 |
| chr6-11663953-11664454 | 8 | 0.992239422831528 |

| | | |
|---|---|---|
| chr2-243139546-243140047 | 3 | 0.912392302109713 |
| chr5-122377564-122378065 | 8 | 0.884509517076317 |
| chr2-241927342-241927843 | 5 | 0.944724125770107 |
| chr4-126463755-126464256 | 1 | 0.85425565074409 |
| chr8-6786170-6786671 | 6 | 0.988636053125861 |
| chr1-180697793-180698294 | 3 | 0.991049879916805 |
| chr9-54143101-54143602 | 3 | 0.925474503578548 |
| chr8-60447956-60448457 | 6 | 0.994005789543049 |
| chr3-55607344-55607845 | 3 | 0.968927993739081 |
| chr10-78870747-78871248 | 5 | 0.983919978840557 |
| chr1-243038327-243038828 | 5 | 0.987858516318369 |
| chr19-45399142-45399643 | 2 | 0.871770854460922 |
| chr15-32075094-32075595 | 8 | 0.932209349564471 |
| chr13-98982711-98983212 | 4 | 0.959820627970283 |
| chr8-98098745-98099246 | 3 | 0.99593553545041 |
| chr7-86522521-86523022 | 3 | 0.857021356672557 |
| chr2-187581374-187581875 | 8 | 0.858930537135423 |
| chr6-37915365-37915866 | 3 | 0.996673985358561 |
| chr3-10625129-10625630 | 7 | 0.931696644548715 |
| chr9-88773397-88773898 | 4 | 0.919370663997399 |
| chr2-21032380-21032881 | 8 | 0.840344303212795 |
| chr4-139682508-139683009 | 8 | 0.988909035016646 |
| chr2-188603792-188604293 | 8 | 0.9569645612717 |
| chr4-44322048-44322549 | 3 | 0.996567725009369 |
| chr1-26704640-26705141 | 4 | 0.900299702460984 |
| chr14-11070157-11070658 | 8 | 0.993493681128311 |
| chr14-41626397-41626898 | 1 | 0.94179180632724 |
| chr12-29190142-29190643 | 5 | 0.990826990423641 |
| chr2-40066325-40066826 | 2 | 0.964535296100025 |
| chr10-60084406-60084907 | 8 | 0.854958476954745 |
| chr14-17526496-17526997 | 8 | 0.867329876548906 |
| chr5-166108322-166108823 | 8 | 0.875388381237676 |
| chr14-4686017-4686518 | 6 | 0.99196612170179 |
| chr13-97591843-97592344 | 2 | 0.855071199563693 |
| chr6-27021853-27022354 | 8 | 0.876951162882693 |
| chr11-36080542-36081043 | 6 | 0.857714156823588 |
| chr1-93944708-93945209 | 8 | 0.86325438665696 |
| chr6-39907348-39907849 | 8 | 0.934029183948269 |
| chr10-76811339-76811840 | 3 | 0.989767975041356 |
| chr8-110811306-110811807 | 3 | 0.9968533788726 |
| chr6-74505751-74506252 | 8 | 0.993938520334104 |
| chr4-127420927-127421428 | 8 | 0.832969289091824 |
| chr14-95910272-95910773 | 4 | 0.946886956171395 |

| | | |
|---|---|---|
| chr11-1021831-1022332 | 3 | 0.871519446074782 |
| chr12-33093539-33094040 | 8 | 0.890417154676065 |
| chr6-81401244-81401745 | 3 | 0.99683347887038 |
| chr14-16342316-16342817 | 6 | 0.991206925481043 |
| chr3-39212393-39212894 | 3 | 0.99593553545041 |
| chr10-71747058-71747559 | 8 | 0.980553747839372 |
| chr2-237715158-237715659 | 6 | 0.886110946294857 |
| chr2-228749796-228750297 | 8 | 0.935464669528169 |
| chr14-77586828-77587329 | 8 | 0.927539679262665 |
| chr5-140186186-140186687 | 3 | 0.877768287173433 |
| chr10-45805622-45806123 | 5 | 0.94373925940891 |
| chr5-38632381-38632882 | 3 | 0.997418424915639 |
| chr15-2750982-2751483 | 8 | 0.995214054427918 |
| chr14-18745873-18746374 | 6 | 0.991599113809853 |
| chr17-24734346-24734847 | 2 | 0.975301225605417 |
| chr3-106917892-106918393 | 8 | 0.877791688334168 |
| chr20-20848393-20848894 | 2 | 0.966799143811998 |
| chr10-98572397-98572898 | 5 | 0.988960232326881 |
| chr8-104824048-104824549 | 3 | 0.998836363932831 |
| chr11-16987345-16987846 | 8 | 0.828794319421897 |
| chr3-117976186-117976687 | 8 | 0.882817032243436 |
| chr11-35798165-35798666 | 8 | 0.873733274468959 |
| chr10-20557220-20557721 | 3 | 0.976832923141503 |
| chr4-41165954-41166455 | 5 | 0.922750115568194 |
| chr5-24893033-24893534 | 5 | 0.98736061991614 |
| chr13-38470527-38471028 | 5 | 0.983701963334581 |
| chr17-31963398-31963899 | 2 | 0.965695842917391 |
| chr14-31894678-31895179 | 3 | 0.987079723982201 |
| chr17-54087262-54087763 | 8 | 0.992913792242369 |
| chr16-21801663-21802164 | 6 | 0.926005594318338 |
| chr8-90690222-90690723 | 1 | 0.886583704578154 |
| chr1-9089835-9090336 | 8 | 0.9922209526523 |
| chr7-113469083-113469584 | 8 | 0.861532388143933 |
| chr7-115054640-115055141 | 2 | 0.848466830168968 |
| chr18-41472762-41473263 | 3 | 0.995102292032559 |
| chr20-31639108-31639609 | 5 | 0.948155968493589 |
| chr7-20173618-20174119 | 8 | 0.934614817305202 |
| chr8-63863404-63863905 | 4 | 0.902236093827304 |
| chrX-103589287-103589788 | 5 | 0.892695162342041 |
| chr11-3673108-3673609 | 3 | 0.99593553545041 |
| chr14-73527673-73528174 | 3 | 0.930855568526647 |
| chr6-10763435-10763936 | 8 | 0.821540290068289 |
| chr13-99361627-99362128 | 8 | 0.837944820530126 |

| | | |
|---|---|---|
| chr20-25212677-25213178 | 5 | 0.983919978840557 |
| chr2-248594859-248595360 | 5 | 0.947522562769065 |
| chr20-9253267-9253768 | 5 | 0.922826010881162 |
| chr4-170425577-170426078 | 3 | 0.982669660071819 |
| chr15-85944516-85945017 | 3 | 0.998004220112959 |
| chr11-57534352-57534853 | 8 | 0.974385320653963 |
| chr10-28189131-28189632 | 2 | 0.896334875224103 |
| chr11-62478874-62479375 | 8 | 0.994903467980769 |
| chr19-53734897-53735398 | 3 | 0.996020717054012 |
| chr8-55792245-55792746 | 4 | 0.876687493361729 |
| chr2-149645550-149646051 | 4 | 0.896214568326173 |
| chr10-3350196-3350697 | 3 | 0.996287960618165 |
| chr16-13670566-13671067 | 3 | 0.996492125681892 |
| chrX-78341781-78342282 | 4 | 0.974470388298243 |
| chr10-13995572-13996073 | 2 | 0.940288092929988 |
| chr5-29352082-29352583 | 2 | 0.861623356529489 |
| chr14-9068199-9068700 | 3 | 0.81636376312041 |
| chr4-160706890-160707391 | 8 | 0.990189672866193 |
| chr2-224960598-224961099 | 3 | 0.996743853635269 |
| chr2-60259470-60259971 | 2 | 0.933438086942827 |
| chr2-186030972-186031473 | 3 | 0.993486055827079 |
| chr2-169715411-169715912 | 6 | 0.955375553644158 |
| chr5-147300357-147300858 | 4 | 0.862699411567909 |
| chr14-75263840-75264341 | 8 | 0.982272031595499 |
| chr13-10741151-10741652 | 3 | 0.991495885785884 |
| chr3-8545061-8545562 | 5 | 0.970111251244366 |
| chr9-67909464-67909965 | 8 | 0.992054796314277 |
| chr1-116803806-116804307 | 3 | 0.8377391659182 |
| chrX-115386047-115386548 | 6 | 0.980272397742582 |
| chr14-3121365-3121866 | 8 | 0.907060068362185 |
| chr7-124317163-124317664 | 3 | 0.99747506079086 |
| chr5-90585940-90586441 | 3 | 0.992961312103787 |
| chr20-28782342-28782843 | 8 | 0.993493681128311 |
| chr4-47003051-47003552 | 8 | 0.986009004194582 |
| chr14-77471646-77472147 | 8 | 0.97764638379703 |
| chr13-66382606-66383107 | 8 | 0.952493761252481 |
| chr6-126400639-126401140 | 5 | 0.975489422852602 |
| chr2-95334242-95334743 | 8 | 0.849508348485397 |
| chr2-116546736-116547237 | 5 | 0.983919978840557 |
| chr3-47185437-47185938 | 3 | 0.996175567561389 |
| chr6-124324538-124325039 | 2 | 0.835649225347194 |
| chr2-169915600-169916101 | 6 | 0.987112249307304 |
| chr1-40114795-40115296 | 3 | 0.895982159091847 |

| | | |
|---|---|---|
| chr17-81556392-81556893 | 3 | 0.998691080957329 |
| chr9-113898215-113898716 | 8 | 0.993990009280908 |
| chr10-79960790-79961291 | 2 | 0.949857246940921 |
| chr17-63514972-63515473 | 3 | 0.977259069598613 |
| chr18-82579750-82580251 | 6 | 0.995822816422042 |
| chr13-30906251-30906752 | 2 | 0.951396034555251 |
| chr4-36091782-36092283 | 6 | 0.990267142472808 |
| chr2-22203544-22204045 | 4 | 0.961337946577903 |
| chr20-32712579-32713080 | 8 | 0.842147470494282 |
| chr14-81134816-81135317 | 2 | 0.862121778154339 |
| chr3-130028796-130029297 | 7 | 0.831143936540811 |
| chr8-49629944-49630445 | 6 | 0.831811402238461 |
| chr9-661513-662014 | 1 | 0.918419077005448 |
| chr2-173084970-173085471 | 3 | 0.99591553056496 |
| chr2-214566988-214567489 | 3 | 0.893295780793919 |
| chr2-104483796-104484297 | 8 | 0.9873560926087 |
| chr18-39773165-39773666 | 5 | 0.960089858893583 |
| chr13-99741345-99741846 | 3 | 0.990596475802716 |
| chr2-213262694-213263195 | 3 | 0.996861943228489 |
| chr6-21599410-21599911 | 8 | 0.833647506800295 |
| chr5-89206644-89207145 | 3 | 0.995884026282164 |
| chr15-8672263-8672764 | 8 | 0.853904872554731 |
| chr16-67048869-67049370 | 1 | 0.897845155764653 |
| chr2-174246386-174246887 | 2 | 0.958124152530419 |
| chr7-8120440-8120941 | 5 | 0.838485647071621 |
| chr18-51928246-51928747 | 8 | 0.99361621433776 |
| chr1-181963215-181963716 | 8 | 0.834910333640885 |
| chr20-755729-756230 | 3 | 0.967924658233334 |
| chr14-1521345-1521846 | 3 | 0.99593553545041 |
| chr20-49769933-49770434 | 6 | 0.993196127754323 |
| chr6-57878352-57878853 | 4 | 0.958089681899318 |
| chr4-118184660-118185161 | 3 | 0.996522406984913 |
| chr1-230574513-230575014 | 6 | 0.993228076316781 |
| chr8-49599612-49600113 | 3 | 0.99672948079131 |
| chr2-36663776-36664277 | 6 | 0.979657329632631 |
| chr8-44495004-44495505 | 5 | 0.978341477624344 |
| chr6-62490500-62491001 | 8 | 0.95158781384672 |
| chr5-110250567-110251068 | 8 | 0.975300668680687 |
| chr13-85022638-85023139 | 6 | 0.984224970839577 |
| chr6-9455632-9456133 | 6 | 0.991785640341767 |
| chr9-3342567-3343068 | 3 | 0.991951590699143 |
| chr13-14163560-14164061 | 6 | 0.995231223853014 |
| chr17-78236479-78236980 | 3 | 0.988787471404976 |

| | | |
|---|---|---|
| chr5-72191775-72192276 | 8 | 0.993493681128311 |
| chr5-69288358-69288859 | 6 | 0.991206925481043 |
| chr2-44718661-44719162 | 3 | 0.970034703962465 |
| chr9-79744886-79745387 | 5 | 0.880331289836497 |
| chr18-66953080-66953581 | 3 | 0.993199667011356 |
| chr1-128378651-128379152 | 8 | 0.94024608100645 |
| chr6-40278951-40279452 | 8 | 0.930203115456203 |
| chr1-206877232-206877733 | 5 | 0.945864672349037 |
| chr10-103769925-103770426 | 3 | 0.996011057902977 |
| chrX-9001062-9001563 | 6 | 0.98547408336658 |
| chr1-2361209-2361710 | 6 | 0.955164362169529 |
| chr5-45389143-45389644 | 6 | 0.992713778899664 |
| chr5-21958263-21958764 | 7 | 0.958342916494143 |
| chr6-110273253-110273754 | 8 | 0.964114325390371 |
| chr14-9398599-9399100 | 5 | 0.947880335722047 |
| chr9-21538887-21539388 | 3 | 0.970074469736232 |
| chr5-58858956-58859457 | 5 | 0.928313927770005 |
| chr13-7640983-7641484 | 3 | 0.970008932443071 |
| chr5-34490180-34490681 | 8 | 0.828550572968016 |
| chr5-57579251-57579752 | 3 | 0.993925437831499 |
| chr5-63047373-63047874 | 3 | 0.993077222009108 |
| chr11-8288065-8288566 | 8 | 0.889512679950956 |
| chr11-29433728-29434229 | 3 | 0.995460757618915 |
| chr3-120887376-120887877 | 8 | 0.957115013634167 |
| chr2-121856578-121857079 | 3 | 0.995664082176581 |
| chr13-98637761-98638262 | 5 | 0.956956153999708 |
| chr13-96886297-96886798 | 5 | 0.985451974494029 |
| chr7-33235363-33235864 | 8 | 0.947878368480785 |
| chr5-48237531-48238032 | 3 | 0.928497018800307 |
| chr17-23244271-23244772 | 8 | 0.866545098416517 |
| chr10-18798804-18799305 | 8 | 0.993493681128311 |
| chr15-49117744-49118245 | 1 | 0.890479082205539 |
| chr3-134689734-134690235 | 1 | 0.850662617094822 |
| chr7-96727486-96727987 | 3 | 0.981568481744728 |
| chr1-173109994-173110495 | 3 | 0.99593553545041 |
| chr1-180492528-180493029 | 8 | 0.953558959293478 |
| chr3-114220817-114221318 | 8 | 0.96060500661586 |
| chr7-96001123-96001624 | 8 | 0.975075045511837 |
| chr4-146556424-146556925 | 4 | 0.90208995989691 |
| chr11-48967106-48967607 | 5 | 0.947432309072396 |
| chr19-18480394-18480895 | 3 | 0.979205047746253 |
| chr9-13611493-13611994 | 8 | 0.964972893279495 |
| chr1-88506658-88507159 | 5 | 0.982669152035559 |

| | | |
|---|---|---|
| chr9-39520843-39521344 | 2 | 0.95719028106053 |
| chr16-7425334-7425835 | 4 | 0.957261197533336 |
| chr9-80709625-80710126 | 3 | 0.945990946928157 |
| chr1-197217482-197217983 | 8 | 0.933112227724597 |
| chr13-68073178-68073679 | 8 | 0.945584530012093 |
| chr2-242306954-242307455 | 1 | 0.871708030055681 |
| chr5-146782507-146783008 | 8 | 0.952377916879344 |
| chr4-36543856-36544357 | 3 | 0.996813148288468 |
| chr16-78440563-78441064 | 8 | 0.961231352337589 |
| chr13-36635915-36636416 | 5 | 0.83469740400658 |
| chr2-29635756-29636257 | 3 | 0.996622360627541 |
| chr1-184078502-184079003 | 1 | 0.894100073020271 |
| chr7-17472740-17473241 | 1 | 0.951603303501582 |
| chr20-24686245-24686746 | 3 | 0.996604275060945 |
| chr13-20908727-20909228 | 3 | 0.995141269445606 |
| chr19-37404282-37404783 | 8 | 0.995197408643903 |
| chr1-91354702-91355203 | 6 | 0.991206925481043 |
| chr5-67982796-67983297 | 3 | 0.993103446017234 |
| chr4-14622106-14622607 | 3 | 0.995573922422031 |
| chr10-35970394-35970895 | 6 | 0.847806105779392 |
| chr17-84551432-84551933 | 6 | 0.972853327294729 |
| chr11-36078894-36079395 | 7 | 0.939883408270203 |
| chr1-190005400-190005901 | 6 | 0.87176685210972 |
| chr15-39764269-39764770 | 8 | 0.852259376784956 |
| chr8-97591168-97591669 | 5 | 0.937723696817816 |
| chr1-13059212-13059713 | 7 | 0.833396543214999 |
| chr16-19978280-19978781 | 3 | 0.830993719310256 |
| chr2-113142618-113143119 | 5 | 0.965841464293082 |
| chr5-104909997-104910498 | 3 | 0.9946267288425 |
| chr16-65496271-65496772 | 8 | 0.92669218382598 |
| chr15-36986860-36987361 | 3 | 0.949389213537389 |
| chr2-232314532-232315033 | 8 | 0.993493681128311 |
| chr1-123797048-123797549 | 6 | 0.991206925481043 |
| chr11-40315213-40315714 | 3 | 0.974922559971343 |
| chr16-11549821-11550322 | 3 | 0.99593553545041 |
| chr2-52118724-52119225 | 2 | 0.949020373373298 |
| chr6-138530491-138530992 | 8 | 0.978225812789746 |
| chr16-10121045-10121546 | 6 | 0.943884669438949 |
| chr2-28696215-28696716 | 7 | 0.829907290099247 |
| chr1-11126960-11127461 | 8 | 0.995227048954799 |
| chr9-18824108-18824609 | 5 | 0.983919978840557 |
| chr4-30313869-30314370 | 3 | 0.99593553545041 |
| chr19-32229039-32229540 | 6 | 0.991206925481043 |

| | | |
|---|---|---|
| chr16-71361979-71362480 | 3 | 0.99593553545041 |
| chr20-21275881-21276382 | 5 | 0.914933004163626 |
| chr14-15347523-15348024 | 8 | 0.860766327009424 |
| chr17-20533527-20534028 | 4 | 0.893299043831436 |
| chr20-52766307-52766808 | 8 | 0.946098234382888 |
| chr18-78919637-78920138 | 3 | 0.832836071178932 |
| chr12-29751098-29751599 | 5 | 0.987428294251812 |
| chr7-40748942-40749443 | 6 | 0.953179632611427 |
| chr1-151614910-151615411 | 3 | 0.996745636166007 |
| chr2-25646588-25647089 | 6 | 0.901525296232909 |
| chr3-115255583-115256084 | 5 | 0.970029147389461 |
| chr8-102128663-102129164 | 2 | 0.963417245460374 |
| chr2-174275265-174275766 | 8 | 0.85479791419703 |
| chr1-185020005-185020506 | 8 | 0.847094048206909 |
| chr19-17837872-17838373 | 5 | 0.8712798318705 |
| chr6-59803191-59803692 | 3 | 0.983564869768527 |
| chr1-204093491-204093992 | 8 | 0.889485968908228 |
| chr5-91470776-91471277 | 3 | 0.99336454344445 |
| chr6-129568138-129568639 | 8 | 0.987451527634634 |
| chr19-11221810-11222311 | 3 | 0.926301109623447 |
| chr4-64871334-64871835 | 1 | 0.942395700064525 |
| chr10-43780311-43780812 | 3 | 0.992944525422242 |
| chr8-58641192-58641693 | 8 | 0.943144222716504 |
| chr5-108153688-108154189 | 3 | 0.99593553545041 |
| chr13-35518318-35518819 | 3 | 0.994889889497659 |
| chr2-3400001-3400502 | 3 | 0.99593553545041 |
| chr6-37862135-37862636 | 3 | 0.992886291574434 |
| chr8-65350734-65351235 | 3 | 0.995218414501872 |
| chr18-67651555-67652056 | 5 | 0.890589789099511 |
| chr5-14267415-14267916 | 3 | 0.990065864562589 |
| chr17-79928625-79929126 | 3 | 0.99593553545041 |
| chr7-30884894-30885395 | 8 | 0.986693596817272 |
| chr2-184147173-184147674 | 8 | 0.978909700860773 |
| chr8-50085669-50086170 | 4 | 0.967484276341656 |
| chr14-4887797-4888298 | 3 | 0.991441274222938 |
| chr1-221246833-221247334 | 5 | 0.988054580054712 |
| chr17-75676239-75676740 | 6 | 0.852674758452672 |
| chr15-100939187-100939688 | 3 | 0.92517110498404 |
| chr3-130218441-130218942 | 3 | 0.99593553545041 |
| chr19-32036894-32037395 | 1 | 0.863471767434919 |
| chrX-106354537-106355038 | 5 | 0.982626838859047 |
| chr18-37026153-37026654 | 8 | 0.893552654654948 |
| chr4-145353538-145354039 | 3 | 0.976034596806218 |

| | | |
|---|---|---|
| chr15-78644331-78644832 | 1 | 0.898867904674046 |
| chr9-38454153-38454654 | 8 | 0.995092676648594 |
| chr3-81813116-81813617 | 3 | 0.99593553545041 |
| chr20-27020799-27021300 | 3 | 0.997123690696114 |
| chr7-11288820-11289321 | 8 | 0.943468283203773 |
| chr10-46826922-46827423 | 2 | 0.937598064653615 |
| chr6-118408026-118408527 | 6 | 0.989815015046739 |
| chr10-66130640-66131141 | 3 | 0.907192415609424 |
| chr10-10038705-10039206 | 4 | 0.933257366718804 |
| chr13-62267103-62267604 | 3 | 0.996615796050092 |
| chr1-10468444-10468945 | 8 | 0.967782274892209 |
| chr1-21774136-21774637 | 3 | 0.996676228639208 |
| chr1-120219082-120219583 | 1 | 0.922136991764456 |
| chr16-42572346-42572847 | 6 | 0.949766958182101 |
| chr14-9300244-9300745 | 8 | 0.995182933774798 |
| chr4-176448700-176449201 | 6 | 0.901680332338862 |
| chrX-9667287-9667788 | 2 | 0.861020757933594 |
| chr16-79958414-79958915 | 4 | 0.908324519585237 |
| chr2-211752535-211753036 | 6 | 0.969757838685787 |
| chr13-42062277-42062778 | 8 | 0.995235858342984 |
| chr8-110865566-110866067 | 3 | 0.884753667946664 |
| chr2-74517213-74517714 | 3 | 0.996888857997728 |
| chr6-87805774-87806275 | 6 | 0.967463770343322 |
| chr19-9868183-9868684 | 5 | 0.976292297770444 |
| chr15-13267576-13268077 | 5 | 0.98020870731814 |
| chr16-58736601-58737102 | 3 | 0.995368606396181 |
| chr2-42280282-42280783 | 7 | 0.843857176435693 |
| chr20-8951211-8951712 | 5 | 0.94748458615853 |
| chr12-40943203-40943704 | 4 | 0.915954441954737 |
| chr19-13305553-13306054 | 5 | 0.983919978840557 |
| chr1-128315209-128315710 | 8 | 0.900497809597073 |
| chr8-57586922-57587423 | 3 | 0.896939295886057 |
| chr11-13398696-13399197 | 7 | 0.901200801955026 |
| chr18-68373333-68373834 | 8 | 0.918144311757578 |
| chr11-78664274-78664775 | 8 | 0.877412159568862 |
| chr1-141541068-141541569 | 6 | 0.988257055277845 |
| chr15-75132002-75132503 | 8 | 0.914786478874531 |
| chr9-67600959-67601460 | 3 | 0.99593553545041 |
| chr4-123531086-123531587 | 8 | 0.853068868582601 |
| chr11-37427438-37427939 | 4 | 0.922666848003052 |
| chr10-97199944-97200445 | 3 | 0.997259147191747 |
| chr8-73859775-73860276 | 8 | 0.965229586829466 |
| chr1-249270427-249270928 | 1 | 0.9223987250913 |

| | | |
|---|---|---|
| chr1-249298716-249299217 | 8 | 0.987559121676707 |
| chr15-543003-543504 | 6 | 0.990584566887248 |
| chr20-13980516-13981017 | 3 | 0.897094951998791 |
| chr5-105850678-105851179 | 3 | 0.987379404578523 |
| chr5-16163700-16164201 | 5 | 0.991202217536389 |
| chr5-158763430-158763931 | 5 | 0.934238678565091 |
| chr4-7633774-7634275 | 5 | 0.897203441119654 |
| chr6-57790415-57790916 | 8 | 0.928413947071179 |
| chr18-69159130-69159631 | 8 | 0.820386195110673 |
| chr6-77025158-77025659 | 6 | 0.974719909334836 |
| chr2-56457884-56458385 | 3 | 0.992812383606831 |
| chr17-57406342-57406843 | 8 | 0.846924196572218 |
| chr14-83185274-83185775 | 1 | 0.86731247235581 |
| chr13-25768799-25769300 | 3 | 0.948702120145092 |
| chr4-146701108-146701609 | 3 | 0.90358808849924 |
| chr14-71296009-71296510 | 1 | 0.843322763877137 |
| chr9-49944791-49945292 | 4 | 0.892861032074397 |
| chr13-63291680-63292181 | 8 | 0.960657313501617 |
| chr9-35309445-35309946 | 3 | 0.817496093303875 |
| chr13-69523123-69523624 | 2 | 0.881174983260104 |
| chr7-17636878-17637379 | 2 | 0.945481148264996 |
| chr11-57090083-57090584 | 5 | 0.92641610280561 |
| chrX-90384904-90385405 | 3 | 0.99593553545041 |
| chr2-44862933-44863434 | 8 | 0.910579365419145 |
| chr5-83176653-83177154 | 6 | 0.990117906291205 |
| chr2-111514680-111515181 | 3 | 0.922309840283263 |
| chr7-129335646-129336147 | 3 | 0.996898025068606 |
| chr7-41151766-41152267 | 6 | 0.897955414426622 |
| chr18-10808180-10808681 | 6 | 0.98127924823251 |
| chr7-93522388-93522889 | 3 | 0.99593553545041 |
| chr14-31123175-31123676 | 8 | 0.86421135697592 |
| chr8-115282378-115282879 | 6 | 0.971377798470626 |
| chr13-12424422-12424923 | 6 | 0.888116320813617 |
| chr15-78192074-78192575 | 5 | 0.849850364701113 |
| chr4-12580108-12580609 | 6 | 0.980102784678594 |
| chr18-22987319-22987820 | 3 | 0.99593553545041 |
| chr3-153276645-153277146 | 7 | 0.967693499507306 |
| chrX-29443415-29443916 | 1 | 0.820320376065167 |
| chr6-11972066-11972567 | 1 | 0.915577362523594 |
| chr1-137940713-137941214 | 4 | 0.898980667440019 |
| chr9-91456889-91457390 | 3 | 0.879623238530138 |
| chr10-71971579-71972080 | 8 | 0.958639551954843 |
| chr7-81093859-81094360 | 5 | 0.9895745689108 |

| | | |
|---|---|---|
| chr5-109980167-109980668 | 6 | 0.933252607126989 |
| chr7-127493541-127494042 | 8 | 0.829760689389993 |
| chr10-73418825-73419326 | 7 | 0.927055268611094 |
| chr20-32710243-32710744 | 3 | 0.996927029049571 |
| chr1-32382371-32382872 | 3 | 0.871938989219421 |
| chr1-175443121-175443622 | 6 | 0.992658159324192 |
| chr8-56704211-56704712 | 7 | 0.950538551187691 |
| chr19-20580141-20580642 | 6 | 0.992935485787459 |
| chr2-72557589-72558090 | 6 | 0.926972441391106 |
| chr9-67046000-67046501 | 1 | 0.960331792968783 |
| chr3-60945323-60945824 | 8 | 0.884645593400175 |
| chr1-26756707-26757208 | 3 | 0.976262110339233 |
| chr19-35327167-35327668 | 8 | 0.867403283731397 |
| chr1-2842015-2842516 | 2 | 0.906579419500192 |
| chr8-119193298-119193799 | 6 | 0.928786600701413 |
| chr2-190487030-190487531 | 8 | 0.988386593826384 |
| chr1-242422673-242423174 | 1 | 0.891491634146202 |
| chr11-37430390-37430891 | 6 | 0.991206925481043 |
| chr18-58503859-58504360 | 8 | 0.911707486726059 |
| chr6-21614584-21615085 | 1 | 0.975279629720229 |
| chr15-14452265-14452766 | 3 | 0.972481772185424 |
| chr3-17633476-17633977 | 3 | 0.986736876915339 |
| chr6-119949552-119950053 | 8 | 0.993493681128311 |
| chrX-58577319-58577820 | 5 | 0.931710739273321 |
| chr18-3438232-3438733 | 3 | 0.995813820413183 |
| chr1-91285360-91285861 | 6 | 0.987899593371307 |
| chr10-103263242-103263743 | 1 | 0.903765081362896 |
| chr13-80196209-80196710 | 8 | 0.864216919120528 |
| chr2-164503319-164503820 | 3 | 0.98493688966329 |
| chr15-95510480-95510981 | 3 | 0.895517172555872 |
| chr6-97038293-97038794 | 3 | 0.99593553545041 |
| chr2-184768639-184769140 | 8 | 0.989395016959515 |
| chr10-19196440-19196941 | 8 | 0.897209870872973 |
| chr5-45839461-45839962 | 4 | 0.866576707676161 |
| chr17-59144136-59144637 | 3 | 0.996632633799964 |
| chr19-52316492-52316993 | 6 | 0.990863706423598 |
| chr2-37616182-37616683 | 1 | 0.904212497571577 |
| chr4-180520057-180520558 | 8 | 0.895194715238815 |
| chr17-73142738-73143239 | 5 | 0.994026535559566 |
| chr2-193244653-193245154 | 2 | 0.895315064318727 |
| chr4-49899812-49900313 | 3 | 0.984147136516339 |
| chr16-20079190-20079691 | 2 | 0.879282667575543 |
| chr11-36868383-36868884 | 5 | 0.983862149883347 |

| | | |
|---|---|---|
| chr7-98194871-98195372 | 8 | 0.995248390296068 |
| chr5-119708871-119709372 | 6 | 0.987492542508704 |
| chr8-63399074-63399575 | 3 | 0.99663796996471 |
| chr12-44492518-44493019 | 7 | 0.945775795916412 |
| chr3-140207806-140208307 | 3 | 0.982299149003764 |
| chr1-17160002-17160503 | 6 | 0.838390993436906 |
| chr6-126663667-126664168 | 8 | 0.915306820363305 |
| chr11-57684325-57684826 | 3 | 0.988531680072905 |
| chr3-32743738-32744239 | 1 | 0.86441540577185 |
| chr4-116274875-116275376 | 4 | 0.887824685412878 |
| chr1-6353581-6354082 | 4 | 0.901213516498521 |
| chr17-62476028-62476529 | 3 | 0.99593553545041 |
| chr13-38010663-38011164 | 5 | 0.992265265262827 |
| chr2-34528202-34528703 | 4 | 0.933850094819496 |
| chr15-20465995-20466496 | 1 | 0.906037011325171 |
| chr2-241359363-241359864 | 8 | 0.826087769895381 |
| chr4-114263735-114264236 | 3 | 0.98934544075366 |
| chr12-16675135-16675636 | 4 | 0.87856632324243 |
| chr5-99091760-99092261 | 8 | 0.912175613123471 |
| chr2-184985850-184986351 | 3 | 0.994039963344947 |
| chr16-55998343-55998844 | 8 | 0.86443576312046 |
| chr16-25061544-25062045 | 8 | 0.887703175475312 |
| chr6-62016980-62017481 | 6 | 0.991206925481043 |
| chr1-48607022-48607523 | 3 | 0.997213338569316 |
| chr17-51695453-51695954 | 5 | 0.981666865683646 |
| chr5-154192190-154192691 | 8 | 0.942122604944083 |
| chr16-5976164-5976665 | 6 | 0.989697719437991 |
| chr19-40623670-40624171 | 6 | 0.89244886512191 |
| chr6-70034811-70035312 | 8 | 0.820017588315503 |
| chr17-17918002-17918503 | 2 | 0.845814033819987 |
| chr1-233217625-233218126 | 8 | 0.989251609350921 |
| chr2-167348292-167348793 | 8 | 0.900605265657881 |
| chr3-148227968-148228469 | 3 | 0.996830526013921 |
| chr3-131222403-131222904 | 3 | 0.857502808116957 |
| chr3-78483751-78484252 | 4 | 0.843274266115396 |
| chr4-77722727-77723228 | 3 | 0.864659909708481 |
| chr7-106614535-106615036 | 6 | 0.98935572997718 |
| chr11-76701123-76701624 | 8 | 0.955291228496399 |
| chr1-30496363-30496864 | 4 | 0.961399883552657 |
| chr15-6293101-6293602 | 3 | 0.982926059889555 |
| chr10-4554863-4555364 | 8 | 0.885591903305262 |
| chr8-3958633-3959134 | 3 | 0.997169853195974 |
| chr18-59334210-59334711 | 8 | 0.87725530074677 |

| | | |
|---|---|---|
| chr9-23172013-23172514 | 6 | 0.992692885091862 |
| chr8-98248003-98248504 | 2 | 0.924487771501278 |
| chr10-82501630-82502131 | 3 | 0.832990808380678 |
| chr3-44221322-44221823 | 2 | 0.914561722092647 |
| chr16-40994394-40994895 | 3 | 0.968657309135174 |
| chr3-129010996-129011497 | 4 | 0.902156542781104 |
| chr5-60910947-60911448 | 7 | 0.862773779520865 |
| chr3-35440119-35440620 | 8 | 0.914567073982904 |
| chr1-54056416-54056917 | 6 | 0.833357082183051 |
| chr20-28164150-28164651 | 8 | 0.898805210448306 |
| chr6-110567341-110567842 | 8 | 0.993493681128311 |
| chr1-106917633-106918134 | 1 | 0.915984108554591 |
| chr13-91348518-91349019 | 4 | 0.966614660470684 |
| chr2-204911693-204912194 | 1 | 0.834771463860511 |
| chr10-35377815-35378316 | 3 | 0.919994177967462 |
| chr5-150820850-150821351 | 8 | 0.988667697812139 |
| chr15-94608771-94609272 | 3 | 0.994325469396148 |
| chr8-83341698-83342199 | 5 | 0.822597518459396 |
| chr8-122114820-122115321 | 1 | 0.826925815132916 |
| chr15-47560715-47561216 | 6 | 0.996135222612157 |
| chr9-93070944-93071445 | 8 | 0.853554946845074 |
| chr9-83977199-83977700 | 3 | 0.996529490141634 |
| chr9-45604549-45605050 | 3 | 0.996660084377339 |
| chr12-22336434-22336935 | 3 | 0.973739745559978 |
| chr16-26307431-26307932 | 2 | 0.887801171993424 |
| chr7-105049474-105049975 | 4 | 0.819708062934644 |
| chr17-16208337-16208838 | 3 | 0.996477831957315 |
| chr9-70270896-70271397 | 6 | 0.966476808344223 |
| chr5-77853888-77854389 | 3 | 0.981933778506816 |
| chr17-20613274-20613775 | 5 | 0.850227248221329 |
| chr1-256249145-256249646 | 1 | 0.837150672525408 |
| chr13-96187339-96187840 | 4 | 0.974053803467726 |
| chr2-43249243-43249744 | 3 | 0.996736578581218 |
| chr20-8840209-8840710 | 5 | 0.988723068149632 |
| chr4-59828930-59829431 | 8 | 0.987458258319873 |
| chr1-177976152-177976653 | 6 | 0.988744449472269 |
| chr2-201088883-201089384 | 3 | 0.955941373587641 |
| chr6-26262778-26263279 | 3 | 0.996863613959495 |
| chr5-130325632-130326133 | 8 | 0.888946412413793 |
| chr18-53913252-53913753 | 6 | 0.991206925481043 |
| chr13-67691557-67692058 | 1 | 0.888006083049033 |
| chr1-105280168-105280669 | 3 | 0.995032888430768 |
| chr10-15175299-15175800 | 8 | 0.975726398953019 |

| | | |
|---|---|---|
| chr9-6721236-6721737 | 8 | 0.931177369263235 |
| chr14-11572979-11573480 | 8 | 0.962990463101633 |
| chr1-189689820-189690321 | 1 | 0.859646770405536 |
| chr2-120316704-120317205 | 4 | 0.955328998138825 |
| chr12-20291583-20292084 | 4 | 0.854072553256704 |
| chr10-83210063-83210564 | 5 | 0.876227961532022 |
| chr3-163415485-163415986 | 8 | 0.970328534275025 |
| chr5-108256001-108256502 | 4 | 0.822402002438308 |
| chr2-112676981-112677482 | 8 | 0.922654633074592 |
| chr2-215179562-215180063 | 5 | 0.920378312905782 |
| chr3-56269863-56270364 | 3 | 0.997228780290097 |
| chr9-104485823-104486324 | 5 | 0.972562752078129 |
| chr14-57775819-57776320 | 5 | 0.986162988141772 |
| chr5-137214842-137215343 | 6 | 0.993226814866666 |
| chr4-116779073-116779574 | 3 | 0.955477132072191 |
| chr7-103413687-103414188 | 8 | 0.935686593308155 |
| chr7-95077830-95078331 | 8 | 0.825581052769736 |
| chr15-98078308-98078809 | 7 | 0.85331681285562 |
| chr7-99252137-99252638 | 8 | 0.980535745778913 |
| chr7-52713951-52714452 | 4 | 0.845471512090779 |
| chr15-49637132-49637633 | 8 | 0.993493681128311 |
| chr3-159607287-159607788 | 8 | 0.853980615068533 |
| chr6-102298547-102299048 | 3 | 0.99593553545041 |
| chrX-71741995-71742496 | 3 | 0.996623841393587 |
| chr17-19243821-19244322 | 8 | 0.882741651362632 |
| chr10-19258728-19259229 | 6 | 0.991206925481043 |
| chr2-33529201-33529702 | 6 | 0.986173400948905 |
| chr11-84292035-84292536 | 2 | 0.836680605699612 |
| chr7-121648554-121649055 | 4 | 0.929432315024849 |
| chr2-148869237-148869738 | 3 | 0.992361044850461 |
| chr4-103927891-103928392 | 5 | 0.985582459893043 |
| chr2-146801851-146802352 | 6 | 0.955191953441909 |
| chr8-115235188-115235689 | 7 | 0.825414648442911 |
| chr5-102396960-102397461 | 6 | 0.993127283833996 |
| chr3-62345425-62345926 | 5 | 0.821372574736932 |
| chr3-113771132-113771633 | 6 | 0.903320413762098 |
| chr9-65394170-65394671 | 3 | 0.984423966802849 |
| chr4-130129404-130129905 | 2 | 0.863566113241947 |
| chr9-76442078-76442579 | 5 | 0.985013536347357 |
| chr18-38559174-38559675 | 1 | 0.924519843451664 |
| chr11-8566726-8567227 | 3 | 0.99593553545041 |
| chr17-55719016-55719517 | 3 | 0.862435815747618 |
| chr1-192294463-192294964 | 8 | 0.84642838312672 |

| | | |
|---|---|---|
| chr14-93131134-93131635 | 3 | 0.996821906698611 |
| chr3-38046804-38047305 | 6 | 0.90316849956378 |
| chr2-12418950-12419451 | 3 | 0.99593553545041 |
| chr11-23724280-23724781 | 2 | 0.93286729767671 |
| chr10-6551930-6552431 | 3 | 0.898317824615556 |
| chr16-25471592-25472093 | 3 | 0.864086691632562 |
| chr2-192791967-192792468 | 8 | 0.938240193712633 |
| chr5-27037971-27038472 | 8 | 0.870392078364831 |
| chr4-48903877-48904378 | 3 | 0.99593553545041 |
| chr4-147709244-147709745 | 5 | 0.956176042718301 |
| chr17-69888983-69889484 | 6 | 0.976835310325491 |
| chr17-7664274-7664775 | 8 | 0.831683108733063 |
| chr11-51073531-51074032 | 8 | 0.834251249521806 |
| chr7-127279699-127280200 | 8 | 0.971914744563549 |
| chr17-69306258-69306759 | 3 | 0.995577065606226 |
| chr17-19273182-19273683 | 5 | 0.972639185709298 |
| chr16-19840685-19841186 | 3 | 0.995534147101952 |
| chr4-160333473-160333974 | 3 | 0.996585699990252 |
| chr6-17290549-17291050 | 8 | 0.968856113063216 |
| chr14-7470933-7471434 | 5 | 0.983592928149057 |
| chrX-148823719-148824220 | 6 | 0.991206925481043 |
| chr16-71384851-71385352 | 7 | 0.851496816276232 |
| chr15-76112913-76113414 | 2 | 0.937369036021518 |
| chr1-217427906-217428407 | 3 | 0.995189149855547 |
| chr13-77940227-77940728 | 7 | 0.889758659283632 |
| chr6-126206142-126206643 | 3 | 0.978868599140771 |
| chr5-56163873-56164374 | 5 | 0.949110297300076 |
| chr1-178575482-178575983 | 6 | 0.992961967094357 |
| chr16-79145061-79145562 | 8 | 0.850629559799956 |
| chr13-91134026-91134527 | 6 | 0.990627589206779 |
| chr16-66926-67427 | 5 | 0.988052175431667 |
| chr10-13598977-13599478 | 4 | 0.824377725742367 |
| chrX-32323754-32324255 | 8 | 0.993984583892088 |
| chr5-54340405-54340906 | 6 | 0.988280827240831 |
| chr2-73326422-73326923 | 3 | 0.948561256507585 |
| chrX-33796769-33797270 | 5 | 0.976979568549661 |
| chr19-38260294-38260795 | 6 | 0.993132013048084 |
| chr14-92987343-92987844 | 2 | 0.922992918189376 |
| chr11-25781764-25782265 | 8 | 0.964474004729617 |
| chr2-78095875-78096376 | 6 | 0.991754163801822 |
| chr6-3974893-3975394 | 8 | 0.878112681993718 |
| chr18-44819442-44819943 | 8 | 0.89720740418041 |
| chr14-40469043-40469544 | 3 | 0.995389865671708 |

| | | |
|---|---|---|
| chr11-2916790-2917291 | 6 | 0.847512809252826 |
| chr4-76107906-76108407 | 3 | 0.939307829958166 |
| chr3-33475377-33475878 | 5 | 0.910571310728497 |
| chr10-19005674-19006175 | 6 | 0.985965631202709 |
| chr6-7314496-7314997 | 6 | 0.962660949903488 |
| chr3-44093028-44093529 | 8 | 0.933664745561049 |
| chr2-68869852-68870353 | 6 | 0.991206925481043 |
| chr9-57337388-57337889 | 6 | 0.886746858139425 |
| chr2-26519955-26520456 | 5 | 0.892092782789442 |
| chr14-35344872-35345373 | 5 | 0.993727052684009 |
| chr10-19060044-19060545 | 8 | 0.939862954517414 |
| chr3-1018385-1018886 | 3 | 0.97954868744565 |
| chr8-122464718-122465219 | 6 | 0.994063202787396 |
| chr1-106790196-106790697 | 8 | 0.826987409592326 |
| chr4-139656490-139656991 | 3 | 0.99593553545041 |
| chr6-42648808-42649309 | 6 | 0.993170387114212 |
| chr4-18943792-18944293 | 3 | 0.877147946852817 |
| chr6-131566367-131566868 | 3 | 0.851970372495655 |
| chr8-73553610-73554111 | 5 | 0.968799689881396 |
| chr10-57782905-57783406 | 7 | 0.838364420422354 |
| chr9-54905054-54905555 | 8 | 0.984990323102351 |
| chr4-107860721-107861222 | 5 | 0.983919978840557 |
| chr1-46397555-46398056 | 6 | 0.970326264659254 |
| chr2-235893536-235894037 | 8 | 0.966987351421173 |
| chr4-14745993-14746494 | 3 | 0.99593553545041 |
| chr1-234741780-234742281 | 2 | 0.950577135602226 |
| chr2-143210603-143211104 | 3 | 0.99593553545041 |
| chr1-100179222-100179723 | 3 | 0.99593553545041 |
| chr1-219660272-219660773 | 3 | 0.953595431935332 |
| chr1-47353175-47353676 | 6 | 0.923176169675312 |
| chr13-45693400-45693901 | 5 | 0.94505706390985 |
| chr7-87184983-87185484 | 5 | 0.991696714930914 |
| chr10-28168478-28168979 | 5 | 0.976748974657077 |
| chr17-18148964-18149465 | 8 | 0.917643551436176 |
| chr3-105407898-105408399 | 6 | 0.951334924232499 |
| chr1-225733728-225734229 | 8 | 0.943146274197982 |
| chr19-47704730-47705231 | 5 | 0.970314725151366 |
| chr17-9740699-9741200 | 6 | 0.990122420292357 |
| chr7-100866520-100867021 | 8 | 0.819758987067008 |
| chr1-233946425-233946926 | 8 | 0.959831477725413 |
| chr3-128850130-128850631 | 6 | 0.967744039547832 |
| chr1-142014001-142014502 | 6 | 0.990141763499022 |
| chr4-113389309-113389810 | 3 | 0.995854809388033 |

| | | |
|---|---|---|
| chr12-40630878-40631379 | 7 | 0.871060405405142 |
| chr12-32713727-32714228 | 8 | 0.970869795826826 |
| chr4-126047627-126048128 | 6 | 0.992430194294401 |
| chr9-63874816-63875317 | 5 | 0.991520280920899 |
| chr13-5689579-5690080 | 3 | 0.99484662893667 |
| chr1-258739602-258740103 | 2 | 0.86520798555132 |
| chr7-31491339-31491840 | 3 | 0.99703332160348 |
| chr6-25685481-25685982 | 7 | 0.946854769722555 |
| chr15-45575273-45575774 | 4 | 0.879368653454974 |
| chr15-70927321-70927822 | 4 | 0.972557835783282 |
| chr1-40341382-40341883 | 5 | 0.825617682198992 |
| chr1-170857589-170858090 | 3 | 0.924965232391061 |
| chr12-24842204-24842705 | 5 | 0.980074995245714 |
| chr3-33245407-33245908 | 8 | 0.859431670136825 |
| chr2-197364427-197364928 | 2 | 0.814909077884457 |
| chr14-55422471-55422972 | 3 | 0.99689813110344 |
| chr3-33626667-33627168 | 7 | 0.868468885715864 |
| chr9-16932769-16933270 | 3 | 0.996386938770266 |
| chr8-86820817-86821318 | 8 | 0.985176112122661 |
| chr15-76957011-76957512 | 8 | 0.95658313738144 |
| chr15-38546195-38546696 | 8 | 0.955031629954921 |
| chr2-89955086-89955587 | 3 | 0.989075512512415 |
| chr3-62565726-62566227 | 3 | 0.973121701748517 |
| chr4-78044804-78045305 | 6 | 0.98227681894521 |
| chr13-66456436-66456937 | 6 | 0.993048638346238 |
| chr7-112349446-112349947 | 8 | 0.976647167979814 |
| chr6-62759757-62760258 | 6 | 0.991206925481043 |
| chr5-164481996-164482497 | 8 | 0.935293837491931 |
| chr1-3006908-3007409 | 6 | 0.996208308203142 |
| chr7-27373887-27374388 | 5 | 0.987368861743345 |
| chr8-116227738-116228239 | 3 | 0.995933287465569 |
| chr6-68667597-68668098 | 5 | 0.957610035520271 |
| chr11-4637822-4638323 | 3 | 0.994533205588537 |
| chr17-18257517-18258018 | 8 | 0.982109609784884 |
| chr6-127873623-127874124 | 4 | 0.881382250102335 |
| chr3-2403312-2403813 | 3 | 0.965890481948734 |
| chr5-32022499-32023000 | 6 | 0.929559853507956 |
| chr15-4688278-4688779 | 3 | 0.849524017132115 |
| chr4-176339323-176339824 | 8 | 0.875956077651291 |
| chr3-93470458-93470959 | 5 | 0.923891181905834 |
| chr5-16717889-16718390 | 7 | 0.835129388625634 |
| chr6-137531874-137532375 | 6 | 0.969128078337683 |
| chr9-88027094-88027595 | 5 | 0.989069390509453 |

| | | |
|---|---|---|
| chr17-65915377-65915878 | 8 | 0.83188937385298 |
| chr20-32200044-32200545 | 2 | 0.898503121981699 |
| chr11-83535664-83536165 | 8 | 0.902018702412153 |
| chr14-48467991-48468492 | 3 | 0.99593553545041 |
| chr1-1159321-1159822 | 6 | 0.992961734580369 |
| chr2-14488097-14488598 | 3 | 0.98770152863202 |
| chr13-56095306-56095807 | 2 | 0.900390651273667 |
| chr13-6980048-6980549 | 3 | 0.996897086900689 |
| chr3-1102762-1103263 | 3 | 0.988667936902676 |
| chr2-125962560-125963061 | 3 | 0.99593553545041 |
| chr2-134132316-134132817 | 3 | 0.9543673814963 |
| chr3-12107105-12107606 | 4 | 0.821338994030735 |
| chr3-48081043-48081544 | 8 | 0.990147016190704 |
| chr17-68522579-68523080 | 6 | 0.992489824795435 |
| chr2-172509272-172509773 | 3 | 0.989633578468294 |
| chr6-38012019-38012520 | 4 | 0.960384918097642 |
| chr10-47392275-47392776 | 6 | 0.974416379714148 |
| chr2-229169462-229169963 | 5 | 0.968049348638846 |
| chr4-163297193-163297694 | 3 | 0.908916197225966 |
| chr14-95864346-95864847 | 3 | 0.915401639533243 |
| chr17-5240714-5241215 | 7 | 0.868527357366146 |
| chr10-74829707-74830208 | 8 | 0.982237455304409 |
| chr19-45586506-45587007 | 8 | 0.916894104287771 |
| chr2-169518424-169518925 | 8 | 0.980206812956699 |
| chr2-45288856-45289357 | 5 | 0.837994601706497 |
| chr10-18320990-18321491 | 3 | 0.995764330718344 |
| chr18-16866870-16867371 | 5 | 0.959119255993931 |
| chr17-39221807-39222308 | 8 | 0.993493681128311 |
| chr8-116446000-116446501 | 3 | 0.973891729792405 |
| chr13-25231181-25231682 | 1 | 0.890479082205539 |
| chr5-109372206-109372707 | 6 | 0.990098393971035 |
| chr13-85792331-85792832 | 3 | 0.973056720896985 |
| chr8-11617493-11617994 | 3 | 0.953960889729367 |
| chr15-75113719-75114220 | 5 | 0.956693844124034 |
| chr1-179657652-179658153 | 8 | 0.982523783657423 |
| chr11-62144065-62144566 | 3 | 0.926073096096921 |
| chr5-87814235-87814736 | 8 | 0.896684750281737 |
| chr5-76009896-76010397 | 3 | 0.99593553545041 |
| chr16-79909850-79910351 | 1 | 0.845913914605564 |
| chr4-12472859-12473360 | 6 | 0.992033744261436 |
| chr10-9915871-9916372 | 6 | 0.992634476999244 |
| chr5-132179599-132180100 | 6 | 0.991773945880776 |
| chr4-74471366-74471867 | 3 | 0.979333310678558 |

| | | |
|---|---|---|
| chr7-125430227-125430728 | 2 | 0.946041969315769 |
| chr3-81287845-81288346 | 6 | 0.990079742444675 |
| chr14-102093378-102093879 | 1 | 0.81941390659204 |
| chr5-137755571-137756072 | 3 | 0.823012245400293 |
| chr4-35338355-35338856 | 2 | 0.969735672942073 |
| chr2-96872306-96872807 | 8 | 0.993440701383319 |
| chr10-106585124-106585625 | 8 | 0.955853759536256 |
| chr5-10374161-10374662 | 5 | 0.989180495871052 |
| chrX-137299122-137299623 | 8 | 0.895713684721068 |
| chr2-219798379-219798880 | 3 | 0.996273349386733 |
| chr7-130541595-130542096 | 8 | 0.928283927694144 |
| chr2-9757490-9757991 | 3 | 0.996542069359078 |
| chr19-50320693-50321194 | 5 | 0.849431919069089 |
| chr17-81075048-81075549 | 5 | 0.982342228444311 |
| chr1-181143350-181143851 | 4 | 0.842067324087502 |
| chr14-8913377-8913878 | 4 | 0.899513217203558 |
| chr18-24629766-24630267 | 6 | 0.965389019959621 |
| chr16-60278366-60278867 | 2 | 0.858760677056239 |
| chr13-106172406-106172907 | 2 | 0.87037038412415 |
| chr3-102486510-102487011 | 5 | 0.960807568199146 |
| chr1-166262489-166262990 | 3 | 0.98346847276499 |
| chr2-97183867-97184368 | 3 | 0.983118755716854 |
| chr7-128819528-128820029 | 8 | 0.878199721987678 |
| chr1-241340803-241341304 | 4 | 0.86156969346054 |
| chr19-55506789-55507290 | 5 | 0.876593522501334 |
| chr1-51901307-51901808 | 8 | 0.974733629051213 |
| chr15-57752837-57753338 | 8 | 0.878411455306995 |
| chr19-36525949-36526450 | 7 | 0.917381632267589 |
| chr20-1520228-1520729 | 8 | 0.903227293520908 |
| chr18-45815389-45815890 | 2 | 0.837501188474441 |
| chr10-64478037-64478538 | 8 | 0.9401387595276 |
| chr4-156320079-156320580 | 6 | 0.990054362343865 |
| chr8-21827945-21828446 | 4 | 0.922256555140061 |
| chr18-11445700-11446201 | 8 | 0.975363896699321 |
| chr17-65962132-65962633 | 3 | 0.985050057031252 |
| chr9-12036508-12037009 | 8 | 0.979160199551542 |
| chr2-242463751-242464252 | 2 | 0.9362033974779 |
| chr2-75215988-75216489 | 3 | 0.996844768644608 |
| chr5-38493939-38494440 | 8 | 0.97602577508859 |
| chr20-5864186-5864687 | 5 | 0.918406420065026 |
| chr4-177528037-177528538 | 8 | 0.881115314285253 |
| chr9-40419514-40420015 | 5 | 0.97819742812404 |
| chr10-23451389-23451890 | 8 | 0.855489505181334 |

| | | |
|---|---|---|
| chr3-128511609-128512110 | 3 | 0.967510355328424 |
| chr5-104191180-104191681 | 1 | 0.860852650284121 |
| chr4-53575548-53576049 | 3 | 0.986048121692886 |
| chr3-120018846-120019347 | 1 | 0.890479082205539 |
| chr12-12349106-12349607 | 2 | 0.92772805374875 |
| chr5-39929853-39930354 | 3 | 0.950644689570449 |
| chr1-43789395-43789896 | 4 | 0.874935614941411 |
| chr1-154683122-154683623 | 3 | 0.933087433109894 |
| chr5-60944091-60944592 | 6 | 0.993630861449006 |
| chr2-95640900-95641401 | 3 | 0.918418581330199 |
| chr3-1250340-1250841 | 6 | 0.968282966389047 |
| chr11-29651655-29652156 | 8 | 0.987886864480805 |
| chr15-8302709-8303210 | 3 | 0.996344122443885 |
| chrX-104692176-104692677 | 1 | 0.879514303927598 |
| chr1-224148179-224148680 | 8 | 0.882644398335275 |
| chr3-35749464-35749965 | 5 | 0.878053985815015 |
| chr14-80048255-80048756 | 6 | 0.996833730450051 |
| chr3-117282888-117283389 | 4 | 0.93278918443826 |
| chr7-106948498-106948999 | 2 | 0.817364019078378 |
| chr3-51836642-51837143 | 6 | 0.992489260680241 |
| chr8-115140043-115140544 | 3 | 0.997035305882847 |
| chr18-73710545-73711046 | 1 | 0.928657388436242 |
| chr1-162436655-162437156 | 6 | 0.991206925481043 |
| chr8-98920862-98921363 | 6 | 0.992239971799875 |
| chr19-50054110-50054611 | 8 | 0.993125491721698 |
| chr8-87265423-87265924 | 8 | 0.841043989980638 |
| chr1-173384795-173385296 | 6 | 0.957858757396396 |
| chr10-68640182-68640683 | 5 | 0.991145307524352 |
| chr8-102495131-102495632 | 3 | 0.997659612276279 |
| chr4-178546253-178546754 | 4 | 0.953528310190811 |
| chr9-61174458-61174959 | 6 | 0.995558056884735 |
| chr16-55522619-55523120 | 8 | 0.839405046220724 |
| chr7-96780938-96781439 | 3 | 0.98041009871642 |
| chr18-40992151-40992652 | 3 | 0.996429762097045 |
| chr5-160798983-160799484 | 8 | 0.983152151162393 |
| chr6-124637951-124638452 | 6 | 0.991206925481043 |
| chr8-111029639-111030140 | 4 | 0.837334039989284 |
| chr5-143362565-143363066 | 4 | 0.851391862309568 |
| chr13-87793507-87794008 | 5 | 0.987915412630052 |
| chr10-96192481-96192982 | 5 | 0.908114976179398 |
| chr13-58994326-58994827 | 2 | 0.954866558114714 |
| chr6-82514604-82515105 | 3 | 0.996766843660291 |
| chr2-120877260-120877761 | 5 | 0.92763372307859 |

| | | |
|---|---|---|
| chr1-15217319-15217820 | 3 | 0.996634484265569 |
| chr18-42662612-42663113 | 6 | 0.971269599813363 |
| chr4-126601726-126602227 | 1 | 0.847170445686763 |
| chr16-66175425-66175926 | 6 | 0.83967394182043 |
| chr1-118727005-118727506 | 6 | 0.917773802314089 |
| chr7-21547713-21548214 | 8 | 0.873062015460969 |
| chr3-159076861-159077362 | 6 | 0.938837133078255 |
| chr2-111425657-111426158 | 4 | 0.887439020128233 |
| chr4-66773546-66774047 | 6 | 0.841778836109262 |
| chr8-468029-468530 | 3 | 0.962509054073763 |
| chr8-101582765-101583266 | 8 | 0.872487021030051 |
| chr4-66489549-66490050 | 3 | 0.887832767832579 |
| chr4-105328727-105329228 | 3 | 0.996607370367141 |
| chr15-100028492-100028993 | 8 | 0.949502411478853 |
| chr16-53750122-53750623 | 3 | 0.986951081889471 |
| chr2-232486474-232486975 | 7 | 0.839198702984923 |
| chr6-137434505-137435006 | 1 | 0.945120451220887 |
| chr1-145777698-145778199 | 3 | 0.900639333196504 |
| chr9-78604831-78605332 | 6 | 0.96576284946286 |
| chr8-110453592-110454093 | 3 | 0.859703158492495 |
| chr1-81845830-81846331 | 3 | 0.979024921460308 |
| chr5-144861340-144861841 | 8 | 0.992341674444386 |
| chr1-58700073-58700574 | 4 | 0.870215737468059 |
| chr1-37948128-37948629 | 5 | 0.989385339706377 |
| chr1-84811743-84812244 | 8 | 0.993553061723642 |
| chr4-179426579-179427080 | 4 | 0.862257216230571 |
| chr6-140675114-140675615 | 3 | 0.99593553545041 |
| chr4-125033537-125034038 | 3 | 0.941474478918388 |
| chr12-11221912-11222413 | 3 | 0.923224358229002 |
| chr14-1224411-1224912 | 2 | 0.963706567678162 |
| chr7-19990320-19990821 | 4 | 0.864387775373462 |
| chr2-143882866-143883367 | 8 | 0.993030985931743 |
| chr8-110617700-110618201 | 6 | 0.997661393598459 |
| chr15-9068336-9068837 | 5 | 0.981262358729705 |
| chr13-99185355-99185856 | 5 | 0.827759023356259 |
| chr6-123846781-123847282 | 5 | 0.988458405860579 |
| chr4-8001459-8001960 | 6 | 0.902132509461902 |
| chr4-79537005-79537506 | 4 | 0.954030806778467 |
| chr8-78578800-78579301 | 4 | 0.924419096953864 |
| chr20-29297014-29297515 | 5 | 0.974662605060297 |
| chr10-67344717-67345218 | 3 | 0.989719576036757 |
| chr16-73876103-73876604 | 6 | 0.946446126781799 |
| chr4-132940872-132941373 | 8 | 0.952421151212563 |

| | | |
|---|---|---|
| chrX-34168680-34169181 | 8 | 0.895062244582705 |
| chr19-40554559-40555060 | 6 | 0.908345854048095 |
| chr8-109568953-109569454 | 8 | 0.980121444952933 |
| chr1-17986145-17986646 | 6 | 0.870401117794709 |
| chr18-7497165-7497666 | 3 | 0.995235970460932 |
| chr14-50498844-50499345 | 3 | 0.984807694491023 |
| chr13-65690566-65691067 | 8 | 0.90872078829106 |
| chr1-155514258-155514759 | 8 | 0.846466860726435 |
| chr7-97690928-97691429 | 3 | 0.996483474163357 |
| chr15-87331899-87332400 | 1 | 0.815973276773937 |
| chr1-120401842-120402343 | 8 | 0.966681504651185 |
| chr7-69727896-69728397 | 3 | 0.827557637983449 |
| chr7-99024447-99024948 | 8 | 0.992673936571447 |
| chr2-137088564-137089065 | 2 | 0.83819088254556 |
| chr1-260178270-260178771 | 6 | 0.992672684374015 |
| chr3-11902159-11902660 | 8 | 0.952500052234168 |
| chr17-65709008-65709509 | 3 | 0.994254800671352 |
| chr11-76668563-76669064 | 8 | 0.994692369725109 |
| chr2-15647542-15648043 | 8 | 0.854463175642681 |
| chr3-144971045-144971546 | 8 | 0.950702909577373 |
| chr15-75750840-75751341 | 3 | 0.995336194498118 |
| chr13-105873081-105873582 | 2 | 0.927574529189447 |
| chr5-5157248-5157749 | 6 | 0.991958000655001 |
| chr7-22018769-22019270 | 3 | 0.863850841104434 |
| chr1-135232209-135232710 | 8 | 0.815645298167872 |
| chr4-105349549-105350050 | 3 | 0.993675334897635 |
| chr16-4821538-4822039 | 3 | 0.91493499353051 |
| chr17-77428457-77428958 | 8 | 0.98483658812676 |
| chr18-55932437-55932938 | 3 | 0.998115789492222 |
| chr12-35545393-35545894 | 6 | 0.996001426907494 |
| chr9-106325685-106326186 | 1 | 0.933250205809471 |
| chr17-13811171-13811672 | 5 | 0.98428018905242 |
| chr3-37160042-37160543 | 5 | 0.991567934452719 |
| chr16-12685402-12685903 | 8 | 0.81570975040311 |
| chr6-125742863-125743364 | 3 | 0.996852459747498 |
| chr4-29675159-29675660 | 6 | 0.976888302478332 |
| chr18-67387919-67388420 | 3 | 0.996896564246578 |
| chr7-98382805-98383306 | 3 | 0.985357093829984 |
| chr10-70960704-70961205 | 5 | 0.969659329027158 |
| chr20-20164309-20164810 | 8 | 0.847188377228814 |
| chr9-65241341-65241842 | 8 | 0.959538552612027 |
| chr14-13679597-13680098 | 3 | 0.996692550282711 |
| chr13-21421226-21421727 | 8 | 0.905097654856902 |

| | | |
|---|---|---|
| chr17-72482471-72482972 | 8 | 0.925645571408353 |
| chr1-157410871-157411372 | 3 | 0.993614178310186 |
| chr4-86138338-86138839 | 6 | 0.972686942721047 |
| chr6-53008387-53008888 | 7 | 0.88328492527475 |
| chr16-71375809-71376310 | 3 | 0.894808411014162 |
| chr5-138986016-138986517 | 3 | 0.993783598103843 |
| chr4-180312556-180313057 | 2 | 0.972758379773532 |
| chrX-97339543-97340044 | 2 | 0.950456264887598 |
| chr5-27009289-27009790 | 4 | 0.968387352020452 |
| chr14-46738031-46738532 | 3 | 0.996857959027645 |
| chr4-150573793-150574294 | 1 | 0.899856041117049 |
| chr14-82655984-82656485 | 7 | 0.856586052010698 |
| chr20-46311922-46312423 | 5 | 0.981226744318095 |
| chr2-178364635-178365136 | 3 | 0.975935283983127 |
| chr7-79662840-79663341 | 8 | 0.862393186375386 |
| chr3-7873970-7874471 | 8 | 0.992375818829731 |
| chr1-170304290-170304791 | 3 | 0.956488719186406 |
| chr13-55737478-55737979 | 5 | 0.897958034141906 |
| chr6-41515621-41516122 | 8 | 0.994957745108651 |
| chr14-74731742-74732243 | 7 | 0.930478508131724 |
| chr18-5251924-5252425 | 1 | 0.909490031225672 |
| chr7-73441993-73442494 | 8 | 0.994662307528206 |
| chr1-30915816-30916317 | 8 | 0.941250415987664 |
| chr1-1386586-1387087 | 5 | 0.977875549336638 |
| chr4-129450184-129450685 | 8 | 0.976192991542012 |
| chr3-163606571-163607072 | 8 | 0.931494037604327 |
| chr9-37219354-37219855 | 3 | 0.99593553545041 |
| chr9-77109287-77109788 | 8 | 0.971199488718079 |
| chr2-192300961-192301462 | 8 | 0.905795049060422 |
| chr8-13246108-13246609 | 3 | 0.978216929991212 |
| chr3-117574157-117574658 | 7 | 0.908997380927528 |
| chr2-174198502-174199003 | 8 | 0.816597623661676 |
| chr1-156787992-156788493 | 8 | 0.99416324180089 |
| chr6-53246136-53246637 | 1 | 0.890479082205539 |
| chr14-103008609-103009110 | 6 | 0.834792994083816 |
| chr2-55934343-55934844 | 3 | 0.96308452833802 |
| chr18-22592030-22592531 | 6 | 0.990861495268164 |
| chr3-83556977-83557478 | 4 | 0.828169536842708 |
| chr3-155777422-155777923 | 8 | 0.881310812339916 |
| chr16-67180227-67180728 | 8 | 0.892222407610206 |
| chr19-38907567-38908068 | 3 | 0.927679460476449 |
| chr4-37921032-37921533 | 8 | 0.971054703518117 |
| chr4-87891509-87892010 | 3 | 0.947903798816052 |

| | | |
|---|---|---|
| chr5-149701372-149701873 | 3 | 0.95218746832151 |
| chr4-44028927-44029428 | 3 | 0.830778113442913 |
| chrX-85373688-85374189 | 6 | 0.83251852605406 |
| chr11-58920069-58920570 | 4 | 0.931618683605403 |
| chrX-149479604-149480105 | 3 | 0.989206059869866 |
| chr4-31785883-31786384 | 8 | 0.905844595546325 |
| chrY-9131621-9132122 | 3 | 0.950667916182718 |
| chr1-181775133-181775634 | 5 | 0.989789925123421 |
| chr4-123840127-123840628 | 3 | 0.932915942914655 |
| chr15-34494011-34494512 | 8 | 0.931694639569575 |
| chr14-90447454-90447955 | 3 | 0.996228644961423 |
| chr2-215239469-215239970 | 8 | 0.89562756789291 |
| chr2-32768633-32769134 | 7 | 0.823379296247927 |
| chr14-33291110-33291611 | 3 | 0.991964606522585 |
| chr13-81897732-81898233 | 8 | 0.887582674713862 |
| chr2-241436681-241437182 | 8 | 0.987527492351938 |
| chr4-11692024-11692525 | 3 | 0.821025752583498 |
| chr14-71433847-71434348 | 1 | 0.916578234570511 |
| chr16-74488856-74489357 | 3 | 0.996804264068549 |
| chr7-126219924-126220425 | 3 | 0.996839280326312 |
| chr7-71084748-71085249 | 8 | 0.983004396067971 |
| chrX-14284515-14285016 | 6 | 0.979335353809204 |
| chr7-117327485-117327986 | 6 | 0.972490101908005 |
| chr14-17706868-17707369 | 2 | 0.830759257931191 |
| chr3-132029518-132030019 | 2 | 0.817778494611158 |
| chr15-53203880-53204381 | 8 | 0.993493681128311 |
| chr20-42346249-42346750 | 8 | 0.938940960481802 |
| chr6-125935635-125936136 | 8 | 0.993493681128311 |
| chr9-4094929-4095430 | 5 | 0.968157999341881 |
| chr4-2401428-2401929 | 6 | 0.964588804510512 |
| chr2-137117520-137118021 | 8 | 0.852618827773705 |
| chr17-67905037-67905538 | 3 | 0.985350513883978 |
| chr14-13592680-13593181 | 8 | 0.972398281839688 |
| chr12-11519287-11519788 | 3 | 0.965784313263398 |
| chr17-22143197-22143698 | 8 | 0.943298033083849 |
| chr9-60679276-60679777 | 5 | 0.991695738285673 |
| chr15-90212605-90213106 | 3 | 0.996050013200828 |
| chr4-44727320-44727821 | 4 | 0.93361506495257 |
| chr14-101026662-101027163 | 1 | 0.892550561390958 |
| chr2-39687293-39687794 | 6 | 0.953364333597679 |
| chr19-38067817-38068318 | 5 | 0.988672237783389 |
| chr4-176885425-176885926 | 3 | 0.986972167603975 |
| chr7-103997601-103998102 | 6 | 0.914168538884634 |

| | | |
|---|---|---|
| chr4-173439705-173440206 | 4 | 0.938426144436945 |
| chr3-113224190-113224691 | 7 | 0.863946871873411 |
| chr17-30985442-30985943 | 6 | 0.957833075016025 |
| chr18-37835519-37836020 | 8 | 0.995232582982055 |
| chr8-114718941-114719442 | 3 | 0.98909379682882 |
| chr4-81997121-81997622 | 6 | 0.921138341881282 |
| chr18-44925453-44925954 | 5 | 0.966429018175685 |
| chr18-35690223-35690724 | 2 | 0.949790207195086 |
| chr15-97629899-97630400 | 8 | 0.905660324292103 |
| chr8-62533999-62534500 | 6 | 0.838629091848795 |
| chr8-98397937-98398438 | 3 | 0.957839183889242 |
| chr4-161617537-161618038 | 8 | 0.91154573039939 |
| chr10-101016521-101017022 | 8 | 0.890098929122051 |
| chr9-45003886-45004387 | 8 | 0.943008918409314 |
| chr2-243062044-243062545 | 3 | 0.932047776315044 |
| chr15-32968438-32968939 | 3 | 0.843104572434826 |
| chr13-43092502-43093003 | 8 | 0.958529954783294 |
| chr9-99528236-99528737 | 3 | 0.876961464502444 |
| chrX-29433005-29433506 | 3 | 0.970298633768412 |
| chr14-34452154-34452655 | 4 | 0.867308334212945 |
| chr13-95095661-95096162 | 3 | 0.978408051986877 |
| chr1-249637966-249638467 | 3 | 0.988843610141029 |
| chr7-115255804-115256305 | 4 | 0.86367714482688 |
| chr4-120151940-120152441 | 8 | 0.990504991215483 |
| chr3-147998334-147998835 | 6 | 0.976047648679667 |
| chr8-120879727-120880228 | 7 | 0.937530938674688 |
| chrX-134566473-134566974 | 7 | 0.856739085879248 |
| chr1-43200037-43200538 | 3 | 0.995450040997645 |
| chr19-55167875-55168376 | 6 | 0.980848717689672 |
| chr9-109005295-109005796 | 3 | 0.984338898232592 |
| chr1-94941861-94942362 | 6 | 0.99182113406123 |
| chr2-174923531-174924032 | 5 | 0.906307575489939 |
| chr10-8414484-8414985 | 5 | 0.924982846714682 |
| chr14-83168065-83168566 | 6 | 0.961423407258591 |
| chr14-32461993-32462494 | 8 | 0.982799743963734 |
| chr3-149989263-149989764 | 6 | 0.99221893660334 |
| chr14-27037315-27037816 | 8 | 0.896845395757503 |
| chr9-17798964-17799465 | 4 | 0.93943545095405 |
| chr20-6070618-6071119 | 5 | 0.96403629870307 |
| chr8-119326337-119326838 | 7 | 0.859759493606781 |
| chr8-113316695-113317196 | 6 | 0.980588243797953 |
| chr15-21119701-21120202 | 7 | 0.91373022792477 |
| chr2-136954217-136954718 | 3 | 0.996737567815445 |

| | | |
|---|---|---|
| chr5-143982989-143983490 | 3 | 0.844893500846911 |
| chr2-79841494-79841995 | 5 | 0.957848436010465 |
| chr10-60820487-60820988 | 8 | 0.868806339718608 |
| chr1-85166983-85167484 | 2 | 0.815037169795388 |
| chr8-52824595-52825096 | 3 | 0.992394798260756 |
| chr14-103216785-103217286 | 3 | 0.889693972956607 |
| chr17-10718214-10718715 | 2 | 0.885457648987278 |
| chr17-60494124-60494625 | 2 | 0.848016087600993 |
| chr15-60106301-60106802 | 3 | 0.984127372551939 |
| chr1-45428531-45429032 | 2 | 0.923630864848473 |
| chr16-6920000-6920501 | 8 | 0.924649702800439 |
| chr6-47300821-47301322 | 3 | 0.995858423754998 |
| chr11-40132175-40132676 | 3 | 0.9966444510962 |
| chr20-35831834-35832335 | 4 | 0.836212603449991 |
| chr1-99382387-99382888 | 2 | 0.934279704258079 |
| chr1-131962316-131962817 | 6 | 0.991206925481043 |
| chr15-35008736-35009237 | 8 | 0.994584354059844 |
| chr9-75127712-75128213 | 2 | 0.881779499468065 |
| chr7-54411589-54412090 | 6 | 0.991276583089416 |
| chr2-53472679-53473180 | 5 | 0.828088373435207 |
| chr5-6746828-6747329 | 3 | 0.867141608353364 |
| chr1-201789784-201790285 | 8 | 0.98942275823941 |
| chr8-46212006-46212507 | 3 | 0.984463568437817 |
| chr4-9320509-9321010 | 3 | 0.824398282331689 |
| chr10-73475139-73475640 | 5 | 0.98766043261992 |
| chr7-43460403-43460904 | 6 | 0.991206925481043 |
| chr2-40324710-40325211 | 6 | 0.925014724769086 |
| chr5-133710958-133711459 | 8 | 0.826195922658849 |
| chr5-143616643-143617144 | 3 | 0.996726913511187 |
| chr5-58064132-58064633 | 8 | 0.871058985394031 |
| chr16-15211208-15211709 | 3 | 0.987868718476225 |
| chr8-55321570-55322071 | 6 | 0.960847835910197 |
| chr5-100610639-100611140 | 3 | 0.907443036911885 |
| chr9-29585330-29585831 | 3 | 0.99593553545041 |
| chr10-60747835-60748336 | 7 | 0.876722556427951 |
| chr2-105134966-105135467 | 3 | 0.99593553545041 |
| chr18-67151130-67151631 | 1 | 0.975511067051073 |
| chr13-91024998-91025499 | 5 | 0.918365512733395 |
| chr14-31750604-31751105 | 8 | 0.994832542749841 |
| chr1-43321355-43321856 | 3 | 0.996740517013679 |
| chr15-68105654-68106155 | 3 | 0.99593553545041 |
| chr18-76220157-76220658 | 6 | 0.985605455000483 |
| chr19-32900585-32901086 | 3 | 0.996919630853497 |

| | | |
|---|---|---|
| chr2-17933893-17934394 | 3 | 0.957569564659344 |
| chr8-123087692-123088193 | 3 | 0.90677086943991 |
| chr9-27168917-27169418 | 6 | 0.937689521986351 |
| chr12-262250-262751 | 3 | 0.987899558291309 |
| chr16-72647438-72647939 | 8 | 0.993493681128311 |
| chr1-167110920-167111421 | 6 | 0.993110362080818 |
| chr2-15544255-15544756 | 6 | 0.991206925481043 |
| chr13-90006867-90007368 | 8 | 0.845629480627113 |
| chr11-18104844-18105345 | 6 | 0.943184333704361 |
| chr3-132790433-132790934 | 3 | 0.99593553545041 |
| chr8-118291814-118292315 | 8 | 0.919055869726613 |
| chr17-3201713-3202214 | 3 | 0.996047031575608 |
| chr2-237105904-237106405 | 4 | 0.903640155473868 |
| chr3-78599363-78599864 | 8 | 0.85577321774117 |
| chr4-72147825-72148326 | 6 | 0.964144110844289 |
| chr1-93909692-93910193 | 8 | 0.984485446326024 |
| chr5-2437901-2438402 | 3 | 0.99593553545041 |
| chr1-109039176-109039677 | 3 | 0.988517043301823 |
| chr2-28043947-28044448 | 4 | 0.864838935411475 |
| chr9-72354866-72355367 | 3 | 0.972375663508681 |
| chr8-62028551-62029052 | 6 | 0.966112588308963 |
| chr13-74036486-74036987 | 3 | 0.993382811408602 |
| chr2-106523430-106523931 | 3 | 0.993719953003487 |
| chr6-42897495-42897996 | 3 | 0.977636971349458 |
| chrX-111400034-111400535 | 5 | 0.869846149650229 |
| chr20-3111231-3111732 | 8 | 0.969447911579601 |
| chr13-59164114-59164615 | 6 | 0.984687886028943 |
| chr16-48822390-48822891 | 6 | 0.985573914112115 |
| chr6-9780577-9781078 | 7 | 0.902317798876521 |
| chr1-151990096-151990597 | 5 | 0.907943532955476 |
| chr3-147210900-147211401 | 8 | 0.87325860489247 |
| chr6-41696191-41696692 | 3 | 0.995443319094357 |
| chr8-54756532-54757033 | 6 | 0.982887695090371 |
| chr6-6887002-6887503 | 5 | 0.99127098694368 |
| chr5-64005755-64006256 | 3 | 0.995190519079887 |
| chr2-165047319-165047820 | 7 | 0.893110743028742 |
| chr2-237233318-237233819 | 6 | 0.991206925481043 |
| chr2-226033127-226033628 | 8 | 0.89478625114912 |
| chr8-56422856-56423357 | 4 | 0.873042535731182 |
| chr17-402363-402864 | 3 | 0.82473305391611 |
| chr17-1168318-1168819 | 8 | 0.85651842770645 |
| chr10-43647486-43647987 | 8 | 0.867492971862291 |
| chr1-134906720-134907221 | 8 | 0.965071431584247 |

| | | |
|---|---|---|
| chr17-27316152-27316653 | 4 | 0.870690424962463 |
| chr3-77646002-77646503 | 8 | 0.939249023386338 |
| chr5-112098137-112098638 | 2 | 0.917114942845337 |
| chr1-121532417-121532918 | 4 | 0.914986830819402 |
| chr12-26364399-26364900 | 1 | 0.862204874692174 |
| chr1-169945165-169945666 | 3 | 0.956003425428269 |
| chr6-124260221-124260722 | 2 | 0.933371935808877 |
| chr1-98115900-98116401 | 3 | 0.99593553545041 |
| chr2-203906140-203906641 | 5 | 0.828881436753049 |
| chr12-24094034-24094535 | 6 | 0.962932775428651 |
| chr11-16032943-16033444 | 5 | 0.985596493141918 |
| chr16-38627978-38628479 | 6 | 0.964327478601243 |
| chr4-173553622-173554123 | 4 | 0.901935048992152 |
| chr6-79060594-79061095 | 6 | 0.904517584539482 |
| chr3-116067059-116067560 | 8 | 0.972295930416976 |
| chr1-108848574-108849075 | 3 | 0.99593553545041 |
| chr8-107251685-107252186 | 6 | 0.897127713723804 |
| chr7-97787857-97788358 | 3 | 0.8675521343568 |
| chr17-52708234-52708735 | 7 | 0.926004914869958 |
| chr7-16747088-16747589 | 6 | 0.981391760166351 |
| chr1-241697487-241697988 | 6 | 0.9733047340525 |
| chr3-159318994-159319495 | 8 | 0.951333852638544 |
| chr12-8070052-8070553 | 2 | 0.854465120903948 |
| chr9-102318977-102319478 | 2 | 0.89630218400357 |
| chr11-43520283-43520784 | 3 | 0.99593553545041 |
| chr11-25779051-25779552 | 4 | 0.943540392431173 |
| chr7-54956644-54957145 | 3 | 0.99593553545041 |
| chr6-94324759-94325260 | 4 | 0.921554112759988 |
| chr10-45149754-45150255 | 2 | 0.882177966697726 |
| chr12-36302312-36302813 | 2 | 0.94302723093517 |
| chr15-15113708-15114209 | 6 | 0.985714045739361 |
| chr10-39208580-39209081 | 8 | 0.864736494415803 |
| chr10-35463066-35463567 | 4 | 0.939914727331546 |
| chr18-50525797-50526298 | 6 | 0.902033401758141 |
| chr3-56121603-56122104 | 6 | 0.991206925481043 |
| chr7-24137981-24138482 | 8 | 0.939345926531545 |
| chr14-9629189-9629690 | 8 | 0.823118135407068 |
| chr5-111822713-111823214 | 3 | 0.993585077309922 |
| chr10-38829664-38830165 | 8 | 0.901490178937223 |
| chr3-157953972-157954473 | 7 | 0.958630912178689 |
| chr4-154575114-154575615 | 4 | 0.823417373789094 |
| chr4-139979899-139980400 | 8 | 0.993493681128311 |
| chr9-79395965-79396466 | 4 | 0.889061314776125 |

| | | |
|---|---|---|
| chr1-165883378-165883879 | 3 | 0.996182367023837 |
| chr5-152867917-152868418 | 3 | 0.997858913145612 |
| chr18-40876639-40877140 | 8 | 0.864991121815159 |
| chr7-42826803-42827304 | 2 | 0.927543256281582 |
| chr7-129237847-129238348 | 8 | 0.993493681128311 |
| chrX-151565933-151566434 | 4 | 0.941137371886687 |
| chr10-7384599-7385100 | 3 | 0.996918187671592 |
| chr4-174082147-174082648 | 5 | 0.907667977136367 |
| chr6-131058951-131059452 | 4 | 0.908789656431069 |
| chr2-110360133-110360634 | 8 | 0.946603080513462 |
| chr2-127046250-127046751 | 6 | 0.909841982794661 |
| chr13-48725369-48725870 | 3 | 0.969018877877567 |
| chr4-162826007-162826508 | 4 | 0.969526868982368 |
| chr7-116451338-116451839 | 1 | 0.859783931578783 |
| chr6-20282995-20283496 | 1 | 0.886721861455367 |
| chr12-12257948-12258449 | 3 | 0.996221664108907 |
| chr11-23827449-23827950 | 1 | 0.889377023361581 |
| chr17-74377663-74378164 | 8 | 0.933164545269101 |
| chr6-8416309-8416810 | 6 | 0.985316911359602 |
| chr8-69830161-69830662 | 8 | 0.94495988150346 |
| chr13-93056662-93057163 | 3 | 0.92529989052237 |
| chr12-37482629-37483130 | 5 | 0.992816446350324 |
| chr9-76303240-76303741 | 8 | 0.993493681128311 |
| chr13-73703715-73704216 | 5 | 0.957647541295506 |
| chr4-129437218-129437719 | 3 | 0.996623575541892 |
| chr5-33135237-33135738 | 6 | 0.983442141932048 |
| chr13-47490720-47491221 | 3 | 0.998184868315157 |
| chr9-88611497-88611998 | 2 | 0.959273755148298 |
| chr8-53790506-53791007 | 6 | 0.842093027241136 |
| chrX-106770745-106771246 | 6 | 0.987652558998252 |
| chr3-112476989-112477490 | 3 | 0.990223278178522 |
| chr13-93667568-93668069 | 2 | 0.919266954516841 |
| chr1-100165038-100165539 | 8 | 0.981610445423393 |
| chrX-40136620-40137121 | 6 | 0.988829268011728 |
| chr10-88378174-88378675 | 2 | 0.942350292323345 |
| chr15-78228664-78229165 | 1 | 0.946268622200749 |
| chr15-45182701-45183202 | 3 | 0.966130320042974 |
| chr1-5174582-5175083 | 6 | 0.90200972912148 |
| chr2-110178042-110178543 | 3 | 0.895300023063294 |
| chr5-9458419-9458920 | 3 | 0.99593553545041 |
| chr5-113682594-113683095 | 8 | 0.937538191565934 |
| chr13-55724854-55725355 | 3 | 0.99593553545041 |
| chr6-101966698-101967199 | 8 | 0.963675940762278 |

| | | |
|---|---|---|
| chr17-34160072-34160573 | 8 | 0.844610397235402 |
| chr4-159957026-159957527 | 7 | 0.971457116892495 |
| chr15-219513-220014 | 3 | 0.993227864001792 |
| chr16-61077490-61077991 | 3 | 0.975109717877255 |
| chr1-78633128-78633629 | 8 | 0.852097019963946 |
| chr12-43169825-43170326 | 8 | 0.904609974110035 |
| chr4-37148629-37149130 | 6 | 0.993225138273038 |
| chr7-103529225-103529726 | 3 | 0.82796612089181 |
| chr8-10461333-10461834 | 5 | 0.870613806961658 |
| chr1-191666874-191667375 | 2 | 0.886128420881701 |
| chr14-30766614-30767115 | 1 | 0.825534525582536 |
| chr1-197526934-197527435 | 3 | 0.909166552066166 |
| chr6-45256261-45256762 | 4 | 0.880334873480437 |
| chr13-71540402-71540903 | 4 | 0.911637839525636 |
| chr13-76819987-76820488 | 5 | 0.969637425130879 |
| chr15-51466892-51467393 | 3 | 0.930870500470593 |
| chr3-14924984-14925485 | 8 | 0.903502522192352 |
| chr12-7125304-7125805 | 1 | 0.93361595891528 |
| chr5-107553609-107554110 | 3 | 0.995303987733676 |
| chr6-50734819-50735320 | 3 | 0.99604143718807 |
| chrX-126523109-126523610 | 6 | 0.993036397251266 |
| chr14-101947801-101948302 | 8 | 0.898413688324873 |
| chr3-7114369-7114870 | 6 | 0.845938613198788 |
| chr7-111757029-111757530 | 8 | 0.878569176010071 |
| chr10-68148318-68148819 | 1 | 0.948562742915898 |
| chr12-6535820-6536321 | 4 | 0.872721349705963 |
| chr1-248893796-248894297 | 3 | 0.933131859111648 |
| chr11-13538108-13538609 | 4 | 0.959259550342869 |
| chr17-65269318-65269819 | 8 | 0.838399210680089 |
| chr13-93749588-93750089 | 8 | 0.990335352107542 |
| chr9-59267417-59267918 | 3 | 0.997187236379835 |
| chr1-256882884-256883385 | 3 | 0.955886414100439 |
| chr1-84080641-84081142 | 4 | 0.899123467639152 |
| chr8-8700229-8700730 | 3 | 0.912671877017743 |
| chr1-159685055-159685556 | 3 | 0.99593553545041 |
| chr7-93773337-93773838 | 6 | 0.819350264937338 |
| chr1-211756286-211756787 | 3 | 0.99930613706984 |
| chr4-130136531-130137032 | 3 | 0.99593553545041 |
| chr17-61196343-61196844 | 7 | 0.828408377133885 |
| chr17-70382332-70382833 | 8 | 0.905944085994368 |
| chr16-51790880-51791381 | 1 | 0.844762991312406 |
| chr17-25147176-25147677 | 6 | 0.992412479201559 |
| chr1-190928605-190929106 | 8 | 0.822839149817603 |

| | | |
|---|---|---|
| chr4-103753333-103753834 | 5 | 0.831287792035459 |
| chr7-31701985-31702486 | 2 | 0.835745997329505 |
| chr8-58474858-58475359 | 8 | 0.988191135464992 |
| chr9-16290738-16291239 | 8 | 0.98529620119264 |
| chr19-57326233-57326734 | 5 | 0.983919978840557 |
| chr9-55978122-55978623 | 3 | 0.994674996734332 |
| chr8-22326846-22327347 | 6 | 0.880834943955296 |
| chr9-42545520-42546021 | 1 | 0.859831118142762 |
| chr6-63576461-63576962 | 8 | 0.991341810303552 |
| chr10-4597495-4597996 | 6 | 0.9925735184735 |
| chr16-31453599-31454100 | 6 | 0.988029262696525 |
| chr9-59371941-59372442 | 8 | 0.892914254185655 |
| chr11-43806394-43806895 | 4 | 0.869619721836622 |
| chr3-158858629-158859130 | 8 | 0.994426166568867 |
| chr8-70984026-70984527 | 2 | 0.841458394471806 |
| chr3-132458470-132458971 | 5 | 0.894694339225545 |
| chr6-129740708-129741209 | 5 | 0.951480882072563 |
| chr6-35538394-35538895 | 3 | 0.989938979309618 |
| chr19-39051092-39051593 | 5 | 0.987355687927997 |
| chr18-40878286-40878787 | 3 | 0.997025260055271 |
| chr12-24759424-24759925 | 8 | 0.931476995467517 |
| chr18-49901471-49901972 | 5 | 0.973878393309492 |
| chr11-46482324-46482825 | 6 | 0.978274951588218 |
| chr14-42638583-42639084 | 8 | 0.90359217895067 |
| chr10-62676909-62677410 | 3 | 0.954352903846261 |
| chr19-7024595-7025096 | 3 | 0.996785546352204 |
| chr15-4624096-4624597 | 3 | 0.857597679212327 |
| chr19-19384020-19384521 | 5 | 0.978666704166629 |
| chr1-218561436-218561937 | 1 | 0.859624027596727 |
| chr14-37157819-37158320 | 3 | 0.99593553545041 |
| chr15-34707742-34708243 | 7 | 0.853748103568348 |
| chr2-171597643-171598144 | 8 | 0.961955047704525 |
| chr2-185950670-185951171 | 5 | 0.983513038227565 |
| chr15-76361819-76362320 | 3 | 0.973324170290077 |
| chr3-117794627-117795128 | 5 | 0.874192395863145 |
| chr8-110352003-110352504 | 8 | 0.883308960502673 |
| chr2-134283303-134283804 | 8 | 0.987766659206114 |
| chr9-82169660-82170161 | 6 | 0.996211185907813 |
| chr2-166235030-166235531 | 6 | 0.939024909565603 |
| chr2-194616333-194616834 | 6 | 0.90308094544987 |
| chr3-37900626-37901127 | 3 | 0.84615055333825 |
| chr16-12030514-12031015 | 6 | 0.918058552028277 |
| chr12-37912674-37913175 | 3 | 0.883014895038185 |

| | | |
|---|---|---|
| chr9-39024623-39025124 | 7 | 0.907213167288334 |
| chr8-48102383-48102884 | 8 | 0.890514165489995 |
| chr11-74851366-74851867 | 6 | 0.99312136012491 |
| chr16-177510-178011 | 3 | 0.997412562856244 |
| chr14-88755852-88756353 | 6 | 0.991070724487121 |
| chrX-135877197-135877698 | 3 | 0.973665389858722 |
| chr3-445765-446266 | 3 | 0.979472800380297 |
| chr1-147042389-147042890 | 7 | 0.941182184038029 |
| chr1-218475339-218475840 | 3 | 0.961445929230228 |
| chr1-96551215-96551716 | 4 | 0.834797115433662 |
| chr7-26523117-26523618 | 6 | 0.89532630197826 |
| chr17-81397525-81398026 | 5 | 0.967800199750309 |
| chr18-31036553-31037054 | 1 | 0.83485921008953 |
| chr1-184041173-184041674 | 2 | 0.83968318292291 |
| chr5-82532676-82533177 | 3 | 0.987446140865604 |
| chr1-136393563-136394064 | 3 | 0.979081986772685 |
| chr3-35847449-35847950 | 6 | 0.958627265579953 |
| chr14-99995619-99996120 | 3 | 0.99593553545041 |
| chr1-171399263-171399764 | 3 | 0.996454101419406 |
| chr14-83242791-83243292 | 6 | 0.833380941148905 |
| chr14-98402979-98403480 | 8 | 0.966369492265877 |
| chr14-21280269-21280770 | 6 | 0.928513105416543 |
| chr3-1000619-1001120 | 3 | 0.919068878931848 |
| chr4-62965493-62965994 | 5 | 0.940300054910645 |
| chr2-147988349-147988850 | 3 | 0.987087450606104 |
| chr13-87297857-87298358 | 3 | 0.992026514691252 |
| chr3-14460136-14460637 | 8 | 0.995150720034934 |
| chr15-99637994-99638495 | 2 | 0.851338567933896 |
| chr14-83994338-83994839 | 3 | 0.991815131946383 |
| chr7-114363711-114364212 | 1 | 0.926626442749575 |
| chr17-54609605-54610106 | 3 | 0.973394967915745 |
| chr13-39161930-39162431 | 8 | 0.993493681128311 |
| chr1-49515299-49515800 | 8 | 0.994376850406724 |
| chr17-79598247-79598748 | 8 | 0.965769284725387 |
| chr16-13030625-13031126 | 6 | 0.991206925481043 |
| chr19-18156206-18156707 | 3 | 0.995374394955516 |
| chr3-121148531-121149032 | 3 | 0.99593553545041 |
| chr1-52006814-52007315 | 6 | 0.993228139258085 |
| chr19-36592494-36592995 | 4 | 0.866905532378782 |
| chr7-132672932-132673433 | 8 | 0.986745057081105 |
| chr6-84587115-84587616 | 4 | 0.970033094537274 |
| chr6-39733030-39733531 | 2 | 0.92296757004484 |
| chr4-19368306-19368807 | 5 | 0.903616793694924 |

| | | |
|---|---|---|
| chr3-50206177-50206678 | 7 | 0.842155837265279 |
| chr10-6853090-6853591 | 2 | 0.927924810246797 |
| chr8-81840980-81841481 | 6 | 0.996263575895951 |
| chr16-76693586-76694087 | 2 | 0.967915536415174 |
| chr7-51711103-51711604 | 5 | 0.885643363266184 |
| chr3-17534428-17534929 | 8 | 0.984734106261241 |
| chr3-69286743-69287244 | 3 | 0.983690084625085 |
| chr1-165847569-165848070 | 3 | 0.896161850557992 |
| chr8-6090988-6091489 | 5 | 0.987737886321472 |
| chr6-46661374-46661875 | 1 | 0.924237465179727 |
| chr1-122468591-122469092 | 4 | 0.855234269094613 |
| chr2-231030717-231031218 | 1 | 0.910409491808585 |
| chr14-62419586-62420087 | 2 | 0.914392570157497 |
| chr17-82388991-82389492 | 6 | 0.890766191852439 |
| chr1-218406815-218407316 | 5 | 0.837936275100437 |
| chr5-24295676-24296177 | 8 | 0.934177819237152 |
| chr2-238141598-238142099 | 8 | 0.991454061514827 |
| chr5-137094704-137095205 | 3 | 0.980152597246798 |
| chr9-50970583-50971084 | 4 | 0.952515163654421 |
| chr5-71933635-71934136 | 8 | 0.967933466527167 |
| chr3-28656418-28656919 | 8 | 0.990794576310252 |
| chr8-68647241-68647742 | 5 | 0.818244853073798 |
| chr3-128670686-128671187 | 4 | 0.942258260030707 |
| chr9-69646042-69646543 | 8 | 0.993493681128311 |
| chr8-42953674-42954175 | 2 | 0.897619534763384 |
| chr19-39896626-39897127 | 3 | 0.938778248048281 |
| chr5-139408240-139408741 | 8 | 0.99436499292547 |
| chr10-15829200-15829701 | 8 | 0.840932960854854 |
| chr5-140562086-140562587 | 3 | 0.99593553545041 |
| chr8-122372320-122372821 | 8 | 0.865083386382658 |
| chr8-64814903-64815404 | 8 | 0.903945238800549 |
| chr7-96704367-96704868 | 6 | 0.959438831513054 |
| chr6-108522006-108522507 | 8 | 0.928482620144329 |
| chr10-65314630-65315131 | 8 | 0.989374602358093 |
| chr19-28594408-28594909 | 8 | 0.993385100146646 |
| chr8-44110696-44111197 | 8 | 0.993493681128311 |
| chr14-38816117-38816618 | 3 | 0.99549792421669 |
| chr9-18100764-18101265 | 3 | 0.826383965608963 |
| chr6-33577195-33577696 | 8 | 0.835659114009813 |
| chr2-81627221-81627722 | 6 | 0.991206925481043 |
| chr3-60389658-60390159 | 1 | 0.947976693425008 |
| chr5-71774453-71774954 | 5 | 0.823821895869155 |
| chr5-160350988-160351489 | 1 | 0.884100056580716 |

| | | |
|---|---|---|
| chr7-30272346-30272847 | 1 | 0.869607250674374 |
| chr5-102156041-102156542 | 8 | 0.933359509593913 |
| chr13-70505007-70505508 | 3 | 0.99039716630123 |
| chr8-118412281-118412782 | 8 | 0.944114243980475 |
| chr10-15877061-15877562 | 8 | 0.994608612539784 |
| chr14-58076220-58076721 | 6 | 0.984498248597368 |
| chr11-32632272-32632773 | 7 | 0.815989232504611 |
| chr5-121492696-121493197 | 8 | 0.992069037781126 |
| chr8-99904541-99905042 | 3 | 0.987240234554931 |
| chrX-128671021-128671522 | 6 | 0.991666034039401 |
| chr10-103643252-103643753 | 8 | 0.963260652473243 |
| chr13-92723245-92723746 | 5 | 0.990671268006156 |
| chr3-89704477-89704978 | 4 | 0.848150147682585 |
| chr11-2673318-2673819 | 3 | 0.99593553545041 |
| chr10-49649803-49650304 | 3 | 0.996488272295582 |
| chr19-32374988-32375489 | 3 | 0.99295902798943 |
| chr1-187295373-187295874 | 3 | 0.847700704081276 |
| chr7-96944701-96945202 | 6 | 0.991206925481043 |
| chr1-43379191-43379692 | 6 | 0.973839753929376 |
| chr1-143505528-143506029 | 1 | 0.821646860739465 |
| chr2-187184321-187184822 | 6 | 0.977925103650182 |
| chr3-162715696-162716197 | 8 | 0.816428707445072 |
| chr2-121291965-121292466 | 5 | 0.82502173389781 |
| chr14-82332478-82332979 | 3 | 0.99593553545041 |
| chr14-1557869-1558370 | 3 | 0.986822178500149 |
| chr4-70502241-70502742 | 6 | 0.978209526890835 |
| chr8-60228053-60228554 | 3 | 0.997221677731803 |
| chr9-81621796-81622297 | 8 | 0.915966508286587 |
| chr8-114391020-114391521 | 8 | 0.921731252320835 |
| chr4-88571423-88571924 | 3 | 0.907827216490554 |
| chr14-19524508-19525009 | 2 | 0.852060624176885 |
| chr6-3674059-3674560 | 5 | 0.955640295480821 |
| chr6-107510200-107510701 | 4 | 0.843652854490596 |
| chr5-2532205-2532706 | 3 | 0.99593553545041 |
| chr3-53220719-53221220 | 3 | 0.99593553545041 |
| chr1-255099888-255100389 | 6 | 0.990655336267055 |
| chr3-24613294-24613795 | 3 | 0.972196954639727 |
| chr10-56953589-56954090 | 3 | 0.996865116705331 |
| chr4-88042767-88043268 | 3 | 0.981035913558275 |
| chr2-101642545-101643046 | 8 | 0.940460712221817 |
| chr20-44380310-44380811 | 2 | 0.833639784584424 |
| chr15-33833685-33834186 | 8 | 0.979972307196901 |
| chr19-50449146-50449647 | 8 | 0.945605057711829 |

| | | |
|---|---|---|
| chr1-256477023-256477524 | 8 | 0.829414292986669 |
| chrX-41891979-41892480 | 3 | 0.99593553545041 |
| chr4-37296432-37296933 | 3 | 0.971744649135668 |
| chr19-36883218-36883719 | 5 | 0.926555448619082 |
| chr13-65030804-65031305 | 8 | 0.950937559413842 |
| chr2-192140730-192141231 | 8 | 0.851038133835581 |
| chr18-15510636-15511137 | 8 | 0.816474026791293 |
| chr1-184606656-184607157 | 3 | 0.995955731005696 |
| chr1-32168471-32168972 | 8 | 0.993493681128311 |
| chr5-102507927-102508428 | 3 | 0.99009329493924 |
| chr4-130339235-130339736 | 8 | 0.87789484561066 |
| chr8-107063080-107063581 | 2 | 0.926431294287823 |
| chr18-61824413-61824914 | 5 | 0.934281224712379 |
| chr12-11077940-11078441 | 5 | 0.989788964738658 |
| chr10-6651070-6651571 | 8 | 0.816615234411308 |
| chr1-180786965-180787466 | 3 | 0.990577141267259 |
| chr1-189221158-189221659 | 2 | 0.887739998388639 |
| chr16-23393198-23393699 | 3 | 0.99534643105137 |
| chr2-102043961-102044462 | 5 | 0.989303683468272 |
| chr18-67916798-67917299 | 8 | 0.822136145139461 |
| chr18-19484573-19485074 | 3 | 0.99593553545041 |
| chr5-66764105-66764606 | 5 | 0.983919978840557 |
| chr1-49126361-49126862 | 6 | 0.977488628817379 |
| chr6-32090769-32091270 | 3 | 0.988726793191445 |
| chr13-77718391-77718892 | 2 | 0.900476266939798 |
| chr4-50552372-50552873 | 5 | 0.879490009434268 |
| chr2-225264436-225264937 | 8 | 0.866641250932488 |
| chr19-42983566-42984067 | 5 | 0.956479057665793 |
| chr4-95851479-95851980 | 3 | 0.935793071100407 |
| chrX-64325738-64326239 | 4 | 0.82317016306034 |
| chr12-14466483-14466984 | 4 | 0.896368180795359 |
| chr14-81325395-81325896 | 8 | 0.840628017364737 |
| chr3-117720722-117721223 | 7 | 0.963807471783675 |
| chr5-83913354-83913855 | 6 | 0.93637774763011 |
| chr11-71517722-71518223 | 8 | 0.993493681128311 |
| chr1-21204673-21205174 | 8 | 0.941080433540385 |
| chr2-10109299-10109800 | 3 | 0.996327151981704 |
| chr3-86207104-86207605 | 2 | 0.878165408486091 |
| chr5-118756562-118757063 | 7 | 0.846259772524507 |
| chr10-57741298-57741799 | 5 | 0.983919978840557 |
| chr5-123766872-123767373 | 8 | 0.915239166710514 |
| chr15-89723725-89724226 | 3 | 0.843390389630196 |
| chr17-28856995-28857496 | 2 | 0.912093533991493 |

| | | |
|---|---|---|
| chr2-159434083-159434584 | 3 | 0.99689603148413 |
| chr16-6000237-6000738 | 6 | 0.993194557331912 |
| chr6-71574606-71575107 | 8 | 0.919151968237383 |
| chr3-152592933-152593434 | 3 | 0.993336423121894 |
| chr13-102405181-102405682 | 6 | 0.983303834023665 |
| chr4-57491107-57491608 | 3 | 0.818679942917414 |
| chr1-7348205-7348706 | 8 | 0.834028044652161 |
| chr20-8930522-8931023 | 5 | 0.973138743631072 |
| chr2-164652445-164652946 | 8 | 0.874572790144592 |
| chr17-78341071-78341572 | 5 | 0.954491429219076 |
| chr3-66196991-66197492 | 5 | 0.973439802877108 |
| chr2-235838057-235838558 | 8 | 0.995170173605043 |
| chrX-16171531-16172032 | 3 | 0.965122564527932 |
| chr16-63518744-63519245 | 5 | 0.960206515236318 |
| chr9-61842926-61843427 | 3 | 0.998053551427915 |
| chr11-51316670-51317171 | 3 | 0.981236379056888 |
| chr16-54866366-54866867 | 3 | 0.986765263871361 |
| chr5-3187908-3188409 | 8 | 0.822788636871862 |
| chr6-39188070-39188571 | 6 | 0.876907366543469 |
| chrX-31071665-31072166 | 5 | 0.983919978840557 |
| chr10-64706870-64707371 | 6 | 0.901771588035699 |
| chr8-101150118-101150619 | 6 | 0.927888001993894 |
| chr2-29016331-29016832 | 8 | 0.99482998129982 |
| chr13-103908360-103908861 | 4 | 0.882591600371903 |
| chr18-72933766-72934267 | 8 | 0.994925520594554 |
| chr16-44913901-44914402 | 1 | 0.859862844004061 |
| chr4-151824085-151824586 | 4 | 0.90200726878411 |
| chr3-123290478-123290979 | 7 | 0.937682026341207 |
| chrX-134943284-134943785 | 8 | 0.994834092646108 |
| chr7-39188449-39188950 | 2 | 0.851618335357104 |
| chr8-102958415-102958916 | 3 | 0.97310738743154 |
| chr20-20028860-20029361 | 5 | 0.871350431662737 |
| chr4-49665004-49665505 | 3 | 0.986189299503124 |
| chr2-171481043-171481544 | 8 | 0.988834005281023 |
| chr2-53212231-53212732 | 3 | 0.99593553545041 |
| chr1-174302432-174302933 | 3 | 0.996844361773958 |
| chr13-97521549-97522050 | 8 | 0.993493681128311 |
| chr14-75523170-75523671 | 3 | 0.997217598094481 |
| chr15-38745157-38745658 | 3 | 0.996036367020219 |
| chr17-38338371-38338872 | 4 | 0.899061018100242 |
| chr6-94455867-94456368 | 5 | 0.816812400110682 |
| chr7-109879608-109880109 | 2 | 0.902156633872889 |
| chr13-31783302-31783803 | 3 | 0.890259603142949 |

| | | |
|---|---|---|
| chr4-181744799-181745300 | 6 | 0.905841081080902 |
| chr1-222453186-222453687 | 1 | 0.861047227804179 |
| chr18-62374934-62375435 | 8 | 0.984357039015282 |
| chr5-86494036-86494537 | 6 | 0.989284911459703 |
| chr7-55199864-55200365 | 8 | 0.959808489386324 |
| chr4-65680295-65680796 | 6 | 0.957431787454463 |
| chr2-160375098-160375599 | 3 | 0.997041117839259 |
| chr3-129246810-129247311 | 3 | 0.996281754246812 |
| chr10-50557377-50557878 | 4 | 0.861440047563359 |
| chr7-91697199-91697700 | 3 | 0.956827912111837 |
| chr2-243888786-243889287 | 6 | 0.992219913037387 |
| chr4-15344884-15345385 | 8 | 0.961386932621959 |
| chr15-30891123-30891624 | 2 | 0.829504033772306 |
| chr4-13062680-13063181 | 8 | 0.923262981618962 |
| chr10-44970872-44971373 | 8 | 0.991818565165092 |
| chr20-45801666-45802167 | 3 | 0.990966868024534 |
| chr19-53400148-53400649 | 8 | 0.984412996505131 |
| chr16-32523530-32524031 | 5 | 0.971876994235983 |
| chr1-239376320-239376821 | 8 | 0.944032406978699 |
| chr3-7472644-7473145 | 3 | 0.928730462239476 |
| chr8-15125506-15126007 | 5 | 0.991749805330763 |
| chr14-40016425-40016926 | 3 | 0.875389561636751 |
| chr1-194830966-194831467 | 5 | 0.88421025208913 |
| chr11-58014846-58015347 | 1 | 0.863187590932163 |
| chr1-154930071-154930572 | 5 | 0.98260505944286 |
| chr11-31968830-31969331 | 3 | 0.997912151729429 |
| chr10-64538674-64539175 | 7 | 0.914782428797832 |
| chr14-6386242-6386743 | 8 | 0.880023435487322 |
| chr2-162829941-162830442 | 3 | 0.99665613555611 |
| chr16-77722087-77722588 | 4 | 0.879112558160772 |
| chr1-215481391-215481892 | 3 | 0.995597232597422 |
| chr11-50397197-50397698 | 3 | 0.996855464397222 |
| chr9-7443864-7444365 | 6 | 0.991206925481043 |
| chr4-175774173-175774674 | 8 | 0.825710666987949 |
| chr1-100326614-100327115 | 4 | 0.865221974712973 |
| chr13-95364059-95364560 | 3 | 0.996580337834397 |
| chr4-80477093-80477594 | 6 | 0.991219908154868 |
| chr3-104711517-104712018 | 8 | 0.859985392684846 |
| chr19-15223-15724 | 6 | 0.815801074424695 |
| chr1-40279200-40279701 | 1 | 0.841318679504519 |
| chr16-17803304-17803805 | 5 | 0.990095619341524 |
| chr19-30570937-30571438 | 4 | 0.832464233323398 |
| chr10-63621812-63622313 | 6 | 0.982953078799906 |

| | | |
|---|---|---|
| chrX-15916877-15917378 | 3 | 0.83116569349511 |
| chr6-32149094-32149595 | 5 | 0.984731682571546 |
| chr4-60770186-60770687 | 8 | 0.891877077370738 |
| chr7-18300340-18300841 | 4 | 0.971888562060124 |
| chr14-84946103-84946604 | 3 | 0.996250401852696 |
| chr16-8326866-8327367 | 8 | 0.995231914228476 |
| chr17-62149799-62150300 | 3 | 0.982818995027033 |
| chr12-21993124-21993625 | 2 | 0.927024901120181 |
| chr1-178827818-178828319 | 6 | 0.975063998512214 |
| chr10-26271580-26272081 | 3 | 0.99593553545041 |
| chr5-120275587-120276088 | 3 | 0.822001907946077 |
| chr15-6586836-6587337 | 6 | 0.986388698090803 |
| chr12-38284829-38285330 | 8 | 0.913753478711762 |
| chr7-50965854-50966355 | 3 | 0.829970951284314 |
| chr2-144536994-144537495 | 3 | 0.971887894128314 |
| chr7-116987273-116987774 | 4 | 0.902591256573261 |
| chr9-64497898-64498399 | 8 | 0.874227186126575 |
| chr1-170287635-170288136 | 3 | 0.995884879761859 |
| chr7-126590692-126591193 | 3 | 0.994422064901807 |
| chr5-78465257-78465758 | 8 | 0.989510521402507 |
| chr9-4532554-4533055 | 3 | 0.99593553545041 |
| chr6-99339889-99340390 | 2 | 0.944458520221565 |
| chr13-78521216-78521717 | 5 | 0.891055705492833 |
| chr4-61438625-61439126 | 6 | 0.98927078315381 |
| chr13-8096503-8097004 | 3 | 0.981437937460425 |
| chr7-119578811-119579312 | 6 | 0.974980697315377 |
| chr13-50472172-50472673 | 5 | 0.983685294347808 |
| chr5-57246488-57246989 | 8 | 0.993493681128311 |
| chr12-26202329-26202830 | 3 | 0.957447989342057 |
| chr19-51350163-51350664 | 6 | 0.871676794184985 |
| chr3-9451267-9451768 | 3 | 0.836756766003372 |
| chr1-77271871-77272372 | 6 | 0.927062637933918 |
| chr5-21404801-21405302 | 3 | 0.93335020065331 |
| chr8-53023770-53024271 | 8 | 0.99471650766406 |
| chr8-65685206-65685707 | 5 | 0.849044461304222 |
| chr15-87018462-87018963 | 3 | 0.99593553545041 |
| chr18-41168357-41168858 | 1 | 0.939694362846073 |
| chr18-75449394-75449895 | 3 | 0.989951844470789 |
| chr16-51421246-51421747 | 8 | 0.870337572515919 |
| chr9-11278495-11278996 | 6 | 0.859799730949952 |
| chr5-151585296-151585797 | 5 | 0.987687450514175 |
| chr2-137850919-137851420 | 8 | 0.921429663189822 |
| chr10-40253780-40254281 | 6 | 0.947247376591727 |

| | | |
|---|---|---|
| chr8-106582215-106582716 | 8 | 0.899003583889977 |
| chr3-104878912-104879413 | 3 | 0.996376673985161 |
| chr14-78576722-78577223 | 3 | 0.958493515568532 |
| chr18-42661978-42662479 | 6 | 0.995868439374935 |
| chr1-240773085-240773586 | 2 | 0.857563748941979 |
| chr1-92109787-92110288 | 1 | 0.911643213861473 |
| chr1-196292847-196293348 | 5 | 0.927786938489627 |
| chr6-58405385-58405886 | 5 | 0.952949601899341 |
| chr15-69812248-69812749 | 1 | 0.892331878308199 |
| chr3-115314900-115315401 | 8 | 0.993547047602088 |
| chr15-9835113-9835614 | 3 | 0.990239646860684 |
| chr12-14797441-14797942 | 5 | 0.988121897286662 |
| chr1-4574160-4574661 | 6 | 0.992294026368259 |
| chr15-4973048-4973549 | 3 | 0.99593553545041 |
| chr5-157184415-157184916 | 4 | 0.890151812132507 |
| chr13-63485492-63485993 | 6 | 0.899546679062785 |
| chr14-91731001-91731502 | 8 | 0.995243878640541 |
| chrX-150745456-150745957 | 8 | 0.993493681128311 |
| chr18-61241653-61242154 | 8 | 0.917719953047033 |
| chr6-87915531-87916032 | 8 | 0.931220190274117 |
| chr7-8614594-8615095 | 4 | 0.851298149967364 |
| chr6-111934613-111935114 | 3 | 0.99517737980011 |
| chr5-59106495-59106996 | 2 | 0.935986137007972 |
| chr10-35748420-35748921 | 2 | 0.924331923002812 |
| chr7-94531371-94531872 | 6 | 0.992848151280515 |
| chr2-20947597-20948098 | 2 | 0.837731045603032 |
| chr4-9581432-9581933 | 7 | 0.843409040892112 |
| chr4-154207172-154207673 | 8 | 0.915169100521981 |
| chr9-69730634-69731135 | 3 | 0.994578754569023 |
| chrX-151730329-151730830 | 6 | 0.99319517759353 |
| chr3-53689111-53689612 | 4 | 0.820963406426822 |
| chr3-60175416-60175917 | 1 | 0.872047663416609 |
| chr4-135721605-135722106 | 8 | 0.993493681128311 |
| chr18-29211615-29212116 | 5 | 0.91845785772759 |
| chr11-29075876-29076377 | 5 | 0.889213924052605 |
| chr17-54193010-54193511 | 5 | 0.987443818275542 |
| chr16-43244404-43244905 | 5 | 0.957781267228905 |
| chr12-3858365-3858866 | 6 | 0.991206925481043 |
| chr2-145714166-145714667 | 3 | 0.990569691512789 |
| chr12-38339222-38339723 | 8 | 0.903175708720469 |
| chr18-72532772-72533273 | 8 | 0.862679527908814 |
| chr14-81128191-81128692 | 8 | 0.852987274240159 |
| chr12-41043484-41043985 | 7 | 0.875597741635281 |

| | | |
|---|---|---|
| chr11-164830-165331 | 6 | 0.946684469184864 |
| chr1-10754974-10755475 | 3 | 0.997227886400447 |
| chr19-39771815-39772316 | 2 | 0.860567264074309 |
| chr1-106864732-106865233 | 4 | 0.818533534457851 |
| chr15-22732439-22732940 | 3 | 0.994769729811805 |
| chr16-74860660-74861161 | 4 | 0.838698777500142 |
| chr4-115970111-115970612 | 5 | 0.883698043223109 |
| chr9-52139195-52139696 | 3 | 0.99593553545041 |
| chr19-28859162-28859663 | 3 | 0.997177734762705 |
| chr12-943746-944247 | 3 | 0.989358545050487 |
| chr8-51815427-51815928 | 3 | 0.993120688242283 |
| chr2-39745754-39746255 | 3 | 0.819006492916651 |
| chr13-50630976-50631477 | 3 | 0.857254047957592 |
| chr18-82025954-82026455 | 5 | 0.944817005313307 |
| chr18-26312810-26313311 | 8 | 0.910695353487705 |
| chr1-91008806-91009307 | 3 | 0.987694583700802 |
| chrX-152210217-152210718 | 3 | 0.992785883055569 |
| chr7-58611490-58611991 | 3 | 0.99422121004857 |
| chr4-60713056-60713557 | 8 | 0.99129826389929 |
| chr1-177141813-177142314 | 8 | 0.968474683946067 |
| chr3-148553314-148553815 | 8 | 0.951169621234833 |
| chr2-110798099-110798600 | 1 | 0.923837993503101 |
| chr2-242340199-242340700 | 6 | 0.990862504736536 |
| chr14-76607077-76607578 | 2 | 0.910769127328687 |
| chr2-173199429-173199930 | 8 | 0.930881978282207 |
| chr3-133807891-133808392 | 6 | 0.974552273136132 |
| chr1-149806507-149807008 | 3 | 0.984799894858663 |
| chr2-141202297-141202798 | 6 | 0.951330779058176 |
| chr6-117153619-117154120 | 8 | 0.993493681128311 |
| chr18-21830794-21831295 | 8 | 0.92688673458058 |
| chr17-84866128-84866629 | 3 | 0.972261923301413 |
| chr17-19604668-19605169 | 3 | 0.986711824716247 |
| chr18-21099700-21100201 | 6 | 0.980972206693889 |
| chr6-16988380-16988881 | 3 | 0.987607836176822 |
| chr2-242369935-242370436 | 1 | 0.953914210946065 |
| chr1-87776345-87776846 | 1 | 0.836846548521145 |
| chr13-55712082-55712583 | 8 | 0.99507903785339 |
| chr3-125041340-125041841 | 1 | 0.901168366699068 |
| chr6-28382200-28382701 | 6 | 0.988213213092036 |
| chr11-35678107-35678608 | 8 | 0.994994896059724 |
| chr8-116552605-116553106 | 6 | 0.992590457625773 |
| chr18-67150342-67150843 | 1 | 0.857982279782371 |
| chr14-6725711-6726212 | 8 | 0.994711811339885 |

| | | |
|---|---|---|
| chr5-146996436-146996937 | 8 | 0.980730491078551 |
| chr2-122703473-122703974 | 8 | 0.923253031795711 |
| chr16-73016260-73016761 | 3 | 0.99593553545041 |
| chrX-100118403-100118904 | 1 | 0.83565746536099 |
| chr10-44545298-44545799 | 8 | 0.961039448538517 |
| chr9-87729229-87729730 | 6 | 0.952142191747506 |
| chr4-99863367-99863868 | 3 | 0.960317481386315 |
| chr3-163010165-163010666 | 8 | 0.832411264487157 |
| chr15-74015854-74016355 | 3 | 0.936954200961472 |
| chr14-66571405-66571906 | 3 | 0.994231726892649 |
| chr4-9439177-9439678 | 2 | 0.90225888361881 |
| chr3-134812269-134812770 | 8 | 0.981583577849844 |
| chr5-33547969-33548470 | 8 | 0.856261004761806 |
| chr2-153926456-153926957 | 6 | 0.992824940994439 |
| chr11-62307202-62307703 | 6 | 0.963933012394115 |
| chr3-8830717-8831218 | 8 | 0.866104173347001 |
| chr7-85085507-85086008 | 8 | 0.848943851183634 |
| chr2-183377338-183377839 | 8 | 0.857589868854018 |
| chr6-39248945-39249446 | 6 | 0.963535712464387 |
| chr3-107211569-107212070 | 3 | 0.995627513757265 |
| chr1-232237334-232237835 | 3 | 0.99593553545041 |
| chr1-103620598-103621099 | 6 | 0.943156368610764 |
| chr1-239038191-239038692 | 3 | 0.968612728176108 |
| chr5-145446499-145447000 | 7 | 0.839711117365831 |
| chr5-21335950-21336451 | 3 | 0.936461678784172 |
| chr5-146155168-146155669 | 8 | 0.842545687151124 |
| chr10-19835064-19835565 | 8 | 0.99077151987179 |
| chr5-97038387-97038888 | 5 | 0.896425773751093 |
| chr14-49073136-49073637 | 3 | 0.994359687468769 |
| chr17-80945265-80945766 | 6 | 0.968604203610844 |
| chr7-53028302-53028803 | 2 | 0.957478804204768 |
| chr4-57199576-57200077 | 6 | 0.981234502194079 |
| chr9-37061472-37061973 | 2 | 0.952180704706487 |
| chr1-80709774-80710275 | 7 | 0.864480980479416 |
| chr6-117951344-117951845 | 6 | 0.930127656349171 |
| chr7-65089839-65090340 | 1 | 0.901460599123799 |
| chr14-97996291-97996792 | 8 | 0.992250822001742 |
| chr1-246515567-246516068 | 2 | 0.960772437142609 |
| chr1-239810801-239811302 | 8 | 0.993493681128311 |
| chr17-61784581-61785082 | 3 | 0.951453427561739 |
| chr5-2938174-2938675 | 3 | 0.995711583084896 |
| chr19-10136564-10137065 | 8 | 0.940667575466298 |
| chr16-21276925-21277426 | 7 | 0.944240643360846 |

| | | |
|---|---|---|
| chr2-6923600-6924101 | 5 | 0.915013746794551 |
| chr4-144367284-144367785 | 6 | 0.834922752047957 |
| chr2-139951523-139952024 | 3 | 0.996783183274627 |
| chr2-50388472-50388973 | 6 | 0.915166879063135 |
| chr1-88565623-88566124 | 8 | 0.991115813879552 |
| chr3-75949977-75950478 | 3 | 0.978348515793154 |
| chr11-47279966-47280467 | 6 | 0.992104555162191 |
| chr9-17986195-17986696 | 6 | 0.953528989047147 |
| chrX-35090360-35090861 | 5 | 0.887711956380155 |
| chr3-29999611-30000112 | 8 | 0.923584437365066 |
| chr3-111728957-111729458 | 6 | 0.955142897332381 |
| chr15-24982138-24982639 | 8 | 0.986933940948958 |
| chr20-25211046-25211547 | 5 | 0.985822522696782 |
| chr9-56997182-56997683 | 8 | 0.906600582696211 |
| chr7-79081224-79081725 | 3 | 0.969328207598692 |
| chr10-50478414-50478915 | 8 | 0.993461303389523 |
| chr4-151356741-151357242 | 1 | 0.913447105680102 |
| chr1-236040298-236040799 | 6 | 0.990012471706501 |
| chr16-68950537-68951038 | 4 | 0.904764598761301 |
| chr5-113070530-113071031 | 6 | 0.958381163374532 |
| chr18-46752383-46752884 | 6 | 0.997288370880902 |
| chr19-12058091-12058592 | 3 | 0.868206613912568 |
| chr18-60592092-60592593 | 5 | 0.988010655513418 |
| chr8-117052852-117053353 | 3 | 0.99593553545041 |
| chr6-44086542-44087043 | 8 | 0.968540636902099 |
| chr13-3238145-3238646 | 6 | 0.991206925481043 |
| chr8-52701912-52702413 | 3 | 0.897489421041426 |
| chr2-241669227-241669728 | 3 | 0.996551261921726 |
| chr14-5647679-5648180 | 1 | 0.958398795466706 |
| chr3-159940932-159941433 | 3 | 0.995749803619722 |
| chr3-18640925-18641426 | 6 | 0.981290132301585 |
| chr19-44316434-44316935 | 5 | 0.896130411598378 |
| chr13-81754445-81754946 | 8 | 0.989370920657919 |
| chr5-69851414-69851915 | 3 | 0.994796530642161 |
| chrX-4848701-4849202 | 8 | 0.834325491764698 |
| chr13-48229651-48230152 | 6 | 0.876616634385374 |
| chr3-147957467-147957968 | 3 | 0.99593553545041 |
| chr6-40731190-40731691 | 8 | 0.872165270529623 |
| chr1-85283662-85284163 | 1 | 0.926143788867232 |
| chr13-38302863-38303364 | 3 | 0.98042064758154 |
| chr6-101690805-101691306 | 8 | 0.964911690953666 |
| chr4-90238192-90238693 | 5 | 0.903822912622392 |
| chrX-78753094-78753595 | 6 | 0.983679057970096 |

| | | |
|---|---|---|
| chr6-96886296-96886797 | 8 | 0.914614155915591 |
| chr10-21992460-21992961 | 5 | 0.958920698787925 |
| chr5-40308535-40309036 | 6 | 0.991206925481043 |
| chr4-76479318-76479819 | 8 | 0.855419596956013 |
| chr17-45917625-45918126 | 3 | 0.91417400714676 |
| chr10-73886573-73887074 | 3 | 0.918174922281455 |
| chr9-104257731-104258232 | 8 | 0.959220663432971 |
| chr2-657710-658211 | 5 | 0.870873641752071 |
| chr6-23234045-23234546 | 8 | 0.892650009118166 |
| chr20-49559739-49560240 | 3 | 0.953026648653092 |
| chr1-9484304-9484805 | 3 | 0.99593553545041 |
| chr4-110234246-110234747 | 8 | 0.993493681128311 |
| chr9-66255178-66255679 | 8 | 0.993493681128311 |
| chr6-123103048-123103549 | 3 | 0.996090644617532 |
| chr6-137353766-137354267 | 5 | 0.977871861194234 |
| chr3-104488946-104489447 | 5 | 0.983919978840557 |
| chr13-83240688-83241189 | 8 | 0.994117291900686 |
| chr2-174192660-174193161 | 5 | 0.975887737776337 |
| chr19-55709166-55709667 | 3 | 0.973789239374427 |
| chr3-44629202-44629703 | 6 | 0.987820845788914 |
| chr1-106572907-106573408 | 3 | 0.996609536548854 |
| chr1-199239411-199239912 | 8 | 0.861842923844425 |
| chr4-148637441-148637942 | 7 | 0.927550072359429 |
| chr7-74040230-74040731 | 3 | 0.963707623163664 |
| chrX-75811282-75811783 | 3 | 0.99593553545041 |
| chr12-31972865-31973366 | 8 | 0.963175447624867 |
| chr7-56812108-56812609 | 5 | 0.845662399017603 |
| chr10-94189343-94189844 | 8 | 0.965402519896212 |
| chr2-188067203-188067704 | 6 | 0.96305437921303 |
| chr4-18357154-18357655 | 5 | 0.956887874669723 |
| chr13-20506813-20507314 | 5 | 0.946921550217936 |
| chr1-62087087-62087588 | 8 | 0.868843152436305 |
| chr6-72120524-72121025 | 3 | 0.991804797016749 |
| chr17-5559010-5559511 | 3 | 0.99593553545041 |
| chr14-26847638-26848139 | 8 | 0.993493681128311 |
| chr8-116466068-116466569 | 8 | 0.928521750613188 |
| chr3-88213403-88213904 | 8 | 0.89886911398727 |
| chr2-111848202-111848703 | 6 | 0.991206925481043 |
| chr6-39132014-39132515 | 8 | 0.82293532197827 |
| chr8-123630942-123631443 | 8 | 0.939976037575245 |
| chr14-97173239-97173740 | 6 | 0.824168750018501 |
| chr11-15666493-15666994 | 8 | 0.92200113086139 |
| chr5-29789944-29790445 | 8 | 0.919297453521659 |

| | | |
|---|---|---|
| chr15-24465014-24465515 | 1 | 0.926389728075427 |
| chr8-6707381-6707882 | 3 | 0.99593553545041 |
| chr2-200130967-200131468 | 8 | 0.964115891949228 |
| chr1-118096475-118096976 | 1 | 0.867884666877164 |
| chr2-82347123-82347624 | 7 | 0.817182062723859 |
| chr17-64037979-64038480 | 8 | 0.96879956711415 |
| chr1-258437742-258438243 | 5 | 0.93074334849931 |
| chr1-80727213-80727714 | 3 | 0.997402763385638 |
| chr6-140473799-140474300 | 3 | 0.841488209920996 |
| chr19-36406602-36407103 | 6 | 0.861656479823119 |
| chr7-90903562-90904063 | 5 | 0.988409014588037 |
| chr13-29698600-29699101 | 2 | 0.836012917542894 |
| chr2-232379916-232380417 | 3 | 0.992678631808281 |
| chr5-108747549-108748050 | 6 | 0.985045299136747 |
| chr4-6073381-6073882 | 3 | 0.9890893840833 |
| chr5-8089388-8089889 | 5 | 0.847520189196657 |
| chr11-67193267-67193768 | 6 | 0.895338727060431 |
| chr11-71597660-71598161 | 6 | 0.881000660798235 |
| chr4-40447805-40448306 | 6 | 0.881845411381203 |
| chr8-103912654-103913155 | 3 | 0.986295095031222 |
| chr17-69718597-69719098 | 8 | 0.976233890993223 |
| chr3-151967751-151968252 | 8 | 0.943563053528393 |
| chr1-40376376-40376877 | 5 | 0.837428486557694 |
| chr1-236059078-236059579 | 3 | 0.996770959801594 |
| chr18-47054241-47054742 | 8 | 0.994808084810861 |
| chr18-83105709-83106210 | 3 | 0.973124690123143 |
| chr7-102178098-102178599 | 3 | 0.944165404074125 |
| chr11-12705336-12705837 | 5 | 0.963738720415276 |
| chr19-21239461-21239962 | 8 | 0.930835800089304 |
| chr3-14006633-14007134 | 8 | 0.889054981986913 |
| chr2-30169366-30169867 | 7 | 0.864021412781602 |
| chr15-45800419-45800920 | 6 | 0.98665860184869 |
| chr2-51393045-51393546 | 3 | 0.996878736509615 |
| chr6-118160255-118160756 | 5 | 0.914121713585811 |
| chr14-82682875-82683376 | 3 | 0.924550755129478 |
| chr2-28078437-28078938 | 8 | 0.903001769743988 |
| chr10-39300437-39300938 | 7 | 0.920176333892829 |
| chr13-25610672-25611173 | 3 | 0.942450996417394 |
| chr1-13833479-13833980 | 1 | 0.957034249400248 |
| chr9-111404376-111404877 | 8 | 0.993493681128311 |
| chr8-98268724-98269225 | 5 | 0.991484638435694 |
| chr8-20033685-20034186 | 4 | 0.837373870627592 |
| chr1-186515049-186515550 | 4 | 0.930533753177343 |

| | | |
|---|---|---|
| chr2-215664178-215664679 | 3 | 0.996016220947232 |
| chr5-69573248-69573749 | 3 | 0.99593553545041 |
| chr9-31246518-31247019 | 6 | 0.956229257555024 |
| chr14-78512105-78512606 | 8 | 0.993493681128311 |
| chr2-220643273-220643774 | 5 | 0.991969047194226 |
| chr2-90869935-90870436 | 6 | 0.996149796397165 |
| chr19-5724597-5725098 | 3 | 0.942280969209459 |
| chr17-16055602-16056103 | 3 | 0.911081394419365 |
| chr14-79303260-79303761 | 7 | 0.904917975569954 |
| chr2-231950595-231951096 | 1 | 0.855699837383869 |
| chr3-120130981-120131482 | 6 | 0.946446337517272 |
| chr13-32747461-32747962 | 1 | 0.920511150112943 |
| chr18-59292807-59293308 | 1 | 0.877865611484707 |
| chr11-50607643-50608144 | 7 | 0.822638899349946 |
| chr1-94454240-94454741 | 6 | 0.97867596413076 |
| chr1-184968762-184969263 | 8 | 0.978625969515794 |
| chr8-101154339-101154840 | 6 | 0.992570293296734 |
| chr18-59128358-59128859 | 6 | 0.992308121502758 |
| chr11-72113791-72114292 | 3 | 0.894423784319408 |
| chr2-30384849-30385350 | 3 | 0.996621916837236 |
| chr9-41632992-41633493 | 3 | 0.996573182886535 |
| chr3-122486220-122486721 | 1 | 0.869899107270847 |
| chr3-136334841-136335342 | 1 | 0.815215259710285 |
| chr20-10399940-10400441 | 5 | 0.988674313178982 |
| chr10-76789514-76790015 | 8 | 0.959026695790963 |
| chrX-68392609-68393110 | 8 | 0.986143531802064 |
| chr5-82729655-82730156 | 3 | 0.996248005883041 |
| chr5-5494136-5494637 | 3 | 0.993507328834002 |
| chr6-75035678-75036179 | 3 | 0.93954365800498 |
| chr18-6741176-6741677 | 3 | 0.989102907471267 |
| chr10-25293798-25294299 | 3 | 0.997773321459897 |
| chr13-50367246-50367747 | 6 | 0.960772351760874 |
| chr7-114311866-114312367 | 8 | 0.934120404369593 |
| chr2-20787372-20787873 | 8 | 0.849739828650461 |
| chr1-9051120-9051621 | 3 | 0.99593553545041 |
| chr5-100116773-100117274 | 3 | 0.988634545919351 |
| chr14-56888635-56889136 | 8 | 0.865081673251573 |
| chr4-133748786-133749287 | 6 | 0.914208700219763 |
| chr14-7300257-7300758 | 6 | 0.992357724242289 |
| chr1-215123373-215123874 | 3 | 0.99593553545041 |
| chr1-24591733-24592234 | 4 | 0.885938370009376 |
| chr8-2665392-2665893 | 4 | 0.821706385852467 |
| chr13-10061568-10062069 | 3 | 0.99593553545041 |

| | | |
|---|---|---|
| chr11-56240318-56240819 | 8 | 0.993493681128311 |
| chr12-25408699-25409200 | 8 | 0.993493681128311 |
| chr1-227524599-227525100 | 5 | 0.962952143612508 |
| chr2-215762972-215763473 | 8 | 0.953238491580698 |
| chr5-118831644-118832145 | 3 | 0.82201583850165 |
| chr15-31264721-31265222 | 8 | 0.91676285770415 |
| chr11-75445958-75446459 | 6 | 0.930939126765968 |
| chr1-22196292-22196793 | 8 | 0.933936333653879 |
| chr13-30245902-30246403 | 8 | 0.886071618735294 |
| chr7-130710454-130710955 | 8 | 0.995088349048109 |
| chr15-98411077-98411578 | 4 | 0.972442064394892 |
| chr18-23674748-23675249 | 8 | 0.87849620068867 |
| chr5-57101963-57102464 | 5 | 0.950204084645158 |
| chr1-88606415-88606916 | 6 | 0.968038468014935 |
| chr4-19155479-19155980 | 3 | 0.99593553545041 |
| chr14-42372648-42373149 | 8 | 0.85359112180125 |
| chr4-92425288-92425789 | 6 | 0.94560955054727 |
| chr1-158182339-158182840 | 6 | 0.993002551996752 |
| chr3-87814777-87815278 | 3 | 0.998014219506909 |
| chr13-61298338-61298839 | 3 | 0.99593553545041 |
| chr10-27547503-27548004 | 3 | 0.996892249314123 |
| chr2-24019412-24019913 | 2 | 0.92631933304799 |
| chr19-7654220-7654721 | 3 | 0.99593553545041 |
| chr6-5526116-5526617 | 2 | 0.95753489063926 |
| chr5-46215779-46216280 | 8 | 0.929160618485019 |
| chr11-65634657-65635158 | 6 | 0.926190160985276 |
| chr6-94587982-94588483 | 8 | 0.993493681128311 |
| chr10-58833294-58833795 | 3 | 0.973686569482088 |
| chr3-85785964-85786465 | 6 | 0.924458651765206 |
| chrX-26615895-26616396 | 8 | 0.843342045257751 |
| chr2-194587607-194588108 | 3 | 0.995353555289805 |
| chr9-4838356-4838857 | 6 | 0.991206925481043 |
| chr4-65973287-65973788 | 8 | 0.989907572414047 |
| chrX-12193862-12194363 | 2 | 0.955623887254831 |
| chr4-13616810-13617311 | 3 | 0.994898616726165 |
| chr15-52907418-52907919 | 6 | 0.987375334335991 |
| chr1-170026244-170026745 | 5 | 0.991273196051225 |
| chr8-43971947-43972448 | 8 | 0.979230473396999 |
| chr8-8080188-8080689 | 3 | 0.99593553545041 |
| chr7-45140327-45140828 | 8 | 0.936546270641102 |
| chr8-86840246-86840747 | 3 | 0.941784036289509 |
| chr13-95781438-95781939 | 5 | 0.960571469601409 |
| chr19-51224489-51224990 | 8 | 0.877330703611146 |

| | | |
|---|---|---|
| chr15-84121078-84121579 | 3 | 0.993631487816059 |
| chr13-23791810-23792311 | 8 | 0.993493681128311 |
| chr7-71905840-71906341 | 6 | 0.99068495909299 |
| chr20-24440755-24441256 | 2 | 0.818777385964396 |
| chr14-95632413-95632914 | 6 | 0.954385252692445 |
| chr4-178347781-178348282 | 6 | 0.993086516281334 |
| chr2-207028934-207029435 | 3 | 0.875443130924794 |
| chr2-196964298-196964799 | 8 | 0.962011458996953 |
| chr20-5288137-5288638 | 5 | 0.973426371789235 |
| chr5-85917745-85918246 | 1 | 0.91412609946102 |
| chr5-159300804-159301305 | 8 | 0.915982069838616 |
| chr2-151284976-151285477 | 8 | 0.913495931342434 |
| chr14-7291330-7291831 | 8 | 0.949998651192006 |
| chr14-3212366-3212867 | 8 | 0.892556137847509 |
| chr9-58388092-58388593 | 6 | 0.99215399075921 |
| chr17-29052028-29052529 | 8 | 0.909529446267958 |
| chr16-57815561-57816062 | 8 | 0.985632463572391 |
| chr2-164881883-164882384 | 4 | 0.97159569789223 |
| chr18-39923972-39924473 | 6 | 0.971894871995097 |
| chr15-12600999-12601500 | 8 | 0.857066840846366 |
| chr7-126671892-126672393 | 5 | 0.991233625833302 |
| chr20-42281408-42281909 | 8 | 0.865333939483218 |
| chr4-175591520-175592021 | 3 | 0.996355382284399 |
| chr17-56894078-56894579 | 3 | 0.99593553545041 |
| chr9-94653154-94653655 | 5 | 0.983919978840557 |
| chr1-70065261-70065762 | 3 | 0.99593553545041 |
| chr1-90478765-90479266 | 4 | 0.969164617739367 |
| chr20-29259627-29260128 | 5 | 0.986117554247537 |
| chr4-28252389-28252890 | 6 | 0.991206925481043 |
| chr6-96465944-96466445 | 5 | 0.923003988496599 |
| chr5-55027803-55028304 | 6 | 0.991206925481043 |
| chrX-116530421-116530922 | 3 | 0.99593553545041 |
| chr5-63617327-63617828 | 8 | 0.989200151199667 |
| chr18-12603277-12603778 | 5 | 0.958705745816459 |
| chr1-59024062-59024563 | 3 | 0.986723133406696 |
| chr9-4781883-4782384 | 6 | 0.871087684320994 |
| chr11-32832066-32832567 | 5 | 0.991515042040004 |
| chr1-144735681-144736182 | 4 | 0.882735546555358 |
| chr7-58505058-58505559 | 5 | 0.986229886139787 |
| chr1-141691473-141691974 | 6 | 0.991206925481043 |
| chr13-48930596-48931097 | 5 | 0.983919978840557 |
| chr1-122683713-122684214 | 6 | 0.890072434027565 |
| chr13-41891432-41891933 | 4 | 0.944777352585045 |

| | | |
|---|---|---|
| chr8-117441055-117441556 | 3 | 0.995935535450410 |
| chr6-22504772-22505273 | 8 | 0.96556843863981 |
| chr1-103215200-103215701 | 6 | 0.940640363376555 |
| chr1-123591075-123591576 | 8 | 0.993493681128311 |
| chr1-82592402-82592903 | 3 | 0.87593621469629 |
| chr12-13004933-13005434 | 6 | 0.866086460198993 |
| chr14-62729129-62729630 | 6 | 0.991206925481043 |
| chr4-82124625-82125126 | 6 | 0.993464804676328 |
| chr13-102732867-102733368 | 4 | 0.858290362989053 |
| chr17-83314188-83314689 | 8 | 0.879685098342139 |
| chr9-60149374-60149875 | 4 | 0.957065733269025 |
| chr16-55566369-55566870 | 3 | 0.853068928637624 |
| chr18-44834630-44835131 | 1 | 0.894922344162332 |
| chr7-94150880-94151381 | 5 | 0.987007991645894 |
| chr5-144681012-144681513 | 4 | 0.969058731545873 |
| chr4-133043086-133043587 | 8 | 0.8537074347749 |
| chr19-36154406-36154907 | 3 | 0.974084876499341 |
| chr13-42711840-42712341 | 3 | 0.878830652801535 |
| chr1-174521450-174521951 | 3 | 0.987586246583528 |
| chr19-9491222-9491723 | 6 | 0.821223167301071 |
| chr8-69975296-69975797 | 3 | 0.994096077100591 |
| chr20-3277461-3277962 | 3 | 0.9517948883178 |
| chr5-150362978-150363479 | 4 | 0.960962792743714 |
| chr14-69149746-69150247 | 8 | 0.994902783830972 |
| chr10-60322428-60322929 | 5 | 0.982580548237176 |
| chr5-33082759-33083260 | 3 | 0.957051228527766 |
| chr19-34423755-34424256 | 8 | 0.984602709707178 |
| chr4-125841872-125842373 | 2 | 0.939391452837684 |
| chr3-22601947-22602448 | 5 | 0.930081127533067 |
| chr10-28611061-28611562 | 6 | 0.989235683048753 |
| chr6-71849888-71850389 | 3 | 0.988878620916818 |
| chr13-70617739-70618240 | 6 | 0.990955195848775 |
| chr4-133680800-133681301 | 8 | 0.975946773952603 |
| chr2-110943310-110943811 | 8 | 0.963339392847072 |
| chr9-50759671-50760172 | 2 | 0.852020395623038 |
| chr7-86939610-86940111 | 5 | 0.986978880454724 |
| chr14-11949513-11950014 | 3 | 0.99593553545041 |
| chr7-53970445-53970946 | 3 | 0.982193859985014 |
| chr13-36419025-36419526 | 3 | 0.896345602744057 |
| chr3-149473165-149473666 | 6 | 0.899420425187726 |
| chr17-37849622-37850123 | 6 | 0.995445129649368 |
| chr13-91510792-91511293 | 4 | 0.907187132290477 |
| chr17-31974455-31974956 | 5 | 0.992001390144415 |

| | | |
|---|---|---|
| chr14-21751213-21751714 | 5 | 0.98210211971649 |
| chr4-109139914-109140415 | 3 | 0.994691788234247 |
| chr13-39804247-39804748 | 4 | 0.965333951269265 |
| chr10-4965228-4965729 | 8 | 0.96421405825364 |
| chr8-120701980-120702481 | 2 | 0.920602231025676 |
| chr3-149893260-149893761 | 4 | 0.919234810822906 |
| chr5-20747777-20748278 | 3 | 0.981707580080169 |
| chr6-137764092-137764593 | 3 | 0.972962090473919 |
| chr6-91184582-91185083 | 8 | 0.985175503059545 |
| chr12-23142704-23143205 | 3 | 0.996760948525419 |
| chr19-18312558-18313059 | 5 | 0.982086936574575 |
| chr4-151921650-151922151 | 3 | 0.909422544832125 |
| chr15-16680479-16680980 | 6 | 0.89339271961113 |
| chr2-135752327-135752828 | 8 | 0.831888732449899 |
| chr6-44717108-44717609 | 4 | 0.901363815081004 |
| chr14-40860522-40861023 | 5 | 0.975127172236608 |
| chr14-74636867-74637368 | 5 | 0.987669804887942 |
| chr6-133714506-133715007 | 6 | 0.969632955354852 |
| chr1-173989196-173989697 | 3 | 0.99593553545041 |
| chr4-80767005-80767506 | 2 | 0.925343280044718 |
| chr7-117269537-117270038 | 3 | 0.884858608156224 |
| chr17-63444685-63445186 | 5 | 0.954069988636539 |
| chr13-97062961-97063462 | 3 | 0.955074072783468 |
| chr8-62757896-62758397 | 8 | 0.916667644642443 |
| chr20-8334389-8334890 | 1 | 0.821880780003472 |
| chr3-163390015-163390516 | 5 | 0.985350871699586 |
| chr14-33933215-33933716 | 5 | 0.952609047019658 |
| chr2-124253858-124254359 | 8 | 0.993493681128311 |
| chr1-170196859-170197360 | 7 | 0.842766344439008 |
| chr10-66800938-66801439 | 8 | 0.994661083245695 |
| chr15-49577236-49577737 | 3 | 0.870735392217513 |
| chr2-121690131-121690632 | 5 | 0.925446047274286 |
| chr7-40178208-40178709 | 6 | 0.991206925481043 |
| chr10-35902302-35902803 | 7 | 0.886826477802356 |
| chr10-42113931-42114432 | 8 | 0.826482982102493 |
| chr1-252244382-252244883 | 3 | 0.996062588204552 |
| chrX-35506921-35507422 | 1 | 0.8848321808951 |
| chr5-20526322-20526823 | 5 | 0.968693471568722 |
| chr11-31599621-31600122 | 7 | 0.934527660418674 |
| chr13-103161577-103162078 | 8 | 0.993288957763184 |
| chr15-4623284-4623785 | 3 | 0.987873910886015 |
| chr7-27506567-27507068 | 5 | 0.987842485765861 |
| chr14-63978026-63978527 | 3 | 0.996535956405822 |

| | | |
|---|---|---|
| chr8-41719184-41719685 | 3 | 0.954315609763623 |
| chrX-32564370-32564871 | 8 | 0.983756331028408 |
| chr13-77932829-77933330 | 4 | 0.856911311942654 |
| chr16-26725579-26726080 | 6 | 0.991799855543663 |
| chr11-35534178-35534679 | 5 | 0.937248449078906 |
| chr9-18952830-18953331 | 3 | 0.99593553545041 |
| chr5-19866442-19866943 | 6 | 0.982995251493829 |
| chr15-37273834-37274335 | 8 | 0.986601246784911 |
| chr4-108575479-108575980 | 3 | 0.98141851529577 |
| chr15-39449780-39450281 | 4 | 0.837236384062148 |
| chr2-209945148-209945649 | 2 | 0.899414458877856 |
| chr19-14313597-14314098 | 4 | 0.867546083083214 |
| chr12-5200553-5201054 | 6 | 0.988945515803836 |
| chr13-75026882-75027383 | 5 | 0.929122545637709 |
| chr4-9659267-9659768 | 5 | 0.922243731681385 |
| chr10-64073794-64074295 | 5 | 0.916736423365382 |
| chr1-3040799-3041300 | 8 | 0.908210964463687 |
| chr2-136094133-136094634 | 6 | 0.881135762677982 |
| chr6-88763285-88763786 | 4 | 0.947279613076776 |
| chr5-41647794-41648295 | 3 | 0.954641913653104 |
| chr6-58381123-58381624 | 6 | 0.987087896442776 |
| chr19-19979166-19979667 | 6 | 0.892436802642262 |
| chr3-149553975-149554476 | 2 | 0.85563652994722 |
| chr10-66800417-66800918 | 8 | 0.991847157213728 |
| chr18-14264766-14265267 | 3 | 0.958477909398133 |
| chr16-31992557-31993058 | 5 | 0.960085098623076 |
| chr7-134514069-134514570 | 3 | 0.975050224990148 |
| chr18-7173034-7173535 | 5 | 0.991211582673838 |
| chr19-13740549-13741050 | 2 | 0.837862526445436 |
| chr5-147610017-147610518 | 8 | 0.878242334717231 |
| chr14-92845567-92846068 | 6 | 0.991852949074357 |
| chr4-119310569-119311070 | 8 | 0.912573556110588 |
| chr14-76261365-76261866 | 5 | 0.834009714343131 |
| chr8-113551904-113552405 | 6 | 0.985524577014944 |
| chr10-27564987-27565488 | 5 | 0.989020929280765 |
| chr2-187974790-187975291 | 5 | 0.991741258943485 |
| chr18-63916632-63917133 | 3 | 0.940035812450235 |
| chr4-9321707-9322208 | 6 | 0.993118898620847 |
| chr3-158453157-158453658 | 8 | 0.866542237987578 |
| chr13-64306486-64306987 | 5 | 0.882976809117361 |
| chr1-233936363-233936864 | 7 | 0.918021171177564 |
| chr17-22390578-22391079 | 5 | 0.991528961563049 |
| chrX-117567877-117568378 | 6 | 0.98657088166979 |

| | | |
|---|---|---|
| chr15-48397163-48397664 | 8 | 0.912293595847301 |
| chr5-110495029-110495530 | 6 | 0.974845788893557 |
| chr11-18190533-18191034 | 5 | 0.917912728587024 |
| chr3-22362858-22363359 | 8 | 0.848532627773329 |
| chr2-121738891-121739392 | 3 | 0.995658500243068 |
| chr17-10853927-10854428 | 5 | 0.815810393276549 |
| chr10-40689503-40690004 | 3 | 0.993078832646977 |
| chr6-31101816-31102317 | 8 | 0.931440196797773 |
| chr18-72151292-72151793 | 8 | 0.823280452540602 |
| chr2-29000700-29001201 | 8 | 0.867089976905249 |
| chr2-136055432-136055933 | 4 | 0.862707733829553 |
| chr10-96822537-96823038 | 8 | 0.971103440619919 |
| chr2-25011381-25011882 | 8 | 0.995246369147472 |
| chr15-642385-642886 | 2 | 0.937188492286618 |
| chr2-35706633-35707134 | 5 | 0.893782571409148 |
| chr5-165998129-165998630 | 6 | 0.917250260796241 |
| chr1-171405942-171406443 | 3 | 0.975016862084672 |
| chr7-77113308-77113809 | 5 | 0.913476280835327 |
| chr2-199068843-199069344 | 3 | 0.957176239033275 |
| chr1-31944823-31945324 | 3 | 0.995121555430929 |
| chr15-93056823-93057324 | 6 | 0.992976357591605 |
| chr14-35887076-35887577 | 2 | 0.862832013599525 |
| chr3-160532783-160533284 | 6 | 0.86988170290108 |
| chr16-73416308-73416809 | 6 | 0.937206924999345 |
| chr16-51344681-51345182 | 8 | 0.825336923637703 |
| chr18-14957704-14958205 | 6 | 0.978628789657264 |
| chr1-170534273-170534774 | 4 | 0.81579711958211 |
| chr3-125518464-125518965 | 5 | 0.968990759326063 |
| chr13-83572587-83573088 | 3 | 0.995012334904379 |
| chr7-82574612-82575113 | 8 | 0.993493681128311 |
| chr6-134607732-134608233 | 6 | 0.85754341499167 |
| chr6-15463796-15464297 | 4 | 0.969252897760632 |
| chr8-89400088-89400589 | 8 | 0.991048449753537 |
| chr3-119946265-119946766 | 5 | 0.988500991112312 |
| chr14-41477574-41478075 | 3 | 0.994062864216931 |
| chr3-90247011-90247512 | 8 | 0.918160137115212 |
| chr8-37125139-37125640 | 5 | 0.904277897884655 |
| chr12-40803689-40804190 | 4 | 0.953408080867256 |
| chr1-12564553-12565054 | 5 | 0.954801426715669 |
| chr17-51679231-51679732 | 2 | 0.955800323580541 |
| chr1-235088558-235089059 | 1 | 0.930616096988749 |
| chr2-187775369-187775870 | 3 | 0.981050840483353 |
| chr7-35266430-35266931 | 6 | 0.991206925481043 |

| | | |
|---|---|---|
| chr18-27130290-27130791 | 8 | 0.993428195206512 |
| chrX-37225931-37226432 | 3 | 0.996677613298887 |
| chr4-171074956-171075457 | 6 | 0.991878722639659 |
| chr2-145588797-145589298 | 6 | 0.991847583424952 |
| chr15-4793554-4794055 | 4 | 0.816761118995801 |
| chr9-24505128-24505629 | 4 | 0.896661268644437 |
| chr7-21913084-21913585 | 6 | 0.90608422091847 |
| chr3-152138024-152138525 | 8 | 0.845962879759972 |
| chr3-65152189-65152690 | 5 | 0.942413284356249 |
| chr1-2129472-2129973 | 8 | 0.954663486308488 |
| chr7-107156187-107156688 | 5 | 0.871292103109954 |
| chr10-98608478-98608979 | 8 | 0.991376142406292 |
| chr13-104844891-104845392 | 8 | 0.940133435392725 |
| chr3-85487933-85488434 | 6 | 0.972937033685302 |
| chr16-125212-125713 | 8 | 0.888601456398728 |
| chr12-11505226-11505727 | 8 | 0.993493681128311 |
| chr1-123090345-123090846 | 3 | 0.975546578557766 |
| chr16-68480555-68481056 | 2 | 0.929263947877599 |
| chr17-26851581-26852082 | 8 | 0.886355655723387 |
| chr9-107734955-107735456 | 2 | 0.887393774298729 |
| chr3-163759342-163759843 | 8 | 0.981257056604278 |
| chr5-114051639-114052140 | 8 | 0.990402192888524 |
| chr1-155640444-155640945 | 5 | 0.933036464306224 |
| chr5-24636437-24636938 | 3 | 0.998084556957411 |
| chr1-172877181-172877682 | 7 | 0.921668681074998 |
| chr8-59636462-59636963 | 8 | 0.893431313655003 |
| chr1-194762960-194763461 | 5 | 0.977079577035985 |
| chr9-61566652-61567153 | 4 | 0.866482209164968 |
| chr2-214069770-214070271 | 5 | 0.981296245459117 |
| chr4-104016028-104016529 | 7 | 0.900231816462594 |
| chr20-19059804-19060305 | 5 | 0.896631395280566 |
| chr11-16326590-16327091 | 3 | 0.990022479533388 |
| chr9-74395331-74395832 | 8 | 0.938051448965096 |
| chr1-20515970-20516471 | 6 | 0.912009360096161 |
| chr1-243269070-243269571 | 4 | 0.954796447356417 |
| chrX-50024372-50024873 | 6 | 0.988704048699626 |
| chr19-14686336-14686837 | 6 | 0.983722289466875 |
| chr8-77731197-77731698 | 8 | 0.962639171163304 |
| chr13-63600149-63600650 | 2 | 0.832364667463844 |
| chr9-63719750-63720251 | 6 | 0.981629798109028 |
| chr1-255337842-255338343 | 7 | 0.855091249567197 |
| chr3-126412619-126413120 | 8 | 0.892584425176997 |
| chr7-102250264-102250765 | 3 | 0.996878737140735 |

| | | |
|---|---|---|
| chr1-6853220-6853721 | 2 | 0.920222271986541 |
| chr17-49555354-49555855 | 2 | 0.900074962943046 |
| chr6-51732395-51732896 | 3 | 0.984488792819768 |
| chr16-74784906-74785407 | 6 | 0.982316964477675 |
| chr7-28848284-28848785 | 8 | 0.930318851262477 |
| chr13-45431202-45431703 | 1 | 0.851051191859943 |
| chr1-91206076-91206577 | 8 | 0.919148115428636 |
| chr8-10189256-10189757 | 3 | 0.994965745444061 |
| chr2-106295986-106296487 | 3 | 0.995201261003603 |
| chr17-46365745-46366246 | 3 | 0.923143028480032 |
| chr14-32945135-32945636 | 8 | 0.927351450323545 |
| chr12-15708664-15709165 | 5 | 0.988519612973255 |
| chr14-104191646-104192147 | 6 | 0.879755009175701 |
| chr17-64309215-64309716 | 6 | 0.996311927188382 |
| chr4-33541776-33542277 | 5 | 0.923342866150905 |
| chr8-69078567-69079068 | 6 | 0.982568473715442 |
| chr6-27402072-27402573 | 3 | 0.969642388181495 |
| chr15-56603206-56603707 | 5 | 0.989786565057046 |
| chr2-139113259-139113760 | 8 | 0.847654318197751 |
| chr7-107587386-107587887 | 8 | 0.989319847177828 |
| chr2-49090090-49090591 | 6 | 0.881022025033172 |
| chr8-122068068-122068569 | 8 | 0.953461133852316 |
| chr6-98460645-98461146 | 8 | 0.914263119215859 |
| chr15-96600955-96601456 | 3 | 0.996898502271943 |
| chr17-31692959-31693460 | 5 | 0.862843981088339 |
| chr14-93688402-93688903 | 4 | 0.956257084081129 |
| chr13-105755515-105756016 | 7 | 0.915654969633683 |
| chr2-80741821-80742322 | 3 | 0.996832327058927 |
| chr14-79404477-79404978 | 6 | 0.981419234950145 |
| chr16-17440664-17441165 | 4 | 0.970483843860683 |
| chr11-74757285-74757786 | 6 | 0.850033348500044 |
| chr4-39683612-39684113 | 3 | 0.978044321083922 |
| chr8-38226473-38226974 | 4 | 0.890175139805679 |
| chr8-66144742-66145243 | 8 | 0.984548475912102 |
| chr19-21408248-21408749 | 6 | 0.99016617382673 |
| chr1-133636218-133636719 | 2 | 0.908675877528741 |
| chr3-61189861-61190362 | 3 | 0.986325938362775 |
| chr15-87735125-87735626 | 6 | 0.966383827623141 |
| chr8-123074884-123075385 | 3 | 0.937534125409074 |
| chr2-34333003-34333504 | 2 | 0.908141172569394 |
| chr9-48493312-48493813 | 3 | 0.994681012183812 |
| chr1-114536499-114537000 | 6 | 0.976969172240063 |
| chr2-92238959-92239460 | 5 | 0.992773674291437 |

| | | |
|---|---|---|
| chr18-71389867-71390368 | 6 | 0.993210220977896 |
| chr19-17570783-17571284 | 3 | 0.996889791511486 |
| chrX-53229565-53230066 | 3 | 0.99593553545041 |
| chr4-10128307-10128808 | 8 | 0.931512065237189 |
| chr2-113330280-113330781 | 1 | 0.907887762146298 |
| chr3-133047363-133047864 | 5 | 0.983767718257003 |
| chr7-111190551-111191052 | 6 | 0.934232227116455 |
| chr8-78293256-78293757 | 4 | 0.817990398695815 |
| chr13-44915808-44916309 | 5 | 0.931742961243496 |
| chrX-130767831-130768332 | 3 | 0.848049810129117 |
| chr15-78500944-78501445 | 6 | 0.991206925481043 |
| chr17-46587338-46587839 | 8 | 0.885368122809253 |
| chr12-28947233-28947734 | 4 | 0.854494561077273 |
| chr10-67685883-67686384 | 8 | 0.867849755668056 |
| chr2-143132427-143132928 | 7 | 0.957992242504498 |
| chr11-80079870-80080371 | 8 | 0.993493681128311 |
| chr10-35816823-35817324 | 5 | 0.857581993872525 |
| chr17-45279084-45279585 | 3 | 0.967897957481732 |
| chr20-19373800-19374301 | 5 | 0.978384960002478 |
| chr14-79396969-79397470 | 7 | 0.914203518274238 |
| chr1-5834643-5835144 | 8 | 0.954106648576155 |
| chr7-17283886-17284387 | 3 | 0.918789063236981 |
| chr14-62918476-62918977 | 3 | 0.997247494270889 |
| chr14-6154148-6154649 | 7 | 0.888966766502994 |
| chr15-36248728-36249229 | 8 | 0.926079733935119 |
| chr1-156516488-156516989 | 2 | 0.944388857326994 |
| chr1-190827774-190828275 | 8 | 0.960041745006976 |
| chr7-129861042-129861543 | 3 | 0.935634931845687 |
| chr5-111707925-111708426 | 8 | 0.964455148076796 |
| chr17-35109636-35110137 | 1 | 0.919662421011823 |
| chr10-10387597-10388098 | 4 | 0.855379938730597 |
| chr7-5600850-5601351 | 6 | 0.982807064462274 |
| chrX-76648579-76649080 | 3 | 0.982823893287189 |
| chr5-54163574-54164075 | 3 | 0.971647870205258 |
| chr17-24690790-24691291 | 2 | 0.87330145772423 |
| chr19-127540-128041 | 3 | 0.99593553545041 |
| chr15-98337528-98338029 | 8 | 0.961868107778302 |
| chr1-142325753-142326254 | 8 | 0.993493681128311 |
| chr14-83240930-83241431 | 7 | 0.864985212713753 |
| chr8-26531455-26531956 | 8 | 0.962703271516717 |
| chr11-68096410-68096911 | 5 | 0.985489622265516 |
| chr1-41394638-41395139 | 8 | 0.960639467907386 |
| chr9-69013978-69014479 | 1 | 0.898539349027734 |

| | | |
|---|---|---|
| chr1-154987942-154988443 | 3 | 0.973162514888713 |
| chr1-46468427-46468928 | 7 | 0.89104895942606 |
| chr8-80845465-80845966 | 3 | 0.979217428917516 |
| chr1-126153272-126153773 | 6 | 0.897063448308673 |
| chr19-5219098-5219599 | 8 | 0.98563701334585 |
| chr8-73025609-73026110 | 8 | 0.930114156884382 |
| chr3-150325791-150326292 | 1 | 0.854204707077581 |
| chr19-31485963-31486464 | 1 | 0.902105569886578 |
| chr1-120292302-120292803 | 6 | 0.99097843020027 |
| chr3-159640240-159640741 | 6 | 0.880693360075314 |
| chr3-157703798-157704299 | 6 | 0.989910252402077 |
| chr1-225378390-225378891 | 6 | 0.84195085266604 |
| chr8-29982106-29982607 | 3 | 0.965941579581635 |
| chr5-8212952-8213453 | 3 | 0.841432630211331 |
| chr16-68863457-68863958 | 1 | 0.84876127438259 |
| chr7-20465142-20465643 | 8 | 0.993493681128311 |
| chr19-46844625-46845126 | 1 | 0.831565033059997 |
| chr10-55567820-55568321 | 2 | 0.831886305322657 |
| chr7-124144004-124144505 | 8 | 0.94858143999246 |
| chr2-122724854-122725355 | 2 | 0.895745374844181 |
| chr4-2089593-2090094 | 3 | 0.834606194009092 |
| chr4-88216382-88216883 | 3 | 0.99593553545041 |
| chr20-36150323-36150824 | 5 | 0.986862251906479 |
| chr17-19875216-19875717 | 5 | 0.983399996643676 |
| chr10-54985766-54986267 | 2 | 0.904015577631738 |
| chr15-76378980-76379481 | 8 | 0.92434664677709 |
| chr3-79392260-79392761 | 1 | 0.943002142857355 |
| chr1-236749884-236750385 | 3 | 0.980230605356196 |
| chr6-45898631-45899132 | 8 | 0.89533025832212 |
| chr7-95165973-95166474 | 8 | 0.909353307145423 |
| chr15-98790885-98791386 | 7 | 0.833538035348455 |
| chr13-75635118-75635619 | 8 | 0.994402155833666 |
| chr1-85023510-85024011 | 3 | 0.95986299551824 |
| chr4-116873279-116873780 | 8 | 0.992973364648099 |
| chr17-36909751-36910252 | 3 | 0.924055019256278 |
| chr4-121821272-121821773 | 5 | 0.935357208664741 |
| chr4-101024567-101025068 | 3 | 0.99593553545041 |
| chr17-56721751-56722252 | 3 | 0.976560370798664 |
| chr7-124777715-124778216 | 6 | 0.993184833611141 |
| chr12-8045860-8046361 | 5 | 0.923516789493662 |
| chr9-106099076-106099577 | 3 | 0.996458599406201 |
| chr9-88531447-88531948 | 2 | 0.867206328683746 |
| chr20-12510562-12511063 | 3 | 0.995248160797613 |

| | | |
|---|---|---|
| chr6-3533327-3533828 | 8 | 0.985286467212762 |
| chr3-165523478-165523979 | 3 | 0.991481355865153 |
| chr10-26897286-26897787 | 3 | 0.99593553545041 |
| chr3-152641183-152641684 | 1 | 0.883549759932263 |
| chr18-67160901-67161402 | 8 | 0.872872150544059 |
| chr5-67642893-67643394 | 6 | 0.959695404874532 |
| chr12-12540081-12540582 | 6 | 0.95734542274856 |
| chr7-115035804-115036305 | 3 | 0.996520407437809 |
| chr1-206715787-206716288 | 1 | 0.925945934423581 |
| chr15-15596389-15596890 | 8 | 0.993493681128311 |
| chr5-164303915-164304416 | 4 | 0.923588557662683 |
| chr17-15582445-15582946 | 8 | 0.993493681128311 |
| chr9-3447059-3447560 | 5 | 0.896446269433841 |
| chr3-48415318-48415819 | 2 | 0.864079592156103 |
| chr20-32184707-32185208 | 2 | 0.864400616617779 |
| chr6-26757166-26757667 | 1 | 0.898256979517632 |
| chr4-133820545-133821046 | 2 | 0.840183190527377 |
| chr2-226865147-226865648 | 3 | 0.99593553545041 |
| chr19-12217629-12218130 | 8 | 0.887883092352085 |
| chr4-77674866-77675367 | 3 | 0.995946209728609 |
| chr3-55080183-55080684 | 5 | 0.978764146194793 |
| chr5-43303824-43304325 | 6 | 0.976553100452288 |
| chr6-140139727-140140228 | 3 | 0.887994862140014 |
| chr3-120617772-120618273 | 2 | 0.90038896076017 |
| chrX-135120996-135121497 | 6 | 0.991206925481043 |
| chr18-600401-600902 | 5 | 0.96818082073367 |
| chr20-37056845-37057346 | 3 | 0.99593553545041 |
| chr13-90178602-90179103 | 5 | 0.949635049609332 |
| chr10-55324664-55325165 | 8 | 0.83487568419015 |
| chr9-89058717-89059218 | 8 | 0.846516479076287 |
| chr9-86282339-86282840 | 2 | 0.852920904397423 |
| chr8-90495273-90495774 | 8 | 0.923915201711123 |
| chr11-44522438-44522939 | 3 | 0.989329072028003 |
| chr10-96312277-96312778 | 8 | 0.946285898811901 |
| chr2-43429465-43429966 | 3 | 0.994152432930071 |
| chr17-32814796-32815297 | 7 | 0.894662607378909 |
| chr15-72851790-72852291 | 3 | 0.958762112592454 |
| chr18-77310615-77311116 | 5 | 0.984382413210502 |
| chr20-44680177-44680678 | 2 | 0.817464728160086 |
| chr15-68588784-68589285 | 6 | 0.977579645905608 |
| chr2-235067038-235067539 | 5 | 0.931824018257284 |
| chr11-5234205-5234706 | 3 | 0.984769060758446 |
| chr4-92342987-92343488 | 3 | 0.920296749343774 |

| | | |
|---|---|---|
| chr10-80131301-80131802 | 4 | 0.931089247453443 |
| chr3-79623750-79624251 | 6 | 0.991363620327941 |
| chr8-96751831-96752332 | 2 | 0.940890892910013 |
| chr4-163890769-163891270 | 3 | 0.988400763066701 |
| chr2-180059912-180060413 | 1 | 0.890479082205539 |
| chr19-5131174-5131675 | 3 | 0.99593553545041 |
| chr8-101072919-101073420 | 4 | 0.899420218674554 |
| chr19-24066528-24067029 | 8 | 0.864505029665189 |
| chr1-177855686-177856187 | 5 | 0.8986363241637 |
| chr8-23342236-23342737 | 6 | 0.992925033506098 |
| chr15-47986199-47986700 | 3 | 0.995666878674608 |
| chr13-64635279-64635780 | 1 | 0.847498157776851 |
| chr20-35259314-35259815 | 6 | 0.842773026690922 |
| chr5-17962574-17963075 | 3 | 0.988303930615724 |
| chr4-177229832-177230333 | 7 | 0.870026058072005 |
| chr7-37539186-37539687 | 3 | 0.975982216499465 |
| chr20-25290406-25290907 | 3 | 0.995087256862556 |
| chr9-14006959-14007460 | 8 | 0.978789669453075 |
| chr11-20924375-20924876 | 6 | 0.906798278071459 |
| chr2-174552244-174552745 | 4 | 0.827628835460929 |
| chr6-13055198-13055699 | 8 | 0.976527621578923 |
| chr1-236538642-236539143 | 6 | 0.848339780015297 |
| chr15-20094874-20095375 | 8 | 0.958755055773804 |
| chr2-25304433-25304934 | 1 | 0.948761505978958 |
| chr2-193085818-193086319 | 5 | 0.959659814640399 |
| chr19-31607951-31608452 | 3 | 0.961337823008739 |
| chr7-27023584-27024085 | 3 | 0.995934885021401 |
| chr13-97756391-97756892 | 4 | 0.942717555167448 |
| chr14-82675019-82675520 | 6 | 0.972342417788926 |
| chr3-119717382-119717883 | 6 | 0.988793153016861 |
| chr3-107442104-107442605 | 1 | 0.859981006724453 |
| chr20-39655573-39656074 | 6 | 0.991206925481043 |
| chr12-40945098-40945599 | 4 | 0.871926769760094 |
| chr8-6359290-6359791 | 6 | 0.979048380054461 |
| chr5-144006012-144006513 | 4 | 0.915709856857478 |
| chr1-187149622-187150123 | 3 | 0.928790033376274 |
| chr8-55482510-55483011 | 5 | 0.967771652140558 |
| chr4-46802541-46803042 | 3 | 0.99593553545041 |
| chr11-66212380-66212881 | 6 | 0.940905074965943 |
| chr16-45395179-45395680 | 6 | 0.945606023274461 |
| chrX-111590607-111591108 | 8 | 0.911037359505309 |
| chrX-58950195-58950696 | 3 | 0.996813359494031 |
| chr1-167916144-167916645 | 5 | 0.926857598852907 |

| | | |
|---|---|---|
| chr2-191206519-191207020 | 3 | 0.985745290860906 |
| chr6-97807197-97807698 | 3 | 0.942428891339443 |
| chr4-31591236-31591737 | 3 | 0.954910274833079 |
| chr11-25841252-25841753 | 1 | 0.860776718507451 |
| chr14-44571901-44572402 | 1 | 0.918207804767271 |
| chr15-18656531-18657032 | 8 | 0.957322341123792 |
| chr9-103819446-103819947 | 8 | 0.993327170229729 |
| chr6-52860782-52861283 | 3 | 0.974177887353377 |
| chr20-16255868-16256369 | 8 | 0.817514049454156 |
| chr9-85495784-85496285 | 8 | 0.917043714601423 |
| chrX-127276767-127277268 | 5 | 0.824656480453288 |
| chr12-6851121-6851622 | 3 | 0.924329377141753 |
| chr5-24379438-24379939 | 6 | 0.9932197997063 |
| chr5-18757202-18757703 | 3 | 0.996194481285706 |
| chr5-56495895-56496396 | 5 | 0.950953488323017 |
| chr1-91360083-91360584 | 8 | 0.993493681128311 |
| chr16-37477293-37477794 | 3 | 0.987039668730557 |
| chr13-16793159-16793660 | 8 | 0.927484706305402 |
| chr3-26032528-26033029 | 6 | 0.931946202994074 |
| chr1-26291956-26292457 | 5 | 0.821078524045661 |
| chr9-35405247-35405748 | 8 | 0.978622296038626 |
| chr5-5351023-5351524 | 3 | 0.849816813032897 |
| chr18-45725901-45726402 | 3 | 0.967103180847431 |
| chr3-167030609-167031110 | 8 | 0.934474676739656 |
| chr3-90719314-90719815 | 3 | 0.996806620412997 |
| chr3-54883282-54883783 | 2 | 0.935010120374039 |
| chr11-62586369-62586870 | 8 | 0.984075612201811 |
| chr5-73108558-73109059 | 6 | 0.855305617219939 |
| chr2-140253722-140254223 | 5 | 0.836725685495833 |
| chr6-56485494-56485995 | 6 | 0.967903894021581 |
| chr5-18877357-18877858 | 3 | 0.995243090063785 |
| chr13-101206092-101206593 | 3 | 0.990458187908277 |
| chr6-27569367-27569868 | 5 | 0.959784442142102 |
| chr10-72355023-72355524 | 7 | 0.822704543431123 |
| chr9-82200053-82200554 | 6 | 0.992215183653475 |
| chr10-43909942-43910443 | 3 | 0.997039803511685 |
| chr11-17626245-17626746 | 3 | 0.99593553545041 |
| chr2-167366679-167367180 | 2 | 0.917012156783826 |
| chr8-44477002-44477503 | 5 | 0.870729155194726 |
| chr13-99299686-99300187 | 5 | 0.914511692788882 |
| chr3-160185696-160186197 | 2 | 0.822166152559797 |
| chr9-21626502-21627003 | 4 | 0.921587003677735 |
| chr17-21363121-21363622 | 8 | 0.976818121029176 |

| | | |
|---|---|---|
| chr6-99000877-99001378 | 1 | 0.862474881170228 |
| chr2-51398761-51399262 | 5 | 0.972498799671921 |
| chr3-111691317-111691818 | 8 | 0.993493681128311 |
| chrX-108703638-108704139 | 6 | 0.991206925481043 |
| chr5-124575622-124576123 | 6 | 0.993192917948365 |
| chr10-73241629-73242130 | 5 | 0.987133365627442 |
| chr4-178393914-178394415 | 3 | 0.993348457527483 |
| chr14-74412056-74412557 | 8 | 0.932138170198162 |
| chr13-97048458-97048959 | 8 | 0.929435998065712 |
| chr8-61195491-61195992 | 8 | 0.983588094640056 |
| chr19-16496597-16497098 | 1 | 0.921202976877915 |
| chr1-196518595-196519096 | 8 | 0.868684012416063 |
| chr18-40988355-40988856 | 3 | 0.992705284947041 |
| chr2-219420584-219421085 | 8 | 0.993493681128311 |
| chr18-73137214-73137715 | 3 | 0.997130564385875 |
| chr4-62742493-62742994 | 3 | 0.997535247424283 |
| chr10-92059325-92059826 | 8 | 0.920976712931987 |
| chr14-79401585-79402086 | 5 | 0.967925254056284 |
| chr6-39639209-39639710 | 7 | 0.86221567037832 |
| chr8-67347953-67348454 | 2 | 0.90570774885785 |
| chr9-61448016-61448517 | 3 | 0.907746685678627 |
| chr14-22035908-22036409 | 3 | 0.951767993784755 |
| chr3-71613939-71614440 | 8 | 0.982570727515409 |
| chr5-6761389-6761890 | 8 | 0.948486613199308 |
| chr4-121066263-121066764 | 3 | 0.993384327821882 |
| chr10-32217129-32217630 | 3 | 0.987071128872868 |
| chr10-73144076-73144577 | 3 | 0.933563602403102 |
| chr1-234990626-234991127 | 8 | 0.937049349492323 |
| chr16-13421964-13422465 | 3 | 0.994437330720707 |
| chr2-105509059-105509560 | 3 | 0.995017762676604 |
| chr15-4397259-4397760 | 8 | 0.874541532135389 |
| chr11-3682400-3682901 | 5 | 0.892374999350186 |
| chr14-32351965-32352466 | 8 | 0.908730000706271 |
| chr9-83815487-83815988 | 3 | 0.995305905923833 |
| chr5-6444667-6445168 | 1 | 0.835119650051326 |
| chrX-58700186-58700687 | 3 | 0.99593553545041 |
| chr16-26055689-26056190 | 8 | 0.87438757024368 |
| chr7-78647379-78647880 | 1 | 0.892376802479955 |
| chr1-254340667-254341168 | 7 | 0.844355092498728 |
| chr4-149068600-149069101 | 3 | 0.995840852921495 |
| chr15-35318155-35318656 | 8 | 0.872551761925019 |
| chr4-145518993-145519494 | 5 | 0.974641577252115 |
| chr5-56797448-56797949 | 6 | 0.992489104879513 |

| | | |
|---|---|---|
| chr8-90209301-90209802 | 8 | 0.954837846904685 |
| chr1-27396999-27397500 | 6 | 0.968675301462157 |
| chr4-112388234-112388735 | 6 | 0.993203084190338 |
| chr6-87591210-87591711 | 6 | 0.99291294071897 |
| chr2-43056636-43057137 | 8 | 0.886038643044506 |
| chr3-33364071-33364572 | 8 | 0.977801248361411 |
| chr3-104594213-104594714 | 4 | 0.820586057754657 |
| chr2-136937557-136938058 | 3 | 0.991625526800115 |
| chr1-163960226-163960727 | 4 | 0.882014119367668 |
| chr13-103470222-103470723 | 5 | 0.989236528696746 |
| chr7-113582174-113582675 | 8 | 0.977875033066087 |
| chr4-147610579-147611080 | 2 | 0.924544006277537 |
| chr15-67068289-67068790 | 3 | 0.99593553545041 |
| chr13-105264101-105264602 | 3 | 0.981931770417145 |
| chr10-14524073-14524574 | 4 | 0.847609721362813 |
| chr15-37247571-37248072 | 8 | 0.954506677348304 |
| chr2-183034603-183035104 | 6 | 0.835384543698643 |
| chr18-23081371-23081872 | 8 | 0.974953079427083 |
| chr19-36376923-36377424 | 8 | 0.8790211379187 |
| chr19-37364594-37365095 | 5 | 0.893452501141032 |
| chr2-182173400-182173901 | 4 | 0.818933604421746 |
| chr17-38306386-38306887 | 8 | 0.857235050889447 |
| chr2-42168477-42168978 | 8 | 0.993493681128311 |
| chr6-97301578-97302079 | 6 | 0.931084684277307 |
| chr8-30189718-30190219 | 3 | 0.984957769332464 |
| chr2-161515621-161516122 | 3 | 0.99593553545041 |
| chr16-67831859-67832360 | 3 | 0.995223908158298 |
| chr1-179462293-179462794 | 5 | 0.991218592663647 |
| chr3-142099257-142099758 | 2 | 0.818046603257315 |
| chr5-145618204-145618705 | 3 | 0.94612051129813 |
| chr15-56483639-56484140 | 3 | 0.990967150451632 |
| chr2-26187796-26188297 | 5 | 0.892717488003225 |
| chr14-19320421-19320922 | 2 | 0.879597280346788 |
| chr1-132883325-132883826 | 5 | 0.981886678054947 |
| chr1-108853442-108853943 | 3 | 0.992567247856683 |
| chr9-43681986-43682487 | 3 | 0.989752889974843 |
| chr16-15940219-15940720 | 3 | 0.996890776610263 |
| chr9-87448541-87449042 | 3 | 0.965096812353534 |
| chr17-70040026-70040527 | 4 | 0.956163100819943 |
| chr5-146517641-146518142 | 2 | 0.856995025595226 |
| chr3-72502350-72502851 | 3 | 0.841095310972024 |
| chr2-202727084-202727585 | 2 | 0.943363609253309 |
| chr1-137274704-137275205 | 5 | 0.961676546609519 |

| | | |
|---|---|---|
| chr8-45804445-45804946 | 6 | 0.991144152450612 |
| chr6-71529770-71530271 | 3 | 0.996266003027604 |
| chr19-19476917-19477418 | 3 | 0.945254584539918 |
| chr16-56813664-56814165 | 8 | 0.897790942369013 |
| chr16-49855167-49855668 | 3 | 0.991674984603655 |
| chr9-99061563-99062064 | 8 | 0.861578552539099 |
| chr17-16665360-16665861 | 6 | 0.930215593792039 |
| chr13-103906266-103906767 | 4 | 0.866488013618079 |
| chr11-68605079-68605580 | 1 | 0.826569448677409 |
| chr6-36529826-36530327 | 6 | 0.985440517078562 |
| chr4-57518303-57518804 | 8 | 0.844046979481041 |
| chr20-41897455-41897956 | 5 | 0.906947122515292 |
| chr13-46112307-46112808 | 8 | 0.932149354720491 |
| chr1-84975487-84975988 | 6 | 0.991206925481043 |
| chr15-3607957-3608458 | 5 | 0.982270705962439 |
| chr5-24744712-24745213 | 7 | 0.869585115593272 |
| chr3-113847244-113847745 | 8 | 0.907305756348682 |
| chr7-86679025-86679526 | 8 | 0.983254614936399 |
| chr3-25260137-25260638 | 4 | 0.844266807019159 |
| chr2-22370022-22370523 | 3 | 0.991555331980348 |
| chr10-3853473-3853974 | 2 | 0.829866435378611 |
| chr18-27384637-27385138 | 8 | 0.988489132587553 |
| chr7-65514988-65515489 | 3 | 0.992791207430702 |
| chr2-81322330-81322831 | 3 | 0.996805531485491 |
| chr2-98895678-98896179 | 3 | 0.959201128429845 |
| chr3-30379526-30380027 | 8 | 0.843220453830265 |
| chr6-25386013-25386514 | 4 | 0.863365370655677 |
| chr12-4677762-4678263 | 8 | 0.986239762586816 |
| chr4-173741288-173741789 | 6 | 0.977987554132111 |
| chr15-2013757-2014258 | 8 | 0.954332251702795 |
| chr16-34837470-34837971 | 8 | 0.958218610164256 |
| chr2-24220058-24220559 | 8 | 0.851820955256689 |
| chr1-227955578-227956079 | 3 | 0.967722278586745 |
| chr4-105078884-105079385 | 3 | 0.96780575969537 |
| chr1-22514156-22514657 | 8 | 0.941486785854796 |
| chr4-58325895-58326396 | 8 | 0.904228252950975 |
| chr1-5081682-5082183 | 3 | 0.996677775597142 |
| chr14-71884292-71884793 | 3 | 0.983971048766084 |
| chr5-64385161-64385662 | 3 | 0.99593553545041 |
| chrX-106423032-106423533 | 3 | 0.886145847908051 |
| chr7-44737979-44738480 | 5 | 0.88433505539108 |
| chr9-15555080-15555581 | 3 | 0.989658291058861 |
| chr7-62554199-62554700 | 3 | 0.996786701272962 |

| | | |
|---|---|---|
| chr3-19324303-19324804 | 8 | 0.830387298746894 |
| chr7-9783118-9783619 | 5 | 0.906918550879416 |
| chr2-191947920-191948421 | 5 | 0.986039026660709 |
| chr6-1653323-1653824 | 3 | 0.995985247144793 |
| chr4-125076422-125076923 | 8 | 0.992459940150411 |
| chr4-96248854-96249355 | 1 | 0.83277902876428 |
| chr17-13964068-13964569 | 4 | 0.967149493791843 |
| chr20-35264444-35264945 | 7 | 0.839861713486903 |
| chr13-8879941-8880442 | 3 | 0.996821895708701 |
| chr15-35108429-35108930 | 4 | 0.885838365190134 |
| chr14-71119230-71119731 | 6 | 0.989462254105253 |
| chr8-73554135-73554636 | 5 | 0.994351937332814 |
| chr8-54123797-54124298 | 5 | 0.95861535695513 |
| chr5-48468468-48468969 | 5 | 0.952892389636687 |
| chr15-46774956-46775457 | 6 | 0.988495286514156 |
| chr13-43053015-43053516 | 3 | 0.995069899799163 |
| chrX-27234845-27235346 | 6 | 0.956001647916643 |
| chr13-105571074-105571575 | 3 | 0.984932410165347 |
| chr1-219786150-219786651 | 3 | 0.950044892578825 |
| chr13-91773159-91773660 | 5 | 0.899103049900226 |
| chr20-92680-93181 | 8 | 0.967788806297786 |
| chr10-78770794-78771295 | 6 | 0.948075940275309 |
| chr14-19897809-19898310 | 3 | 0.955130719552539 |
| chr13-76586346-76586847 | 3 | 0.995015409546495 |
| chr4-80829190-80829691 | 4 | 0.923991755217852 |
| chr1-55069147-55069648 | 2 | 0.858430200355947 |
| chr1-150753933-150754434 | 3 | 0.99571890801685 |
| chr5-130865126-130865627 | 2 | 0.939225437460728 |
| chr9-15726291-15726792 | 8 | 0.910511804221347 |
| chr3-19192421-19192922 | 8 | 0.881483405390638 |
| chr4-162604800-162605301 | 6 | 0.993228279193384 |
| chr1-86900796-86901297 | 3 | 0.830365199271827 |
| chr2-12807383-12807884 | 8 | 0.989308552450199 |
| chr15-832630-833131 | 7 | 0.894081085483309 |
| chr19-47463116-47463617 | 8 | 0.938098671372204 |
| chr9-71022369-71022870 | 8 | 0.969126249851908 |
| chr3-88817096-88817597 | 5 | 0.983919978840557 |
| chr7-127282123-127282624 | 8 | 0.995231428299854 |
| chr4-177837243-177837744 | 3 | 0.874840400815808 |
| chr17-74737768-74738269 | 5 | 0.986742969285917 |
| chr5-22477482-22477983 | 1 | 0.872185676975698 |
| chr15-8621640-8622141 | 6 | 0.986164574289357 |
| chr5-8288095-8288596 | 1 | 0.865152599020692 |

| | | |
|---|---|---|
| chr12-11218726-11219227 | 8 | 0.950356672256456 |
| chr5-58763301-58763802 | 8 | 0.939699599321006 |
| chr5-164465104-164465605 | 6 | 0.985458029143926 |
| chr17-86400599-86401100 | 3 | 0.996403288462141 |
| chr1-17949593-17950094 | 8 | 0.96439323737654 |
| chrX-136088852-136089353 | 5 | 0.983919978840557 |
| chr8-55633215-55633716 | 4 | 0.963253129891826 |
| chr4-11326344-11326845 | 5 | 0.930731069141431 |
| chr8-76937126-76937627 | 6 | 0.961969835915609 |
| chr3-142352534-142353035 | 3 | 0.991348109653365 |
| chr7-90661558-90662059 | 8 | 0.988784878208232 |
| chr3-100178787-100179288 | 7 | 0.920886149210766 |
| chr11-79904969-79905470 | 3 | 0.994859590290088 |
| chr3-119789530-119790031 | 3 | 0.884842997712034 |
| chr1-228172261-228172762 | 5 | 0.824373505188177 |
| chr6-106932330-106932831 | 5 | 0.963234437810482 |
| chr3-130332398-130332899 | 3 | 0.994665445686273 |
| chr4-179755480-179755981 | 3 | 0.99641052254019 |
| chr11-47720977-47721478 | 6 | 0.982056364665141 |
| chr11-35520537-35521038 | 8 | 0.988832984198516 |
| chr13-80077908-80078409 | 5 | 0.988721183822435 |
| chr3-56686593-56687094 | 8 | 0.957277004152819 |
| chr1-207024350-207024851 | 8 | 0.939907767836555 |
| chr10-8943200-8943701 | 3 | 0.900347876575394 |
| chr2-86980879-86981380 | 8 | 0.993493681128311 |
| chr19-28331908-28332409 | 4 | 0.950032954020311 |
| chr4-55529642-55530143 | 3 | 0.995953993055754 |
| chr9-50269035-50269536 | 8 | 0.994866478527759 |
| chr10-31801393-31801894 | 8 | 0.993493681128311 |
| chr1-116580394-116580895 | 6 | 0.990693683818644 |
| chr3-119411605-119412106 | 3 | 0.927586090346804 |
| chr19-16177113-16177614 | 5 | 0.982323266517659 |
| chr2-122813362-122813863 | 5 | 0.983919978840557 |
| chr18-4373037-4373538 | 2 | 0.824398752608238 |
| chr16-55666871-55667372 | 1 | 0.825727053447515 |
| chr1-171208869-171209370 | 3 | 0.973466828530369 |
| chr20-19237328-19237829 | 8 | 0.924839441059114 |
| chr1-52638978-52639479 | 6 | 0.991248730007621 |
| chr4-132213752-132214253 | 6 | 0.984772345833995 |
| chr12-44305565-44306066 | 5 | 0.990706261166045 |
| chr1-183740244-183740745 | 3 | 0.875767647764608 |
| chr5-134040514-134041015 | 3 | 0.900912226201675 |
| chr7-38450868-38451369 | 3 | 0.98203946891168 |

| | | |
|---|---|---|
| chr4-128379015-128379516 | 6 | 0.988764403229561 |
| chr10-50229506-50230007 | 8 | 0.988865758055956 |
| chr4-74450619-74451120 | 3 | 0.980312609646898 |
| chr10-72423869-72424370 | 8 | 0.894769627992834 |
| chr15-79724931-79725432 | 7 | 0.882131322751744 |
| chr17-64038908-64039409 | 3 | 0.99593553545041 |
| chr1-97208994-97209495 | 2 | 0.937181114525283 |
| chr6-40729721-40730222 | 8 | 0.909891619726916 |
| chr1-58575453-58575954 | 6 | 0.944927936399774 |
| chr2-84710623-84711124 | 8 | 0.816996354479258 |
| chr7-62787531-62788032 | 2 | 0.839126247082564 |
| chr2-195973477-195973978 | 5 | 0.945290939606419 |
| chr4-146993894-146994395 | 6 | 0.975415639393346 |
| chr19-19216553-19217054 | 8 | 0.959951078606777 |
| chr12-41582945-41583446 | 5 | 0.994753897802664 |
| chr15-31797456-31797957 | 8 | 0.880088873085938 |
| chr12-31305303-31305804 | 8 | 0.857699687645576 |
| chr8-89321632-89322133 | 4 | 0.924974781536388 |
| chr6-13585417-13585918 | 8 | 0.991982475990422 |
| chr5-3972773-3973274 | 6 | 0.908472187803099 |
| chr3-45402617-45403118 | 6 | 0.978854453495949 |
| chr17-18667461-18667962 | 2 | 0.978620388105242 |
| chr17-60014221-60014722 | 2 | 0.916983283169412 |
| chr6-25214214-25214715 | 3 | 0.989453479903656 |
| chr2-165888749-165889250 | 3 | 0.875347533470413 |
| chr19-9573120-9573621 | 8 | 0.988751138731994 |
| chr19-34910422-34910923 | 8 | 0.918611453046004 |
| chr12-45970088-45970589 | 3 | 0.973653370743961 |
| chr19-42455375-42455876 | 5 | 0.830776016181814 |
| chr18-75976043-75976544 | 3 | 0.995821925536532 |
| chr20-11499274-11499775 | 1 | 0.950835274265764 |
| chr1-160894028-160894529 | 5 | 0.983919978840557 |
| chr13-46210462-46210963 | 6 | 0.976353595537914 |
| chr2-111135861-111136362 | 3 | 0.963927838869932 |
| chr16-47673992-47674493 | 3 | 0.971598526287901 |
| chr18-62316478-62316979 | 4 | 0.898230589489443 |
| chr8-49174953-49175454 | 8 | 0.965062718756162 |
| chr7-38303066-38303567 | 3 | 0.989790770165388 |
| chr1-83782465-83782966 | 4 | 0.888433432990074 |
| chr7-37980091-37980592 | 3 | 0.995210291428128 |
| chr10-59994715-59995216 | 8 | 0.841983368670711 |
| chrX-130333022-130333523 | 8 | 0.969483652055193 |
| chr1-93572372-93572873 | 3 | 0.834389658723571 |

| | | |
|---|---|---|
| chr10-50199005-50199506 | 6 | 0.936973596762631 |
| chr2-136326260-136326761 | 2 | 0.8455413706567 |
| chr4-180503364-180503865 | 6 | 0.927072972683907 |
| chr12-25015015-25015516 | 8 | 0.940036316941925 |
| chr3-80481884-80482385 | 5 | 0.971424353503016 |
| chr19-48119566-48120067 | 3 | 0.998076739028436 |
| chr1-109336615-109337116 | 3 | 0.996721370667899 |
| chr13-67499078-67499579 | 3 | 0.921450941047957 |
| chr1-5021569-5022070 | 6 | 0.988927653473905 |
| chr7-47261885-47262386 | 8 | 0.861266414770062 |
| chr2-169711763-169712264 | 8 | 0.847940045274686 |
| chr14-98796480-98796981 | 4 | 0.88794124102652 |
| chr9-106022601-106023102 | 6 | 0.980779434384718 |
| chr3-10544707-10545208 | 5 | 0.991721807107564 |
| chr1-45439322-45439823 | 3 | 0.894701515605788 |
| chr13-46774305-46774806 | 8 | 0.979329208868638 |
| chr15-11193650-11194151 | 5 | 0.986079735625148 |
| chr2-8473403-8473904 | 6 | 0.994028424132074 |
| chr20-47295279-47295780 | 8 | 0.990350910685438 |
| chr2-49497195-49497696 | 5 | 0.987227116058279 |
| chr3-159376321-159376822 | 7 | 0.824659540957102 |
| chr3-46895141-46895642 | 8 | 0.898435187666413 |
| chr13-54803506-54804007 | 3 | 0.876784672282611 |
| chr1-100536367-100536868 | 8 | 0.886166713173809 |
| chr2-41215794-41216295 | 3 | 0.988512308368875 |
| chr10-29506635-29507136 | 8 | 0.906592540423492 |
| chr15-92466133-92466634 | 3 | 0.880320138008859 |
| chr2-148675084-148675585 | 3 | 0.931864421296473 |
| chr19-30820175-30820676 | 3 | 0.983350487378462 |
| chr14-27096181-27096682 | 4 | 0.957082918204961 |
| chr9-111182283-111182784 | 8 | 0.829012146544761 |
| chr19-12549681-12550182 | 3 | 0.99593553545041 |
| chr8-45301752-45302253 | 6 | 0.986623780458944 |
| chr2-188713660-188714161 | 4 | 0.908658653749819 |
| chr8-69967524-69968025 | 2 | 0.92664612946001 |
| chr15-26459884-26460385 | 8 | 0.953003623919247 |
| chr7-58161966-58162467 | 5 | 0.88621572794843 |
| chr7-94172655-94173156 | 4 | 0.899454698248551 |
| chr1-120879786-120880287 | 2 | 0.925529618732685 |
| chr15-53073161-53073662 | 4 | 0.917824864257413 |
| chr14-56747177-56747678 | 8 | 0.995375853994712 |
| chr13-74155813-74156314 | 6 | 0.991206925481043 |
| chr7-50449617-50450118 | 3 | 0.931406143299057 |

| | | |
|---|---|---|
| chr3-41187769-41188270 | 6 | 0.983325037361701 |
| chr3-130023444-130023945 | 4 | 0.854989996359691 |
| chr10-37315881-37316382 | 8 | 0.825410450756163 |
| chr9-92688451-92688952 | 8 | 0.923858213838563 |
| chr4-68942802-68943303 | 6 | 0.855091587979433 |
| chr19-18527071-18527572 | 5 | 0.99271890899594 |
| chr10-104989591-104990092 | 8 | 0.937145413396219 |
| chr9-88852743-88853244 | 8 | 0.878983779254915 |
| chr9-23778958-23779459 | 7 | 0.868951851063257 |
| chr1-171779757-171780258 | 6 | 0.930755174842048 |
| chr20-10158993-10159494 | 8 | 0.945026322009606 |
| chr14-54146686-54147187 | 3 | 0.99679175779585 |
| chr5-120197592-120198093 | 6 | 0.891871622700538 |
| chr19-4925651-4926152 | 3 | 0.821115775968452 |
| chr5-68285314-68285815 | 3 | 0.995510053470719 |
| chr10-71272135-71272636 | 8 | 0.988242657812884 |
| chr5-25076541-25077042 | 1 | 0.850266434432549 |
| chr7-16480898-16481399 | 3 | 0.989790595377599 |
| chr7-19795682-19796183 | 4 | 0.84628209891725 |
| chr5-163800908-163801409 | 4 | 0.889108248987899 |
| chr5-18917639-18918140 | 8 | 0.912048780343791 |
| chr5-165187737-165188238 | 8 | 0.951492138440243 |
| chr3-132976340-132976841 | 8 | 0.964174 |
| chr2-236960437-236960938 | 3 | 0.974365611681727 |
| chr4-161001054-161001555 | 7 | 0.87169419288892 |
| chr4-104829542-104830043 | 8 | 0.826523039429062 |
| chr13-80648055-80648556 | 3 | 0.995594655228677 |
| chr10-71565917-71566418 | 8 | 0.969153687226696 |
| chr4-13301948-13302449 | 6 | 0.96906702128409 |
| chr14-7195779-7196280 | 8 | 0.993493681128311 |
| chr5-43142946-43143447 | 8 | 0.854708707273047 |
| chr9-43279884-43280385 | 6 | 0.992945747779593 |
| chr16-79581861-79582362 | 3 | 0.996418027030115 |
| chr4-148337824-148338325 | 3 | 0.996875853559562 |
| chr6-10296299-10296800 | 6 | 0.991206925481043 |
| chr14-71516098-71516599 | 8 | 0.959064854639053 |
| chr6-73392555-73393056 | 3 | 0.986199813707826 |
| chr5-149859311-149859812 | 8 | 0.822108519140873 |
| chr1-188834230-188834731 | 3 | 0.951153870304501 |
| chr2-20907524-20908025 | 3 | 0.99593553545041 |
| chr14-75614040-75614541 | 5 | 0.901767811068897 |
| chr9-103429266-103429767 | 4 | 0.943398824400568 |
| chr3-41250952-41251453 | 8 | 0.962339551230054 |

| | | |
|---|---|---|
| chr16-45295219-45295720 | 7 | 0.918281511311867 |
| chr13-21266944-21267445 | 3 | 0.995010760810788 |
| chr3-118358854-118359355 | 7 | 0.822944214747881 |
| chr3-151399230-151399731 | 3 | 0.87785461881118 |
| chr1-167250315-167250816 | 2 | 0.938862215570934 |
| chr2-135261429-135261930 | 5 | 0.990938350573003 |
| chr10-37031826-37032327 | 4 | 0.863593040045058 |
| chr15-53758868-53759369 | 8 | 0.944838432178119 |
| chr8-89190918-89191419 | 3 | 0.99593553545041 |
| chr6-57881138-57881639 | 6 | 0.901199843612476 |
| chr15-16388586-16389087 | 8 | 0.960084344343353 |
| chr14-58222103-58222604 | 8 | 0.992085721669391 |
| chr1-187294363-187294864 | 6 | 0.982730021132768 |
| chr10-4908743-4909244 | 1 | 0.92677243154758 |
| chr8-63003007-63003508 | 5 | 0.852972955012098 |
| chr7-113520963-113521464 | 8 | 0.982687583656404 |
| chr1-16981808-16982309 | 1 | 0.896905722445186 |
| chr4-142018502-142019003 | 1 | 0.881305501085205 |
| chr19-34695407-34695908 | 3 | 0.951101838572652 |
| chr16-67733019-67733520 | 5 | 0.986235147769794 |
| chr14-64519272-64519773 | 3 | 0.99593553545041 |
| chrX-116902637-116903138 | 8 | 0.84987616640956 |
| chr17-30445060-30445561 | 8 | 0.956955610891306 |
| chr16-22402963-22403464 | 3 | 0.945207653607496 |
| chr2-79510429-79510930 | 3 | 0.995346114700231 |
| chr6-95447013-95447514 | 3 | 0.910617922713078 |
| chr20-46035618-46036119 | 5 | 0.978533502545205 |
| chr14-67082762-67083263 | 5 | 0.972686028469589 |
| chr3-82477652-82478153 | 3 | 0.994287484338095 |
| chr18-59918606-59919107 | 8 | 0.82770978957503 |
| chr11-42366265-42366766 | 6 | 0.934267429625418 |
| chr20-32463161-32463662 | 3 | 0.99593553545041 |
| chr17-48195950-48196451 | 4 | 0.832168736451809 |
| chr12-27630519-27631020 | 6 | 0.868779268390339 |
| chr2-76165821-76166322 | 7 | 0.824492646751557 |
| chr2-243855771-243856272 | 3 | 0.985215533768353 |
| chr7-106661908-106662409 | 6 | 0.914875659870423 |
| chr1-212376075-212376576 | 8 | 0.965059042074353 |
| chr5-143340812-143341313 | 6 | 0.993040217050868 |
| chrX-66849819-66850320 | 2 | 0.868713018968968 |
| chr4-121835253-121835754 | 1 | 0.819825668987807 |
| chr15-74600621-74601122 | 3 | 0.992345900750862 |
| chr20-1430639-1431140 | 1 | 0.935808029774883 |

| | | |
|---|---|---|
| chr17-67865065-67865566 | 3 | 0.99593553545041 |
| chr13-80150144-80150645 | 6 | 0.991206925481043 |
| chr5-20417225-20417726 | 8 | 0.910536505498701 |
| chr19-5805023-5805524 | 3 | 0.99593553545041 |
| chr1-53890677-53891178 | 8 | 0.821458669866813 |
| chr4-175842121-175842622 | 5 | 0.933829123892329 |
| chr17-5435531-5436032 | 8 | 0.968205128908181 |
| chr13-101593094-101593595 | 6 | 0.947993164443422 |
| chr3-60297809-60298310 | 1 | 0.883175504674075 |
| chr6-138906280-138906781 | 3 | 0.989954406921269 |
| chr15-16621323-16621824 | 3 | 0.929362007346763 |
| chr6-112664297-112664798 | 6 | 0.991206925481043 |
| chr4-10353117-10353618 | 5 | 0.840456394462581 |
| chr8-65416349-65416850 | 2 | 0.913680199461218 |
| chr5-46728929-46729430 | 8 | 0.985813827256651 |
| chr4-82887173-82887674 | 8 | 0.825121080338685 |
| chr1-244770066-244770567 | 5 | 0.987730201483308 |
| chr4-93076306-93076807 | 6 | 0.969004455857886 |
| chr9-75911085-75911586 | 7 | 0.969996135500341 |
| chr9-88814741-88815242 | 6 | 0.976509322758922 |
| chr6-46084287-46084788 | 3 | 0.962572051655828 |
| chr2-19954157-19954658 | 8 | 0.931196458095326 |
| chr4-124573903-124574404 | 2 | 0.820711447726279 |
| chr4-120178389-120178890 | 6 | 0.993029871174098 |
| chr12-23098331-23098832 | 8 | 0.918175137997066 |
| chr16-45885729-45886230 | 8 | 0.995204657613716 |
| chr2-241447369-241447870 | 8 | 0.925610519740879 |
| chr2-175399610-175400111 | 3 | 0.90075731462804 |
| chr10-45098104-45098605 | 5 | 0.989934752889006 |
| chr3-114863809-114864310 | 8 | 0.848490674466047 |
| chr13-64742801-64743302 | 3 | 0.922522445359259 |
| chr19-49692755-49693256 | 7 | 0.948792034080471 |
| chr9-42942015-42942516 | 6 | 0.990955632055628 |
| chr2-99377208-99377709 | 8 | 0.983851067509458 |
| chr2-149598783-149599284 | 8 | 0.910278009437883 |
| chr9-106966899-106967400 | 5 | 0.956517582879197 |
| chr5-109681397-109681898 | 8 | 0.974259824041293 |
| chr6-90098465-90098966 | 6 | 0.855452811281858 |
| chr4-33539055-33539556 | 5 | 0.964881212538877 |
| chr20-28498144-28498645 | 6 | 0.982201514113214 |
| chr20-17030045-17030546 | 3 | 0.981286262734634 |
| chr4-19039044-19039545 | 2 | 0.912218499076395 |
| chr2-240956304-240956805 | 3 | 0.992275022886988 |

| | | |
|---|---|---|
| chr7-88168706-88169207 | 8 | 0.990584874104544 |
| chr17-20494538-20495039 | 6 | 0.98448905051072 |
| chr14-28671341-28671842 | 3 | 0.982810311242845 |
| chr15-40856649-40857150 | 6 | 0.986483410442562 |
| chr8-4294648-4295149 | 6 | 0.954665035415292 |
| chr12-35437234-35437735 | 6 | 0.960398321294277 |
| chr8-66586764-66587265 | 3 | 0.988321740073589 |
| chr6-30665351-30665852 | 3 | 0.907082314746218 |
| chr4-60876896-60877397 | 3 | 0.816735245493121 |
| chr17-24476796-24477297 | 8 | 0.890191932633579 |
| chrX-117261297-117261798 | 6 | 0.959955727011313 |
| chr15-55104724-55105225 | 3 | 0.827668517494738 |
| chr13-104326144-104326645 | 8 | 0.90318436610781 |
| chr3-50170069-50170570 | 3 | 0.990420412135465 |
| chr9-79922798-79923299 | 3 | 0.926927725822911 |
| chr4-57090547-57091048 | 3 | 0.983445993118503 |
| chr16-84262405-84262906 | 5 | 0.86932900304416 |
| chr9-73368075-73368576 | 6 | 0.949766975083828 |
| chr18-49498488-49498989 | 3 | 0.995030065149055 |
| chr5-92562702-92563203 | 1 | 0.918642977908946 |
| chr15-37724760-37725261 | 2 | 0.928398725784688 |
| chr17-53669366-53669867 | 8 | 0.989273589450196 |
| chr18-75337893-75338394 | 3 | 0.99083317815464 |
| chr14-83376628-83377129 | 8 | 0.944153245035413 |
| chr2-13974157-13974658 | 3 | 0.99593553545041 |
| chr5-150342572-150343073 | 4 | 0.941058743332941 |
| chr6-108271651-108272152 | 2 | 0.884096903094187 |
| chr2-144603766-144604267 | 8 | 0.931789460179216 |
| chr13-100633309-100633810 | 8 | 0.993107734510986 |
| chr10-28830258-28830759 | 8 | 0.882006462918492 |
| chr3-143364717-143365218 | 1 | 0.887882384496219 |
| chr2-60780510-60781011 | 5 | 0.826765881673989 |
| chr2-157938623-157939124 | 8 | 0.993493681128311 |
| chr11-36231547-36232048 | 3 | 0.869430586783066 |
| chr19-35750637-35751138 | 8 | 0.989134469330397 |
| chr4-118495164-118495665 | 3 | 0.95340519177707 |
| chr8-102003404-102003905 | 3 | 0.947322315873505 |
| chrX-96981964-96982465 | 3 | 0.996192448472295 |
| chr7-95228665-95229166 | 8 | 0.862194042286112 |
| chr1-183361136-183361637 | 3 | 0.996506322640294 |
| chr14-48038806-48039307 | 3 | 0.99593553545041 |
| chr2-142350726-142351227 | 8 | 0.972805559972021 |
| chr2-243403490-243403991 | 5 | 0.978838492396425 |

| | | |
|---|---|---|
| chr20-49691678-49692179 | 8 | 0.987978707066004 |
| chr10-104139841-104140342 | 8 | 0.984712308175873 |
| chr15-94555236-94555737 | 5 | 0.933988802956376 |
| chr5-137076740-137077241 | 8 | 0.921807651053994 |
| chr18-65054025-65054526 | 3 | 0.992122848639319 |
| chr13-78811006-78811507 | 6 | 0.908382400394935 |
| chr2-109720425-109720926 | 5 | 0.983919978840557 |
| chr8-56107547-56108048 | 6 | 0.992021696948434 |
| chr5-42950863-42951364 | 2 | 0.816531586090713 |
| chr2-229472449-229472950 | 3 | 0.996200278455491 |
| chr10-50122674-50123175 | 2 | 0.96854399985493 |
| chr3-161462706-161463207 | 8 | 0.995189079786298 |
| chr13-32212077-32212578 | 5 | 0.966286710904433 |
| chr1-231325691-231326192 | 7 | 0.912077010390253 |
| chr5-162022116-162022617 | 3 | 0.992868377805589 |
| chr8-70872527-70873028 | 8 | 0.865405958110056 |
| chr17-30699645-30700146 | 3 | 0.997160404691593 |
| chr3-117530224-117530725 | 8 | 0.877664704454928 |
| chr20-9268999-9269500 | 6 | 0.992457114114228 |
| chr19-56411986-56412487 | 1 | 0.842381361481893 |
| chr1-128424935-128425436 | 3 | 0.998541973144968 |
| chr1-49585973-49586474 | 3 | 0.995609025606054 |
| chr15-96710207-96710708 | 6 | 0.989619108361735 |
| chr8-97327658-97328159 | 4 | 0.915715092347614 |
| chr13-92919750-92920251 | 8 | 0.927274715593802 |
| chr1-15148442-15148943 | 5 | 0.97801635729946 |
| chr14-101940779-101941280 | 3 | 0.945851188699725 |
| chr12-27470420-27470921 | 8 | 0.825545490912126 |
| chr3-23142042-23142543 | 3 | 0.996470503861321 |
| chr7-131960425-131960926 | 5 | 0.965590062978528 |
| chr1-165307935-165308436 | 8 | 0.840593898740268 |
| chr13-50126807-50127308 | 8 | 0.919621729801879 |
| chr9-52080838-52081339 | 8 | 0.882835766588344 |
| chr17-17867311-17867812 | 8 | 0.917152361523017 |
| chr12-10155575-10156076 | 8 | 0.909327864373297 |
| chr18-82516447-82516948 | 3 | 0.868423820563336 |
| chr12-41800718-41801219 | 6 | 0.971451242783689 |
| chr3-28946671-28947172 | 3 | 0.996894863243938 |
| chr8-30497884-30498385 | 8 | 0.980631366714687 |
| chr14-97388996-97389497 | 8 | 0.950395940955966 |
| chr1-39376120-39376621 | 6 | 0.988917866659592 |
| chr11-59499712-59500213 | 8 | 0.993606237578643 |
| chr8-92896197-92896698 | 3 | 0.84334562179349 |

| | | |
|---|---|---|
| chr4-178775809-178776310 | 5 | 0.971431187909655 |
| chr3-140985912-140986413 | 5 | 0.986950234271898 |
| chr15-16345008-16345509 | 6 | 0.991206925481043 |
| chr2-61380266-61380767 | 5 | 0.901076809140482 |
| chr1-7029394-7029895 | 7 | 0.836930438596335 |
| chr4-43639008-43639509 | 3 | 0.981443600889701 |
| chr14-52467893-52468394 | 5 | 0.983207763715534 |
| chr14-83844790-83845291 | 3 | 0.992744666770676 |
| chr13-35845682-35846183 | 3 | 0.99593553545041 |
| chr3-147257115-147257616 | 2 | 0.955352660456419 |
| chr1-163763619-163764120 | 3 | 0.882176600822795 |
| chr7-43633468-43633969 | 3 | 0.99593553545041 |
| chr7-64869601-64870102 | 6 | 0.992278898762908 |
| chr17-24231891-24232392 | 6 | 0.953497986443158 |
| chr8-48388310-48388811 | 7 | 0.855308131162988 |
| chr4-177633900-177634401 | 8 | 0.86106658640417 |
| chr9-64375620-64376121 | 7 | 0.87436320496017 |
| chr1-221274606-221275107 | 3 | 0.995360795832981 |
| chr4-117734286-117734787 | 3 | 0.975803857547287 |
| chr8-36915405-36915906 | 8 | 0.971232854425484 |
| chr18-55982566-55983067 | 3 | 0.834074841301272 |
| chr11-21891179-21891680 | 5 | 0.990261727192761 |
| chr8-23698998-23699499 | 3 | 0.965375417122793 |
| chr7-69010843-69011344 | 3 | 0.999197816021703 |
| chr1-186342369-186342870 | 6 | 0.985976824981788 |
| chr9-2395857-2396358 | 3 | 0.993096446730213 |
| chr4-133688431-133688932 | 8 | 0.995238041711041 |
| chr20-18128001-18128502 | 1 | 0.909215520398565 |
| chr3-23912872-23913373 | 5 | 0.985750482106667 |
| chr11-29042600-29043101 | 4 | 0.841535832609923 |
| chr5-14100701-14101202 | 3 | 0.99593553545041 |
| chr6-45407051-45407552 | 6 | 0.996960511440807 |
| chr3-52305761-52306262 | 3 | 0.986165531759578 |
| chr1-117953629-117954130 | 8 | 0.993493681128311 |
| chr5-124140484-124140985 | 3 | 0.996056325924105 |
| chr5-141333781-141334282 | 8 | 0.881451105125977 |
| chr8-87703181-87703682 | 2 | 0.940710749320683 |
| chr16-74727316-74727817 | 8 | 0.915020494580671 |
| chr5-62429678-62430179 | 6 | 0.968981636590759 |
| chr1-123991297-123991798 | 5 | 0.981644301830801 |
| chr3-135085628-135086129 | 5 | 0.956622519807534 |
| chr16-5552818-5553319 | 6 | 0.9140119052554 |
| chr4-179730993-179731494 | 5 | 0.991451872600602 |

| | | |
|---|---|---|
| chr9-83697388-83697889 | 6 | 0.996036742869878 |
| chr9-86324207-86324708 | 3 | 0.973104977202597 |
| chr3-199623-200124 | 3 | 0.918661397163018 |
| chr14-77903905-77904406 | 8 | 0.956987946673142 |
| chr1-126017053-126017554 | 6 | 0.992401577759655 |
| chr13-23424967-23425468 | 3 | 0.99593553545041 |
| chr5-153794588-153795089 | 8 | 0.926755854773694 |
| chr2-190347947-190348448 | 3 | 0.992312975889346 |
| chr8-115365880-115366381 | 6 | 0.949218999212746 |
| chr3-148673851-148674352 | 3 | 0.97227310919944 |
| chr13-68421903-68422404 | 4 | 0.940604228416494 |
| chr2-40248210-40248711 | 8 | 0.993493681128311 |
| chr18-21380454-21380955 | 2 | 0.913528242493566 |
| chr12-33825590-33826091 | 4 | 0.966507739830463 |
| chr5-96629521-96630022 | 8 | 0.947836050972391 |
| chr4-84256222-84256723 | 8 | 0.925871864179073 |
| chr1-53342653-53343154 | 3 | 0.902078486122825 |
| chr1-8387913-8388414 | 5 | 0.988425729263765 |
| chr17-44089054-44089555 | 3 | 0.996889806606285 |
| chrX-79973192-79973693 | 1 | 0.826756867937238 |
| chr7-122806348-122806849 | 6 | 0.987177149387313 |
| chr3-77669717-77670218 | 5 | 0.927708888865791 |
| chr16-43720924-43721425 | 5 | 0.986237637729397 |
| chr14-41604144-41604645 | 3 | 0.968716202486512 |
| chr5-54304321-54304822 | 4 | 0.844937 |
| chr7-97901724-97902225 | 3 | 0.996760023301225 |
| chr8-43881065-43881566 | 5 | 0.957246983802414 |
| chr3-12750001-12750502 | 5 | 0.94814828444514 |
| chr3-12677873-12678374 | 4 | 0.854608944350519 |
| chr4-74668858-74669359 | 8 | 0.8834727493314 |
| chr8-47874692-47875193 | 6 | 0.961753309581762 |
| chr14-18715394-18715895 | 4 | 0.833219192029078 |
| chr12-28320878-28321379 | 8 | 0.955420881364293 |
| chr10-90594101-90594602 | 8 | 0.882223899223009 |
| chr9-74858526-74859027 | 4 | 0.964581107379884 |
| chr6-20876304-20876805 | 8 | 0.993659763895484 |
| chr18-67073354-67073855 | 5 | 0.8573905542775 |
| chrX-23571510-23572011 | 8 | 0.94922235115499 |
| chr16-44144445-44144946 | 4 | 0.954221483121158 |
| chr5-113701412-113701913 | 5 | 0.883716008149802 |
| chrX-141997539-141998040 | 3 | 0.99593553545041 |
| chr14-2873464-2873965 | 6 | 0.978071304965444 |
| chr15-31859835-31860336 | 6 | 0.846289948100747 |

| | | |
|---|---|---|
| chr5-133130554-133131055 | 5 | 0.974880899197845 |
| chr1-153433988-153434489 | 8 | 0.87916357220713 |
| chr18-15335262-15335763 | 4 | 0.853562719142912 |
| chr9-69417047-69417548 | 3 | 0.956750473079355 |
| chr11-11438519-11439020 | 8 | 0.82584231038925 |
| chr2-44377340-44377841 | 1 | 0.842610353723178 |
| chr4-46939098-46939599 | 8 | 0.887016965847097 |
| chr2-104364954-104365455 | 3 | 0.877935172030624 |
| chr3-79185720-79186221 | 5 | 0.932400690154292 |
| chr5-109711386-109711887 | 3 | 0.950428991162673 |
| chr7-69416000-69416501 | 3 | 0.872636128453714 |
| chr7-115889118-115889619 | 4 | 0.840843886866761 |
| chr10-44317568-44318069 | 2 | 0.827522005635674 |
| chr1-217154261-217154762 | 6 | 0.897477403553028 |
| chr4-117652720-117653221 | 3 | 0.912678362748464 |
| chr4-146489089-146489590 | 8 | 0.930779899851916 |
| chr1-149695071-149695572 | 4 | 0.934535199252262 |
| chr1-85854561-85855062 | 8 | 0.993493681128311 |
| chr4-150145743-150146244 | 6 | 0.969577664008232 |
| chr16-56221016-56221517 | 8 | 0.929863047611582 |
| chr4-57420423-57420924 | 8 | 0.959990192763582 |
| chr4-105952651-105953152 | 3 | 0.99593553545041 |
| chr6-20220580-20221081 | 8 | 0.987478348496242 |
| chr5-68196806-68197307 | 6 | 0.975001875767592 |
| chr1-11530924-11531425 | 3 | 0.99593553545041 |
| chr1-191808546-191809047 | 8 | 0.913113703738281 |
| chr6-68617583-68618084 | 3 | 0.993900356757547 |
| chr16-77929299-77929800 | 2 | 0.854824090447686 |
| chr6-131095637-131096138 | 5 | 0.902558386143023 |
| chr6-51901871-51902372 | 8 | 0.968460537381743 |
| chr2-29149042-29149543 | 1 | 0.877272346016801 |
| chr2-219918184-219918685 | 8 | 0.862958484915253 |
| chr14-1865368-1865869 | 8 | 0.990128884235818 |
| chr1-224236036-224236537 | 8 | 0.848202102434445 |
| chr2-134594621-134595122 | 4 | 0.967727152376057 |
| chr3-42336389-42336890 | 3 | 0.99593553545041 |
| chr14-56826174-56826675 | 8 | 0.929702368889548 |
| chr1-151988139-151988640 | 3 | 0.996870320260068 |
| chr2-53209399-53209900 | 2 | 0.950735855932072 |
| chr18-67368795-67369296 | 8 | 0.992459323919938 |
| chr17-40118497-40118998 | 3 | 0.988194459009504 |
| chr4-83266255-83266756 | 5 | 0.979840400366239 |
| chr12-1299250-1299751 | 8 | 0.829277382621981 |

| | | |
|---|---|---|
| chr10-65626824-65627325 | 8 | 0.884307712979504 |
| chr11-63050863-63051364 | 2 | 0.857594470060037 |
| chr17-65103299-65103800 | 3 | 0.997140368803633 |
| chr4-163652187-163652688 | 5 | 0.983919978840557 |
| chr10-60415808-60416309 | 5 | 0.927465741293874 |
| chr17-77778770-77779271 | 8 | 0.993493681128311 |
| chr7-123441755-123442256 | 8 | 0.935160558638335 |
| chr5-121488909-121489410 | 5 | 0.965608736832821 |
| chr2-87966878-87967379 | 8 | 0.939684965215197 |
| chr5-150917187-150917688 | 6 | 0.978627711481427 |
| chr5-121474027-121474528 | 8 | 0.815568514837701 |
| chr3-127118924-127119425 | 5 | 0.941278245382898 |
| chr4-146872839-146873340 | 5 | 0.885988926762857 |
| chr4-158318568-158319069 | 6 | 0.875992347661121 |
| chr10-35987807-35988308 | 3 | 0.978095817925989 |
| chr2-240238453-240238954 | 1 | 0.949250847835032 |
| chr8-7992604-7993105 | 3 | 0.996887519527629 |
| chr11-31269487-31269988 | 2 | 0.93850448804877 |
| chr4-104686297-104686798 | 6 | 0.960918893338851 |
| chr2-81344715-81345216 | 3 | 0.987574419788903 |
| chr9-17711377-17711878 | 3 | 0.852367167611758 |
| chr14-12211215-12211716 | 1 | 0.912841829346531 |
| chr3-1547590-1548091 | 2 | 0.847189687494299 |
| chrX-32944117-32944618 | 6 | 0.953238195177898 |
| chr11-34502861-34503362 | 1 | 0.913992853570173 |
| chrX-97018370-97018871 | 8 | 0.993493681128311 |
| chr11-33434421-33434922 | 3 | 0.959354374110837 |
| chr8-61319634-61320135 | 5 | 0.945307686499269 |
| chr4-132041799-132042300 | 8 | 0.992348615092774 |
| chr12-26144120-26144621 | 5 | 0.989667416037098 |
| chr1-252275082-252275583 | 8 | 0.992745819206162 |
| chr5-51430991-51431492 | 8 | 0.870122564152015 |
| chr4-129007705-129008206 | 3 | 0.99416114756515 |
| chr3-32015345-32015846 | 6 | 0.991206925481043 |
| chr10-100013790-100014291 | 8 | 0.886274555304731 |
| chr6-50451265-50451766 | 3 | 0.995446033579843 |
| chr2-173272434-173272935 | 5 | 0.972909733562985 |
| chr8-54931511-54932012 | 4 | 0.880827796014407 |
| chr10-45333486-45333987 | 8 | 0.849141347596959 |
| chr5-27825251-27825752 | 8 | 0.952872599479799 |
| chr2-96423840-96424341 | 8 | 0.995242226511646 |
| chr4-97889112-97889613 | 3 | 0.99593553545041 |
| chr2-59206262-59206763 | 3 | 0.982965484689979 |

| | | |
|---|---|---|
| chr3-18438805-18439306 | 8 | 0.953718204860825 |
| chr2-527915-528416 | 3 | 0.99593553545041 |
| chr17-63903604-63904105 | 3 | 0.979558690754411 |
| chr5-165080538-165081039 | 8 | 0.910006618893222 |
| chr4-46245101-46245602 | 6 | 0.944558099024833 |
| chr9-107009678-107010179 | 3 | 0.918461074779312 |
| chr14-71939309-71939810 | 8 | 0.994466262502284 |
| chr1-126695171-126695672 | 3 | 0.859725816046756 |
| chr6-55443651-55444152 | 3 | 0.933739614901685 |
| chr1-213903990-213904491 | 3 | 0.99593553545041 |
| chr2-114298634-114299135 | 3 | 0.984742057185845 |
| chr1-225723068-225723569 | 3 | 0.945877576804028 |
| chr5-34319171-34319672 | 3 | 0.982000324321146 |
| chr2-149575039-149575540 | 6 | 0.990851359070655 |
| chr1-108557356-108557857 | 5 | 0.835235712291012 |
| chr1-184725113-184725614 | 8 | 0.927774568116386 |
| chr2-27152554-27153055 | 4 | 0.885966020885834 |
| chr16-30979485-30979986 | 3 | 0.995502885898944 |
| chr2-209901435-209901936 | 8 | 0.82924394382352 |
| chr6-40054028-40054529 | 6 | 0.939976630996193 |
| chr2-79661783-79662284 | 3 | 0.824697368550448 |
| chr5-163758795-163759296 | 3 | 0.996777995124012 |
| chr2-166607016-166607517 | 3 | 0.995672815894577 |
| chr3-140951790-140952291 | 4 | 0.855279414578836 |
| chr14-3505671-3506172 | 1 | 0.923782137459299 |
| chr11-82323405-82323906 | 4 | 0.865470222525141 |
| chr12-22589789-22590290 | 4 | 0.821466953533679 |
| chr5-124137732-124138233 | 3 | 0.995293227196014 |
| chr1-16837509-16838010 | 5 | 0.958548537867619 |
| chr9-72612454-72612955 | 8 | 0.821073400117077 |
| chr17-15553961-15554462 | 5 | 0.995647938869809 |
| chr2-196966458-196966959 | 8 | 0.844278831182556 |
| chr6-70888612-70889113 | 7 | 0.82539581166189 |
| chr5-107993183-107993684 | 6 | 0.911252080670569 |
| chr8-72389916-72390417 | 5 | 0.988459184858326 |
| chr8-72489172-72489673 | 5 | 0.867405798848836 |
| chr14-34300426-34300927 | 3 | 0.99689857652917 |
| chr17-64055436-64055937 | 3 | 0.996865285585664 |
| chr17-52711495-52711996 | 8 | 0.888826984970114 |
| chr8-51140226-51140727 | 8 | 0.993493681128311 |
| chr4-87383398-87383899 | 3 | 0.954728281975931 |
| chr12-26210552-26211053 | 3 | 0.976571937360667 |
| chr7-133677400-133677901 | 4 | 0.956483005264272 |

| | | |
|---|---|---|
| chr13-49965272-49965773 | 5 | 0.987073151319587 |
| chr2-184240851-184241352 | 5 | 0.989095122215376 |
| chr1-221095282-221095783 | 5 | 0.985294819336759 |
| chr19-56124023-56124524 | 5 | 0.980957460784852 |
| chr9-96727450-96727951 | 6 | 0.993218244958568 |
| chr1-52963259-52963760 | 5 | 0.982446185389934 |
| chr9-61071844-61072345 | 8 | 0.958311943255975 |
| chr18-38396374-38396875 | 6 | 0.983505925789762 |
| chr15-3375113-3375614 | 3 | 0.972370539200919 |
| chr13-902037-902538 | 5 | 0.983919978840557 |
| chr4-161744196-161744697 | 3 | 0.9718404337505 |
| chr6-35735248-35735749 | 3 | 0.982900559378584 |
| chr3-108306208-108306709 | 5 | 0.991440678740938 |
| chr19-11111213-11111714 | 8 | 0.82966740114276 |
| chr16-8425037-8425538 | 3 | 0.901723397065403 |
| chr14-13653169-13653670 | 3 | 0.99655688302576 |
| chr3-149203493-149203994 | 3 | 0.995040318300007 |
| chr9-59428380-59428881 | 8 | 0.931478895441819 |
| chr5-150883302-150883803 | 5 | 0.98989525240303 |
| chr6-49244335-49244836 | 8 | 0.980409265752807 |
| chr2-197328793-197329294 | 8 | 0.936032964493625 |
| chr6-87578326-87578827 | 8 | 0.991125374090313 |
| chr6-130546056-130546557 | 6 | 0.995360893391302 |
| chr10-1922292-1922793 | 8 | 0.975346074341037 |
| chr18-71329048-71329549 | 6 | 0.947233623904586 |
| chr13-77604396-77604897 | 4 | 0.96155442969504 |
| chr2-29653443-29653944 | 4 | 0.904182878395857 |
| chr13-96330474-96330975 | 5 | 0.929636508170428 |
| chr5-10990402-10990903 | 3 | 0.965277308158837 |
| chr17-34405731-34406232 | 8 | 0.921344444043496 |
| chr18-73109891-73110392 | 6 | 0.86496819575086 |
| chr2-27009304-27009805 | 3 | 0.996860707488835 |
| chr7-65817660-65818161 | 3 | 0.865765650620152 |
| chr1-171492908-171493409 | 3 | 0.990744160759904 |
| chr3-127143681-127144182 | 3 | 0.995890865144074 |
| chr4-133633119-133633620 | 8 | 0.903087390552315 |
| chr14-37611586-37612087 | 3 | 0.99593553545041 |
| chr2-192304978-192305479 | 2 | 0.856092904771041 |
| chr11-75156851-75157352 | 3 | 0.996881481934351 |
| chr4-154431093-154431594 | 8 | 0.989867900994858 |
| chr4-28326721-28327222 | 4 | 0.89077576463854 |
| chr9-81291047-81291548 | 2 | 0.959493506286909 |
| chr2-137380489-137380990 | 5 | 0.989508118859161 |

| | | |
|---|---|---|
| chr4-91024636-91025137 | 6 | 0.993178675731963 |
| chr3-154256990-154257491 | 8 | 0.984520007876621 |
| chr1-221520663-221521164 | 6 | 0.963908851031901 |
| chr11-81019474-81019975 | 8 | 0.853527790727558 |
| chr1-45804476-45804977 | 2 | 0.842856827427708 |
| chr5-63633048-63633549 | 8 | 0.975778091791736 |
| chr3-37085230-37085731 | 8 | 0.989708690555973 |
| chr15-77110240-77110741 | 6 | 0.878758385236266 |
| chr7-37590768-37591269 | 3 | 0.996861487624464 |
| chr4-13444032-13444533 | 3 | 0.996780562862761 |
| chr16-32054712-32055213 | 6 | 0.997595567114982 |
| chr4-160708807-160709308 | 4 | 0.94251126459357 |
| chr17-8725840-8726341 | 5 | 0.96721846464012 |
| chr4-65309619-65310120 | 3 | 0.991857967832128 |
| chr4-7995748-7996249 | 6 | 0.991620747601191 |
| chr1-127817217-127817718 | 8 | 0.921939340122317 |
| chr2-206666145-206666646 | 3 | 0.96507100479441 |
| chr16-513384-513885 | 4 | 0.932955478994572 |
| chr6-121314229-121314730 | 3 | 0.988754518372958 |
| chr13-69441320-69441821 | 3 | 0.915909924693837 |
| chr1-87066642-87067143 | 8 | 0.986735038357052 |
| chr19-18505156-18505657 | 8 | 0.896290718390973 |
| chr17-68888976-68889477 | 3 | 0.995255025966654 |
| chr2-220425732-220426233 | 7 | 0.929531662380819 |
| chr11-31001708-31002209 | 6 | 0.97915135311479 |
| chr2-160630337-160630838 | 6 | 0.973790892428284 |
| chr18-25378448-25378949 | 8 | 0.942364535590559 |
| chr16-10433046-10433547 | 6 | 0.954873256595071 |
| chr17-69701473-69701974 | 1 | 0.916628629797512 |
| chr2-31448020-31448521 | 2 | 0.820669584963708 |
| chr10-66514222-66514723 | 5 | 0.988465268211314 |
| chr3-122022163-122022664 | 3 | 0.973940974666521 |
| chr16-23032411-23032912 | 3 | 0.99593553545041 |
| chr3-134089849-134090350 | 8 | 0.989307167762968 |
| chrX-7692770-7693271 | 6 | 0.991206925481043 |
| chr10-41418087-41418588 | 1 | 0.853720439682541 |
| chr4-82846506-82847007 | 5 | 0.966190410528458 |
| chr15-34051508-34052009 | 3 | 0.974840986275937 |
| chrX-28039510-28040011 | 5 | 0.983919978840557 |
| chr1-7372685-7373186 | 6 | 0.992180148191733 |
| chr7-8627894-8628395 | 8 | 0.841269479127487 |
| chr14-81614058-81614559 | 5 | 0.969506652490432 |
| chr4-43487157-43487658 | 6 | 0.991432358635437 |

| | | |
|---|---|---|
| chr9-73395792-73396293 | 8 | 0.892233392342127 |
| chr1-154816826-154817327 | 5 | 0.942310568692291 |
| chr8-42820334-42820835 | 5 | 0.983919978840557 |
| chr1-223747331-223747832 | 8 | 0.983909590523229 |
| chr16-56275008-56275509 | 6 | 0.842753239769554 |
| chr12-45235277-45235778 | 8 | 0.981098454169334 |
| chr19-6048916-6049417 | 6 | 0.977869905191459 |
| chr5-53815687-53816188 | 5 | 0.840064927048903 |
| chr9-7329402-7329903 | 6 | 0.979409568327597 |
| chr8-104409039-104409540 | 4 | 0.842850297013496 |
| chr2-226982271-226982772 | 8 | 0.883755649741442 |
| chr2-245192924-245193425 | 6 | 0.907899395633642 |
| chr11-11977738-11978239 | 6 | 0.992601637312221 |
| chr13-30333057-30333558 | 1 | 0.91289142671838 |
| chr10-62531408-62531909 | 2 | 0.900579167640428 |
| chr7-99220879-99221380 | 7 | 0.888458425764033 |
| chr11-20808746-20809247 | 3 | 0.99593553545041 |
| chr1-6797681-6798182 | 8 | 0.815287388897371 |
| chr10-33904720-33905221 | 1 | 0.823657372634536 |
| chr5-101080291-101080792 | 8 | 0.960008543855244 |
| chr14-56977647-56978148 | 4 | 0.955895546250423 |
| chr20-44307395-44307896 | 6 | 0.991206925481043 |
| chr7-72684775-72685276 | 6 | 0.943665975925239 |
| chr7-22603456-22603957 | 2 | 0.869583037785241 |
| chr1-96921668-96922169 | 5 | 0.977499542366359 |
| chr6-138387341-138387842 | 7 | 0.840564760012334 |
| chr1-120542385-120542886 | 7 | 0.927733532367141 |
| chr10-20269483-20269984 | 5 | 0.820444594116453 |
| chr18-72445426-72445927 | 8 | 0.994988275665063 |
| chr17-56723592-56724093 | 4 | 0.818248613514193 |
| chr15-32720257-32720758 | 8 | 0.956301198967046 |
| chr9-88360448-88360949 | 2 | 0.890976251659595 |
| chr2-219722491-219722992 | 8 | 0.994956004735606 |
| chr20-12736533-12737034 | 5 | 0.981920696606733 |
| chr14-35306138-35306639 | 8 | 0.975475775184662 |
| chrX-54963281-54963782 | 5 | 0.983919978840557 |
| chr5-1557217-1557718 | 6 | 0.97586937143649 |
| chr6-52985067-52985568 | 5 | 0.869043499028272 |
| chrX-127095013-127095514 | 6 | 0.967636143236886 |
| chr15-4871085-4871586 | 6 | 0.976589482510641 |
| chr1-124892201-124892702 | 6 | 0.97878587863442 |
| chr8-58268774-58269275 | 4 | 0.910446862224278 |
| chrX-11441884-11442385 | 1 | 0.902830536880742 |

| | | |
|---|---|---|
| chr3-90655545-90656046 | 6 | 0.980534787327377 |
| chr16-47050995-47051496 | 6 | 0.991206925481043 |
| chr3-123806118-123806619 | 8 | 0.860184394949114 |
| chr9-88926629-88927130 | 8 | 0.993195362626959 |
| chr8-11079737-11080238 | 8 | 0.884935777437582 |
| chr15-60849557-60850058 | 6 | 0.991206925481043 |
| chr18-72655734-72656235 | 3 | 0.976784817340053 |
| chr5-152778865-152779366 | 4 | 0.884270802097004 |
| chr17-53451579-53452080 | 3 | 0.892321733932407 |
| chr1-11536754-11537255 | 8 | 0.995143712195928 |
| chr16-23052966-23053467 | 8 | 0.909778646366924 |
| chr18-224112-224613 | 3 | 0.995642522914036 |
| chr13-92899616-92900117 | 5 | 0.839291159082842 |
| chr14-73749153-73749654 | 6 | 0.969117595945754 |
| chr15-66215665-66216166 | 6 | 0.971696642443934 |
| chr7-22907577-22908078 | 8 | 0.882363154474309 |
| chr18-35736913-35737414 | 6 | 0.9877848026967 |
| chr5-71628955-71629456 | 5 | 0.95177817642912 |
| chr1-130606612-130607113 | 6 | 0.974953248925112 |
| chr2-141656426-141656927 | 8 | 0.834196720882926 |
| chr2-143833775-143834276 | 7 | 0.81467217419402 |
| chr2-241426369-241426870 | 5 | 0.946839929092533 |
| chr13-43617864-43618365 | 4 | 0.969368264699795 |
| chr1-23069657-23070158 | 5 | 0.888386705563906 |
| chr7-11738493-11738994 | 8 | 0.881536367850057 |
| chr17-39729786-39730287 | 3 | 0.976944520252684 |
| chr20-10750146-10750647 | 3 | 0.976168546512939 |
| chr13-38963843-38964344 | 5 | 0.989675832471756 |
| chr6-123805586-123806087 | 8 | 0.993493681128311 |
| chr13-66007900-66008401 | 3 | 0.936567427023882 |
| chr16-50431272-50431773 | 3 | 0.995320945494369 |
| chr1-85900987-85901488 | 8 | 0.828241752766722 |
| chr1-103409149-103409650 | 3 | 0.976585314381061 |
| chr16-38619164-38619665 | 6 | 0.981791562474542 |
| chr13-74794573-74795074 | 8 | 0.892005293122617 |
| chr17-52193868-52194369 | 8 | 0.883694582882268 |
| chr8-40955194-40955695 | 5 | 0.991148815516101 |
| chr6-71847895-71848396 | 1 | 0.939541748817275 |
| chr5-60508806-60509307 | 3 | 0.996864884134633 |
| chr15-37040394-37040895 | 3 | 0.867827860546663 |
| chr17-78223232-78223733 | 3 | 0.994848581527115 |
| chr7-72011743-72012244 | 3 | 0.987339440198957 |
| chr14-93930475-93930976 | 6 | 0.911772692767366 |

| | | |
|---|---|---|
| chr14-2397669-2398170 | 8 | 0.91057187470908 |
| chr2-210821036-210821537 | 3 | 0.996555330341232 |
| chr1-49238457-49238958 | 7 | 0.933508010873637 |
| chr2-104741318-104741819 | 4 | 0.913404804980835 |
| chr16-58112996-58113497 | 7 | 0.866026468759015 |
| chr9-27017836-27018337 | 3 | 0.984042987912035 |
| chr5-60960771-60961272 | 5 | 0.927102782942414 |
| chr18-39782277-39782778 | 6 | 0.965183526118535 |
| chr10-38048695-38049196 | 8 | 0.979252536245195 |
| chr6-5680809-5681310 | 8 | 0.94599130954064 |
| chr9-98248233-98248734 | 8 | 0.838553217437832 |
| chr6-72911366-72911867 | 5 | 0.915750652787726 |
| chr5-104112274-104112775 | 8 | 0.993493681128311 |
| chr1-178653077-178653578 | 6 | 0.95456427702196 |
| chr19-7494654-7495155 | 3 | 0.834387751018595 |
| chr3-60505334-60505835 | 2 | 0.969388967887219 |
| chr3-13178804-13179305 | 8 | 0.899574975048733 |
| chr3-140791674-140792175 | 2 | 0.836300724124129 |
| chr3-120669645-120670146 | 8 | 0.993493681128311 |
| chrX-132969032-132969533 | 7 | 0.834542116040536 |
| chr5-18778693-18779194 | 8 | 0.848551122805855 |
| chr1-89619195-89619696 | 6 | 0.991206925481043 |
| chr10-33322974-33323475 | 3 | 0.996494852306111 |
| chr11-52612978-52613479 | 6 | 0.966808720190087 |
| chr12-21333875-21334376 | 6 | 0.817486772916858 |
| chr14-43245612-43246113 | 8 | 0.981009768116322 |
| chr2-170300925-170301426 | 7 | 0.936977958037878 |
| chr11-84050642-84051143 | 3 | 0.964842783730147 |
| chr1-167722179-167722680 | 3 | 0.996270714867447 |
| chr18-47203345-47203846 | 8 | 0.986818294129267 |
| chr2-27136083-27136584 | 2 | 0.867356208280254 |
| chr18-7454864-7455365 | 3 | 0.988074342863458 |
| chr2-223013612-223014113 | 6 | 0.96841241028248 |
| chr1-246943585-246944086 | 8 | 0.957983905571056 |
| chr3-2682236-2682737 | 5 | 0.983919978840557 |
| chr17-46883705-46884206 | 6 | 0.992483359533977 |
| chr2-60017398-60017899 | 3 | 0.997552368209919 |
| chr7-89698544-89699045 | 4 | 0.930783683488106 |
| chr6-45588891-45589392 | 3 | 0.991196103815202 |
| chr6-51792439-51792940 | 8 | 0.919457575933744 |
| chr9-66109271-66109772 | 6 | 0.973181046558085 |
| chr2-137020751-137021252 | 6 | 0.930279824954786 |
| chr3-165852622-165853123 | 4 | 0.851619899135337 |

| | | |
|---|---|---|
| chr2-153579975-153580476 | 3 | 0.997342576802159 |
| chr6-107102744-107103245 | 8 | 0.908430248312848 |
| chr2-30260581-30261082 | 8 | 0.935861421098087 |
| chr16-55841629-55842130 | 6 | 0.981223613245203 |
| chr2-45587613-45588114 | 8 | 0.823668292563383 |
| chr18-62123410-62123911 | 8 | 0.889002815098895 |
| chr14-95954793-95955294 | 2 | 0.918567311258606 |
| chr15-7619653-7620154 | 5 | 0.938875026466139 |
| chr9-106265822-106266323 | 7 | 0.924860746916244 |
| chr16-10518885-10519386 | 4 | 0.94602284937221 |
| chr2-46755444-46755945 | 8 | 0.882853905884699 |
| chr20-8596789-8597290 | 8 | 0.836061765594448 |
| chr16-31802564-31803065 | 6 | 0.963089942633183 |
| chr5-119915100-119915601 | 8 | 0.877237168868484 |
| chr4-131864600-131865101 | 5 | 0.986956521177615 |
| chr6-21685370-21685871 | 7 | 0.909907704382567 |
| chr18-66848875-66849376 | 6 | 0.894610408167842 |
| chr2-75972738-75973239 | 8 | 0.993493681128311 |
| chr6-105396597-105397098 | 8 | 0.963284064528234 |
| chr2-145282091-145282592 | 7 | 0.923424197323661 |
| chr3-88904329-88904830 | 3 | 0.99569288723196 |
| chr19-555768-556269 | 8 | 0.831989380585827 |
| chr9-85215116-85215617 | 8 | 0.993493681128311 |
| chr3-157415725-157416226 | 6 | 0.971440660397298 |
| chr7-76513188-76513689 | 3 | 0.99593553545041 |
| chr4-147274013-147274514 | 6 | 0.974129983195711 |
| chr10-51621029-51621530 | 7 | 0.95178386028819 |
| chr2-191436015-191436516 | 8 | 0.975363759480483 |
| chrX-151021460-151021961 | 3 | 0.998253820661255 |
| chr1-240442014-240442515 | 4 | 0.833856117407564 |
| chr15-81886187-81886688 | 3 | 0.994276213019206 |
| chr13-39510008-39510509 | 8 | 0.88622145021517 |
| chr11-35401518-35402019 | 3 | 0.994637605329851 |
| chr4-159932080-159932581 | 8 | 0.995117439303815 |
| chr7-46754617-46755118 | 8 | 0.992864655142423 |
| chr5-110769331-110769832 | 1 | 0.966106737298632 |
| chr1-240800438-240800939 | 1 | 0.90512755742852 |
| chr2-115421579-115422080 | 4 | 0.884572307383732 |
| chr13-35389330-35389831 | 3 | 0.996804079724976 |
| chr15-47998731-47999232 | 7 | 0.834456235078429 |
| chr9-81074288-81074789 | 3 | 0.992289806540502 |
| chr5-121532607-121533108 | 8 | 0.986375673124173 |
| chr16-16915893-16916394 | 6 | 0.865912390631911 |

| | | |
|---|---|---|
| chr1-236995270-236995771 | 3 | 0.995935535545041 |
| chr18-41585761-41586262 | 2 | 0.943229069532699 |
| chr17-63525455-63525956 | 3 | 0.995379163302737 |
| chr1-92039192-92039693 | 6 | 0.979653956014893 |
| chr4-62622539-62623040 | 6 | 0.991206925481043 |
| chr9-74944272-74944773 | 8 | 0.951509482579387 |
| chr10-21175666-21176167 | 3 | 0.996088941750714 |
| chr5-134193279-134193780 | 6 | 0.991260263662952 |
| chr17-22071797-22072298 | 8 | 0.950244471470016 |
| chr13-82909296-82909797 | 5 | 0.989158401694397 |
| chr4-112625872-112626373 | 3 | 0.996289421448298 |
| chr17-17298840-17299341 | 3 | 0.879926947882565 |
| chr10-55051979-55052480 | 4 | 0.89505157531503 |
| chr8-12067159-12067660 | 5 | 0.988703817209031 |
| chr3-96020628-96021129 | 6 | 0.974803028269926 |
| chr11-65156066-65156567 | 3 | 0.957277913504434 |
| chr10-106676869-106677370 | 4 | 0.968505856584262 |
| chr5-5382608-5383109 | 8 | 0.960771596832643 |
| chr3-93174375-93174876 | 8 | 0.994619546351472 |
| chr9-89679896-89680397 | 8 | 0.96799811206907 |
| chr5-151800764-151801265 | 3 | 0.963119809803599 |
| chr6-138260337-138260838 | 8 | 0.982306904737349 |
| chr10-71024033-71024534 | 5 | 0.910268672650313 |
| chr17-52974526-52975027 | 1 | 0.855724130741507 |
| chr3-14690343-14690844 | 2 | 0.961780557962408 |
| chr4-145844650-145845151 | 2 | 0.898482852510117 |
| chr1-192751329-192751830 | 8 | 0.990692907503224 |
| chr10-10856020-10856521 | 3 | 0.908203418407557 |
| chrX-22209207-22209708 | 1 | 0.890479082205539 |
| chr14-33924475-33924976 | 3 | 0.99537590070444 |
| chr6-16837328-16837829 | 3 | 0.838506634870642 |
| chr1-196245346-196245847 | 5 | 0.986154339619484 |
| chr18-69739167-69739668 | 8 | 0.990218140980915 |
| chr17-21730173-21730674 | 3 | 0.902517309335264 |
| chr8-111220733-111221234 | 2 | 0.887158901529389 |
| chr4-147357832-147358333 | 6 | 0.993650315983508 |
| chrX-73997000-73997501 | 6 | 0.991206925481043 |
| chr10-60354859-60355360 | 3 | 0.984379054858119 |
| chr3-28463484-28463985 | 5 | 0.8754530727769 |
| chr1-190079517-190080018 | 3 | 0.998010347936834 |
| chr14-81110112-81110613 | 8 | 0.978645415010594 |
| chrX-112839743-112840244 | 6 | 0.991206925481043 |
| chr13-43648938-43649439 | 8 | 0.993540378466341 |

| | | |
|---|---|---|
| chr10-36750453-36750954 | 3 | 0.996654648009155 |
| chr6-115806046-115806547 | 3 | 0.849881871245217 |
| chr6-125800417-125800918 | 8 | 0.941385074627779 |
| chr1-207373065-207373566 | 4 | 0.846616912680852 |
| chr8-64705555-64706056 | 5 | 0.858238614116855 |
| chr19-44618241-44618742 | 8 | 0.864462809108284 |
| chr9-26203394-26203895 | 5 | 0.917554183027839 |
| chrX-25050665-25051166 | 4 | 0.838088123614196 |
| chr9-63193515-63194016 | 6 | 0.888113769202161 |
| chr5-105781470-105781971 | 6 | 0.982294689153672 |
| chr15-83620226-83620727 | 5 | 0.983919978840557 |
| chr20-11231549-11232050 | 3 | 0.898787551696604 |
| chr12-25090006-25090507 | 8 | 0.993493681128311 |
| chr10-90765672-90766173 | 3 | 0.996863425805592 |
| chr1-167034280-167034781 | 3 | 0.920543426208829 |
| chr8-43090083-43090584 | 7 | 0.854730019703967 |
| chr4-45079546-45080047 | 3 | 0.996583216587041 |
| chr8-96699385-96699886 | 6 | 0.93180959150665 |
| chr2-221364322-221364823 | 8 | 0.893195185079114 |
| chr19-38255098-38255599 | 3 | 0.993973024170784 |
| chr12-26600780-26601281 | 4 | 0.816820307647075 |
| chr16-4597179-4597680 | 5 | 0.983068429614373 |
| chr7-8999175-8999676 | 8 | 0.981380847326418 |
| chr18-6619895-6620396 | 3 | 0.997254770815554 |
| chr16-34283690-34284191 | 3 | 0.974661527771438 |
| chr8-79104716-79105217 | 4 | 0.845878378017165 |
| chr9-98594884-98595385 | 4 | 0.856480643149041 |
| chr10-78770013-78770514 | 6 | 0.981991644976612 |
| chr7-19815570-19816071 | 8 | 0.995170844119766 |
| chr3-107160905-107161406 | 1 | 0.868452881957433 |
| chr6-109549053-109549554 | 3 | 0.995760801709403 |
| chr12-24967296-24967797 | 4 | 0.961912196566357 |
| chr2-118892720-118893221 | 6 | 0.940237622462515 |
| chr17-34563376-34563877 | 2 | 0.902654300468597 |
| chr7-8604000-8604501 | 8 | 0.890514947799196 |
| chr20-13276523-13277024 | 4 | 0.852376230847268 |
| chr16-12311315-12311816 | 5 | 0.90266591361004 |
| chr17-54507712-54508213 | 8 | 0.930792475172554 |
| chr7-57543036-57543537 | 5 | 0.84985259397602 |
| chr9-4829248-4829749 | 7 | 0.843293440179036 |
| chr16-51158313-51158814 | 3 | 0.996541911333753 |
| chr8-108573198-108573699 | 3 | 0.826296073556683 |
| chr18-48424981-48425482 | 5 | 0.971448501537596 |

| | | |
|---|---|---|
| chr9-77649959-77650460 | 8 | 0.894846278654669 |
| chr17-23307003-23307504 | 1 | 0.921383087455174 |
| chr1-173356869-173357370 | 6 | 0.977258689130233 |
| chr16-33606845-33607346 | 8 | 0.993493681128311 |
| chr10-11717077-11717578 | 8 | 0.840519122860032 |
| chr11-24556477-24556978 | 3 | 0.987550970285603 |
| chr11-28629311-28629812 | 3 | 0.99593553545041 |
| chr18-51005301-51005802 | 3 | 0.99593553545041 |
| chr17-46884735-46885236 | 6 | 0.978832203273259 |
| chr17-85836342-85836843 | 3 | 0.942690808997203 |
| chr1-232360570-232361071 | 6 | 0.817847825983487 |
| chr20-46338315-46338816 | 2 | 0.840945774167613 |
| chr4-75021346-75021847 | 3 | 0.992846840196127 |
| chr3-165815208-165815709 | 6 | 0.991206925481043 |
| chr3-93146671-93147172 | 4 | 0.880302074125962 |
| chr3-166464518-166465019 | 7 | 0.874201064570651 |
| chr1-36554051-36554552 | 6 | 0.954392647820098 |
| chr2-113322550-113323051 | 5 | 0.980633548807044 |
| chr5-132656823-132657324 | 5 | 0.984951475655139 |
| chr6-59825062-59825563 | 3 | 0.998199063286559 |
| chr6-84067349-84067850 | 2 | 0.947003844876348 |
| chr1-12548727-12549228 | 5 | 0.972661315028768 |
| chr2-173041868-173042369 | 6 | 0.990840480698981 |
| chr4-24974491-24974992 | 3 | 0.987526365467747 |
| chr4-132727531-132728032 | 1 | 0.862351853609553 |
| chr2-230357927-230358428 | 8 | 0.986361830111035 |
| chr13-95238692-95239193 | 6 | 0.992618622252053 |
| chr7-34377677-34378178 | 7 | 0.818702966907194 |
| chr12-6773478-6773979 | 6 | 0.983841837756519 |
| chr14-94393402-94393903 | 5 | 0.857031549783701 |
| chr8-104322964-104323465 | 8 | 0.829392852269299 |
| chr9-58470617-58471118 | 3 | 0.99685528402667 |
| chr8-37394991-37395492 | 5 | 0.94231220578614 |
| chr8-926196-926697 | 6 | 0.993196351202581 |
| chr17-84293698-84294199 | 4 | 0.87638452162152 |
| chr15-79538611-79539112 | 8 | 0.871536178210911 |
| chr4-74167862-74168363 | 3 | 0.8148427767887 |
| chr2-246049440-246049941 | 5 | 0.967271035154252 |
| chr5-24551876-24552377 | 5 | 0.990358810042229 |
| chr13-37820380-37820881 | 5 | 0.823725825487533 |
| chr15-76820051-76820552 | 6 | 0.991359955265644 |
| chr20-32687452-32687953 | 5 | 0.966916319733644 |
| chr4-179141042-179141543 | 7 | 0.919131944663405 |

| | | |
|---|---|---|
| chr4-69709006-69709507 | 3 | 0.993009785243468 |
| chr3-86722181-86722682 | 5 | 0.97290874083926 |
| chr5-118113254-118113755 | 8 | 0.843345863279062 |
| chr4-58529866-58530367 | 8 | 0.934374465723448 |
| chr4-84398238-84398739 | 3 | 0.864559066769803 |
| chr7-96657084-96657585 | 6 | 0.994688791169747 |
| chr1-207931046-207931547 | 1 | 0.965746925408173 |
| chr5-113917221-113917722 | 8 | 0.91935847909025 |
| chr2-45569564-45570065 | 6 | 0.985797997927368 |
| chr1-164995288-164995789 | 7 | 0.870458082166907 |
| chr17-66139278-66139779 | 8 | 0.907079504268039 |
| chrX-20230452-20230953 | 8 | 0.817119309495856 |
| chr14-66266722-66267223 | 3 | 0.995109473121581 |
| chr9-60623881-60624382 | 8 | 0.990654108065944 |
| chr6-119021914-119022415 | 8 | 0.896472069265883 |
| chr18-21035404-21035905 | 6 | 0.909017082149806 |
| chr18-78131884-78132385 | 3 | 0.933682898479005 |
| chr4-129625707-129626208 | 5 | 0.985088640760035 |
| chr10-5810857-5811358 | 6 | 0.905933646798959 |
| chr13-83083592-83084093 | 5 | 0.936302840826569 |
| chr2-77742306-77742807 | 2 | 0.818668100156213 |
| chr15-54828555-54829056 | 1 | 0.980765362449379 |
| chr13-50627856-50628357 | 5 | 0.884287533419057 |
| chr3-69666307-69666808 | 3 | 0.987341819412781 |
| chr5-6265729-6266230 | 3 | 0.998293123750517 |
| chr17-24664116-24664617 | 3 | 0.903221128447175 |
| chr2-170381678-170382179 | 7 | 0.973420838839592 |
| chr3-21741302-21741803 | 5 | 0.967881990056233 |
| chr2-223102628-223103129 | 3 | 0.965229678423043 |
| chr1-196225524-196226025 | 8 | 0.983026091792841 |
| chr3-29277235-29277736 | 5 | 0.954006258138196 |
| chr14-9093709-9094210 | 5 | 0.975524560494812 |
| chr1-117798244-117798745 | 2 | 0.925450861021332 |
| chr13-61980135-61980636 | 3 | 0.99593553545041 |
| chr9-16542790-16543291 | 3 | 0.974309056646099 |
| chr1-151229653-151230154 | 8 | 0.995247071504269 |
| chr6-88550975-88551476 | 3 | 0.998460369689049 |
| chr1-11489524-11490025 | 6 | 0.991206925481043 |
| chr2-104736146-104736647 | 6 | 0.991336170829158 |
| chr10-47512162-47512663 | 6 | 0.936784884495486 |
| chr14-58877382-58877883 | 3 | 0.992671465777398 |
| chr6-58057416-58057917 | 3 | 0.892363779707749 |
| chr2-216228697-216229198 | 4 | 0.881707019660958 |

| | | |
|---|---|---|
| chr12-5384794-5385295 | 5 | 0.971702677366517 |
| chr14-86130465-86130966 | 8 | 0.968177138662857 |
| chr3-47214298-47214799 | 6 | 0.992690152841551 |
| chr17-35137326-35137827 | 8 | 0.939823384801688 |
| chr6-28761928-28762429 | 1 | 0.888827080441819 |
| chr2-224209347-224209848 | 8 | 0.993493681128311 |
| chr3-79443616-79444117 | 8 | 0.944774184728567 |
| chr14-34285689-34286190 | 5 | 0.989103242314085 |
| chr9-64447325-64447826 | 3 | 0.985897233293319 |
| chr14-94164954-94165455 | 5 | 0.916468076543677 |
| chr1-91336514-91337015 | 3 | 0.935520172843245 |
| chr7-117028790-117029291 | 8 | 0.841465621000247 |
| chr8-92232131-92232632 | 3 | 0.978519178200773 |
| chr5-99722367-99722868 | 1 | 0.901283851406012 |
| chr1-133748383-133748884 | 5 | 0.991007897431889 |
| chr1-19018221-19018722 | 3 | 0.889302003941101 |
| chr7-69186100-69186601 | 8 | 0.869013738863257 |
| chrX-131893539-131894040 | 6 | 0.991206925481043 |
| chr13-77742245-77742746 | 3 | 0.996672148745067 |
| chr2-42486438-42486939 | 8 | 0.851550767874732 |
| chr2-247600867-247601368 | 6 | 0.904210619419362 |
| chr7-50475244-50475745 | 3 | 0.99593553545041 |
| chr1-215813532-215814033 | 3 | 0.995700926242037 |
| chr13-71896028-71896529 | 3 | 0.974654709048834 |
| chr4-27461297-27461798 | 5 | 0.983919978840557 |
| chr6-69324698-69325199 | 6 | 0.997210738292162 |
| chr7-87816877-87817378 | 8 | 0.822858763104372 |
| chr11-37440417-37440918 | 3 | 0.997221695313496 |
| chr8-114578933-114579434 | 3 | 0.990186311419089 |
| chr5-152926043-152926544 | 5 | 0.959183032104402 |
| chr11-17585745-17586246 | 8 | 0.930597154863887 |
| chr6-110638579-110639080 | 6 | 0.95551778560958 |
| chr4-160264571-160265072 | 5 | 0.961122909499369 |
| chr5-19839248-19839749 | 6 | 0.948756235881213 |
| chr3-157643015-157643516 | 3 | 0.997471334804661 |
| chr1-55795081-55795582 | 8 | 0.994938222734225 |
| chr7-109057341-109057842 | 1 | 0.893182947202107 |
| chr14-71385305-71385806 | 6 | 0.981039018520464 |
| chrX-31939151-31939652 | 6 | 0.871254642946353 |
| chr20-29444923-29445424 | 8 | 0.818829378815489 |
| chr10-11094007-11094508 | 8 | 0.995104729218893 |
| chr14-94088933-94089434 | 8 | 0.994877513047097 |
| chr3-22930911-22931412 | 3 | 0.992395588609225 |

| | | |
|---|---|---|
| chr3-125563992-125564493 | 6 | 0.980958294181985 |
| chr11-75277275-75277776 | 4 | 0.922889134012093 |
| chr6-15099118-15099619 | 5 | 0.862159048181969 |
| chr15-27788939-27789440 | 3 | 0.98988777395988 |
| chr5-129280328-129280829 | 3 | 0.996163219422379 |
| chr1-135548439-135548940 | 7 | 0.9463872158983 |
| chr17-55364392-55364893 | 3 | 0.943561006956984 |
| chr18-66827303-66827804 | 6 | 0.991235090077594 |
| chr7-126371971-126372472 | 6 | 0.990794793996324 |
| chr16-74943968-74944469 | 1 | 0.820787754486369 |
| chr7-28607606-28608107 | 2 | 0.965943606623417 |
| chr14-93191402-93191903 | 8 | 0.897614213100795 |
| chr2-243778138-243778639 | 3 | 0.991758224398377 |
| chr6-43680874-43681375 | 2 | 0.952606845700267 |
| chr7-108961792-108962293 | 8 | 0.836677305848088 |
| chr6-57857291-57857792 | 5 | 0.900653970899991 |
| chr13-12763854-12764355 | 1 | 0.940912593930236 |
| chrX-128146597-128147098 | 5 | 0.959030735427334 |
| chr10-89606711-89607212 | 8 | 0.821150678112503 |
| chr15-40321535-40322036 | 8 | 0.874024787710032 |
| chr7-70822882-70823383 | 8 | 0.944094921264504 |
| chr13-48506831-48507332 | 7 | 0.906782150495721 |
| chr15-100190011-100190512 | 8 | 0.993493681128311 |
| chr2-24757807-24758308 | 7 | 0.941073478665392 |
| chr1-103566735-103567236 | 3 | 0.985321132236568 |
| chr9-64887343-64887844 | 8 | 0.888884265378215 |
| chr1-133562105-133562606 | 8 | 0.992244890723887 |
| chr20-4630886-4631387 | 6 | 0.991938298874192 |
| chr8-25159540-25160041 | 8 | 0.986431318482432 |
| chr19-13528778-13529279 | 8 | 0.981181216852706 |
| chr5-131651649-131652150 | 8 | 0.955989337638139 |
| chr1-137647204-137647705 | 3 | 0.996171271242012 |
| chr3-52007256-52007757 | 2 | 0.948704932450663 |
| chr19-42394295-42394796 | 8 | 0.993493681128311 |
| chr6-41699451-41699952 | 8 | 0.871782986624419 |
| chr14-97381988-97382489 | 2 | 0.952658143440438 |
| chr10-23042254-23042755 | 6 | 0.943281390134081 |
| chr18-59123465-59123966 | 8 | 0.907765298496389 |
| chr20-29359241-29359742 | 3 | 0.992099834757904 |
| chr2-110999194-110999695 | 8 | 0.866339288253248 |
| chr20-2042722-2043223 | 6 | 0.987391266325744 |
| chr1-239221097-239221598 | 1 | 0.936139439368316 |
| chr6-114398238-114398739 | 8 | 0.976543978618743 |

| | | |
|---|---|---|
| chr2-218909849-218910350 | 2 | 0.913084472731852 |
| chr2-189597151-189597652 | 6 | 0.848783 |
| chr2-215774402-215774903 | 8 | 0.974040394667742 |
| chr2-26857777-26858278 | 5 | 0.988650594684744 |
| chr14-63921178-63921679 | 5 | 0.867691187797344 |
| chr4-13563658-13564159 | 8 | 0.881055277190551 |
| chr1-198206246-198206747 | 6 | 0.992594152753138 |
| chr19-16744665-16745166 | 7 | 0.829945507632969 |
| chr9-11883711-11884212 | 8 | 0.907651532621836 |
| chr3-141563132-141563633 | 8 | 0.902867856115667 |
| chr18-46990263-46990764 | 8 | 0.840000226478225 |
| chr6-94481054-94481555 | 6 | 0.978304151452863 |
| chr3-91167639-91168140 | 3 | 0.997134152501093 |
| chr7-130881556-130882057 | 2 | 0.948839879488261 |
| chr6-105016795-105017296 | 3 | 0.911875691925835 |
| chr5-149735904-149736405 | 3 | 0.861127396838078 |
| chr6-99096254-99096755 | 7 | 0.828910365731017 |
| chr1-218124994-218125495 | 3 | 0.967673610110039 |
| chr2-71466575-71467076 | 3 | 0.828391926154936 |
| chr4-36033333-36033834 | 6 | 0.991666989098466 |
| chr5-154698736-154699237 | 3 | 0.996714063895179 |
| chr7-60599581-60600082 | 6 | 0.993223605853396 |
| chrX-93318627-93319128 | 6 | 0.979097206124589 |
| chr6-123660624-123661125 | 8 | 0.941913273157271 |
| chr17-5307502-5308003 | 3 | 0.99488578756055 |
| chr3-39779731-39780232 | 6 | 0.969517755040733 |
| chr2-207505697-207506198 | 8 | 0.994009610929548 |
| chr18-8486062-8486563 | 5 | 0.983919978840557 |
| chr8-46954476-46954977 | 8 | 0.874565686696658 |
| chr6-23150038-23150539 | 8 | 0.99266329240679 |
| chr12-17199818-17200319 | 1 | 0.824620677178724 |
| chr6-35985777-35986278 | 3 | 0.99706779720281 |
| chr16-58271465-58271966 | 1 | 0.853099846004886 |
| chr18-75179278-75179779 | 6 | 0.988993820262966 |
| chr6-11559762-11560263 | 7 | 0.889609754645567 |
| chr14-9278543-9279044 | 5 | 0.990977231798049 |
| chr7-93899031-93899532 | 3 | 0.953911663089536 |
| chr14-8205373-8205874 | 2 | 0.870747479016883 |
| chr17-66047544-66048045 | 6 | 0.970948920224881 |
| chr17-66609349-66609850 | 4 | 0.972831491988396 |
| chr1-44211773-44212274 | 3 | 0.99593553545041 |
| chr1-215522350-215522851 | 8 | 0.99517979638415 |
| chr17-63721479-63721980 | 1 | 0.884229705449561 |

| | | |
|---|---|---|
| chr4-154278414-154278915 | 8 | 0.857489035659439 |
| chr17-26768839-26769340 | 2 | 0.859032094789811 |
| chr18-78474975-78475476 | 5 | 0.931786564992776 |
| chr7-106374637-106375138 | 8 | 0.924915823451678 |
| chr3-91007987-91008488 | 6 | 0.975637950641268 |
| chr15-37998041-37998542 | 6 | 0.993055347956789 |
| chr6-117676081-117676582 | 8 | 0.991760531253517 |
| chr7-95866966-95867467 | 3 | 0.863516576439375 |
| chr3-27214730-27215231 | 3 | 0.99593553545041 |
| chr16-81821702-81822203 | 6 | 0.821512343842328 |
| chr2-187612247-187612748 | 8 | 0.995207866691641 |
| chr13-31503857-31504358 | 4 | 0.949869942597848 |
| chrX-50219463-50219964 | 4 | 0.840498411771932 |
| chr5-126419415-126419916 | 3 | 0.954878803097101 |
| chr7-43439400-43439901 | 6 | 0.875258575199103 |
| chr2-241462332-241462833 | 3 | 0.856984756180829 |
| chr13-105835464-105835965 | 3 | 0.98745784164977 |
| chr1-124571755-124572256 | 6 | 0.989612947046013 |
| chr9-72726637-72727138 | 3 | 0.978725452838651 |
| chr15-26187969-26188470 | 7 | 0.951689440470133 |
| chr4-137485914-137486415 | 5 | 0.986747215791506 |
| chr15-55686573-55687074 | 3 | 0.912027089896085 |
| chr8-66504533-66505034 | 7 | 0.886850002577812 |
| chr9-91834364-91834865 | 8 | 0.923301154985336 |
| chr10-53096744-53097245 | 3 | 0.850133868860616 |
| chr9-53678532-53679033 | 5 | 0.902715457784388 |
| chr12-12555484-12555985 | 8 | 0.835554083169031 |
| chr8-20964897-20965398 | 5 | 0.969767034080667 |
| chr14-47128691-47129192 | 6 | 0.991206925481043 |
| chr20-18964955-18965456 | 3 | 0.852609324144596 |
| chr4-703537-704038 | 5 | 0.963469686139985 |
| chr2-41537969-41538470 | 8 | 0.847041521979438 |
| chr4-128992086-128992587 | 8 | 0.846893732319078 |
| chr5-112880949-112881450 | 3 | 0.996898544393807 |
| chr2-24915241-24915742 | 6 | 0.992318866098506 |
| chr1-118939677-118940178 | 5 | 0.981812297681871 |
| chr6-23230378-23230879 | 5 | 0.97776944400766 |
| chr6-38629492-38629993 | 3 | 0.918527738110024 |
| chr18-82733990-82734491 | 2 | 0.945833133923947 |
| chr19-15837015-15837516 | 5 | 0.983919978840557 |
| chr7-53604190-53604691 | 4 | 0.873723788548398 |
| chr14-71297115-71297616 | 1 | 0.915919862673111 |
| chr17-56556110-56556611 | 3 | 0.965489883888934 |

| | | |
|---|---|---|
| chr7-85855692-85856193 | 4 | 0.902387149032641 |
| chr10-70652104-70652605 | 8 | 0.979153494211519 |
| chr1-156457280-156457781 | 3 | 0.990647110698882 |
| chr1-90674764-90675265 | 8 | 0.994640684929723 |
| chr7-45249267-45249768 | 5 | 0.962467605167832 |
| chr8-113031860-113032361 | 3 | 0.846681620009368 |
| chr20-2454458-2454959 | 2 | 0.954515769194911 |
| chr2-171273869-171274370 | 3 | 0.993891690936645 |
| chr1-12372971-12373472 | 1 | 0.890721283865511 |
| chr14-99051363-99051864 | 8 | 0.956532098104645 |
| chr13-91472495-91472996 | 1 | 0.907290276763266 |
| chr17-47216116-47216617 | 8 | 0.969712279739268 |
| chr3-80880969-80881470 | 6 | 0.991206925481043 |
| chr17-18674221-18674722 | 8 | 0.947734382153129 |
| chr19-52597372-52597873 | 8 | 0.822394603717081 |
| chr7-100137733-100138234 | 8 | 0.941322336153973 |
| chr6-129176846-129177347 | 6 | 0.991206925481043 |
| chr6-98067020-98067521 | 6 | 0.958956027261324 |
| chr10-92986060-92986561 | 2 | 0.964372973341801 |
| chrX-4600243-4600744 | 3 | 0.994989255654951 |
| chr5-129611747-129612248 | 8 | 0.923264361674616 |
| chr18-19098879-19099380 | 6 | 0.881735474592969 |
| chr20-18190971-18191472 | 3 | 0.973243864407587 |
| chrX-85676770-85677271 | 7 | 0.870919176485629 |
| chr7-75578665-75579166 | 5 | 0.971906173843746 |
| chr16-56625411-56625912 | 8 | 0.890158348276365 |
| chr5-83991136-83991637 | 3 | 0.995953401451714 |
| chr17-19438443-19438944 | 3 | 0.994749627767208 |
| chr10-76346298-76346799 | 3 | 0.830522656531805 |
| chr6-45015142-45015643 | 3 | 0.996607954949174 |
| chr7-30281035-30281536 | 8 | 0.830478496327721 |
| chr3-26458819-26459320 | 3 | 0.99593553545041 |
| chr17-70176916-70177417 | 5 | 0.972197694342599 |
| chr10-72402934-72403435 | 8 | 0.885800982431111 |
| chr14-2726841-2727342 | 6 | 0.979163695798889 |
| chr15-29033050-29033551 | 6 | 0.959995441664809 |
| chr4-41599449-41599950 | 3 | 0.996867125909769 |
| chr18-53854070-53854571 | 3 | 0.99593553545041 |
| chr11-15271281-15271782 | 3 | 0.943889873133582 |
| chr9-24530491-24530992 | 3 | 0.976387171837959 |
| chr7-25613223-25613724 | 8 | 0.86067729485922 |
| chr1-246687951-246688452 | 2 | 0.876732299467358 |
| chr4-63427657-63428158 | 8 | 0.982861753644156 |

| | | |
|---|---|---|
| chr11-17099239-17099740 | 6 | 0.823537401130328 |
| chr19-14276924-14277425 | 5 | 0.98616326445835 |
| chr11-37931452-37931953 | 6 | 0.991206925481043 |
| chr5-76853081-76853582 | 6 | 0.994092778887306 |
| chr16-45945767-45946268 | 5 | 0.985508565213842 |
| chr12-22957861-22958362 | 4 | 0.888807902269082 |
| chr13-66592114-66592615 | 5 | 0.979158826688158 |
| chr2-73860266-73860767 | 3 | 0.984412798579276 |
| chr7-48415146-48415647 | 3 | 0.99586580520001 |
| chr1-246517171-246517672 | 2 | 0.90251570217618 |
| chr4-160919260-160919761 | 5 | 0.871378666477785 |
| chr2-191502272-191502773 | 3 | 0.969192885276382 |
| chr14-52720463-52720964 | 3 | 0.996347004192518 |
| chr15-30483119-30483620 | 3 | 0.8959272930232 |
| chr8-101222586-101223087 | 8 | 0.848394755396508 |
| chr16-64832958-64833459 | 2 | 0.968095034117614 |
| chr5-38891584-38892085 | 4 | 0.822001854878374 |
| chr13-23961882-23962383 | 6 | 0.99195706800853 |
| chr5-149453318-149453819 | 3 | 0.975000008113167 |
| chr4-7692058-7692559 | 6 | 0.993197289069718 |
| chr7-89990106-89990607 | 3 | 0.993097378931625 |
| chr3-37066179-37066680 | 4 | 0.948972508819407 |
| chr3-17453607-17454108 | 8 | 0.914220748872916 |
| chr5-46035709-46036210 | 6 | 0.991206925481043 |
| chrX-9686767-9687268 | 5 | 0.989456278083481 |
| chr10-11084013-11084514 | 8 | 0.95412969197098 |
| chr11-69684290-69684791 | 3 | 0.903542535803981 |
| chr10-98043324-98043825 | 8 | 0.912933754301891 |
| chr8-61795258-61795759 | 8 | 0.935752330123111 |
| chr17-26590257-26590758 | 4 | 0.818083126802307 |
| chr6-56788606-56789107 | 2 | 0.932219739318835 |
| chr8-109293838-109294339 | 8 | 0.995018385370297 |
| chr1-199407257-199407758 | 7 | 0.903460467063869 |
| chr6-83533011-83533512 | 5 | 0.968560006181054 |
| chr16-63339475-63339976 | 3 | 0.930159475453227 |
| chr3-80129605-80130106 | 3 | 0.986910187985819 |
| chr7-129355299-129355800 | 8 | 0.983044854403442 |
| chr3-91922010-91922511 | 6 | 0.966035454624498 |
| chr7-67538614-67539115 | 6 | 0.974094907793708 |
| chr5-3197496-3197997 | 4 | 0.897897485545118 |
| chr11-50057415-50057916 | 3 | 0.995697261434417 |
| chr1-48290901-48291402 | 8 | 0.991158712918663 |
| chr2-175209253-175209754 | 8 | 0.858567138533817 |

| | | |
|---|---|---|
| chr8-48228480-48228981 | 2 | 0.893876194626894 |
| chr15-63111648-63112149 | 3 | 0.996807178983891 |
| chr3-106247328-106247829 | 5 | 0.988479759864519 |
| chr4-67478806-67479307 | 8 | 0.984032154144278 |
| chrX-132728585-132729086 | 6 | 0.909141096829443 |
| chr17-19443977-19444478 | 8 | 0.82597136702756 |
| chr3-140110231-140110732 | 6 | 0.991206925481043 |
| chr11-20531867-20532368 | 6 | 0.987677384081882 |
| chr2-185211731-185212232 | 5 | 0.900428249003527 |
| chr13-73332269-73332770 | 8 | 0.955791456711708 |
| chr10-61968954-61969455 | 7 | 0.912374647761916 |
| chr2-215787239-215787740 | 4 | 0.82933415382722 |
| chr16-3321150-3321651 | 8 | 0.993493681128311 |
| chr13-89789363-89789864 | 5 | 0.959747765029148 |
| chr9-99447261-99447762 | 4 | 0.969651571478333 |
| chr18-19222027-19222528 | 3 | 0.995084350453341 |
| chr13-30594811-30595312 | 1 | 0.927095618520564 |
| chr2-167859421-167859922 | 6 | 0.86093857173983 |
| chr9-69271401-69271902 | 5 | 0.981650902242602 |
| chr3-96766738-96767239 | 8 | 0.876543313731305 |
| chr7-47043076-47043577 | 5 | 0.969822459702897 |
| chr4-131145974-131146475 | 2 | 0.942685991415314 |
| chr2-101504391-101504892 | 3 | 0.994252438966628 |
| chr10-28701282-28701783 | 8 | 0.992544271135702 |
| chr2-188904643-188905144 | 8 | 0.950946618804451 |
| chr18-69519498-69519999 | 4 | 0.85040556975996 |
| chr16-55089571-55090072 | 3 | 0.95444175836305 |
| chr1-260201146-260201647 | 3 | 0.996352063755145 |
| chr3-106755022-106755523 | 6 | 0.989191275153205 |
| chr7-33001899-33002400 | 5 | 0.977759376360735 |
| chr3-10700338-10700839 | 8 | 0.841025167890407 |
| chr7-89755285-89755786 | 6 | 0.979105688180577 |
| chr2-38399993-38400494 | 5 | 0.843770307391363 |
| chr3-130982575-130983076 | 3 | 0.964316544161242 |
| chr7-116724187-116724688 | 8 | 0.89133824897839 |
| chr4-63890996-63891497 | 8 | 0.988459812867379 |
| chr16-71912296-71912797 | 3 | 0.967480340177322 |
| chr12-40359687-40360188 | 8 | 0.902036536163913 |
| chr1-155081560-155082061 | 6 | 0.992196925248502 |
| chr4-68291492-68291993 | 5 | 0.990137820969938 |
| chr13-97885132-97885633 | 8 | 0.858372987380127 |
| chr18-7345072-7345573 | 5 | 0.983919978840557 |
| chr3-32164492-32164993 | 4 | 0.95556723347547 |

| | | |
|---|---|---|
| chr8-55132997-55133498 | 8 | 0.978799531693456 |
| chr2-92707318-92707819 | 8 | 0.984399302482084 |
| chr7-86163401-86163902 | 3 | 0.991822394995626 |
| chr4-57370941-57371442 | 6 | 0.991227977006005 |
| chr12-39911530-39912031 | 3 | 0.993341786866055 |
| chr1-56402043-56402544 | 3 | 0.996063983087663 |
| chr4-7589709-7590210 | 8 | 0.834367648150398 |
| chr8-64882780-64883281 | 8 | 0.994933764222414 |
| chr3-169019472-169019973 | 5 | 0.963291409037624 |
| chr18-12648824-12649325 | 2 | 0.969377281253218 |
| chr1-96309911-96310412 | 5 | 0.991675713767143 |
| chr16-7266126-7266627 | 1 | 0.879839176258094 |
| chr3-114948922-114949423 | 3 | 0.971542671634918 |
| chr15-6140545-6141046 | 3 | 0.997202938846344 |
| chr4-162385527-162386028 | 8 | 0.922903783167055 |
| chr4-129804763-129805264 | 4 | 0.818291245385605 |
| chr2-117227379-117227880 | 3 | 0.994377863194357 |
| chr14-15471142-15471643 | 5 | 0.970017550127255 |
| chr2-184687520-184688021 | 8 | 0.960100648501179 |
| chr6-15883215-15883716 | 5 | 0.989554486278761 |
| chr9-18164835-18165336 | 6 | 0.883779376371179 |
| chr9-106955619-106956120 | 8 | 0.856973755788892 |
| chr7-72405594-72406095 | 8 | 0.988933606301426 |
| chr6-132504735-132505236 | 3 | 0.979608911546773 |
| chr4-118354624-118355125 | 3 | 0.996177834259748 |
| chr4-145279257-145279758 | 3 | 0.989144060523918 |
| chr2-92788320-92788821 | 8 | 0.901468151827626 |
| chr11-81581838-81582339 | 3 | 0.960327840711401 |
| chr2-113373965-113374466 | 3 | 0.829634326928652 |
| chr4-122844023-122844524 | 6 | 0.977905203871875 |
| chr2-187515679-187516180 | 6 | 0.90847625355339 |
| chr1-7848778-7849279 | 3 | 0.981197378104162 |
| chr1-34424840-34425341 | 8 | 0.909072874163446 |
| chr4-121011088-121011589 | 2 | 0.872489170012808 |
| chr1-25938803-25939304 | 3 | 0.978353039388653 |
| chr14-86982981-86983482 | 6 | 0.991206925481043 |
| chr4-89859175-89859676 | 3 | 0.884315066241903 |
| chr11-52015636-52016137 | 3 | 0.994979021646864 |
| chr7-111140780-111141281 | 4 | 0.968753043445123 |
| chr18-13713149-13713650 | 6 | 0.992791369510864 |
| chr3-108492496-108492997 | 3 | 0.909401814147503 |
| chr6-37604716-37605217 | 3 | 0.979371530295535 |
| chr17-12263901-12264402 | 5 | 0.903971411772676 |

| | | |
|---|---|---|
| chr3-25550461-25550962 | 6 | 0.99248621092564 |
| chr10-77392442-77392943 | 3 | 0.996853090082961 |
| chr2-230465794-230466295 | 4 | 0.969886210409365 |
| chr3-122830501-122831002 | 6 | 0.971726625176285 |
| chrX-140317809-140318310 | 3 | 0.99593553545041 |
| chr4-167584954-167585455 | 3 | 0.981549949711215 |
| chr6-22709003-22709504 | 6 | 0.912774118750787 |
| chr1-171979166-171979667 | 3 | 0.888083018570124 |
| chr5-79422890-79423391 | 8 | 0.940145012934622 |
| chr5-22529823-22530324 | 5 | 0.990295777235509 |
| chr4-6975514-6976015 | 4 | 0.949674106765911 |
| chr4-104017692-104018193 | 5 | 0.975341308115937 |
| chr3-50081618-50082119 | 8 | 0.822606353599219 |
| chr3-9474434-9474935 | 3 | 0.966563448982301 |
| chr1-153999584-154000085 | 5 | 0.907815900876345 |
| chr14-77054285-77054786 | 5 | 0.961757841254073 |
| chr10-51984480-51984981 | 3 | 0.986753404855152 |
| chr2-223627978-223628479 | 8 | 0.937161658145184 |
| chr3-11497810-11498311 | 8 | 0.893518267291445 |
| chr17-70479256-70479757 | 6 | 0.852054484195682 |
| chr2-204073985-204074486 | 8 | 0.940557909400849 |
| chr1-133750349-133750850 | 3 | 0.931408125947366 |
| chr10-39940295-39940796 | 6 | 0.95291611186994 |
| chr1-47403672-47404173 | 6 | 0.978122096552085 |
| chr9-24739201-24739702 | 6 | 0.991206925481043 |
| chr17-32588891-32589392 | 8 | 0.979520308427669 |
| chr12-12429535-12430036 | 3 | 0.99593553545041 |
| chr2-70423558-70424059 | 3 | 0.997568917245164 |
| chr3-139779874-139780375 | 6 | 0.932381186786407 |
| chr2-42504508-42505009 | 8 | 0.840980072685771 |
| chr17-52460074-52460575 | 5 | 0.948018382057919 |
| chr4-24630931-24631432 | 3 | 0.992733779998909 |
| chr9-84250949-84251450 | 7 | 0.964081346981634 |
| chr6-72786602-72787103 | 3 | 0.997548682814887 |
| chr7-14373551-14374052 | 8 | 0.901762381751914 |
| chr10-91291417-91291918 | 5 | 0.954405524309523 |
| chr1-137624994-137625495 | 8 | 0.988849165745074 |
| chr3-48158254-48158755 | 3 | 0.988849984333466 |
| chr2-30576381-30576882 | 4 | 0.82370737447876 |
| chr6-51177025-51177526 | 8 | 0.993493681128311 |
| chr18-35639948-35640449 | 6 | 0.993018917218407 |
| chr1-78344847-78345348 | 6 | 0.882857058758764 |
| chr4-77386057-77386558 | 6 | 0.874184677227263 |

| | | |
|---|---|---|
| chr9-65416331-65416832 | 5 | 0.990650724624147 |
| chr3-41850770-41851271 | 3 | 0.945045753391501 |
| chr9-18221677-18222178 | 4 | 0.822599644830125 |
| chr17-3800724-3801225 | 8 | 0.929592321646895 |
| chrX-24637730-24638231 | 5 | 0.971573088017372 |
| chr19-55744319-55744820 | 6 | 0.994789855677432 |
| chr5-118192949-118193450 | 7 | 0.88709559692271 |
| chr10-7765845-7766346 | 8 | 0.978835252154333 |
| chr11-52524315-52524816 | 4 | 0.848125815732174 |
| chr4-85699519-85700020 | 3 | 0.996807806021989 |
| chr7-60480188-60480689 | 6 | 0.995785596987476 |
| chr14-60566954-60567455 | 6 | 0.992426068299594 |
| chr9-8118568-8119069 | 3 | 0.820515131751081 |
| chr2-50288760-50289261 | 6 | 0.989950526576485 |
| chr13-41418978-41419479 | 6 | 0.992112572565498 |
| chr5-93653191-93653692 | 6 | 0.96711374886286 |
| chr2-33495623-33496124 | 8 | 0.981152658152916 |
| chr10-4204613-4205114 | 3 | 0.983984924485243 |
| chr5-47784877-47785378 | 3 | 0.997158680082806 |
| chr13-95022371-95022872 | 6 | 0.992475788612243 |
| chr11-33271347-33271848 | 6 | 0.824903238376062 |
| chr5-162922550-162923051 | 2 | 0.828281999497311 |
| chr7-33613167-33613668 | 3 | 0.991201826921022 |
| chr1-189560105-189560606 | 3 | 0.970143499248915 |
| chr4-146899705-146900206 | 5 | 0.952202621815295 |
| chr1-85122025-85122526 | 3 | 0.942665208070544 |
| chr5-77940969-77941470 | 8 | 0.97232815690506 |
| chr2-154208293-154208794 | 3 | 0.973891859936266 |
| chr4-59005170-59005671 | 8 | 0.900525141479802 |
| chr3-148008819-148009320 | 1 | 0.874389203824565 |
| chr9-1226459-1226960 | 8 | 0.88495598470887 |
| chr5-71390687-71391188 | 5 | 0.901106102392817 |
| chr9-49392746-49393247 | 8 | 0.97780506739401 |
| chr1-106238446-106238947 | 5 | 0.892421865152905 |
| chr10-87834655-87835156 | 4 | 0.862692983080708 |
| chr6-20657142-20657643 | 6 | 0.893226322725047 |
| chr3-144599123-144599624 | 3 | 0.995914072899412 |
| chr10-72623142-72623643 | 3 | 0.99593553545041 |
| chr8-12410535-12411036 | 4 | 0.897477303495532 |
| chr5-149992818-149993319 | 8 | 0.970536306324264 |
| chr7-65459312-65459813 | 3 | 0.996673424158244 |
| chr4-130910452-130910953 | 6 | 0.991206925481043 |
| chr17-34115634-34116135 | 3 | 0.994367643417981 |

| | | |
|---|---|---|
| chr13-67138372-67138873 | 2 | 0.943189510691377 |
| chr9-40681991-40682492 | 7 | 0.911000754788266 |
| chr6-30949232-30949733 | 3 | 0.995976173872864 |
| chr14-98498527-98499028 | 8 | 0.815658857833243 |
| chr1-252946769-252947270 | 5 | 0.852444596734421 |
| chr18-70804411-70804912 | 3 | 0.996439165522765 |
| chr5-144578146-144578647 | 8 | 0.887607532214243 |
| chr6-31031942-31032443 | 3 | 0.958803998438713 |
| chr11-82990776-82991277 | 3 | 0.99593553545041 |
| chr20-9310265-9310766 | 2 | 0.930283072334326 |
| chr3-22708203-22708704 | 1 | 0.884650551851933 |
| chr5-43364171-43364672 | 6 | 0.828904830640137 |
| chr8-66799683-66800184 | 3 | 0.981992640047904 |
| chr8-23812710-23813211 | 5 | 0.986235350677889 |
| chr15-34768528-34769029 | 2 | 0.865942613109321 |
| chr3-32745125-32745626 | 2 | 0.837869462441596 |
| chr3-158927317-158927818 | 8 | 0.944608367879244 |
| chr3-91418684-91419185 | 3 | 0.994548837664631 |
| chr4-63245905-63246406 | 6 | 0.993220617764008 |
| chr6-91419806-91420307 | 4 | 0.958744462346082 |
| chr20-52002141-52002642 | 5 | 0.934068796550375 |
| chr4-36945897-36946398 | 6 | 0.991206925481043 |
| chrX-52147132-52147633 | 6 | 0.991206925481043 |
| chr4-66510966-66511467 | 6 | 0.991016453006814 |
| chr5-35533667-35534168 | 3 | 0.99691011166391 |
| chr2-143710676-143711177 | 8 | 0.994150535417652 |
| chr7-111341183-111341684 | 6 | 0.993392374727116 |
| chr15-3031596-3032097 | 8 | 0.98699056979484 |
| chr6-112933861-112934362 | 6 | 0.970256900754281 |
| chr2-26326389-26326890 | 6 | 0.997002562223526 |
| chr8-44197998-44198499 | 8 | 0.982717260814512 |
| chr2-66071283-66071784 | 8 | 0.993493681128311 |
| chr19-48394954-48395455 | 8 | 0.906268927464167 |
| chr10-83242761-83243262 | 8 | 0.954046345568022 |
| chr4-80588221-80588722 | 5 | 0.961055177451729 |
| chr12-20775938-20776439 | 8 | 0.896384685113659 |
| chr10-71804381-71804882 | 3 | 0.880167160049085 |
| chr3-78552405-78552906 | 5 | 0.882668644277254 |
| chr7-78812066-78812567 | 3 | 0.995620631856231 |
| chr4-10516234-10516735 | 6 | 0.990080030280503 |
| chr18-14081112-14081613 | 3 | 0.834032793143927 |
| chr19-35610324-35610825 | 3 | 0.997140977263647 |
| chr19-49357084-49357585 | 6 | 0.996950714540477 |

| | | |
|---|---|---|
| chr5-60031809-60032310 | 7 | 0.932500909237937 |
| chr5-161528565-161529066 | 4 | 0.897026788760035 |
| chr8-121602556-121603057 | 6 | 0.993919697929272 |
| chrX-115122809-115123310 | 8 | 0.836243274991454 |
| chr13-77800850-77801351 | 5 | 0.983919978840557 |
| chr16-67755730-67756231 | 6 | 0.893994739948254 |
| chr6-99221580-99222081 | 3 | 0.995715332571648 |
| chr10-57207801-57208302 | 3 | 0.979988238836714 |
| chr1-139376081-139376582 | 6 | 0.841264267204292 |
| chr2-52094611-52095112 | 6 | 0.993215674538441 |
| chr2-11263262-11263763 | 8 | 0.909275392270909 |
| chrX-714342-714843 | 3 | 0.968662256405731 |
| chr13-72375540-72376041 | 8 | 0.993493681128311 |
| chr3-133686356-133686857 | 3 | 0.938930824372852 |
| chr17-18142240-18142741 | 8 | 0.977761467519826 |
| chr3-48419251-48419752 | 6 | 0.990375209956296 |
| chr16-55596844-55597345 | 6 | 0.99144856461115 |
| chr20-19027765-19028266 | 6 | 0.990066151654673 |
| chr1-129399610-129400111 | 8 | 0.817541252163197 |
| chr3-57455337-57455838 | 8 | 0.965702952845787 |
| chr4-116995632-116996133 | 4 | 0.914480916046597 |
| chr4-169245136-169245637 | 3 | 0.960494850514228 |
| chr16-20240606-20241107 | 3 | 0.823518196905401 |
| chr15-40361355-40361856 | 8 | 0.921346973115451 |
| chr4-146419138-146419639 | 3 | 0.970890823378196 |
| chr2-92932769-92933270 | 6 | 0.991594087198995 |
| chr3-13841195-13841696 | 8 | 0.817433805528305 |
| chr17-48153959-48154460 | 6 | 0.972171311217179 |
| chr6-101327454-101327955 | 1 | 0.95480819293037 |
| chr11-45651524-45652025 | 3 | 0.99593553545041 |
| chr3-30055655-30056156 | 3 | 0.941986968800459 |
| chr15-91987886-91988387 | 6 | 0.976312257194744 |
| chr16-76042967-76043468 | 5 | 0.977846782769843 |
| chr4-701404-701905 | 3 | 0.994774792984102 |
| chr14-97160669-97161170 | 3 | 0.984398154524483 |
| chr2-35814675-35815176 | 3 | 0.979974473253524 |
| chr10-18423235-18423736 | 8 | 0.995213035482481 |
| chr3-102312029-102312530 | 3 | 0.996511597281583 |
| chr16-17048689-17049190 | 3 | 0.996653205015777 |
| chr19-20813892-20814393 | 1 | 0.884028270428618 |
| chr3-64774618-64775119 | 3 | 0.997222745165288 |
| chr13-63742350-63742851 | 3 | 0.980654506942375 |
| chr17-10013657-10014158 | 8 | 0.895884733855983 |

| | | |
|---|---|---|
| chr3-100633815-100634316 | 6 | 0.990554054207488 |
| chrX-9537229-9537730 | 3 | 0.938494157926733 |
| chr2-216538207-216538708 | 3 | 0.918716718577299 |
| chr2-208756075-208756576 | 8 | 0.988352140236899 |
| chr18-5020965-5021466 | 6 | 0.892680354962685 |
| chr19-14348379-14348880 | 8 | 0.995207785817953 |
| chr2-64634914-64635415 | 3 | 0.987864378207076 |
| chr3-115913228-115913729 | 3 | 0.89350401958807 |
| chr2-29693465-29693966 | 3 | 0.927625426919885 |
| chr16-21397494-21397995 | 3 | 0.996757967797594 |
| chr13-28180948-28181449 | 6 | 0.991206925481043 |
| chr8-116038652-116039153 | 2 | 0.936508271652221 |
| chr3-144412615-144413116 | 3 | 0.964602751755521 |
| chr15-60939108-60939609 | 3 | 0.99593553545041 |
| chr7-109924973-109925474 | 1 | 0.888812965803224 |
| chr1-176801325-176801826 | 8 | 0.943170703039985 |
| chr7-27633493-27633994 | 6 | 0.89156449954668 |
| chr15-4736742-4737243 | 3 | 0.966188919743602 |
| chr3-152413930-152414431 | 1 | 0.846233813188162 |
| chr9-68627069-68627570 | 8 | 0.993493681128311 |
| chr11-73370161-73370662 | 8 | |
| chr1-189689820-189690321 | 1 | |
| chr4-6750900-6751401 | 8 | |
| chr1-151442902-151443403 | 8 | |
| chr9-65297370-65297871 | 8 | |
| chr5-163817177-163817678 | 5 | |
| chr10-83210063-83210564 | 5 | |
| chr9-45220592-45221093 | 8 | |
| chr3-163415485-163415986 | 8 | |
| chr2-26710392-26710893 | 1 | |
| chr11-67865233-67865734 | 8 | |
| chr18-26585898-26586399 | 8 | |
| chr5-79825917-79826418 | 6 | |
| chr1-202350231-202350732 | 8 | |
| chr2-112676981-112677482 | 8 | |
| chr2-215179562-215180063 | 5 | |
| chr9-9196070-9196571 | 8 | |
| chr9-104485823-104486324 | 5 | |
| chr14-57775819-57776320 | 5 | |
| chr4-31298832-31299333 | 8 | |
| chr15-50759370-50759871 | 1 | |
| chr10-18512155-18512656 | 1 | |
| chr5-137214842-137215343 | 6 | |

| | |
|---|---|
| chr6-98273237-98273738 | 8 |
| chr7-103413687-103414188 | 8 |
| chr7-95077830-95078331 | 8 |
| chr7-99252137-99252638 | 8 |
| chr11-17671584-17672085 | 8 |
| chr15-49637132-49637633 | 8 |
| chr3-159607287-159607788 | 8 |
| chr17-19243821-19244322 | 8 |
| chr10-19258728-19259229 | 6 |
| chr2-33529201-33529702 | 6 |
| chr12-27773164-27773665 | 6 |
| chrX-112965916-112966417 | 6 |
| chr17-57181700-57182201 | 1 |
| chr15-906736-907237 | 1 |
| chr2-142349932-142350433 | 8 |
| chr4-103927891-103928392 | 5 |
| chr2-146801851-146802352 | 6 |
| chr8-29874953-29875454 | 1 |
| chr2-216370250-216370751 | 8 |
| chr20-8421900-8422401 | 5 |
| chr5-102396960-102397461 | 6 |
| chr2-153759016-153759517 | 6 |
| chr3-62345425-62345926 | 5 |
| chr7-24121900-24122401 | 8 |
| chr1-207653371-207653872 | 1 |
| chr3-113771132-113771633 | 6 |
| chr12-22408072-22408573 | 8 |
| chr4-72532595-72533096 | 8 |
| chr15-53792625-53793126 | 8 |
| chrX-67571976-67572477 | 8 |
| chr9-76442078-76442579 | 5 |
| chr18-38559174-38559675 | 1 |
| chr1-192294463-192294964 | 8 |
| chr3-38046804-38047305 | 6 |
| chr2-78202599-78203100 | 8 |
| chr5-38191715-38192216 | 1 |
| chr7-31621619-31622120 | 1 |
| chr8-69483588-69484089 | 5 |
| chr2-192791967-192792468 | 8 |
| chr5-27037971-27038472 | 8 |
| chr3-7979330-7979831 | 5 |
| chr4-147709244-147709745 | 5 |
| chr11-77710656-77711157 | 1 |

| | |
|---|---|
| chr17-69888983-69889484 | 6 |
| chr7-53635434-53635935 | 1 |
| chr17-7664274-7664775 | 8 |
| chr11-51073531-51074032 | 8 |
| chr7-127279699-127280200 | 8 |
| chr17-19273182-19273683 | 5 |
| chr6-17290549-17291050 | 8 |
| chr14-7470933-7471434 | 5 |
| chrX-148823719-148824220 | 6 |
| chr5-42247112-42247613 | 6 |
| chr4-124555861-124556362 | 1 |
| chr5-56163873-56164374 | 5 |
| chr1-178575482-178575983 | 6 |
| chr16-79145061-79145562 | 8 |
| chr13-91134026-91134527 | 6 |
| chr20-12942254-12942755 | 8 |
| chr16-66926-67427 | 5 |
| chrX-32323754-32324255 | 8 |
| chr2-141598545-141599046 | 5 |
| chr5-54340405-54340906 | 6 |
| chr9-41466899-41467400 | 1 |
| chrX-33796769-33797270 | 5 |
| chr19-38260294-38260795 | 6 |
| chr5-128575338-128575839 | 8 |
| chr11-25781764-25782265 | 8 |
| chr2-78095875-78096376 | 6 |
| chr2-157872931-157873432 | 5 |
| chr3-127701238-127701739 | 5 |
| chr11-12101125-12101626 | 1 |
| chr6-3974893-3975394 | 8 |
| chr18-3403624-3404125 | 8 |
| chr18-44819442-44819943 | 8 |
| chr11-2916790-2917291 | 6 |
| chr14-94449694-94450195 | 8 |
| chr3-33475377-33475878 | 5 |
| chr10-19005674-19006175 | 6 |
| chr6-7314496-7314997 | 6 |
| chr2-153197504-153198005 | 8 |
| chr3-44093028-44093529 | 8 |
| chr2-68869852-68870353 | 6 |
| chr9-57337388-57337889 | 6 |
| chr17-69191559-69192060 | 8 |
| chr2-26519955-26520456 | 5 |

| | |
|---|---|
| chr14-35344872-35345373 | 5 |
| chr10-19060044-19060545 | 8 |
| chr7-94718995-94719496 | 1 |
| chr3-78689377-78689878 | 8 |
| chr8-93270956-93271457 | 8 |
| chr1-164317889-164318390 | 8 |
| chr2-222935962-222936463 | 6 |
| chr8-122464718-122465219 | 6 |
| chr1-106790196-106790697 | 8 |
| chr9-7339190-7339691 | 5 |
| chr6-42648808-42649309 | 6 |
| chr2-115851742-115852243 | 6 |
| chr8-73553610-73554111 | 5 |
| chr14-5373922-5374423 | 8 |
| chr6-100013390-100013891 | 8 |
| chr4-148026795-148027296 | 8 |
| chr9-54905054-54905555 | 8 |
| chr9-84681426-84681927 | 8 |
| chr4-107860721-107861222 | 5 |
| chrX-11850991-11851492 | 1 |
| chr1-46397555-46398056 | 6 |
| chr2-235893536-235894037 | 8 |
| chr2-77380089-77380590 | 1 |
| chr1-47353175-47353676 | 6 |
| chr13-45693400-45693901 | 5 |
| chr18-49215352-49215853 | 1 |
| chr9-33308317-33308818 | 8 |
| chr7-87184983-87185484 | 5 |
| chr10-28168478-28168979 | 5 |
| chr17-18148964-18149465 | 8 |
| chr2-11319241-11319742 | 8 |
| chr7-31096596-31097097 | 6 |
| chr9-56524377-56524878 | 8 |
| chr3-105407898-105408399 | 6 |
| chr1-225733728-225734229 | 8 |
| chr19-47704730-47705231 | 5 |
| chr17-9740699-9741200 | 6 |
| chr7-100866520-100867021 | 8 |
| chr1-233946425-233946926 | 8 |
| chr2-145505725-145506226 | 1 |
| chr3-128850130-128850631 | 6 |
| chr1-142014001-142014502 | 6 |
| chr12-32713727-32714228 | 8 |

| | |
|---|---|
| chr4-126047627-126048128 | 6 |
| chr9-63874816-63875317 | 5 |
| chr2-104280421-104280922 | 5 |
| chr5-136759503-136760004 | 8 |
| chr13-101203950-101204451 | 8 |
| chr7-44922125-44922626 | 1 |
| chr11-76053129-76053630 | 8 |
| chr1-40341382-40341883 | 5 |
| chr15-78378960-78379461 | 8 |
| chr3-68852048-68852549 | 8 |
| chr12-24842204-24842705 | 5 |
| chr3-33245407-33245908 | 8 |
| chr18-32178408-32178909 | 1 |
| chr13-65776884-65777385 | 6 |
| chrX-96836891-96837392 | 8 |
| chr4-18018283-18018784 | 6 |
| chr10-76615163-76615664 | 5 |
| chr12-32304080-32304581 | 8 |
| chr8-104707203-104707704 | 6 |
| chr2-94885079-94885580 | 5 |
| chr8-86820817-86821318 | 8 |
| chr15-76957011-76957512 | 8 |
| chr15-38546195-38546696 | 8 |
| chr17-15899629-15900130 | 5 |
| chr4-78044804-78045305 | 6 |
| chr13-66456436-66456937 | 6 |
| chr8-102968626-102969127 | 8 |
| chr7-112349446-112349947 | 8 |
| chr7-110801197-110801698 | 5 |
| chr6-62759757-62760258 | 6 |
| chr5-164481996-164482497 | 8 |
| chr1-3006908-3007409 | 6 |
| chr7-27373887-27374388 | 5 |
| chr2-143134138-143134639 | 8 |
| chr6-68667597-68668098 | 5 |
| chr18-49001678-49002179 | 5 |
| chr18-62920453-62920954 | 8 |
| chr17-18257517-18258018 | 8 |
| chr2-106245634-106246135 | 6 |
| chr5-32022499-32023000 | 6 |
| chr1-92049690-92050191 | 8 |
| chr4-176339323-176339824 | 8 |
| chr3-152183995-152184496 | 8 |

| | |
|---|---|
| chr3-93470458-93470959 | 5 |
| chr6-102876585-102877086 | 8 |
| chr6-137531874-137532375 | 6 |
| chr15-48738460-48738961 | 8 |
| chr9-88027094-88027595 | 5 |
| chr9-20567559-20568060 | 8 |
| chr17-65915377-65915878 | 8 |
| chr3-144151575-144152076 | 8 |
| chrX-115142072-115142573 | 6 |
| chr4-172143888-172144389 | 8 |
| chr8-92652521-92653022 | 8 |
| chr17-51670090-51670591 | 5 |
| chr11-83535664-83536165 | 8 |
| chr10-104886541-104887042 | 8 |
| chr4-147570630-147571131 | 1 |
| chr2-178805814-178806315 | 8 |
| chr18-23686798-23687299 | 1 |
| chrX-10202934-10203435 | 8 |
| chr7-117224045-117224546 | 8 |
| chr1-1159321-1159822 | 6 |
| chr4-162182535-162183036 | 1 |
| chr8-111382574-111383075 | 8 |
| chr6-90626831-90627332 | 1 |
| chr13-92089432-92089933 | 8 |
| chr3-48081043-48081544 | 8 |
| chr1-14999819-15000320 | 6 |
| chr17-68522579-68523080 | 6 |
| chr10-47392275-47392776 | 6 |
| chr2-229169462-229169963 | 5 |
| chr18-33802225-33802726 | 8 |
| chr16-28701492-28701993 | 1 |
| chr7-109569105-109569606 | 8 |
| chrX-49496961-49497462 | 8 |
| chr16-70525197-70525698 | 5 |
| chr4-174280165-174280666 | 8 |
| chr17-40612188-40612689 | 1 |
| chr10-74829707-74830208 | 8 |
| chr19-45586506-45587007 | 8 |
| chr2-169518424-169518925 | 8 |
| chr3-90897024-90897525 | 5 |
| chr2-45288856-45289357 | 5 |
| chr19-19890398-19890899 | 1 |
| chr1-40135660-40136161 | 8 |

| | |
|---|---|
| chr18-16866870-16867371 | 5 |
| chr3-70541248-70541749 | 5 |
| chr17-39221807-39222308 | 8 |
| chr13-25231181-25231682 | 1 |
| chr5-109372206-109372707 | 6 |
| chr10-97962180-97962681 | 8 |
| chr15-75113719-75114220 | 5 |
| chr3-80354879-80355380 | 8 |
| chr1-179657652-179658153 | 8 |
| chr8-67673357-67673858 | 8 |
| chr4-145034657-145035158 | 8 |
| chr19-830400-830901 | 1 |
| chr5-87814235-87814736 | 8 |
| chrX-108619078-108619579 | 1 |
| chr7-93005838-93006339 | 5 |
| chr16-79909850-79910351 | 1 |
| chr4-12472859-12473360 | 6 |
| chr10-9915871-9916372 | 6 |
| chr16-45505130-45505631 | 1 |
| chr5-132179599-132180100 | 6 |
| chr3-81287845-81288346 | 6 |
| chr14-102093378-102093879 | 1 |
| chr2-96872306-96872807 | 8 |
| chr10-106585124-106585625 | 8 |
| chr5-10374161-10374662 | 5 |
| chrX-137299122-137299623 | 8 |
| chr9-104473985-104474486 | 5 |
| chr7-130541595-130542096 | 8 |
| chr16-80896143-80896644 | 8 |
| chr2-72375864-72376365 | 5 |
| chr19-50320693-50321194 | 5 |
| chr4-179429066-179429567 | 8 |
| chr17-81075048-81075549 | 5 |
| chr1-243052083-243052584 | 8 |
| chr4-160204334-160204835 | 8 |
| chr18-24629766-24630267 | 6 |
| chr19-50577430-50577931 | 8 |
| chrX-149790536-149791037 | 8 |
| chr2-45586771-45587272 | 8 |
| chr3-102486510-102487011 | 5 |
| chr6-99031724-99032225 | 6 |
| chrX-107241354-107241855 | 5 |
| chr6-93578466-93578967 | 6 |

| | |
|---|---|
| chr2-231202214-231202715 | 6 |
| chr12-4860098-4860599 | 8 |
| chr1-14397388-14397889 | 8 |
| chr7-128819528-128820029 | 8 |
| chr19-55506789-55507290 | 5 |
| chr1-51901307-51901808 | 8 |
| chr15-57752837-57753338 | 8 |
| chr20-1520228-1520729 | 8 |
| chr2-121331335-121331836 | 5 |
| chr13-102292532-102293033 | 5 |
| chr6-116110070-116110571 | 8 |
| chr10-64478037-64478538 | 8 |
| chr4-156320079-156320580 | 6 |
| chr18-62306780-62307281 | 8 |
| chr13-29928614-29929115 | 8 |
| chr16-52119998-52120499 | 8 |
| chr18-11445700-11446201 | 8 |
| chr3-14064427-14064928 | 8 |
| chr9-12036508-12037009 | 8 |
| chr16-45533754-45534255 | 8 |
| chr11-49197395-49197896 | 1 |
| chr4-169484715-169485216 | 8 |
| chr5-38493939-38494440 | 8 |
| chr20-5864186-5864687 | 5 |
| chr4-177528037-177528538 | 8 |
| chr9-40419514-40420015 | 5 |
| chr10-23451389-23451890 | 8 |
| chrX-97840450-97840951 | 8 |
| chr10-68864959-68865460 | 8 |
| chr15-4195325-4195826 | 5 |
| chr1-151339787-151340288 | 6 |
| chr5-104191180-104191681 | 1 |
| chr17-60837164-60837665 | 6 |
| chr19-19387270-19387771 | 8 |
| chr2-237733439-237733940 | 5 |
| chr3-120018846-120019347 | 1 |
| chr2-15687372-15687873 | 5 |
| chr4-44242572-44243073 | 8 |
| chr5-60944091-60944592 | 6 |
| chr3-1250340-1250841 | 6 |
| chr13-59417298-59417799 | 6 |
| chr11-29651655-29652156 | 8 |
| chrX-104692176-104692677 | 1 |

| | |
|---|---|
| chr1-224148179-224148680 | 8 |
| chr3-35749464-35749965 | 5 |
| chr14-80048255-80048756 | 6 |
| chr14-78819443-78819944 | 8 |
| chr3-51836642-51837143 | 6 |
| chr13-94701503-94702004 | 5 |
| chr18-73710545-73711046 | 1 |
| chr4-17361344-17361845 | 1 |
| chr13-42858440-42858941 | 8 |
| chr1-162436655-162437156 | 6 |
| chr8-98920862-98921363 | 6 |
| chr7-90674137-90674638 | 8 |
| chr19-50054110-50054611 | 8 |
| chr8-87265423-87265924 | 8 |
| chr15-6016959-6017460 | 6 |
| chr1-173384795-173385296 | 6 |
| chr5-23314238-23314739 | 6 |
| chr10-68640182-68640683 | 5 |
| chr9-112801079-112801580 | 5 |
| chr4-120226320-120226821 | 6 |
| chr5-148823531-148824032 | 8 |
| chr3-45886026-45886527 | 1 |
| chr7-56787051-56787552 | 6 |
| chr9-61174458-61174959 | 6 |
| chr16-55522619-55523120 | 8 |
| chr6-51396554-51397055 | 8 |
| chr3-58468595-58469096 | 6 |
| chr17-65548185-65548686 | 6 |
| chr5-160798983-160799484 | 8 |
| chr6-124637951-124638452 | 6 |
| chr2-187313418-187313919 | 1 |
| chr18-31223321-31223822 | 8 |
| chr3-122519130-122519631 | 1 |
| chr4-147583340-147583841 | 1 |
| chr13-87793507-87794008 | 5 |
| chr10-96192481-96192982 | 5 |
| chr9-91599119-91599620 | 1 |
| chr2-120877260-120877761 | 5 |
| chr18-42662612-42663113 | 6 |
| chr4-126186434-126186935 | 5 |
| chr18-19966466-19966967 | 8 |
| chr4-126601726-126602227 | 1 |
| chr16-66175425-66175926 | 6 |

| | |
|---|---|
| chrX-17260462-17260963 | 6 |
| chr1-118727005-118727506 | 6 |
| chr7-21547713-21548214 | 8 |
| chr7-118788641-118789142 | 6 |
| chrX-78145833-78146334 | 8 |
| chr15-48799187-48799688 | 8 |
| chr3-159076861-159077362 | 6 |
| chr4-66773546-66774047 | 6 |
| chr13-41902708-41903209 | 1 |
| chr8-101582765-101583266 | 8 |
| chr15-100028492-100028993 | 8 |
| chr8-54611885-54612386 | 6 |
| chr4-14538562-14539063 | 8 |
| chr6-137434505-137435006 | 1 |
| chr6-9446092-9446593 | 8 |
| chr9-78604831-78605332 | 6 |
| chr17-25587482-25587983 | 6 |
| chr3-158449248-158449749 | 8 |
| chr7-57547706-57548207 | 5 |
| chr7-43813431-43813932 | 1 |
| chr5-144861340-144861841 | 8 |
| chr2-124316803-124317304 | 8 |
| chr6-137046186-137046687 | 6 |
| chr2-77954376-77954877 | 8 |
| chr3-155769176-155769677 | 8 |
| chr1-37948128-37948629 | 5 |
| chr1-118095451-118095952 | 6 |
| chr1-84811743-84812244 | 8 |
| chr19-52828208-52828709 | 8 |
| chr2-143882866-143883367 | 8 |
| chr1-142195065-142195566 | 8 |
| chr8-110617700-110618201 | 6 |
| chr15-9068336-9068837 | 5 |
| chr12-38568831-38569332 | 8 |
| chr13-99185355-99185856 | 5 |
| chr10-12816244-12816745 | 8 |
| chr6-123846781-123847282 | 5 |
| chr4-8001459-8001960 | 6 |
| chr8-12720217-12720718 | 5 |
| chr8-67189792-67190293 | 1 |
| chr5-144893259-144893760 | 8 |
| chr20-29297014-29297515 | 5 |
| chr16-44852199-44852700 | 1 |

| | |
|---|---|
| chr16-73876103-73876604 | 6 |
| chr4-132940872-132941373 | 8 |
| chrX-34168680-34169181 | 8 |
| chr19-40554559-40555060 | 6 |
| chr8-109568953-109569454 | 8 |
| chr1-17986145-17986646 | 6 |
| chr2-25349750-25350251 | 1 |
| chr11-64743533-64744034 | 8 |
| chr3-99398370-99398871 | 6 |
| chr13-65690566-65691067 | 8 |
| chr1-204060778-204061279 | 1 |
| chr1-155514258-155514759 | 8 |
| chr18-59678824-59679325 | 8 |
| chr15-87331899-87332400 | 1 |
| chr1-218425208-218425709 | 8 |
| chr1-120401842-120402343 | 8 |
| chr5-56359032-56359533 | 8 |
| chr4-160479877-160480378 | 8 |
| chr7-99024447-99024948 | 8 |
| chr10-64653227-64653728 | 8 |
| chr18-73674476-73674977 | 8 |
| chr1-260178270-260178771 | 6 |
| chr3-11902159-11902660 | 8 |
| chr11-76668563-76669064 | 8 |
| chr2-15647542-15648043 | 8 |
| chr3-144971045-144971546 | 8 |
| chr10-95942051-95942552 | 5 |
| chr13-70154711-70155212 | 5 |
| chrX-16084539-16085040 | 1 |
| chr5-5157248-5157749 | 6 |
| chr1-135232209-135232710 | 8 |
| chr10-41004796-41005297 | 6 |
| chr5-67897942-67898443 | 8 |
| chr4-12749196-12749697 | 8 |
| chr17-77428457-77428958 | 8 |
| chr17-35708121-35708622 | 1 |
| chr2-30335198-30335699 | 5 |
| chr12-35545393-35545894 | 6 |
| chr9-106325685-106326186 | 1 |
| chr12-4630885-4631386 | 8 |
| chr17-13811171-13811672 | 5 |
| chr3-37160042-37160543 | 5 |
| chr16-12685402-12685903 | 8 |

| | |
|---|---|
| chr8-74857714-74858215 | 6 |
| chr10-24628230-24628731 | 5 |
| chr9-57351467-57351968 | 6 |
| chr17-66379471-66379972 | 6 |
| chr16-804352-804853 | 8 |
| chr3-116323678-116324179 | 8 |
| chr4-29675159-29675660 | 6 |
| chr10-95452090-95452591 | 6 |
| chr9-89263385-89263886 | 6 |
| chr10-70960704-70961205 | 5 |
| chr20-20164309-20164810 | 8 |
| chr9-65241341-65241842 | 8 |
| chr4-78202954-78203455 | 8 |
| chr13-21421226-21421727 | 8 |
| chr17-72482471-72482972 | 8 |
| chr4-86138338-86138839 | 6 |
| chr19-44724857-44725358 | 1 |
| chr19-19152049-19152550 | 6 |
| chr17-37661442-37661943 | 6 |
| chr9-89368613-89369114 | 8 |
| chr4-150573793-150574294 | 1 |
| chr20-46311922-46312423 | 5 |
| chr7-79662840-79663341 | 8 |
| chr3-7873970-7874471 | 8 |
| chr6-29686417-29686918 | 6 |
| chr12-24995764-24996265 | 5 |
| chr19-27623119-27623620 | 6 |
| chr13-55737478-55737979 | 5 |
| chr6-41515621-41516122 | 8 |
| chr15-7012720-7013221 | 8 |
| chr18-5251924-5252425 | 1 |
| chr7-73441993-73442494 | 8 |
| chr1-30915816-30916317 | 8 |
| chr1-1386586-1387087 | 5 |
| chr4-129450184-129450685 | 8 |
| chr7-55642752-55643253 | 8 |
| chr3-163606571-163607072 | 8 |
| chr9-77109287-77109788 | 8 |
| chr4-168095401-168095902 | 6 |
| chr11-44847122-44847623 | 1 |
| chr2-192300961-192301462 | 8 |
| chr7-92972875-92973376 | 1 |
| chr6-15416152-15416653 | 1 |

| | |
|---|---|
| chr7-7567177-7567678 | 1 |
| chr19-812245-812746 | 6 |
| chr2-174198502-174199003 | 8 |
| chr5-101240305-101240806 | 8 |
| chr6-21645182-21645683 | 8 |
| chr5-105934113-105934614 | 8 |
| chr1-98778970-98779471 | 6 |
| chr1-156787992-156788493 | 8 |
| chr6-53246136-53246637 | 1 |
| chr14-103008609-103009110 | 6 |
| chr18-34219832-34220333 | 8 |
| chr6-121598167-121598668 | 1 |
| chr8-101700574-101701075 | 8 |
| chr18-22592030-22592531 | 6 |
| chr14-65040736-65041237 | 5 |
| chr3-144144475-144144976 | 8 |
| chr3-155777422-155777923 | 8 |
| chr16-67180227-67180728 | 8 |
| chr4-37921032-37921533 | 8 |
| chr13-99498122-99498623 | 8 |
| chrX-85373688-85374189 | 6 |
| chr19-50934950-50935451 | 8 |
| chr2-247746256-247746757 | 5 |
| chr5-124656271-124656772 | 6 |
| chr4-31785883-31786384 | 8 |
| chr14-15154456-15154957 | 5 |
| chr1-181775133-181775634 | 5 |
| chr15-34494011-34494512 | 8 |
| chr2-215239469-215239970 | 8 |
| chr13-67404399-67404900 | 8 |
| chr20-7620471-7620972 | 6 |
| chrX-41478517-41479018 | 8 |
| chr13-81897732-81898233 | 8 |
| chr2-241436681-241437182 | 8 |
| chr14-71433847-71434348 | 1 |
| chr7-71084748-71085249 | 8 |
| chr18-24148230-24148731 | 5 |
| chr9-40610057-40610558 | 6 |
| chrX-14284515-14285016 | 6 |
| chr7-117327485-117327986 | 6 |
| chr15-53203880-53204381 | 8 |
| chr20-42346249-42346750 | 8 |
| chr6-125935635-125936136 | 8 |

| | |
|---|---|
| chr9-4094929-4095430 | 5 |
| chr4-2401428-2401929 | 6 |
| chr2-137117520-137118021 | 8 |
| chr14-13592680-13593181 | 8 |
| chr17-22143197-22143698 | 8 |
| chr9-60679276-60679777 | 5 |
| chr2-228506124-228506625 | 6 |
| chr16-57457634-57458135 | 8 |
| chr2-57285117-57285618 | 8 |
| chr7-116650804-116651305 | 8 |
| chr14-101026662-101027163 | 1 |
| chr17-59305672-59306173 | 1 |
| chr2-39687293-39687794 | 6 |
| chr19-38067817-38068318 | 5 |
| chr7-103997601-103998102 | 6 |
| chr6-125857587-125858088 | 8 |
| chr3-139518129-139518630 | 5 |
| chr17-30985442-30985943 | 6 |
| chr5-47296657-47297158 | 8 |
| chr18-37835519-37836020 | 8 |
| chr13-39892194-39892695 | 8 |
| chr16-25499114-25499615 | 8 |
| chr4-81997121-81997622 | 6 |
| chr15-8190034-8190535 | 8 |
| chr18-44925453-44925954 | 5 |
| chr5-128454663-128455164 | 8 |
| chr7-126278506-126279007 | 6 |
| chr15-97629899-97630400 | 8 |
| chr8-62533999-62534500 | 6 |
| chr17-48697463-48697964 | 8 |
| chr16-69376983-69377484 | 8 |
| chr4-161617537-161618038 | 8 |
| chr4-145357678-145358179 | 8 |
| chr10-101016521-101017022 | 8 |
| chr9-45003886-45004387 | 8 |
| chr13-43092502-43093003 | 8 |
| chr17-26106803-26107304 | 8 |
| chr7-35810665-35811166 | 8 |
| chr7-46300779-46301280 | 8 |
| chr17-1475969-1476470 | 8 |
| chr1-133210320-133210821 | 8 |
| chr3-18740164-18740665 | 8 |
| chr10-4922871-4923372 | 8 |

| | |
|---|---|
| chr4-120151940-120152441 | 8 |
| chr3-147998334-147998835 | 6 |
| chr19-55167875-55168376 | 6 |
| chr1-94941861-94942362 | 6 |
| chr2-174923531-174924032 | 5 |
| chr14-40755642-40756143 | 8 |
| chr3-78987602-78988103 | 6 |
| chr17-79790910-79791411 | 6 |
| chr7-124342834-124343335 | 8 |
| chr10-8414484-8414985 | 5 |
| chr14-83168065-83168566 | 6 |
| chr14-32461993-32462494 | 8 |
| chr3-149989263-149989764 | 6 |
| chr14-27037315-27037816 | 8 |
| chr20-6070618-6071119 | 5 |
| chr8-113316695-113317196 | 6 |
| chr10-101527945-101528446 | 6 |
| chr2-79841494-79841995 | 5 |
| chr2-38113748-38114249 | 1 |
| chr2-59489788-59490289 | 8 |
| chr10-60820487-60820988 | 8 |
| chr5-158905695-158906196 | 1 |
| chr1-53720278-53720779 | 8 |
| chr8-86459459-86459960 | 8 |
| chr15-37612405-37612906 | 8 |
| chr16-6920000-6920501 | 8 |
| chr3-127070877-127071378 | 8 |
| chr17-49075763-49076264 | 6 |
| chr14-82837928-82838429 | 8 |
| chr1-131962316-131962817 | 6 |
| chr1-134536367-134536868 | 1 |
| chr15-35008736-35009237 | 8 |
| chr2-104279406-104279907 | 8 |
| chr7-54411589-54412090 | 6 |
| chr2-53472679-53473180 | 5 |
| chr1-201789784-201790285 | 8 |
| chr7-115367627-115368128 | 8 |
| chr4-133179768-133180269 | 8 |
| chr15-28668401-28668902 | 8 |
| chr10-73475139-73475640 | 5 |
| chr7-43460403-43460904 | 6 |
| chr2-40324710-40325211 | 6 |
| chr5-133710958-133711459 | 8 |

| | |
|---|---|
| chr14-32622138-32622639 | 1 |
| chr5-58064132-58064633 | 8 |
| chr3-40667075-40667576 | 1 |
| chrX-26960811-26961312 | 8 |
| chr8-55321570-55322071 | 6 |
| chr4-9806122-9806623 | 8 |
| chr18-67151130-67151631 | 1 |
| chr14-94868876-94869377 | 5 |
| chr13-91024998-91025499 | 5 |
| chr14-31750604-31751105 | 8 |
| chr18-53271400-53271901 | 8 |
| chr4-153022376-153022877 | 8 |
| chr20-21506899-21507400 | 8 |
| chr6-25600857-25601358 | 8 |
| chr18-76220157-76220658 | 6 |
| chr7-133696687-133697188 | 8 |
| chr1-41963804-41964305 | 1 |
| chr4-168516225-168516726 | 8 |
| chr12-26776622-26777123 | 8 |
| chr18-33889140-33889641 | 1 |
| chr1-87821604-87822105 | 8 |
| chr9-27168917-27169418 | 6 |
| chr9-65334854-65335355 | 1 |
| chr16-72647438-72647939 | 8 |
| chr1-167110920-167111421 | 6 |
| chr2-15544255-15544756 | 6 |
| chr1-83441429-83441930 | 5 |
| chr1-29634727-29635228 | 1 |
| chr13-90006867-90007368 | 8 |
| chr11-18104844-18105345 | 6 |
| chr8-118291814-118292315 | 8 |
| chr3-78599363-78599864 | 8 |
| chr16-48912932-48913433 | 8 |
| chr4-72147825-72148326 | 6 |
| chr1-93909692-93910193 | 8 |
| chr10-5030542-5031043 | 5 |
| chr17-570738-571239 | 1 |
| chr8-62028551-62029052 | 6 |
| chrX-111400034-111400535 | 5 |
| chr20-3111231-3111732 | 8 |
| chr13-59164114-59164615 | 6 |
| chr16-76443933-76444434 | 8 |
| chr7-46192268-46192769 | 8 |

| | |
|---|---|
| chr16-48822390-48822891 | 6 |
| chr18-27990331-27990832 | 8 |
| chr1-151990096-151990597 | 5 |
| chr3-147210900-147211401 | 8 |
| chr2-239793887-239794388 | 1 |
| chr8-54756532-54757033 | 6 |
| chr2-8461663-8462164 | 1 |
| chr18-12256438-12256939 | 1 |
| chr6-63596775-63597276 | 6 |
| chr12-7398920-7399421 | 6 |
| chr3-56950275-56950776 | 1 |
| chr6-6887002-6887503 | 5 |
| chr2-237233318-237233819 | 6 |
| chr15-95863329-95863830 | 1 |
| chr2-226033127-226033628 | 8 |
| chr9-88296693-88297194 | 1 |
| chr17-1168318-1168819 | 8 |
| chr10-43647486-43647987 | 8 |
| chr1-134906720-134907221 | 8 |
| chr10-21445955-21446456 | 1 |
| chr3-77646002-77646503 | 8 |
| chr12-26364399-26364900 | 1 |
| chr16-44389304-44389805 | 8 |
| chr2-11456689-11457190 | 1 |
| chr15-33187992-33188493 | 1 |
| chr1-201642096-201642597 | 8 |
| chr3-94590099-94590600 | 1 |
| chr2-203906140-203906641 | 5 |
| chr12-24094034-24094535 | 6 |
| chr9-54053766-54054267 | 1 |
| chr11-16032943-16033444 | 5 |
| chr16-38627978-38628479 | 6 |
| chr2-113223890-113224391 | 1 |
| chr15-12379743-12380244 | 6 |
| chr6-21638072-21638573 | 8 |
| chr6-79060594-79061095 | 6 |
| chr3-116067059-116067560 | 8 |
| chr12-24880646-24881147 | 6 |
| chr16-79253497-79253998 | 6 |
| chr8-107251685-107252186 | 6 |
| chr10-85255239-85255740 | 8 |
| chr4-12486023-12486524 | 1 |
| chr6-130004706-130005207 | 6 |

| | |
|---|---|
| chr1-258581737-258582238 | 6 |
| chr4-126506354-126506855 | 1 |
| chr14-83719149-83719650 | 8 |
| chr7-16747088-16747589 | 6 |
| chr1-241697487-241697988 | 6 |
| chr7-126019353-126019854 | 8 |
| chr3-159318994-159319495 | 8 |
| chr2-52670264-52670765 | 6 |
| chr5-51633126-51633627 | 6 |
| chr17-79167638-79168139 | 8 |
| chr5-22572962-22573463 | 8 |
| chr6-98365638-98366139 | 1 |
| chr10-8587628-8588129 | 8 |
| chr5-99810625-99811126 | 1 |
| chr1-207664989-207665490 | 8 |
| chr13-91095520-91096021 | 8 |
| chr17-4604304-4604805 | 8 |
| chr3-165909292-165909793 | 8 |
| chr18-24999683-25000184 | 6 |
| chr15-74287250-74287751 | 1 |
| chr11-82239525-82240026 | 6 |
| chr9-106800537-106801038 | 8 |
| chr15-15113708-15114209 | 6 |
| chrX-131414087-131414588 | 8 |
| chr4-35852478-35852979 | 8 |
| chr10-39208580-39209081 | 8 |
| chr18-50525797-50526298 | 6 |
| chr3-56121603-56122104 | 6 |
| chr18-33931684-33932185 | 5 |
| chr7-24137981-24138482 | 8 |
| chr5-1882747-1883248 | 8 |
| chr2-43784616-43785117 | 8 |
| chr14-9629189-9629690 | 8 |
| chr4-94248290-94248791 | 5 |
| chr17-34899981-34900482 | 5 |
| chr10-38829664-38830165 | 8 |
| chr5-82481989-82482490 | 5 |
| chr4-139979899-139980400 | 8 |
| chr1-151230178-151230679 | 8 |
| chr18-40876639-40877140 | 8 |
| chr4-148292535-148293036 | 6 |
| chr7-129237847-129238348 | 8 |
| chr8-118124960-118125461 | 8 |

| | |
|---|---|
| chr4-174082147-174082648 | 5 |
| chr1-239708300-239708801 | 5 |
| chr2-110360133-110360634 | 8 |
| chr16-78902204-78902705 | 5 |
| chr2-127046250-127046751 | 6 |
| chr7-116451338-116451839 | 1 |
| chr7-124373101-124373602 | 1 |
| chr3-140682259-140682760 | 6 |
| chr1-56090911-56091412 | 6 |
| chr6-20282995-20283496 | 1 |
| chr11-23827449-23827950 | 1 |
| chr17-74377663-74378164 | 8 |
| chr6-8416309-8416810 | 6 |
| chr5-7638870-7639371 | 1 |
| chr20-9743957-9744458 | 8 |
| chr8-69830161-69830662 | 8 |
| chr20-25584074-25584575 | 6 |
| chr11-82689475-82689976 | 8 |
| chr12-37482629-37483130 | 5 |
| chr9-76303240-76303741 | 8 |
| chr13-73703715-73704216 | 5 |
| chr5-33135237-33135738 | 6 |
| chr9-88561747-88562248 | 1 |
| chr2-203560035-203560536 | 1 |
| chr8-53790506-53791007 | 6 |
| chrX-106770745-106771246 | 6 |
| chr5-140334550-140335051 | 5 |
| chr6-103835617-103836118 | 1 |
| chr11-68720905-68721406 | 8 |
| chr14-30758886-30759387 | 1 |
| chr1-100165038-100165539 | 8 |
| chrX-40136620-40137121 | 6 |
| chr5-96772599-96773100 | 8 |
| chr15-78228664-78229165 | 1 |
| chr5-15168146-15168647 | 1 |
| chr1-5174582-5175083 | 6 |
| chr5-113682594-113683095 | 8 |
| chr8-99735062-99735563 | 1 |
| chr6-101966698-101967199 | 8 |
| chr17-34160072-34160573 | 8 |
| chr15-13299556-13300057 | 8 |
| chr1-135862436-135862937 | 8 |
| chr1-78633128-78633629 | 8 |

| | |
|---|---|
| chr6-37969514-37970015 | 6 |
| chr12-43169825-43170326 | 8 |
| chr4-37148629-37149130 | 6 |
| chr8-10461333-10461834 | 5 |
| chr14-30766614-30767115 | 1 |
| chr13-76819987-76820488 | 5 |
| chr3-14924984-14925485 | 8 |
| chr12-7125304-7125805 | 1 |
| chrX-126523109-126523610 | 6 |
| chr14-101947801-101948302 | 8 |
| chr3-7114369-7114870 | 6 |
| chr7-111757029-111757530 | 8 |
| chr10-68148318-68148819 | 1 |
| chr17-53096992-53097493 | 8 |
| chr2-224512481-224512982 | 5 |
| chr3-37036422-37036923 | 5 |
| chr17-65269318-65269819 | 8 |
| chr5-154452697-154453198 | 6 |
| chr13-93749588-93750089 | 8 |
| chr12-10677231-10677732 | 8 |
| chr20-16505942-16506443 | 6 |
| chr1-136244964-136245465 | 5 |
| chr19-11231522-11232023 | 8 |
| chr11-84596289-84596790 | 8 |
| chr13-82362100-82362601 | 8 |
| chr15-90411667-90412168 | 8 |
| chr9-107516218-107516719 | 1 |
| chr4-29340838-29341339 | 8 |
| chr2-233383344-233383845 | 8 |
| chr7-93773337-93773838 | 6 |
| chr18-48845828-48846329 | 5 |
| chr7-92032568-92033069 | 6 |
| chr19-9329241-9329742 | 8 |
| chr17-70382332-70382833 | 8 |
| chr8-36747758-36748259 | 8 |
| chr16-51790880-51791381 | 1 |
| chr3-30874694-30875195 | 5 |
| chr2-26060296-26060797 | 8 |
| chr17-25147176-25147677 | 6 |
| chr1-190928605-190929106 | 8 |
| chr17-67338888-67339389 | 8 |
| chr2-163333089-163333590 | 8 |
| chrX-41678416-41678917 | 8 |

| | |
|---|---|
| chr4-68137463-68137964 | 5 |
| chr4-103753333-103753834 | 5 |
| chr8-58474858-58475359 | 8 |
| chr9-16290738-16291239 | 8 |
| chr19-57326233-57326734 | 5 |
| chr8-22326846-22327347 | 6 |
| chr9-42545520-42546021 | 1 |
| chr6-63576461-63576962 | 8 |
| chr10-4597495-4597996 | 6 |
| chr16-31453599-31454100 | 6 |
| chr3-143751417-143751918 | 8 |
| chr9-59371941-59372442 | 8 |
| chr5-47276616-47277117 | 8 |
| chr16-34025277-34025778 | 8 |
| chr3-158858629-158859130 | 8 |
| chr18-56472464-56472965 | 8 |
| chr3-132458470-132458971 | 5 |
| chr7-95874098-95874599 | 8 |
| chr8-120678632-120679133 | 8 |
| chr6-129740708-129741209 | 5 |
| chr11-13447087-13447588 | 6 |
| chr3-109639178-109639679 | 6 |
| chr8-37262486-37262987 | 5 |
| chr19-39051092-39051593 | 5 |
| chr12-24759424-24759925 | 8 |
| chr20-30211722-30212223 | 8 |
| chr18-49901471-49901972 | 5 |
| chr10-37087350-37087851 | 1 |
| chr7-116551204-116551705 | 5 |
| chrX-12415920-12416421 | 8 |
| chr11-46482324-46482825 | 6 |
| chr14-42638583-42639084 | 8 |
| chr3-29082841-29083342 | 8 |
| chr2-6954421-6954922 | 8 |
| chr8-45691910-45692411 | 8 |
| chr19-19384020-19384521 | 5 |
| chr2-81356207-81356708 | 5 |
| chr6-32664804-32665305 | 6 |
| chr20-45481466-45481967 | 6 |
| chr1-218561436-218561937 | 1 |
| chr10-40988404-40988905 | 1 |
| chr18-24424327-24424828 | 8 |
| chr12-34108021-34108522 | 8 |

| | |
|---|---|
| chr3-34443890-34444391 | 8 |
| chr11-66335627-66336128 | 1 |
| chr2-171597643-171598144 | 8 |
| chr2-185950670-185951171 | 5 |
| chr7-129706451-129706952 | 8 |
| chr3-117794627-117795128 | 5 |
| chr15-19829769-19830270 | 8 |
| chr8-110352003-110352504 | 8 |
| chr2-134283303-134283804 | 8 |
| chr9-82169660-82170161 | 6 |
| chr5-95961693-95962194 | 5 |
| chr2-166235030-166235531 | 6 |
| chr8-78844447-78844948 | 8 |
| chr2-194616333-194616834 | 6 |
| chr1-17174679-17175180 | 8 |
| chr16-12030514-12031015 | 6 |
| chr2-241336108-241336609 | 8 |
| chr8-48972224-48972725 | 8 |
| chr3-153668124-153668625 | 8 |
| chr3-161813115-161813616 | 8 |
| chr5-122867360-122867861 | 8 |
| chr6-69511776-69512277 | 6 |
| chr5-141692640-141693141 | 8 |
| chr8-48102383-48102884 | 8 |
| chr11-74851366-74851867 | 6 |
| chr3-56088811-56089312 | 8 |
| chr5-160882421-160882922 | 8 |
| chr14-88755852-88756353 | 6 |
| chr1-231894863-231895364 | 1 |
| chr8-4982214-4982715 | 5 |
| chr7-26523117-26523618 | 6 |
| chr20-47124876-47125377 | 8 |
| chr17-81397525-81398026 | 5 |
| chr15-20767754-20768255 | 8 |
| chr17-45712022-45712523 | 8 |
| chr18-31036553-31037054 | 1 |
| chr13-38512842-38513343 | 8 |
| chr3-35847449-35847950 | 6 |
| chr1-46867725-46868226 | 6 |
| chr2-125727043-125727544 | 5 |
| chr14-83242791-83243292 | 6 |
| chr14-98402979-98403480 | 8 |
| chr14-21280269-21280770 | 6 |

| | |
|---|---|
| chr20-3447685-3448186 | 5 |
| chr4-62965493-62965994 | 5 |
| chr6-69323476-69323977 | 6 |
| chr1-234017404-234017905 | 6 |
| chr4-59131217-59131718 | 6 |
| chr3-14460136-14460637 | 8 |
| chr2-238032325-238032826 | 1 |
| chr1-136555336-136555837 | 8 |
| chr7-114363711-114364212 | 1 |
| chr8-86496934-86497435 | 8 |
| chr13-39161930-39162431 | 8 |
| chr1-49515299-49515800 | 8 |
| chr17-79598247-79598748 | 8 |
| chr16-13030625-13031126 | 6 |
| chr1-194244440-194244941 | 5 |
| chr1-52006814-52007315 | 6 |
| chr1-100004348-100004849 | 6 |
| chr17-15385516-15386017 | 8 |
| chr4-161548397-161548898 | 1 |
| chr7-132672932-132673433 | 8 |
| chr4-19368306-19368807 | 5 |
| chr8-81840980-81841481 | 6 |
| chr6-77318139-77318640 | 1 |
| chr12-618174-618675 | 8 |
| chr7-51711103-51711604 | 5 |
| chr3-17534428-17534929 | 8 |
| chr16-16964134-16964635 | 8 |
| chr1-225070897-225071398 | 8 |
| chr2-105672394-105672895 | 5 |
| chr8-6090988-6091489 | 5 |
| chr9-38671368-38671869 | 1 |
| chr6-46661374-46661875 | 1 |
| chr3-78144824-78145325 | 8 |
| chr7-108285638-108286139 | 8 |
| chr4-126653394-126653895 | 6 |
| chr14-77548419-77548920 | 8 |
| chr3-154845995-154846496 | 6 |
| chr8-23028869-23029370 | 8 |
| chr16-20873616-20874117 | 8 |
| chr16-47465077-47465578 | 8 |
| chr2-231030717-231031218 | 1 |
| chr1-193963201-193963702 | 8 |
| chr1-125070007-125070508 | 1 |

| | |
|---|---|
| chrX-10282368-10282869 | 1 |
| chr12-20504387-20504888 | 6 |
| chr17-82388991-82389492 | 6 |
| chr1-218406815-218407316 | 5 |
| chr5-24295676-24296177 | 8 |
| chr2-238141598-238142099 | 8 |
| chr18-68249632-68250133 | 5 |
| chr5-71933635-71934136 | 8 |
| chr3-28656418-28656919 | 8 |
| chr13-44663165-44663666 | 8 |
| chr8-68647241-68647742 | 5 |
| chr1-11492487-11492988 | 6 |
| chr18-26869410-26869911 | 6 |
| chr9-69646042-69646543 | 8 |
| chr6-28120268-28120769 | 8 |
| chr5-139408240-139408741 | 8 |
| chr10-15829200-15829701 | 8 |
| chr17-7318851-7319352 | 5 |
| chr8-122372320-122372821 | 8 |
| chr8-64814903-64815404 | 8 |
| chrX-3984142-3984643 | 1 |
| chr5-15817782-15818283 | 8 |
| chr15-10683844-10684345 | 8 |
| chr13-79021126-79021627 | 6 |
| chr7-96704367-96704868 | 6 |
| chr6-108522006-108522507 | 8 |
| chr19-25019259-25019760 | 8 |
| chr2-190650260-190650761 | 8 |
| chr10-65314630-65315131 | 8 |
| chr19-28594408-28594909 | 8 |
| chr8-44110696-44111197 | 8 |
| chr10-39230848-39231349 | 5 |
| chr17-70714791-70715292 | 6 |
| chr7-17457748-17458249 | 5 |
| chr6-33577195-33577696 | 8 |
| chr15-39217323-39217824 | 8 |
| chr2-81627221-81627722 | 6 |
| chr3-60389658-60390159 | 1 |
| chr5-71774453-71774954 | 5 |
| chr5-160350988-160351489 | 1 |
| chr7-30272346-30272847 | 1 |
| chr5-102156041-102156542 | 8 |
| chr8-118412281-118412782 | 8 |

| | |
|---|---|
| chr10-15877061-15877562 | 8 |
| chr14-58076220-58076721 | 6 |
| chr3-149460778-149461279 | 1 |
| chr5-121492696-121493197 | 8 |
| chr5-158547769-158548270 | 8 |
| chrX-128671021-128671522 | 6 |
| chr10-103643252-103643753 | 8 |
| chr13-92723245-92723746 | 5 |
| chr5-134787350-134787851 | 8 |
| chr20-8031011-8031512 | 8 |
| chr8-44868672-44869173 | 8 |
| chr7-96944701-96945202 | 6 |
| chr1-43379191-43379692 | 6 |
| chr19-45550773-45551274 | 8 |
| chr1-143505528-143506029 | 1 |
| chr15-32098639-32099140 | 8 |
| chr14-34788534-34789035 | 8 |
| chr2-187184321-187184822 | 6 |
| chr3-162715696-162716197 | 8 |
| chr2-121291965-121292466 | 5 |
| chr4-70502241-70502742 | 6 |
| chr6-399440-399941 | 8 |
| chr9-75497027-75497528 | 5 |
| chr6-35055751-35056252 | 8 |
| chr9-81621796-81622297 | 8 |
| chr8-114391020-114391521 | 8 |
| chr2-225893037-225893538 | 1 |
| chr6-3674059-3674560 | 5 |
| chr4-18739407-18739908 | 8 |
| chr10-103995218-103995719 | 8 |
| chr5-21437914-21438415 | 5 |
| chr14-96722310-96722811 | 5 |
| chr12-42048328-42048829 | 5 |
| chr1-255099888-255100389 | 6 |
| chr19-49401088-49401589 | 5 |
| chr6-21690810-21691311 | 5 |
| chr4-77819907-77820408 | 8 |
| chr2-101642545-101643046 | 8 |
| chr15-33833685-33834186 | 8 |
| chr8-118174133-118174634 | 8 |
| chr19-50449146-50449647 | 8 |
| chr1-256477023-256477524 | 8 |
| chr7-57561506-57562007 | 5 |

| | |
|---|---|
| chr1-121106645-121107146 | 1 |
| chr19-36883218-36883719 | 5 |
| chr13-65030804-65031305 | 8 |
| chr2-192140730-192141231 | 8 |
| chr3-59446530-59447031 | 8 |
| chr18-15510636-15511137 | 8 |
| chr1-32168471-32168972 | 8 |
| chr4-130339235-130339736 | 8 |
| chr18-61824413-61824914 | 5 |
| chr12-11077940-11078441 | 5 |
| chr10-6651070-6651571 | 8 |
| chr2-102043961-102044462 | 5 |
| chr1-171784719-171785220 | 6 |
| chr18-67916798-67917299 | 8 |
| chr5-66764105-66764606 | 5 |
| chr1-49126361-49126862 | 6 |
| chr6-14499410-14499911 | 8 |
| chr2-153831931-153832432 | 8 |
| chr12-3697685-3698186 | 6 |
| chr7-43759663-43760164 | 6 |
| chr15-74808645-74809146 | 1 |
| chr4-50552372-50552873 | 5 |
| chr2-225264436-225264937 | 8 |
| chr19-42983566-42984067 | 5 |
| chr14-81325395-81325896 | 8 |
| chr5-83913354-83913855 | 6 |
| chr16-65177776-65178277 | 1 |
| chr6-31781472-31781973 | 6 |
| chr2-190902147-190902648 | 5 |
| chr11-71517722-71518223 | 8 |
| chr1-21204673-21205174 | 8 |
| chr7-67943065-67943566 | 8 |
| chr7-92974652-92975153 | 6 |
| chr15-869434-869935 | 8 |
| chr13-33236201-33236702 | 5 |
| chr8-110555204-110555705 | 8 |
| chr15-13345476-13345977 | 5 |
| chr6-124121613-124122114 | 8 |
| chr5-121471944-121472445 | 8 |
| chr10-57741298-57741799 | 5 |
| chr18-49847640-49848141 | 8 |
| chrX-28021602-28022103 | 8 |
| chr5-123766872-123767373 | 8 |

| | |
|---|---|
| chr10-68910350-68910851 | 8 |
| chr2-175589870-175590371 | 8 |
| chr14-4044827-4045328 | 1 |
| chr16-6000237-6000738 | 6 |
| chr19-28340778-28341279 | 6 |
| chr5-21278807-21279308 | 6 |
| chr4-7341534-7342035 | 8 |
| chr6-71574606-71575107 | 8 |
| chr6-21896038-21896539 | 1 |
| chr13-102405181-102405682 | 6 |
| chr7-108636743-108637244 | 8 |
| chr1-7348205-7348706 | 8 |
| chr14-86742436-86742937 | 8 |
| chr20-8930522-8931023 | 5 |
| chr2-164652445-164652946 | 8 |
| chr17-78341071-78341572 | 5 |
| chr12-1550619-1551120 | 5 |
| chr3-89371527-89372028 | 8 |
| chr3-66196991-66197492 | 5 |
| chr2-235838057-235838558 | 8 |
| chr16-63518744-63519245 | 5 |
| chr15-62403832-62404333 | 1 |
| chr15-30455699-30456200 | 1 |
| chr5-3187908-3188409 | 8 |
| chr6-39188070-39188571 | 6 |
| chrX-31071665-31072166 | 5 |
| chr18-36659865-36660366 | 1 |
| chr10-64706870-64707371 | 6 |
| chr8-101150118-101150619 | 6 |
| chr2-113651280-113651781 | 8 |
| chr2-29016331-29016832 | 8 |
| chr3-117678000-117678501 | 1 |
| chr3-148872841-148873342 | 8 |
| chr6-93913186-93913687 | 8 |
| chr10-2604745-2605246 | 8 |
| chr1-84684224-84684725 | 8 |
| chr16-80381079-80381580 | 8 |
| chr6-84298457-84298958 | 6 |
| chr18-72933766-72934267 | 8 |
| chr16-44913901-44914402 | 1 |
| chr2-221033761-221034262 | 8 |
| chr9-9430115-9430616 | 6 |
| chrX-134943284-134943785 | 8 |

| | |
|---|---|
| chr18-68632318-68632819 | 8 |
| chr11-77276410-77276911 | 1 |
| chr3-88610604-88611105 | 8 |
| chr20-20028860-20029361 | 5 |
| chr2-171481043-171481544 | 8 |
| chr5-128131301-128131802 | 5 |
| chr13-97521549-97522050 | 8 |
| chr1-28539643-28540144 | 6 |
| chr7-55097024-55097525 | 5 |
| chr17-1966458-1966959 | 8 |
| chr5-85849940-85850441 | 8 |
| chr6-94455867-94456368 | 5 |
| chr4-181744799-181745300 | 6 |
| chr1-222453186-222453687 | 1 |
| chr14-34653756-34654257 | 8 |
| chr18-62374934-62375435 | 8 |
| chr1-258105382-258105883 | 8 |
| chr5-86494036-86494537 | 6 |
| chr7-55199864-55200365 | 8 |
| chr16-80624898-80625399 | 1 |
| chr4-65680295-65680796 | 6 |
| chr11-69743250-69743751 | 8 |
| chr8-31250517-31251018 | 8 |
| chr10-95971229-95971730 | 1 |
| chr2-243888786-243889287 | 6 |
| chr4-15344884-15345385 | 8 |
| chr4-13062680-13063181 | 8 |
| chr10-44970872-44971373 | 8 |
| chr19-53400148-53400649 | 8 |
| chr16-32523530-32524031 | 5 |
| chr16-45195093-45195594 | 5 |
| chr1-239376320-239376821 | 8 |
| chr9-78740278-78740779 | 1 |
| chr15-18433471-18433972 | 8 |
| chr3-141952178-141952679 | 5 |
| chr1-240805099-240805600 | 8 |
| chr9-96039766-96040267 | 6 |
| chr8-15125506-15126007 | 5 |
| chr13-104261447-104261948 | 8 |
| chr13-103339265-103339766 | 1 |
| chrX-2483309-2483810 | 6 |
| chr8-122141867-122142368 | 1 |
| chr3-69619435-69619936 | 5 |

| | |
|---|---|
| chr1-194830966-194831467 | 5 |
| chr12-44715695-44716196 | 8 |
| chrX-41160243-41160744 | 1 |
| chr11-58014846-58015347 | 1 |
| chr1-154930071-154930572 | 5 |
| chr9-91153530-91154031 | 8 |
| chr2-61929482-61929983 | 5 |
| chr14-6386242-6386743 | 8 |
| chr2-112960610-112961111 | 1 |
| chr4-153017150-153017651 | 8 |
| chr9-7443864-7444365 | 6 |
| chr4-175774173-175774674 | 8 |
| chr10-35822950-35823451 | 8 |
| chr4-80477093-80477594 | 6 |
| chr3-104711517-104712018 | 8 |
| chr2-44001833-44002334 | 8 |
| chr19-15223-15724 | 6 |
| chr1-40279200-40279701 | 1 |
| chr5-161126314-161126815 | 8 |
| chr6-139324238-139324739 | 8 |
| chr16-17803304-17803805 | 5 |
| chr10-63621812-63622313 | 6 |
| chr6-32149094-32149595 | 5 |
| chr10-57702521-57703022 | 8 |
| chr4-60770186-60770687 | 8 |
| chr10-63657323-63657824 | 8 |
| chr16-8326866-8327367 | 8 |
| chr16-43050329-43050830 | 8 |
| chr9-16135017-16135518 | 8 |
| chr2-136784569-136785070 | 6 |
| chr1-178827818-178828319 | 6 |
| chr15-4201686-4202187 | 8 |
| chr4-57111446-57111947 | 8 |
| chr1-53923176-53923677 | 5 |
| chr1-136203493-136203994 | 6 |
| chr15-6586836-6587337 | 6 |
| chr2-82543433-82543934 | 8 |
| chr12-38284829-38285330 | 8 |
| chr5-104556839-104557340 | 6 |
| chr15-16474417-16474918 | 8 |
| chr17-67849745-67850246 | 1 |
| chr9-64497898-64498399 | 8 |
| chr1-13924835-13925336 | 1 |

| | |
|---|---|
| chr17-35689163-35689664 | 1 |
| chr5-78465257-78465758 | 8 |
| chr9-34210049-34210550 | 6 |
| chr18-55941549-55942050 | 6 |
| chr13-78521216-78521717 | 5 |
| chr1-259217374-259217875 | 8 |
| chr1-214308862-214309363 | 6 |
| chr4-61438625-61439126 | 6 |
| chr7-119578811-119579312 | 6 |
| chr14-91892231-91892732 | 8 |
| chr13-50472172-50472673 | 5 |
| chr5-57246488-57246989 | 8 |
| chr8-64298249-64298750 | 5 |
| chr7-113502881-113503382 | 8 |
| chr2-232487493-232487994 | 5 |
| chr1-211622784-211623285 | 8 |
| chr4-174036915-174037416 | 8 |
| chr19-51350163-51350664 | 6 |
| chr15-45835413-45835914 | 8 |
| chr2-54913748-54914249 | 8 |
| chr13-92130097-92130598 | 1 |
| chr5-137750348-137750849 | 6 |
| chr1-77271871-77272372 | 6 |
| chr7-29105598-29106099 | 1 |
| chr8-53023770-53024271 | 8 |
| chr8-65685206-65685707 | 5 |
| chr18-41168357-41168858 | 1 |
| chr16-51421246-51421747 | 8 |
| chr9-11278495-11278996 | 6 |
| chr5-151585296-151585797 | 5 |
| chr2-137850919-137851420 | 8 |
| chr10-40253780-40254281 | 6 |
| chr8-106582215-106582716 | 8 |
| chr5-151297834-151298335 | 8 |
| chr14-103341003-103341504 | 8 |
| chr18-42661978-42662479 | 6 |
| chr3-104467024-104467525 | 6 |
| chr1-92109787-92110288 | 1 |
| chr13-62982659-62983160 | 5 |
| chr1-196292847-196293348 | 5 |
| chr6-58405385-58405886 | 5 |
| chr15-69812248-69812749 | 1 |
| chr10-44077462-44077963 | 8 |

| | |
|---|---|
| chr3-115314900-115315401 | 8 |
| chr3-55363593-55364094 | 8 |
| chr1-50169812-50170313 | 8 |
| chr12-14797441-14797942 | 5 |
| chr1-4574160-4574661 | 6 |
| chr19-1464134-1464635 | 5 |
| chr6-87556471-87556972 | 8 |
| chr13-63485492-63485993 | 6 |
| chr4-131578294-131578795 | 8 |
| chr14-91731001-91731502 | 8 |
| chr13-82082157-82082658 | 8 |
| chrX-150745456-150745957 | 8 |
| chr18-61241653-61242154 | 8 |
| chr6-87915531-87916032 | 8 |
| chr14-66316454-66316955 | 6 |
| chr7-94531371-94531872 | 6 |
| chr1-123897996-123898497 | 8 |
| chr3-166256112-166256613 | 6 |
| chr1-190924790-190925291 | 1 |
| chr13-32404571-32405072 | 8 |
| chr4-154207172-154207673 | 8 |
| chr10-22810038-22810539 | 6 |
| chr10-60704731-60705232 | 5 |
| chr2-120157750-120158251 | 8 |
| chr3-167238412-167238913 | 5 |
| chrX-151730329-151730830 | 6 |
| chr1-136155289-136155790 | 6 |
| chr16-72086466-72086967 | 8 |
| chr1-229458433-229458934 | 1 |
| chr3-60175416-60175917 | 1 |
| chr4-135721605-135722106 | 8 |
| chr18-29211615-29212116 | 5 |
| chr8-121747891-121748392 | 1 |
| chr11-29075876-29076377 | 5 |
| chr17-54193010-54193511 | 5 |
| chr16-43244404-43244905 | 5 |
| chr11-74753377-74753878 | 6 |
| chr12-3858365-3858866 | 6 |
| chr6-13928235-13928736 | 1 |
| chr10-69147029-69147530 | 1 |
| chr12-38339222-38339723 | 8 |
| chr18-72532772-72533273 | 8 |
| chr14-81128191-81128692 | 8 |

| | |
|---|---|
| chr1-153979862-153980363 | 5 |
| chr17-34851697-34852198 | 1 |
| chr11-164830-165331 | 6 |
| chr3-108642021-108642522 | 6 |
| chr10-105008419-105008920 | 8 |
| chr4-115970111-115970612 | 5 |
| chr11-3331302-3331803 | 6 |
| chr11-50620750-50621251 | 8 |
| chr3-70199527-70200028 | 8 |
| chr9-107411555-107412056 | 1 |
| chr4-47886484-47886985 | 5 |
| chr2-113668455-113668956 | 1 |
| chr18-82025954-82026455 | 5 |
| chr9-94334474-94334975 | 8 |
| chr18-26312810-26313311 | 8 |
| chr8-59359500-59360001 | 6 |
| chr19-48211475-48211976 | 8 |
| chr4-60713056-60713557 | 8 |
| chr1-177141813-177142314 | 8 |
| chr3-148553314-148553815 | 8 |
| chr2-110798099-110798600 | 1 |
| chr2-242340199-242340700 | 6 |
| chr5-147534366-147534867 | 1 |
| chr2-224581775-224582276 | 8 |
| chr20-1503772-1504273 | 8 |
| chr8-16141981-16142482 | 6 |
| chr1-30171016-30171517 | 5 |
| chr1-20766325-20766826 | 6 |
| chr2-173199429-173199930 | 8 |
| chr4-65573706-65574207 | 6 |
| chr3-133807891-133808392 | 6 |
| chr4-50443750-50444251 | 8 |
| chr2-132586845-132587346 | 6 |
| chr2-141202297-141202798 | 6 |
| chr5-76374438-76374939 | 8 |
| chr6-117153619-117154120 | 8 |
| chr18-21830794-21831295 | 8 |
| chr18-21099700-21100201 | 6 |
| chr13-48664170-48664671 | 6 |
| chr1-79069810-79070311 | 8 |
| chr14-78066529-78067030 | 8 |
| chr2-242369935-242370436 | 1 |
| chr3-115934096-115934597 | 6 |

| | |
|---|---|
| chr14-35391847-35392348 | 1 |
| chr1-87776345-87776846 | 1 |
| chr5-150007773-150008274 | 1 |
| chr1-232296893-232297394 | 1 |
| chr13-55712082-55712583 | 8 |
| chr3-125041340-125041841 | 1 |
| chr6-28382200-28382701 | 6 |
| chr11-35678107-35678608 | 8 |
| chr20-28794175-28794676 | 5 |
| chr8-116552605-116553106 | 6 |
| chr18-67150342-67150843 | 1 |
| chr17-8851944-8852445 | 5 |
| chr14-6725711-6726212 | 8 |
| chr5-146996436-146996937 | 8 |
| chr2-122703473-122703974 | 8 |
| chr3-128657168-128657669 | 6 |
| chr9-66146993-66147494 | 5 |
| chr11-22914766-22915267 | 5 |
| chrX-100118403-100118904 | 1 |
| chr10-44545298-44545799 | 8 |
| chr9-87729229-87729730 | 6 |
| chr1-249124174-249124675 | 8 |
| chr3-163010165-163010666 | 8 |
| chr8-55954464-55954965 | 1 |
| chr3-134812269-134812770 | 8 |
| chr5-33547969-33548470 | 8 |
| chr2-220300371-220300872 | 1 |
| chr8-20640891-20641392 | 8 |
| chr13-106482841-106483342 | 1 |
| chr17-82167522-82168023 | 8 |
| chr2-153926456-153926957 | 6 |
| chr11-62307202-62307703 | 6 |
| chr1-127910660-127911161 | 8 |
| chr18-55170242-55170743 | 8 |
| chr3-8830717-8831218 | 8 |
| chr7-85085507-85086008 | 8 |
| chr2-183377338-183377839 | 8 |
| chr8-37225374-37225875 | 8 |
| chr2-210731329-210731830 | 8 |
| chr6-39248945-39249446 | 6 |
| chr1-103620598-103621099 | 6 |
| chr5-6635857-6636358 | 1 |
| chr5-146155168-146155669 | 8 |

| | |
|---|---|
| chr10-19835064-19835565 | 8 |
| chr15-34837979-34838480 | 8 |
| chr5-97038387-97038888 | 5 |
| chr17-80945265-80945766 | 6 |
| chr2-123072916-123073417 | 8 |
| chr4-57199576-57200077 | 6 |
| chr19-9832884-9833385 | 8 |
| chr6-117951344-117951845 | 6 |
| chr7-65089839-65090340 | 1 |
| chr14-97996291-97996792 | 8 |
| chr1-239810801-239811302 | 8 |
| chr19-11061607-11062108 | 1 |
| chr2-134459182-134459683 | 8 |
| chr17-53039667-53040168 | 1 |
| chr19-10136564-10137065 | 8 |
| chr3-22609359-22609860 | 8 |
| chr1-227297317-227297818 | 8 |
| chr2-6923600-6924101 | 5 |
| chr2-136689530-136690031 | 8 |
| chr4-86188273-86188774 | 8 |
| chr4-144367284-144367785 | 6 |
| chrX-12530422-12530923 | 8 |
| chr4-45561476-45561977 | 1 |
| chr18-5021706-5022207 | 8 |
| chr2-50388472-50388973 | 6 |
| chr1-88565623-88566124 | 8 |
| chr8-87136833-87137334 | 8 |
| chr2-104273593-104274094 | 5 |
| chr11-47279966-47280467 | 6 |
| chr9-17986195-17986696 | 6 |
| chrX-35090360-35090861 | 5 |
| chr3-29999611-30000112 | 8 |
| chr16-75594795-75595296 | 5 |
| chr3-111728957-111729458 | 6 |
| chr15-24982138-24982639 | 8 |
| chr20-25211046-25211547 | 5 |
| chr2-189346384-189346885 | 8 |
| chr18-30822754-30823255 | 6 |
| chr6-139710595-139711096 | 8 |
| chr9-56997182-56997683 | 8 |
| chr10-50478414-50478915 | 8 |
| chr4-151356741-151357242 | 1 |
| chr1-236040298-236040799 | 6 |

| | |
|---|---|
| chr1-201313774-201314275 | 1 |
| chr14-66238026-66238527 | 8 |
| chr4-133749918-133750419 | 8 |
| chr3-50258829-50259330 | 6 |
| chr5-113070530-113071031 | 6 |
| chr18-46752383-46752884 | 6 |
| chr18-60592092-60592593 | 5 |
| chr6-44086542-44087043 | 8 |
| chr13-3238145-3238646 | 6 |
| chr6-130652664-130653165 | 1 |
| chr14-5647679-5648180 | 1 |
| chr15-1272128-1272629 | 8 |
| chr3-18640925-18641426 | 6 |
| chr19-44316434-44316935 | 5 |
| chr13-81754445-81754946 | 8 |
| chrX-4848701-4849202 | 8 |
| chr5-166375635-166376136 | 8 |
| chr13-48229651-48230152 | 6 |
| chr6-40731190-40731691 | 8 |
| chr1-85283662-85284163 | 1 |
| chr6-101690805-101691306 | 8 |
| chr4-90238192-90238693 | 5 |
| chr5-122614126-122614627 | 8 |
| chrX-78753094-78753595 | 6 |
| chr6-96886296-96886797 | 8 |
| chr4-130871788-130872289 | 8 |
| chr5-72714719-72715220 | 8 |
| chr7-66216797-66217298 | 1 |
| chr10-21992460-21992961 | 5 |
| chr5-40308535-40309036 | 6 |
| chr4-76479318-76479819 | 8 |
| chr4-154234061-154234562 | 5 |
| chr18-5753248-5753749 | 5 |
| chr7-46672752-46673253 | 8 |
| chr4-54238514-54239015 | 8 |
| chr5-137293688-137294189 | 8 |
| chr11-56576525-56577026 | 8 |
| chr15-13190462-13190963 | 8 |
| chr5-42952829-42953330 | 8 |
| chr9-104257731-104258232 | 8 |
| chr2-657710-658211 | 5 |
| chr6-23234045-23234546 | 8 |
| chr1-86474370-86474871 | 8 |

| | |
|---|---|
| chr7-105897045-105897546 | 8 |
| chr4-110234246-110234747 | 8 |
| chr9-66255178-66255679 | 8 |
| chr6-137353766-137354267 | 5 |
| chr3-104488946-104489447 | 5 |
| chr3-111497934-111498435 | 1 |
| chr13-83240688-83241189 | 8 |
| chr2-174192660-174193161 | 5 |
| chr2-46141332-46141833 | 8 |
| chr12-26761536-26762037 | 1 |
| chr3-44629202-44629703 | 6 |
| chr7-100011846-100012347 | 5 |
| chr1-199239411-199239912 | 8 |
| chr3-158446982-158447483 | 6 |
| chr9-43792821-43793322 | 8 |
| chr12-31972865-31973366 | 8 |
| chr7-56812108-56812609 | 5 |
| chr10-94189343-94189844 | 8 |
| chr2-188067203-188067704 | 6 |
| chr4-18357154-18357655 | 5 |
| chr13-20506813-20507314 | 5 |
| chr1-62087087-62087588 | 8 |
| chr1-214611544-214612045 | 8 |
| chr14-26847638-26848139 | 8 |
| chr8-116466068-116466569 | 8 |
| chr3-88213403-88213904 | 8 |
| chr2-111848202-111848703 | 6 |
| chr2-206966192-206966693 | 5 |
| chr6-39132014-39132515 | 8 |
| chr17-37847627-37848128 | 6 |
| chr8-123630942-123631443 | 8 |
| chr14-97173239-97173740 | 6 |
| chr11-15666493-15666994 | 8 |
| chr5-29789944-29790445 | 8 |
| chr15-24465014-24465515 | 1 |
| chr3-99279494-99279995 | 8 |
| chr1-192513260-192513761 | 8 |
| chr2-200130967-200131468 | 8 |
| chr18-54825730-54826231 | 8 |
| chr1-118096475-118096976 | 1 |
| chr1-199561264-199561765 | 8 |
| chr4-121371625-121372126 | 8 |
| chr17-64037979-64038480 | 8 |

| | |
|---|---|
| chr1-245531513-245532014 | 8 |
| chr1-258437742-258438243 | 5 |
| chr19-36406602-36407103 | 6 |
| chr7-90903562-90904063 | 5 |
| chr2-234523187-234523688 | 5 |
| chr5-108747549-108748050 | 6 |
| chr1-255902593-255903094 | 8 |
| chr5-8089388-8089889 | 5 |
| chr11-67193267-67193768 | 6 |
| chr11-71597660-71598161 | 6 |
| chr9-34536699-34537200 | 8 |
| chr4-40447805-40448306 | 6 |
| chr18-21180290-21180791 | 8 |
| chr17-69718597-69719098 | 8 |
| chr8-78280089-78280590 | 1 |
| chr1-96140245-96140746 | 8 |
| chr3-151967751-151968252 | 8 |
| chr13-93989605-93990106 | 8 |
| chr1-40376376-40376877 | 5 |
| chr6-95514059-95514560 | 5 |
| chr11-82275059-82275560 | 5 |
| chr4-160914303-160914804 | 8 |
| chr18-47054241-47054742 | 8 |
| chr11-12705336-12705837 | 5 |
| chr17-19912912-19913413 | 8 |
| chr19-21239461-21239962 | 8 |
| chr15-38641494-38641995 | 8 |
| chr3-14006633-14007134 | 8 |
| chr15-12698503-12699004 | 1 |
| chr15-45800419-45800920 | 6 |
| chr6-118160255-118160756 | 5 |
| chr9-103040542-103041043 | 8 |
| chr2-107632500-107633001 | 8 |
| chr10-52312416-52312917 | 1 |
| chr2-247038726-247039227 | 8 |
| chr6-42095316-42095817 | 1 |
| chr9-39729736-39730237 | 1 |
| chr2-28078437-28078938 | 8 |
| chr8-62780685-62781186 | 8 |
| chr4-162164125-162164626 | 1 |
| chr10-7931511-7932012 | 5 |
| chr1-13833479-13833980 | 1 |
| chr10-55306289-55306790 | 8 |

| | |
|---|---|
| chr9-111404376-111404877 | 8 |
| chr8-98268724-98269225 | 5 |
| chr16-29359624-29360125 | 5 |
| chrX-111617563-111618064 | 8 |
| chr2-114368129-114368630 | 6 |
| chr15-75434716-75435217 | 6 |
| chr9-31246518-31247019 | 6 |
| chr9-38366541-38367042 | 8 |
| chr19-29015093-29015594 | 6 |
| chr14-78512105-78512606 | 8 |
| chr10-72724065-72724566 | 8 |
| chr2-220643273-220643774 | 5 |
| chr2-90869935-90870436 | 6 |
| chr1-176748155-176748656 | 8 |
| chr6-92693871-92694372 | 5 |
| chr2-231950595-231951096 | 1 |
| chr3-120130981-120131482 | 6 |
| chr13-32747461-32747962 | 1 |
| chr18-59292807-59293308 | 1 |
| chr13-61596914-61597415 | 6 |
| chr18-59333674-59334175 | 8 |
| chr3-60026696-60027197 | 1 |
| chr1-94454240-94454741 | 6 |
| chr1-184968762-184969263 | 8 |
| chr8-101154339-101154840 | 6 |
| chr5-147873945-147874446 | 8 |
| chr6-22642947-22643448 | 8 |
| chr18-59128358-59128859 | 6 |
| chr4-120881308-120881809 | 5 |
| chr2-11336577-11337078 | 5 |
| chr3-122486220-122486721 | 1 |
| chr2-223646783-223647284 | 1 |
| chr2-210233135-210233636 | 8 |
| chr3-136334841-136335342 | 1 |
| chr10-34702983-34703484 | 8 |
| chr9-63923308-63923809 | 1 |
| chr9-17849121-17849622 | 8 |
| chr20-10399940-10400441 | 5 |
| chr10-76789514-76790015 | 8 |
| chrX-68392609-68393110 | 8 |
| chr6-92567654-92568155 | 6 |
| chr7-31949986-31950487 | 1 |
| chr14-35601162-35601663 | 8 |

| | |
|---|---|
| chr1-1010693-1011194 | 5 |
| chr1-102869824-102870325 | 8 |
| chr13-50367246-50367747 | 6 |
| chr7-114311866-114312367 | 8 |
| chr1-93629009-93629510 | 8 |
| chr2-20787372-20787873 | 8 |
| chr1-79523700-79524201 | 8 |
| chr14-56888635-56889136 | 8 |
| chr4-133748786-133749287 | 6 |
| chr9-38811261-38811762 | 8 |
| chr14-7300257-7300758 | 6 |
| chr3-112695433-112695934 | 1 |
| chr8-114813072-114813573 | 8 |
| chr1-219355124-219355625 | 8 |
| chr6-101941040-101941541 | 8 |
| chr11-56240318-56240819 | 8 |
| chr12-25408699-25409200 | 8 |
| chr1-227524599-227525100 | 5 |
| chr13-96053154-96053655 | 8 |
| chr2-215762972-215763473 | 8 |
| chr15-31264721-31265222 | 8 |
| chr11-75445958-75446459 | 6 |
| chr10-82283553-82284054 | 6 |
| chr1-22196292-22196793 | 8 |
| chr13-30245902-30246403 | 8 |
| chr7-130710454-130710955 | 8 |
| chr18-23674748-23675249 | 8 |
| chr2-122800076-122800577 | 6 |
| chr5-57101963-57102464 | 5 |
| chr5-58315846-58316347 | 8 |
| chr5-27094668-27095169 | 1 |
| chr6-110498156-110498657 | 8 |
| chr2-113179285-113179786 | 8 |
| chr2-166814066-166814567 | 6 |
| chr1-88606415-88606916 | 6 |
| chr2-219003803-219004304 | 8 |
| chr5-147279323-147279824 | 6 |
| chr14-42372648-42373149 | 8 |
| chr4-92425288-92425789 | 6 |
| chr1-158182339-158182840 | 6 |
| chr19-9305530-9306031 | 8 |
| chr1-134737307-134737808 | 8 |
| chr6-19885558-19886059 | 6 |

| | |
|---|---|
| chr1-86968105-86968606 | 5 |
| chr4-120956045-120956546 | 6 |
| chr3-65738072-65738573 | 5 |
| chr5-46215779-46216280 | 8 |
| chr11-65634657-65635158 | 6 |
| chr6-94587982-94588483 | 8 |
| chr3-85785964-85786465 | 6 |
| chrX-26615895-26616396 | 8 |
| chr11-35628710-35629211 | 8 |
| chr9-4838356-4838857 | 6 |
| chr4-65973287-65973788 | 8 |
| chr12-45324473-45324974 | 1 |
| chr1-96549979-96550480 | 5 |
| chr15-52907418-52907919 | 6 |
| chr14-66290017-66290518 | 1 |
| chr1-170026244-170026745 | 5 |
| chr11-33154143-33154644 | 8 |
| chr8-43971947-43972448 | 8 |
| chr7-45140327-45140828 | 8 |
| chr7-23970502-23971003 | 8 |
| chr1-77027701-77028202 | 8 |
| chr13-95781438-95781939 | 5 |
| chr19-51224489-51224990 | 8 |
| chr13-23791810-23792311 | 8 |
| chr7-71905840-71906341 | 6 |
| chr4-33173056-33173557 | 8 |
| chr1-192750121-192750622 | 5 |
| chr14-95632413-95632914 | 6 |
| chr4-178347781-178348282 | 6 |
| chr3-133611718-133612219 | 8 |
| chr2-11556142-11556643 | 5 |
| chr2-196964298-196964799 | 8 |
| chr20-5288137-5288638 | 5 |
| chr19-17389450-17389951 | 1 |
| chr1-118853168-118853669 | 8 |
| chr13-49609380-49609881 | 8 |
| chr14-82580301-82580802 | 5 |
| chr5-85917745-85918246 | 1 |
| chr5-159300804-159301305 | 8 |
| chr2-151284976-151285477 | 8 |
| chr14-7291330-7291831 | 8 |
| chr3-12917522-12918023 | 1 |
| chr5-8178446-8178947 | 1 |

| | |
|---|---|
| chr14-3212366-3212867 | 8 |
| chr9-58388092-58388593 | 6 |
| chr4-125452874-125453375 | 1 |
| chr17-29052028-29052529 | 8 |
| chr16-57815561-57816062 | 8 |
| chr18-39923972-39924473 | 6 |
| chr5-22198016-22198517 | 8 |
| chr15-12600999-12601500 | 8 |
| chr11-33774644-33775145 | 5 |
| chr3-117697774-117698275 | 8 |
| chr4-149008281-149008782 | 1 |
| chr7-126671892-126672393 | 5 |
| chr9-36536596-36537097 | 6 |
| chr20-42281408-42281909 | 8 |
| chr2-143932399-143932900 | 5 |
| chr14-23018538-23019039 | 5 |
| chr7-91468695-91469196 | 8 |
| chr2-54232668-54233169 | 1 |
| chr3-160582011-160582512 | 6 |
| chr9-94653154-94653655 | 5 |
| chr20-29259627-29260128 | 5 |
| chr4-28252389-28252890 | 6 |
| chr6-96465944-96466445 | 5 |
| chr5-55027803-55028304 | 6 |
| chr5-63617327-63617828 | 8 |
| chr7-50190006-50190507 | 6 |
| chr18-12603277-12603778 | 5 |
| chr9-4781883-4782384 | 6 |
| chr11-32832066-32832567 | 5 |
| chr7-55583555-55584056 | 8 |
| chrX-149687613-149688114 | 1 |
| chr7-58505058-58505559 | 5 |
| chr3-46112343-46112844 | 5 |
| chr1-141691473-141691974 | 6 |
| chr13-48930596-48931097 | 5 |
| chr1-18531008-18531509 | 8 |
| chr1-122683713-122684214 | 6 |
| chr1-33783524-33784025 | 6 |
| chr1-124985672-124986173 | 8 |
| chr6-22504772-22505273 | 8 |
| chr19-46611427-46611928 | 5 |
| chr1-103215200-103215701 | 6 |
| chr1-123591075-123591576 | 8 |

| | |
|---|---|
| chr3-131057106-131057607 | 5 |
| chr12-13004933-13005434 | 6 |
| chr14-62729129-62729630 | 6 |
| chr4-82124625-82125126 | 6 |
| chr17-83314188-83314689 | 8 |
| chr11-17282992-17283493 | 1 |
| chr5-20139708-20140209 | 1 |
| chr8-122054801-122055302 | 8 |
| chr8-64739745-64740246 | 5 |
| chr18-44834630-44835131 | 1 |
| chr8-110446032-110446533 | 8 |
| chr13-99360353-99360854 | 8 |
| chr7-94150880-94151381 | 5 |
| chr4-133043086-133043587 | 8 |
| chr18-30713494-30713995 | 8 |
| chr6-13573415-13573916 | 1 |
| chr1-257086109-257086610 | 8 |
| chr20-29803800-29804301 | 8 |
| chr19-9491222-9491723 | 6 |
| chr16-76354958-76355459 | 8 |
| chr14-69149746-69150247 | 8 |
| chr10-60322428-60322929 | 5 |
| chr6-7860166-7860667 | 1 |
| chr19-34423755-34424256 | 8 |
| chr3-22601947-22602448 | 5 |
| chr10-28611061-28611562 | 6 |
| chr13-70617739-70618240 | 6 |
| chr4-133680800-133681301 | 8 |
| chr2-110943310-110943811 | 8 |
| chr3-104168935-104169436 | 8 |
| chr8-83498603-83499104 | 6 |
| chr10-85438170-85438671 | 8 |
| chr7-86939610-86940111 | 5 |
| chrX-21321271-21321772 | 1 |
| chr3-149473165-149473666 | 6 |
| chr17-37849622-37850123 | 6 |
| chr17-31974455-31974956 | 5 |
| chr14-21751213-21751714 | 5 |
| chr16-78231746-78232247 | 6 |
| chr10-4965228-4965729 | 8 |
| chr10-16640186-16640687 | 8 |
| chrX-2429755-2430256 | 1 |
| chr6-91184582-91185083 | 8 |

| | |
|---|---|
| chr19-18312558-18313059 | 5 |
| chr1-219882624-219883125 | 8 |
| chr15-16680479-16680980 | 6 |
| chr2-135752327-135752828 | 8 |
| chr14-84402159-84402660 | 6 |
| chr14-40860522-40861023 | 5 |
| chr17-53164717-53165218 | 8 |
| chr14-74636867-74637368 | 5 |
| chr6-133714506-133715007 | 6 |
| chr14-86949405-86949906 | 1 |
| chr2-113186835-113187336 | 8 |
| chr17-36087631-36088132 | 8 |
| chr17-63444685-63445186 | 5 |
| chr10-102248696-102249197 | 5 |
| chr10-91759345-91759846 | 5 |
| chr8-62757896-62758397 | 8 |
| chr20-8334389-8334890 | 1 |
| chr3-163390015-163390516 | 5 |
| chr14-33933215-33933716 | 5 |
| chr2-124253858-124254359 | 8 |
| chr4-171215670-171216171 | 5 |
| chr8-111938066-111938567 | 8 |
| chr10-66800938-66801439 | 8 |
| chr2-121690131-121690632 | 5 |
| chr3-69153705-69154206 | 1 |
| chr7-40178208-40178709 | 6 |
| chr1-187938412-187938913 | 8 |
| chr10-42113931-42114432 | 8 |
| chr7-20119714-20120215 | 8 |
| chr3-55143946-55144447 | 6 |
| chr10-73434967-73435468 | 8 |
| chrX-35506921-35507422 | 1 |
| chr5-20526322-20526823 | 5 |
| chr13-103161577-103162078 | 8 |
| chr7-52204176-52204677 | 8 |
| chr10-38559257-38559758 | 6 |
| chr7-27506567-27507068 | 5 |
| chrX-32564370-32564871 | 8 |
| chr16-26725579-26726080 | 6 |
| chr11-35534178-35534679 | 5 |
| chr5-19866442-19866943 | 6 |
| chr15-37273834-37274335 | 8 |
| chr17-18992720-18993221 | 6 |

| | |
|---|---|
| chr5-16475696-16476197 | 8 |
| chr3-79799418-79799919 | 8 |
| chr10-31381554-31382055 | 8 |
| chr19-6501239-6501740 | 6 |
| chr16-73550945-73551446 | 8 |
| chr12-5200553-5201054 | 6 |
| chr13-75026882-75027383 | 5 |
| chr4-9659267-9659768 | 5 |
| chr2-17838640-17839141 | 8 |
| chr17-81045436-81045937 | 8 |
| chr10-64073794-64074295 | 5 |
| chr10-16962309-16962810 | 5 |
| chr1-3040799-3041300 | 8 |
| chr2-136094133-136094634 | 6 |
| chr17-61490431-61490932 | 8 |
| chr7-84358508-84359009 | 1 |
| chr6-58381123-58381624 | 6 |
| chr13-84105428-84105929 | 8 |
| chr19-19979166-19979667 | 6 |
| chr8-95914932-95915433 | 8 |
| chr15-2498077-2498578 | 1 |
| chr10-66800417-66800918 | 8 |
| chr4-126187057-126187558 | 8 |
| chr18-29904785-29905286 | 8 |
| chr9-91528769-91529270 | 1 |
| chr1-85893230-85893731 | 5 |
| chr16-31992557-31993058 | 5 |
| chr18-7173034-7173535 | 5 |
| chr1-133553709-133554210 | 8 |
| chr10-4441294-4441795 | 8 |
| chr18-55461865-55462366 | 8 |
| chr5-147610017-147610518 | 8 |
| chr1-228979916-228980417 | 5 |
| chr1-82482873-82483374 | 8 |
| chr3-116472822-116473323 | 6 |
| chr4-50384076-50384577 | 1 |
| chr14-92845567-92846068 | 6 |
| chr4-119310569-119311070 | 8 |
| chr18-31037535-31038036 | 1 |
| chr14-76261365-76261866 | 5 |
| chr6-69339374-69339875 | 8 |
| chr8-113551904-113552405 | 6 |
| chr10-27564987-27565488 | 5 |

| | |
|---|---|
| chr8-65881473-65881974 | 6 |
| chr14-9163557-9164058 | 8 |
| chr4-126329803-126330304 | 8 |
| chr2-187974790-187975291 | 5 |
| chr4-9321707-9322208 | 6 |
| chr3-158453157-158453658 | 8 |
| chr13-64306486-64306987 | 5 |
| chr20-20887722-20888223 | 8 |
| chr2-172194201-172194702 | 8 |
| chr12-42541481-42541982 | 1 |
| chr3-88573397-88573898 | 1 |
| chr4-104938792-104939293 | 8 |
| chr6-51780207-51780708 | 8 |
| chr11-62331530-62332031 | 5 |
| chr17-22390578-22391079 | 5 |
| chr1-188788253-188788754 | 8 |
| chr8-2836214-2836715 | 6 |
| chr15-50566339-50566840 | 8 |
| chrX-117567877-117568378 | 6 |
| chr15-48397163-48397664 | 8 |
| chr5-110495029-110495530 | 6 |
| chr2-100619753-100620254 | 6 |
| chr1-154484868-154485369 | 8 |
| chr11-18190533-18191034 | 5 |
| chr14-77125297-77125798 | 8 |
| chr3-149045969-149046470 | 6 |
| chr3-22362858-22363359 | 8 |
| chr14-8664377-8664878 | 5 |
| chr17-10853927-10854428 | 5 |
| chr1-256590614-256591115 | 8 |
| chr8-70239462-70239963 | 8 |
| chr6-31101816-31102317 | 8 |
| chr3-107223157-107223658 | 5 |
| chr18-72151292-72151793 | 8 |
| chr7-125643620-125644121 | 8 |
| chr11-74698844-74699345 | 8 |
| chr2-21592185-21592686 | 1 |
| chr12-7290442-7290943 | 5 |
| chr2-29000700-29001201 | 8 |
| chr10-96822537-96823038 | 8 |
| chr2-25011381-25011882 | 8 |
| chr14-98410579-98411080 | 8 |
| chr16-51858325-51858826 | 1 |

| | |
|---|---|
| chr1-254481528-254482029 | 8 |
| chr2-120040593-120041094 | 8 |
| chr2-35706633-35707134 | 5 |
| chr9-86329312-86329813 | 8 |
| chr1-123783865-123784366 | 6 |
| chrX-97162822-97163323 | 8 |
| chr5-165998129-165998630 | 6 |
| chr7-31054195-31054696 | 1 |
| chr7-77113308-77113809 | 5 |
| chr2-78899406-78899907 | 6 |
| chr4-176203390-176203891 | 8 |
| chr4-114719930-114720431 | 8 |
| chr3-21169889-21170390 | 1 |
| chr15-93056823-93057324 | 6 |
| chr7-27653568-27654069 | 1 |
| chr2-193289140-193289641 | 1 |
| chr3-160532783-160533284 | 6 |
| chr16-73416308-73416809 | 6 |
| chr19-49587713-49588214 | 6 |
| chr13-93518838-93519339 | 8 |
| chr10-22734984-22735485 | 8 |
| chr2-38183328-38183829 | 6 |
| chr20-43001099-43001600 | 8 |
| chr16-51344681-51345182 | 8 |
| chr7-28250956-28251457 | 8 |
| chr18-14957704-14958205 | 6 |
| chr3-125518464-125518965 | 5 |
| chr9-90291265-90291766 | 8 |
| chr1-20312410-20312911 | 8 |
| chr6-106618420-106618921 | 8 |
| chr14-52279442-52279943 | 6 |
| chr7-82574612-82575113 | 8 |
| chr1-219165554-219166055 | 1 |
| chr5-155378539-155379040 | 8 |
| chr6-134607732-134608233 | 6 |
| chr8-89400088-89400589 | 8 |
| chr1-239346424-239346925 | 8 |
| chr12-36806174-36806675 | 8 |
| chr3-119946265-119946766 | 5 |
| chr4-148282748-148283249 | 1 |
| chr2-195595087-195595588 | 8 |
| chr3-90247011-90247512 | 8 |
| chr9-46697081-46697582 | 1 |

| | |
|---|---|
| chr8-37125139-37125640 | 5 |
| chr5-115743640-115744141 | 8 |
| chr3-107104271-107104772 | 8 |
| chr1-12564553-12565054 | 5 |
| chr13-74046923-74047424 | 8 |
| chr1-235088558-235089059 | 1 |
| chr6-35595934-35596435 | 1 |
| chr18-55225449-55225950 | 5 |
| chr7-35266430-35266931 | 6 |
| chr13-61448367-61448868 | 6 |
| chr18-27130290-27130791 | 8 |
| chr18-50324460-50324961 | 1 |
| chr6-121185320-121185821 | 5 |
| chr15-54211111-54211612 | 8 |
| chr4-171074956-171075457 | 6 |
| chr2-145588797-145589298 | 6 |
| chr7-21913084-21913585 | 6 |
| chr14-71192842-71193343 | 1 |
| chr10-18181688-18182189 | 1 |
| chr3-152138024-152138525 | 8 |
| chr20-8484886-8485387 | 8 |
| chr3-65152189-65152690 | 5 |
| chr3-15505033-15505534 | 1 |
| chr1-2129472-2129973 | 8 |
| chr7-107156187-107156688 | 5 |
| chr2-246726324-246726825 | 6 |
| chr7-114457690-114458191 | 1 |
| chr10-98608478-98608979 | 8 |
| chr1-99260793-99261294 | 8 |
| chr13-104844891-104845392 | 8 |
| chr3-85487933-85488434 | 6 |
| chr16-125212-125713 | 8 |
| chr12-11505226-11505727 | 8 |
| chr7-100328537-100329038 | 8 |
| chr1-133271375-133271876 | 8 |
| chr16-66191780-66192281 | 8 |
| chr5-97017619-97018120 | 1 |
| chr17-26851581-26852082 | 8 |
| chr3-163759342-163759843 | 8 |
| chr7-133728644-133729145 | 8 |
| chr6-51618480-51618981 | 8 |
| chr5-114051639-114052140 | 8 |
| chr1-155640444-155640945 | 5 |

| | |
|---|---|
| chr13-21960034-21960535 | 8 |
| chr14-3773611-3774112 | 6 |
| chr4-128405860-128406361 | 5 |
| chr8-59636462-59636963 | 8 |
| chr1-194762960-194763461 | 5 |
| chr1-187815545-187816046 | 8 |
| chr2-214069770-214070271 | 5 |
| chr20-19059804-19060305 | 5 |
| chr14-33164302-33164803 | 8 |
| chr7-40998361-40998862 | 8 |
| chr9-74395331-74395832 | 8 |
| chr1-20515970-20516471 | 6 |
| chrX-50024372-50024873 | 6 |
| chr19-14686336-14686837 | 6 |
| chr6-26607235-26607736 | 8 |
| chr5-143481314-143481815 | 5 |
| chr4-30632715-30633216 | 1 |
| chr8-77731197-77731698 | 8 |
| chr9-63719750-63720251 | 6 |
| chr3-126412619-126413120 | 8 |
| chr8-11748217-11748718 | 5 |
| chr5-113761043-113761544 | 8 |
| chr8-65616875-65617376 | 8 |
| chr19-23853825-23854326 | 5 |
| chr16-74784906-74785407 | 6 |
| chr18-40980028-40980529 | 6 |
| chr13-30996586-30997087 | 6 |
| chr7-28848284-28848785 | 8 |
| chr13-45431202-45431703 | 1 |
| chr2-136778391-136778892 | 1 |
| chr1-91206076-91206577 | 8 |
| chr7-110851805-110852306 | 8 |
| chr7-100509738-100510239 | 5 |
| chr15-54361615-54362116 | 8 |
| chr13-93628294-93628795 | 8 |
| chr14-58482602-58483103 | 1 |
| chr14-32945135-32945636 | 8 |
| chr7-27417154-27417655 | 8 |
| chr5-104305473-104305974 | 8 |
| chr3-12926197-12926698 | 8 |
| chr2-7293635-7294136 | 5 |
| chr9-88584315-88584816 | 8 |
| chr12-15708664-15709165 | 5 |

| | |
|---|---|
| chr14-104191646-104192147 | 6 |
| chr17-64309215-64309716 | 6 |
| chr4-33541776-33542277 | 5 |
| chr6-35313278-35313779 | 8 |
| chr12-40327509-40328010 | 8 |
| chr8-69078567-69079068 | 6 |
| chr8-21899627-21900128 | 8 |
| chr6-16364844-16365345 | 8 |
| chr15-56603206-56603707 | 5 |
| chr2-139113259-139113760 | 8 |
| chr1-255938501-255939002 | 5 |
| chr7-107587386-107587887 | 8 |
| chr6-120887190-120887691 | 6 |
| chr2-49090090-49090591 | 6 |
| chr8-122068068-122068569 | 8 |
| chr6-98460645-98461146 | 8 |
| chr17-31692959-31693460 | 5 |
| chr1-185101443-185101944 | 8 |
| chr14-79404477-79404978 | 6 |
| chr11-74757285-74757786 | 6 |
| chr8-66144742-66145243 | 8 |
| chr19-21408248-21408749 | 6 |
| chr15-87735125-87735626 | 6 |
| chr1-114536499-114537000 | 6 |
| chr13-49852742-49853243 | 1 |
| chr19-30091378-30091879 | 8 |
| chr2-92238959-92239460 | 5 |
| chr18-71389867-71390368 | 6 |
| chr19-27550409-27550910 | 8 |
| chr2-241116884-241117385 | 8 |
| chr4-10128307-10128808 | 8 |
| chr2-113330280-113330781 | 1 |
| chr3-133047363-133047864 | 5 |
| chr7-111190551-111191052 | 6 |
| chr13-44915808-44916309 | 5 |
| chr2-113376184-113376685 | 1 |
| chr15-78500944-78501445 | 6 |
| chr17-46587338-46587839 | 8 |
| chr15-37008693-37009194 | 8 |
| chr5-150442257-150442758 | 8 |
| chr17-48264141-48264642 | 8 |
| chr6-104469399-104469900 | 1 |
| chr10-67685883-67686384 | 8 |

| | |
|---|---|
| chr12-35273644-35274145 | 8 |
| chr1-239143204-239143705 | 1 |
| chr8-95731040-95731541 | 8 |
| chr14-4093967-4094468 | 8 |
| chr11-80079870-80080371 | 8 |
| chr6-26257897-26258398 | 8 |
| chr16-44861672-44862173 | 1 |
| chr4-60215403-60215904 | 5 |
| chr10-35816823-35817324 | 5 |
| chr20-19373800-19374301 | 5 |
| chr1-5834643-5835144 | 8 |
| chr9-37300057-37300558 | 6 |
| chr7-92948071-92948572 | 1 |
| chr15-36248728-36249229 | 8 |
| chr18-65363324-65363825 | 1 |
| chr1-190827774-190828275 | 8 |
| chr13-48377700-48378201 | 1 |
| chr5-111707925-111708426 | 8 |
| chr17-35109636-35110137 | 1 |
| chr7-117445260-117445761 | 6 |
| chr5-126201307-126201808 | 5 |
| chr7-5600850-5601351 | 6 |
| chr2-124983502-124984003 | 8 |
| chr15-98337528-98338029 | 8 |
| chr1-142325753-142326254 | 8 |
| chr9-106752761-106753262 | 5 |
| chr8-26531455-26531956 | 8 |
| chr10-68305037-68305538 | 8 |
| chr11-68096410-68096911 | 5 |
| chr15-90127809-90128310 | 8 |
| chr8-29474342-29474843 | 5 |
| chr7-108266457-108266958 | 5 |
| chr1-41394638-41395139 | 8 |
| chr9-69013978-69014479 | 1 |
| chr1-126153272-126153773 | 6 |
| chr19-5219098-5219599 | 8 |
| chr16-44877460-44877961 | 6 |
| chr13-98675859-98676360 | 5 |
| chr8-73025609-73026110 | 8 |
| chr1-1438836-1439337 | 8 |
| chr3-150325791-150326292 | 1 |
| chr19-31485963-31486464 | 1 |
| chr1-120292302-120292803 | 6 |

| | |
|---|---|
| chr11-50209155-50209656 | 5 |
| chr3-159640240-159640741 | 6 |
| chr3-157703798-157704299 | 6 |
| chr19-53516149-53516650 | 5 |
| chr6-97822393-97822894 | 6 |
| chr4-67103670-67104171 | 8 |
| chr1-225378390-225378891 | 6 |
| chr1-155024893-155025394 | 8 |
| chr3-99381021-99381522 | 8 |
| chr2-124597324-124597825 | 8 |
| chr16-68863457-68863958 | 1 |
| chr7-20465142-20465643 | 8 |
| chr3-17295978-17296479 | 5 |
| chr9-86203918-86204419 | 1 |
| chr19-46844625-46845126 | 1 |
| chr7-124144004-124144505 | 8 |
| chr10-91169309-91169810 | 8 |
| chr5-30486926-30487427 | 5 |
| chr3-147432961-147433462 | 5 |
| chr2-39353113-39353614 | 6 |
| chr8-92116479-92116980 | 8 |
| chr20-36150323-36150824 | 5 |
| chr2-179133575-179134076 | 5 |
| chr17-19875216-19875717 | 5 |
| chr10-18510901-18511402 | 1 |
| chr1-196766204-196766705 | 8 |
| chr13-77689068-77689569 | 8 |
| chr13-46554441-46554942 | 5 |
| chr17-73806561-73807062 | 1 |
| chr15-76378980-76379481 | 8 |
| chr3-79392260-79392761 | 1 |
| chr6-35893441-35893942 | 8 |
| chr6-45898631-45899132 | 8 |
| chr7-95165973-95166474 | 8 |
| chr13-75635118-75635619 | 8 |
| chr5-76605672-76606173 | 8 |
| chr17-80656512-80657013 | 6 |
| chr19-46680080-46680581 | 6 |
| chr17-73159402-73159903 | 5 |
| chr4-116873279-116873780 | 8 |
| chr6-30691656-30692157 | 1 |
| chr11-70118751-70119252 | 1 |
| chr13-103116065-103116566 | 8 |

| | |
|---|---|
| chr10-68025008-68025509 | 6 |
| chr4-121821272-121821773 | 5 |
| chr1-30310222-30310723 | 8 |
| chr3-158444908-158445409 | 1 |
| chr15-61947053-61947554 | 1 |
| chr5-6054047-6054548 | 5 |
| chr13-106612463-106612964 | 8 |
| chr1-177150960-177151461 | 8 |
| chr6-27199843-27200344 | 8 |
| chr7-124777715-124778216 | 6 |
| chr8-19866796-19867297 | 8 |
| chr12-8045860-8046361 | 5 |
| chr1-152223238-152223739 | 8 |
| chr1-235272775-235273276 | 5 |
| chr14-43668489-43668990 | 8 |
| chr6-3533327-3533828 | 8 |
| chr10-50560986-50561487 | 5 |
| chr1-202948506-202949007 | 5 |
| chr3-152641183-152641684 | 1 |
| chr5-150601645-150602146 | 5 |
| chr18-67160901-67161402 | 8 |
| chr5-67642893-67643394 | 6 |
| chr12-12540081-12540582 | 6 |
| chr1-206715787-206716288 | 1 |
| chr15-15596389-15596890 | 8 |
| chr17-15582445-15582946 | 8 |
| chr9-3447059-3447560 | 5 |
| chr5-150881982-150882483 | 6 |
| chr9-86187446-86187947 | 8 |
| chr6-26757166-26757667 | 1 |
| chr12-10215119-10215620 | 5 |
| chr19-12217629-12218130 | 8 |
| chr3-55080183-55080684 | 5 |
| chr5-43303824-43304325 | 6 |
| chr4-68026821-68027322 | 8 |
| chr1-102076176-102076677 | 5 |
| chr10-65717669-65718170 | 1 |
| chr7-89173296-89173797 | 8 |
| chrX-135120996-135121497 | 6 |
| chr8-114823560-114824061 | 8 |
| chr18-600401-600902 | 5 |
| chr3-22938832-22939333 | 8 |
| chr13-90178602-90179103 | 5 |

| | |
|---|---|
| chr10-55324664-55325165 | 8 |
| chr10-46209533-46210034 | 5 |
| chr9-89058717-89059218 | 8 |
| chr8-90495273-90495774 | 8 |
| chr15-16309499-16310000 | 8 |
| chr17-53676745-53677246 | 8 |
| chr10-96312277-96312778 | 8 |
| chr5-67655704-67656205 | 1 |
| chr18-77310615-77311116 | 5 |
| chr15-68588784-68589285 | 6 |
| chr2-235067038-235067539 | 5 |
| chr3-79623750-79624251 | 6 |
| chr7-128820972-128821473 | 1 |
| chrX-15903869-15904370 | 1 |
| chr2-180059912-180060413 | 1 |
| chr19-55852337-55852838 | 5 |
| chr19-24066528-24067029 | 8 |
| chr12-36854103-36854604 | 8 |
| chr16-25817472-25817973 | 6 |
| chr1-177855686-177856187 | 5 |
| chr8-23342236-23342737 | 6 |
| chr13-64635279-64635780 | 1 |
| chr20-35259314-35259815 | 6 |
| chr15-1318400-1318901 | 1 |
| chr13-76467234-76467735 | 6 |
| chr3-103110633-103111134 | 8 |
| chr8-90745304-90745805 | 1 |
| chrX-13100820-13101321 | 6 |
| chr9-14006959-14007460 | 8 |
| chr18-37191991-37192492 | 5 |
| chr10-70834357-70834858 | 1 |
| chr19-23804213-23804714 | 8 |
| chr11-20924375-20924876 | 6 |
| chr1-190154798-190155299 | 1 |
| chr6-13055198-13055699 | 8 |
| chr19-15401645-15402146 | 1 |
| chr1-236538642-236539143 | 6 |
| chr15-20094874-20095375 | 8 |
| chr1-32877101-32877602 | 8 |
| chr2-25304433-25304934 | 1 |
| chr4-73096323-73096824 | 8 |
| chr11-32641533-32642034 | 8 |
| chrX-12998005-12998506 | 5 |

| | |
|---|---|
| chr2-193085818-193086319 | 5 |
| chr7-112944894-112945395 | 8 |
| chr14-82675019-82675520 | 6 |
| chr6-13214733-13215234 | 8 |
| chr18-58646095-58646596 | 1 |
| chr3-119717382-119717883 | 6 |
| chr3-107442104-107442605 | 1 |
| chr20-39655573-39656074 | 6 |
| chr8-6359290-6359791 | 6 |
| chr8-100322369-100322870 | 8 |
| chr20-8434917-8435418 | 5 |
| chr20-42973830-42974331 | 8 |
| chr7-22535748-22536249 | 5 |
| chr8-55482510-55483011 | 5 |
| chr11-66212380-66212881 | 6 |
| chr16-45395179-45395680 | 6 |
| chrX-111590607-111591108 | 8 |
| chr3-19261159-19261660 | 6 |
| chr5-150760359-150760860 | 8 |
| chr1-167916144-167916645 | 5 |
| chr14-81800548-81801049 | 6 |
| chr11-25841252-25841753 | 1 |
| chr14-44571901-44572402 | 1 |
| chr15-18656531-18657032 | 8 |
| chr9-103819446-103819947 | 8 |
| chr14-5791898-5792399 | 8 |
| chr9-110705193-110705694 | 8 |
| chr20-16255868-16256369 | 8 |
| chr9-85495784-85496285 | 8 |
| chrX-127276767-127277268 | 5 |
| chr15-75154550-75155051 | 8 |
| chr5-24379438-24379939 | 6 |
| chr5-70370050-70370551 | 8 |
| chr5-56495895-56496396 | 5 |
| chr18-14671395-14671896 | 8 |
| chr1-91360083-91360584 | 8 |
| chr18-24175779-24176280 | 5 |
| chr13-16793159-16793660 | 8 |
| chr3-26032528-26033029 | 6 |
| chr1-26291956-26292457 | 5 |
| chr6-100730722-100731223 | 5 |
| chr9-36444446-36444947 | 5 |
| chr7-106482554-106483055 | 6 |

| | |
|---|---|
| chr9-35405247-35405748 | 8 |
| chr3-153241884-153242385 | 1 |
| chr1-192013875-192014376 | 1 |
| chr3-167030609-167031110 | 8 |
| chr11-62586369-62586870 | 8 |
| chr5-73108558-73109059 | 6 |
| chr2-140253722-140254223 | 5 |
| chr9-105487232-105487733 | 8 |
| chr18-71519747-71520248 | 1 |
| chr6-56485494-56485995 | 6 |
| chrX-116522941-116523442 | 5 |
| chr1-51291520-51292021 | 1 |
| chr5-133490670-133491171 | 8 |
| chr14-94519958-94520459 | 8 |
| chr19-20588863-20589364 | 6 |
| chr7-20563453-20563954 | 6 |
| chr8-48883207-48883708 | 8 |
| chr4-30266954-30267455 | 8 |
| chr6-27569367-27569868 | 5 |
| chr20-13525480-13525981 | 1 |
| chr17-35702055-35702556 | 1 |
| chr5-61607888-61608389 | 1 |
| chr3-167043102-167043603 | 8 |
| chr17-46172973-46173474 | 8 |
| chr9-82200053-82200554 | 6 |
| chr16-80672980-80673481 | 6 |
| chr1-81334208-81334709 | 6 |
| chr14-15682941-15683442 | 8 |
| chr1-46234407-46234908 | 8 |
| chr7-40253005-40253506 | 6 |
| chr16-6741524-6742025 | 8 |
| chr10-88940437-88940938 | 1 |
| chr15-80557178-80557679 | 5 |
| chr8-44477002-44477503 | 5 |
| chr13-99299686-99300187 | 5 |
| chr1-238243000-238243501 | 5 |
| chr15-4005371-4005872 | 5 |
| chr14-96863919-96864420 | 5 |
| chr17-21363121-21363622 | 8 |
| chr4-122833567-122834068 | 8 |
| chr6-99000877-99001378 | 1 |
| chr2-118640615-118641116 | 5 |
| chrX-111502994-111503495 | 8 |

| | |
|---|---|
| chr8-40589623-40590124 | 6 |
| chr2-51398761-51399262 | 5 |
| chr3-111691317-111691818 | 8 |
| chr7-117110051-117110552 | 6 |
| chrX-108703638-108704139 | 6 |
| chr16-68953630-68954131 | 1 |
| chr5-124575622-124576123 | 6 |
| chr10-73241629-73242130 | 5 |
| chr8-109060042-109060543 | 6 |
| chr14-74412056-74412557 | 8 |
| chr3-13912005-13912506 | 8 |
| chr7-27645135-27645636 | 5 |
| chr15-36692627-36693128 | 5 |
| chr18-76198674-76199175 | 8 |
| chr13-97048458-97048959 | 8 |
| chr7-97537592-97538093 | 8 |
| chr8-61195491-61195992 | 8 |
| chr18-75120686-75121187 | 8 |
| chr19-16496597-16497098 | 1 |
| chr2-120430273-120430774 | 8 |
| chr8-46632428-46632929 | 6 |
| chr1-196518595-196519096 | 8 |
| chr7-128425793-128426294 | 1 |
| chr14-3737046-3737547 | 8 |
| chr4-119512172-119512673 | 5 |
| chr2-172309123-172309624 | 8 |
| chr8-114280734-114281235 | 8 |
| chr2-219420584-219421085 | 8 |
| chr4-124546395-124546896 | 1 |
| chr10-92059325-92059826 | 8 |
| chr4-179393070-179393571 | 1 |
| chr17-22999422-22999923 | 8 |
| chr14-79401585-79402086 | 5 |
| chr18-58749114-58749615 | 8 |
| chr13-79207240-79207741 | 8 |
| chr8-55099627-55100128 | 6 |
| chr1-137823030-137823531 | 8 |
| chr5-146895101-146895602 | 5 |
| chr15-52702859-52703360 | 8 |
| chr3-71613939-71614440 | 8 |
| chr7-111794865-111795366 | 8 |
| chr5-6761389-6761890 | 8 |
| chr4-118426195-118426696 | 8 |

| | |
|---|---|
| chr10-87753972-87754473 | 6 |
| chr2-142146879-142147380 | 8 |
| chr6-119788342-119788843 | 1 |
| chr1-234990626-234991127 | 8 |
| chr2-114204770-114205271 | 8 |
| chr16-42161923-42162424 | 6 |
| chr17-33500946-33501447 | 8 |
| chr15-4397259-4397760 | 8 |
| chr7-124898825-124899326 | 5 |
| chr18-40158315-40158816 | 8 |
| chr11-3682400-3682901 | 5 |
| chr14-32351965-32352466 | 8 |
| chr4-60241201-60241702 | 1 |
| chr7-100173273-100173774 | 1 |
| chr5-6444667-6445168 | 1 |
| chr3-44740730-44741231 | 6 |
| chr16-26055689-26056190 | 8 |
| chr7-78647379-78647880 | 1 |
| chr3-124995220-124995721 | 1 |
| chr4-37133783-37134284 | 8 |
| chr9-90695161-90695662 | 8 |
| chr13-89259086-89259587 | 5 |
| chr3-93249182-93249683 | 8 |
| chr15-35318155-35318656 | 8 |
| chr13-32127693-32128194 | 5 |
| chr4-145518993-145519494 | 5 |
| chr4-62388881-62389382 | 6 |
| chr10-71376660-71377161 | 1 |
| chr5-56797448-56797949 | 6 |
| chr8-90209301-90209802 | 8 |
| chr1-27396999-27397500 | 6 |
| chr13-84664701-84665202 | 5 |
| chr4-112388234-112388735 | 6 |
| chr6-87591210-87591711 | 6 |
| chr19-35648984-35649485 | 8 |
| chr13-22889345-22889846 | 1 |
| chr2-177579162-177579663 | 6 |
| chr14-76362908-76363409 | 8 |
| chr2-43056636-43057137 | 8 |
| chr3-33364071-33364572 | 8 |
| chr10-38478084-38478585 | 6 |
| chr13-103470222-103470723 | 5 |
| chr14-97361694-97362195 | 8 |

| | |
|---|---|
| chr13-87986225-87986726 | 8 |
| chr7-113582174-113582675 | 8 |
| chr5-123428795-123429296 | 8 |
| chr10-86964094-86964595 | 8 |
| chr16-18191166-18191667 | 1 |
| chr15-37247571-37248072 | 8 |
| chr2-183034603-183035104 | 6 |
| chr18-23081371-23081872 | 8 |
| chr19-36376923-36377424 | 8 |
| chr19-37364594-37365095 | 5 |
| chr17-52101475-52101976 | 8 |
| chr8-41563539-41564040 | 8 |
| chr12-31690426-31690927 | 1 |
| chr17-38306386-38306887 | 8 |
| chr10-95940828-95941329 | 1 |
| chr6-7782342-7782843 | 6 |
| chr2-42168477-42168978 | 8 |
| chr6-97301578-97302079 | 6 |
| chr1-179462293-179462794 | 5 |
| chr17-66402764-66403265 | 1 |
| chr1-242957356-242957857 | 6 |
| chr6-52884527-52885028 | 8 |
| chr4-158069203-158069704 | 8 |
| chr9-110969019-110969520 | 8 |
| chr9-65583798-65584299 | 8 |
| chr2-26187796-26188297 | 5 |
| chr2-23924712-23925213 | 1 |
| chr19-10643597-10644098 | 8 |
| chr3-69042055-69042556 | 8 |
| chr1-132883325-132883826 | 5 |
| chr13-40414168-40414669 | 8 |
| chr3-93280530-93281031 | 5 |
| chr2-189208177-189208678 | 1 |
| chr20-3473518-3474019 | 8 |
| chr4-133128086-133128587 | 6 |
| chr7-29585345-29585846 | 5 |
| chr1-192587272-192587773 | 1 |
| chr18-32180509-32181010 | 8 |
| chr14-71184667-71185168 | 6 |
| chr1-137274704-137275205 | 5 |
| chr1-44828631-44829132 | 8 |
| chr8-45804445-45804946 | 6 |
| chr5-104050832-104051333 | 8 |

| | |
|---|---|
| chr5-20552251-20552752 | 1 |
| chr16-56813664-56814165 | 8 |
| chr3-70429236-70429737 | 1 |
| chr11-65298881-65299382 | 8 |
| chr15-34298075-34298576 | 6 |
| chr12-34120557-34121058 | 8 |
| chr9-99061563-99062064 | 8 |
| chr17-16665360-16665861 | 6 |
| chr11-68605079-68605580 | 1 |
| chr6-36529826-36530327 | 6 |
| chr4-57518303-57518804 | 8 |
| chr20-41897455-41897956 | 5 |
| chr13-46112307-46112808 | 8 |
| chr1-84975487-84975988 | 6 |
| chr15-3607957-3608458 | 5 |
| chr3-113847244-113847745 | 8 |
| chr7-86679025-86679526 | 8 |
| chr8-64306764-64307265 | 8 |
| chr5-74377178-74377679 | 8 |
| chr8-45009387-45009888 | 1 |
| chr18-27384637-27385138 | 8 |
| chrX-12540974-12541475 | 1 |
| chr12-22007595-22008096 | 8 |
| chr6-21601555-21602056 | 1 |
| chr11-34543609-34544110 | 6 |
| chr1-234511227-234511728 | 8 |
| chr9-81930714-81931215 | 8 |
| chr16-75352138-75352639 | 1 |
| chr17-23818105-23818606 | 8 |
| chr2-110884771-110885272 | 1 |
| chr3-30379526-30380027 | 8 |
| chr4-62214390-62214891 | 5 |
| chr4-146887932-146888433 | 8 |
| chr12-4677762-4678263 | 8 |
| chr4-59622710-59623211 | 6 |
| chr9-15946838-15947339 | 6 |
| chr4-173741288-173741789 | 6 |
| chr15-2013757-2014258 | 8 |
| chr15-4421522-4422023 | 5 |
| chr16-34837470-34837971 | 8 |
| chr1-93831339-93831840 | 8 |
| chr2-24220058-24220559 | 8 |
| chr1-22514156-22514657 | 8 |

| | |
|---|---|
| chr4-58325895-58326396 | 8 |
| chr8-66744174-66744675 | 1 |
| chr17-22877487-22877988 | 1 |
| chr7-44737979-44738480 | 5 |
| chr10-15046372-15046873 | 8 |
| chr2-242745704-242746205 | 5 |
| chr4-52289806-52290307 | 6 |
| chr2-203725346-203725847 | 5 |
| chr3-19324303-19324804 | 8 |
| chr20-3988134-3988635 | 5 |
| chr7-9783118-9783619 | 5 |
| chr13-97398257-97398758 | 8 |
| chr5-29448085-29448586 | 5 |
| chr18-54119586-54120087 | 1 |
| chr2-191947920-191948421 | 5 |
| chr4-125076422-125076923 | 8 |
| chr4-96248854-96249355 | 1 |
| chr7-86390796-86391297 | 5 |
| chr15-47843517-47844018 | 8 |
| chr8-11287639-11288140 | 8 |
| chr14-71119230-71119731 | 6 |
| chr8-73554135-73554636 | 5 |
| chr8-54123797-54124298 | 5 |
| chr17-23162383-23162884 | 1 |
| chr5-48468468-48468969 | 5 |
| chr15-46774956-46775457 | 6 |
| chr7-40496709-40497210 | 8 |
| chrX-27234845-27235346 | 6 |
| chr17-75051396-75051897 | 5 |
| chrX-15962845-15963346 | 8 |
| chr18-41499630-41500131 | 8 |
| chr13-91773159-91773660 | 5 |
| chr12-44349820-44350321 | 8 |
| chr20-92680-93181 | 8 |
| chr10-78770794-78771295 | 6 |
| chr5-115788992-115789493 | 8 |
| chr1-187843573-187844074 | 5 |
| chr2-39396643-39397144 | 1 |
| chr9-15726291-15726792 | 8 |
| chr9-86492931-86493432 | 8 |
| chr3-19192421-19192922 | 8 |
| chr3-28646643-28647144 | 6 |
| chr4-162604800-162605301 | 6 |

| | |
|---|---|
| chr20-2848352-2848853 | 8 |
| chr2-12807383-12807884 | 8 |
| chr7-95758509-95759010 | 5 |
| chr6-88701736-88702237 | 8 |
| chr19-47463116-47463617 | 8 |
| chr18-61662641-61663142 | 8 |
| chr6-101775268-101775769 | 8 |
| chr9-71022369-71022870 | 8 |
| chr3-88817096-88817597 | 5 |
| chr8-31044083-31044584 | 8 |
| chr7-127282123-127282624 | 8 |
| chr1-120844206-120844707 | 8 |
| chr17-74737768-74738269 | 5 |
| chr1-242960429-242960930 | 6 |
| chr5-22477482-22477983 | 1 |
| chr15-8621640-8622141 | 6 |
| chr17-55963619-55964120 | 8 |
| chr5-8288095-8288596 | 1 |
| chr12-11218726-11219227 | 8 |
| chr18-4334607-4335108 | 6 |
| chr5-58763301-58763802 | 8 |
| chr1-83944338-83944839 | 1 |
| chr5-164465104-164465605 | 6 |
| chr1-17949593-17950094 | 8 |
| chrX-136088852-136089353 | 5 |
| chr1-48048687-48049188 | 8 |
| chr4-11326344-11326845 | 5 |
| chr8-76937126-76937627 | 6 |
| chr2-189739286-189739787 | 8 |
| chr11-50824988-50825489 | 8 |
| chr7-90661558-90662059 | 8 |
| chr4-150682876-150683377 | 1 |
| chr1-228172261-228172762 | 5 |
| chr6-106932330-106932831 | 5 |
| chr20-5114935-5115436 | 1 |
| chr11-67005299-67005800 | 5 |
| chr8-71685158-71685659 | 1 |
| chr10-65269742-65270243 | 8 |
| chr15-39209004-39209505 | 1 |
| chr11-47720977-47721478 | 6 |
| chr11-35520537-35521038 | 8 |
| chr13-80077908-80078409 | 5 |
| chr3-56686593-56687094 | 8 |

| | |
|---|---|
| chr5-5906458-5906959 | 8 |
| chr1-207024350-207024851 | 8 |
| chr2-86980879-86981380 | 8 |
| chr8-123101646-123102147 | 1 |
| chr4-34284910-34285411 | 6 |
| chr9-50269035-50269536 | 8 |
| chr18-60924392-60924893 | 5 |
| chr6-75841180-75841681 | 6 |
| chr18-45894591-45895092 | 5 |
| chr10-31801393-31801894 | 8 |
| chr13-71957491-71957992 | 6 |
| chr8-37117938-37118439 | 1 |
| chr1-116580394-116580895 | 6 |
| chr19-16177113-16177614 | 5 |
| chr2-122813362-122813863 | 5 |
| chr6-75537012-75537513 | 8 |
| chr16-55666871-55667372 | 1 |
| chr2-41136876-41137377 | 1 |
| chr20-19237328-19237829 | 8 |
| chr6-11326394-11326895 | 8 |
| chr2-203796059-203796560 | 8 |
| chr18-58494662-58495163 | 6 |
| chr1-52638978-52639479 | 6 |
| chr4-132213752-132214253 | 6 |
| chr12-44305565-44306066 | 5 |
| chr20-21467128-21467629 | 1 |
| chr10-46053729-46054230 | 8 |
| chr1-50945393-50945894 | 1 |
| chr4-128379015-128379516 | 6 |
| chr1-222979037-222979538 | 8 |
| chr16-43480618-43481119 | 8 |
| chr10-50229506-50230007 | 8 |
| chr9-12546334-12546835 | 5 |
| chr10-72423869-72424370 | 8 |
| chr12-20662024-20662525 | 8 |
| chr6-40729721-40730222 | 8 |
| chr1-58575453-58575954 | 6 |
| chr2-84710623-84711124 | 8 |
| chr2-195973477-195973978 | 5 |
| chr4-146993894-146994395 | 6 |
| chr19-19216553-19217054 | 8 |
| chr5-26261312-26261813 | 8 |
| chr12-41582945-41583446 | 5 |

| | |
|---|---|
| chr15-31797456-31797957 | 8 |
| chr12-31305303-31305804 | 8 |
| chr13-90926351-90926852 | 8 |
| chr6-13585417-13585918 | 8 |
| chr5-3972773-3973274 | 6 |
| chr3-45402617-45403118 | 6 |
| chr13-79391149-79391650 | 1 |
| chr8-61994958-61995459 | 8 |
| chr18-54039355-54039856 | 8 |
| chr14-24575155-24575656 | 6 |
| chr19-9573120-9573621 | 8 |
| chr5-165142465-165142966 | 8 |
| chr19-34910422-34910923 | 8 |
| chr19-42455375-42455876 | 5 |
| chr16-57700444-57700945 | 8 |
| chr15-31822655-31823156 | 8 |
| chr7-53235130-53235631 | 8 |
| chr10-95352917-95353418 | 8 |
| chr20-11499274-11499775 | 1 |
| chr2-174364689-174365190 | 8 |
| chr9-91649564-91650065 | 1 |
| chr1-79670120-79670621 | 5 |
| chr1-160894028-160894529 | 5 |
| chr13-46210462-46210963 | 6 |
| chr2-136321296-136321797 | 8 |
| chr2-54201761-54202262 | 5 |
| chr8-49174953-49175454 | 8 |
| chr9-98056051-98056552 | 1 |
| chr15-76840341-76840842 | 8 |
| chr7-124782976-124783477 | 5 |
| chr16-60878890-60879391 | 8 |
| chr16-71712257-71712758 | 8 |
| chr6-76262348-76262849 | 8 |
| chr10-59994715-59995216 | 8 |
| chrX-130333022-130333523 | 8 |
| chr10-50199005-50199506 | 6 |
| chr14-33211274-33211775 | 8 |
| chr4-180503364-180503865 | 6 |
| chr12-25015015-25015516 | 8 |
| chr3-80481884-80482385 | 5 |
| chr1-18138949-18139450 | 8 |
| chr10-102101594-102102095 | 6 |
| chr7-106132150-106132651 | 8 |

| | |
|---|---|
| chr10-21662737-21663238 | 5 |
| chr1-5021569-5022070 | 6 |
| chr7-47261885-47262386 | 8 |
| chr9-98123637-98124138 | 8 |
| chr2-169711763-169712264 | 8 |
| chrX-11488295-11488796 | 6 |
| chr20-730864-731365 | 8 |
| chr9-106022601-106023102 | 6 |
| chr3-10544707-10545208 | 5 |
| chr2-204915751-204916252 | 1 |
| chr11-80871061-80871562 | 5 |
| chr13-46774305-46774806 | 8 |
| chr15-11193650-11194151 | 5 |
| chr2-8473403-8473904 | 6 |
| chr20-47295279-47295780 | 8 |
| chr2-49497195-49497696 | 5 |
| chr3-46895141-46895642 | 8 |
| chr18-34193875-34194376 | 5 |
| chr1-100536367-100536868 | 8 |
| chr15-98793793-98794294 | 8 |
| chr15-62890648-62891149 | 5 |
| chr7-119743501-119744002 | 8 |
| chr10-29506635-29507136 | 8 |
| chr2-56078737-56079238 | 1 |
| chr1-77066289-77066790 | 5 |
| chr4-7821521-7822022 | 8 |
| chr4-117960765-117961266 | 8 |
| chr9-111182283-111182784 | 8 |
| chr8-45301752-45302253 | 6 |
| chr15-26459884-26460385 | 8 |
| chr7-58161966-58162467 | 5 |
| chr8-114839748-114840249 | 6 |
| chr8-16588154-16588655 | 5 |
| chr2-46063302-46063803 | 8 |
| chr14-56747177-56747678 | 8 |
| chr3-47203302-47203803 | 5 |
| chr3-60464125-60464626 | 6 |
| chr13-74155813-74156314 | 6 |
| chr7-118239437-118239938 | 8 |
| chr10-40340920-40341421 | 1 |
| chr3-41187769-41188270 | 6 |
| chr4-93972423-93972924 | 8 |
| chr10-37315881-37316382 | 8 |

| | |
|---|---|
| chr9-92688451-92688952 | 8 |
| chr4-68942802-68943303 | 6 |
| chr19-18527071-18527572 | 5 |
| chr9-1766814-1767315 | 8 |
| chr3-106372527-106373028 | 5 |
| chr8-4877474-4877975 | 5 |
| chr3-80268902-80269403 | 5 |
| chr10-104989591-104990092 | 8 |
| chr9-88852743-88853244 | 8 |
| chr3-144597239-144597740 | 8 |
| chr1-171779757-171780258 | 6 |
| chr20-10158993-10159494 | 8 |
| chr10-13166872-13167373 | 6 |
| chr5-120197592-120198093 | 6 |
| chr8-92871971-92872472 | 8 |
| chr10-71272135-71272636 | 8 |
| chr5-25076541-25077042 | 1 |
| chr7-47063243-47063744 | 8 |
| chr5-18917639-18918140 | 8 |
| chr5-165187737-165188238 | 8 |
| chr3-132976340-132976841 | 8 |
| chr19-19301065-19301566 | 5 |
| chr4-104829542-104830043 | 8 |
| chr10-71565917-71566418 | 8 |
| chr4-13301948-13302449 | 6 |
| chr14-96692606-96693107 | 1 |
| chr13-89721038-89721539 | 6 |
| chr6-119728290-119728791 | 1 |
| chr4-167012885-167013386 | 8 |
| chr14-7195779-7196280 | 8 |
| chr5-43142946-43143447 | 8 |
| chr10-88778831-88779332 | 6 |
| chr9-43279884-43280385 | 6 |
| chr20-13004976-13005477 | 8 |
| chr6-10296299-10296800 | 6 |
| chr14-71516098-71516599 | 8 |
| chr1-174563406-174563907 | 8 |
| chr1-118933834-118934335 | 5 |
| chr9-10767012-10767513 | 8 |
| chr5-149859311-149859812 | 8 |
| chr4-83559181-83559682 | 1 |
| chr7-23495930-23496431 | 8 |
| chr9-3430000-3430501 | 8 |

| | |
|---|---|
| chr20-26755779-26756280 | 5 |
| chr8-57053612-57054113 | 6 |
| chr14-75614040-75614541 | 5 |
| chr4-142852617-142853118 | 8 |
| chr3-41250952-41251453 | 8 |
| chr14-67249129-67249630 | 8 |
| chr20-21537949-21538450 | 8 |
| chr4-179823554-179824055 | 1 |
| chr9-49476900-49477401 | 1 |
| chr9-90323274-90323775 | 6 |
| chr2-135261429-135261930 | 5 |
| chr15-53758868-53759369 | 8 |
| chr18-877193-877694 | 8 |
| chr2-103532718-103533219 | 8 |
| chr6-57881138-57881639 | 6 |
| chr15-16388586-16389087 | 8 |
| chr2-58031768-58032269 | 8 |
| chr20-35767937-35768438 | 8 |
| chr14-58222103-58222604 | 8 |
| chr6-88184121-88184622 | 8 |
| chr1-187294363-187294864 | 6 |
| chr10-4908743-4909244 | 1 |
| chr12-32204286-32204787 | 8 |
| chr8-63003007-63003508 | 5 |
| chr7-113520963-113521464 | 8 |
| chr1-16981808-16982309 | 1 |
| chr16-55726484-55726985 | 5 |
| chr16-25613877-25614378 | 6 |
| chr4-142018502-142019003 | 1 |
| chr3-128942621-128943122 | 8 |
| chr16-67733019-67733520 | 5 |
| chrX-116902637-116903138 | 8 |
| chr17-30445060-30445561 | 8 |
| chr2-115606751-115607252 | 8 |
| chr1-252004136-252004637 | 6 |
| chr20-46035618-46036119 | 5 |
| chr15-98271639-98272140 | 8 |
| chr14-67082762-67083263 | 5 |
| chr18-59918606-59919107 | 8 |
| chr11-42366265-42366766 | 6 |
| chr10-3782125-3782626 | 6 |
| chr14-32553413-32553914 | 6 |
| chr19-44614103-44614604 | 8 |

| | |
|---|---|
| chr12-27630519-27631020 | 6 |
| chr7-106661908-106662409 | 6 |
| chr1-212376075-212376576 | 8 |
| chr5-143340812-143341313 | 6 |
| chr1-29449500-29450001 | 8 |
| chr18-12318730-12319231 | 8 |
| chr4-121835253-121835754 | 1 |
| chr20-1430639-1431140 | 1 |
| chr17-48398365-48398866 | 6 |
| chr19-36618052-36618553 | 5 |
| chr13-75768765-75769266 | 8 |
| chr13-80150144-80150645 | 6 |
| chr5-20417225-20417726 | 8 |
| chr1-53890677-53891178 | 8 |
| chr4-175842121-175842622 | 5 |
| chr19-29711870-29712371 | 1 |
| chr17-5435531-5436032 | 8 |
| chr16-55900068-55900569 | 1 |
| chr13-101593094-101593595 | 6 |
| chr4-93764332-93764833 | 1 |
| chr3-60297809-60298310 | 1 |
| chr16-76680680-76681181 | 8 |
| chr9-76780512-76781013 | 8 |
| chr6-112664297-112664798 | 6 |
| chr4-10353117-10353618 | 5 |
| chr4-103890833-103891334 | 5 |
| chr2-236816085-236816586 | 6 |
| chr16-45487810-45488311 | 8 |
| chr5-46728929-46729430 | 8 |
| chr4-82887173-82887674 | 8 |
| chr1-244770066-244770567 | 5 |
| chr4-93076306-93076807 | 6 |
| chr6-137575994-137576495 | 5 |
| chr2-139933249-139933750 | 1 |
| chr9-88814741-88815242 | 6 |
| chr4-120953857-120954358 | 6 |
| chr2-19954157-19954658 | 8 |
| chr2-203554236-203554737 | 1 |
| chr4-120178389-120178890 | 6 |
| chr12-23098331-23098832 | 8 |
| chr16-45885729-45886230 | 8 |
| chr2-241447369-241447870 | 8 |
| chrX-132105002-132105503 | 5 |

| | |
|---|---|
| chr5-72856769-72857270 | 8 |
| chr13-94889040-94889541 | 1 |
| chr2-226245824-226246325 | 5 |
| chr10-45098104-45098605 | 5 |
| chr3-114863809-114864310 | 8 |
| chr8-105659555-105660056 | 8 |
| chr9-42942015-42942516 | 6 |
| chr20-27388603-27389104 | 8 |
| chr2-99377208-99377709 | 8 |
| chr8-119302547-119303048 | 6 |
| chr2-149598783-149599284 | 8 |
| chr8-45210910-45211411 | 8 |
| chr17-72348698-72349199 | 8 |
| chr9-106966899-106967400 | 5 |
| chr4-68723582-68724083 | 8 |
| chr10-20319403-20319904 | 8 |
| chr1-187748771-187749272 | 8 |
| chr4-172718541-172719042 | 8 |
| chr8-65050596-65051097 | 8 |
| chr7-9949556-9950057 | 8 |
| chr17-5644620-5645121 | 5 |
| chr14-102707997-102708498 | 8 |
| chr1-154304234-154304735 | 6 |
| chr1-127782335-127782836 | 8 |
| chr5-109681397-109681898 | 8 |
| chr6-90098465-90098966 | 6 |
| chr4-33539055-33539556 | 5 |
| chr20-28498144-28498645 | 6 |
| chr15-44314637-44315138 | 6 |
| chr7-88168706-88169207 | 8 |
| chr9-87213786-87214287 | 5 |
| chr11-72140913-72141414 | 8 |
| chr17-20494538-20495039 | 6 |
| chr10-6323990-6324491 | 6 |
| chr13-85831074-85831575 | 8 |
| chr6-43953411-43953912 | 8 |
| chr15-40856649-40857150 | 6 |
| chr6-125570468-125570969 | 5 |
| chr8-4294648-4295149 | 6 |
| chr19-45355183-45355684 | 1 |
| chr12-35437234-35437735 | 6 |
| chr17-24476796-24477297 | 8 |
| chr17-41448332-41448833 | 5 |

| | |
|---|---|
| chrX-117261297-117261798 | 6 |
| chr2-26468723-26469224 | 5 |
| chr4-104403706-104404207 | 6 |
| chr10-57879203-57879704 | 8 |
| chr13-104326144-104326645 | 8 |
| chr16-84262405-84262906 | 5 |
| chr9-73368075-73368576 | 6 |
| chr6-103466478-103466979 | 8 |
| chr5-92562702-92563203 | 1 |
| chr12-25059825-25060326 | 8 |
| chr16-80619797-80620298 | 1 |
| chr17-53669366-53669867 | 8 |
| chr17-69804102-69804603 | 8 |
| chr2-112927157-112927658 | 8 |
| chr6-3192499-3193000 | 8 |
| chr9-55556552-55557053 | 6 |
| chr15-79296369-79296870 | 8 |
| chr14-83376628-83377129 | 8 |
| chr9-30377901-30378402 | 8 |
| chr4-10840884-10841385 | 8 |
| chr2-144603766-144604267 | 8 |
| chr1-180908562-180909063 | 8 |
| chr13-100633309-100633810 | 8 |
| chr10-28830258-28830759 | 8 |
| chr3-15374404-15374905 | 5 |
| chr3-143364717-143365218 | 1 |
| chr2-60780510-60781011 | 5 |
| chr5-109577463-109577964 | 1 |
| chr2-157938623-157939124 | 8 |
| chr3-153810194-153810695 | 5 |
| chr15-2196585-2197086 | 8 |
| chr5-22297239-22297740 | 8 |
| chr19-35750637-35751138 | 8 |
| chr4-81427369-81427870 | 5 |
| chr7-95228665-95229166 | 8 |
| chr2-142350726-142351227 | 8 |
| chr2-243403490-243403991 | 5 |
| chr4-16599154-16599655 | 8 |
| chr20-49691678-49692179 | 8 |
| chr10-104139841-104140342 | 8 |
| chr15-94555236-94555737 | 5 |
| chr2-43588596-43589097 | 8 |
| chr5-137076740-137077241 | 8 |

| | |
|---|---|
| chr1-99119544-99120045 | 1 |
| chr13-78811006-78811507 | 6 |
| chr2-109720425-109720926 | 5 |
| chr8-56107547-56108048 | 6 |
| chr3-161462706-161463207 | 8 |
| chr13-32212077-32212578 | 5 |
| chr1-153410280-153410781 | 8 |
| chr8-70872527-70873028 | 8 |
| chr4-60187712-60188213 | 6 |
| chr5-144073003-144073504 | 1 |
| chr3-89799873-89800374 | 8 |
| chr6-22489161-22489662 | 1 |
| chr3-117530224-117530725 | 8 |
| chr20-9268999-9269500 | 6 |
| chr13-42178336-42178837 | 8 |
| chr19-56411986-56412487 | 1 |
| chr15-20455721-20456222 | 8 |
| chr4-13693825-13694326 | 1 |
| chr12-22318243-22318744 | 5 |
| chr9-40372111-40372612 | 6 |
| chrX-135262620-135263121 | 8 |
| chr3-79014646-79015147 | 6 |
| chr1-138104301-138104802 | 8 |
| chr3-34055546-34056047 | 8 |
| chr15-96710207-96710708 | 6 |
| chr13-92919750-92920251 | 8 |
| chr11-32224814-32225315 | 6 |
| chr2-186589967-186590468 | 8 |
| chr1-13047942-13048443 | 6 |
| chr1-15148442-15148943 | 5 |
| chr3-116594043-116594544 | 8 |
| chr8-19734146-19734647 | 8 |
| chr12-27470420-27470921 | 8 |
| chr17-12839331-12839832 | 8 |
| chr8-111444901-111445402 | 6 |
| chr6-138745747-138746248 | 5 |
| chr4-59894479-59894980 | 1 |
| chr7-131960425-131960926 | 5 |
| chr1-165307935-165308436 | 8 |
| chr4-66333083-66333584 | 1 |
| chr13-50126807-50127308 | 8 |
| chr5-35870756-35871257 | 8 |
| chr9-52080838-52081339 | 8 |

| | |
|---|---|
| chr17-17867311-17867812 | 8 |
| chr12-10155575-10156076 | 8 |
| chr19-25315050-25315551 | 8 |
| chr17-59125099-59125600 | 8 |
| chr9-86243197-86243698 | 6 |
| chr1-159832102-159832603 | 1 |
| chr15-66518910-66519411 | 5 |
| chr12-41800718-41801219 | 6 |
| chr3-121581557-121582058 | 1 |
| chr8-30497884-30498385 | 8 |
| chr14-97388996-97389497 | 8 |
| chr2-223913040-223913541 | 8 |
| chr10-16628087-16628588 | 8 |
| chr1-39376120-39376621 | 6 |
| chr13-89122985-89123486 | 8 |
| chr11-59499712-59500213 | 8 |
| chr4-178775809-178776310 | 5 |
| chr1-177359341-177359842 | 1 |
| chr3-140985912-140986413 | 5 |
| chr15-16345008-16345509 | 6 |
| chr2-32919811-32920312 | 8 |
| chr2-61380266-61380767 | 5 |
| chr3-44134939-44135440 | 8 |
| chr14-52467893-52468394 | 5 |
| chr6-71206304-71206805 | 5 |
| chr5-164823785-164824286 | 8 |
| chr10-41929000-41929501 | 8 |
| chr13-92285386-92285887 | 5 |
| chr20-31456903-31457404 | 5 |
| chr7-93292787-93293288 | 1 |
| chr11-67864190-67864691 | 8 |
| chr7-64869601-64870102 | 6 |
| chr17-24231891-24232392 | 6 |
| chr4-177633900-177634401 | 8 |
| chr2-215404795-215405296 | 6 |
| chr7-129381047-129381548 | 8 |
| chr8-30451215-30451716 | 6 |
| chr8-36915405-36915906 | 8 |
| chr3-127471233-127471734 | 1 |
| chr11-21891179-21891680 | 5 |
| chr11-59082260-59082761 | 8 |
| chr11-72025999-72026500 | 8 |
| chr1-186342369-186342870 | 6 |

| | |
|---|---|
| chr4-133688431-133688932 | 8 |
| chr20-18128001-18128502 | 1 |
| chr3-23912872-23913373 | 5 |
| chr4-168350859-168351360 | 8 |
| chr1-52962745-52963246 | 5 |
| chr6-25872626-25873127 | 8 |
| chr13-101235767-101236268 | 8 |
| chr6-45407051-45407552 | 6 |
| chr8-114570365-114570866 | 6 |
| chr1-117953629-117954130 | 8 |
| chr11-36832791-36833292 | 8 |
| chr5-141333781-141334282 | 8 |
| chr1-216689822-216690323 | 8 |
| chr15-38154359-38154860 | 1 |
| chr16-74727316-74727817 | 8 |
| chr1-143694456-143694957 | 1 |
| chr5-62429678-62430179 | 6 |
| chr1-123991297-123991798 | 5 |
| chr3-111485348-111485849 | 5 |
| chr3-135085628-135086129 | 5 |
| chr16-5552818-5553319 | 6 |
| chr4-179730993-179731494 | 5 |
| chr10-59997282-59997783 | 5 |
| chr9-83697388-83697889 | 6 |
| chr13-65183086-65183587 | 8 |
| chr11-83959104-83959605 | 8 |
| chr7-66639001-66639502 | 8 |
| chr6-62635817-62636318 | 5 |
| chr14-77903905-77904406 | 8 |
| chr1-126017053-126017554 | 6 |
| chr2-81637951-81638452 | 5 |
| chr18-47102042-47102543 | 5 |
| chr5-153794588-153795089 | 8 |
| chr5-102794689-102795190 | 8 |
| chr1-190414704-190415205 | 1 |
| chr7-118706078-118706579 | 8 |
| chr8-115365880-115366381 | 6 |
| chr9-97170811-97171312 | 6 |
| chr2-40248210-40248711 | 8 |
| chrX-106131083-106131584 | 6 |
| chr10-38591803-38592304 | 1 |
| chr10-45117413-45117914 | 1 |
| chr5-96629521-96630022 | 8 |

| | |
|---|---|
| chr7-100132403-100132904 | 5 |
| chr4-84256222-84256723 | 8 |
| chr1-204047057-204047558 | 1 |
| chr1-8387913-8388414 | 5 |
| chr1-146259114-146259615 | 8 |
| chr12-34501024-34501525 | 5 |
| chrX-79973192-79973693 | 1 |
| chr7-122806348-122806849 | 6 |
| chr12-21209565-21210066 | 8 |
| chr2-138600324-138600825 | 8 |
| chr3-77669717-77670218 | 5 |
| chr3-101581638-101582139 | 1 |
| chr5-146888777-146889278 | 5 |
| chr16-43720924-43721425 | 5 |
| chr7-111854459-111854960 | 5 |
| chr3-11758249-11758750 | 8 |
| chr8-43881065-43881566 | 5 |
| chr3-12750001-12750502 | 5 |
| chr3-93023882-93024383 | 8 |
| chr8-11683794-11684295 | 8 |
| chr4-74668858-74669359 | 8 |
| chr3-132543161-132543662 | 5 |
| chr8-47874692-47875193 | 6 |
| chr12-28320878-28321379 | 8 |
| chr10-90594101-90594602 | 8 |
| chr19-5459798-5460299 | 1 |
| chr6-20876304-20876805 | 8 |
| chr18-67073354-67073855 | 5 |
| chr3-133809130-133809631 | 8 |
| chrX-23571510-23572011 | 8 |
| chr5-113701412-113701913 | 5 |
| chr14-2873464-2873965 | 6 |
| chr13-74251151-74251652 | 5 |
| chr15-46609989-46610490 | 8 |
| chr15-31859835-31860336 | 6 |
| chr17-26183601-26184102 | 5 |
| chr9-91477367-91477868 | 6 |
| chr5-133130554-133131055 | 5 |
| chr1-153433988-153434489 | 8 |
| chr4-116002190-116002691 | 8 |
| chr11-11438519-11439020 | 8 |
| chr8-58351245-58351746 | 1 |
| chr18-58754917-58755418 | 1 |

| | |
|---|---|
| chr2-50858314-50858815 | 5 |
| chr11-17170913-17171414 | 6 |
| chr2-44377340-44377841 | 1 |
| chr2-240258317-240258818 | 1 |
| chr4-46939098-46939599 | 8 |
| chr3-79185720-79186221 | 5 |
| chrX-125202556-125203057 | 1 |
| chr10-106680409-106680910 | 8 |
| chr6-99229152-99229653 | 5 |
| chr1-217154261-217154762 | 6 |
| chr4-146489089-146489590 | 8 |
| chr9-89579834-89580335 | 8 |
| chr11-78504297-78504798 | 8 |
| chr1-85854561-85855062 | 8 |
| chr4-150145743-150146244 | 6 |
| chr16-56221016-56221517 | 8 |
| chr17-40136278-40136779 | 6 |
| chr4-57420423-57420924 | 8 |
| chr6-20220580-20221081 | 8 |
| chr15-26702017-26702518 | 5 |
| chr4-77093943-77094444 | 8 |
| chr5-68196806-68197307 | 6 |
| chr1-191808546-191809047 | 8 |
| chr2-235641054-235641555 | 8 |
| chr6-131095637-131096138 | 5 |
| chr6-51901871-51902372 | 8 |
| chr2-29149042-29149543 | 1 |
| chr4-141870573-141871074 | 5 |
| chr5-92670429-92670930 | 5 |
| chr9-111913048-111913549 | 6 |
| chr7-23544841-23545342 | 8 |
| chr2-42114868-42115369 | 8 |
| chr2-219918184-219918685 | 8 |
| chr14-1865368-1865869 | 8 |
| chr3-29380725-29381226 | 8 |
| chr10-73425641-73426142 | 1 |
| chr1-224236036-224236537 | 8 |
| chr9-104050502-104051003 | 8 |
| chr14-56826174-56826675 | 8 |
| chr2-240323600-240324101 | 6 |
| chr18-67368795-67369296 | 8 |
| chr4-83266255-83266756 | 5 |
| chr6-47707577-47708078 | 6 |

| | |
|---|---|
| chr12-1299250-1299751 | 8 |
| chr10-65626824-65627325 | 8 |
| chr11-74006416-74006917 | 8 |
| chr2-221368709-221369210 | 8 |
| chr1-154480995-154481496 | 5 |
| chr4-163652187-163652688 | 5 |
| chr10-60415808-60416309 | 5 |
| chr17-77778770-77779271 | 8 |
| chr7-104750164-104750665 | 8 |
| chr3-7852636-7853137 | 6 |
| chr2-190678001-190678502 | 8 |
| chr7-123441755-123442256 | 8 |
| chr1-239501955-239502456 | 5 |
| chr5-121488909-121489410 | 5 |
| chr2-87966878-87967379 | 8 |
| chr5-150917187-150917688 | 6 |
| chr9-92457410-92457911 | 5 |
| chr9-6890321-6890822 | 8 |
| chr11-3055987-3056488 | 5 |
| chr5-121474027-121474528 | 8 |
| chr15-13000480-13000981 | 1 |
| chr10-35493012-35493513 | 8 |
| chr3-127118924-127119425 | 5 |
| chr4-146872839-146873340 | 5 |
| chr1-242699548-242700049 | 8 |
| chr16-28925551-28926052 | 5 |
| chr4-158318568-158319069 | 6 |
| chr10-101693751-101694252 | 6 |
| chr10-105572478-105572979 | 6 |
| chr2-240238453-240238954 | 1 |
| chr2-30362069-30362570 | 5 |
| chr6-122139348-122139849 | 8 |
| chr6-9970836-9971337 | 6 |
| chr2-210595736-210596237 | 5 |
| chr8-53720635-53721136 | 1 |
| chr4-104686297-104686798 | 6 |
| chr14-12211215-12211716 | 1 |
| chrX-32944117-32944618 | 6 |
| chr11-34502861-34503362 | 1 |
| chrX-97018370-97018871 | 8 |
| chr16-76183772-76184273 | 5 |
| chr8-61319634-61320135 | 5 |
| chr4-132041799-132042300 | 8 |

| | |
|---|---|
| chr1-221827685-221828186 | 8 |
| chr12-26144120-26144621 | 5 |
| chr1-252275082-252275583 | 8 |
| chr2-245010852-245011353 | 8 |
| chr5-51430991-51431492 | 8 |
| chr1-41257428-41257929 | 8 |
| chr3-32015345-32015846 | 6 |
| chr10-100013790-100014291 | 8 |
| chr7-53660915-53661416 | 1 |
| chr2-173272434-173272935 | 5 |
| chr2-175264712-175265213 | 1 |
| chr10-45333486-45333987 | 8 |
| chr11-34201892-34202393 | 6 |
| chr5-27825251-27825752 | 8 |
| chr2-96423840-96424341 | 8 |
| chr6-90764857-90765358 | 6 |
| chr15-18745355-18745856 | 8 |
| chr3-18438805-18439306 | 8 |
| chr5-165080538-165081039 | 8 |
| chr4-46245101-46245602 | 6 |
| chr14-71939309-71939810 | 8 |
| chr9-102820216-102820717 | 5 |
| chr2-149575039-149575540 | 6 |
| chr1-108557356-108557857 | 5 |
| chr1-184725113-184725614 | 8 |
| chr2-209901435-209901936 | 8 |
| chr18-27050283-27050784 | 8 |
| chr6-14668609-14669110 | 5 |
| chr6-40054028-40054529 | 6 |
| chr15-98782182-98782683 | 1 |
| chr2-132893526-132894027 | 8 |
| chr14-3505671-3506172 | 1 |
| chr10-17097903-17098404 | 8 |
| chr1-16837509-16838010 | 5 |
| chr9-72612454-72612955 | 8 |
| chr11-27008544-27009045 | 8 |
| chr17-15553961-15554462 | 5 |
| chr2-196966458-196966959 | 8 |
| chr5-107993183-107993684 | 6 |
| chr8-72389916-72390417 | 5 |
| chr8-72489172-72489673 | 5 |
| chr7-113637420-113637921 | 8 |
| chr1-146983814-146984315 | 1 |

| | |
|---|---|
| chr17-52711495-52711996 | 8 |
| chr1-135804016-135804517 | 5 |
| chr8-51140226-51140727 | 8 |
| chr8-84036748-84037249 | 8 |
| chr8-51417286-51417787 | 5 |
| chr8-12648042-12648543 | 8 |
| chr13-49965272-49965773 | 5 |
| chr2-184240851-184241352 | 5 |
| chr1-221095282-221095783 | 5 |
| chr7-90546087-90546588 | 6 |
| chr2-204129257-204129758 | 1 |
| chr13-71680998-71681499 | 1 |
| chr19-56124023-56124524 | 5 |
| chr6-26455889-26456390 | 8 |
| chr9-96727450-96727951 | 6 |
| chrX-67323743-67324244 | 8 |
| chr1-52963259-52963760 | 5 |
| chr9-61071844-61072345 | 8 |
| chr14-983367-983868 | 8 |
| chr18-38396374-38396875 | 6 |
| chr1-172814653-172815154 | 5 |
| chr13-902037-902538 | 5 |
| chr3-108306208-108306709 | 5 |
| chr6-73238612-73239113 | 1 |
| chr9-109768188-109768689 | 6 |
| chr19-11111213-11111714 | 8 |
| chr6-122567736-122568237 | 6 |
| chr9-59428380-59428881 | 8 |
| chr5-150883302-150883803 | 5 |
| chr6-49244335-49244836 | 8 |
| chr15-38242-38743 | 8 |
| chr17-44781484-44781985 | 8 |
| chr2-197328793-197329294 | 8 |
| chr6-87578326-87578827 | 8 |
| chr6-130546056-130546557 | 6 |
| chr5-49114485-49114986 | 8 |
| chr10-1922292-1922793 | 8 |
| chr18-71329048-71329549 | 6 |
| chr13-96330474-96330975 | 5 |
| chr17-34405731-34406232 | 8 |
| chr18-73109891-73110392 | 6 |
| chr9-74510949-74511450 | 8 |
| chr16-78235768-78236269 | 1 |

| | |
|---|---|
| chr5-8177866-8178367 | 1 |
| chr4-133633119-133633620 | 8 |
| chr4-67886298-67886799 | 1 |
| chr1-83374711-83375212 | 8 |
| chr4-154431093-154431594 | 8 |
| chr16-45534338-45534839 | 8 |
| chr5-106026004-106026505 | 8 |
| chr11-76820597-76821098 | 8 |
| chr3-76577755-76578256 | 8 |
| chr4-79081112-79081613 | 8 |
| chr13-63461313-63461814 | 8 |
| chr2-137380489-137380990 | 5 |
| chr9-65341000-65341501 | 1 |
| chr2-220049787-220050288 | 8 |
| chr4-91024636-91025137 | 6 |
| chr3-154256990-154257491 | 8 |
| chr1-221520663-221521164 | 6 |
| chr14-84570212-84570713 | 6 |
| chr11-81019474-81019975 | 8 |
| chr5-63633048-63633549 | 8 |
| chr3-37085230-37085731 | 8 |
| chr15-77110240-77110741 | 6 |
| chr13-92114105-92114606 | 8 |
| chr10-60942979-60943480 | 1 |
| chr5-145889957-145890458 | 8 |
| chr16-32054712-32055213 | 6 |
| chr17-8725840-8726341 | 5 |
| chr9-100491308-100491809 | 5 |
| chr4-7995748-7996249 | 6 |
| chr16-37375857-37376358 | 8 |
| chr1-176998283-176998784 | 5 |
| chr1-127817217-127817718 | 8 |
| chr1-87066642-87067143 | 8 |
| chr9-85031735-85032236 | 1 |
| chr19-18505156-18505657 | 8 |
| chr19-16190762-16191263 | 5 |
| chrX-105972141-105972642 | 8 |
| chr1-234204278-234204779 | 6 |
| chr1-239371733-239372234 | 8 |
| chr11-31001708-31002209 | 6 |
| chr2-160630337-160630838 | 6 |
| chr18-25378448-25378949 | 8 |
| chr8-70713014-70713515 | 1 |

| | |
|---|---|
| chr16-10433046-10433547 | 6 |
| chr13-65383932-65384433 | 8 |
| chr17-69701473-69701974 | 1 |
| chr10-66514222-66514723 | 5 |
| chr3-134089849-134090350 | 8 |
| chr7-68331536-68332037 | 1 |
| chrX-7692770-7693271 | 6 |
| chr8-43397170-43397671 | 1 |
| chr10-41418087-41418588 | 1 |
| chr16-44456895-44457396 | 8 |
| chr4-82846506-82847007 | 5 |
| chr5-152534347-152534848 | 6 |
| chrX-28039510-28040011 | 5 |
| chr1-7372685-7373186 | 6 |
| chr7-8627894-8628395 | 8 |
| chr14-81614058-81614559 | 5 |
| chr4-43487157-43487658 | 6 |
| chr9-73395792-73396293 | 8 |
| chr1-250481268-250481769 | 5 |
| chr1-154816826-154817327 | 5 |
| chr5-139293303-139293804 | 5 |
| chr6-22995796-22996297 | 5 |
| chr8-42820334-42820835 | 5 |
| chr5-76876366-76876867 | 8 |
| chr1-223747331-223747832 | 8 |
| chr16-56275008-56275509 | 6 |
| chr6-72490182-72490683 | 8 |
| chr12-45235277-45235778 | 8 |
| chr8-96368175-96368676 | 1 |
| chr19-6048916-6049417 | 6 |
| chr5-53815687-53816188 | 5 |
| chr9-7329402-7329903 | 6 |
| chr1-224594675-224595176 | 8 |
| chr1-185952625-185953126 | 8 |
| chr2-226982271-226982772 | 8 |
| chr2-245192924-245193425 | 6 |
| chr15-74570307-74570808 | 8 |
| chr1-228745279-228745780 | 6 |
| chr11-11977738-11978239 | 6 |
| chr13-30333057-30333558 | 1 |
| chr6-106849676-106850177 | 8 |
| chr1-6797681-6798182 | 8 |
| chr10-33904720-33905221 | 1 |

| | |
|---|---|
| chr1-191921599-191922100 | 6 |
| chr5-101080291-101080792 | 8 |
| chr20-44307395-44307896 | 6 |
| chr3-69545568-69546069 | 8 |
| chr10-82339269-82339770 | 8 |
| chr7-72684775-72685276 | 6 |
| chr1-96921668-96922169 | 5 |
| chr14-68010618-68011119 | 8 |
| chr2-238170863-238171364 | 1 |
| chr10-20269483-20269984 | 5 |
| chr18-72445426-72445927 | 8 |
| chr15-32720257-32720758 | 8 |
| chr2-219722491-219722992 | 8 |
| chr20-12736533-12737034 | 5 |
| chr14-35306138-35306639 | 8 |
| chrX-54963281-54963782 | 5 |
| chr5-1557217-1557718 | 6 |
| chr2-78875374-78875875 | 1 |
| chr6-52985067-52985568 | 5 |
| chr13-65299405-65299906 | 1 |
| chrX-127095013-127095514 | 6 |
| chr14-75427330-75427831 | 8 |
| chr2-191072990-191073491 | 8 |
| chr6-22072825-22073326 | 1 |
| chr10-90919538-90920039 | 1 |
| chr15-4871085-4871586 | 6 |
| chr1-124892201-124892702 | 6 |
| chr5-149644808-149645309 | 8 |
| chrX-11441884-11442385 | 1 |
| chr3-90655545-90656046 | 6 |
| chr16-47050995-47051496 | 6 |
| chr3-123806118-123806619 | 8 |
| chr9-88926629-88927130 | 8 |
| chr8-11079737-11080238 | 8 |
| chr18-59401387-59401888 | 1 |
| chr15-60849557-60850058 | 6 |
| chr2-121555028-121555529 | 5 |
| chr17-74235027-74235528 | 6 |
| chr17-20725856-20726357 | 1 |
| chr3-42162366-42162867 | 5 |
| chr1-11536754-11537255 | 8 |
| chr16-23052966-23053467 | 8 |
| chr17-29336665-29337166 | 1 |

| | |
|---|---|
| chr13-92899616-92900117 | 5 |
| chr18-61198554-61199055 | 8 |
| chr2-137019768-137020269 | 6 |
| chr7-51945714-51946215 | 5 |
| chr14-73749153-73749654 | 6 |
| chr15-66215665-66216166 | 6 |
| chr1-86828320-86828821 | 6 |
| chr10-53207107-53207608 | 5 |
| chr15-34915523-34916024 | 5 |
| chr9-14645794-14646295 | 8 |
| chr3-108376379-108376880 | 8 |
| chr7-22907577-22908078 | 8 |
| chr18-32417371-32417872 | 6 |
| chr18-35736913-35737414 | 6 |
| chr13-37515685-37516186 | 8 |
| chr5-71628955-71629456 | 5 |
| chr1-130606612-130607113 | 6 |
| chr3-60021995-60022496 | 1 |
| chr13-38581146-38581647 | 8 |
| chr2-141656426-141656927 | 8 |
| chr2-241426369-241426870 | 5 |
| chr13-106190337-106190838 | 6 |
| chr1-54282790-54283291 | 6 |
| chr1-23069657-23070158 | 5 |
| chr1-138749281-138749782 | 8 |
| chr1-201778447-201778948 | 8 |
| chr7-11738493-11738994 | 8 |
| chr16-33814422-33814923 | 8 |
| chr13-38963843-38964344 | 5 |
| chr13-74811456-74811957 | 1 |
| chr6-123805586-123806087 | 8 |
| chr3-112335441-112335942 | 8 |
| chr11-80454574-80455075 | 8 |
| chr1-85900987-85901488 | 8 |
| chr16-38619164-38619665 | 6 |
| chr10-20239669-20240170 | 8 |
| chr1-43954017-43954518 | 8 |
| chr7-86634275-86634776 | 1 |
| chr3-111917419-111917920 | 1 |
| chr13-74794573-74795074 | 8 |
| chr13-91483564-91484065 | 8 |
| chr13-64896804-64897305 | 6 |
| chr2-197114973-197115474 | 8 |

| | |
|---|---|
| chr17-52193868-52194369 | 8 |
| chr8-40955194-40955695 | 5 |
| chr19-31334311-31334812 | 6 |
| chr6-71847895-71848396 | 1 |
| chr11-33336166-33336667 | 5 |
| chr17-55771455-55771956 | 8 |
| chr13-65542039-65542540 | 8 |
| chr5-78714937-78715438 | 6 |
| chr9-86930393-86930894 | 1 |
| chr14-93930475-93930976 | 6 |
| chr7-41909201-41909702 | 8 |
| chr6-105881929-105882430 | 8 |
| chr6-72151339-72151840 | 5 |
| chr14-2397669-2398170 | 8 |
| chr2-113787809-113788310 | 6 |
| chr2-240384671-240385172 | 1 |
| chr11-50235257-50235758 | 1 |
| chr17-21761600-21762101 | 1 |
| chr2-164465883-164466384 | 8 |
| chr16-82431129-82431630 | 8 |
| chr10-15624622-15625123 | 5 |
| chr5-60960771-60961272 | 5 |
| chr18-39782277-39782778 | 6 |
| chr10-38048695-38049196 | 8 |
| chr14-62502484-62502985 | 8 |
| chr6-5680809-5681310 | 8 |
| chr8-120761894-120762395 | 6 |
| chr1-135209386-135209887 | 8 |
| chr9-98248233-98248734 | 8 |
| chr9-60603497-60603998 | 8 |
| chr13-29770952-29771453 | 8 |
| chr6-72911366-72911867 | 5 |
| chr5-104112274-104112775 | 8 |
| chr6-16898494-16898995 | 6 |
| chr1-178653077-178653578 | 6 |
| chr16-78798545-78799046 | 8 |
| chr7-111973415-111973916 | 1 |
| chr3-13178804-13179305 | 8 |
| chr4-62162617-62163118 | 1 |
| chr16-5662843-5663344 | 8 |
| chr17-16474763-16475264 | 1 |
| chr3-120669645-120670146 | 8 |
| chr5-18778693-18779194 | 8 |

| | |
|---|---|
| chr11-66432803-66433304 | 8 |
| chr1-89619195-89619696 | 6 |
| chr5-77474380-77474881 | 6 |
| chr11-52612978-52613479 | 6 |
| chr12-21333875-21334376 | 6 |
| chr14-43245612-43246113 | 8 |
| chr18-47203345-47203846 | 8 |
| chr5-58839196-58839697 | 1 |
| chr17-1052306-1052807 | 8 |
| chr2-223013612-223014113 | 6 |
| chr1-246943585-246944086 | 8 |
| chr3-114010242-114010743 | 8 |
| chr3-2682236-2682737 | 5 |
| chr17-46883705-46884206 | 6 |
| chr6-57153285-57153786 | 1 |
| chr4-154317487-154317988 | 1 |
| chr6-131312403-131312904 | 8 |
| chr17-1497371-1497872 | 8 |
| chr6-51792439-51792940 | 8 |
| chr9-66109271-66109772 | 6 |
| chr2-137020751-137021252 | 6 |
| chr20-13304964-13305465 | 6 |
| chr19-47129685-47130186 | 1 |
| chr18-33461375-33461876 | 5 |
| chr6-107102744-107103245 | 8 |
| chr8-66769370-66769871 | 8 |
| chr6-51946952-51947453 | 8 |
| chr2-30260581-30261082 | 8 |
| chr16-55841629-55842130 | 6 |
| chr15-30214575-30215076 | 5 |
| chr2-50914421-50914922 | 5 |
| chr1-11539454-11539955 | 8 |
| chr2-45587613-45588114 | 8 |
| chr18-62123410-62123911 | 8 |
| chr15-100165562-100166063 | 8 |
| chr11-79271700-79272201 | 8 |
| chr5-100523253-100523754 | 6 |
| chr15-7619653-7620154 | 5 |
| chr12-31760576-31761077 | 8 |
| chr3-78692739-78693240 | 1 |
| chr13-65077113-65077614 | 8 |
| chr2-137009591-137010092 | 1 |
| chr1-77395133-77395634 | 8 |

| | |
|---|---|
| chr8-46508632-46509133 | 8 |
| chr2-46755444-46755945 | 8 |
| chr20-8596789-8597290 | 8 |
| chr9-113018024-113018525 | 1 |
| chr16-31802564-31803065 | 6 |
| chr5-119915100-119915601 | 8 |
| chr4-131864600-131865101 | 5 |
| chr18-66848875-66849376 | 6 |
| chr2-75972738-75973239 | 8 |
| chr6-105396597-105397098 | 8 |
| chr2-141388216-141388717 | 8 |
| chr2-148488188-148488689 | 8 |
| chrX-65613576-65614077 | 6 |
| chr19-555768-556269 | 8 |
| chr8-53729677-53730178 | 6 |
| chr18-60904324-60904825 | 5 |
| chr9-85215116-85215617 | 8 |
| chr3-157415725-157416226 | 6 |
| chr2-235550347-235550848 | 1 |
| chr4-147274013-147274514 | 6 |
| chr2-191436015-191436516 | 8 |
| chr8-80944701-80945202 | 6 |
| chr5-57996060-57996561 | 8 |
| chr13-39510008-39510509 | 8 |
| chr4-159932080-159932581 | 8 |
| chr2-183626777-183627278 | 8 |
| chr7-46754617-46755118 | 8 |
| chr5-110769331-110769832 | 1 |
| chrX-15947871-15948372 | 1 |
| chr1-240800438-240800939 | 1 |
| chr20-1485616-1486117 | 5 |
| chr2-77094417-77094918 | 5 |
| chr4-40717959-40718460 | 1 |
| chr5-121532607-121533108 | 8 |
| chr6-88434251-88434752 | 6 |
| chr3-2262690-2263191 | 5 |
| chr8-1572664-1573165 | 1 |
| chr3-43059208-43059709 | 1 |
| chr8-48951030-48951531 | 6 |
| chr16-16915893-16916394 | 6 |
| chr8-60885720-60886221 | 8 |
| chr2-56143251-56143752 | 1 |
| chr4-170485535-170486036 | 8 |

| | |
|---|---|
| chr2-237728398-237728899 | 8 |
| chr3-36638257-36638758 | 1 |
| chr1-57588504-57589005 | 6 |
| chr1-92039192-92039693 | 6 |
| chr4-62622539-62623040 | 6 |
| chr9-74944272-74944773 | 8 |
| chr5-134193279-134193780 | 6 |
| chr12-19335219-19335720 | 5 |
| chr17-22071797-22072298 | 8 |
| chr13-82909296-82909797 | 5 |
| chr13-40198417-40198918 | 8 |
| chr18-30527882-30528383 | 8 |
| chr3-151941487-151941988 | 6 |
| chr8-12067159-12067660 | 5 |
| chr3-96020628-96021129 | 6 |
| chr20-11014125-11014626 | 8 |
| chr20-34543447-34543948 | 5 |
| chr5-67771633-67772134 | 8 |
| chr15-20732166-20732667 | 8 |
| chr5-5382608-5383109 | 8 |
| chr3-93174375-93174876 | 8 |
| chr17-76691769-76692270 | 8 |
| chrX-62219945-62220446 | 1 |
| chr9-89679896-89680397 | 8 |
| chr4-123984186-123984687 | 8 |
| chr4-181744049-181744550 | 6 |
| chr10-62975118-62975619 | 5 |
| chr10-71169318-71169819 | 8 |
| chr6-138260337-138260838 | 8 |
| chr10-71024033-71024534 | 5 |
| chr15-54945624-54946125 | 8 |
| chr8-56875142-56875643 | 8 |
| chr13-39506924-39507425 | 1 |
| chr3-104842590-104843091 | 5 |
| chr17-52974526-52975027 | 1 |
| chr13-30398237-30398738 | 8 |
| chr3-47222686-47223187 | 8 |
| chr1-192751329-192751830 | 8 |
| chr5-132871152-132871653 | 1 |
| chrX-22209207-22209708 | 1 |
| chr8-83310114-83310615 | 6 |
| chr10-41112167-41112668 | 1 |
| chr1-196245346-196245847 | 5 |

| | |
|---|---|
| chr18-69739167-69739668 | 8 |
| chr3-121808136-121808637 | 1 |
| chr4-147357832-147358333 | 6 |
| chrX-73997000-73997501 | 6 |
| chr3-28463484-28463985 | 5 |
| chr10-85960116-85960617 | 8 |
| chr16-2405731-2406232 | 8 |
| chrX-97142007-97142508 | 5 |
| chr14-81110112-81110613 | 8 |
| chrX-112839743-112840244 | 6 |
| chr13-43648938-43649439 | 8 |
| chr17-23478034-23478535 | 8 |
| chr6-125800417-125800918 | 8 |
| chr1-23359342-23359843 | 8 |
| chr19-42068890-42069391 | 1 |
| chr9-14097229-14097730 | 1 |
| chr16-77655938-77656439 | 8 |
| chr8-64705555-64706056 | 5 |
| chr19-44618241-44618742 | 8 |
| chr9-26203394-26203895 | 5 |
| chr9-63193515-63194016 | 6 |
| chr5-105781470-105781971 | 6 |
| chr15-83620226-83620727 | 5 |
| chr7-56722934-56723435 | 8 |
| chr17-76255696-76256197 | 8 |
| chr12-25090006-25090507 | 8 |
| chr1-108916243-108916744 | 6 |
| chr5-131932952-131933453 | 8 |
| chr8-96699385-96699886 | 6 |
| chr2-221364322-221364823 | 8 |
| chr16-4597179-4597680 | 5 |
| chr7-8999175-8999676 | 8 |
| chr10-78770013-78770514 | 6 |
| chr5-162435029-162435530 | 8 |
| chr9-107236848-107237349 | 1 |
| chr7-19815570-19816071 | 8 |
| chr16-8571274-8571775 | 8 |
| chr3-107160905-107161406 | 1 |
| chr2-118892720-118893221 | 6 |
| chr5-115923338-115923839 | 8 |
| chr7-89458217-89458718 | 5 |
| chr3-147940798-147941299 | 8 |
| chr7-8604000-8604501 | 8 |

| | |
|---|---|
| chr11-4214293-4214794 | 8 |
| chr17-31556833-31557334 | 8 |
| chr16-12311315-12311816 | 5 |
| chr4-133253627-133254128 | 1 |
| chr17-54507712-54508213 | 8 |
| chr7-57543036-57543537 | 5 |
| chr8-108523604-108524105 | 8 |
| chr18-48424981-48425482 | 5 |
| chr9-77649959-77650460 | 8 |
| chr17-23307003-23307504 | 1 |
| chr1-155859553-155860054 | 5 |
| chr9-40319951-40320452 | 8 |
| chr5-149745275-149745776 | 8 |
| chr5-60735640-60736141 | 8 |
| chr1-173356869-173357370 | 6 |
| chr2-56303543-56304044 | 1 |
| chr4-69771459-69771960 | 8 |
| chr4-117952382-117952883 | 8 |
| chr16-33606845-33607346 | 8 |
| chr10-11717077-11717578 | 8 |
| chr15-99192459-99192960 | 1 |
| chr7-65922347-65922848 | 6 |
| chr17-46884735-46885236 | 6 |
| chr15-8928985-8929486 | 5 |
| chr1-15389366-15389867 | 8 |
| chr1-232360570-232361071 | 6 |
| chr10-41389372-41389873 | 8 |
| chr11-58011796-58012297 | 1 |
| chr3-165815208-165815709 | 6 |
| chr8-13836521-13837022 | 1 |
| chr1-78588571-78589072 | 5 |
| chr1-36554051-36554552 | 6 |
| chr17-3958800-3959301 | 8 |
| chr2-113322550-113323051 | 5 |
| chr5-132656823-132657324 | 5 |
| chr1-138729771-138730272 | 1 |
| chr1-12548727-12549228 | 5 |
| chr8-51355845-51356346 | 8 |
| chr2-173041868-173042369 | 6 |
| chr4-132727531-132728032 | 1 |
| chr2-230357927-230358428 | 8 |
| chr13-95238692-95239193 | 6 |
| chr18-9608039-9608540 | 5 |

| | |
|---|---|
| chr7-65993936-65994437 | 1 |
| chr17-18542865-18543366 | 8 |
| chr12-6773478-6773979 | 6 |
| chr14-94393402-94393903 | 5 |
| chr4-126363917-126364418 | 1 |
| chr8-104322964-104323465 | 8 |
| chr8-37394991-37395492 | 5 |
| chr1-86315105-86315606 | 6 |
| chr8-926196-926697 | 6 |
| chr9-98065742-98066243 | 8 |
| chr5-13143053-13143554 | 1 |
| chr15-79538611-79539112 | 8 |
| chr2-246049440-246049941 | 5 |
| chr2-99334395-99334896 | 6 |
| chr6-83025049-83025550 | 8 |
| chr5-24551876-24552377 | 5 |
| chr12-567360-567861 | 8 |
| chr14-56970598-56971099 | 6 |
| chr13-37820380-37820881 | 5 |
| chr15-26903483-26903984 | 8 |
| chr15-76820051-76820552 | 6 |
| chr20-32687452-32687953 | 5 |
| chr7-51447952-51448453 | 1 |
| chr3-86722181-86722682 | 5 |
| chr19-39038590-39039091 | 8 |
| chr11-27787111-27787612 | 5 |
| chr6-41371677-41372178 | 8 |
| chr6-53809972-53810473 | 1 |
| chr5-118113254-118113755 | 8 |
| chr4-58529866-58530367 | 8 |
| chr4-33564090-33564591 | 5 |
| chr12-37056129-37056630 | 8 |
| chr14-10808111-10808612 | 6 |
| chr3-117224155-117224656 | 8 |
| chr13-91988274-91988775 | 1 |
| chr7-96657084-96657585 | 6 |
| chr3-67845105-67845606 | 6 |
| chr16-25336384-25336885 | 1 |
| chr7-107284553-107285054 | 8 |
| chr19-26388880-26389381 | 1 |
| chr2-139930467-139930968 | 5 |
| chr1-207931046-207931547 | 1 |
| chr5-113917221-113917722 | 8 |

| | |
|---|---|
| chr18-48826562-48827063 | 8 |
| chr2-45569564-45570065 | 6 |
| chr15-82469204-82469705 | 8 |
| chr17-66139278-66139779 | 8 |
| chrX-20230452-20230953 | 8 |
| chr5-9540394-9540895 | 8 |
| chr14-67875396-67875897 | 1 |
| chr16-10894415-10894916 | 6 |
| chr5-129144820-129145321 | 1 |
| chr13-74537708-74538209 | 8 |
| chr15-4717236-4717737 | 8 |
| chr9-60623881-60624382 | 8 |
| chr6-119021914-119022415 | 8 |
| chr18-21035404-21035905 | 6 |
| chr4-129625707-129626208 | 5 |
| chr10-5810857-5811358 | 6 |
| chr1-219150628-219151129 | 8 |
| chr13-83083592-83084093 | 5 |
| chr15-54828555-54829056 | 1 |
| chr10-80441398-80441899 | 5 |
| chr10-54454280-54454781 | 8 |
| chr13-50627856-50628357 | 5 |
| chr2-110141239-110141740 | 8 |
| chr2-150473387-150473888 | 6 |
| chr3-21741302-21741803 | 5 |
| chr1-196225524-196226025 | 8 |
| chr3-29277235-29277736 | 5 |
| chr14-9093709-9094210 | 5 |
| chr3-12570189-12570690 | 8 |
| chr18-54975086-54975587 | 1 |
| chr9-7511839-7512340 | 8 |
| chr1-151229653-151230154 | 8 |
| chr1-11489524-11490025 | 6 |
| chr1-215532703-215533204 | 8 |
| chr2-104736146-104736647 | 6 |
| chr10-47512162-47512663 | 6 |
| chr3-7814613-7815114 | 5 |
| chr14-44640721-44641222 | 1 |
| chr12-5384794-5385295 | 5 |
| chr14-86130465-86130966 | 8 |
| chr4-90662359-90662860 | 5 |
| chr3-47214298-47214799 | 6 |
| chr1-207651322-207651823 | 8 |

| | |
|---|---|
| chr10-104033838-104034339 | 8 |
| chr18-74431156-74431657 | 8 |
| chr17-35137326-35137827 | 8 |
| chr4-33092791-33093292 | 8 |
| chrX-64448262-64448763 | 6 |
| chr6-28761928-28762429 | 1 |
| chr2-224209347-224209848 | 8 |
| chr13-80591256-80591757 | 5 |
| chr3-79443616-79444117 | 8 |
| chr6-124985259-124985760 | 6 |
| chr14-34285689-34286190 | 5 |
| chr9-25602964-25603465 | 1 |
| chr10-57180394-57180895 | 8 |
| chr14-94164954-94165455 | 5 |
| chr7-117028790-117029291 | 8 |
| chr5-99722367-99722868 | 1 |
| chr3-111895468-111895969 | 6 |
| chr8-50372142-50372643 | 1 |
| chr1-133748383-133748884 | 5 |
| chrX-147436129-147436630 | 1 |
| chr7-69186100-69186601 | 8 |
| chrX-131893539-131894040 | 6 |
| chr6-111964904-111965405 | 8 |
| chr2-42486438-42486939 | 8 |
| chr2-247600867-247601368 | 6 |
| chr1-20412920-20413421 | 8 |
| chr18-3722796-3723297 | 1 |
| chr6-126540086-126540587 | 6 |
| chr4-27461297-27461798 | 5 |
| chr6-69324698-69325199 | 6 |
| chr7-87816877-87817378 | 8 |
| chr11-29910826-29911327 | 5 |
| chr5-122861838-122862339 | 8 |
| chr5-152926043-152926544 | 5 |
| chr11-17585745-17586246 | 8 |
| chr6-110638579-110639080 | 6 |
| chr4-160264571-160265072 | 5 |
| chr5-19839248-19839749 | 6 |
| chr1-55795081-55795582 | 8 |
| chr1-41928282-41928783 | 6 |
| chr7-109057341-109057842 | 1 |
| chr14-71385305-71385806 | 6 |
| chrX-31939151-31939652 | 6 |

| | |
|---|---|
| chr6-139698429-139698930 | 1 |
| chr2-27812128-27812629 | 8 |
| chr17-26441105-26441606 | 1 |
| chr20-29444923-29445424 | 8 |
| chr5-146567505-146568006 | 8 |
| chr14-71218386-71218887 | 1 |
| chr4-125422342-125422843 | 8 |
| chr10-11094007-11094508 | 8 |
| chr14-94088933-94089434 | 8 |
| chr6-42128408-42128909 | 8 |
| chr3-125563992-125564493 | 6 |
| chr10-82779721-82780222 | 8 |
| chr6-15099118-15099619 | 5 |
| chr1-571847-572348 | 8 |
| chr11-56895927-56896428 | 5 |
| chr16-24869951-24870452 | 1 |
| chr3-133426499-133427000 | 8 |
| chr18-49843923-49844424 | 8 |
| chr15-41862119-41862620 | 8 |
| chr18-66827303-66827804 | 6 |
| chr7-126371971-126372472 | 6 |
| chr1-50777870-50778371 | 8 |
| chr7-29051438-29051939 | 1 |
| chr8-11947048-11947549 | 8 |
| chr16-74943968-74944469 | 1 |
| chr14-93191402-93191903 | 8 |
| chr20-8275681-8276182 | 8 |
| chr4-116521482-116521983 | 8 |
| chr10-57150542-57151043 | 8 |
| chr16-25553915-25554416 | 5 |
| chr7-108961792-108962293 | 8 |
| chr1-183298794-183299295 | 8 |
| chr7-37653411-37653912 | 5 |
| chr6-57857291-57857792 | 5 |
| chr13-12763854-12764355 | 1 |
| chr6-88532862-88533363 | 1 |
| chr4-125682660-125683161 | 8 |
| chr2-140870743-140871244 | 8 |
| chrX-128146597-128147098 | 5 |
| chr10-89606711-89607212 | 8 |
| chr15-40321535-40322036 | 8 |
| chr7-70822882-70823383 | 8 |
| chr15-100190011-100190512 | 8 |

| | |
|---|---|
| chr8-49984736-49985237 | 8 |
| chr2-77945656-77946157 | 1 |
| chr5-56261438-56261939 | 8 |
| chr9-64887343-64887844 | 8 |
| chr1-133562105-133562606 | 8 |
| chr20-4630886-4631387 | 6 |
| chr8-25159540-25160041 | 8 |
| chr15-7568817-7569318 | 8 |
| chr1-87441229-87441730 | 8 |
| chr19-13528778-13529279 | 8 |
| chr5-131651649-131652150 | 8 |
| chr15-52074372-52074873 | 1 |
| chr10-93100449-93100950 | 8 |
| chr19-42394295-42394796 | 8 |
| chr17-20102354-20102855 | 6 |
| chr6-41699451-41699952 | 8 |
| chr10-23042254-23042755 | 6 |
| chr14-89059697-89060198 | 1 |
| chr18-59123465-59123966 | 8 |
| chr2-110999194-110999695 | 8 |
| chr20-2042722-2043223 | 6 |
| chr1-239221097-239221598 | 1 |
| chr6-114398238-114398739 | 8 |
| chr14-32880110-32880611 | 8 |
| chr2-189597151-189597652 | 6 |
| chr2-215774402-215774903 | 8 |
| chr2-26857777-26858278 | 5 |
| chr14-63921178-63921679 | 5 |
| chr4-13563658-13564159 | 8 |
| chr1-198206246-198206747 | 6 |
| chr6-45377717-45378218 | 8 |
| chr1-46201619-46202120 | 6 |
| chr8-57509516-57510017 | 8 |
| chr1-51806545-51807046 | 8 |
| chr6-52806033-52806534 | 8 |
| chr7-121297453-121297954 | 1 |
| chr9-11883711-11884212 | 8 |
| chr3-141563132-141563633 | 8 |
| chr18-46990263-46990764 | 8 |
| chr13-62513345-62513846 | 8 |
| chr6-94481054-94481555 | 6 |
| chr1-215211729-215212230 | 8 |
| chr13-73693703-73694204 | 8 |

| | |
|---|---|
| chr9-59917210-59917711 | 8 |
| chr4-55164442-55164943 | 1 |
| chr10-75055302-75055803 | 1 |
| chr17-79416338-79416839 | 8 |
| chr9-23584321-23584822 | 5 |
| chr4-170526739-170527240 | 8 |
| chr7-127478888-127479389 | 8 |
| chr13-29818744-29819245 | 8 |
| chr8-31513320-31513821 | 8 |
| chr8-37184428-37184929 | 5 |
| chr4-36033333-36033834 | 6 |
| chr14-30806130-30806631 | 8 |
| chr13-81870833-81871334 | 8 |
| chr7-60599581-60600082 | 6 |
| chrX-93318627-93319128 | 6 |
| chr5-71103440-71103941 | 8 |
| chr6-123660624-123661125 | 8 |
| chr3-39779731-39780232 | 6 |
| chr2-207505697-207506198 | 8 |
| chr18-8486062-8486563 | 5 |
| chr8-46954476-46954977 | 8 |
| chr6-23150038-23150539 | 8 |
| chr6-57037263-57037764 | 8 |
| chr19-9644603-9645104 | 5 |
| chr7-109185082-109185583 | 6 |
| chr12-17199818-17200319 | 1 |
| chr16-58271465-58271966 | 1 |
| chr18-75179278-75179779 | 6 |
| chr8-60406039-60406540 | 8 |
| chr1-69118821-69119322 | 8 |
| chr8-119387202-119387703 | 8 |
| chr3-33693501-33694002 | 1 |
| chr14-15495816-15496317 | 8 |
| chr14-9278543-9279044 | 5 |
| chr17-67023431-67023932 | 1 |
| chr17-66047544-66048045 | 6 |
| chr1-215522350-215522851 | 8 |
| chr17-63721479-63721980 | 1 |
| chr4-154278414-154278915 | 8 |
| chr4-14113798-14114299 | 5 |
| chr13-56193358-56193859 | 6 |
| chr5-159401908-159402409 | 5 |
| chr18-78474975-78475476 | 5 |

| | |
|---|---|
| chr3-37725341-37725842 | 8 |
| chr7-106374637-106375138 | 8 |
| chr3-91007987-91008488 | 6 |
| chr4-130730385-130730886 | 8 |
| chr15-37998041-37998542 | 6 |
| chr2-170244474-170244975 | 1 |
| chr8-72173262-72173763 | 8 |
| chr3-37150961-37151462 | 5 |
| chr6-117676081-117676582 | 8 |
| chr6-70886268-70886769 | 8 |
| chr16-81821702-81822203 | 6 |
| chr2-187612247-187612748 | 8 |
| chr7-131592113-131592614 | 8 |
| chr7-43439400-43439901 | 6 |
| chr3-37590558-37591059 | 8 |
| chr1-124571755-124572256 | 6 |
| chr2-190674546-190675047 | 8 |
| chr13-45669818-45670319 | 5 |
| chr1-239513191-239513692 | 1 |
| chr19-46416584-46417085 | 6 |
| chr4-137485914-137486415 | 5 |
| chr3-126819317-126819818 | 8 |
| chr8-121710621-121711122 | 8 |
| chr4-148928287-148928788 | 1 |
| chr18-56834487-56834988 | 5 |
| chr9-91834364-91834865 | 8 |
| chr4-26039253-26039754 | 8 |
| chr9-53678532-53679033 | 5 |
| chr4-62595323-62595824 | 8 |
| chr12-12555484-12555985 | 8 |
| chr8-20964897-20965398 | 5 |
| chr4-81577657-81578158 | 6 |
| chr14-47128691-47129192 | 6 |
| chr18-13641551-13642052 | 6 |
| chr4-703537-704038 | 5 |
| chr2-41537969-41538470 | 8 |
| chr4-128992086-128992587 | 8 |
| chr8-11541697-11542198 | 8 |
| chr2-24915241-24915742 | 6 |
| chr9-57150669-57151170 | 5 |
| chr1-194343999-194344500 | 1 |
| chr5-71429239-71429740 | 6 |
| chr1-118939677-118940178 | 5 |

| | |
|---|---|
| chr6-23230378-23230879 | 5 |
| chr19-15837015-15837516 | 5 |
| chr12-37798932-37799433 | 8 |
| chr14-71297115-71297616 | 1 |
| chr2-242286444-242286945 | 5 |
| chr1-169657082-169657583 | 1 |
| chr3-55848345-55848846 | 5 |
| chr19-26433360-26433861 | 1 |
| chr10-70652104-70652605 | 8 |
| chr1-90674764-90675265 | 8 |
| chr4-2599915-2600416 | 8 |
| chr7-45249267-45249768 | 5 |
| chr11-54962678-54963179 | 8 |
| chr6-105578463-105578964 | 8 |
| chr3-145225708-145226209 | 5 |
| chr3-56255341-56255842 | 8 |
| chr1-12372971-12373472 | 1 |
| chr14-99051363-99051864 | 8 |
| chr16-45426514-45427015 | 8 |
| chr10-53979056-53979557 | 8 |
| chr13-91472495-91472996 | 1 |
| chr20-33801089-33801590 | 6 |
| chr17-47216116-47216617 | 8 |
| chr5-33038426-33038927 | 8 |
| chr3-80880969-80881470 | 6 |
| chr17-18674221-18674722 | 8 |
| chr19-52597372-52597873 | 8 |
| chr7-100137733-100138234 | 8 |
| chr18-31499917-31500418 | 8 |
| chr10-34005941-34006442 | 8 |
| chr14-96911028-96911529 | 8 |
| chr7-122981576-122982077 | 6 |
| chr14-7763859-7764360 | 8 |
| chr6-129176846-129177347 | 6 |
| chr12-28193121-28193622 | 1 |
| chr6-98067020-98067521 | 6 |
| chr14-8197518-8198019 | 5 |
| chr5-129611747-129612248 | 8 |
| chr2-35576577-35577078 | 8 |
| chr11-67446125-67446626 | 8 |
| chr18-19098879-19099380 | 6 |
| chr2-246175924-246176425 | 6 |
| chr3-166336438-166336939 | 8 |

| | |
|---|---|
| chr17-64159659-64160160 | 1 |
| chr19-19282024-19282525 | 8 |
| chr20-29700717-29701218 | 8 |
| chr7-75578665-75579166 | 5 |
| chr19-53412890-53413391 | 1 |
| chr16-56625411-56625912 | 8 |
| chr9-83878551-83879052 | 8 |
| chr3-107190079-107190580 | 8 |
| chr18-54114769-54115270 | 1 |
| chr14-32219644-32220145 | 5 |
| chr7-30281035-30281536 | 8 |
| chr17-70176916-70177417 | 5 |
| chr14-58082504-58083005 | 5 |
| chr16-9631218-9631719 | 8 |
| chr10-72402934-72403435 | 8 |
| chr4-63036960-63037461 | 5 |
| chr16-13326415-13326916 | 5 |
| chr14-2726841-2727342 | 6 |
| chr2-248737573-248738074 | 6 |
| chr15-29033050-29033551 | 6 |
| chr18-39700168-39700669 | 8 |
| chr5-123163565-123164066 | 8 |
| chr3-124483784-124484285 | 8 |
| chr16-46483810-46484311 | 1 |
| chr12-11770061-11770562 | 8 |
| chr10-91292956-91293457 | 8 |
| chr15-1138558-1139059 | 8 |
| chr15-18496748-18497249 | 6 |
| chr6-58881805-58882306 | 8 |
| chr7-25613223-25613724 | 8 |
| chr4-63427657-63428158 | 8 |
| chr11-17099239-17099740 | 6 |
| chr3-49867775-49868276 | 5 |
| chr19-14276924-14277425 | 5 |
| chr11-37931452-37931953 | 6 |
| chr5-76853081-76853582 | 6 |
| chr13-85333884-85334385 | 6 |
| chr10-33402538-33403039 | 1 |
| chr7-108337003-108337504 | 5 |
| chr16-45945767-45946268 | 5 |
| chr7-92335136-92335637 | 6 |
| chr13-66592114-66592615 | 5 |
| chr2-100192376-100192877 | 5 |

| | |
|---|---|
| chr4-160919260-160919761 | 5 |
| chr2-37987908-37988409 | 1 |
| chr2-162242313-162242814 | 6 |
| chr11-62075463-62075964 | 1 |
| chr8-101222586-101223087 | 8 |
| chr3-157178008-157178509 | 8 |
| chr12-7407889-7408390 | 5 |
| chr1-96453152-96453653 | 8 |
| chr1-170786041-170786542 | 8 |
| chr10-95430440-95430941 | 8 |
| chr2-118320717-118321218 | 8 |
| chr13-23961882-23962383 | 6 |
| chr3-113196342-113196843 | 8 |
| chr9-61726951-61727452 | 8 |
| chr4-7692058-7692559 | 6 |
| chr3-18266125-18266626 | 8 |
| chr2-189397925-189398426 | 8 |
| chr17-24750866-24751367 | 8 |
| chr3-17453607-17454108 | 8 |
| chr5-46035709-46036210 | 6 |
| chrX-9686767-9687268 | 5 |
| chr10-11084013-11084514 | 8 |
| chr1-104707875-104708376 | 6 |
| chr5-76897678-76898179 | 5 |
| chr5-130217303-130217804 | 8 |
| chr10-98043324-98043825 | 8 |
| chr8-61795258-61795759 | 8 |
| chr8-109293838-109294339 | 8 |
| chr1-132042766-132043267 | 6 |
| chr6-83533011-83533512 | 5 |
| chr20-2506431-2506932 | 6 |
| chr13-72000907-72001408 | 6 |
| chr1-174267358-174267859 | 8 |
| chr19-44572650-44573151 | 6 |
| chr7-129355299-129355800 | 8 |
| chr3-43023361-43023862 | 8 |
| chr3-91922010-91922511 | 6 |
| chr17-23789304-23789805 | 8 |
| chr7-67538614-67539115 | 6 |
| chr18-35433083-35433584 | 8 |
| chr1-48290901-48291402 | 8 |
| chr7-76252356-76252857 | 6 |
| chr18-35216086-35216587 | 5 |

| | |
|---|---|
| chr3-18500406-18500907 | 8 |
| chr2-175209253-175209754 | 8 |
| chr5-157019020-157019521 | 6 |
| chr13-48259145-48259646 | 1 |
| chr3-106247328-106247829 | 5 |
| chr4-67478806-67479307 | 8 |
| chr1-145407526-145408027 | 8 |
| chrX-132728585-132729086 | 6 |
| chr17-19443977-19444478 | 8 |
| chr3-140110231-140110732 | 6 |
| chr18-35600251-35600752 | 6 |
| chr17-7784282-7784783 | 8 |
| chr11-20531867-20532368 | 6 |
| chr2-242965306-242965807 | 8 |
| chr18-34411662-34412163 | 8 |
| chr2-185211731-185212232 | 5 |
| chr13-73332269-73332770 | 8 |
| chr13-89690406-89690907 | 1 |
| chr4-124456354-124456855 | 5 |
| chr16-3321150-3321651 | 8 |
| chr13-89789363-89789864 | 5 |
| chr1-146926392-146926893 | 8 |
| chr10-12642940-12643441 | 8 |
| chr13-30594811-30595312 | 1 |
| chr2-167859421-167859922 | 6 |
| chr17-30092294-30092795 | 6 |
| chr9-69271401-69271902 | 5 |
| chr4-121580910-121581411 | 8 |
| chr3-96766738-96767239 | 8 |
| chr2-135400042-135400543 | 6 |
| chr7-47043076-47043577 | 5 |
| chr7-57510674-57511175 | 8 |
| chr7-51320377-51320878 | 1 |
| chr13-103090742-103091243 | 8 |
| chr17-76415633-76416134 | 6 |
| chr2-148429149-148429650 | 8 |
| chr1-32096792-32097293 | 8 |
| chr10-28701282-28701783 | 8 |
| chr2-188904643-188905144 | 8 |
| chr6-35924650-35925151 | 1 |
| chr11-69698373-69698874 | 8 |
| chr16-43815695-43816196 | 8 |
| chr17-22915810-22916311 | 8 |

| | |
|---|---|
| chr8-123417977-123418478 | 5 |
| chr3-106755022-106755523 | 6 |
| chr7-33001899-33002400 | 5 |
| chr3-10700338-10700839 | 8 |
| chr7-89755285-89755786 | 6 |
| chr2-38399993-38400494 | 5 |
| chr1-183798091-183798592 | 5 |
| chr6-14838455-14838956 | 8 |
| chr7-116724187-116724688 | 8 |
| chr4-63890996-63891497 | 8 |
| chr19-25650073-25650574 | 6 |
| chr12-40359687-40360188 | 8 |
| chr19-43373931-43374432 | 6 |
| chr17-42596485-42596986 | 1 |
| chrX-115166046-115166547 | 8 |
| chr1-155081560-155082061 | 6 |
| chr17-26887761-26888262 | 8 |
| chr4-68291492-68291993 | 5 |
| chr13-97885132-97885633 | 8 |
| chr18-7345072-7345573 | 5 |
| chr6-19944560-19945061 | 8 |
| chr19-51238446-51238947 | 1 |
| chr8-55132997-55133498 | 8 |
| chr13-81182119-81182620 | 5 |
| chr2-92707318-92707819 | 8 |
| chr4-57370941-57371442 | 6 |
| chr10-102149490-102149991 | 8 |
| chr9-55970159-55970660 | 8 |
| chr1-225671170-225671671 | 5 |
| chr4-7589709-7590210 | 8 |
| chr8-64882780-64883281 | 8 |
| chr10-57904859-57905360 | 8 |
| chr10-3671435-3671936 | 5 |
| chr13-93376628-93377129 | 5 |
| chr3-169019472-169019973 | 5 |
| chr13-55140997-55141498 | 5 |
| chr1-96309911-96310412 | 5 |
| chr2-165038842-165039343 | 1 |
| chr16-7266126-7266627 | 1 |
| chr4-40525890-40526391 | 6 |
| chr20-18299785-18300286 | 8 |
| chr6-38777226-38777727 | 8 |
| chr4-162385527-162386028 | 8 |

| | |
|---|---|
| chr2-152521415-152521916 | 1 |
| chr12-12218267-12218768 | 8 |
| chr4-114616576-114617077 | 6 |
| chr14-15471142-15471643 | 5 |
| chr2-184687520-184688021 | 8 |
| chr2-190648675-190649176 | 8 |
| chr6-15883215-15883716 | 5 |
| chr5-80386509-80387010 | 6 |
| chr9-18164835-18165336 | 6 |
| chr13-55707698-55708199 | 8 |
| chr9-106955619-106956120 | 8 |
| chr6-127220602-127221103 | 8 |
| chr7-72405594-72406095 | 8 |
| chr13-98453227-98453728 | 6 |
| chr18-5781700-5782201 | 8 |
| chr2-92788320-92788821 | 8 |
| chr9-107388993-107389494 | 8 |
| chr3-116645689-116646190 | 1 |
| chr14-96818577-96819078 | 8 |
| chr4-122844023-122844524 | 6 |
| chr13-43719805-43720306 | 8 |
| chr2-187515679-187516180 | 6 |
| chrX-16191565-16192066 | 1 |
| chr4-58637535-58638036 | 1 |
| chr14-15900114-15900615 | 8 |
| chr1-34424840-34425341 | 8 |
| chr14-86982981-86983482 | 6 |
| chr2-226801182-226801683 | 8 |
| chr18-13713149-13713650 | 6 |
| chr18-49094294-49094795 | 8 |
| chr17-12263901-12264402 | 5 |
| chr16-79938564-79939065 | 6 |
| chr14-41587379-41587880 | 8 |
| chr3-25550461-25550962 | 6 |
| chr3-135585027-135585528 | 8 |
| chr3-122830501-122831002 | 6 |
| chr6-22709003-22709504 | 6 |
| chr5-79422890-79423391 | 8 |
| chr1-77282516-77283017 | 8 |
| chr19-52800603-52801104 | 6 |
| chr5-22529823-22530324 | 5 |
| chr10-37674284-37674785 | 5 |
| chr10-33321623-33322124 | 6 |

| | |
|---|---|
| chr4-104017692-104018193 | 5 |
| chr3-50081618-50082119 | 8 |
| chr1-153999584-154000085 | 5 |
| chr6-92196308-92196809 | 5 |
| chr8-24057332-24057833 | 1 |
| chr14-77054285-77054786 | 5 |
| chr6-88604480-88604981 | 6 |
| chr11-3973787-3974288 | 1 |
| chr2-223627978-223628479 | 8 |
| chrX-108103877-108104378 | 6 |
| chr13-98501178-98501679 | 5 |
| chr3-11497810-11498311 | 8 |
| chr17-70479256-70479757 | 6 |
| chr2-204073985-204074486 | 8 |
| chr17-81905856-81906357 | 5 |
| chr10-39940295-39940796 | 6 |
| chr1-47403672-47404173 | 6 |
| chr4-148593071-148593572 | 5 |
| chr9-24739201-24739702 | 6 |
| chr17-32588891-32589392 | 8 |
| chr3-139779874-139780375 | 6 |
| chr2-46785621-46786122 | 8 |
| chr2-42504508-42505009 | 8 |
| chr17-52460074-52460575 | 5 |
| chr3-27970517-27971018 | 6 |
| chrX-97131594-97132095 | 1 |
| chr3-127032652-127033153 | 5 |
| chr7-14373551-14374052 | 8 |
| chr17-13959533-13960034 | 8 |
| chr17-32022737-32023238 | 6 |
| chr10-91291417-91291918 | 5 |
| chr1-224469864-224470365 | 1 |
| chr1-137624994-137625495 | 8 |
| chr6-51177025-51177526 | 8 |
| chr5-119436929-119437430 | 6 |
| chr18-35639948-35640449 | 6 |
| chr10-57845906-57846407 | 1 |
| chr11-66238402-66238903 | 5 |
| chr1-78344847-78345348 | 6 |
| chr4-77386057-77386558 | 6 |
| chr1-180418636-180419137 | 5 |
| chr9-65416331-65416832 | 5 |
| chr15-92160811-92161312 | 5 |

| | |
|---|---|
| chr17-3800724-3801225 | 8 |
| chrX-24637730-24638231 | 5 |
| chr19-55744319-55744820 | 6 |
| chr17-27205454-27205955 | 1 |
| chr5-59941027-59941528 | 1 |
| chr6-21881267-21881768 | 8 |
| chr1-41942600-41943101 | 8 |
| chr10-7765845-7766346 | 8 |
| chr13-53159317-53159818 | 6 |
| chr7-60480188-60480689 | 6 |
| chr14-60566954-60567455 | 6 |
| chr17-17796008-17796509 | 5 |
| chr2-50288760-50289261 | 6 |
| chr16-7637955-7638456 | 5 |
| chr13-41418978-41419479 | 6 |
| chr4-121494181-121494682 | 5 |
| chr7-84927847-84928348 | 8 |
| chr5-93653191-93653692 | 6 |
| chr2-33495623-33496124 | 8 |
| chr2-77118396-77118897 | 6 |
| chr16-15105227-15105728 | 8 |
| chr4-10246816-10247317 | 8 |
| chr13-76356134-76356635 | 6 |
| chr13-95022371-95022872 | 6 |
| chr11-33271347-33271848 | 6 |
| chr3-76791122-76791623 | 8 |
| chr13-103239335-103239836 | 6 |
| chr1-13196232-13196733 | 6 |
| chr4-146899705-146900206 | 5 |
| chr5-77940969-77941470 | 8 |
| chr4-59005170-59005671 | 8 |
| chr3-148008819-148009320 | 1 |
| chr10-63865624-63866125 | 8 |
| chr9-1226459-1226960 | 8 |
| chr3-94394024-94394525 | 6 |
| chr5-71390687-71391188 | 5 |
| chr2-78083456-78083957 | 6 |
| chr15-36498579-36499080 | 6 |
| chrX-9350717-9351218 | 1 |
| chr9-49392746-49393247 | 8 |
| chr1-106238446-106238947 | 5 |
| chr15-78685842-78686343 | 5 |
| chr4-36446998-36447499 | 8 |

| | |
|---|---|
| chr6-20657142-20657643 | 6 |
| chr11-76728022-76728523 | 5 |
| chr5-149992818-149993319 | 8 |
| chr13-40239278-40239779 | 8 |
| chr8-70753843-70754344 | 1 |
| chr15-75970866-75971367 | 8 |
| chr17-76100285-76100786 | 6 |
| chr5-141850160-141850661 | 5 |
| chr2-76013837-76014338 | 1 |
| chr7-9512628-9513129 | 8 |
| chr7-27342672-27343173 | 8 |
| chr4-130910452-130910953 | 6 |
| chr17-29704647-29705148 | 1 |
| chr20-19269560-19270061 | 6 |
| chr14-98498527-98499028 | 8 |
| chr1-252946769-252947270 | 5 |
| chr5-144578146-144578647 | 8 |
| chr13-46488211-46488712 | 8 |
| chr9-74886421-74886922 | 6 |
| chr4-57023782-57024283 | 8 |
| chr10-16395000-16395501 | 6 |
| chr3-22708203-22708704 | 1 |
| chr15-98810334-98810835 | 1 |
| chr5-43364171-43364672 | 6 |
| chr16-20590889-20591390 | 8 |
| chr14-13764726-13765227 | 6 |
| chr8-23812710-23813211 | 5 |
| chr7-119773254-119773755 | 5 |
| chr3-158927317-158927818 | 8 |
| chr7-48282780-48283281 | 5 |
| chr4-63245905-63246406 | 6 |
| chr20-52002141-52002642 | 5 |
| chr4-36945897-36946398 | 6 |
| chrX-52147132-52147633 | 6 |
| chr19-776596-777097 | 5 |
| chr4-66510966-66511467 | 6 |
| chr2-143710676-143711177 | 8 |
| chr7-111341183-111341684 | 6 |
| chr5-165354522-165355023 | 8 |
| chr15-3031596-3032097 | 8 |
| chr4-83967304-83967805 | 8 |
| chr6-112933861-112934362 | 6 |
| chr2-26326389-26326890 | 6 |

| | |
|---|---|
| chr8-44197998-44198499 | 8 |
| chr1-96470689-96471190 | 8 |
| chr2-66071283-66071784 | 8 |
| chr9-24146236-24146737 | 8 |
| chr19-48394954-48395455 | 8 |
| chr10-83242761-83243262 | 8 |
| chr19-6305344-6305845 | 6 |
| chr4-80588221-80588722 | 5 |
| chr12-20775938-20776439 | 8 |
| chr3-78552405-78552906 | 5 |
| chr4-68809717-68810218 | 8 |
| chr15-50483647-50484148 | 5 |
| chr4-10516234-10516735 | 6 |
| chr10-93265267-93265768 | 8 |
| chr19-49357084-49357585 | 6 |
| chr8-30847372-30847873 | 1 |
| chr13-38184977-38185478 | 6 |
| chr8-121602556-121603057 | 6 |
| chrX-115122809-115123310 | 8 |
| chr13-77800850-77801351 | 5 |
| chr6-101399385-101399886 | 1 |
| chr16-67755730-67756231 | 6 |
| chr1-139376081-139376582 | 6 |
| chr2-52094611-52095112 | 6 |
| chr2-11263262-11263763 | 8 |
| chr7-92711073-92711574 | 8 |
| chr3-147335893-147336394 | 1 |
| chr13-72375540-72376041 | 8 |
| chr7-68523915-68524416 | 8 |
| chr17-18142240-18142741 | 8 |
| chr15-32134045-32134546 | 1 |
| chr3-48419251-48419752 | 6 |
| chr5-47166937-47167438 | 5 |
| chr16-55596844-55597345 | 6 |
| chr20-19027765-19028266 | 6 |
| chr1-129399610-129400111 | 8 |
| chr1-196847831-196848332 | 8 |
| chr3-57455337-57455838 | 8 |
| chr2-163483695-163484196 | 6 |
| chr11-26933179-26933680 | 6 |
| chr8-47917445-47917946 | 8 |
| chr4-77634730-77635231 | 1 |
| chr15-40361355-40361856 | 8 |

| | |
|---|---|
| chr2-92932769-92933270 | 6 |
| chr3-13841195-13841696 | 8 |
| chr1-26931013-26931514 | 1 |
| chr17-48153959-48154460 | 6 |
| chr6-101327454-101327955 | 1 |
| chr12-9733046-9733547 | 1 |
| chr15-74838048-74838549 | 8 |
| chr15-91987886-91988387 | 6 |
| chr16-76042967-76043468 | 5 |
| chr17-3943767-3944268 | 8 |
| chr10-18423235-18423736 | 8 |
| chr1-42031088-42031589 | 1 |
| chr19-20813892-20814393 | 1 |
| chr6-5339930-5340431 | 1 |
| chr16-8819485-8819986 | 8 |
| chr18-58959949-58960450 | 8 |
| chr18-61792844-61793345 | 8 |
| chr7-52763106-52763607 | 8 |
| chr10-97865173-97865674 | 8 |
| chr2-137776263-137776764 | 8 |
| chr17-10013657-10014158 | 8 |
| chr2-56274187-56274688 | 1 |
| chr3-100633815-100634316 | 6 |
| chr1-46330818-46331319 | 1 |
| chr4-150835042-150835543 | 5 |
| chr2-77103688-77104189 | 8 |
| chr19-48418511-48419012 | 6 |
| chr13-103216122-103216623 | 8 |
| chr9-50688426-50688927 | 8 |
| chr2-208756075-208756576 | 8 |
| chr18-5020965-5021466 | 6 |
| chr5-14969813-14970314 | 1 |
| chr19-14348379-14348880 | 8 |
| chr3-122742990-122743491 | 1 |
| chr3-54909618-54910119 | 8 |
| chr14-32291713-32292214 | 6 |
| chr13-28180948-28181449 | 6 |
| chr14-86646648-86647149 | 8 |
| chr7-109924973-109925474 | 1 |
| chr6-14640570-14641071 | 5 |
| chr1-176801325-176801826 | 8 |
| chr8-72673299-72673800 | 8 |
| chr7-27633493-27633994 | 6 |

| | |
|---|---|
| chr1-83802937-83803438 | 8 |
| chr12-15582029-15582530 | 1 |
| chr3-152413930-152414431 | 1 |
| chr9-68627069-68627570 | 8 |
| chr2-229900121-229900622 | 1 |
